# Mako: a graph-based pattern growth approach to detect complex structural variants

**DOI:** 10.1101/2021.03.01.433465

**Authors:** Jiadong Lin, Xiaofei Yang, Walter Kosters, Tun Xu, Yanyan Jia, Songbo Wang, Qihui Zhu, Mallory Ryan, Li Guo, Chengsheng Zhang, The Human Genome Structural Variation Consortium, Charles Lee, Scott E. Devine, Evan E. Eichler, Kai Ye

## Abstract

Complex structural variants (CSVs) are genomic alterations that have more than two breakpoints and are considered as simultaneous occurrence of simple structural variants. However, detecting the compounded mutational signals of CSVs is challenging through a commonly used model-match strategy. As a result, there has been limited progress for CSV discovery compared with simple structural variants. We systematically analyzed the multi-breakpoint connection feature of CSVs, and proposed Mako, utilizing a bottom-up guided model-free strategy, to detect CSVs from paired-end short-read sequencing. Specifically, we implemented a graph-based pattern growth approach, where the graph depicts potential breakpoint connections and pattern growth enables CSV detection without predefined models. Comprehensive evaluations on both simulated and real datasets revealed that Mako outperformed other algorithms. Notably, validation rates of CSV on real data based on experimental and computational validations as well as manual inspections are around 70%, where the medians of experimental and computational breakpoint shift are 13bp and 26bp, respectively. Moreover, Mako CSV subgraph effectively characterized the breakpoint connections of a CSV event and uncovered a total of 15 CSV types, including two novel types of adjacent segments swap and tandem dispersed duplication. Further analysis of these CSVs also revealed impact of sequence homology in the formation of CSVs. Mako is publicly available at https://github.com/jiadong324/Mako.

## Introduction

Computational methods based on next-generation-sequencing (NGS) have provided an increasingly comprehensive discovery and catalog of simple structure variants (SVs) that usually have two breakpoints, such as deletions and inversions [1–7]. In general, these approaches follow a model-match strategy, where a specific SV model and its corresponding mutational signal model is proposed. Afterwards, the mutational signal model is used to match observed signals for the detection (**Figure 1A**). This model-match strategy has been proved effective for detecting simple SVs, providing us with prominent opportunities to study and understand genome evaluation and disease progression [8–11]. However, recent research has revealed that some rearrangements have multiple, compounded mutational signals and usually cannot fit into the simple SV models [8, 12–16] (**Figure 1B**). For example, in 2015, Sudmant et al. systematically categorized 5 types of complex structural variants (CSVs) and found that a remarkable 80% of 229 inversion sites were complex events [8]. Collins et al. used long-insert size whole genome sequencing (liWGS) on autism spectrum disease (ASD) and successfully resolved 16 classes of 9,666 CSVs from 686 patients [17]. In 2019, Lee et al. revealed that 74% of known fusion oncogenes of lung adenocarcinomas were caused by complex genomic rearrangements, including *EML4-ALK* and *CD74-ROS1* [16]. Though less frequently reported compared with simple SVs, these multiple breakpoint rearrangements were considered as punctuated events, leading to severe genome alterations at once [10, 18–21]. This dramatic change of genome provided distinctive evidence to study formation mechanisms of rearrangement and to understand cancer genome evolution [13, 14, 17, 19, 21–25].

**Figure 1.**
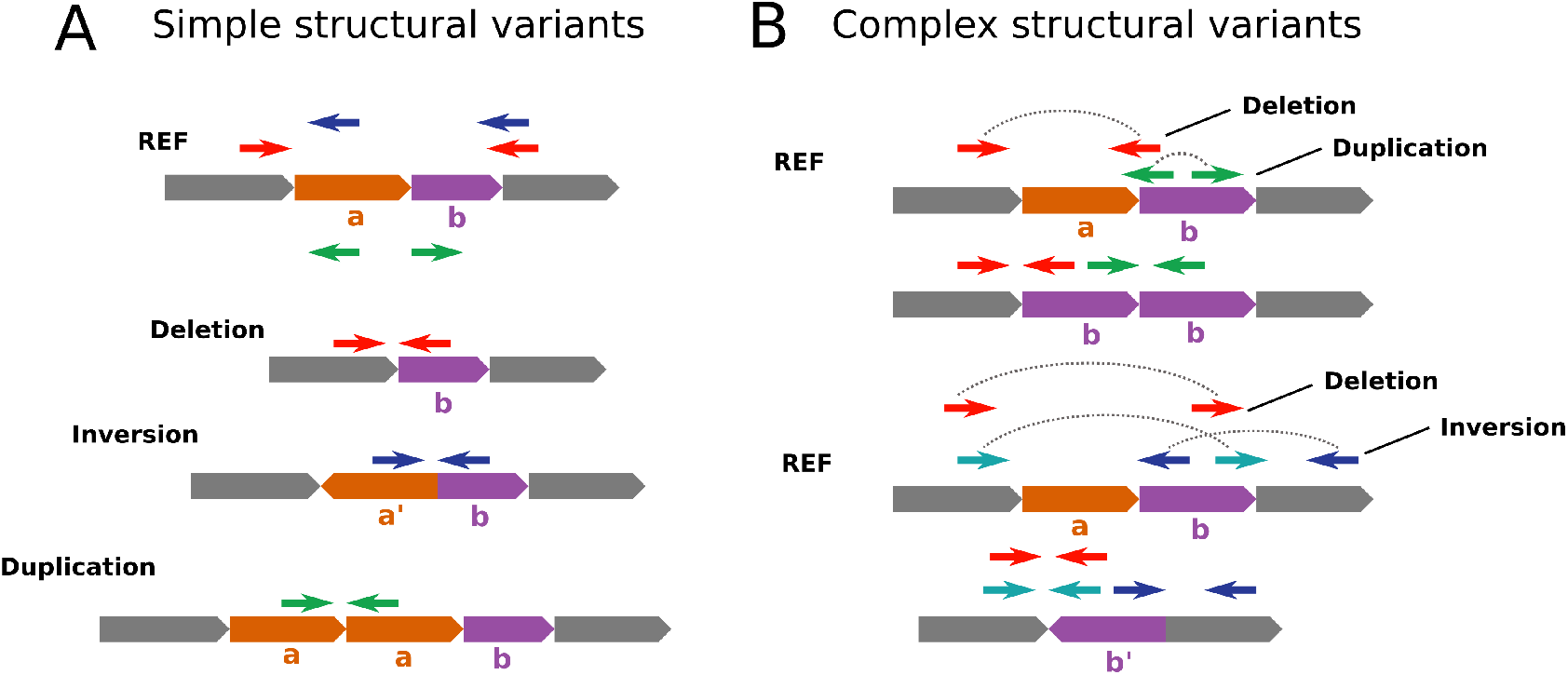
Explanation of simple and complex structure variants alignment models derived from abnormal read pairs. (**A**) Three common simple SV and their corresponding abnormal read pair alignment on the reference genome, representing by red, blue and green arrows. (**B**) The alignment signature of two CSVs, each of them involves two types of signature that can be matched by simple SV alignment model.

However, due to lack of effective CSV detection algorithms, most CSV related studies screen these events from the “sea” of simple SVs through computational expensive contig assembly and realignment, incomplete breakpoints clustering or even targeted manual inspection [8, 12, 16]. In fact, many CSVs have been already neglected or misclassified in this “sea” because of the incompatibility between complicated mutational signals and existing SV models. Although the importance and challenge for CSV detection have been recognized, only a few dedicated algorithms were proposed for CSVs discovery, and they followed two major approaches guided by the model-match strategy. TARDIS and SVelter utilizes the top-down approach, where they attempt to model all the mutational signals of a CSV event instead of modeling specific parts of signals. In particular, TARDIS [26] proposed sophisticated abnormal alignment models to depict the mutational signals reflected by dispersed duplication and inverted duplication. The pre-defined models were then used to fit observed signals from alignments for the detection of the two specific CSV types. Indeed, this was complicated and greatly limited by the diversity types of CSV. To solve this, SVelter [27] replaced the modeling process for specific CSVs with a randomly created virtual rearrangement. And CSVs were detected by minimizing the difference between the virtual rearrangement and the observed signals. Whereas GRIDSS [28] represents the assembly-based approach, which detected CSVs through extra breakpoints discovered from contig-assembly and realignment. Though assembly-based approach is sensitive for breakpoint detection, it lacks certain regulations to constrain or classify these breakpoints and leave them as independent events. As a result, these model-match guided approaches would substantially break-up or misinterpret the CSVs because of partially matched signals (**Figure 1B**). Moreover, graph is another approach that has been widely used for simple [2, 29] and complex [19, 30] SV detection. Notably, ARC-SV [30] uses clustered discordant read-pairs to construct an adjacency graph and adopts a maximum likelihood model to detection complex SVs, showing great potential of using graph to detect complex SVs. Accordingly, there is an urgent demand of a new strategy, enabling CSV detection without predefined models as well as maintaining the completeness of a CSV event.

In this study, we proposed a bottom-up guided model-free strategy, implemented as Mako, to effectively discover CSVs all at once based on short-read sequencing. Specifically, Mako uses a graph to build connections of mutational signals derived from abnormal alignment, providing the potential breakpoint connections of CSVs. Meanwhile, Mako replaces model fitting with the detection of maximal subgraphs through a pattern growth approach. Pattern growth is a bottom-up approach, which captures the natural features of data without sophisticated model generation, allowing CSV detection without predefined models. We benchmarked Mako against five widely used tools on a series of simulated and real data. The results show that Mako is an effective and efficient algorithm for CSV discovery, which will provide more opportunities to study genome evolution and disease progression from large cohorts. Remarkably, the analysis of subgraphs detected by Mako highlights the unique strength of Mako, where Mako was able to effectively characterize the CSV breakpoint connections, confirming the completeness of a CSV event. Moreover, we systematically analyzed the CSVs detected by Mako on three healthy samples, revealing a novel role of sequence homology in CSV formation.

## Results

In this section, we give an overview of the Mako algorithm, with full details available in the Methods section. For performance comparison, we propose all-breakpoint match and unique-interval match to evaluate Mako against five published methods on both simulated and real data. The detailed explanation of the evaluation measurements, CSV simulation and real data CSV benchmarks are described in the Methods section. Additionally, we describe our observations of Mako’s CSV discovery from HG00514, HG00733 and NA19240. These samples are sequenced by Human Structural Variants Consortium (HGSVC) and publicly available.

### Overview of the Mako algorithm

Given the fact that a CSV is a single event with multiple breakpoint connections, we observe that either false positive breakpoints or breakpoints from other events will not have connection with the breakpoints in the current CSV because of weak or non-exist connections. Thus, we formulate the detection of CSVs as maximal subgraph pattern detection in a signal graph. To detect the CSV subgraphs, Mako comprises two major steps (**Figure 2**). Firstly, it collects abnormal aligned reads clusters as nodes and uses two types of edges to build the so-called signal graph. To build the high-quality graph, we filtered discordance alignments based on procedure described in BreakDancer [4] (**Methods**). The resulting signal graph is formally defined as follows:

**Figure 2.**
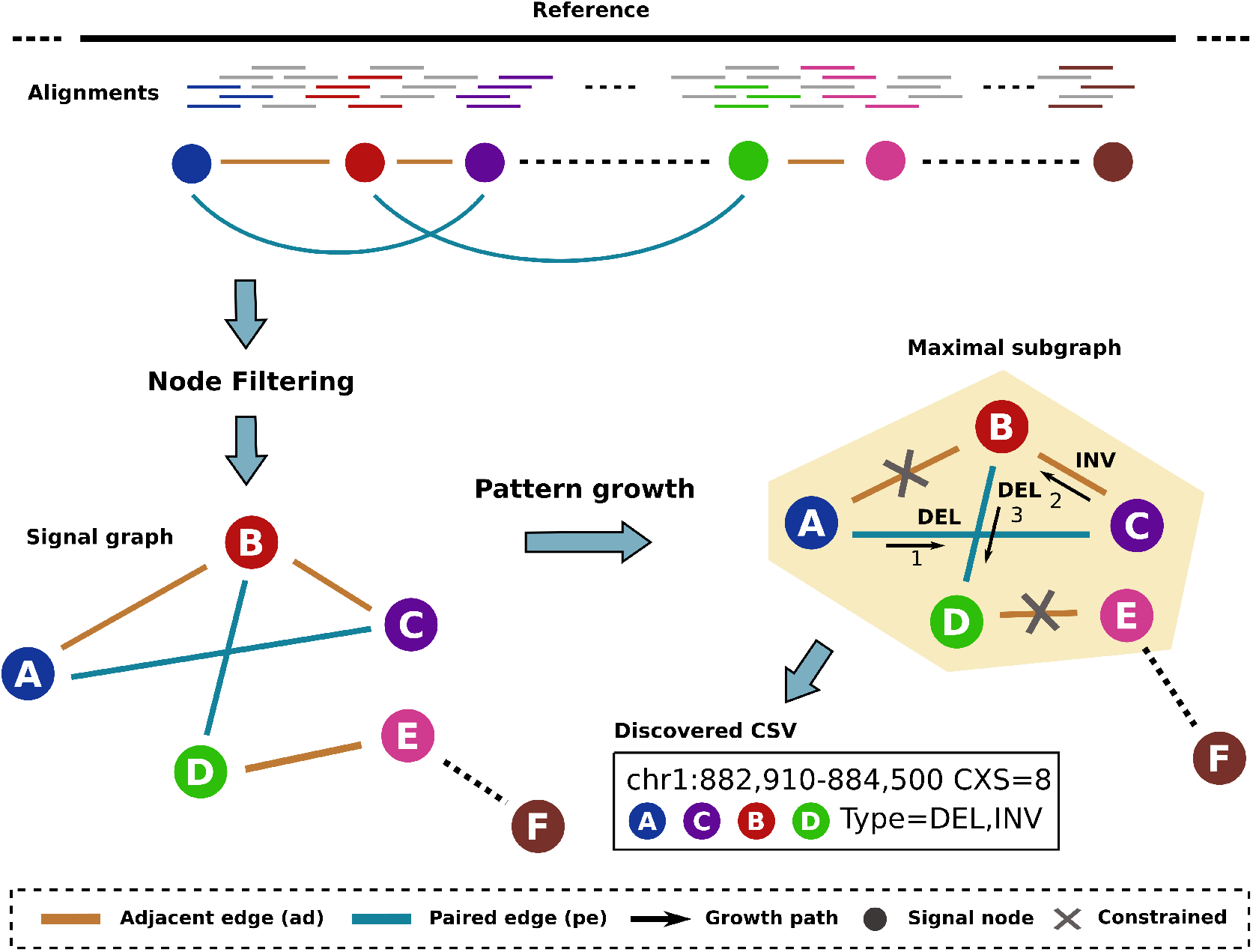
Overview of Mako for identifying CSVs from NGS data. Mako first builds a signal graph by collecting abnormal aligned reads as nodes and their edge connections are provided by paired-end alignment and split alignment. Afterwards, Mako utilizes the pattern growth approach to find a maximal subgraph as potential CSV site. In the example output, the maximal subgraph contains A, B, C, D, whereas F is not able to appended because of none existing edge (dashed line). The CSV is derived from this subgraph with estimate breakpoints and CXS score, where the discovered CSV subgraph contains four different nodes, one *E_ae_* edge and two *E_pe_* edges of type DEL and INV, thus CXS = 8.

*G* = (*V, E*) with *V* = {*ν*_1_, *ν*_2_,…, *ν_n_*} and *E* = {*E_pe_, E_ae_*}, where each node *ν* ∈ *V* is represented as *ν* = (*type, pos, weight*), and each edge in *E_pe_* and *E_ae_* is represented as either *e_pe_* = (*ν_i_, ν_j_, rp*) or *e_ae_* = (*ν_i_, ν_j_, dist*), with *ν_i_, ν_j_* ∈ *V*. In particular, Mako uses weight and the ratio between weight and coverage at pos to filter nodes, which are created separately by clustering discordant read-pairs, clipped reads and split reads (**Methods**). For the edge set, *E_pe_* contains the paired edges that represent connections between two signals on the genome derived from paired-reads or split-alignment, while *E_e_* consists of adjacency edges that indicate distances between signals along the genome. Afterwards, Mako applies a pattern growth search strategy to efficiently discover these subgraphs as potential CSVs at whole genome scale. Meanwhile, the attributes of the subgraph are used to measure the complexity and to define the types of CSVs. Specifically, the CSVs types are given by the edge connection types of the corresponding subgraphs (**Figure 2**).

### Mako effectively characterizes multiple breakpoints of CSV

The most important feature for a CSV is the presence of multiple breakpoints in a single event. Thus, we first examined the performance of breakpoint detection for Mako, Lumpy, Manta, SVelter, TARDIS and GRIDSS. The results were evaluated according to the all-breakpoint match criteria on both reported and randomized CSV type simulations (**Methods**). For convenience, we used the terms reported CSV and randomized CSV throughout this study. Overall, for the heterozygous (HET) (**Figure 3A**) and homozygous (HOM) (**Figure 3B**) simulation, Mako was comparable to GRIDSS and they outperformed other algorithms. For example, GRIDSS, Mako and Lumpy detected 50%, 51% and 46% for reported HET CSV breakpoints, while they reported 53%, 54% and 44% for randomized ones. Because the graph encoded both multiple breakpoints and their substantial connections for each CSV, Mako achieved better performance on randomized events, which included more subcomponents than the reported ones. Indeed, by comparing reported and randomized simulation, the breakpoint detection sensitivity (**Figure 3A** and **Figure 3B**) of Mako increased, while that of other algorithms dropped except for GRIDSS. Although the assembly-based method, GRIDSS, is as effective as Mako for breakpoint detection, it lacks a proper procedure to resolve the connections among breakpoints.

**Figure 3.**
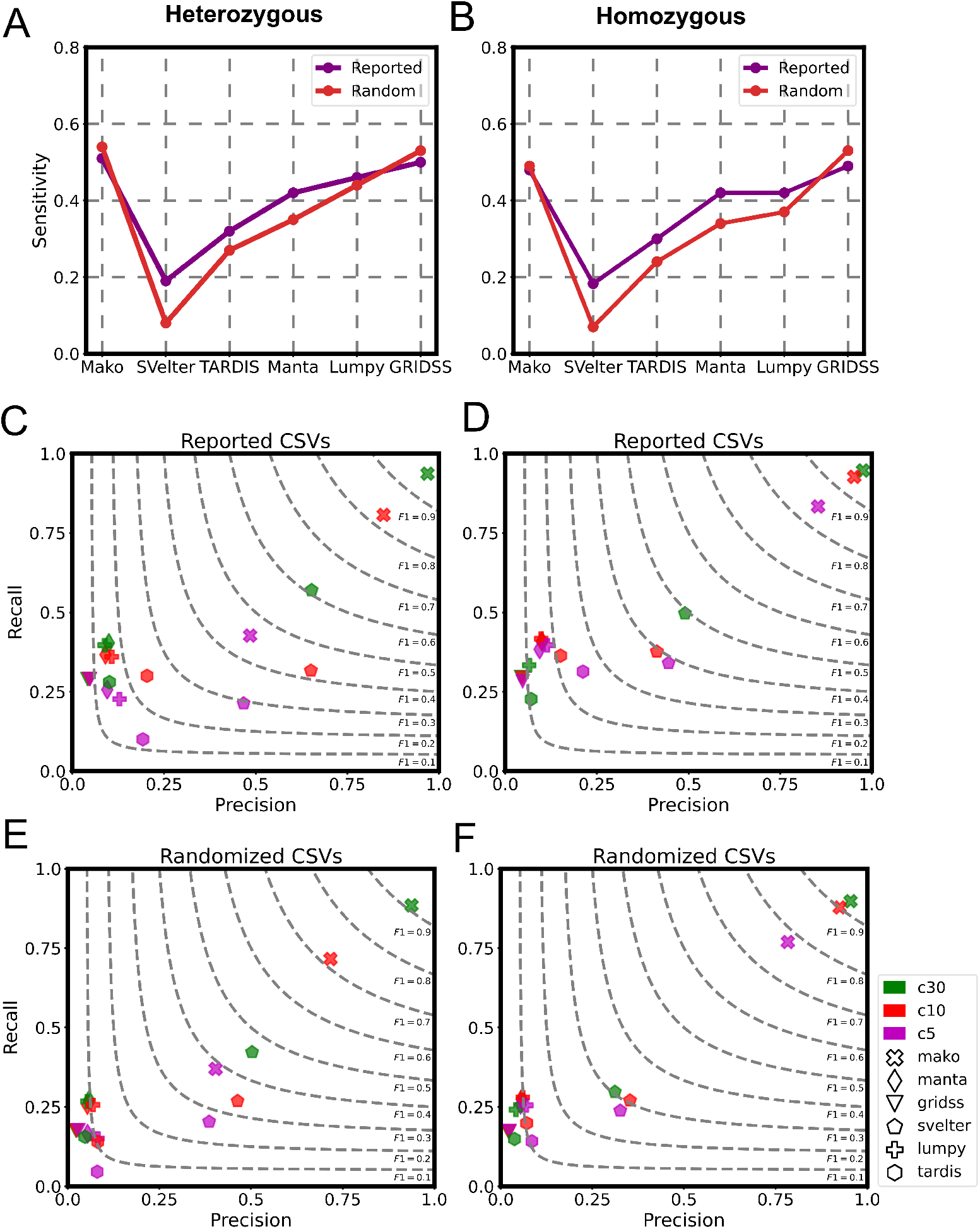
Performance comparison on simulated CSVs with different match criteria. All-breakpoint match (**A** and **B**) and unique-interval match (**C-F**) evaluation of selected tools on simulated CSVs. (**A**) The sensitivity of detecting heterozygous CSVs breakpoints. (**B**) The sensitivity of detecting homozygous CSVs breakpoints. The red and purple curve indicate randomized and reported CSVs, respectively. (**C**) Evaluation of reported heterozygous CSV simulation. (**D**) Evaluation of reported homozygous CSV simulation. (**E**) Evaluation of randomized heterozygous CSV simulation. (**F**) Evaluation of randomized homozygous CSV simulation. From (**C**) to (**F**), the performance is evaluated by recall (y-axis), precision (x-axis) and F1-score (dotted lines). The right top corner of the plot indicates better performance. The c5-c30 indicates coverage, e.g. c5 indicates 5X coverage.

### Mako precisely discovers CSV unique-interval

CSV is considered as a single event consisted of connected breakpoints and we have demonstrated that Mako was able to detect CSV breakpoints effectively. However, the breakpoint detection evaluation only assesses the discovery of basic components for a CSV and lacks examination for CSV completeness. We then investigated whether Mako could precisely capture the entire CSV interval even with missing breakpoints. In general, according to the unique-interval match (**Methods**) criteria, Mako consistently outperformed other algorithms for both reported and randomly created CSVs, while SVelter and GRIDSS ranked second and third, respectively. For the reported CSVs at 30X coverage (**Figure 3C** and **Figure 3D**), the recall of Mako was 94% and 92%, which was significantly higher than SVelter (49% and 57%) for both reported HET and HOM CSVs, respectively. Due to the model guided top-down approach, SVelter was able to discover some complete CSV events. However, the virtual rearrangement generation may not fully explore all possibilities. Remarkably, we noted that Mako’s superior sensitivity was most significant for randomized simulation (**Figure 3E** and **Figure 3F**), which was consistent with our previous observation (**Figure 3A, Figure 3B**). In particular, at 30X coverage, Mako’s recall (88%) was much higher than SVelter (29%) for the HET CSVs (**Figure 3E**). This was due to the complementary nature of the graph edges (adjacent and paired), from which the subgraph can be expanded alternatively through one or the other, enabling the complete CSVs discovery even with missing breakpoints.

### Performance on real data

Since Mako outperformed other methods on simulated data, we further compared Mako with SVelter, GRIDSS and TARDIS on whole genome sequencing data of NA19240 and SKBR3. Firstly, we obtained 6,060, 7,733, 6,426 and 15,358 calls for NA19240, and 2,962, 2,468, 3,077 and 4,010 for SKBR3 predicted by Mako, SVelter, GRIDSS and TARDIS, respectively (**Methods**, **Supplementary Figure S1**-**S2**). By comparing their predictions, we found Mako and GRIDSS showed similar performance (**Figure 4A** and **4B**) which was consistent with our observation in simulated data (**Figure 3**). Furthermore, we examined the discovery completeness of 59 (NA19240) and 21 (SKBR3) benchmark CSVs (**Table 1**, **Supplementary File 1**, **Supplementary Table S1**). Because Manta and Lumpy contributed to the CSV benchmarks, they were excluded from the comparison. The results showed that Mako performed the best for the two benchmarks with different CXS thresholds, while TARDIS ranked second (**Figure 4C**). Given that inverted duplication and dispersed duplication dominated the benchmark set and that TARDIS has designed specific models for these two types, TARDIS detected more events of these two duplication types than others did (**Table 1**). SVelter only detected a few benchmark CSVs for SKBR3, because the procedure of randomly created rearrangement was not optimized, leading to either incorrect events or inaccurate breakpoints. Based on the above observation, we concluded that either randomized model (SVelter) or specific model (TARDIS) was far from comprehensive to cover the large diversity of real CSV types.

**Figure 4.**
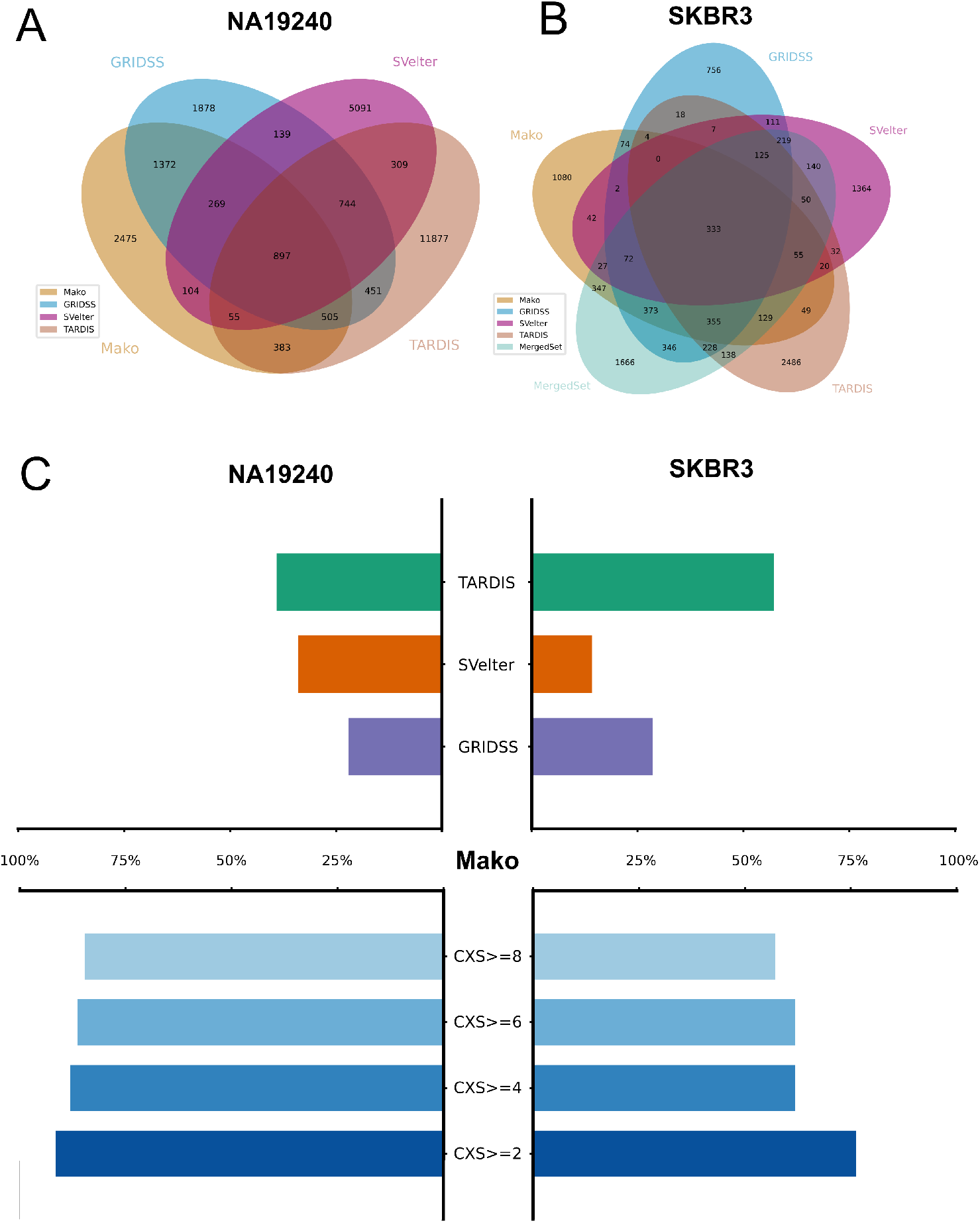
Overview of performance on NA19240 and SKBR3 for Mako, GRIDSS, SVelter and TARDIS. (**A**) and (**B**) are the Venn diagram of 50% reciprocal overlap between callsets for both NA19240 and SKBR3. They are created by a publicly available tool Intervene with – bedtools-options enabled. (**B**) The MergedSet is the callset provided by the publication. (**C**) The percentage of completely and uniquely discovered CSVs from the NA19240 and SKBR3. The results of Mako (bottom panel) are shown according to different CXS threshold.

**Table 1.** Summary of benchmark CSVs. The CSV type abbreviations and their corresponding descriptions are also listed.

| Benchmark summaries |  |  |  |
| --- | --- | --- | --- |
| Type | NA19240 | SKBR3 | Description |
| Disdup | 15 | 12 | Dispersed duplication |
| Invdup | 18 | - | Inverted duplication |
| DelInv | 7 | 5 | Deletion associated with inversion |
| DelDisdup | 5 | 1 | Deletion associated with dispersed duplication |
| DelInvdup | 1 | - | Deletion associated with inverted duplication |
| DisdupInvdup | 2 | 2 | Dispersed duplication with inverted duplication |
| InsInv | 1 | - | Insertion associated with inversion |
| Tantrans | 1 | - | Adjacent segments swap |
| DelSpaDel | 8 | 1 | Two deletions with inverted or non-inverted spacer |
| TanDisdup | 1 | - | Tandem dispersed duplications |

### CSV subgraph illustrates breakpoints connections

CSVs from autosomes were selected from Mako’s callset with more than one edge connection type observed in the subgraph, leading to 403, 609, and 556 events for HG00514, HG00733 and NA19240, respectively (**Figure 5A**, **Supplementary Table S2**). We systemically evaluated all CSV events in HG00733 via experimental and computational validation as well as manual inspection. For experimental validation, we successfully designed primers for 107 CSVs (**Supplementary Table S3**), where 15 out of 21 (71%, **Table 2**) successfully amplified were validated by Sanger sequencing (**Supplementary Table S4**). The computational validation (**Supplementary Figure S3**) showed up to 87% accuracy, indicating a combination of methods and external data is necessary for comprehensive CSV validation (**Table 3**, **Methods**). Further analysis showed that the medians of breakpoint shift were 13bp and 26bp comparing to breakpoints given by experimental and computational evaluation (**Supplementary Figure S4**). We observed that approximately 54% of CSVs were found in either STR or VNTR regions, contributing to 75% of all events inside the repetitive regions (**Figure 5A**). For the connection types, more than half of the events contains DUP and INS edges in the graph, indicating duplication involved sequence insertion. Moreover, around 40% of the events contain DEL edges (**Figure 5A**), showing two distant segment connections derived from either duplication or inversion events. We further examined whether the CSV subgraph depict the connections for each CSV via discordant read-pairs. Interestingly, we observed two representative events with four breakpoints at chr6:128,961,308-128,962,212 (**Figure 5B**) and chr5:151,511,018-151,516,780 (**Figure 5C**) from NA19240 and SKBR3, respectively. Both events were correctly detected by Mako, but missed by SVelter and reported more than once by GRIDSS and TARDIS (**Supplementary Table S5**). In particular, the CSV at chr6:128,961,308-128,962,212 that consists of two deletions and an inverted spacer was reported twice and five times by GRIDSS and TARDIS. The event at chromosome 5 that consists of a deletion and dispersed duplication was reported four and three times by GRDISS and TARDIS. These redundant predictions complicate and mislead downstream functional annotations. On the contrary, Mako was able to completely detect the above two CSV events, and also capable of revealing the breakpoint connections of CSVs encoded in the subgraphs. The above observations suggested that the subgraphs detected through pattern growth are interpretable, from which we can characterize the breakpoint connections for a given CSV event.

**Figure 5.**
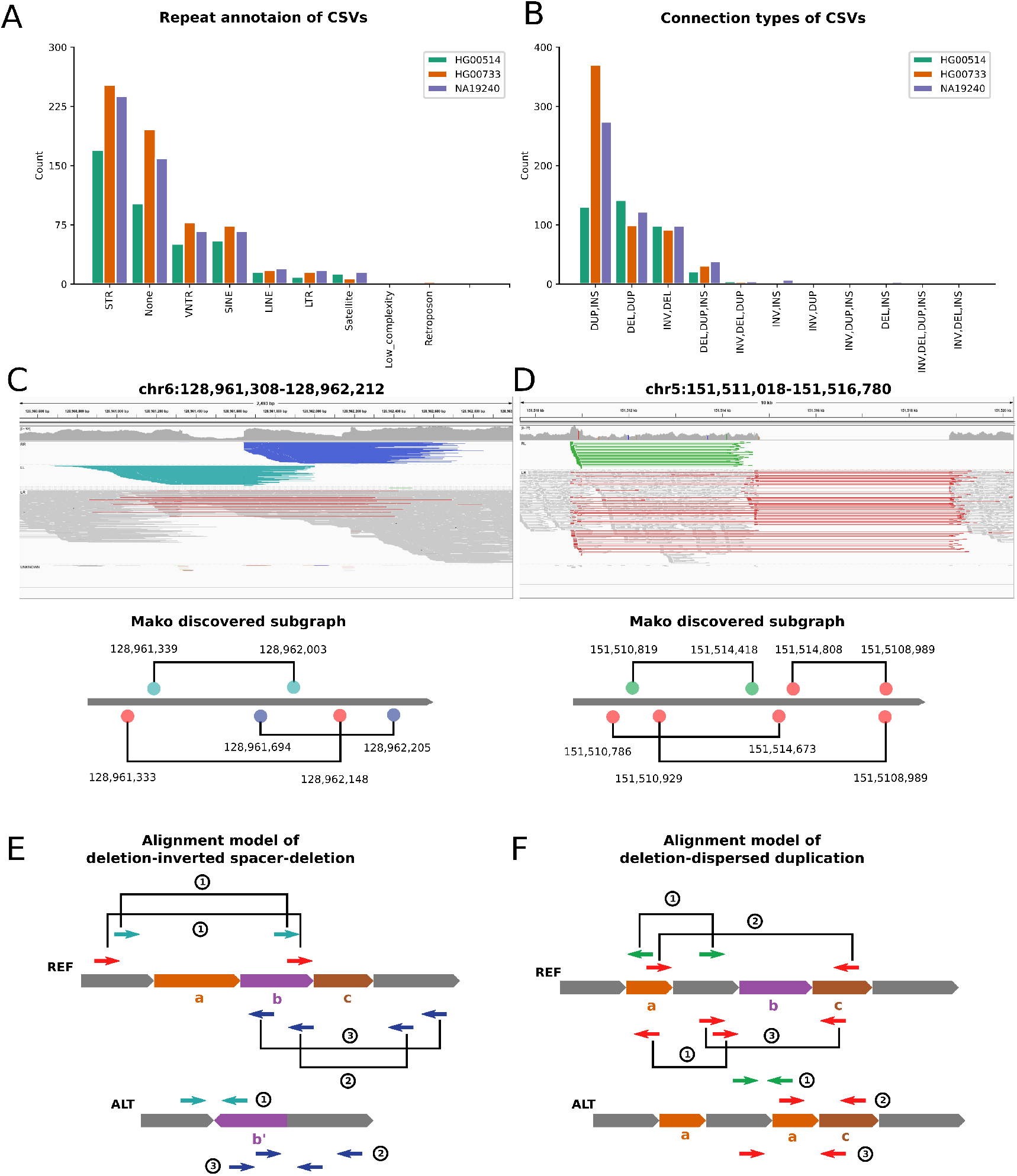
Repeat annotation and types of CSV s with two representative examples identified by Mako. (**A**) is repeat annotation and (**B**) is detected connection types of CSVs, respectively. The top panel of (**C**) and (**D**) are IGV view of the two events and the alignments are grouped by pair orientation. The dark blue shows reverse-reverse alignments, light blue is the forward-forward alignments, green is the reverse-forward alignments and the red indicates alignment of large insert size. The bottom panel of (**C**) and (**D**) are sub-graph structure discovered by Mako. The colored circles and solid lines are nodes and edges in the sub-graph. (**E**) The alignment model of deletions with inverted spacer. (**F**) The alignment model of deletion associated with dispersed duplication. In (**E**) and (**F**), short arrows are paired-end reads that span breakpoint junctions, and their alignment are shown on the reference genome with corresponding ID in circle. Noted that a single ID may have more than one corresponding abnormal alignment types on the reference.

**Table 2.**
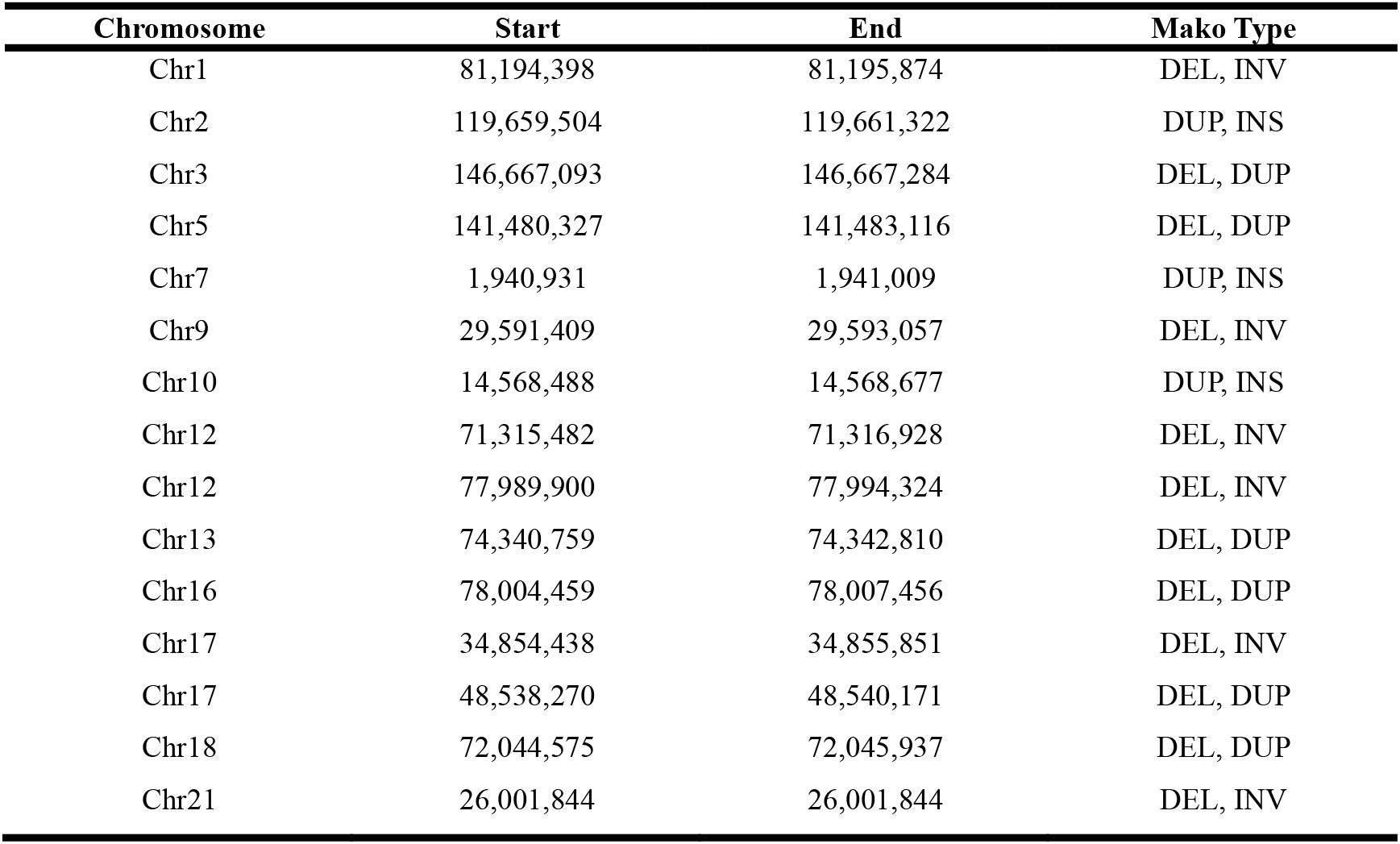
Summary of experimentally validated CSVs.

**Table 3.** Summary of experimental and computational validation as well as manual inspection for CSVs.

| Validation Strategy |  | Total | Valid | Invalid | Inconclusive |
| --- | --- | --- | --- | --- | --- |
| Experimental (PCR succeeded) |  | 21 | 15 (71%) | 6 (29%) | - |
| Computational | ONT reads |  | 256 (42%) | - | 353 (58%) |
|  | HiFi contig | 609 | 414 (68%) | 191 (32%) | - |
|  | ONT reads or HiFi contig |  | 544 (87%) | 76 (13%) | - |
| Manual | HiFi reads | 609 | 440 (72%) | 169 (28%) | - |

### Contribution of homology sequence in CSV formation

Ongoing studies have revealed that genome alterations are mainly caused by the inaccurate DNA repair and the 2-33bp long microhomology sequence at breakpoint junctions plays an important role in CSV formation [18, 31–34]. To further characterize CSVs’ internal structure and examine the impact of homology sequence on CSV formation, we manually reconstructed (**Methods**) 1,052 high-confident CSV calls given by Mako (252/403 from HG00514, 440/609 from HG00733 and 360/556 from NA19240) via PacBio HiFi reads (**Figure 6A**, **Supplementary Table S6, Supplementary Figure S5, Supplementary File 2**). The percentage of successfully reconstructed events was similar to the orthogonal validation rate, showing CSVs detected by Mako were accurate and the validation method was effective. The high-confident CSV callset contains 816 insDup events with both insertion and duplication edge connections. Further investigation revealed that these events contains irregular repeat sequence expansion, making them different from simple insertion or duplications (**Supplementary Figure S6**). Besides, we found two novel types, which were named adjacent segments swap and tandem dispersed duplication (**Figure 6B**, **Supplementary Figure S7-S8**). We inferred that homology sequence mediated inaccuracy replication was the major cause for these two types. Furthermore, we observed that 134 CSVs contains either inverted or dispersed duplications (**Supplementary Table S6**). These duplications involved CSVs were mainly caused by Microhomology Mediated Break-Induced Replication (MMBIR) according to previous studies[18, 32, 35]. It was known that different homology patterns cause distinct CSV types (**Figure 6C** and **Figure 6D**). Surprisingly, one particular pattern of homology sequence yielded multiple CSV types (**Figure 6E**). In particular situations of the three different homology patterns, DNA double strand break (DSB) occurred after replication of fragment *C*. According to the MMBIR mechanism and template switch [23, 32–34], pattern I (**Figure 6C**) and pattern II (**Figure 6D**) can only have one output but pattern III (**Figure 6E**) produces three different outcomes. The results provided additional evidence for understanding the impact of sequence contents on DNA DSB repair, leading to better understanding of diversity variants produced by CRISPR [36, 37].

**Figure 6.**
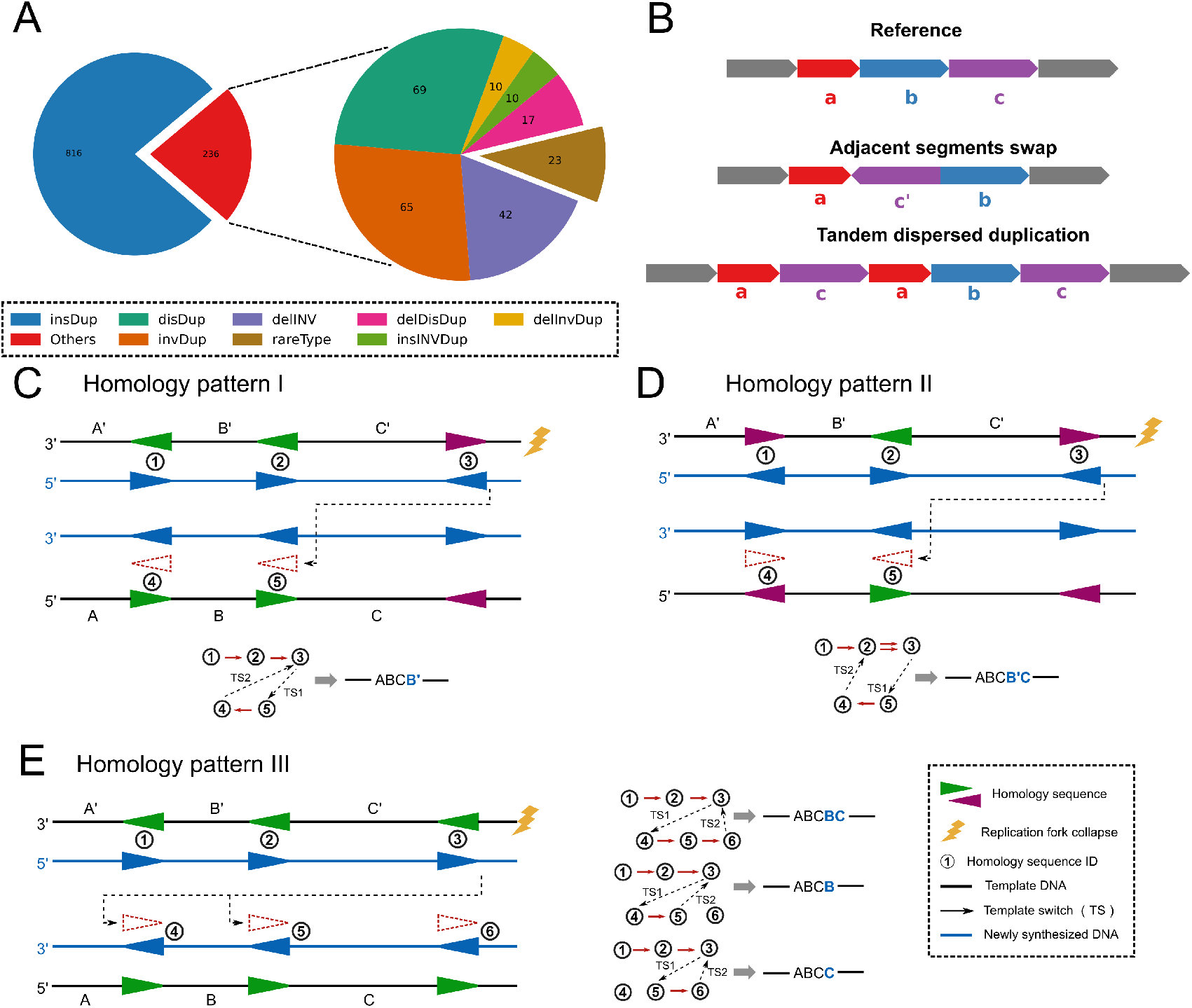
Overview of Mako’s CSV discoveries from three healthy samples and proposed CSV formation mechanisms. (**A**) Summary of discovered CSV types, these types are reconstructed by HiFi PacBio reads, where a type with less than 10 events was summarized as rareType. (**B**) Diagrams of two novel and rare CSV types discovered by Mako. In particular, Mako finds three events of adjacent segments swap and only one tandem dispersed duplication. (**C-E**) Replication diagram explains the impact of homology pattern for MMBIR produced CSVs. In these diagrams, sequence *ABC* has been replicated before the replication fork collapse (flash symbol). The single strand DNA at the DNA double strand break (DSB) starts searching for homology sequence (purple and green triangle) to repair. The above procedure is explicitly explained as a replication graph, from which, nodes are homology sequences and edges keep track of the template switch (dotted arrow lines) as well as the normal replication at different strand (red lines). If there are two red lines between two nodes, the sequence between these two nodes will be replicate twice as shown in (**D**).

## Discussion

Currently, short read sequencing is significantly reduced in cost and has been applied to clinical diagnostics and large cohort studies [16, 38, 39]. However, CSVs from short read data are not fully explored due to the methodology limitations. Though long read sequencing technologies bring us promising opportunities to characterize CSVs [13, 14, 40], their application is currently limited to small-scale projects and the methods for CSV discovery are also underdeveloped. As far as we know, NGMLR combined with Sniffles is the only pipeline that utilizes the model-match strategy to discover two specific forms of CSVs, namely deletion-inversion and inverted duplication. Therefore, there is a strong demand in the genomic community to develop effective and efficient algorithms to detect CSV using short read data. It should be noted that CSV breakpoints might come from either single haplotype or different haplotypes, where two simple SVs from different haplotypes lead to false positives (**Supplementary Figure S9**). This may substantially increase the false discoveries because Mako currently is not able to determine the exact haplotype of each breakpoints. However, Mako can be extended to differentiate such false positives by adding additional features to the graph, e.g. phased reads. Given that short read sequencing is not able to span all breakpoints of a CSV, Mako could only infer the CSV types based on the edge connections from the subgraph, while it is difficult to characterize the exact components of CSVs. Therefore, our next work will integrate both short and long reads to the signal graph for CSV discovery and characterization.

To sum up, we developed Mako, utilizing the graph-based pattern growth approach, to discover CSVs. Meanwhile, the intensive experimental and computational validations as well as manual inspections showed around 70% accuracy and 20bp median breakpoint shift. Besides the improvement of CSV detection performance, the optimized pattern growth algorithm on sequentially constrained subgraph detection is not restricted to CSV detection and can be generalized to other graph problems with similar constraints. Most importantly, to the best of our knowledge, Mako is the first algorithm that utilizes the bottom-up guided model-free strategy for SV discovery, avoiding the complicated model and match procedures. Given the fact that CSVs are largely unexplored, Mako presents opportunities to broaden our knowledge of genome evolution and disease progression.

## Materials and methods

### Materials

The short read aligned BAM files for NA19240, HG00514 and HG00733 were obtained from the HGSVC [9] (**Supplementary Note**). The PacBio HiFi reads were provided by HGSVC and we aligned these reads with pbmm2 (https://github.com/PacificBiosciences/pbmm2) and NGLMR [40] under default settings (**Supplementary Note**). The haploid assembly of HG00733 were obtained from HGSVC and aligned with pbmm2 (**Supplementary Note**). Both short reads and long reads were aligned to the human reference genome GRCh38. The coverage was approximately 70X and 30X for short and long reads, respectively (**Supplementary Note**). The simple SV callset for NA19240 is publicly available from HGSVC, and was contributed by Manta [7], Lumpy [3], Pindel [1] and etc. Alignment files and SV callset for the SK-BR-3 cell line were obtained from a recent publication [13] (**Supplementary Note**). The SK-BR-3 callset (**Supplementary Note**) was merged by SURVIVOR from contributions by Manta [7], Lumpy [3], Delly [2] and PopIns [41], and contains 627 inversions (INV), 2,776 deletions (DEL), 483 duplications (DUP) and 1,160 translocations (TRA).

### Building signal graph

To create the signal graph *G*, Mako collects mutational signals satisfying one of the following criteria from the alignment file to create the signal nodes set *V* of *G*: 1) clipped portion with minimum 10% size fraction of the overall read length; 2) split reads with high mapping quality; 3) discordant read-pairs. Notice that a discordant alignment will create two nodes correspondingly. Meanwhile, each node is represented by a cluster of mutational signals and is given three attributes type, pos and weight. Mako uses two types of signal clusters. One of the clusters is single-nucleotide resolution cluster created by clipped reads or split reads, namely Mako clusters these reads at the same location to create node. Another cluster is formed by discordant read-pairs, where the clustering distance is set as estimated average insert size minus two times read length. To avoid using randomly occurred discordant alignment clusters, we followed the procedure introduced by Chen [4]. Specifically, it assumed one type of discordant alignment at the gnomic location is uniformly distributed under the null hypothesis of no variant. For locations that have more than one type of discordant alignment, the number of such alignments at particular location forms a mixture Poisson distribution with each mixture component representing one of the discordant types. Thus, we summarize the statistics of clustering of a particular type *i* as the probability of having more than observed number of discordant alignments in a given region:

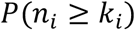

where *n_i_* denotes the Poisson random variable with mean equal to *λ_i_*, and *k_i_* is the number of observed type *i* discordant alignment. The estimation of *λ_i_* can be calculated based on the uniform assumption:

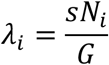

where *s* represents the cumulative size of the regions that discordant alignments anchored, *N_i_* the total number of type *i* alignment in the BAM and *G* the length of reference genome.

It should be noted that some discordant read-pairs may contain two types of signals, e.g. abnormal insert size and incorrect mapping orientation, which are clustered separately to create nodes. Moreover, split reads created nodes not only provide precise location but also complement edges for discordant read-pairs. Therefore, Mako’s performance will not be dramatically affected by the skewed insert size distribution because skewed distribution only affects estimation of abnormal insert size. The attribute weight and pos indicate the number of abnormal reads and approximate position on the genome, respectively; and type denotes the type of abnormal alignment, such as *MS*, indicating the node consists of reads clipped at the right part. Importantly, we consider nodes with the same type as identical nodes. For the edge set *E* = {*E_pe_, E_ae_*} of signal graph *G*, the paired edges from *E_pe_* are derived from read-pairs or split-reads between two signal nodes, where rp indicates the number of paired reads involved. Adjacency edges from *E_ae_* measure the distance dist between two adjacent signals. However, adjacent edges are virtual links compared with the paired edges derived from alignments, thus the pattern growth through adjacent edges is constrained by dist to avoid pointless pattern expansion. It should be noted that both types of edges might coexist between two nodes. To achieve efficient subgraph detection and avoid overlapping subgraphs, we use a linearized database to store the graph and this graph can be built efficiently in linear time by reading the input file once.

### Detecting CSVs with pattern growth

Pattern growth is an efficient heuristic approach for frequent pattern discovery in strings and graphs [42], which has been widely used in many areas [43–48], such as INDEL detection in DNA sequences [1, 24]. Compared with statistical methods, pattern growth discloses the intrinsic features of the data without sophisticated model generation. Meanwhile, the output of the pattern growth approach is usually interpretable, which is very important for specific applications [49].

In the CSV detection, the subgraph pattern starts at a single node and grows by adding more nodes until it cannot find a proper node (**Algorithm I, Supplementary Figure S10**). In addition, to avoid overlapping subgraphs, we only allow the subgraph to grow according to the increasing order of pos value for each node. Meanwhile, backtracking is only allowed for nodes involved in the current subgraph. For example (**Figure 2**), Mako detects the maximal subgraph by visiting nodes *A*, *C*, *B*, and *D*, respectively. Since the edge distances between *A* and *B* as well as D and E is larger than the distance (minDist) threshold, Mako grows the subgraph through *C* and backtrack node *B* to expand the subgraph, whereas edge between D and E is constrained.

Given the fact that the signal graph contains millions of nodes at whole genome scale, we use a strategy similar to “seed-and-extension” that has been utilized by sequence alignment algorithms [50, 51] to accelerate the subgraph detection process. Meanwhile, we only keep the index of each node in the database to save memory for subgraph detection (**Supplementary Figure S11**). Moreover, as we assigned attributes to each node, the discovered subgraphs not only differ in edge connections but also in the type of signal nodes in the subgraph. Therefore, we propose an algorithm that starts at multiple signal nodes of the same type and extends locally for efficient subgraph detection (**Algorithm II**). It should be noted that sequence alignment usually results in one best alignment [50, 51], whereas our algorithm is also encouraged to discover multiple maximal subgraphs that share the same edge connections but different node attributes. To avoid missing subgraphs or incomplete detection, minFreq =1 is a default parameter for subgraph detection, but this could also be time consuming and affected by graph noise. Thus, Mako allows users to set larger minFreq to avoid random subgraphs and detects the connected components of subgraphs to ensure complete detection. In particular, a larger minFreq value allows multiple identical subgraphs to be discovered, and edges between these subgraphs are kept and used to build connections between subgraphs. These edges can be reliably marked, because the frequency of the current subgraph becomes smaller than the minFreq value by adding those edges. Then, a local maximal subgraph represented by a connected component can be discovered from the subgraph connection graph. A significant feature of discovering CSVs from a graph is that it provides the connections between multiple breakpoints of a CSV, so that the attributes of the discovered subgraph can be directly used as a measure for CSV. Namely, if the subgraph contains more non-identical nodes and *E_pe_* edges, this subgraph is more likely to indicate a complex event. Therefore, Mako defines the boundary of CSVs using the leftmost and rightmost pos value of the nodes involved, and utilizes the number of identical node types multiplied by the number of *E_pe_* edges as a complexity score CXS (default=2). For example (**Figure 2**), the discovered CSV subgraph has a CXS score of 8, because of four identical nodes and two paired edges.

**Algorithm I:**
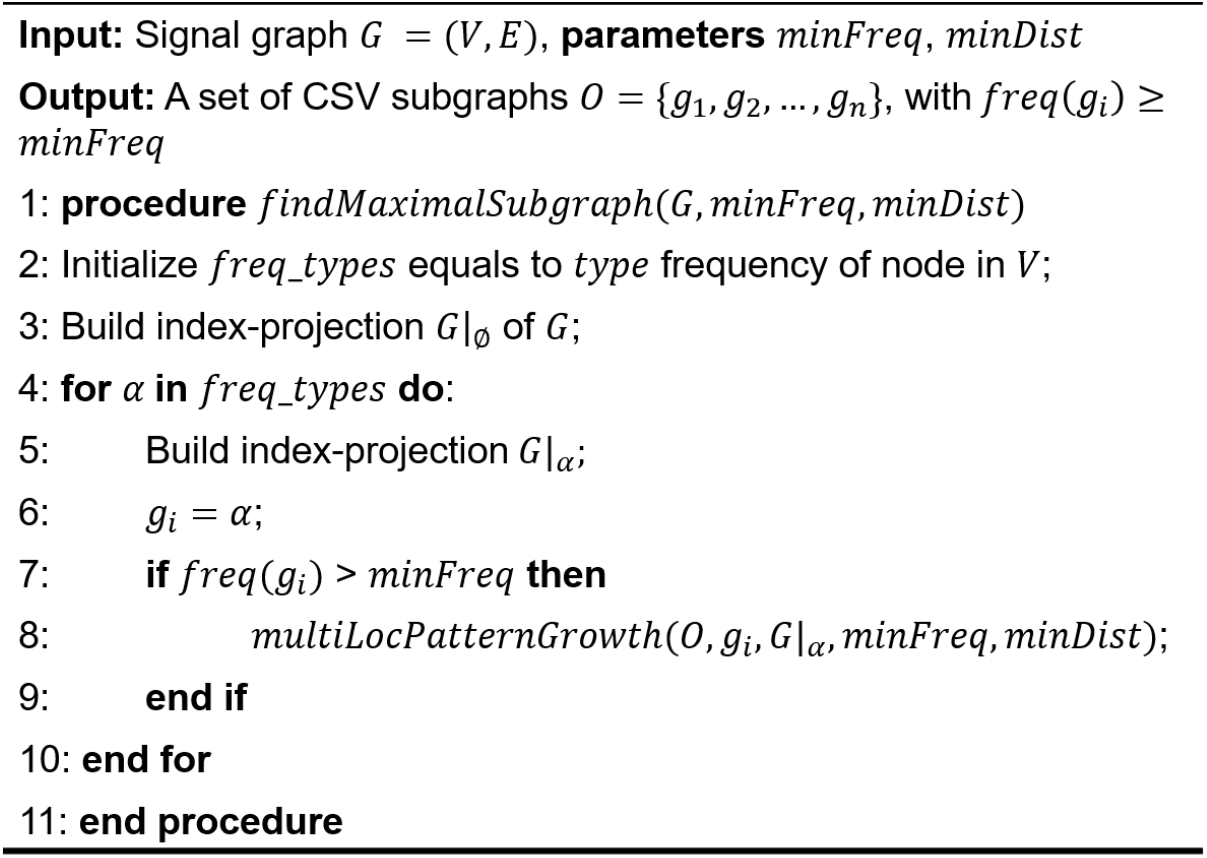
Detect maximal subgraphs

**Algorithm II:**
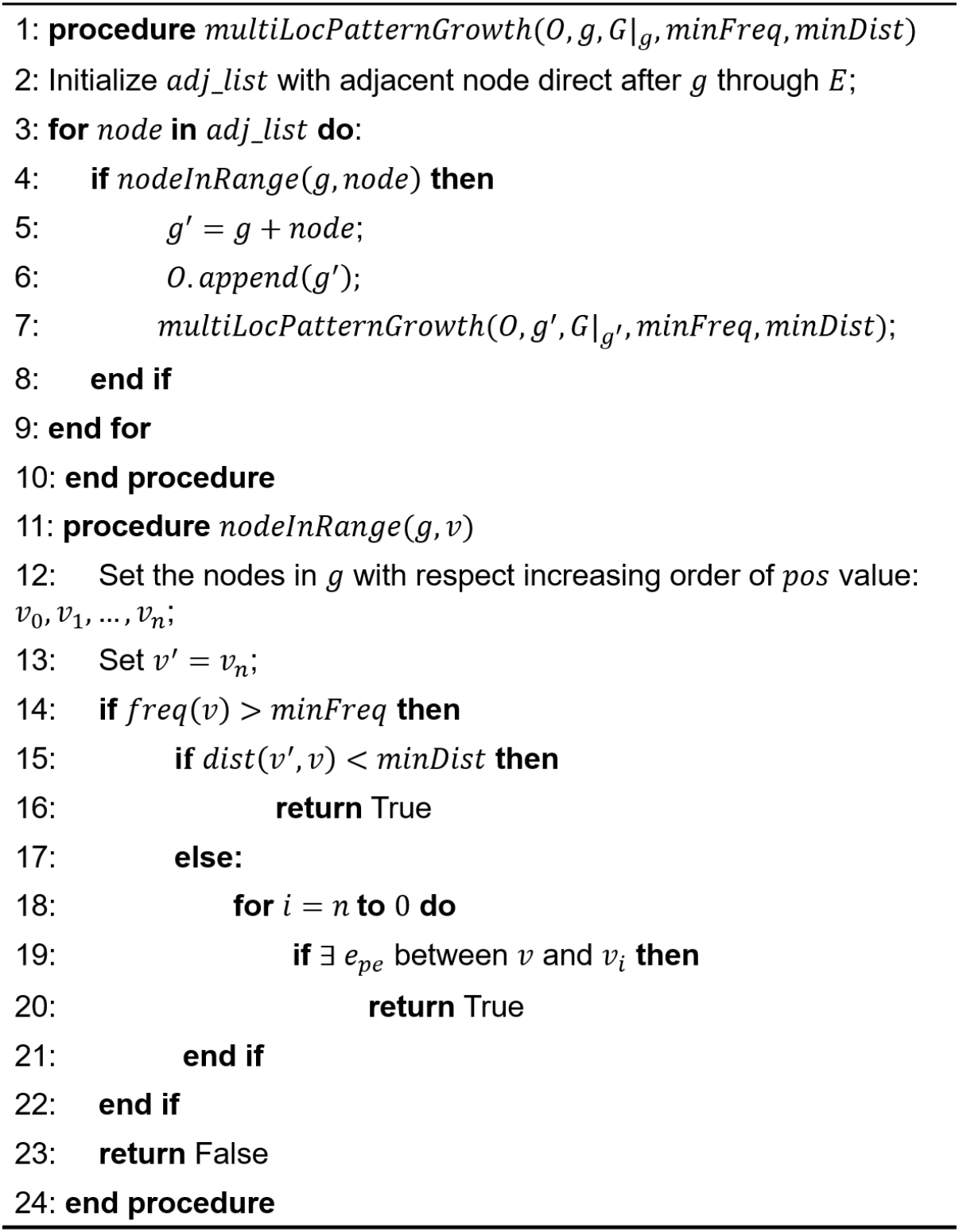
Multi-location subgraph growth

### Design of simulation studies

To create CSVs, we follow the simulation strategy introduced by the Sniffles[40]. In general, simple SVs generated by VISOR[52] are randomly selected and combined to make CSVs (**Supplementary Figure S12**). In this study, we first create deletion, inversion, inverted tandem duplication, tandem duplication and translocation copy-paste with 5000bp average size and 500bp standard deviation (**Supplementary Note**). We only consider focal translocations, where the distance between source sequence and insert position is smaller than 100Kbp. These events are created using reference genome GRCh38 and collected as basic operations for further random combination usage. For example, suppose segments on the reference genome are ABCDE and the following criteria are considered for CSV simulation:

1. The deletion (C) associated with inversion (D’) ABD’DE can be generated by first creating a deletion event and adding the inversion to a flanking region of the deletion.
2. The dispersed duplication and inverted duplication are produced through translocation copy-paste, and the orientations at the paste position distinguish these two types of duplication. For example, if we copy-paste segment B and insert it after D, a dispersed duplication ABCDBE will be created.
3. Additionally, to create translocation copy-paste involved CSVs, we only manipulate segments adjacent to the insert position of the source segment. For instance, a deletion can be associated with the dispersed duplication ABCDBE by removing D or E, leading to ABCBE or ABCDB.

To produce homozygous or heterozygous CSVs, we use the purity parameter introduced by VISOR to control the ratio of reads sequenced from variation genome and reference genome. After the variation genome is created, VISOR used wgsim (https://github.com/lh3/wgsim) to simulate paired-end reads and applied BWA-MEM [51] to align the simulated reads to the reference genome (**Supplementary Note**). Overall, VISOR has efficient functions for creating basic operations, building variation genome with simulated CSVs, simulating reads and alignment. We add the random selection and combination step as part of VISOR.

We first evaluate whether Mako is able to capture reported CSV types published by previous studies [8, 17], such as deletion flanked by inversion, inverted duplication, dispersed duplication and etc. This was termed as reported CSV. For the reported CSV, we only randomly select and combine deletion, inversion, inverted tandem duplication and tandem duplication, but leave translocation copy-paste unchanged (**Supplementary Note**). In total, we simulated 300 reported CSV types on chromosome 1. The reported CSVs usually have four to six breakpoints, which are still feasible to be detected by model-based methods. However, we emphasize that limited knowledge of CSV variety and the complex mutational signals produced by breakpoint connections are the major challenges for CSV discovery. From this perspective, we made another set of randomly simulated CSV types on autosomes, termed as randomized CSV, where we created 4,500 CSVs with 4~10 breakpoints through random combinations of at least two basic operations including translocation copy-paste (**Supplementary Note**).

### Creating CSV benchmark from real data

It has been recognized that the most significant feature of CSVs is simultaneous appearance of multiple breakpoints[8, 12, 27, 53, 54]. However, the development of robust tools for screening complex events is a difficult and unsolved problem because there are currently no well-defined rules for constraining the expected breakpoint patterns[12]. In order to study CSVs, researchers follow four major steps[12, 20] to resolve CSVs from an enormous number of simple SVs: 1) breakpoint clustering; 2) clustered breakpoints enrichment test; 3) contig assembly and realignment; 4) manually inspection from visualization. Fortunately, PacBio reads provide us with the direct evidence to validate and categorize CSVs, which can be used to screen each simple SV site for CSVs. But to avoid the intensive manually investigation of each simple SV, we first cluster simple SVs and only inspect clusters with at least two SVs. In particular, we treat each SV as an interval and apply the hierarchical clustering to find interval clusters. The distance measure for clustering is defined as follows:

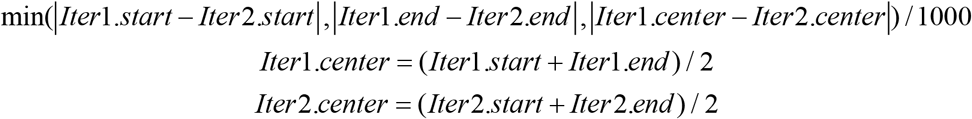

where Iter is an SV breakpoint interval, and Iter. start, Iter. end and Iter. center indicate the start, end and center of the interval, respectively. We then use the average method to calculate distance between intervals in two clusters *u* and *υ*, which is assigned by:

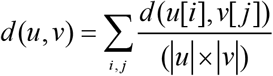

To select a proper threshold for merging clusters from the hierarchical clustering results, we use the threshold from a set of values that could produce the most clusters for each chromosome independently (**Supplementary Table S7**, **Supplementary Note**, **Supplementary Figure S13-S16**).

We further utilize the sequence dot-plot to resolve CSVs based on PacBio long reads. Sequence dot-plot is a classic way to investigate genome rearrangement between species or chromosomes[55]. It applies a k-mer match approach between sequences and keeps matches in a similarity matrix. Thus, we can define the breakpoints and type of a CSV by visualizing the similarity matrix. We use the publicly available interactive sequence dot-plot tool Gepard[56] for this process. Since CSVs are rare and might appear at the minor allele, we create a dot-plot for each long read that spans the corresponding SV cluster. Afterwards, we manually inspect all these dot-plots to identify CSVs, and their breakpoints can be easily obtained from Gepard’s interactive user interface (**Supplementary Figure S17**).

### Parameter selection for Mako and other methods

Mako run with minAf = 0.2, minFreq = 1, minWeight = 10 for real data (NA19240, HG00514, HG00733) and all simulated data (**Supplementary Note**). The minFreq was set to 1 to detect rare events. The minDist is set four times the estimated library fragment average size. And these values are all default settings for Mako. For the cancer cell line (SKBR3), considering the coverage and highly rearranged nature compared with the normal genome, we reduce the cutoff from 0.2 to 0.1 and 10 to 5 for minAf and minWeight, respectively, so that the graph could involve more nodes. Signal nodes satisfying either the minAf or minWeight threshold will be included to create the graph. The other selected tools are run under default settings for both simulated and real data (**Supplementary Note**). We use the latest version of TARDIS [26] and the SVelter callset for NA19240 is provided by HGSVC [9] (**Supplementary Note**). For the CSV detection evaluation, all predictions larger than 50p are involved and additional filtering has been done according to the recommended procedures [26–28]. In particular, GRIDSS’s callset is filtered by a filter field in VCF header such as ASSEMBLY_TOO_FEW_READ and SVs with coordinates like [57] and [p2, p1] are kept only once. The prediction of SVelter is filtered by a validation score of −1 (**Supplementary Note**).

### Performance evaluation

Typically, a correct discovery is defined as a best match between benchmark and predictions, and thus the closest event to the benchmark CSVs with similar size is considered as true positive [58]. However, performance comparison of CSVs is less straightforward than that of simple SVs because of multiple breakpoints involved [27]. To address the demand of detecting CSVs as a single event and avoiding redundant predictions [12], the performance is evaluated from two aspects. For example, a CSV with inversion flanked by two deletions is evaluated as three components. Correct prediction of all breakpoints for the three components is considered as all-breakpoint match. Meanwhile, if only one prediction is close to the leftmost and rightmost breakpoints of the CSV with similar size, this prediction is treated as unique-interval match. In the evaluation, the closeness bpDist and size similarity sim between prediction and benchmark are 500bp and 0.7. For example, assume a benchmark [b. start, b. end, b. size], and a prediction [p. start, p. end, p. size]; then a correct prediction will satisfy the following equations:

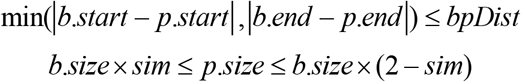

For simulated data, true positive (TP) is defined as the nearest prediction with similar size to the benchmark, while predictions not in the benchmark are treated as false positives (FP). False negatives (FN) are events in the benchmark set that are not matched by predictions (**Supplementary Note**). Then, the usual measurements can be calculated as follows:

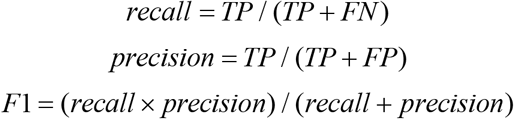

Since it is usually hard to measure the false positives of each tool for real data, we only consider the number of correct discoveries. To fully characterize Mako’s performance, we further evaluate it on NA19240 based on PacBio reads by using sensitivity and specificity (**Supplementary Note**) Additionally, because the breakpoints are not as precise as that in the simulation, we relax the size similarity threshold sim to 0.5 for real data sensitivity evaluation. To examine Mako’s CSV breakpoint offset, we first manually labeled the breakpoints of each CSV from HG00514, HG00733 and NA19240 based on PacBio reads create sequence Doplot (**Supplementary Note**). Secondly, we compare manually labeled breakpoints to Mako reported ones to calculate the offset.

### Orthogonal validation of Mako detected CSVs

To evaluate detected CSVs, we used experimental and computational validation as well as manual inspections of HG00733. The raw CSV calls from HG00733 was obtained by selecting events with more than one link types observed in the subgraph, resulting in 609 CSVs. To design primers, Primer3 (https://github.com/primer3-org/primer3) was used in conjunction with internal software to design and select PCR primers, where the optimal primer size was set to 23bp. In particular, we extend Mako detected breakpoints by 500bp to select primers with average GC contents close to 50% and a predicted melting temperature 60 °C. Primers were then selected within the extended distance but 200bp outside of the boundaries of the breakpoints defined by Mako (**Supplementary Figure S18**). If duplication and inversion like edges were found in the subgraph, primers were also designed on the reverse complementary strand. All primer pairs were tested for their uniqueness across the human genome using In Silico PCR from UCSC Genome Browser. BLAT (https://users.soe.ucsc.edu/~kent/) search was also performed at the same time to make sure all primer candidates have only one hit in the human genome. If the above procedure does not result in a valid primer pair, the size of the regions for which primers are designed was increased from 500bp to 650bp and all process were repeated to search for primers (primers are in **Supplementary Table S3**). PCR amplifications were performed in a volume of 25 ul concentration of reagents, consisting of 1) 1x of 10x *Ex Taq* Buffer (Mg^2+^ Plus); 2) 0.4 mM of dNTP mix, 0.4 uM for each primer; 3) 0.75 units of Ex Taq DNA polymerase (TakaRa, Japan) and 4) 30 ng of DNA. The amplification cycle was performed in Mastercycler^®^ nexus gradient (Eppendorf, Germany), including 1) 5 minutes’ predenaturation at 94°C; 2) 35 cycles of denaturation at 94°C for 45 seconds, annealing 45 seconds according to different TM value of each primer and elongation at 72°C for 90 seconds; 3) followed by 10 minutes’ extension at 72°C. The amplification products were separated by electrophoresis in 1.5% agarose gels with CellPro™ DNA-Red (InCellGene LLC, USA) and bands were visualized under the UV light. Then, we selected products with the expected product size and bright electrophoretic bands (**Supplementary Figure S19**, all results in **Supplementary File 3)**, which were further purified and cloned into the expression vector pEASY-T1 (Transgene, China). The positive clones containing the targeted fragments were send to TsingKe Biological Technology Company for Sanger sequencing. The Sanger sequencing data were aligned against the reference allele of the CSV site and visualized with Gepard for breakpoint inspection.

We used HiFi reads from HGSVC to manually reconstruct each CSVs. Similar to the procedure of creating the benchmark CSV for NA19240 and SKBR3, SAMtools was used to get the HiFi reads spanning the breakpoints. Afterwards, Gepard was applied to create the sequence dotplot between each read and the reference genome. We than go through all the sequence dotpot to validate CSVs detected by Mako (**Supplementary Figure S17, Supplementary Note, Supplementary File 2**). The validation rate measured whether Mako detected subgraphs contained different types of breakpoint connection edges. For dotplots with ‘messy’ regions, they could produce duplication and insertion like breakpoint connections based on short-read sequencing. Therefore, it was difficult or even impossible for short reads to distinguish between distinct complex events and those detected at repeat regions. To characterize these events based on long-read sequencing, we introduced a three steps workflow as follows:

Step 1. Identifying event breakpoints inside the ‘messy’ regions in the dotplots. Those outside the ‘messy’ regions were considered as distinct complex events.
Step 2. We defined 3 dotplot patterns (**Supplementary Figure S20**) to classify ‘messy’ events to CSVs, where the x-axis and y-axis are REF and ALT sequence, respectively. Region 1, 2 and 3 indicates regions where extra segments could be found. Especially, region 2 in each case indicates the ‘messy’ region caused by repeats.

○ Case A: Blue segments indicate an insertion event with single breakpoint on the reference. A CSV should contain at least one duplicated segments (purple) in region 1, 2 or 3. Example events include chr1:206,924,211-206,924,525 (**Supplementary File 2**, **page 89**) and etc.
○ Case B: Blue segments indicate repeat expansion on the ALT sequence. A CSV should contain extra segments in region 1, 2 or 3. Example events include chr1:1,382,295-1,382,470 (**Supplementary File 2**, **page 145**) and etc.
○ Case C: Blue segments overlap on the REF, but have a gap on the ALT sequence. This type of events could be interpreted as insertion with duplications, which is considered as complex event. We also observed some CSV contained segments (purple) in region 1, 2 or 3. Example events include chr3:50,311,835-50,312,092 (**Supplementary File 2**, **page 226**) and etc.
Step 3. We further investigate events that failed the examination in Step 2 according to 2 dotplot patterns (**Supplementary Figure S21**).

○ Case A: A simple insertion event, where the breakpoint locates inside tandem repeats (region 2) and other segments cannot be found.
○ Case B: Regular repeat expansion (purple segments) in ALT sequence.

For computational validation, we obtained ONT reads of HG00733 from HGSVC and applied VaPoR [59], an independent structural variants validation method, to validate these CSVs (**Supplementary Note**). VaPoR is able to validate calls based predicted region and types with a confidence score. VaPoR labeled NAs and 0 to some of the inconclusive events due to highly repetitive sequence and unclear recurrent pattern that can be observed (**Supplementary Figure S22**). We termed the above procedure as ONT validation. Besides, we obtained HiFi assemblies from HGSVC and applied a K-mer based breakpoint examination and calculate the breakpoint shifts. Specifically, CSV spanning H1 or H2 contig sequence (ALT) and reference (REF) sequence were extracted from alignment and GRCh38, respectively. We first identified the matched segments between ATL and REF through K-mer (k=32bp) realignment as well as sorted these segments according to their position on reference. Afterwards, we marked the unmatched or gap regions, from which, we calculated the breakpoints and size similarity. A CSV was considered valid if both left and right breakpoint difference are smaller than 500bp. This constrain was used by Truvari (https://github.com/spiralgenetics/truvari/), a standard benchmarking tool used by Genome In A Bottle (GIAB). The implementation of K-mer validation is available at Mako GitHub site. Breakpoint comparison of experimental and K-mer validation were listed in **Supplementary Table S8**, which was used to calculate the breakpoint resolution. Because VaPoR is able to report Valid, NA and 0 events but not to report the breakpoint based on ONT (**Supplementary Table S9**), we did not include VaPoR’s results in the breakpoint shift analysis.

## Code availability

Mako is implemented in Java 1.8, and it is available at https://github.com/jiadong324/Mako. It is free for non-commercial use by academic, government, andnon-profit/not-for-profit institutions. A commercial version of the software is available and licensed through Xi’an Jiao-tong University. All scripts used in this study are also included in the Github repository, and a detailed description of using these scripts and other tools is provided in **Supplementary Note**.

## Data availability

All materials or datasets used in this study are publicly available and their links are listed in **Supplementary Note**.

## Authors’ contributions

In particular, KY conceived and designed the study; JL, XY and WK developed the graph-based pattern growth algorithm for SV breakpoint discovery; JL and TX created the CSV benchmarks for real data and manually reconstructed CSVs. YJ, CZ, QH and MR performed the wet lab experimental validation; SW performed the computational validation; JL, XY, CZ, LG, WK and KY wrote the manuscript; EE, SD, CL provides the ONT reads and HiFi reads. HGSVC produced the HiFi assembly and all authors contributed to the critical revision of the manuscript and approved the final version.

## Acknowledgments

This study was supported by the National Key R&D Program of China (grand NO. 2018YFC0910400, 2017YFC0907500), the National Science and Technology Major Project of China (grand NO. 2018ZX10302205), the National Science Foundation of China (grand NO. 31671372, 61702406 and 31701739) and the “World-Class Universities and the Characteristic Development Guidance Funds for the Central Universities”. Supported by Shanghai Municipal Science and Technology Major Project (Grant No.2017SHZDZX01).

## Supplementary material

**Supplementary Note** contains supplementary information for MATERIALS and METHODS.

**Supplementary Figures** contains the supplementary figures for this study.

**Supplementary Table S1** provides the benchmark CSVs, SV clustering summary and examples used to illustrate Mako CSV subgraph.

**Supplementary Table S2** provides Mako detected CSVs for HG00733, HG00514 and NA19240.

**Supplementary Table S3** provides events with successfully designed primers.

**Supplementary Table S4** provides the summary of experimental and computational validation as well as manual inspections of HG00733.

**Supplementary Table S5** provides the details of breakpoints for the two examples in Figure 5C to 5F.

**Supplementary Table S6** provides the results of manual inspections of HG00733, HG00514 and NA19240 based on PacBio HiFi reads.

**Supplementary Table S7** provides parameters used for creating the CSV benchmarks for NA19240 and SKBR3.

**Supplementary Table S8** provides experimental and computational evaluated breakpoints, which was used for breakpoint shift analysis.

**Supplementary Table S9** provides the details of VaPoR results of HG00733.

**Supplementary File 1** provides the IGV view and PacBio reads dotplot of each benchmark CSVs.

**Supplementary File 2** provides the PacBio HiFi reads dotplots for manual inspections of HG00733.

**Supplementary File 3** provides the PCR results and visualization of CSV breakpoint validated through Sanger sequencing.

