## Supplementary Figures for "Mako: a graph-based pattern growth approach to detect complex structural variants"

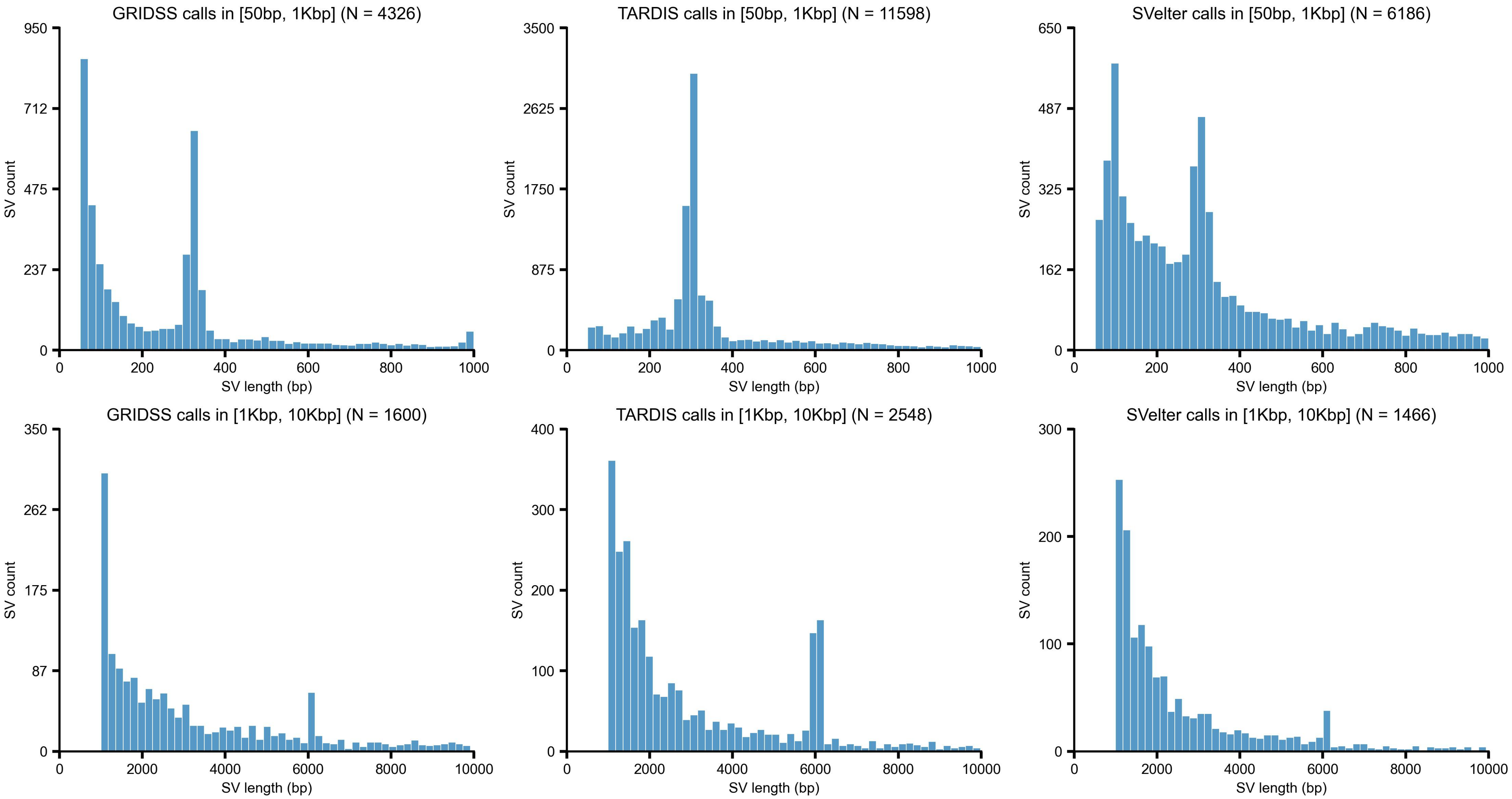


**Supplementary Figure S1** Size distribution of SV in range [50bp, 10Kbp] from NA19240

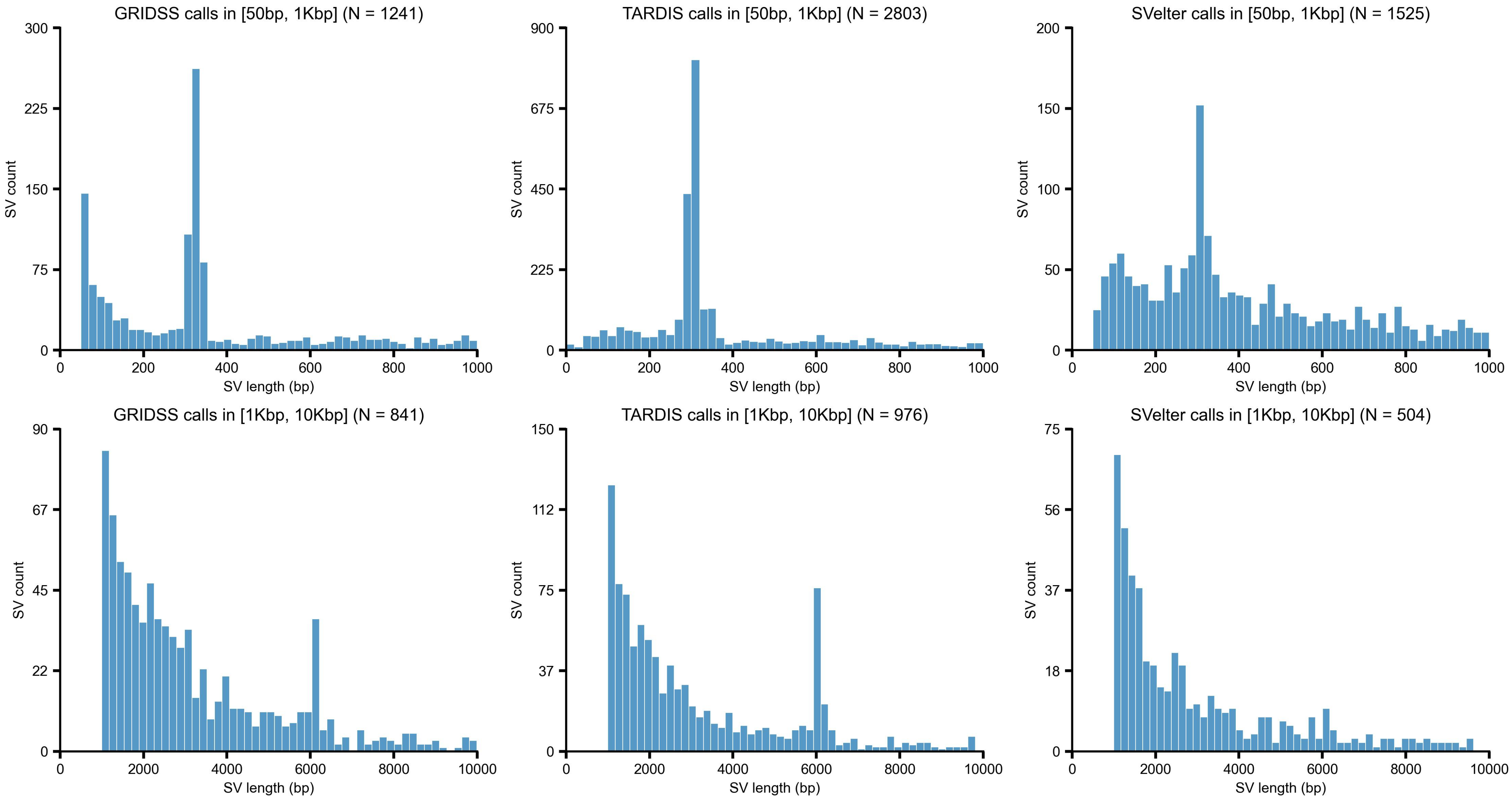


**Supplementary Figure S2** Size distribution of SV in range [50bp, 10Kbp] from SKBR3 breast cancer cell line

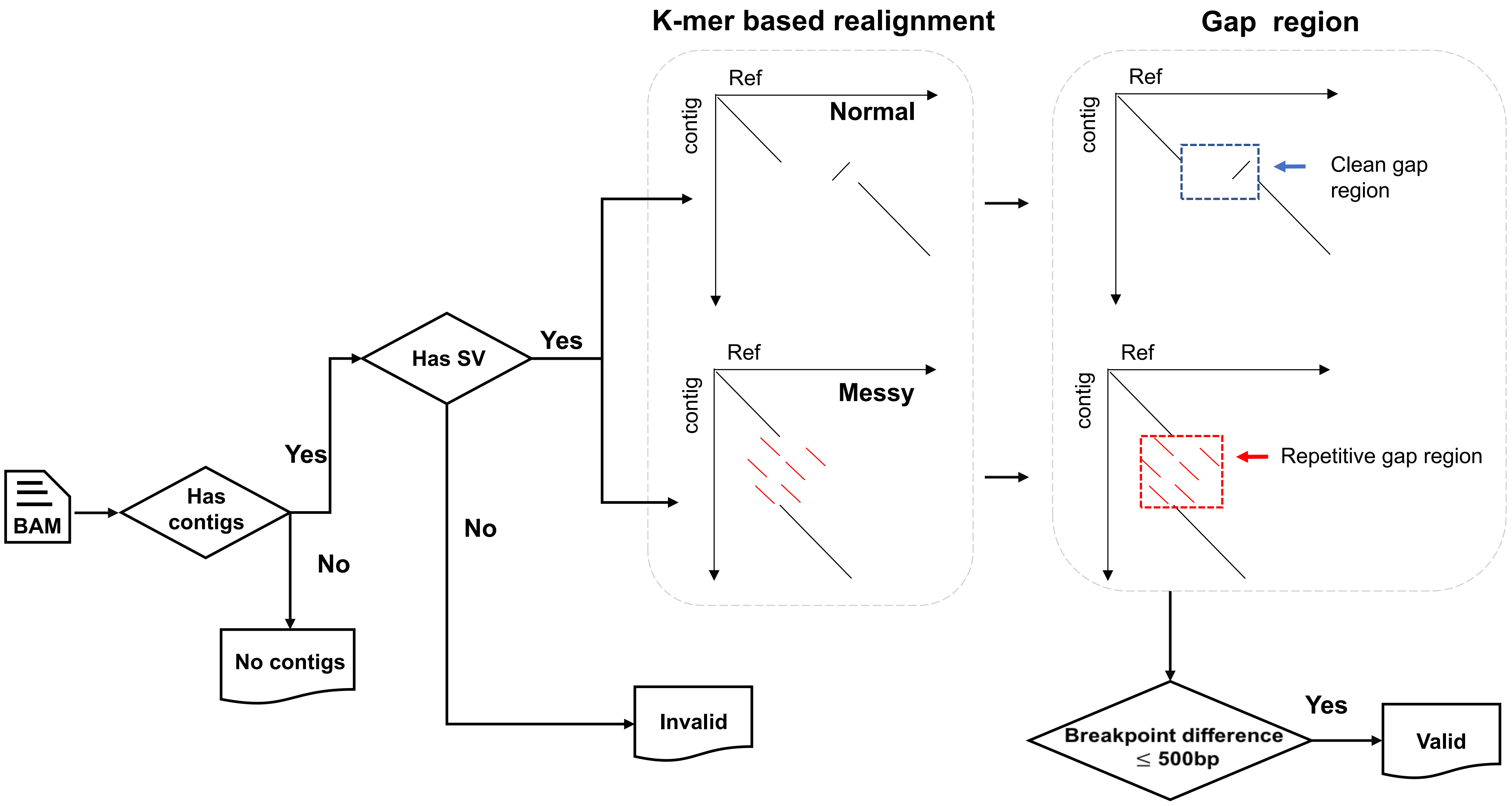


**Supplementary Figure S3** Workflow of HiFi assembly K-mer validation
For dotplots in the workflow, y-axis indicates sequence from HiFi contig and x-axis is the corresponding reference sequence.

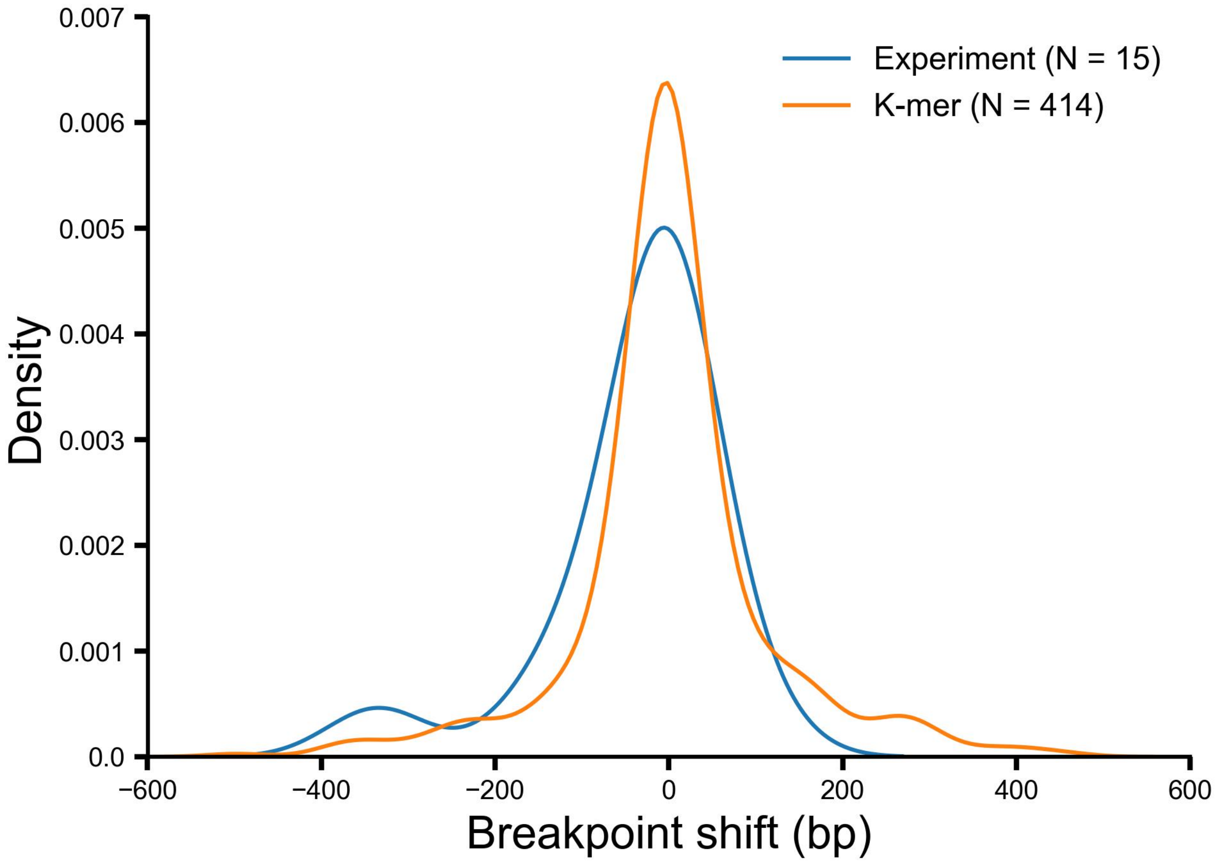


**Supplementary Figure S4** Mako detected CSV breakpoint resolution compared to HiFi contig (K-mer) and experiment

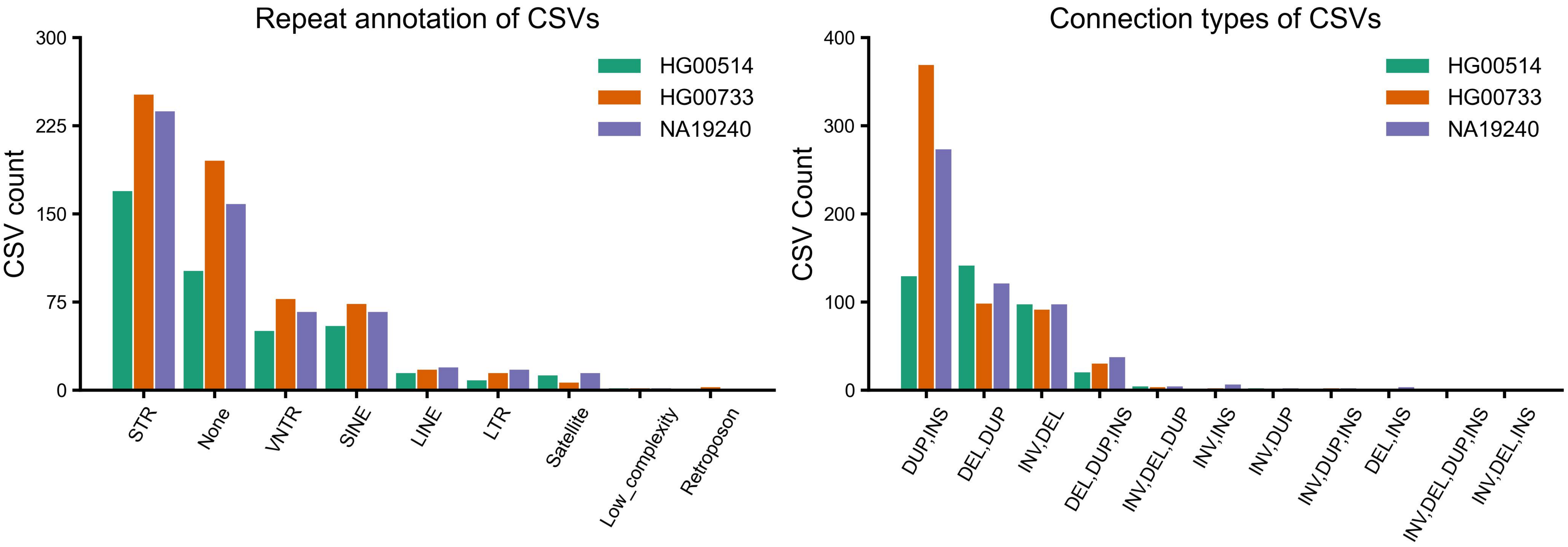


**Supplementary Figure S5** Repeat annotation and connection types of Mako detected CSVs from three samples

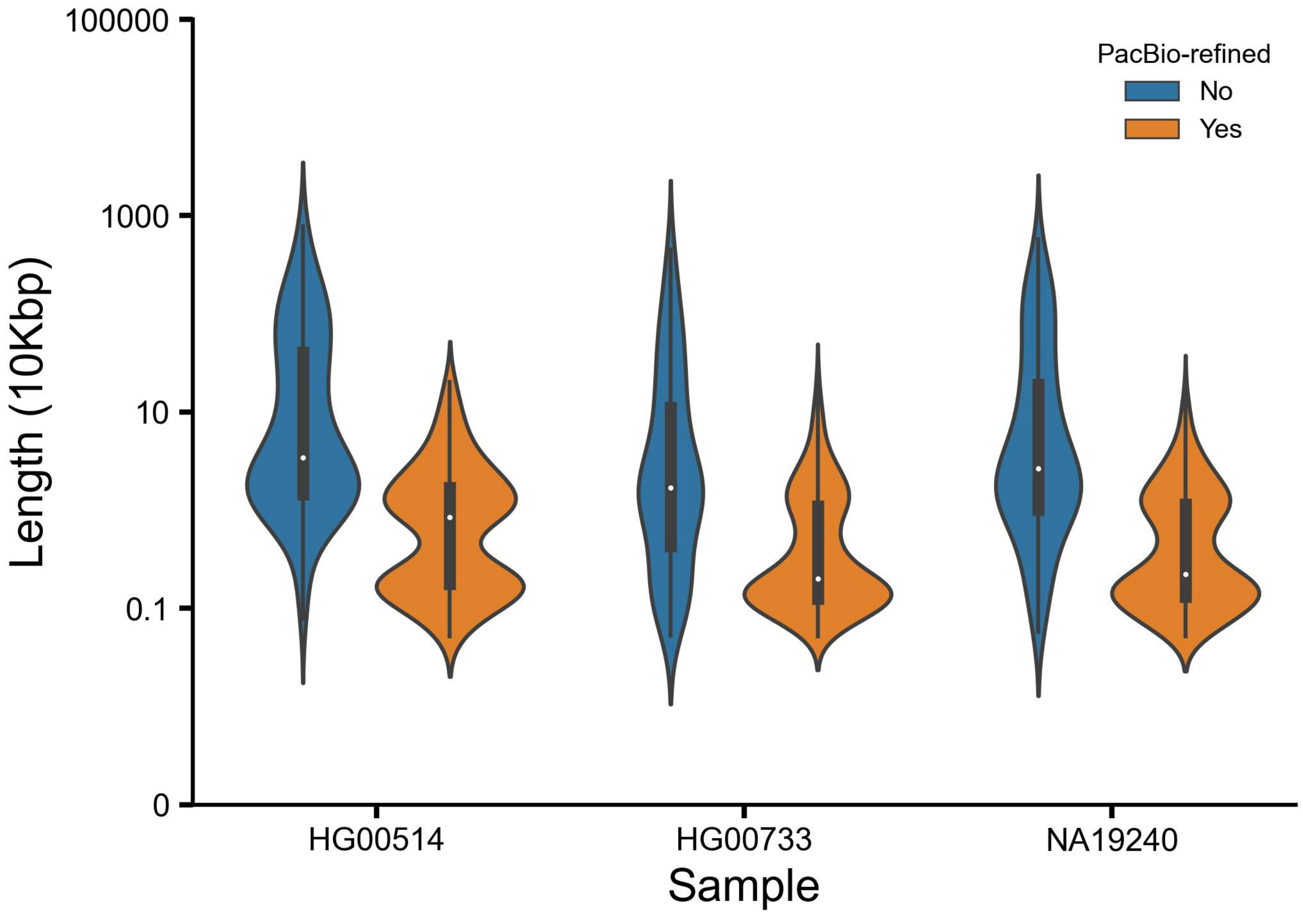


**Supplementary Figure S6** Mako detected CSV and PacBio HiFi read refined CSV size distribution

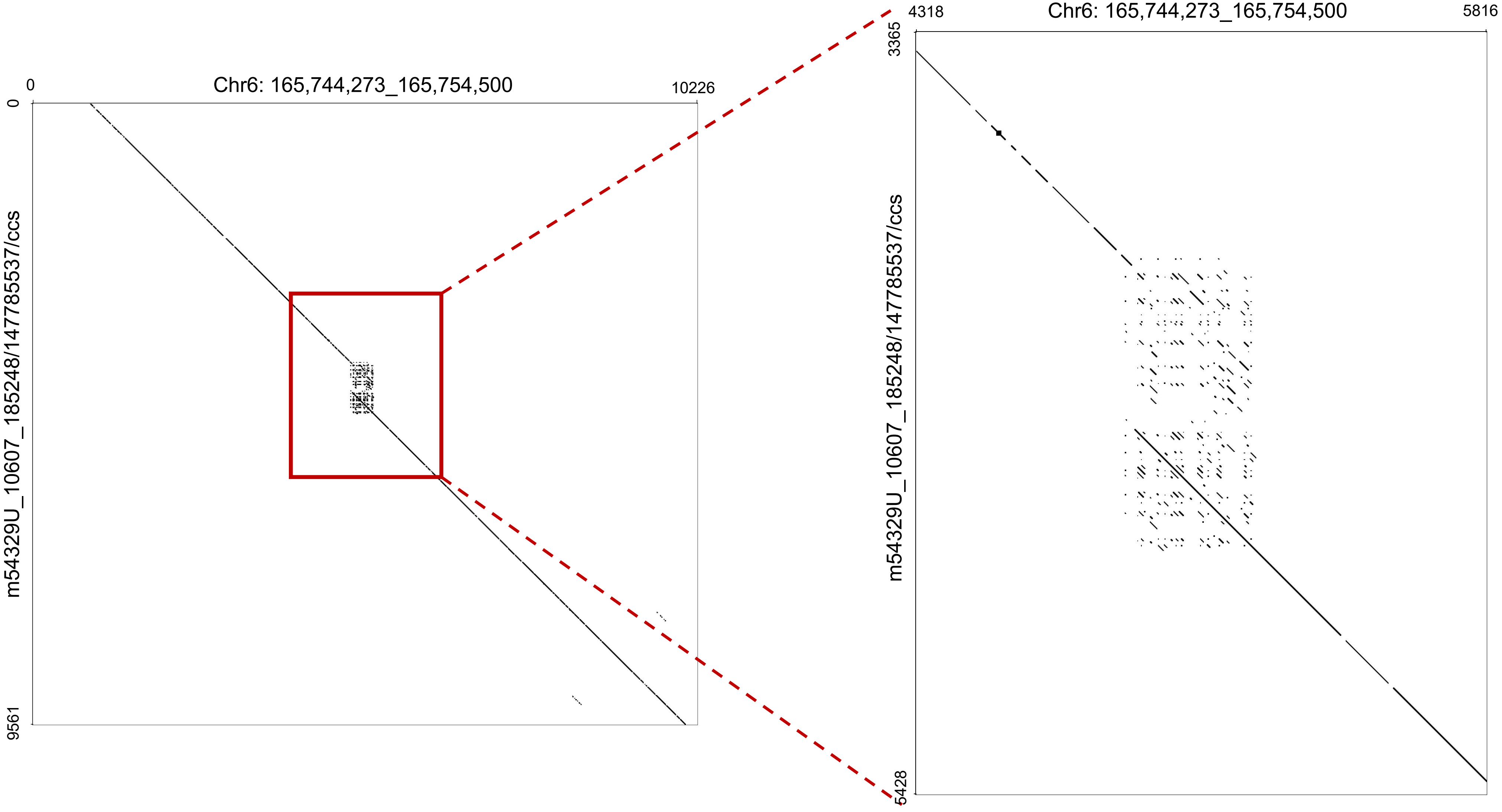


**Supplementary Figure S7** Example of an insertion associated with duplication event (InsDup) at Chr6: 165,749,273_165,749,500

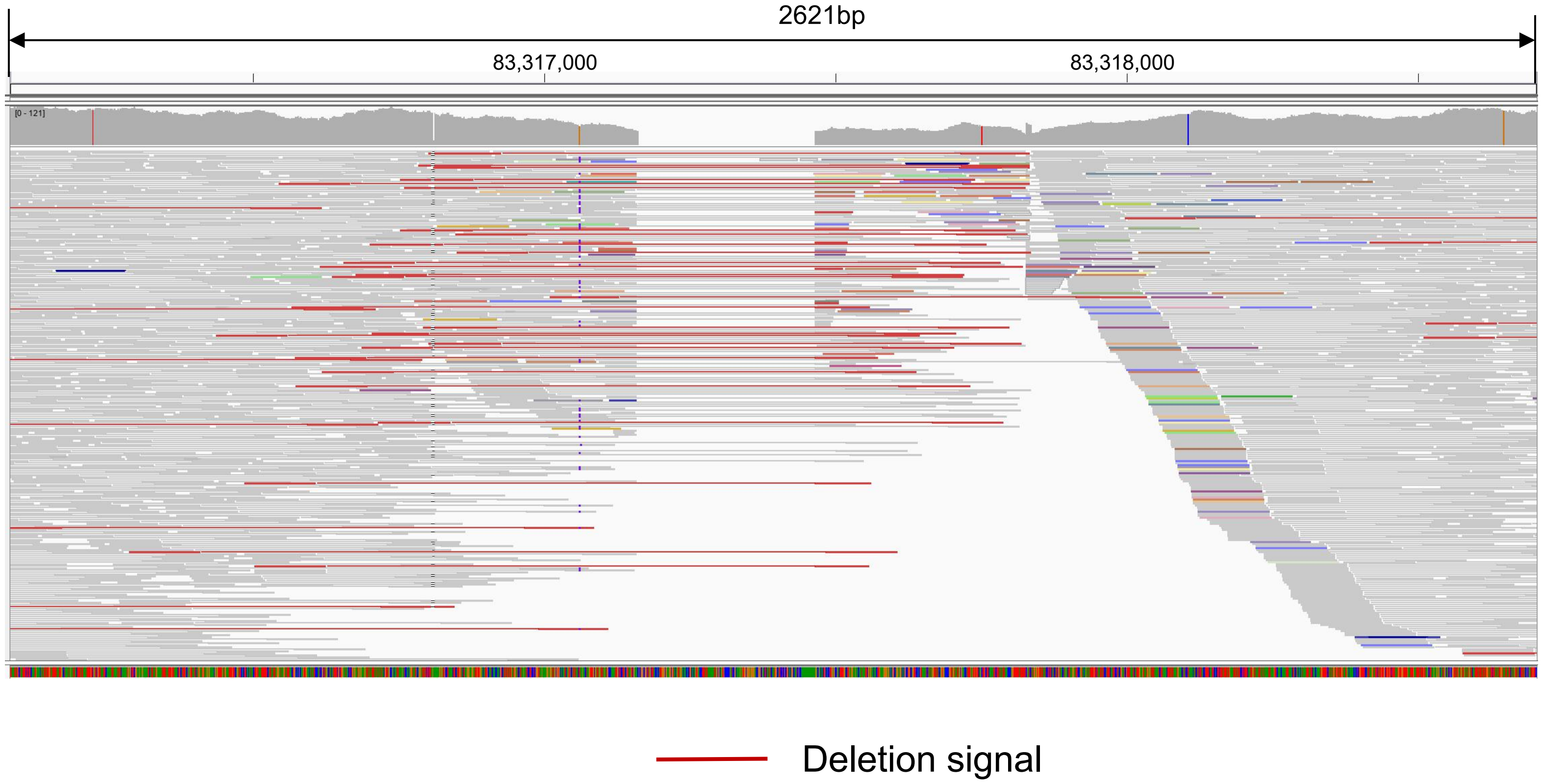


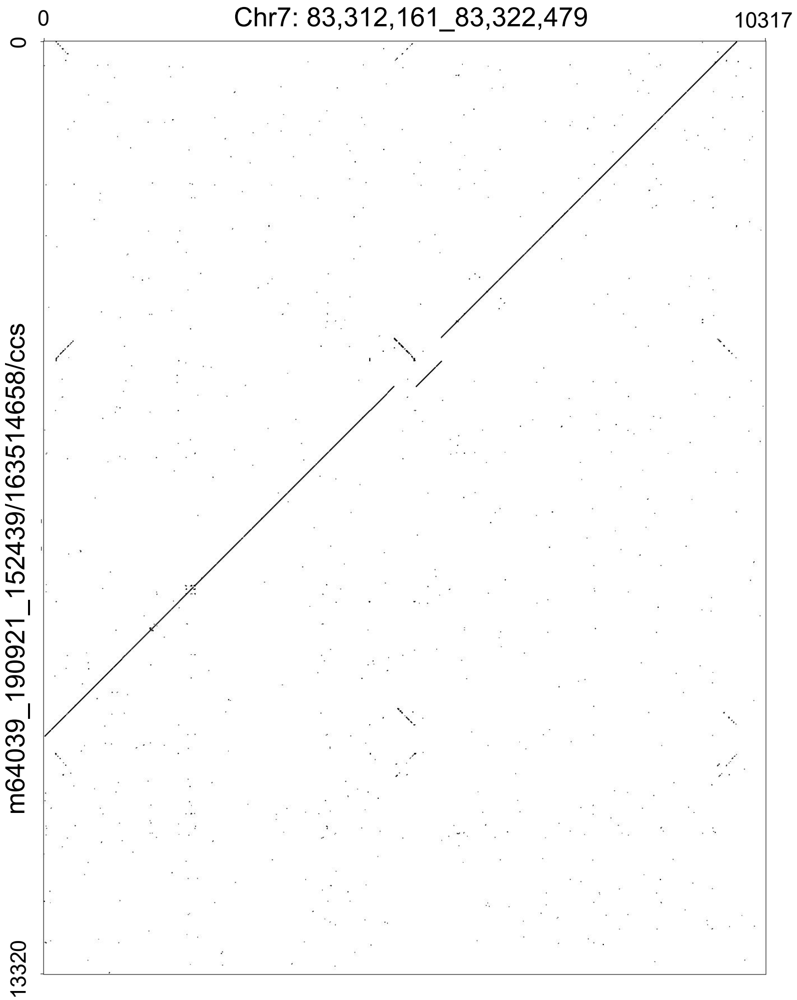


**Supplementary Figure S8** The IGV view and sequence dot-plot of the adjacent segment swap from NA19240 at Chr7: 83,316,809_83,317,466

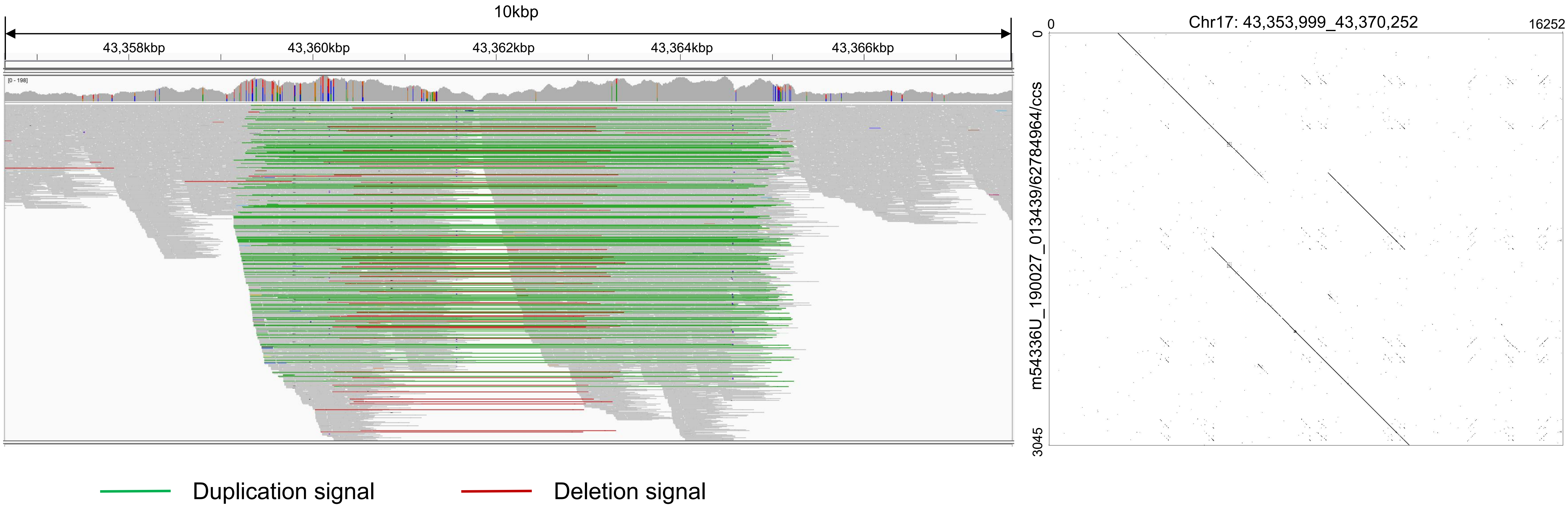


**Supplementary Figure S9** The IGV view and sequence dot-plot of the tandem dispersed duplication from NA19240 at Chr17: 43,359,104_43,365,253

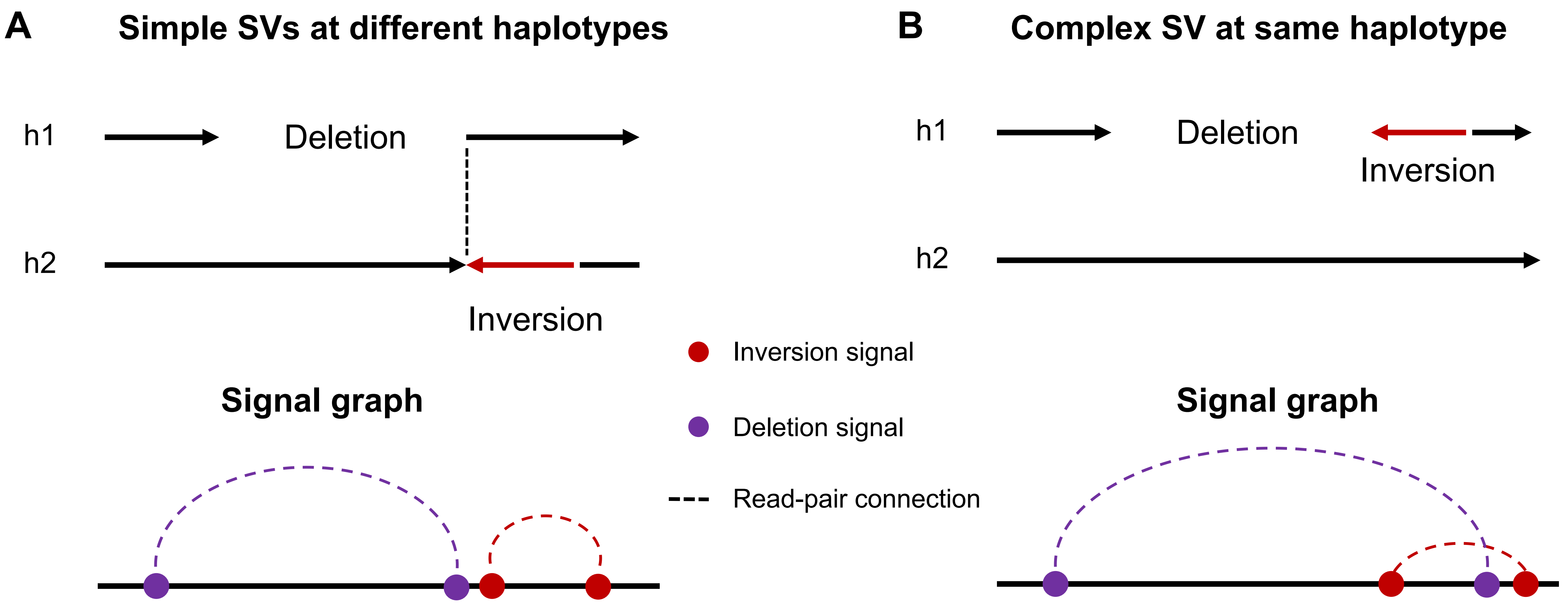


**Supplementary Figure S10** Examples to show the difference of CSV breakpoints from single haplotype or two haplotypes

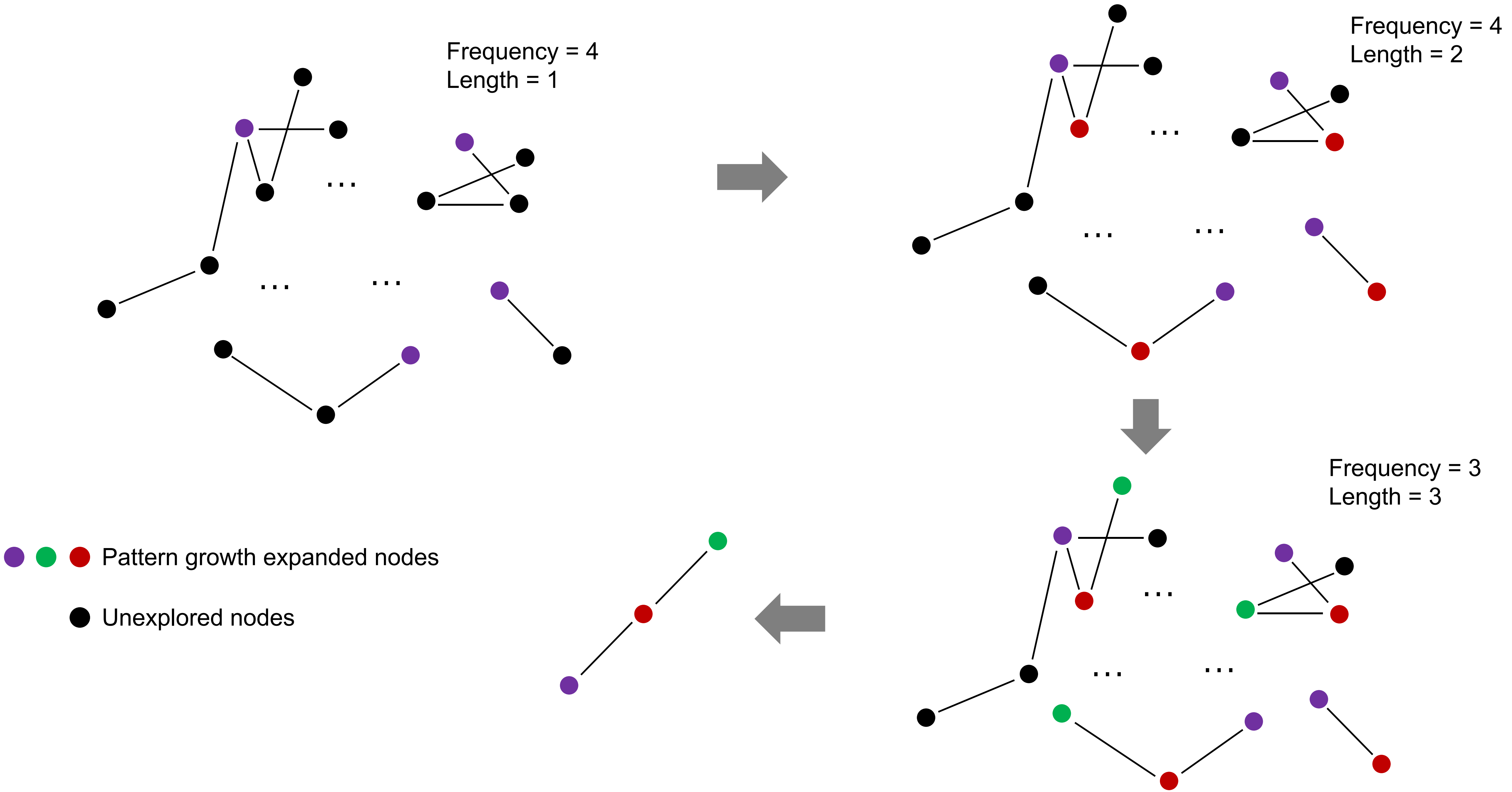


**Supplementary Figure S11** An toy example to explain the pattern growth process

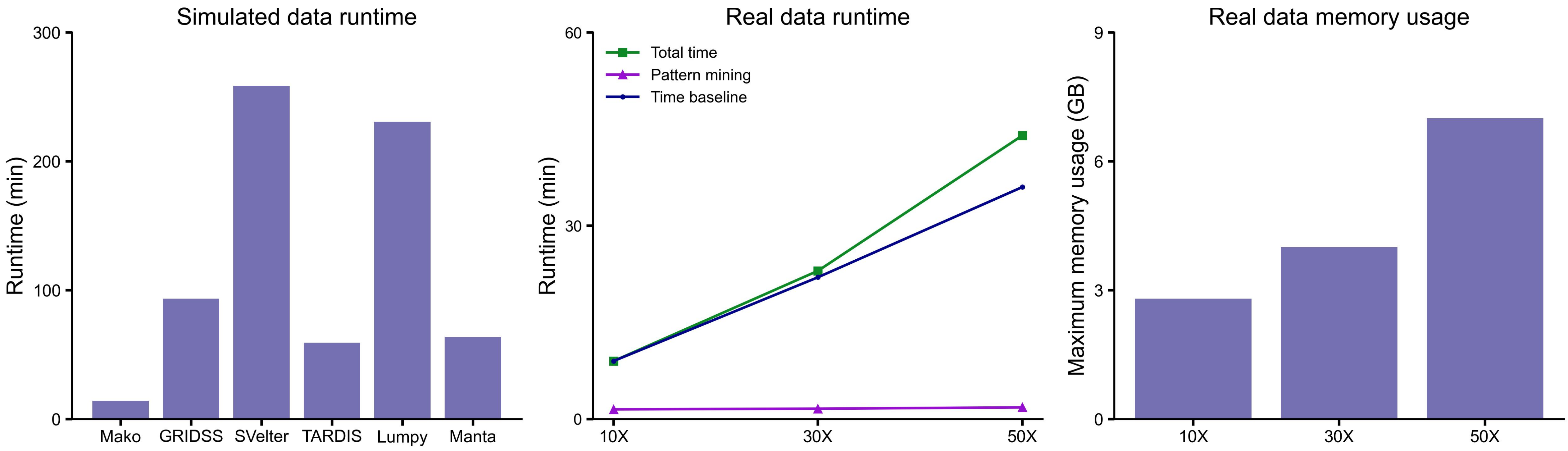


**Supplementary Figure S12** Running time comparison between different methods

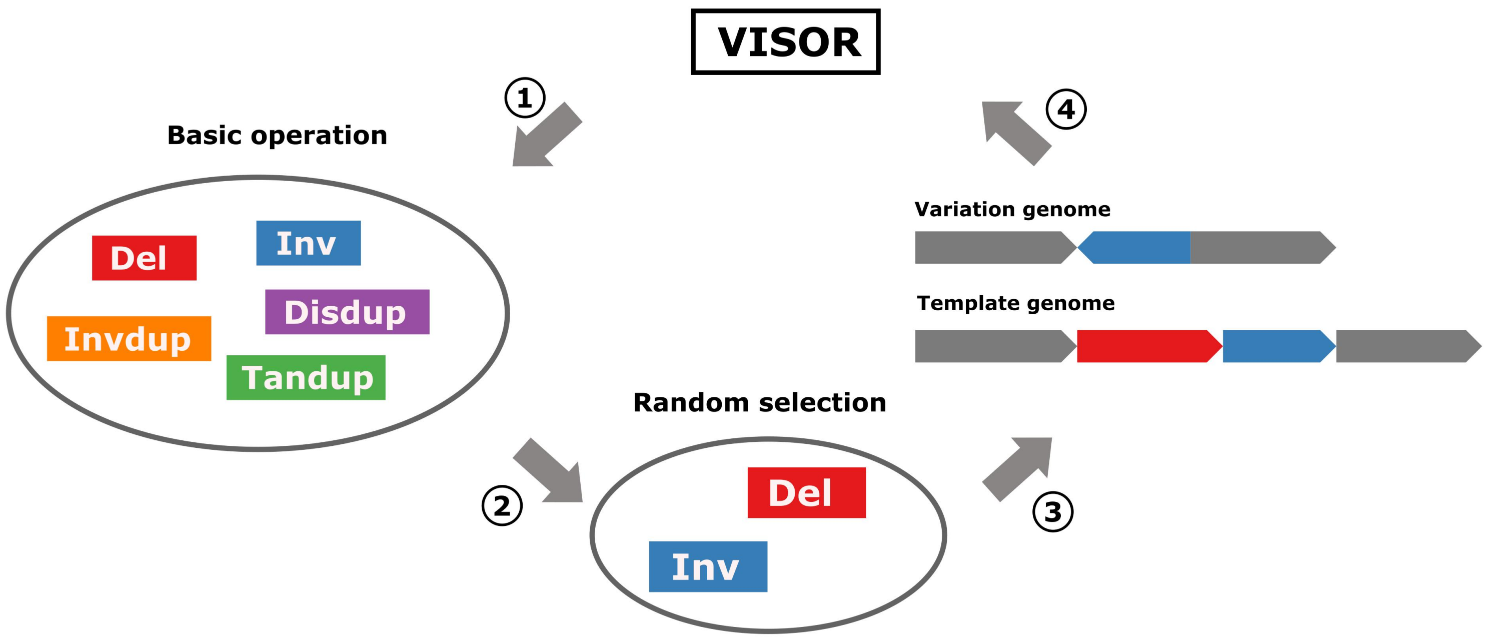


**Supplementary Figure S13** Workflow of CSV simulation

The simulation contains four major steps, including ① basic operation generation; ② operation random selection and combination; ③ variation genome simulation; ④ paired-end reads simulation and alignment. Basic operations contain Del (deletion), Inv (inversion), Disdup (dispersed duplication), Invdup (inverted duplication) and Tandup (tandem duplication).

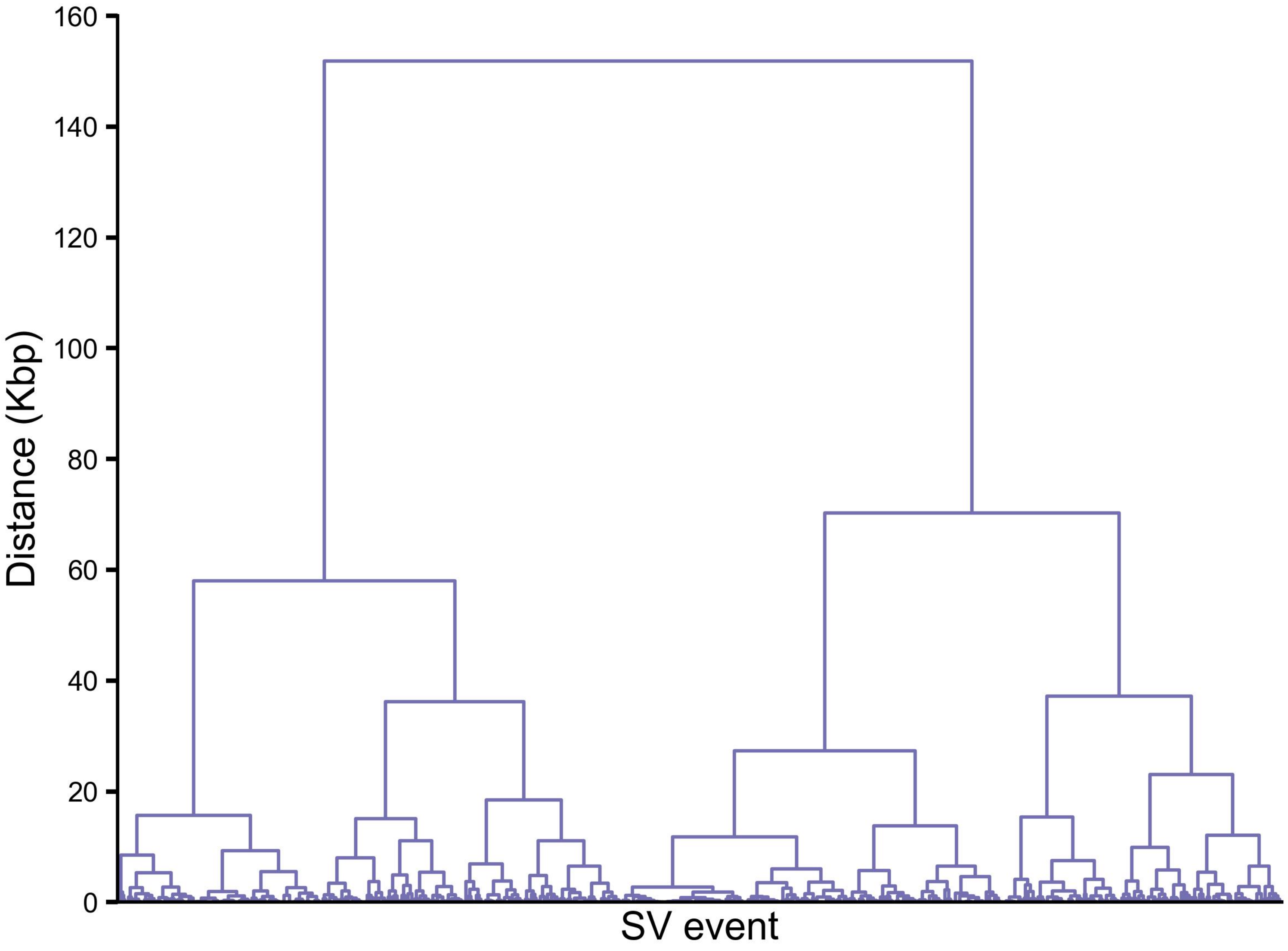


**Supplementary Figure S14** Hierarchical clustering tree view of SVs from NA19240 chromosome 1
The y-axis is the distance to select as cutoff for leaf nodes merge. The x-axis is the SV event, and ticks are omitted due to the large numbers.

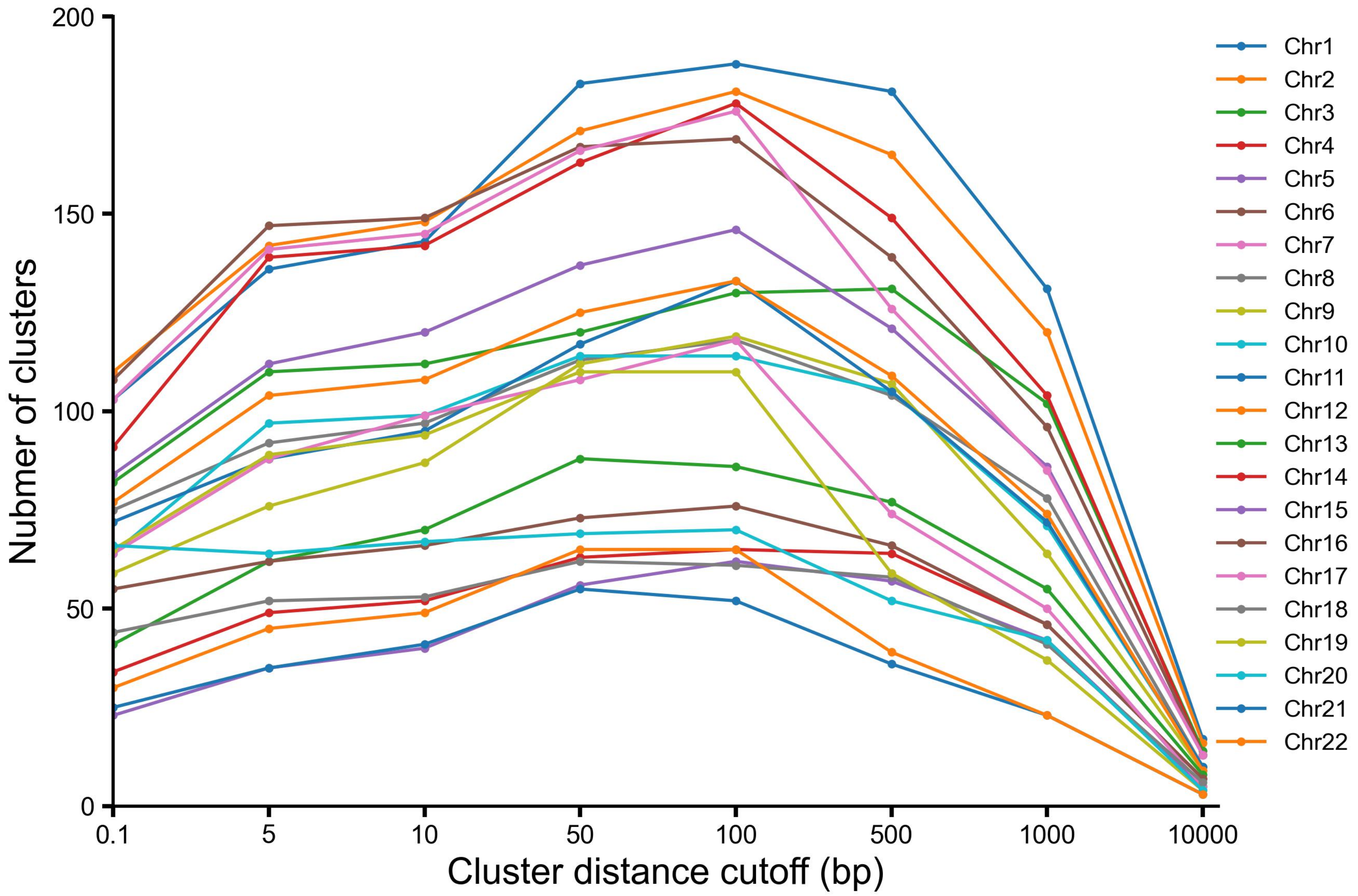


**Supplementary Figure S15** The curve plot between cluster distance cutoff and number of clusters for SVs from NA19240 autosomes

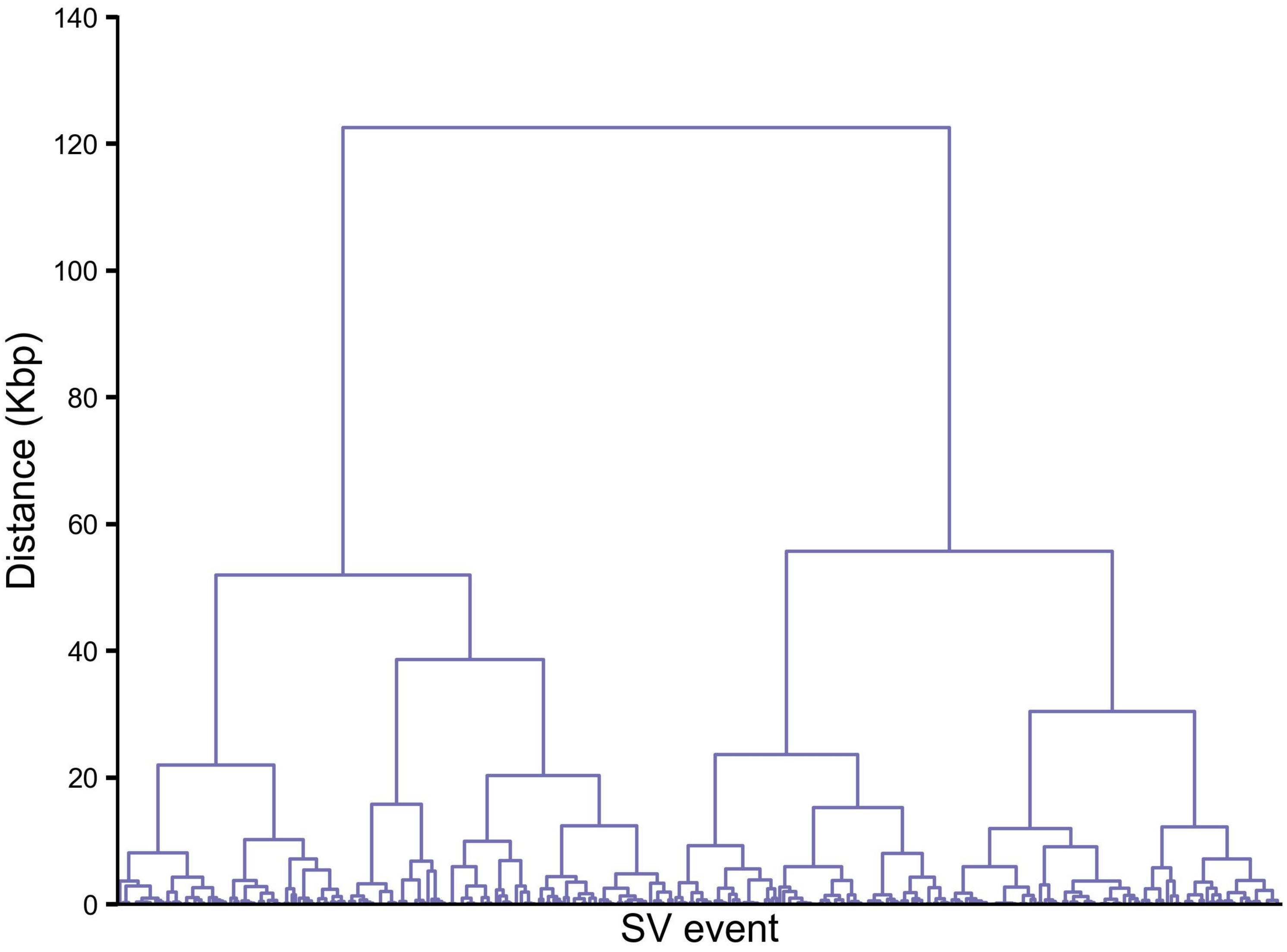


**Supplementary Figure S16** Hierarchical clustering tree view of SVs from SKBR3 chromosome 1
The y-axis is the distance to select as cutoff for leaf nodes merge. The x-axis is the SV event, and ticks are omitted due to the large numbers.

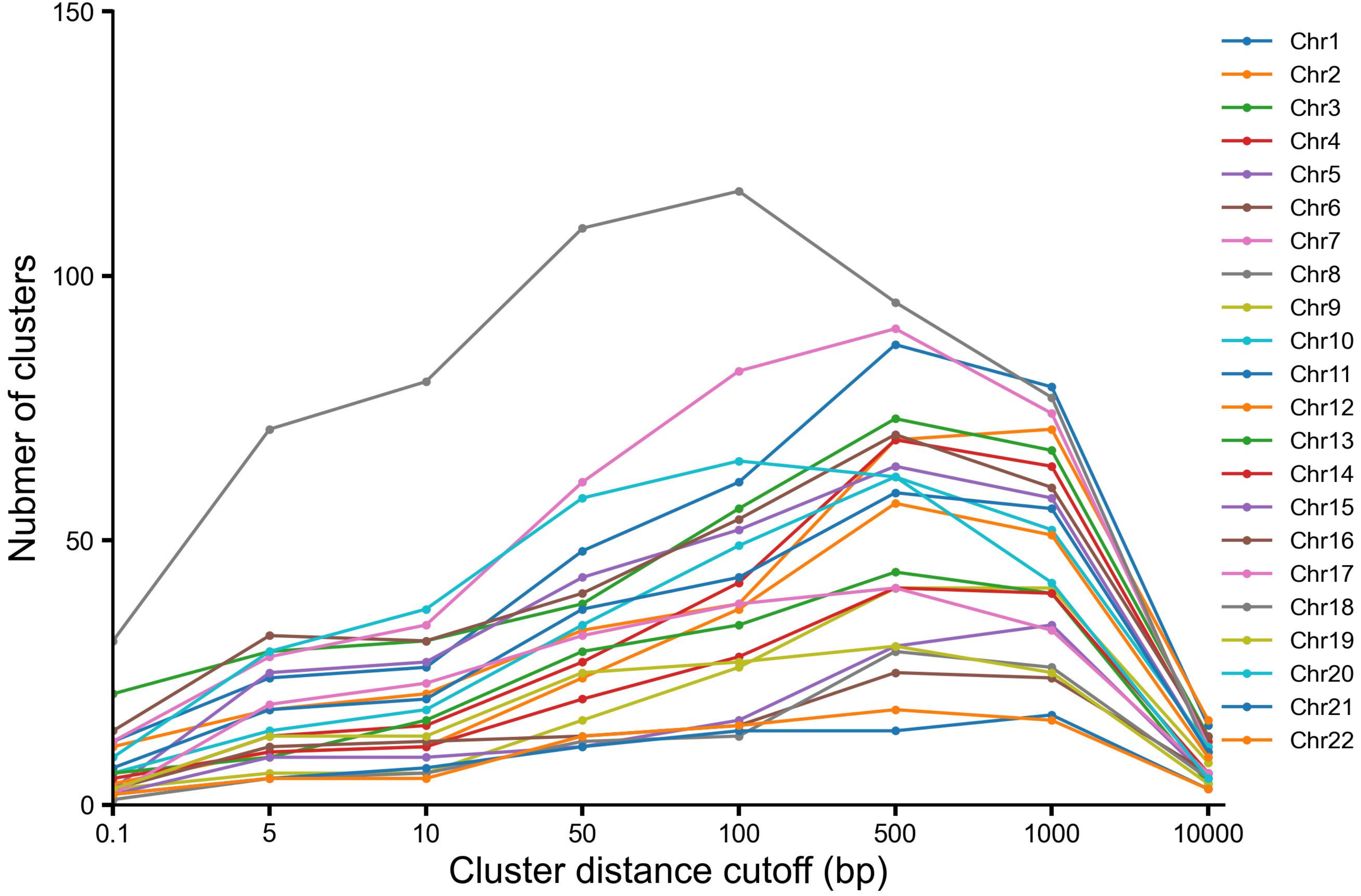


**Supplementary Figure S17** The curve plot between cluster distance cutoff and number of clusters for SVs from SKBR3 autosomes

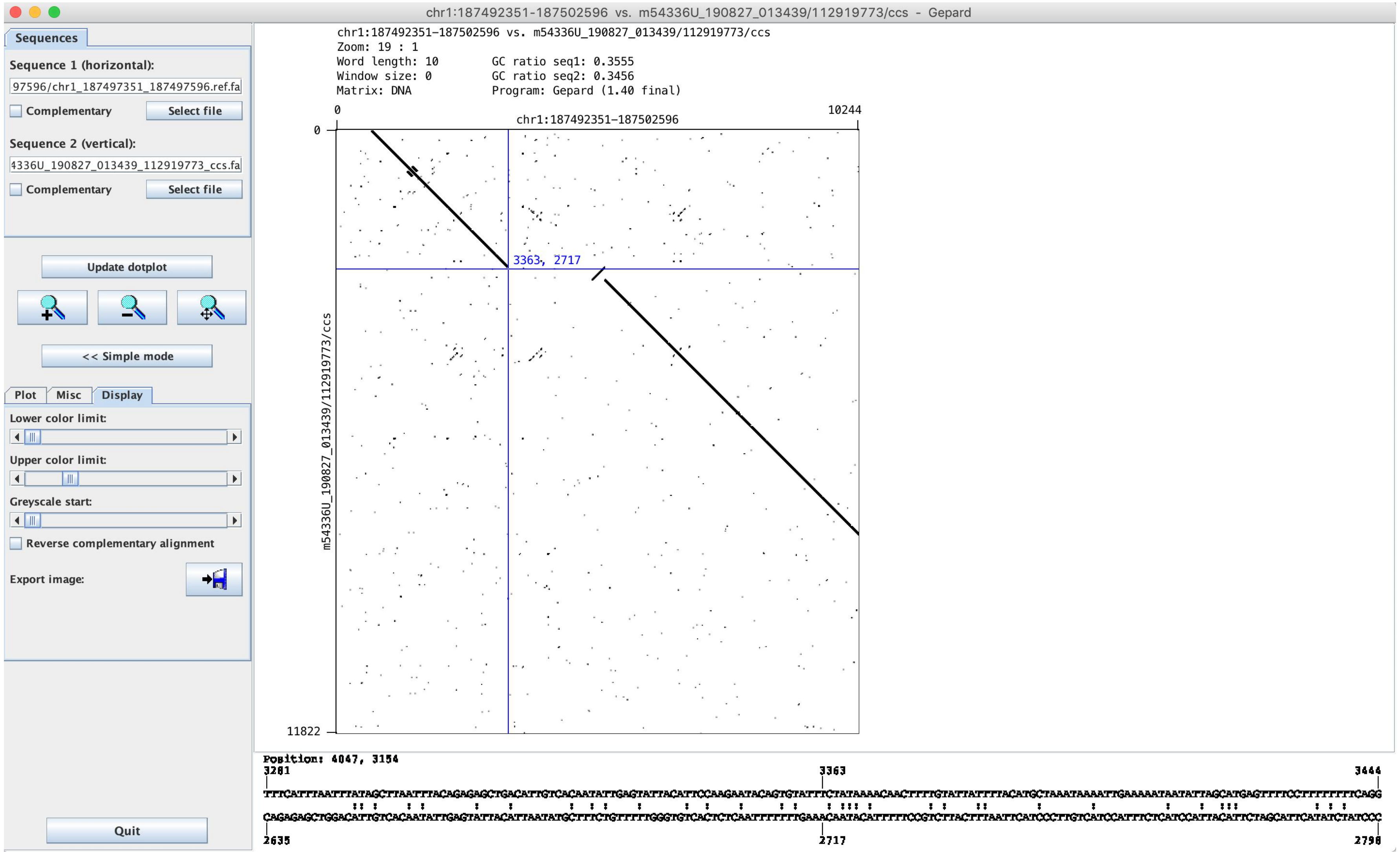


**Supplementary Figure S18** A screenshot using Gepard to investigate a deletion associated with inversion event

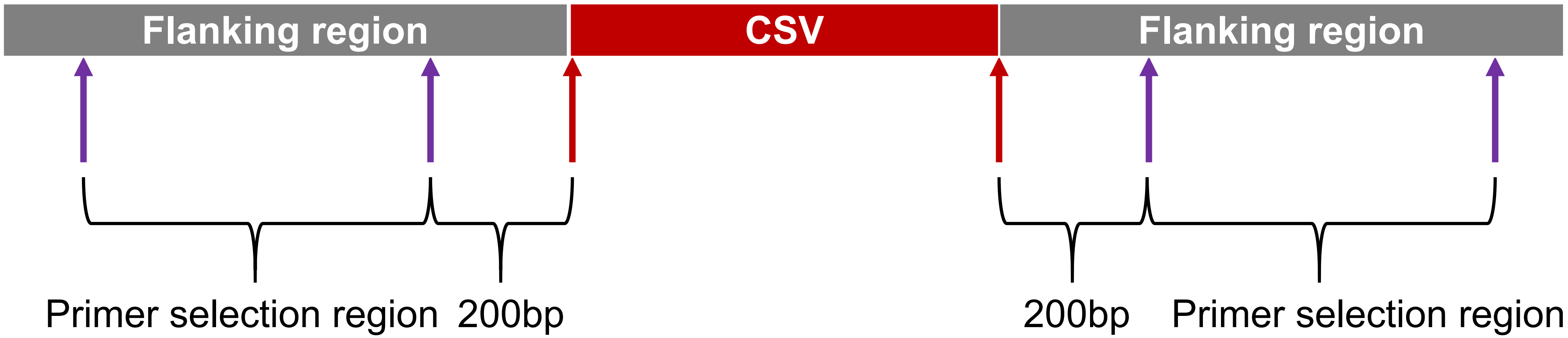


**Supplementary Figure S19** Diagram of selecting primers for each CSV

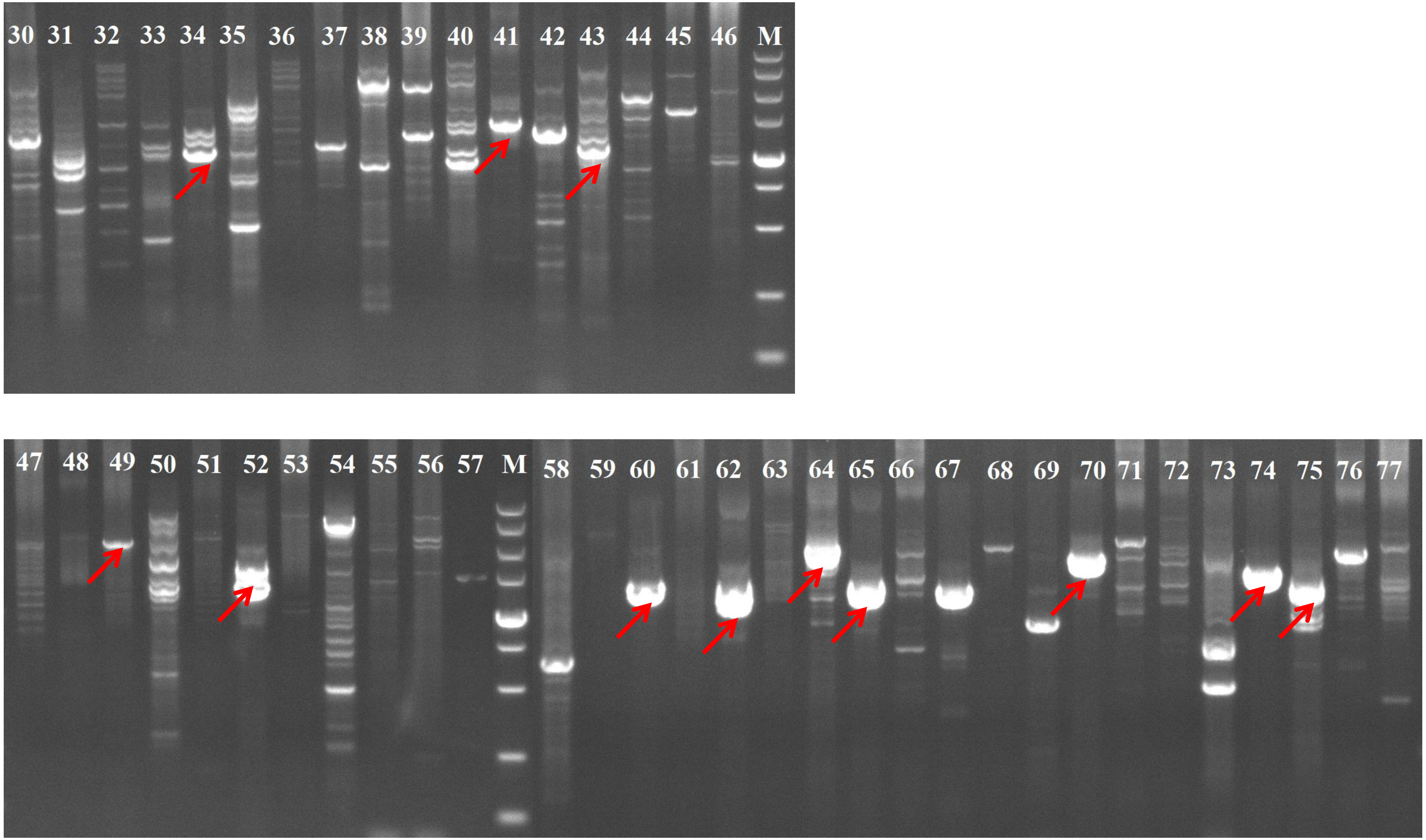


**Supplementary Figure S20** Examples of PCR electrophoretic bands visualized under the UV light
The numbers in each bar indicate variant ID (M is the marker band), the red arrows point to the ones selected for Sanger sequencing.

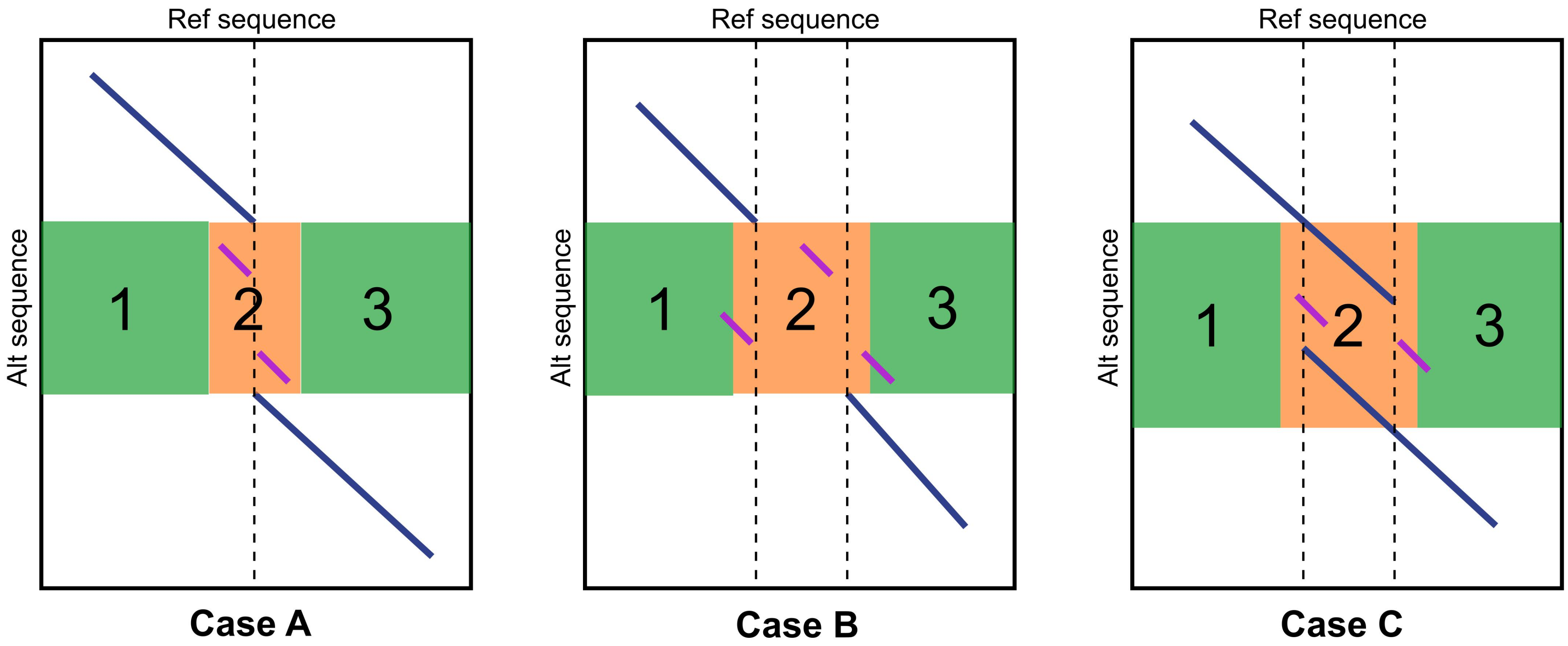


**Supplementary Figure S21** Dotplot patterns of CSVs used in characterization workflow step2
Besides the longest match (blue line), extra segments (purple line) might found in green or orange regions. The orange region contains repeat sequence, while green regions are two flanking regions of the orange one. These three cases are considered as InsDup.

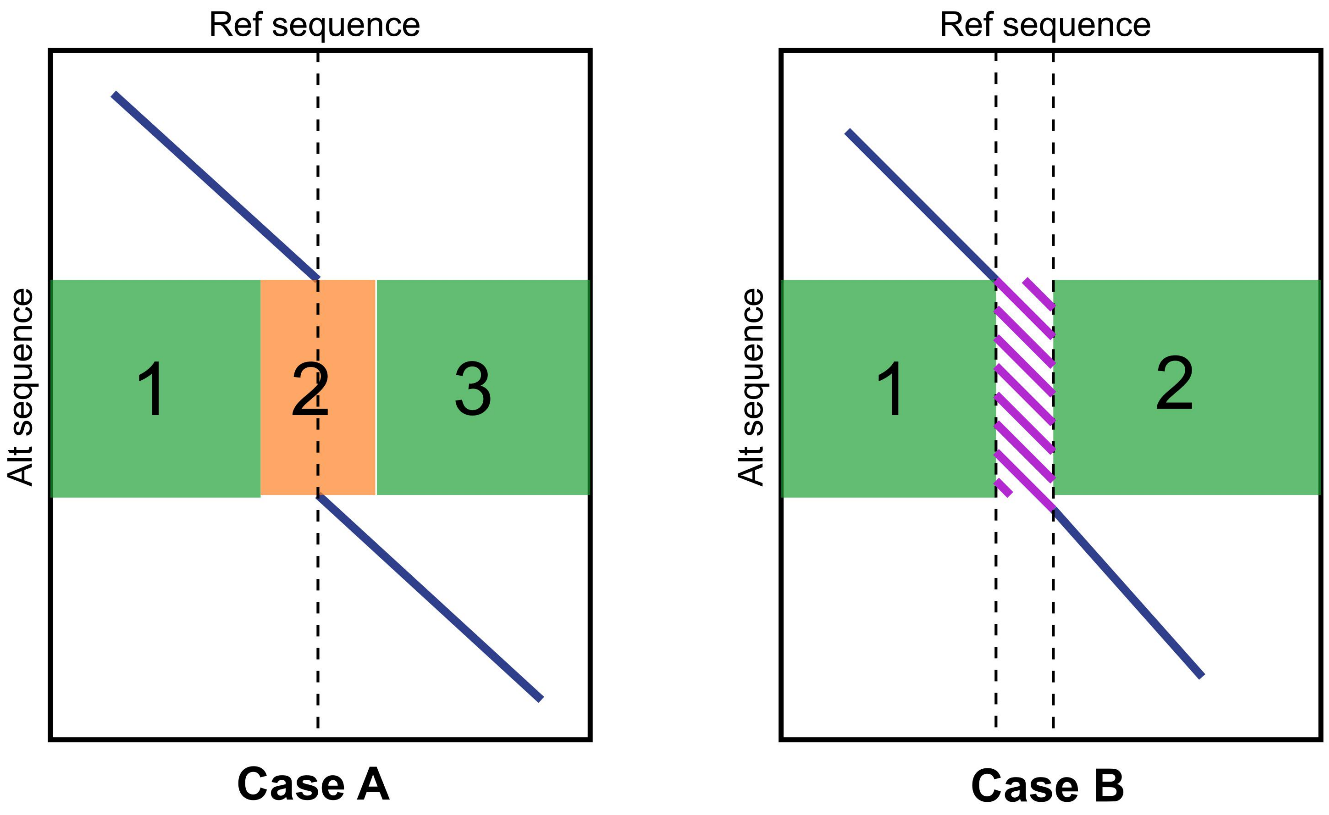


**Supplementary Figure S22** Dotplot patterns of SVs used in characterization workflow step3
Besides the longest match (blue line), extra segments (purple line) are found between two green regions. CaseA indicates insertion, and CaseB is duplication.

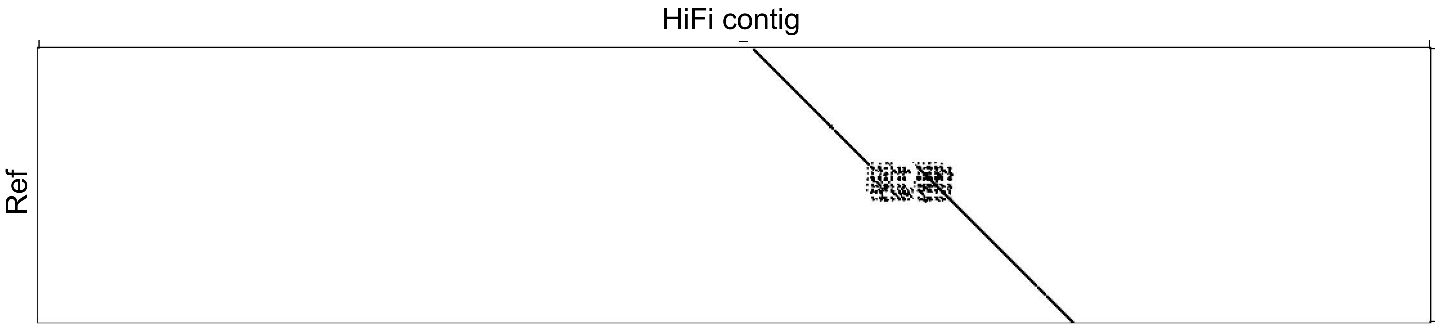


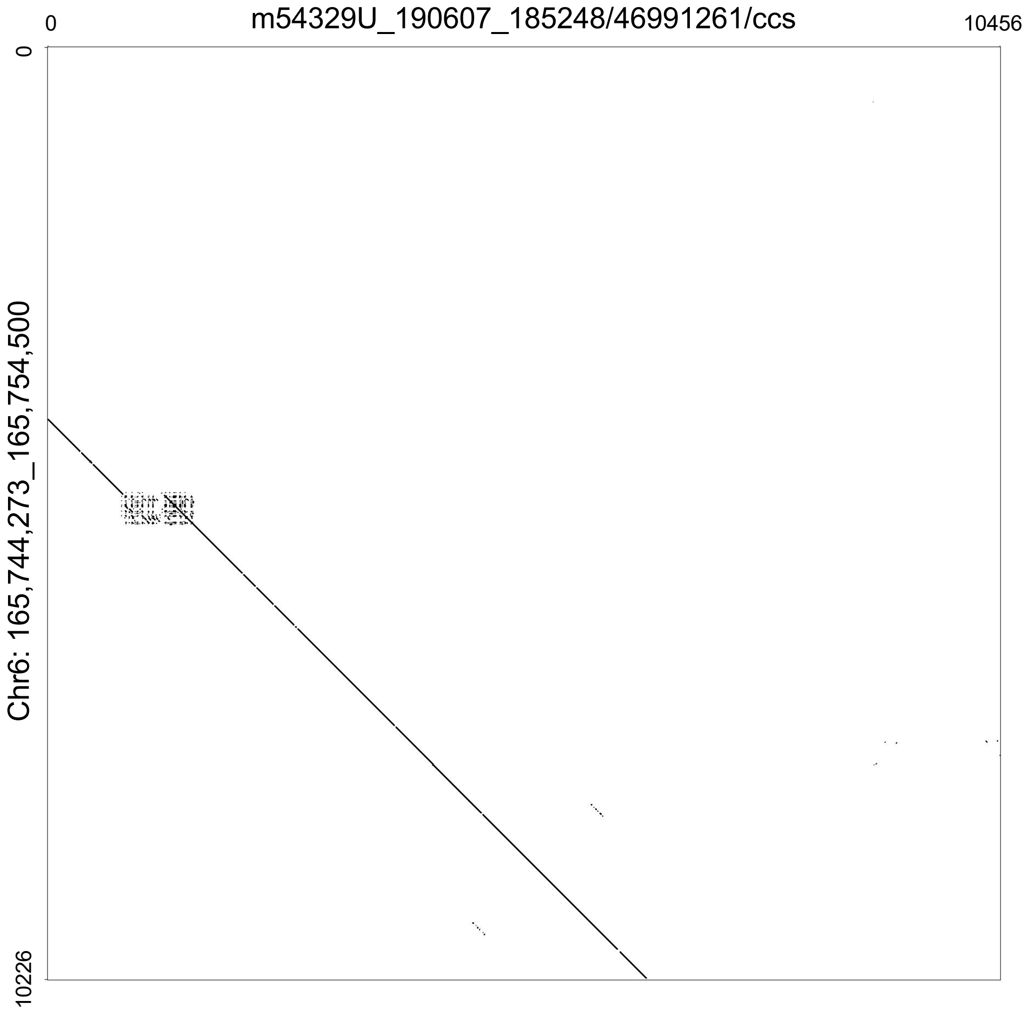


**Supplementary Figure S23** Example call labeled by VaPoR as NA at Chr6: 165,749,273_165,749,500
