## Supplementary File 1 for "Mako: a graph-based pattern growth approach to detect complex structural variants"

### NA19240 benchmark CSVs

### chr1:16081190-16082430

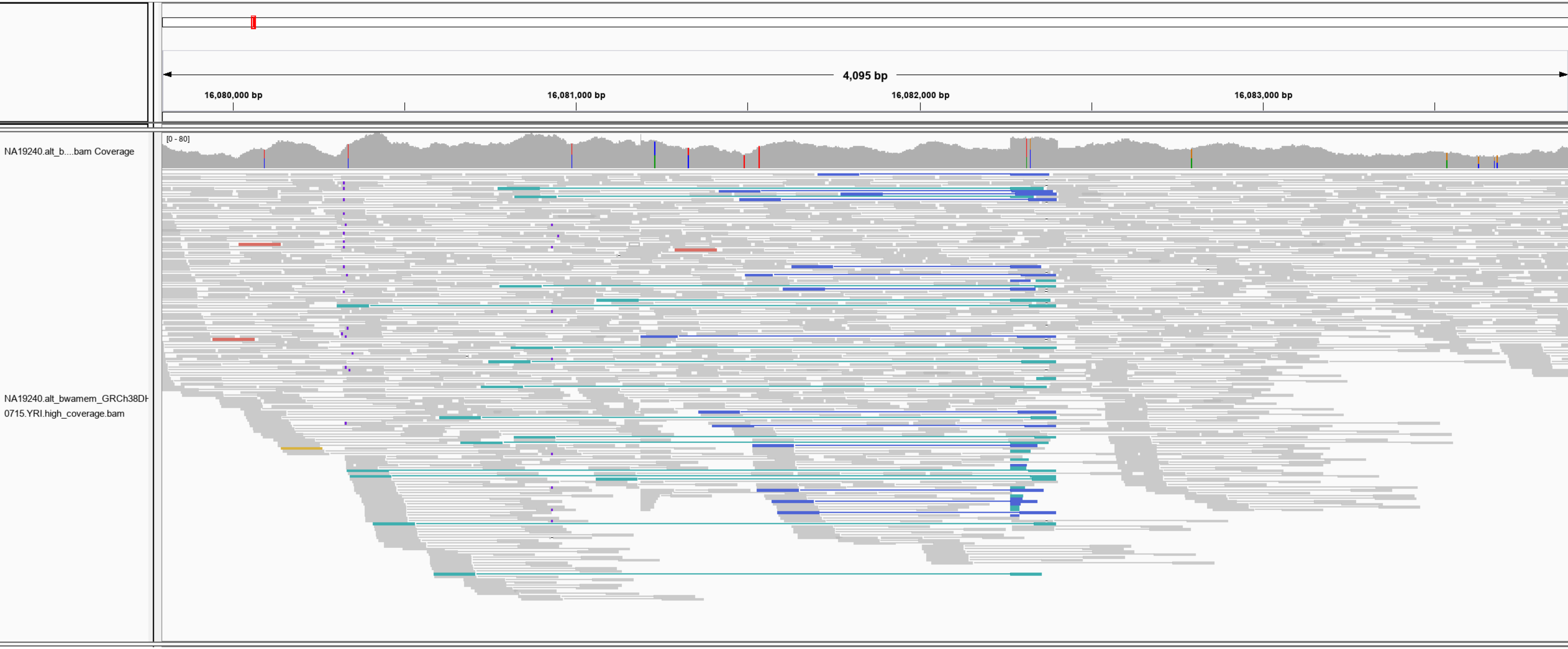

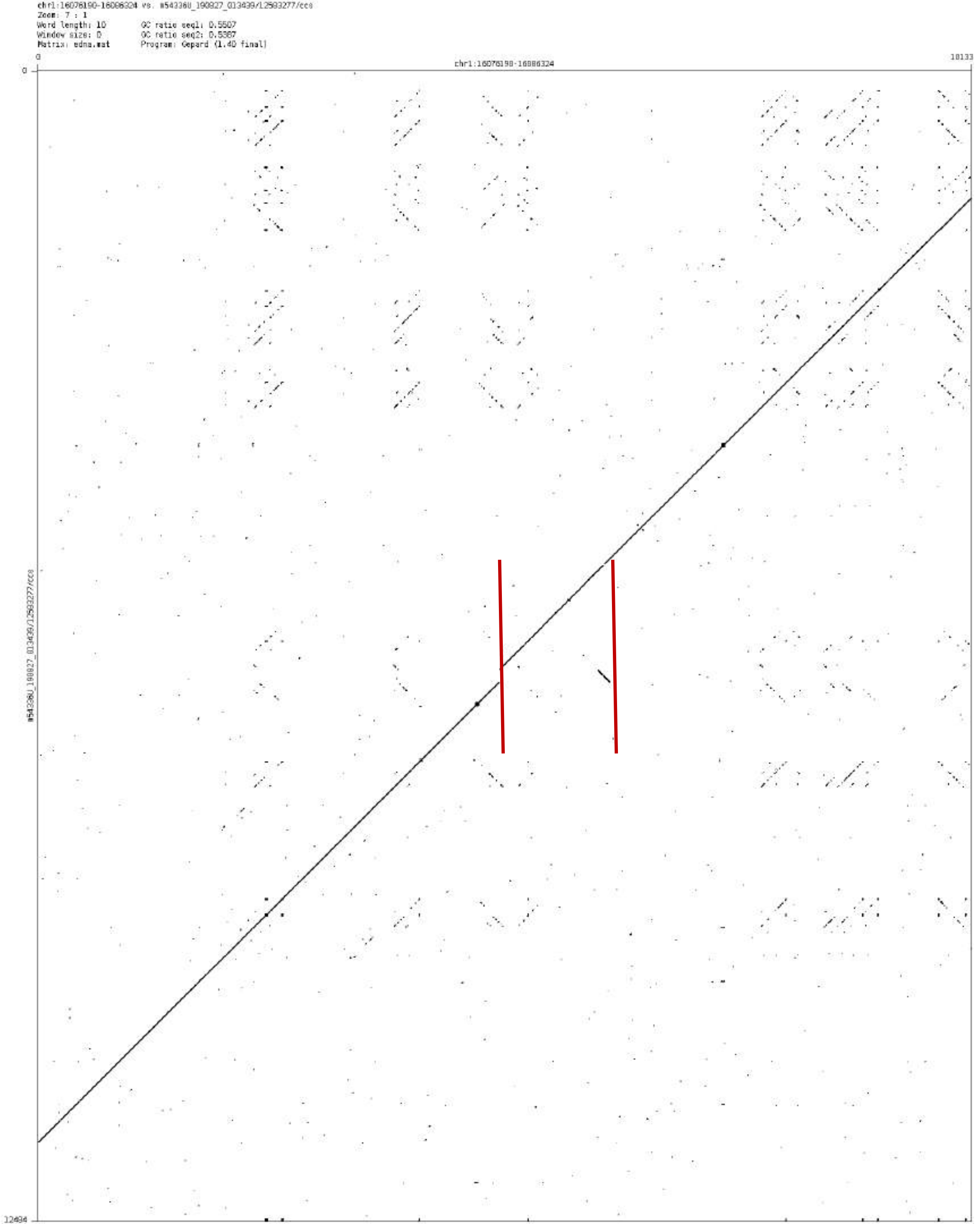
chr1:16076190-16086324 vs. m54336U_190827_013439/12583277/ccs
Zoom: 7 : 1
Word length: 10
Window size: 0
Matrix: edna.mat
GC ratio seq1: 0.5507
GC ratio seq2: 0.5387
Program: Gepard (1.40 final)
chr1:16076190-16086324
m54336U_190827_013439/12583277/ccs
0
18133
12494

chr1:41278200-41279034

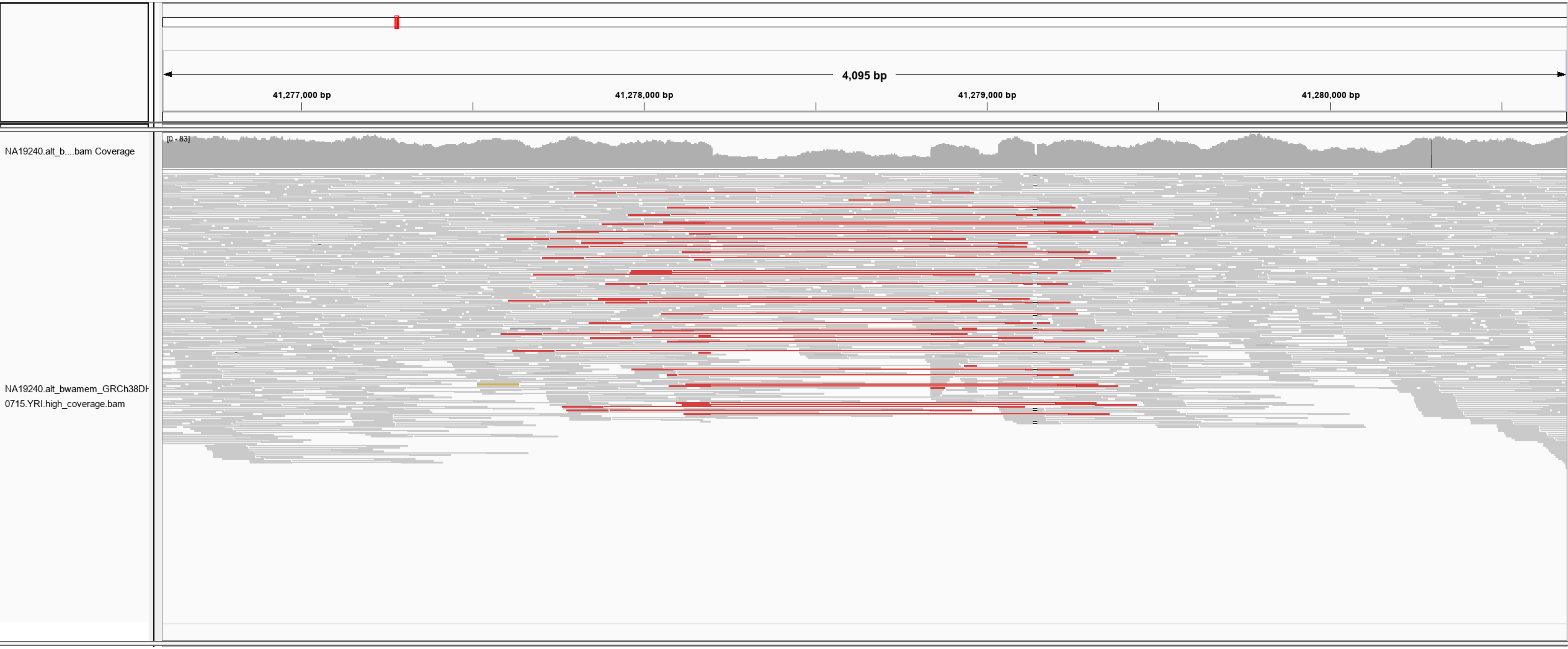

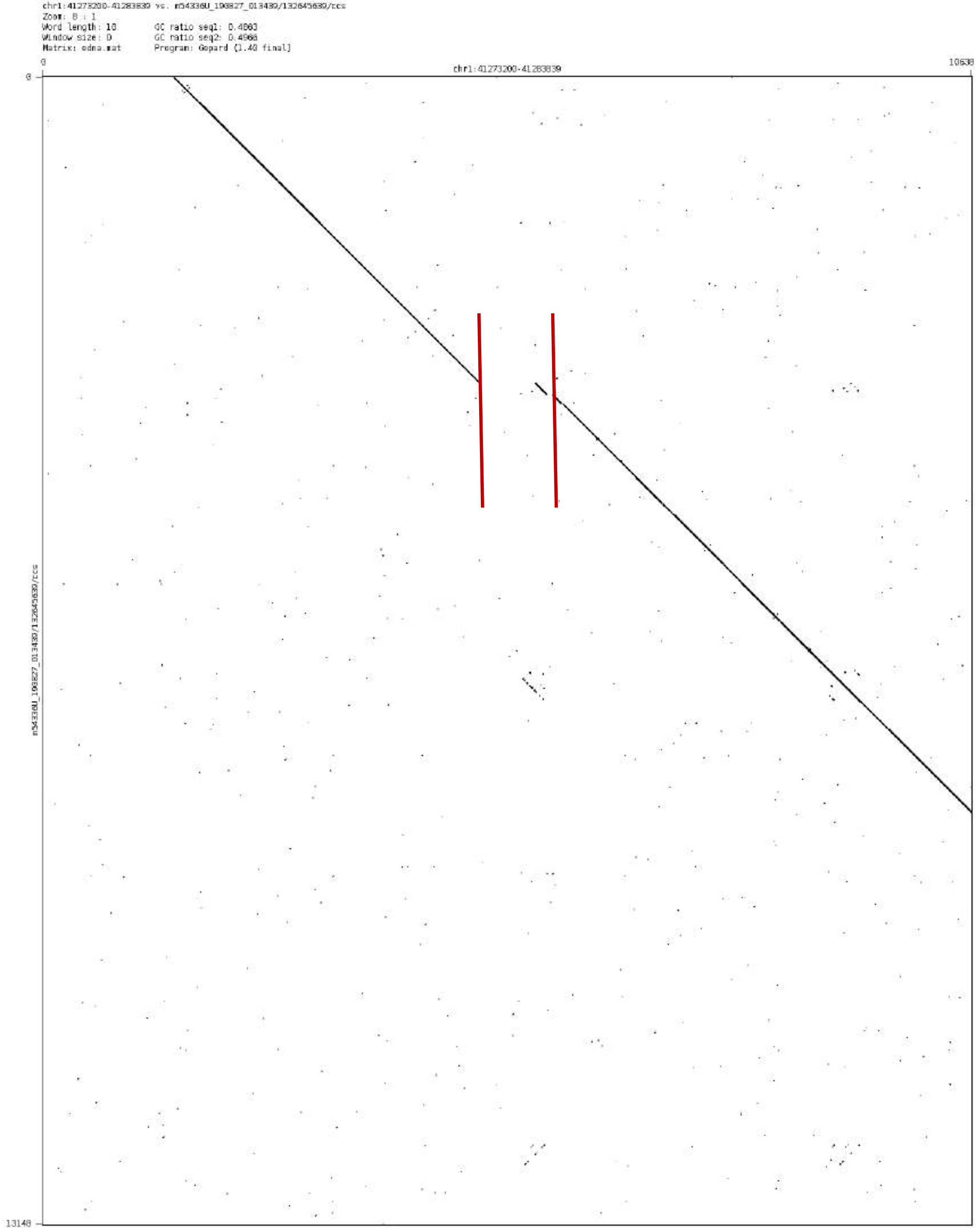
chr1:41273200-41283839 vs. m54336U_190827_013439/132645639/ccs
Zoom: 8 : 1
Word length: 10
Window size: 0
Matrix: edna.mat
GC ratio seq1: 0.4863
GC ratio seq2: 0.4968
Program: Gepard (1.40 final)
chr1:41273200-41283839
0
10638
0
13148
m54336U_190827_013439/132645639/ccs

chr1:43593621-43594291

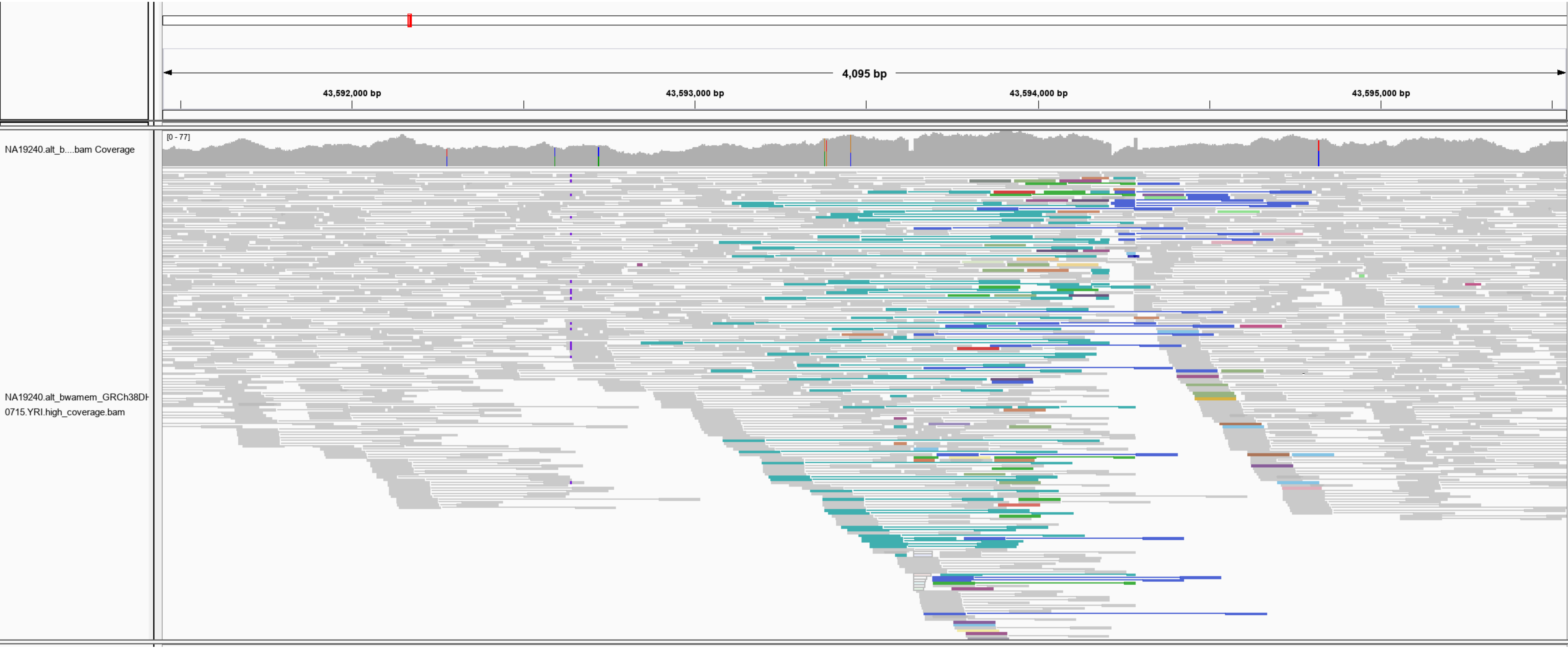

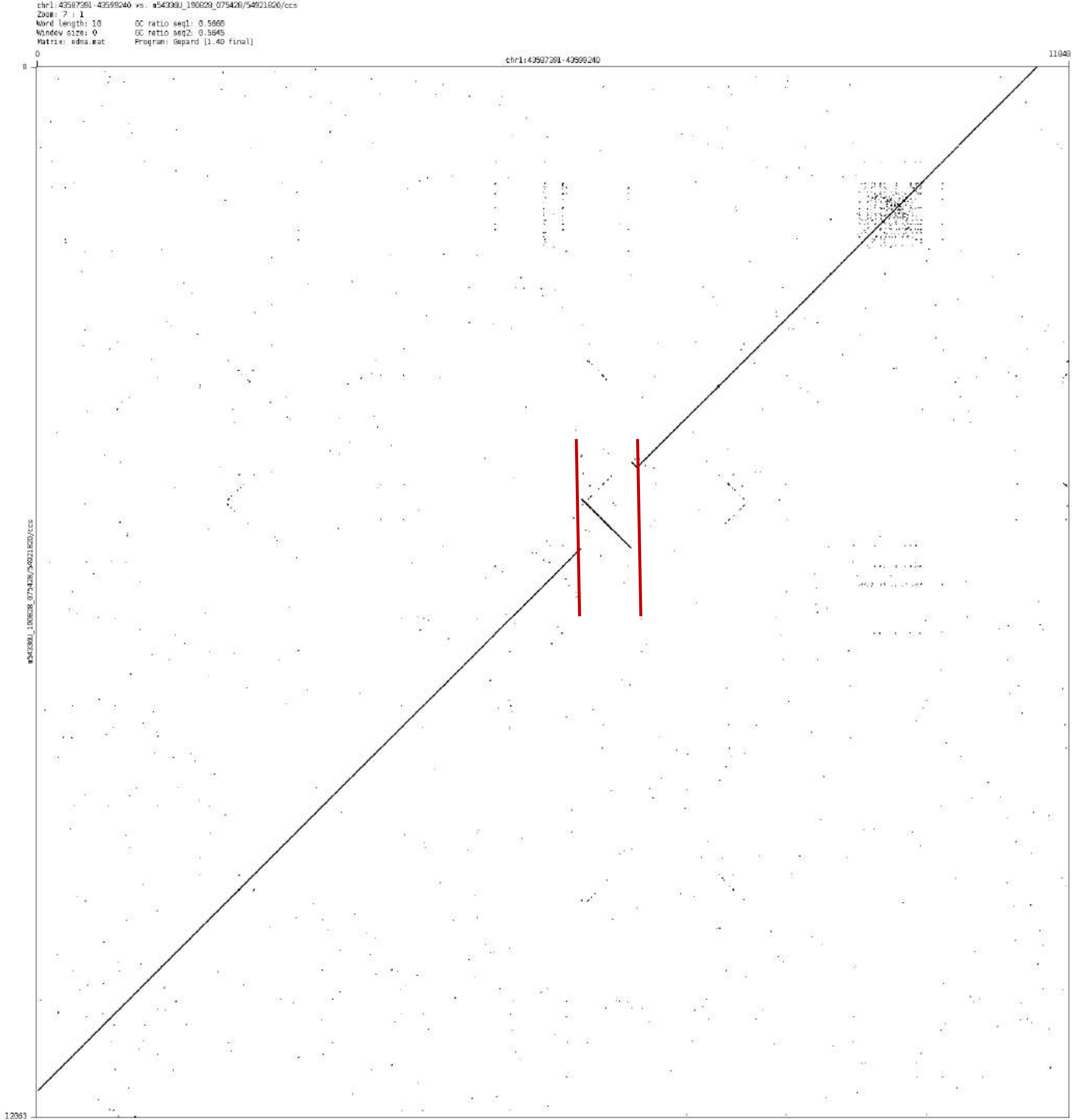
chr1:43587391-43599240 vs. m54338U_190828_075428/54921820/ccs
Zoom: 7 : 1
Word length: 10
Window size: 0
Matrix: edna.mat
GC ratio seq1: 0.5668
GC ratio seq2: 0.5645
Program: Gepard [1.40 final]
0
chr1:43587391-43599240
11849
0
12063

### chr1:175231315-175232795

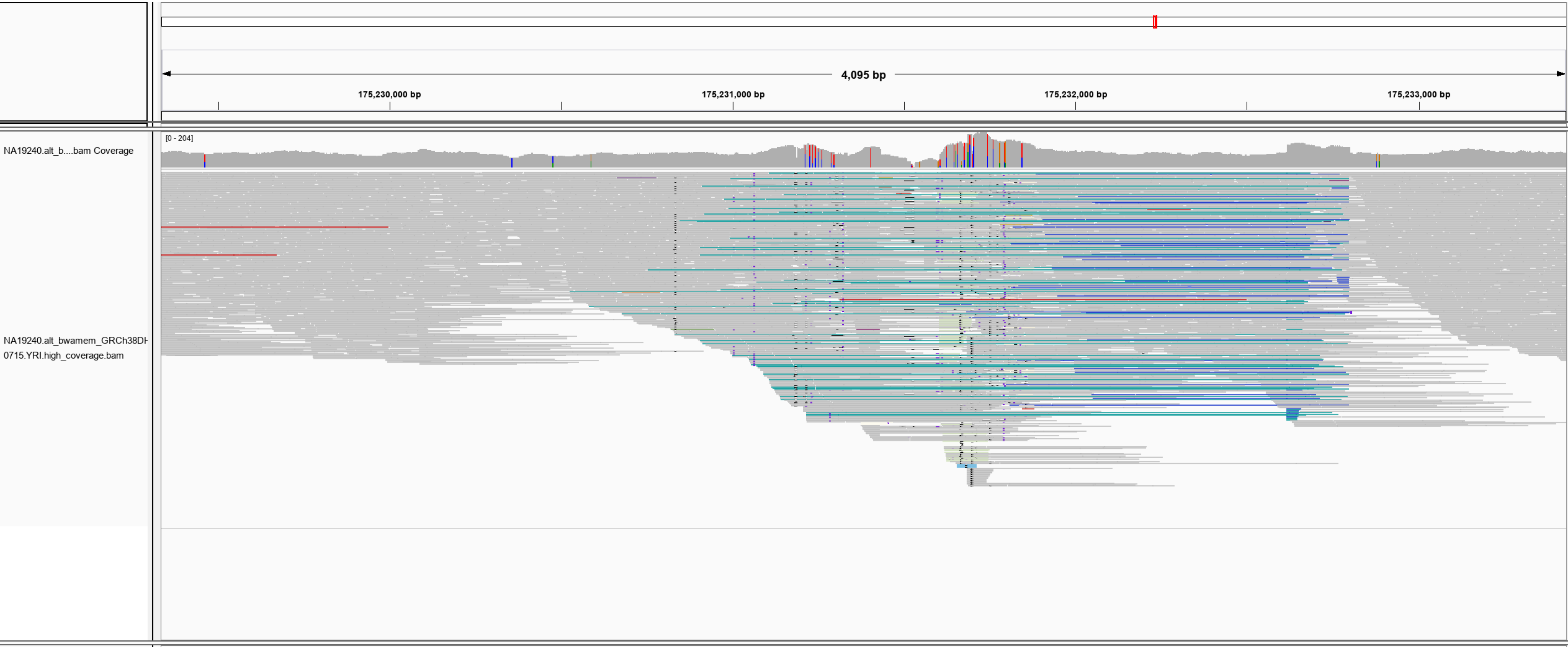

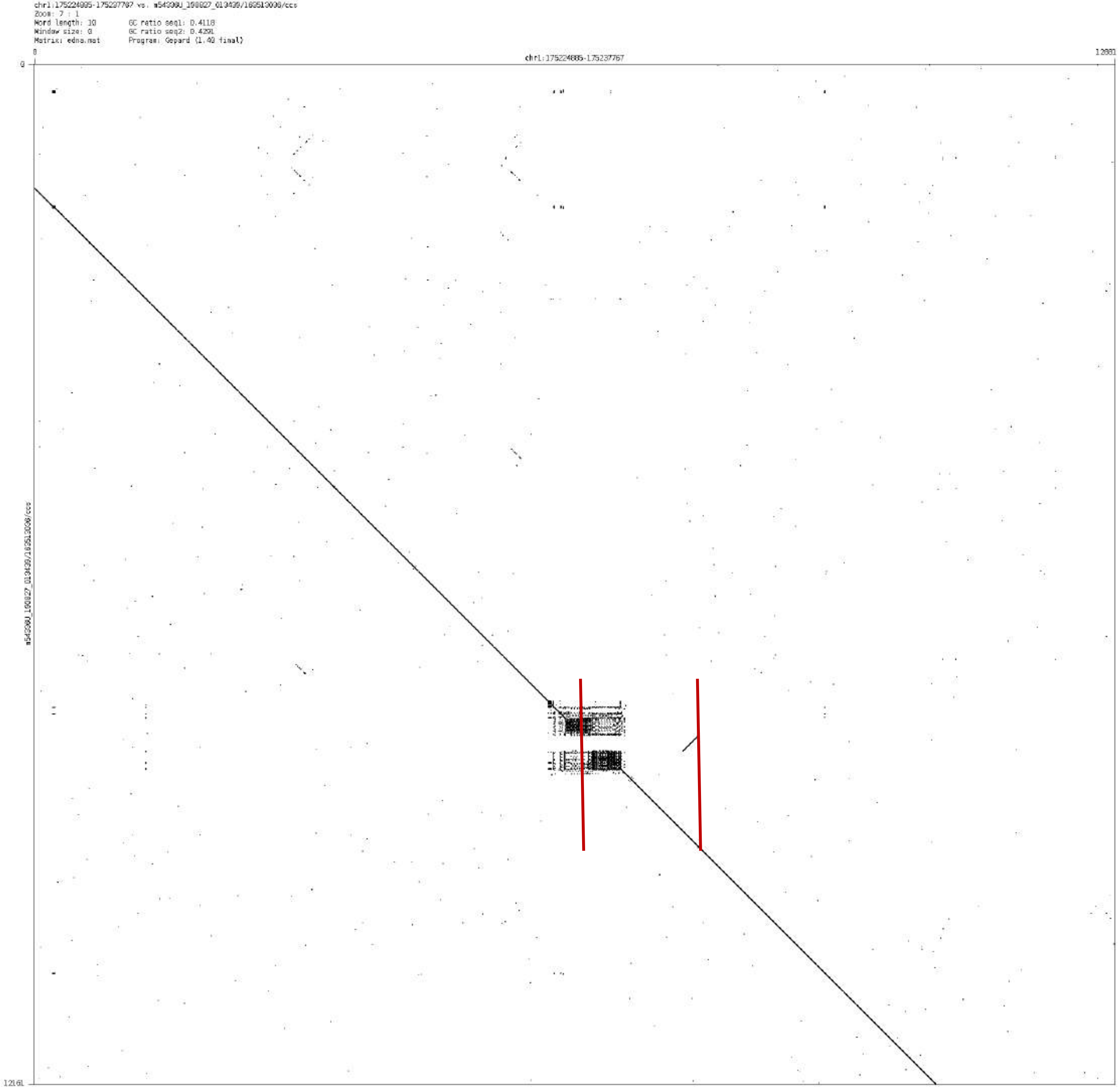
chr1:175224885-175237767 vs. m54330U_190827_013439/163513036/ccs
Zoom: 7 : 1
Word length: 10
Window size: 0
Matrix: edna.mat
GC ratio seq1: 0.4118
GC ratio seq2: 0.4291
Program: Gepard (1.40 final)
0
chr1:175224885-175237767
12881
0
12161

### chr1:187495696-187497571

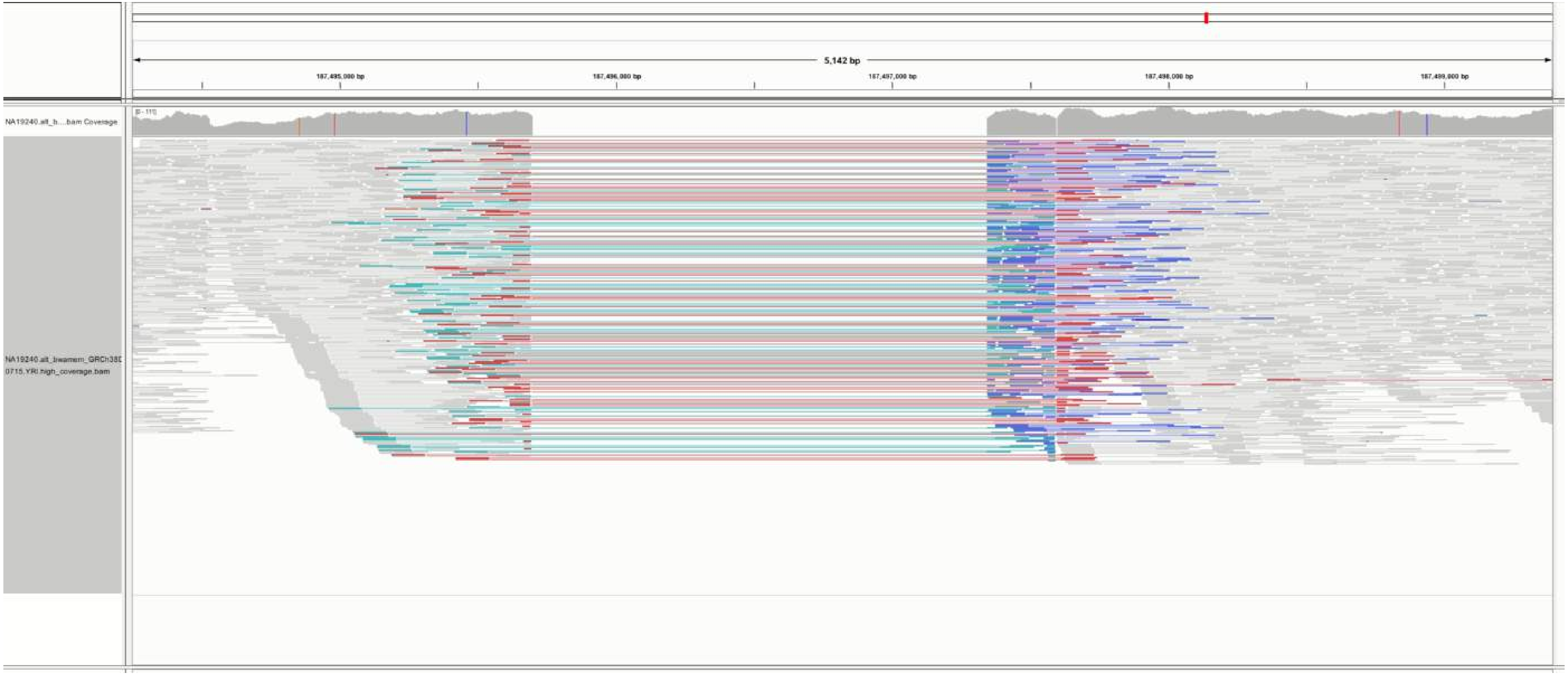

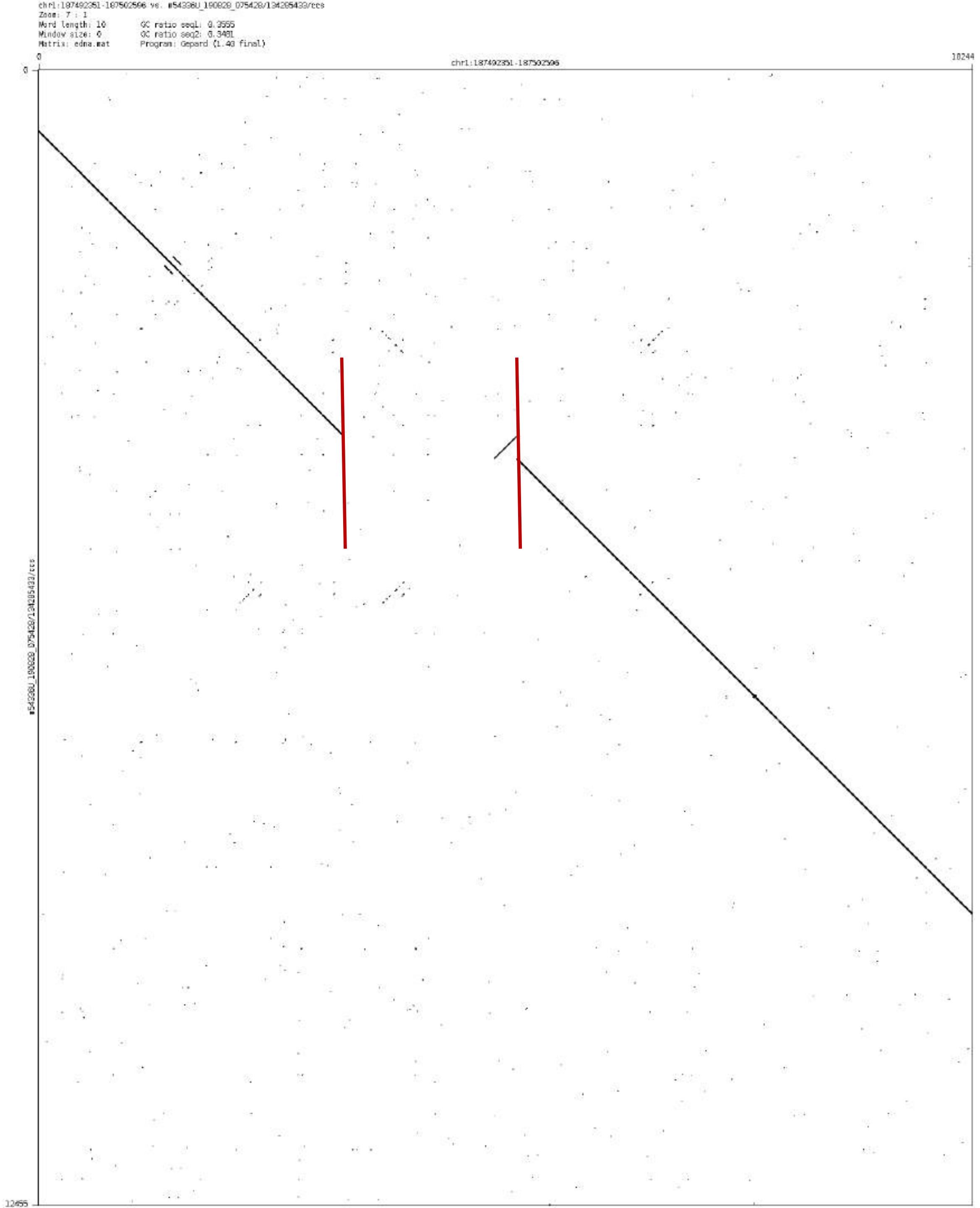
chr1:187492351-187502596 vs. m54336U_190828_075428/134285433/ccs
Zoom: 7 : 1
Word length: 10
Window size: 0
Matrix: edna.mat
GC ratio seq1: 0.3555
GC ratio seq2: 0.3481
Program: Gepard (1.40 final)
0
chr1:187492351-187502596
10244
0
12455

chr1:209761615-209762730

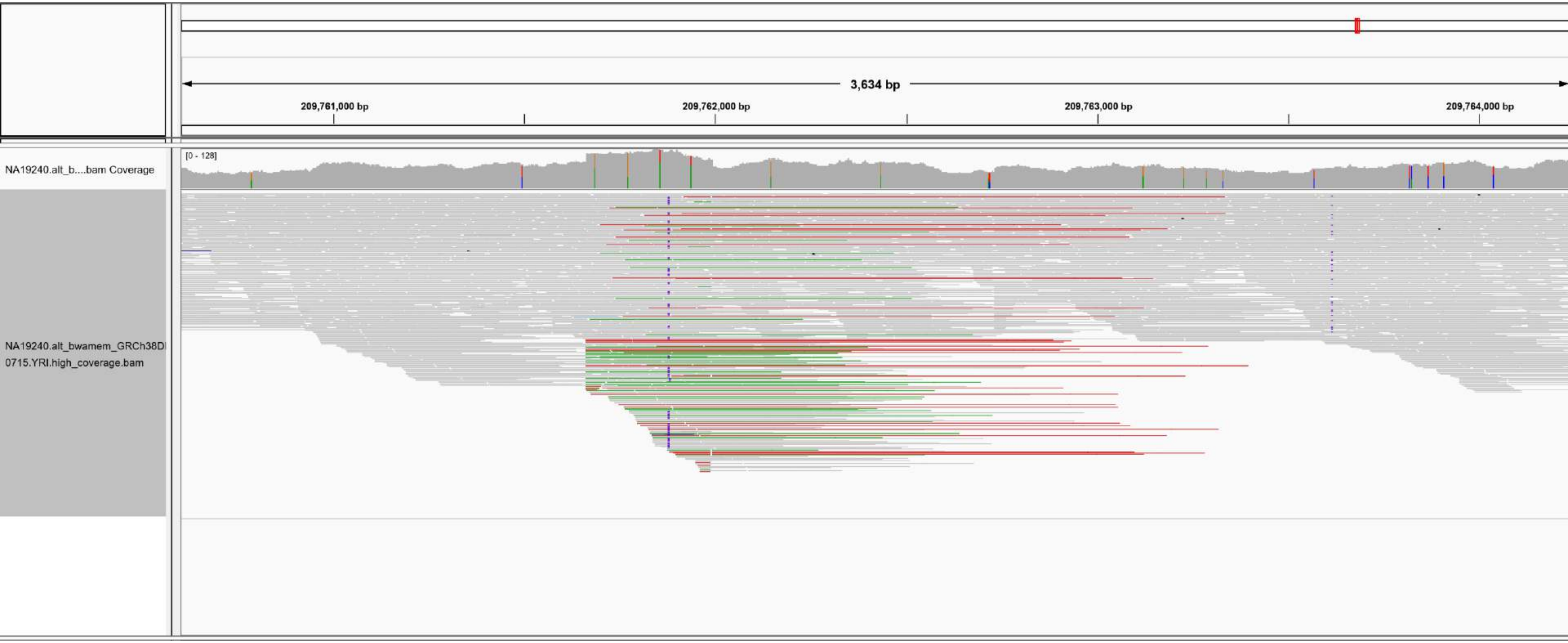

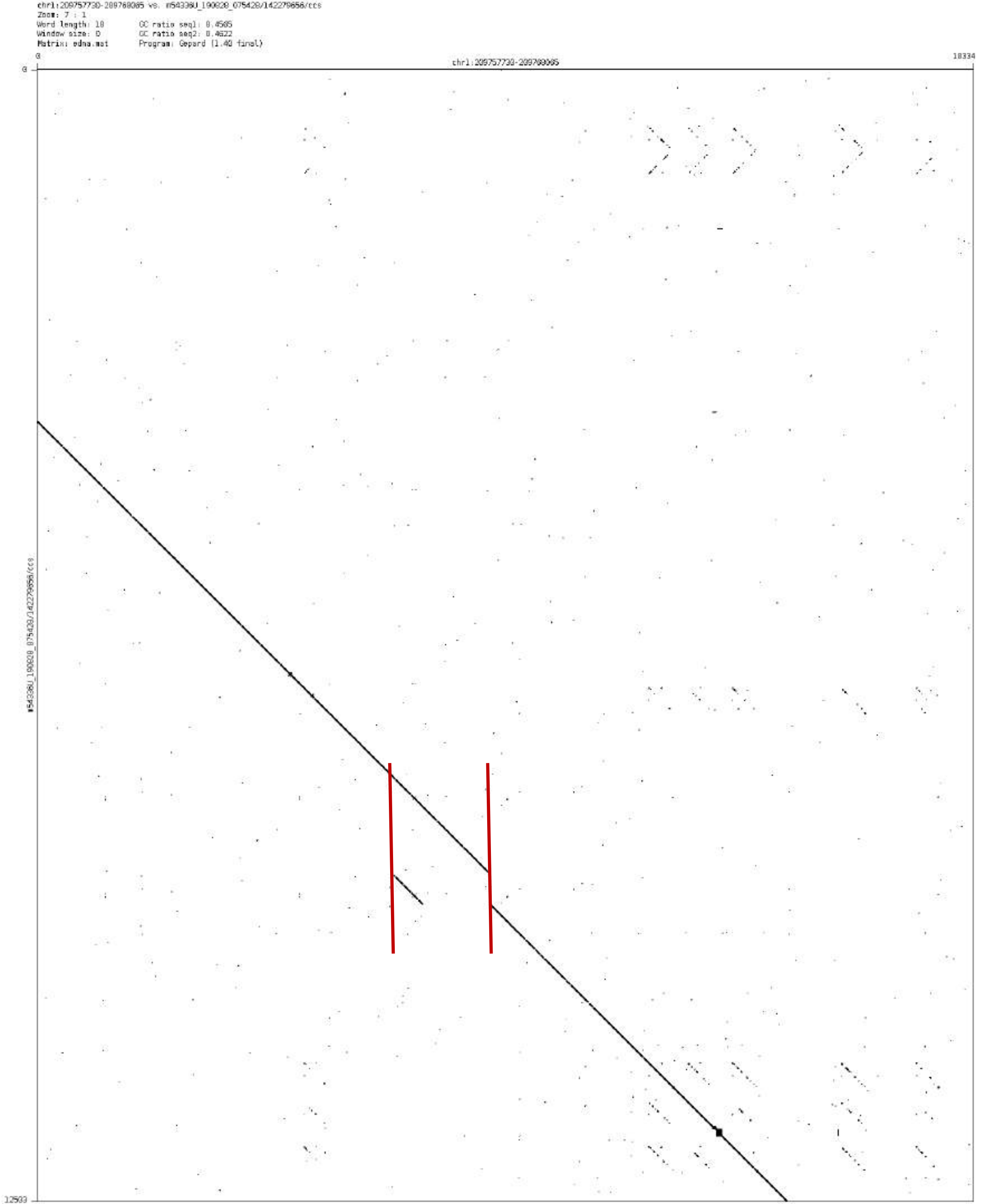

chr1:209757730-209768085 vs. m54336U_190828_075428/142279656/ccs
Zoom: 7 : 1
Word length: 10
Window size: 0
Matrix: edna.mat
GC ratio seq1: 0.4585
GC ratio seq2: 0.4622
Program: Gepard (1.40 final)
0
chr1:209757730-209768085
18334
12503
m54336U_190828_075428/142279656/ccs

chr1:237390170-237392642

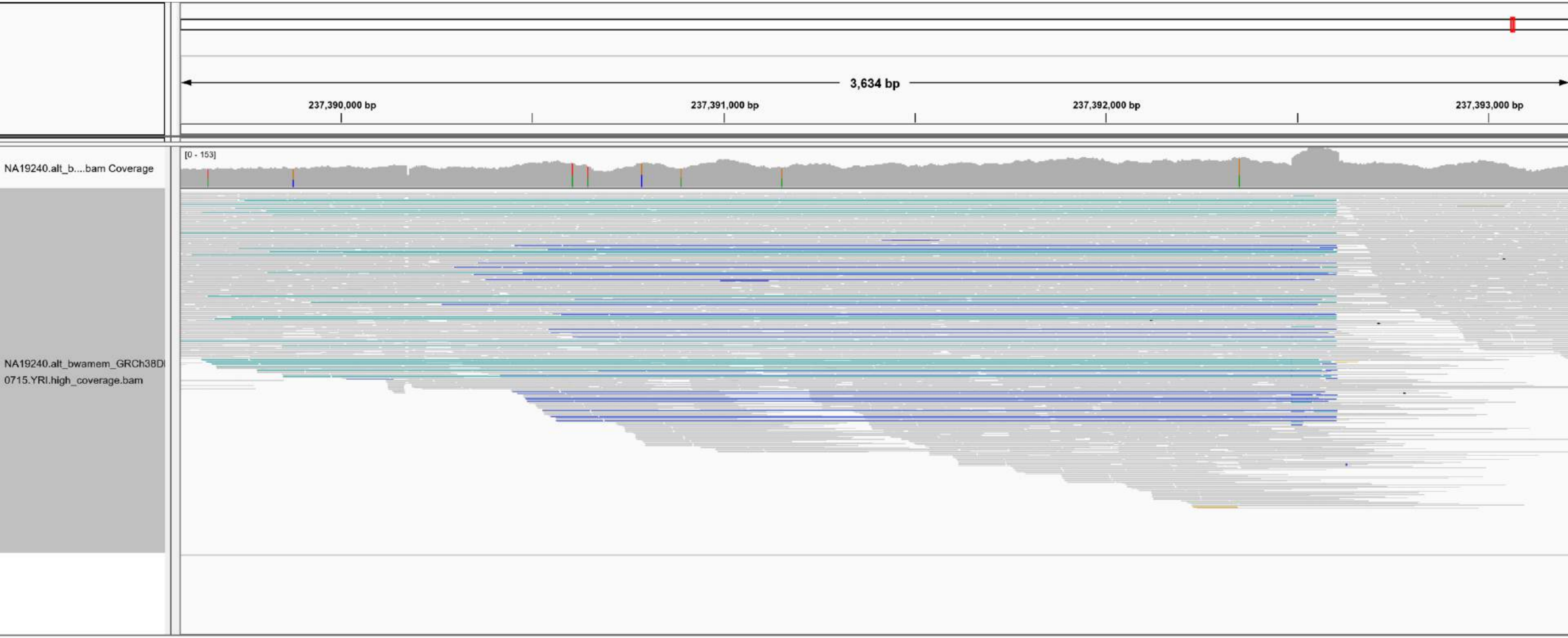

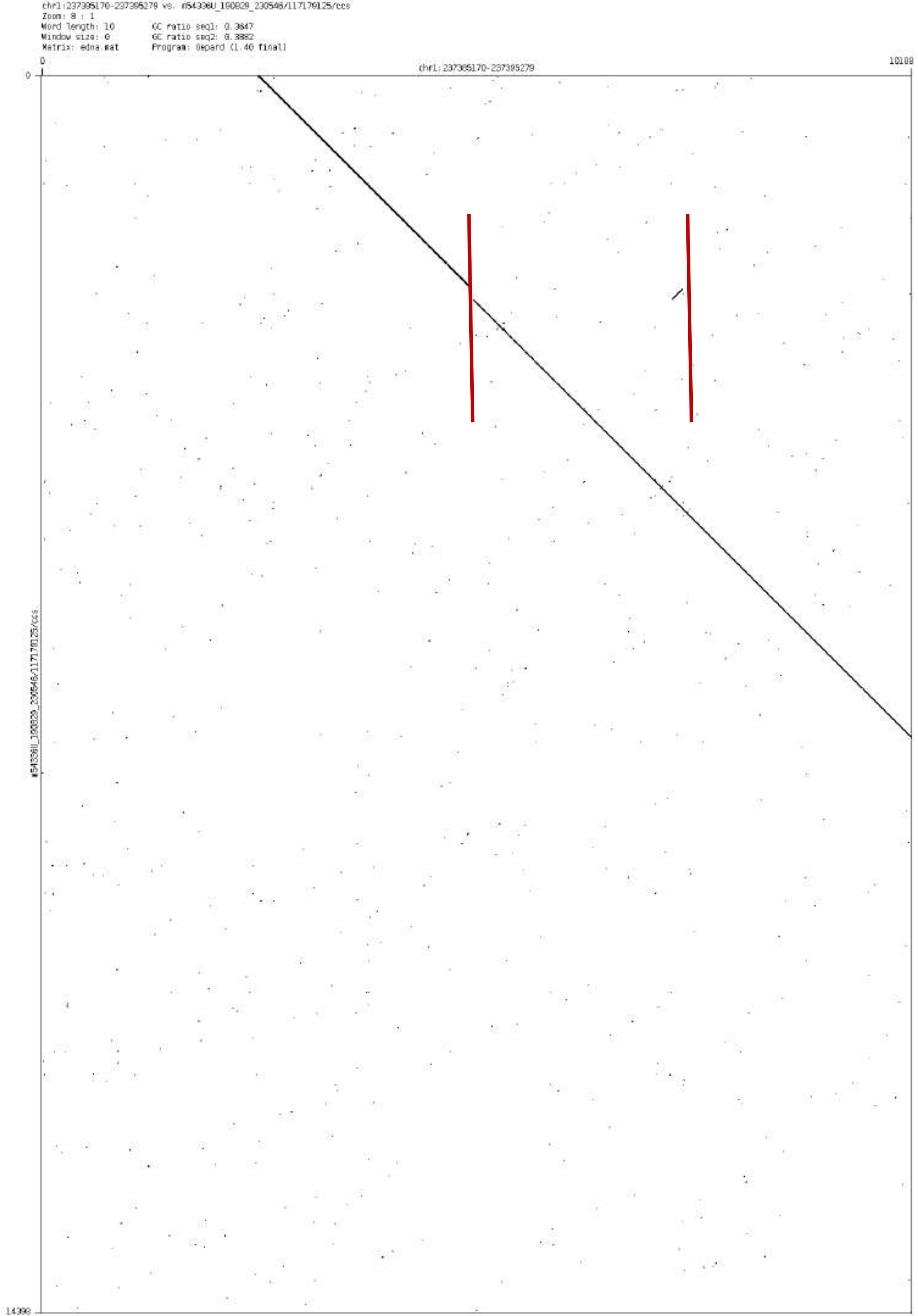

chr1:237395170-237395279 vs. m54336U_190829_230546/117179125/ccs
Zoom: 8 : 1
Word length: 10
GC ratio seq1: 0.3647
Window size: 0
GC ratio seq2: 0.3882
Matrix: edna.mat
Program: Gepard (1.40 final)
0
chr1:237395170-237395279
10108
14390

chr1:239952898-239953380

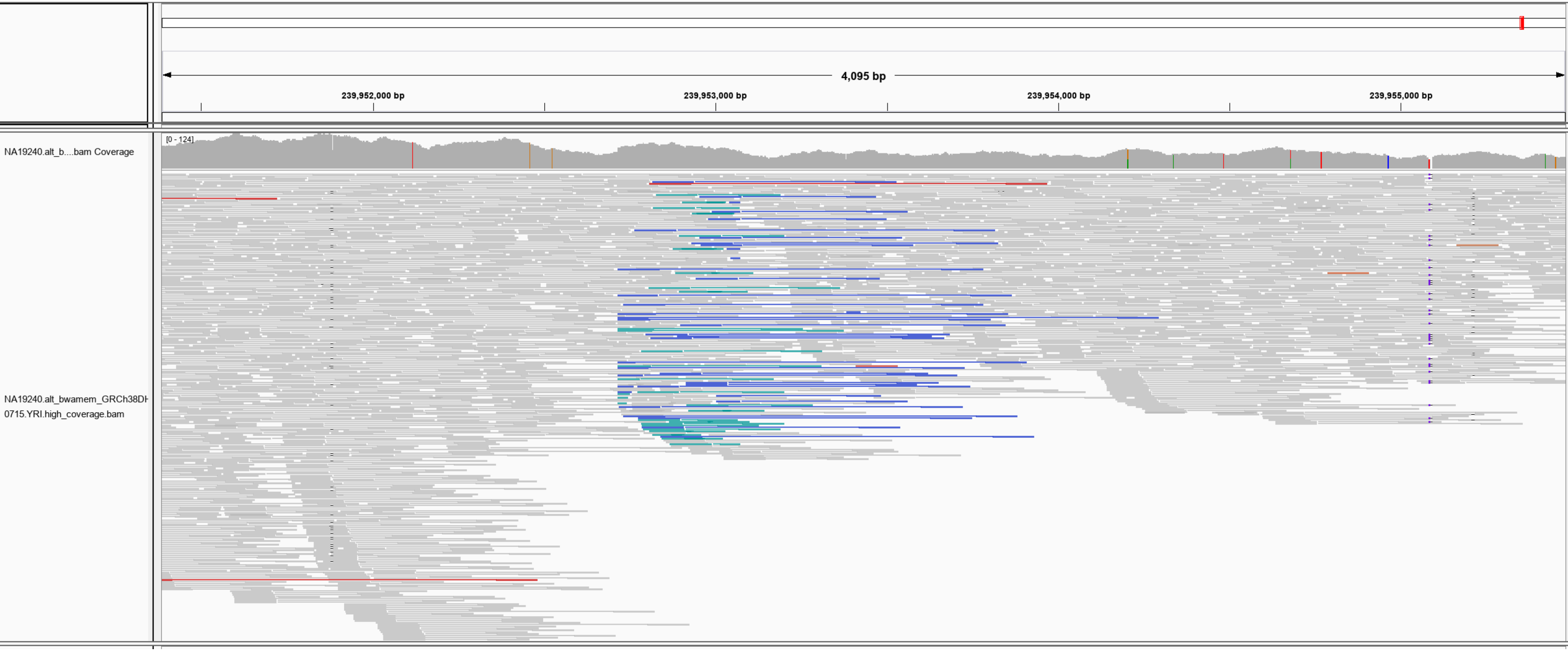

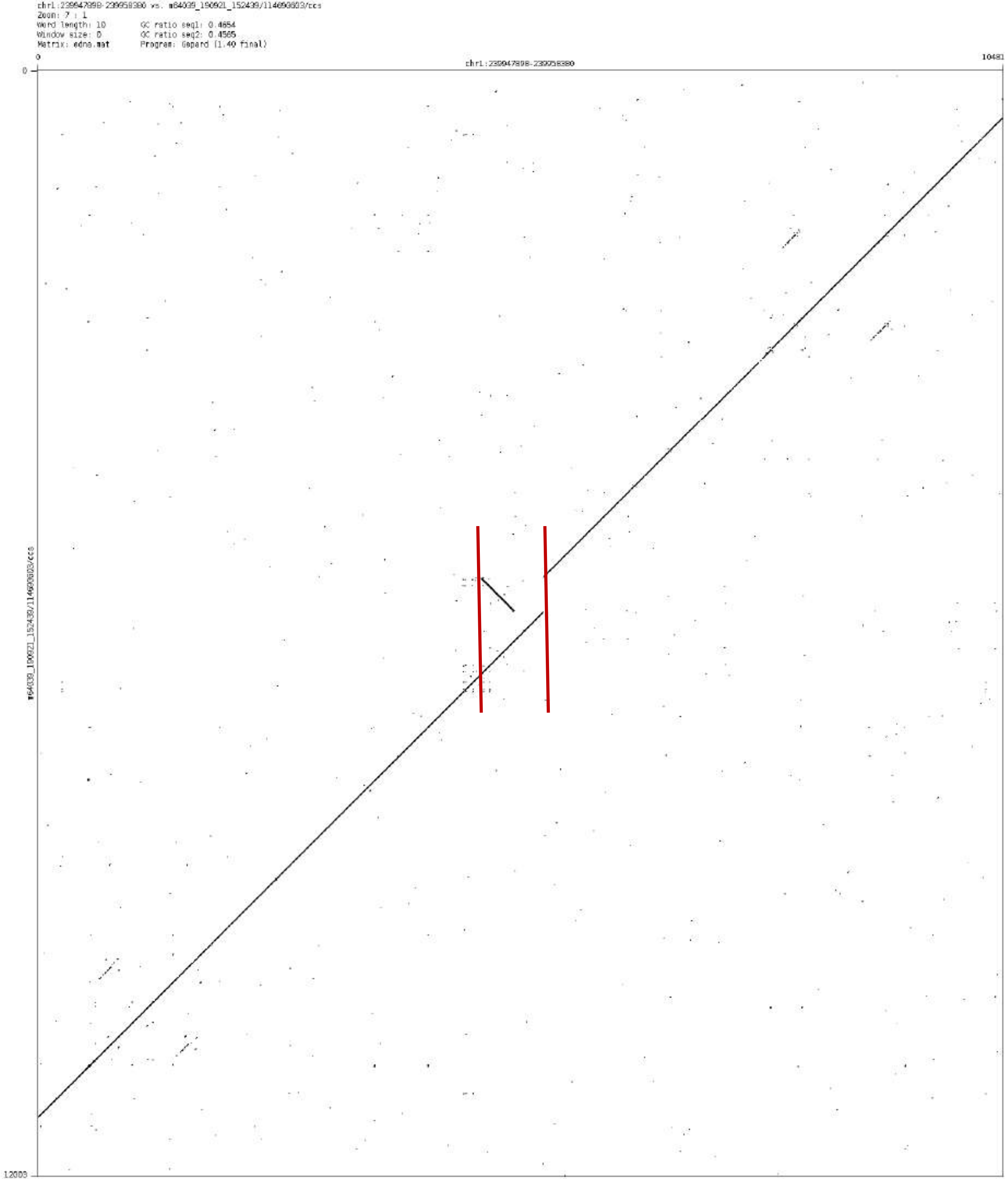
chr1:239947898-239958380 vs. m64039_190921_152439/114690803/ccs
Zoom: 7 : 1
Word length: 10
Window size: 0
Matrix: edna.mat
GC ratio seq1: 0.4654
GC ratio seq2: 0.4565
Program: Gepard (1.40 final)
0
chr1:239947898-239958380
10481
0
m64039_190921_152439/114690803/ccs
12003

chr1:248795414-248799304

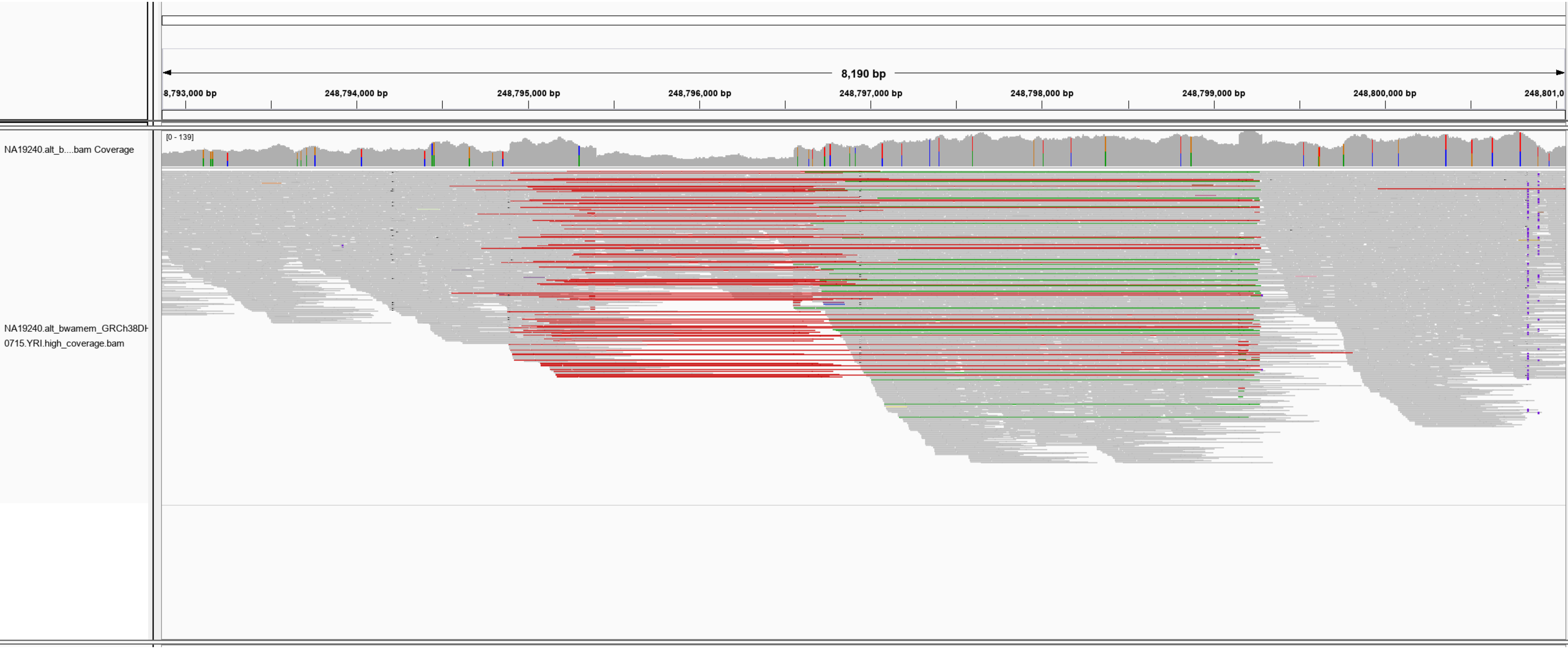

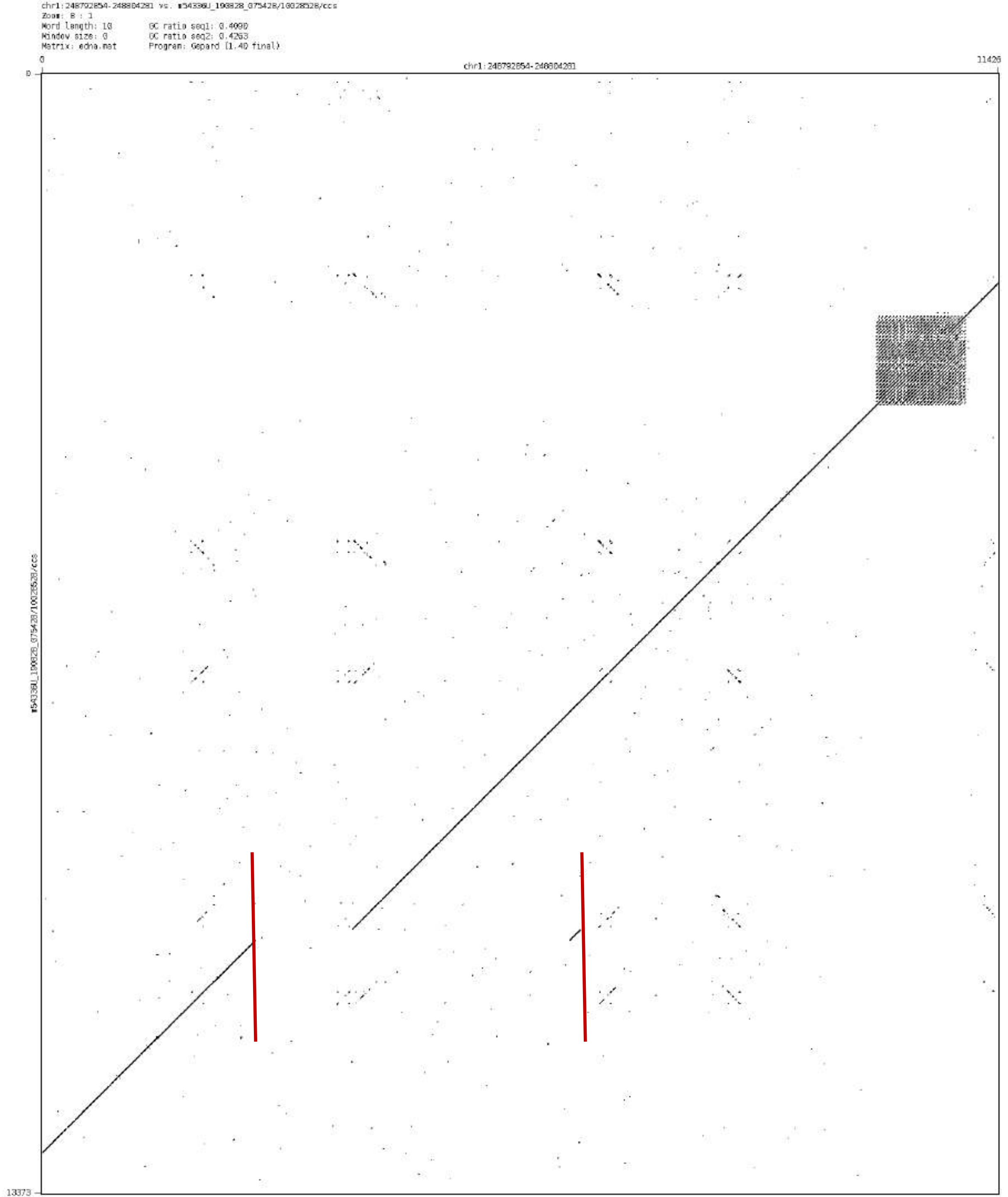
chr1:248792854-248804281 vs. m54336U_190828_075428/10028528/ccs
Zoom: 8 : 1
Word length: 10
Window size: 0
Matrix: edna.mat
GC ratio seq1: 0.4090
GC ratio seq2: 0.4263
Program: Gepard (1.40 final)
0
chr1:248792854-248804281
11426
0
m54336U_190828_075428/10028528/ccs
13373

### chr2:16225125-16226710

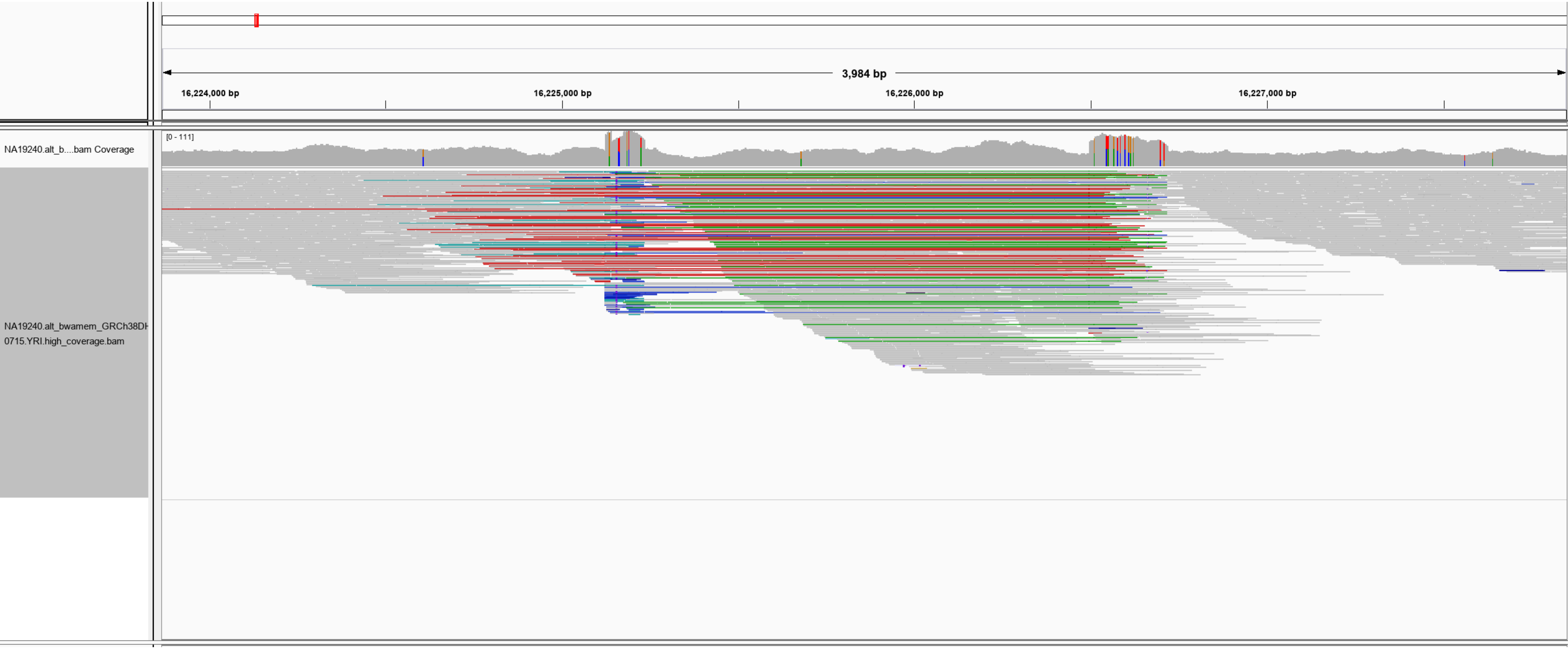

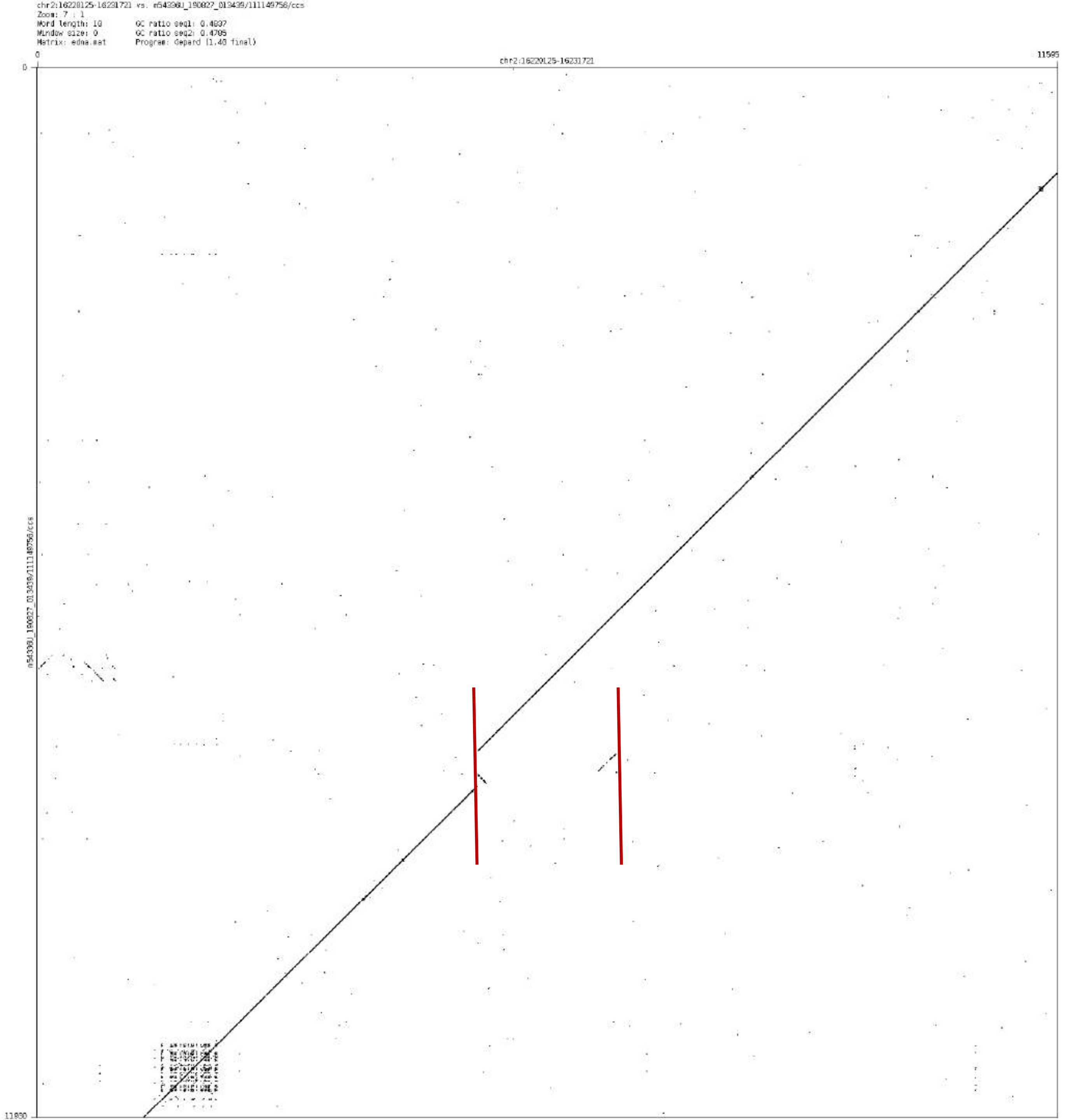
chr2:16220125-16231721 vs. m54398U_190827_013439/111149756/ccs
Zoom: 7 : 1
Word length: 10
Window size: 0
Matrix: edna.mat
GC ratio seq1: 0.4837
GC ratio seq2: 0.4785
Program: depard (1.40 final)
0
chr2:16220125-16231721
11595
0
m54398U_190827_013439/111149756/ccs
11930

chr2:50538785-50539113

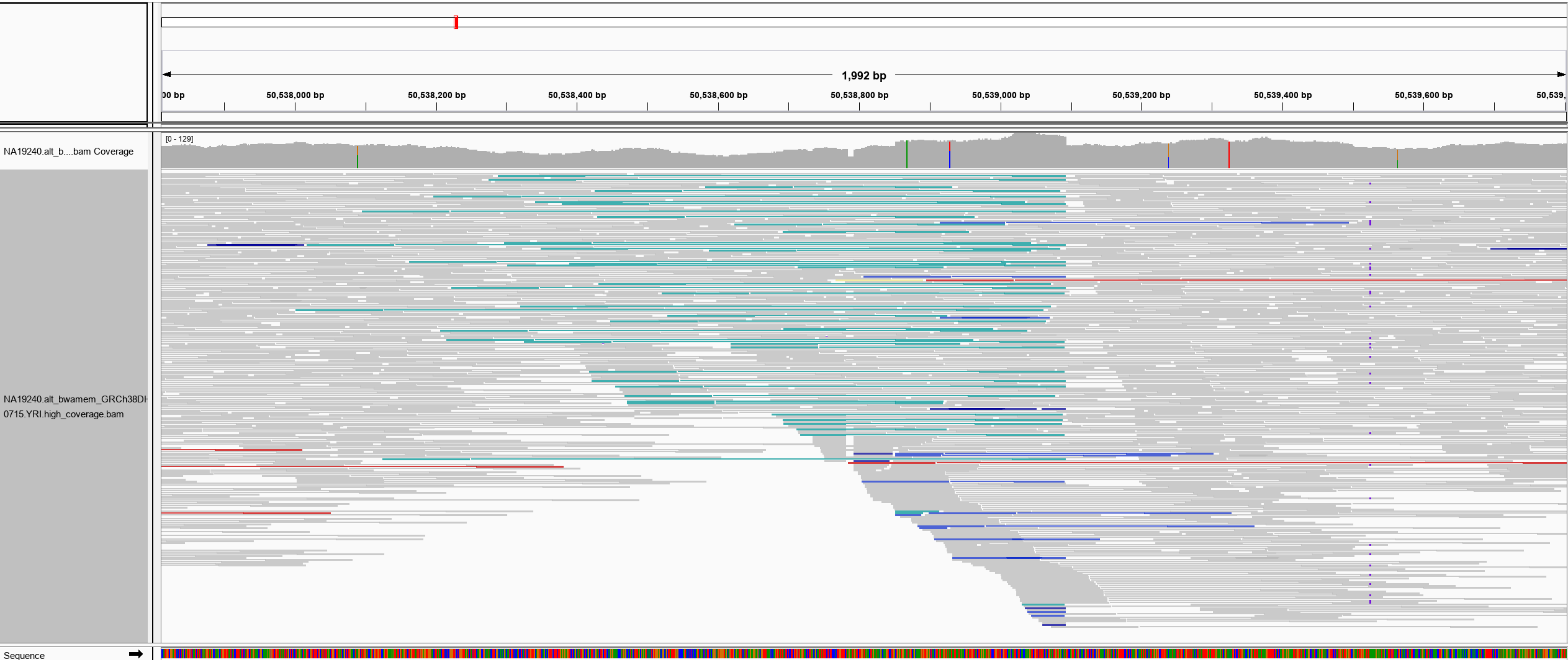

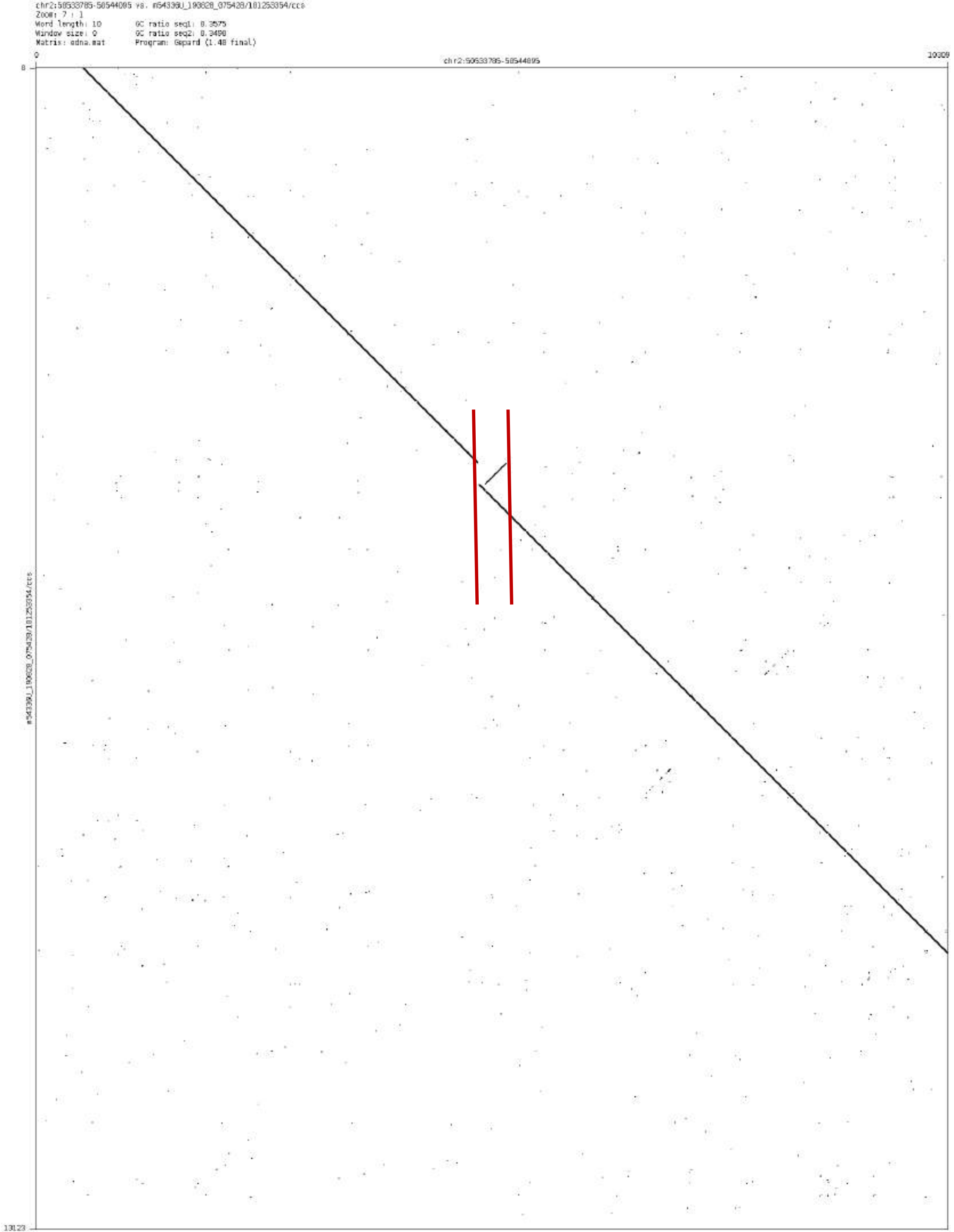

chr2:58533785-58544095 vs. m54336U_190828_075428/101253354/ccs
Zoom: 7 : 1
Word length: 10
Window size: 0
Matrix: edna.mat
GC ratio seq1: 0.3575
GC ratio seq2: 0.3490
Program: Gepard (1.40 final)
chr2:58533785-58544095
10309
13123

### chr2:124294182-124295678

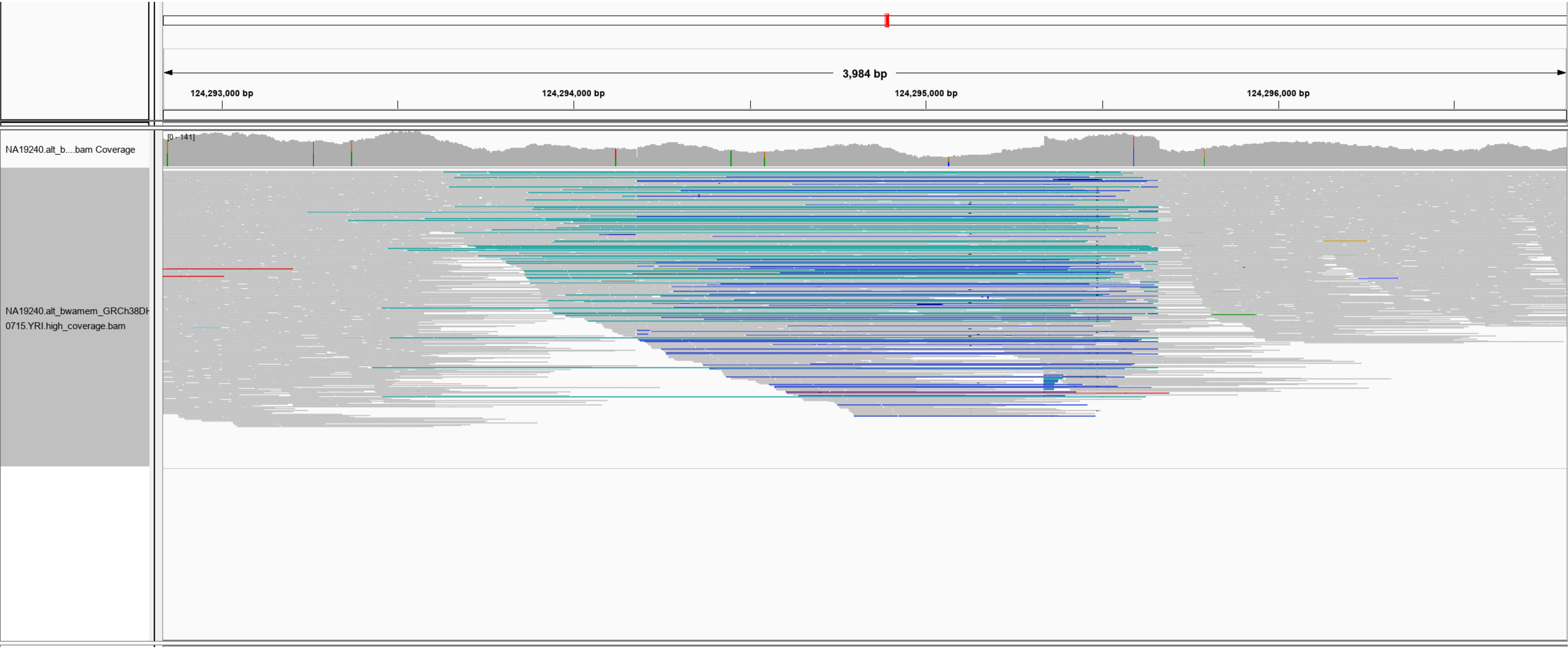

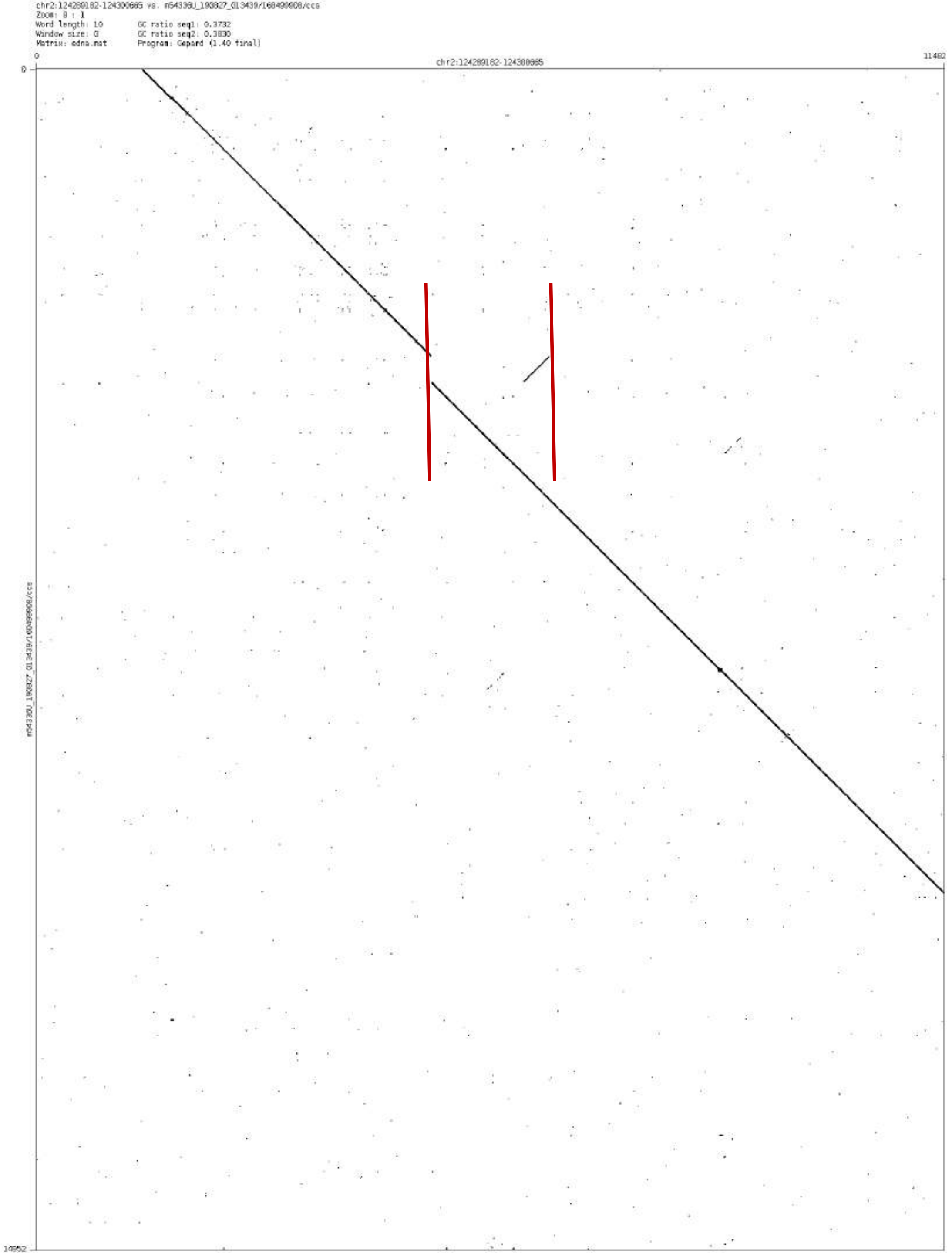

Zoom: 8 : 1
Word length: 10
Window size: 0
Matrix: edna.mat
GC ratio seq1: 0.3732
GC ratio seq2: 0.3830
Program: Gepard (1.40 final)
0
11482
14952

chr2: 129791792-129801213

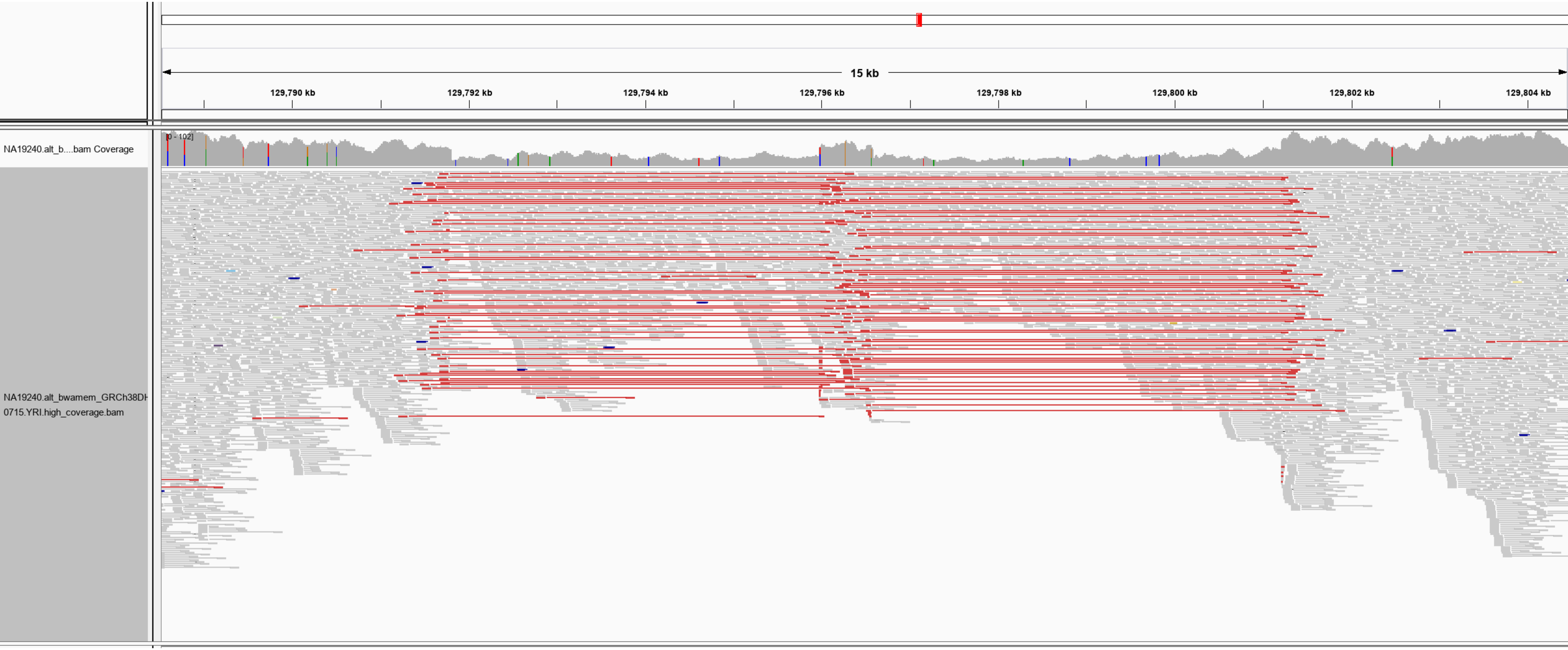

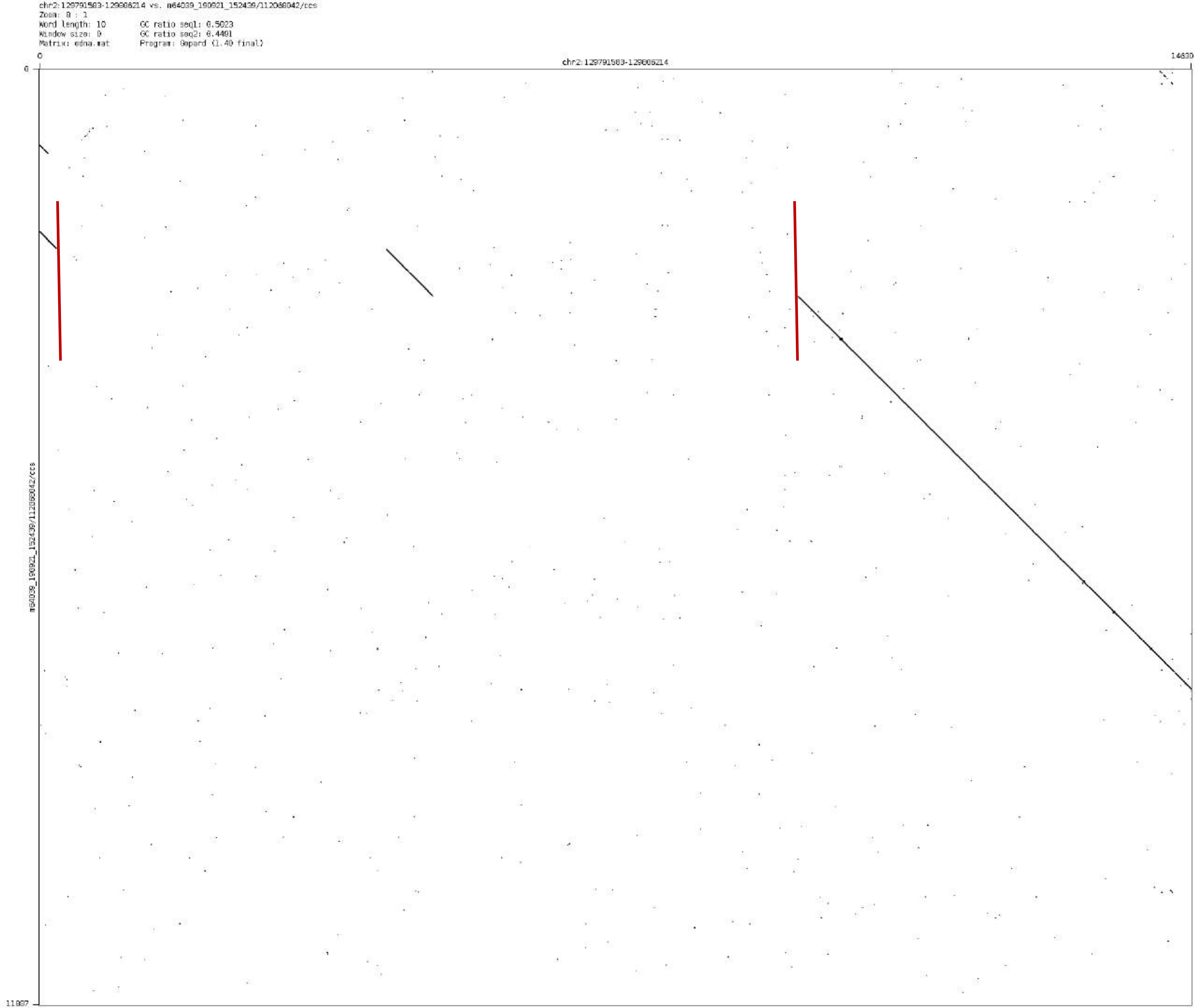
chr2:129791583-129806214 vs. m64039_190921_152439/112088042/ccs
Zoom: 8 : 1
Word length: 10
GC ratio seq1: 0.5023
Window size: 0
GC ratio seq2: 0.4491
Matrix: edna.mat
Program: Gepard (1.40 final)
0
14630
chr2:129791583-129806214
0
11887
m64039_190921_152439/112088042/ccs

chr2:143314745-143316563

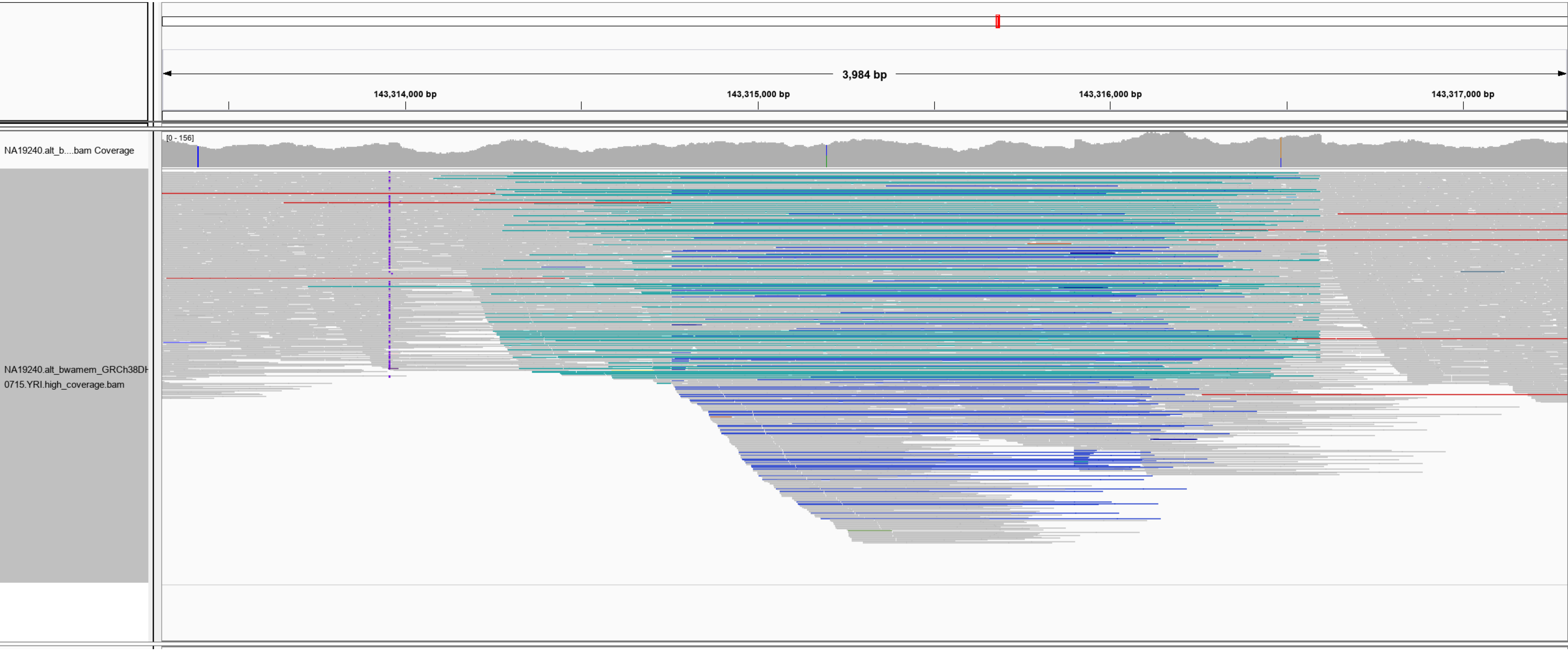

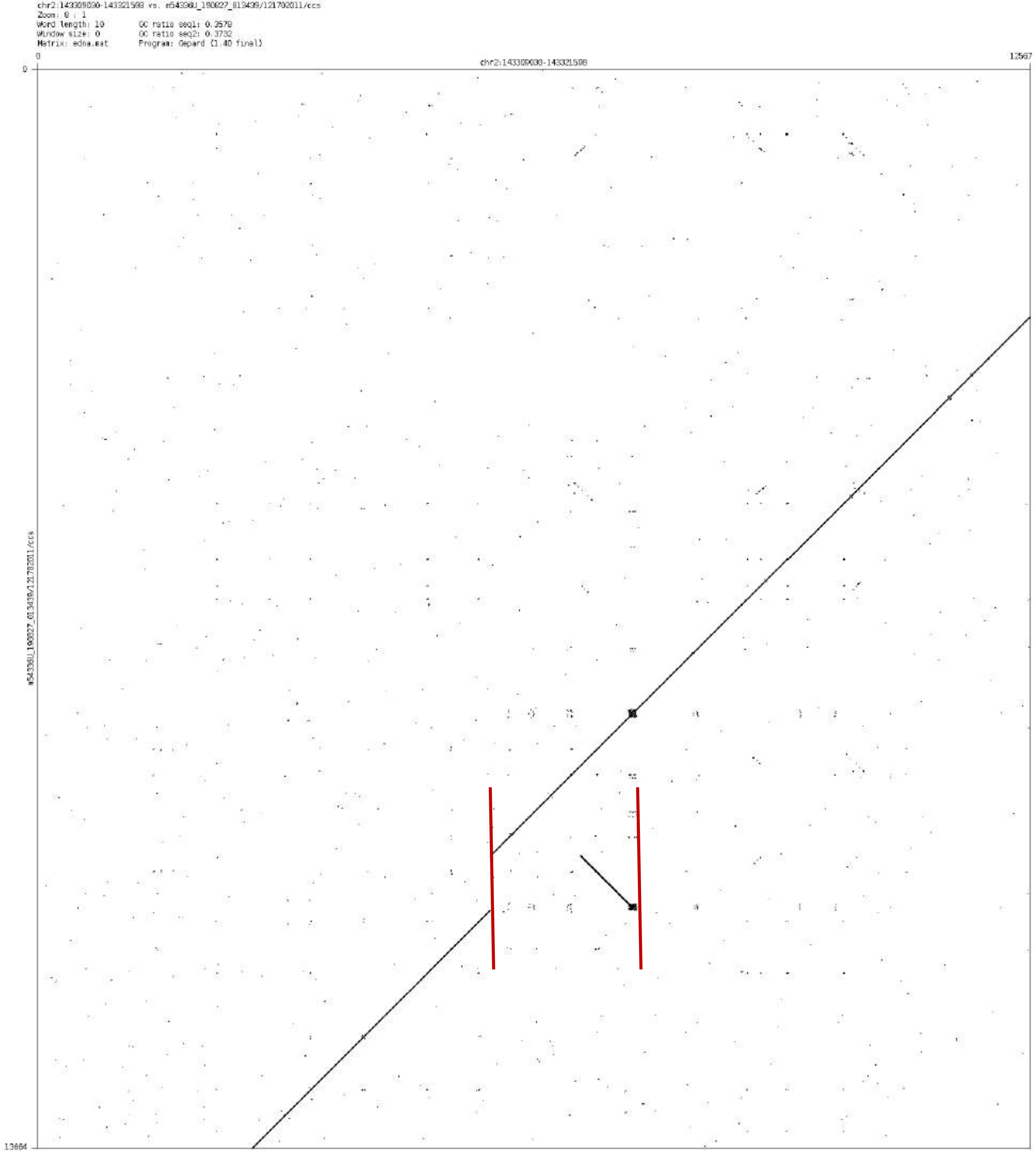

chr2:143309030-143321598 vs. m54336U_190827_013439/121702011/ccs
Zoom: 0 : 1
Word length: 10
Window size: 0
Matrix: edna.mat
GC ratio seq1: 0.3578
GC ratio seq2: 0.3732
Program: Gepard (1.40 final)
0
chr2:143309030-143321598
12567
0
13664

chr2:156813519-156815219

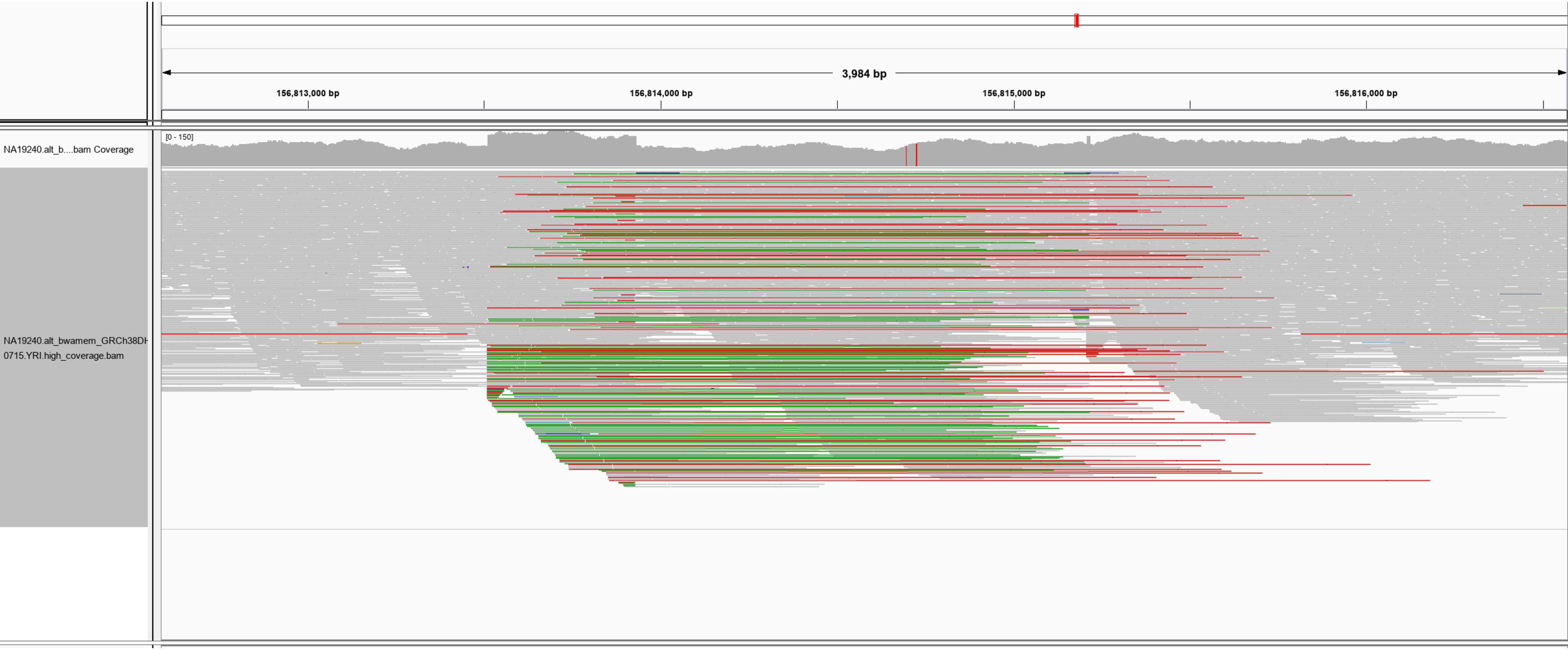

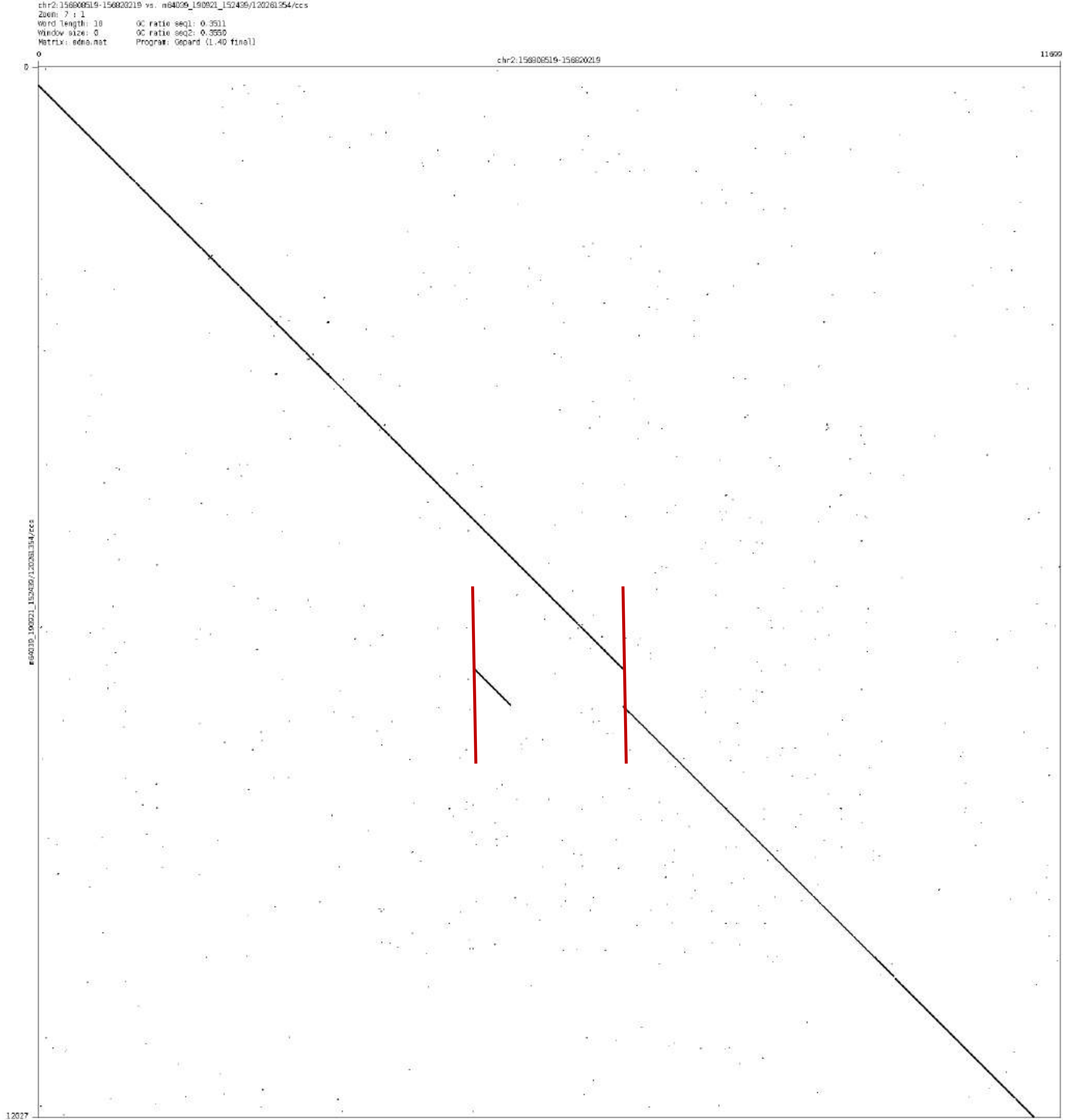

chr2:156808519-156820219 vs. m64039_190921_152439/120261354/ccs
Zoom: 7 : 1
Word length: 10
GC ratio seq1: 0.3511
Window size: 0
GC ratio seq2: 0.3550
Matrix: edna.mat
Program: Gepard (1.40 final)
0
chr2:156808519-156820219
11699
0
m64039_190921_152439/120261354/ccs
12027

### chr2:216225220-216228380

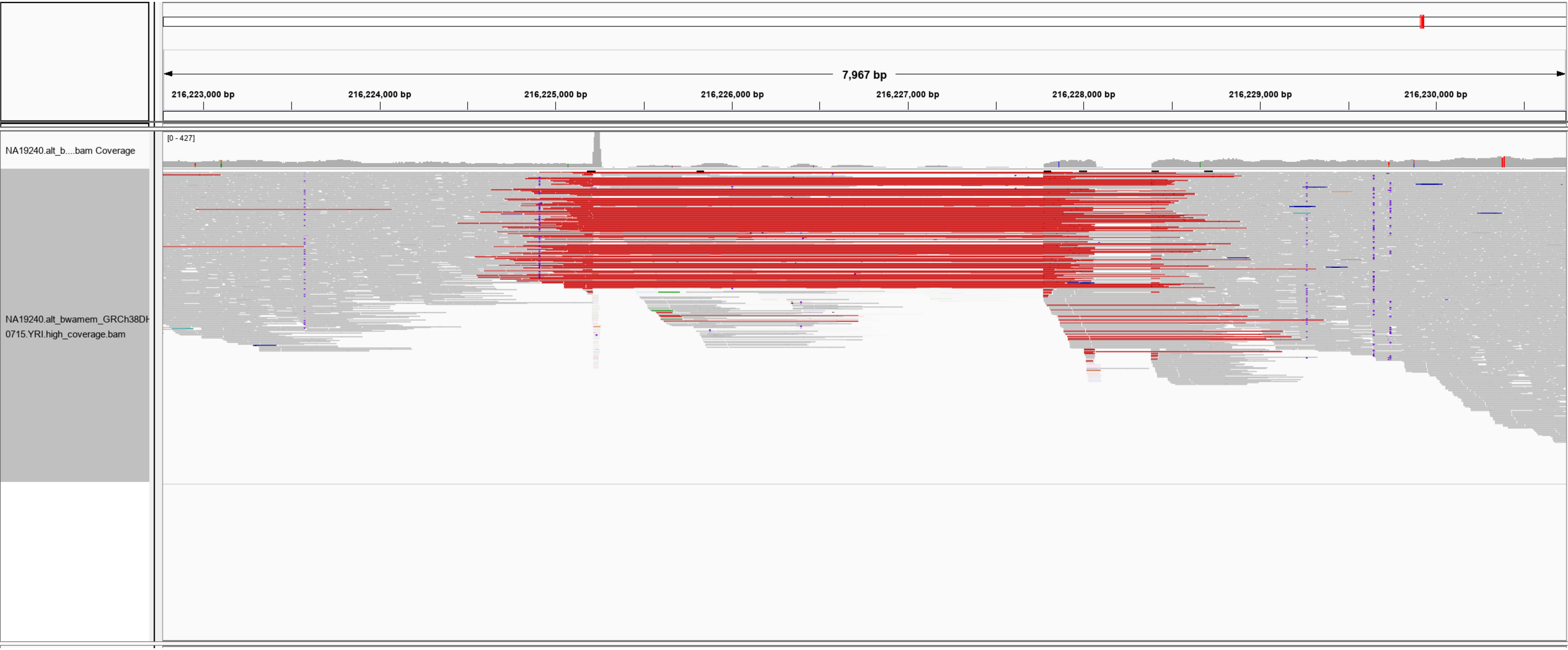

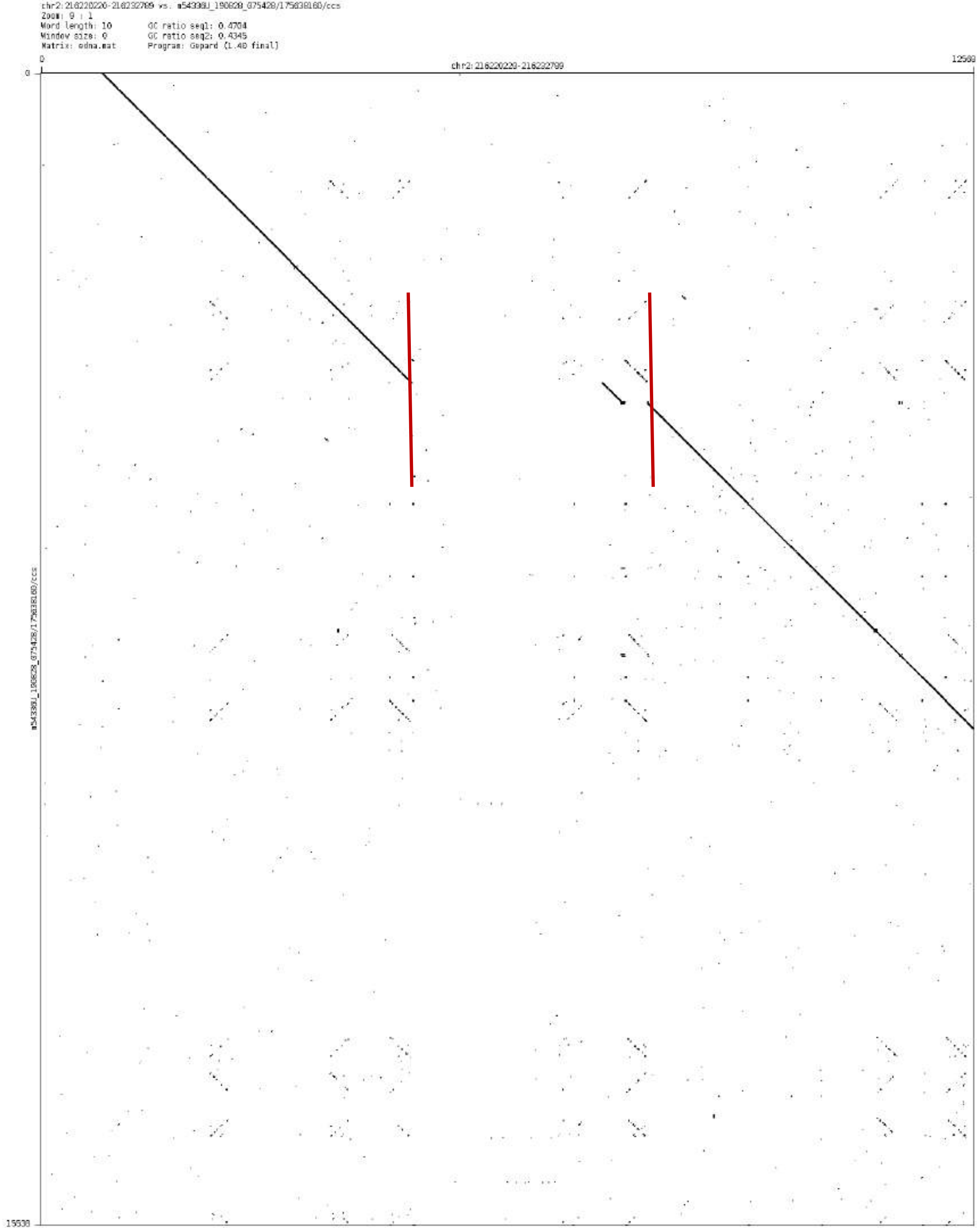
chr2:216220220-216232789 vs. m54338U_190828_075428/175838160/ccs
Zoom: 9 : 1
Word length: 10
Window size: 0
Matrix: edna.mat
GC ratio seq1: 0.4704
GC ratio seq2: 0.4345
Program: Gepard (1.40 final)
0
chr2:216220220-216232789
12569
0
m54338U_190828_075428/175838160/ccs
15538

chr3:95746925-95752145

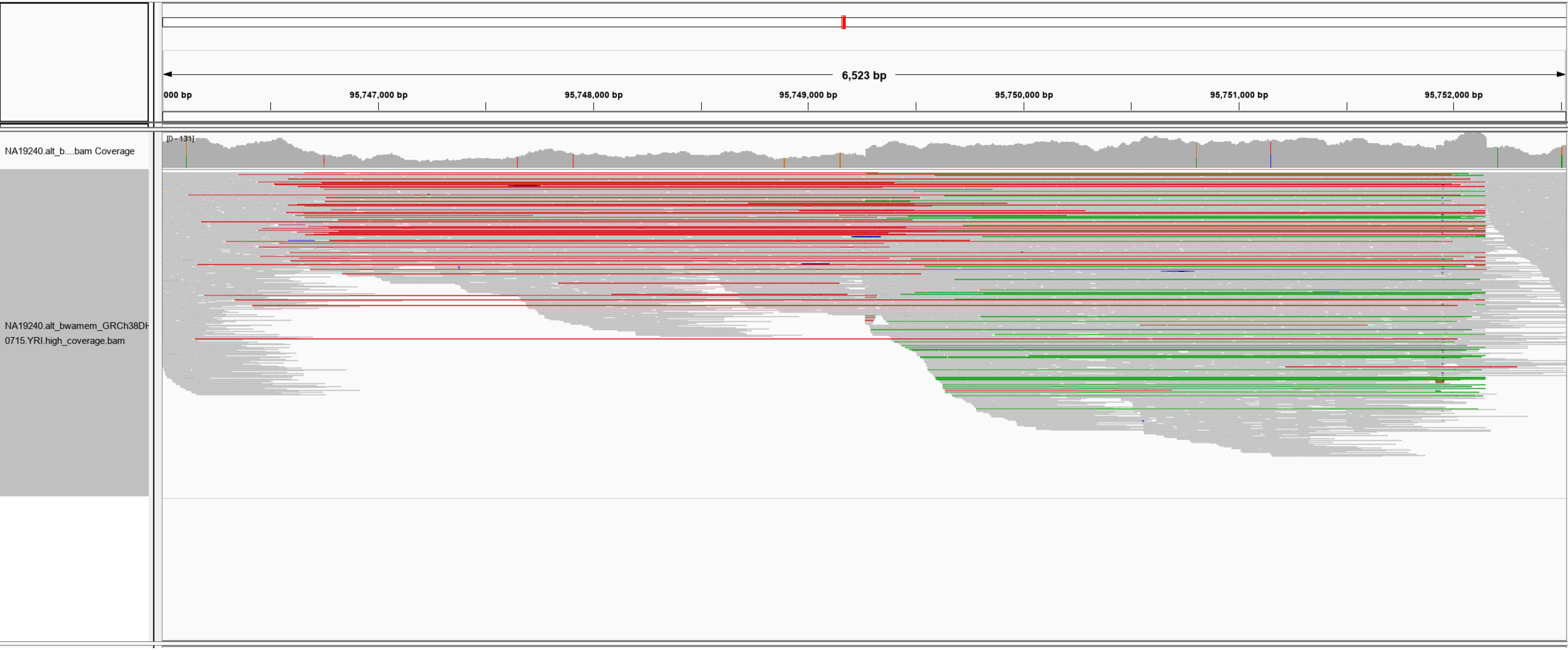

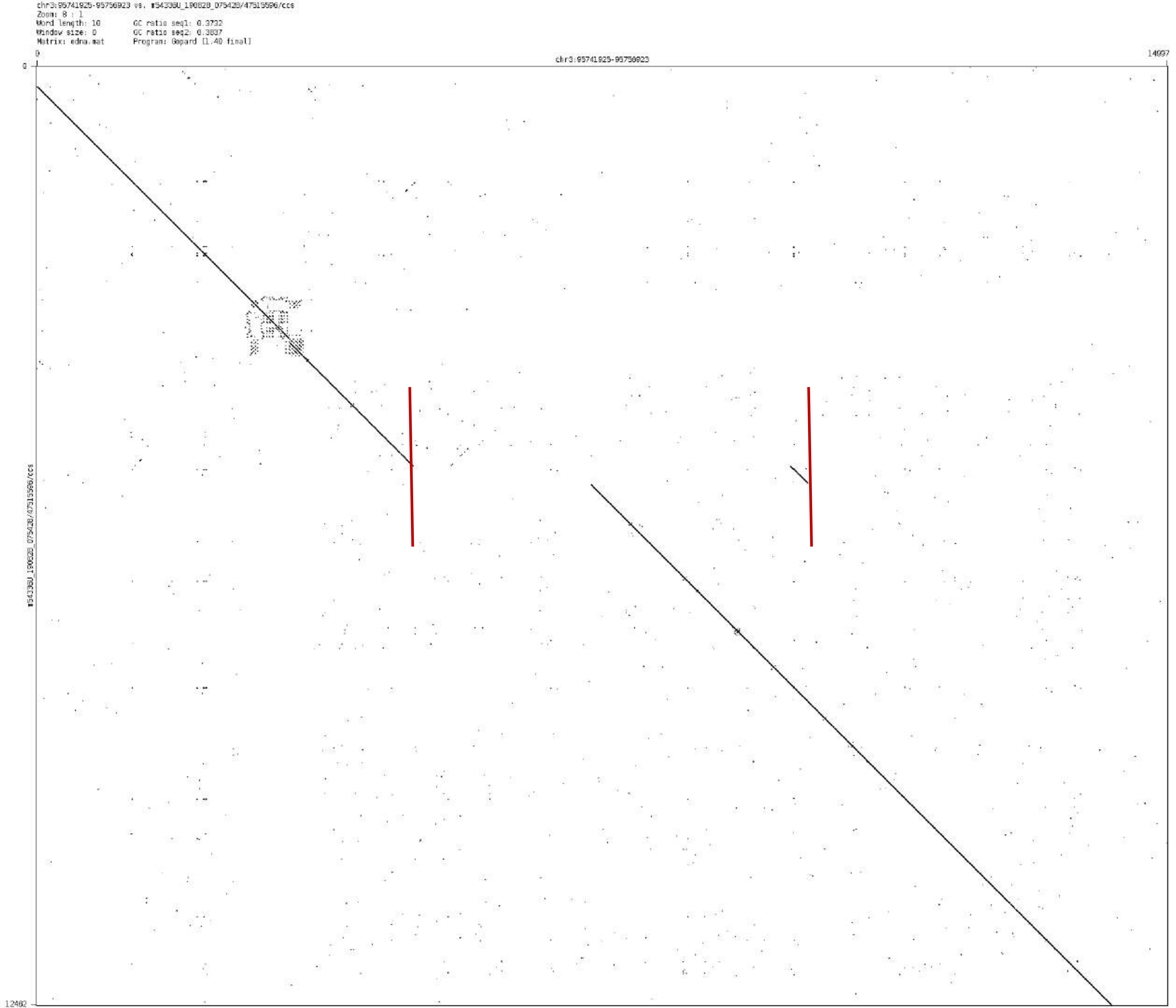
chr3:95741925-95756923 vs. m54336U_190828_075428/47515596/ccs
Zoom: 8 : 1
Word length: 10
GC ratio seq1: 0.3732
Window size: 0
GC ratio seq2: 0.3837
Matrix: edna.mat
Program: Gepard [1.40 final]
0
14997
chr3:95741925-95756923
0
12462
m54336U_190828_075428/47515596/ccs

### chr3:146667406-146677351

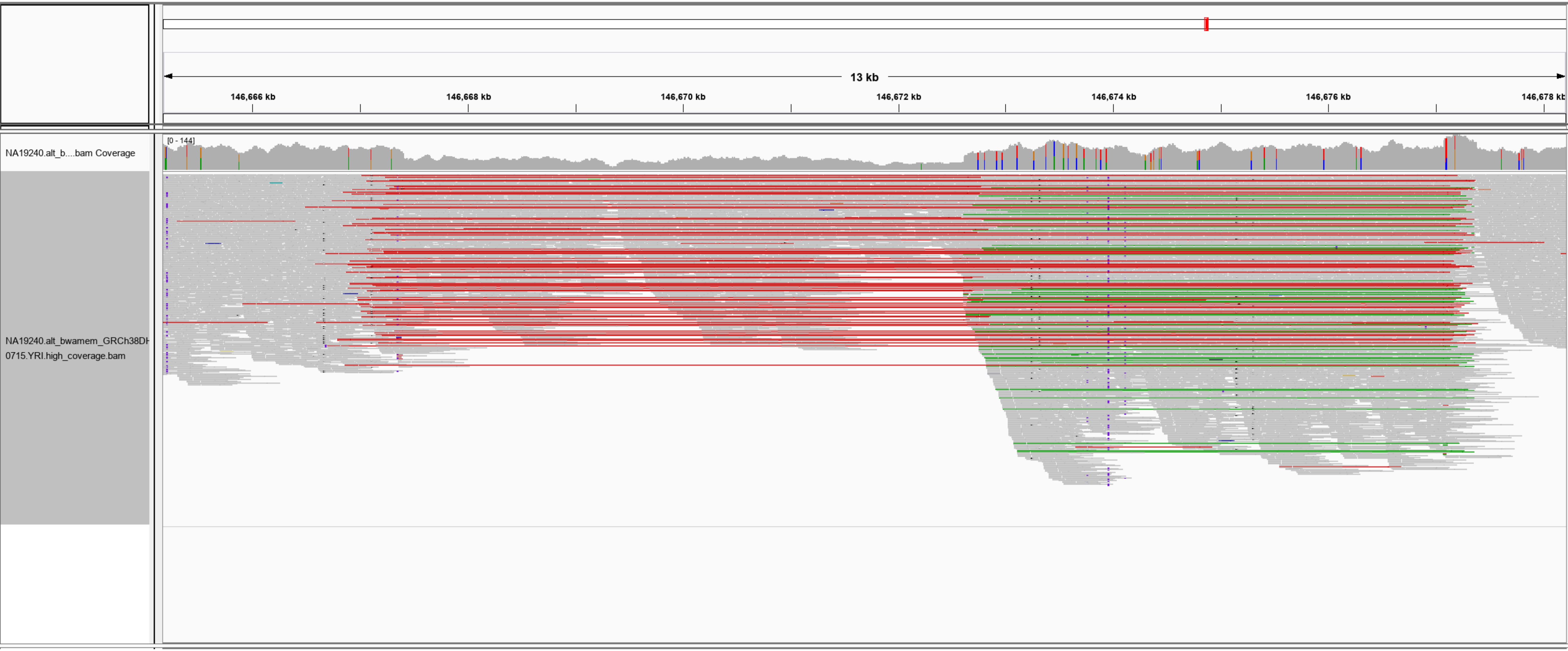

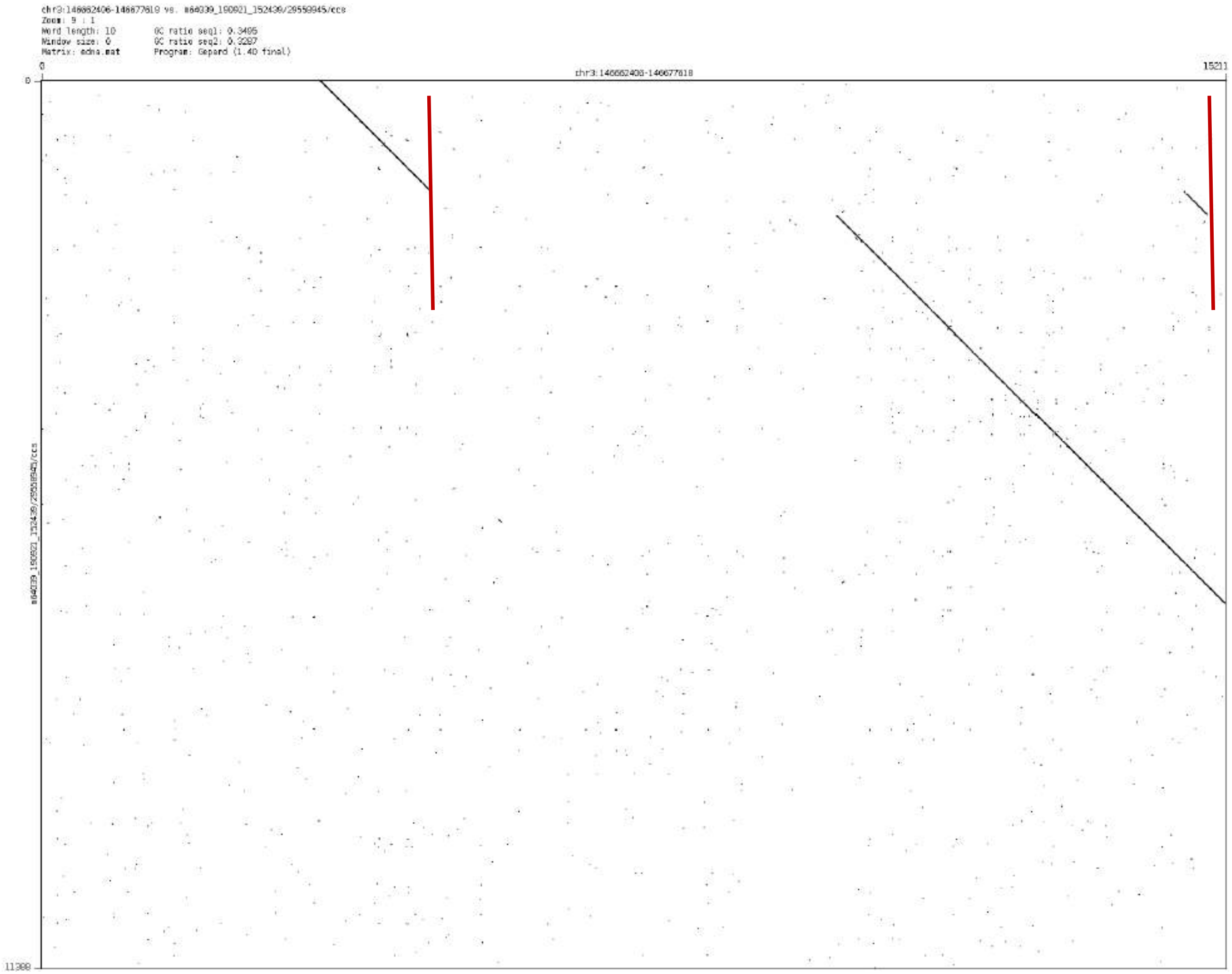

chr3:146662406-146677618 vs. m64039_190921_152439/29559945/ccs
Zoom: 9 : 1
Word length: 10
Window size: 0
Matrix: edna.mat
GC ratio seq1: 0.3405
GC ratio seq2: 0.3287
Program: Gepard (1.40 final)
0
15211
chr3:146662406-146677618
0
11388
m64039_190921_152439/29559945/ccs

chr4: 92646201-92649031

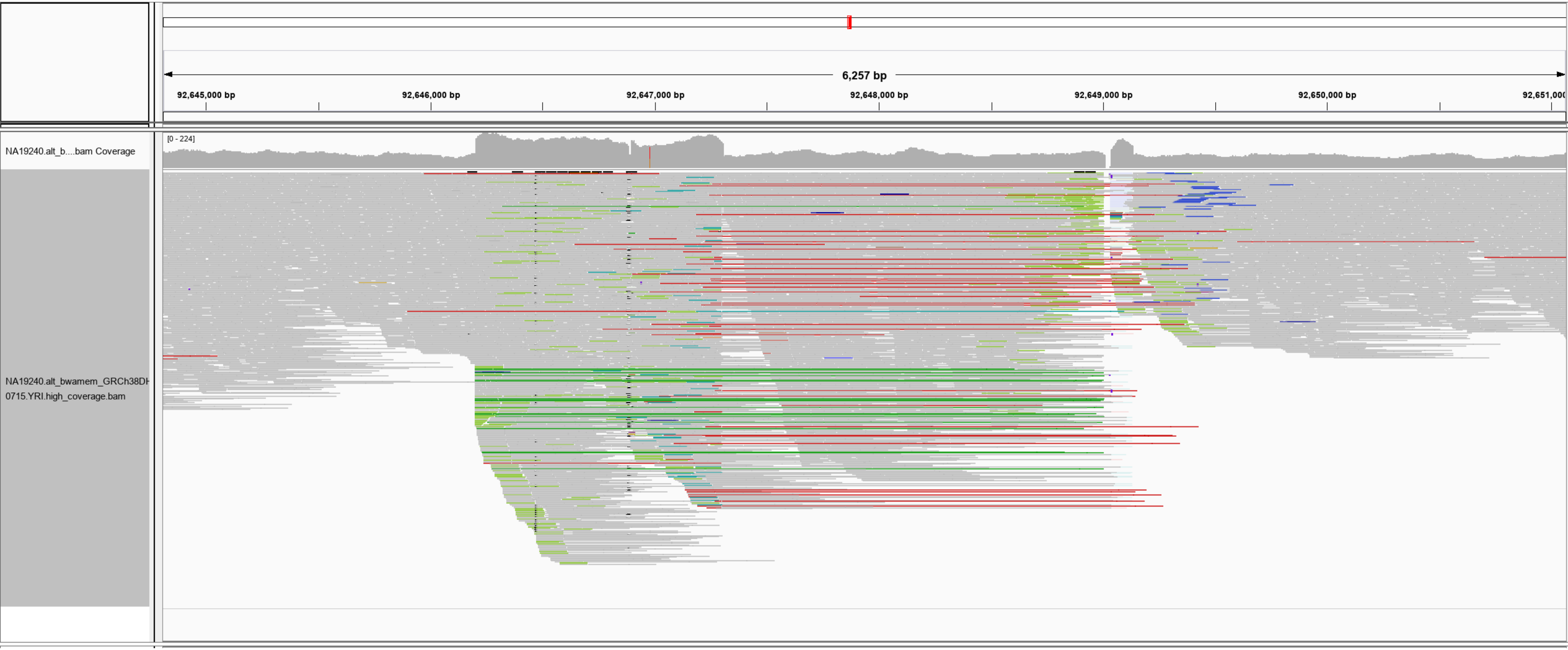

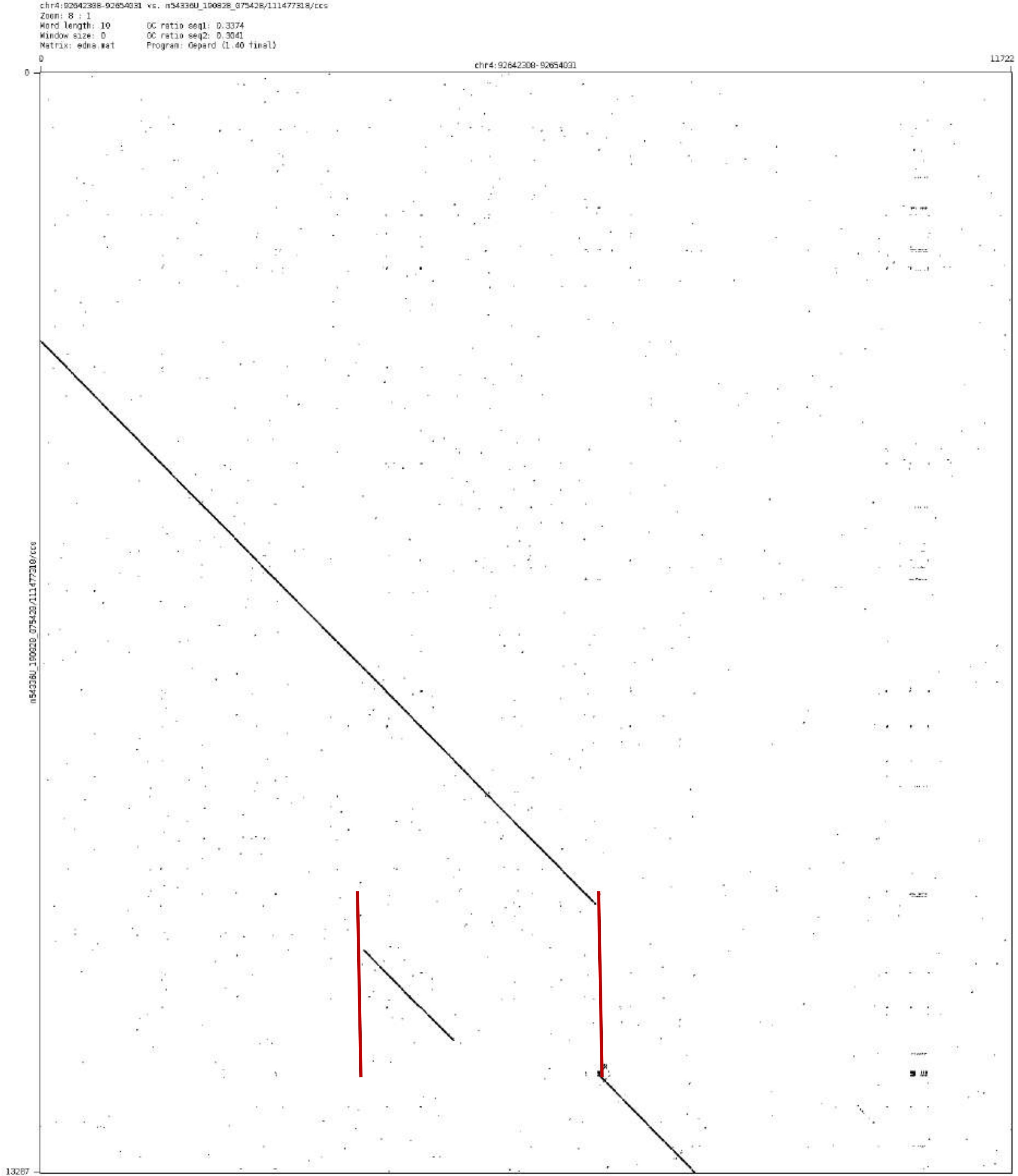

chr4:92642308-92654031 vs. m54336U_190828_075428/111477318/ccs
Zoom: 8 : 1
Word length: 10
Window size: 0
Matrix: edna.mat
GC ratio seq1: 0.3374
GC ratio seq2: 0.3041
Program: Gepard (1.40 final)
0
chr4:92642308-92654031
11722
0
13287
m54336U_190828_075428/111477318/ccs

chr5:25821671-25826900

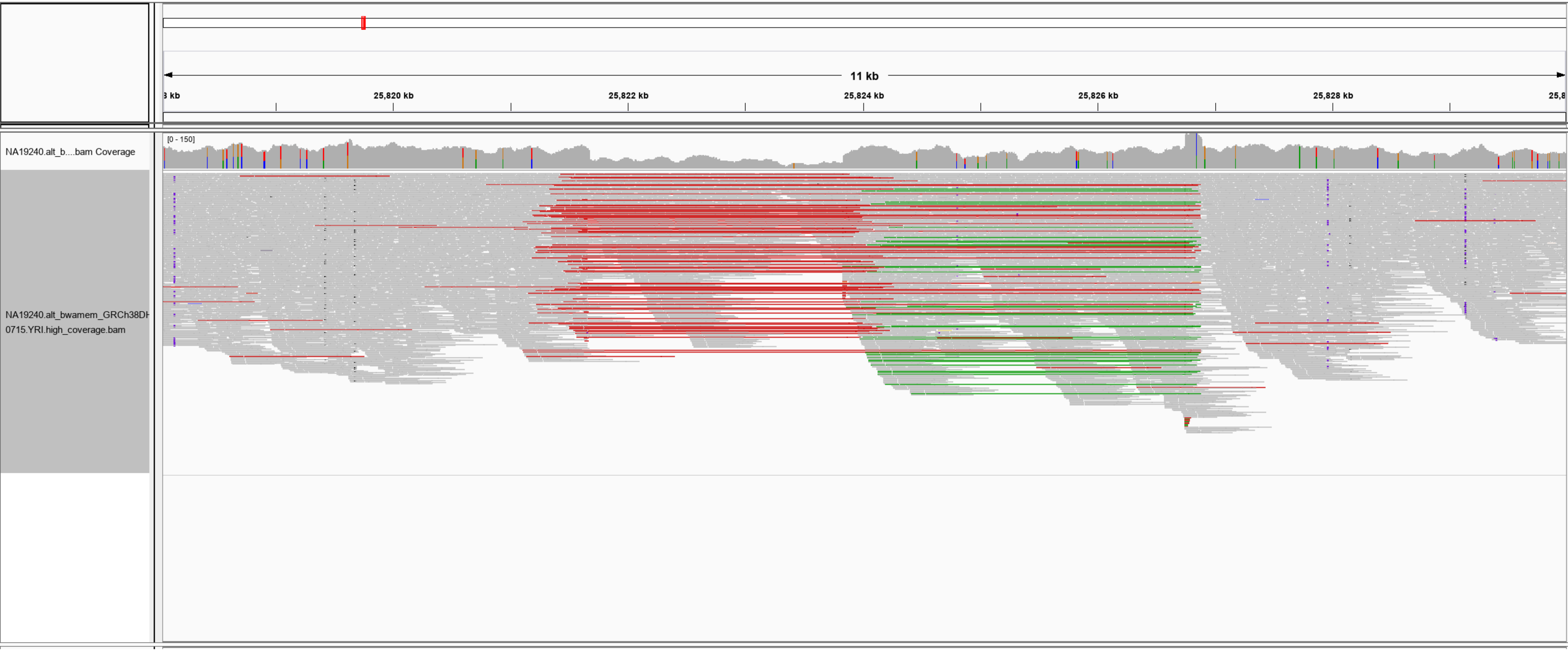

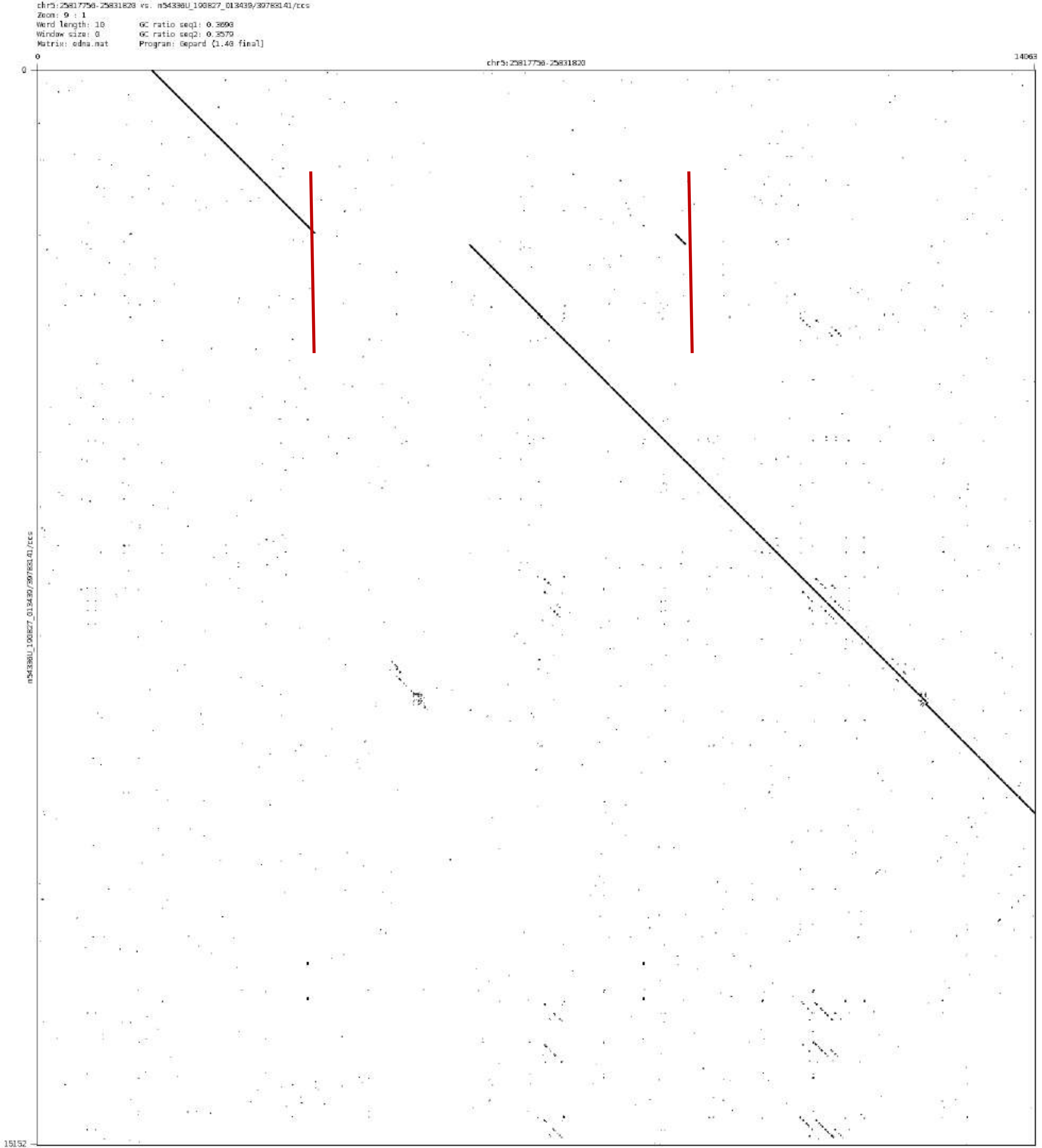
chr5:25817756-25831820 vs. m54336U_190827_013439/39783141/ccs
Zoom: 9 : 1
Word length: 10
Window size: 0
Matrix: edna.mat
GC ratio seq1: 0.3690
GC ratio seq2: 0.3579
Program: Gepard (1.40 final)
0
14063
chr5:25817756-25831820
0
15152
m54336U_190827_013439/39783141/ccs

chr5:114919235-114925727

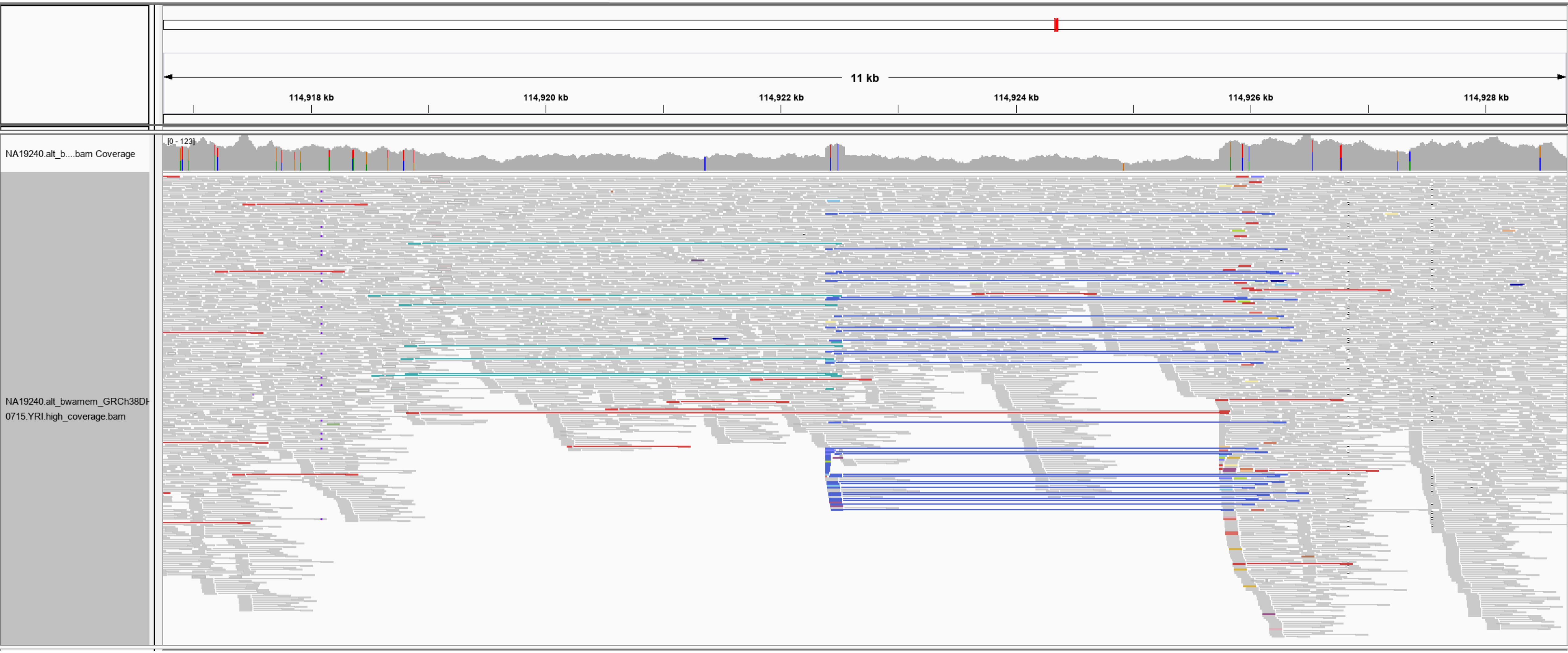

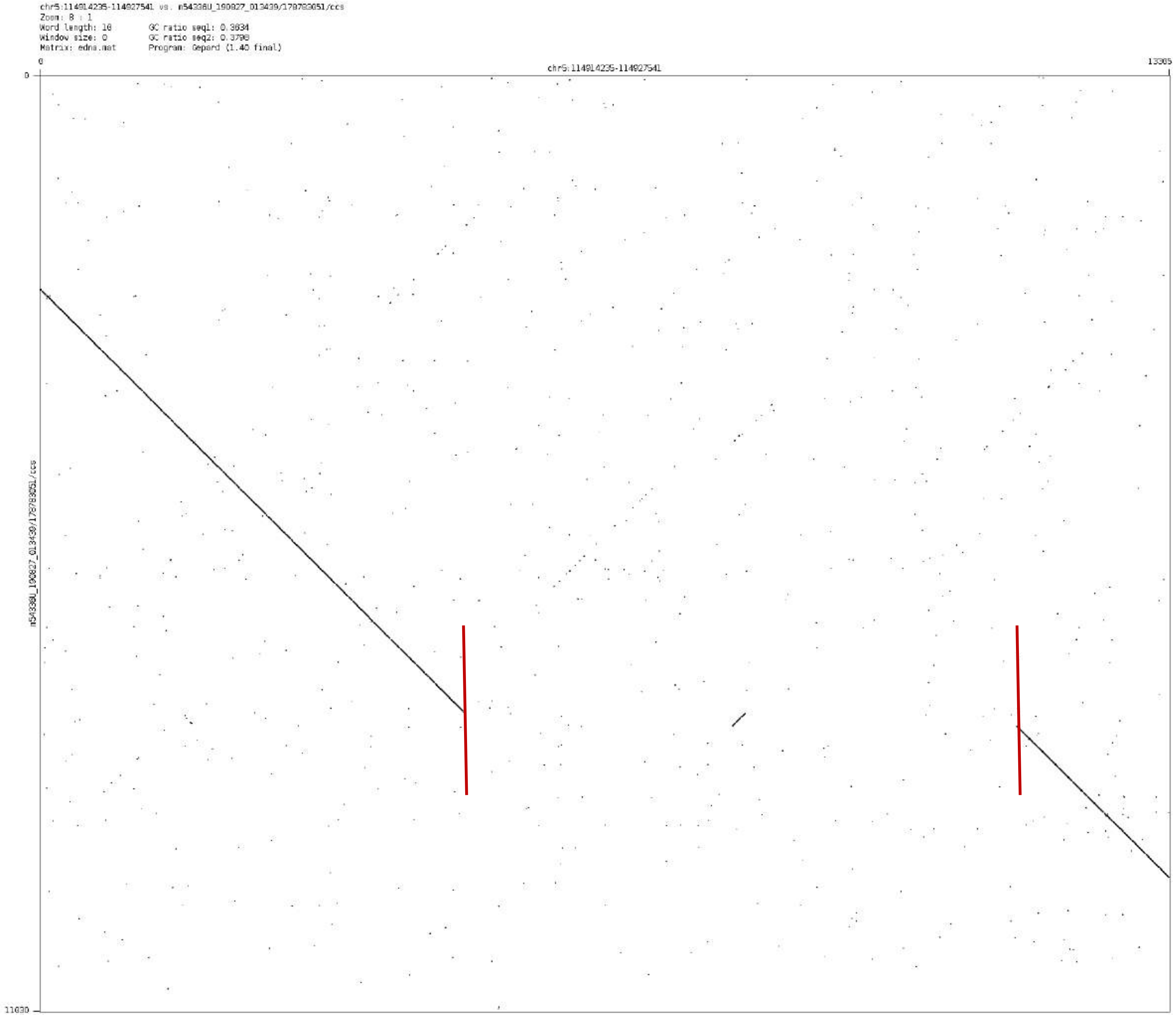

chr5:114914235-114927541 vs. m54336U_190827_013439/178783051/ccs
Zoom: 8 : 1
Word length: 10
Window size: 0
Matrix: edna.mat
GC ratio seq1: 0.3634
GC ratio seq2: 0.3798
Program: Gepard (1.40 final)
0
chr5:114914235-114927541
13305
0
m54336U_190827_013439/178783051/ccs
11030

chr5:116011020-116015362

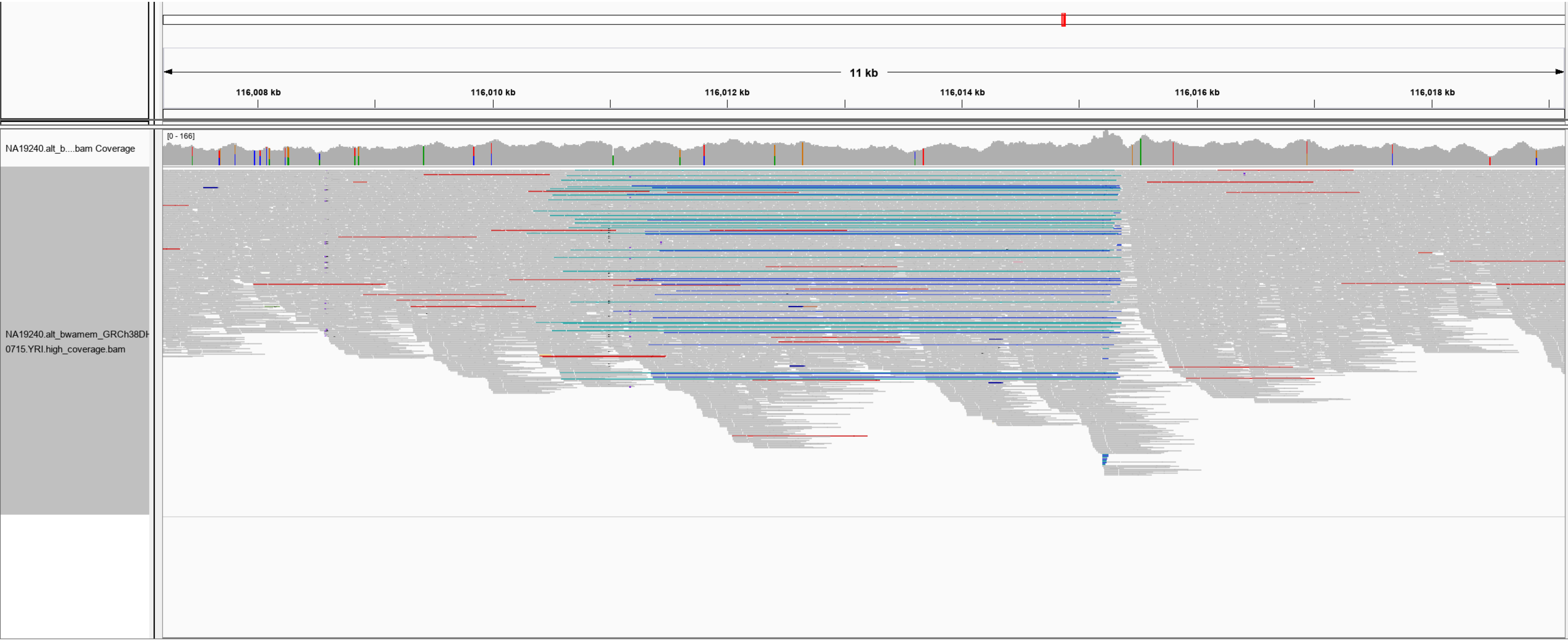

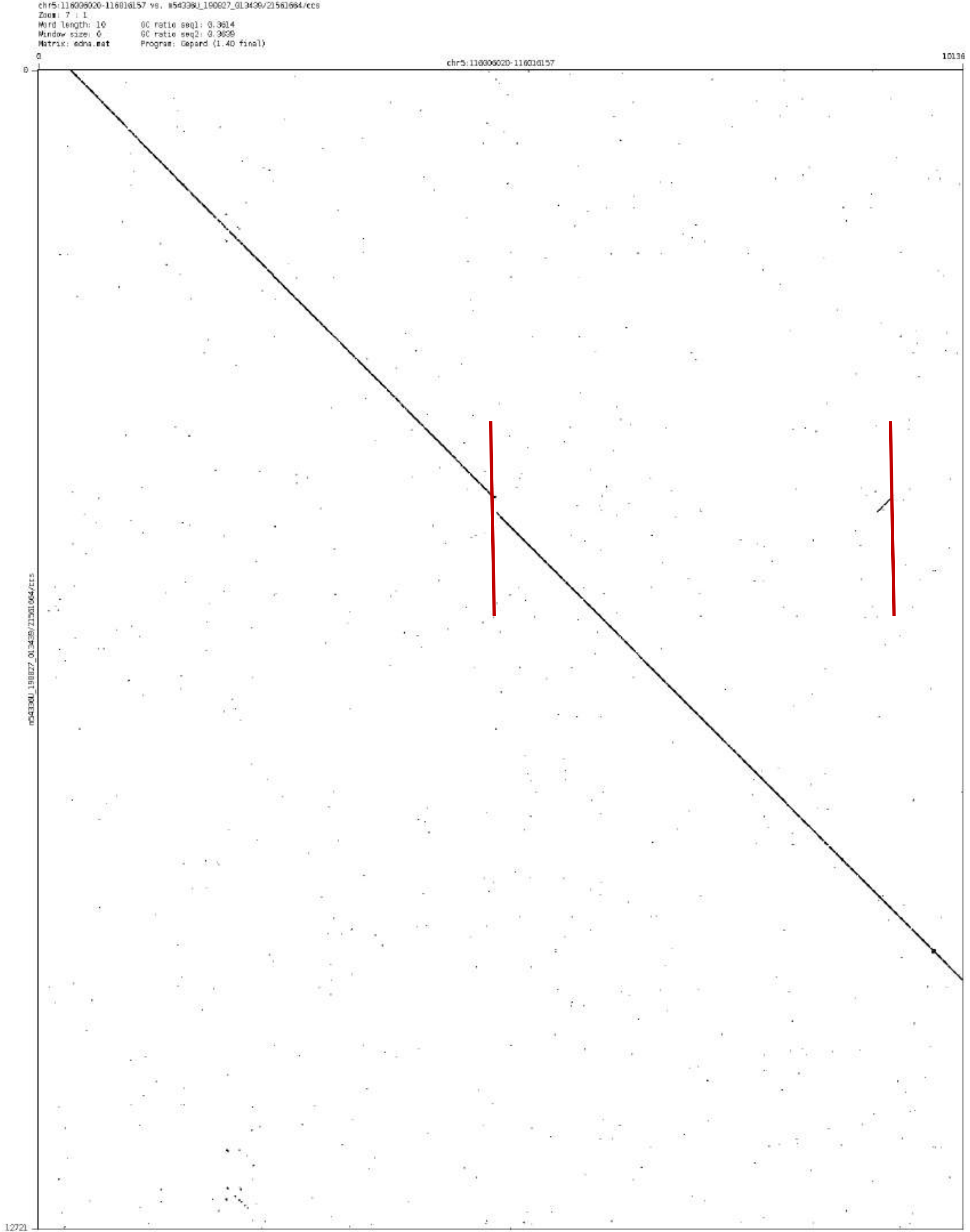
Zoom: 7 : 1
Word length: 10
Window size: 0
Matrix: edna.mat
Program: Gepard (1.40 final)
0
chr5:116006020-116016157
10136
12721

### chr5:144132930-144135491

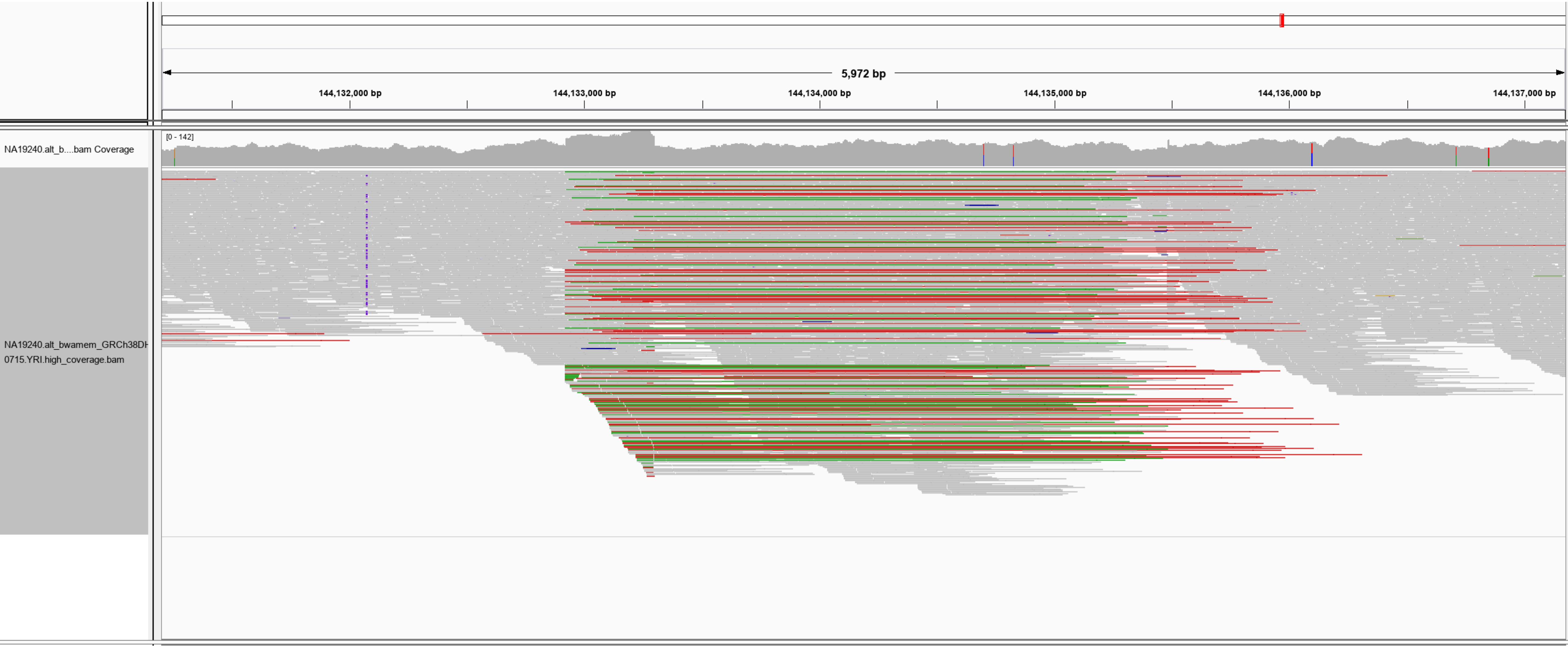

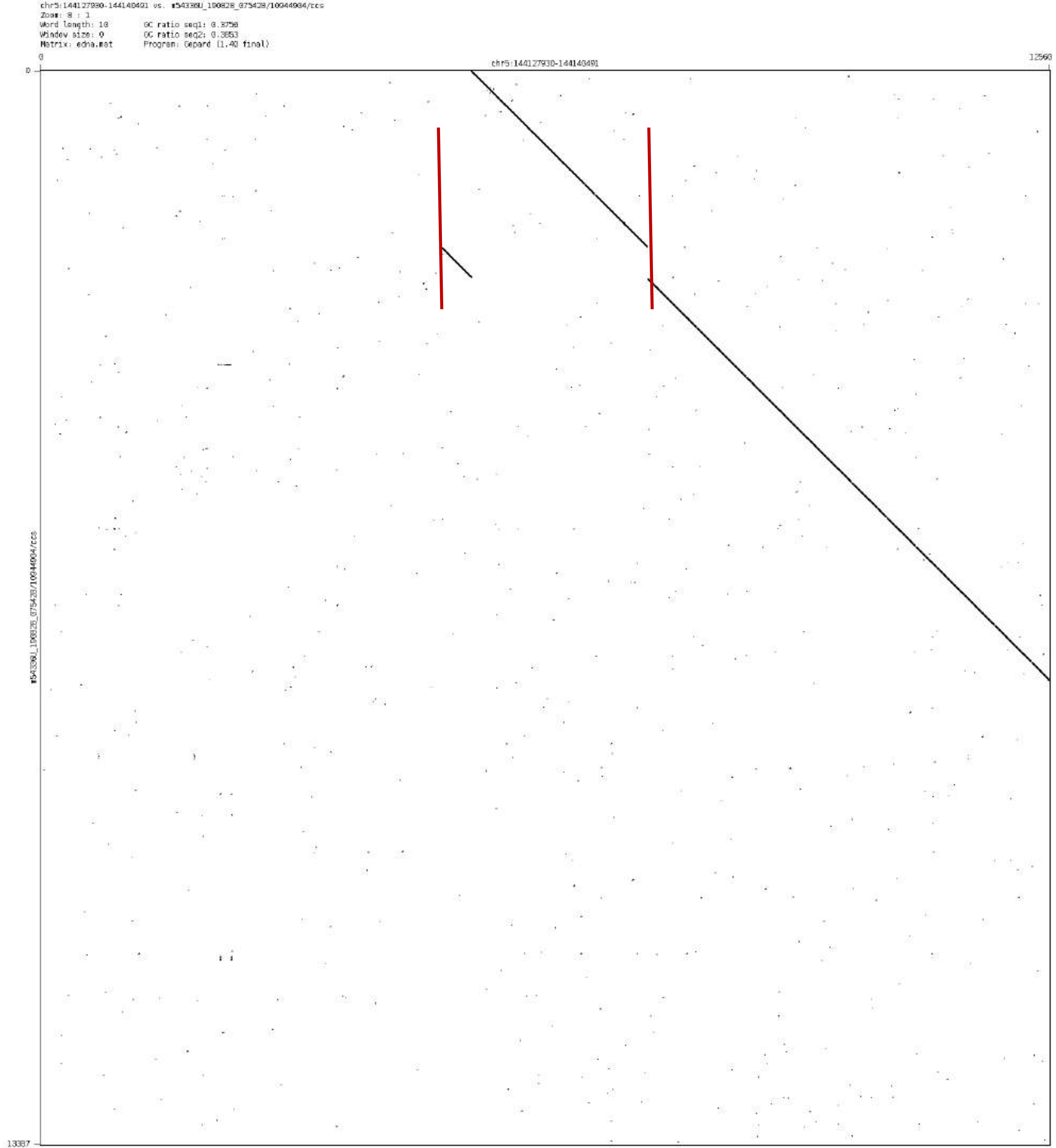
chr5:144127930-144140491 vs. m54336U_190828_075428/10944934/ccs
Zoom: 8 : 1
Word length: 10
Window size: 0
Matrix: edna.mat
GC ratio seq1: 0.3758
GC ratio seq2: 0.3853
Program: Gepard (1.40 final)
0
chr5:144127930-144140491
12560
0
m54336U_190828_075428/10944934/ccs
13387

chr6:102801951-102802684

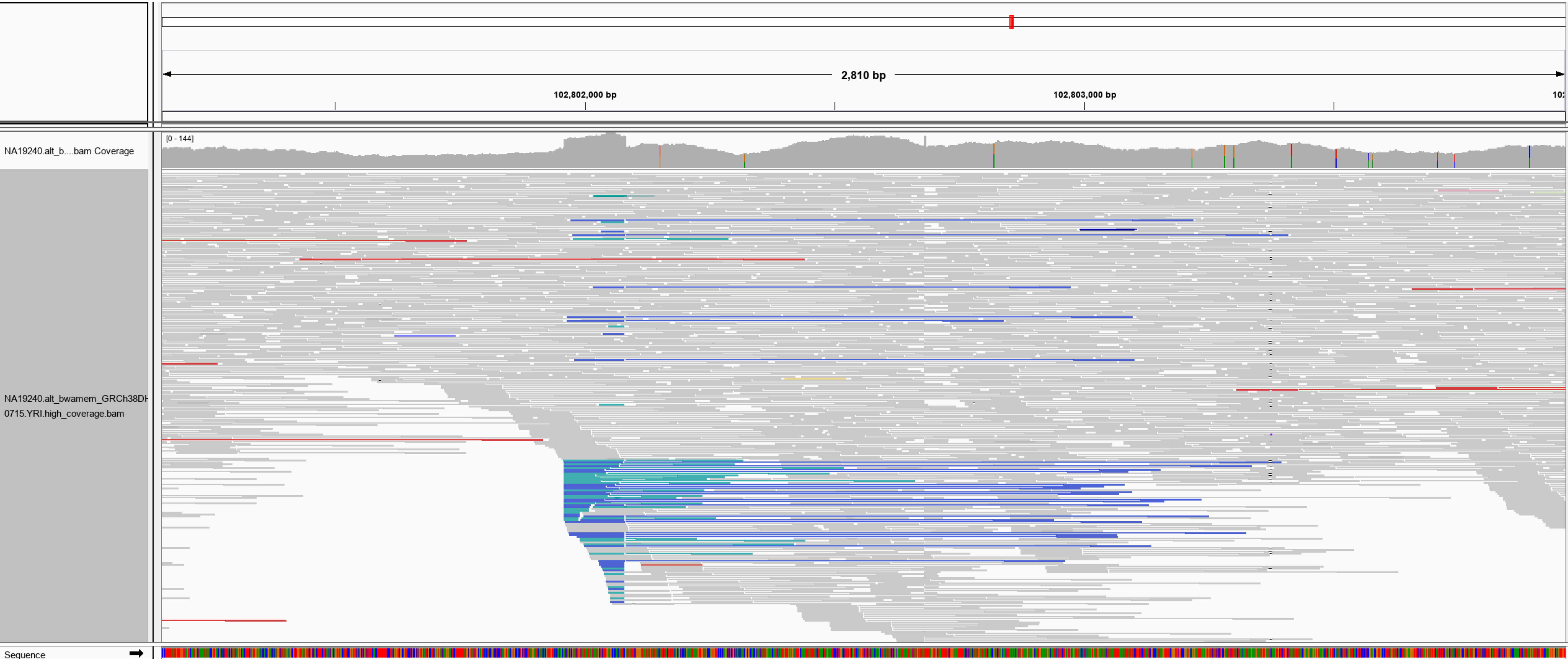

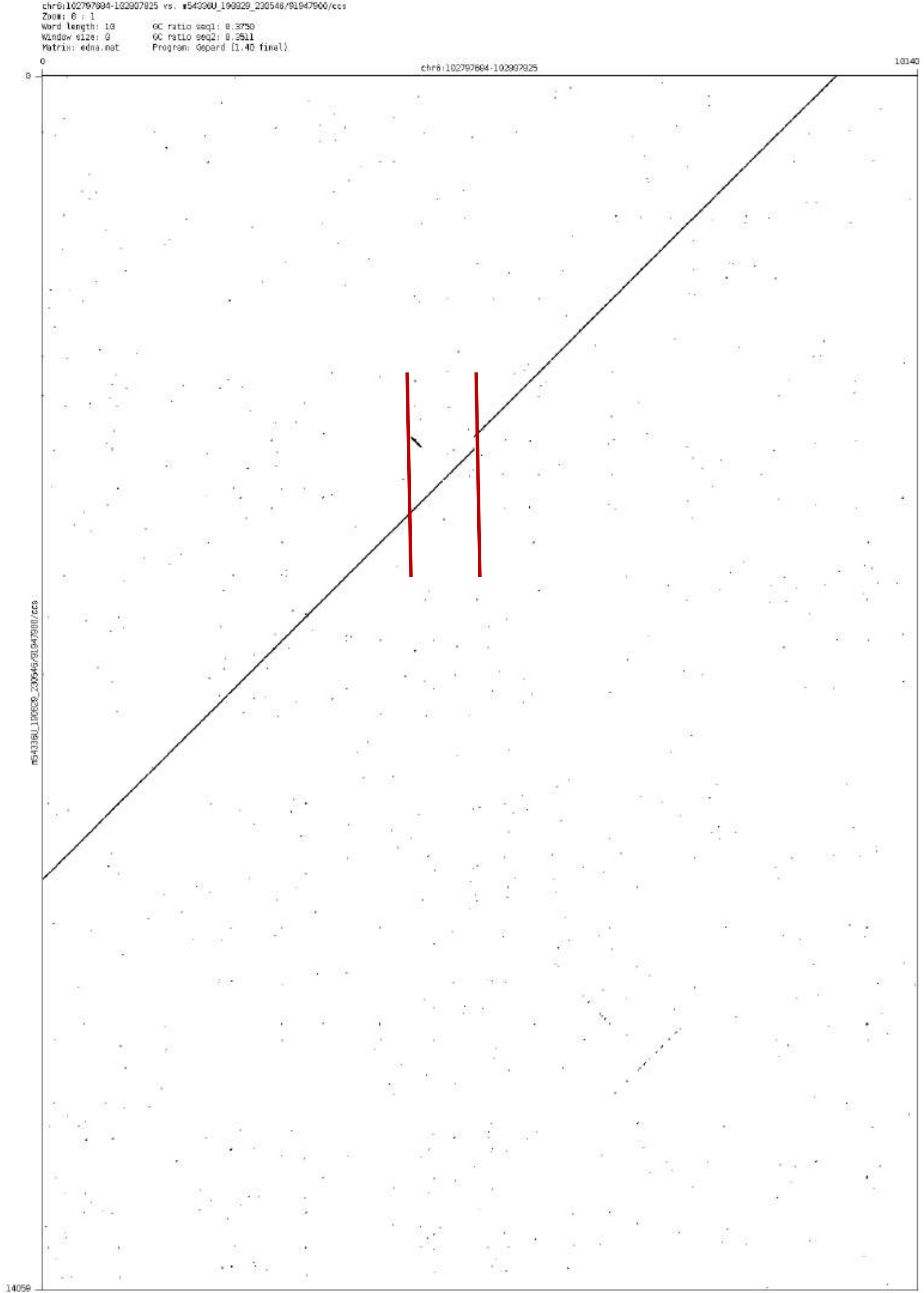

chr8:102797804-102907825 vs. m54336U_190829_230546/91947906/ccs
Zoom: 8 : 1
Word length: 10
Window size: 0
Matrix: edna.mat
GC ratio seq1: 0.3750
GC ratio seq2: 0.3511
Program: Gepard [1.40 final]
0
10140
chr8:102797804-102907825
0
14059
m54336U_190829_230546/91947906/ccs

### chr6:128961308-128962212

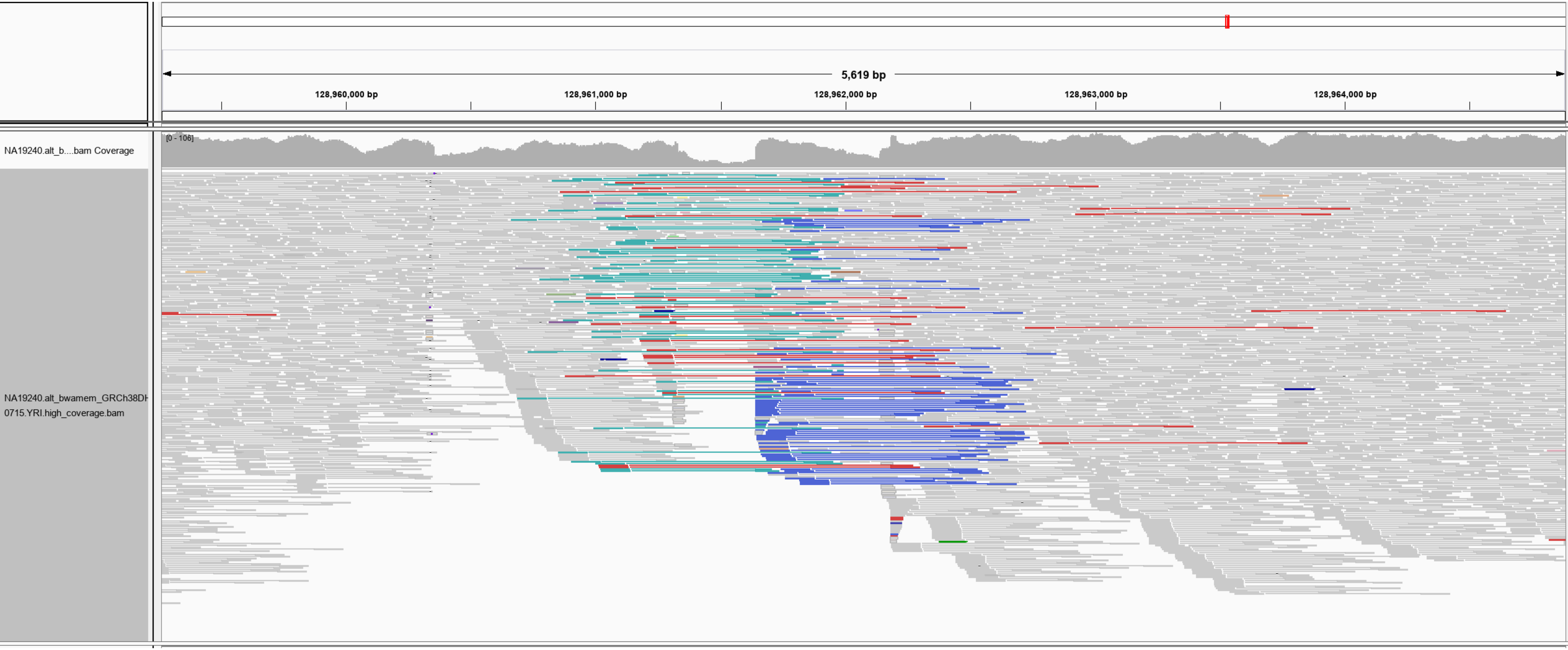

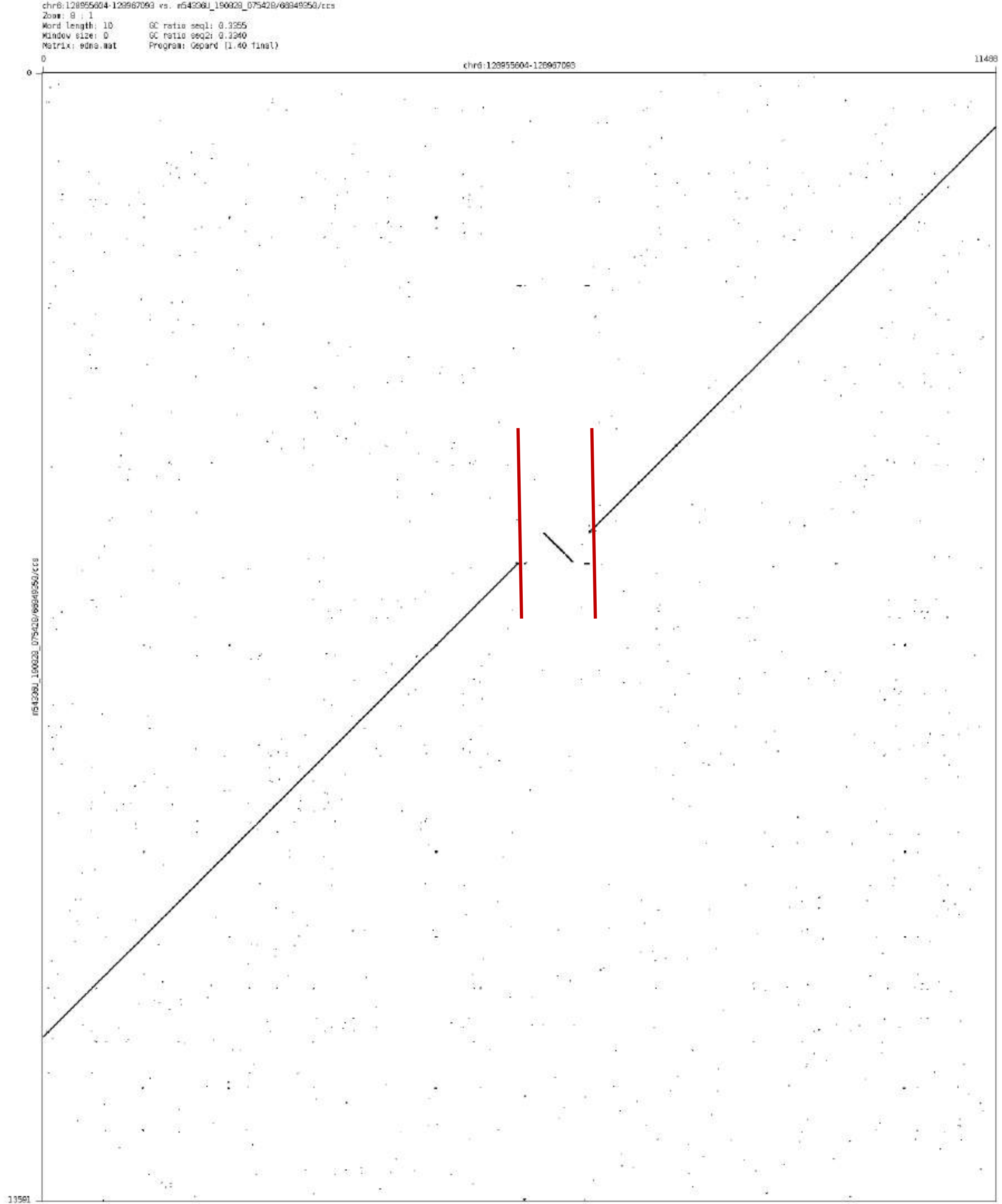

chr6:128955604-128967093 vs. m54336U_190828_075428/66949350/ccs
Zoom: 8 : 1
Word length: 10
Window size: 0
Matrix: edna.mat
GC ratio seq1: 0.3355
GC ratio seq2: 0.3340
Program: Gepard (1.40 final)
0
chr6:128955604-128967093
11488
0
m54336U_190828_075428/66949350/ccs
13591

chr6:139281530-139285938

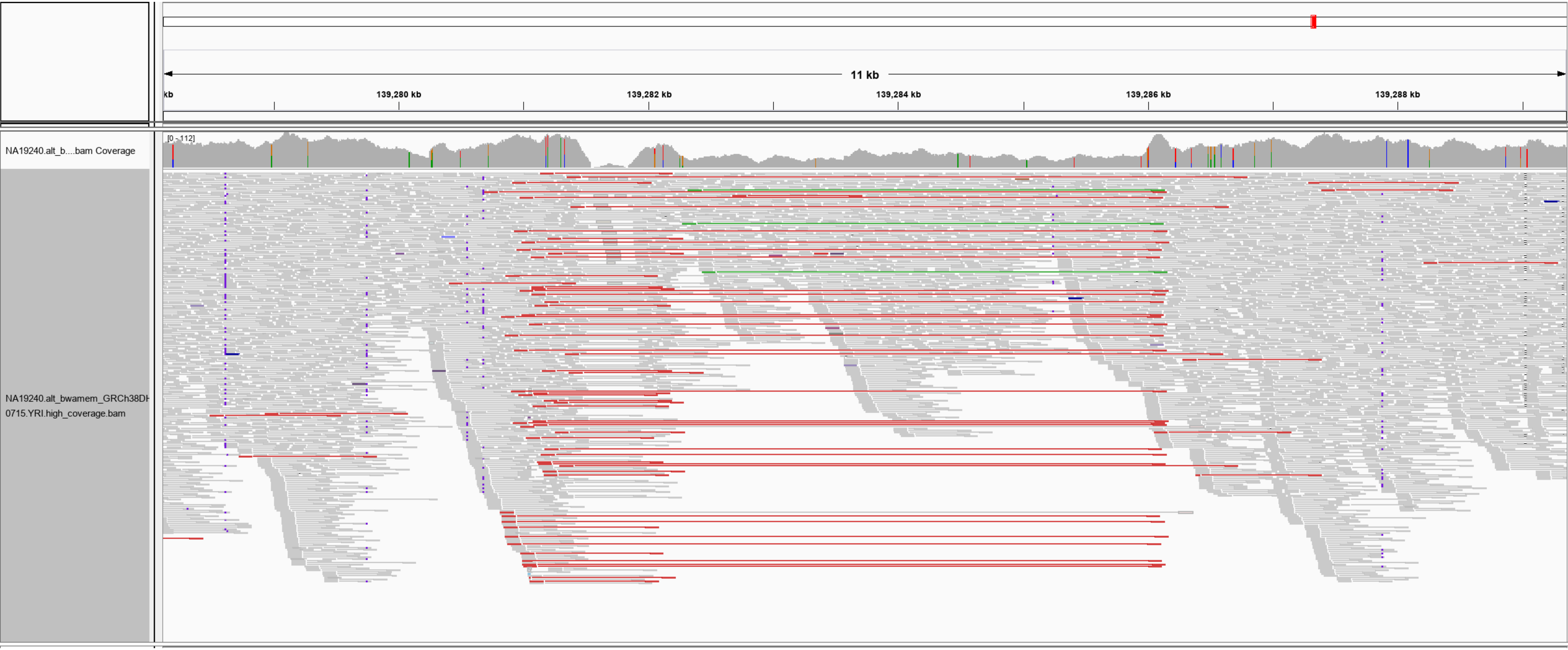

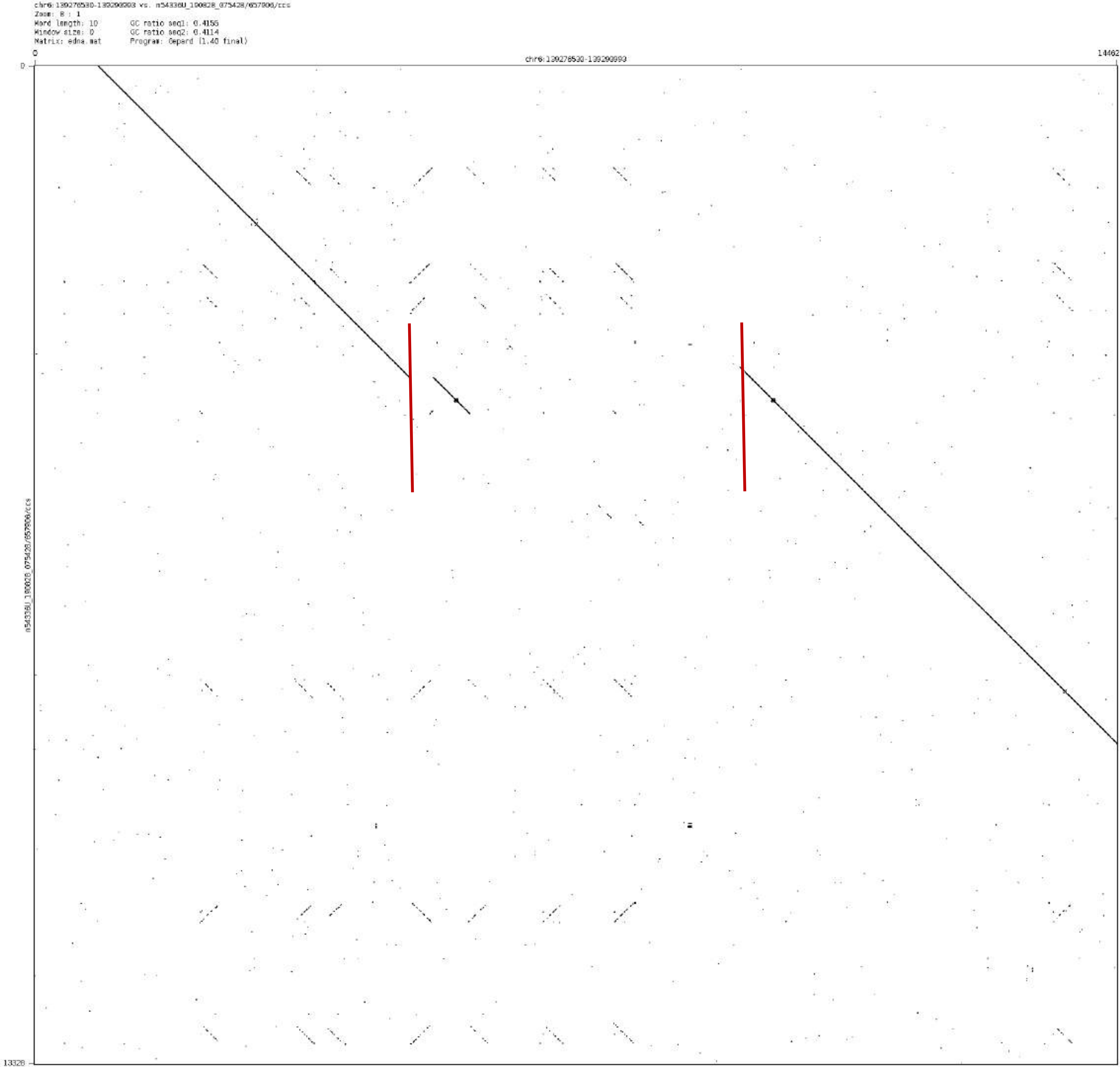
chr6:139276530-139290993 vs. m54336U_190828_075428/657906/ccs
Zoom: 8 : 1
Word length: 10
Window size: 0
Matrix: edna.mat
GC ratio seq1: 0.4155
GC ratio seq2: 0.4114
Program: Gepard (1.40 final)
chr6:139276530-139290993
0
14462
0
13328
m54336U_190828_075428/657906/ccs

chr7:83317161-83318171

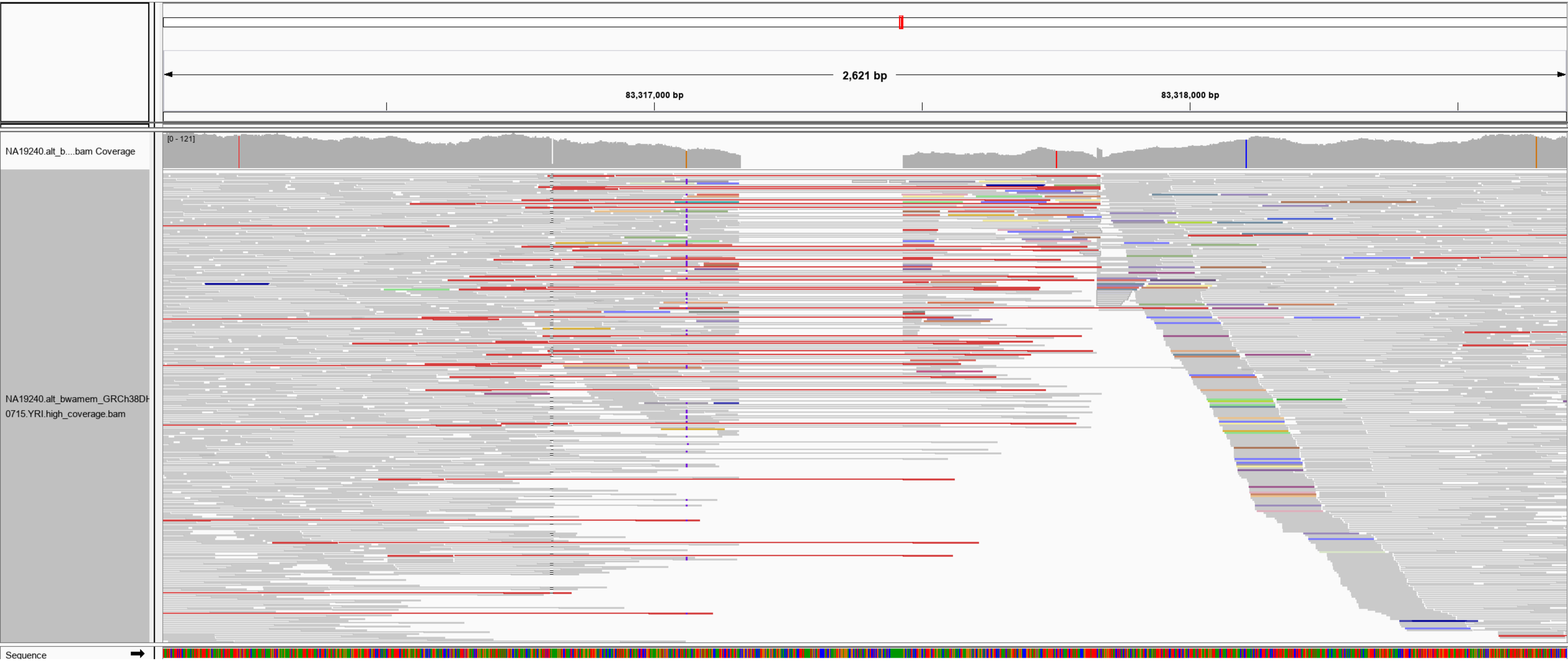

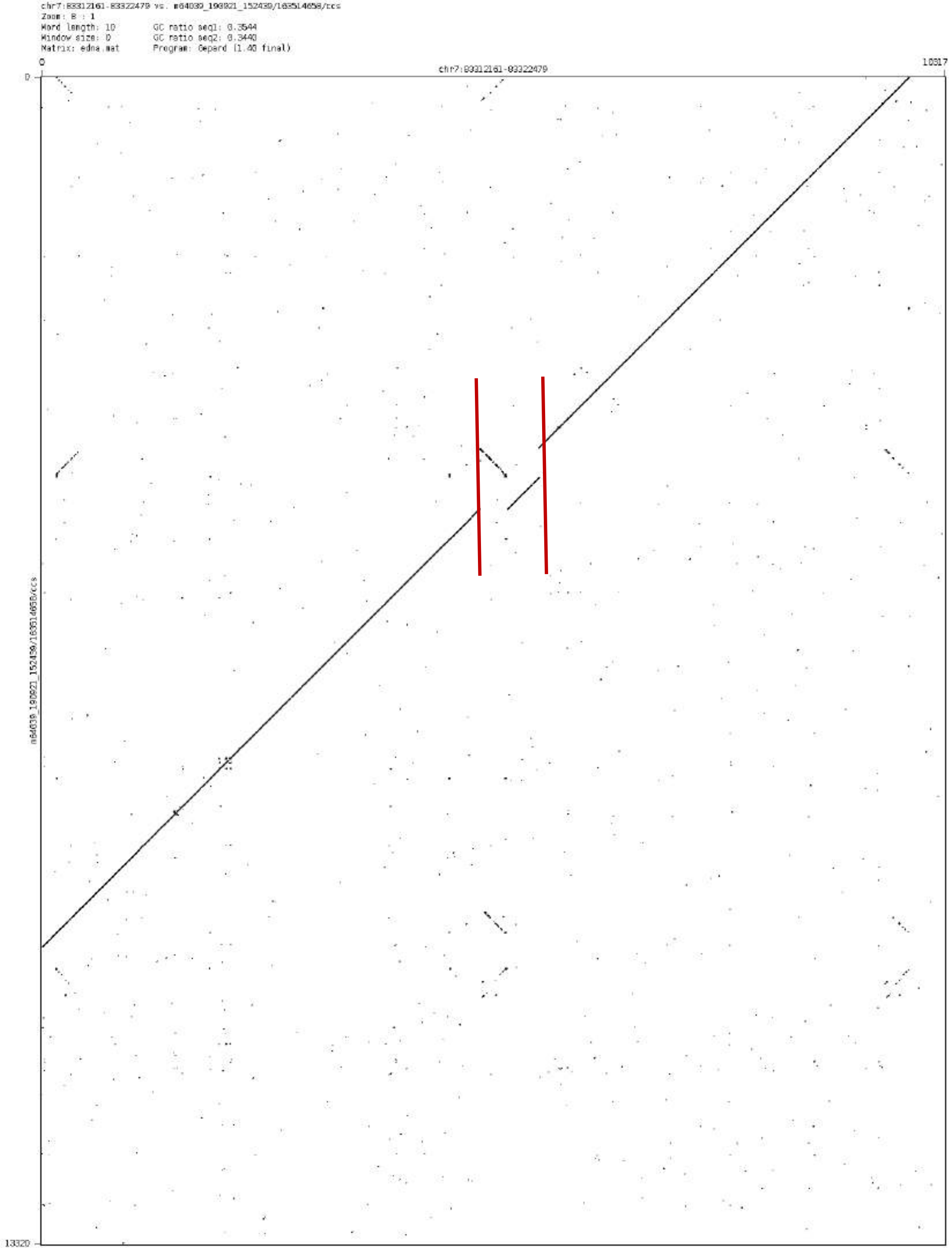

chr7:83312161-83322479 vs. m64039_190921_152439/163514658/ccs
Zoom: 8 : 1
Word length: 10
GC ratio seq1: 0.3544
Window size: 0
GC ratio seq2: 0.3440
Matrix: edna.mat
Program: Gepard (1.40 final)
0
10317
chr7:83312161-83322479
0
13320
m64039_190921_152439/163514658/ccs

chr7: 89324124-89324962

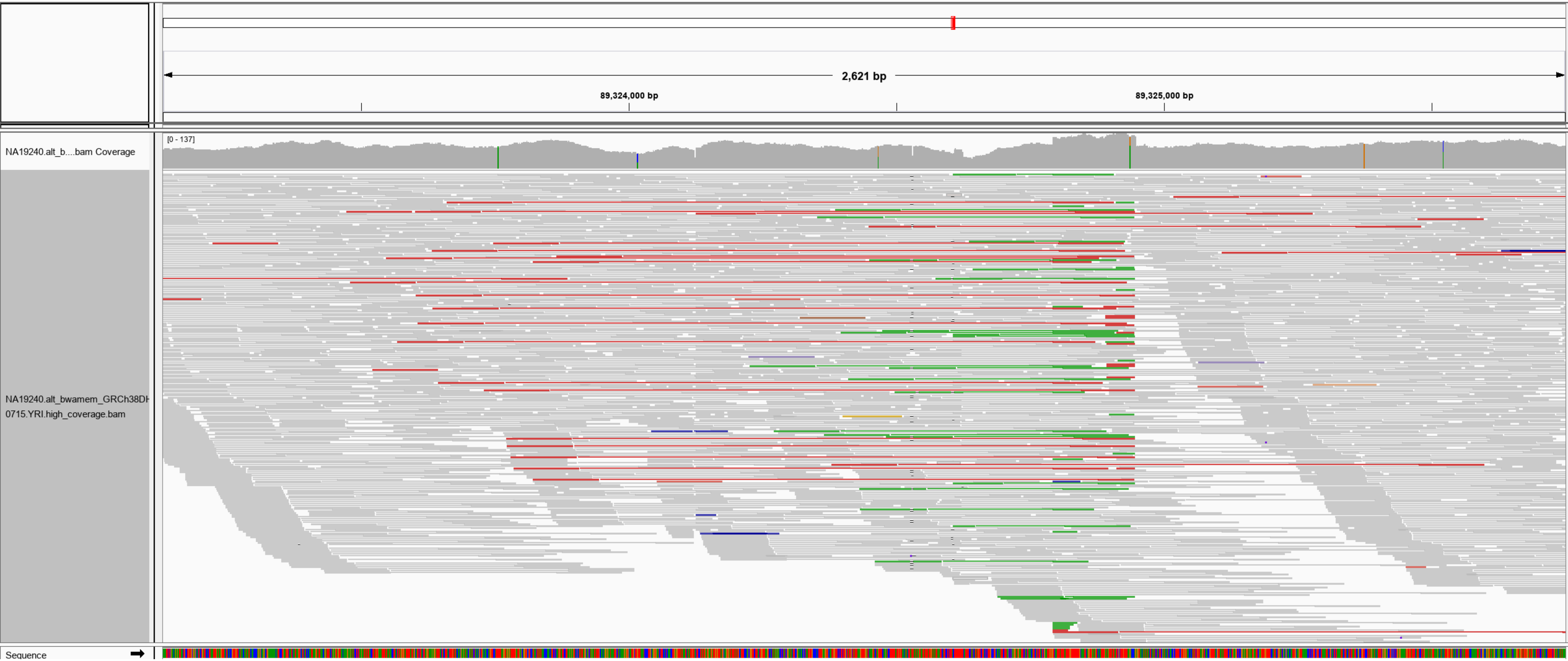

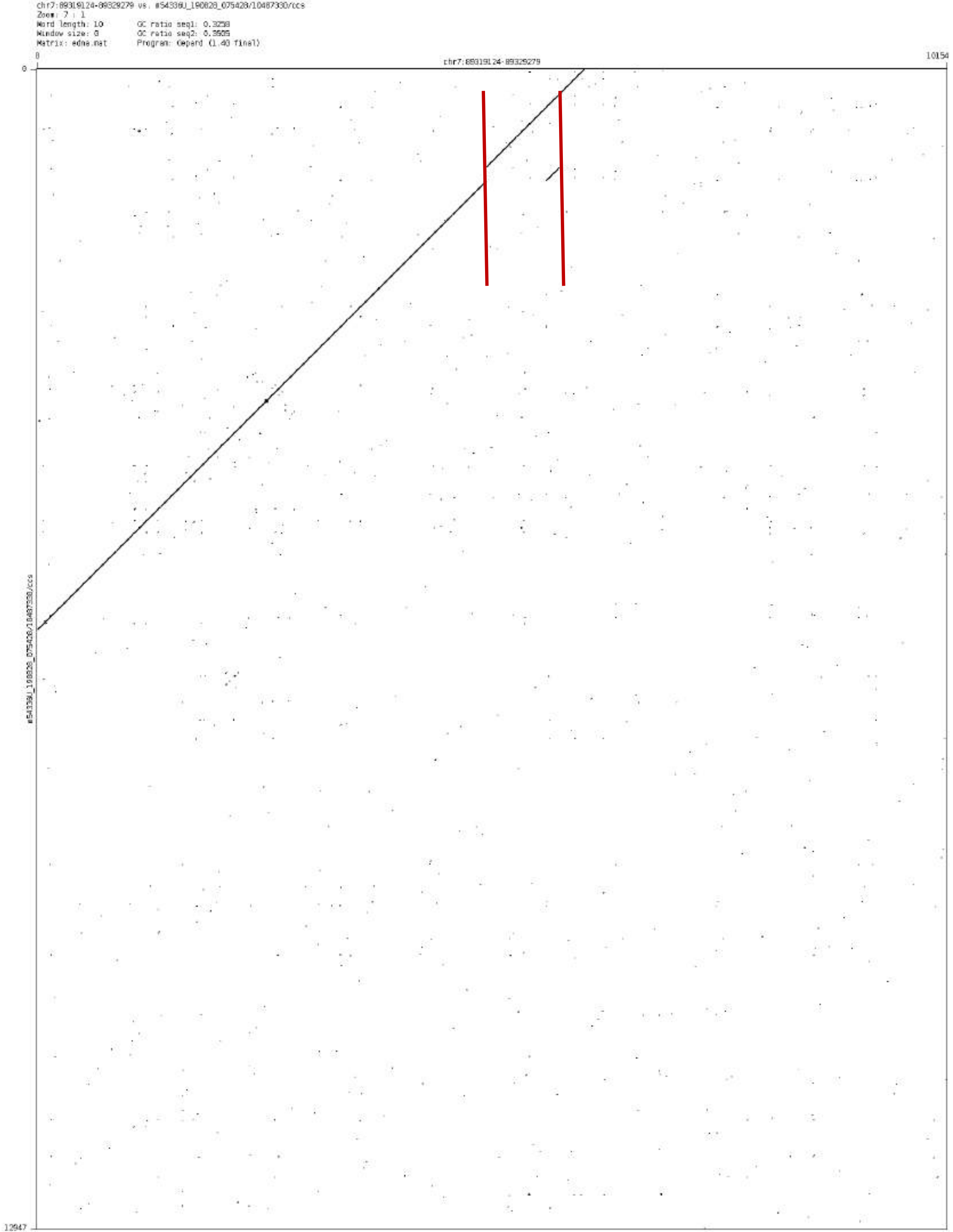
chr7:89319124-89329279 vs. #54338U_190828_075428/10487330/ccs
Zoom: 7 : 1
Word length: 10
Window size: 0
Matrix: edna.mat
GC ratio seq1: 0.3258
GC ratio seq2: 0.3505
Program: Gepard (1.40 final)
chr7:89319124-89329279
0
10154
0
12947

chr7:151312949-151314989

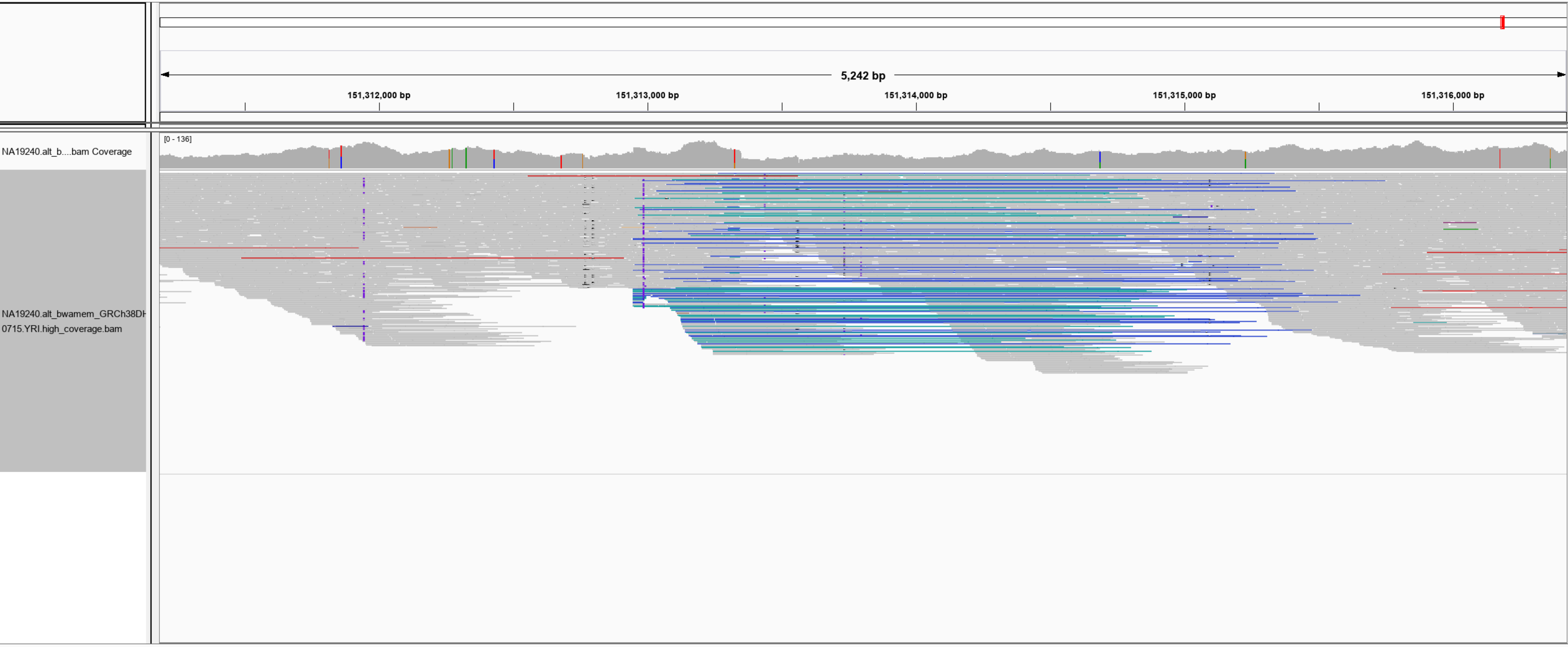

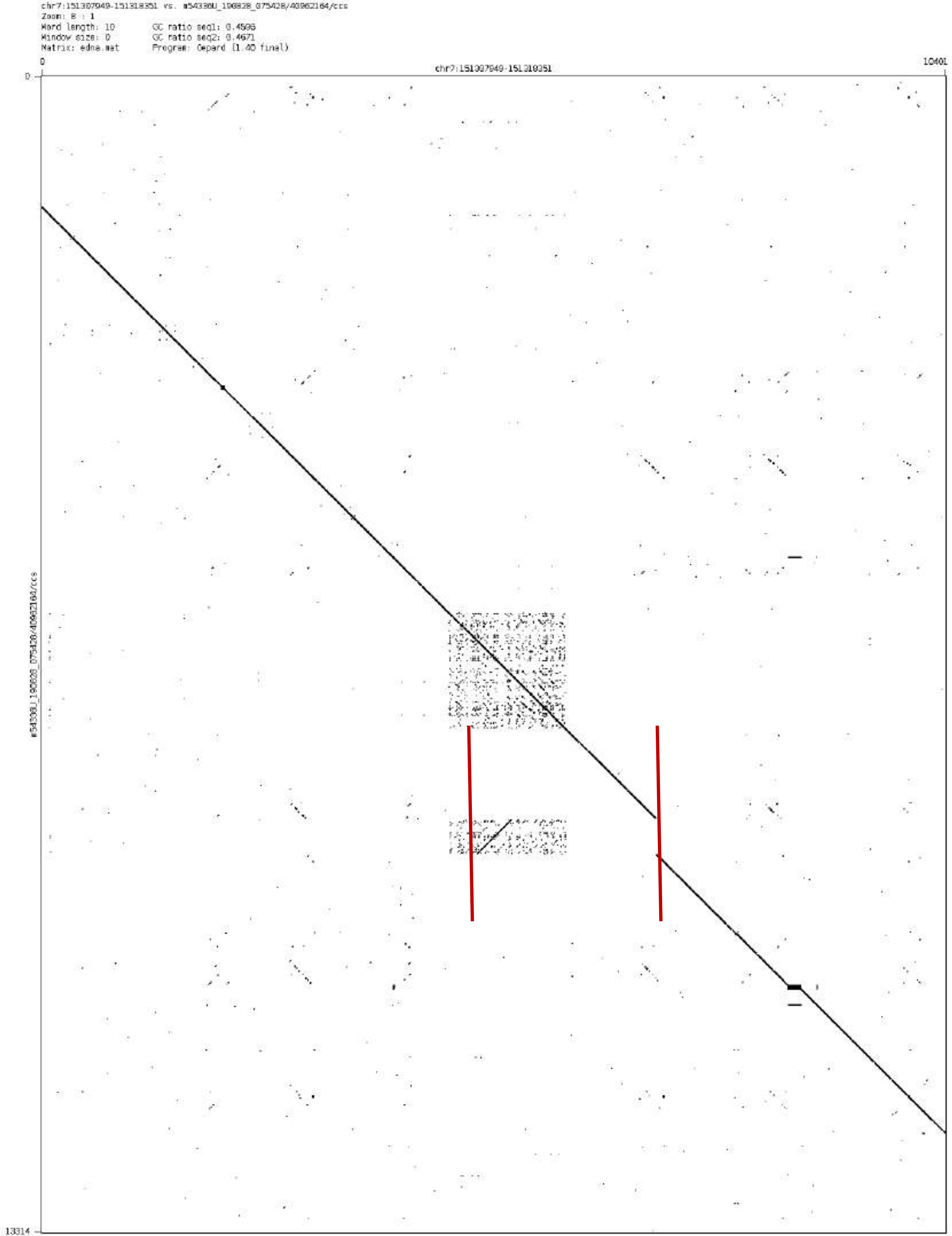

chr7:151307949-151318351 vs. m54330U_190828_075428/40962164/ccs
Zoom: 8 : 1
Word length: 10
GC ratio seq1: 0.4596
Window size: 0
GC ratio seq2: 0.4671
Matrix: edna.mat
Program: Gepard (1.40 final)
0
chr7:151307949-151318351
10401
0
m54330U_190828_075428/40962164/ccs
13314

chr9:69280304-69281825

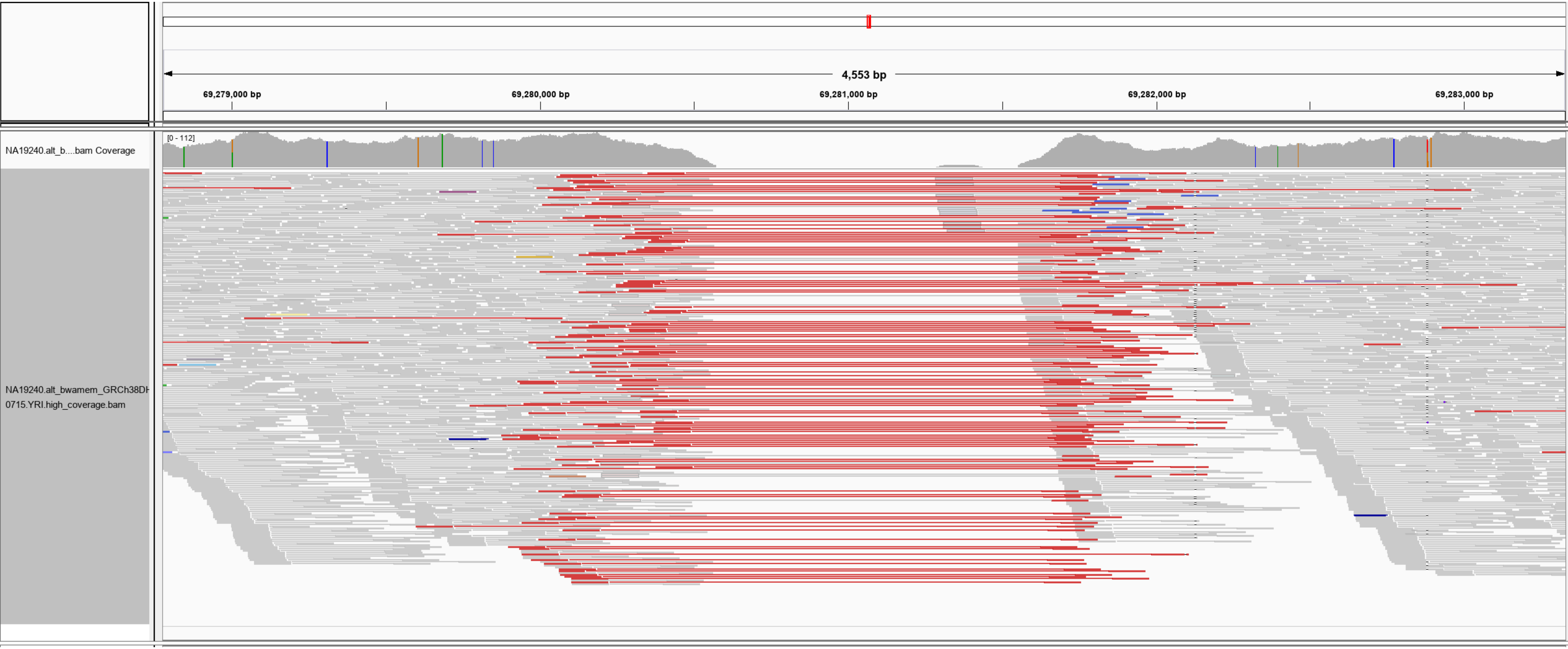

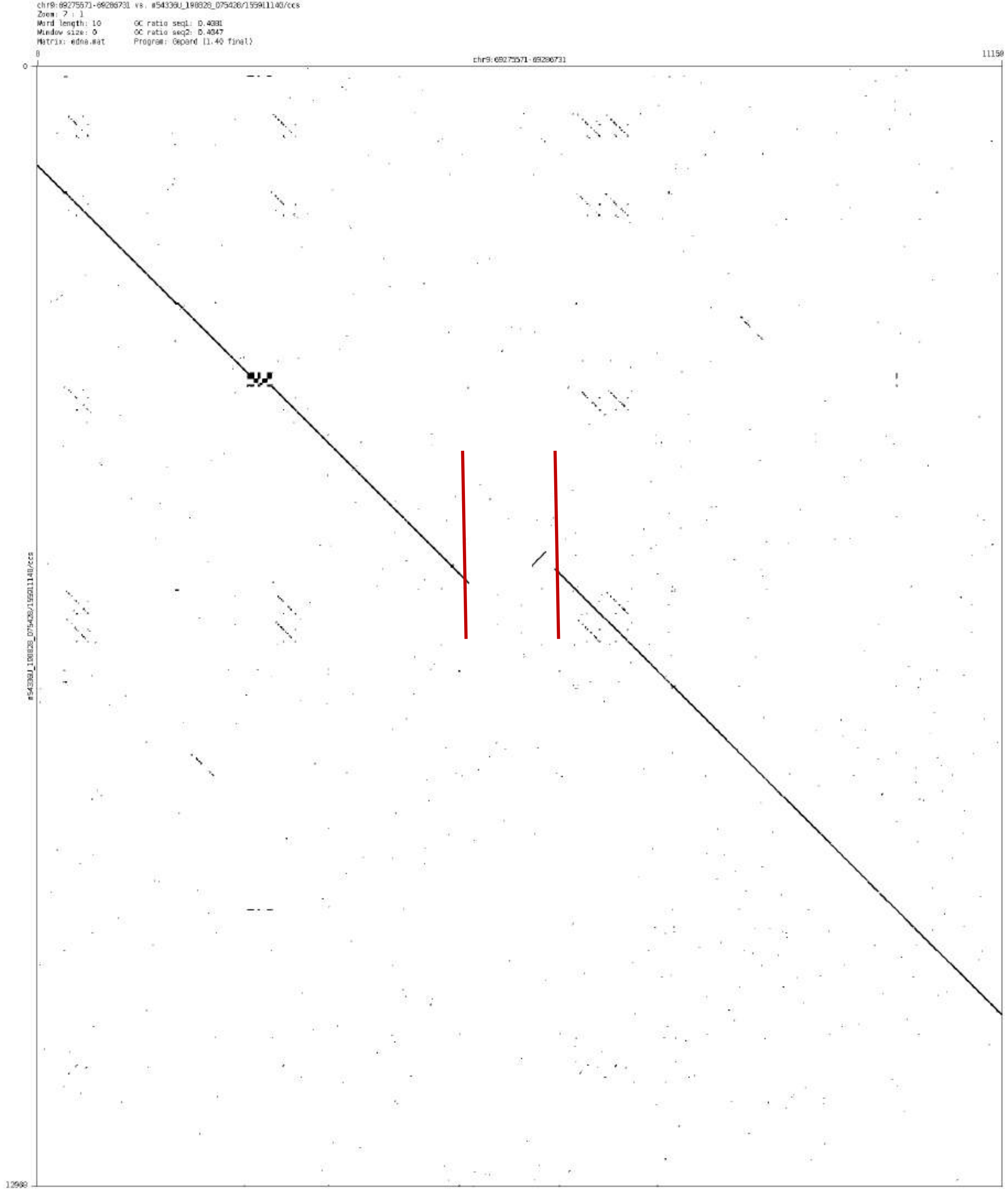

Zoom: 7 : 1
Word length: 10
Window size: 0
Matrix: edna.mat
Program: Gepard (1.40 final)

chr9:86539584-86541086

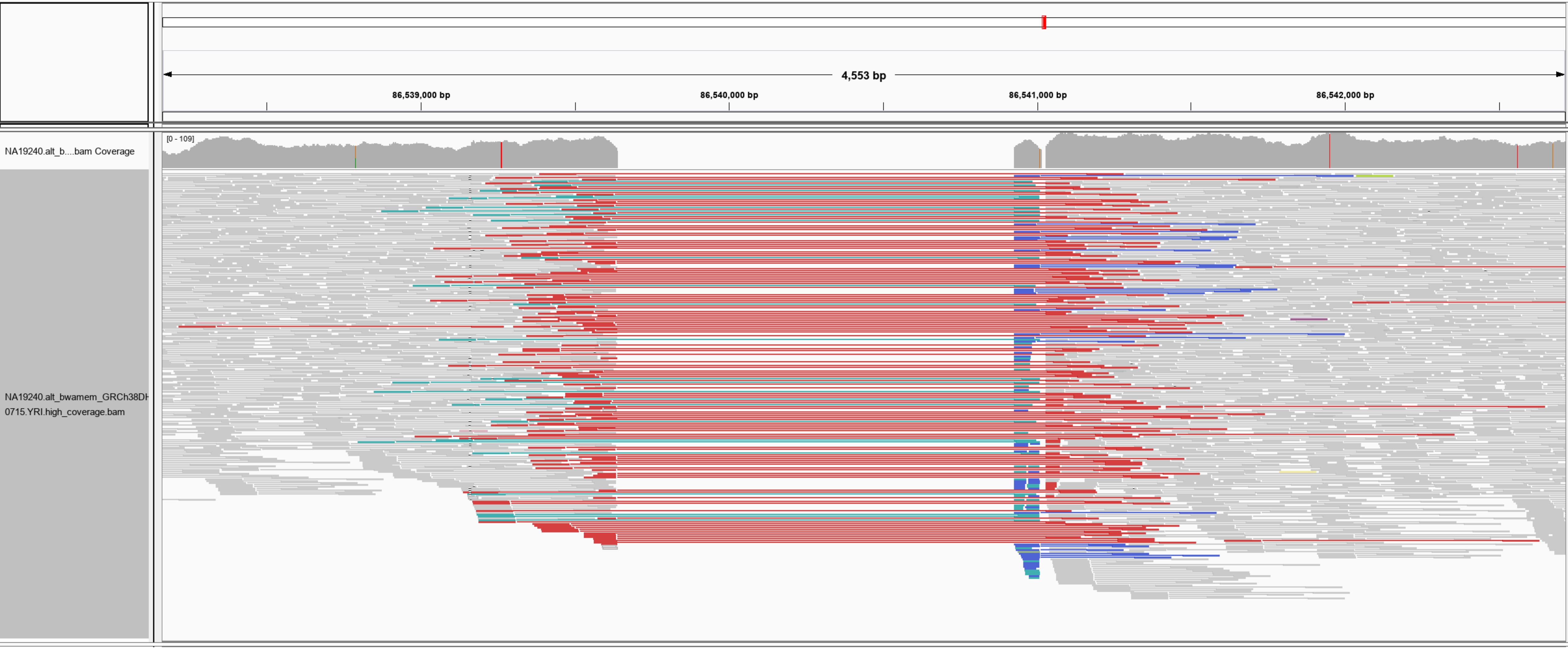

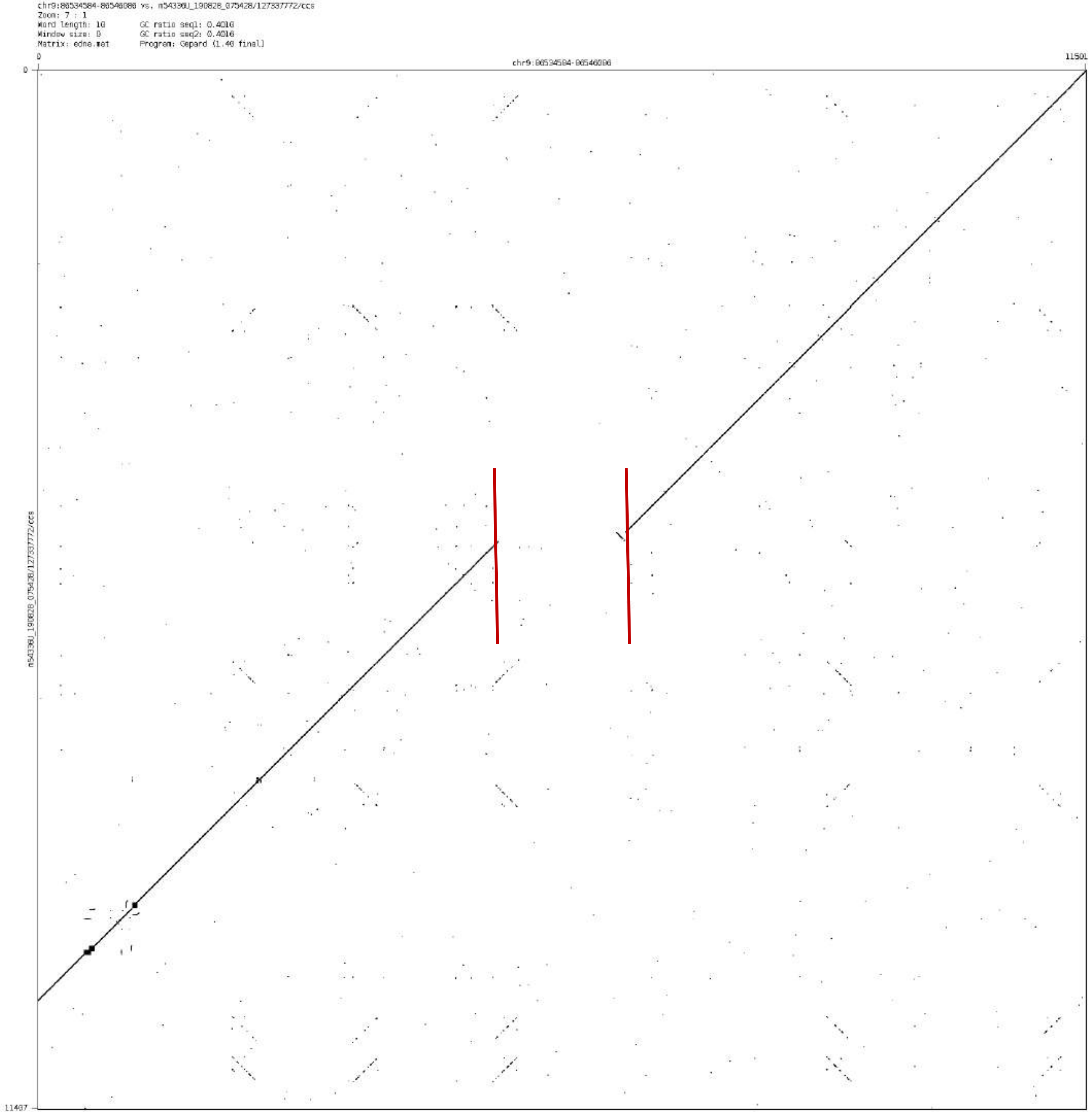
chr9:86534584-86546086 vs. m54330U_190828_075428/127337772/ccs
Zoom: 7 : 1
Word length: 10
Window size: 0
Matrix: edna.mat
GC ratio seq1: 0.4016
GC ratio seq2: 0.4016
Program: Gepard (1.40 final)
0
chr9:86534584-86546086
11501
11407
m54330U_190828_075428/127337772/ccs

chr9:105054355-105055067

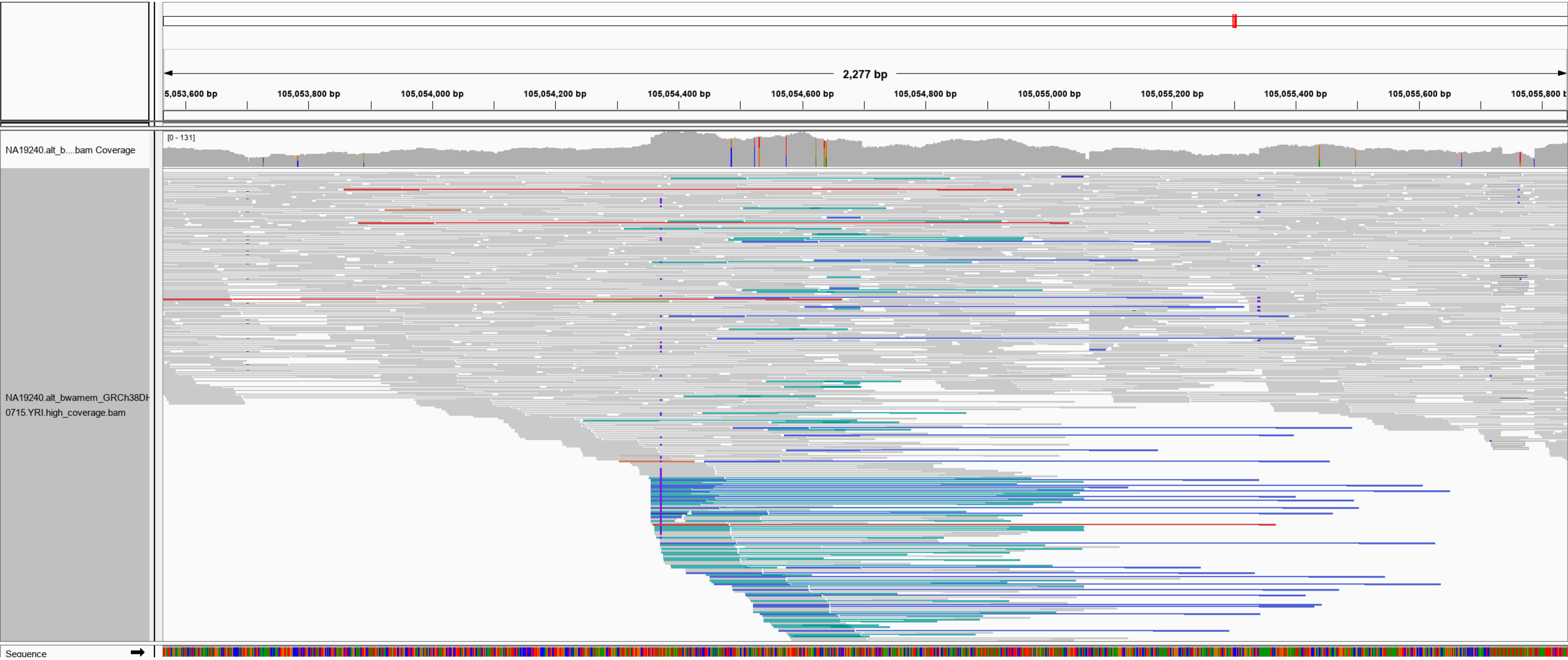

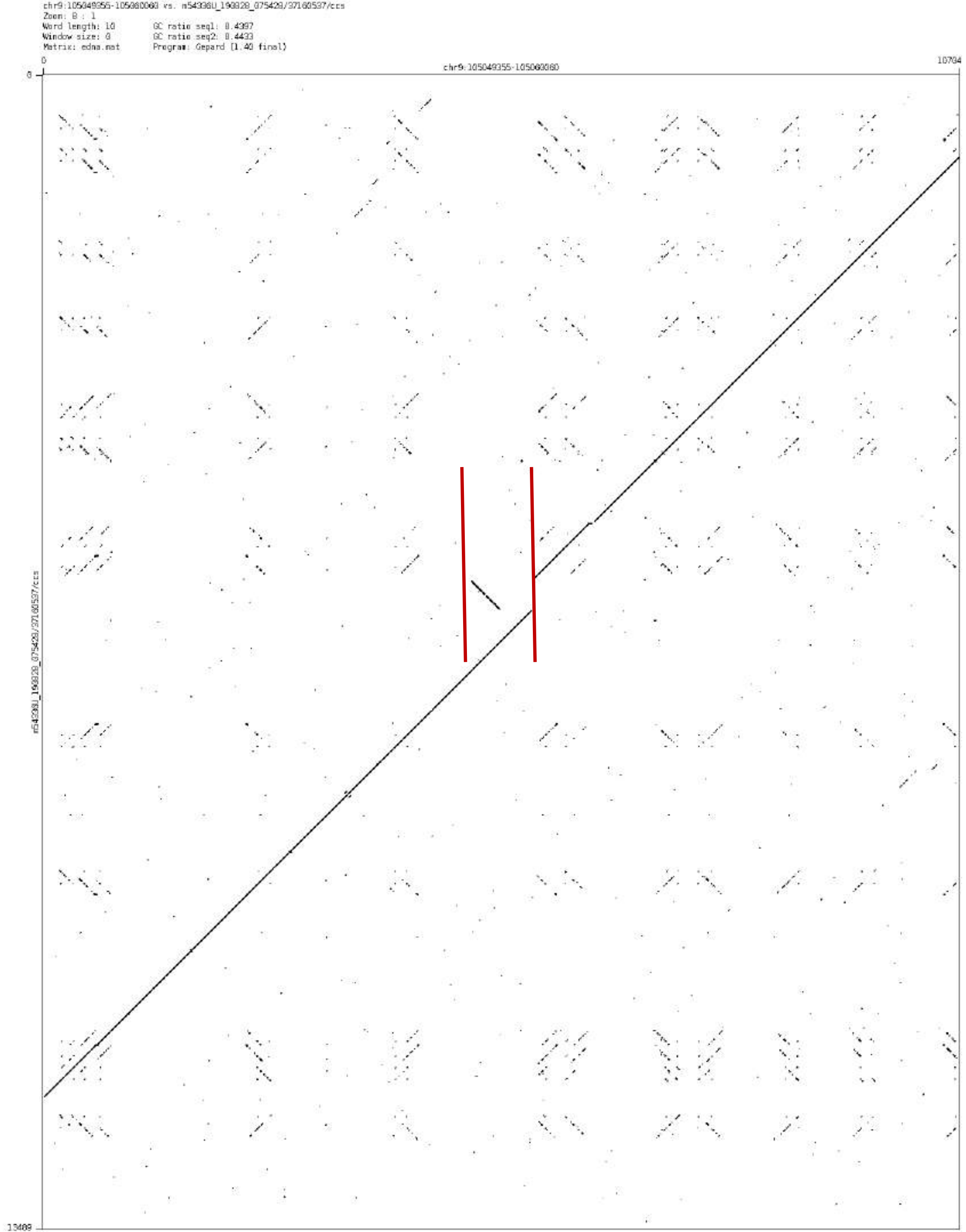
chr9:105049355-105060060 vs. m54338U_190828_075429/37160537/ccs
Zoom: 8 : 1
Word length: 10
Window size: 0
Matrix: edna.mat
GC ratio seq1: 0.4397
GC ratio seq2: 0.4433
Program: Gepard [1.40 final]
0
10704
chr9:105049355-105060060
0
13409
m54338U_190828_075429/37160537/ccs

chr9:109523550-109524392

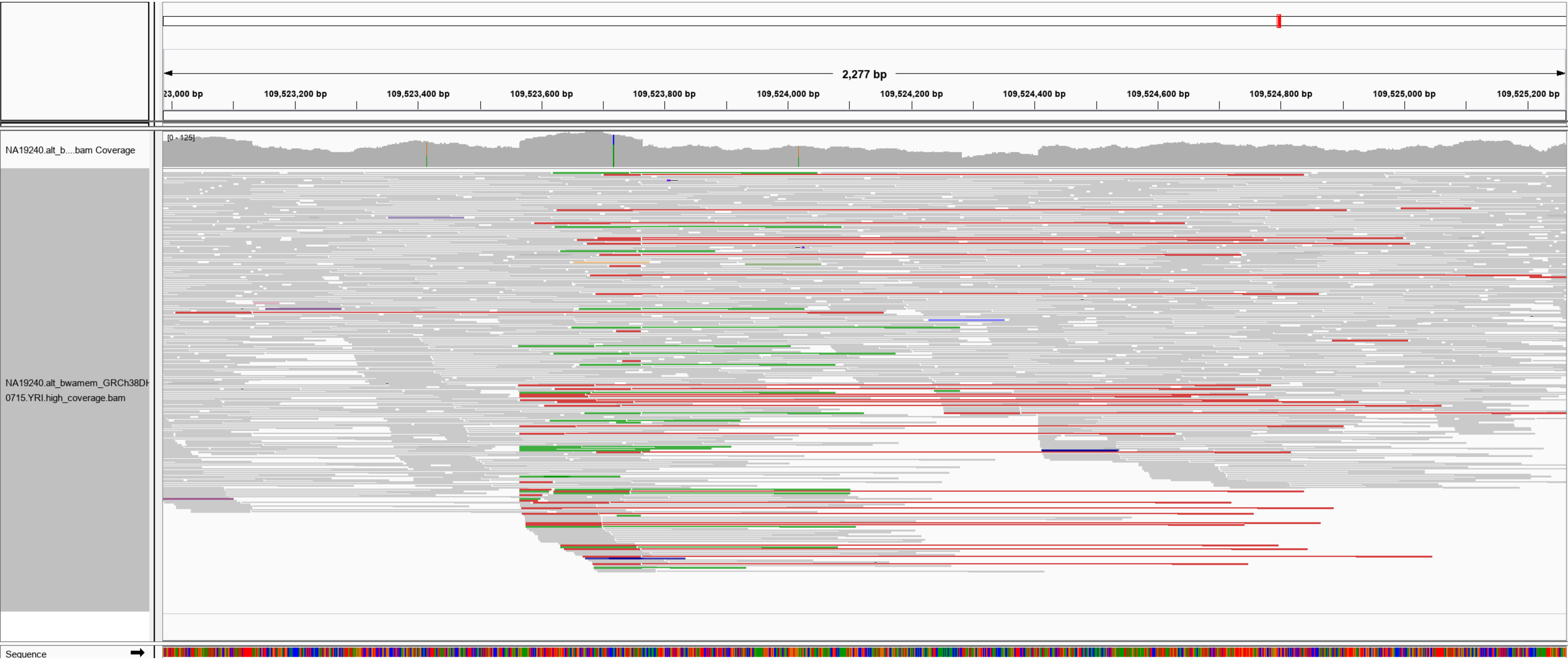

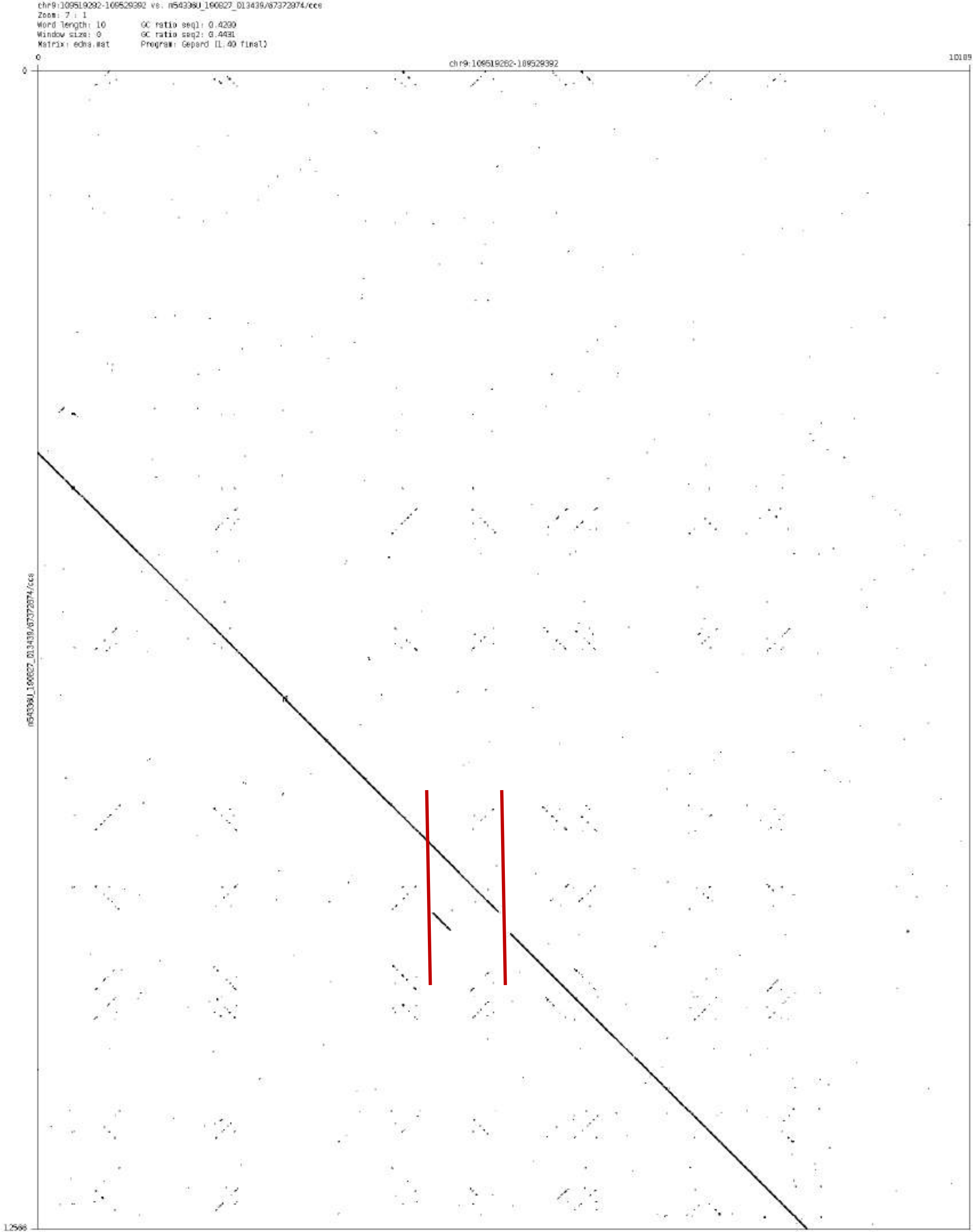
chr9:109519292-109529392 vs. m54336U_190827_013439/67372974/ccs
Zoom: 7 : 1
Word length: 10
Window size: 0
Matrix: edna.mat
GC ratio seq1: 0.4200
GC ratio seq2: 0.4431
Program: Gepard (1.40 final)
0
chr9:109519292-109529392
10109
0
m54336U_190827_013439/67372974/ccs
12566

### chr9:132883700-132884812

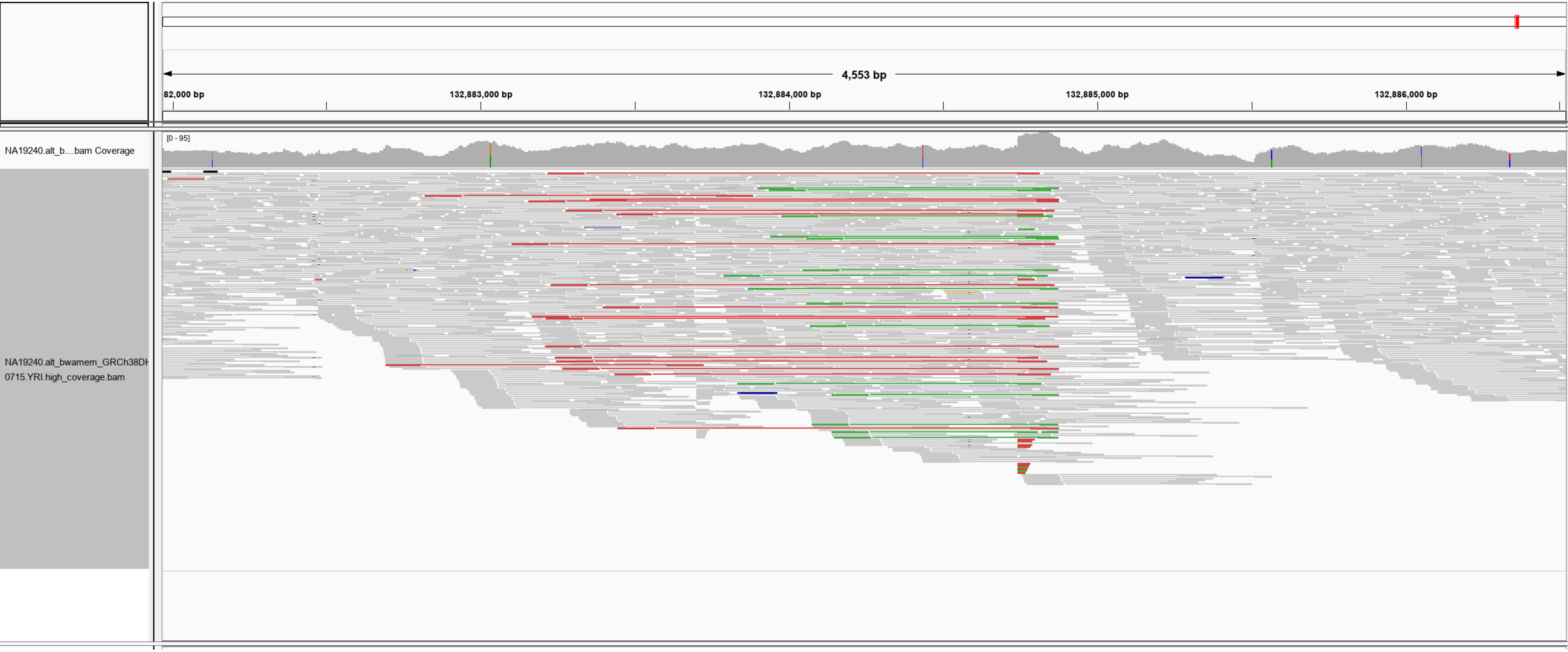

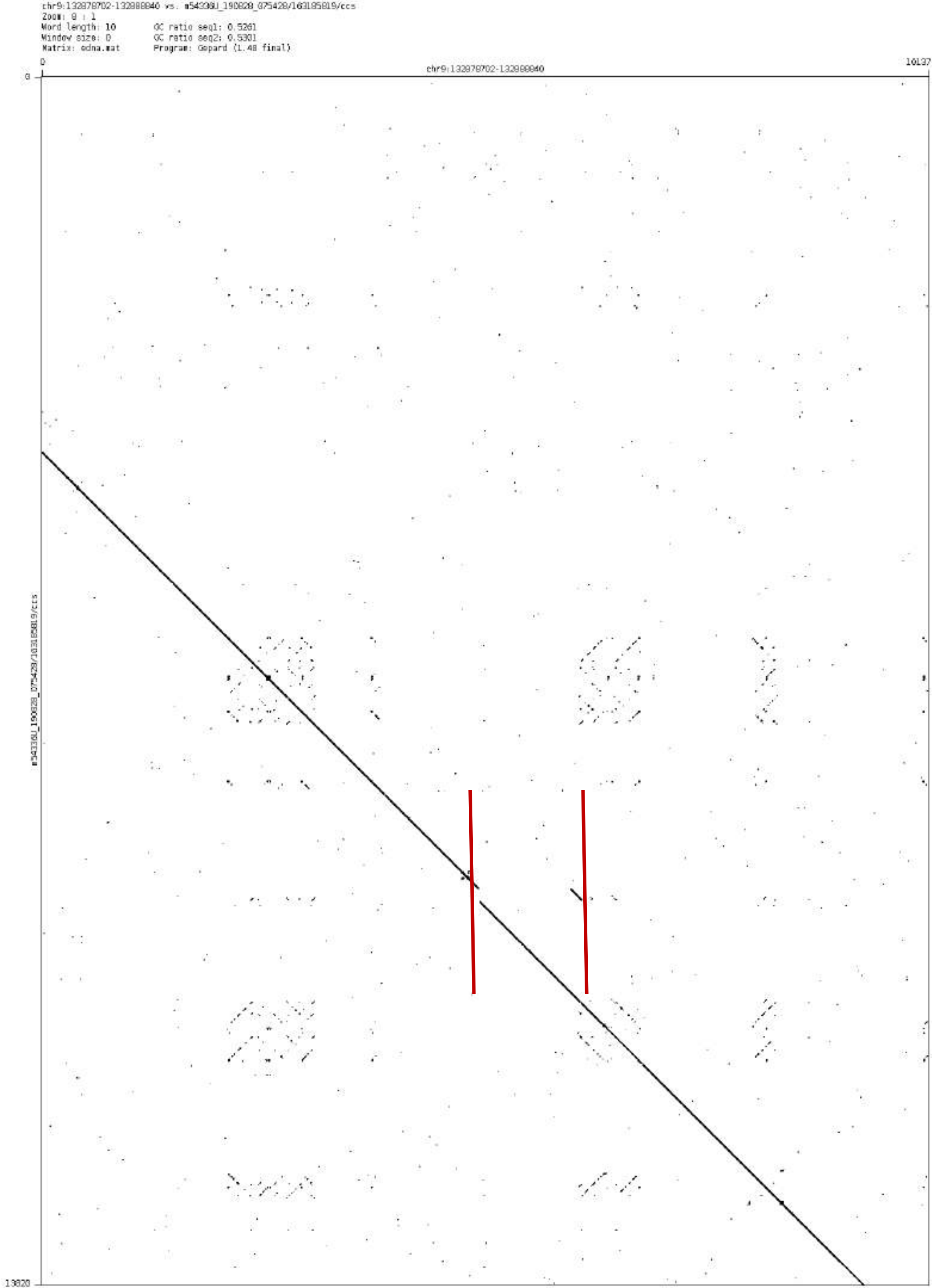
chr9:132878702-132888840 vs. m54336U_190828_075428/163185819/ccs
Zoom: 8 : 1
Word length: 10
GC ratio seq1: 0.5261
Window size: 0
GC ratio seq2: 0.5301
Matrix: ednа.mat
Program: Gepard (1.40 final)
0
chr9:132878702-132888840
10137
0
13820

### chr10:44927495-44928688

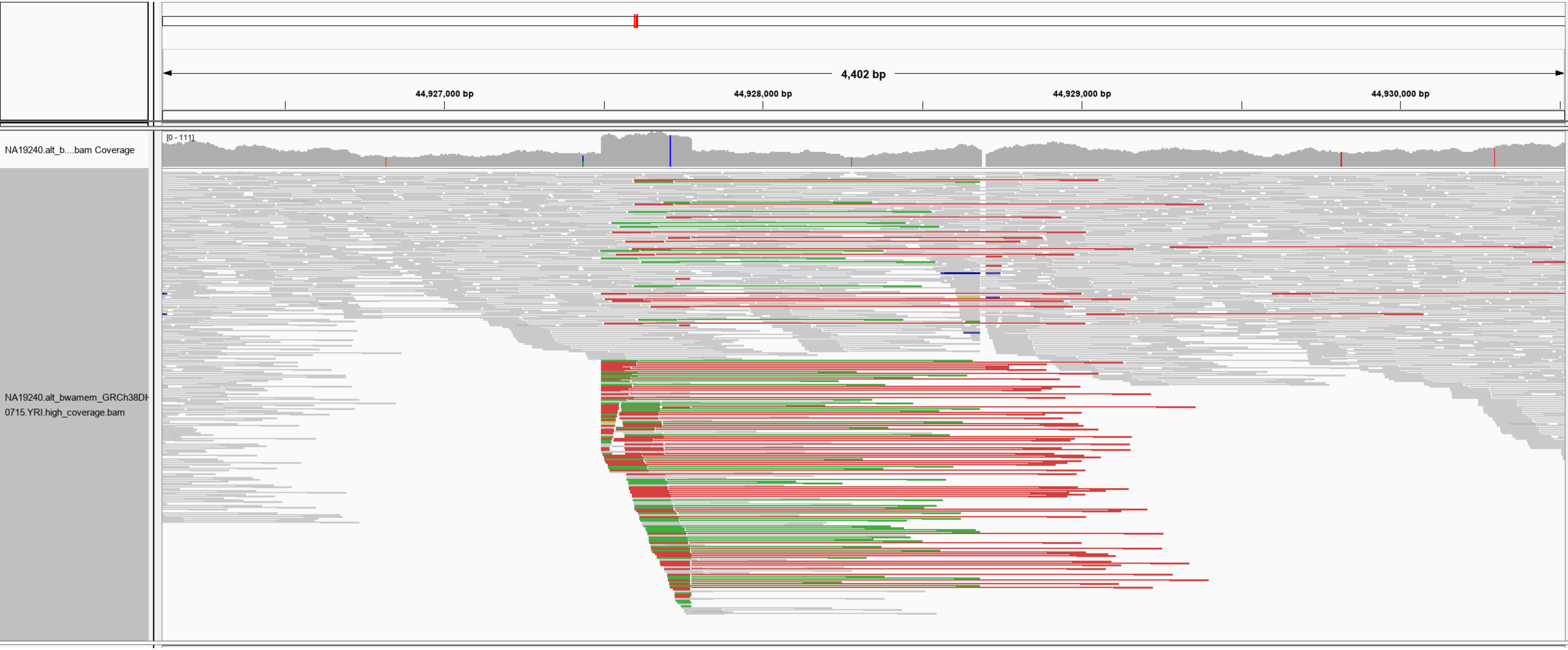

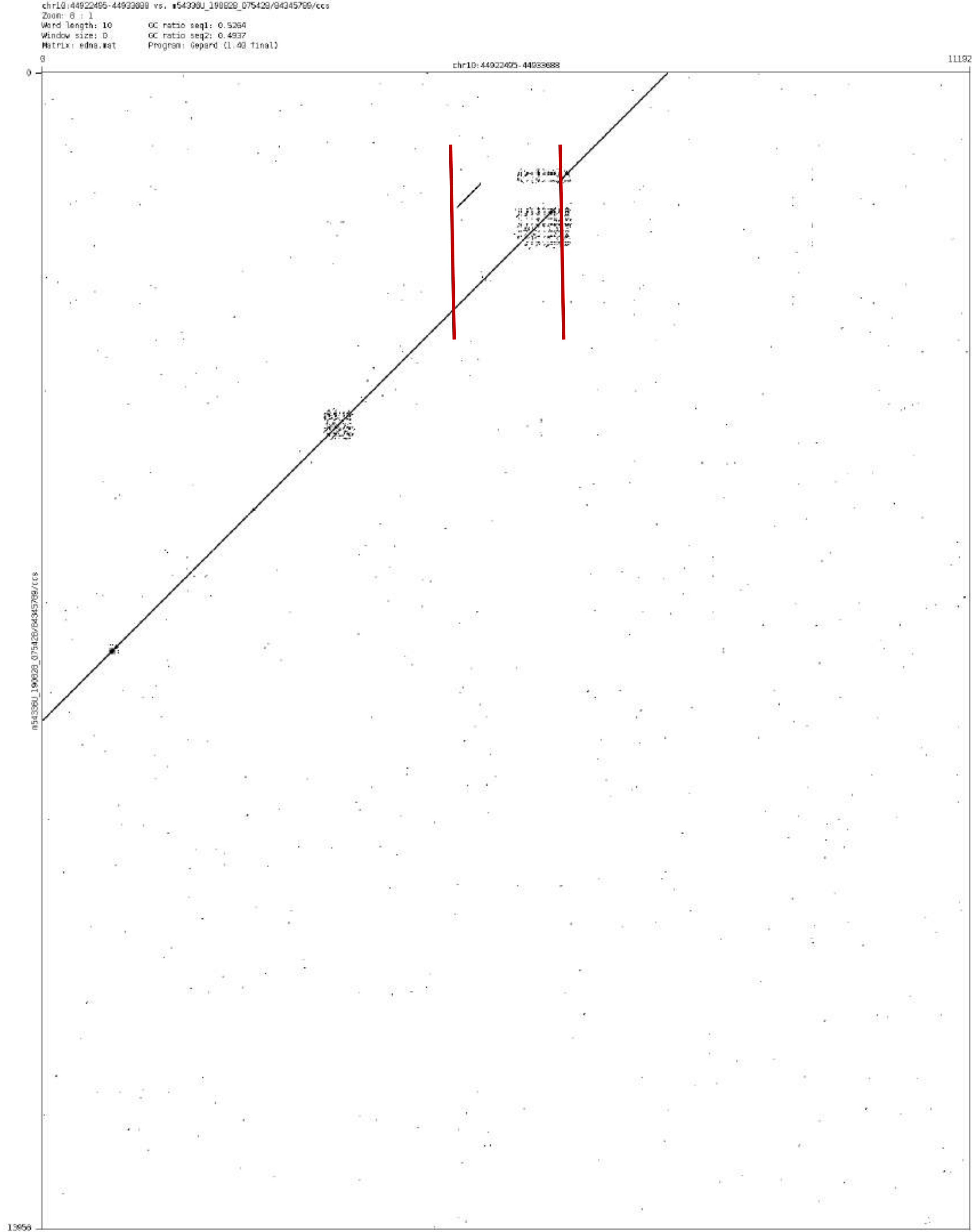
Zoom: 8 : 1
Word length: 10
Window size: 0
Matrix: edna.mat
GC ratio seq1: 0.5264
GC ratio seq2: 0.4937
Program: Gepard (1.40 final)
0
11192
0
13956

chr11:6042862-6044585

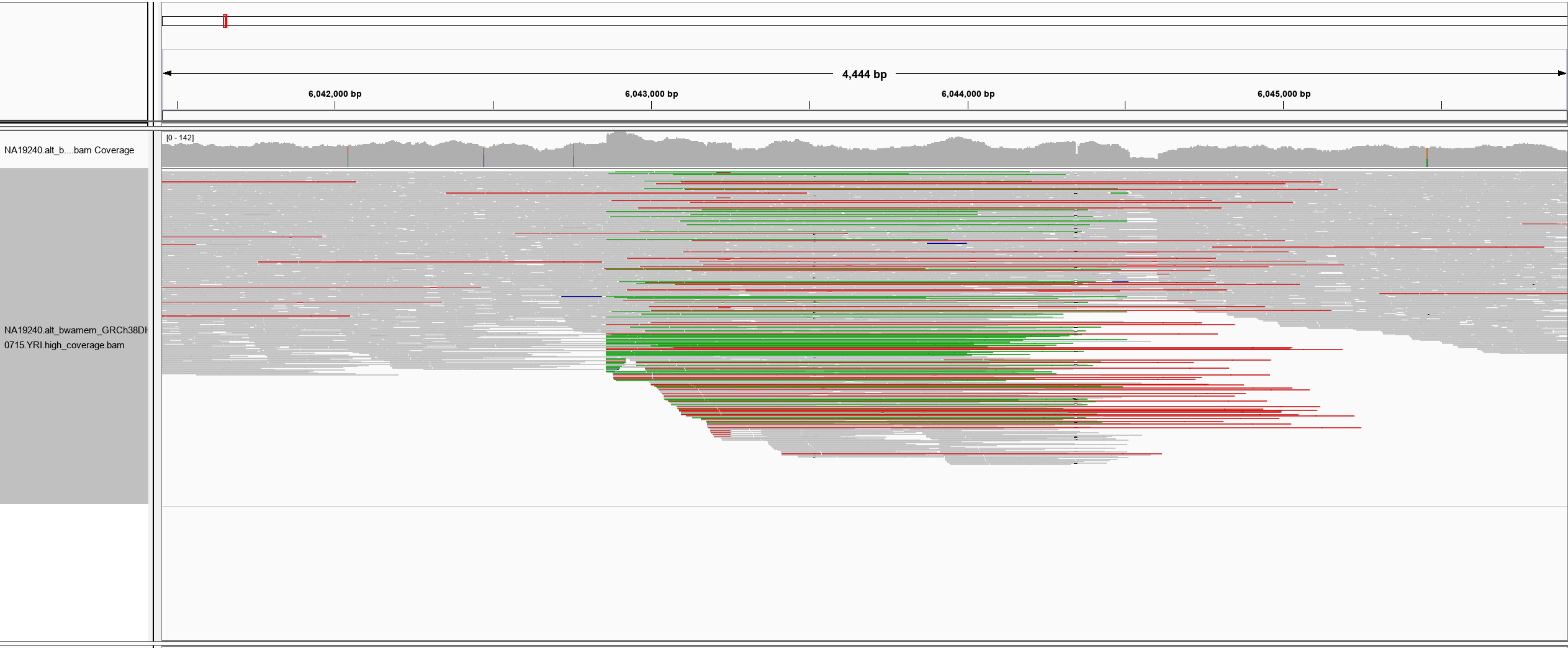

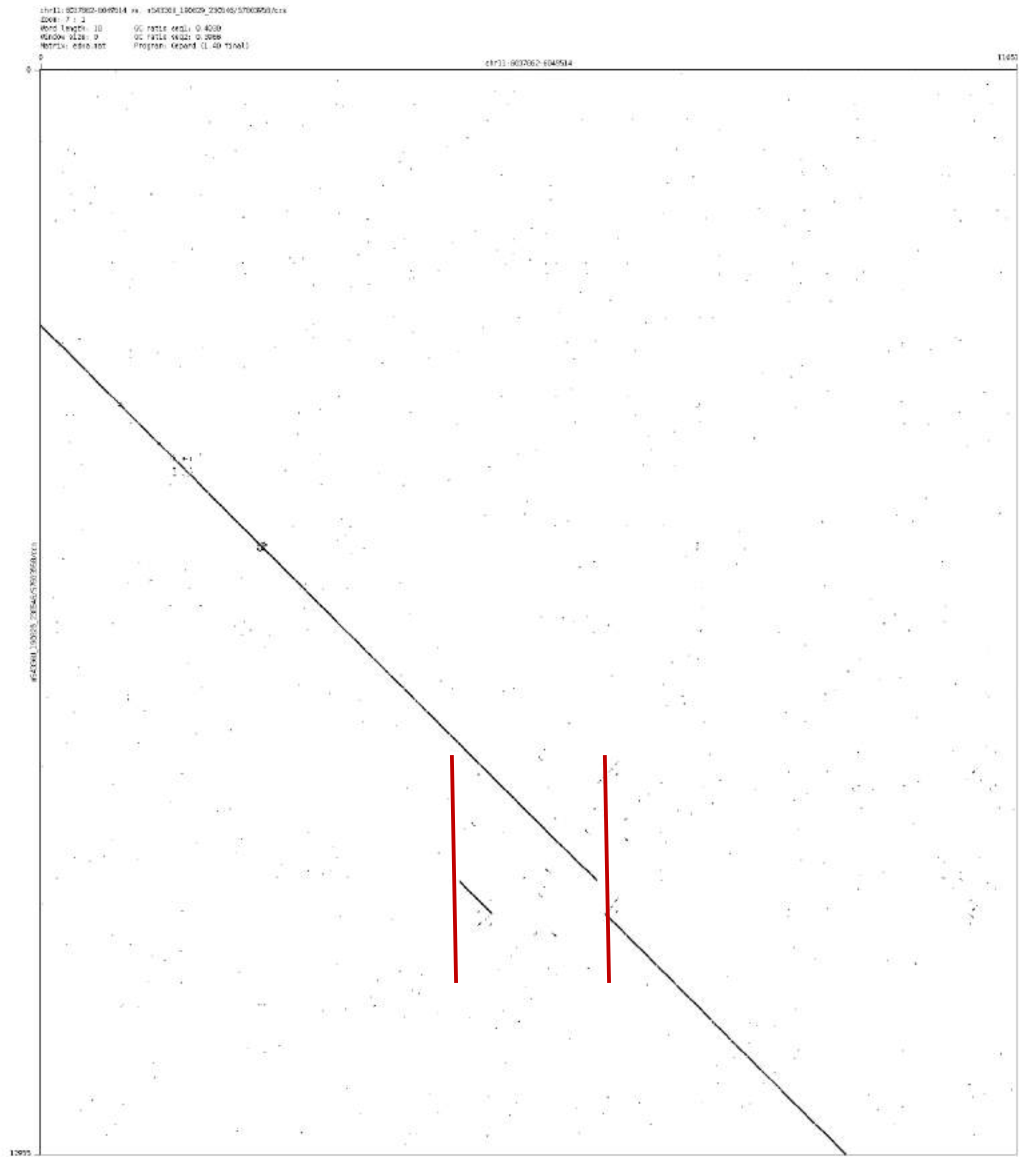

Word length: 10
Program: Gepard (1.40 final)

### chr11:24882412-24883333

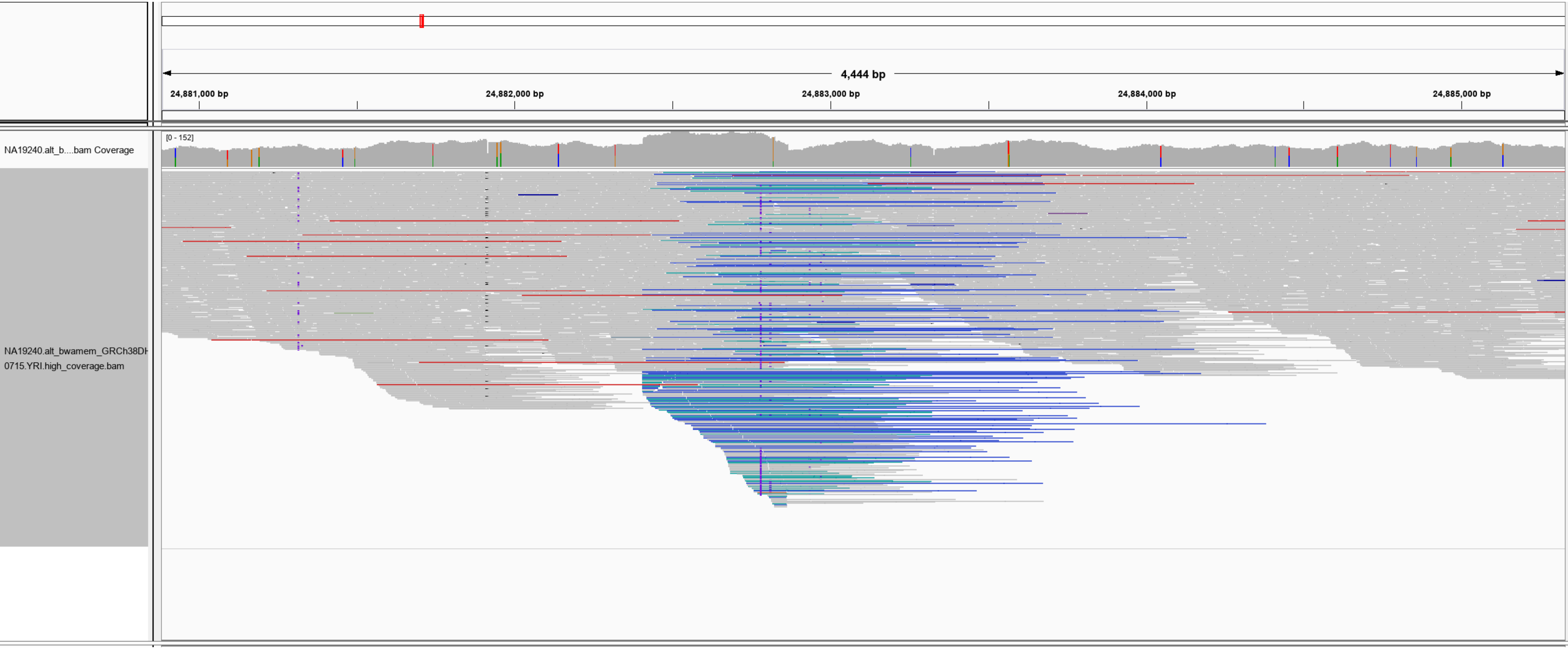

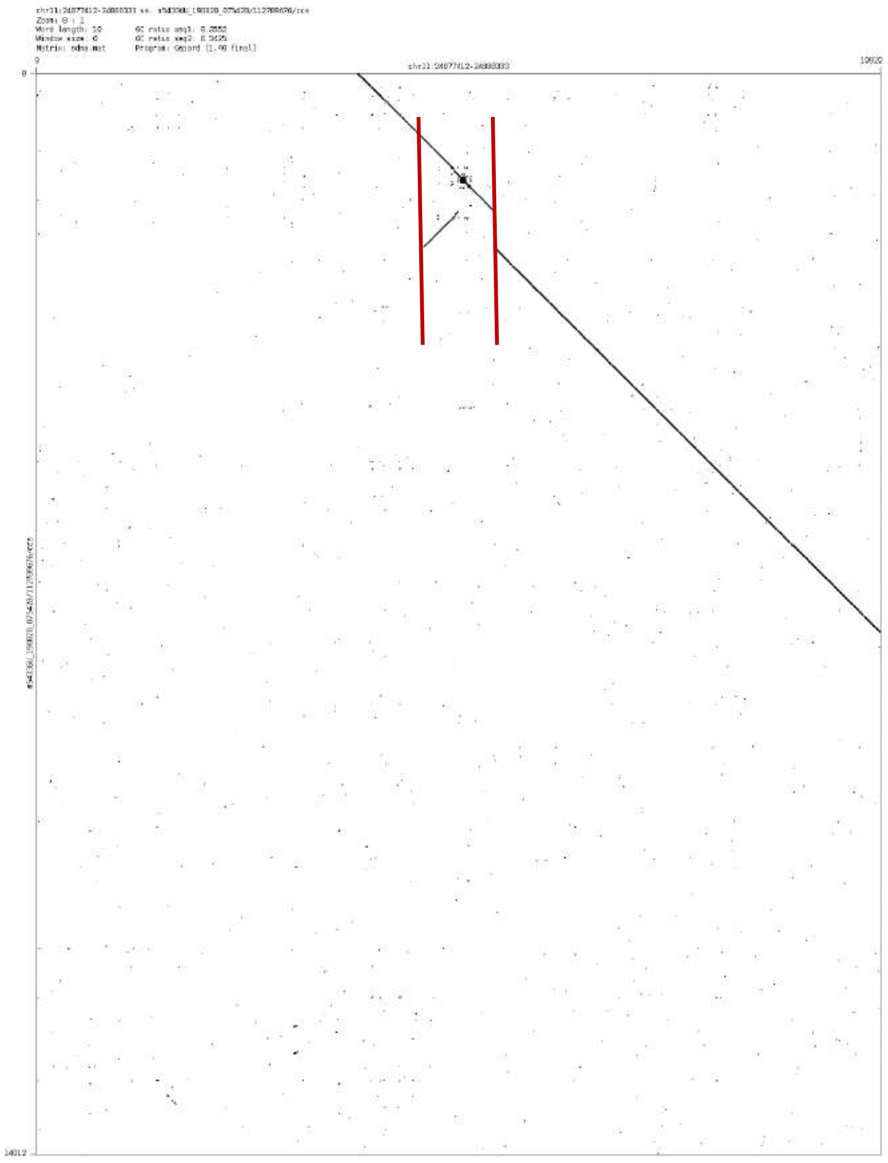
Zoom: 8 : 1
Word length: 10
Window size: 0
Matrix: edna.mat
Program: Gepard [1.40 final]
0
0
14012

chr11:99819304-99820578

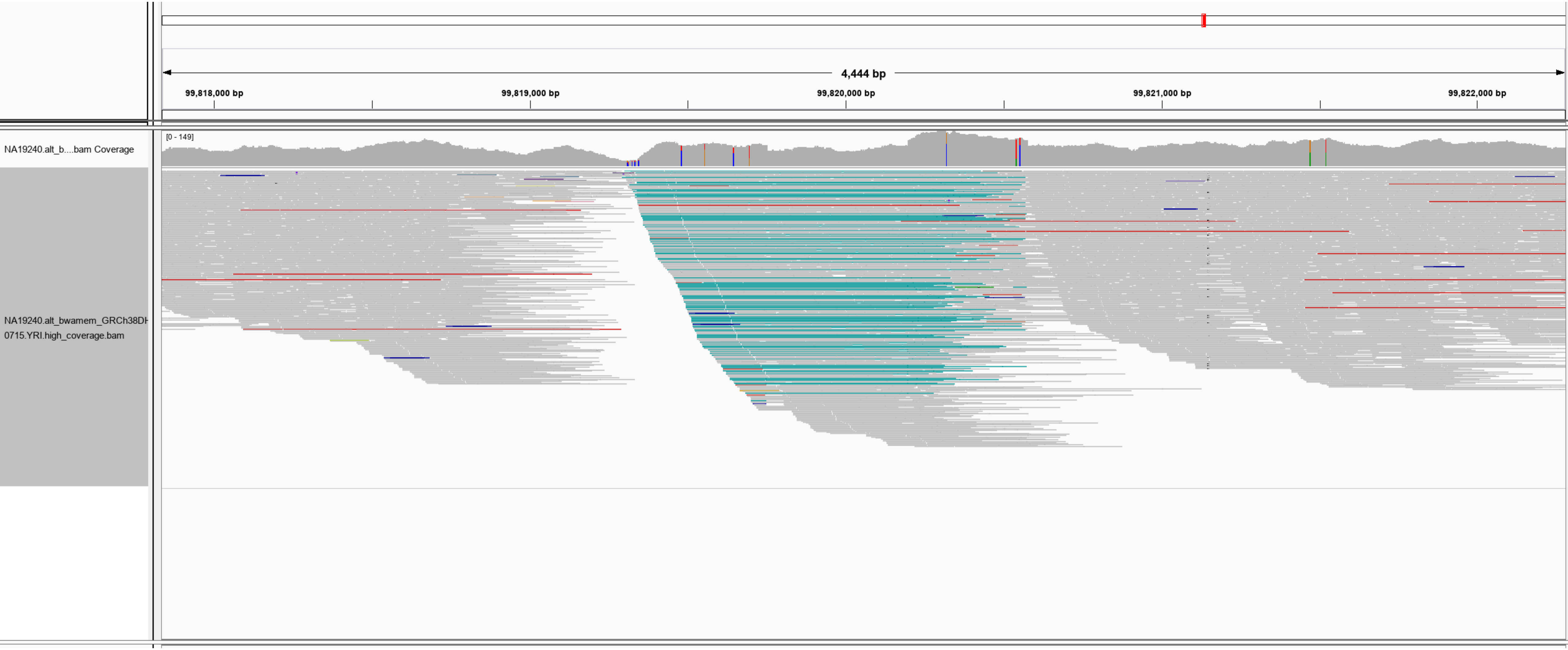

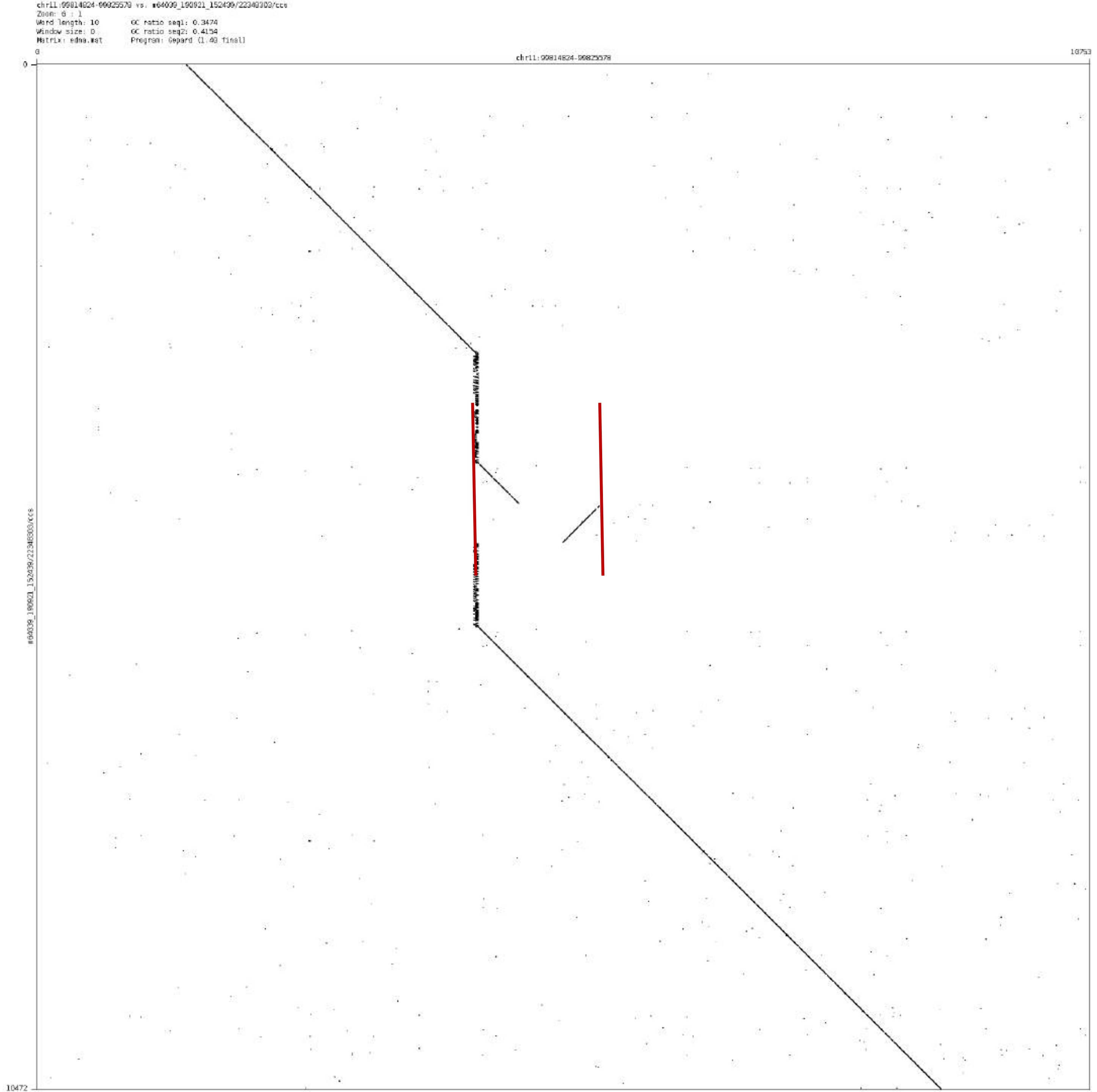

chr11:99814824-99825578 vs. m64039_190921_152439/22348303/ccs
Zoom: 6 : 1
Word length: 10
Window size: 0
Matrix: edna.mat
GC ratio seq1: 0.3474
GC ratio seq2: 0.4154
Program: Gepard (1.40 final)
chr11:99814824-99825578
0
10753
10472
m64039_190921_152439/22348303/ccs

### chr11:120061454-120062063

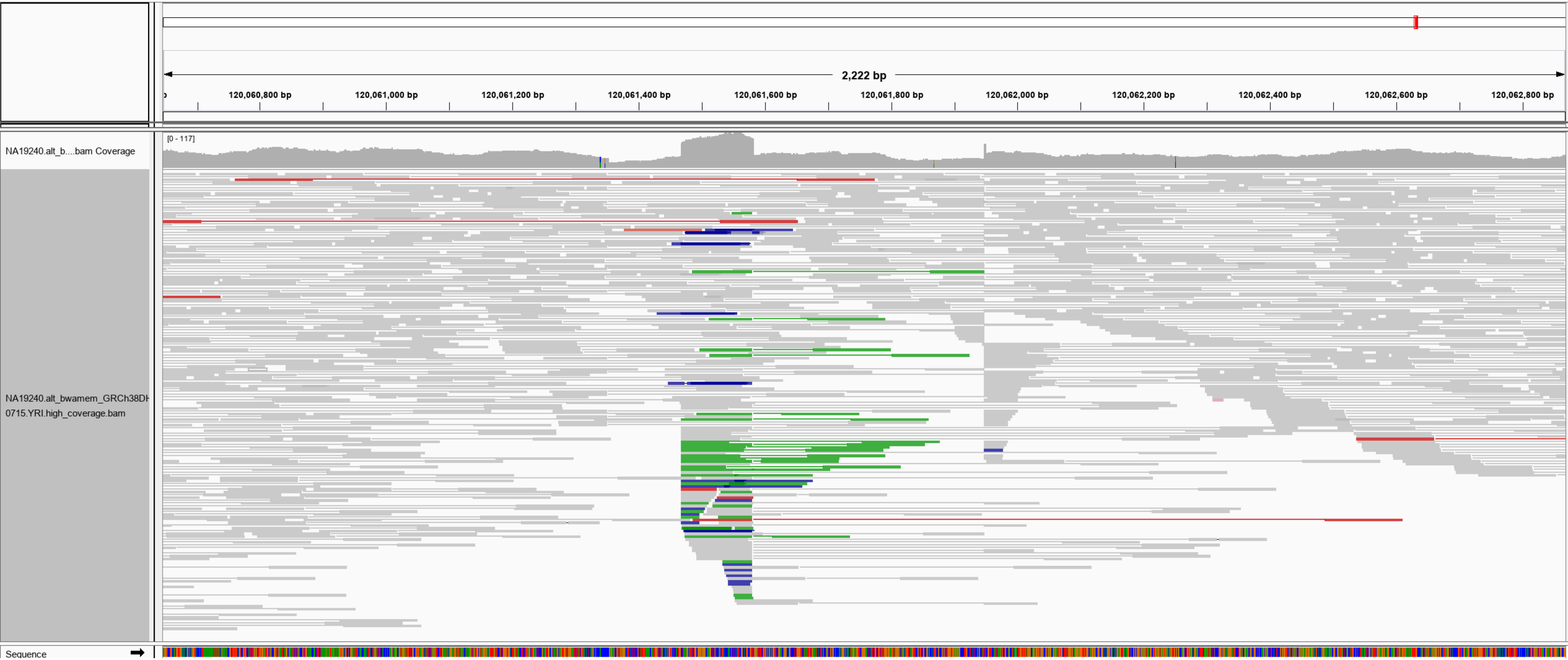

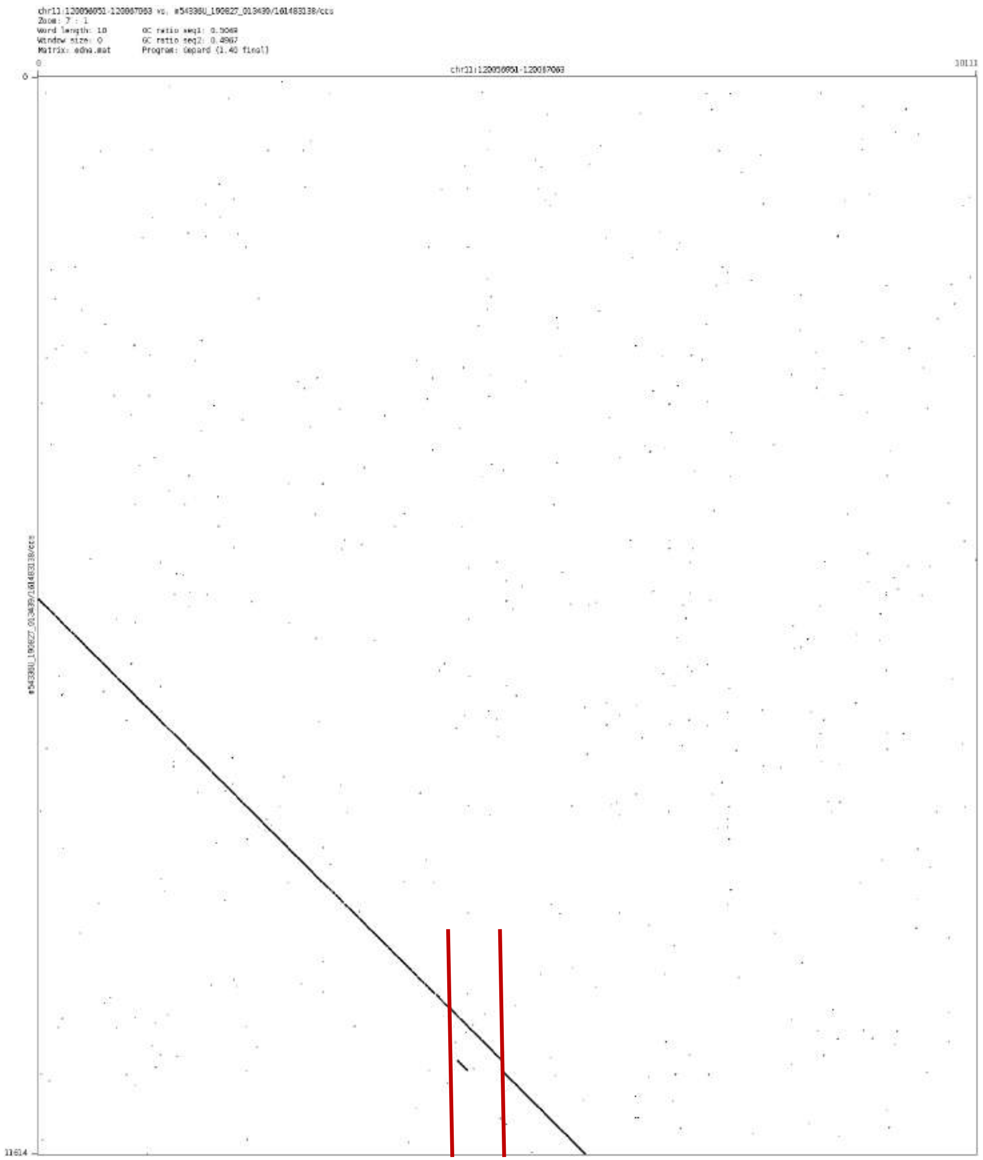
chr11:120056051-120067063 vs. m54336U_190827_013439/161483138/ccs
Zoom: 7 : 1
Word length: 10
Window size: 0
Matrix: edna.mat
GC ratio seq1: 0.5069
GC ratio seq2: 0.4967
Program: Gepard (1.40 final)
0
chr11:120056051-120067063
10111
0
11614

### chr11:129143073-129145383

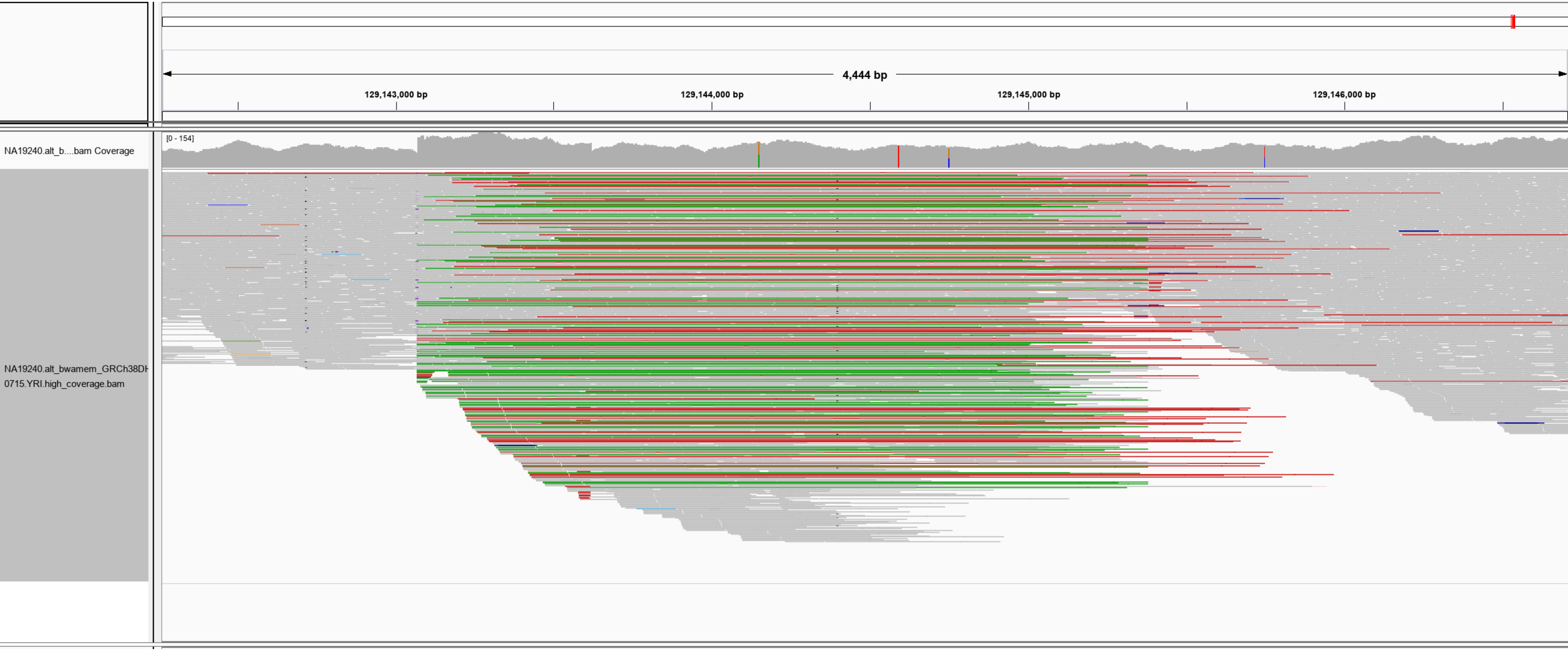

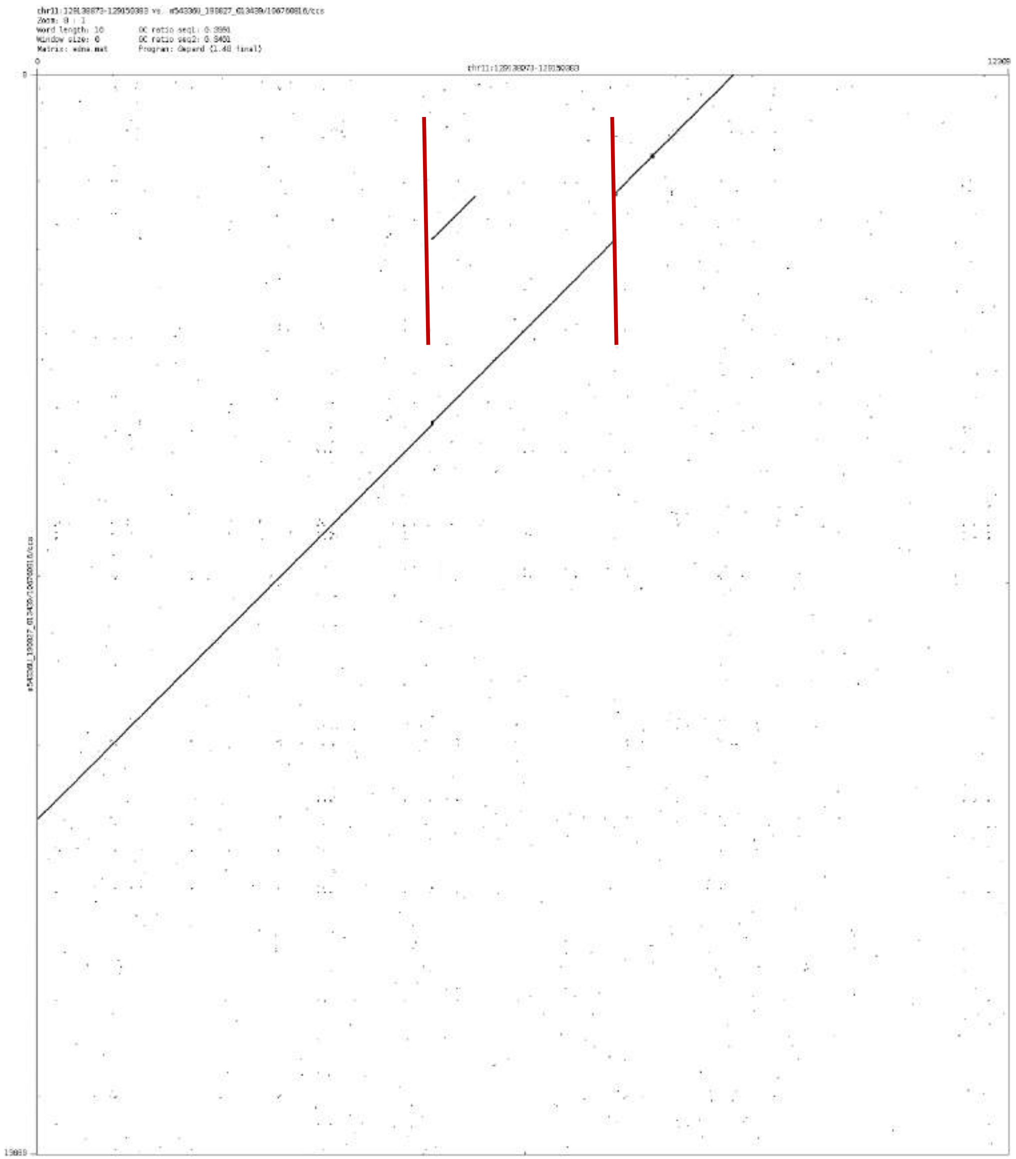

Zoom: 8 : 1
Word length: 10
Window size: 0
Matrix: edna.mat
Program: Gepard (1.40 final)

chr12:46634787-46639521

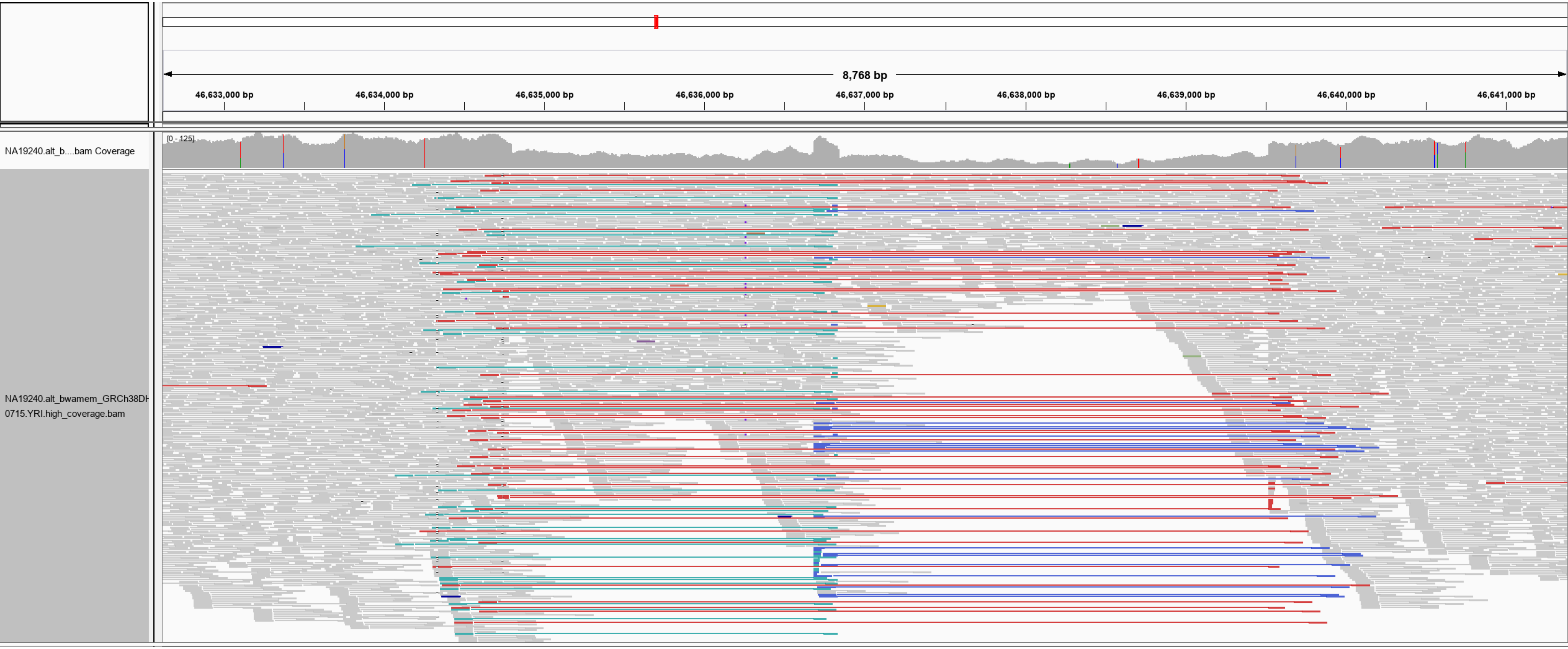

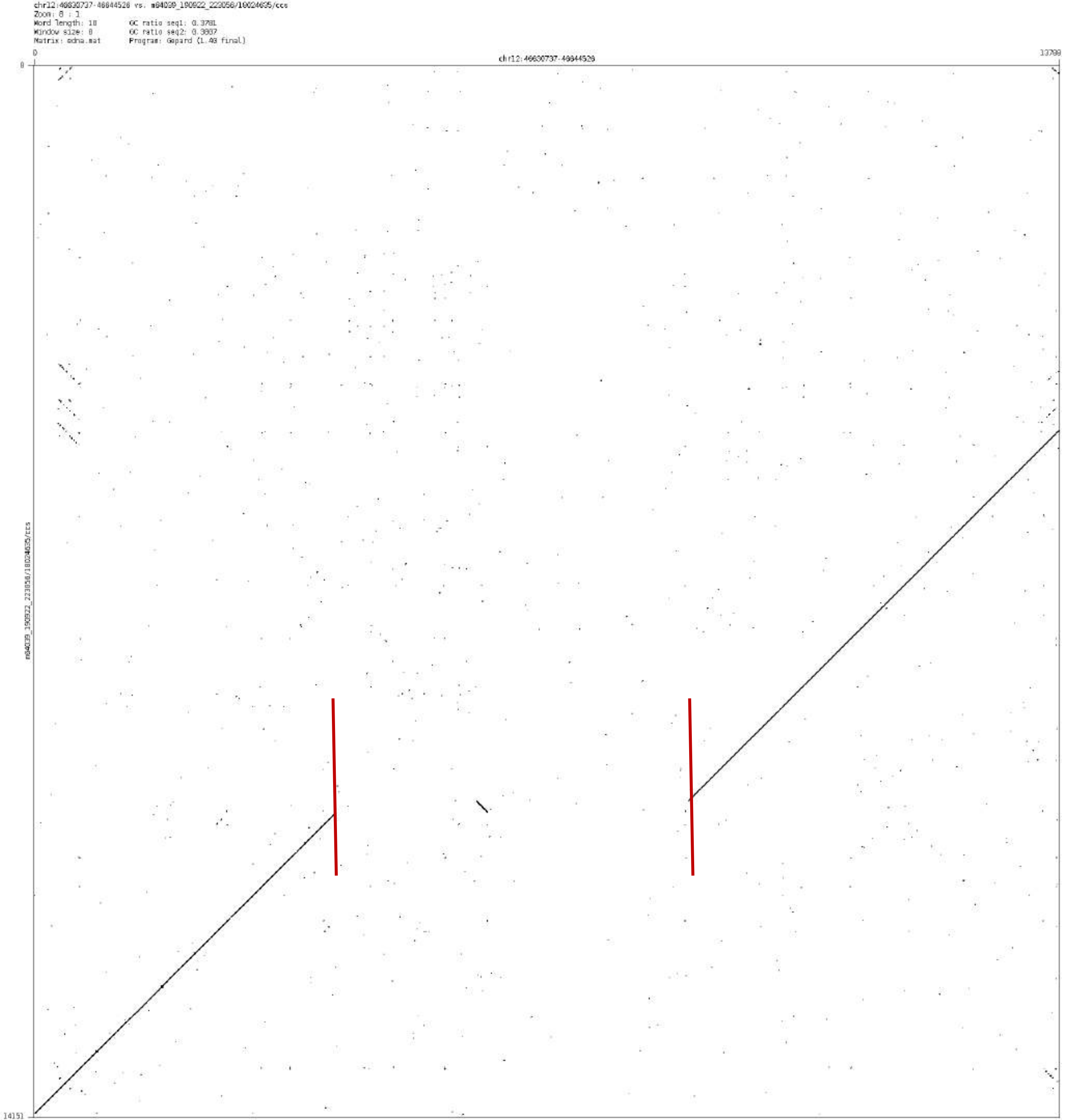
chr12:46630737-46644528 vs. m64039_190922_223056/10024635/ccs
Zoom: 8 : 1
Word length: 10
Window size: 0
Matrix: edna.mat
GC ratio seq1: 0.3781
GC ratio seq2: 0.3807
Program: Gepard (1.40 final)
0
chr12:46630737-46644528
13790
0
m64039_190922_223056/10024635/ccs
14151

### chr12:71139013-71139972

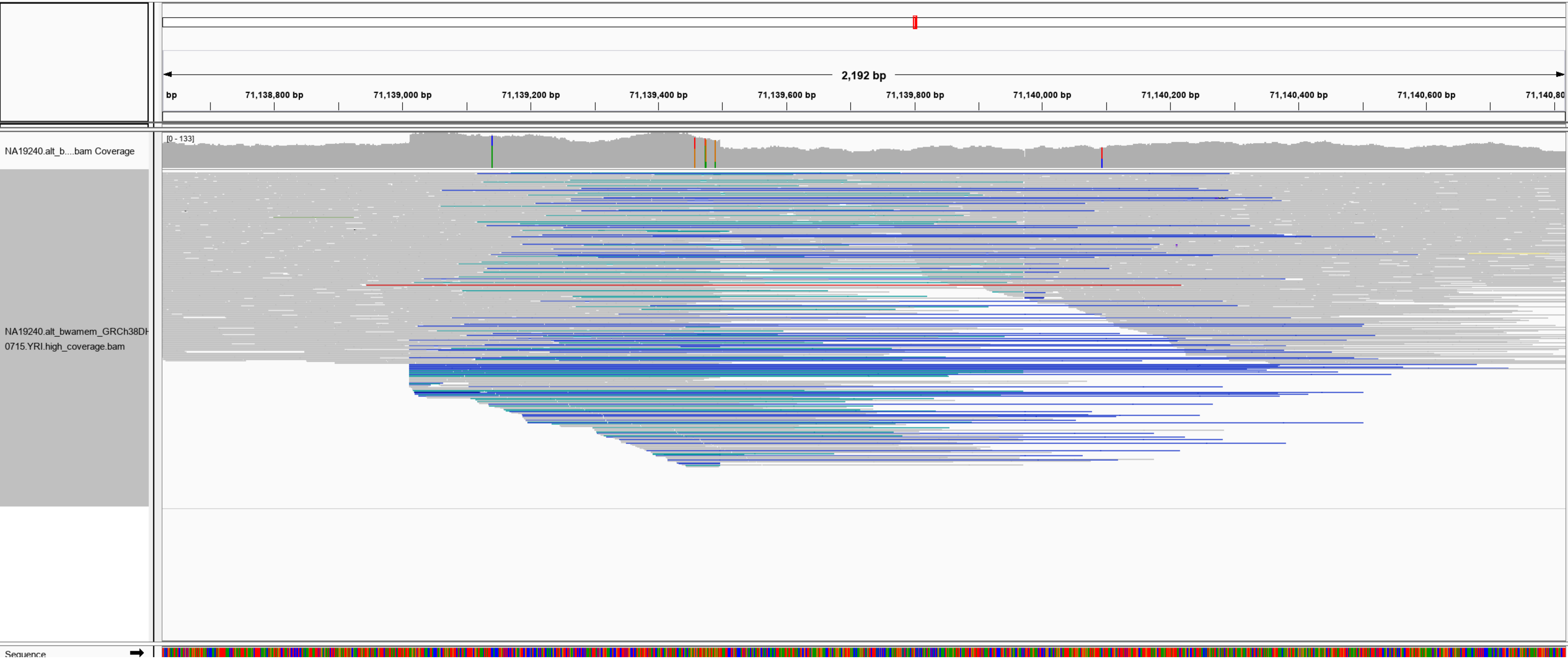

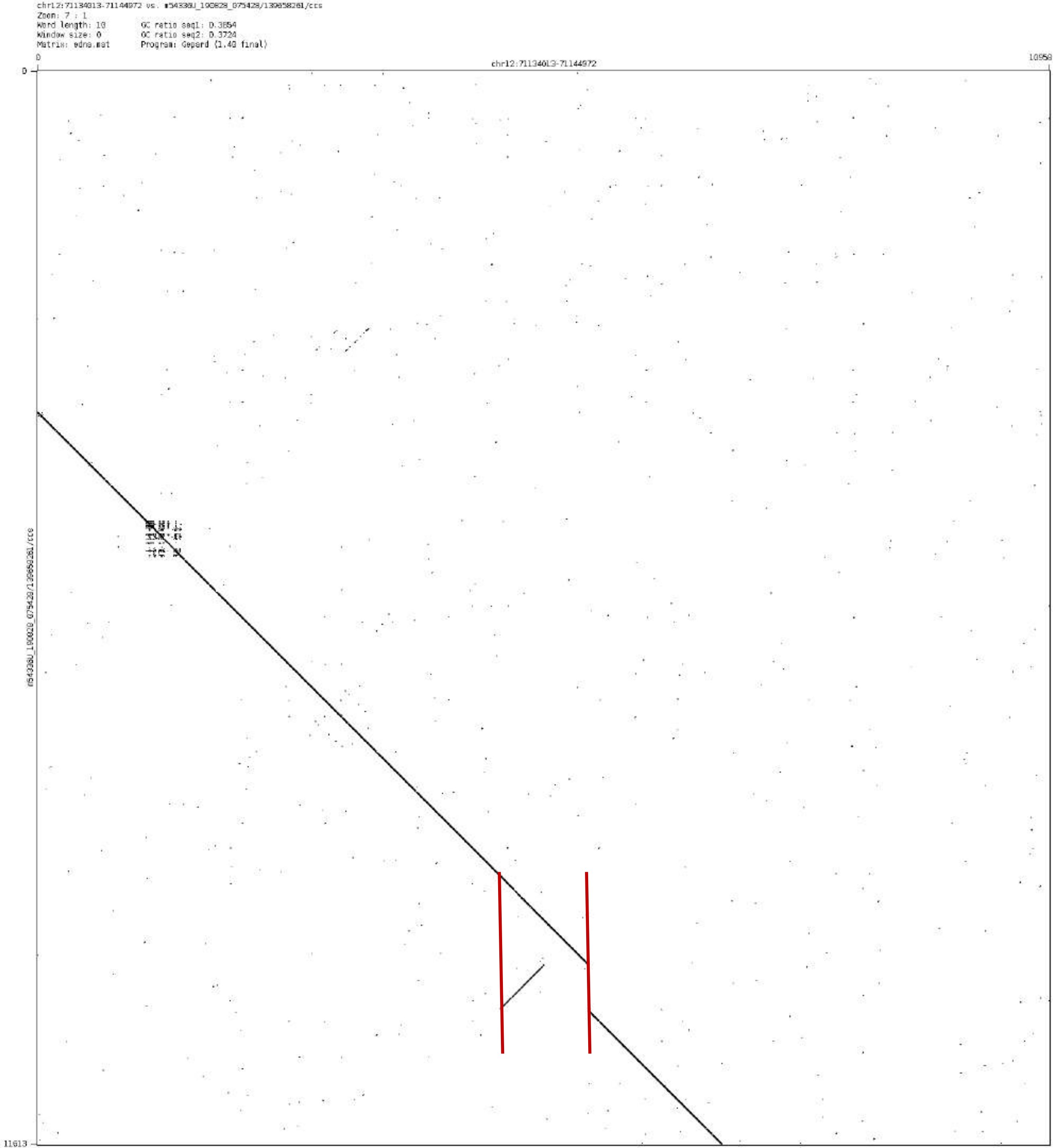
chr12:71134013-71144972 vs. m54338U_190828_075428/139858261/ccs
Zoom: 7 : 1
Word length: 10
Window size: 0
Matrix: edna.mat
GC ratio seq1: 0.3854
GC ratio seq2: 0.3724
Program: Gepard (1.40 final)
chr12:71134013-71144972
0
10958
11613

chr12:77990108-77994493

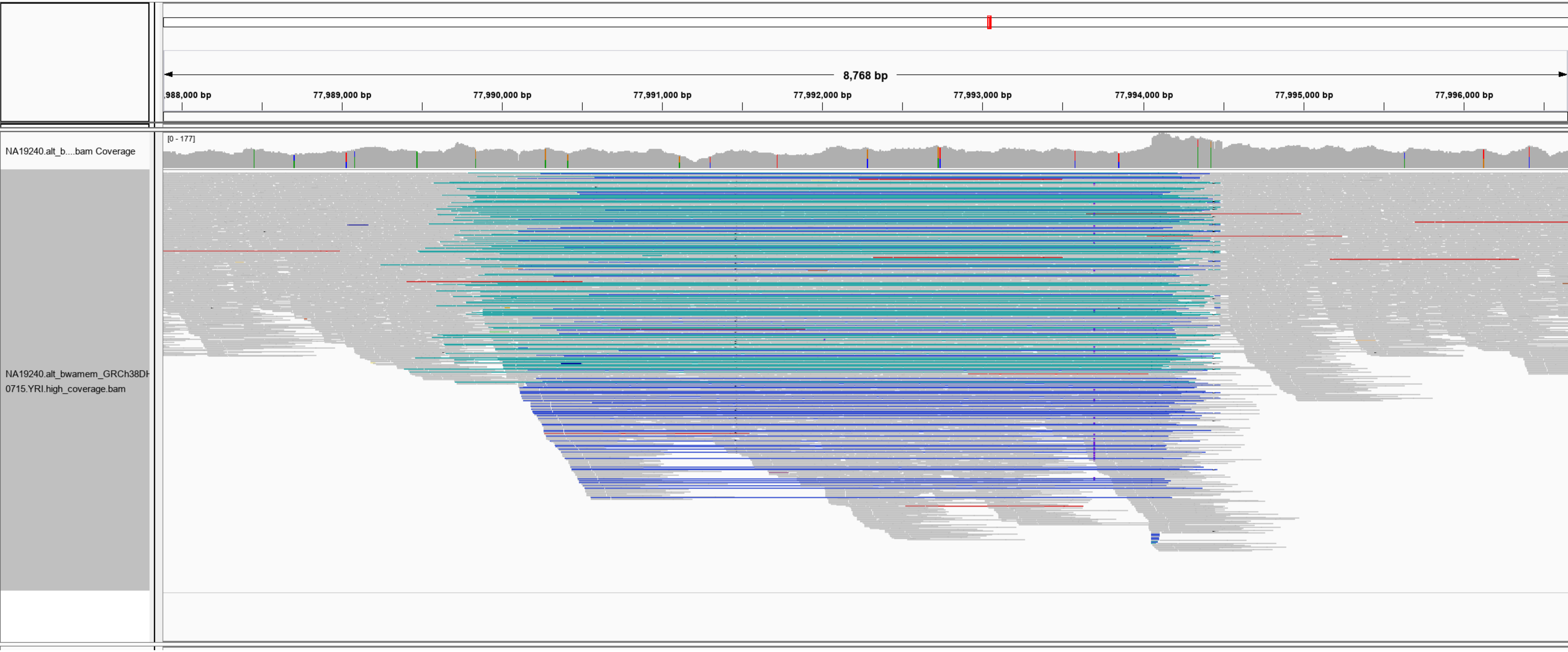

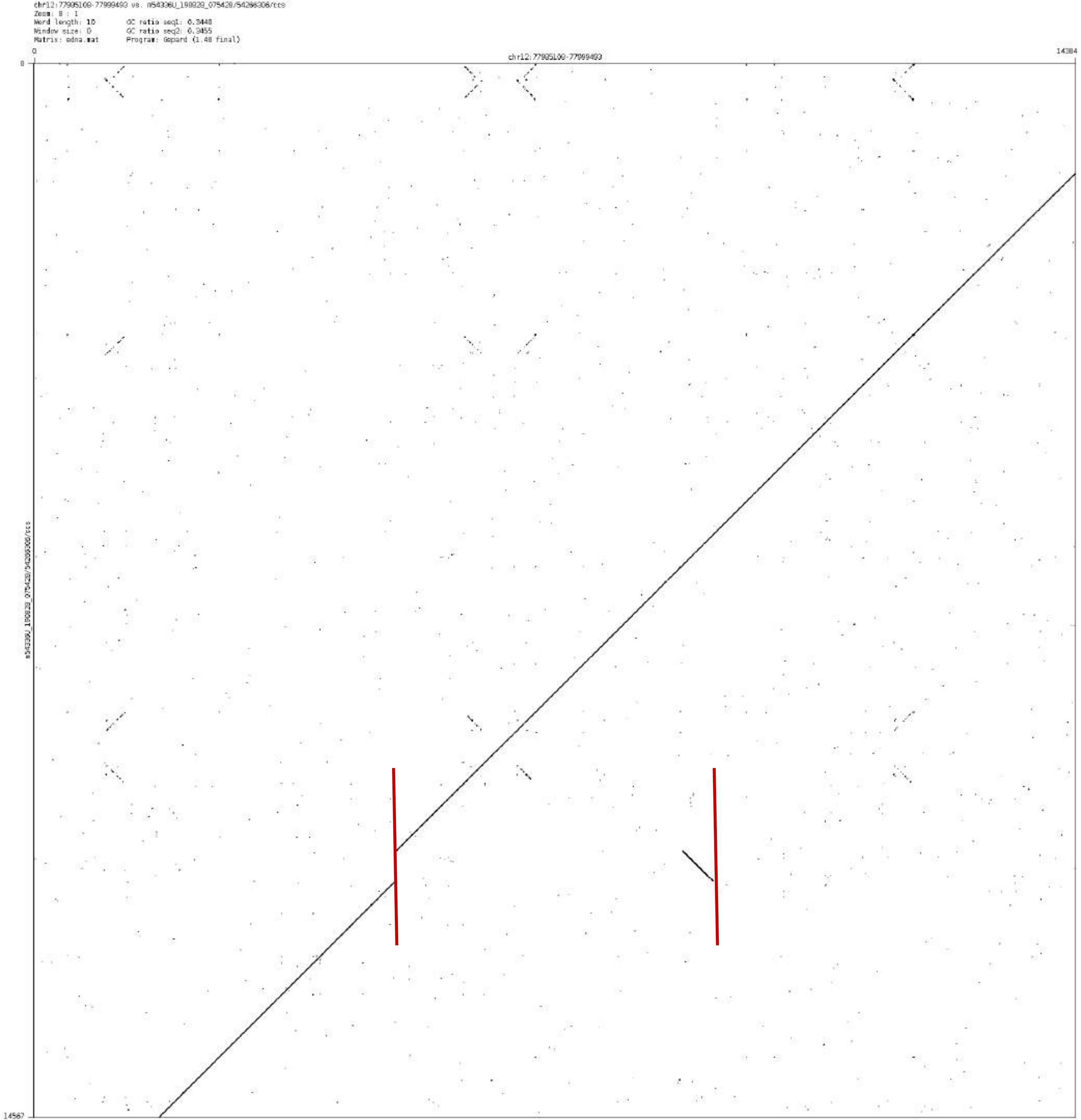
chr12:77995100-77999493 vs. m54336U_190829_075428/54206306/ccs
Zoom: 8 : 1
Word length: 10
Window size: 0
Matrix: edna.mat
GC ratio seq1: 0.3448
GC ratio seq2: 0.3455
Program: Gepard (1.40 final)
0
chr12:77995100-77999493
14384
0
m54336U_190829_075428/54206306/ccs
14567

### chr12:92745382-92748451

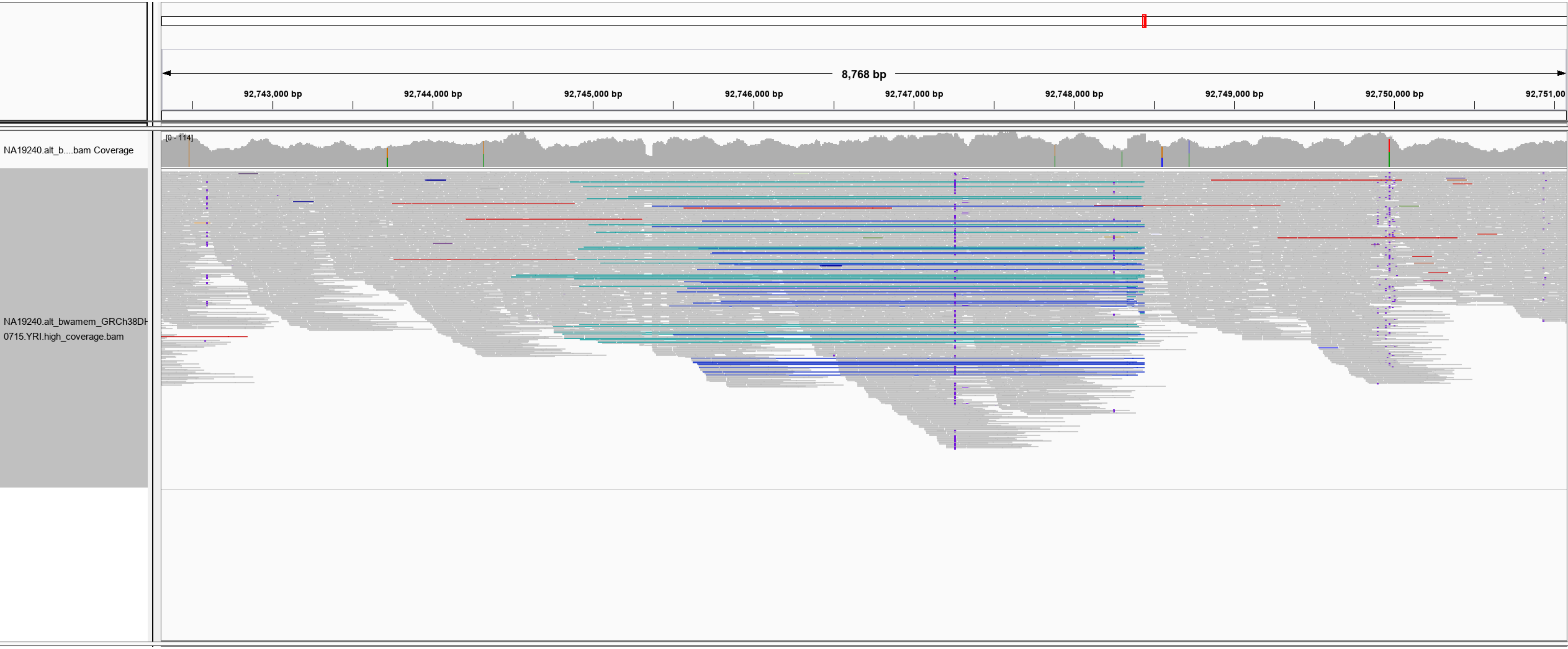

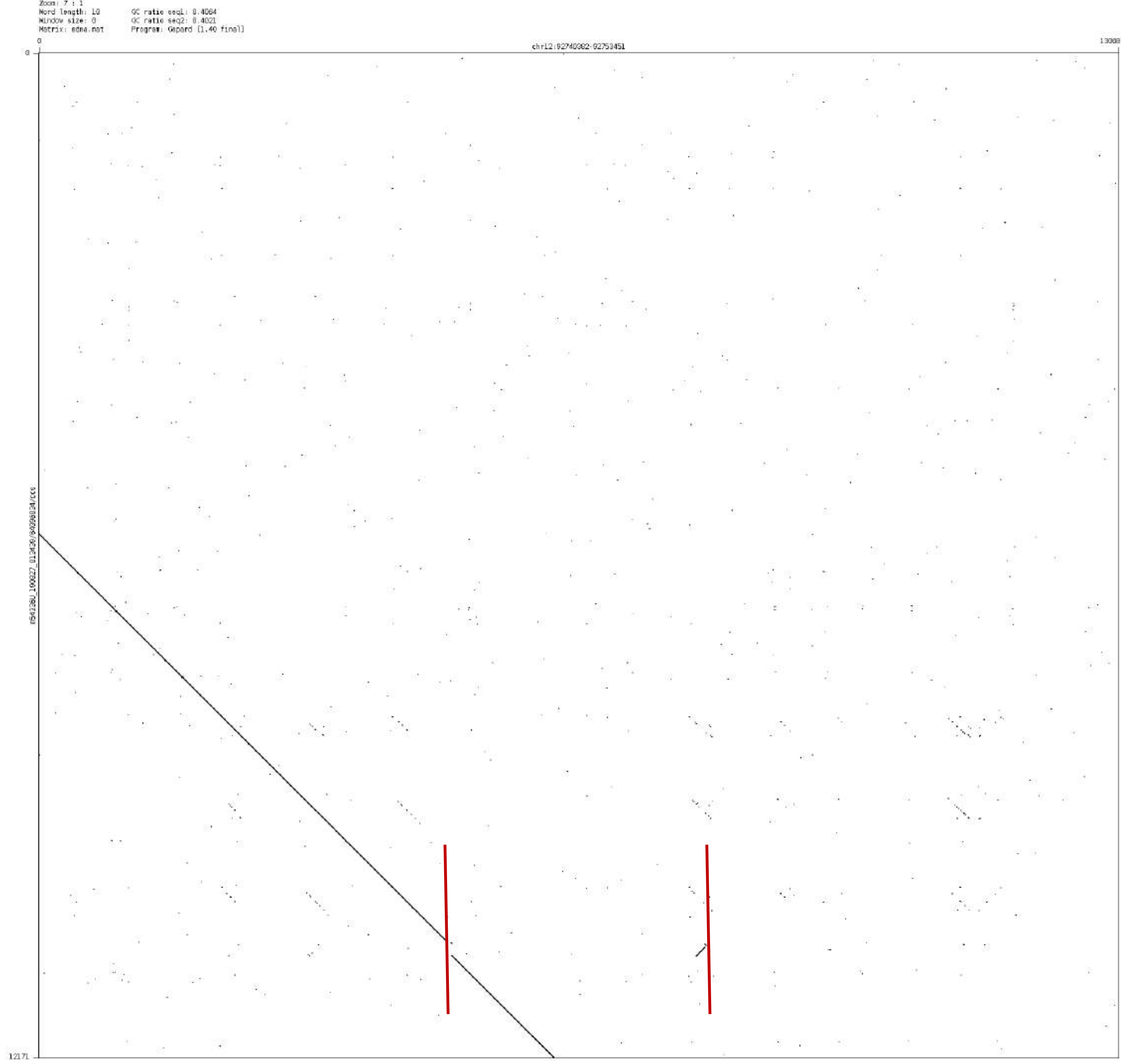

Zoom: 7 : 1
Word length: 10
Window size: 0
Matrix: edna.mat
GC ratio seq1: 0.4064
GC ratio seq2: 0.4021
Program: Gepard [1.40 final]
0
13088
chr12:92740382-92753451
12171

chr12:95558627-95559959

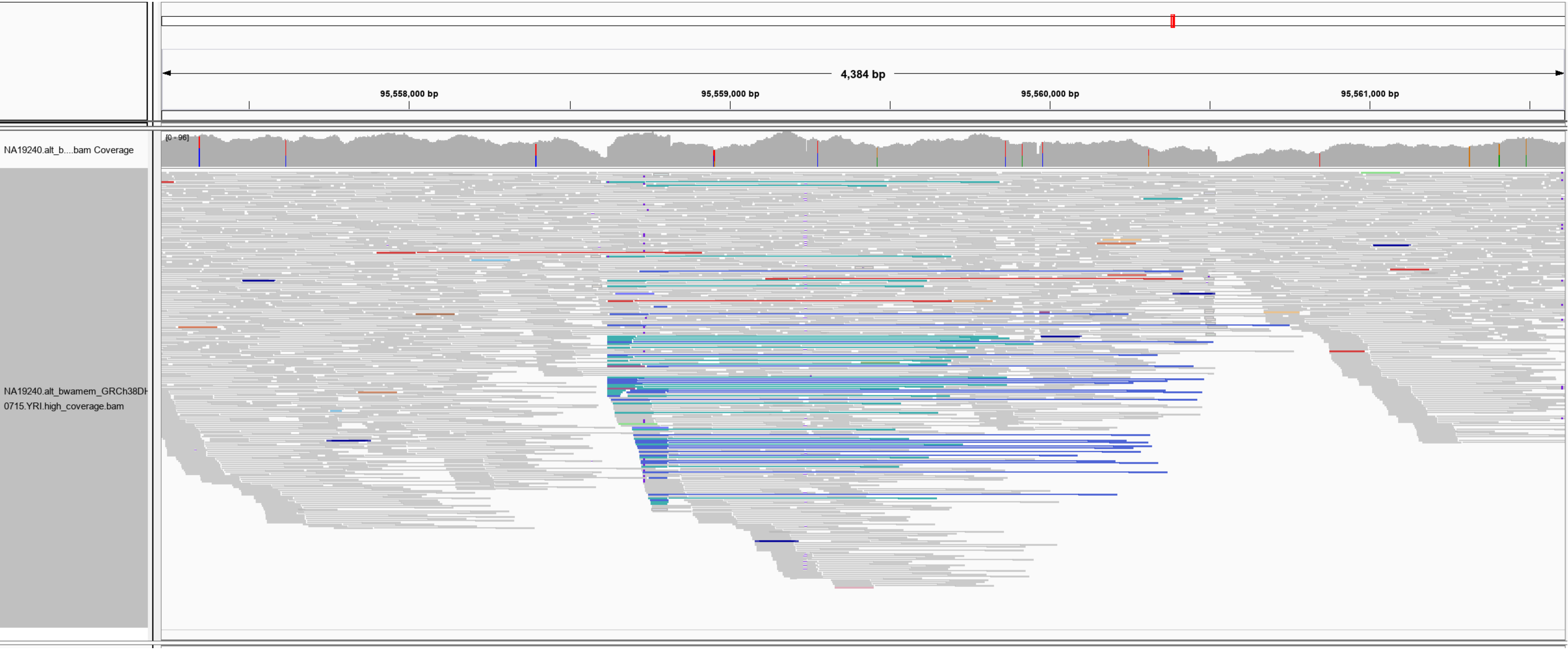

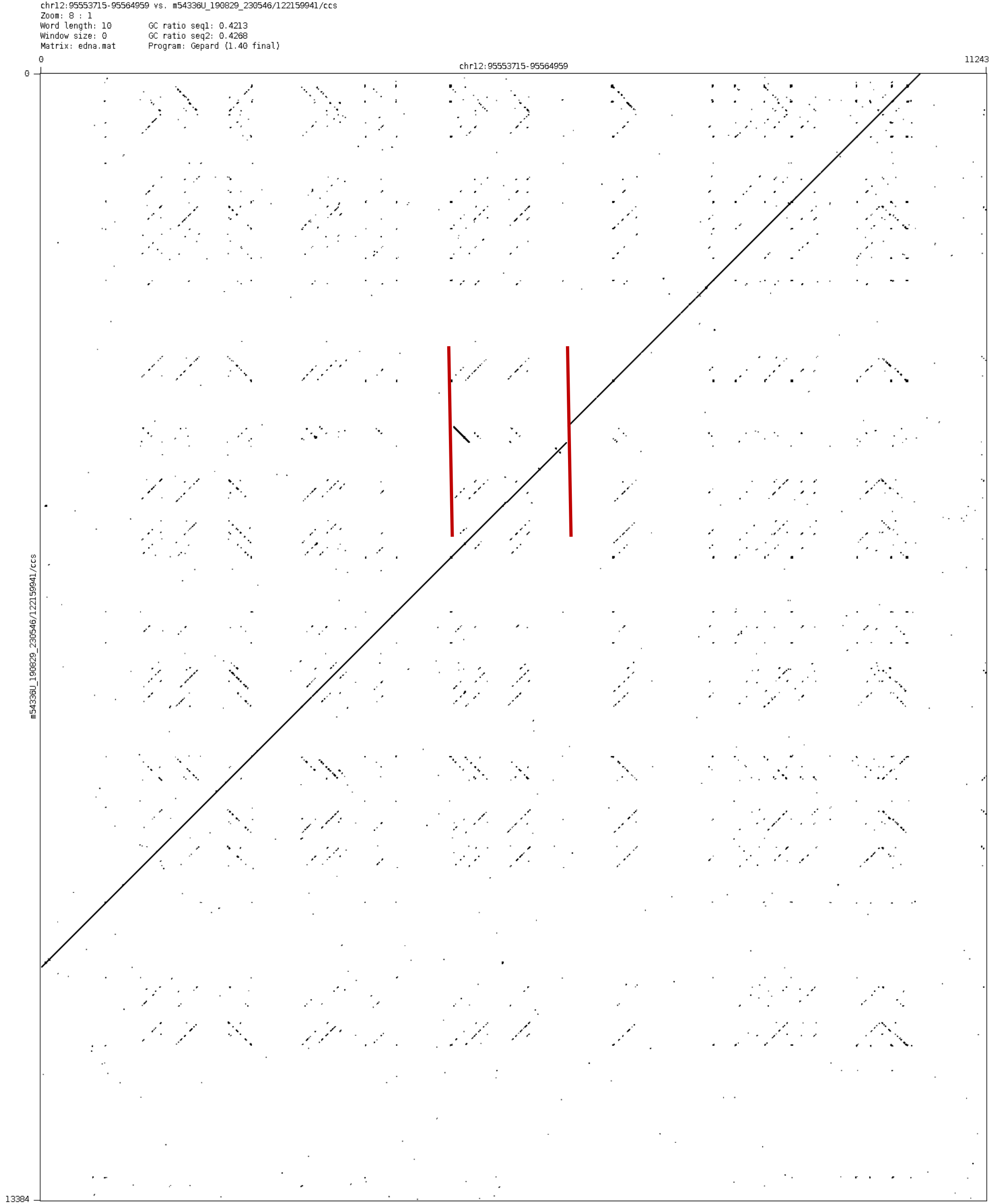
chr12:95553715-95564959 vs. m54336U_190829_230546/122159941/ccs
Zoom: 8 : 1
Word length: 10
Window size: 0
Matrix: edna.mat
GC ratio seq1: 0.4213
GC ratio seq2: 0.4268
Program: Gepard (1.40 final)
0
11243
chr12:95553715-95564959
0
13384
m54336U_190829_230546/122159941/ccs

chr14:20082944-20087060

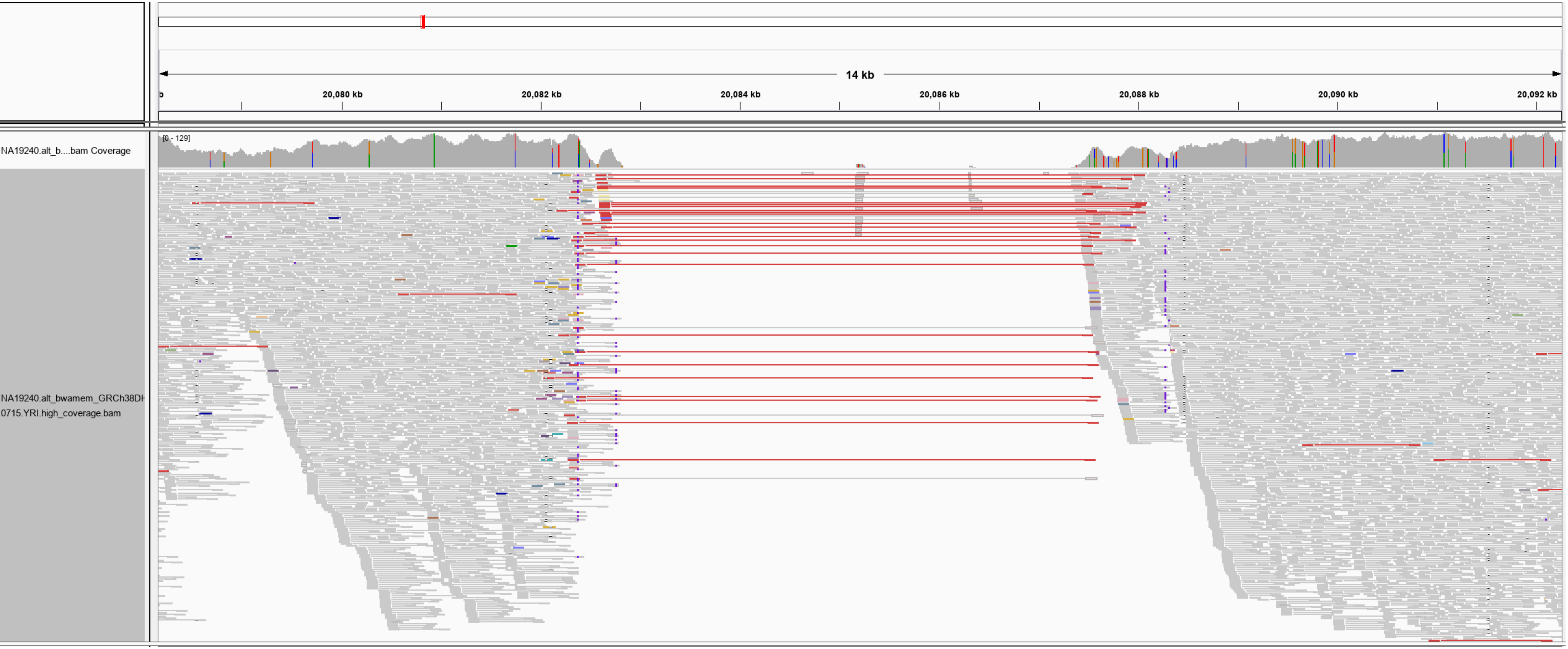

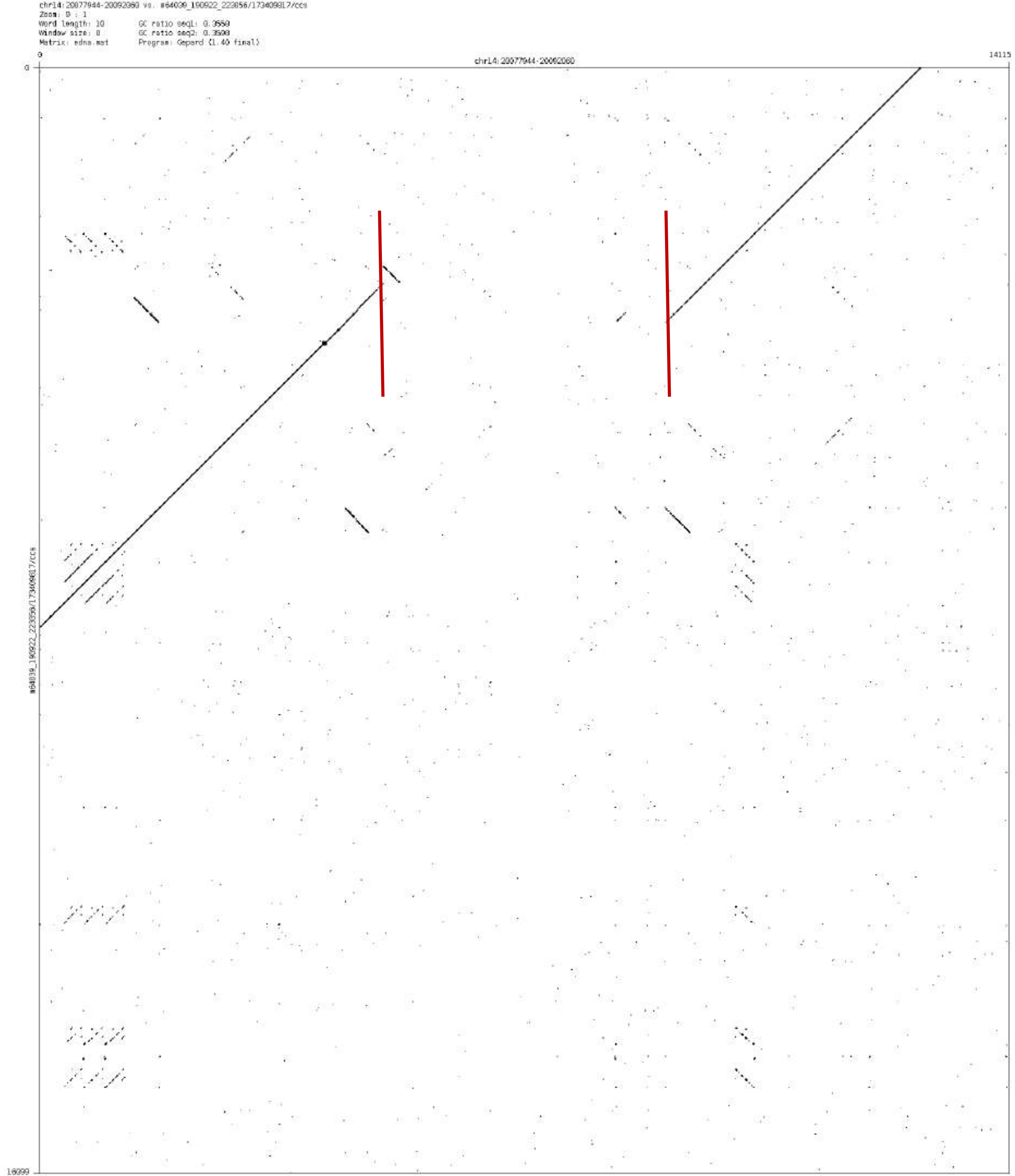

chr14:20077944-20092060 vs. m64039_190922_223056/173408817/ccs
Zoom: 0 : 1
Word length: 10
Window size: 0
Matrix: edna.mat
GC ratio seq1: 0.3558
GC ratio seq2: 0.3508
Program: Gepard (1.40 final)
0
14115
chr14:20077944-20092060
16099
m64039_190922_223056/173408817/ccs

### chr14:22636224-22640024

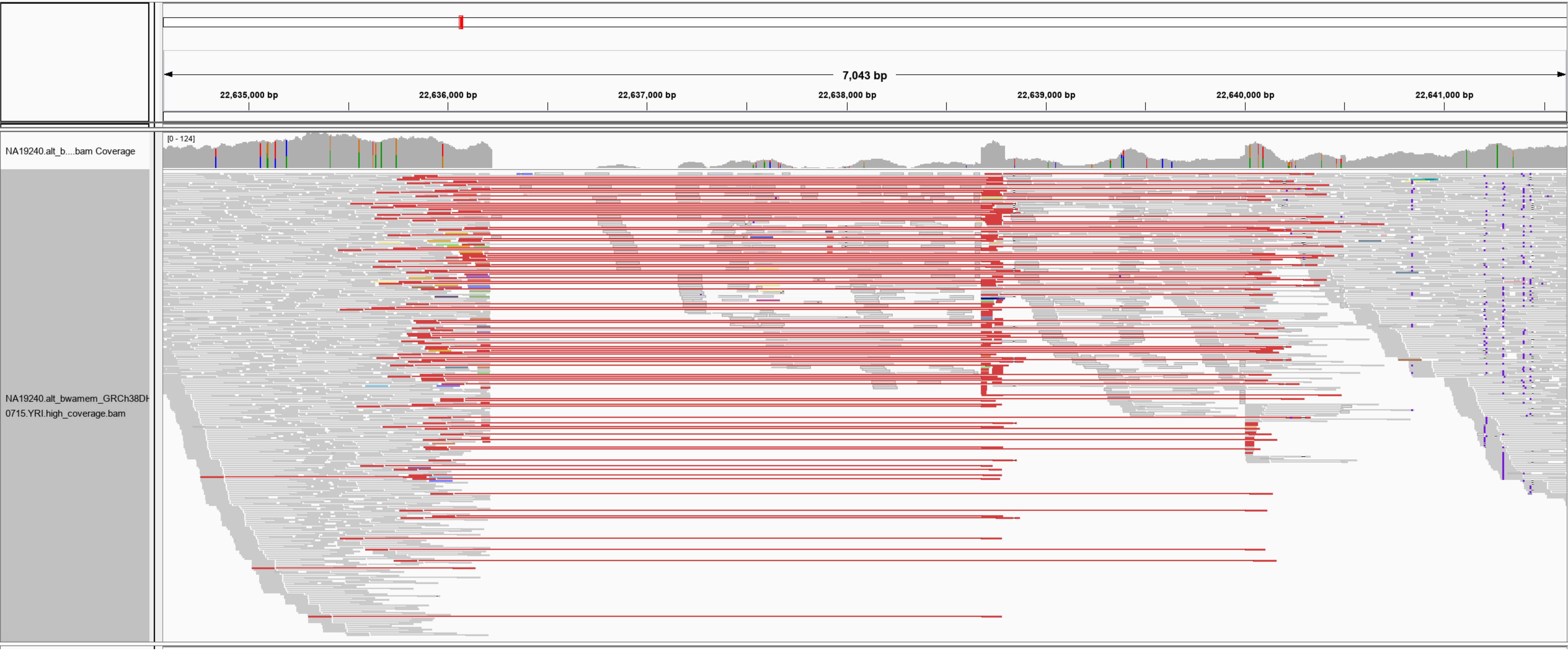

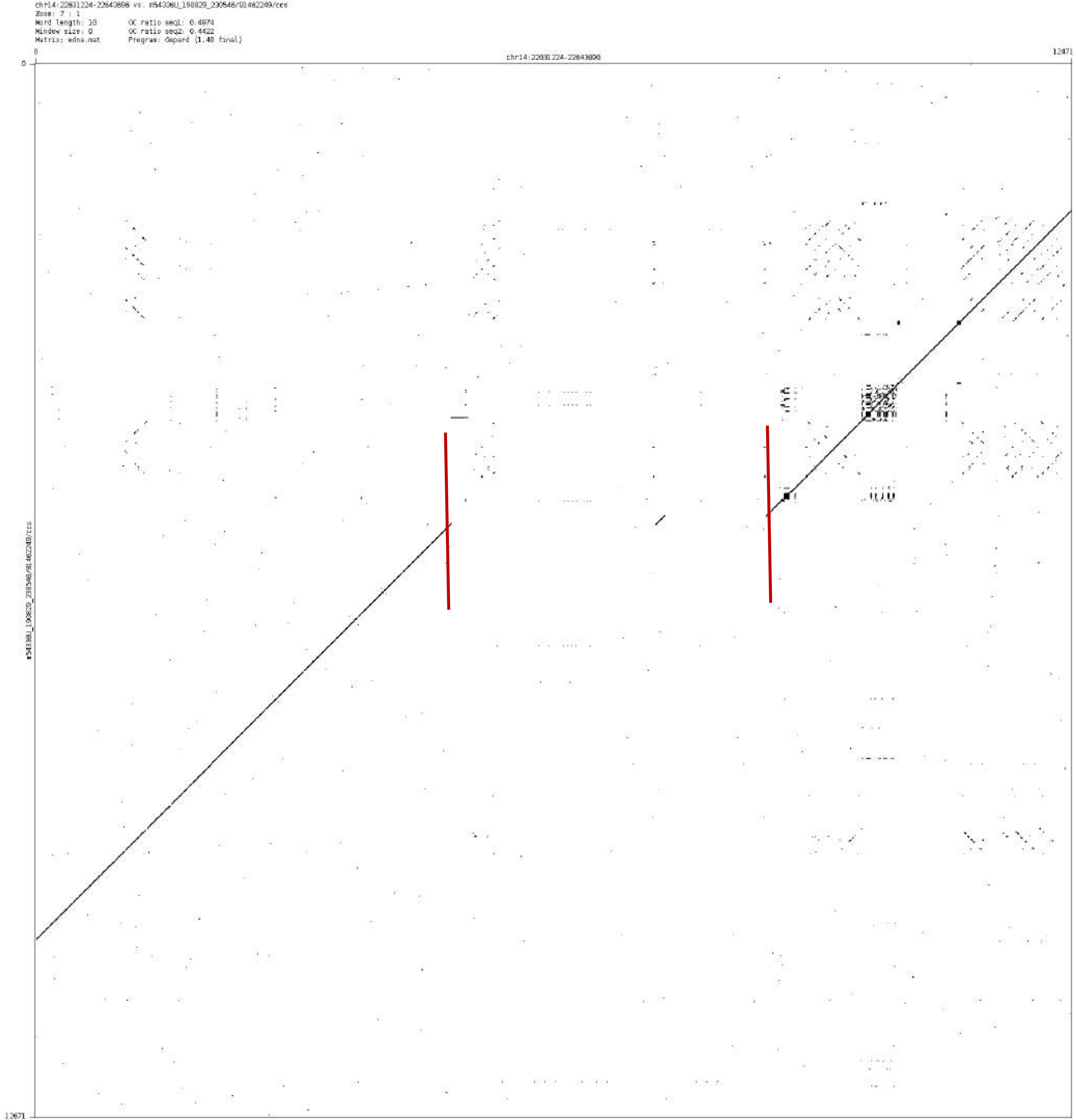
chr14:22631224-22643696 vs. m54336U_190829_230546/81462249/ccs
Zoom: 7 : 1
Word length: 10
Window size: 0
Matrix: edna.mat
GC ratio seq1: 0.4974
GC ratio seq2: 0.4422
Program: Gepard (1.40 final)
0
chr14:22631224-22643696
12471
0
12671

chr14:65375854-65376595

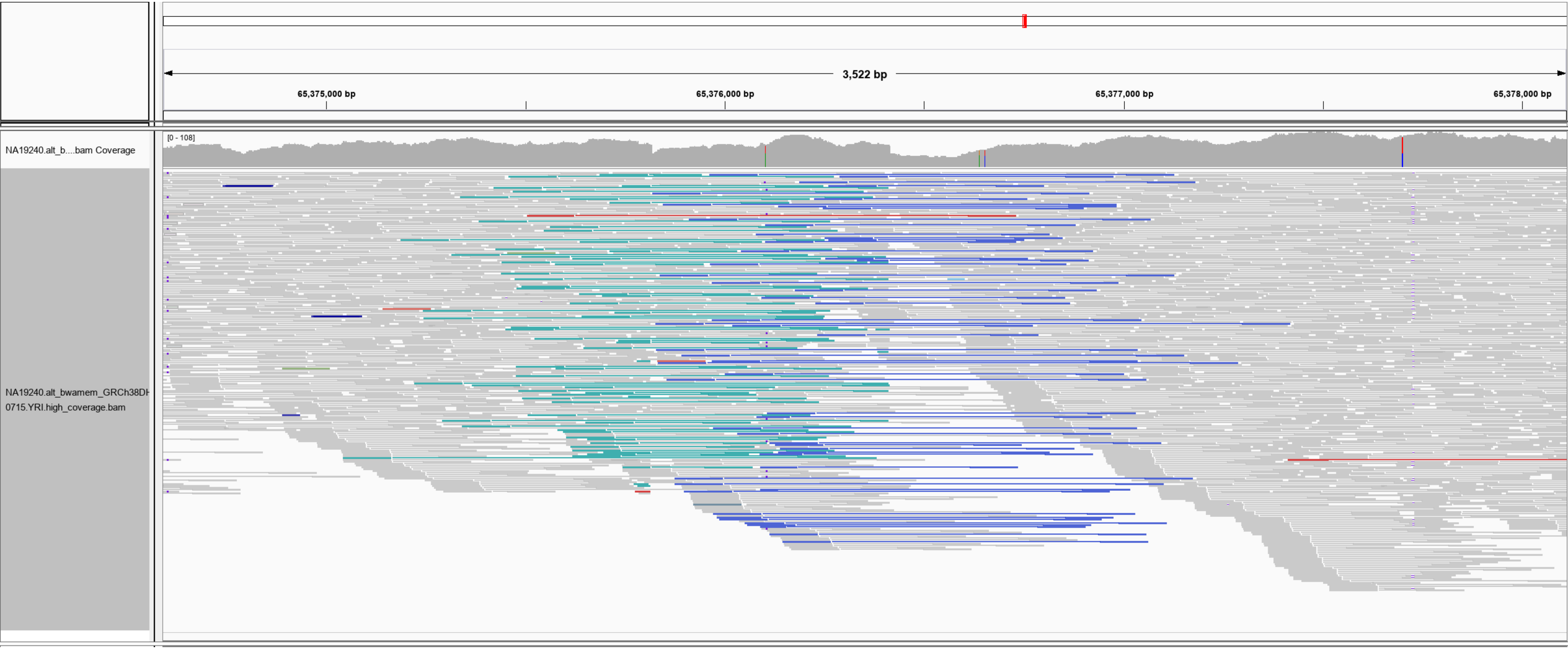

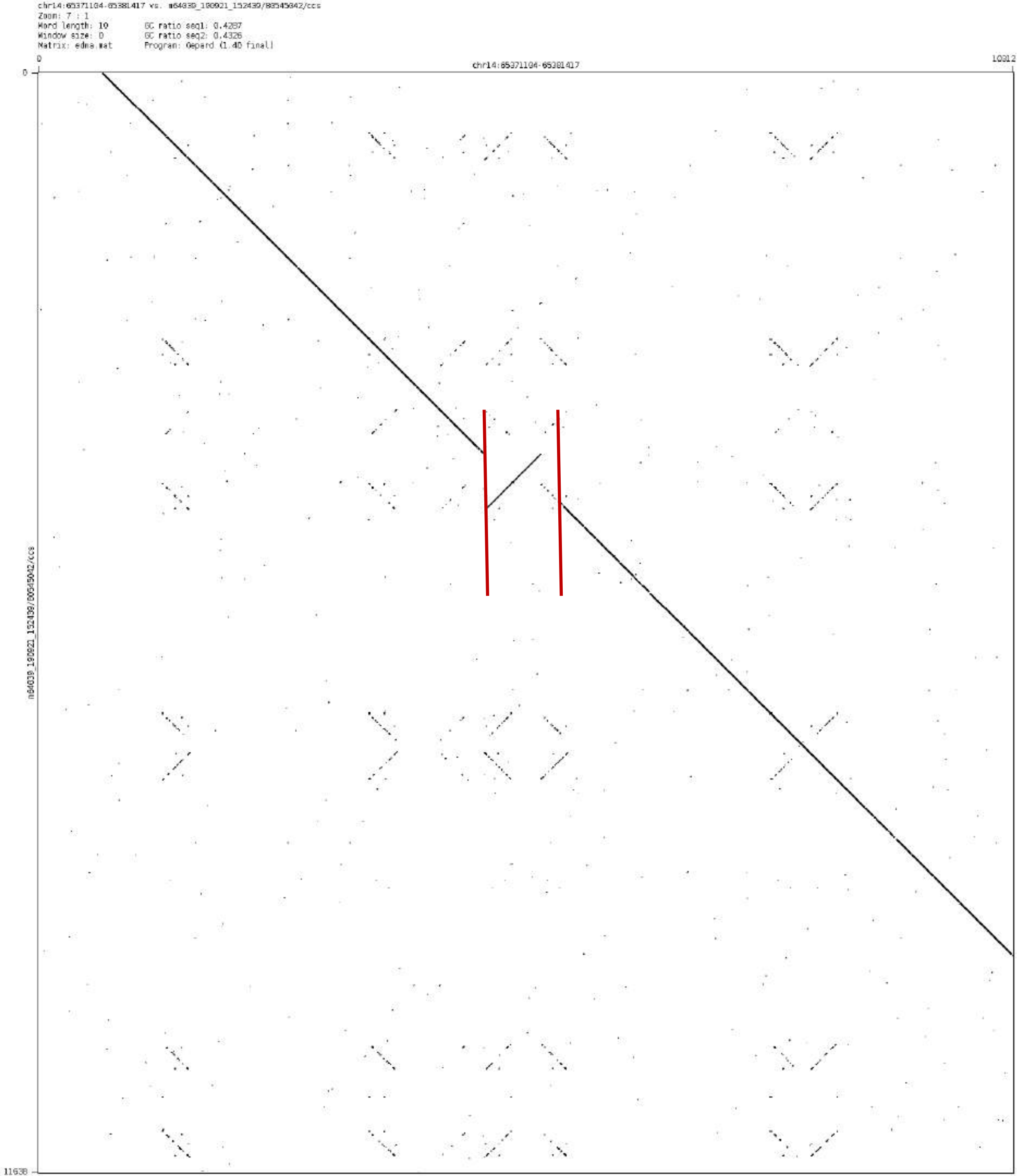
chr14:65371104-65381417 vs. m64039_190921_152439/80545342/ccs
Zoom: 7 : 1
Word length: 10
Window size: 0
Matrix: edna.mat
GC ratio seq1: 0.4287
GC ratio seq2: 0.4326
Program: Gepard (1.40 final)
0
chr14:65371104-65381417
10312
0
11638

chr14:66703613-66705301

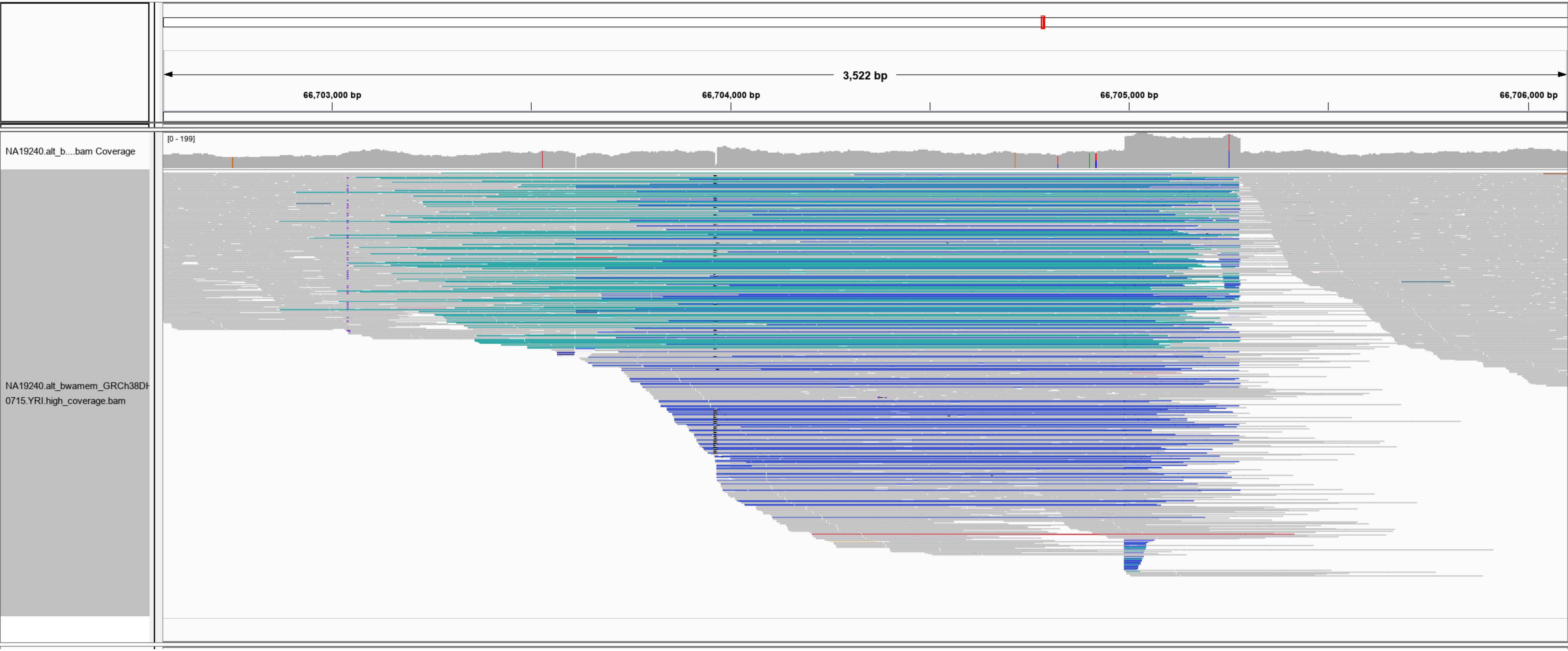

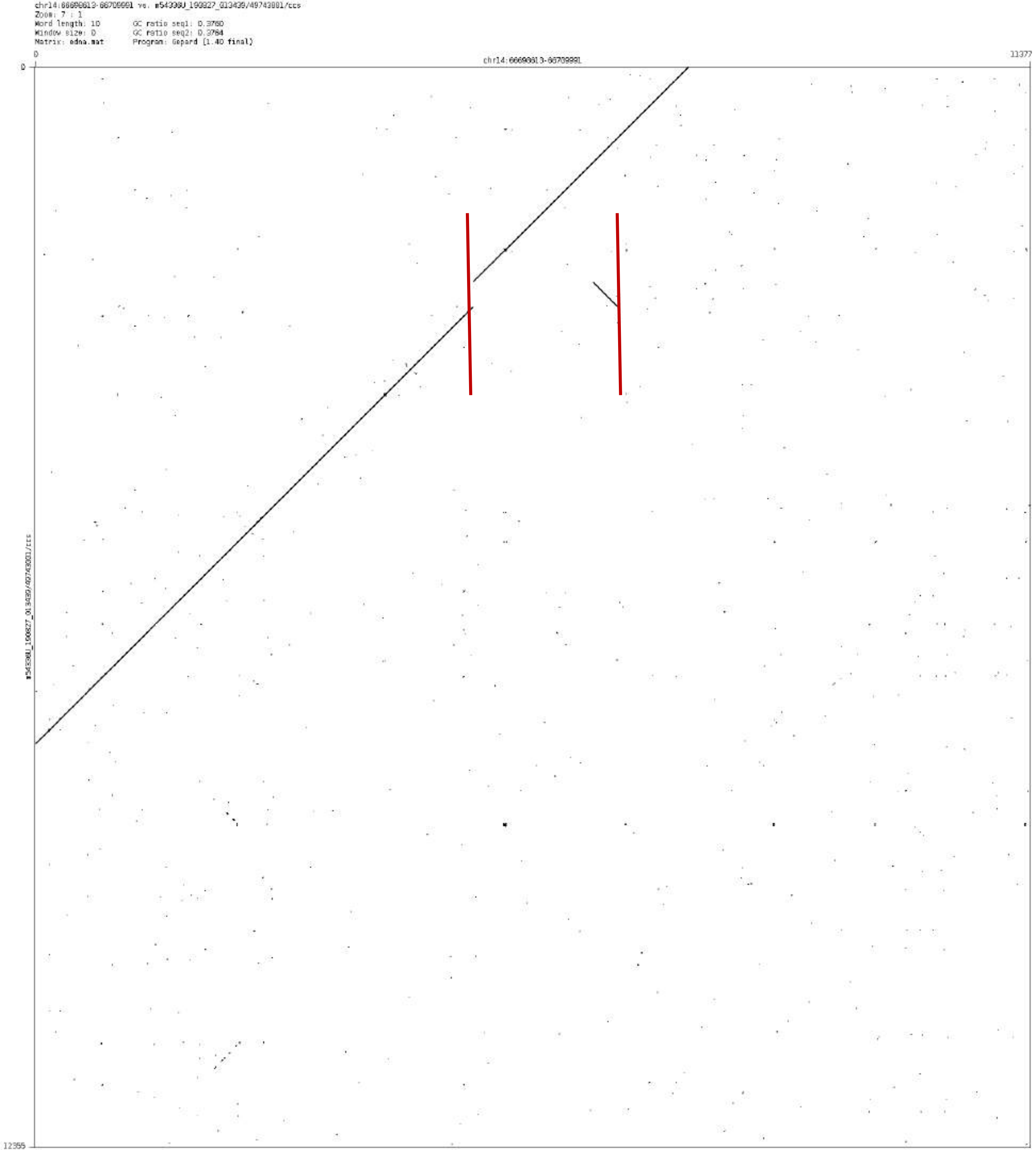
Zoom: 7 : 1
Word length: 10
Window size: 0
Matrix: edna.mat
GC ratio seq1: 0.3760
GC ratio seq2: 0.3764
Program: Gepard (1.40 final)
0
11377
12355

chr15:95148441-95149540

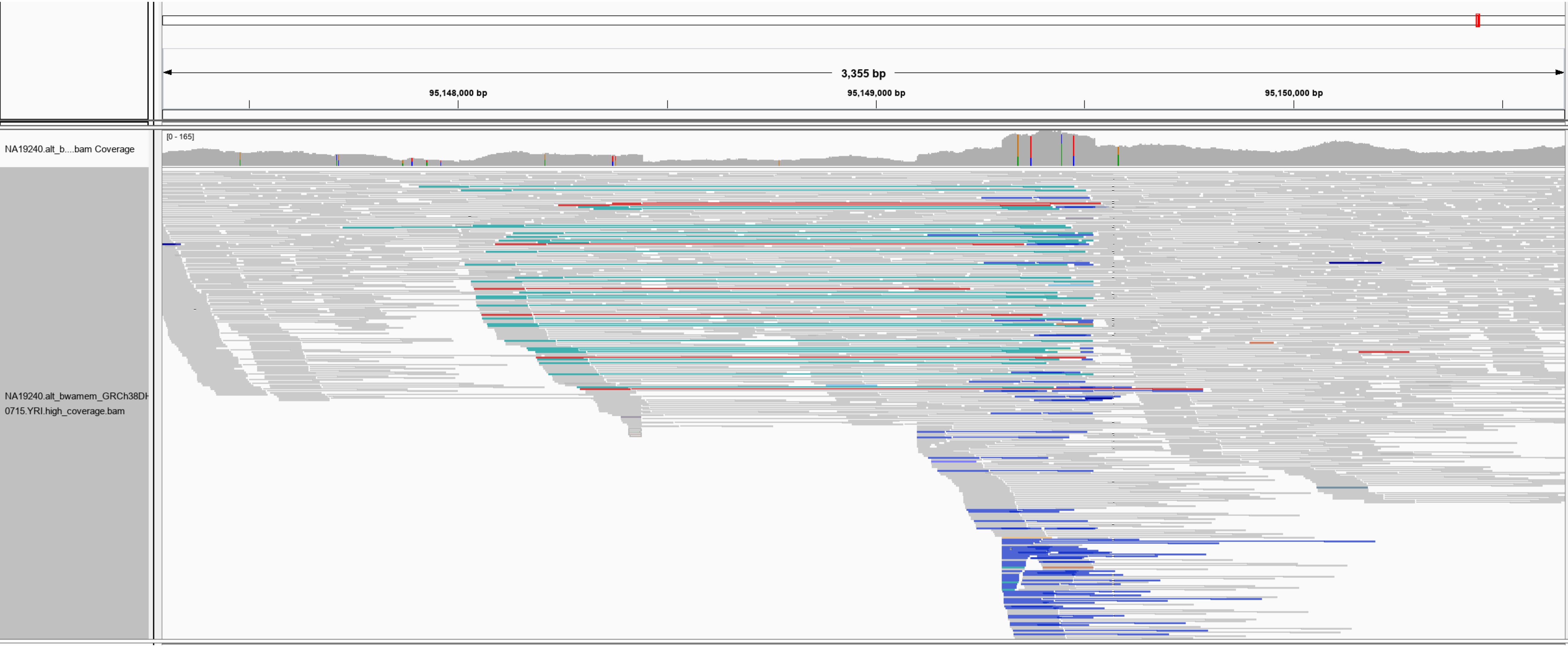

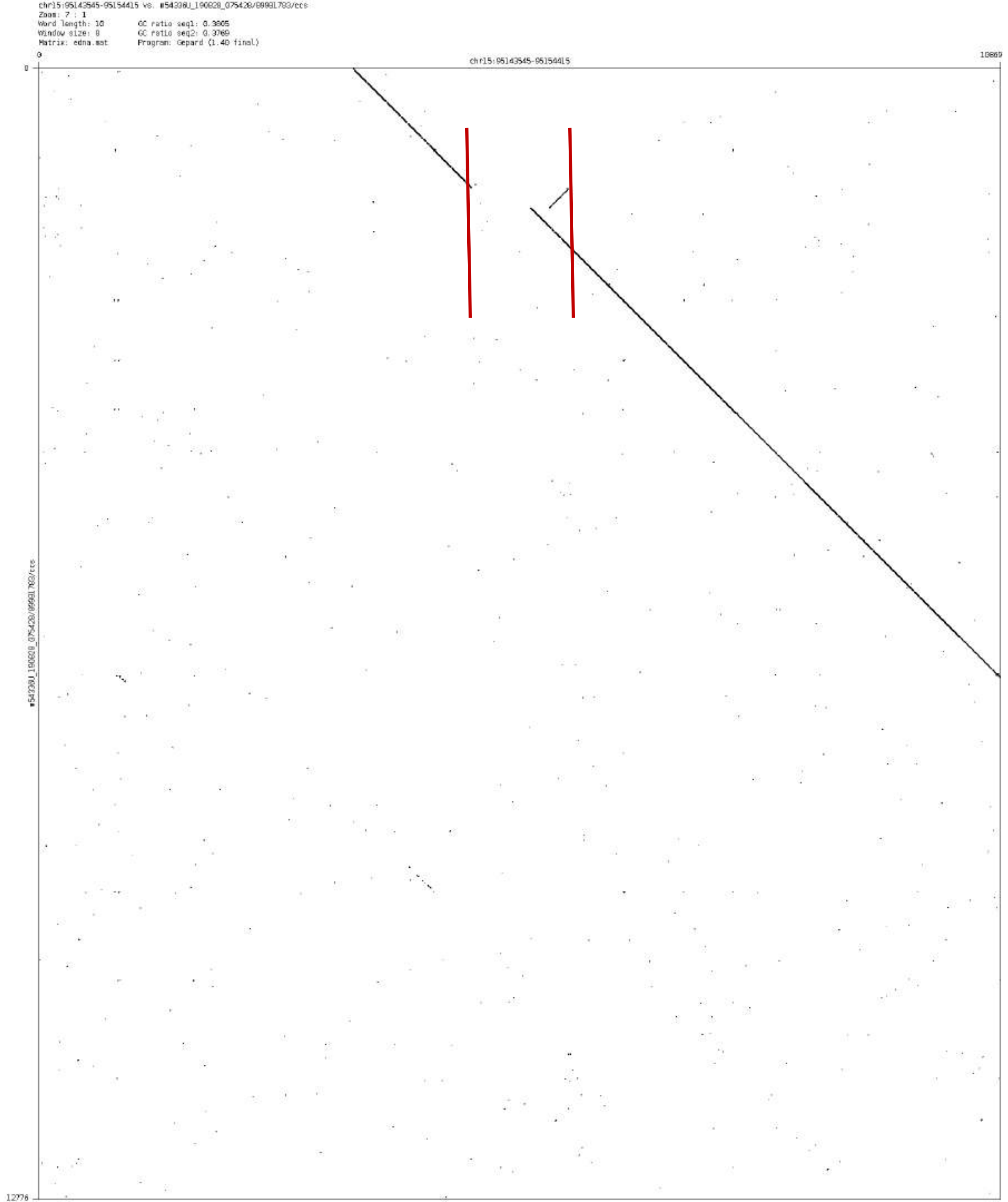
chr15:95143545-95154415 vs. m54338U_190828_075420/89981783/ccs
Zoom: 7 : 1
Word length: 10
Window size: 0
Matrix: edna.mat
GC ratio seq1: 0.3805
GC ratio seq2: 0.3769
Program: Gepard (1.40 final)
0
10869
chr15:95143545-95154415
0
12776

chr16:10676974-10677779

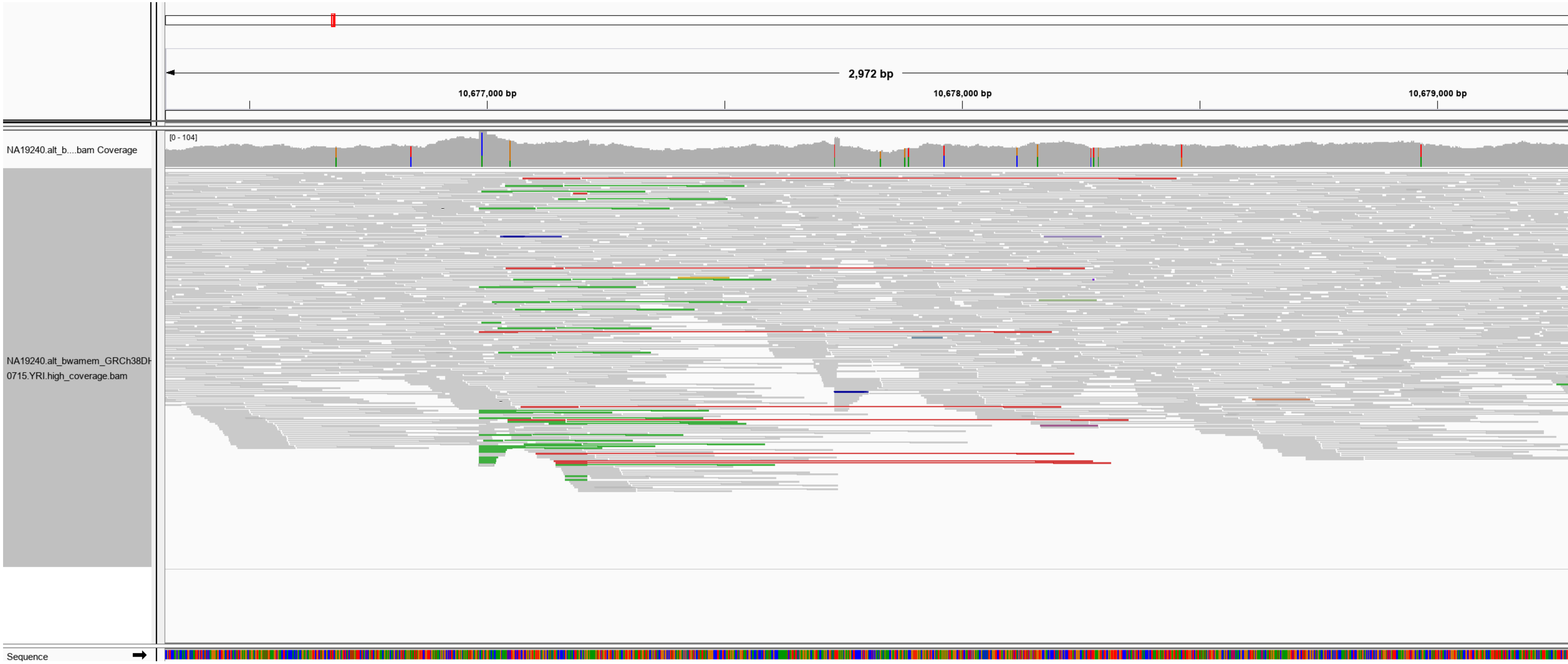

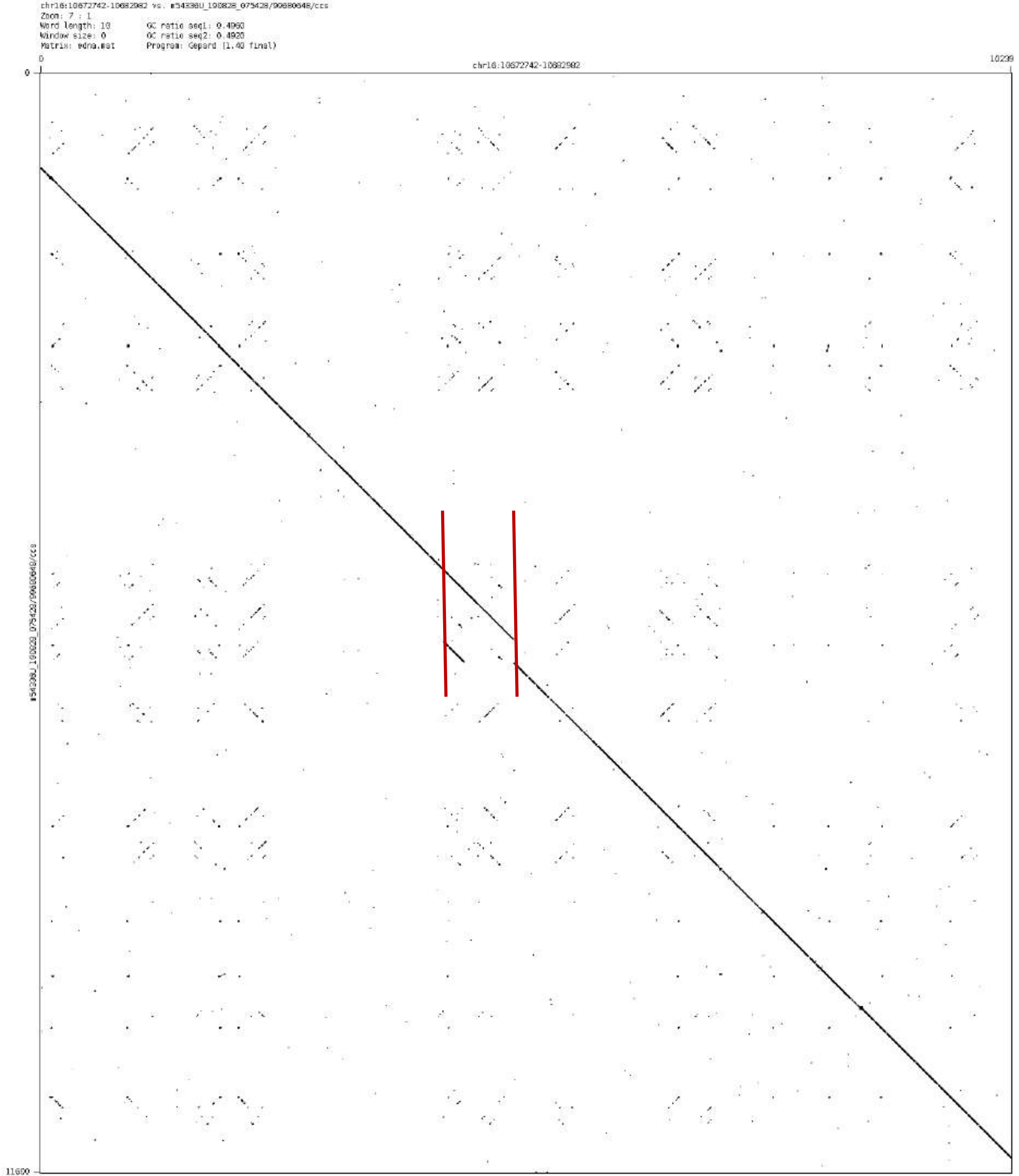
chr16:10672742-10682982 vs. m54330U_190828_075428/99680648/ccs
Zoom: 7 : 1
Word length: 10
Window size: 0
Matrix: edna.mat
GC ratio seq1: 0.4960
GC ratio seq2: 0.4920
Program: Gepard (1.40 final)
0
10239
chr16:10672742-10682982
0
11600
m54330U_190828_075428/99680648/ccs

chr16:69727929-69728986

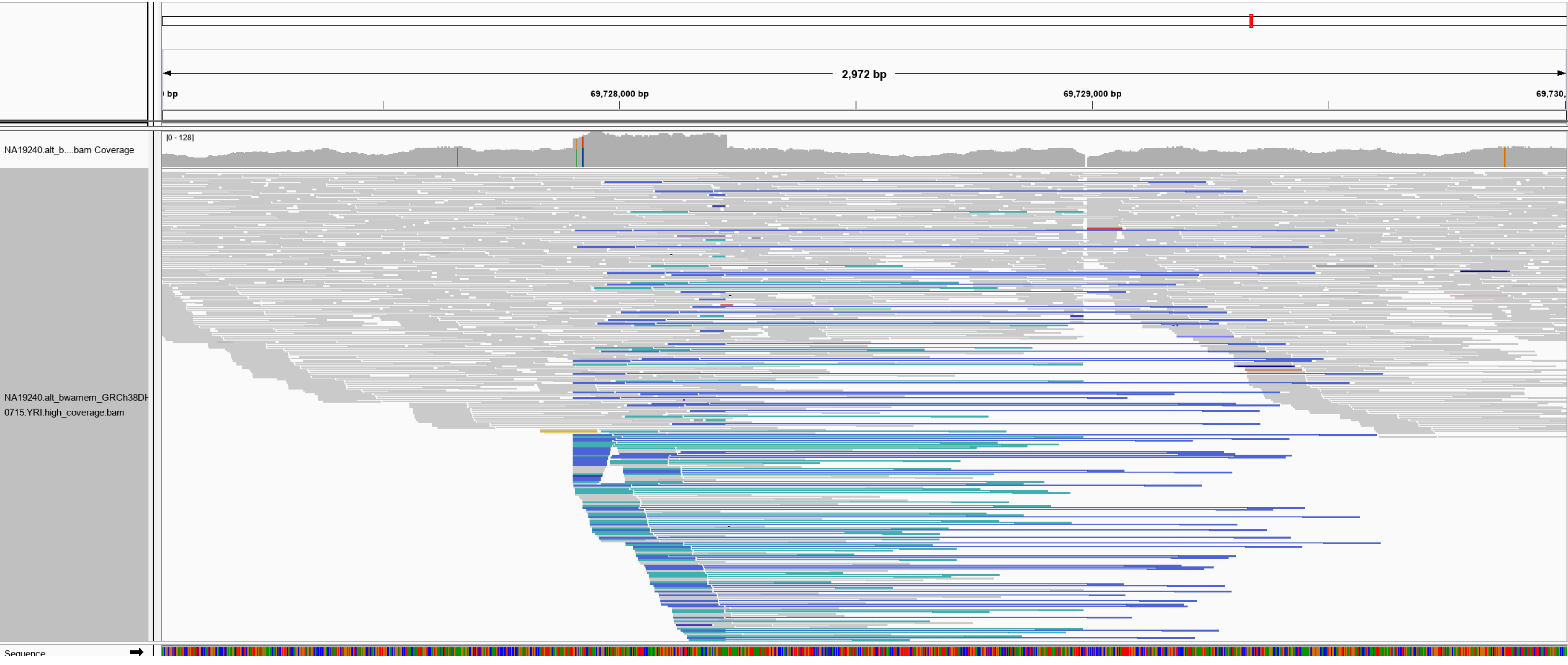

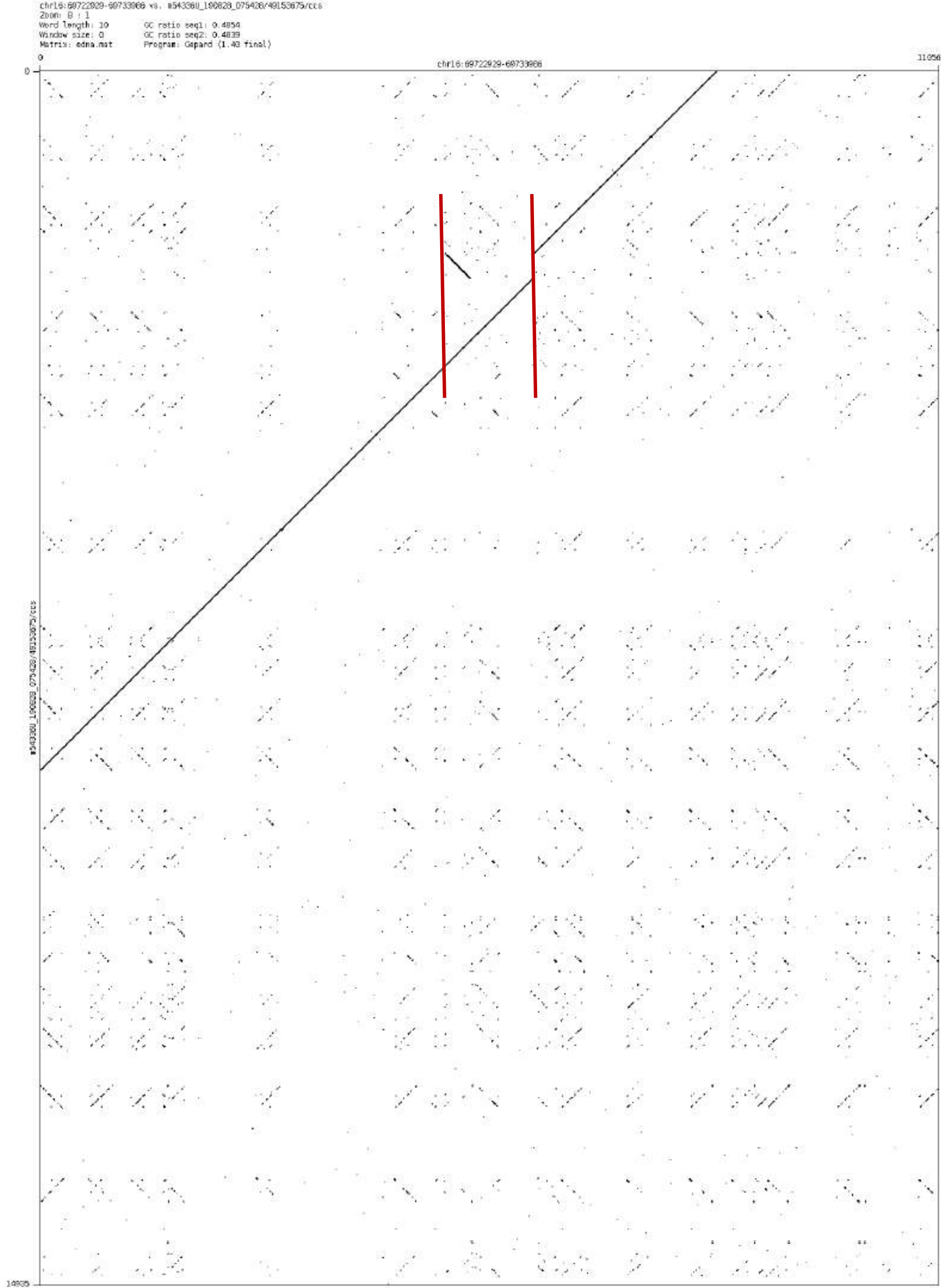

chr16:60722929-60733986 vs. m54336U_190828_075426/49153675/ccs
Zoom: 8 : 1
Word length: 10
Window size: 0
Matrix: edna.mat
GC ratio seq1: 0.4854
GC ratio seq2: 0.4839
Program: Gepard (1.40 final)
0
chr16:60722929-60733986
11056
0
14935

chr17:10983538-10992414

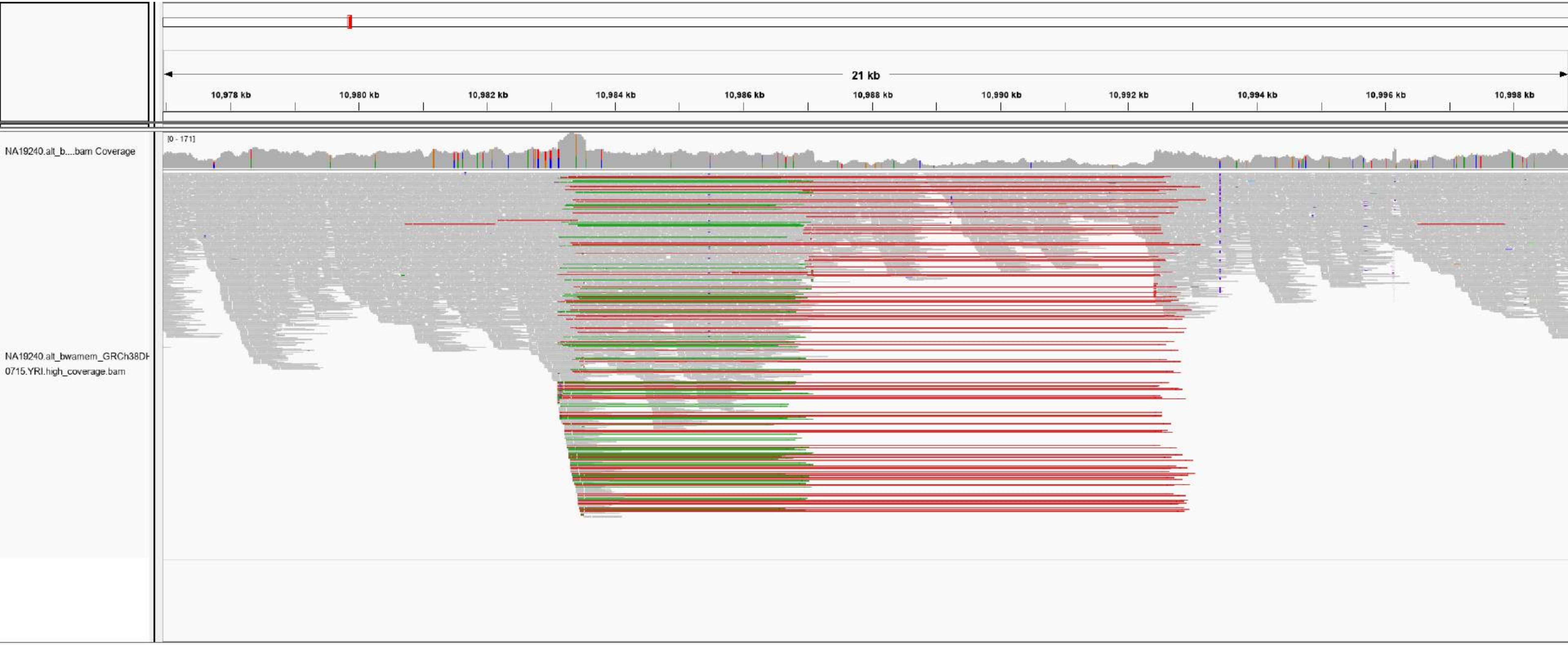

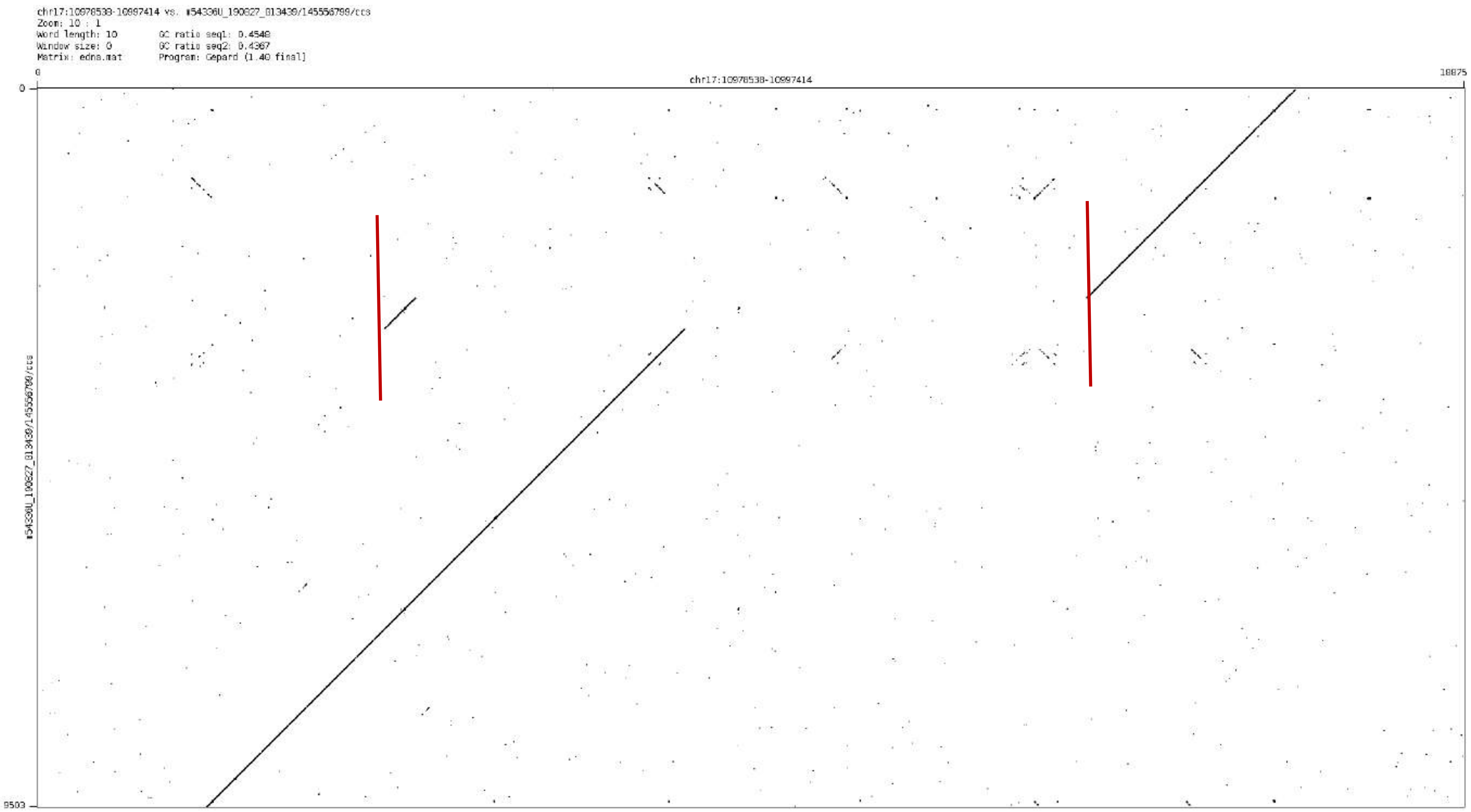

chr17:10978538-10997414 vs. #54336U_190827_013439/145556799/ccs
Zoom: 10 : 1
Word length: 10
GC ratio seq1: 0.4548
Window size: 0
GC ratio seq2: 0.4367
Matrix: edna.mat
Program: Gepard (1.40 final)
0
18875
chr17:10978538-10997414
0
9503
#54336U_190827_013439/145556799/ccs

### chr17:43358999-43365252

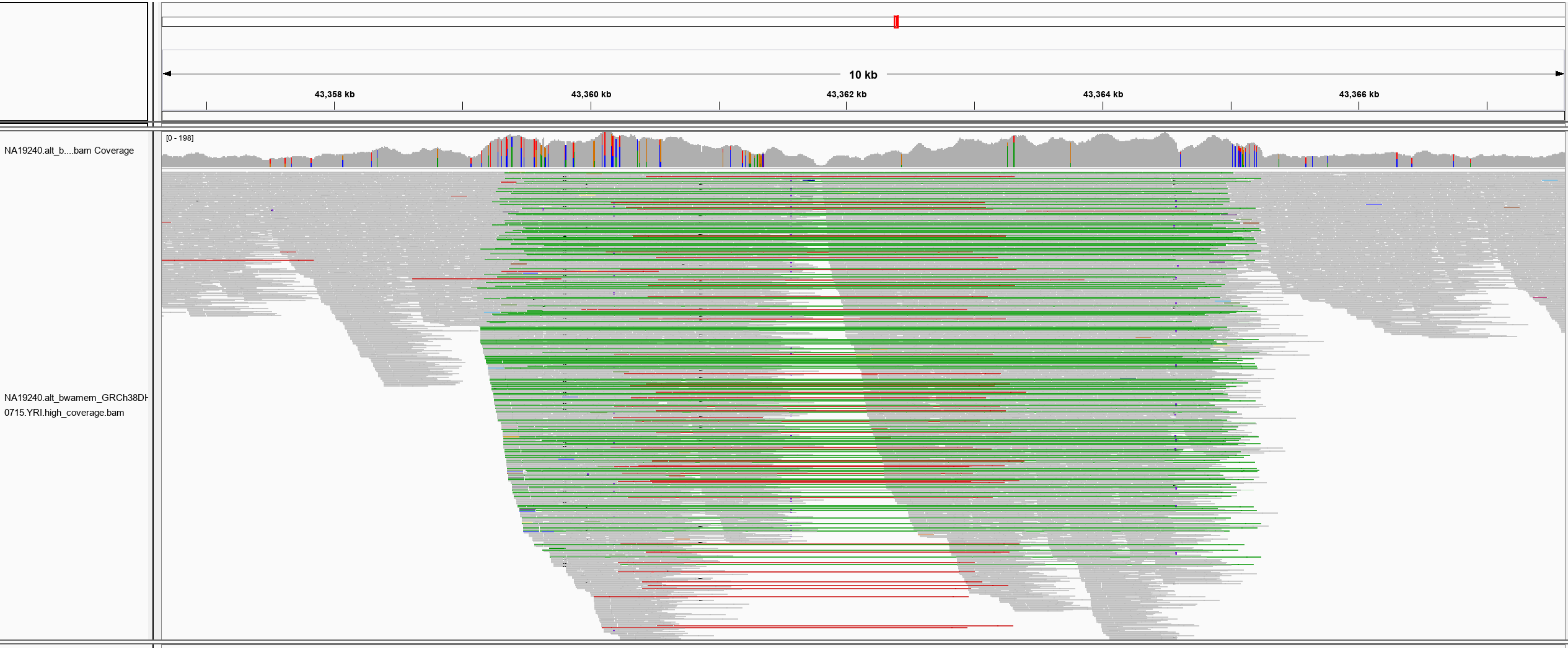

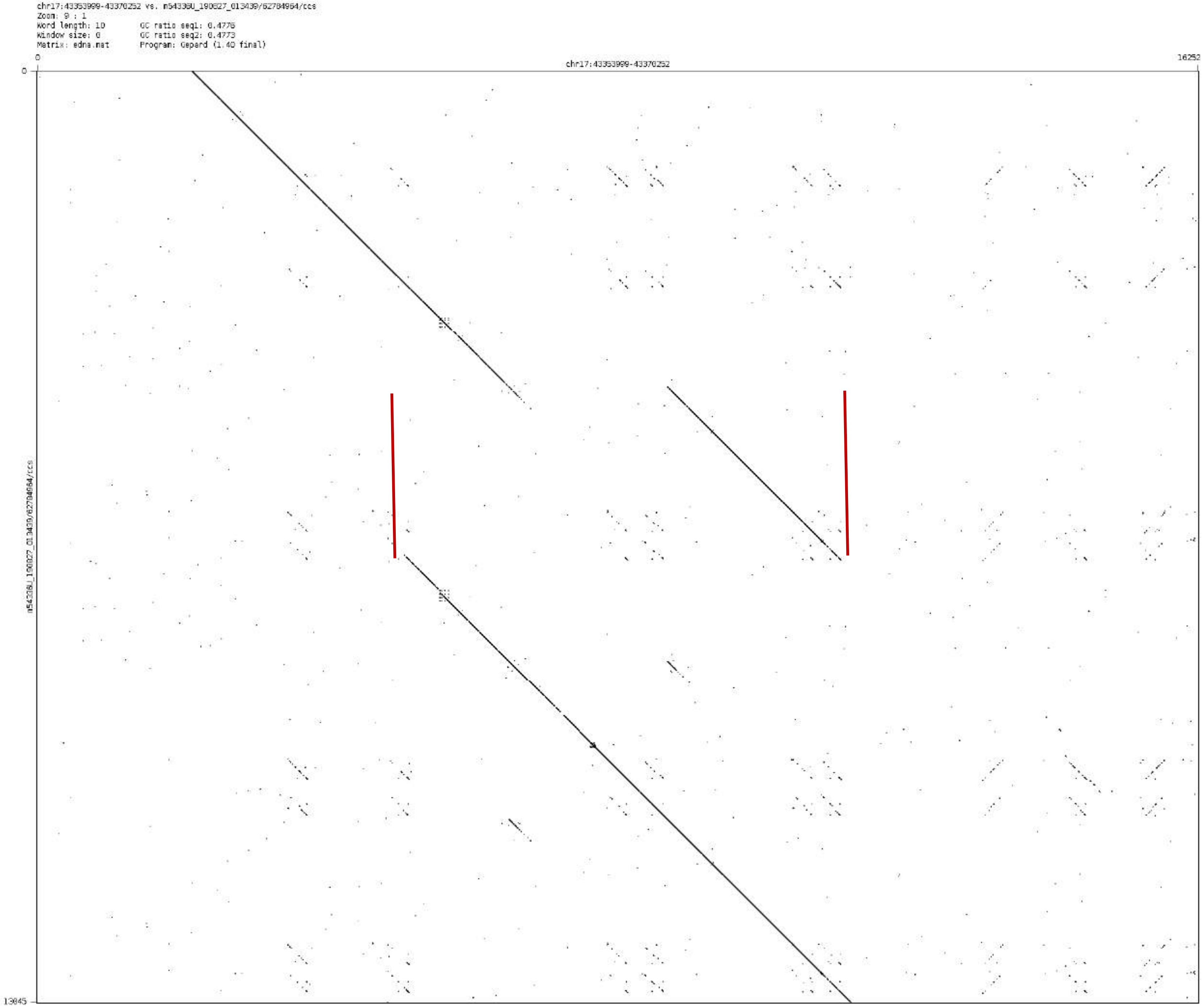
chr17:43353999-43370252 vs. m54336U_190827_013439/62784964/ccs
Zoom: 9 : 1
Word length: 10
Window size: 0
Matrix: edna.mat
GC ratio seq1: 0.4776
GC ratio seq2: 0.4773
Program: Gepard (1.40 final)
0
16252
chr17:43353999-43370252
0
13845
m54336U_190827_013439/62784964/ccs

### chr18:11509284-11513469

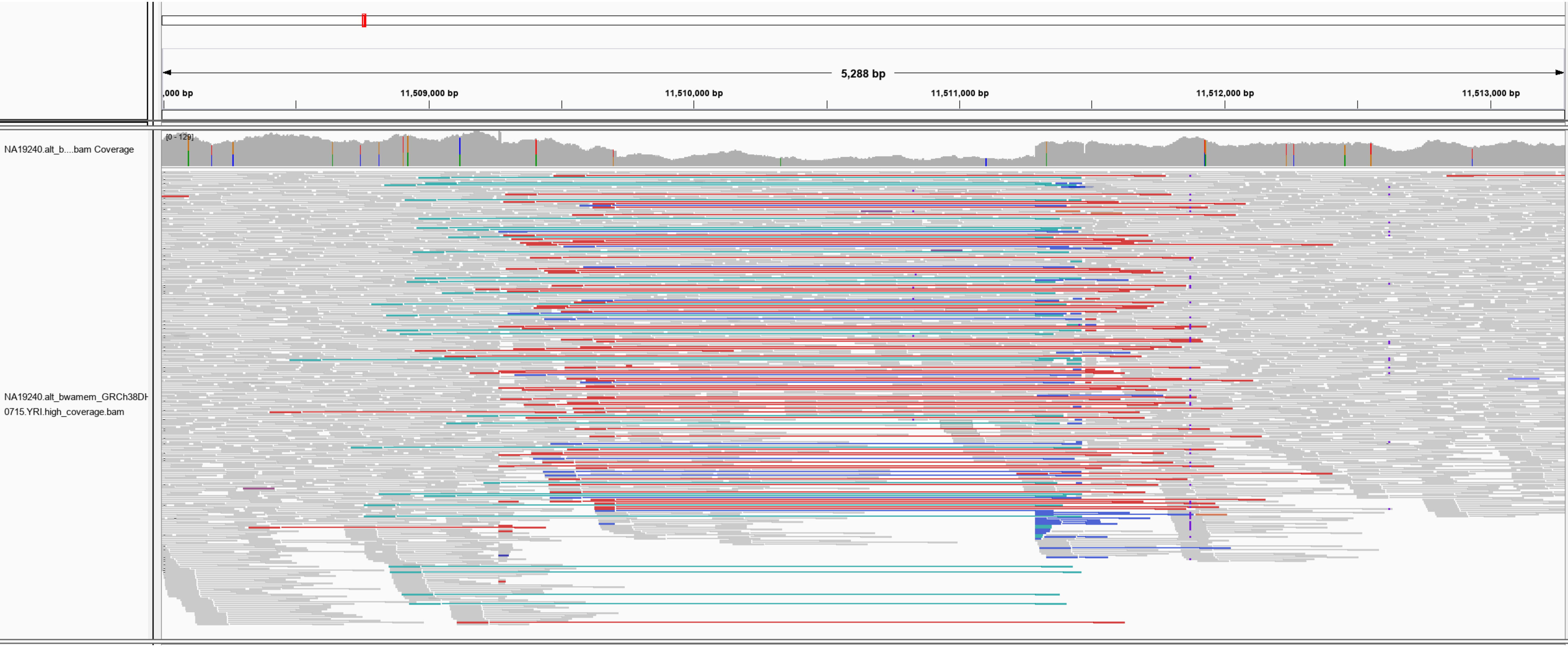

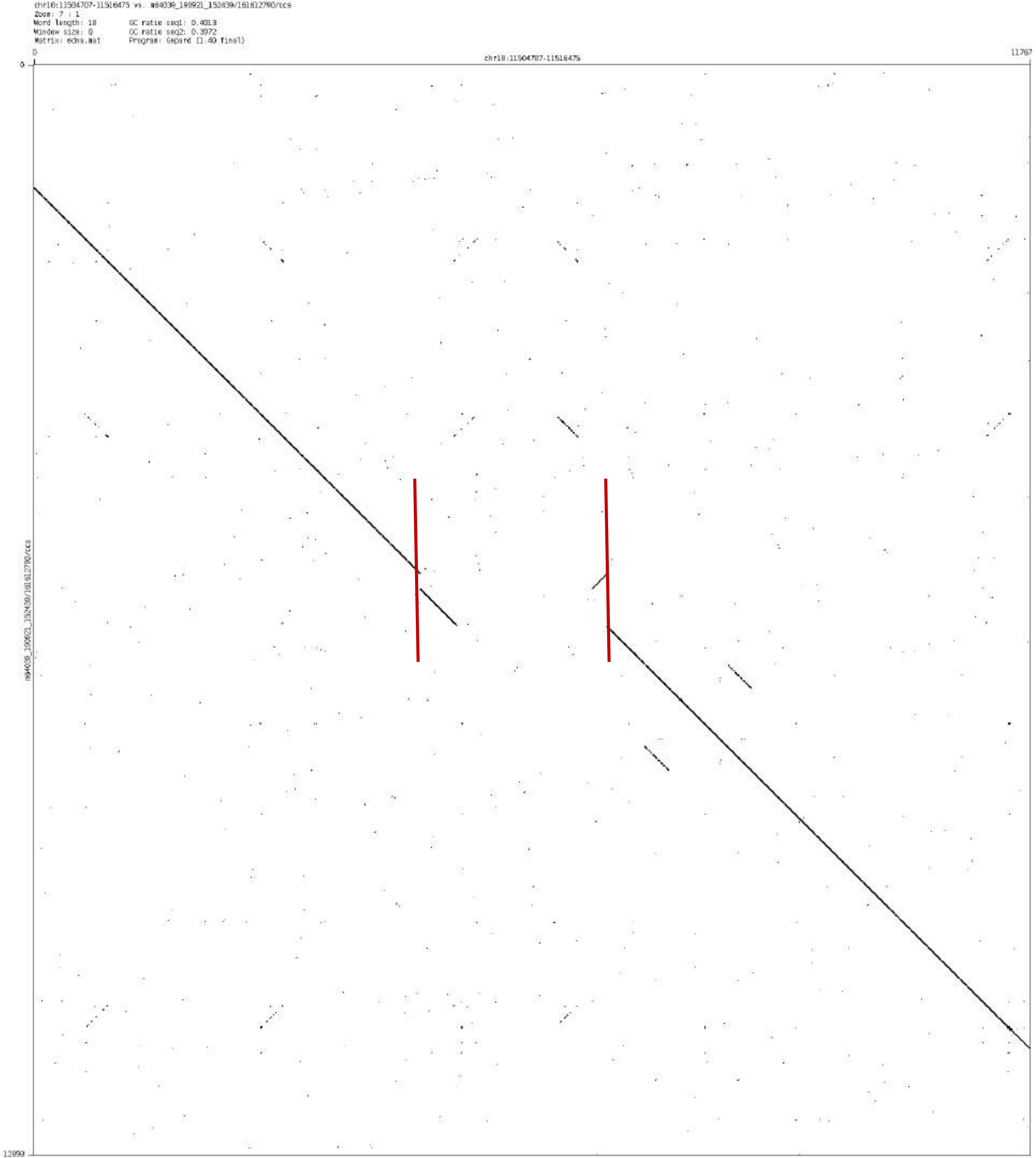
chr18:11504707-11516475 vs. m64039_190921_152439/161612790/ccs
Zoom: 7 : 1
Word length: 18
Window size: 0
Matrix: edna.mat
GC ratio seq1: 0.4013
GC ratio seq2: 0.3972
Program: Gepard (1.40 final)
0
11767
chr18:11504707-11516475
0
12899
m64039_190921_152439/161612790/ccs

chr19:56640093-56640958

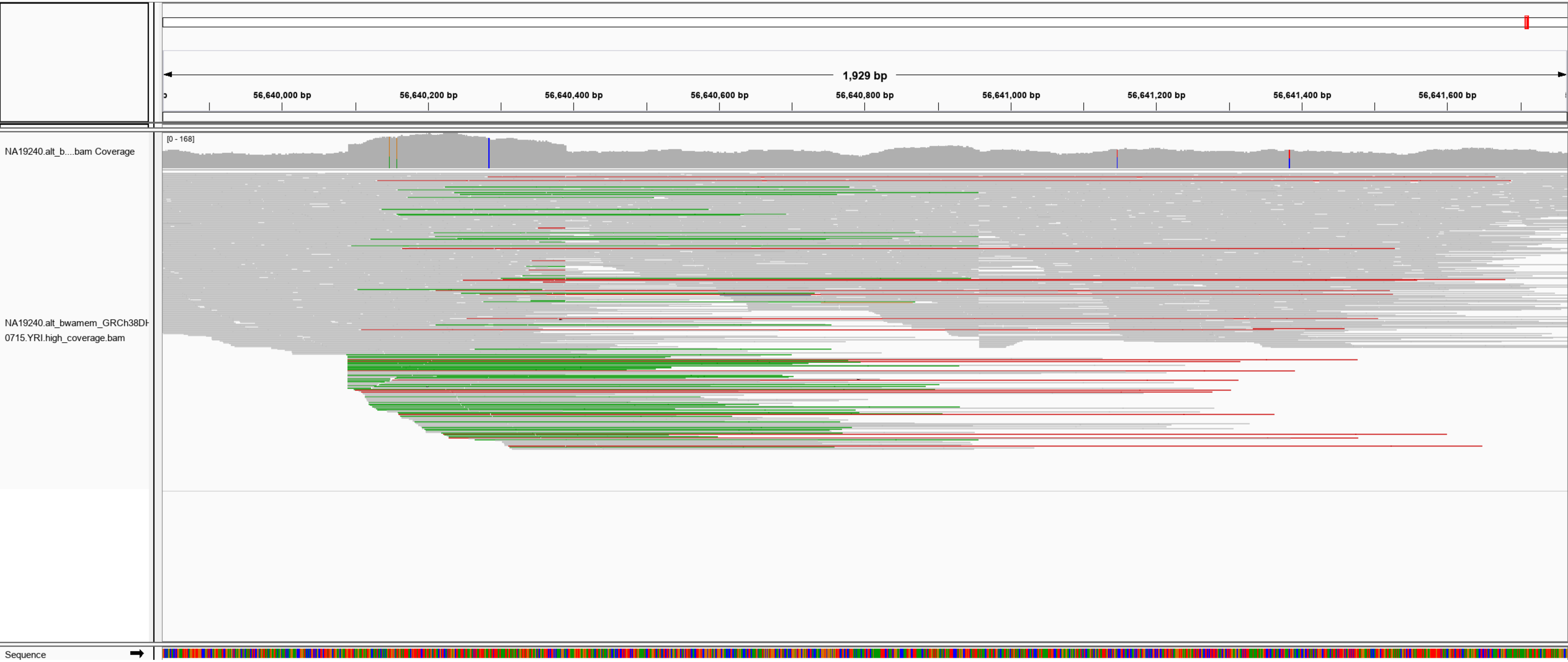

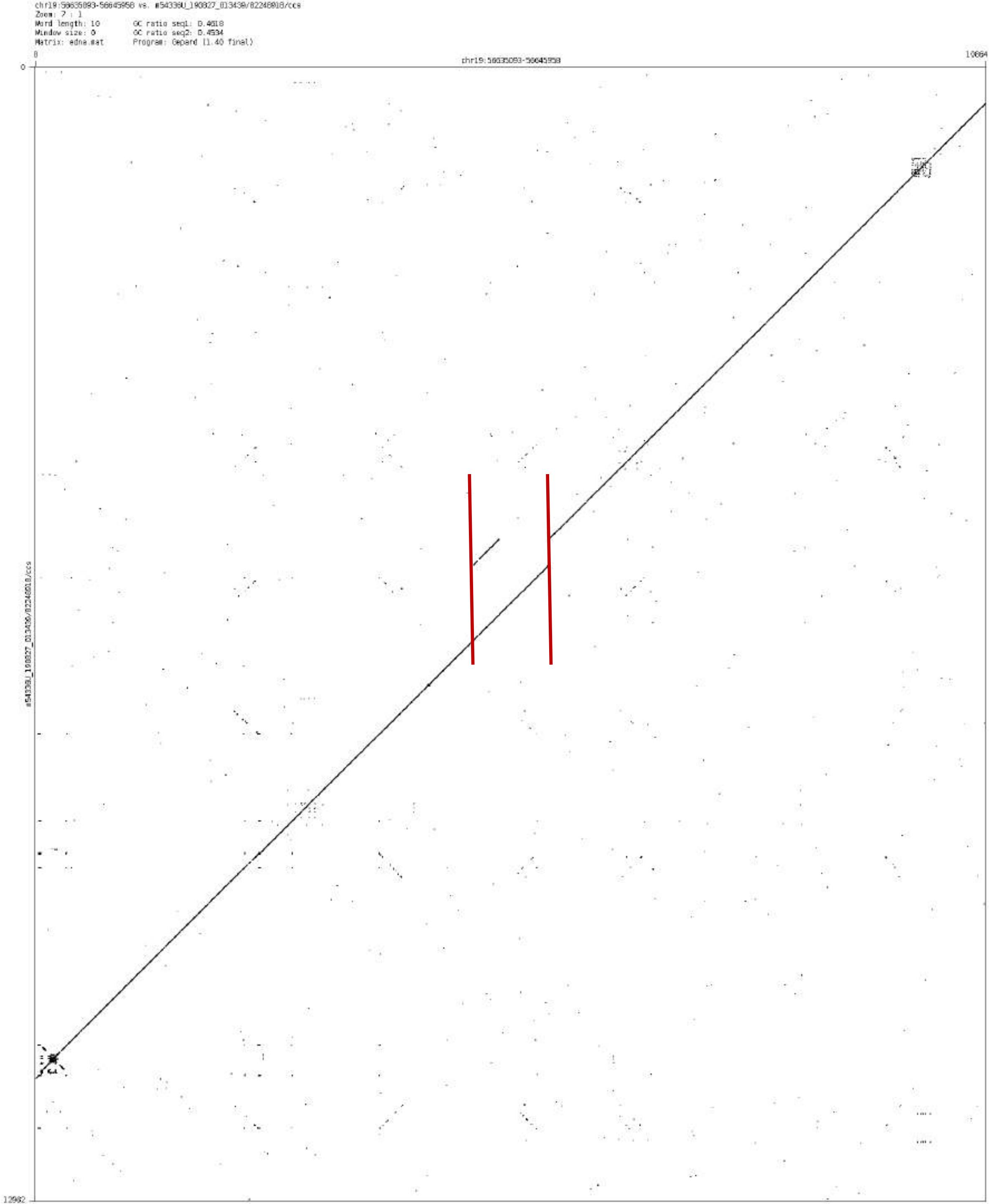

chr19:56635093-56645958 vs. m54338U_190827_013430/82248018/ccs
Zoom: 7 : 1
Word length: 10
Window size: 0
Matrix: edna.mat
GC ratio seq1: 0.4618
GC ratio seq2: 0.4534
Program: Gepard (1.40 final)
chr19:56635093-56645958

chr20:2379153-2379968

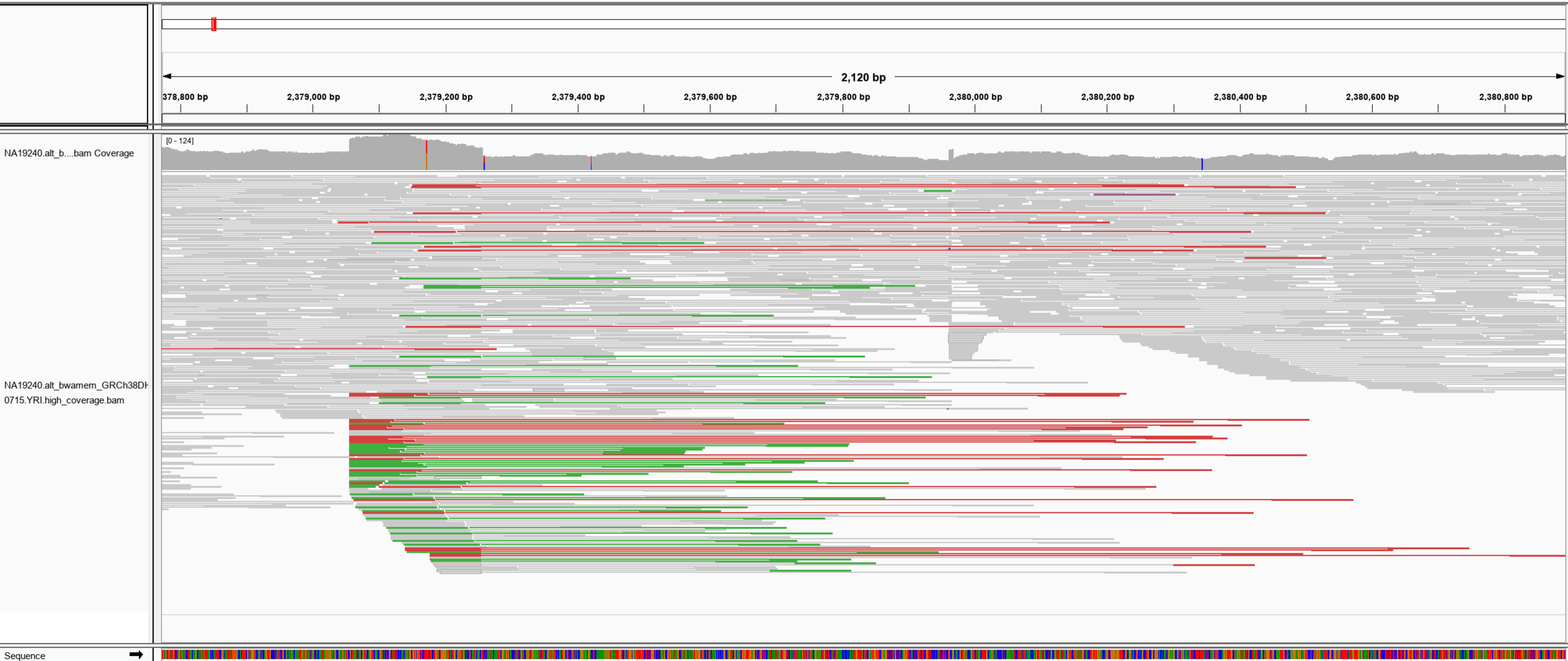

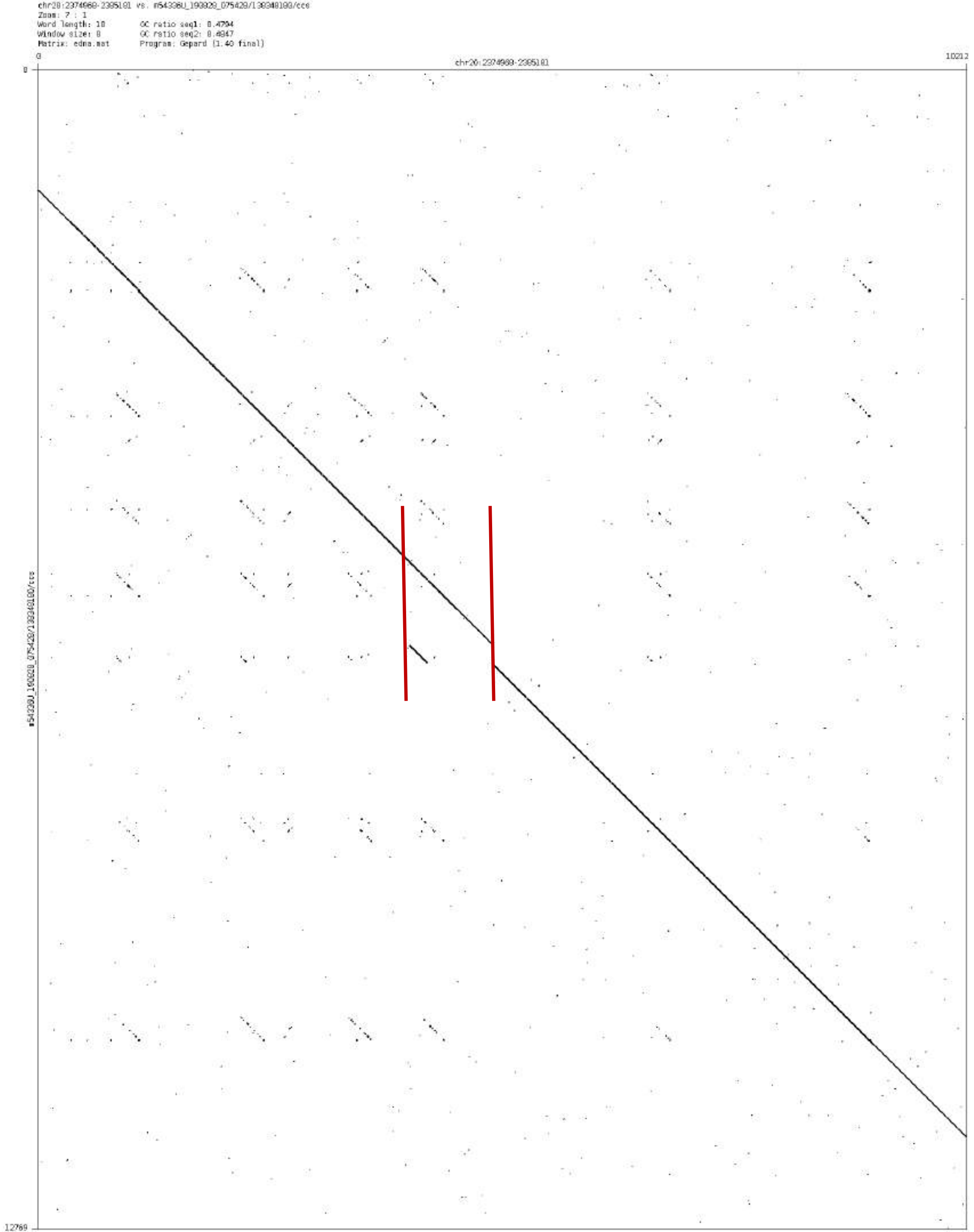

chr20:2374960-2385181 vs. m54336U_190920_075420/130340180/ccs
Zoom: 7 : 1
Word length: 10
Window size: 0
Matrix: edna.mat
GC ratio seq1: 0.4794
GC ratio seq2: 0.4847
Program: Gepard (1.40 final)
0
chr20:2374960-2385181
10212
0
12769

chr20:23122542-23122996

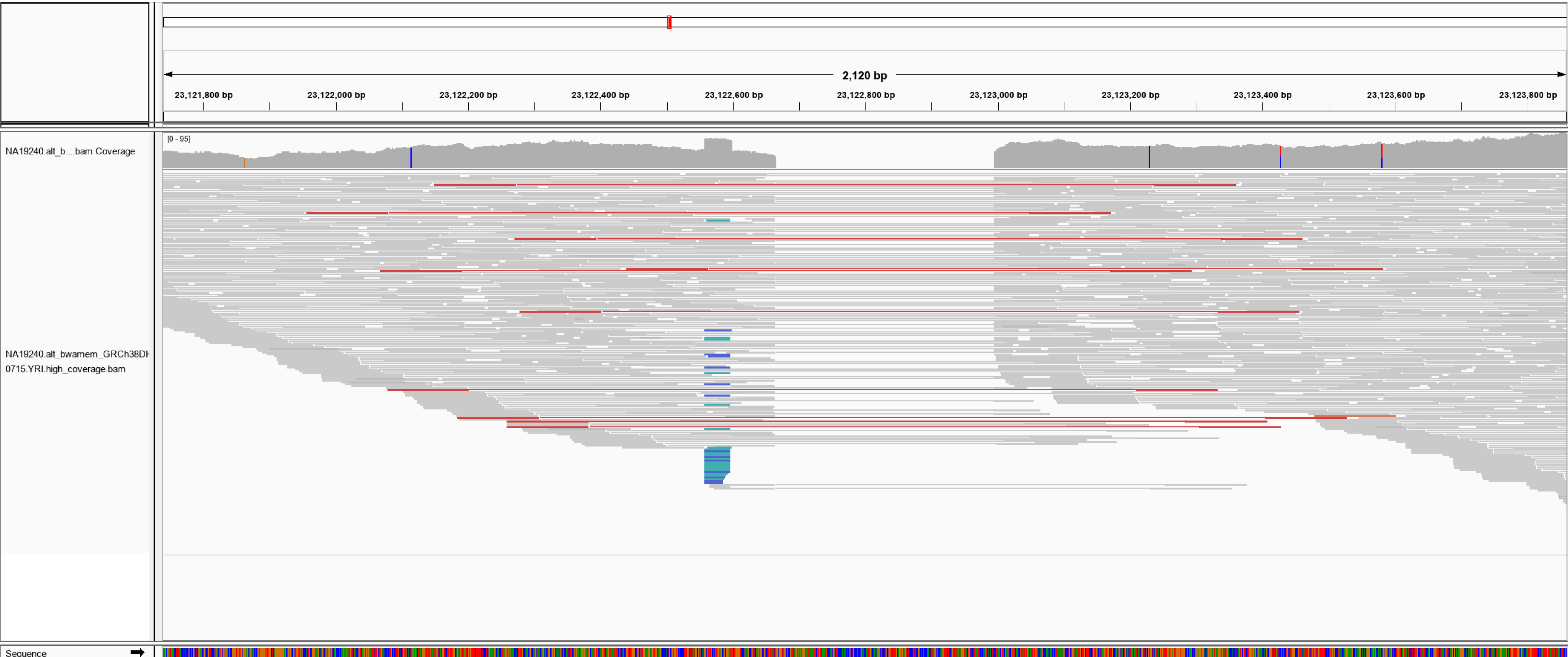

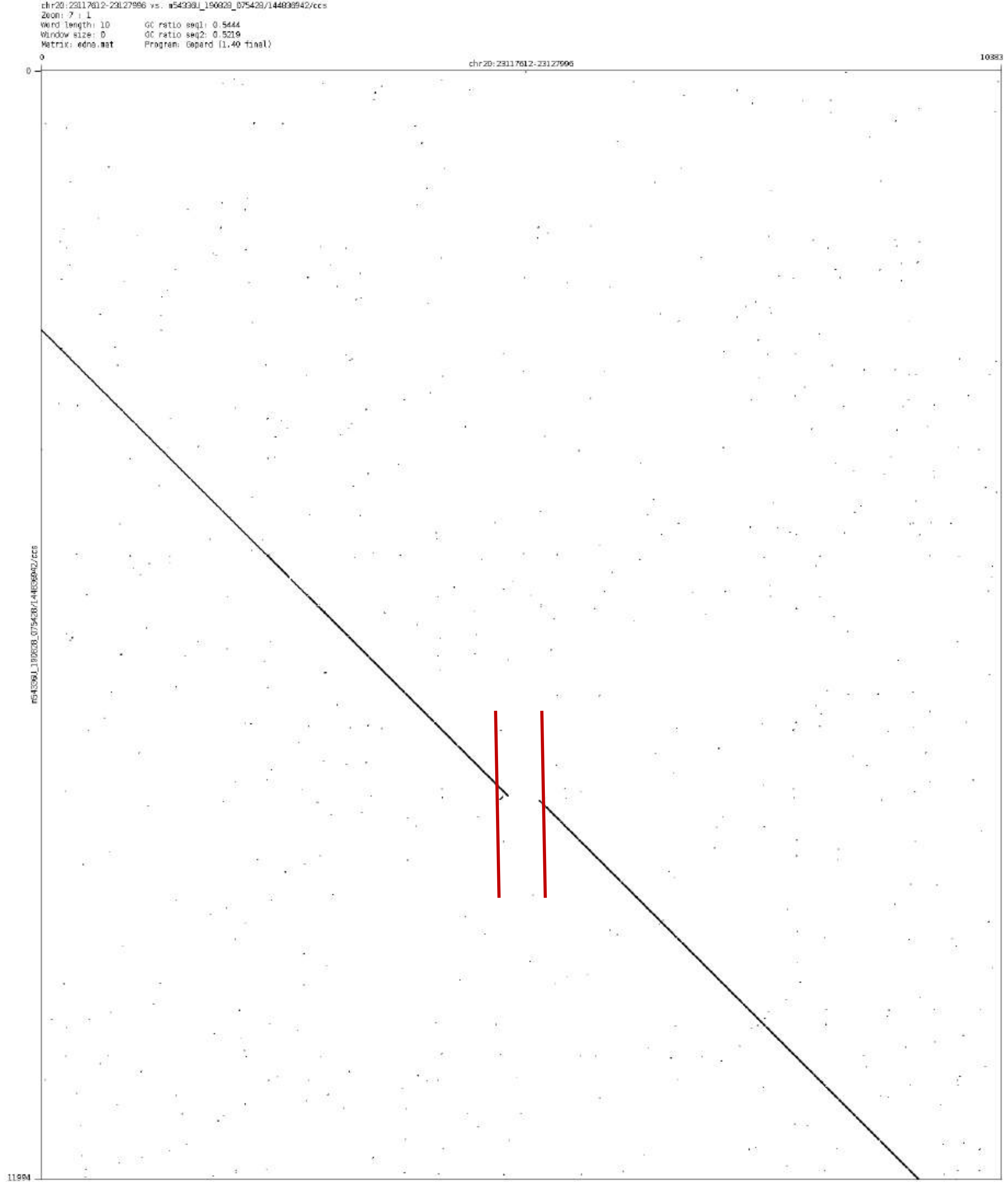
chr20:23117612-23127996 vs. m54338U_190828_075428/144838942/ccs
Zoom: 7 : 1
Word length: 10
Window size: 0
Matrix: edna.mat
GC ratio seq1: 0.5444
GC ratio seq2: 0.5219
Program: Gepard (1.40 final)
0
chr20:23117612-23127996
10383
0
11994

chr22:49368449-49373094

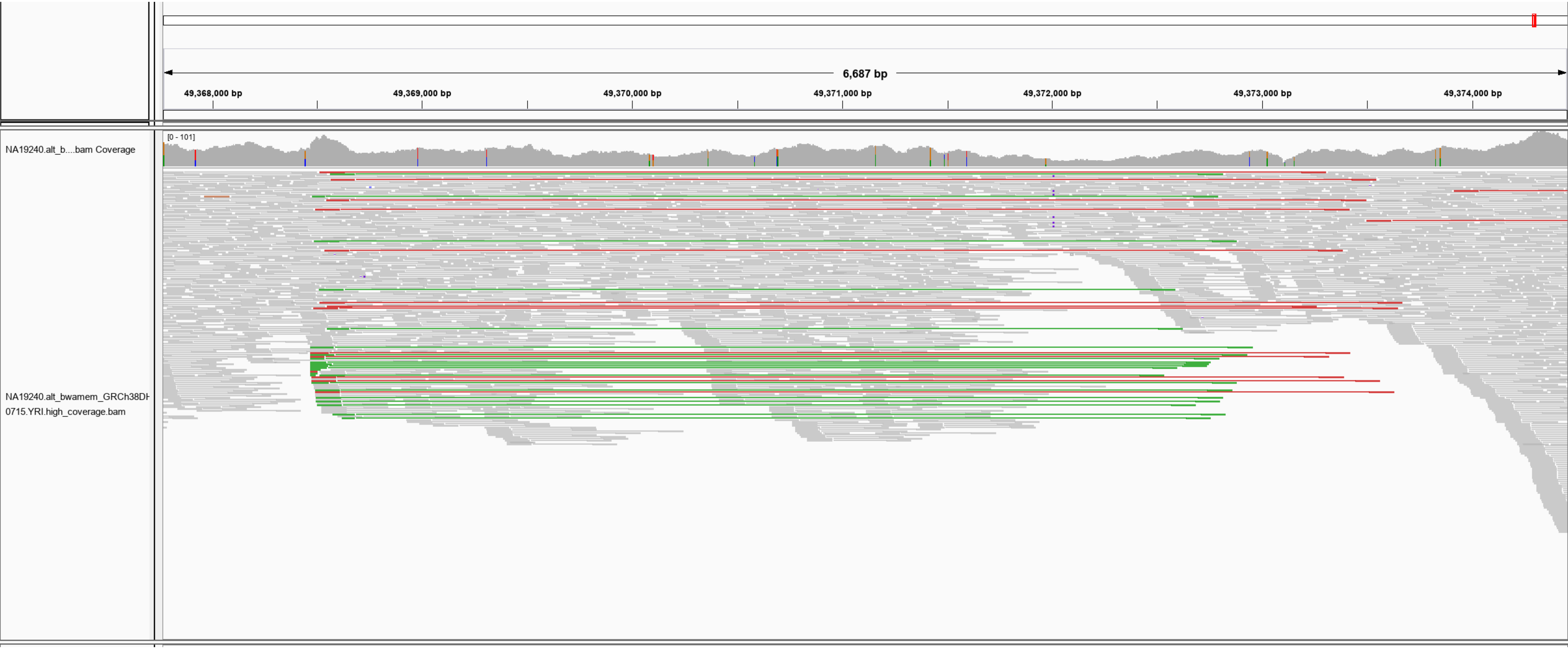

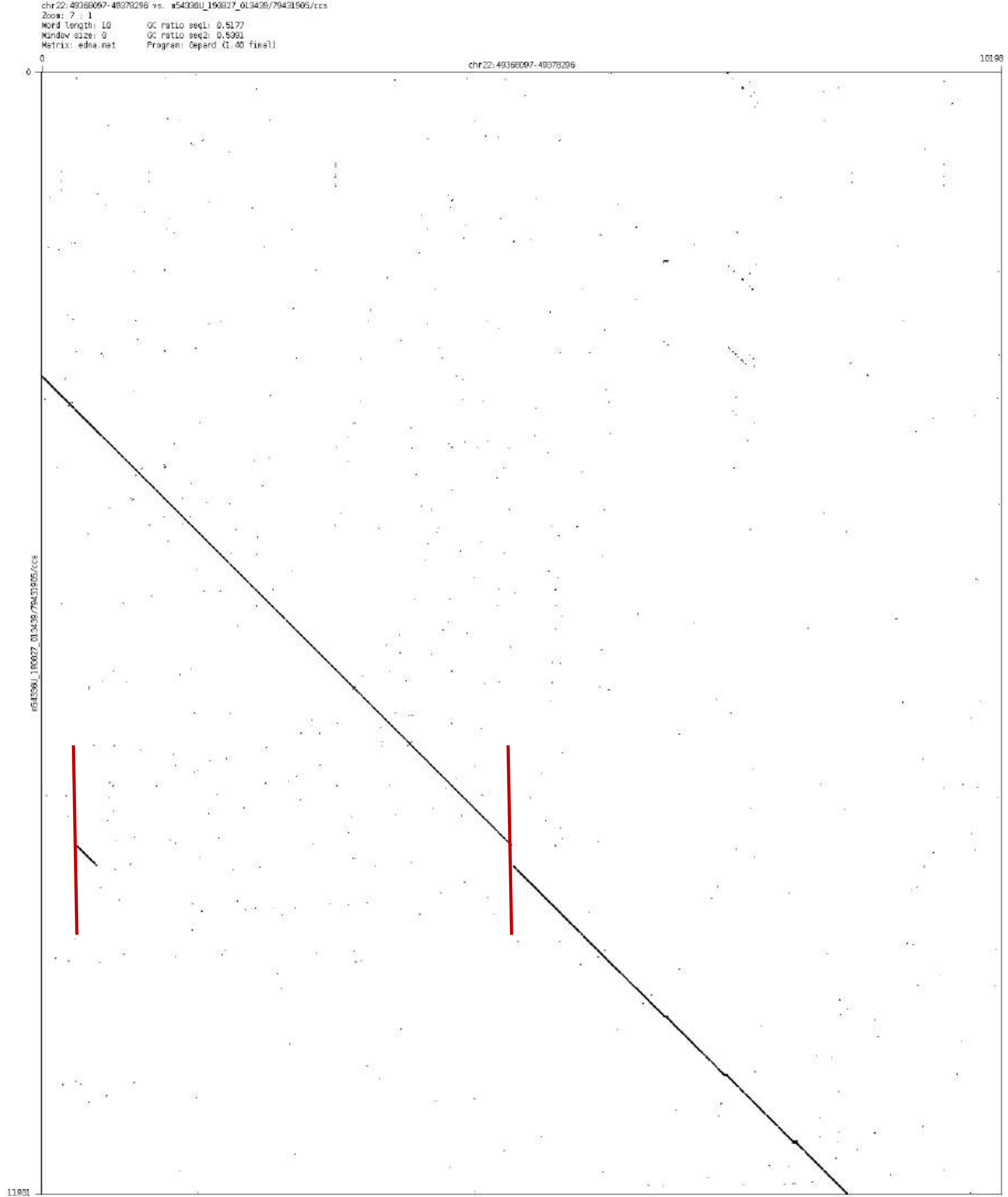

chr22:49368097-49378296 vs. m54336U_190827_013439/79431905/ccs
Zoom: 7 : 1
Word length: 10
Window size: 0
Matrix: edna.mat
GC ratio seq1: 0.5177
GC ratio seq2: 0.5391
Program: Gepard (1.40 final)
0
chr22:49368097-49378296
10198
0
m54336U_190827_013439/79431905/ccs
11981

### SK-BR-3 benchmark CSVs

### chr1:61060495-61062387

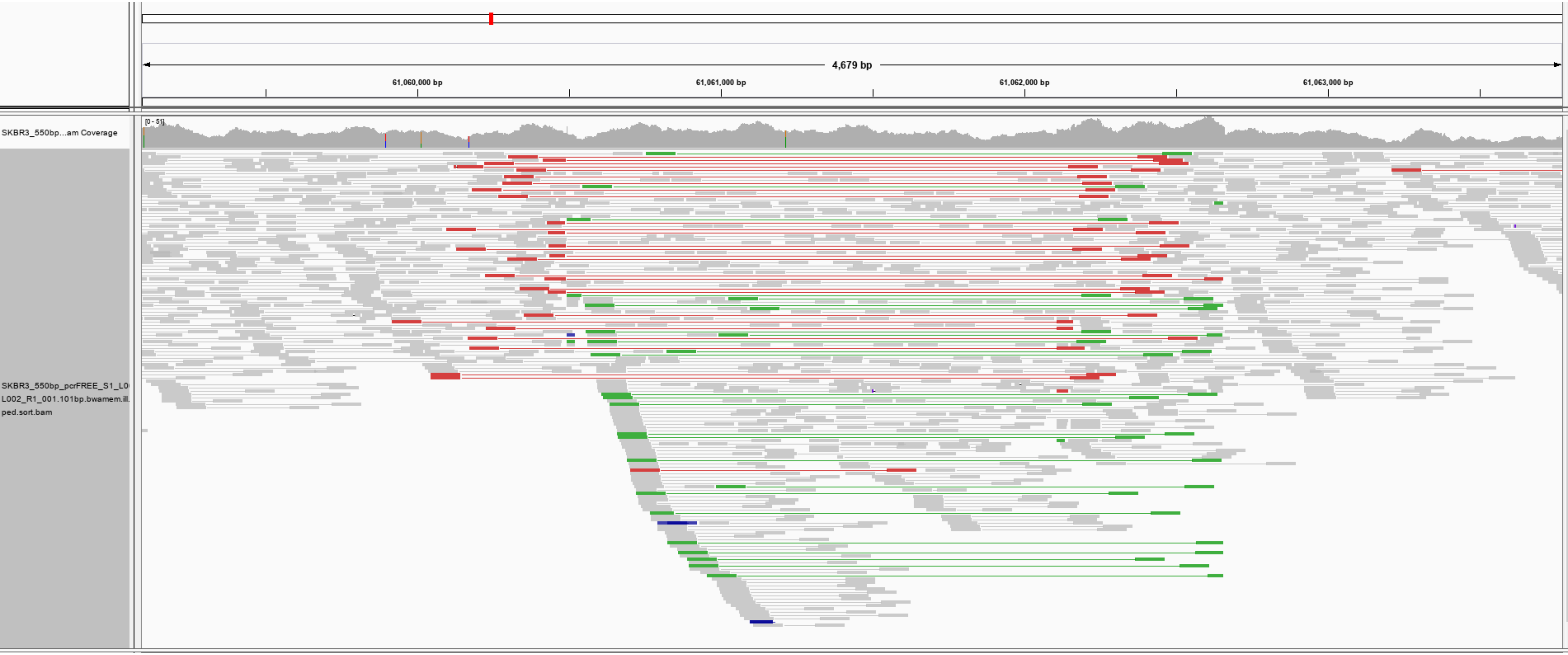

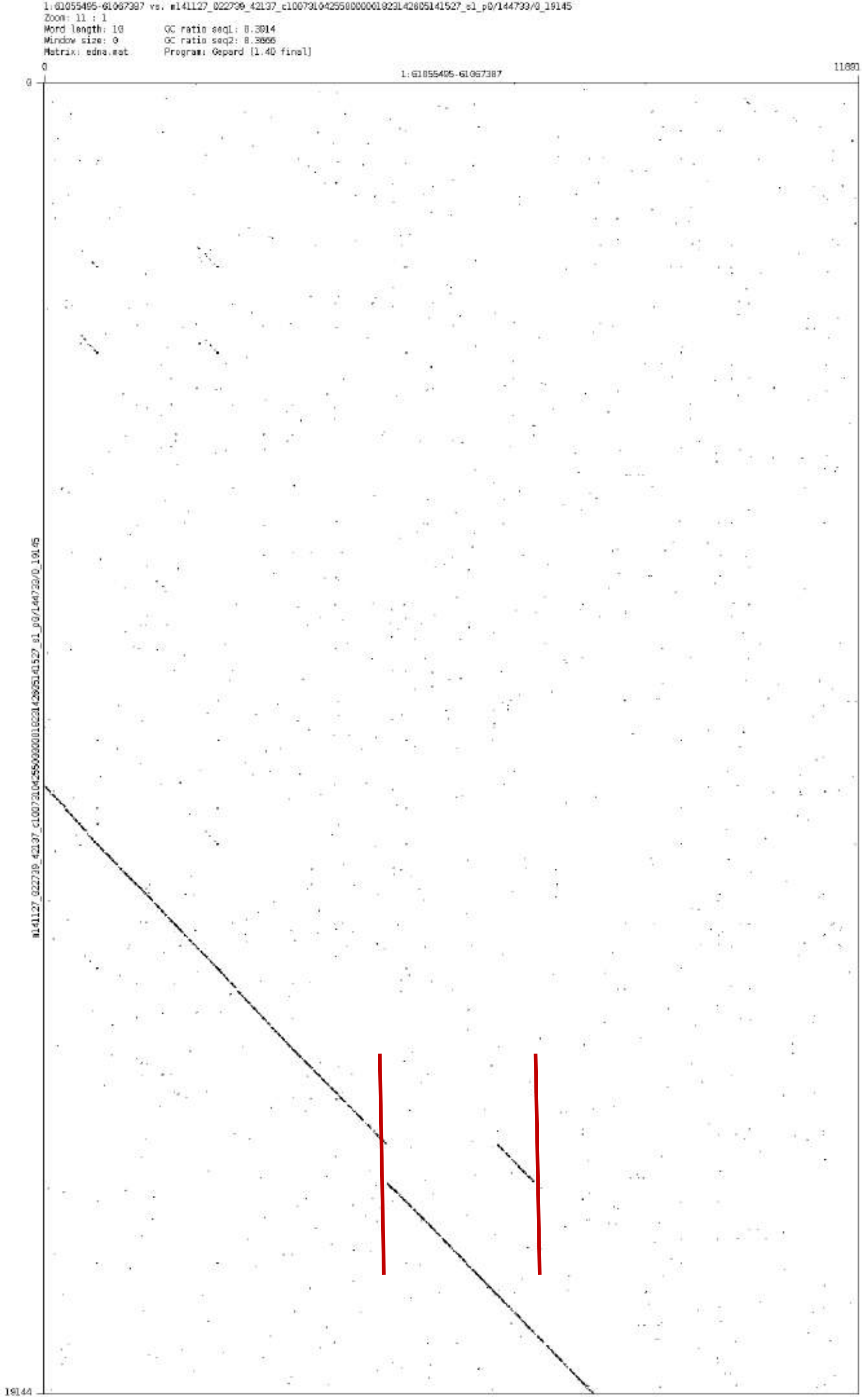
1:61055495-61067387 vs. m141127_022739_42137_c100731042550000001823142805141527_s1_p0/144733/0_19145
Zoom: 11 : 1
Word length: 10
Window size: 0
Matrix: edna.mat
GC ratio seq1: 0.3814
GC ratio seq2: 0.3866
Program: Gepard [1.40 final]
1:61055495-61067387
m141127_022739_42137_c100731042550000001823142805141527_s1_p0/144733/0_19145
0
11881
0
19144

chr1:84056441-84057506

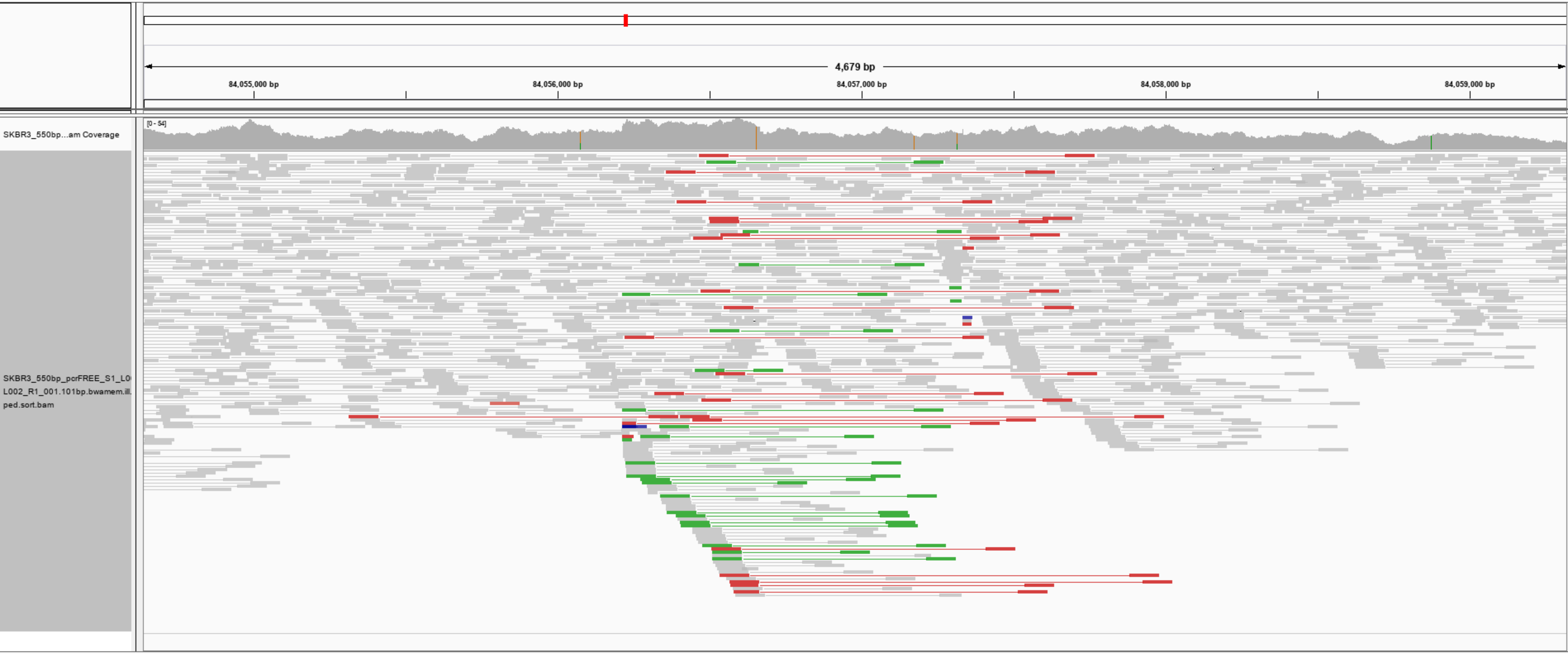

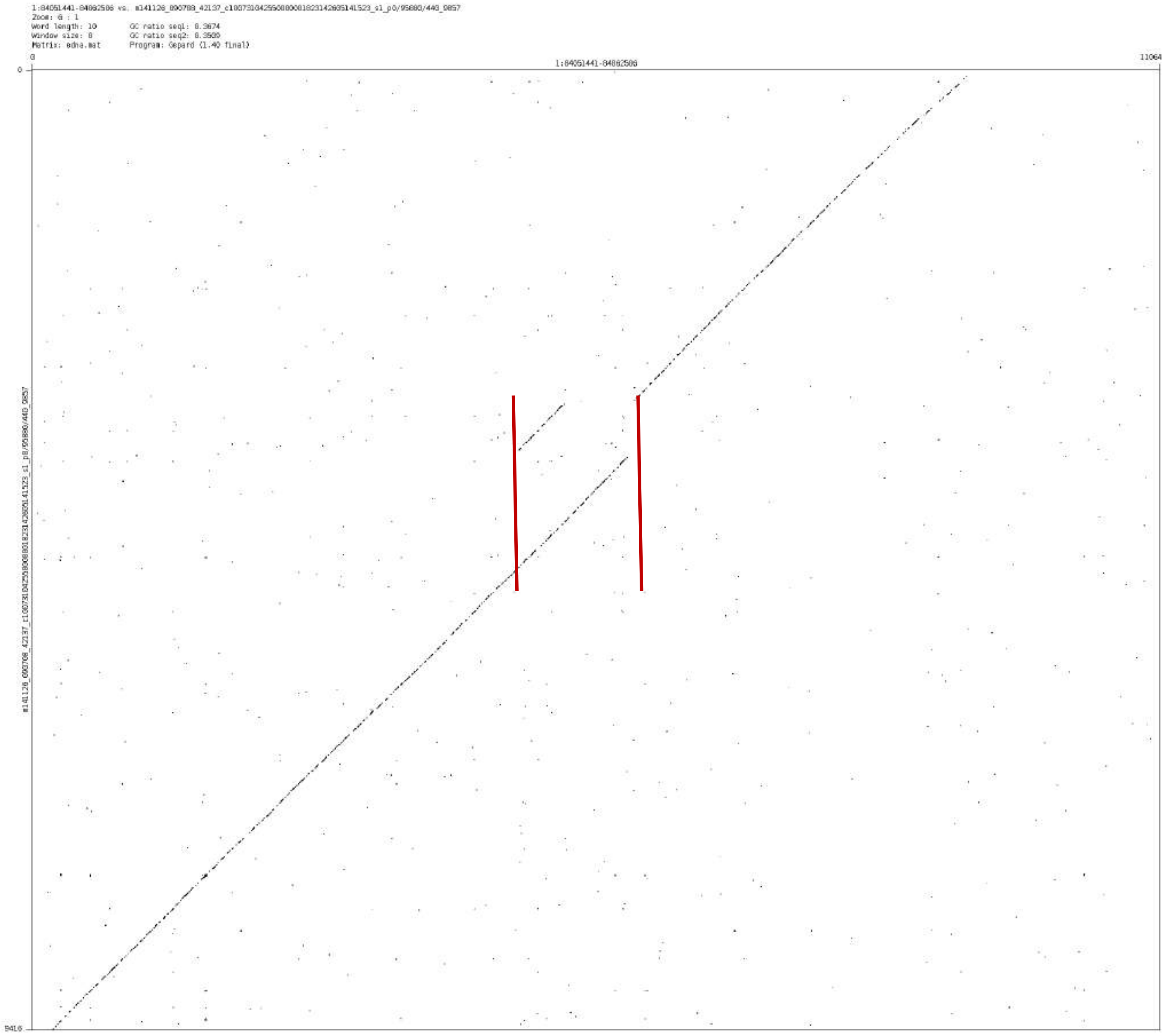

1:84051441-84062506 vs. m141126_090708_42137_c100731042550000001823142035141523_s1_p0/95880/440_9857
Zoom: 6 : 1
Word length: 10
Window size: 8
Matrix: edna.mat
GC ratio seq1: 0.3674
GC ratio seq2: 0.3500
Program: Gepard (1.40 final)
0
1:84051441-84062506
11064
9416

### chr1:144898739-144906810

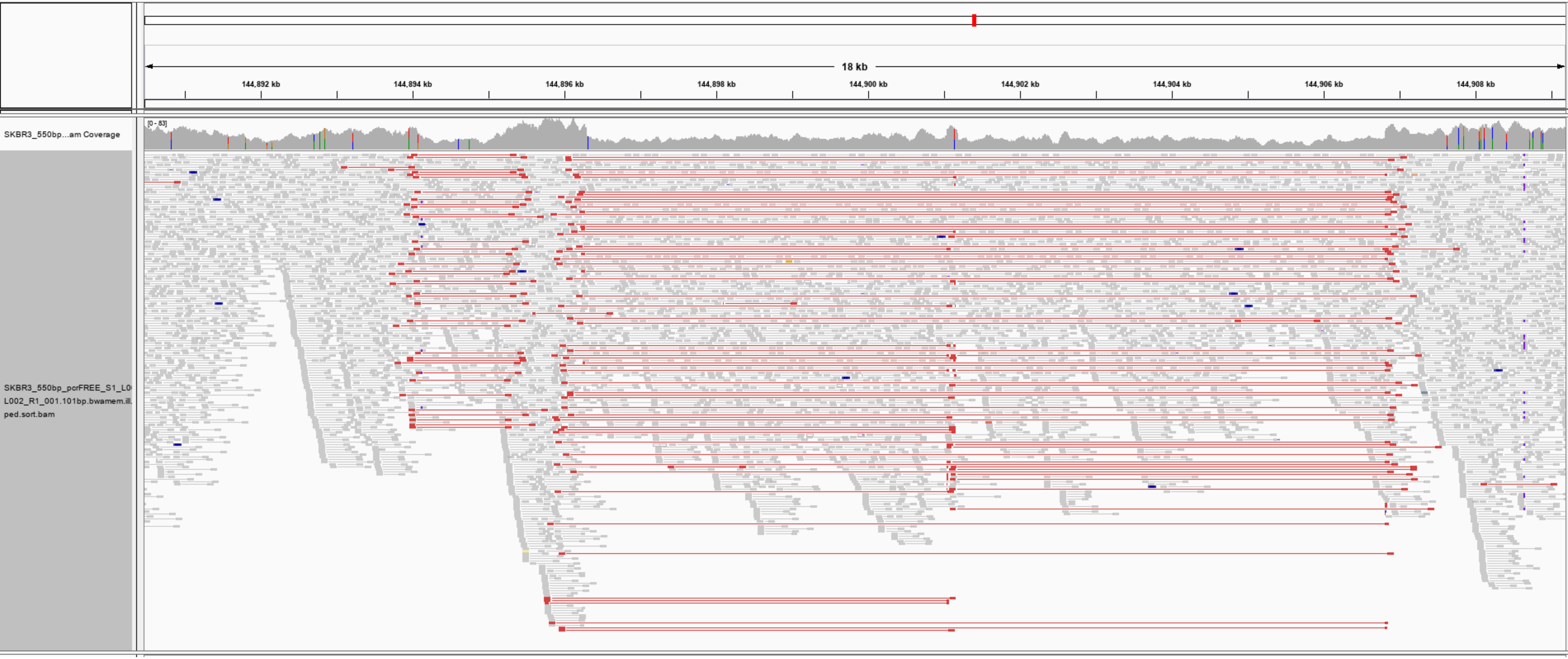

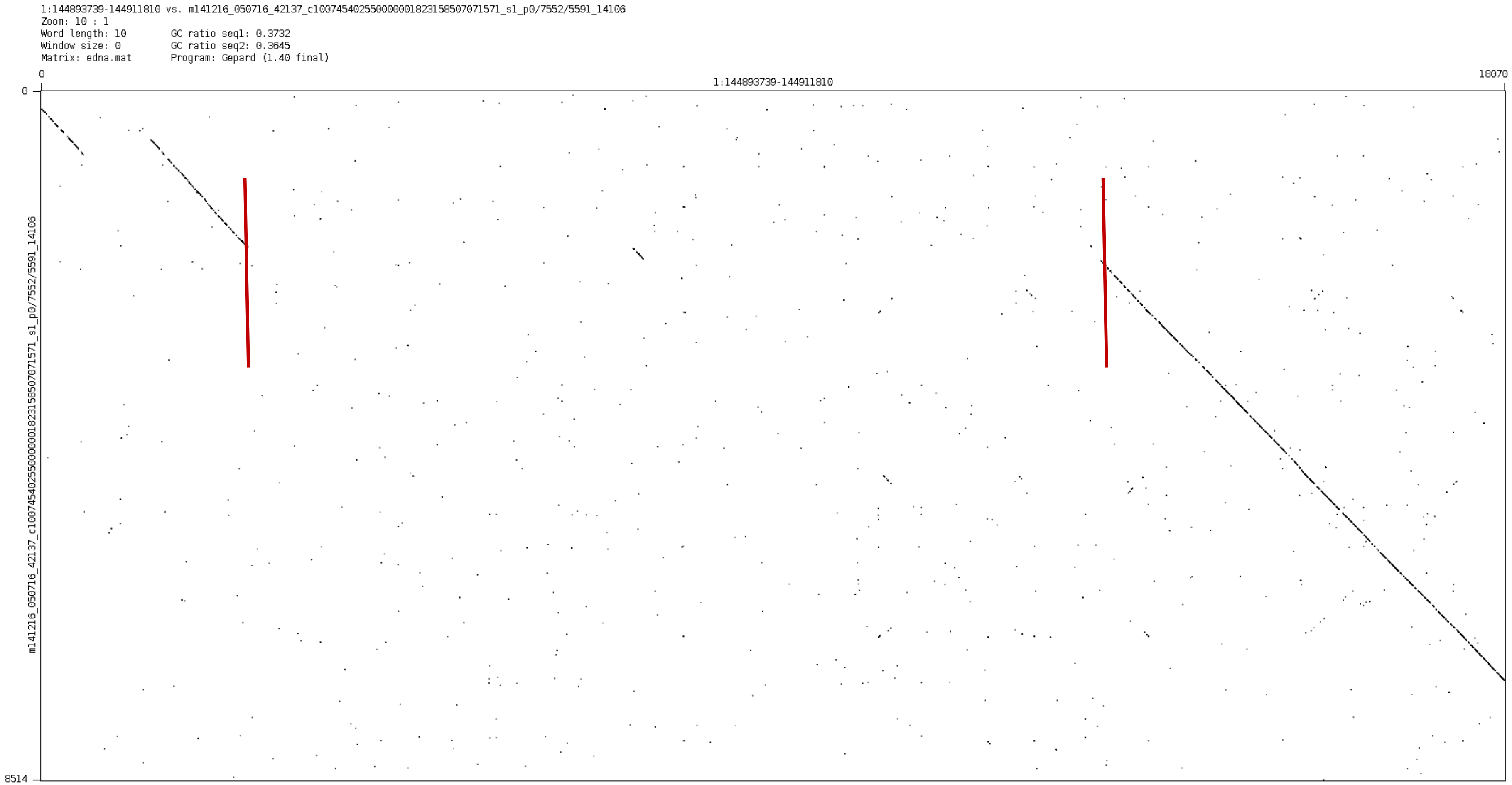
1:144893739-144911810 vs. m141216_050716_42137_c100745402550000001823158507071571_s1_p0/7552/5591_14106
Zoom: 10 : 1
Word length: 10
GC ratio seq1: 0.3732
Window size: 0
GC ratio seq2: 0.3645
Matrix: edna.mat
Program: Gepard (1.40 final)
0
18070
1:144893739-144911810
0
8514
m141216_050716_42137_c100745402550000001823158507071571_s1_p0/7552/5591_14106

chr1:187464829-187466729

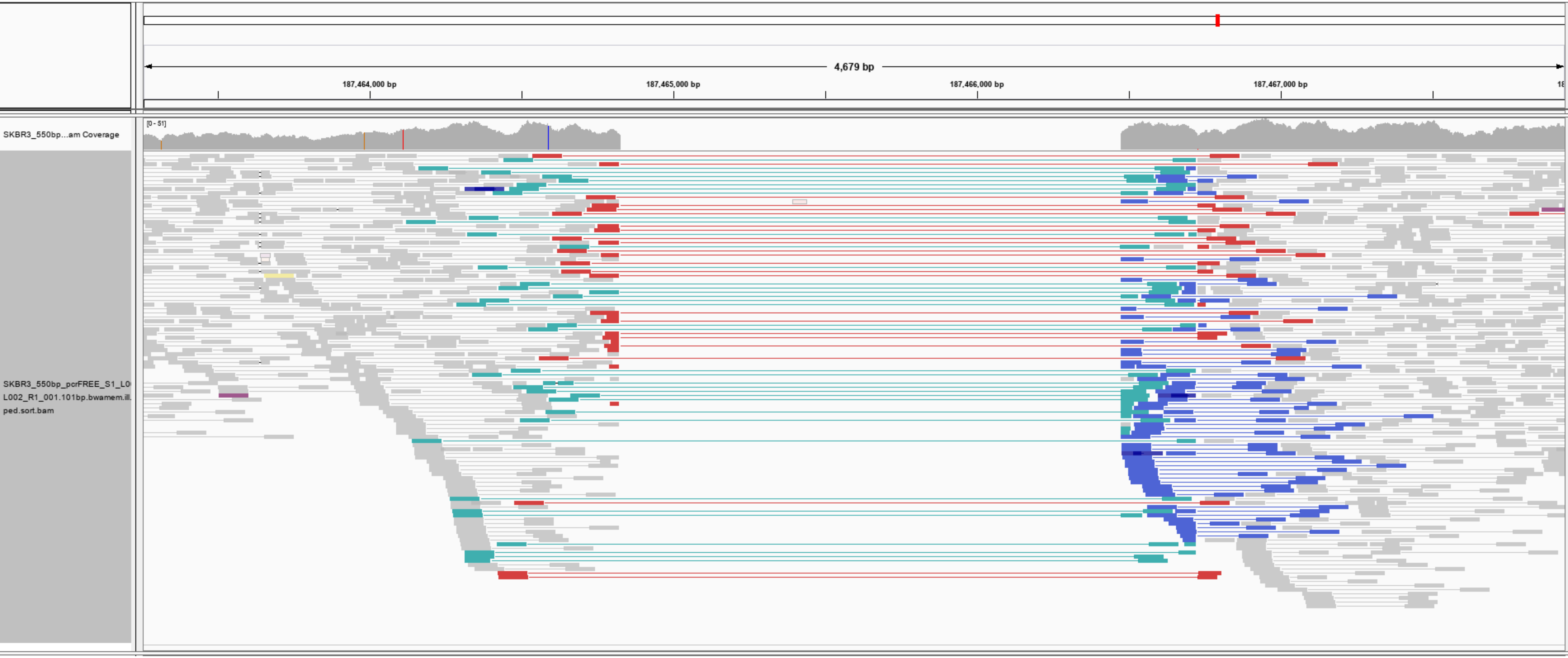

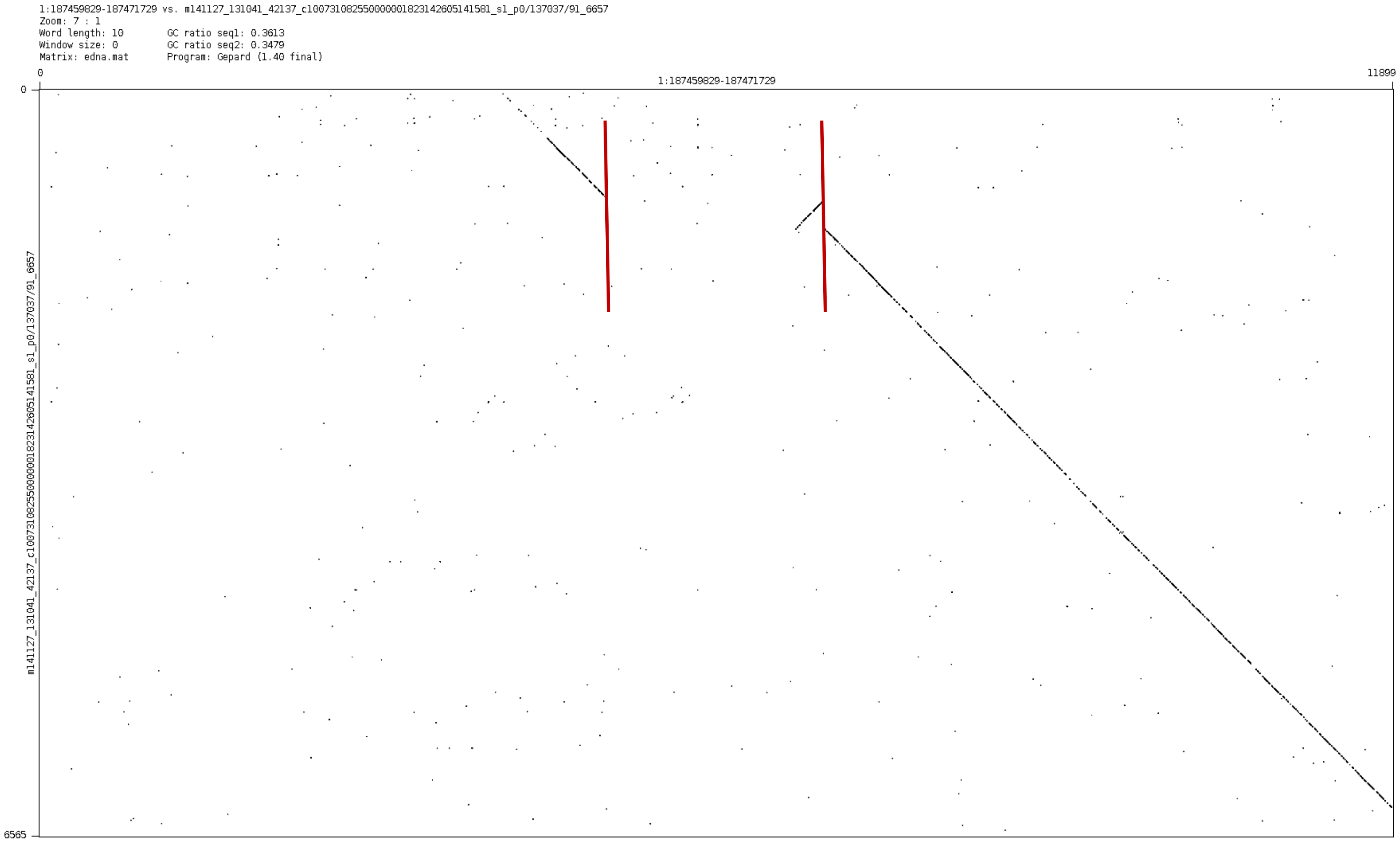
1:187459829-187471729 vs. m141127_131041_42137_c100731082550000001823142605141581_s1_p0/137037/91_6657
Zoom: 7 : 1
Word length: 10
GC ratio seq1: 0.3613
Window size: 0
GC ratio seq2: 0.3479
Matrix: edna.mat
Program: Gepard (1.40 final)
0
11899
1:187459829-187471729
0
6565
m141127_131041_42137_c100731082550000001823142605141581_s1_p0/137037/91_6657

chr1:207292347-207293196

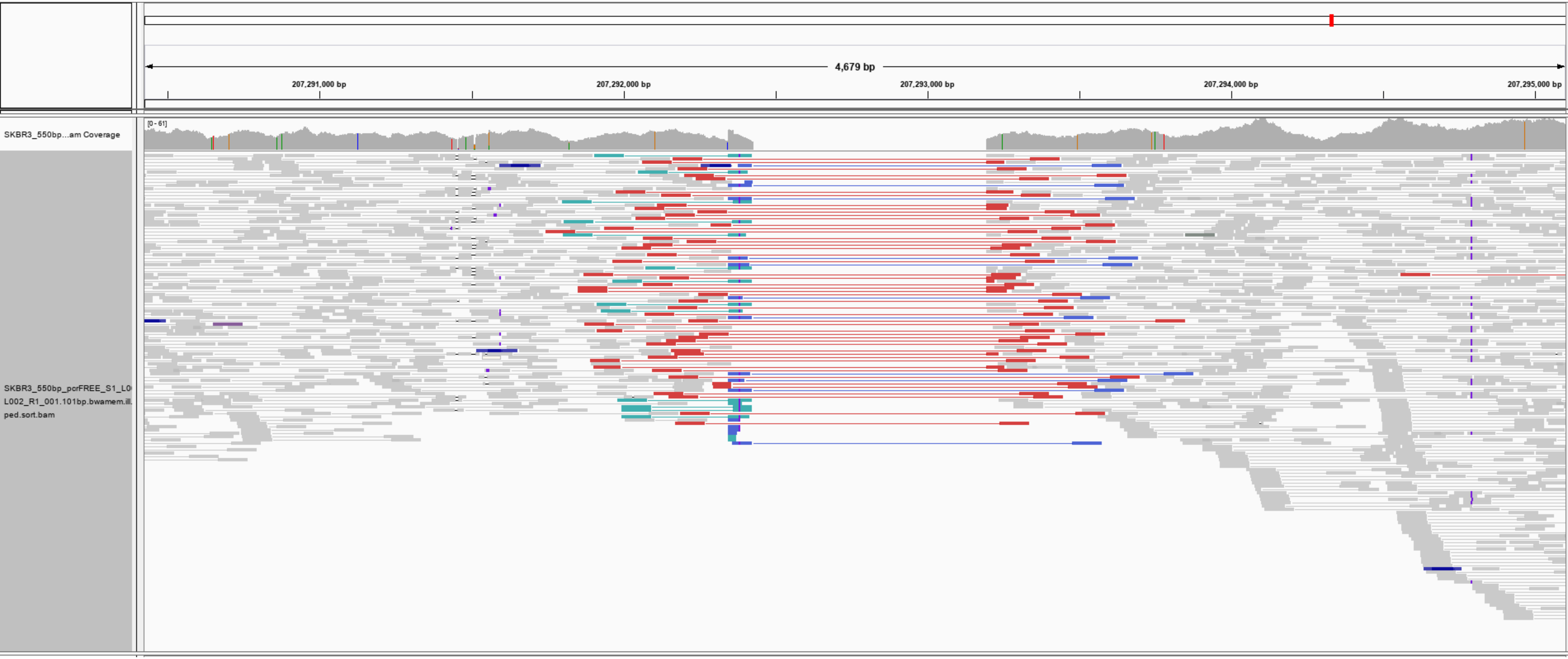

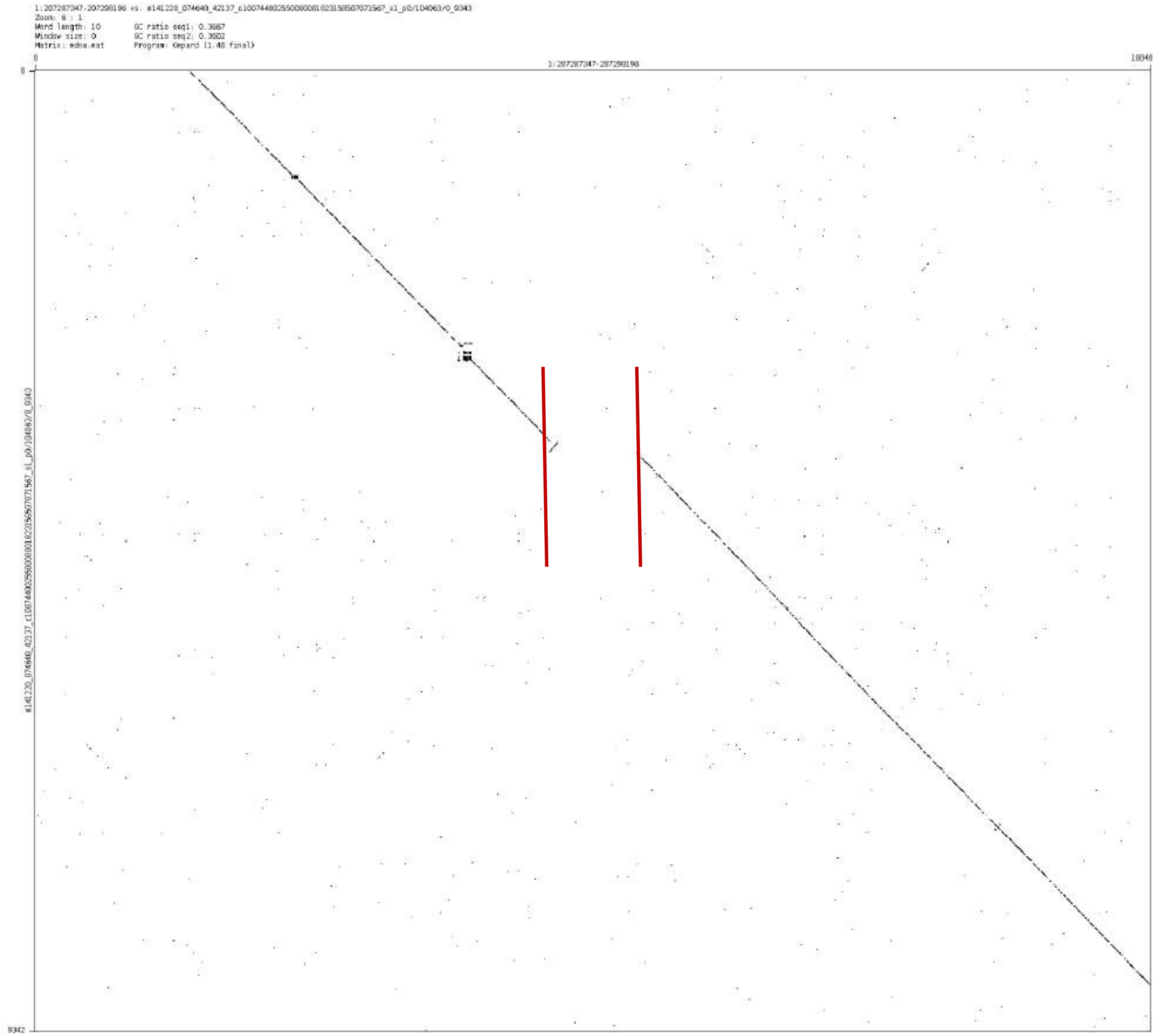
1:207287347-207298196 vs. m141228_074648_42137_c100744832550000001823158507071567_s1_p0/104063/0_9343
Zoom: 6 : 1
Word length: 10
Window size: 0
Matrix: edna.mat
GC ratio seq1: 0.3667
GC ratio seq2: 0.3602
Program: Gepard (1.40 final)
0
1:207287347-207298196
10848
0
9342

chr2:16406412-16407796

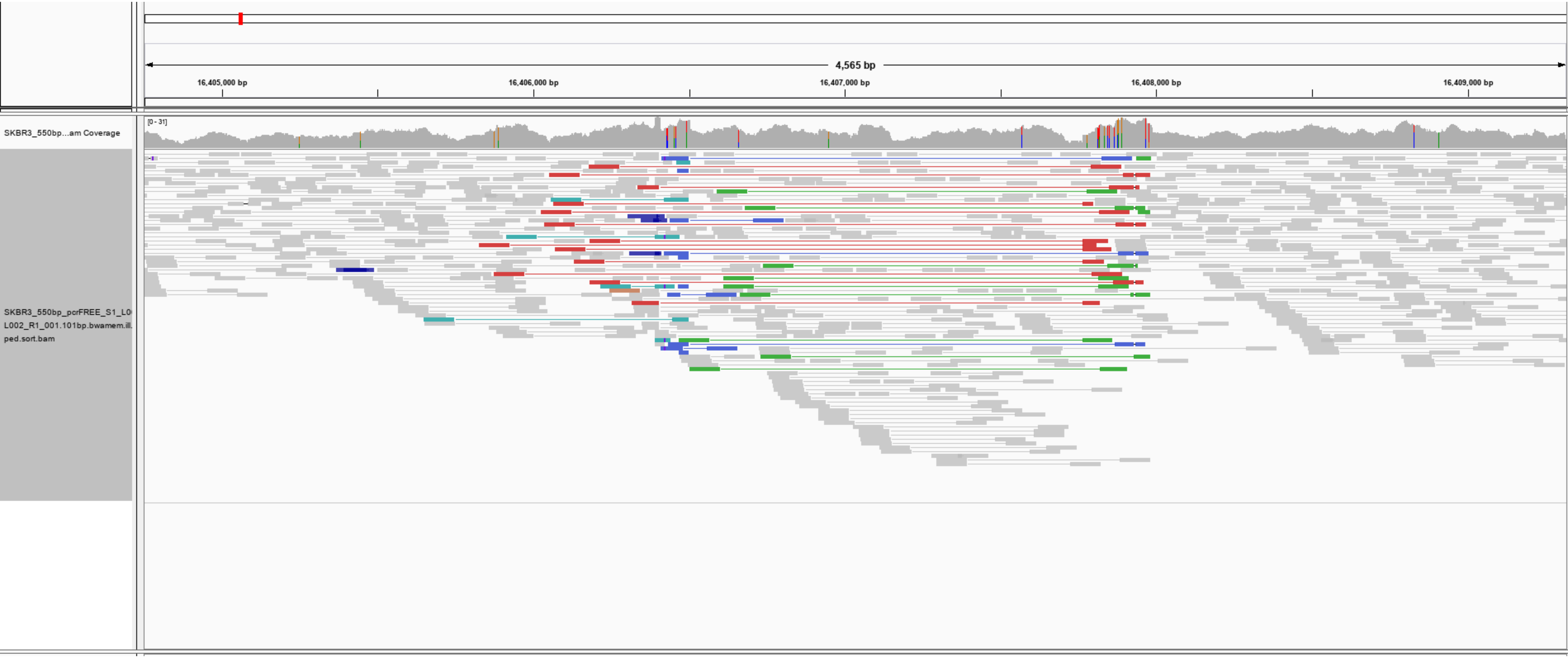

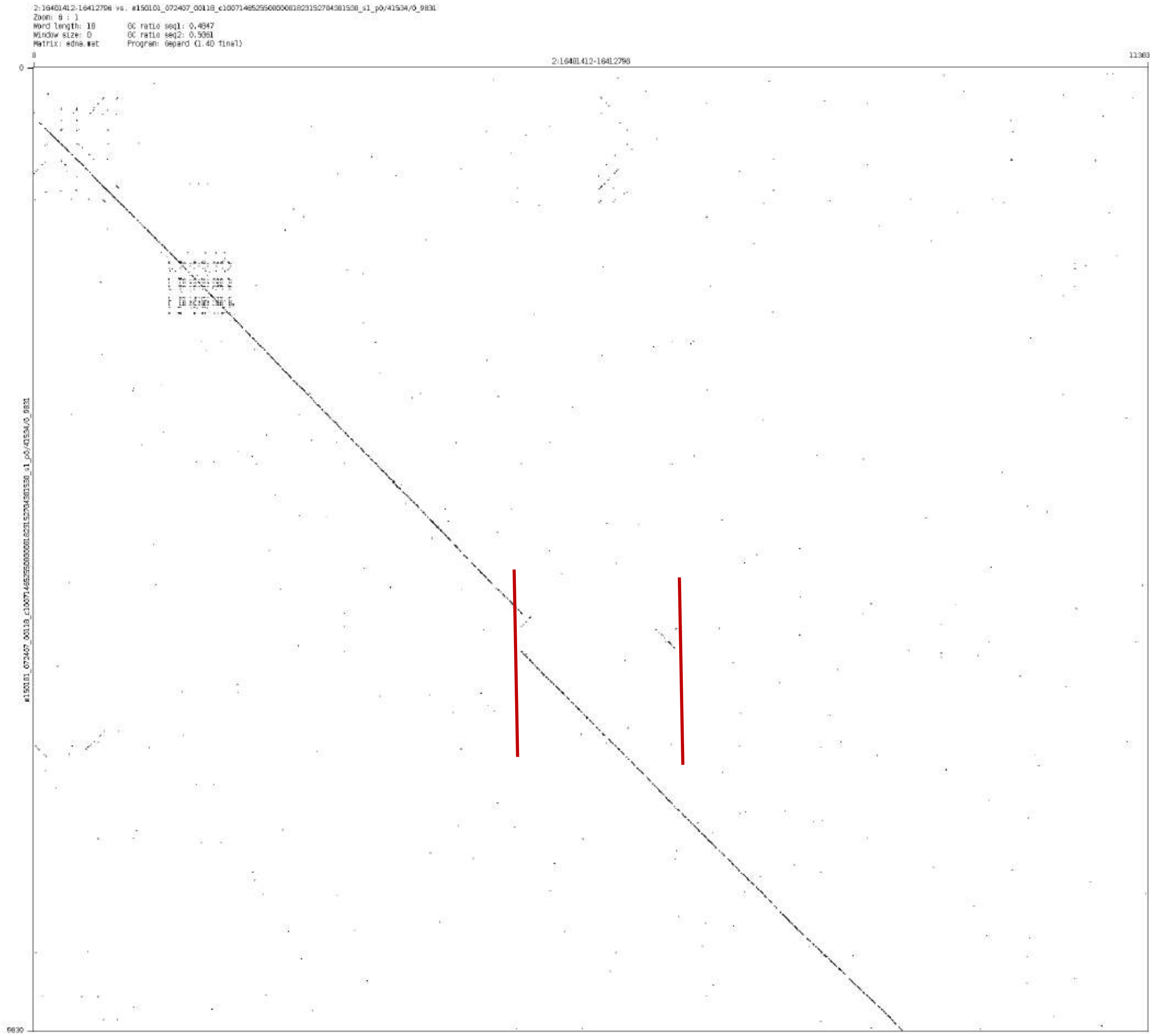
Zoom: 8 : 1
Word length: 10
GC ratio seq1: 0.4847
Window size: 0
GC ratio seq2: 0.5361
Matrix: edna.mat
Program: Gepard (1.40 final)

chr2:120417142-120418497

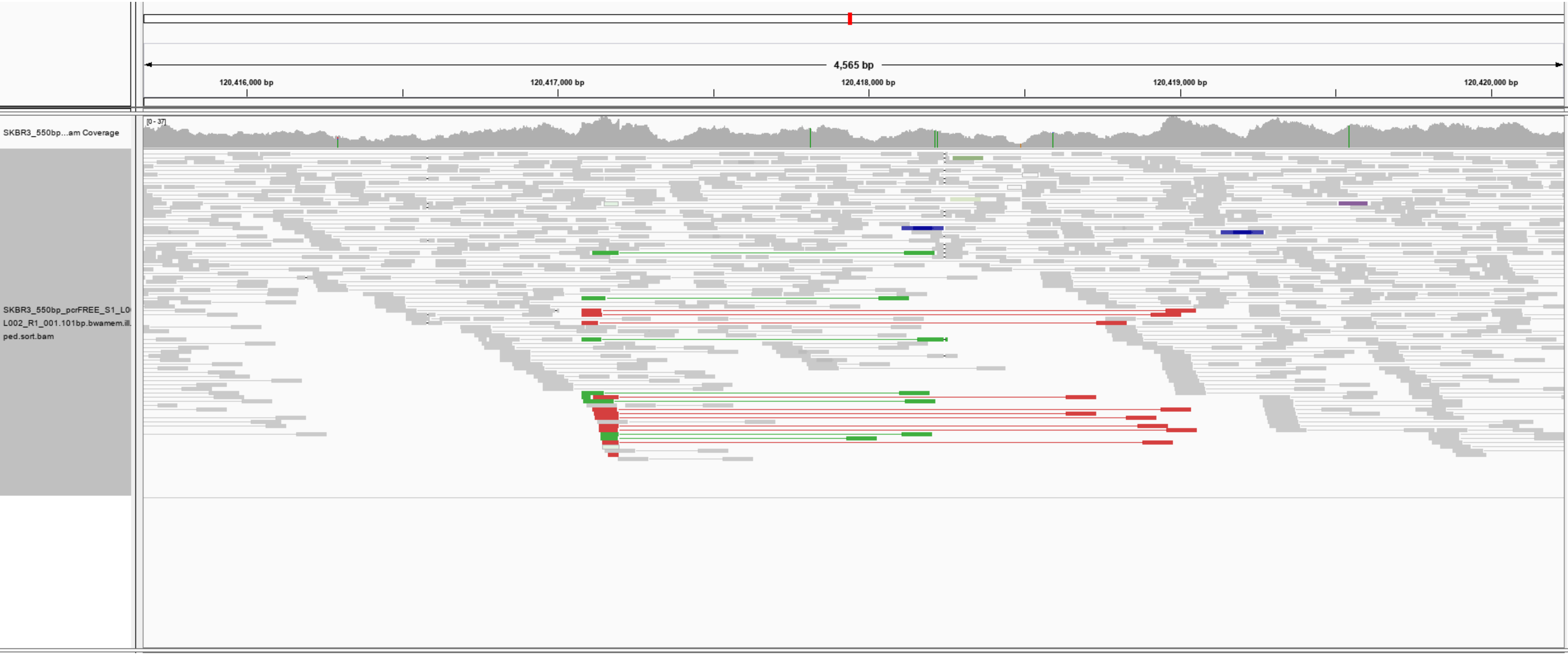

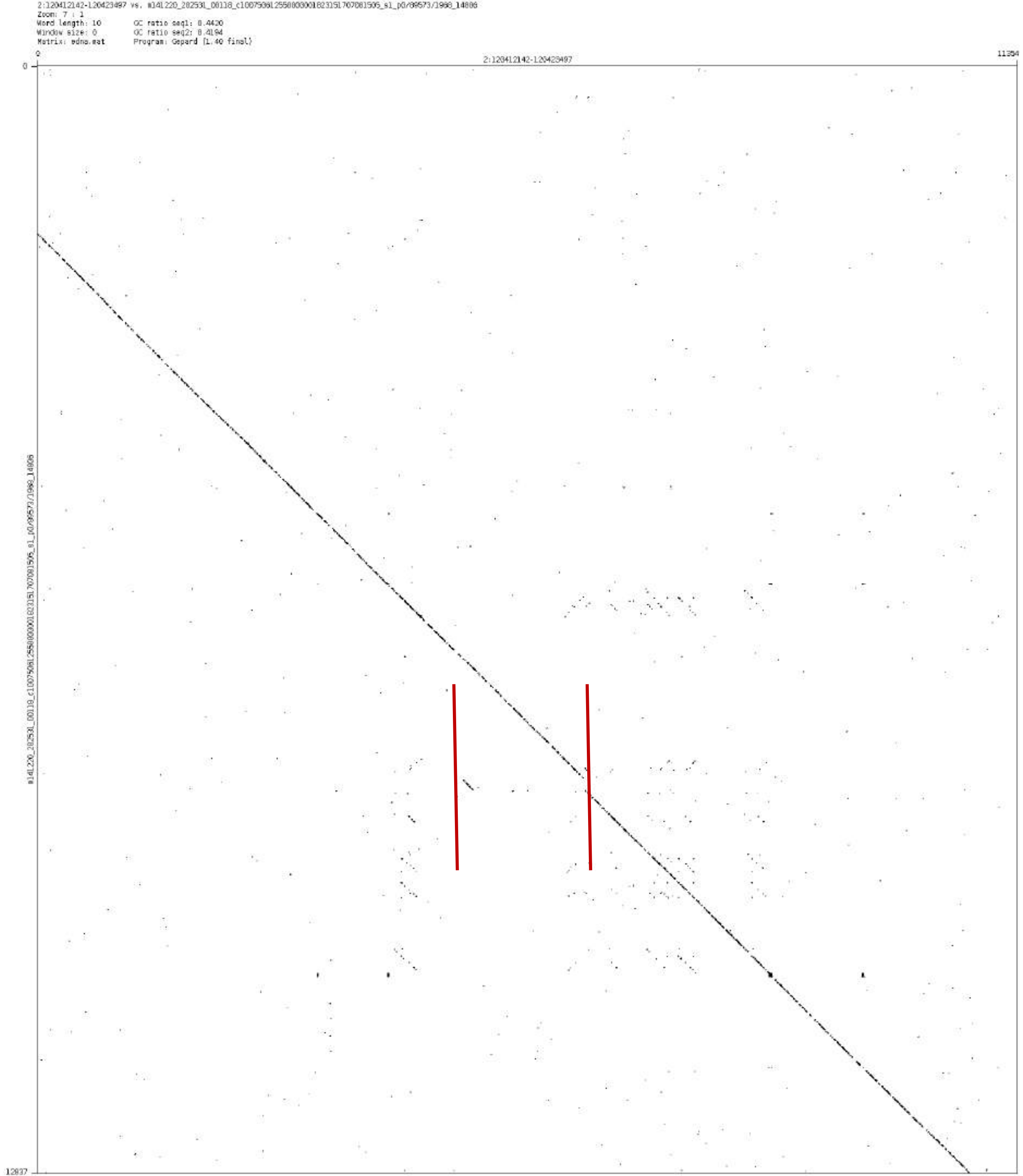

Zoom: 7 : 1
Word length: 10
Window size: 0
Matrix: edna.mat
GC ratio seq1: 0.4420
GC ratio seq2: 0.4194
Program: Gepard (1.40 final)
0
2:120412142-120423497
11354
0
12837

### chr2:225292798-225293577

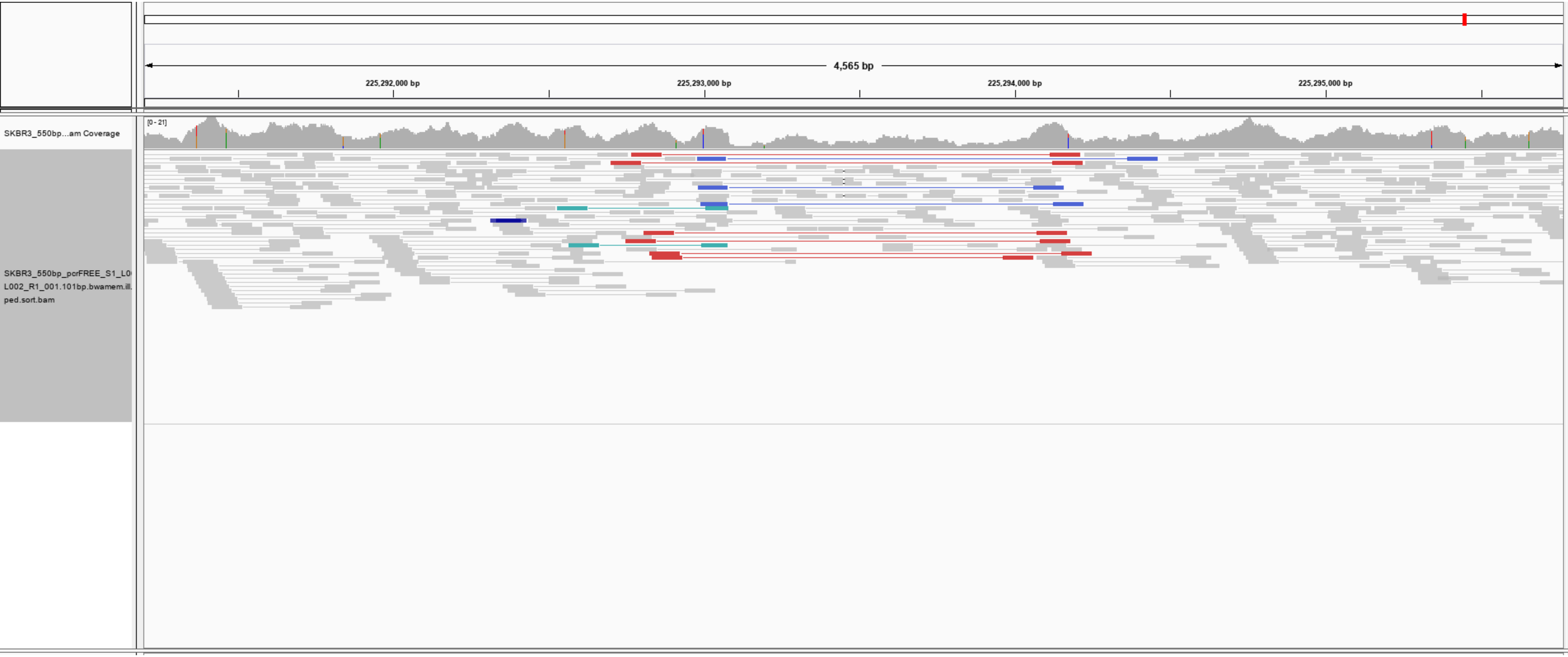

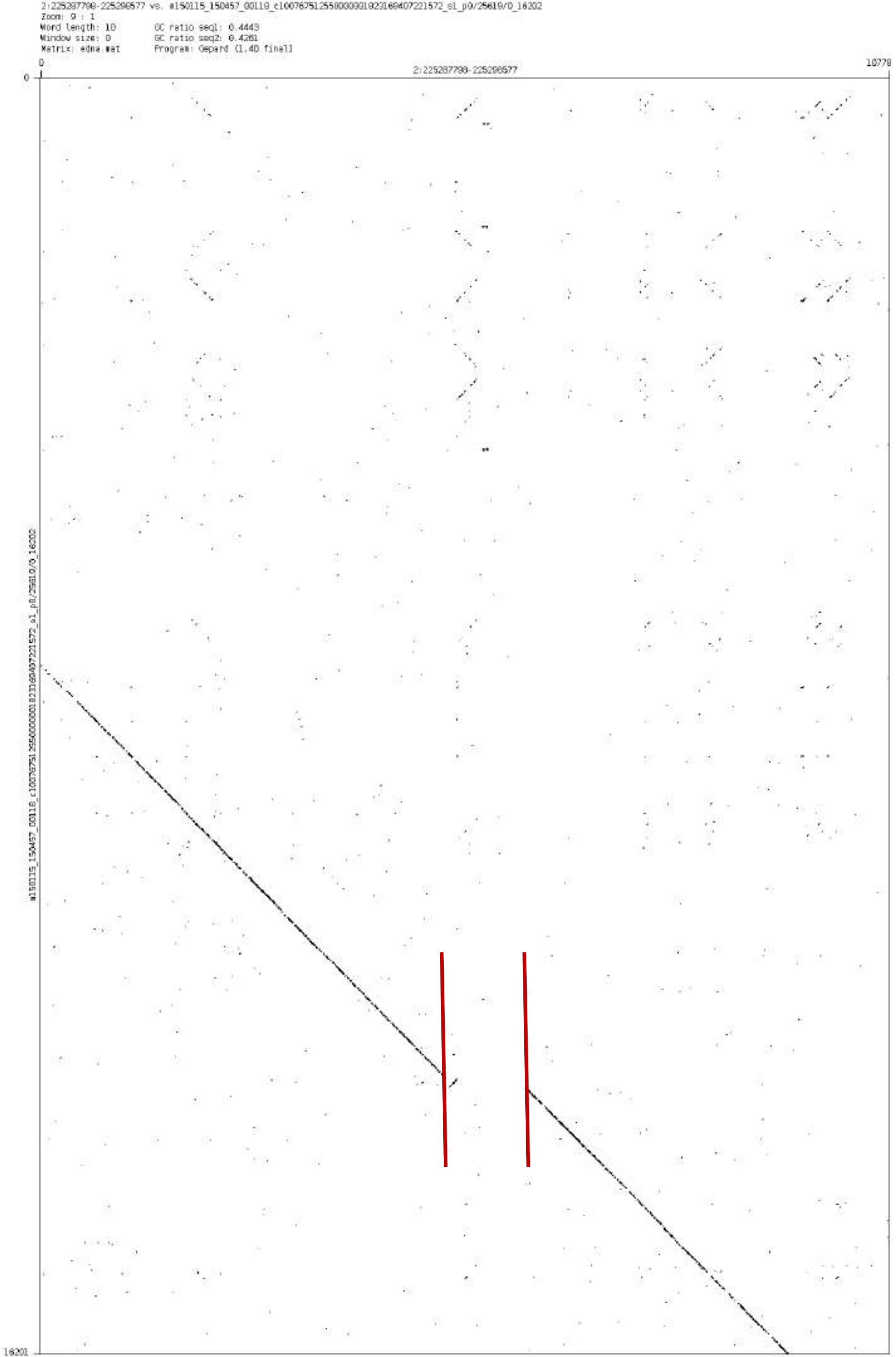
2:225287798-225298577 vs. m150115_150457_00118_c100767512550000001823169407221572_s1_p0/25619/0_16202
Zoom: 9 : 1
Word length: 10
Window size: 0
Matrix: edna.mat
GC ratio seq1: 0.4443
GC ratio seq2: 0.4261
Program: Gepard (1.40 final)
0
10779
2:225287798-225298577
0
16201

chr4:32065823-32071044

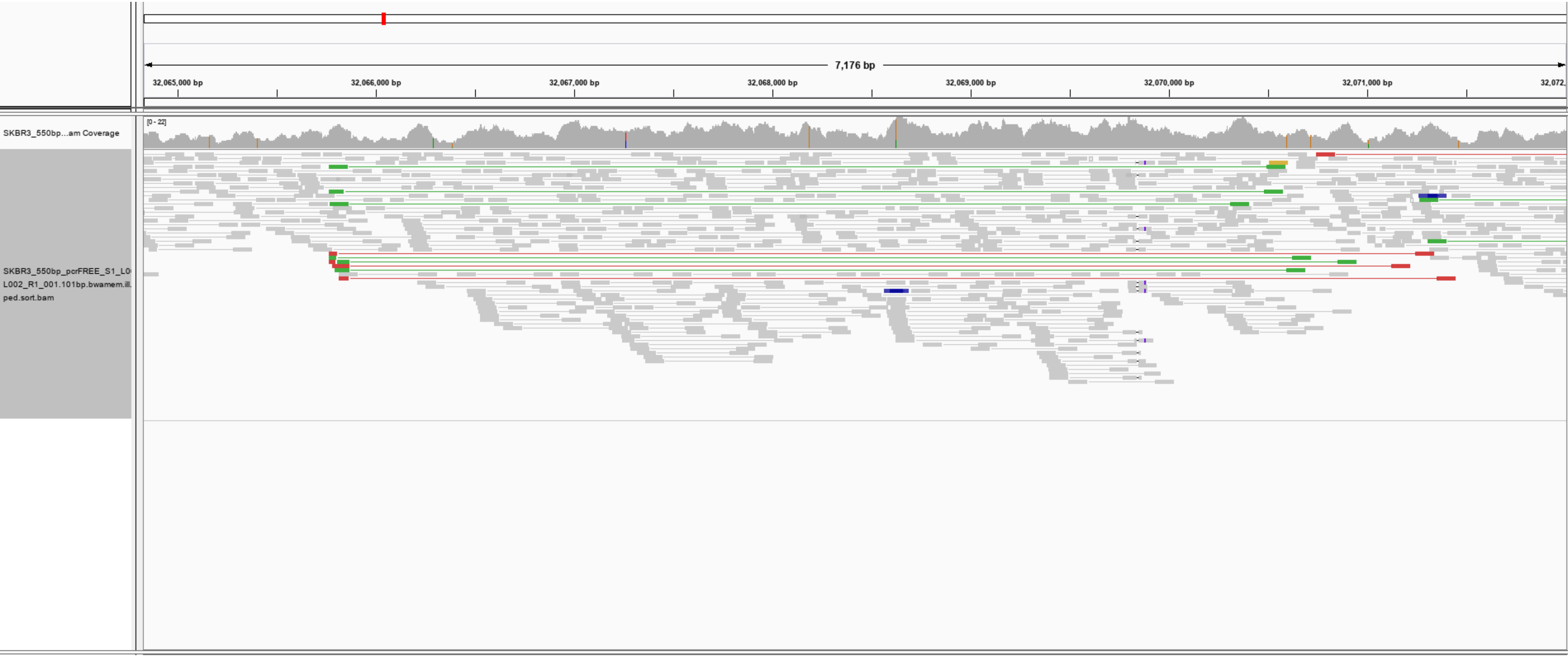

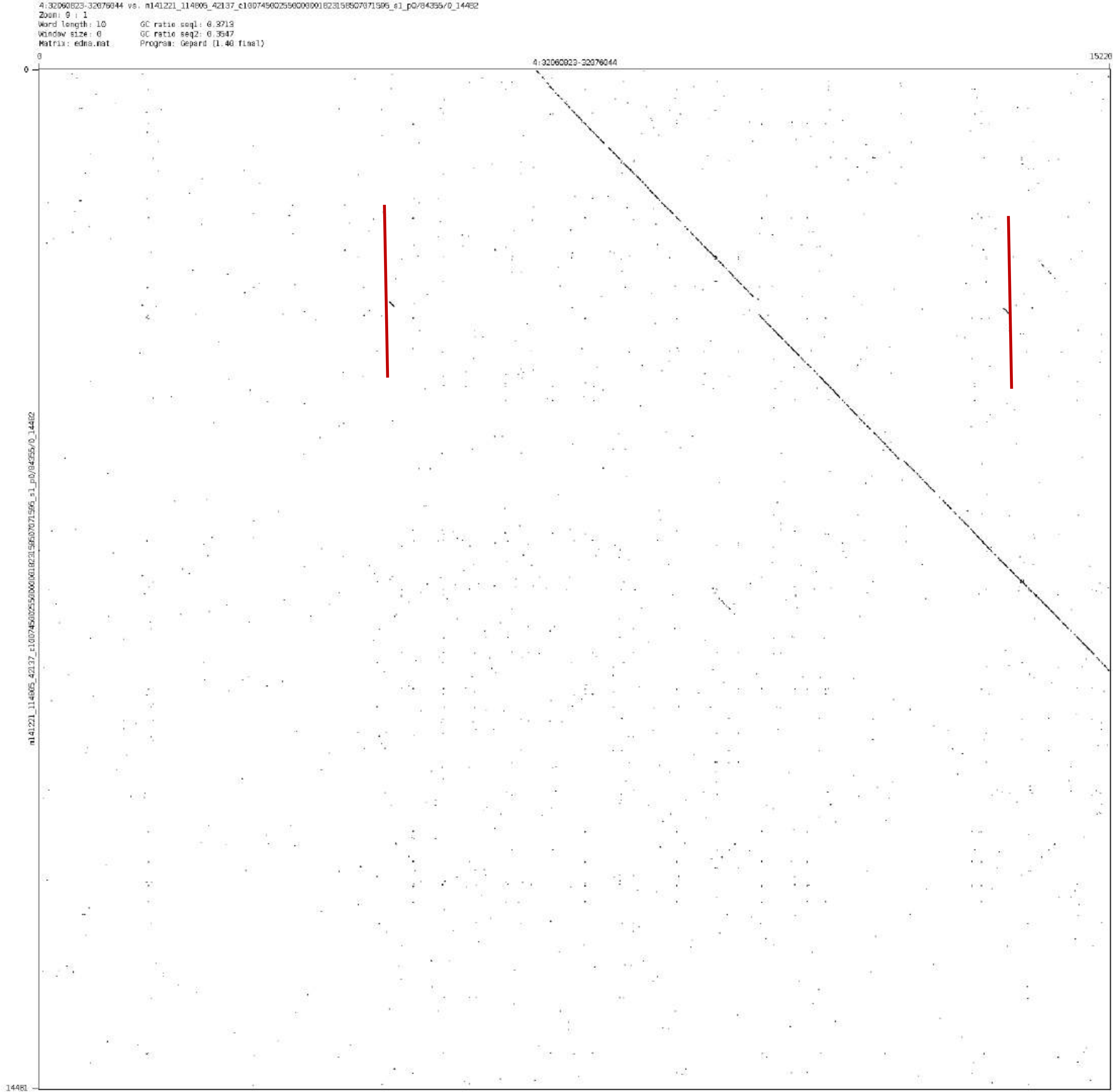
4:32060823-32076044 vs. m141221_114806_42137_c100745002550000001823158507071596_s1_p0/84355/0_14482
Zoom: 9 : 1
Word length: 10
Window size: 0
Matrix: edna.mat
GC ratio seq1: 0.3713
GC ratio seq2: 0.3547
Program: Gepard (1.40 final)
4:32060823-32076044
0
15220
0
14481
m141221_114806_42137_c100745002550000001823158507071596_s1_p0/84355/0_14482

chr4:93568459-93570189

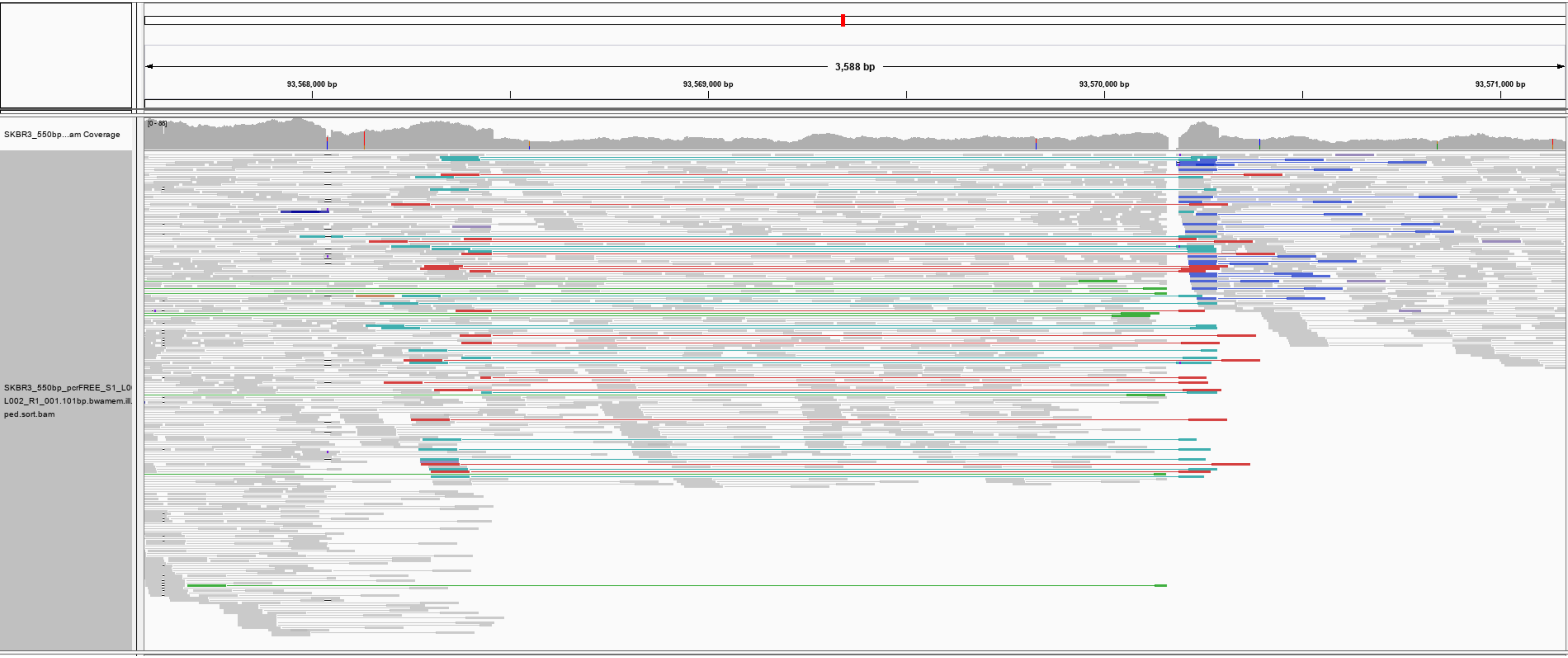

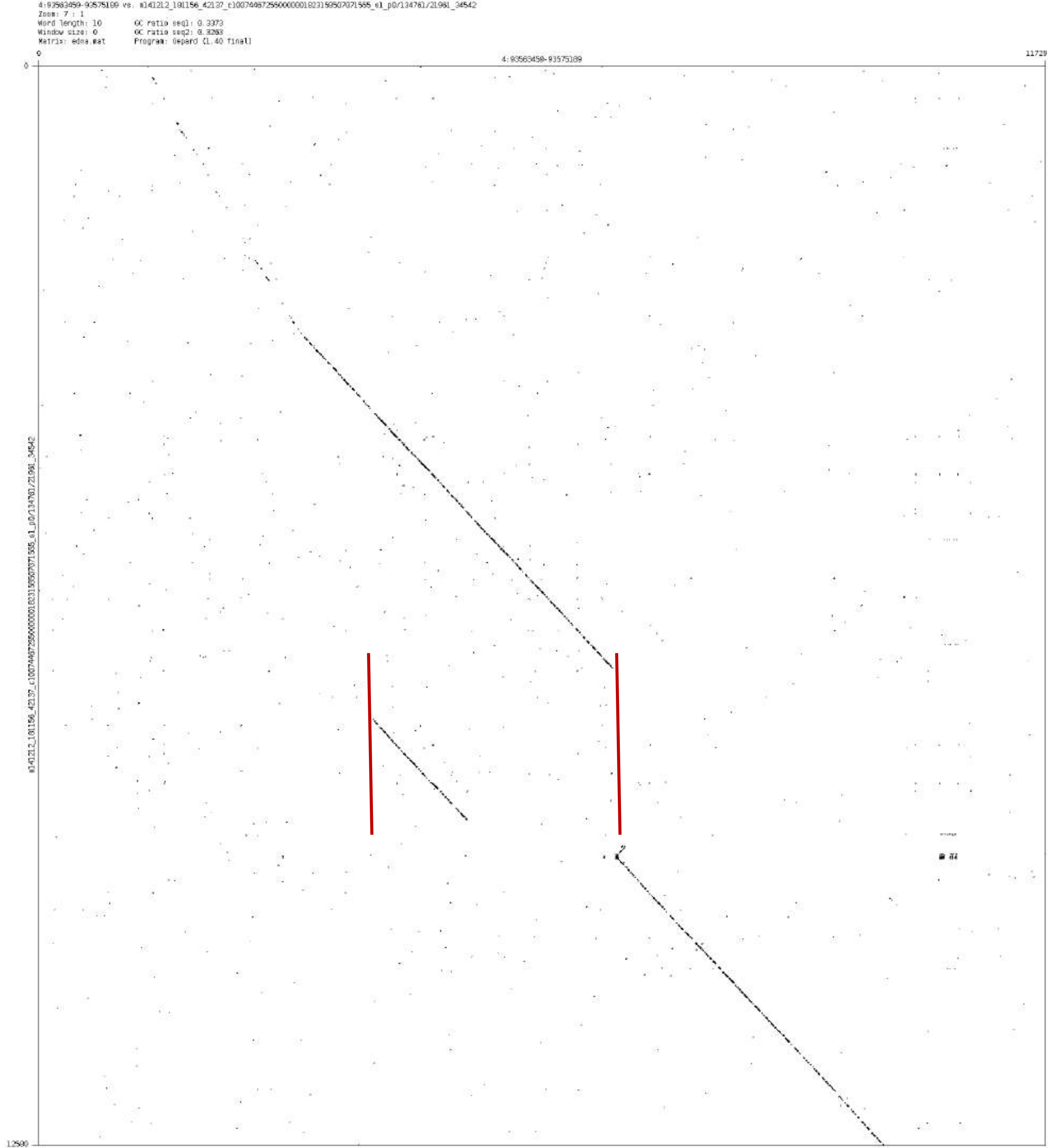

4:93563459-93575189 vs. m141212_181156_42137_c100744672550000001823159507071565_s1_p0/134761/21961_34542
Zoom: 7 : 1
Word length: 10
Window size: 0
Matrix: edna.mat
GC ratio seq1: 0.3373
GC ratio seq2: 0.3263
Program: Gepard (1.40 final)
0
4:93563459-93575189
11729
12500
m141212_181156_42137_c100744672550000001823159507071565_s1_p0/134761/21961_34542

chr5:151511018-151516780

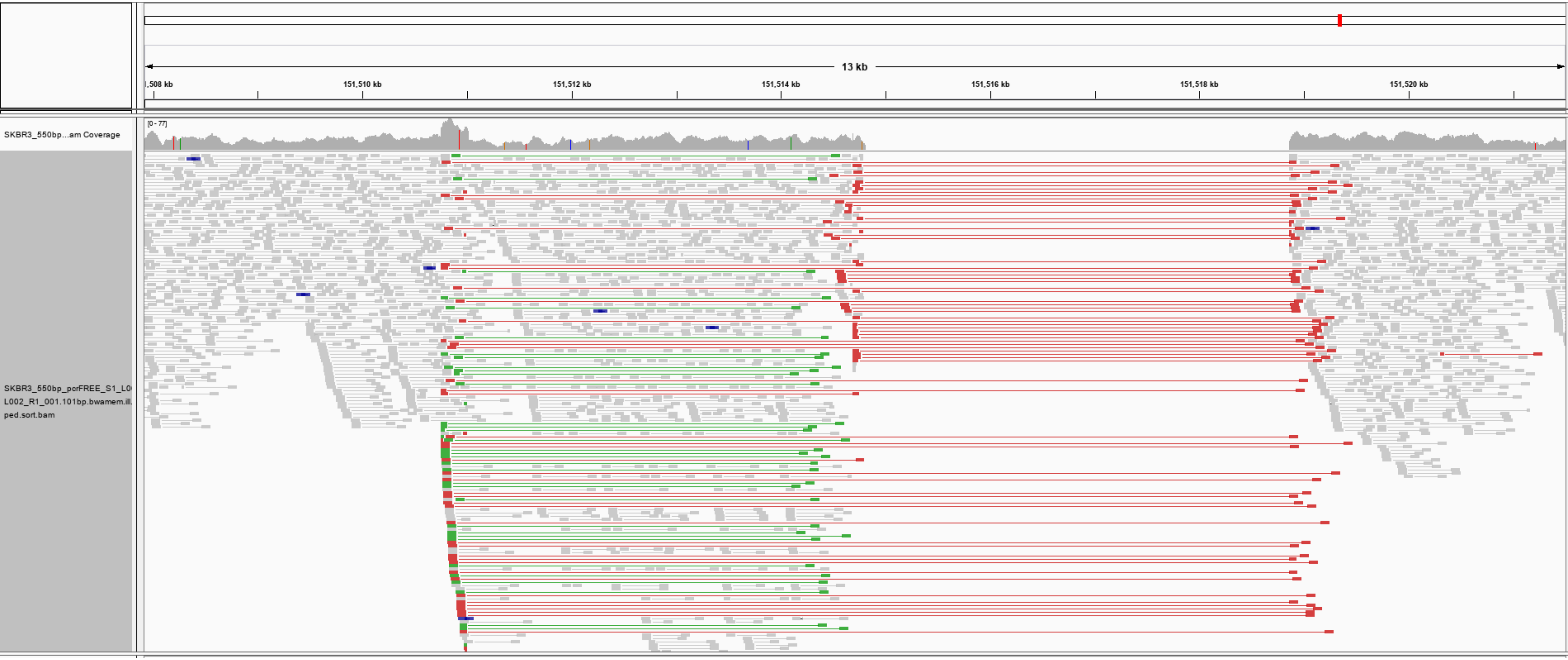

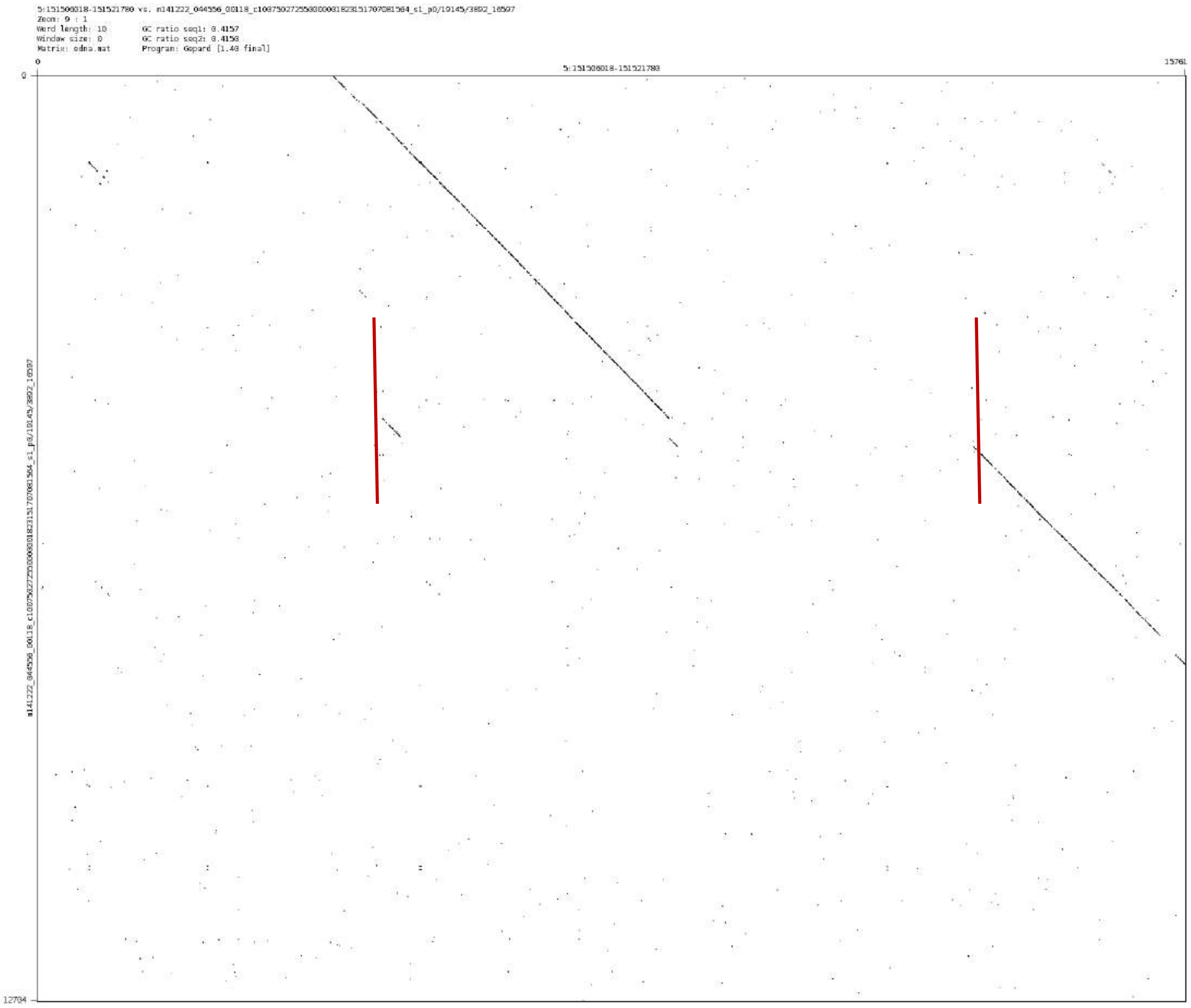
Zoom: 9 : 1
Word length: 10
Window size: 0
Matrix: edna.mat
GC ratio seq1: 0.4157
GC ratio seq2: 0.4158
Program: Gepard [1.40 final]
0
5:151506018-151521780
15761
12734

chr7:154876776-154880629

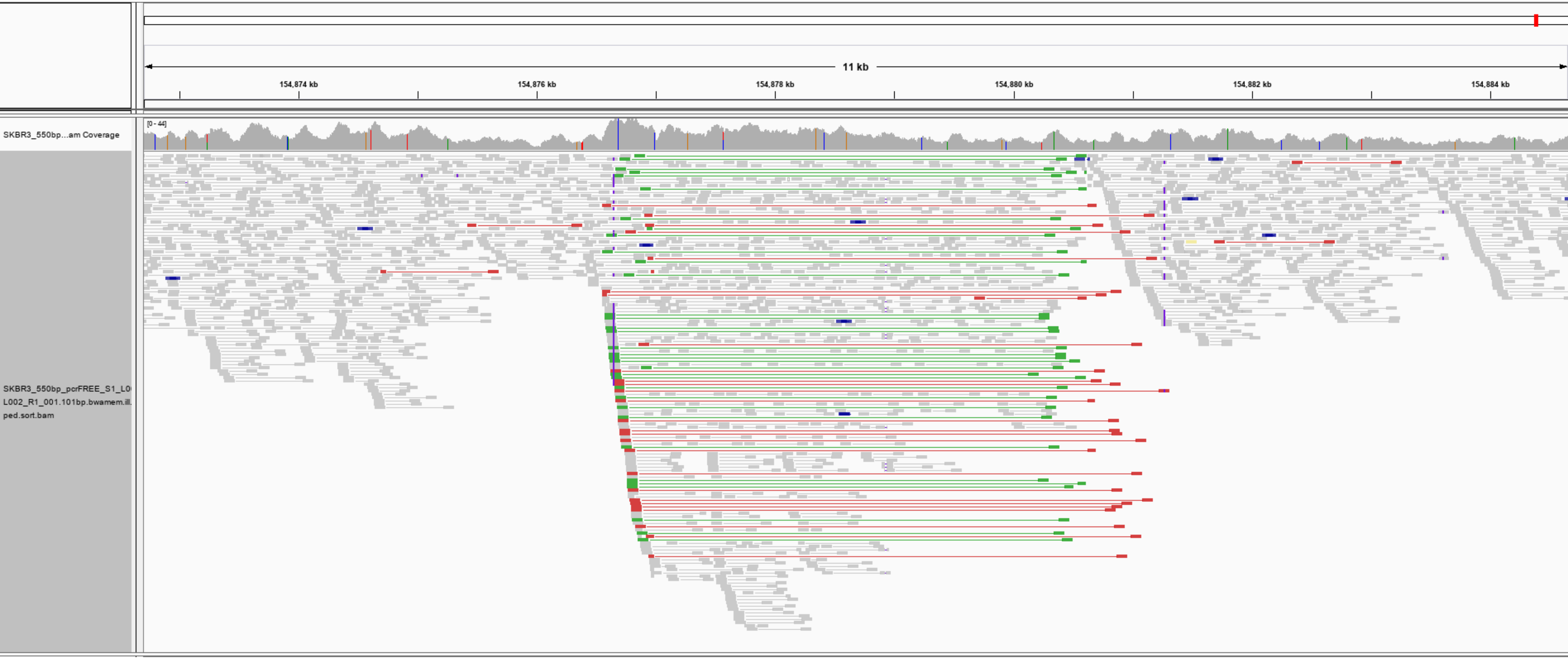

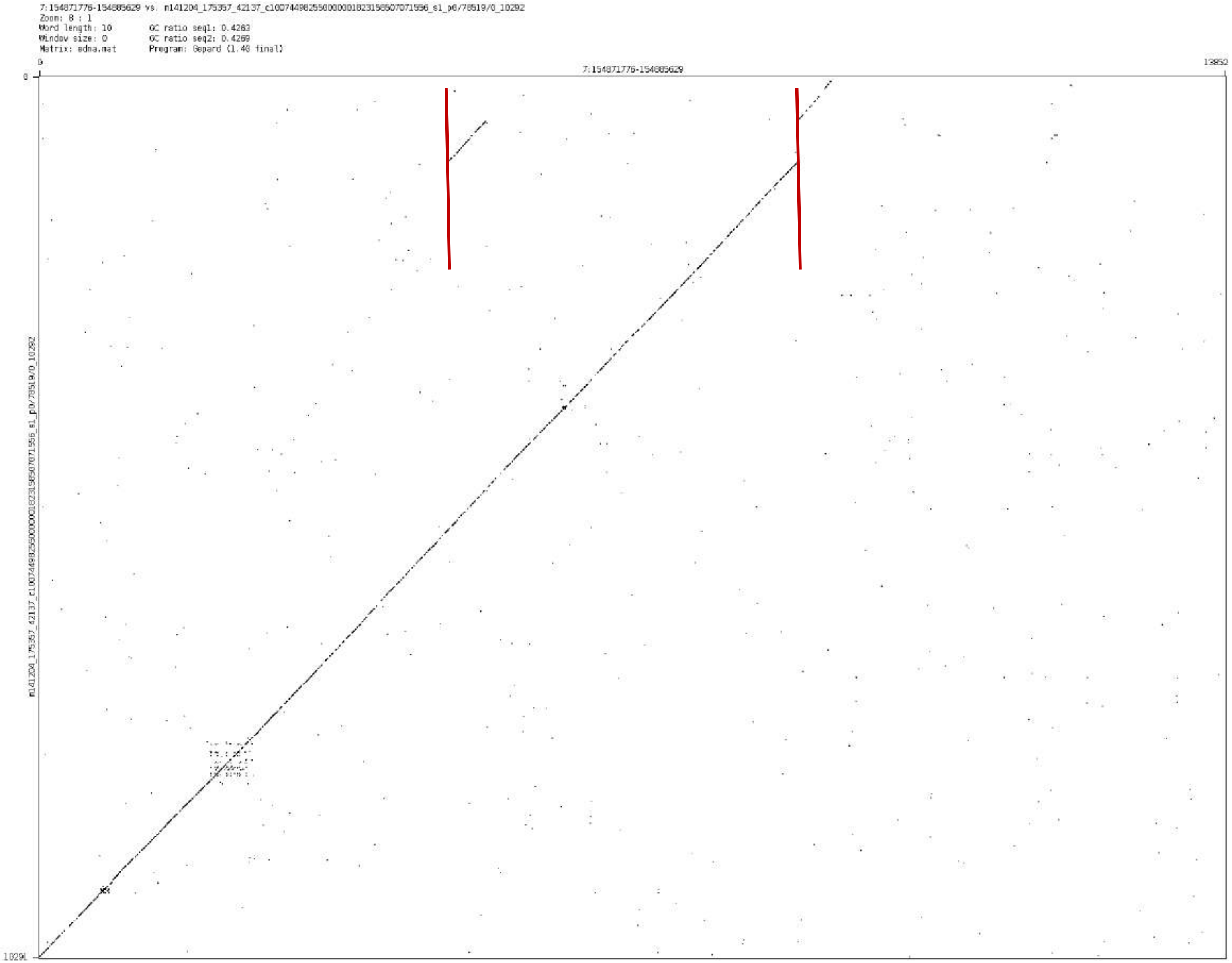
7:154871776-154885629 vs. m141204_175357_42137_c100744982550000001823156507071556_s1_p0/78519/0_10292
Zoom: 8 : 1
Word length: 10
GC ratio seq1: 0.4263
Window size: 0
GC ratio seq2: 0.4269
Matrix: edna.mat
Program: Gepard (1.40 final)
0
7:154871776-154885629
13852
0
10291
m141204_175357_42137_c100744982550000001823156507071556_s1_p0/78519/0_10292

chr9:89154436-89155939

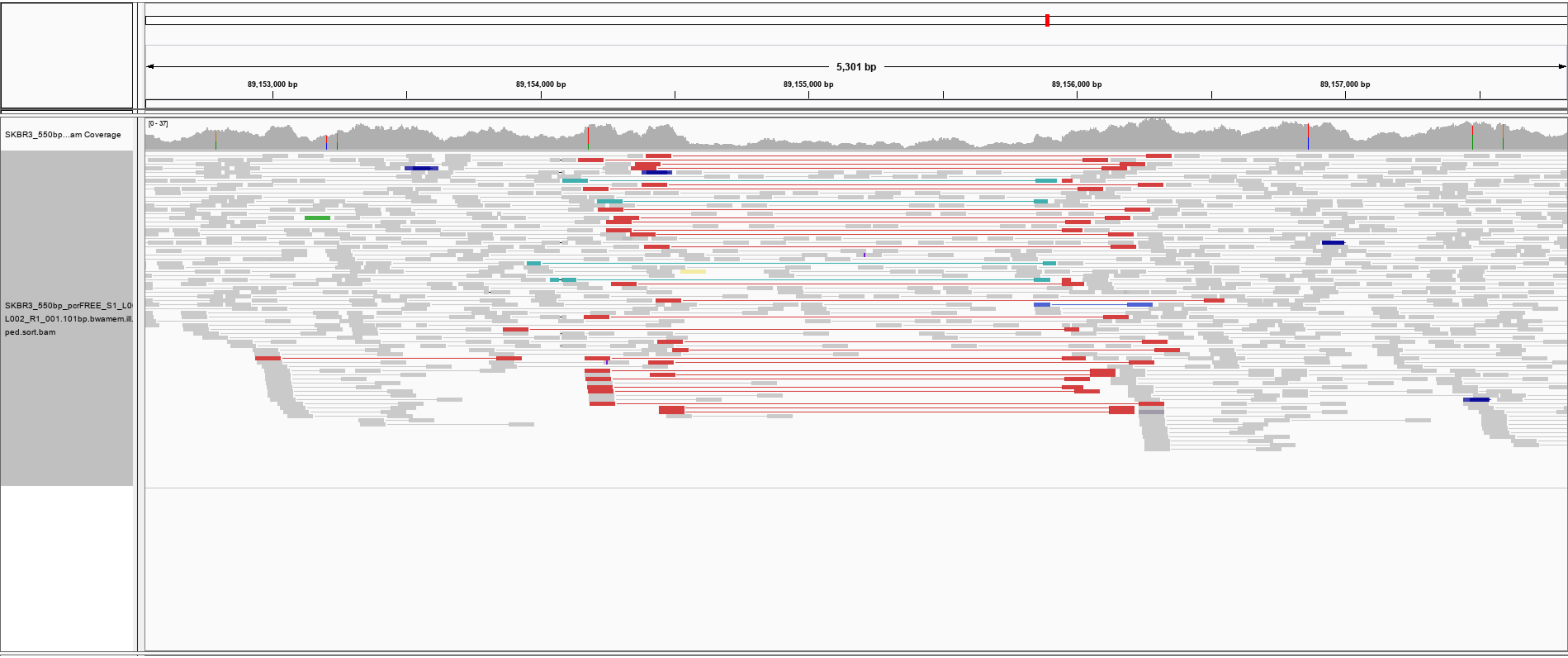

9:89149436-89160939 vs. m150110_025818_42137_c100744742550000001823165807071567_s1_p0/13189/0_6833
Zoom: 7 : 1
Word length: 10 GC ratio seq1: 0.4011
Window size: 0 GC ratio seq2: 0.3767
Matrix: edna.mat Program: Gepard (1.40 final)

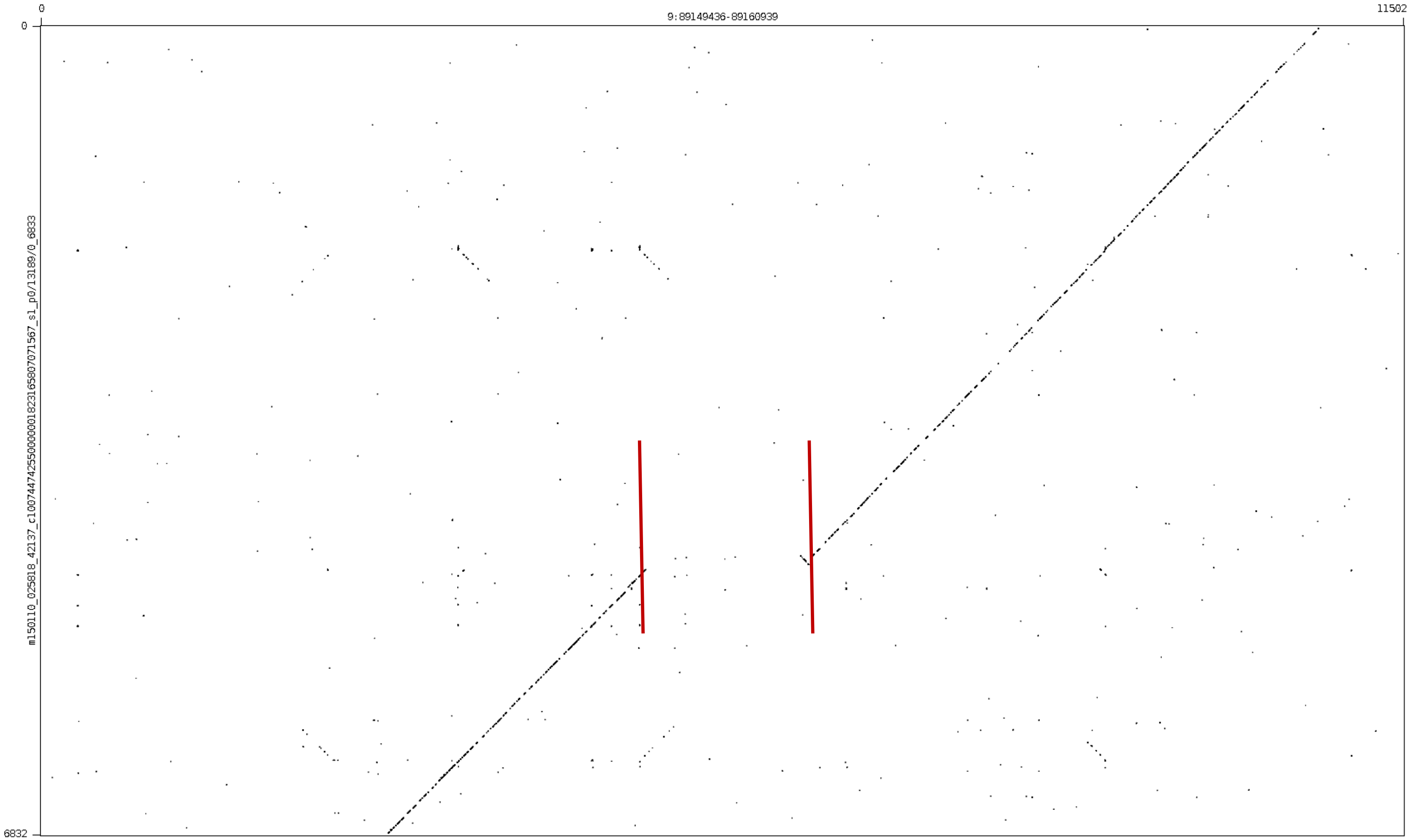

chr9:112285937-112286755

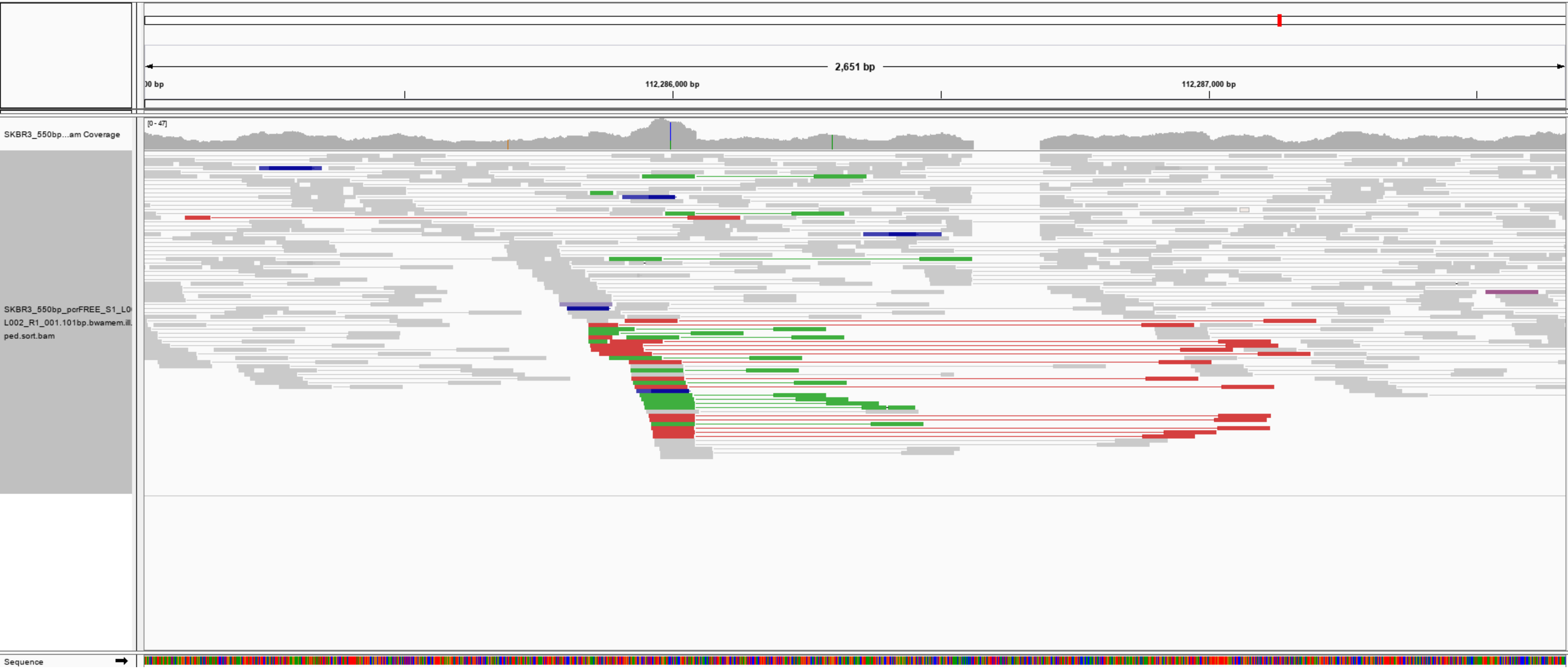

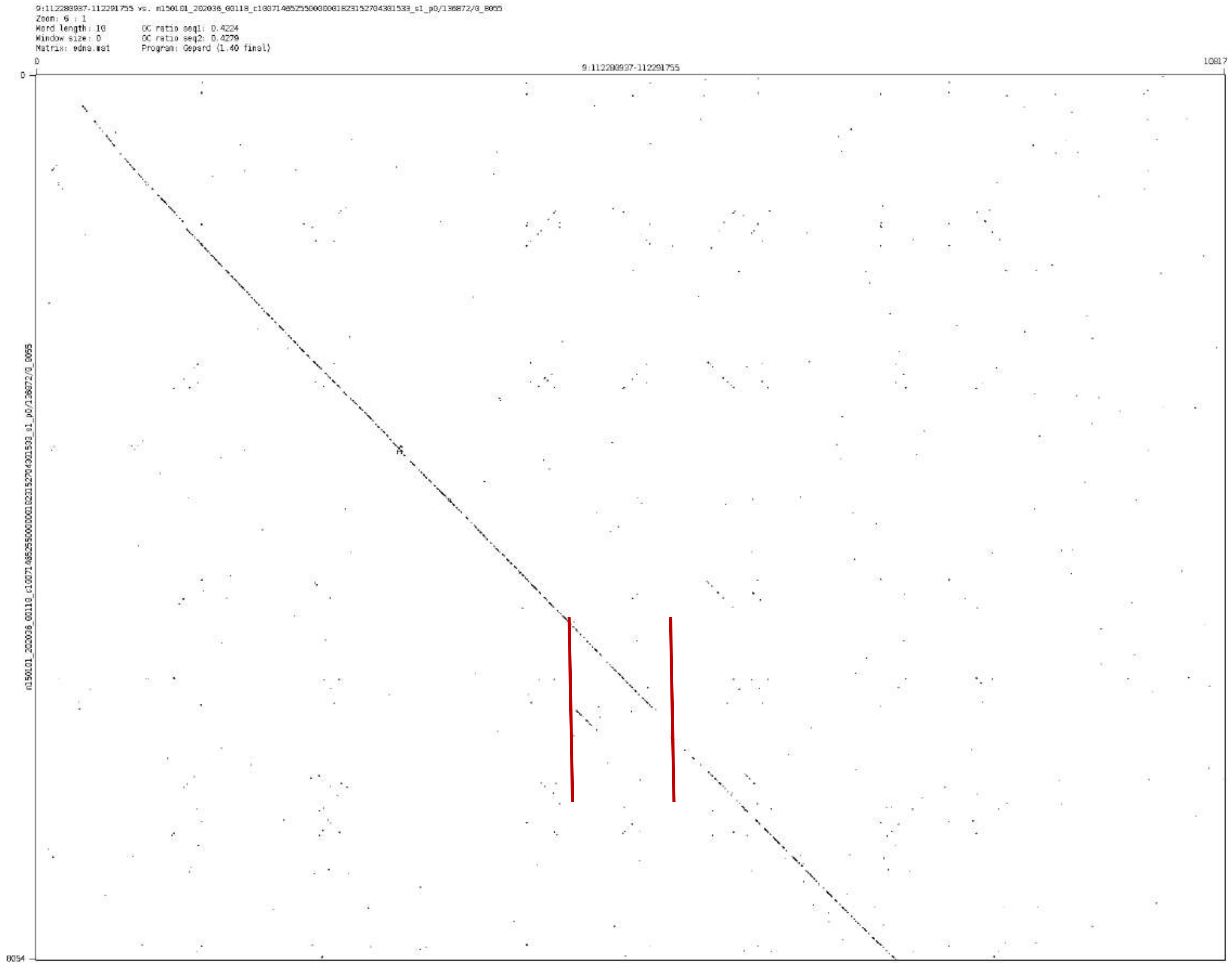

9:112288937-112291755 vs. m150101_202036_00118_c100714852550000001823152704301533_s1_p0/136872/0_8055
Zoom: 6 : 1
Word length: 10
Window size: 0
Matrix: edna.mat
GC ratio seq1: 0.4224
GC ratio seq2: 0.4279
Program: Gepard (1.40 final)
9:112288937-112291755
0
10817
8054
m150101_202036_00118_c100714852550000001823152704301533_s1_p0/136872/0_8055

chr10:127191013-127197233

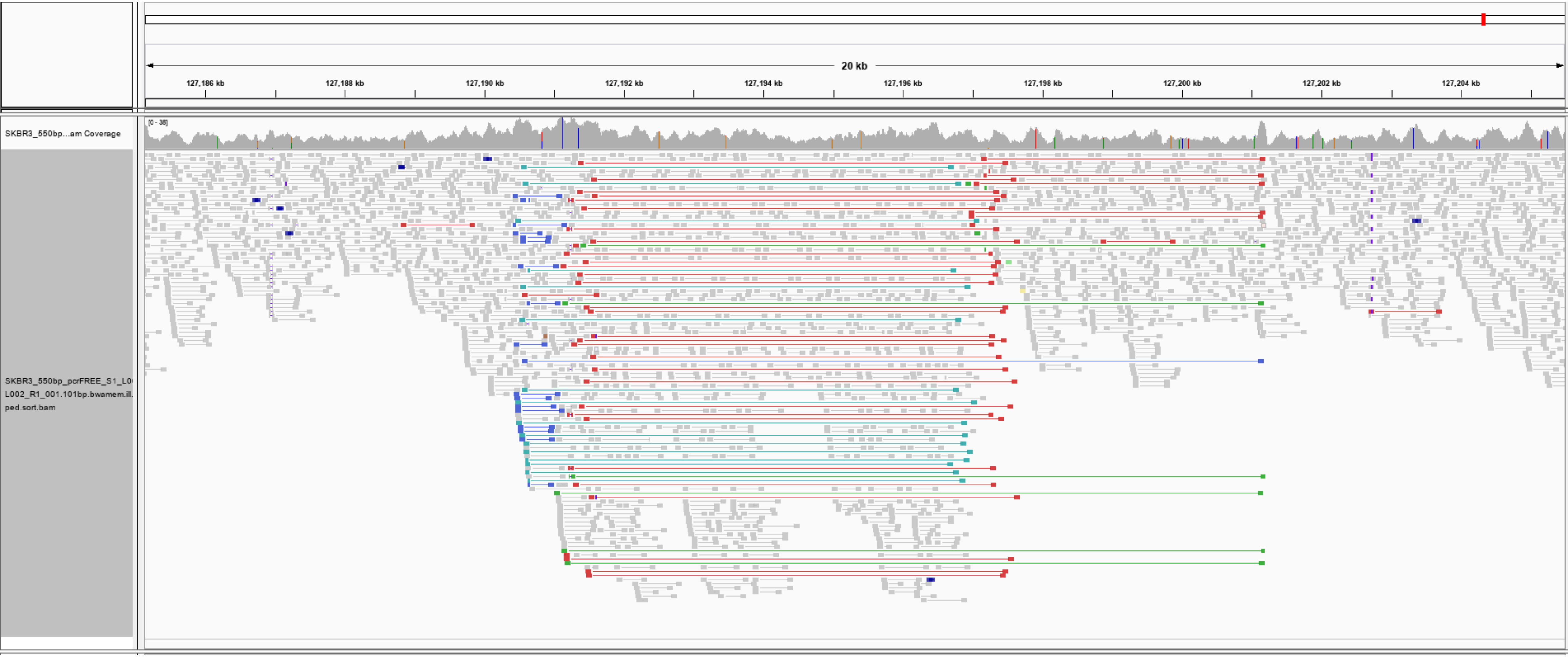

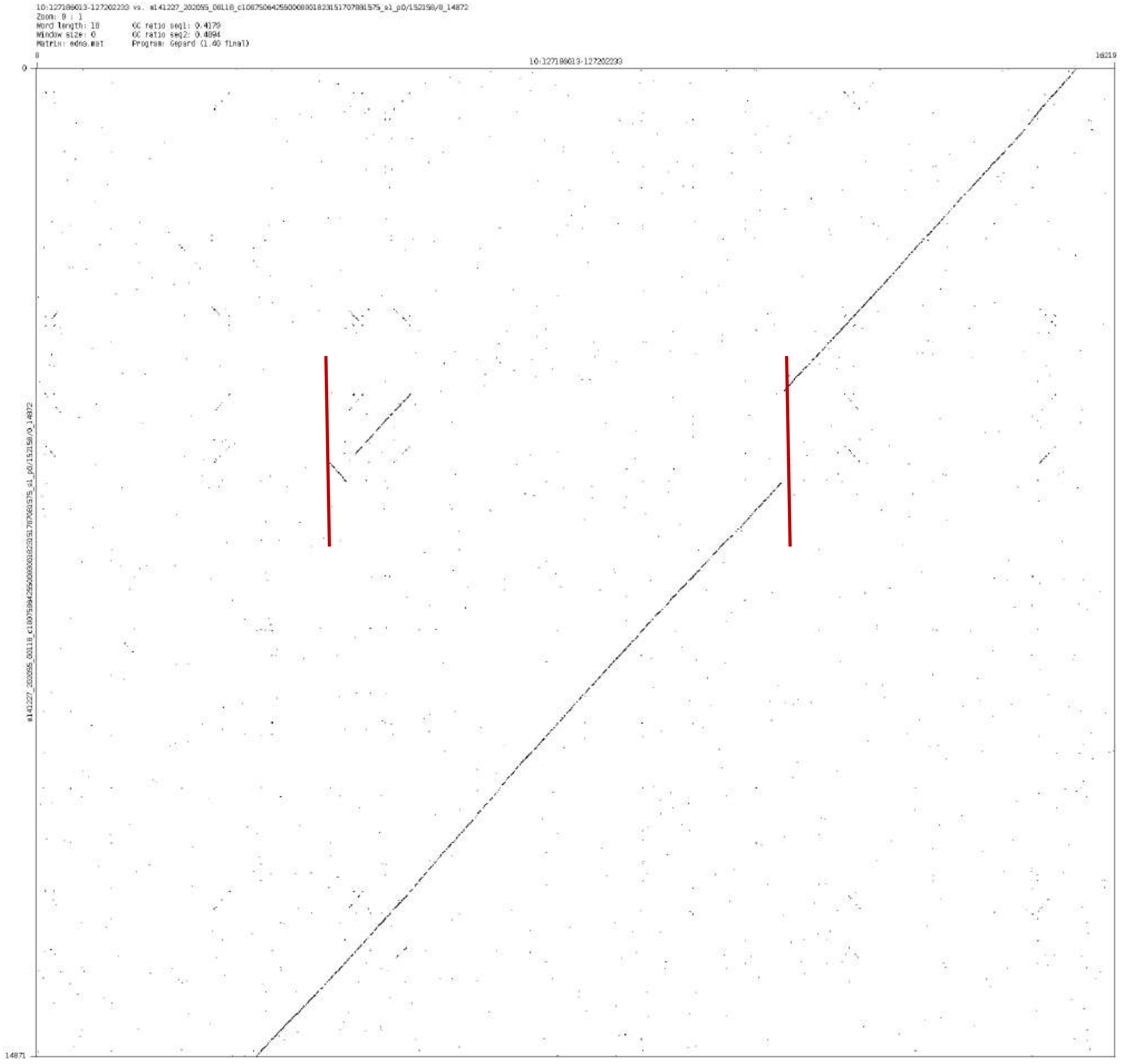

Zoom: 9 : 1
Word length: 10
Window size: 0
Matrix: edna.mat
GC ratio seq1: 0.4179
GC ratio seq2: 0.4894
Program: Gepard (1.40 final)
10:127186013-127202233
0
16219
14871

### chr11:63698921-63701800

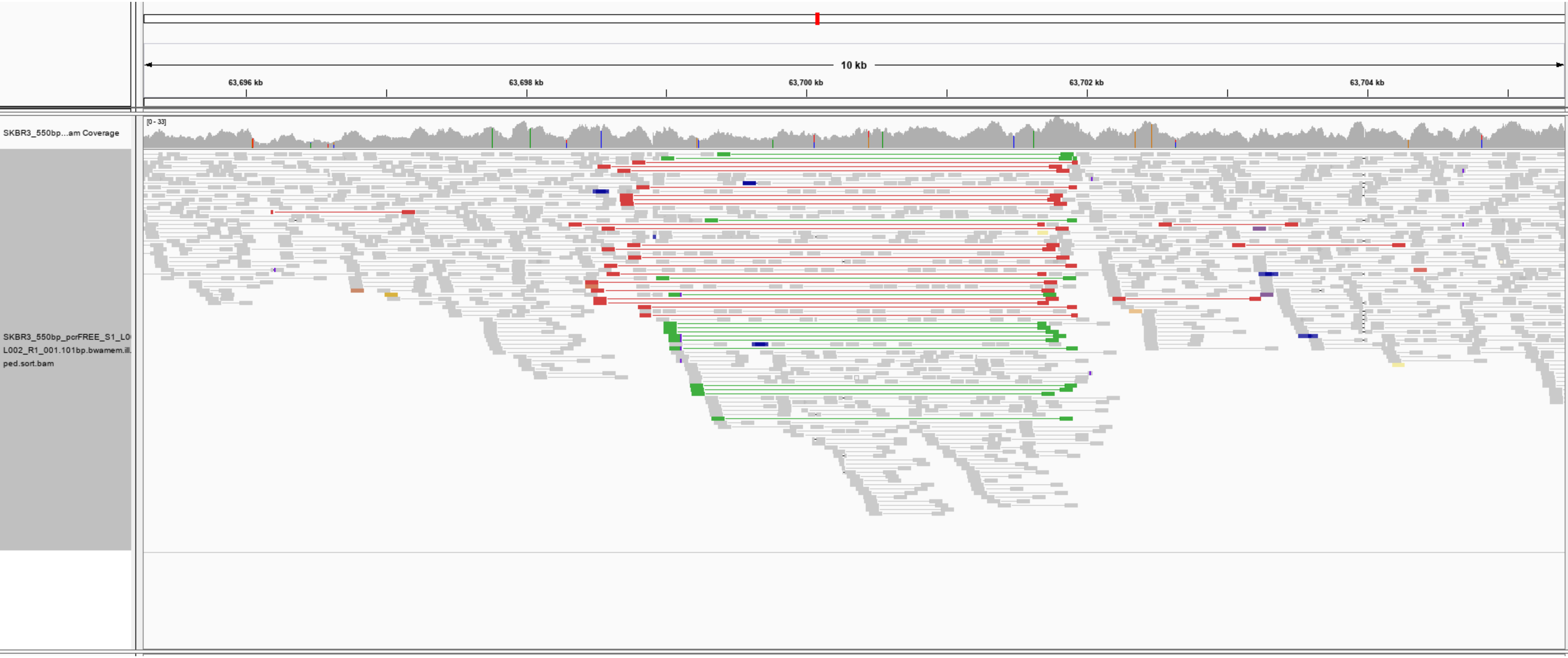

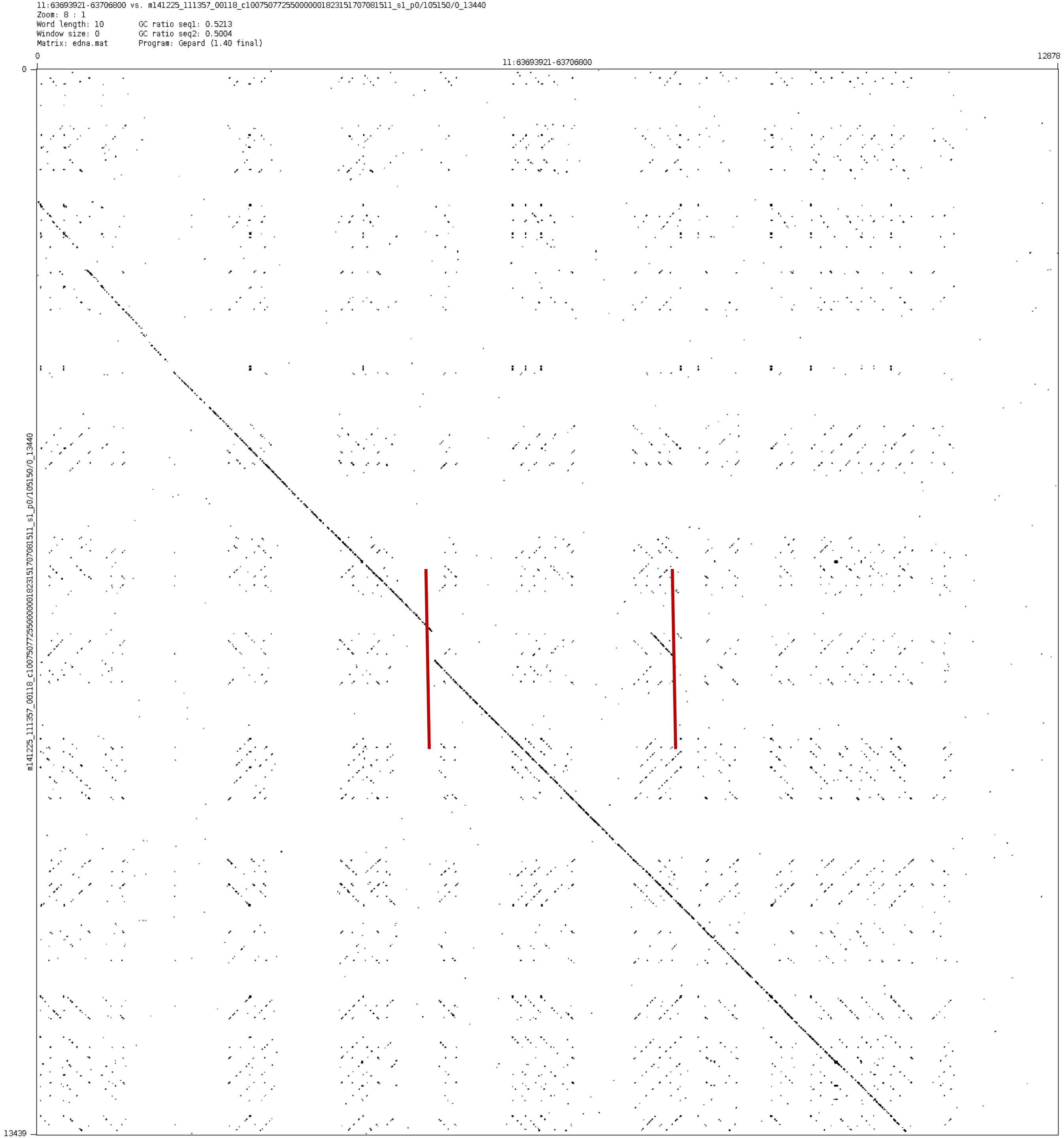
11:63693921-63706800 vs. m141225_111357_00118_c100750772550000001823151707081511_s1_p0/105150/0_13440
Zoom: 8 : 1
Word length: 10
Window size: 0
Matrix: edna.mat
GC ratio seq1: 0.5213
GC ratio seq2: 0.5004
Program: Gepard (1.40 final)
0
11:63693921-63706800
12878
0
m141225_111357_00118_c100750772550000001823151707081511_s1_p0/105150/0_13440
13439

chr11:78039089-78042394

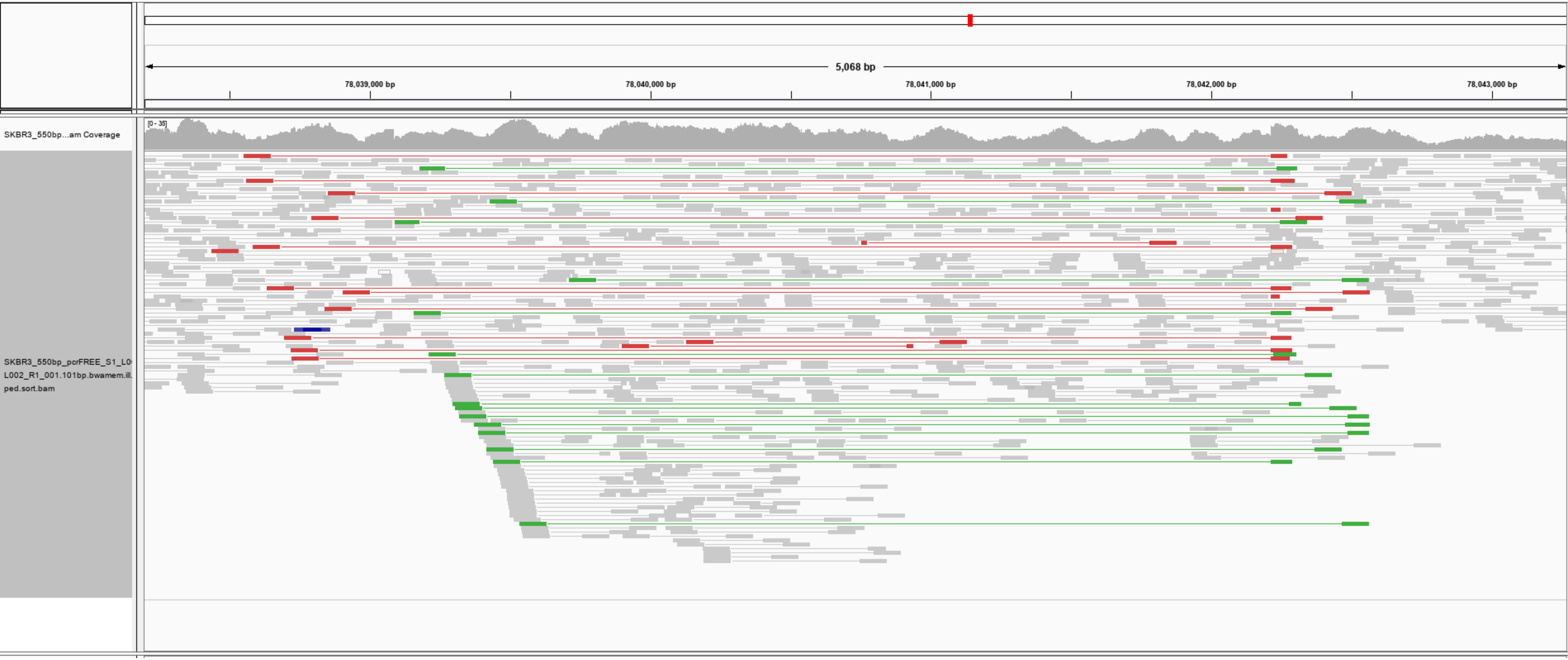

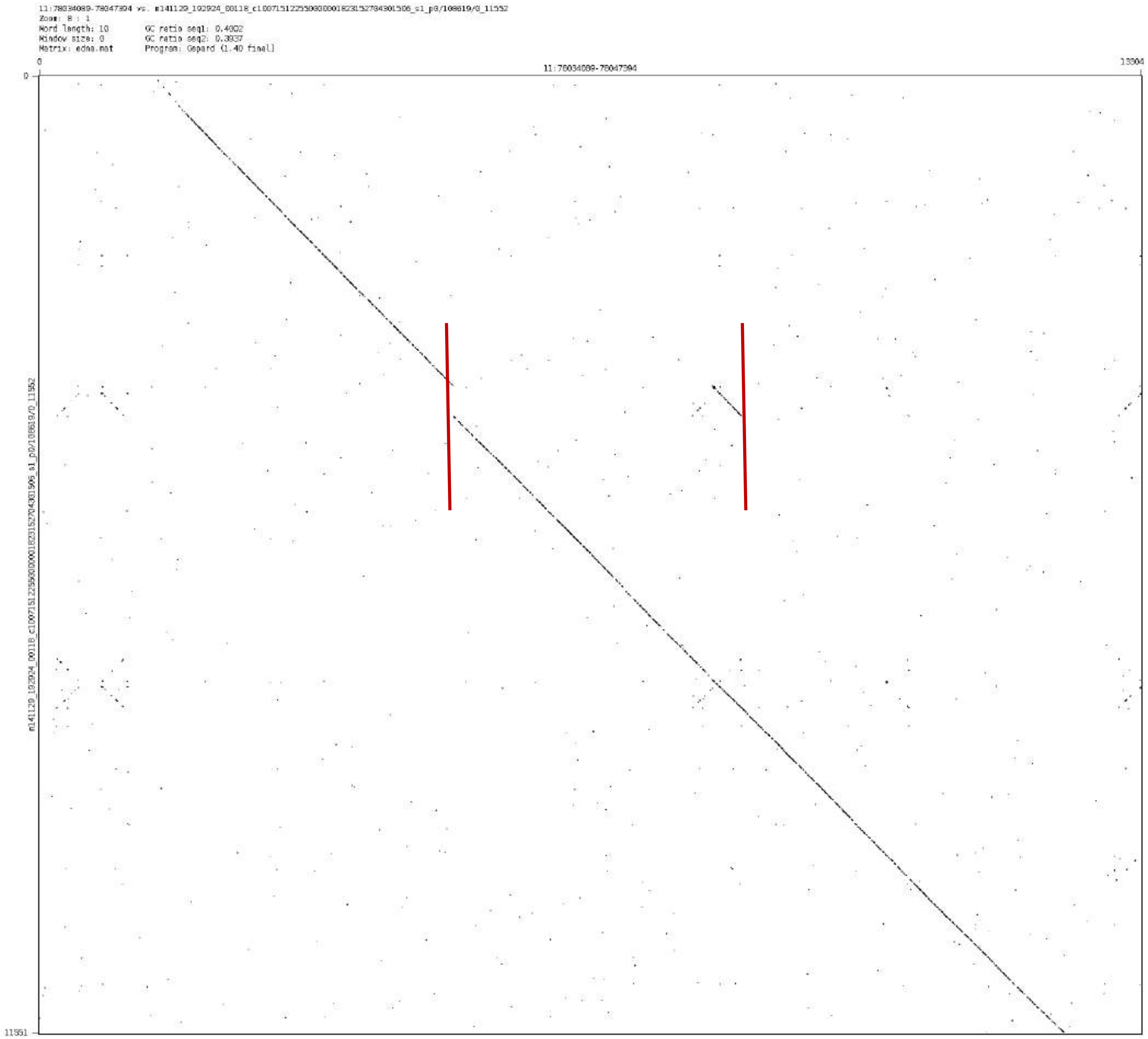

Word length: 10
Window size: 0
Matrix: edna.mat
GC ratio seq1: 0.4002
GC ratio seq2: 0.3937
Program: Gepard (1.40 final)
0
13304
11551

chr14:40113933-40114921

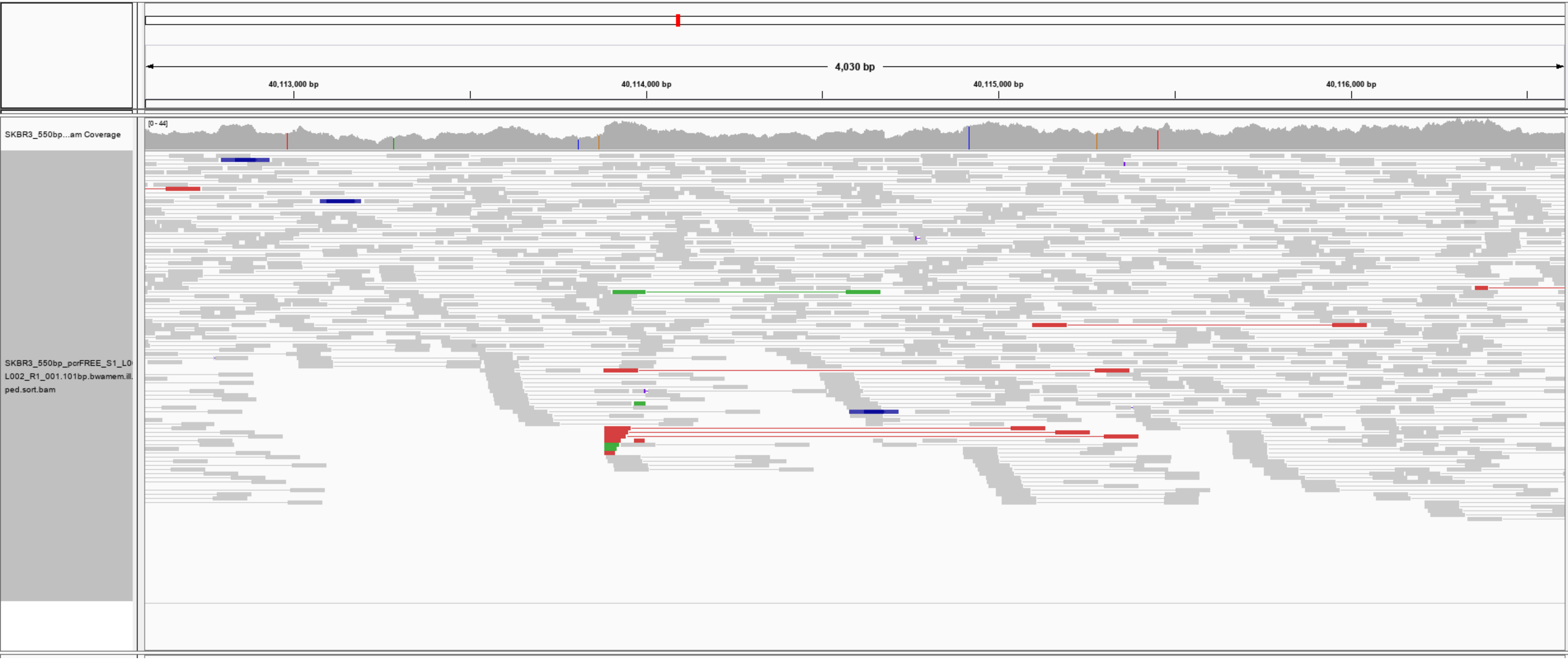

# 14_40113933_40114921.27

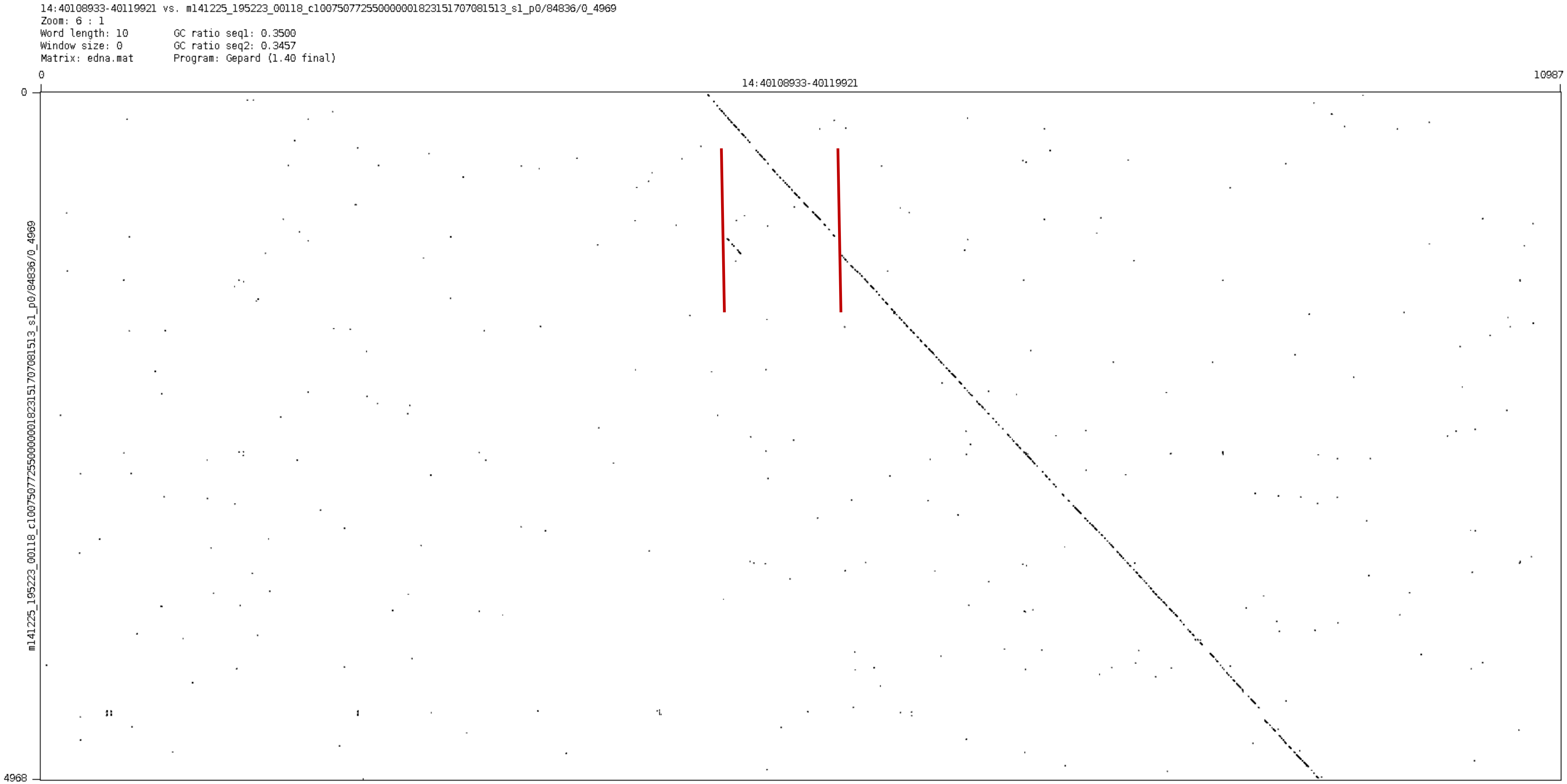

### chr18:11509472-11511497

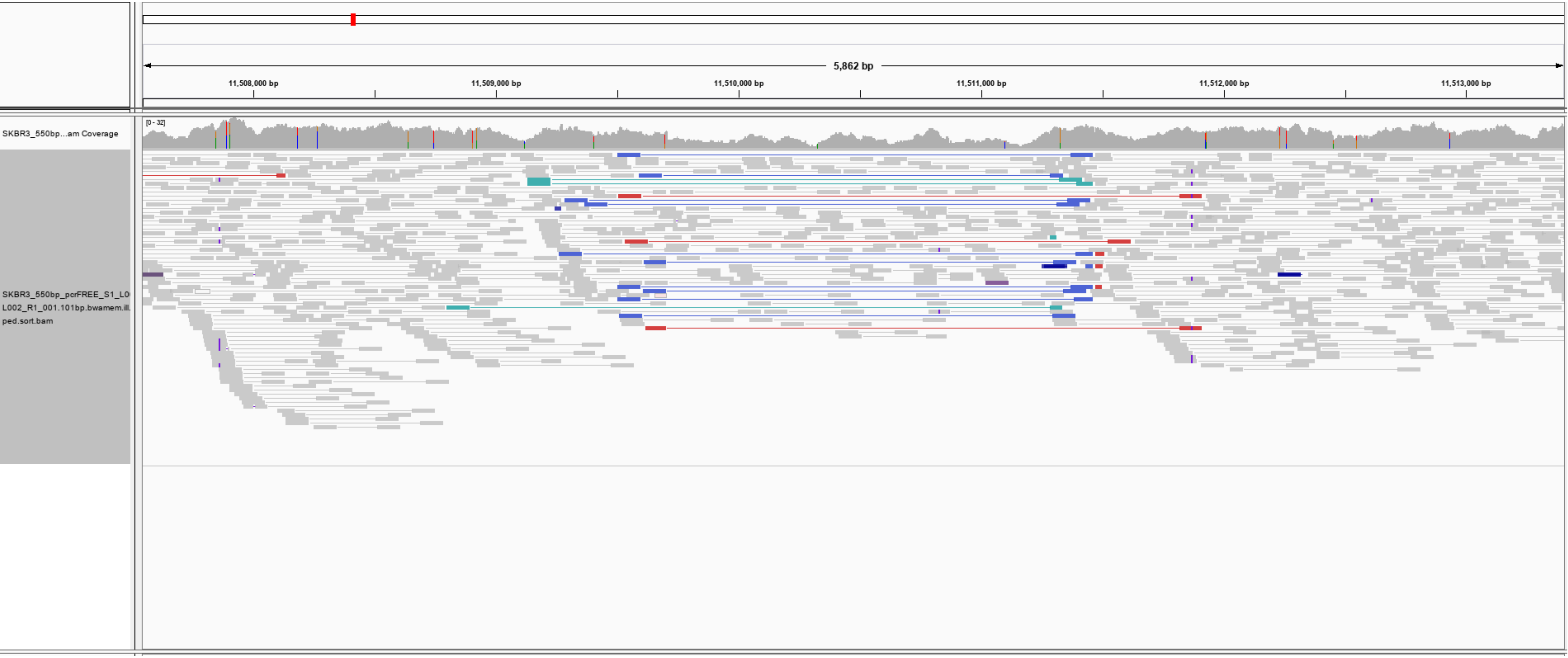

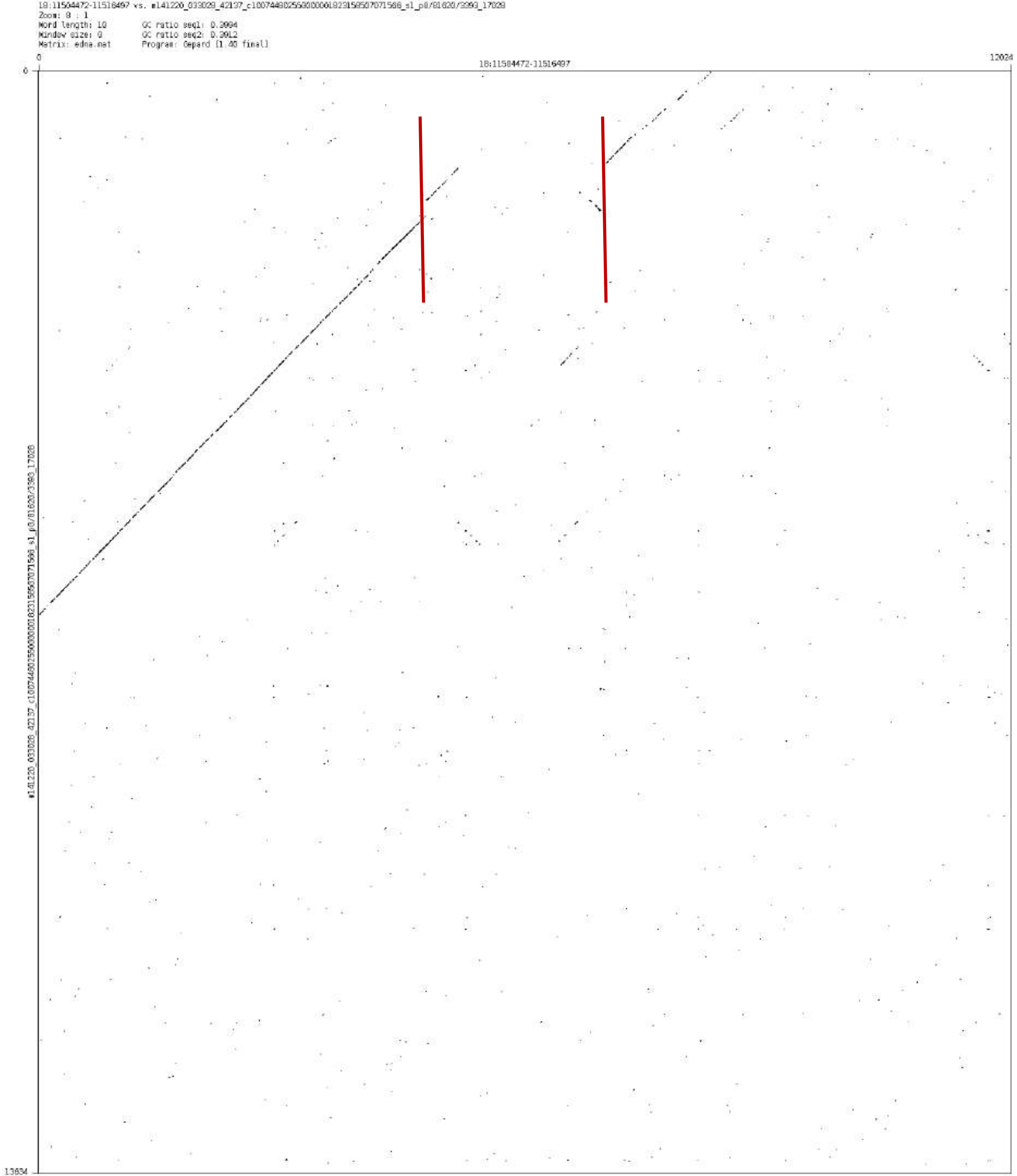

Zoom: 8 : 1
Word length: 10
Window size: 0
Matrix: edna.mat
GC ratio seq1: 0.3994
GC ratio seq2: 0.3912
Program: Gepard [1.40 final]
0
18:11504472-11516497
12024
0
13634

### chr18:69711908-69712979

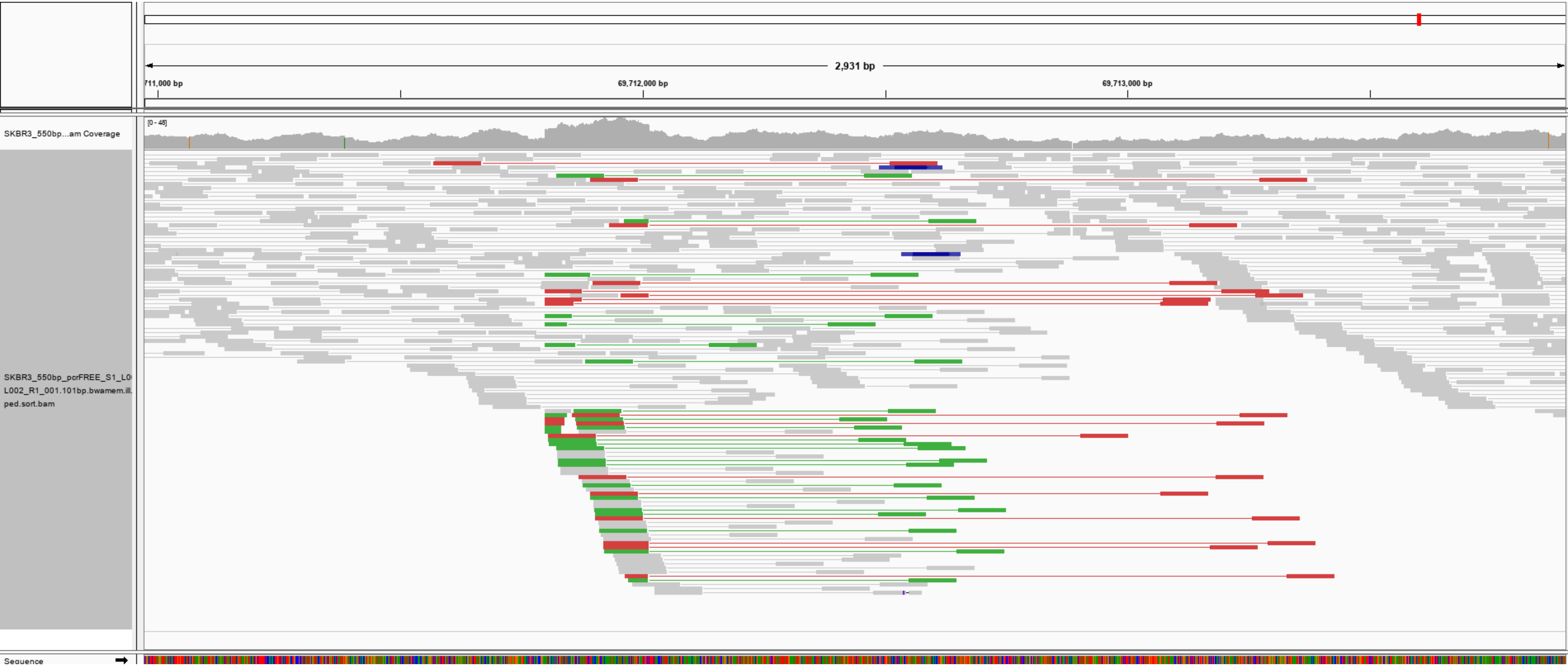

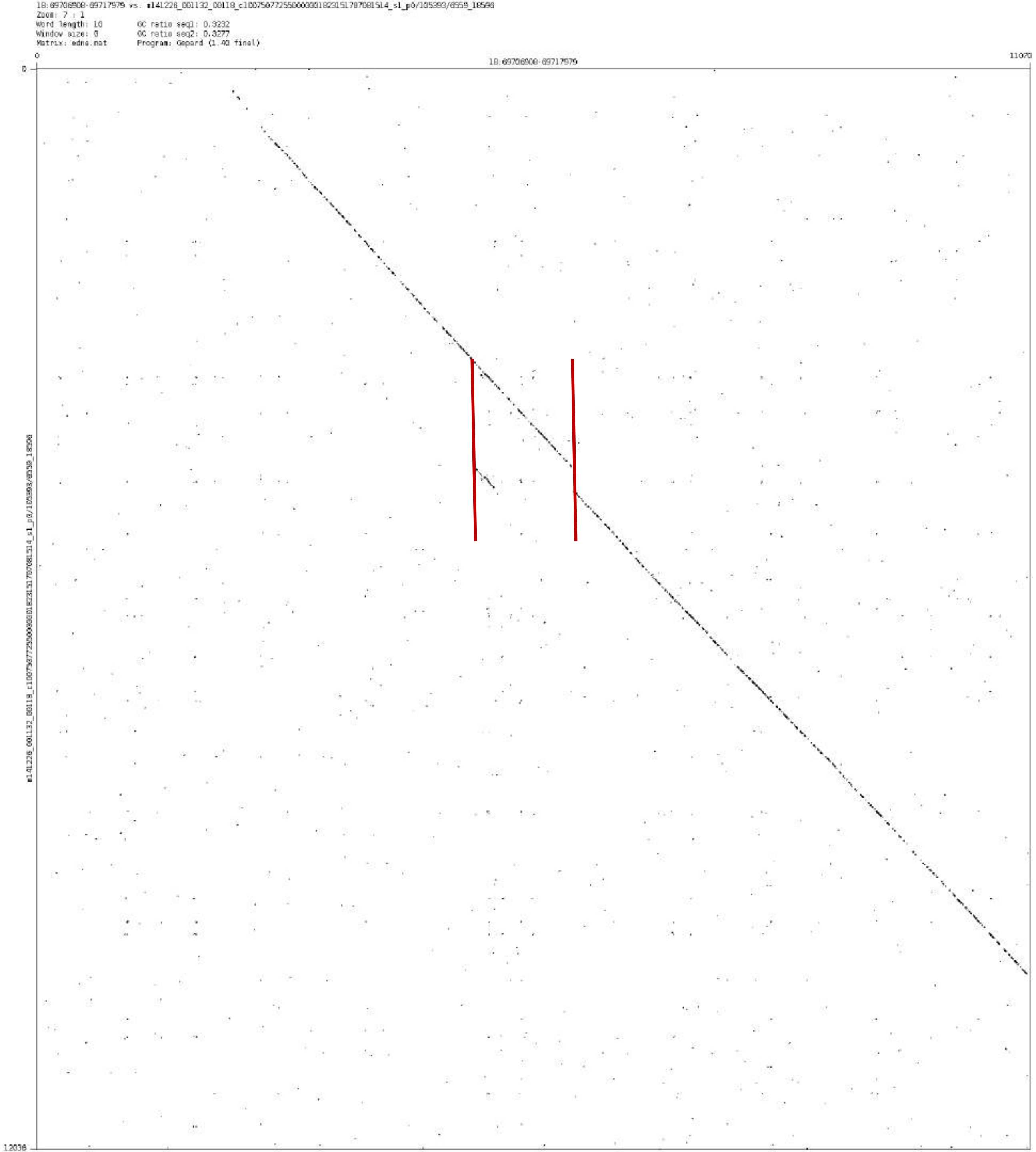
18:69706908-69717979 vs. m141226_001132_00118_c100750772550000001823151707081514_s1_p0/105393/8559_18596
Zoom: 7 : 1
Word length: 10
Window size: 0
Matrix: edna.mat
GC ratio seq1: 0.3232
GC ratio seq2: 0.3277
Program: Gepard (1.40 final)
0
18:69706908-69717979
11070
0
12036

chr21:22783456-22784516

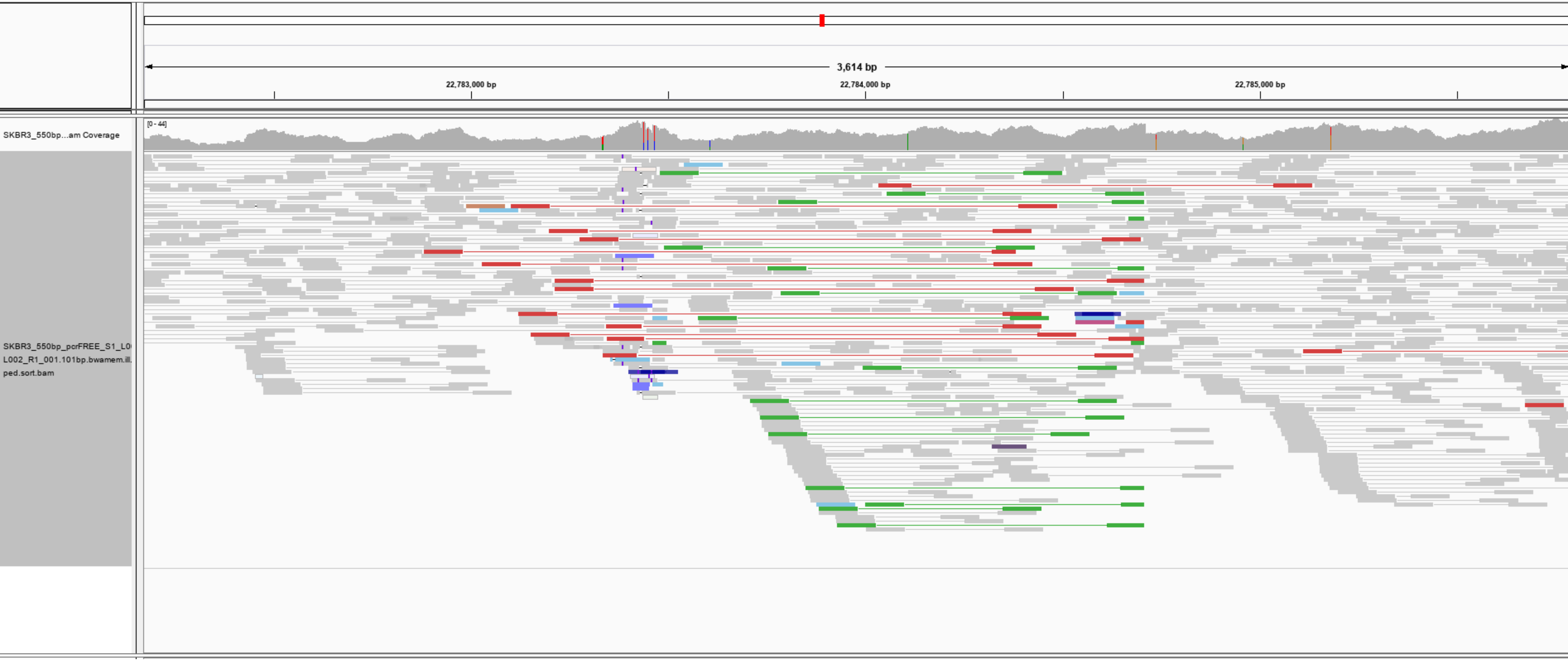

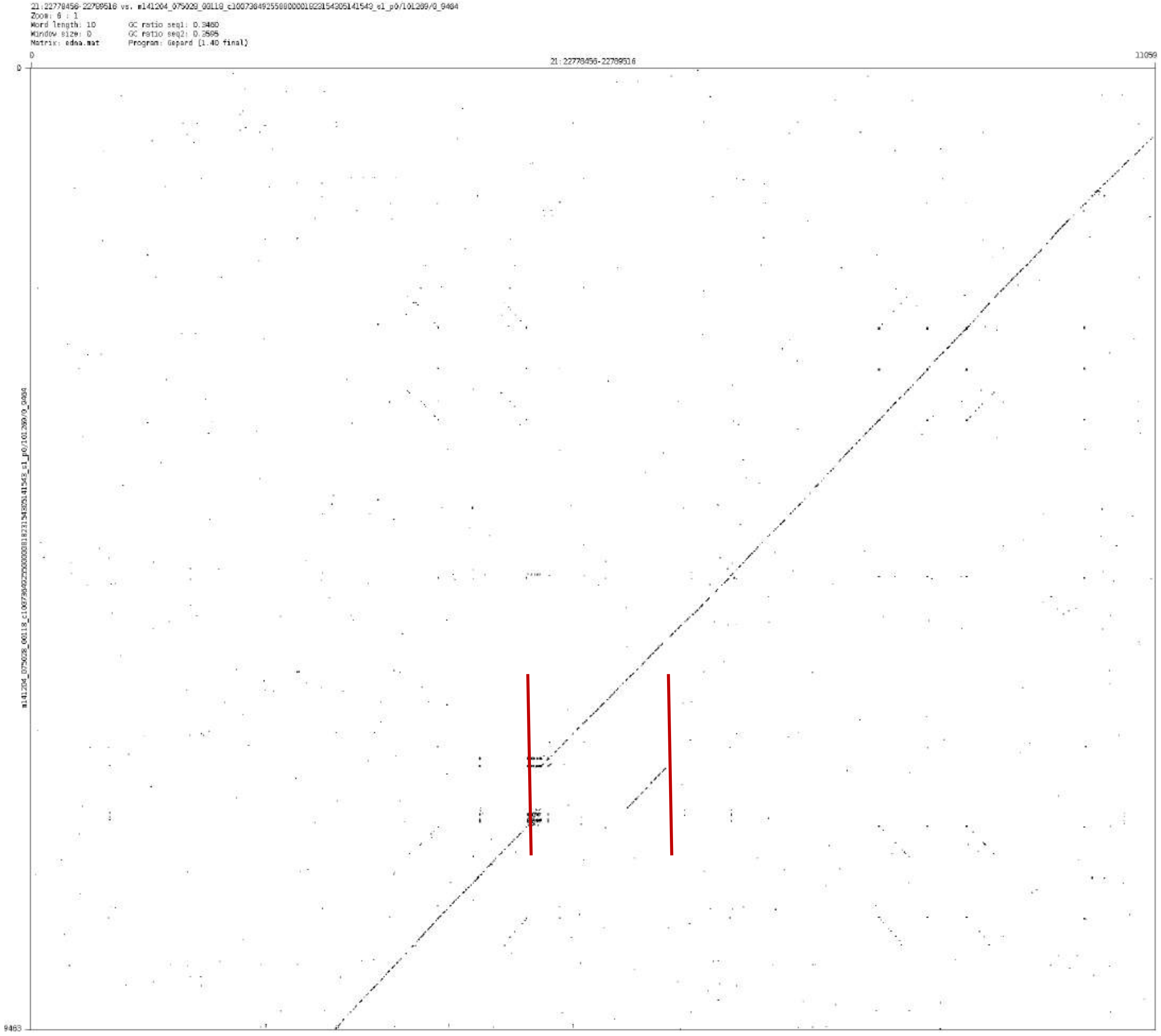
21:22778456-22789516 vs. m141204_075029_00118_c100738492550000001823154305141543_s1_p0/101289/0_9484
Zoom: 6 : 1
Word length: 10
Window size: 0
Matrix: edna.mat
GC ratio seq1: 0.3480
GC ratio seq2: 0.3585
Program: Gepard (1.40 final)
0
21:22778456-22789516
11059
0
9483
