## Supplementary File 2 for "Mako: a graph-based pattern growth approach to detect complex structural variants"

| Classifications | | | Count | Total |
|---|---|---|---|---|
| CSV | Distinct | | 84 | 233 |
| | Messy | Case A | 56 | |
| | | Case B | 80 | |
| | | Case C | 13 | |
| Simple SV | Case A | | 4 | 19 |
| | Case B | | 15 | |
| Inconclusive | | | | 188 |

### Distinct complex events

➢ chr1_14109802_14112516

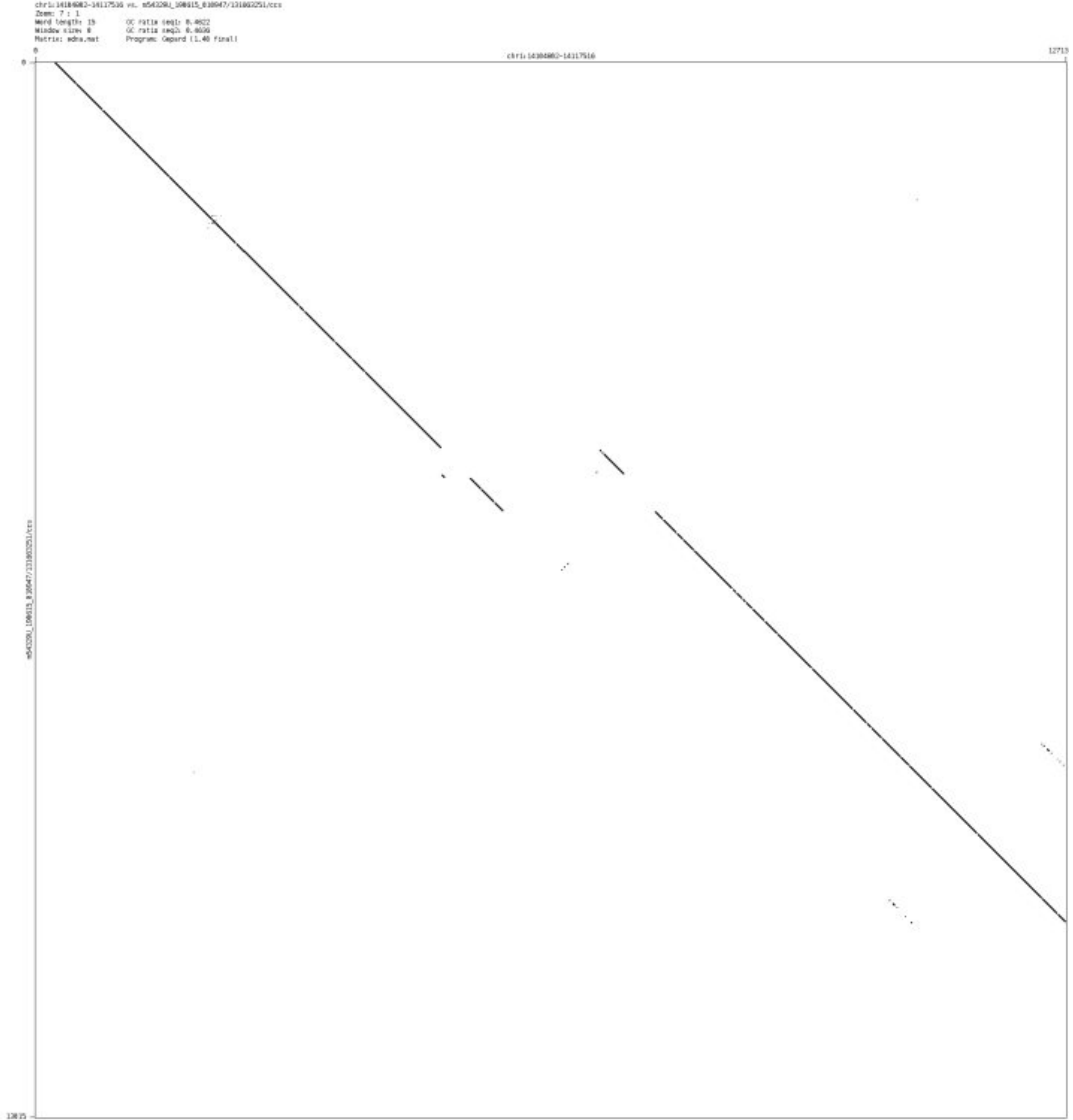

- chr1_16050024_16059805

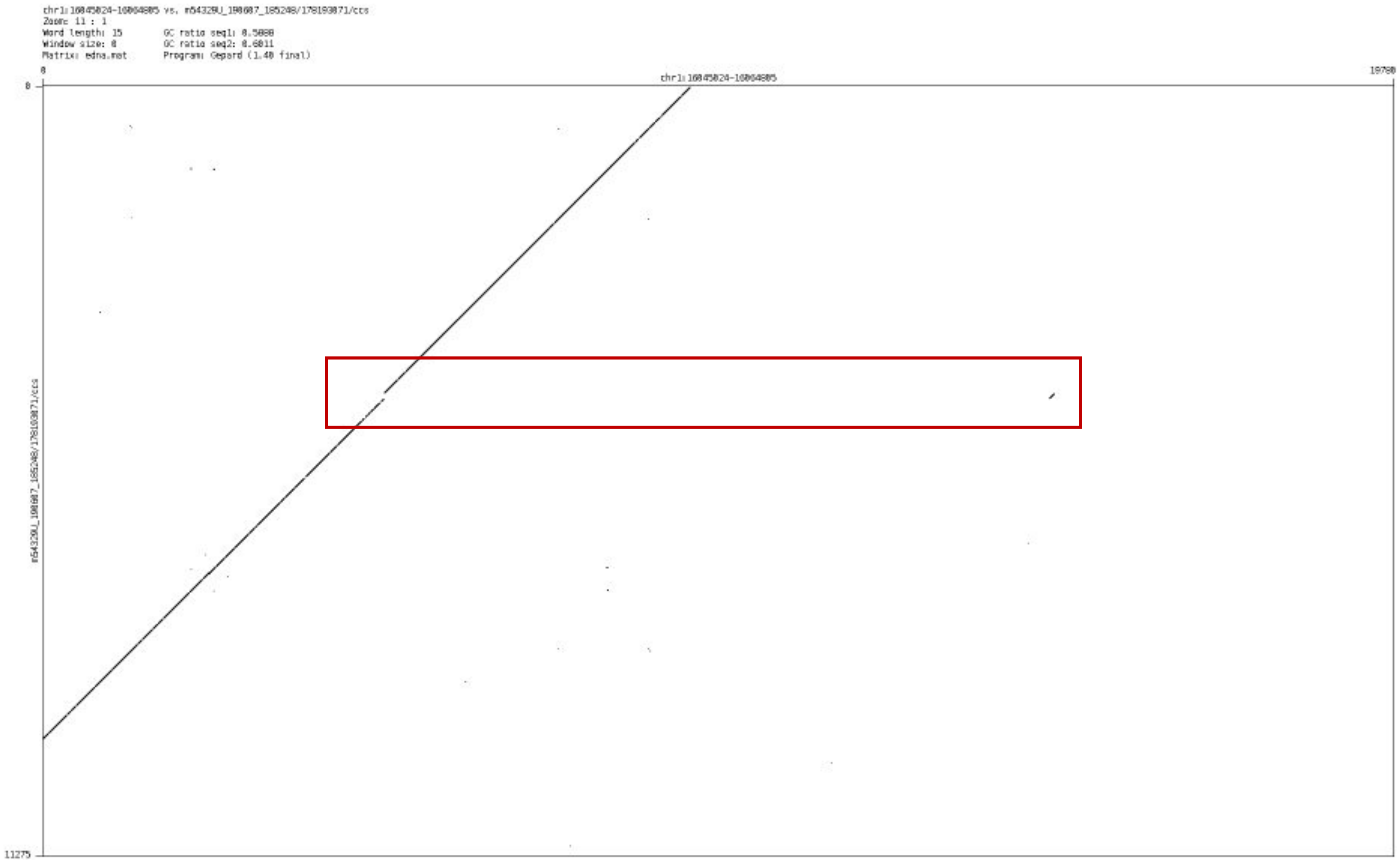

➢ chr1_36267369_36269040

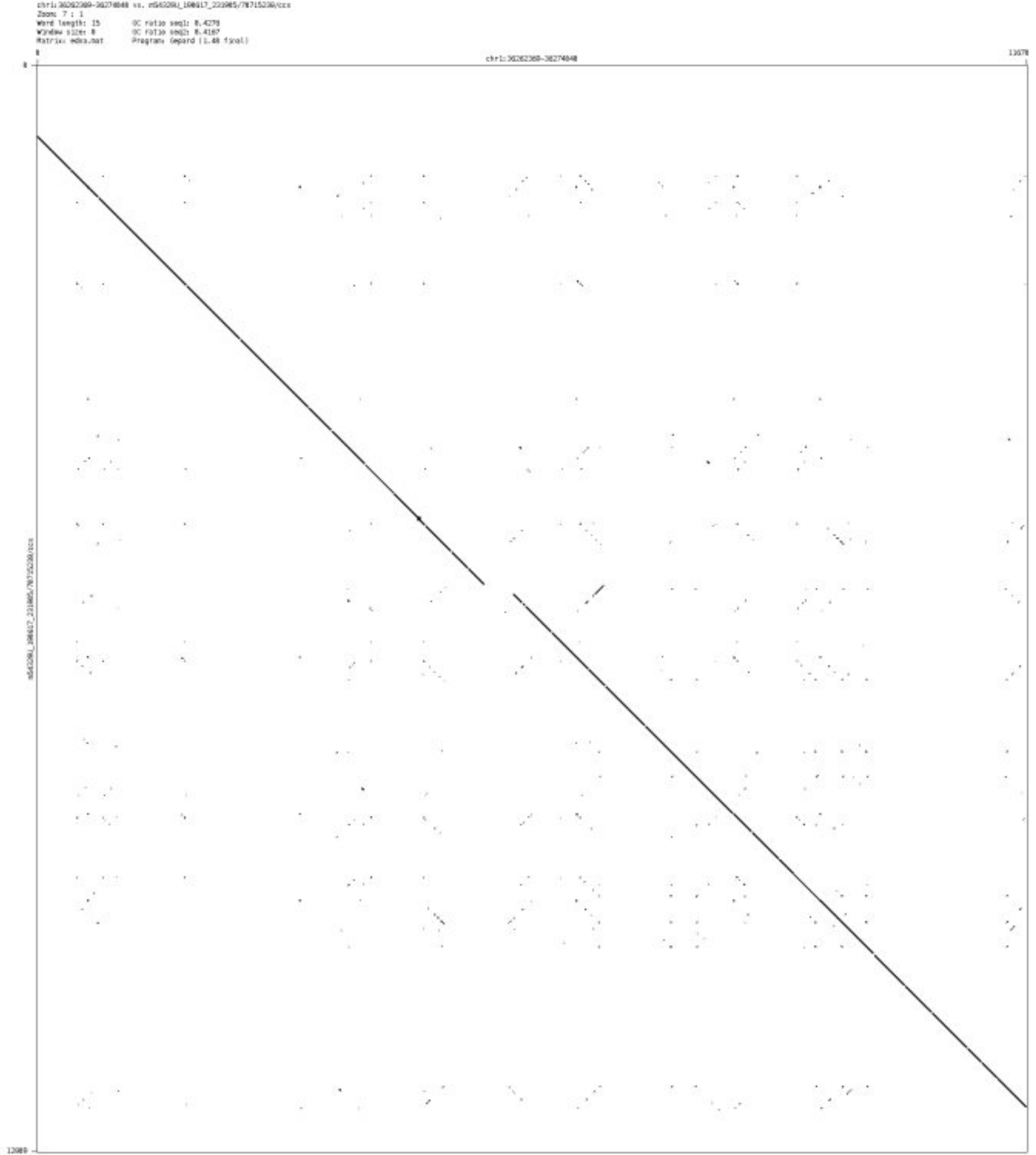

- chr1_43593650_43594474

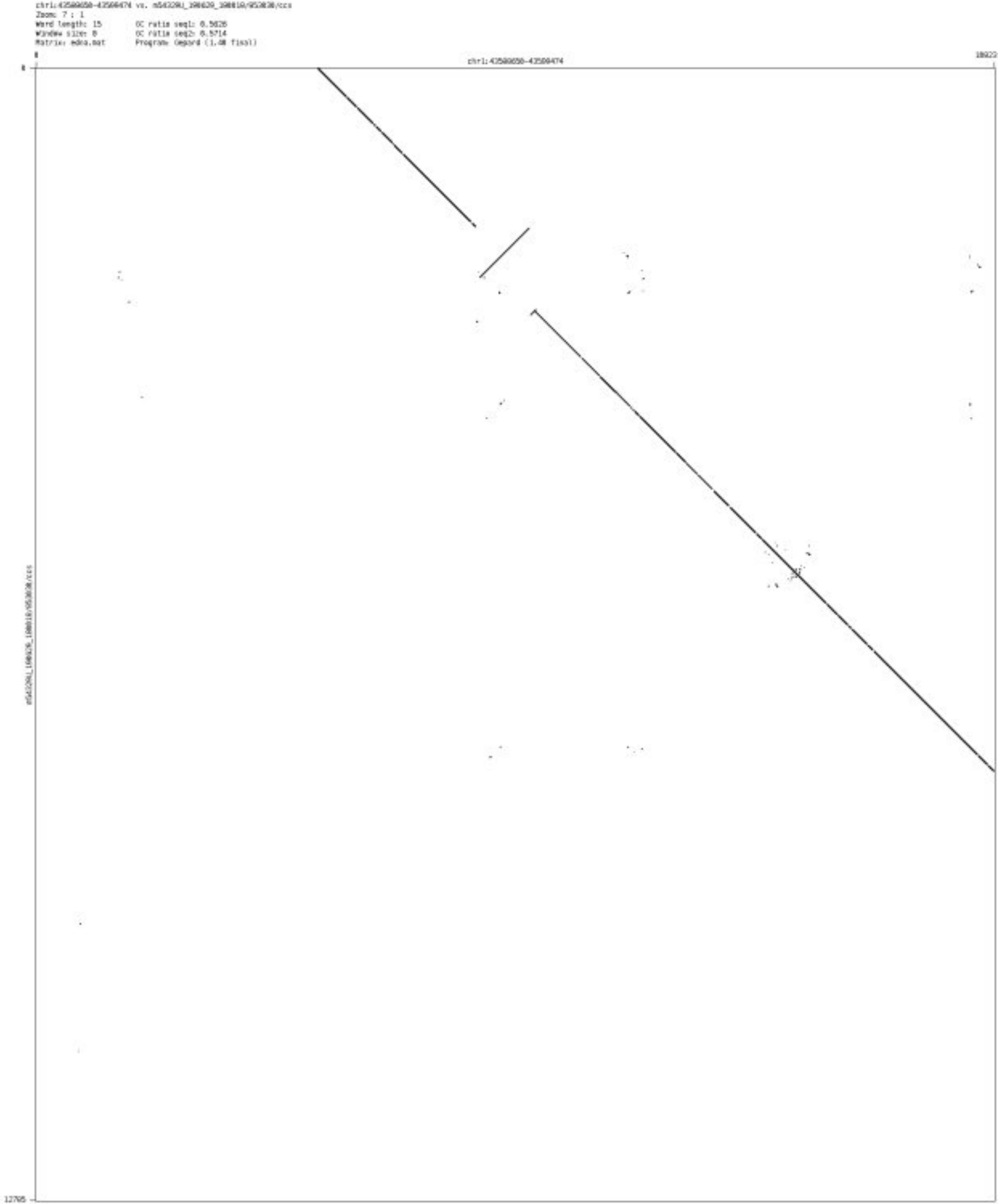

➢ chr1_81194398_81195874

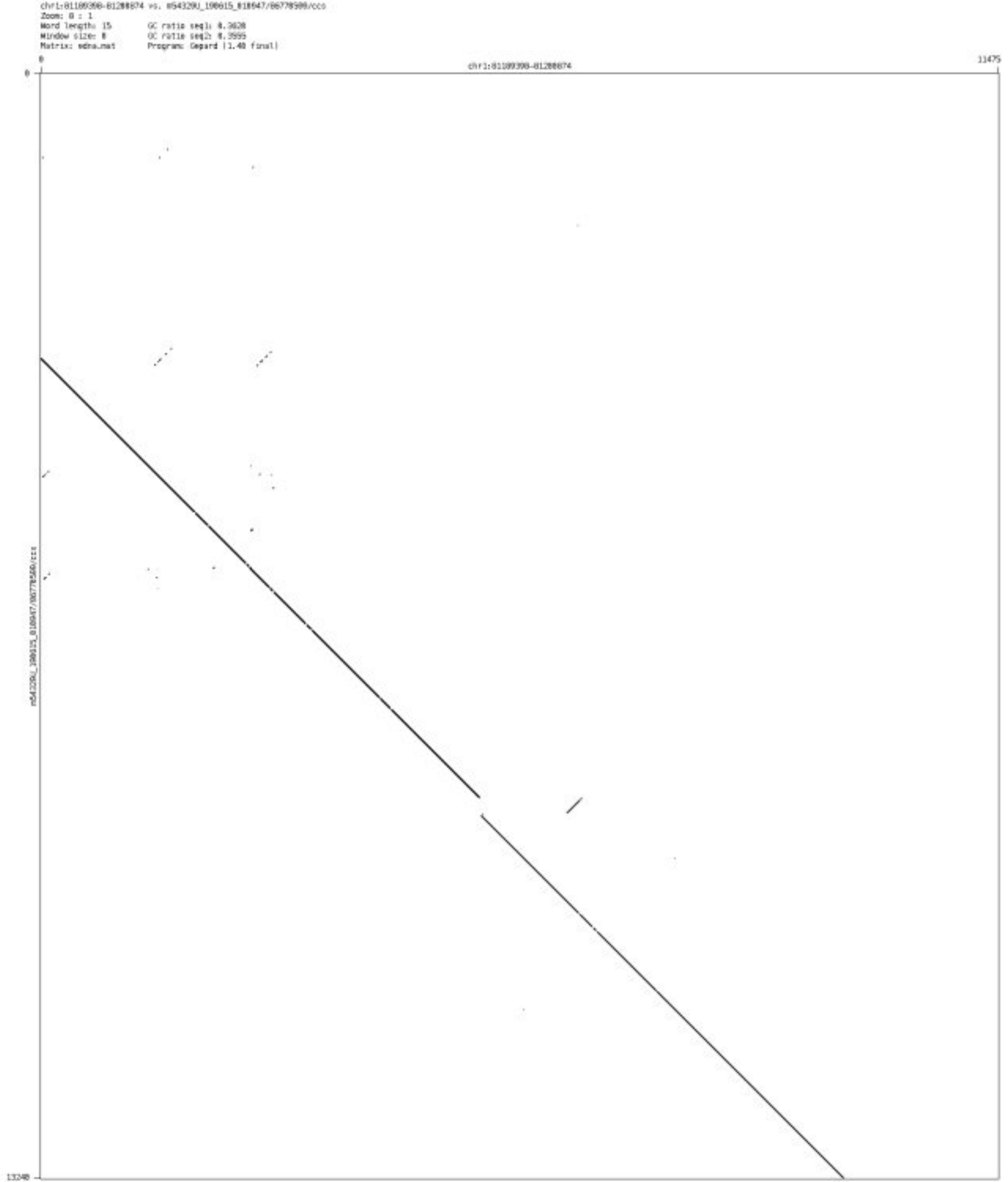

- chr1_175231051_175232802

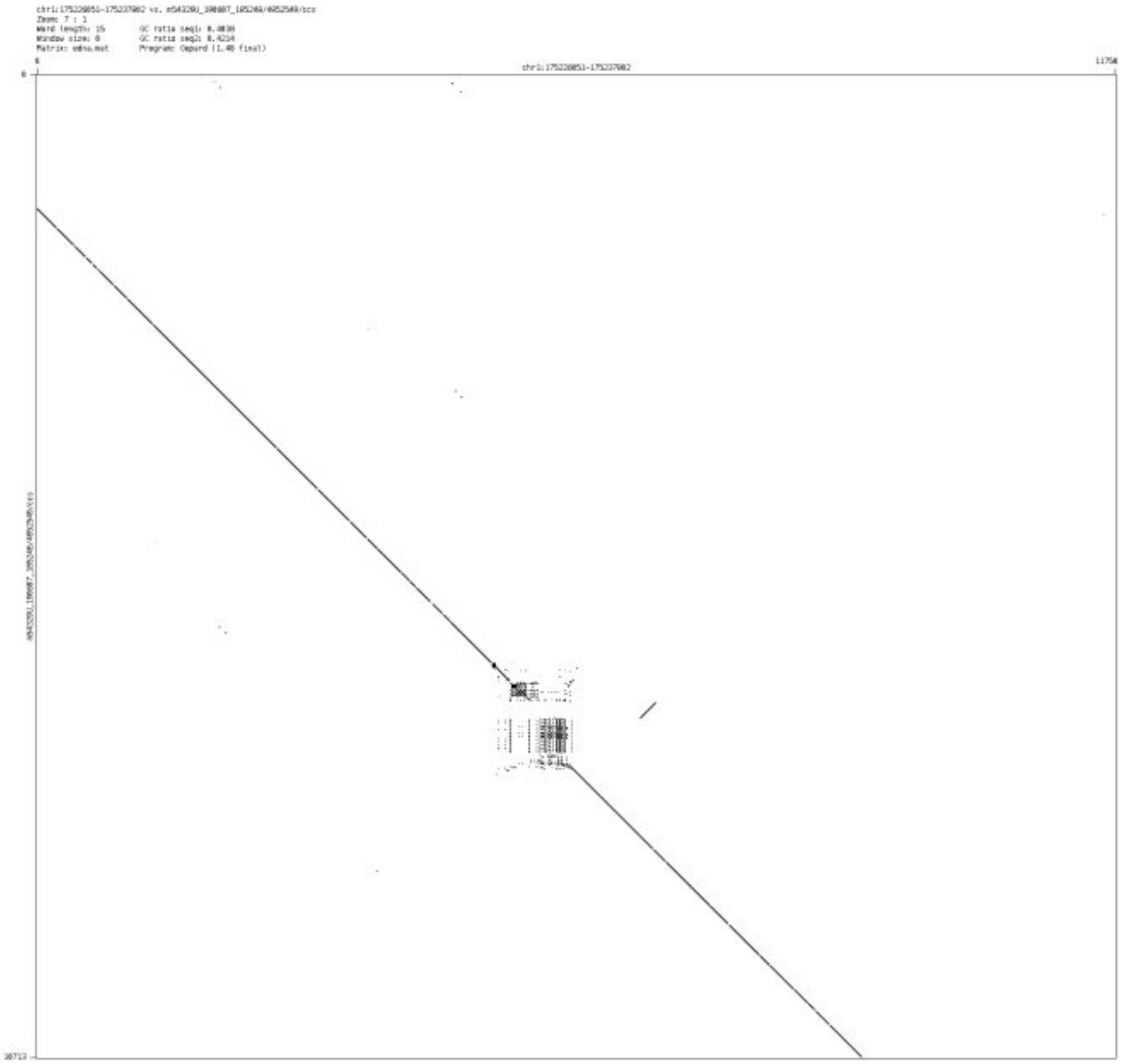

- chr1_187495434_187497604

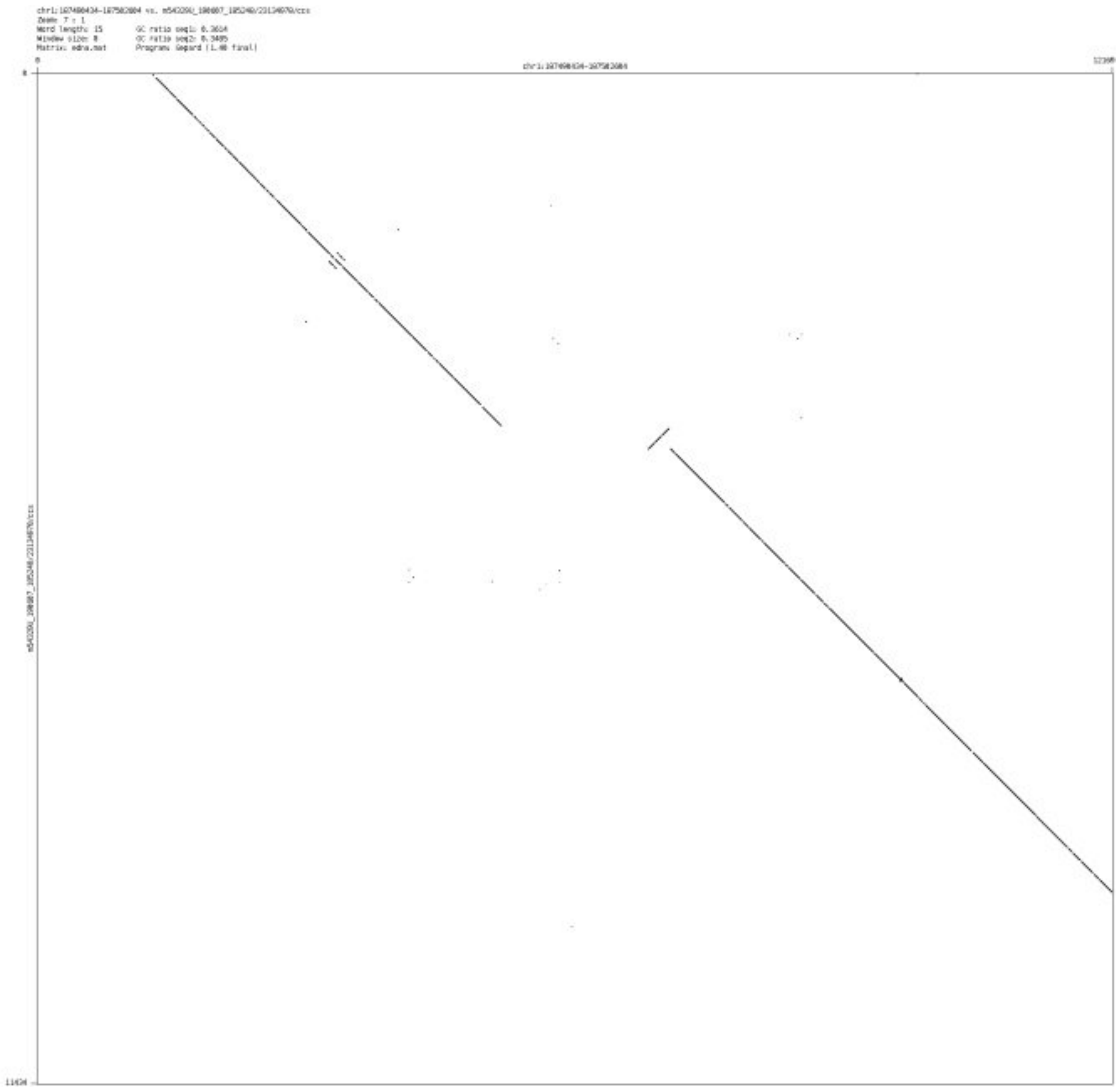

- chr1_207118600_207120000

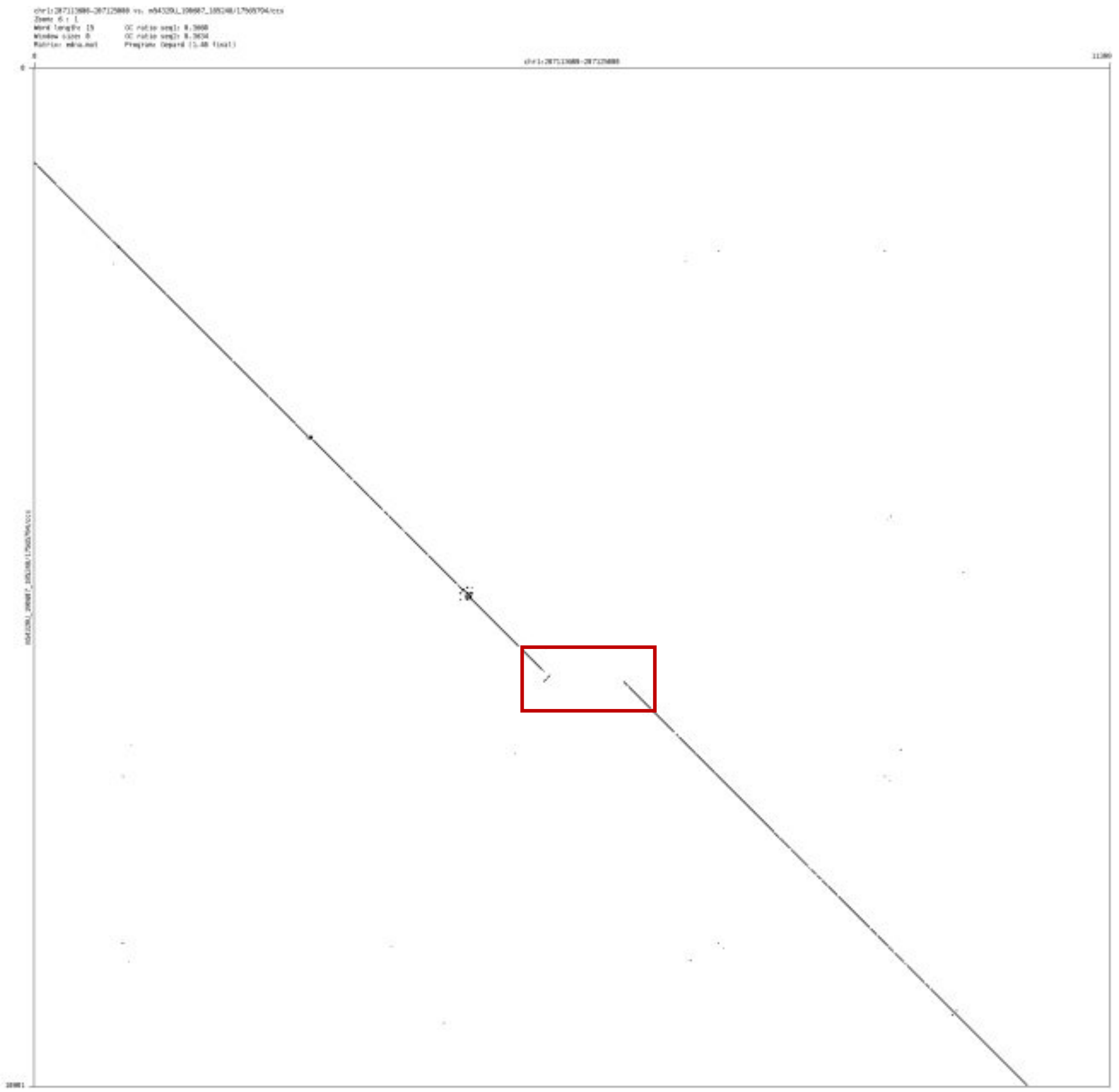

- chr1_246244294_246247399

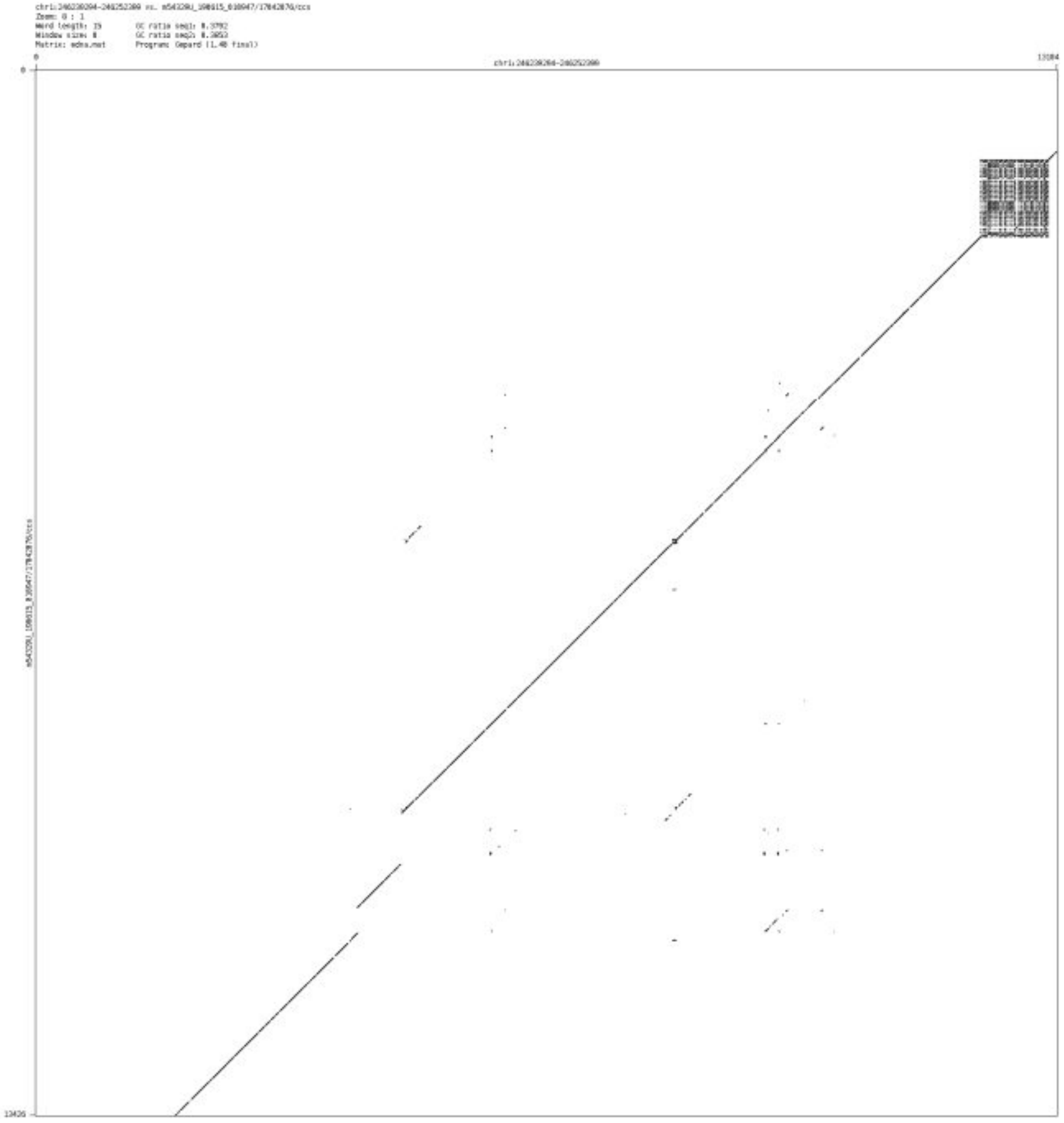

➢ chr2_16224687_16226666

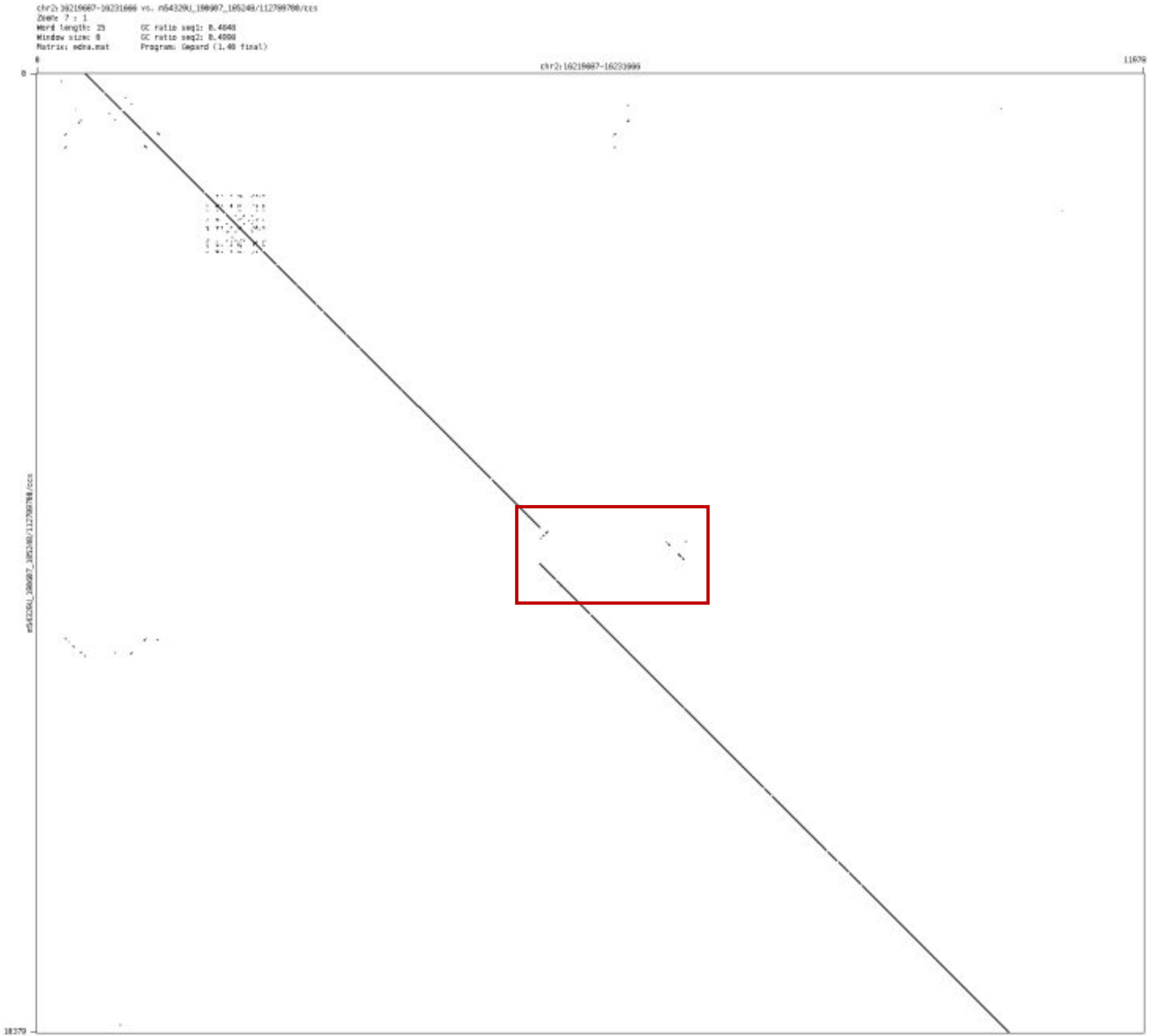

- chr2_61473303_61476318

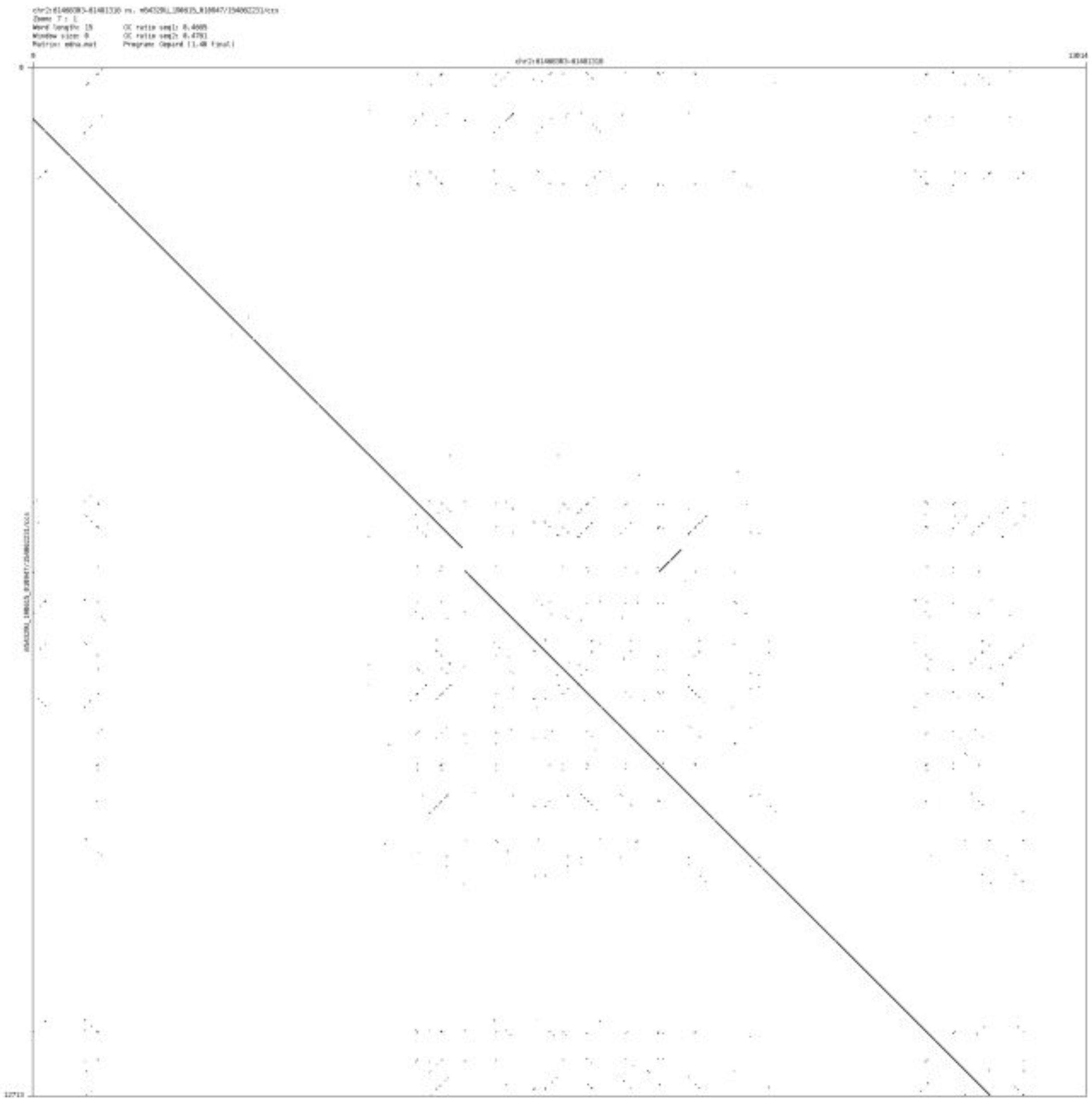

➢ chr2_95544970_95546071

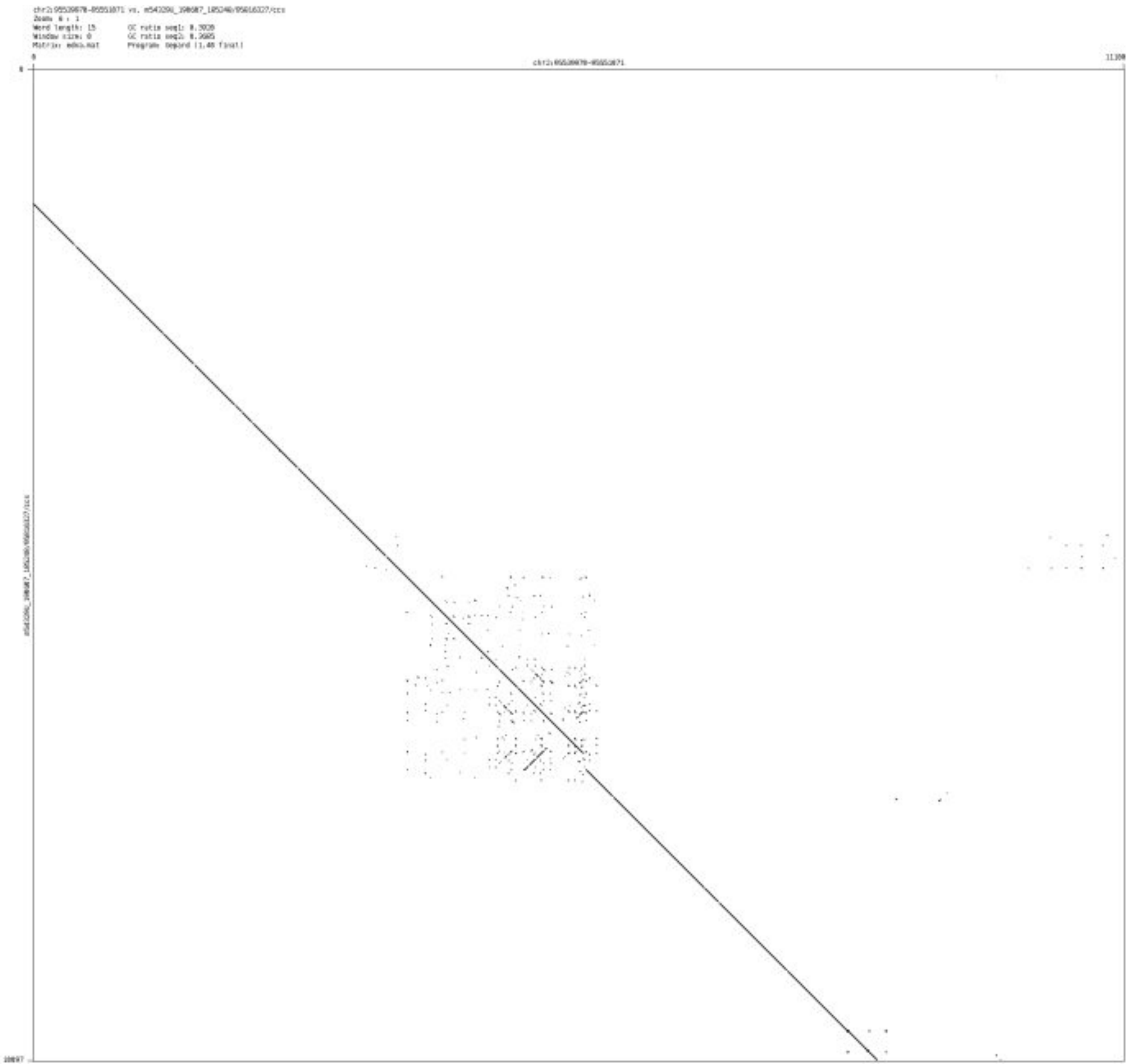

### ➢ chr2_116216764_116224735

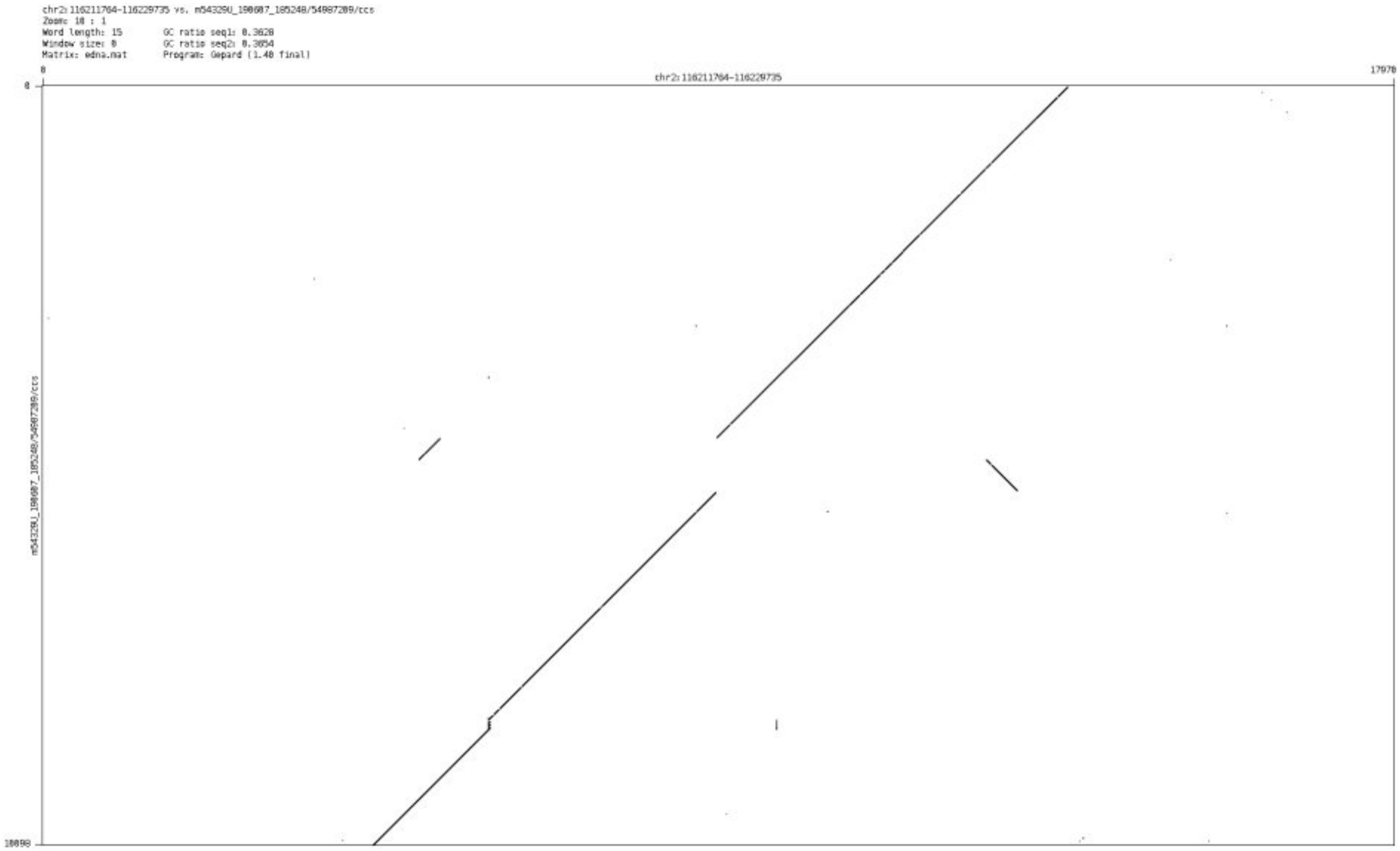

- chr2_119659504_119661322

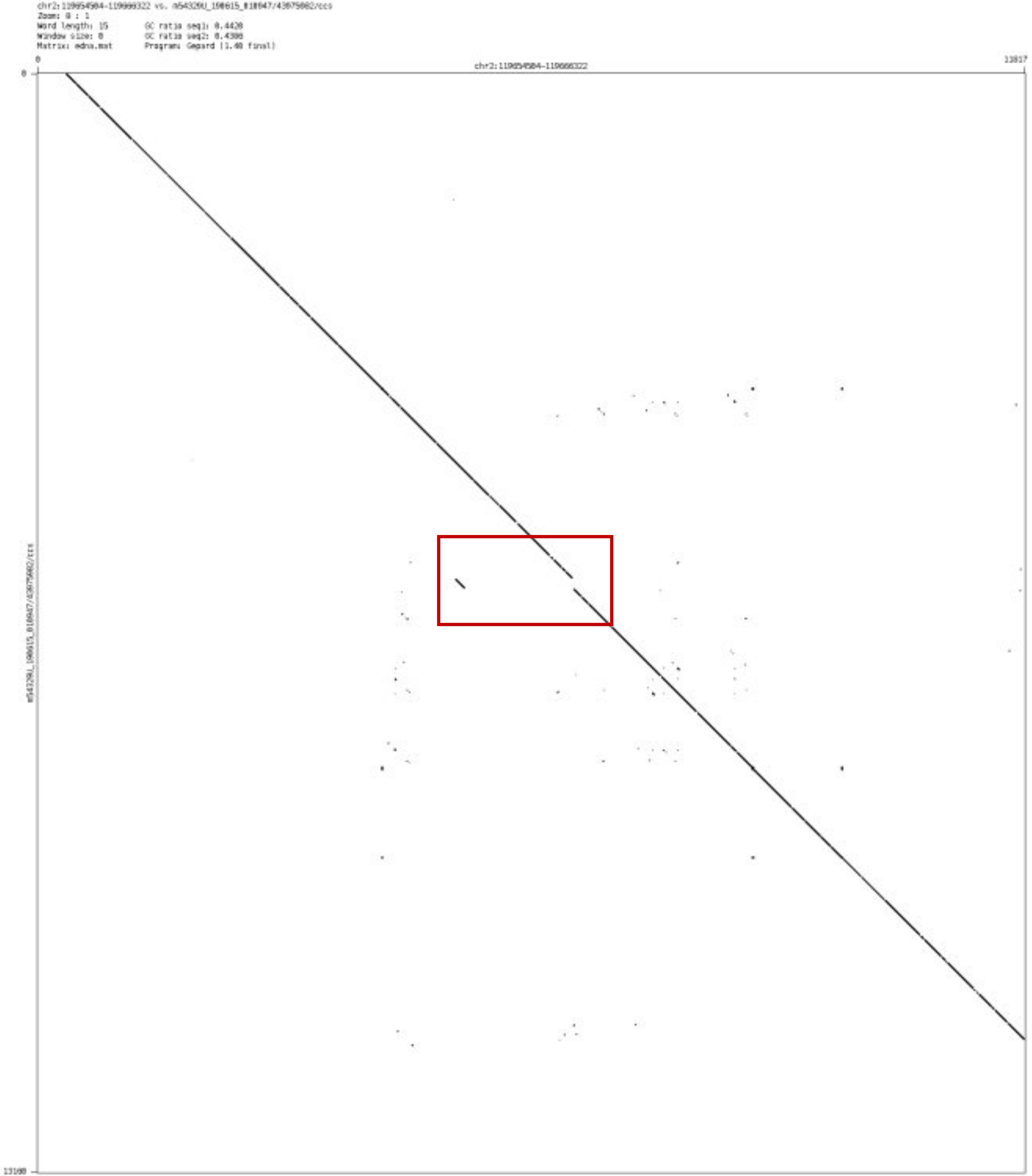

➢ chr2_122612259_122613019

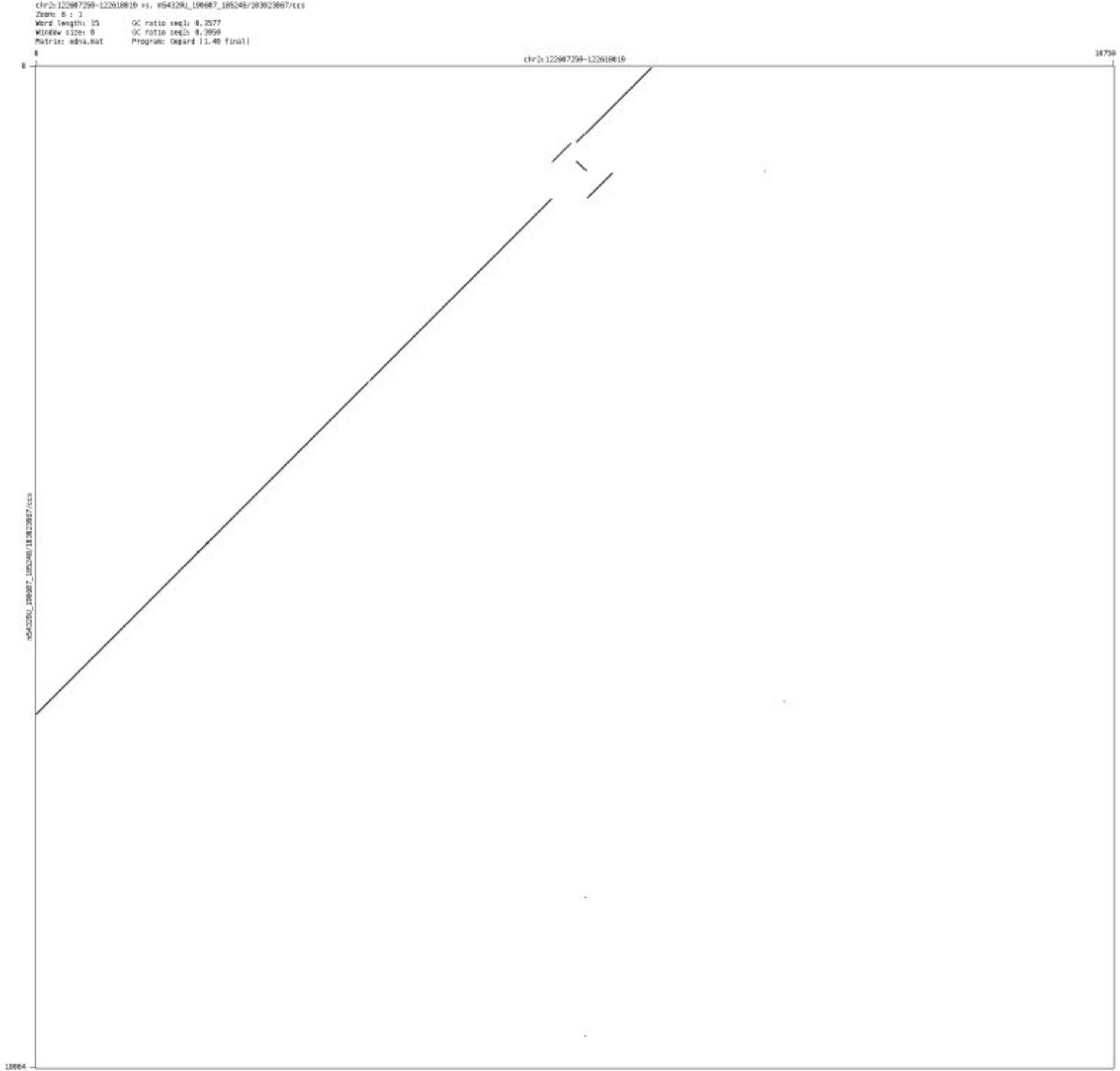

- chr2_124293968_124295664

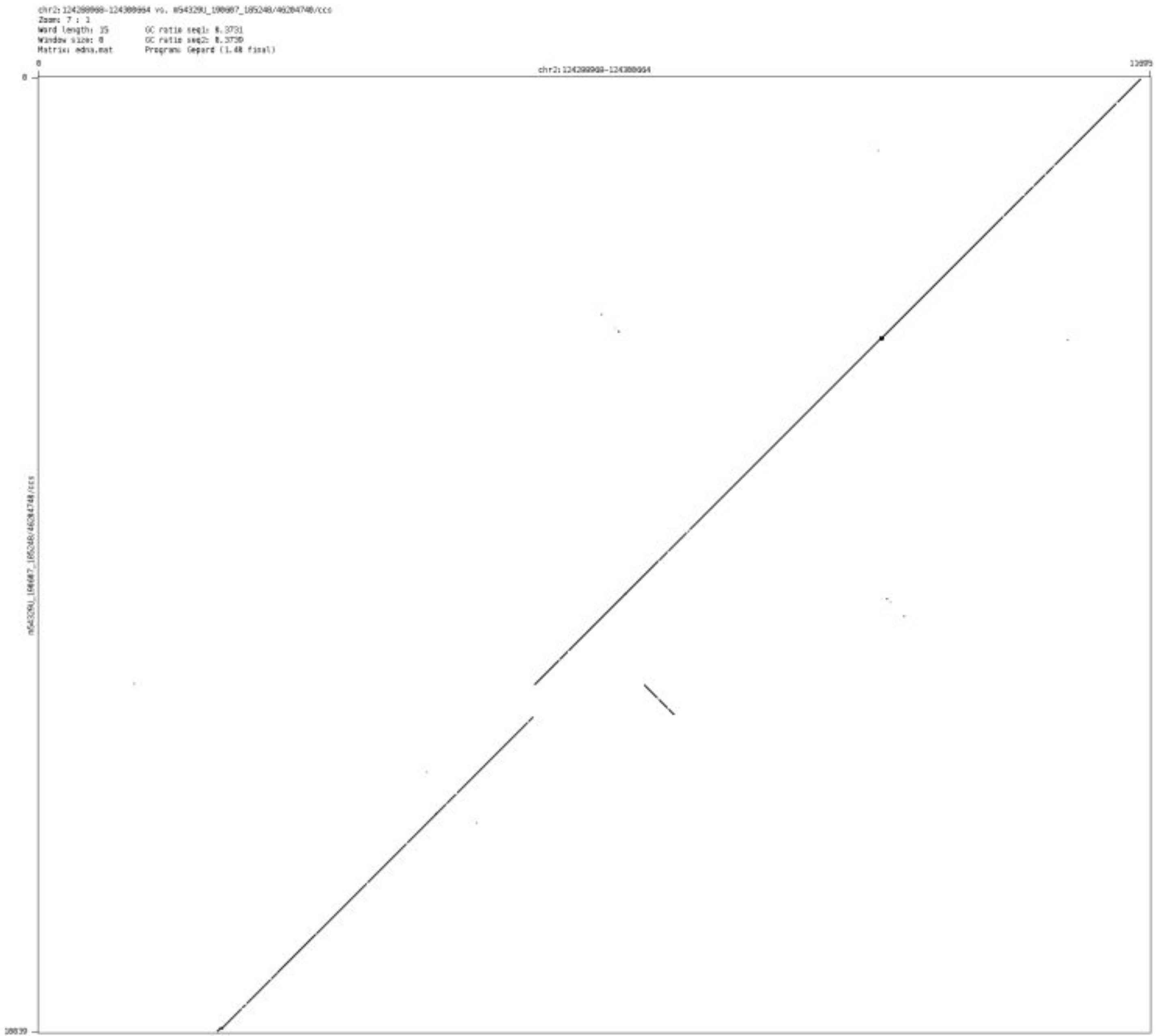

- chr2_125008685_125010825

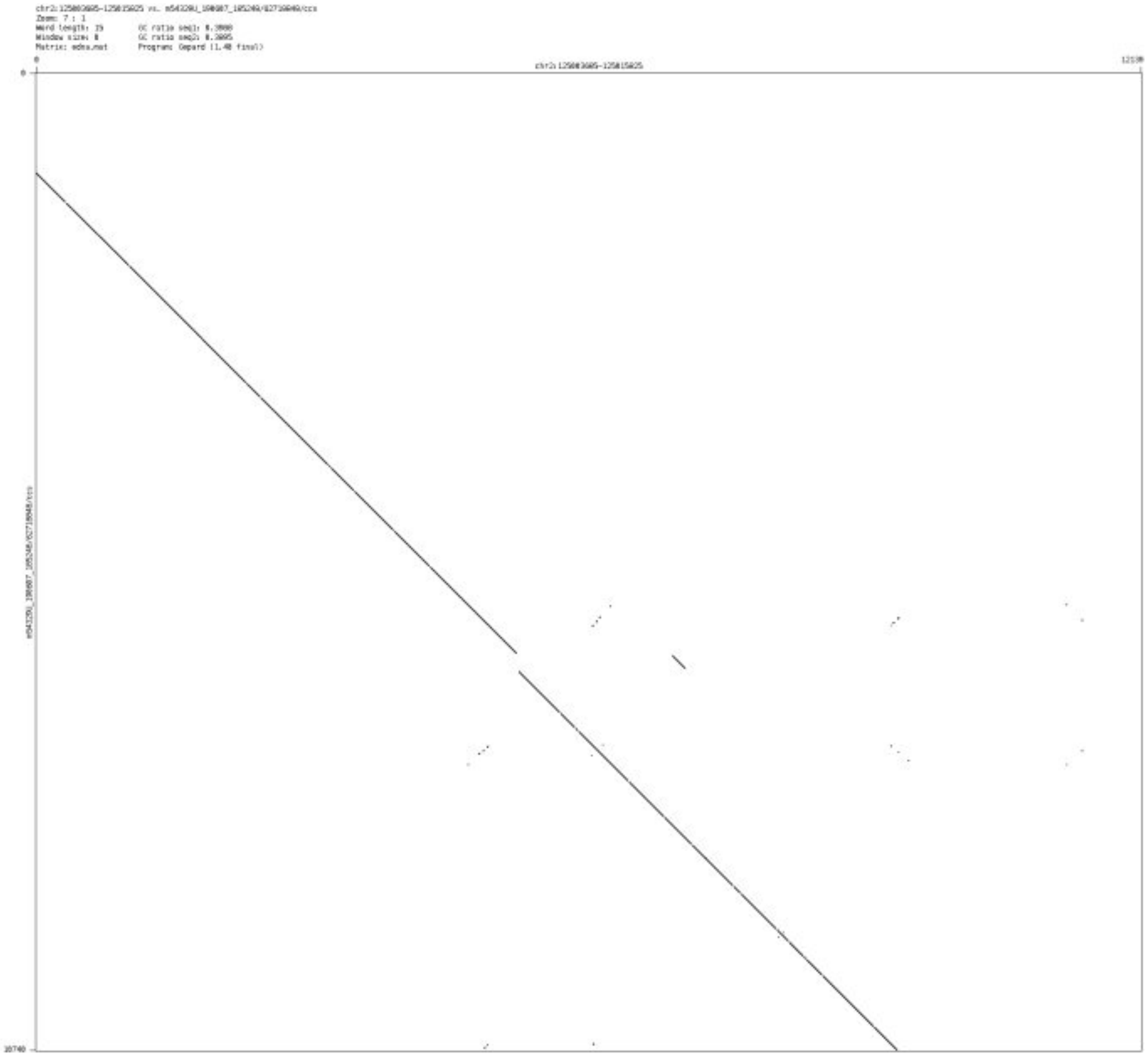

➢ chr2_161074754_161075236

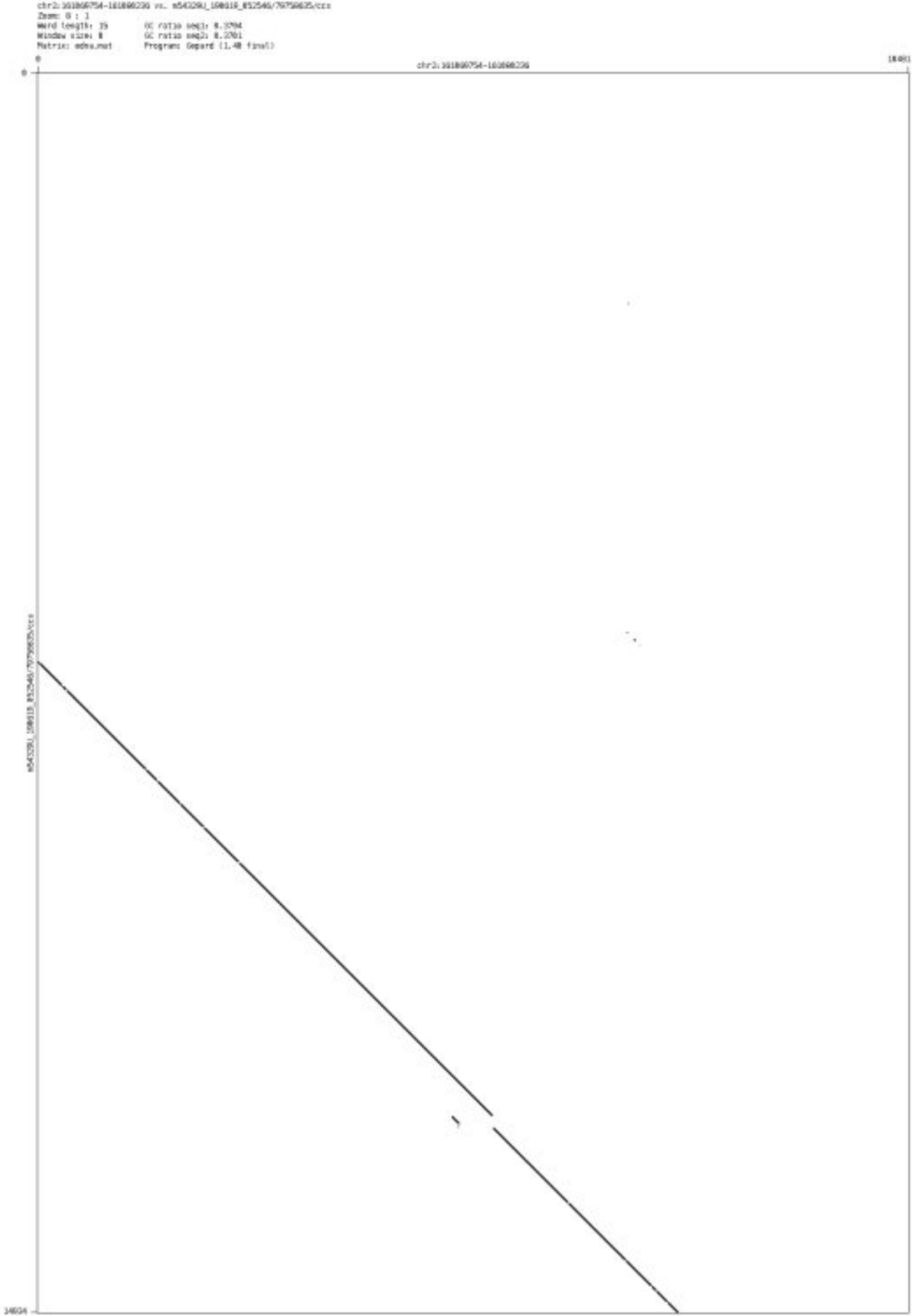

### ➢ chr3_6604527_6613044

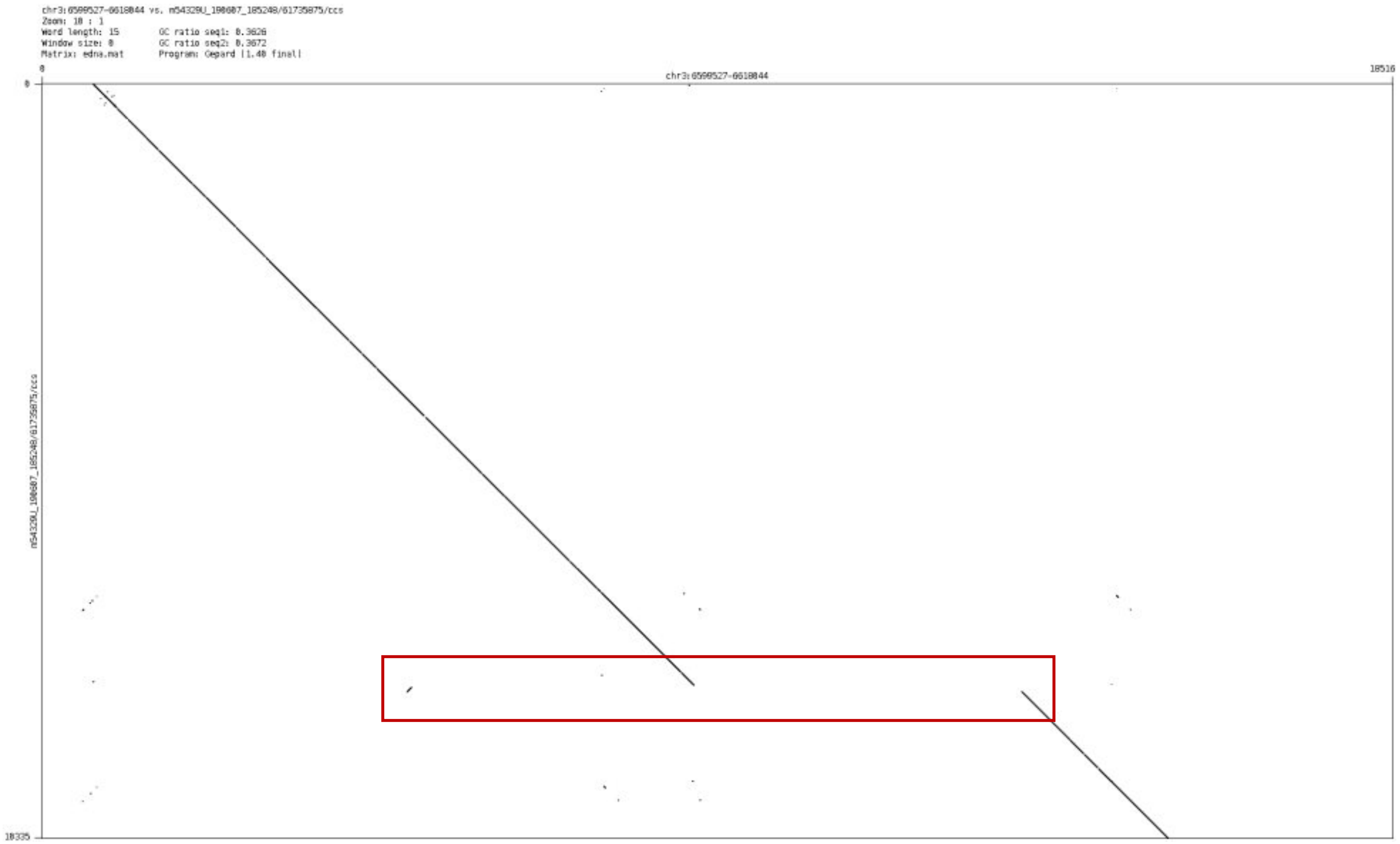

➢ chr3_41320442_41322556

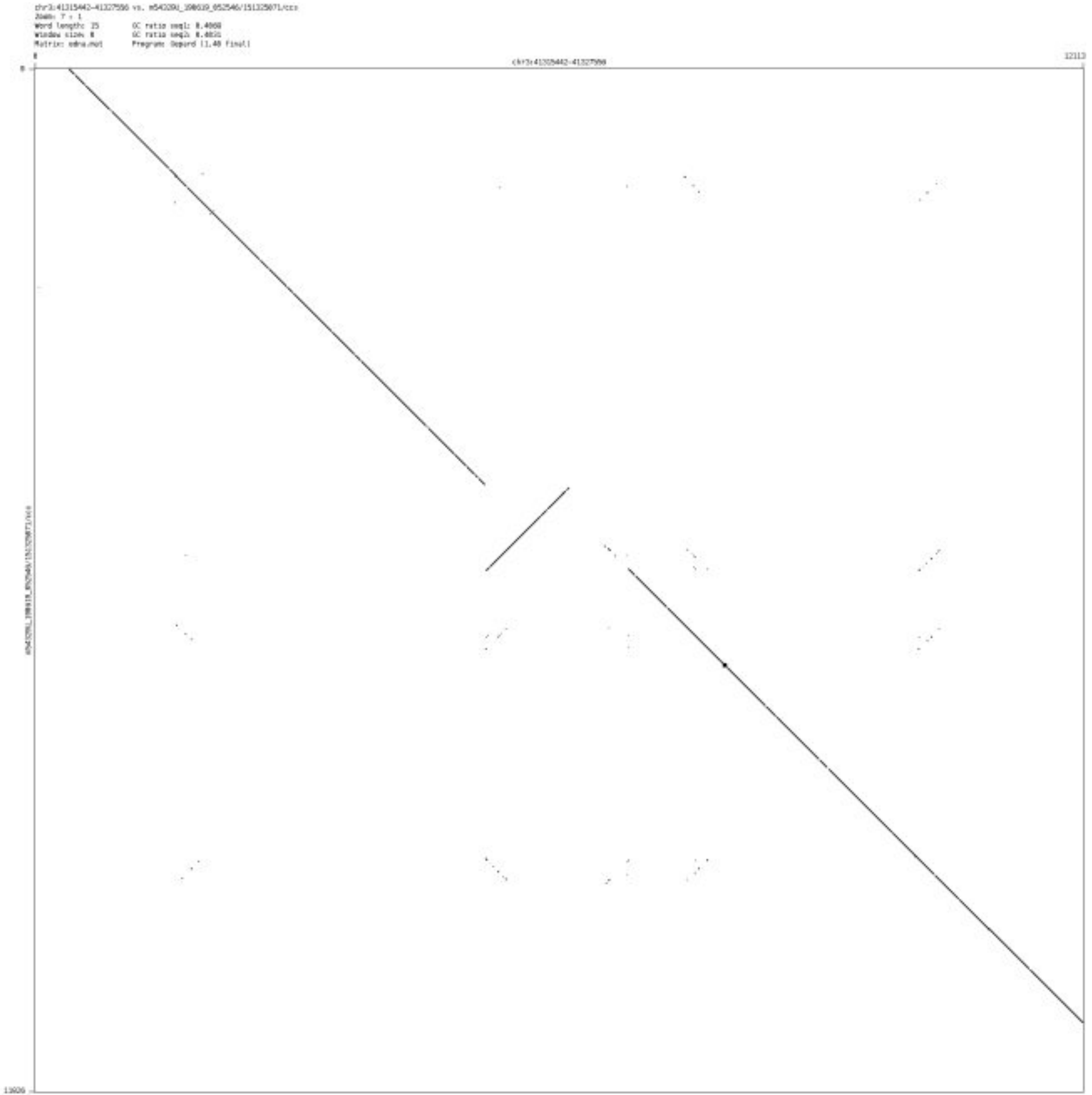

➢ chr3_80013338_80016105

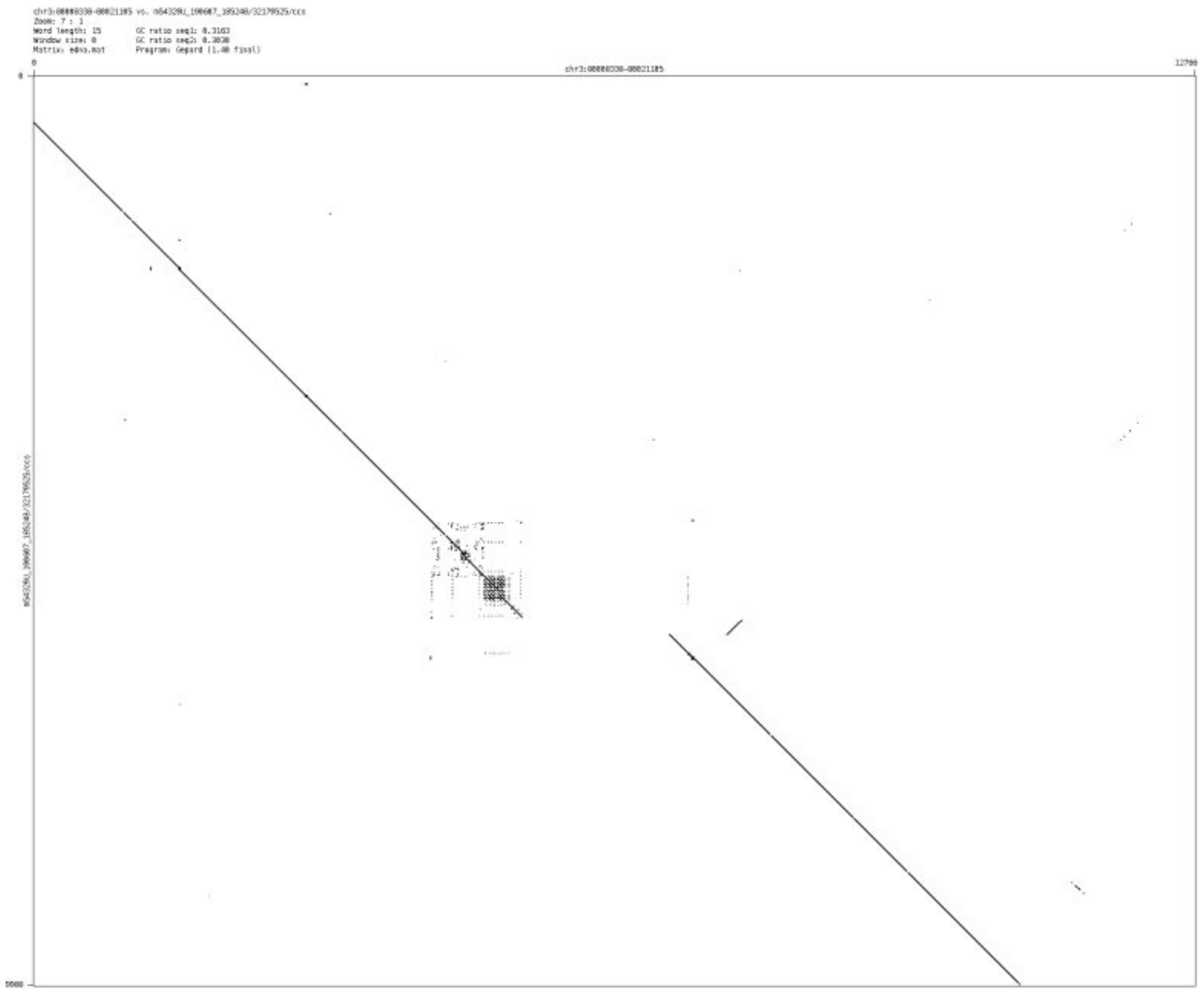

- chr3_95749273_95752156

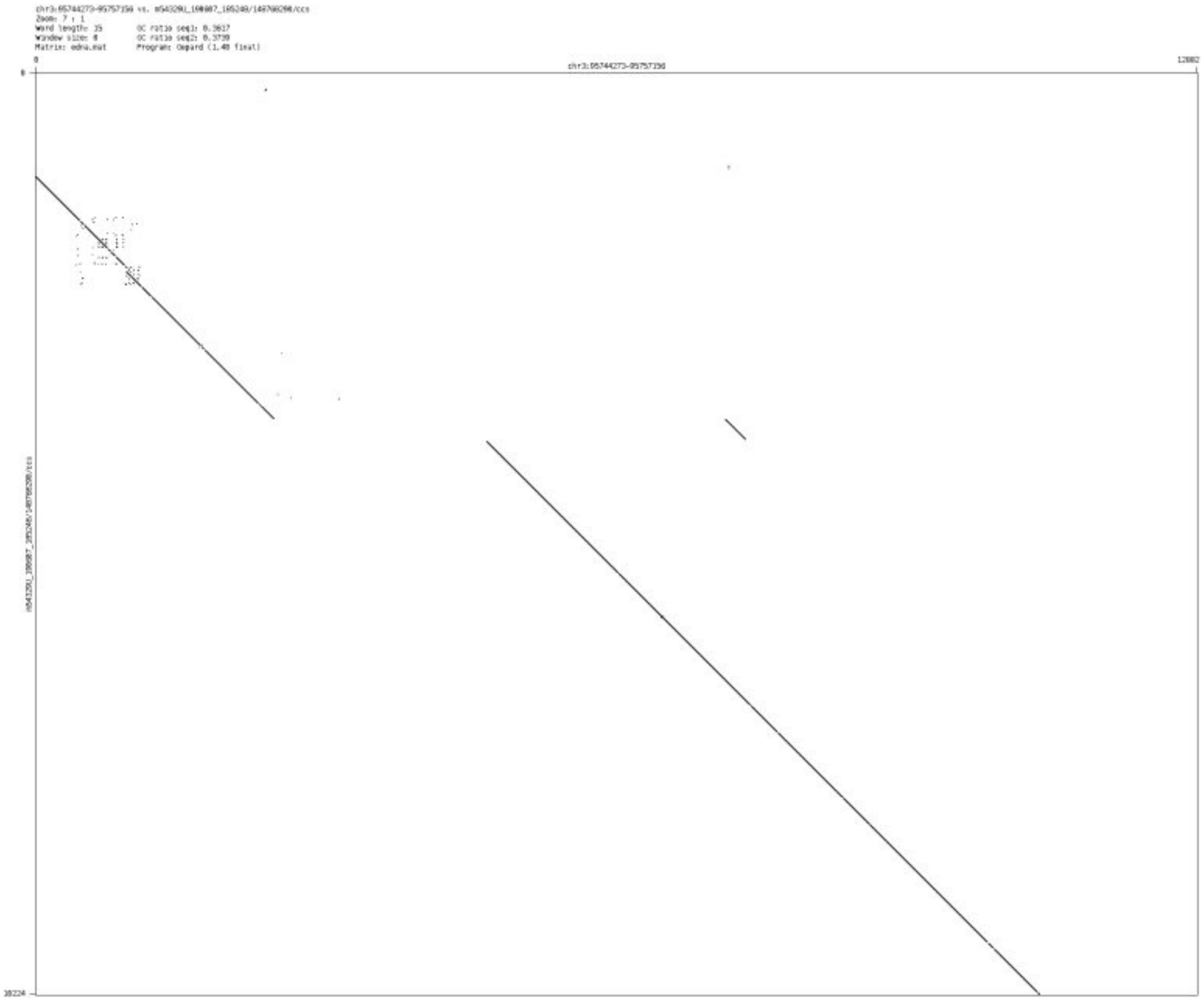

- chr3_131988873_131994701

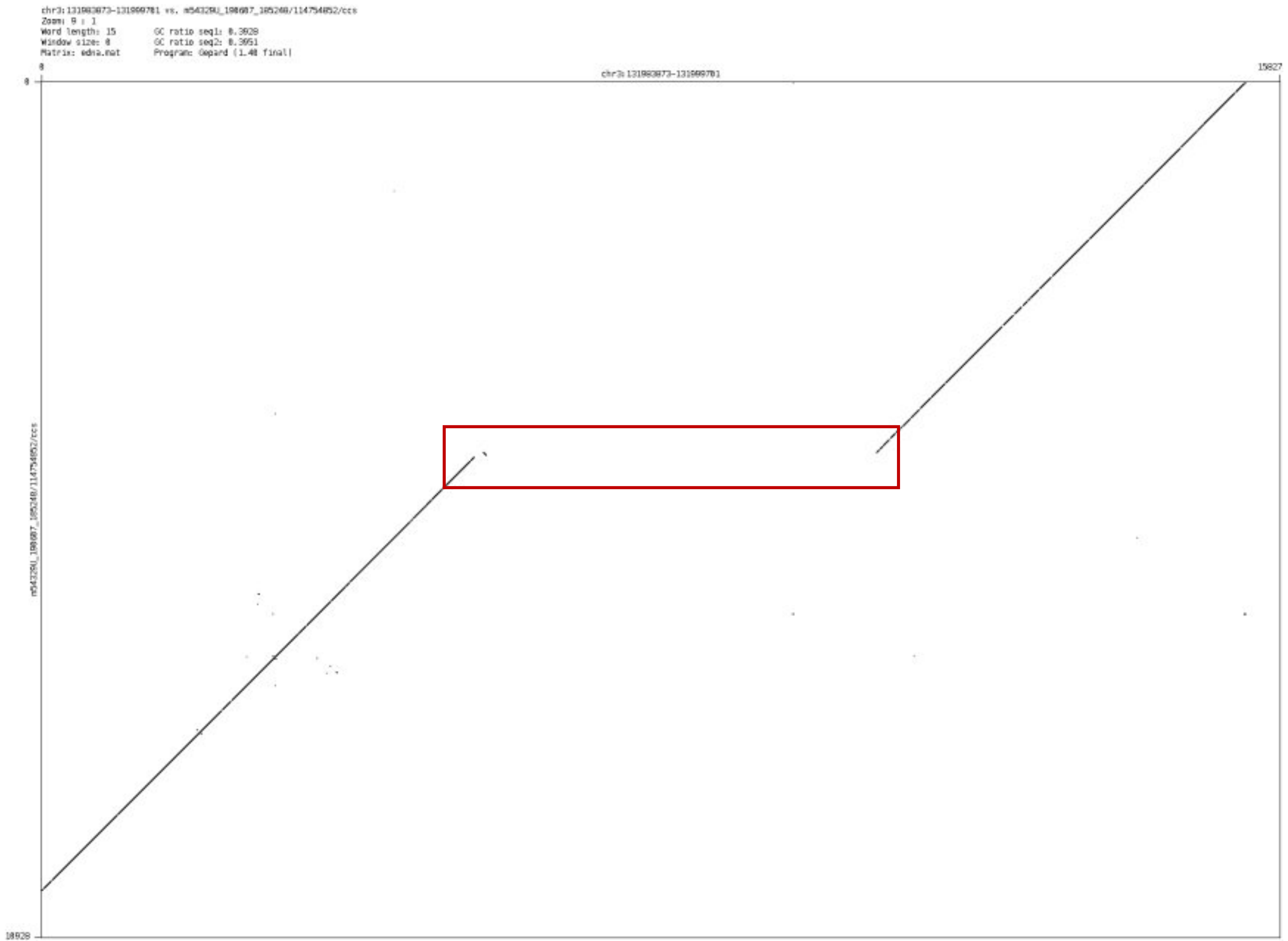

- chr3_146667093_146677284

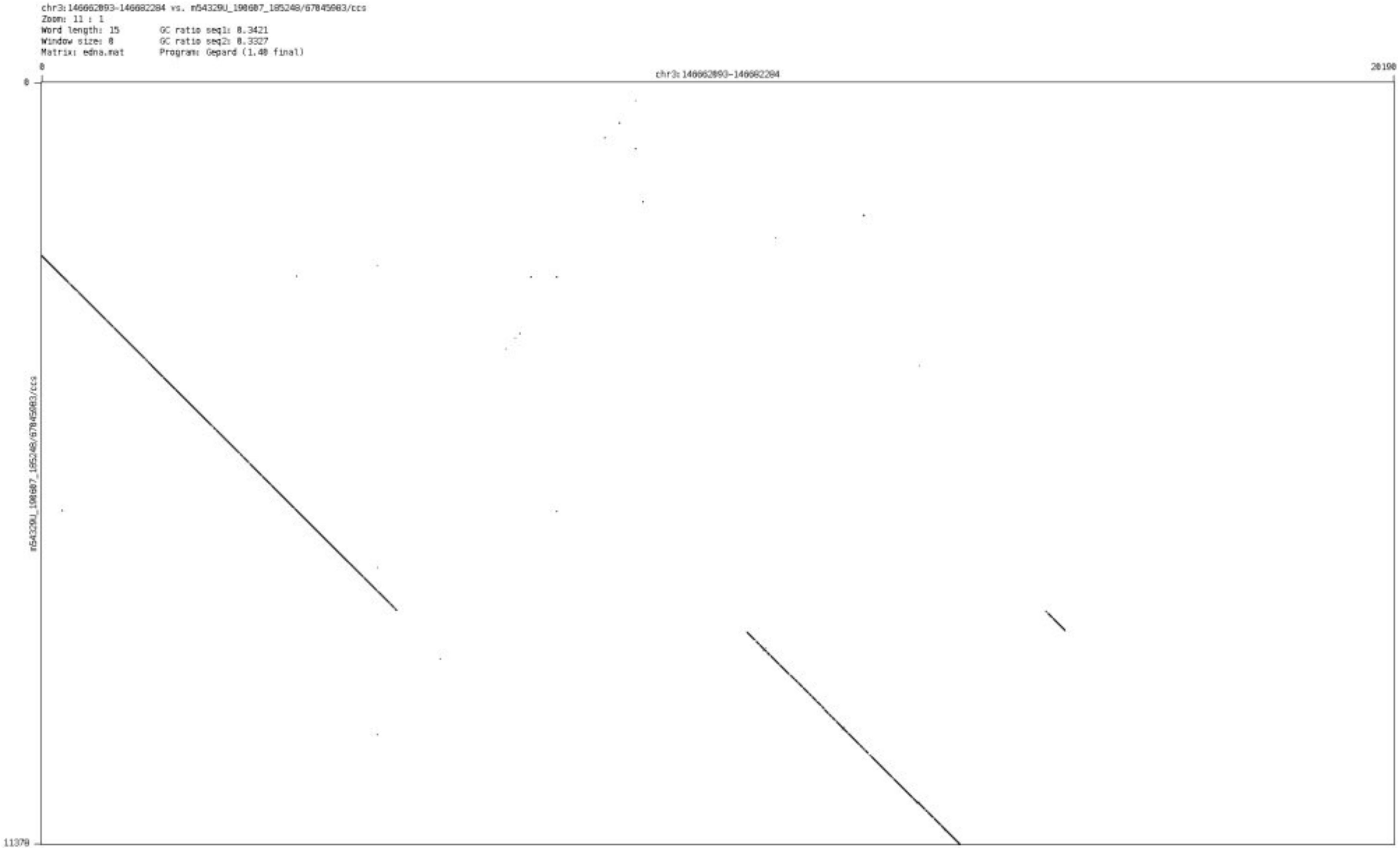

➢ chr3_162807802_162830163

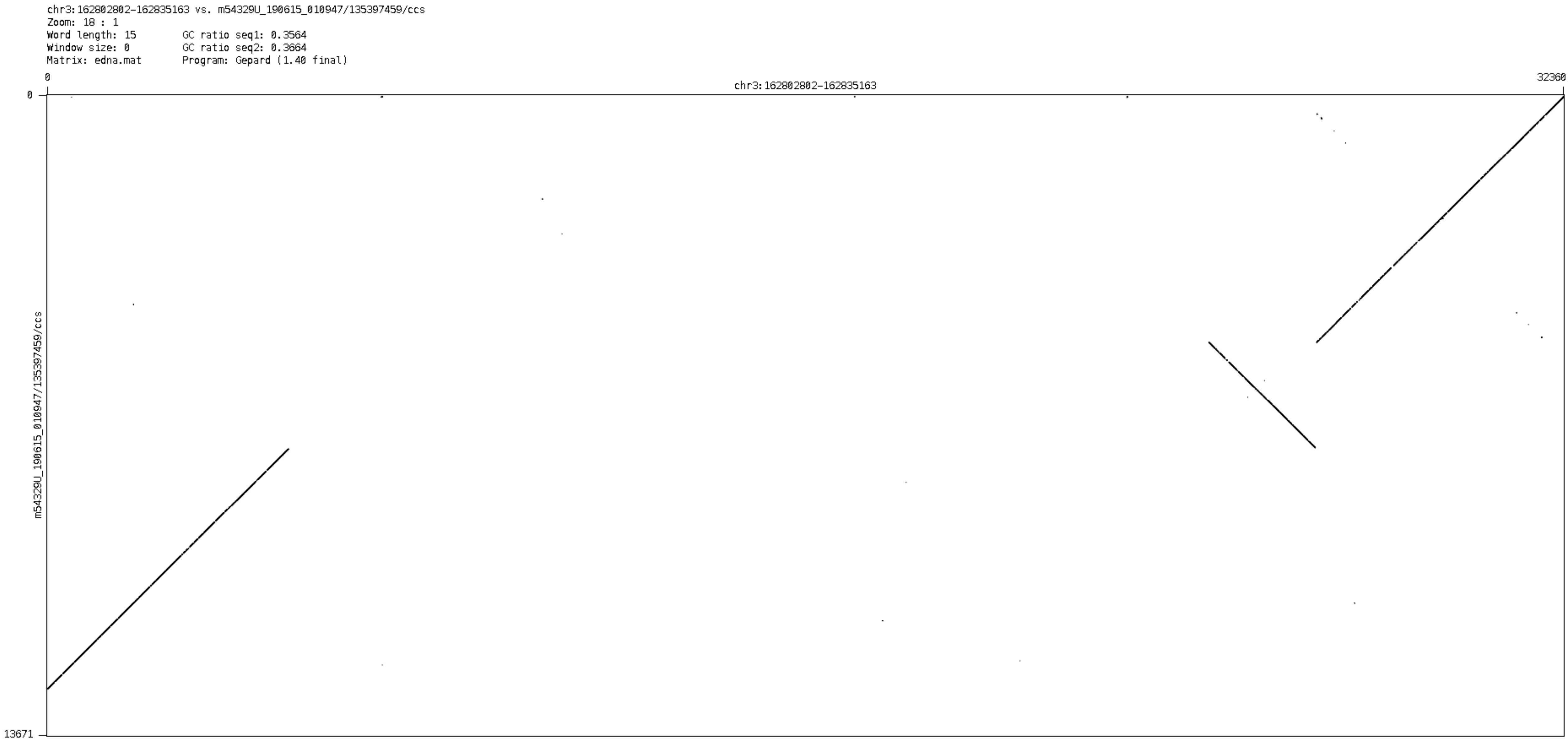

- chr4_58562600_58569633

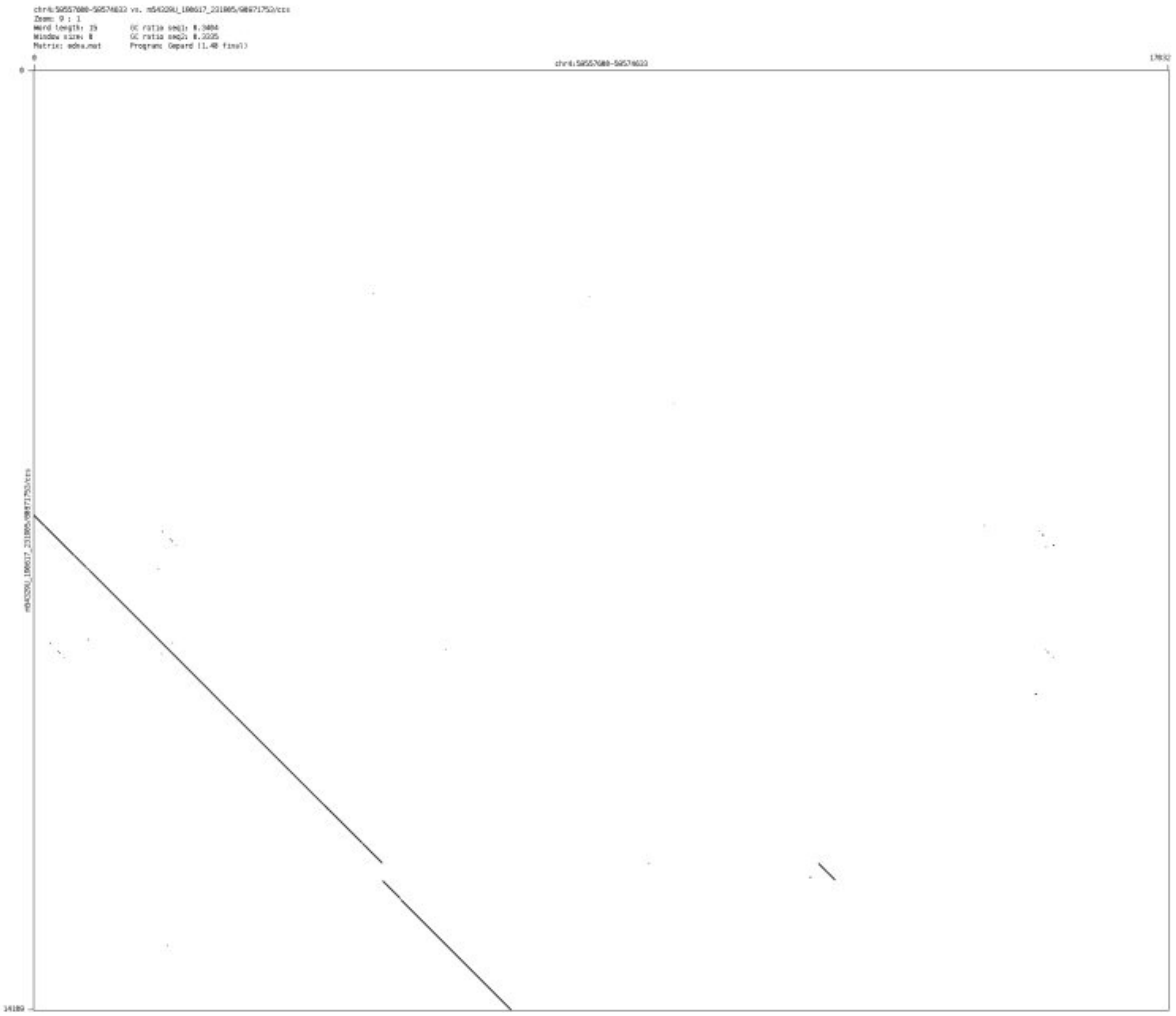

- chr4_92646227_92649130

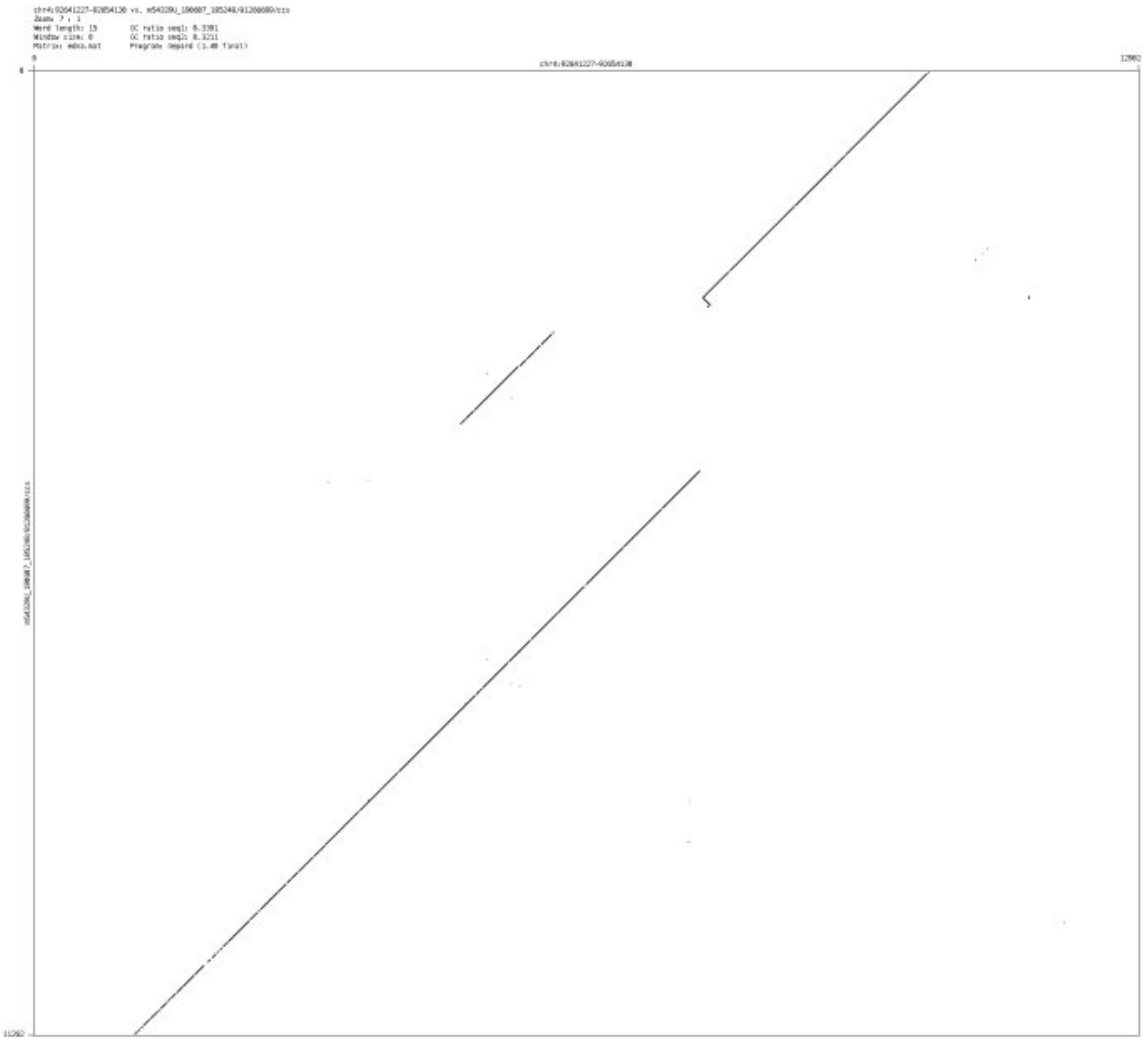

- chr4_162657226_162658438

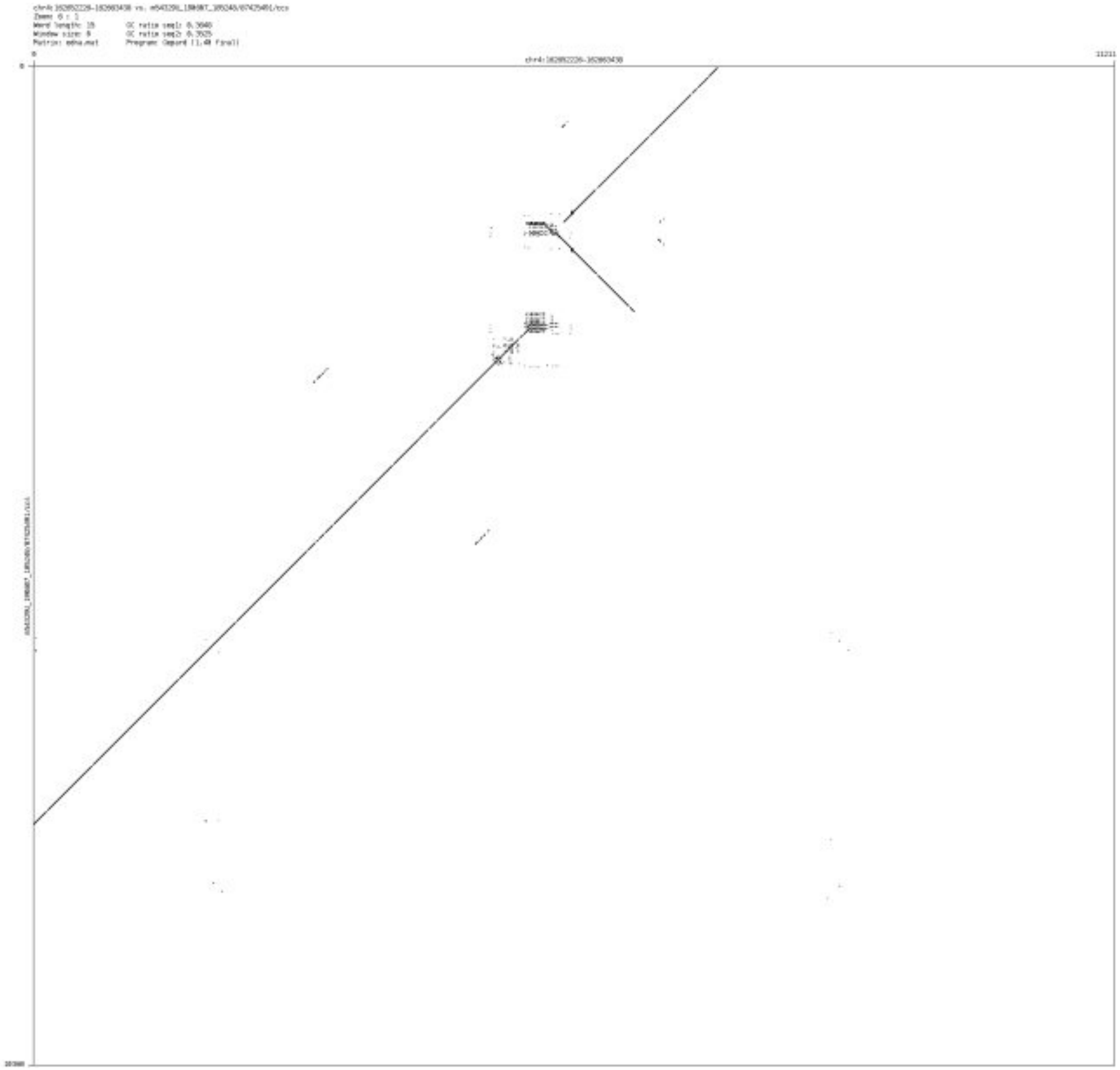

➢ chr5_28932185_28934937

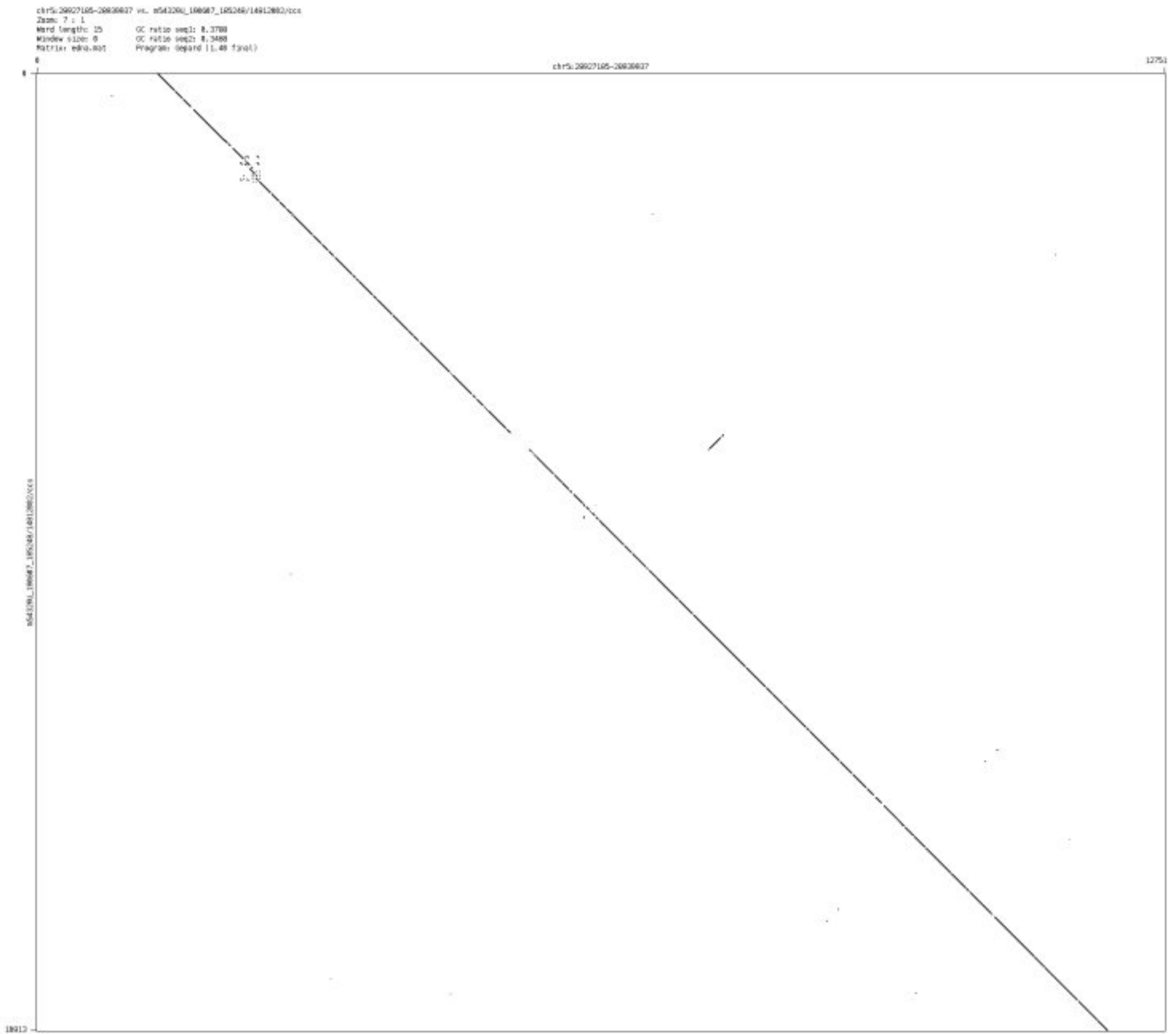

➢ chr5_53826784_53830121

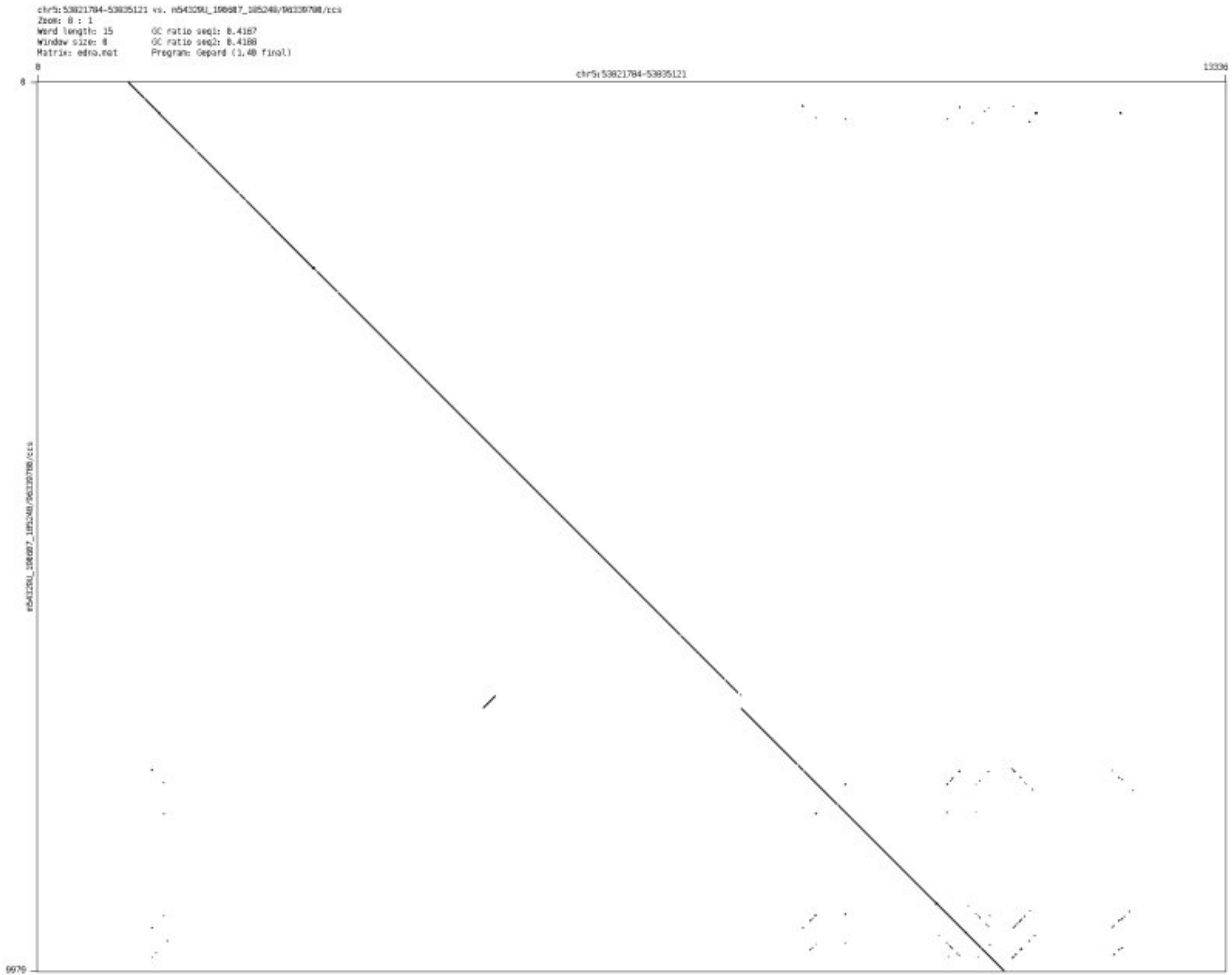

➢ chr5_116010763_116015366

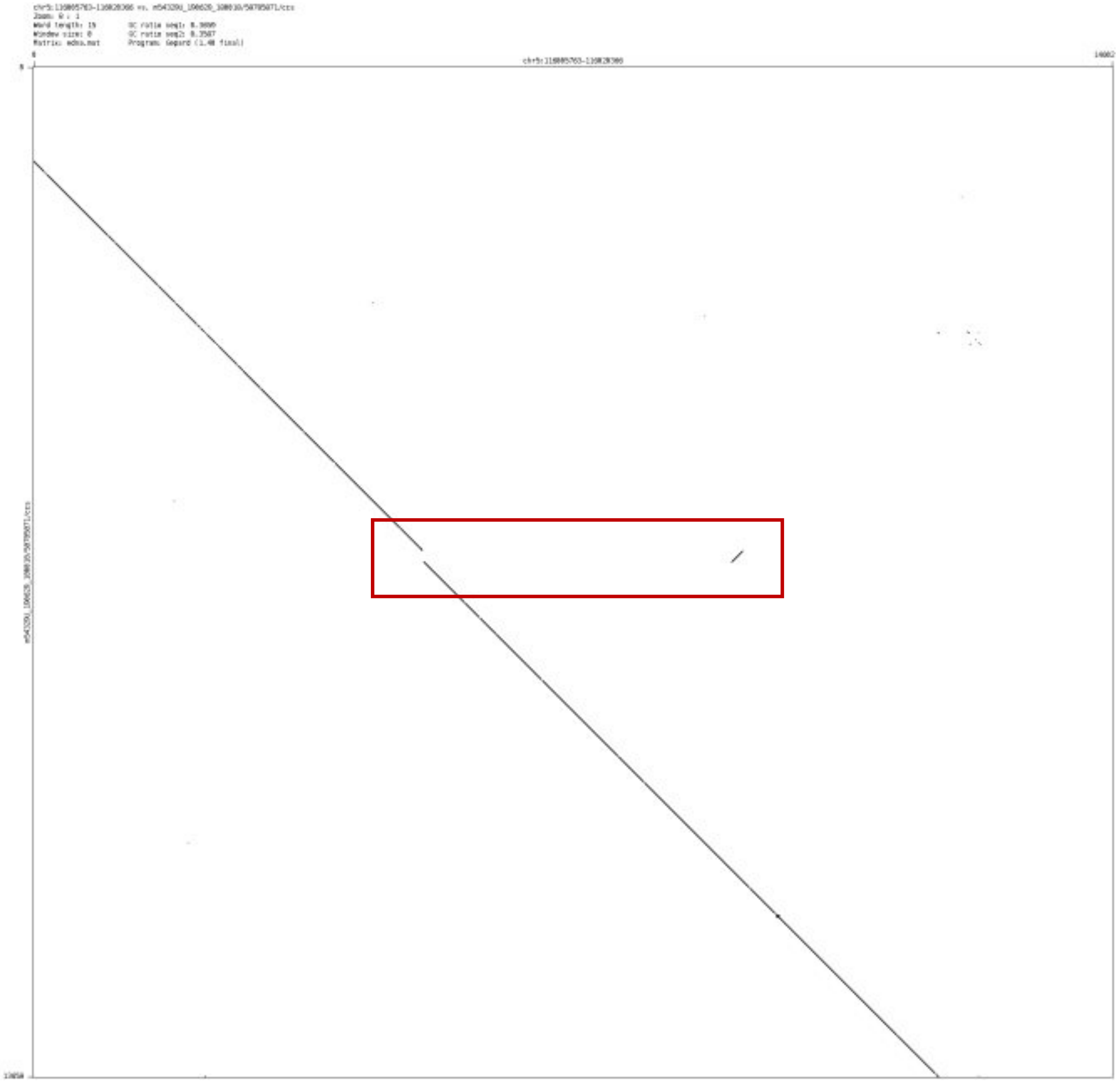

- chr5_141480327_141483116

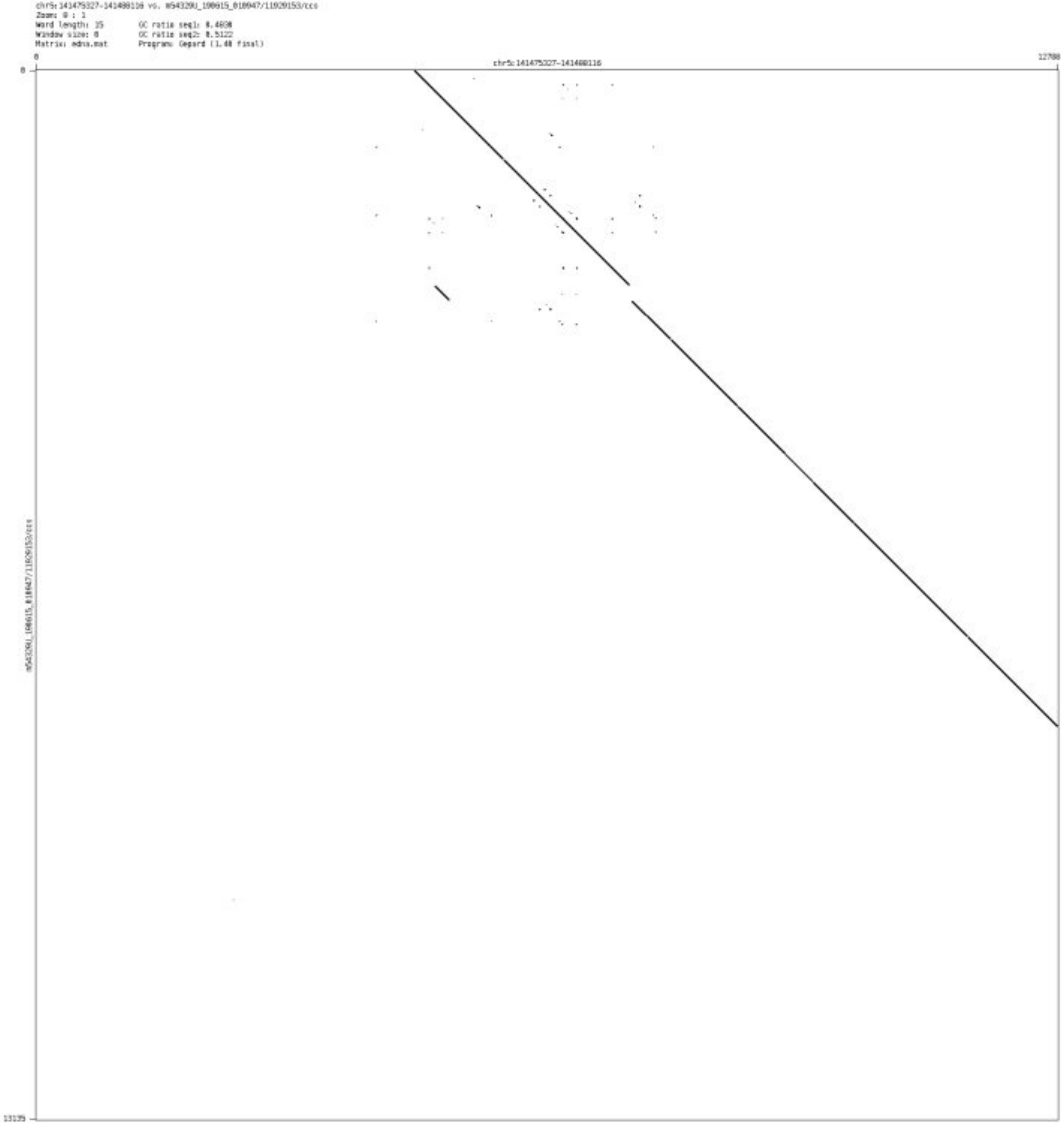

- chr5_144132924_144135827

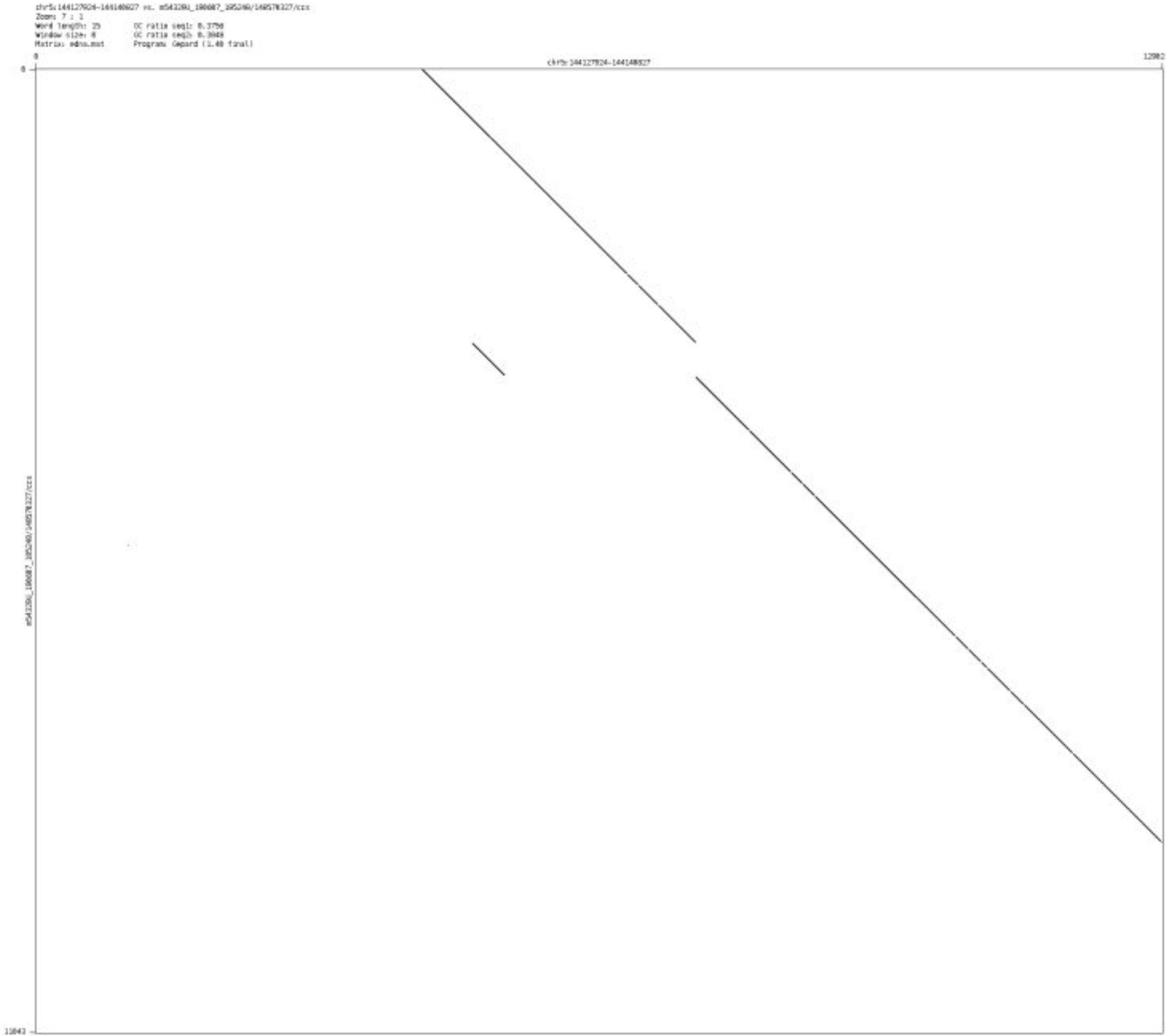

- chr5_148173301_148175687

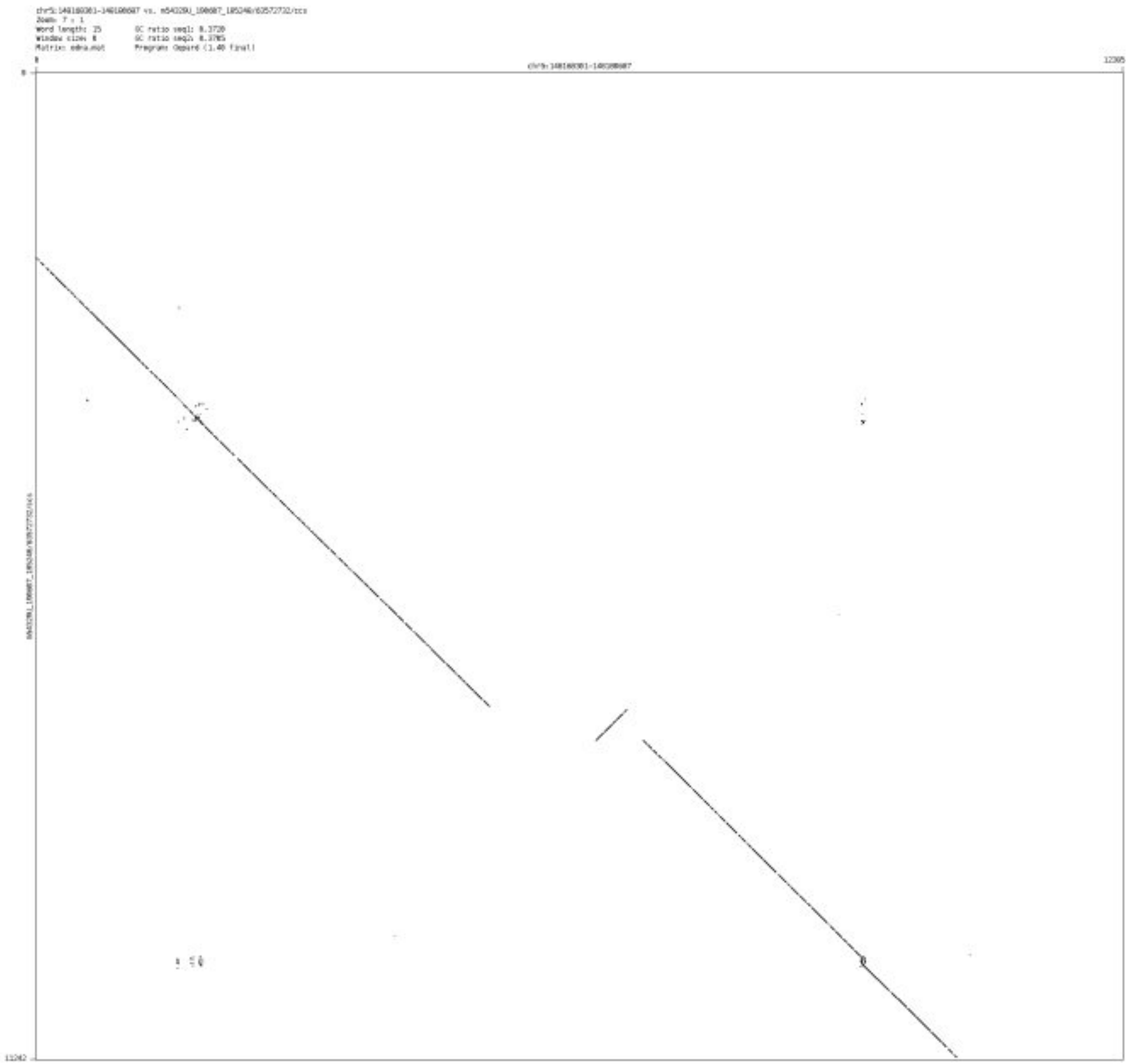

- chr6_54059863_54070121

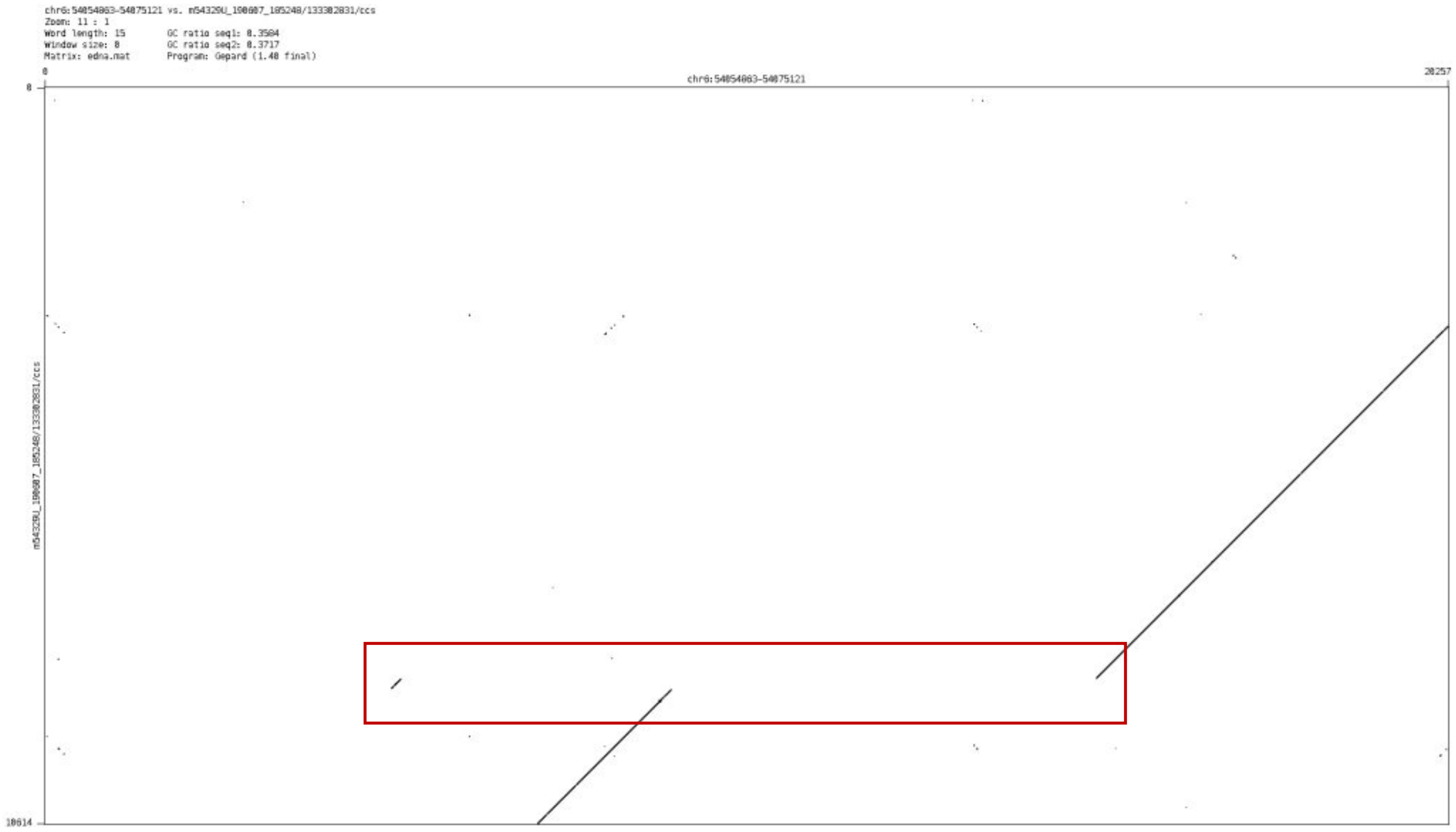

- chr6_89211609_89214231

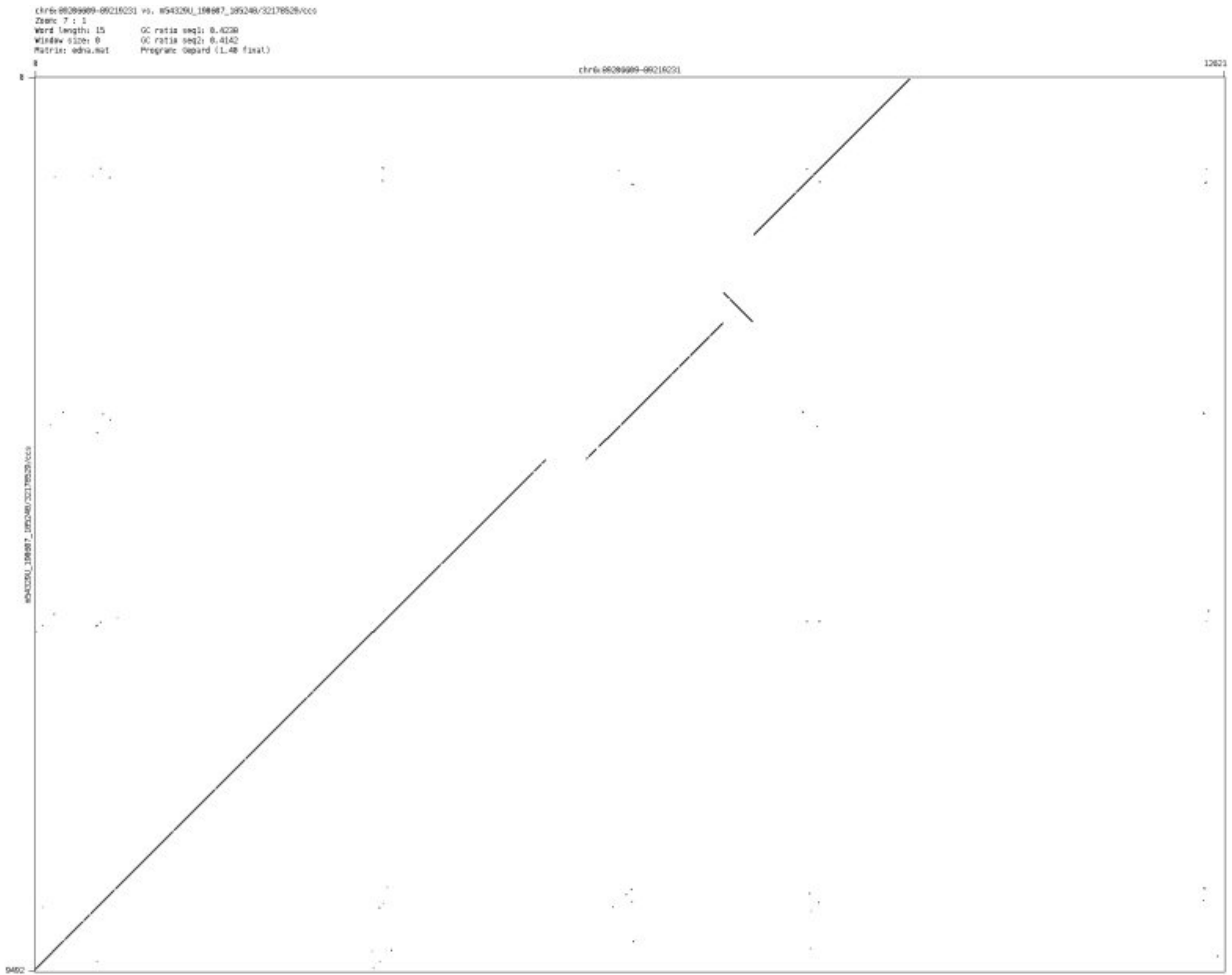

➢ chr6_118690556_118692766

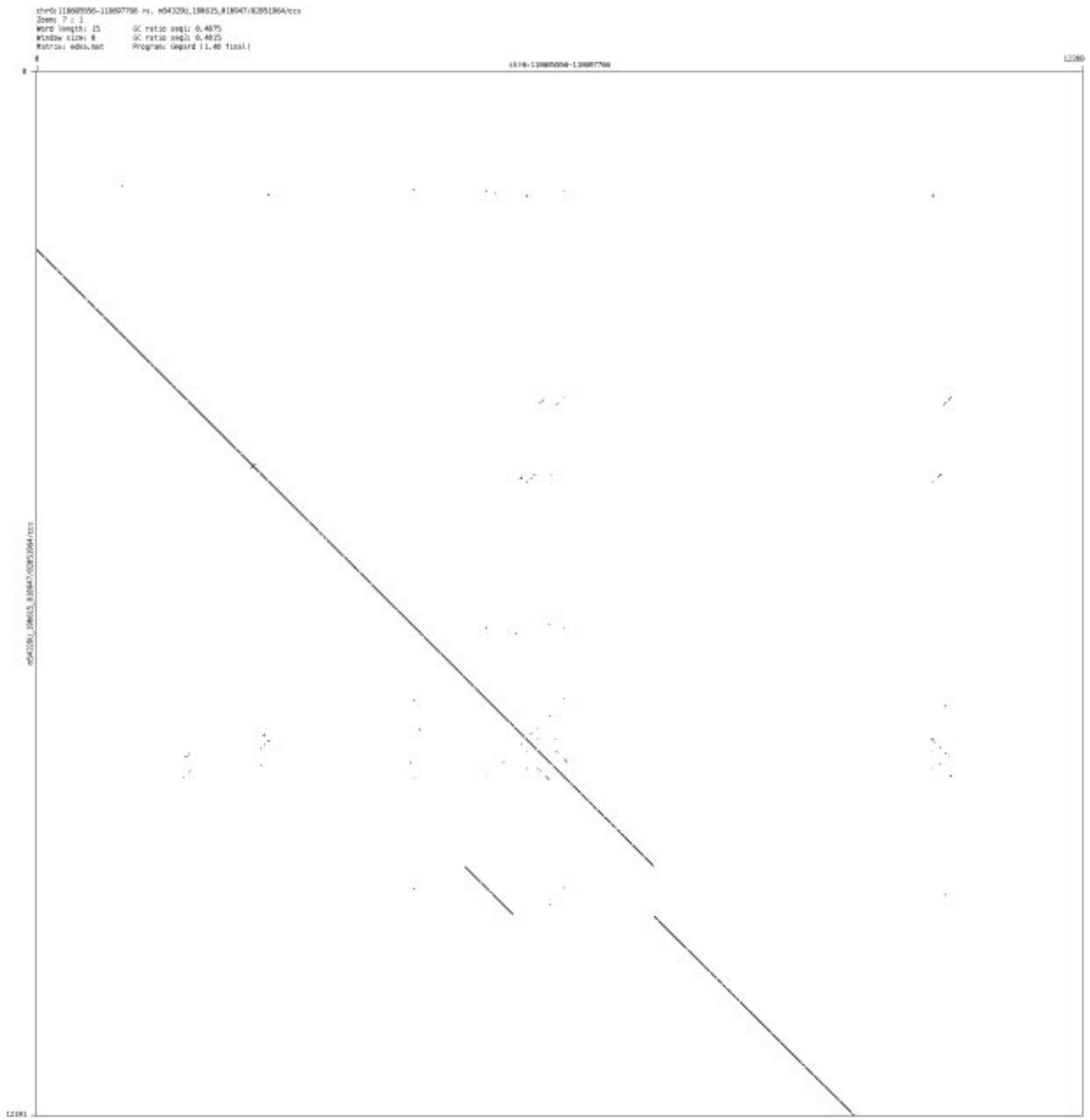

- chr6_150946253_150948473

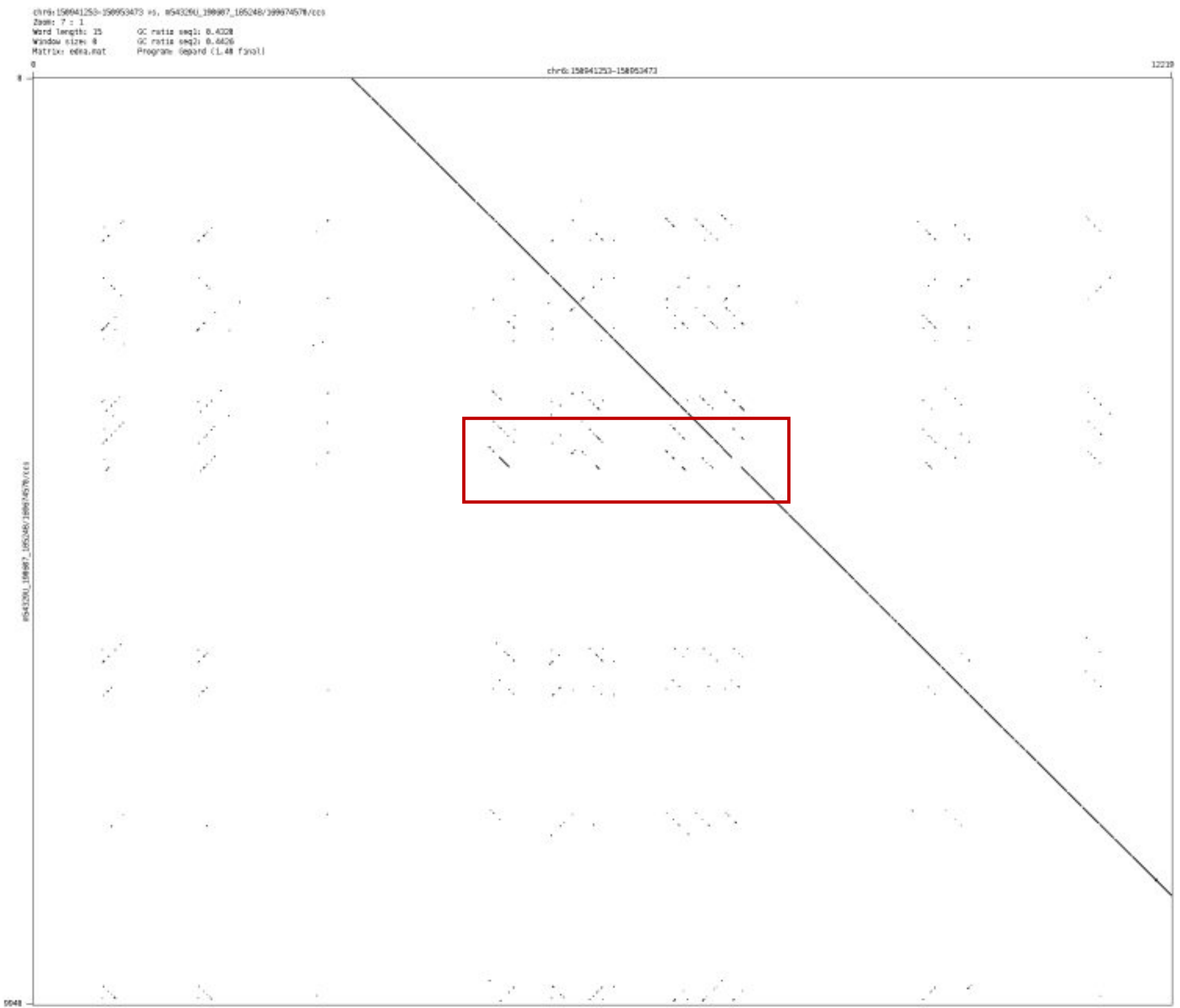

### ➢ chr7_70955852_70974154

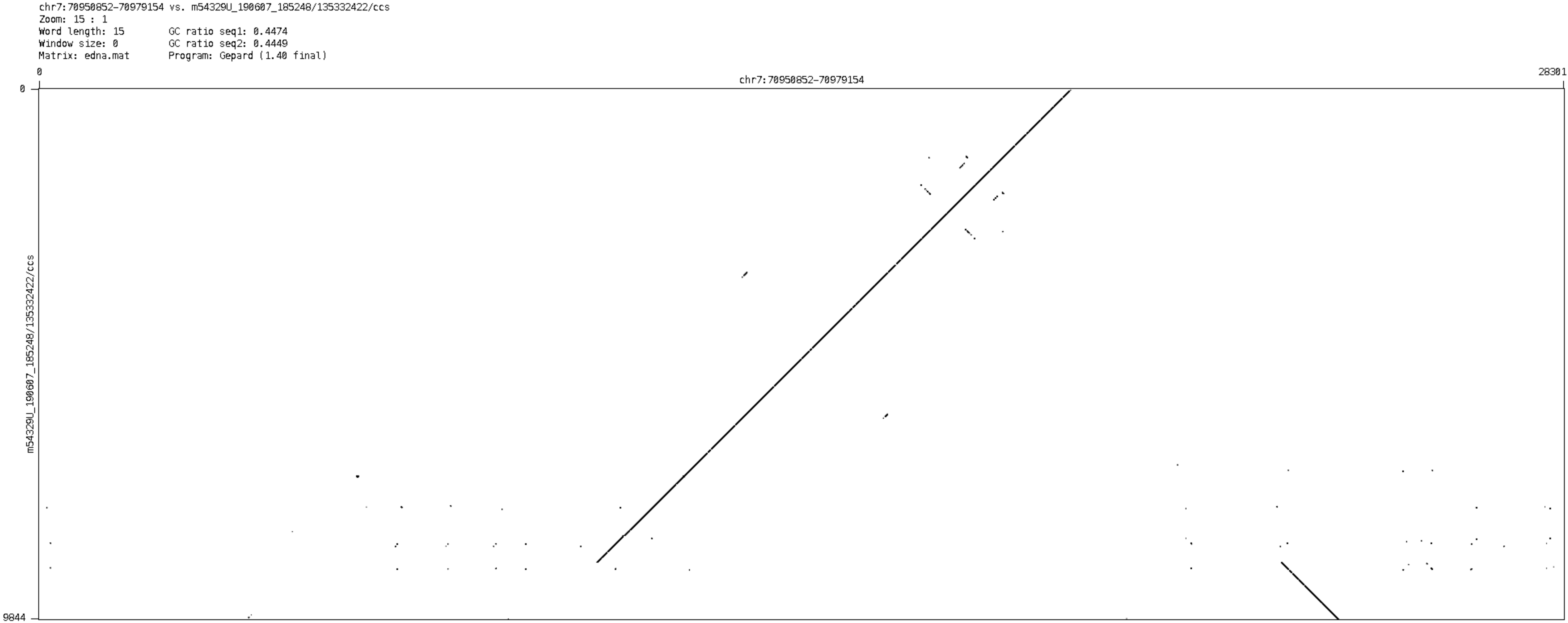

➢ chr7_85460059_85462014

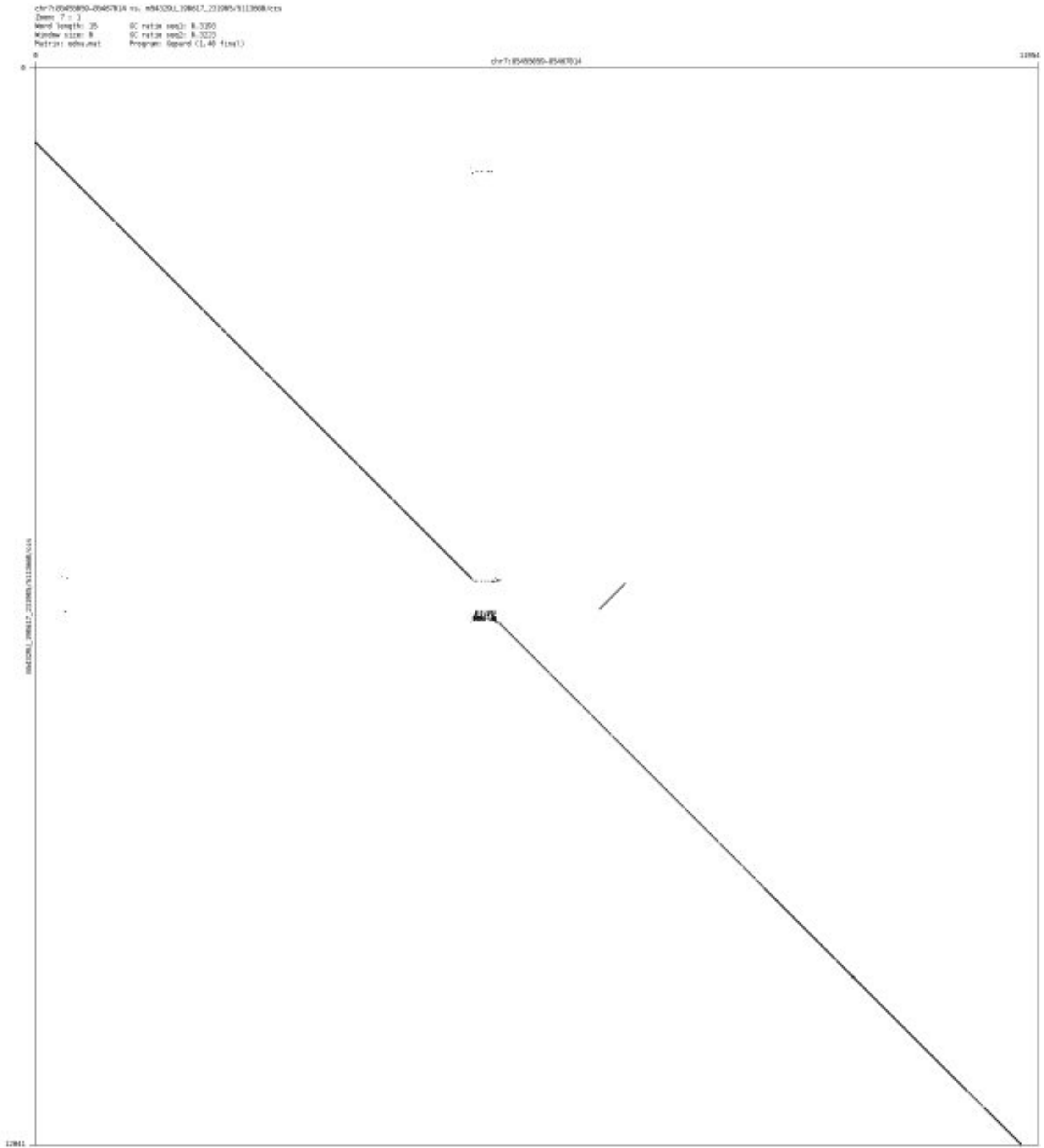

- chr7_102080906_102085205

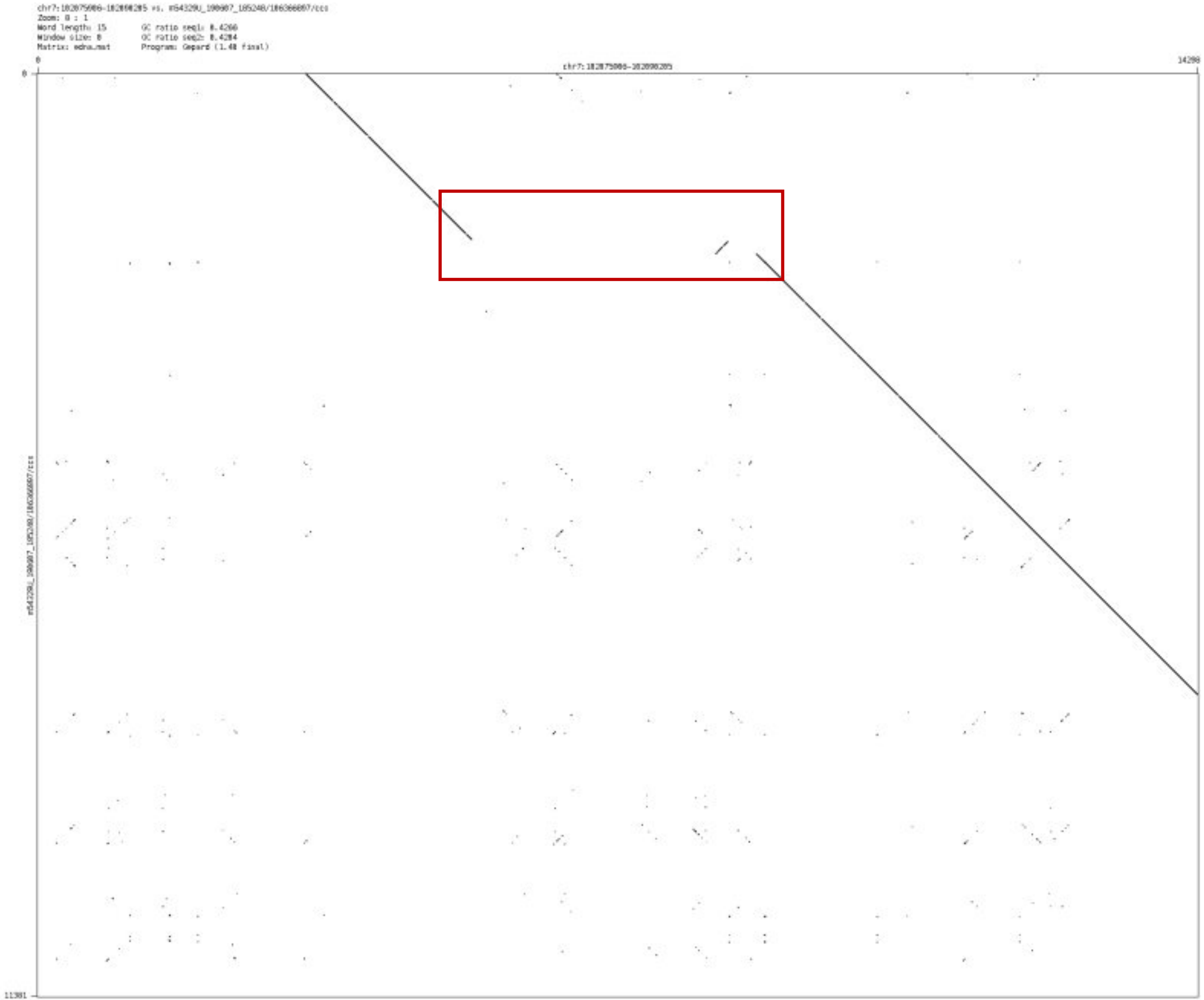

➢ chr8_92178203_92183376

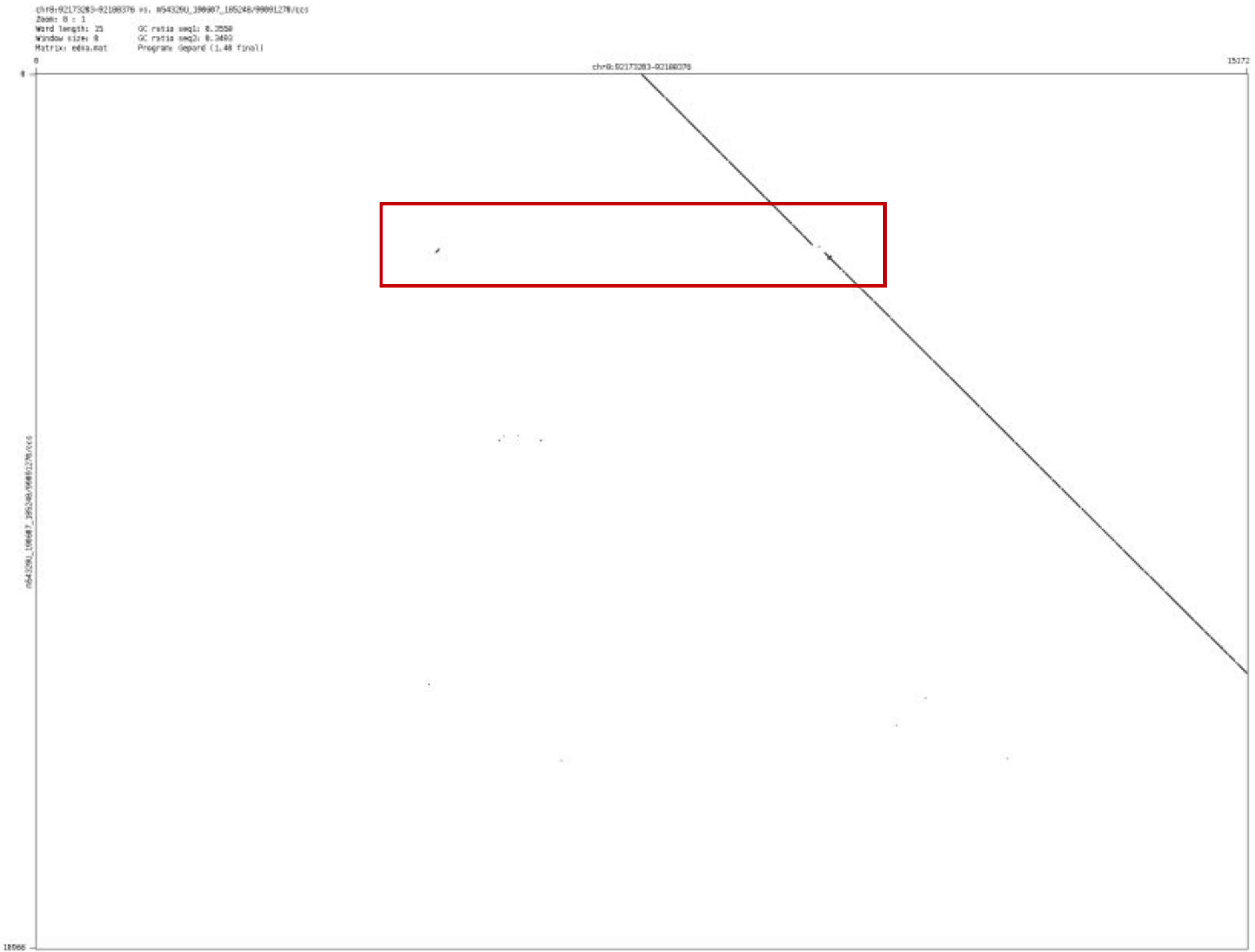

- ➢ chr9_29591409_29593057

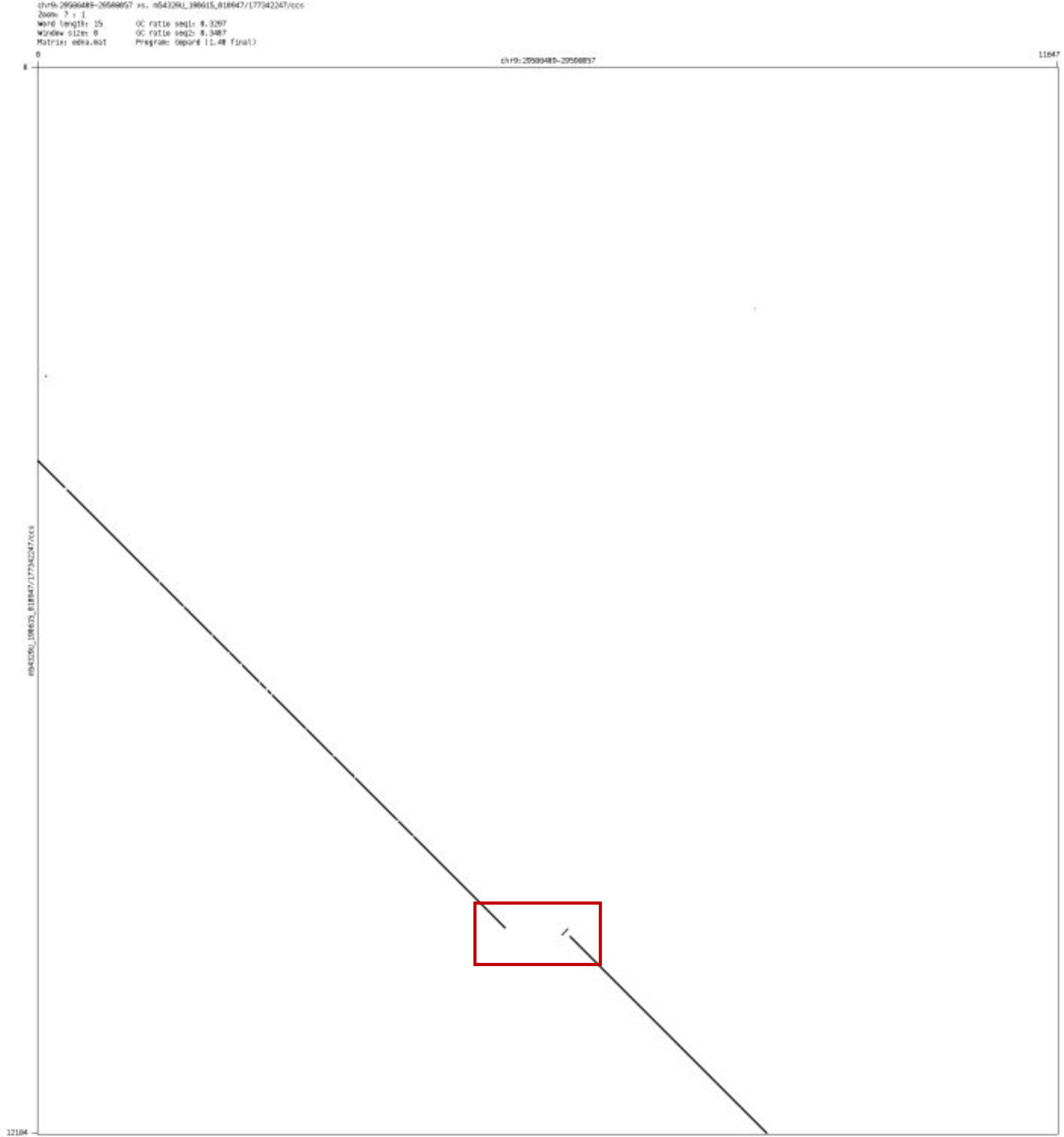

#### ➢ chr9_62806317_62823001

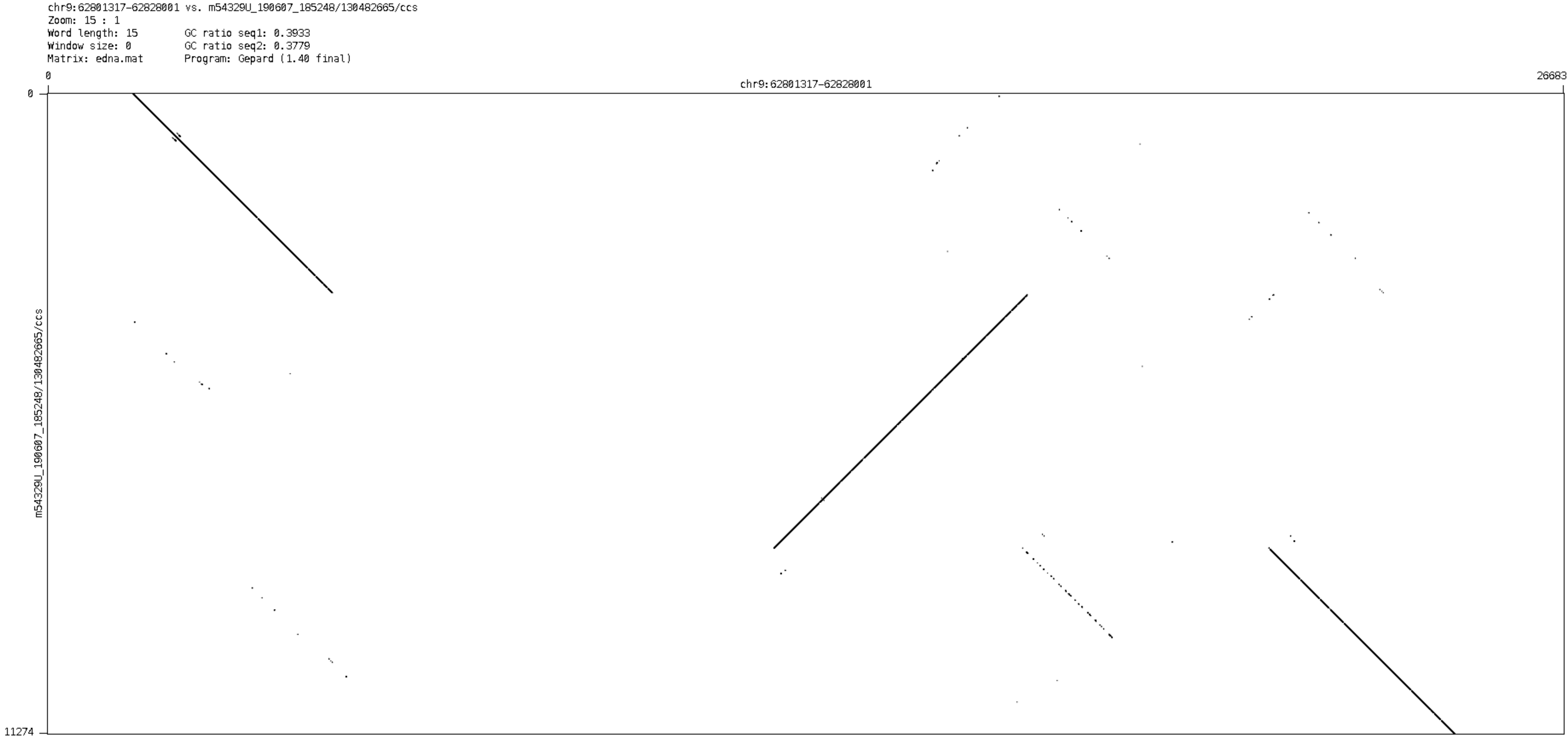

- chr9_109523566_109524653

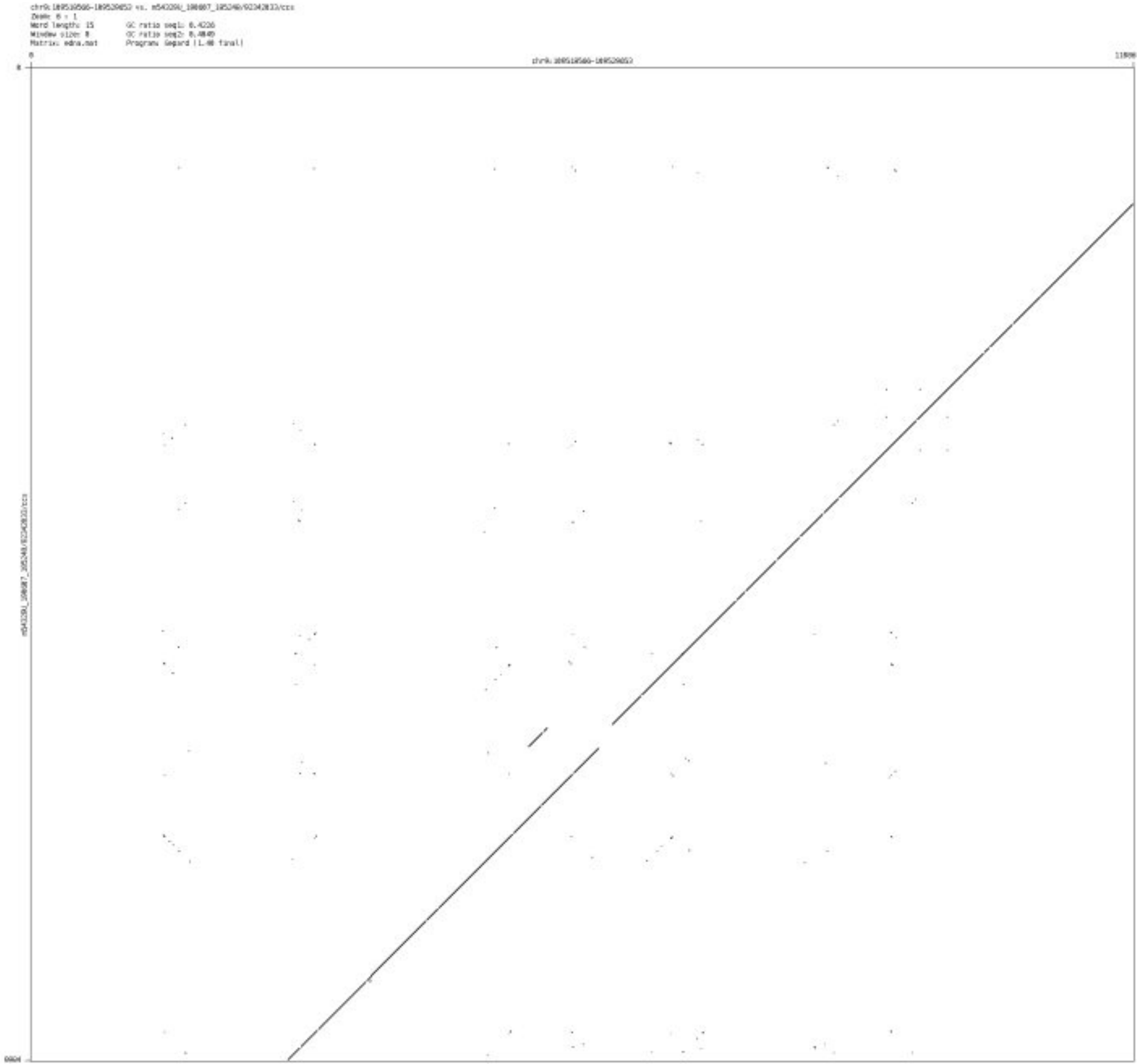

- chr10_44927520_44928982

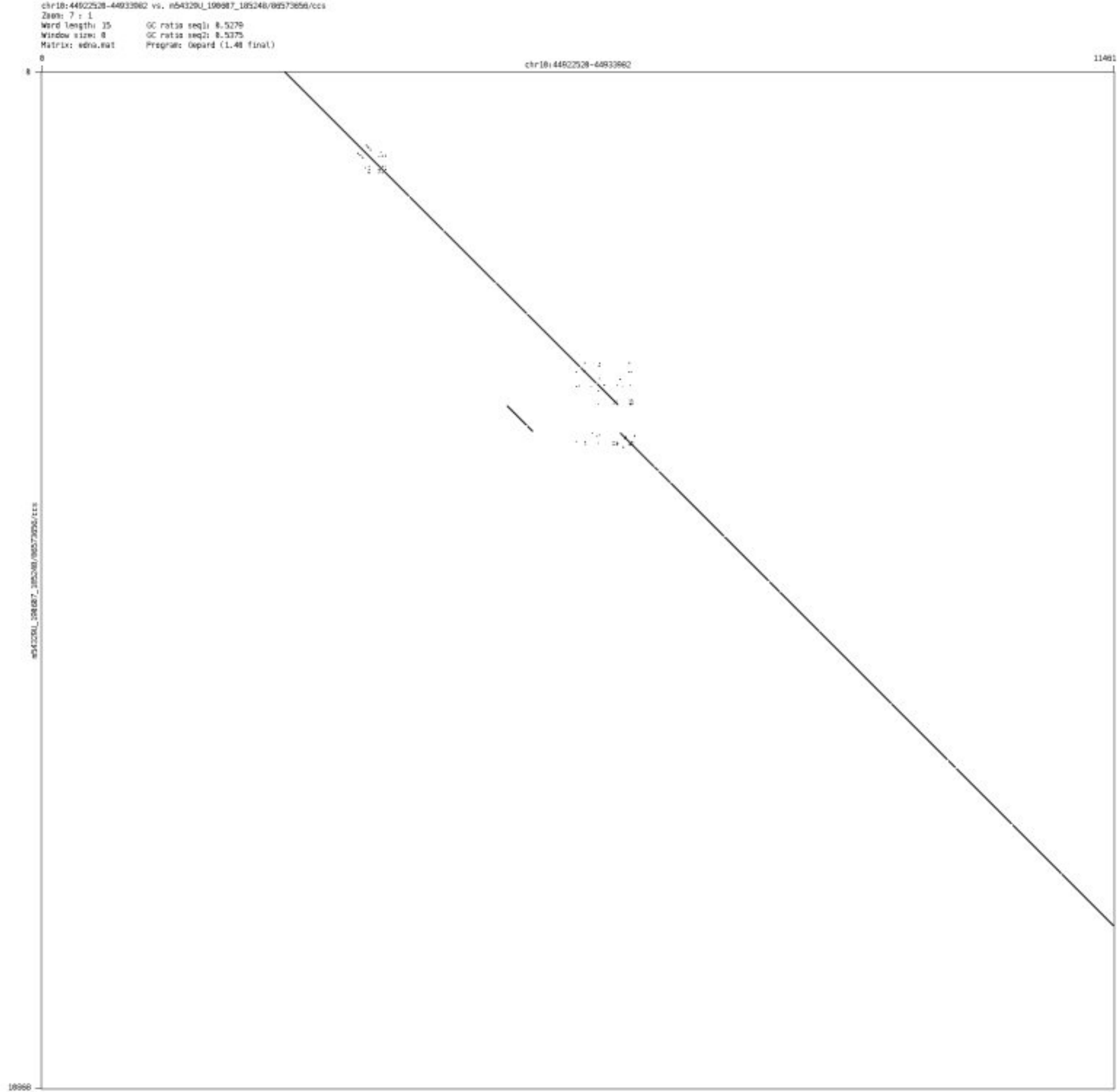

- chr10_57496913_57498588

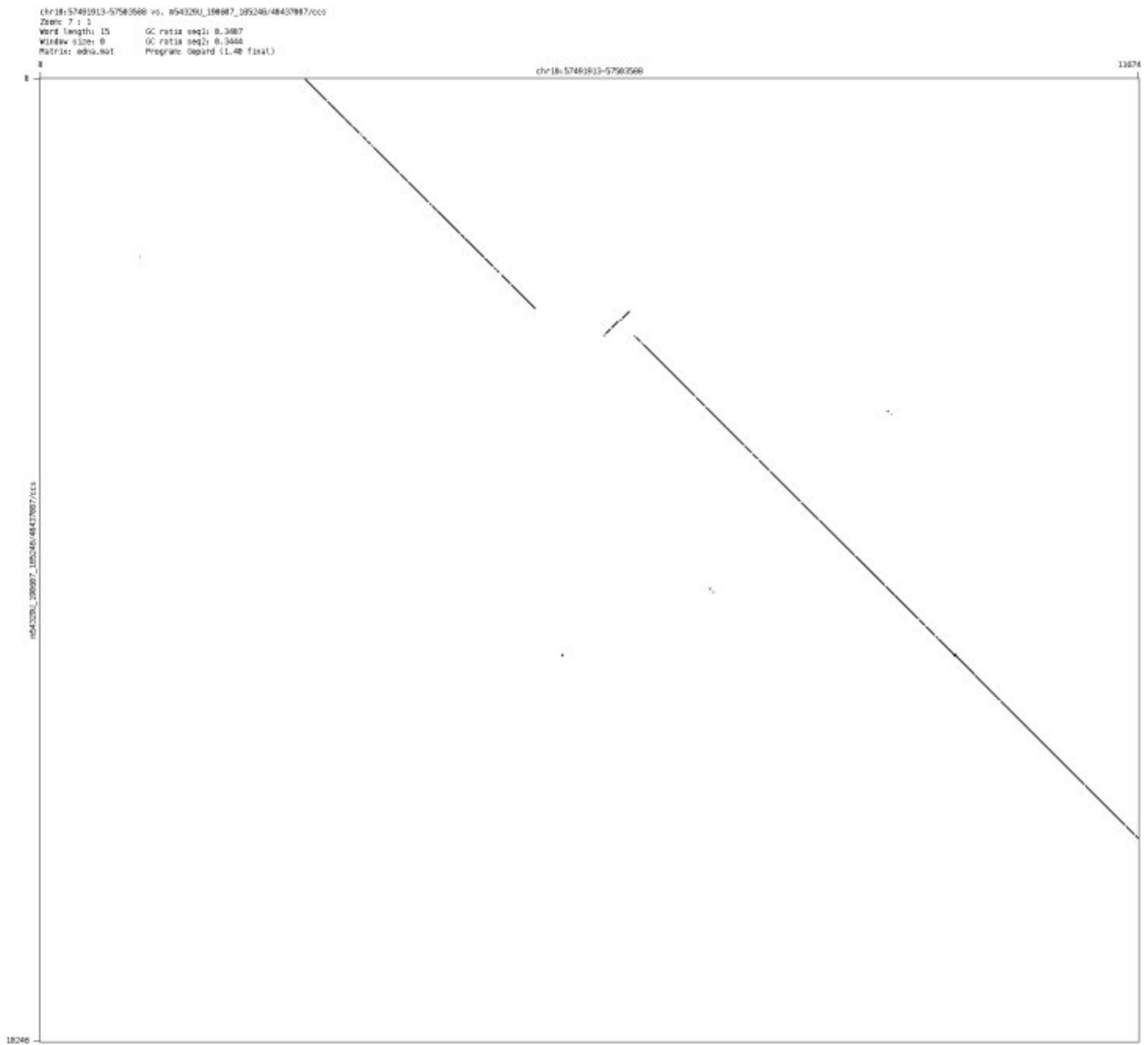

### ➢ chr10_95447029_95448567

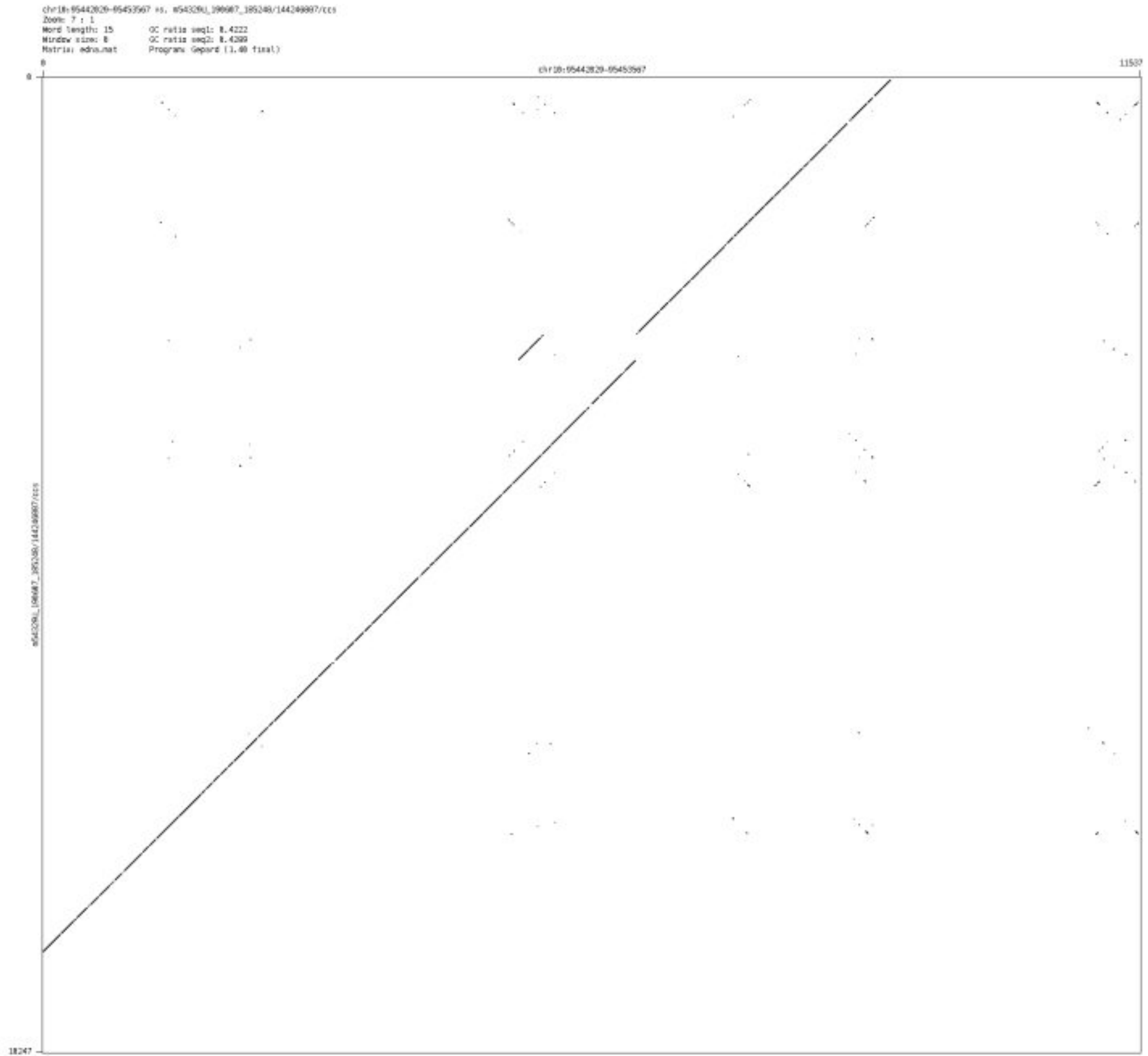

- chr10_125501894_125512533

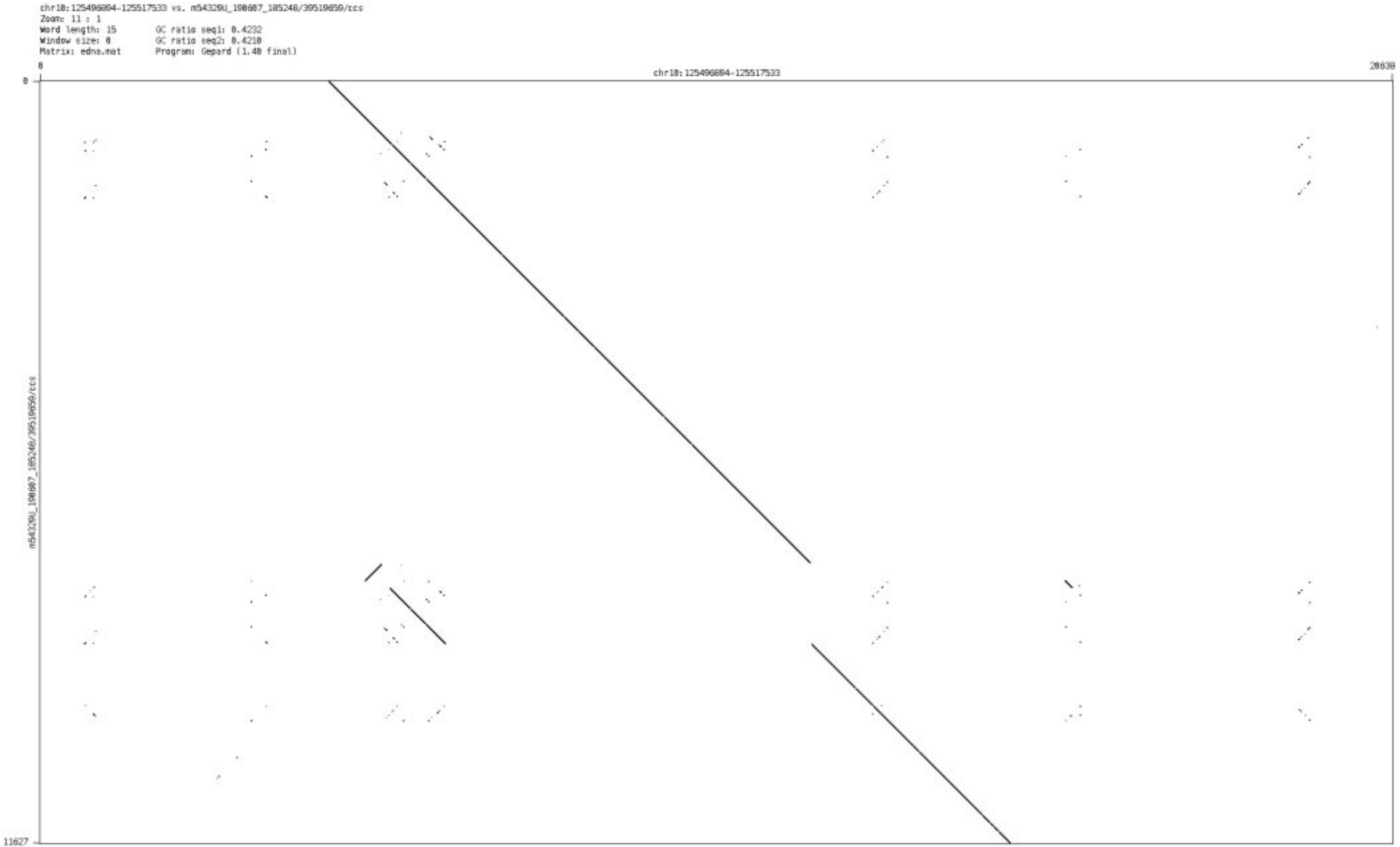

### ➢ chr11_59277818_59282964

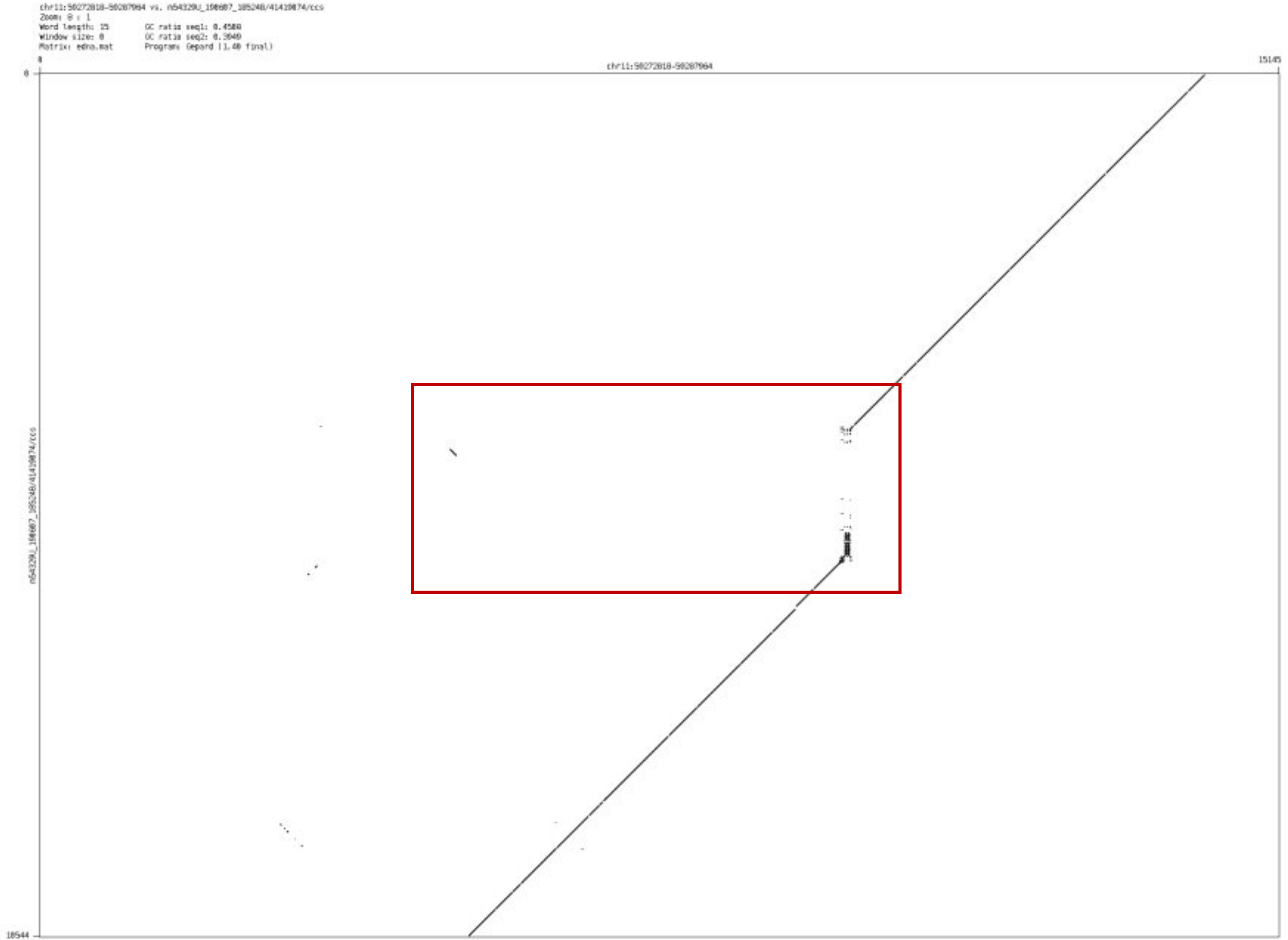

➢ chr11_63930944_63934473

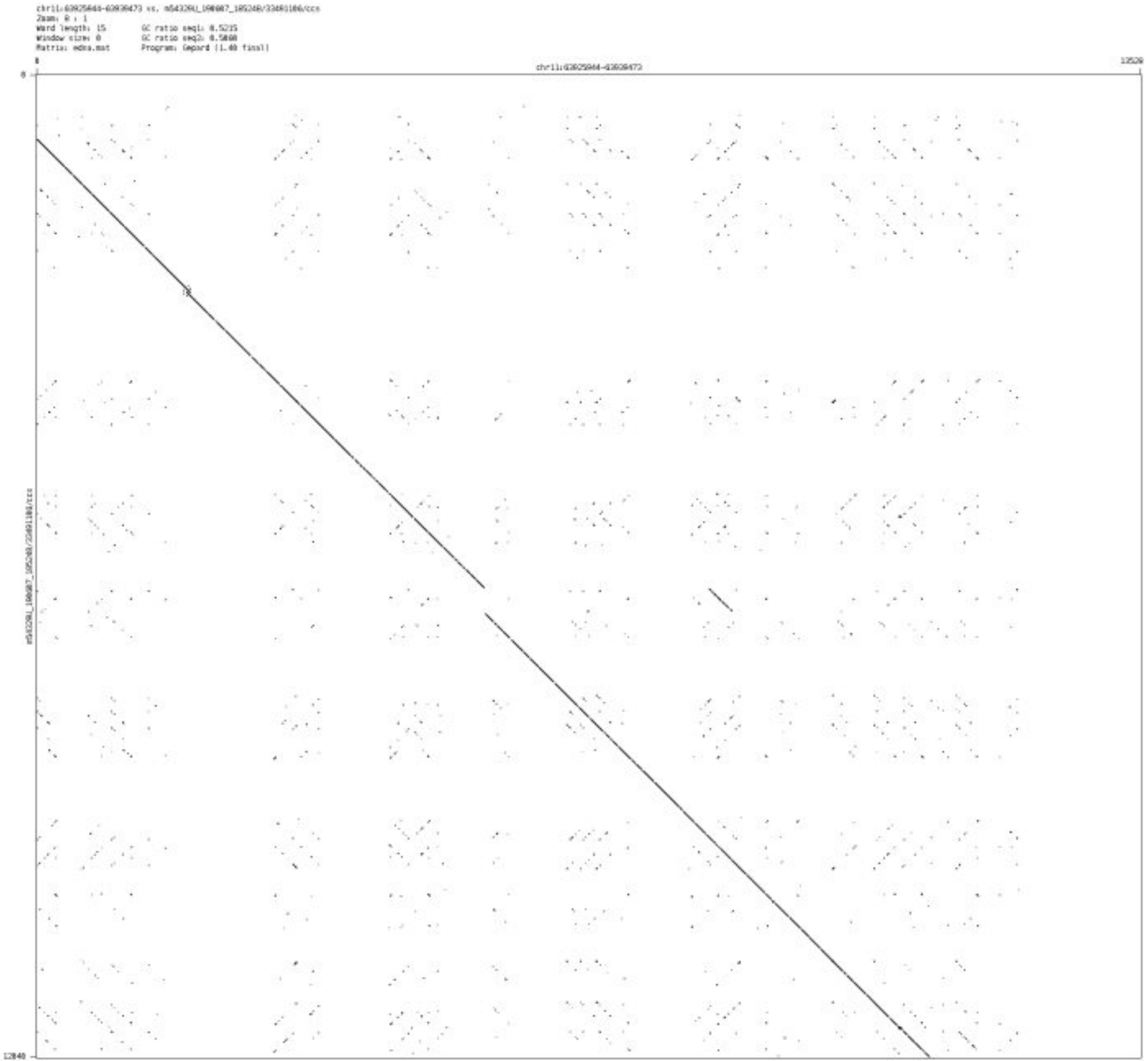

- chr11_99819490_99820566

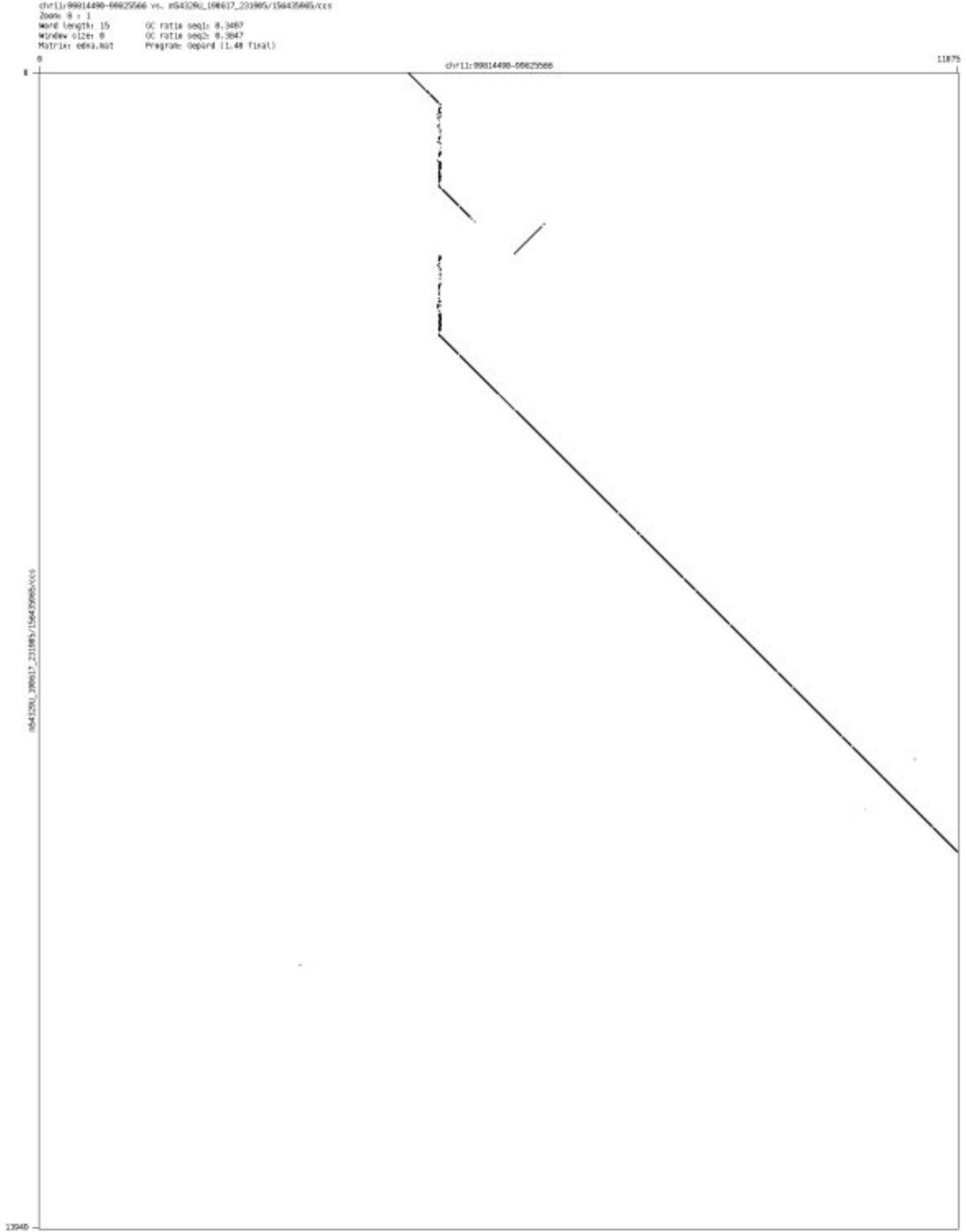

- chr12_25106390_25107673

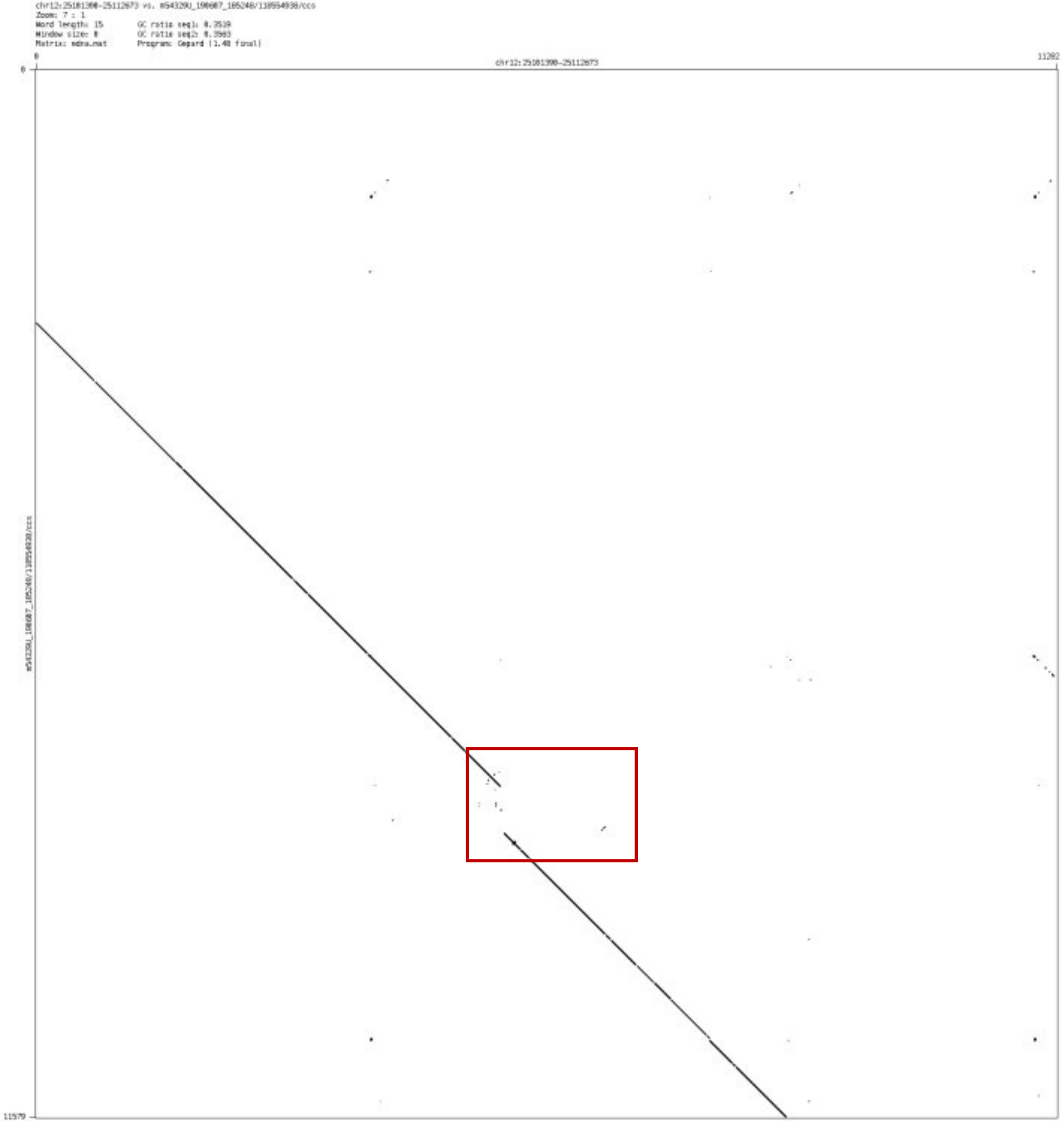

➢ chr12_40480640_40490425

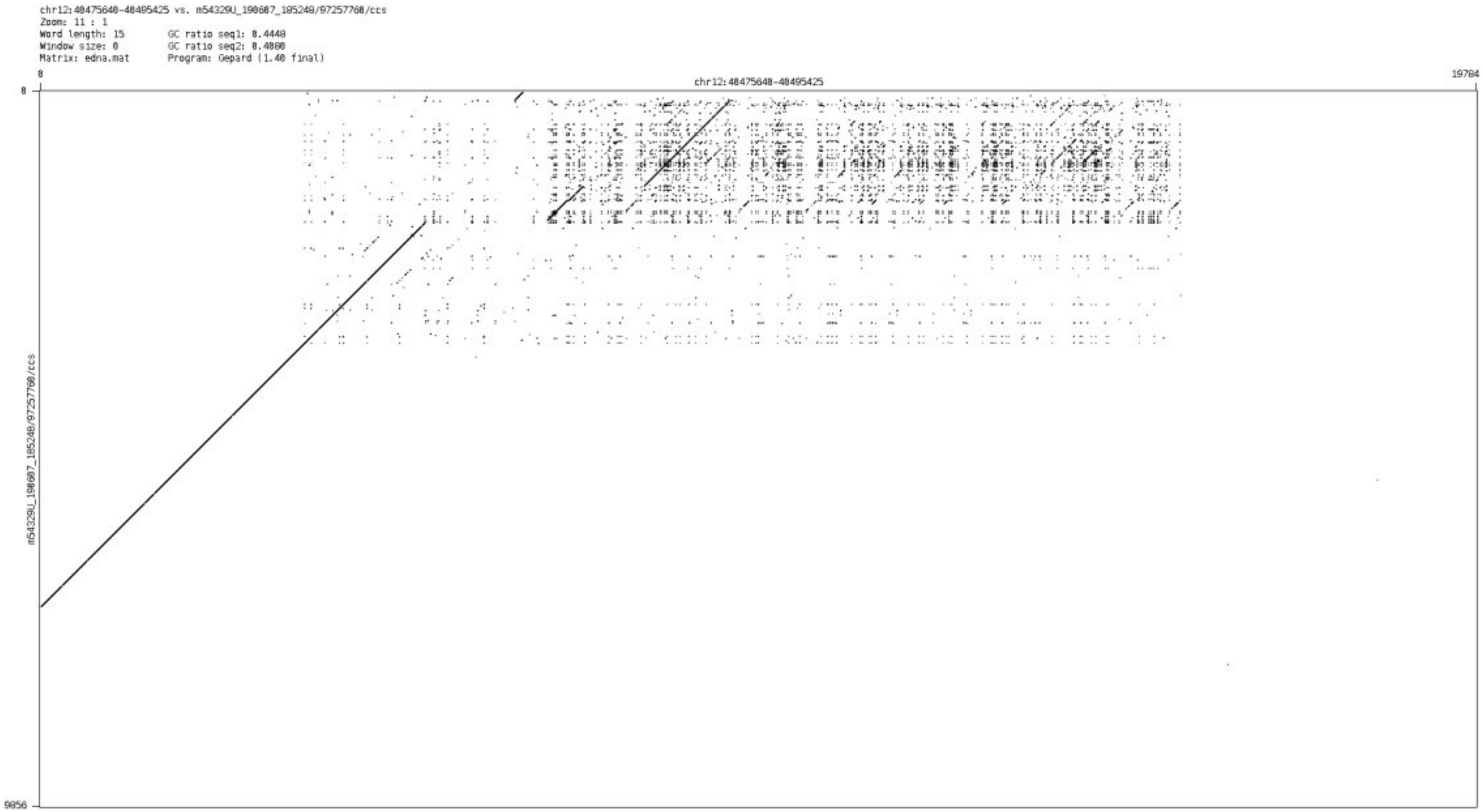

➢ chr12_65424697_65425168

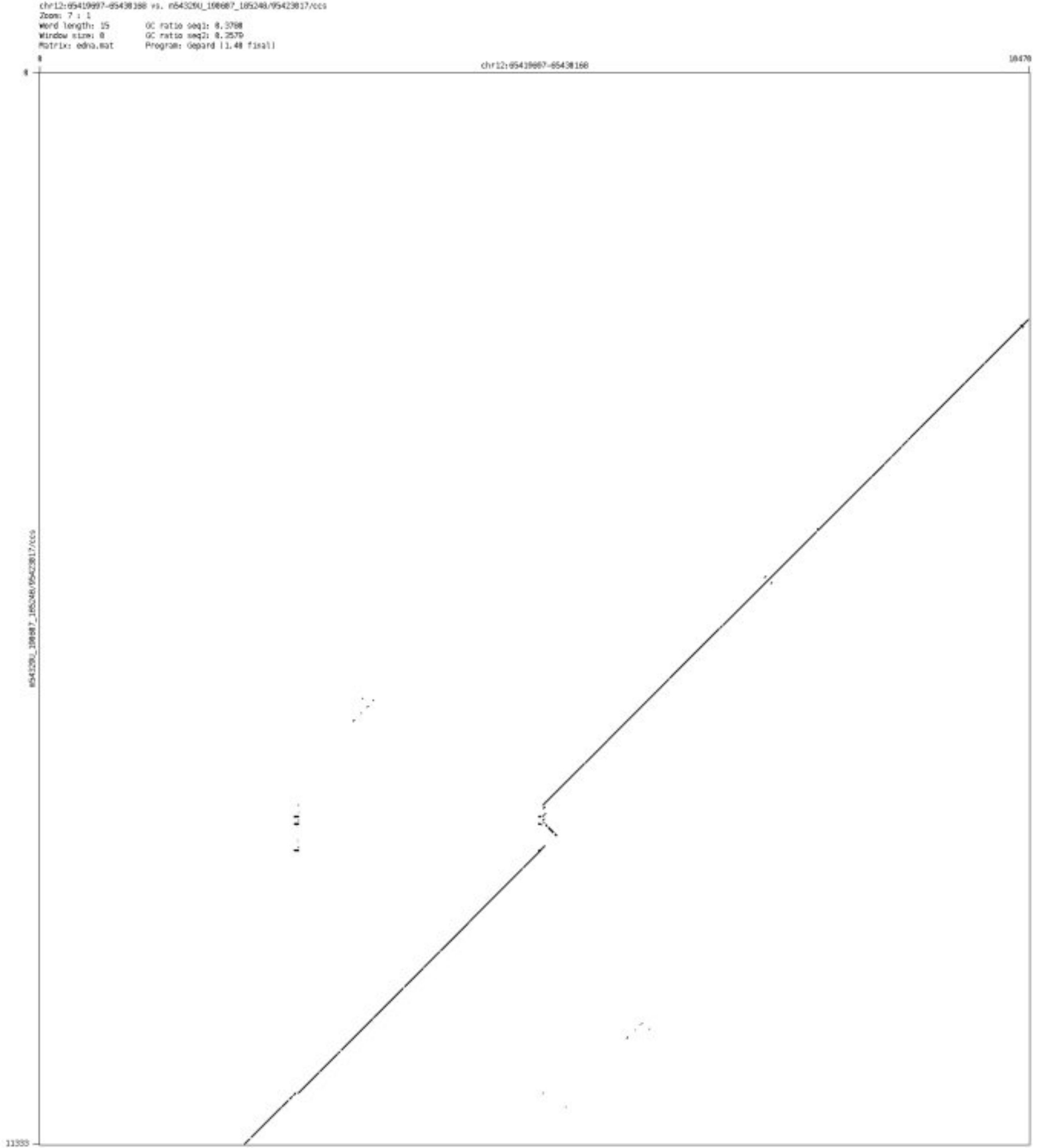

- ➢ chr12_71315482_71316928

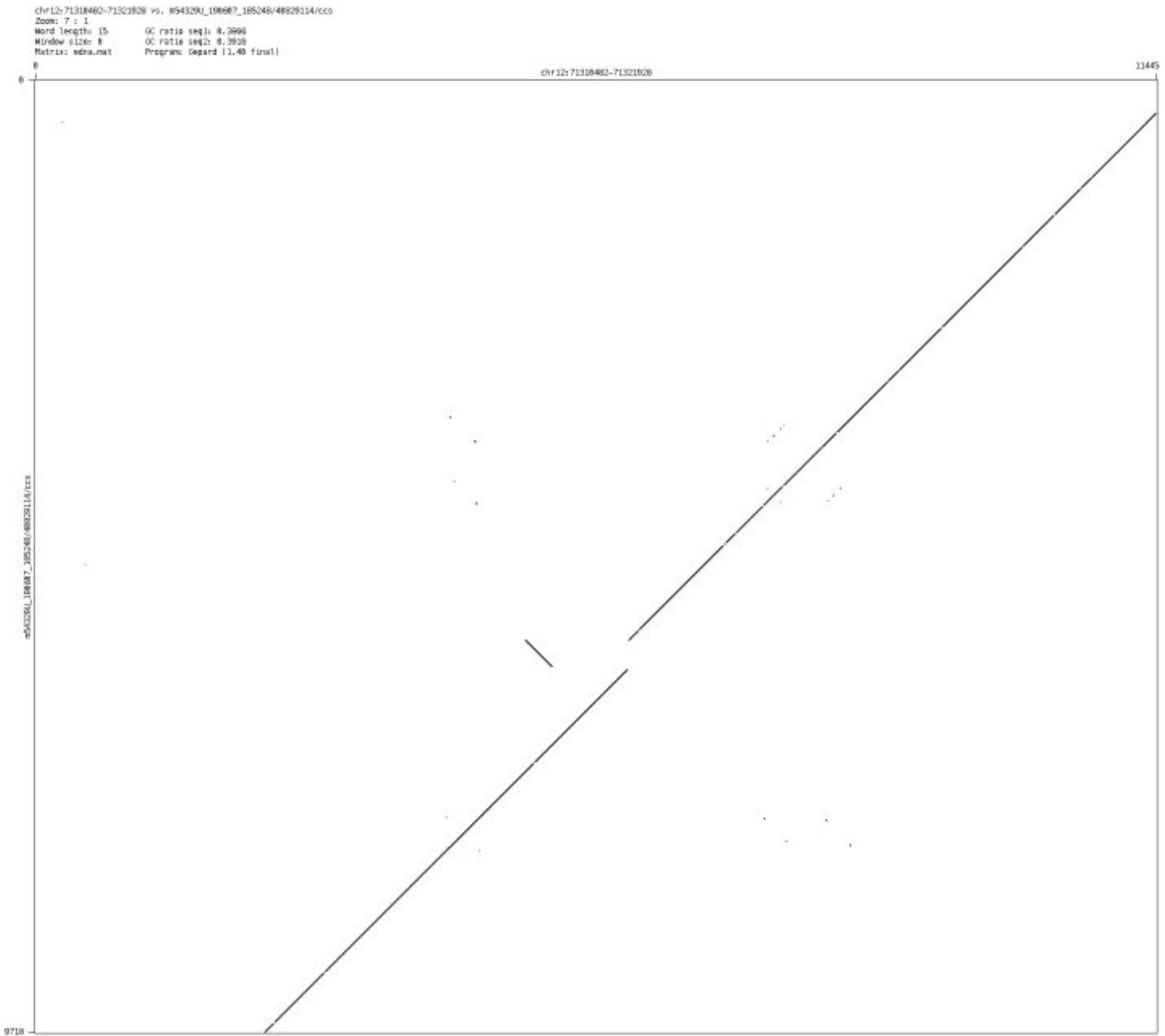

- chr12_77989900_77994324

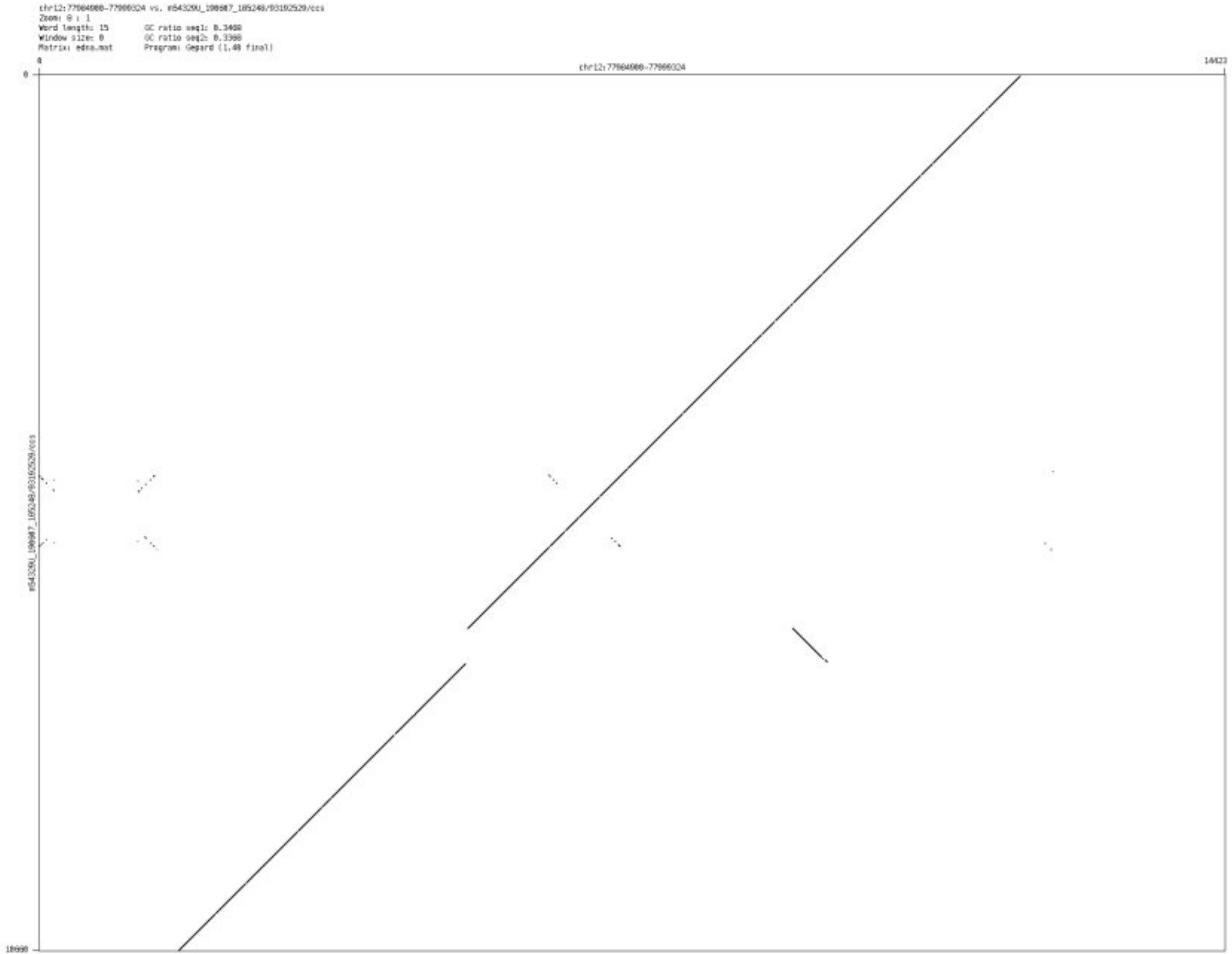

- ➢ chr12_86810684_86812040

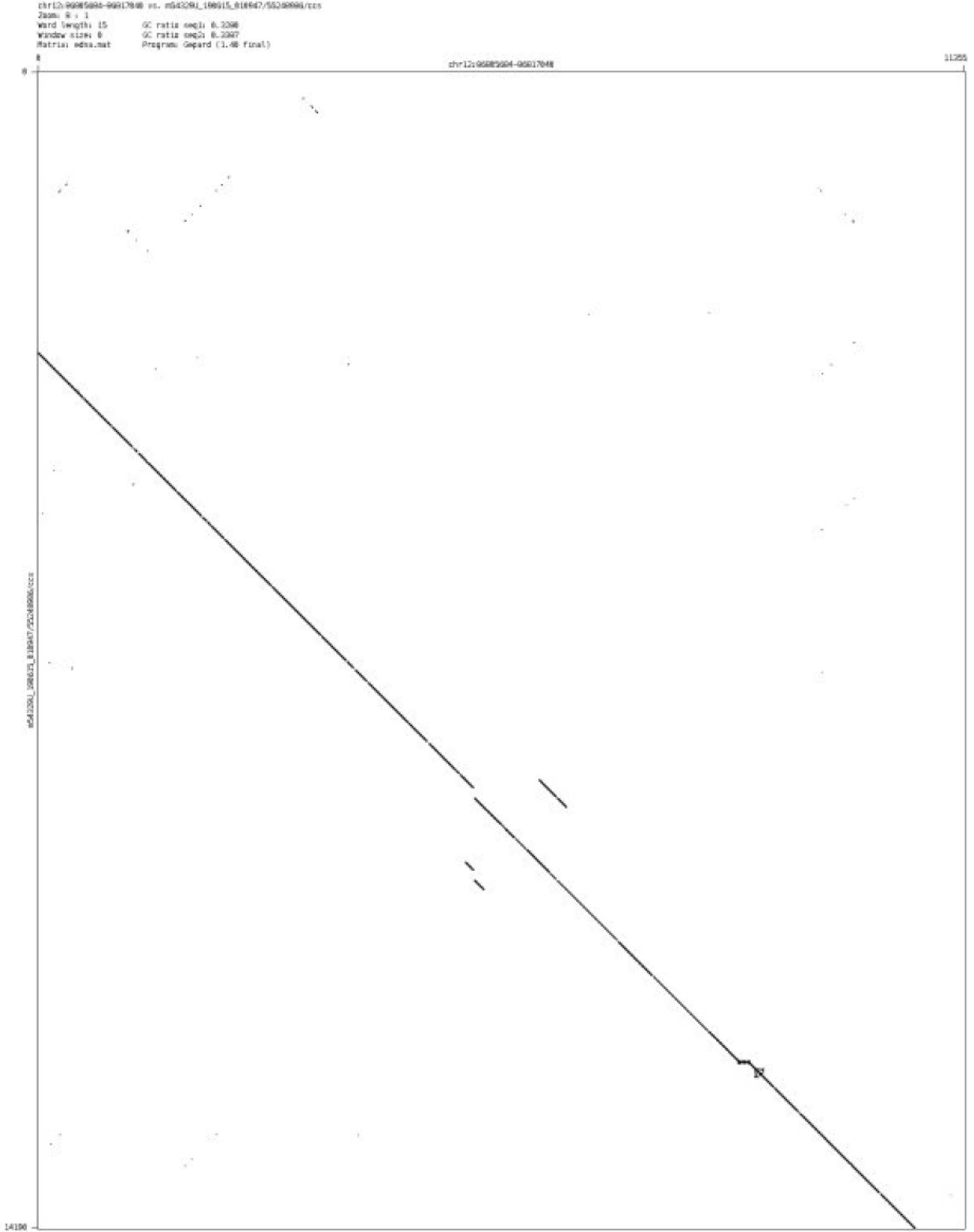

- chr12_115639648_115642432

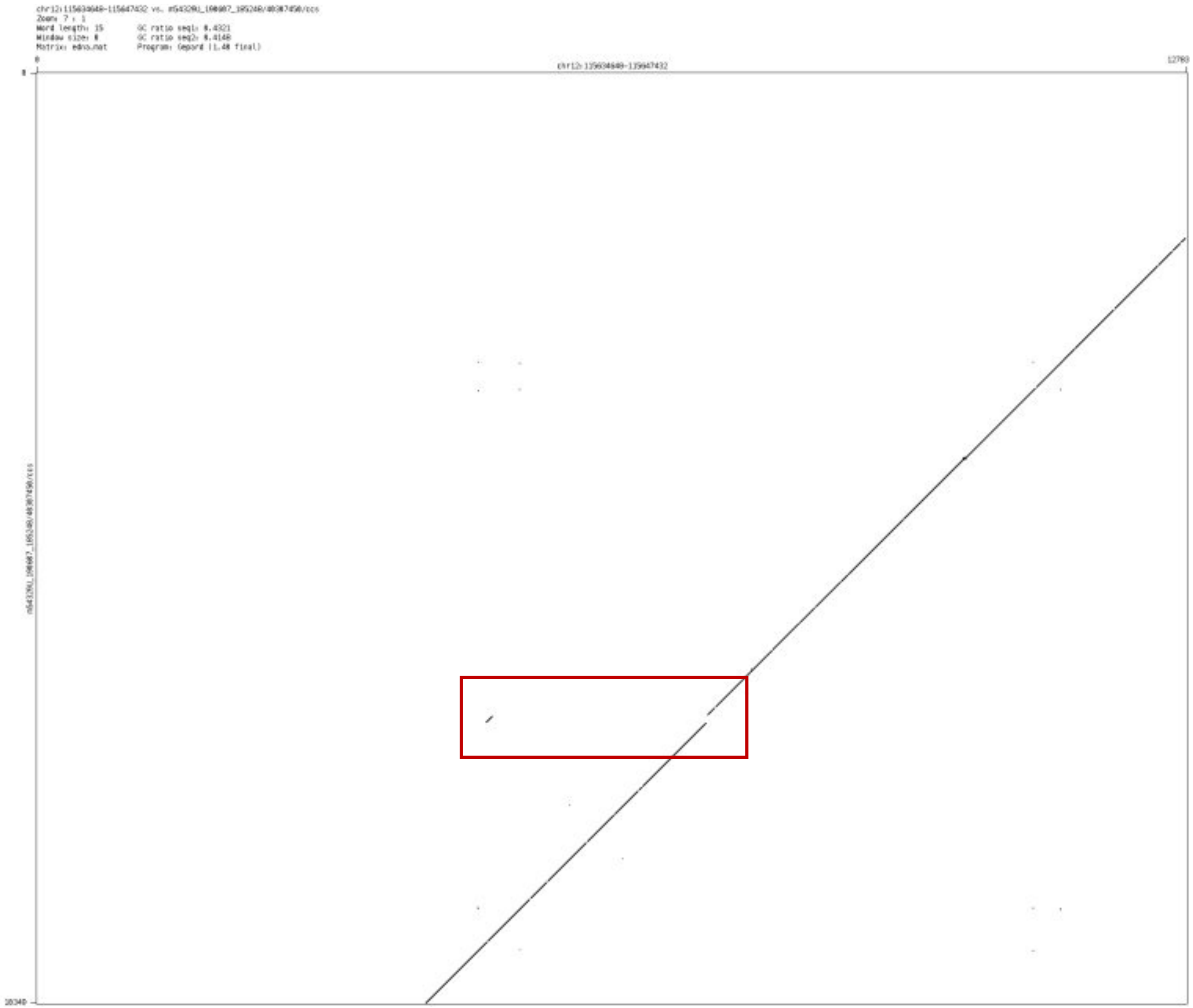

➢ chr12_127906184_127907400

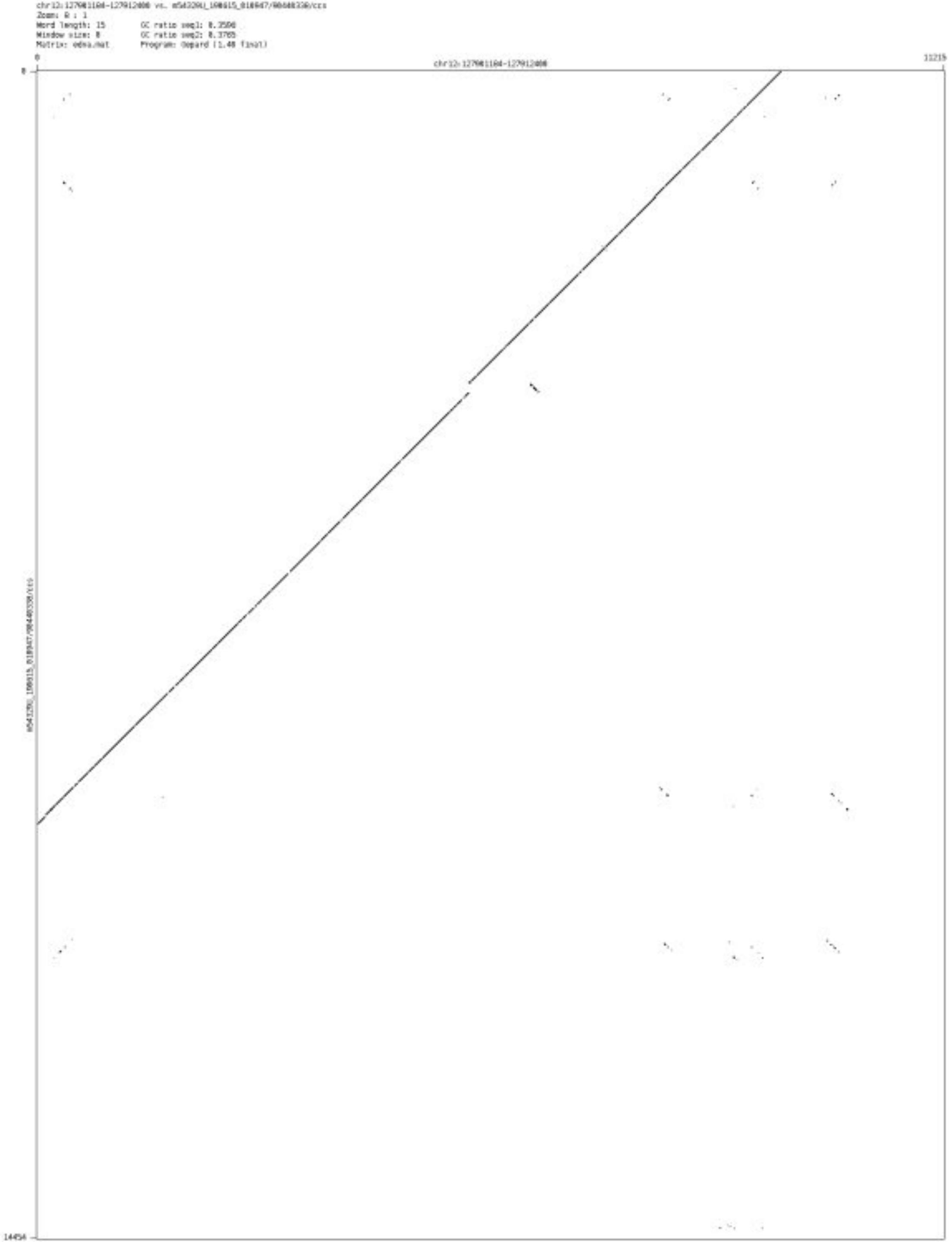

- chr13_74340759_74342810

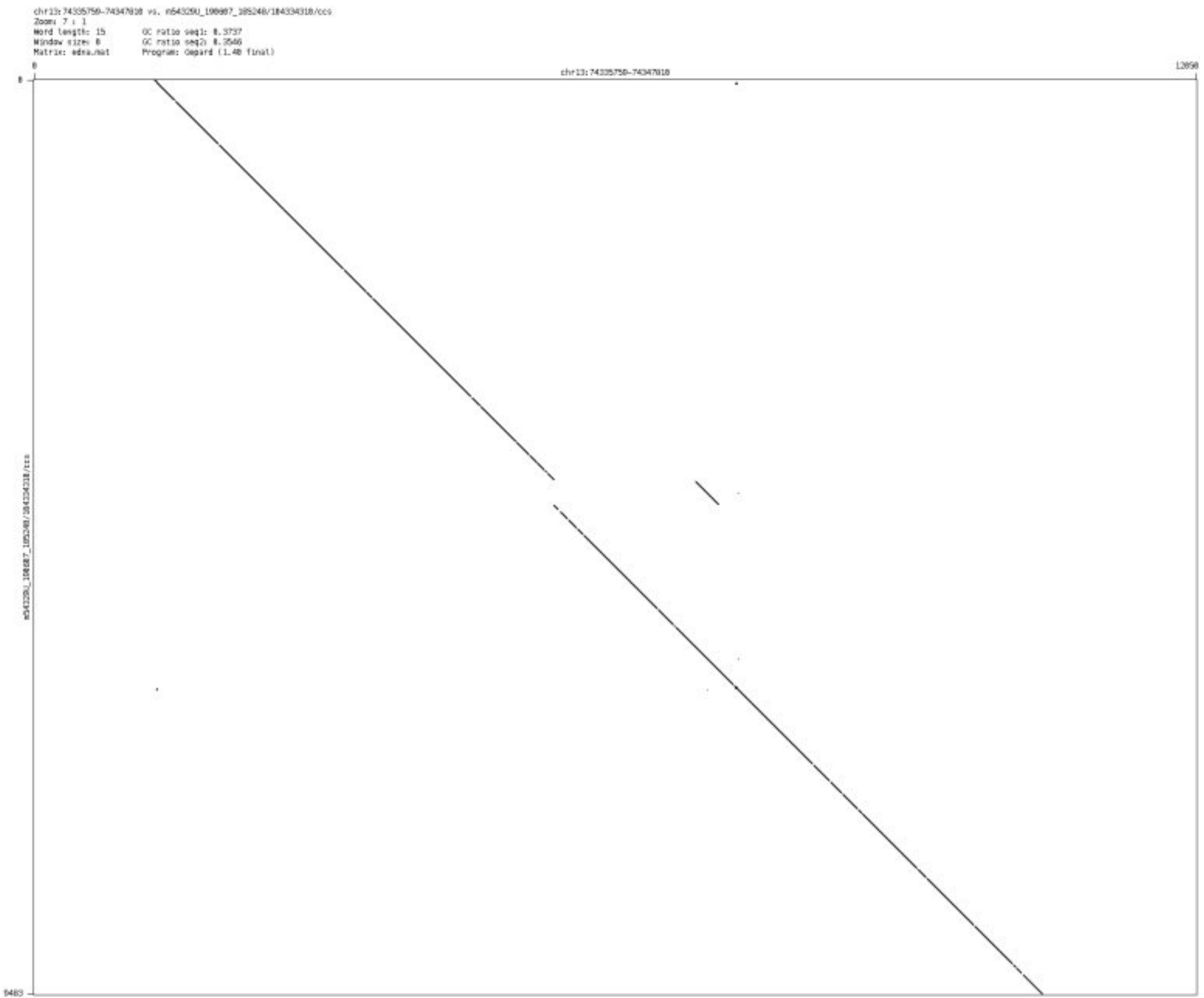

➢ chr13_88458439_88460627

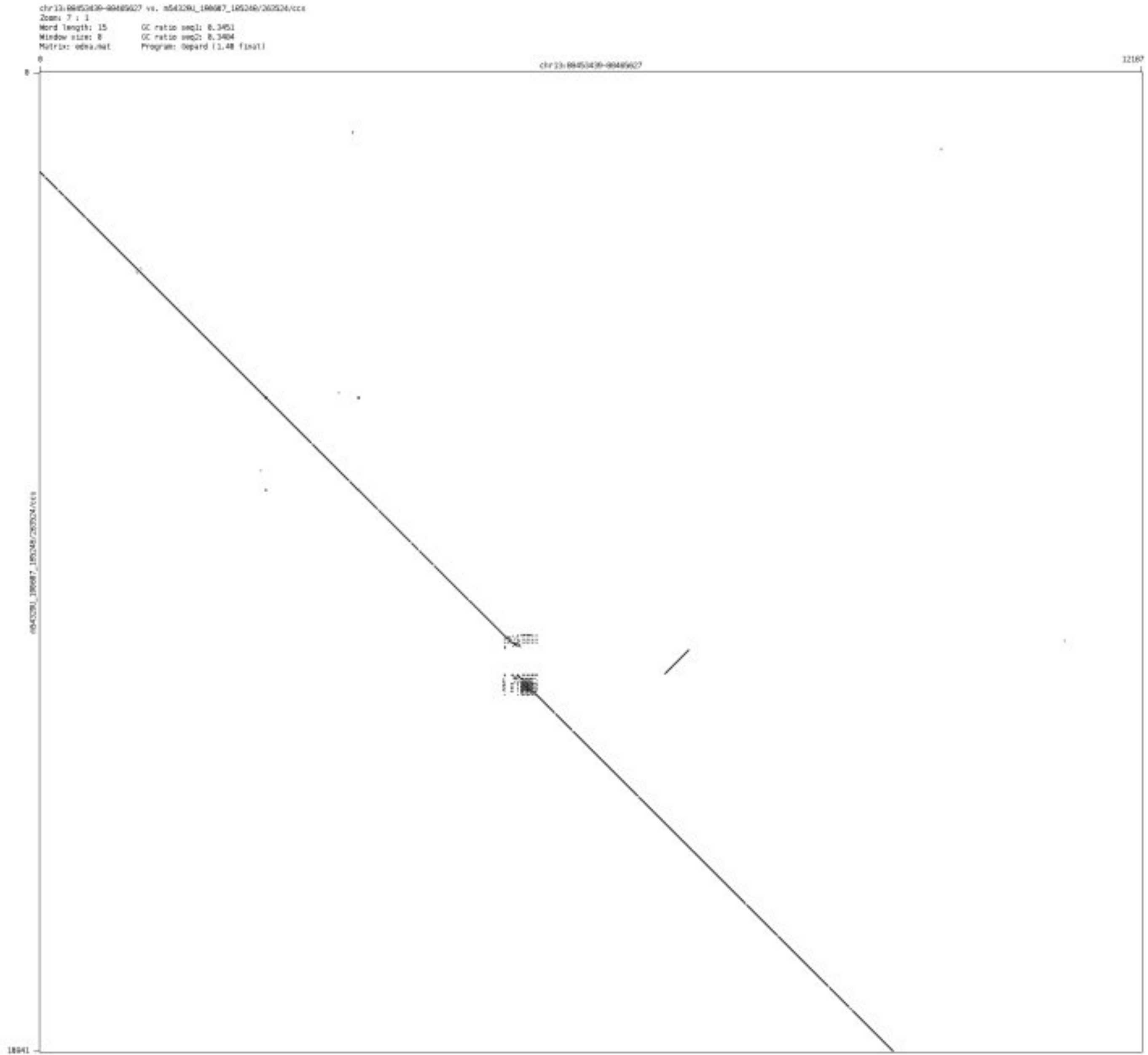

➢ chr14_25141909_25144901

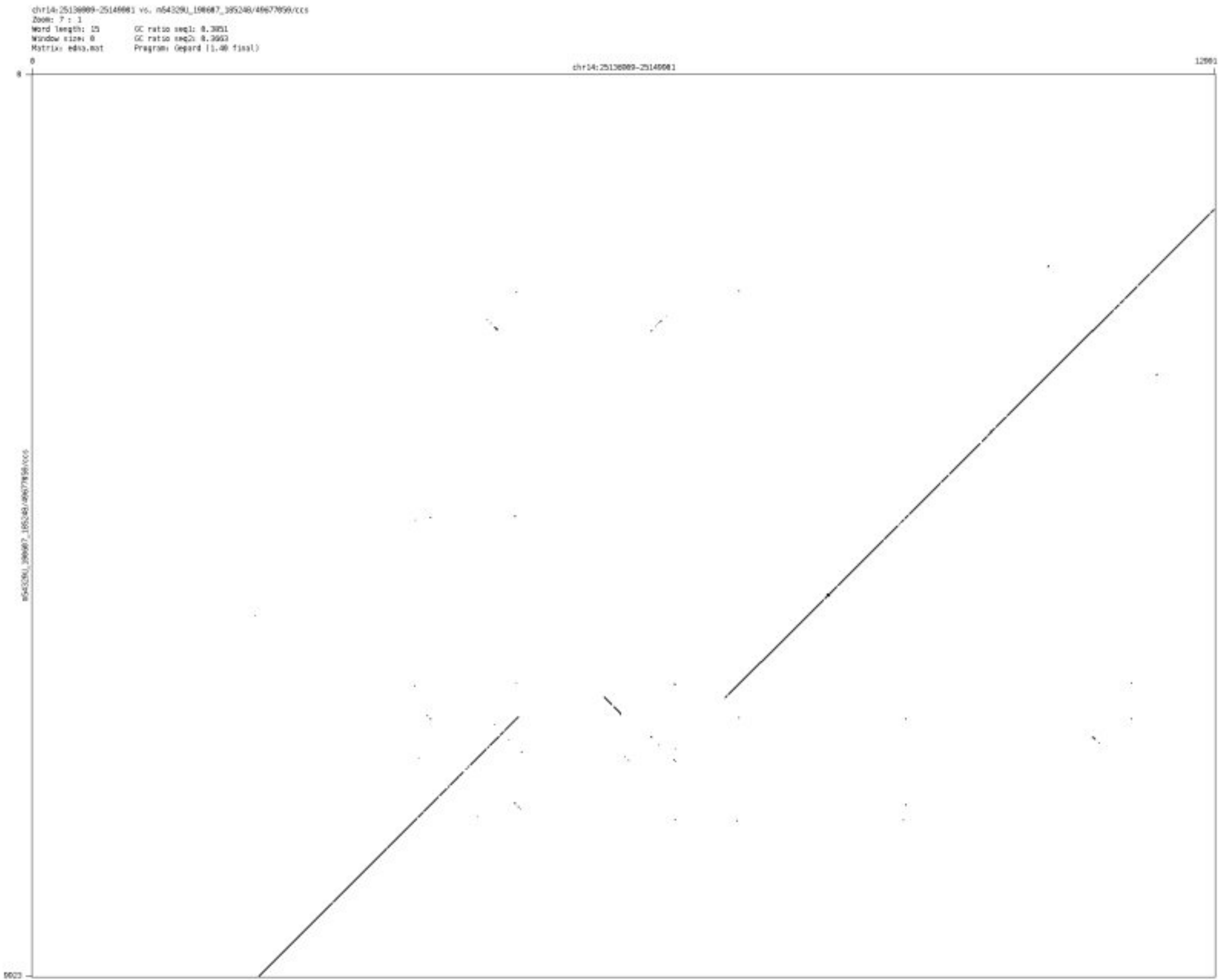

- chr14_39644680_39645896

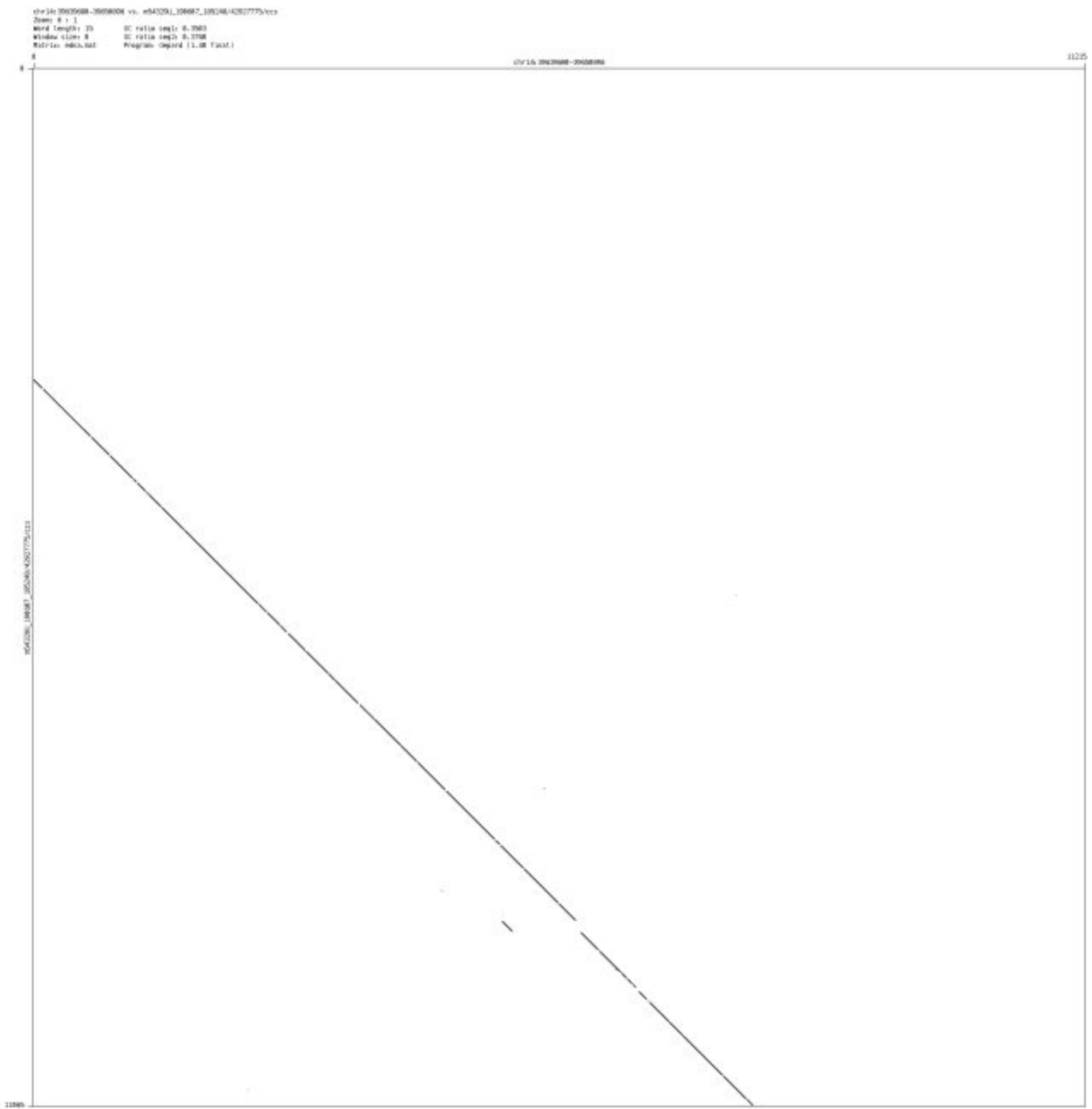

- chr14_40906700_40907954

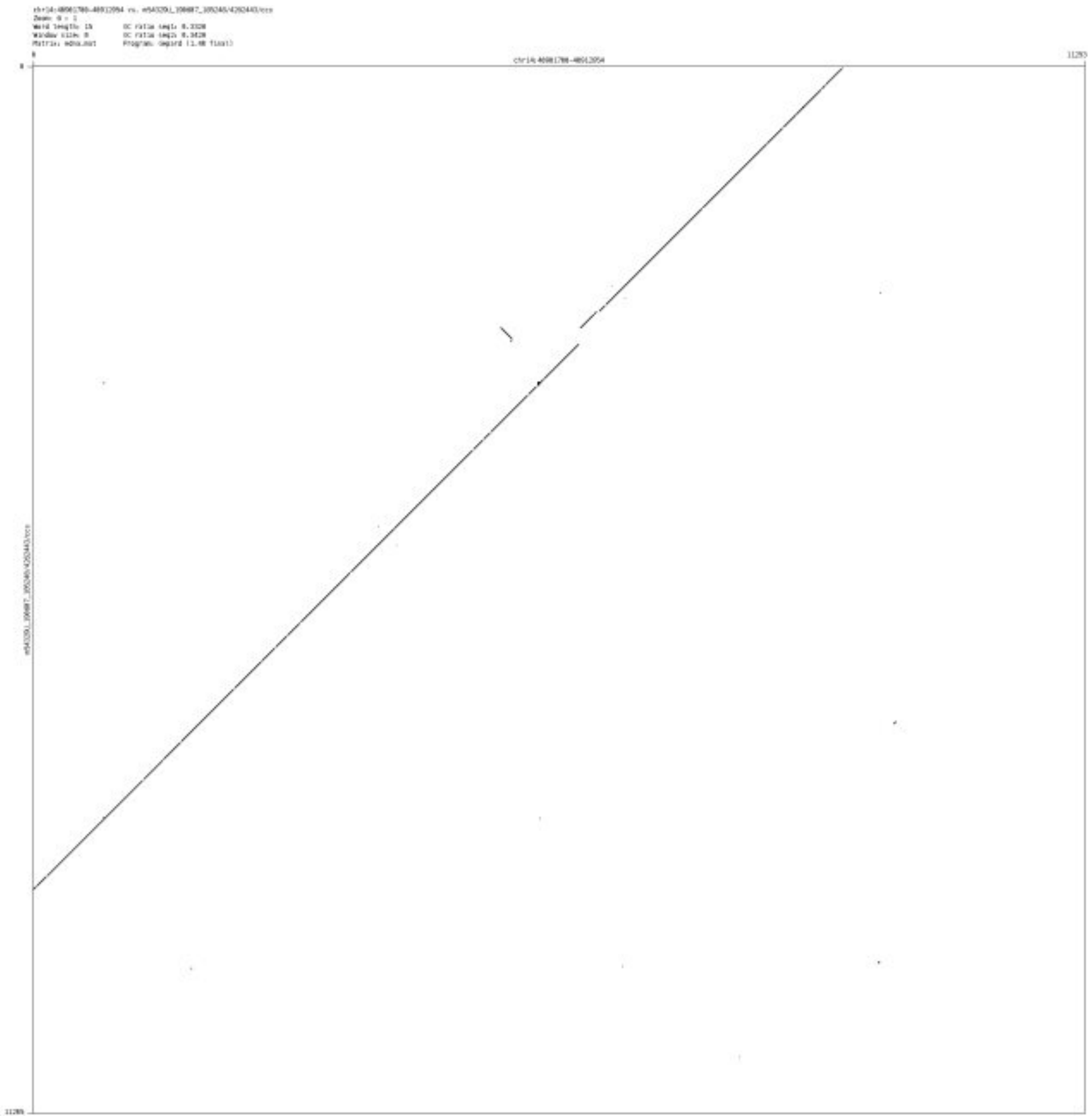

➢ chr14_47854498_47856278

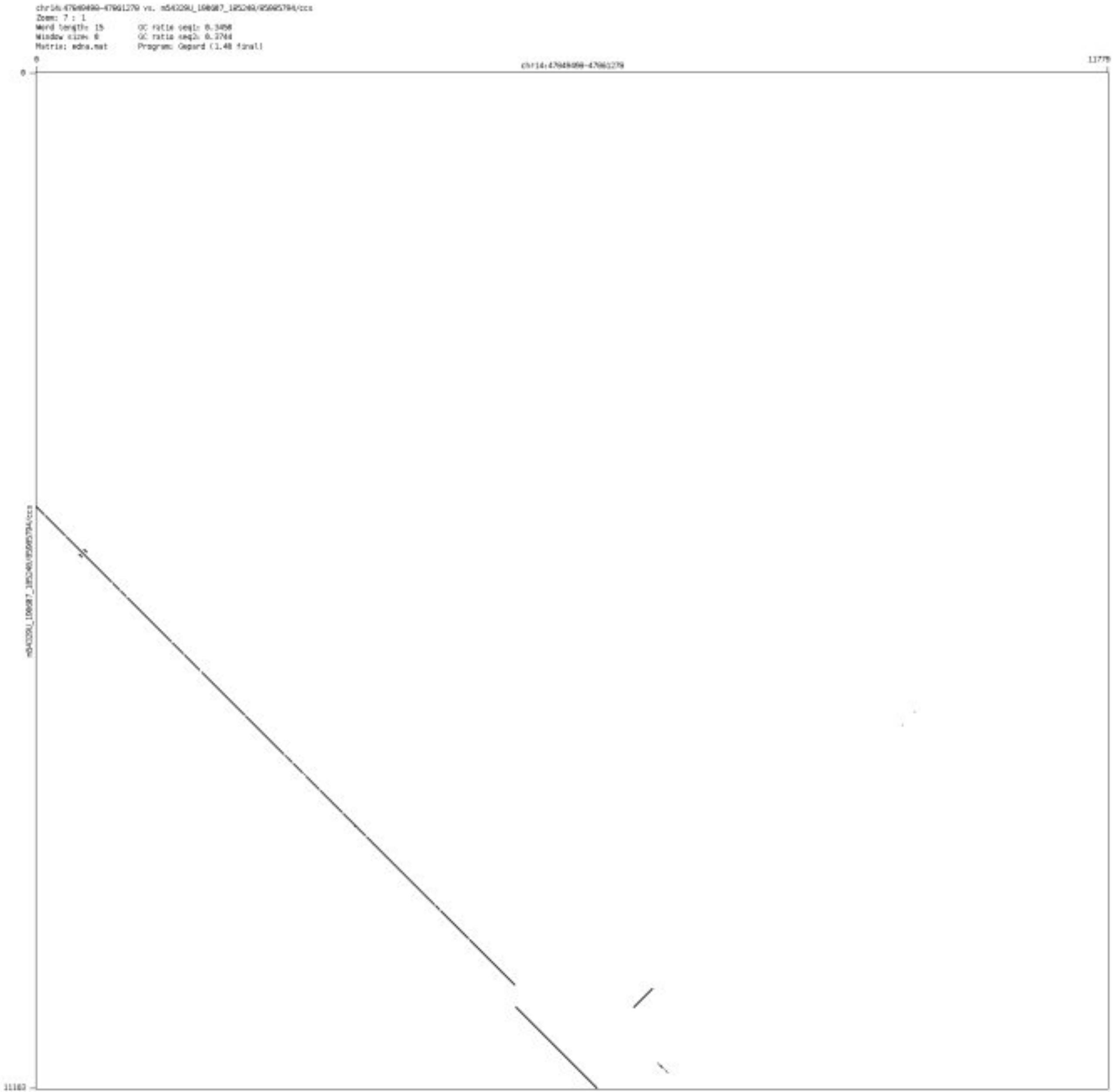

- chr14_65375836_65376948

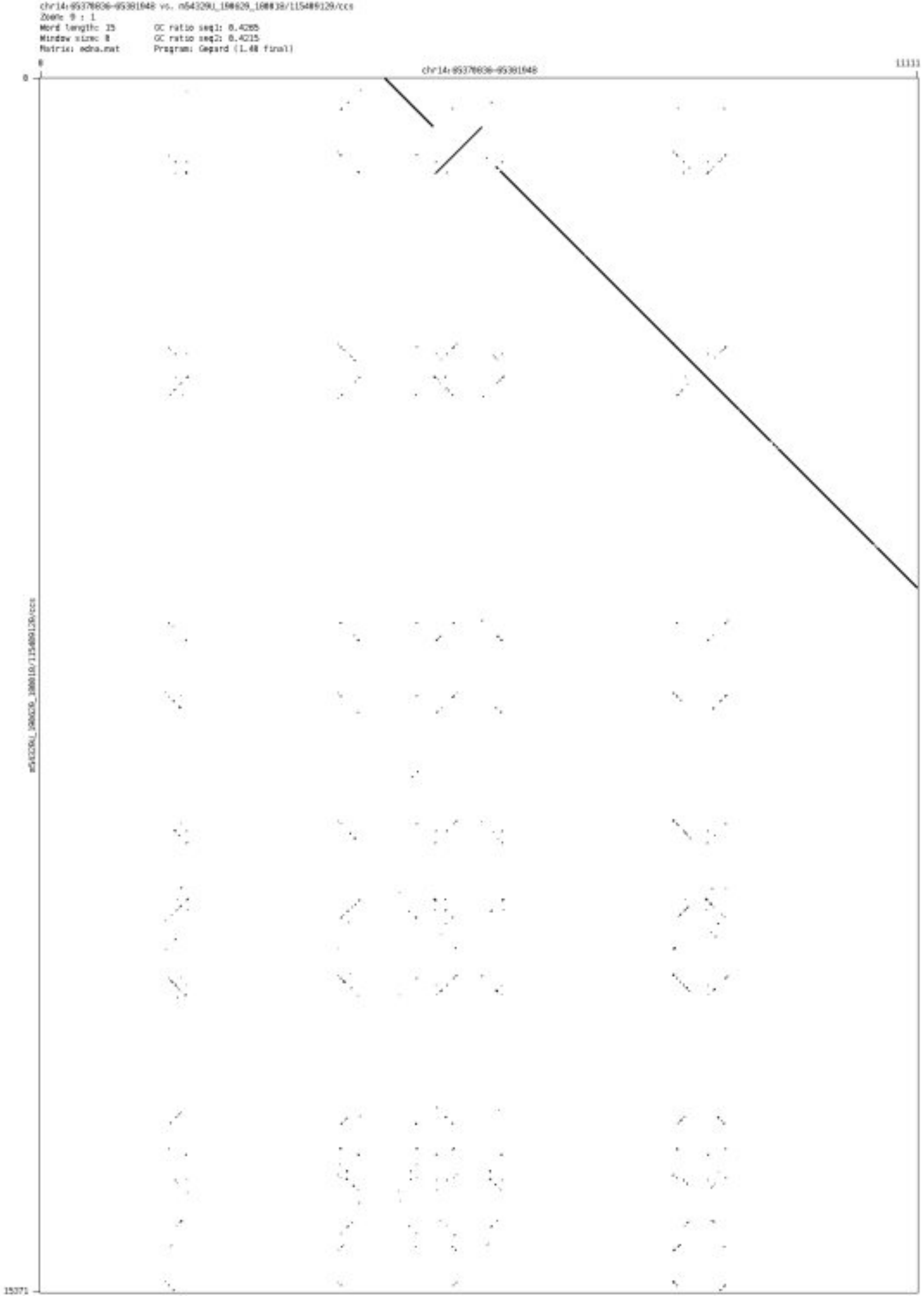

- chr15_37003075_37005371

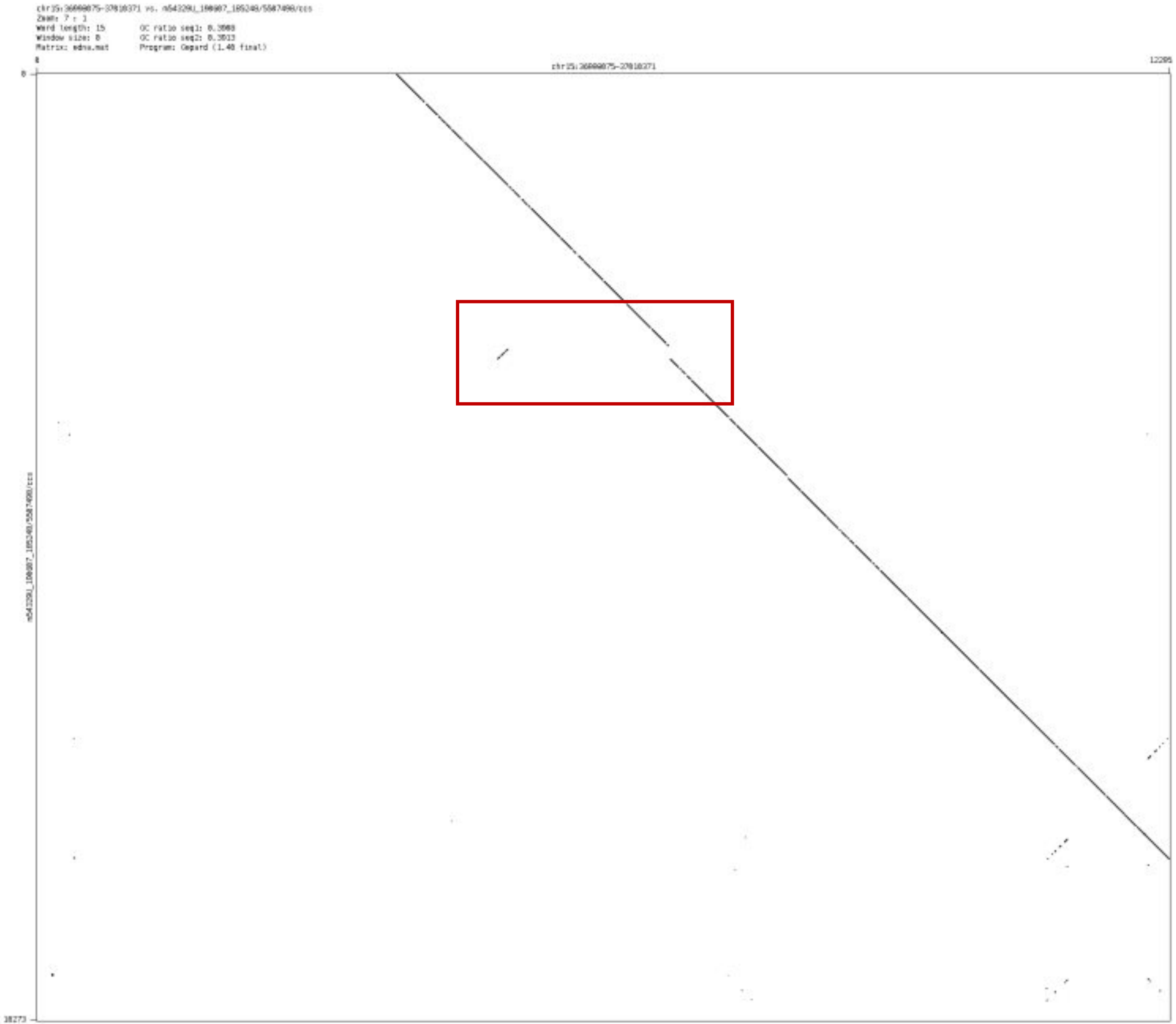

- chr15_91437836_91446359

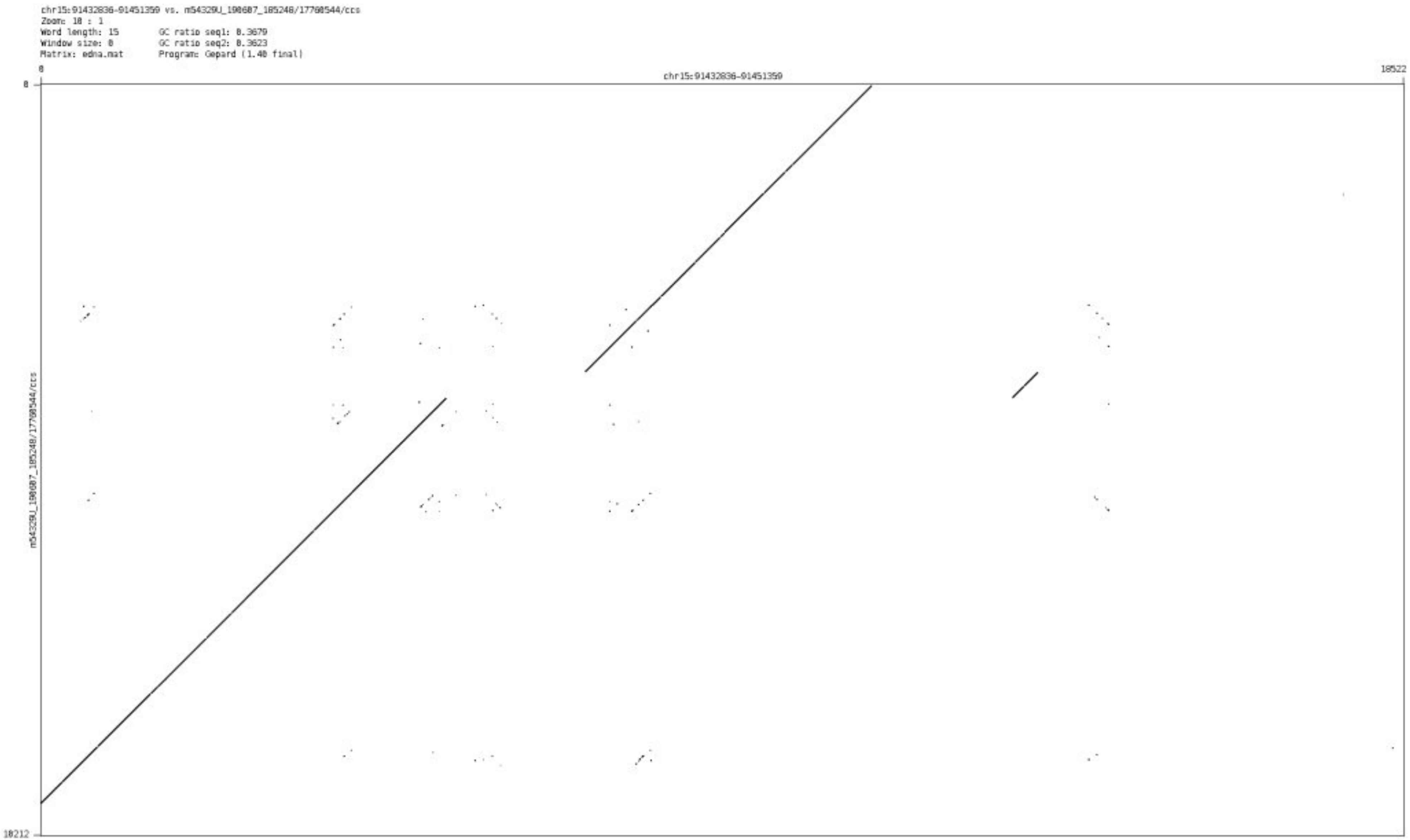

➢ chr15_95148202_95149527

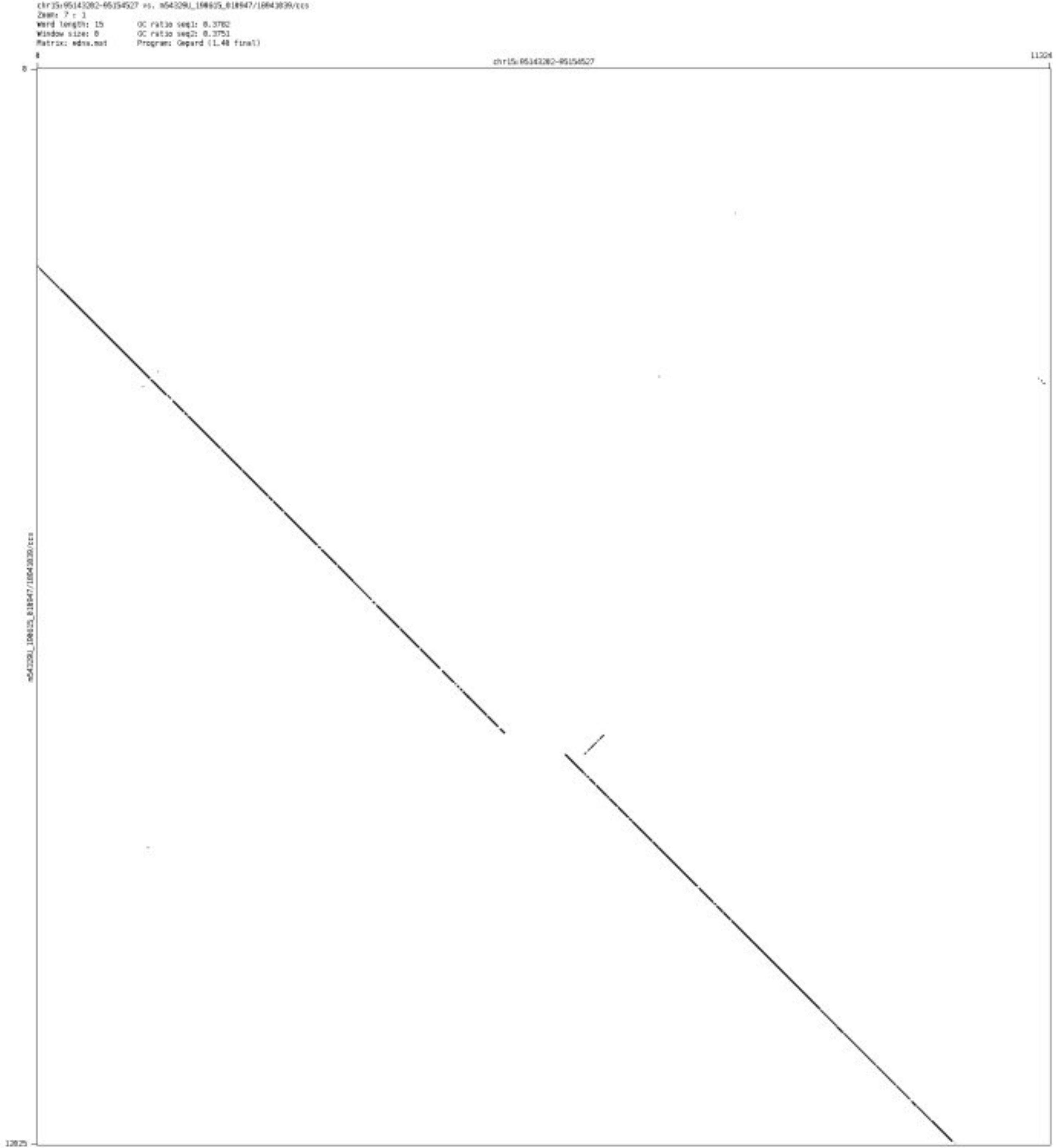

➢ chr16_48871190_48872586

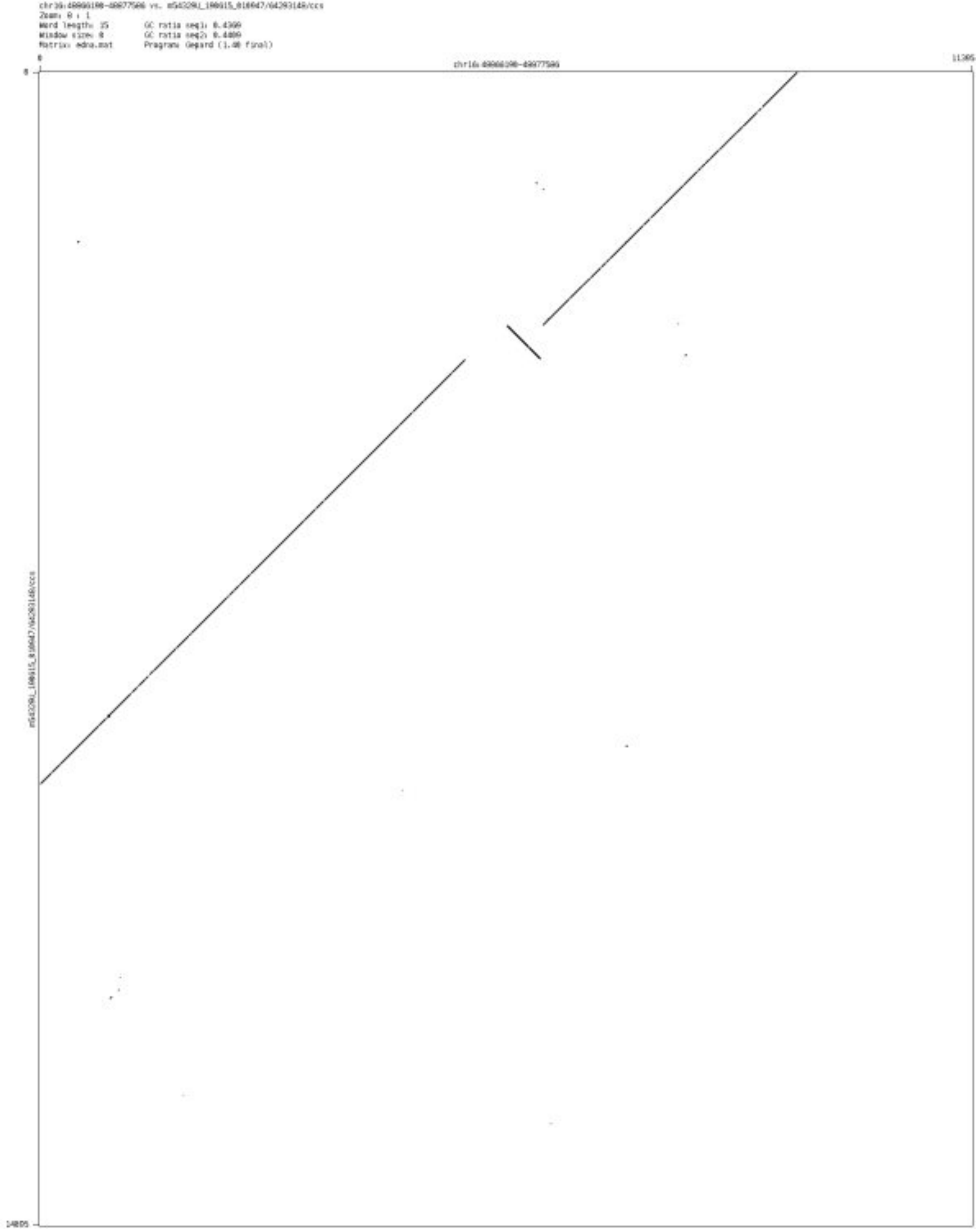

➢ chr16_69727976_69729327

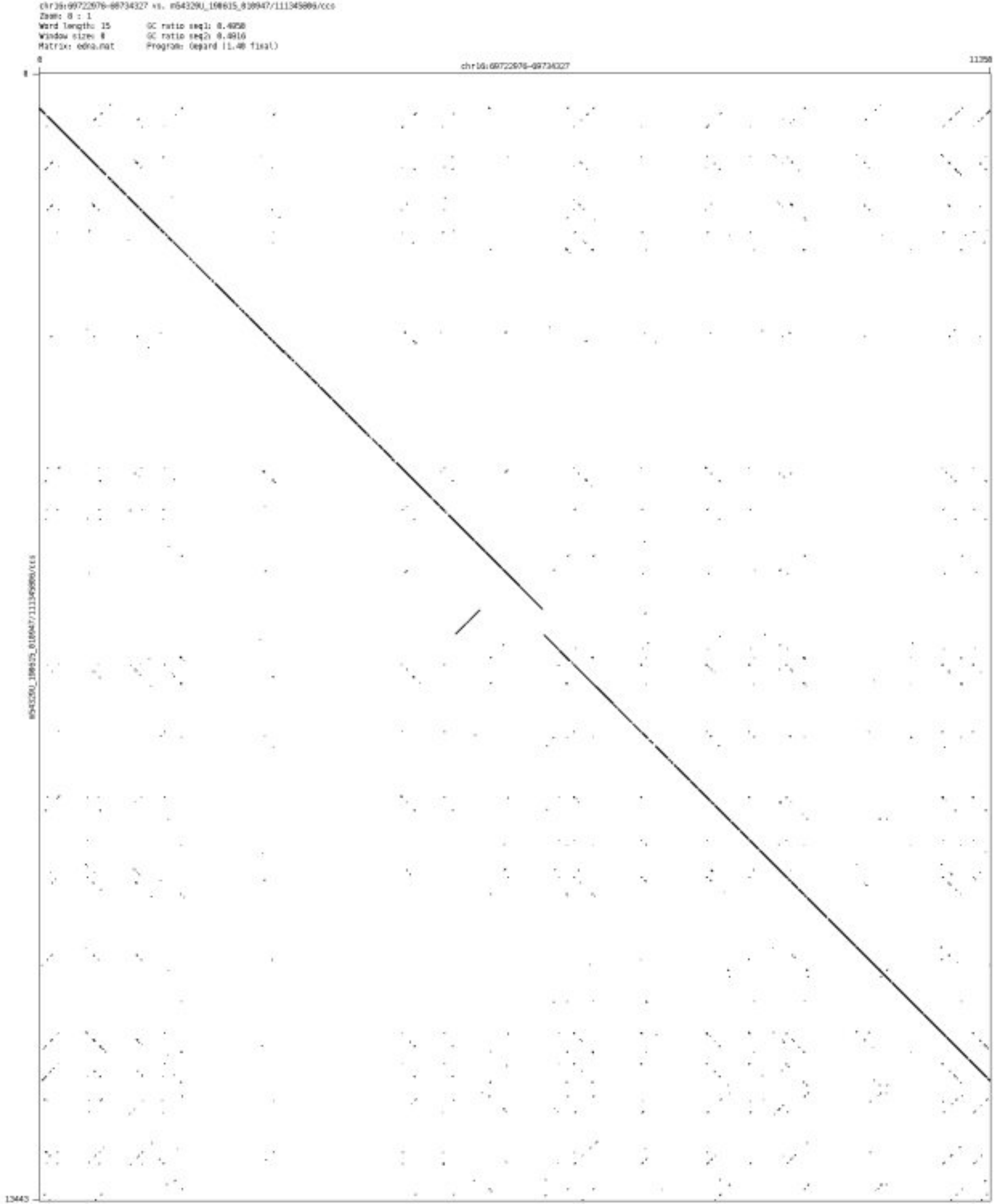

➢ chr16_78004459_78007456

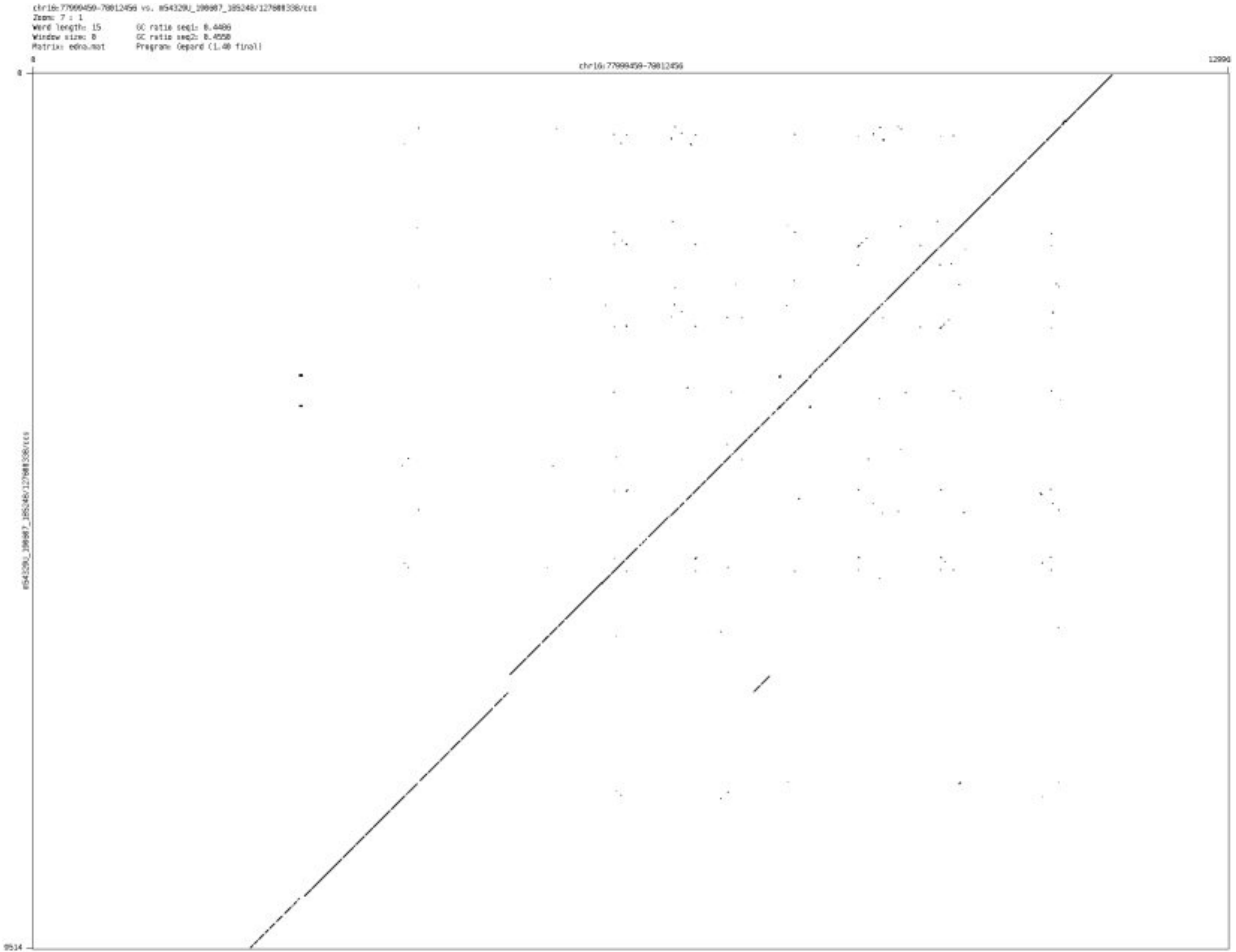

- chr17_34854438_34855851

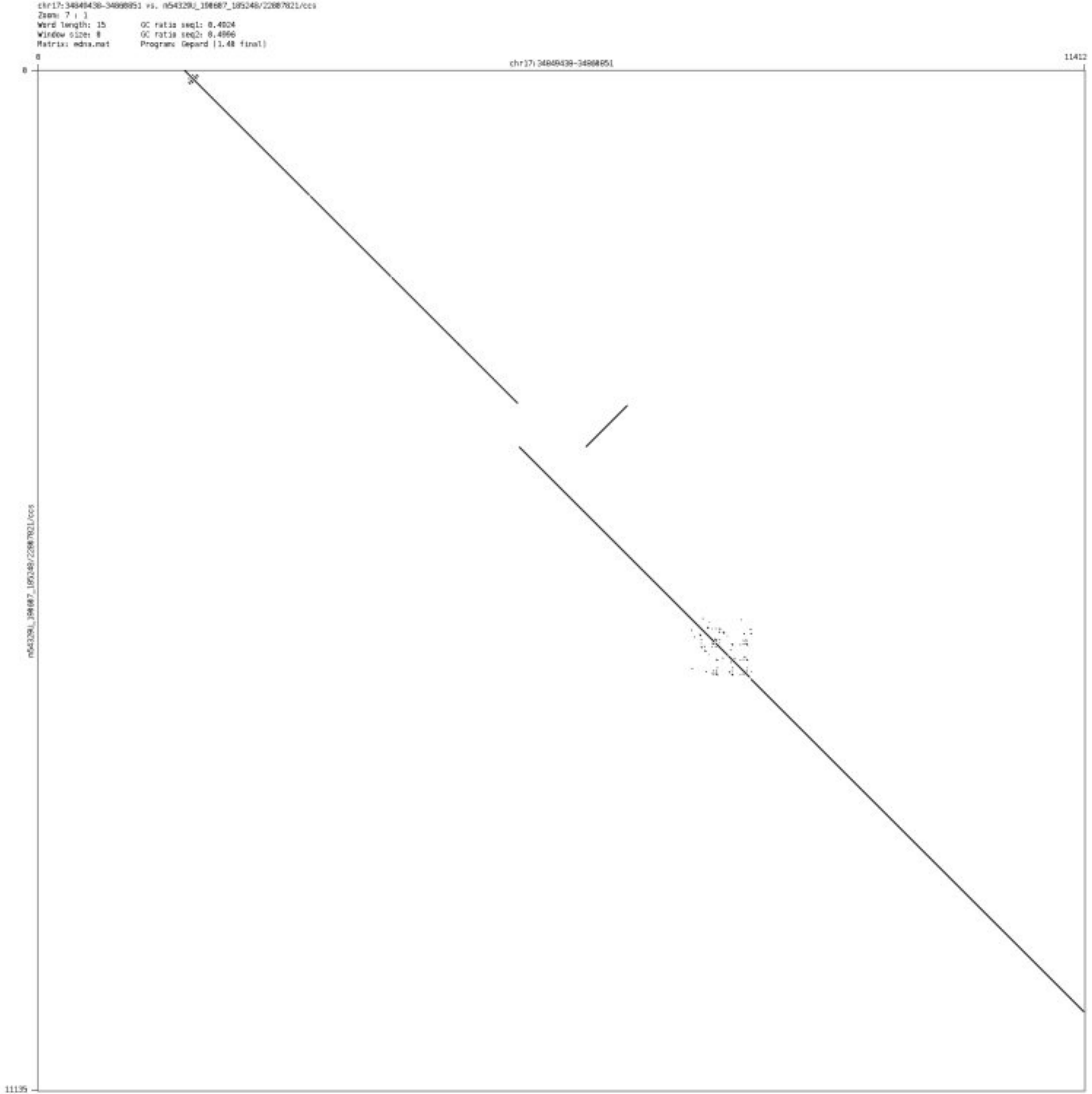

➢ chr17_48538270_48540171

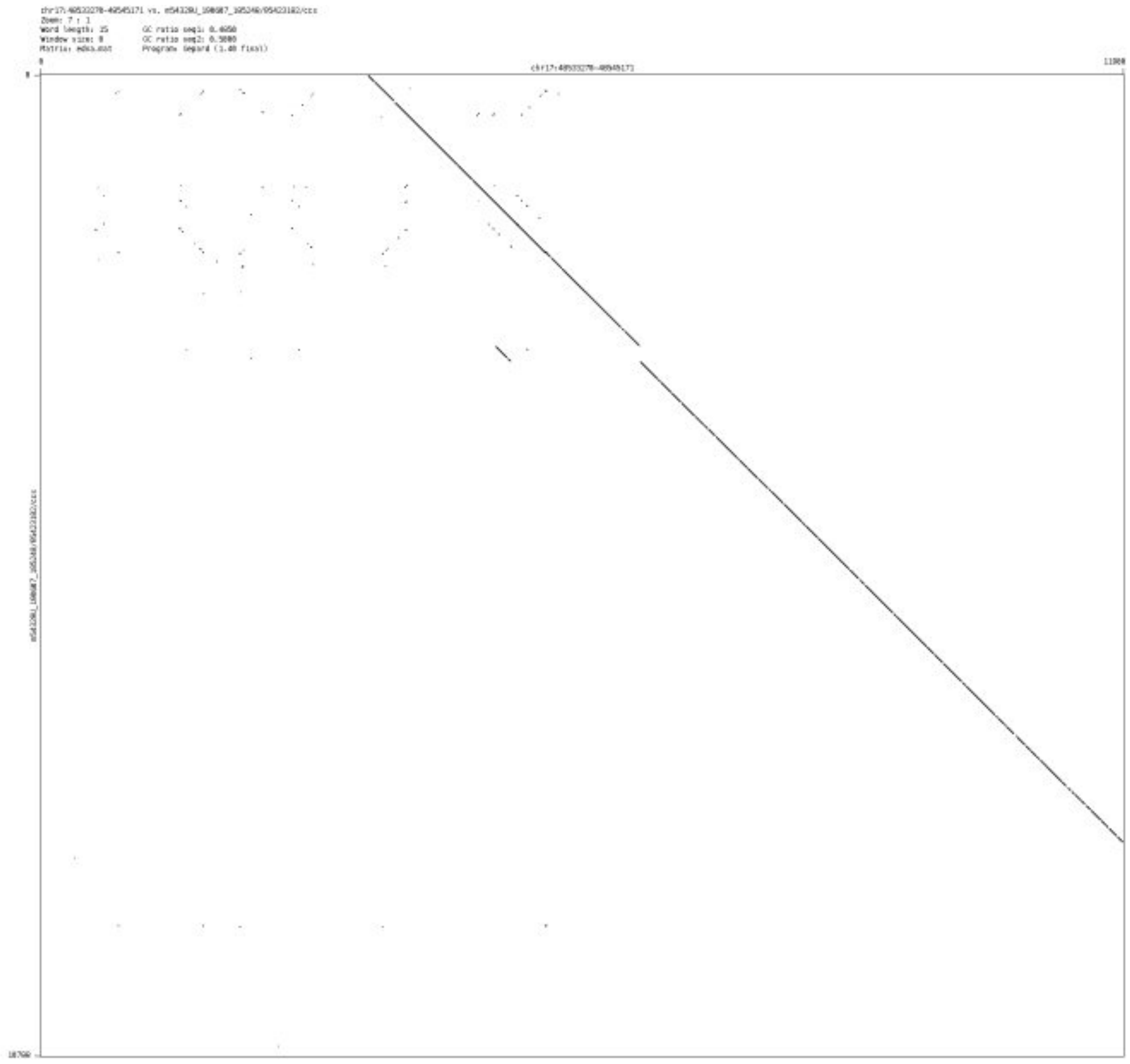

- ➢ chr17_73219884_73221595

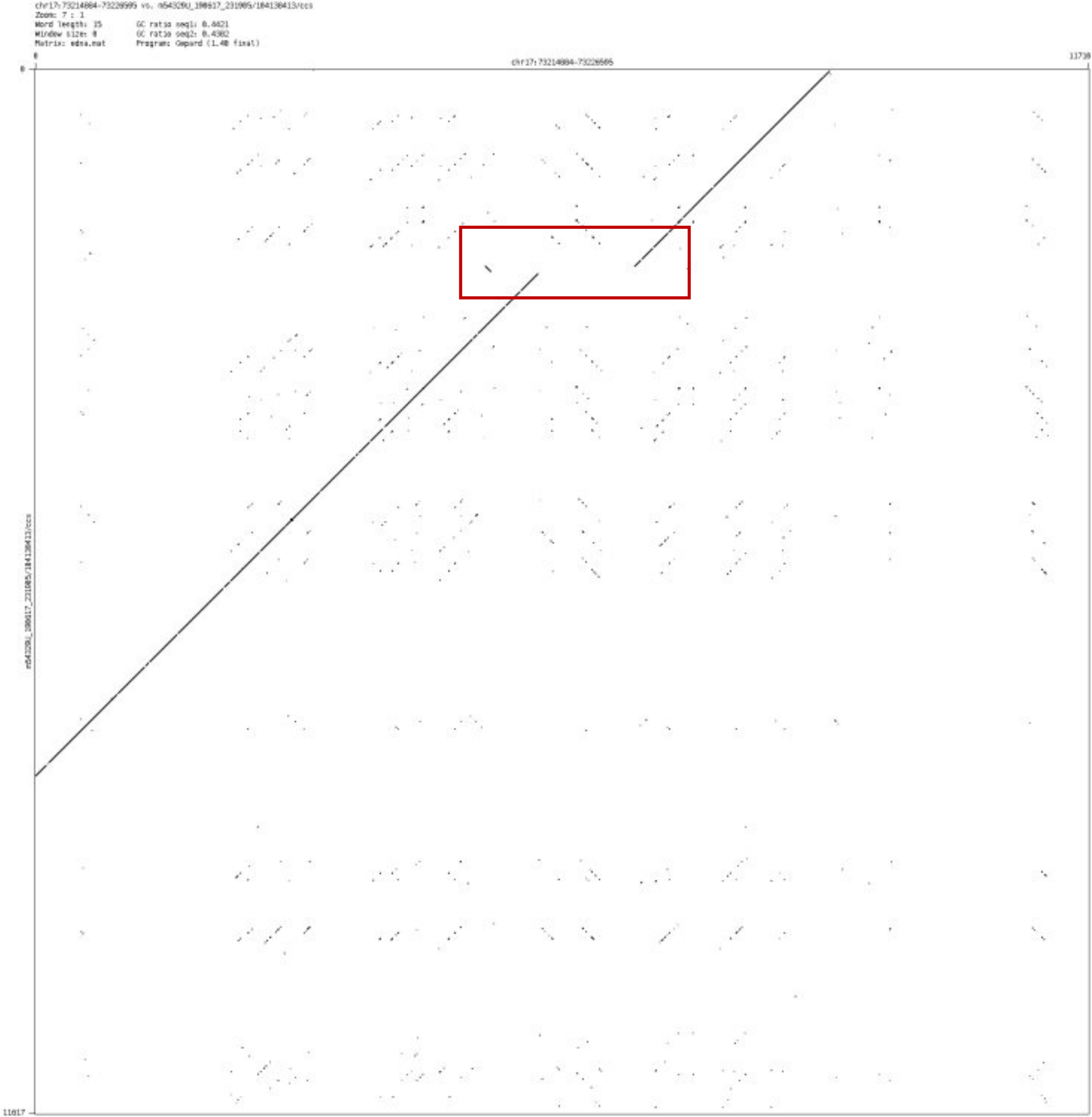

- chr18_11508929_11511476

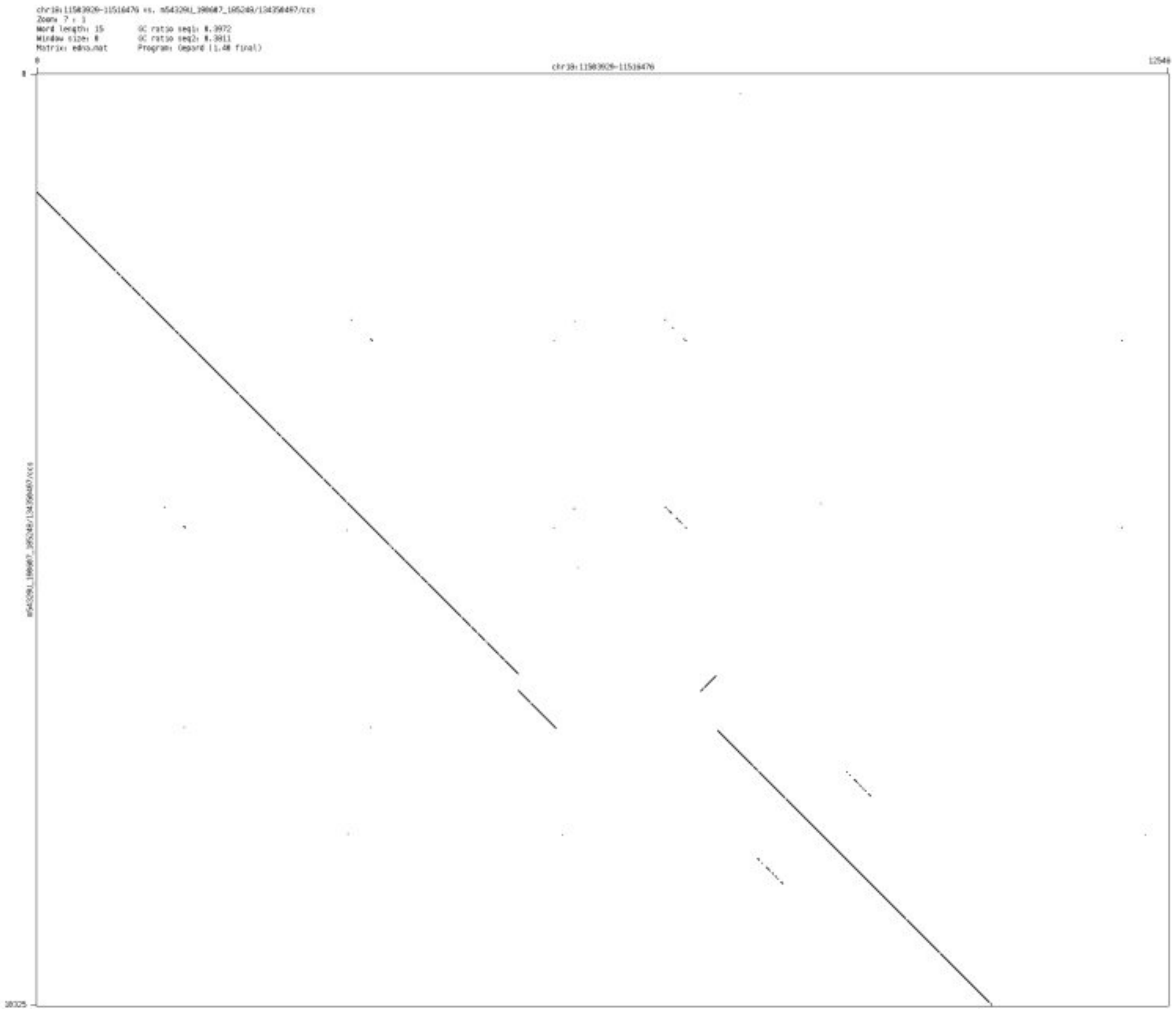

➢ chr18_72044575_72045937

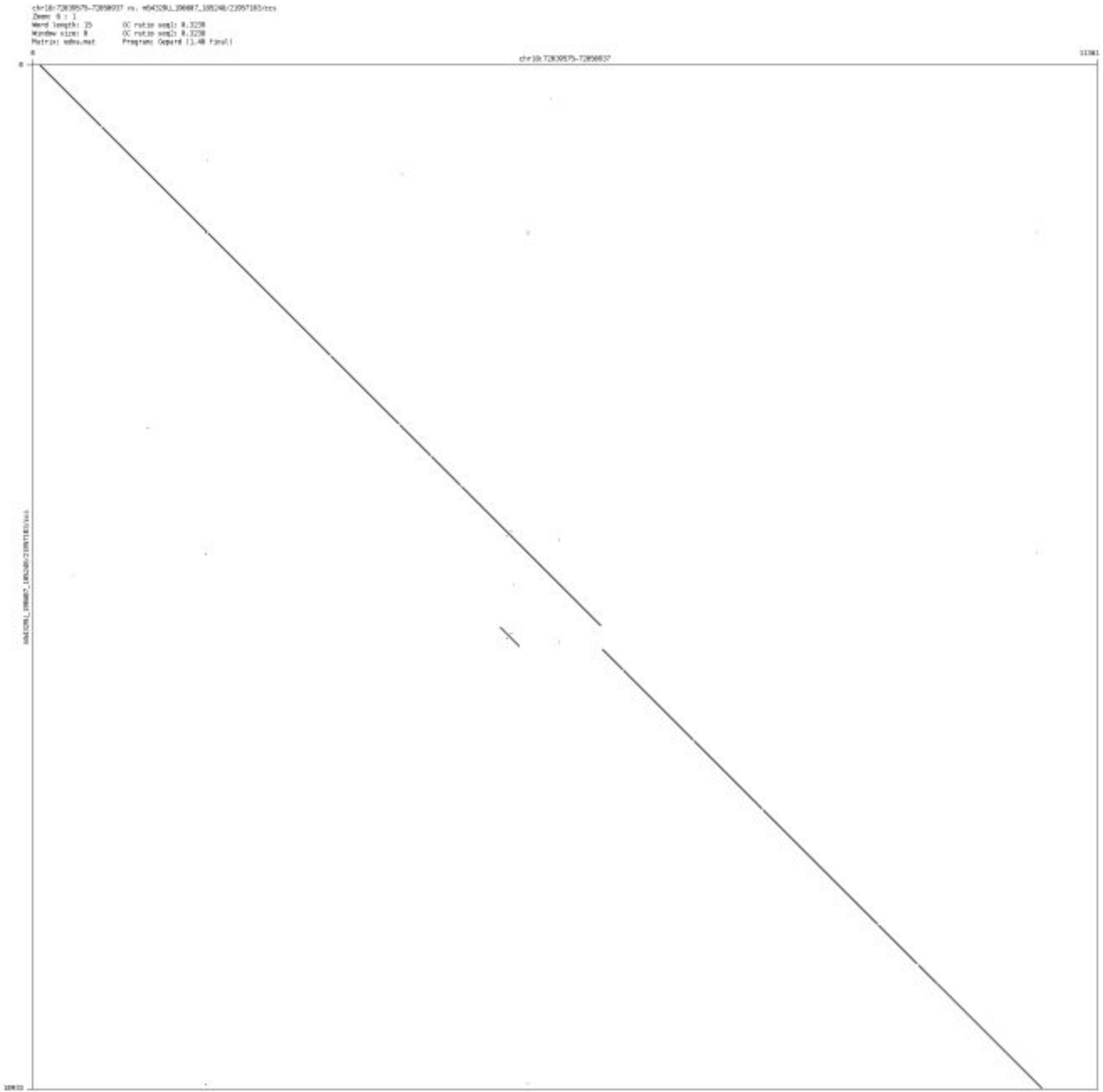

- chr19_41803069_41807384

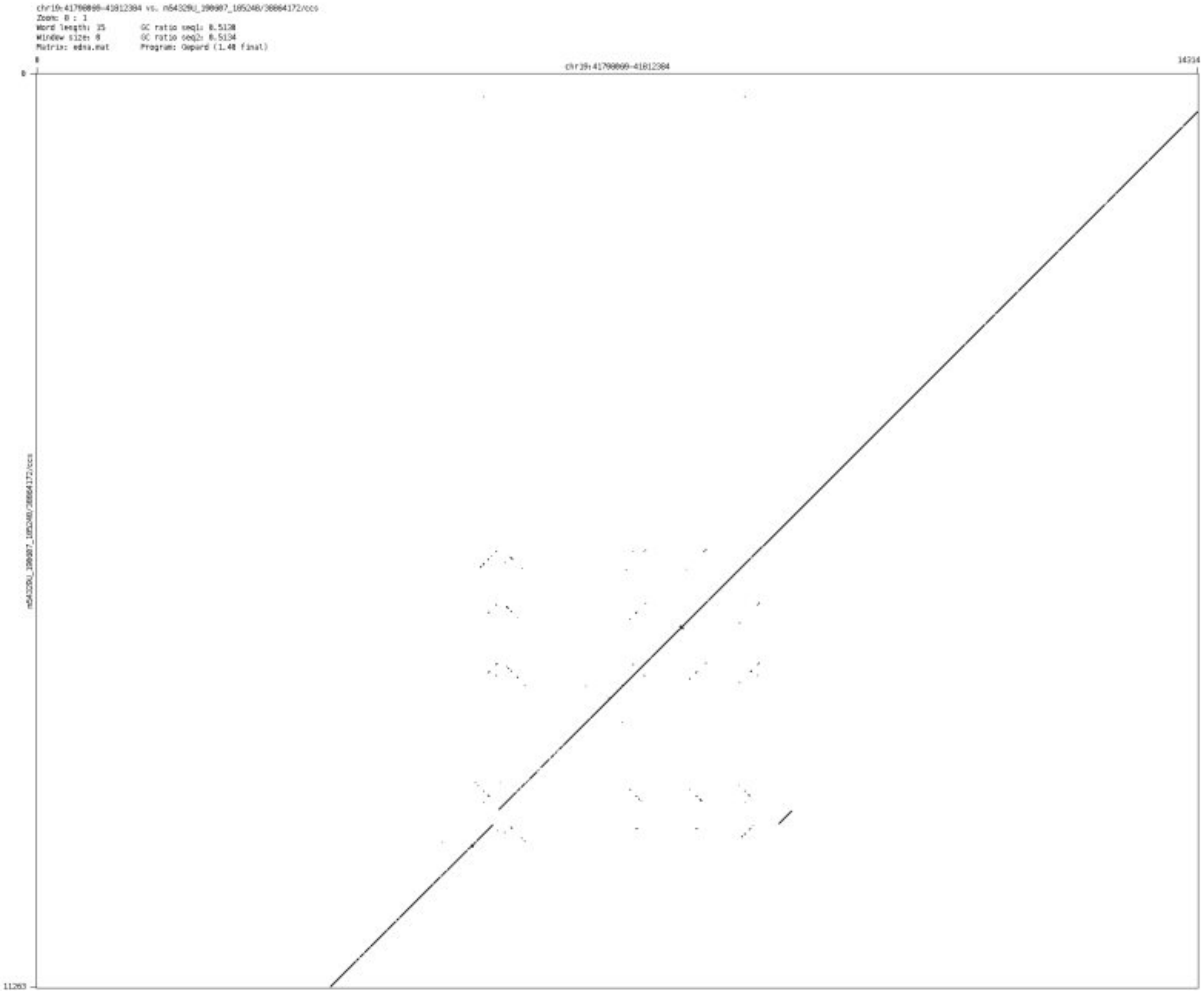

- chr20_2379056_2380273

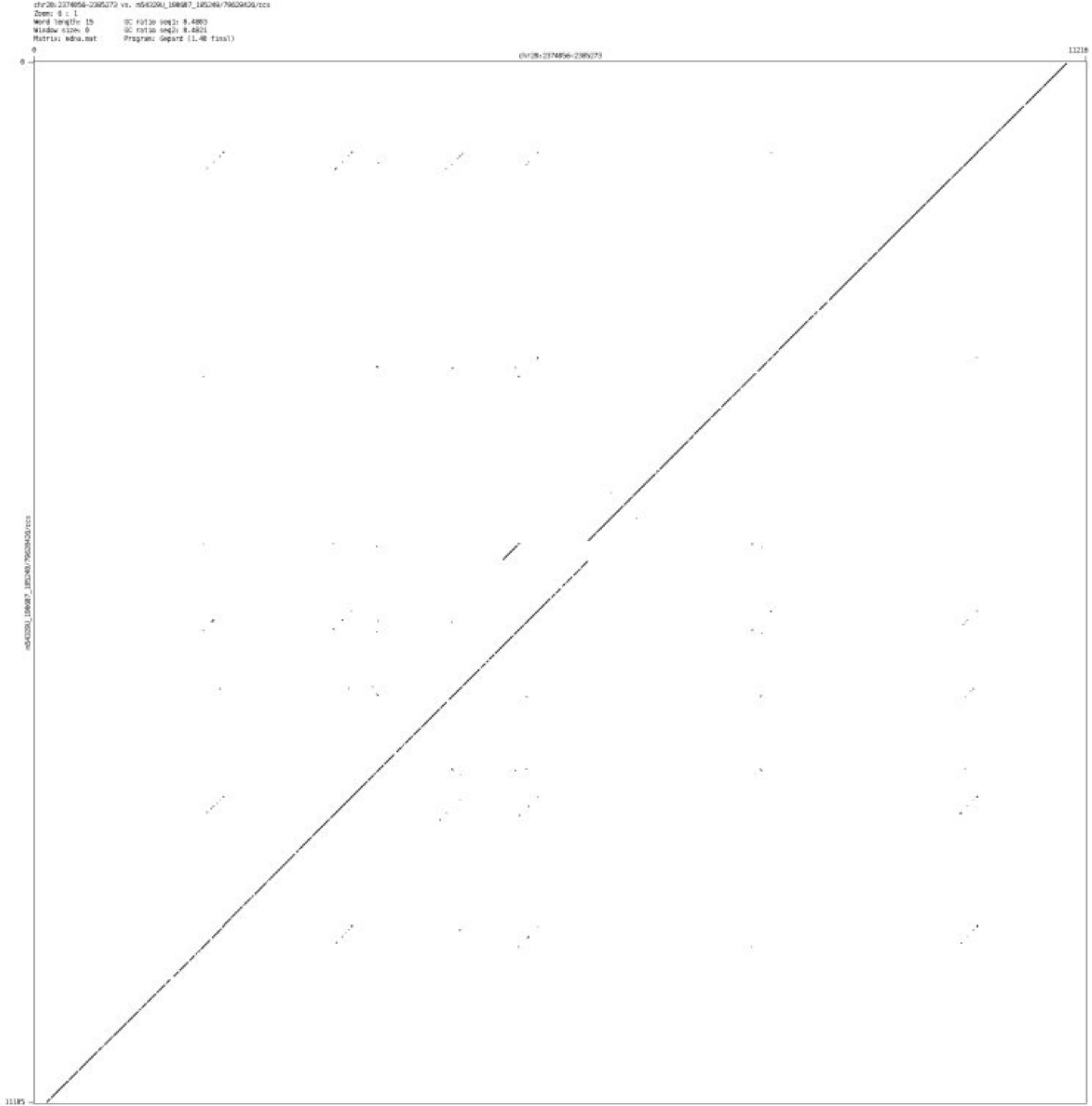

➢ chr20_44676947_44683491

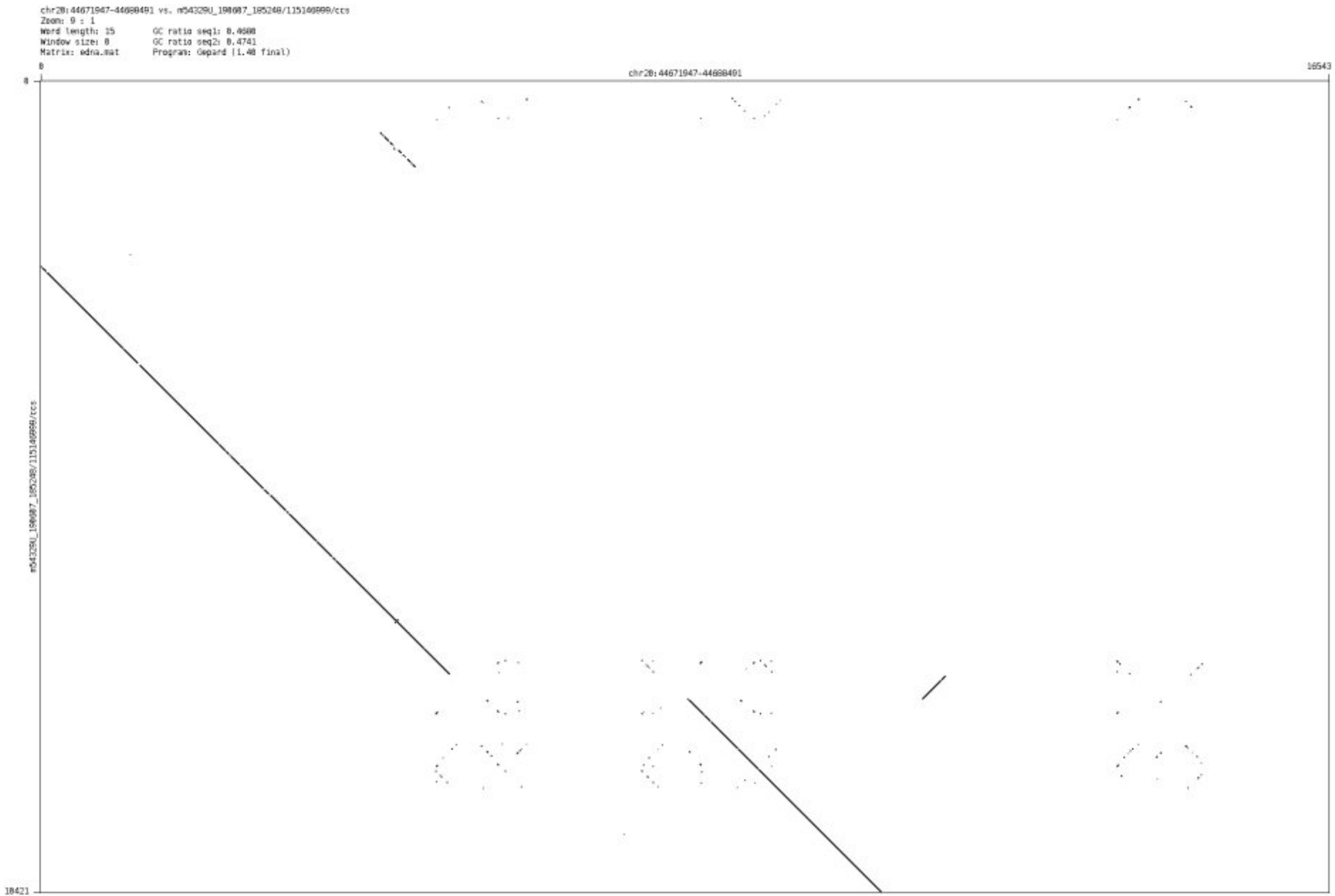

- chr21_24391466_24392454

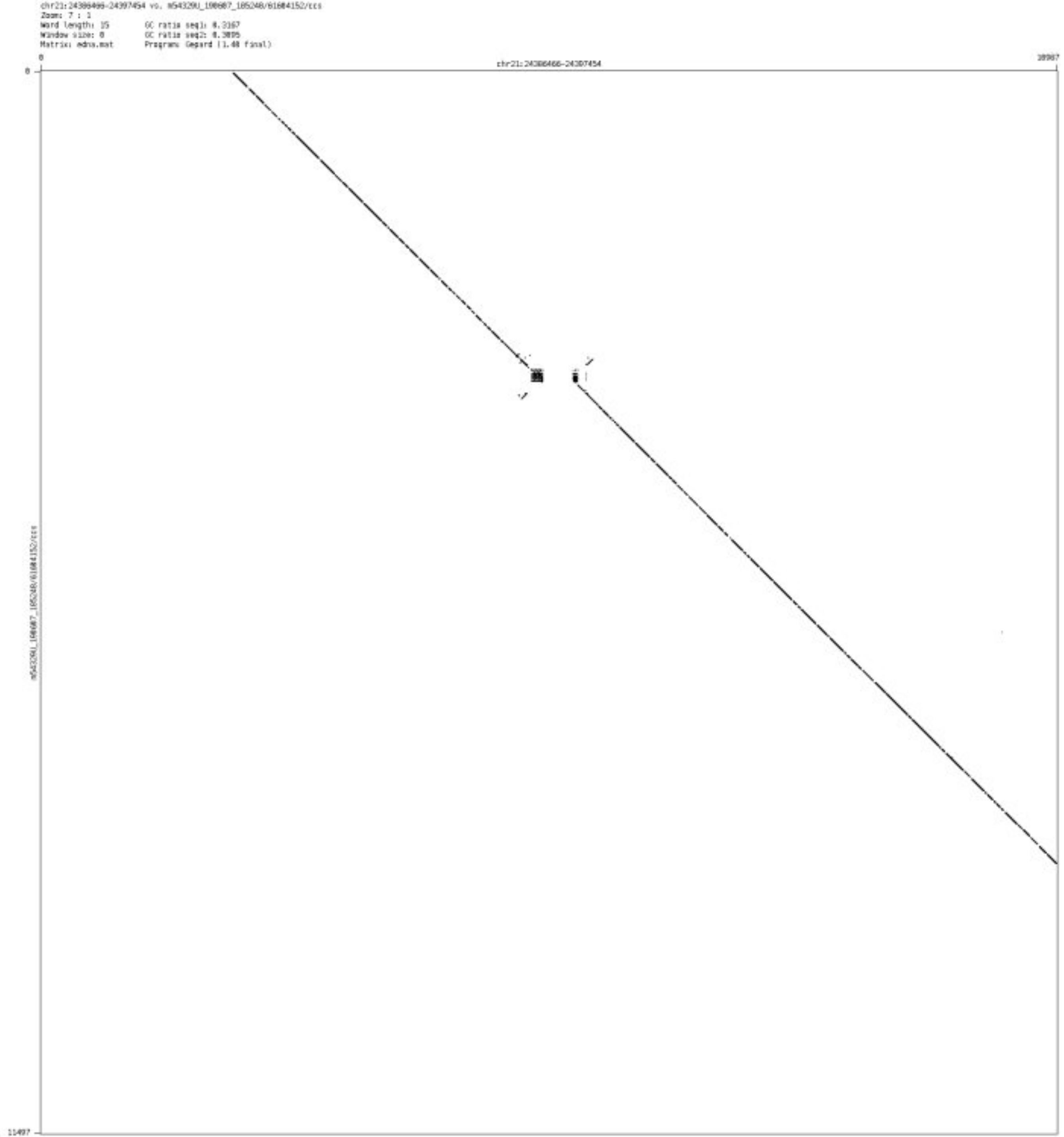

- chr22_20075270_20075522

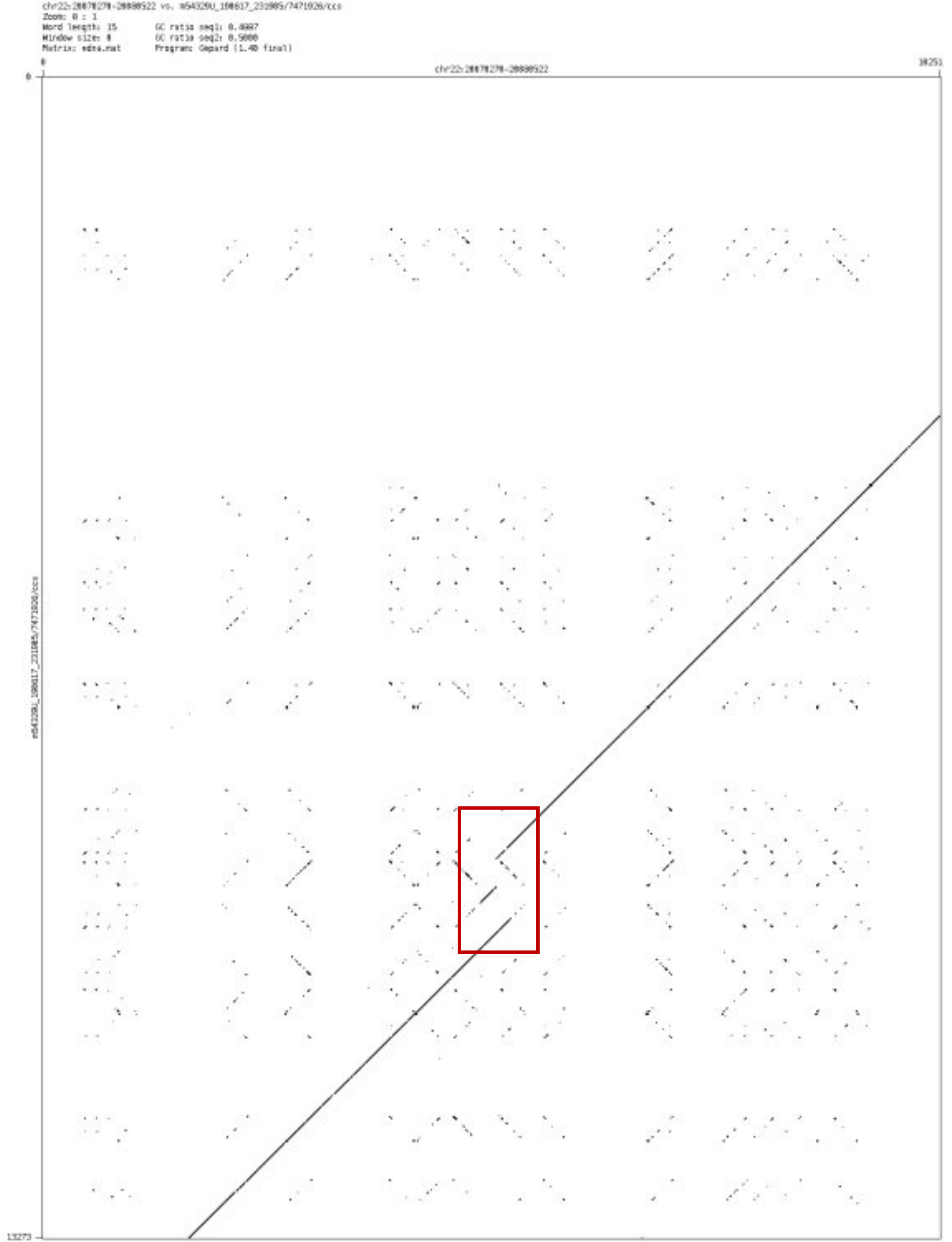

- chr22_45349173_45350498

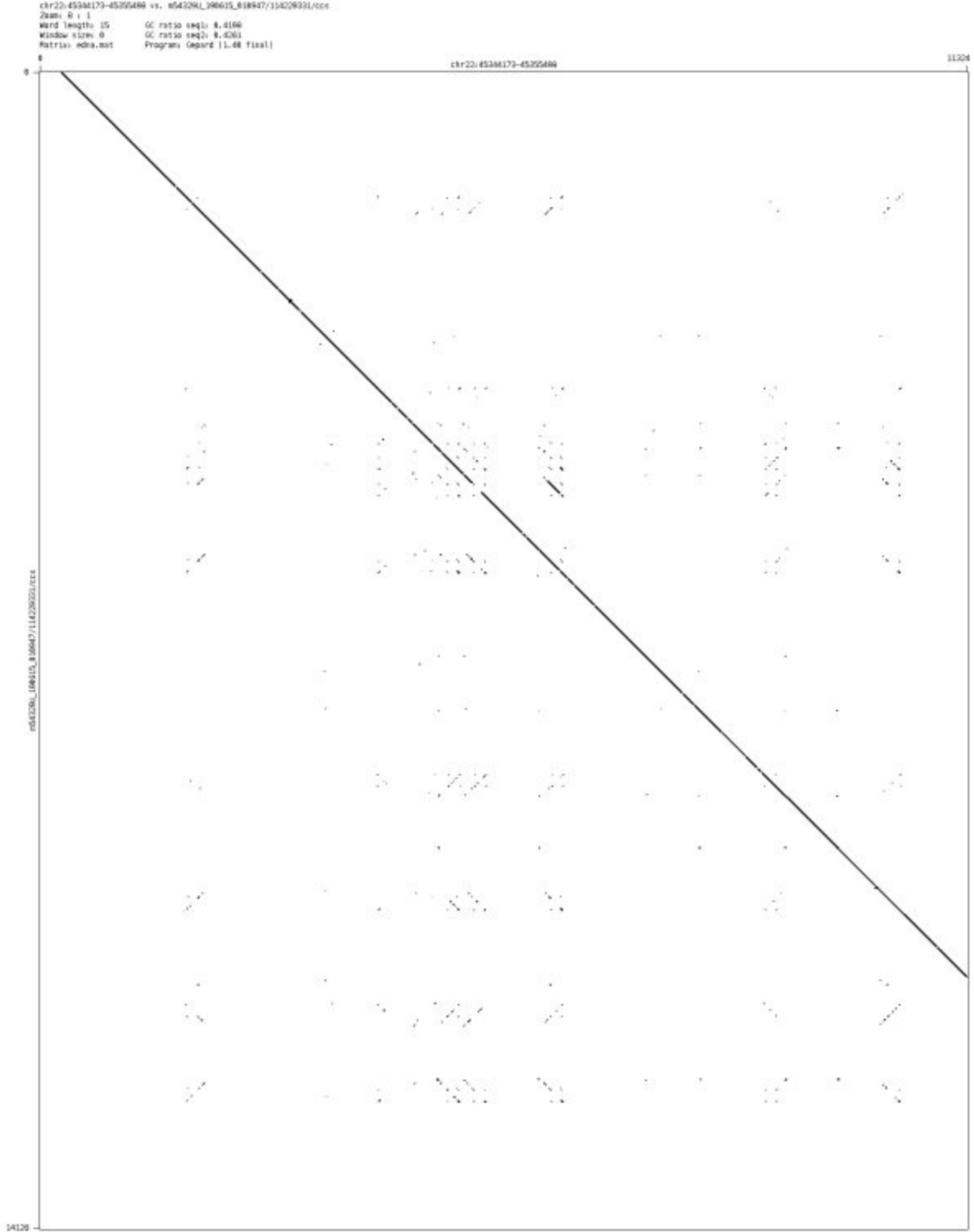

### ’Messy’ CSV-Case A

➢ chr1_231202060_231202180

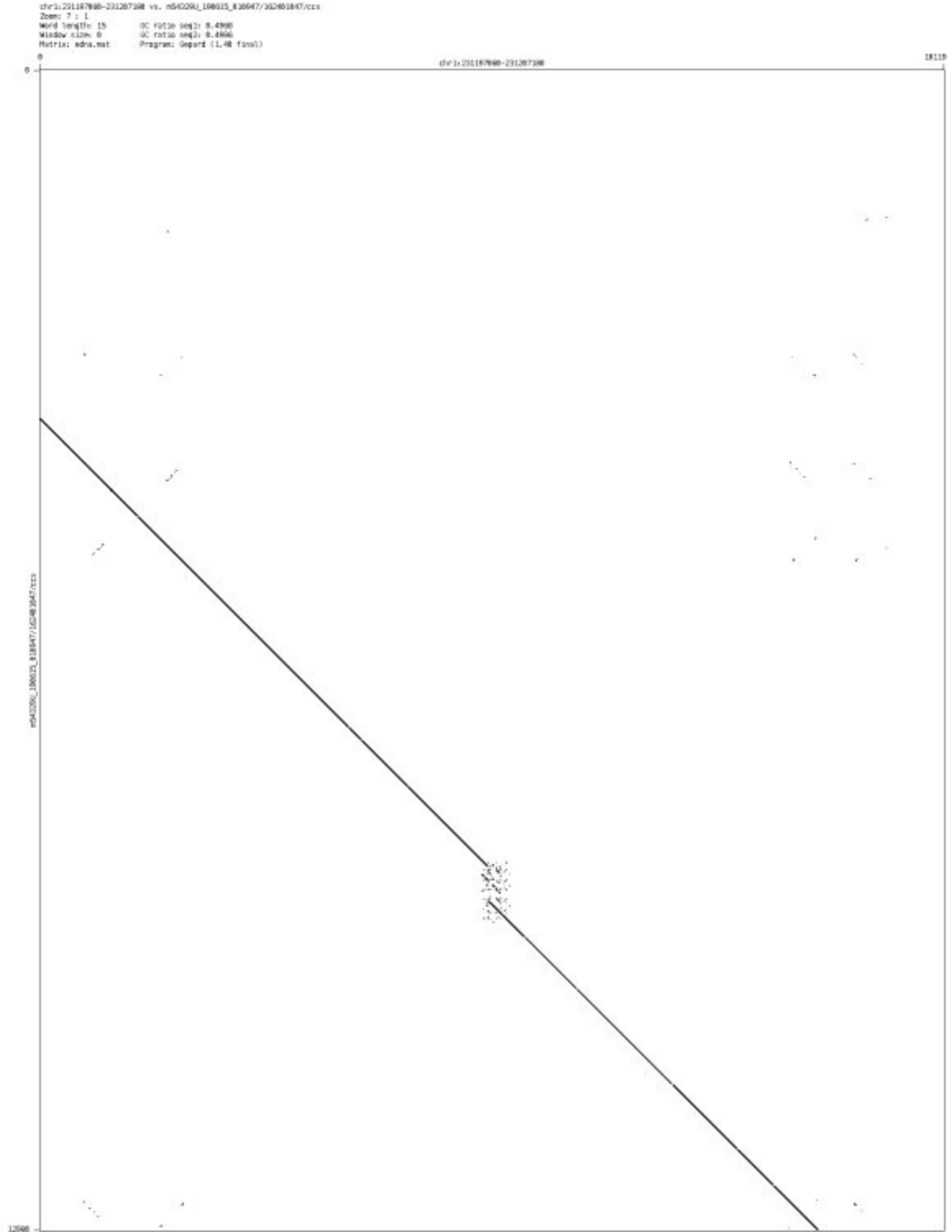

- chr1_206924211_206924525

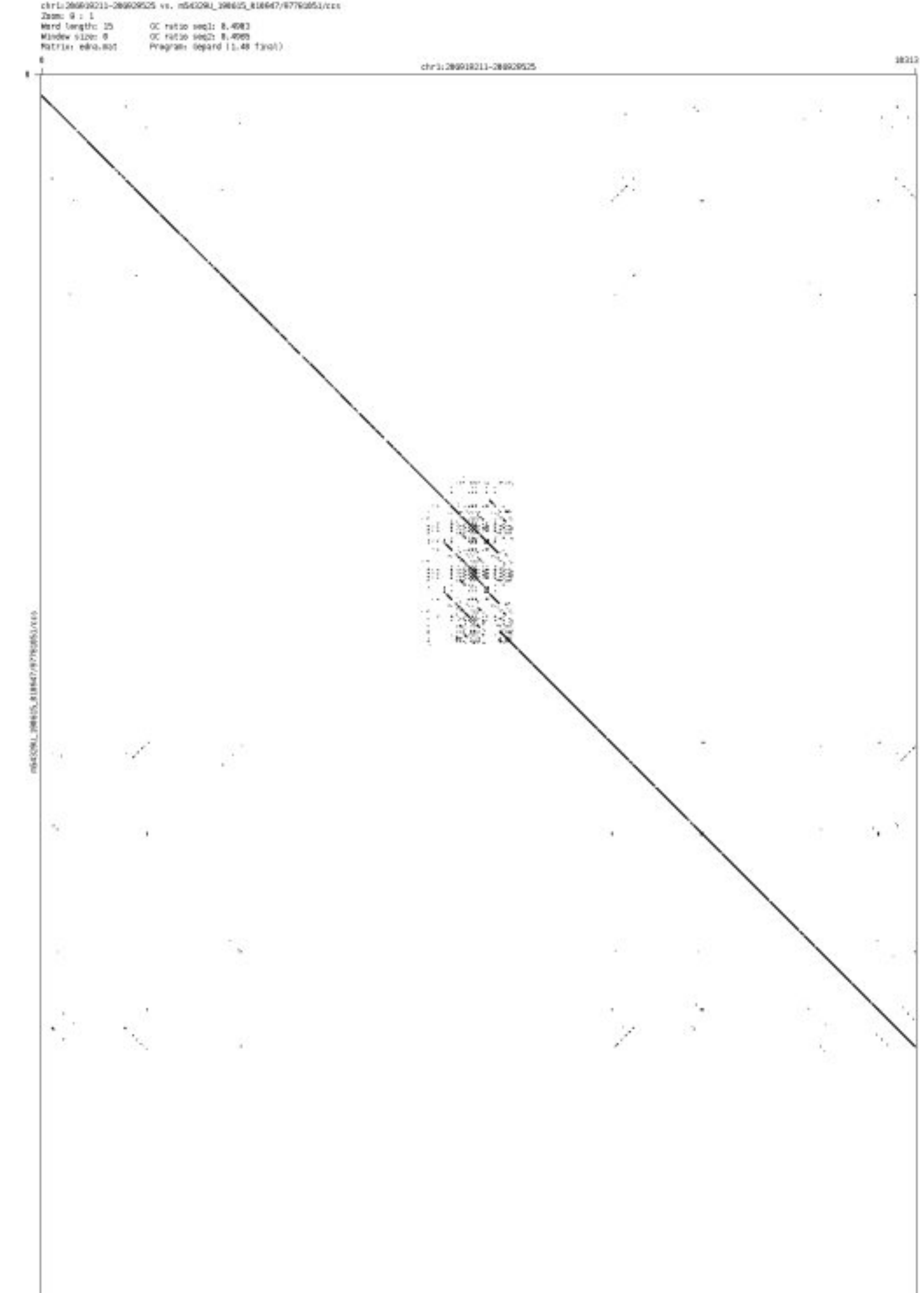

➢ chr1_154714108_154714439

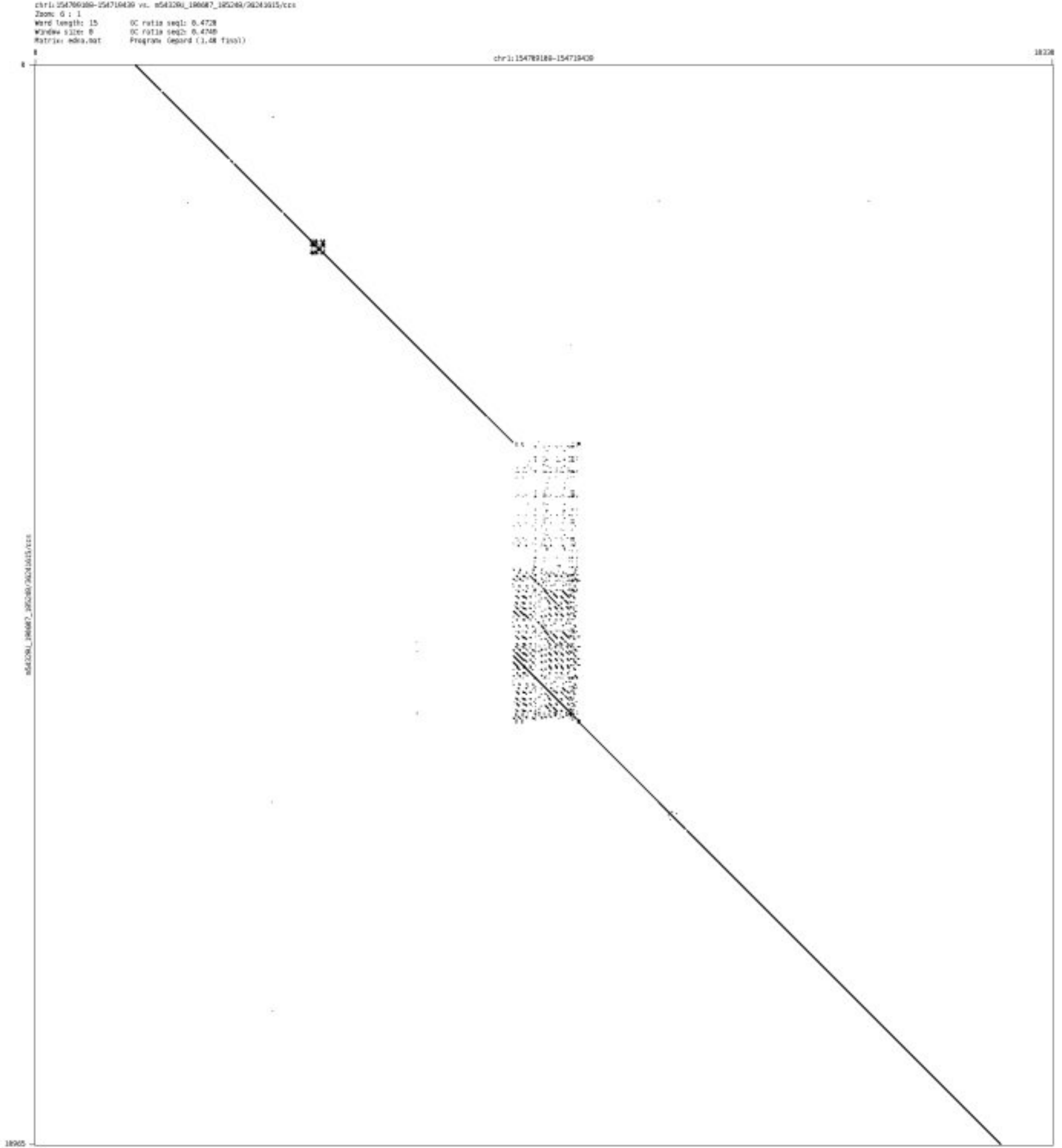

- chr1_234247922_234248122

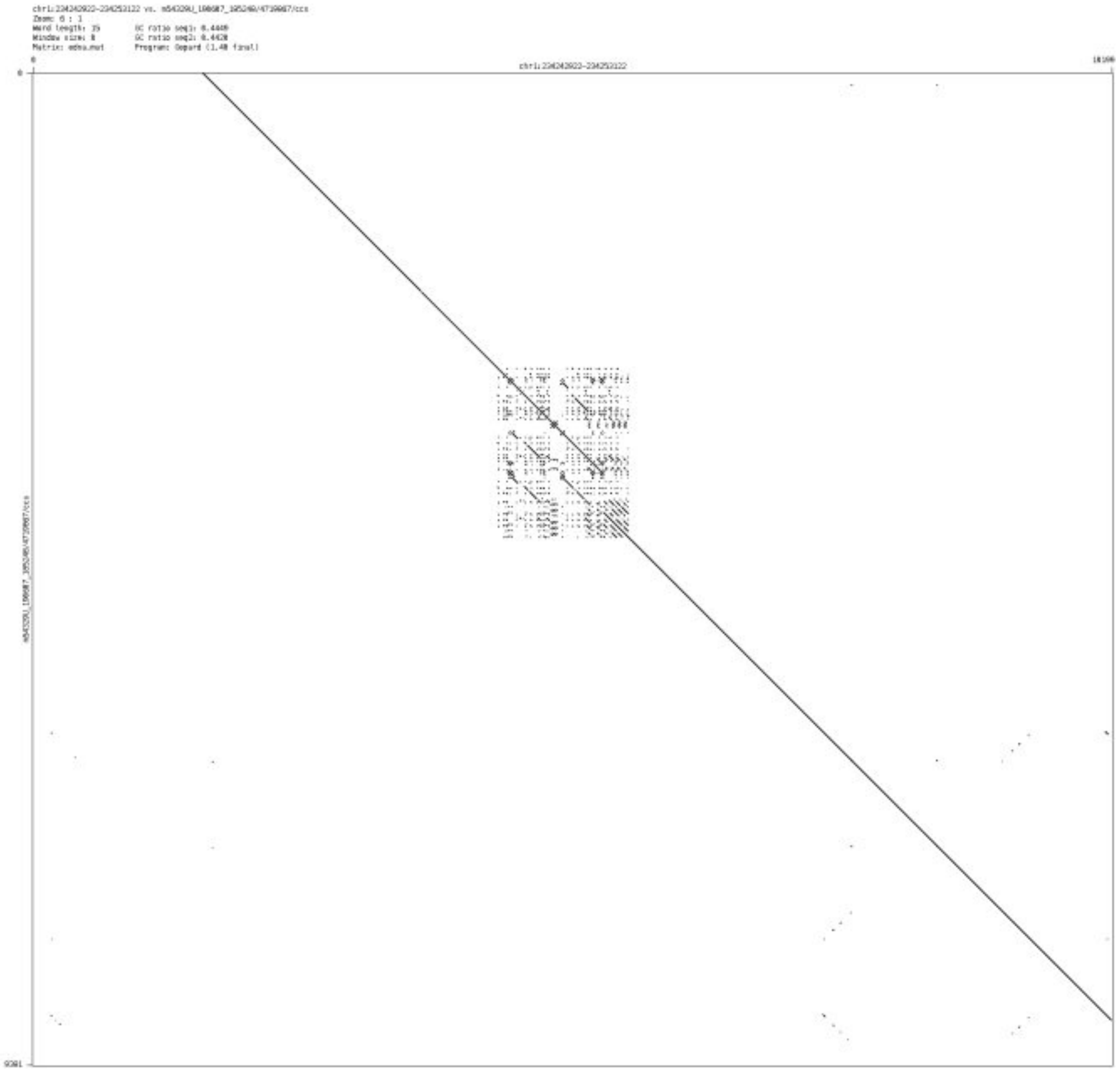

- chr1_235078305_235078561

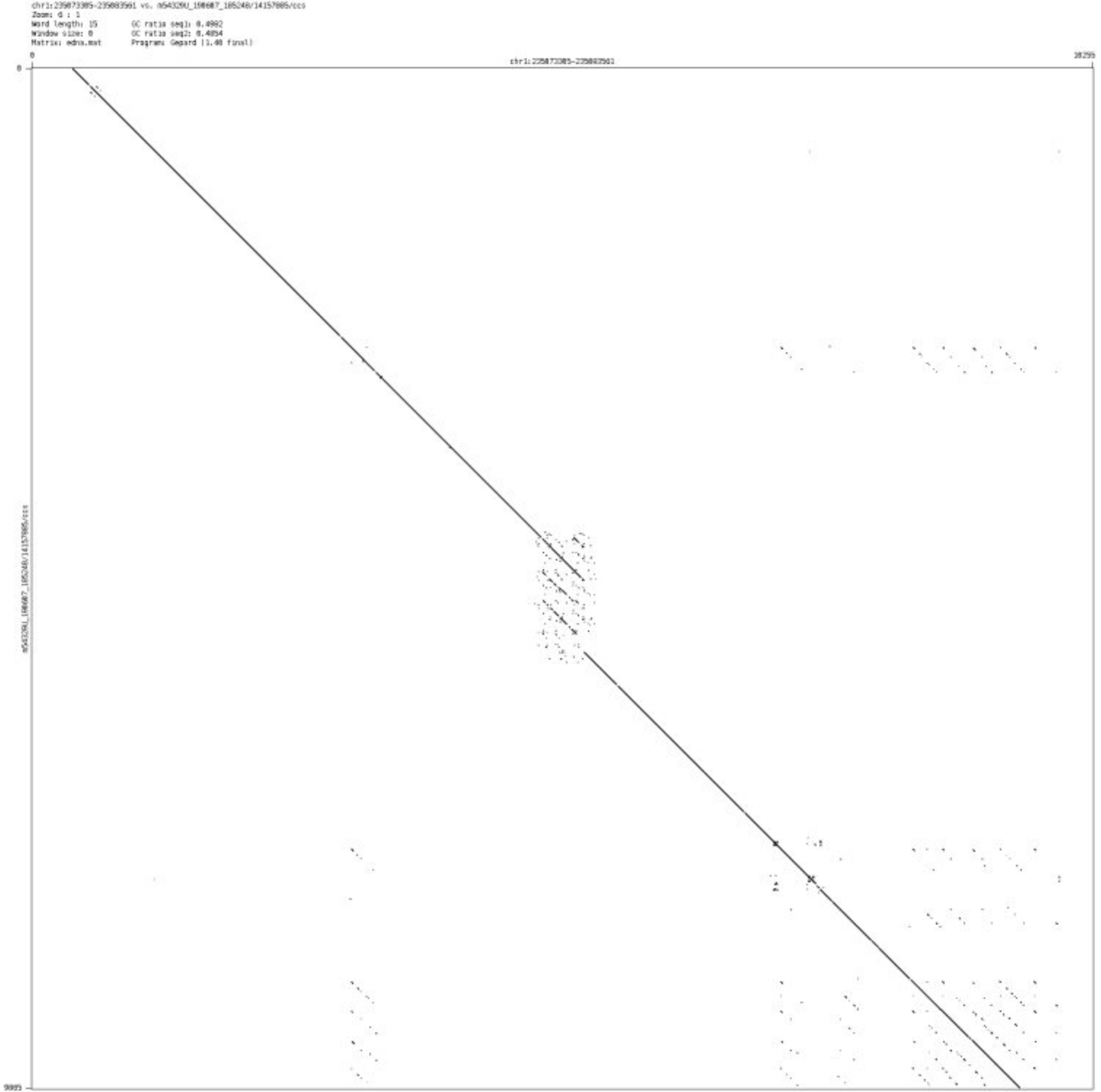

- chr2_1281271_1281469

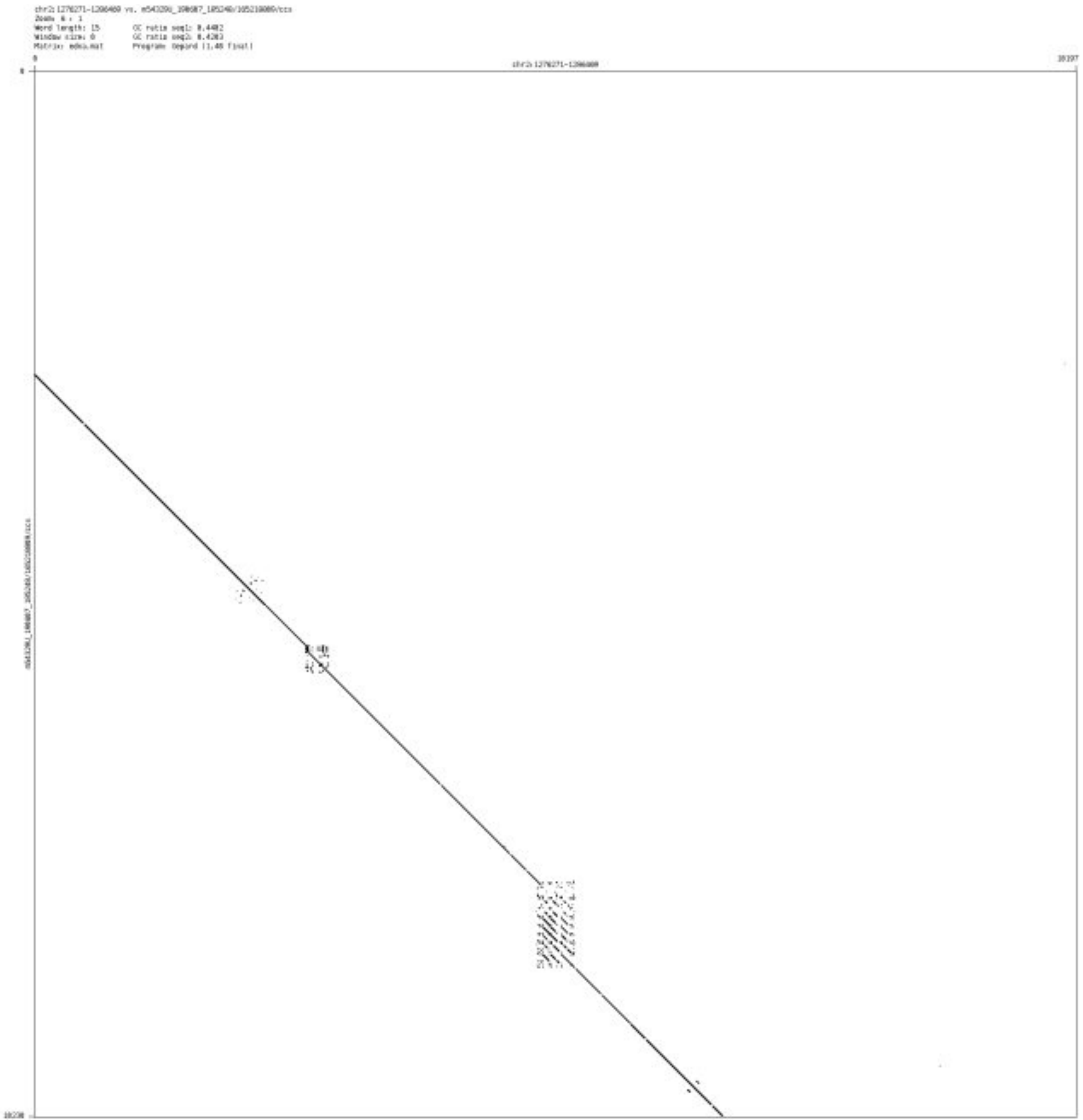

- chr2_1861782_1861968

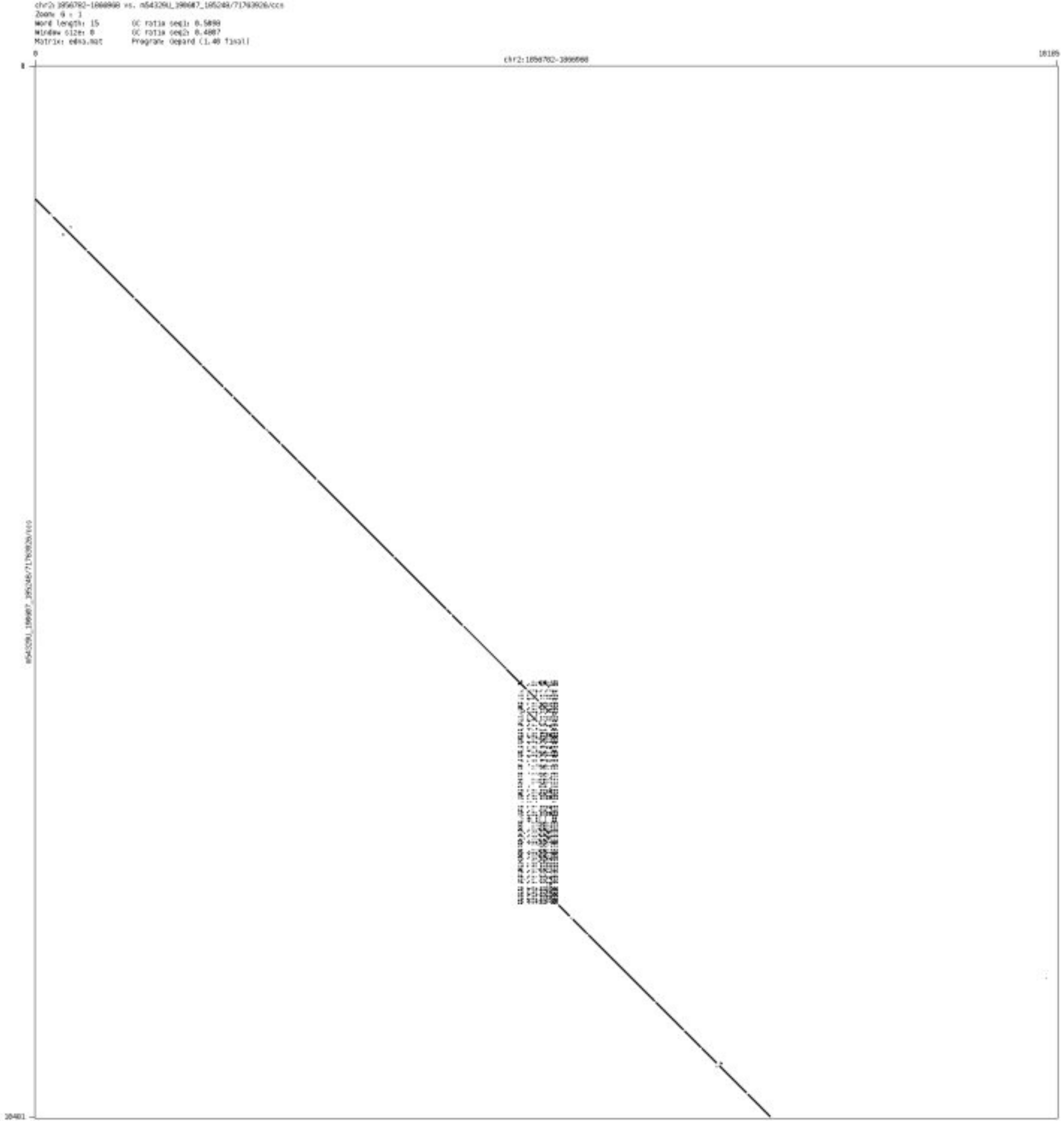

➢ chr2_4974106_4974189

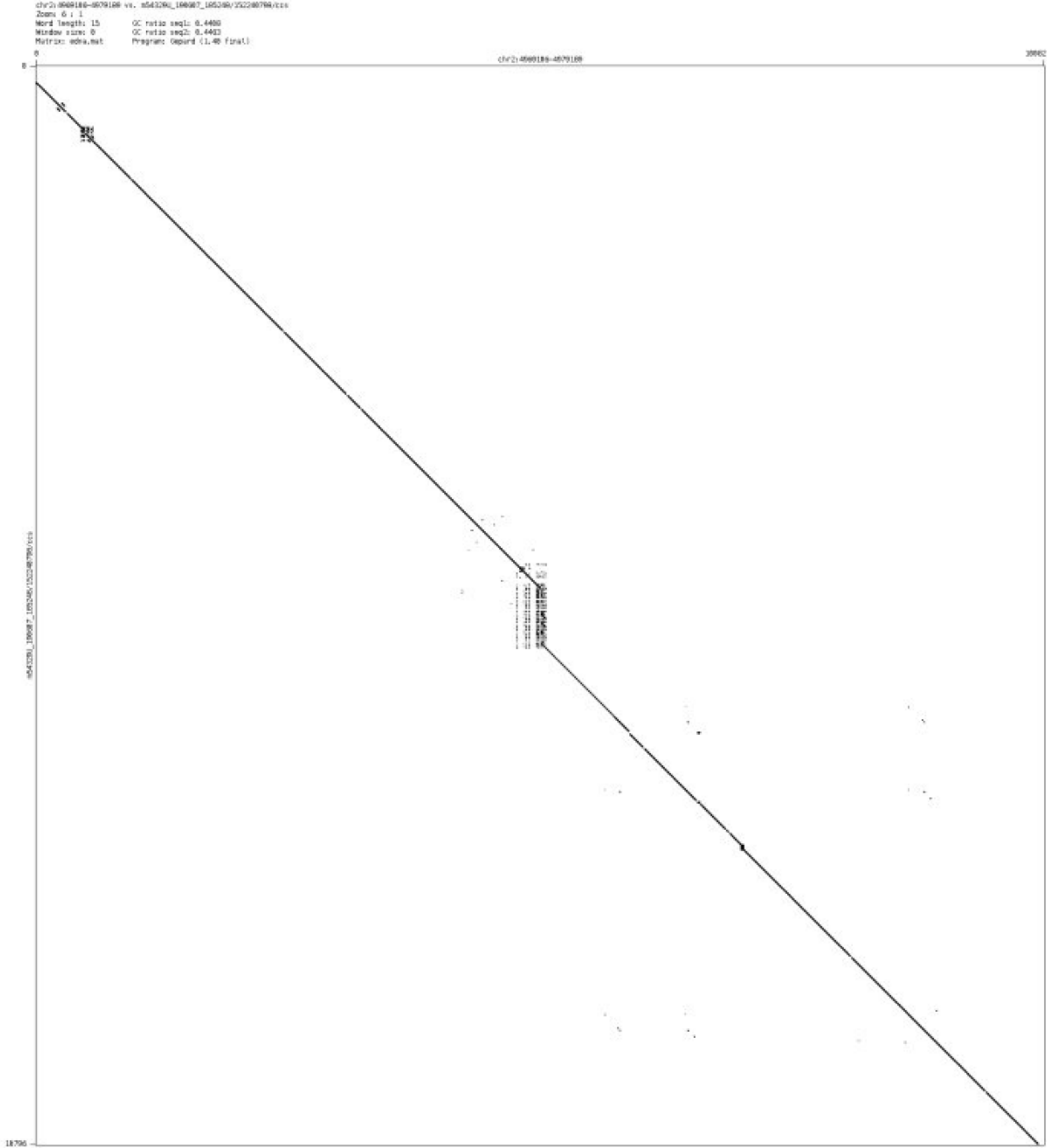

- chr2_10011727_10015474

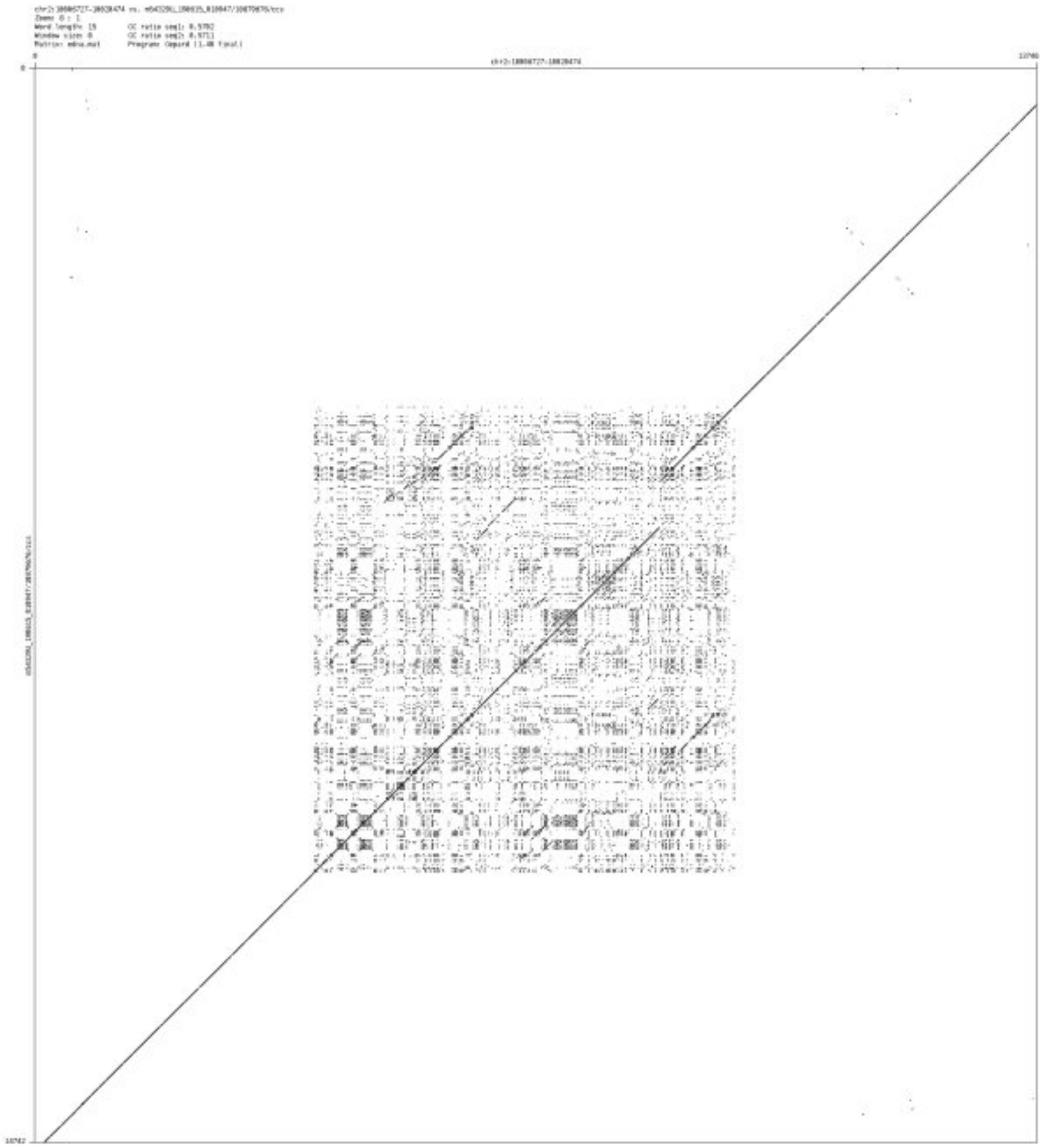

- chr3_195055517_195059786

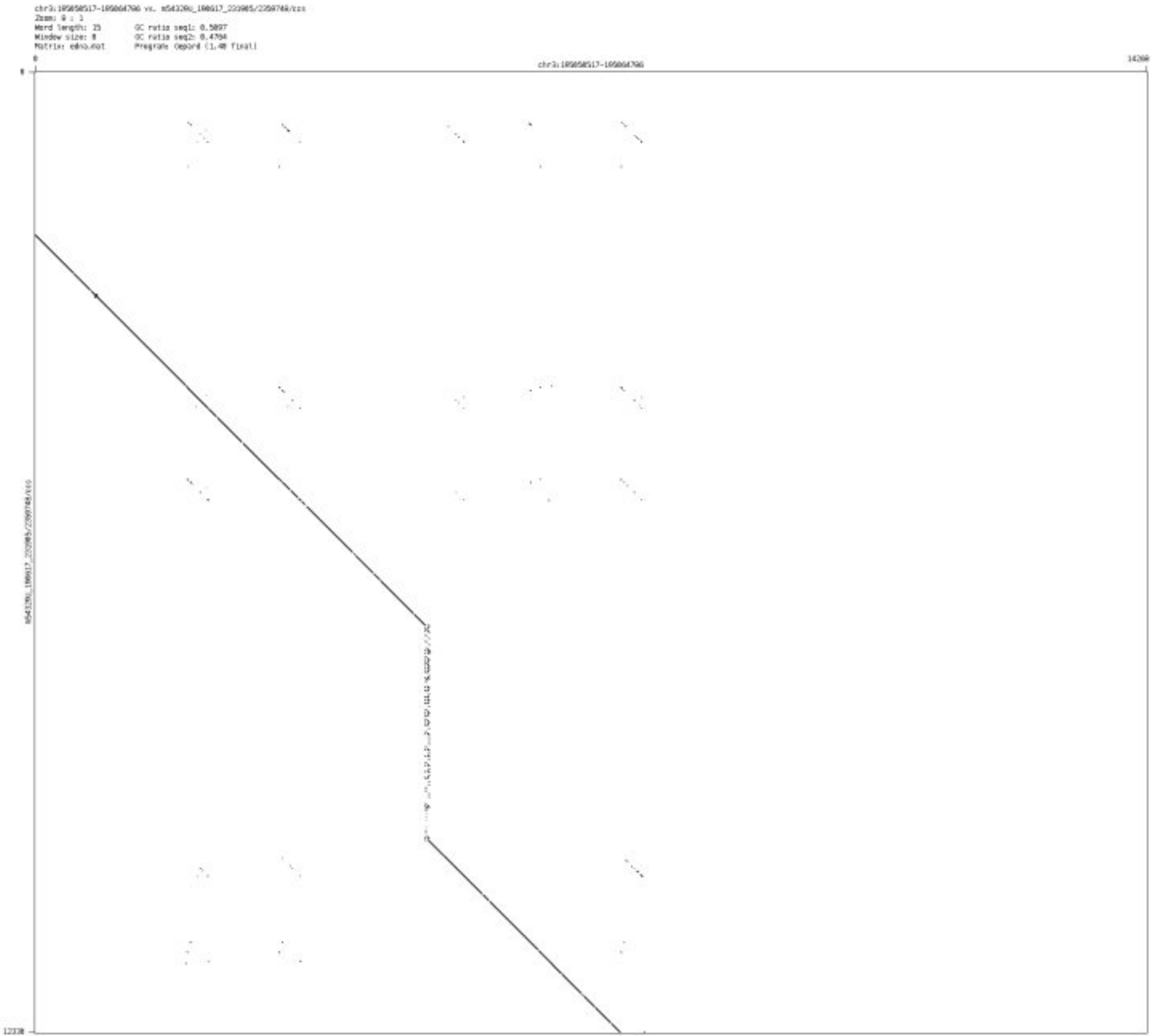

➢ chr5_1044925_1045186

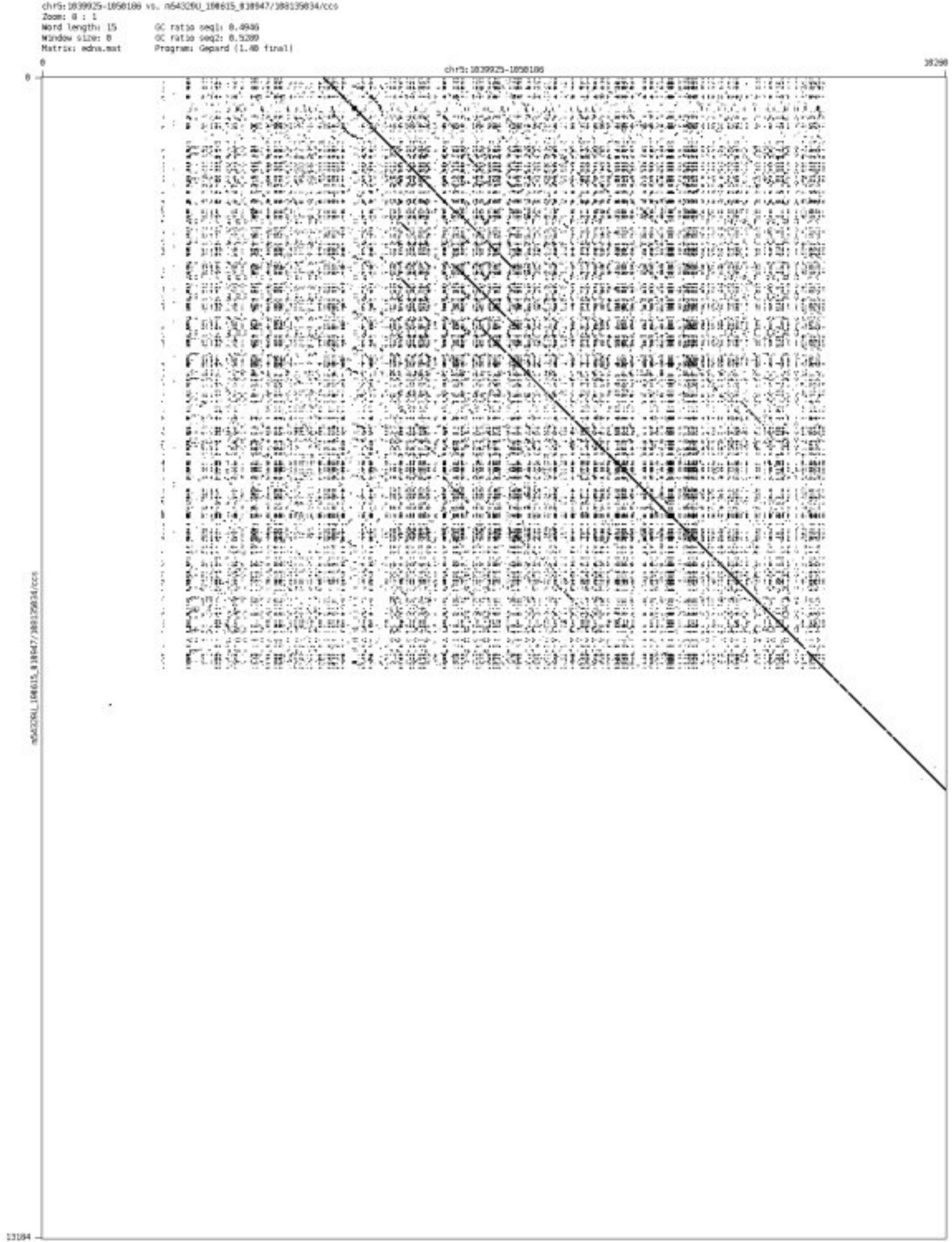

- chr5_1845637_1845877

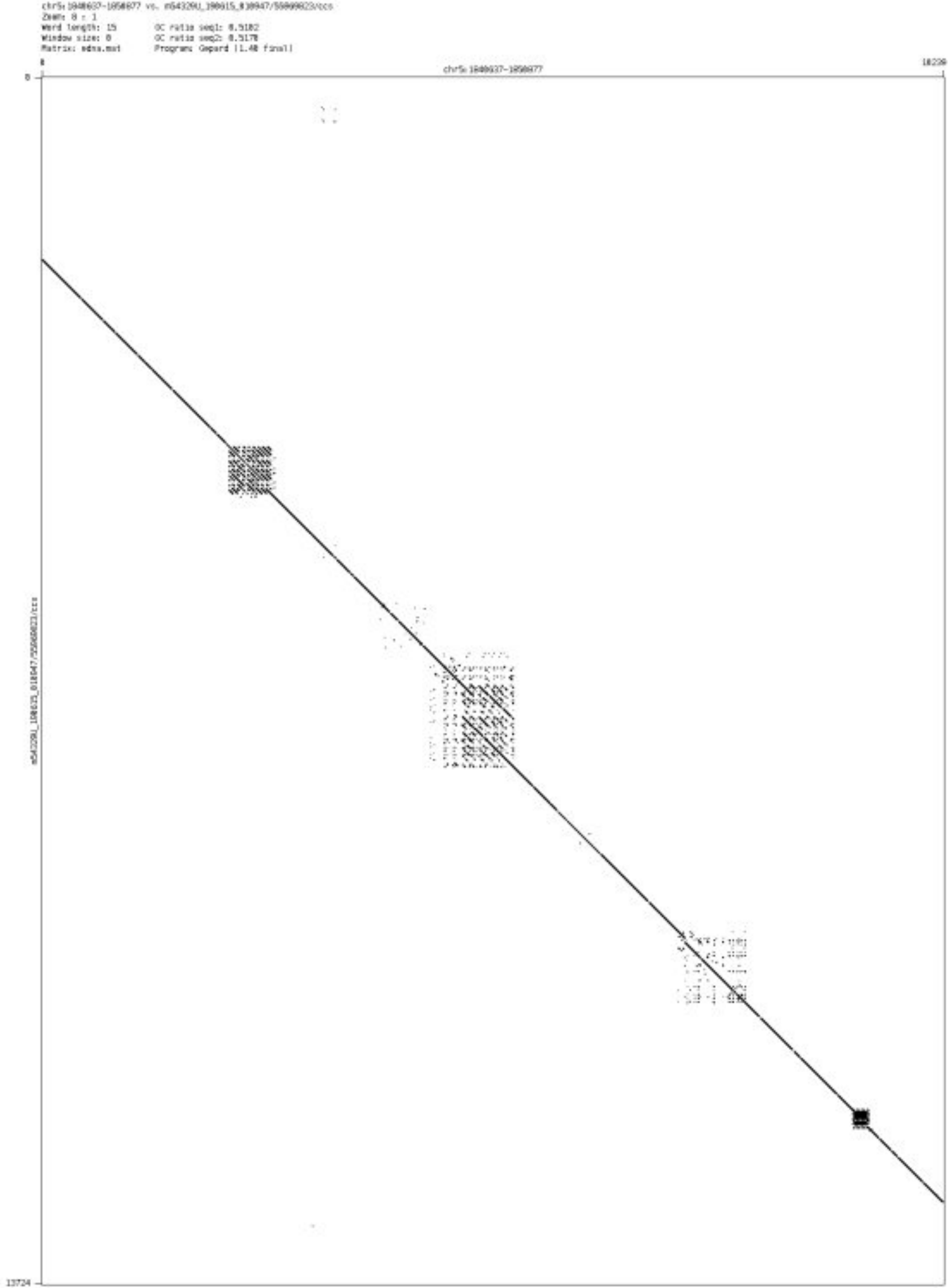

- chr5_3659963_3660133

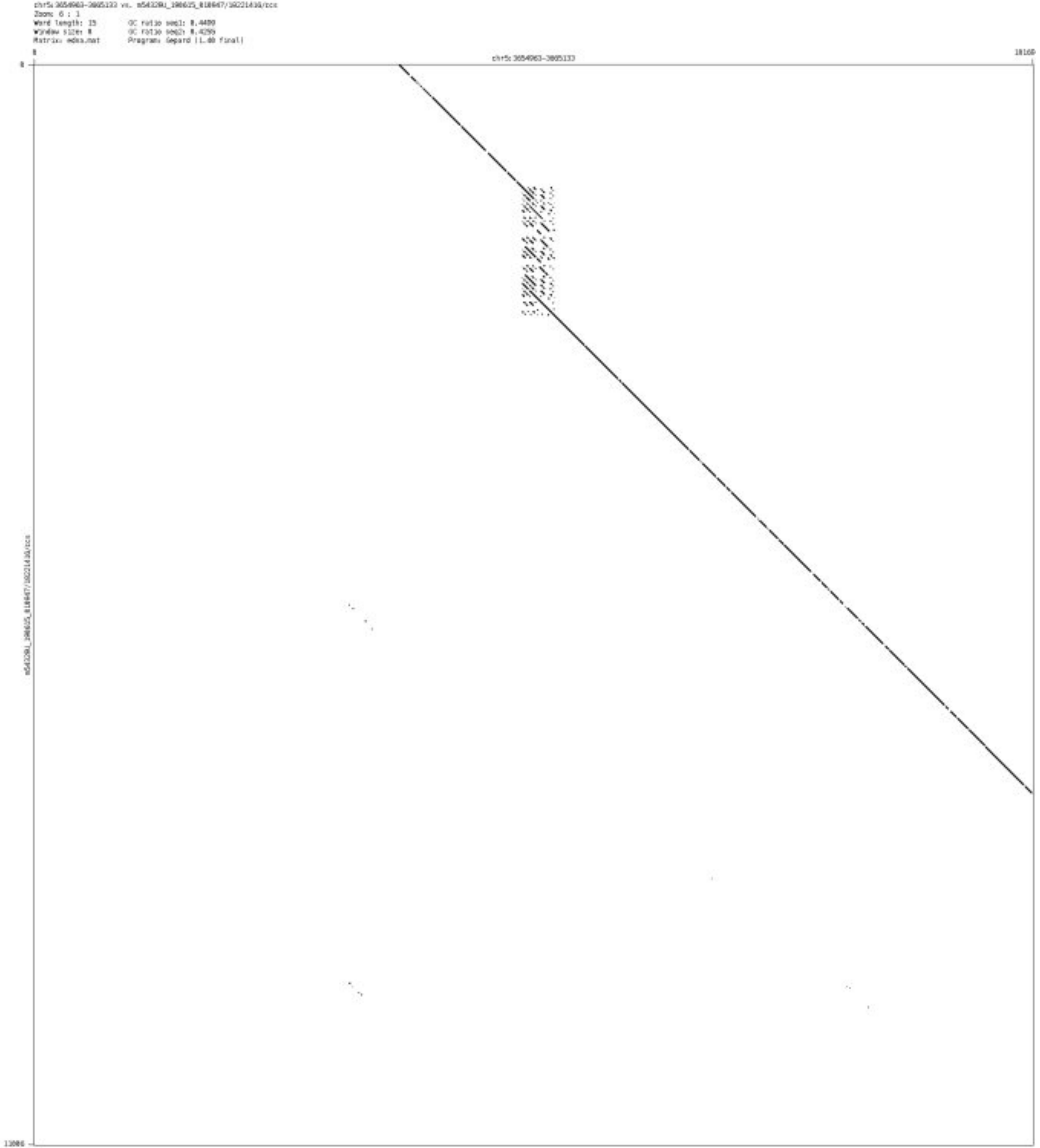

➢ chr6_2504753_2504887

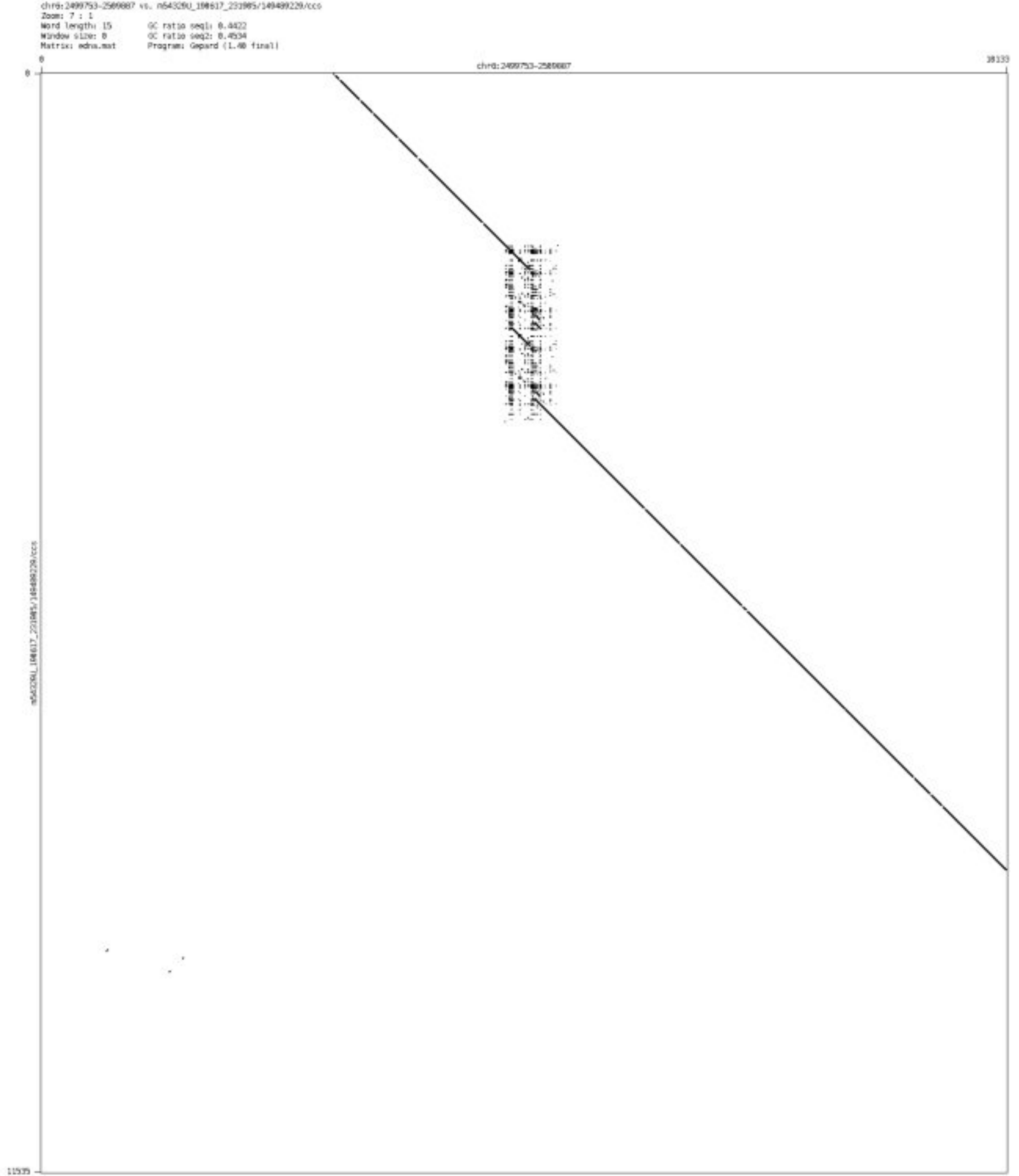

- chr6_157238333_157238398

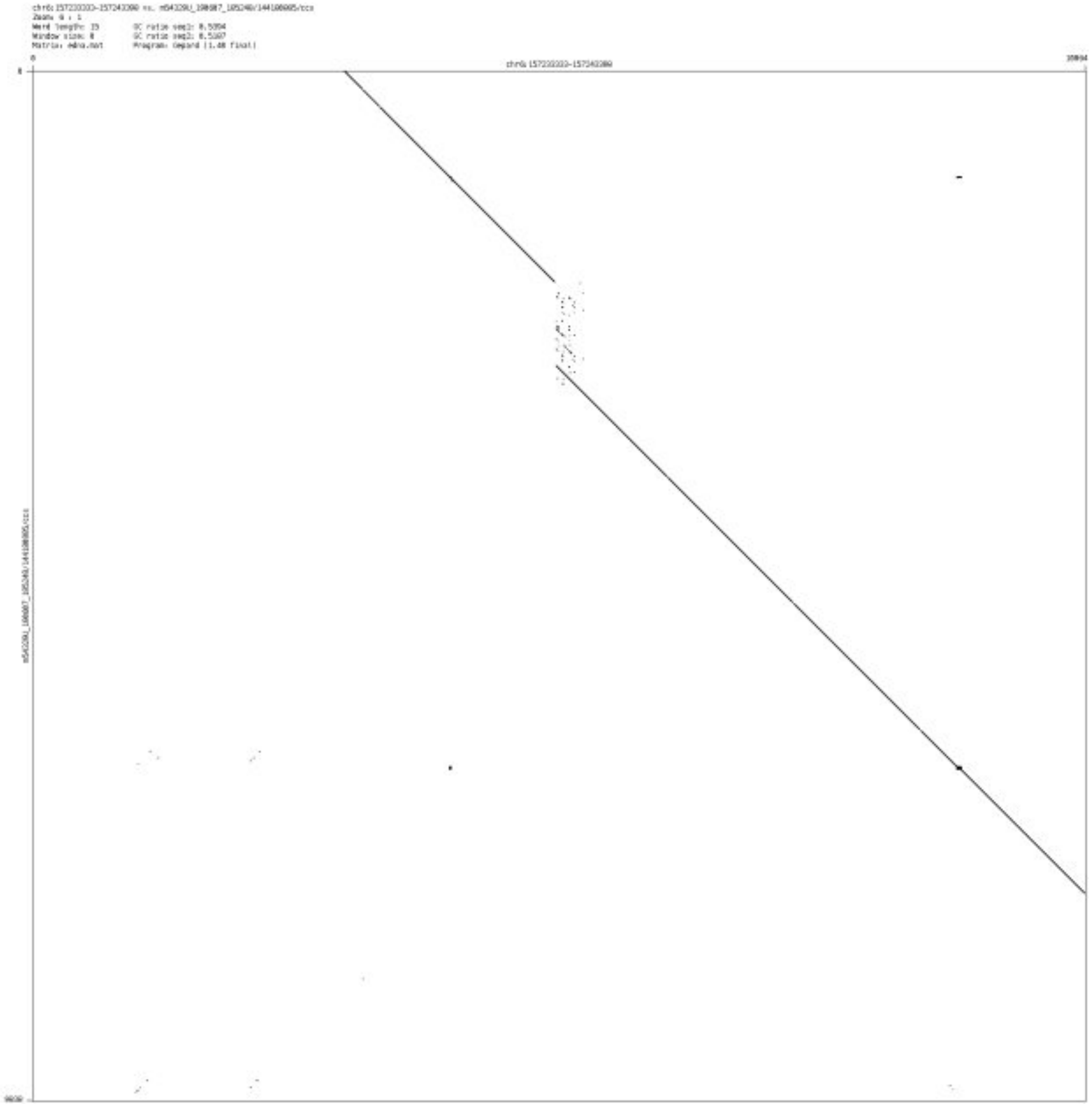

➢ chr6_165749273_165749500

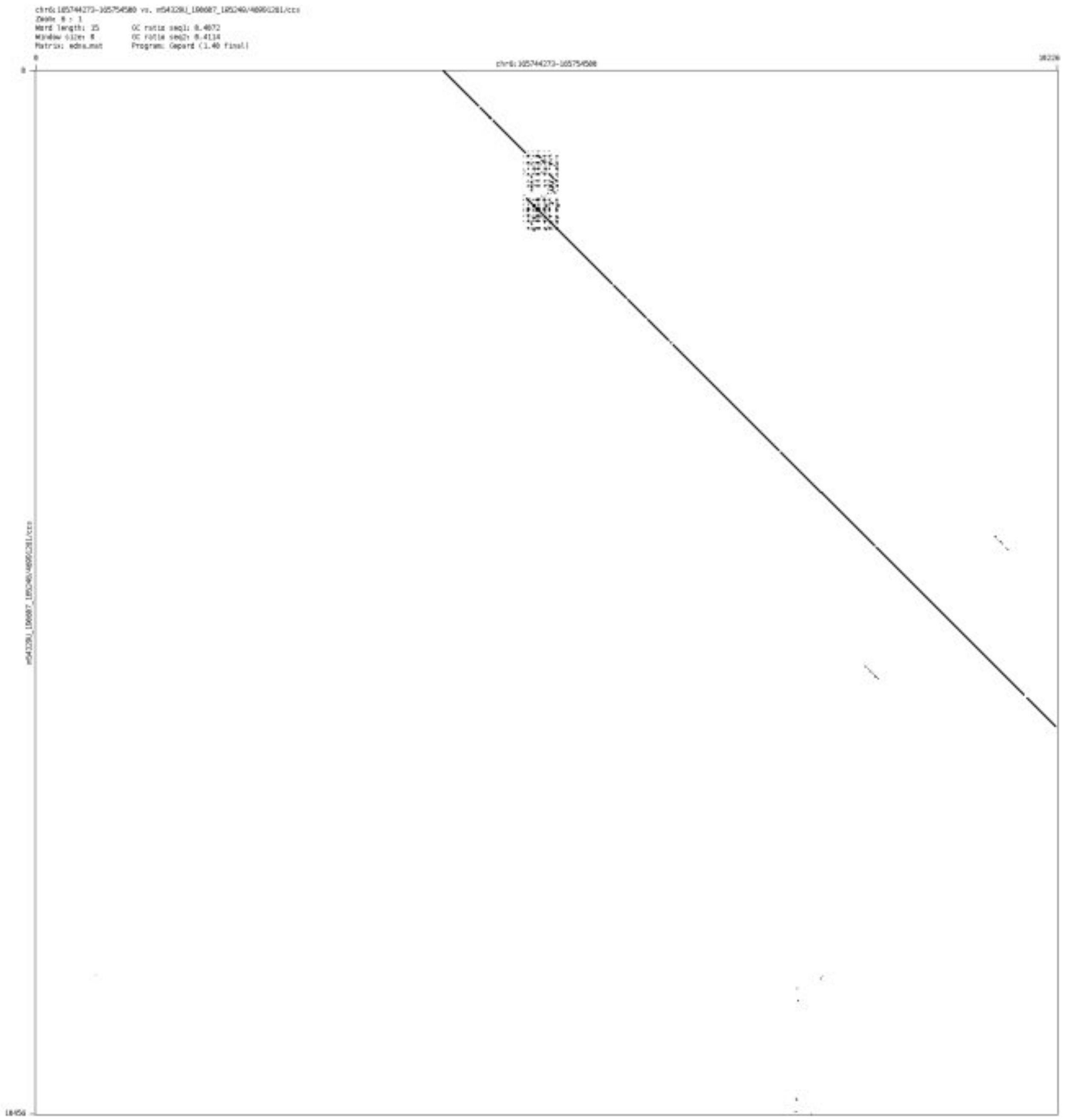

- ➢ chr6_165839937_165840008

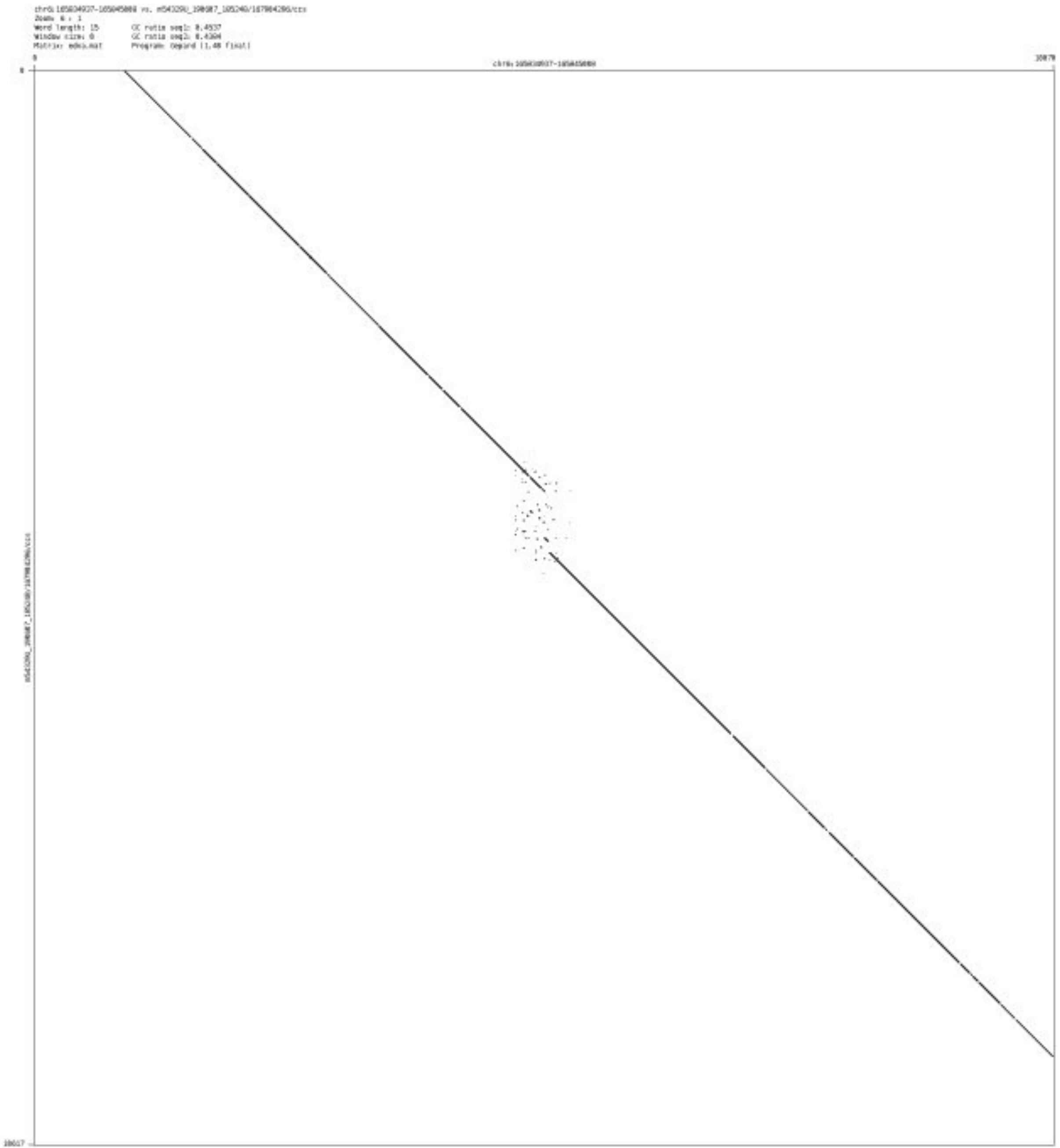

➢ chr6_166416140_166416256

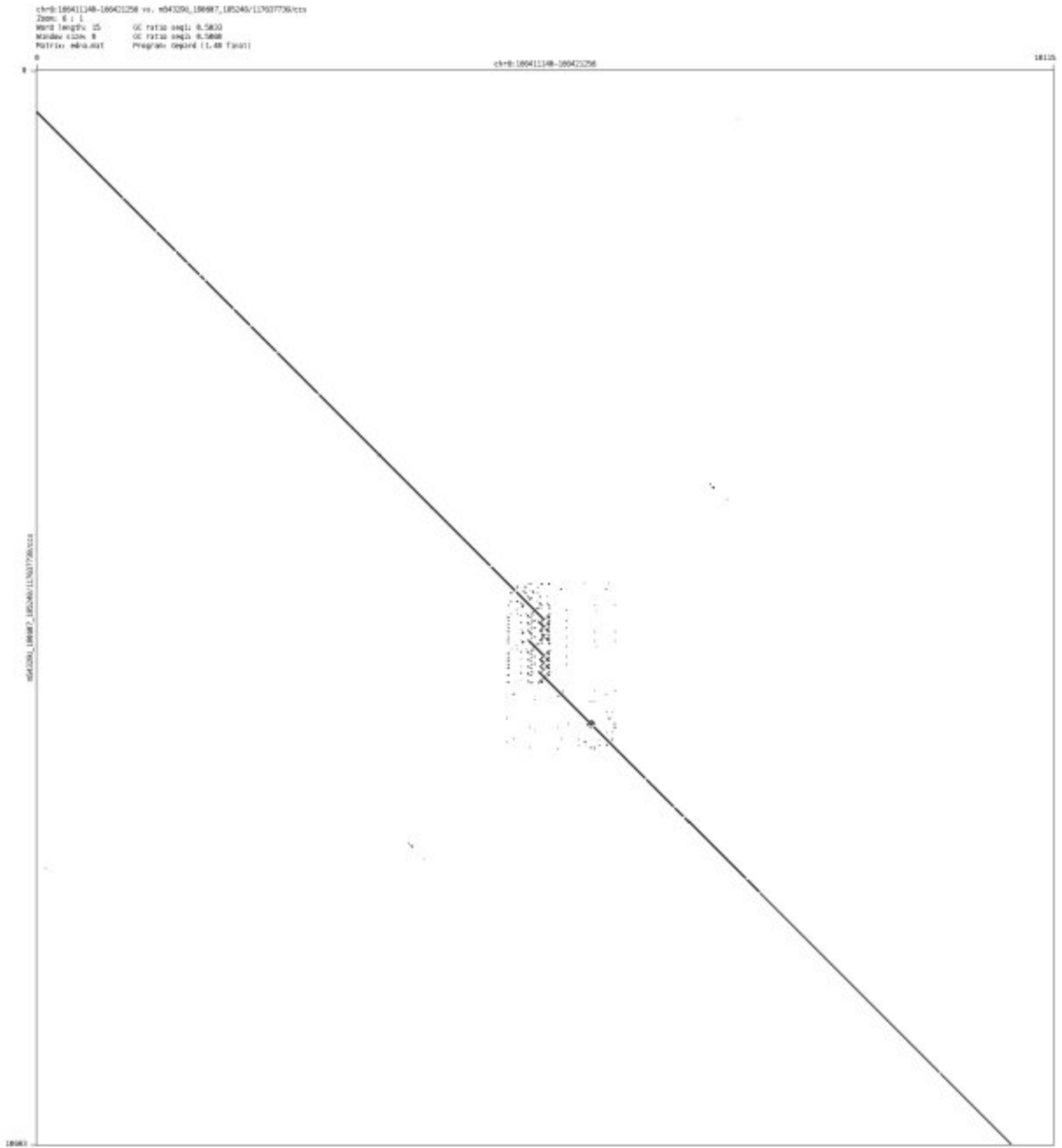

- chr6_167651465_167651832

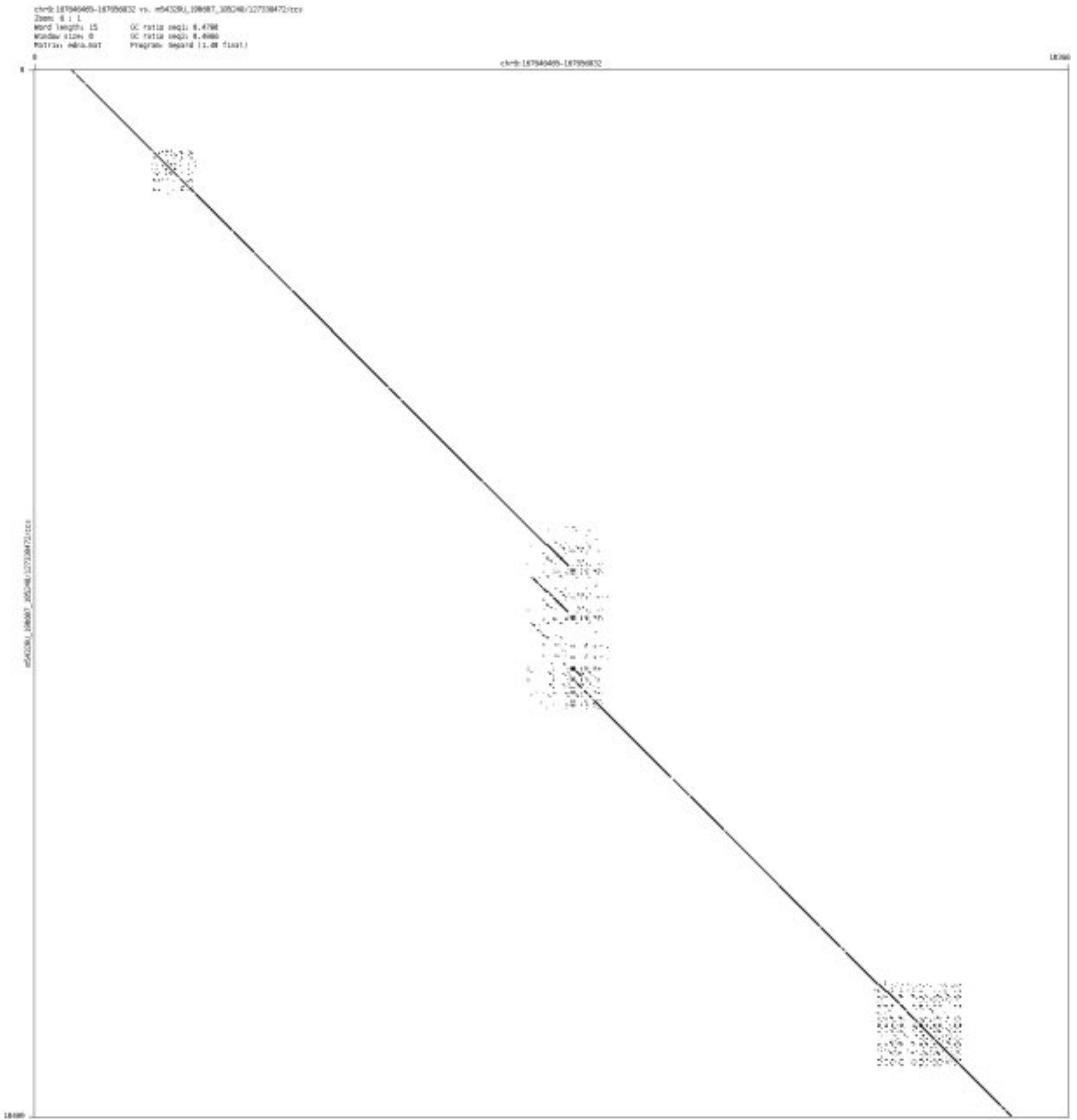

- chr6_167735329_167735534

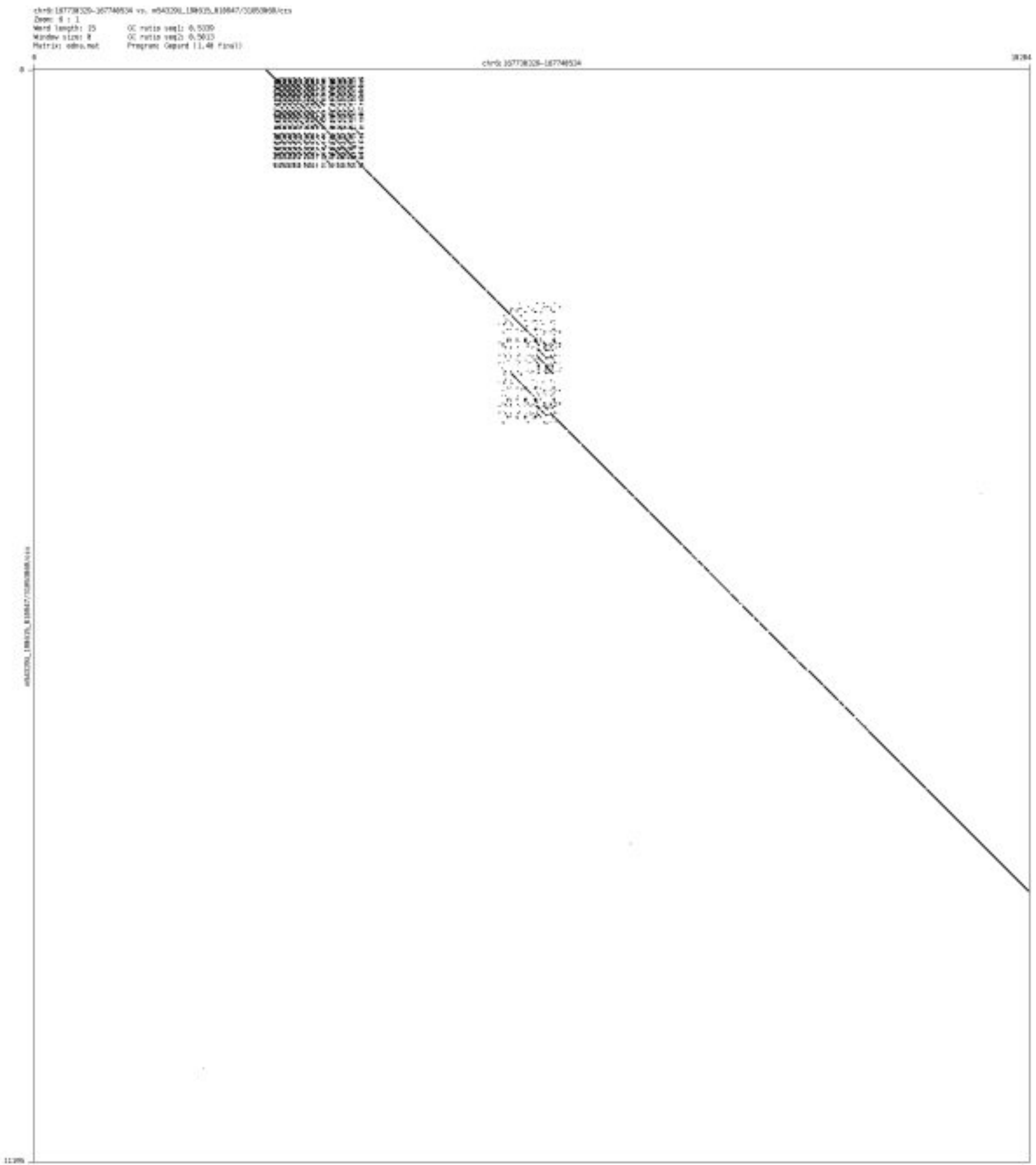

- ➢ chr6_170113638_170114174

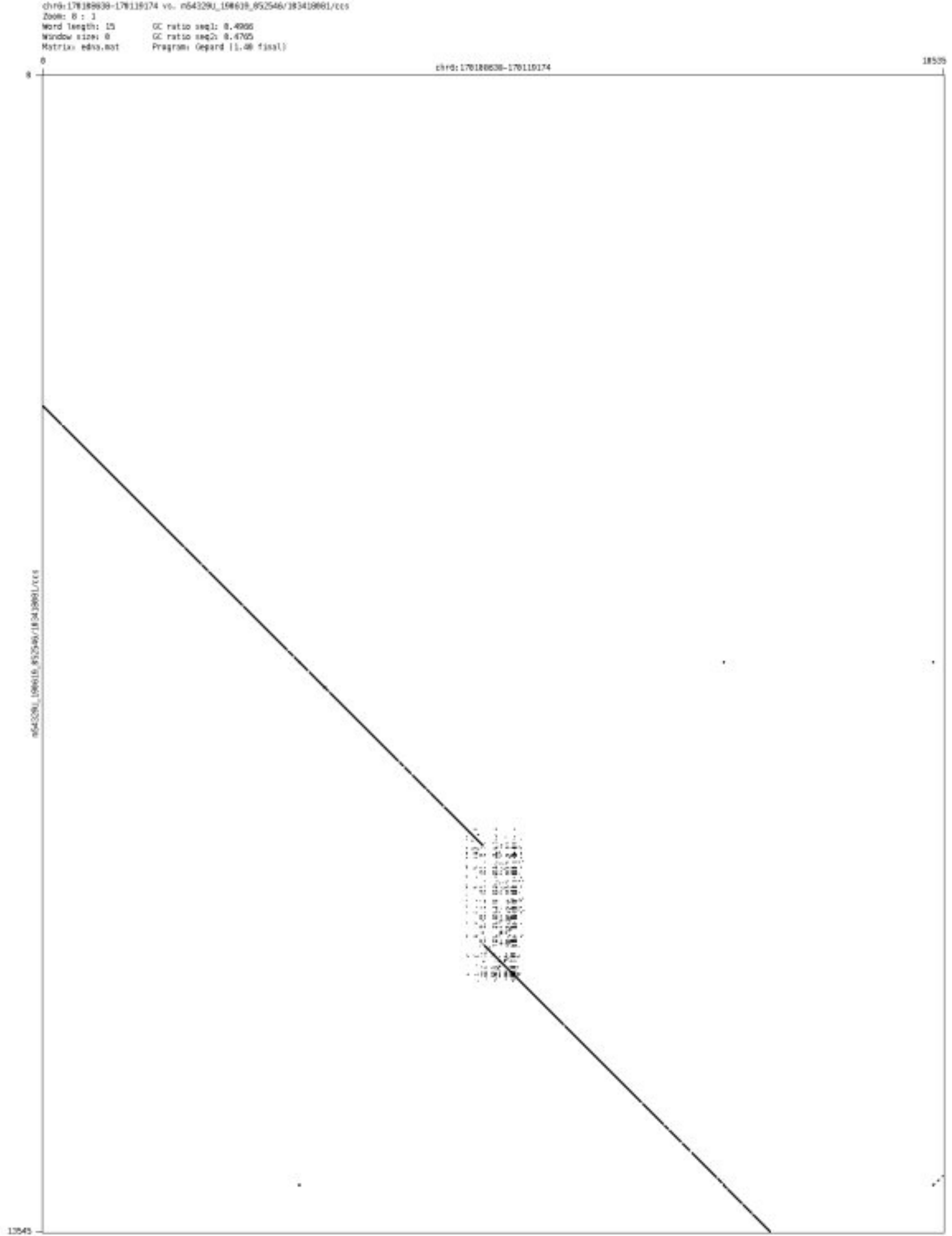

➢ chr7_45610820_45611071

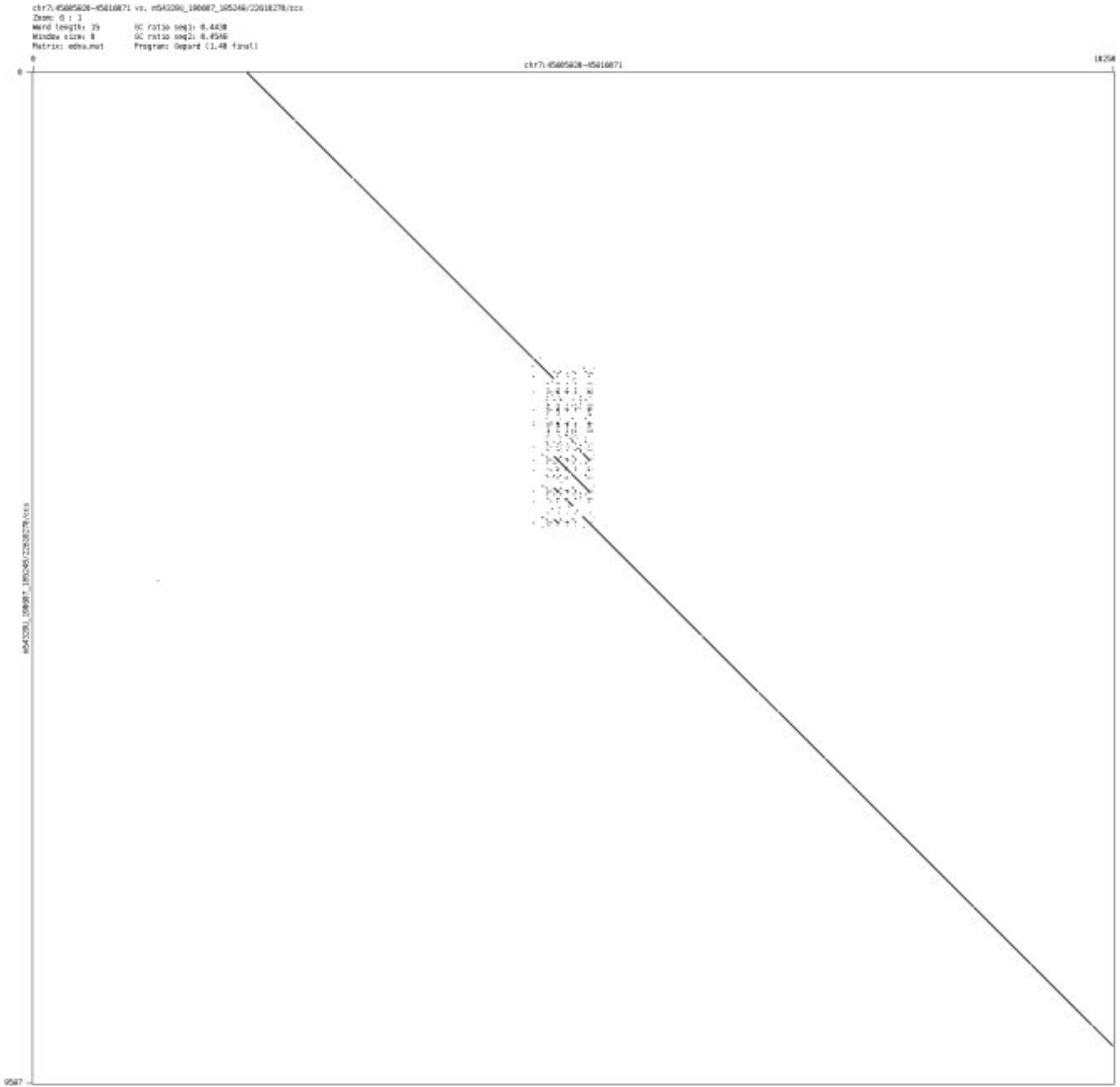

- chr7_76997407_76998955

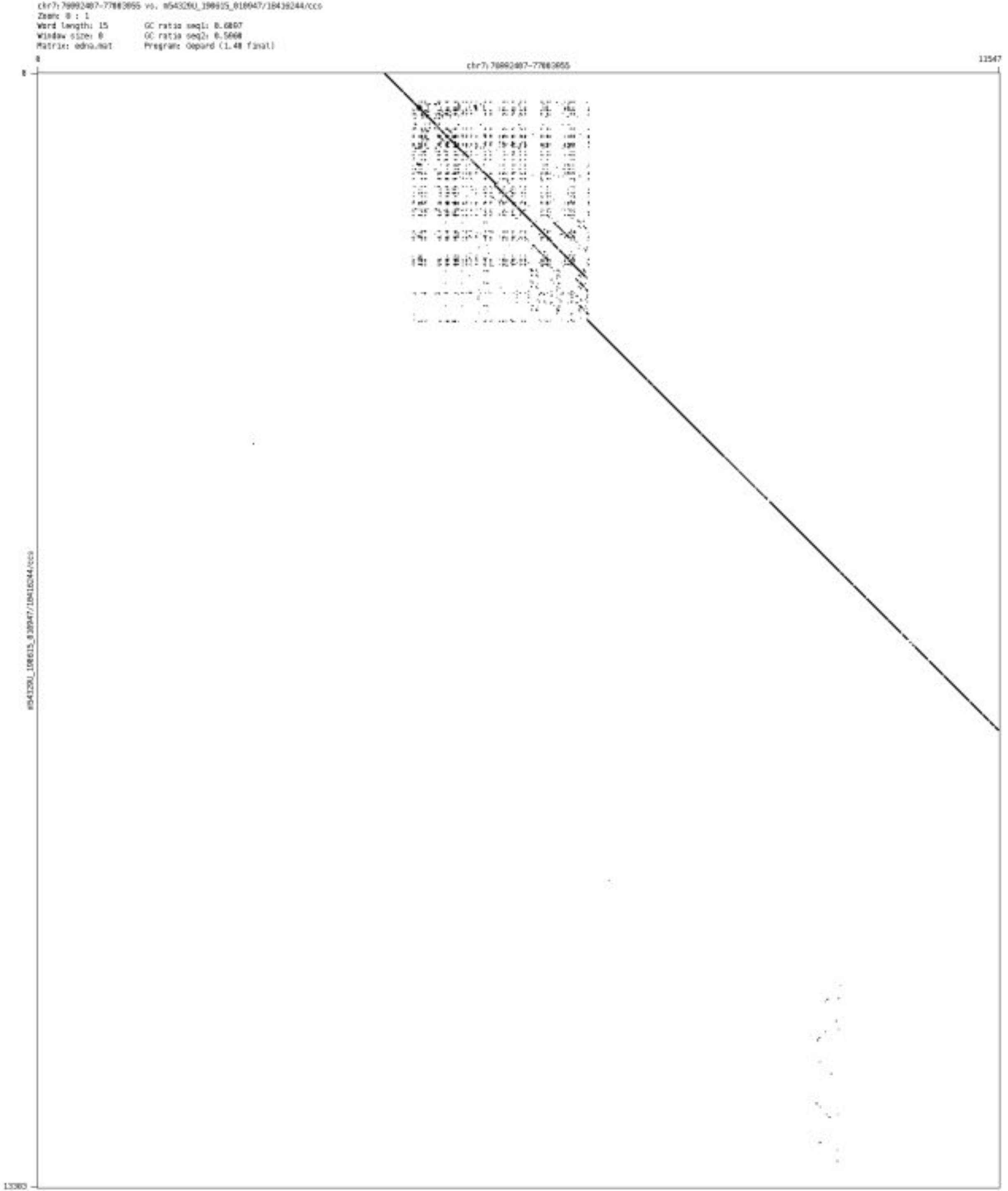

- chr7_98811382_98811519

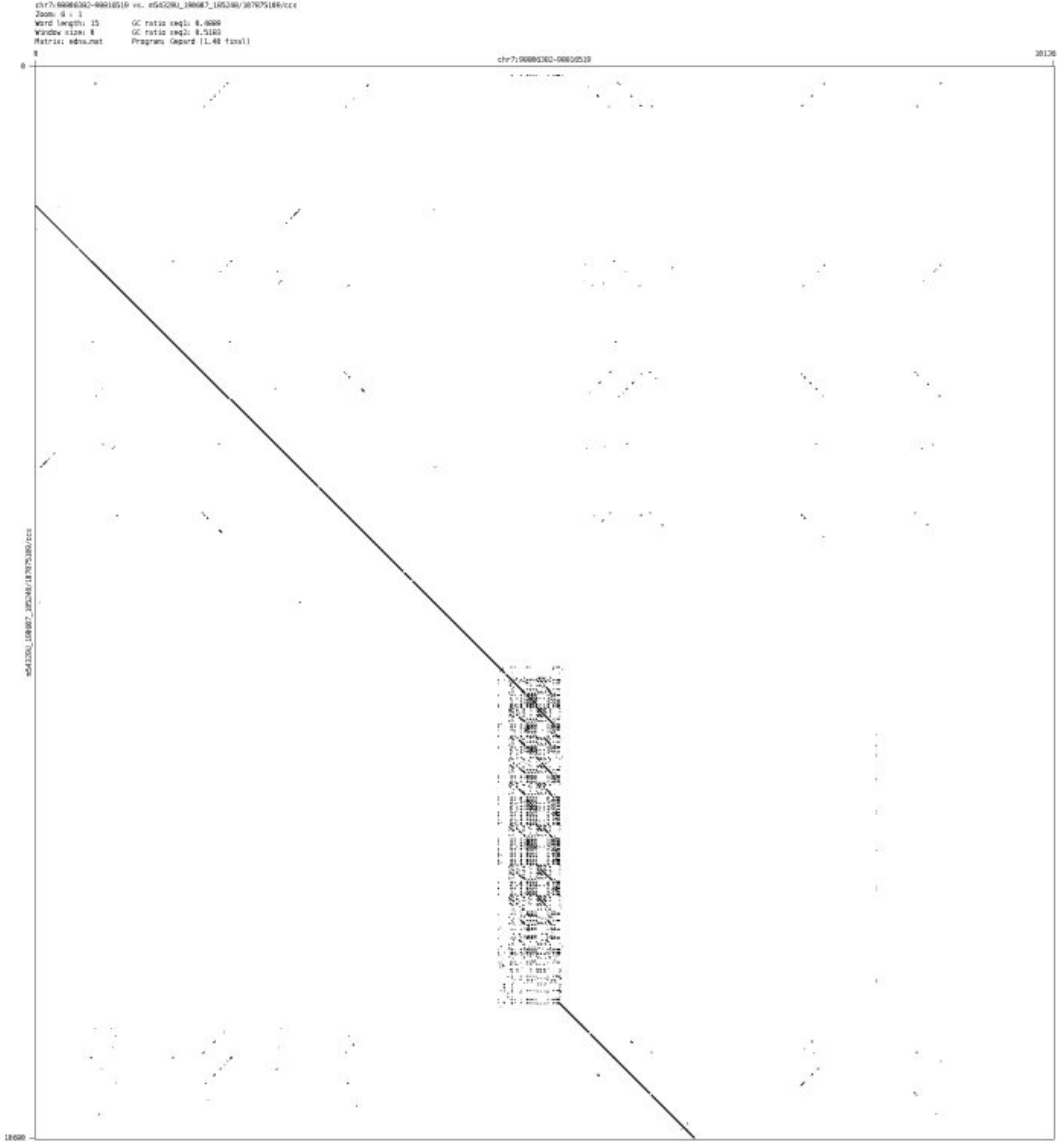

- ➢ chr7_100957703_100958508

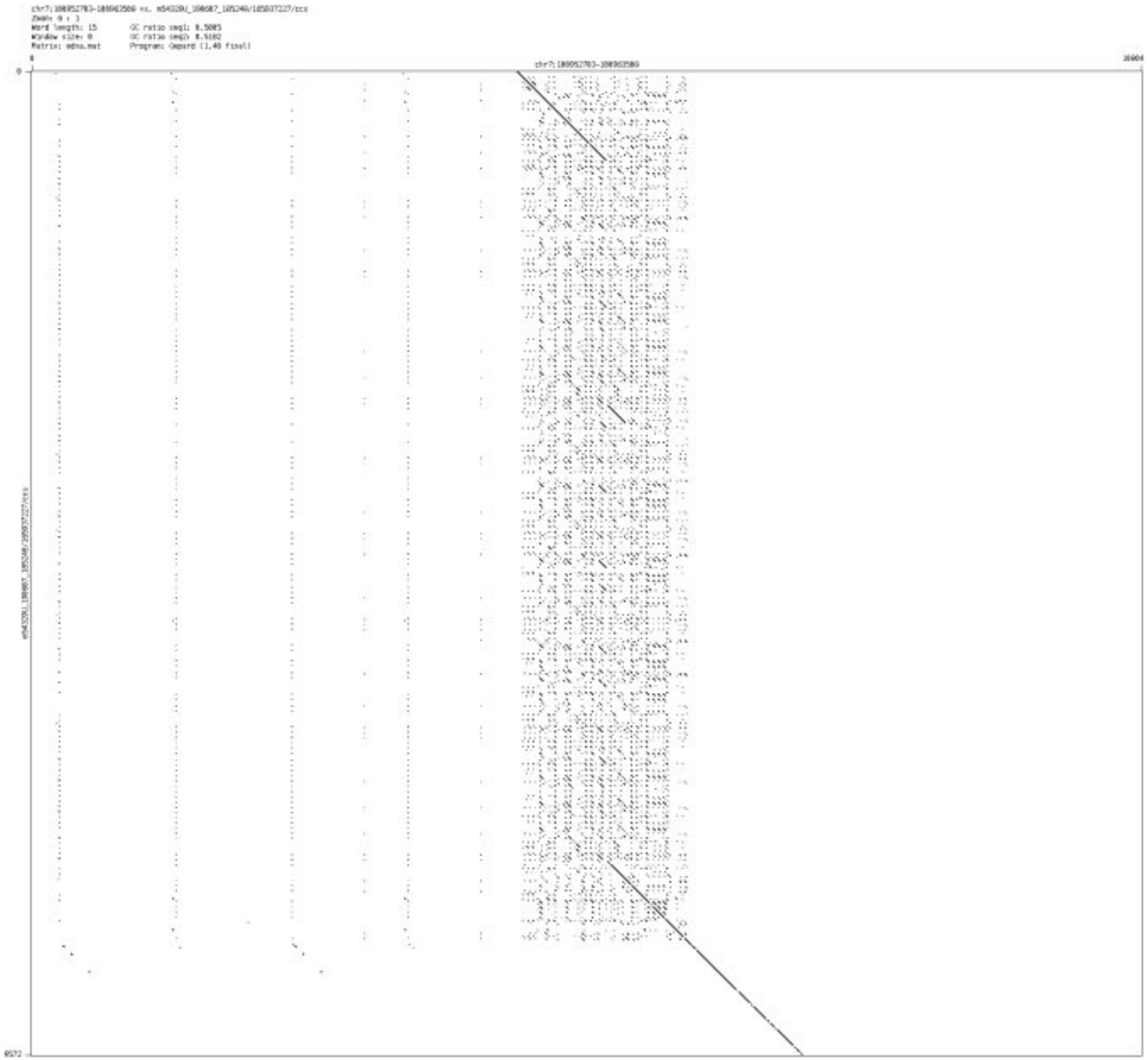

- chr7_155207777_155208913

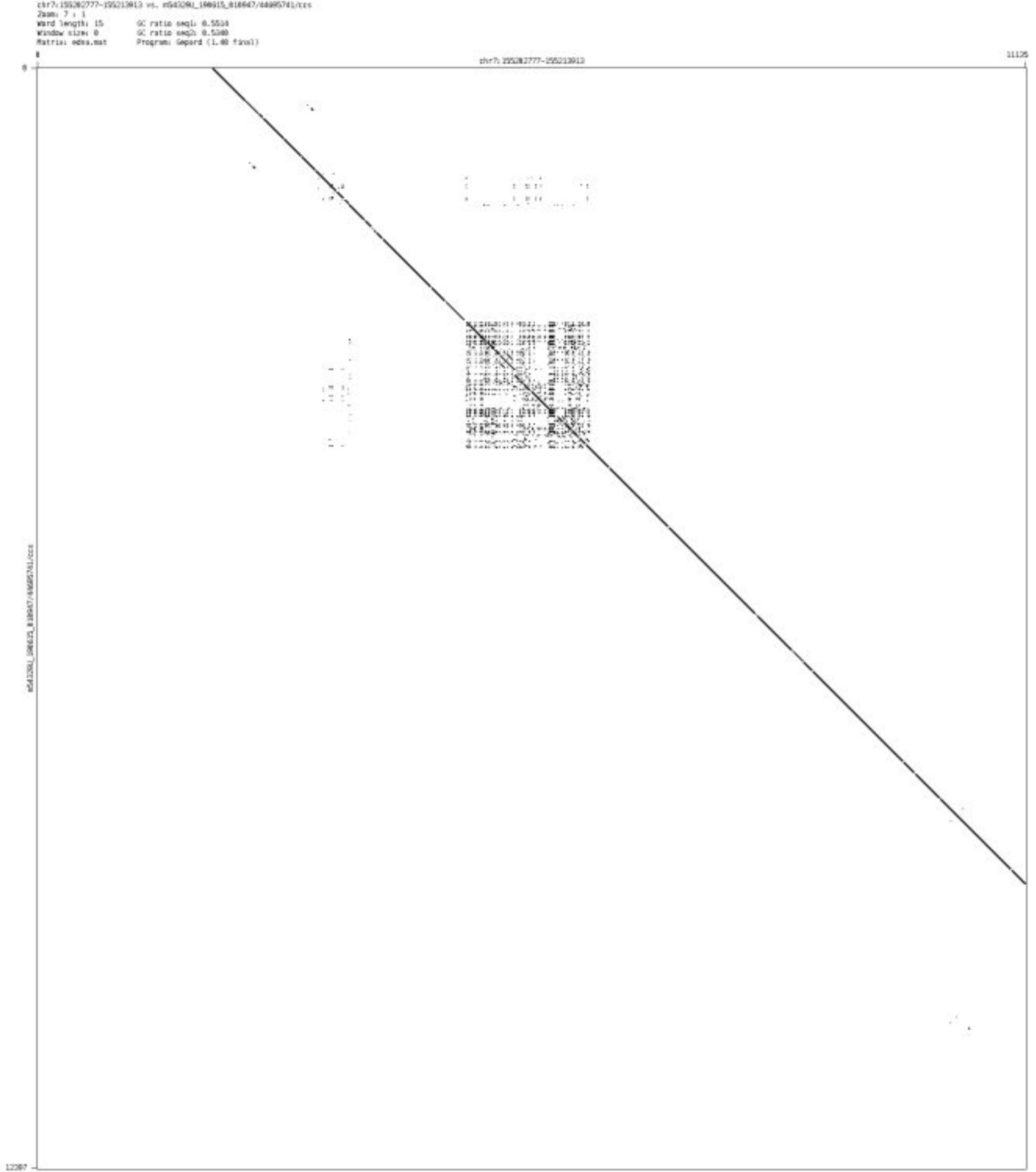

➢ chr7_155792027_155792109

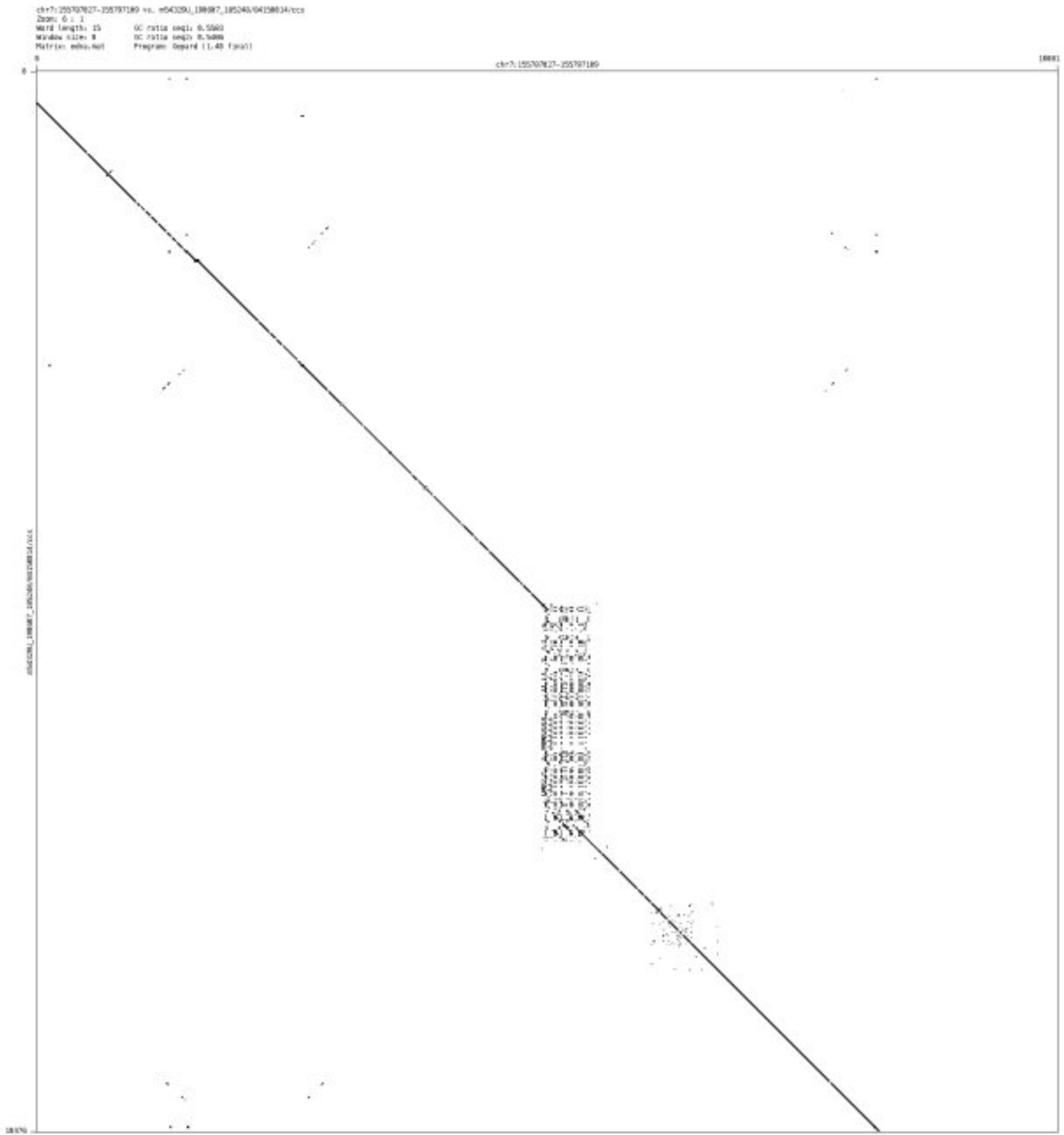

➢ chr8_138206296_138206471

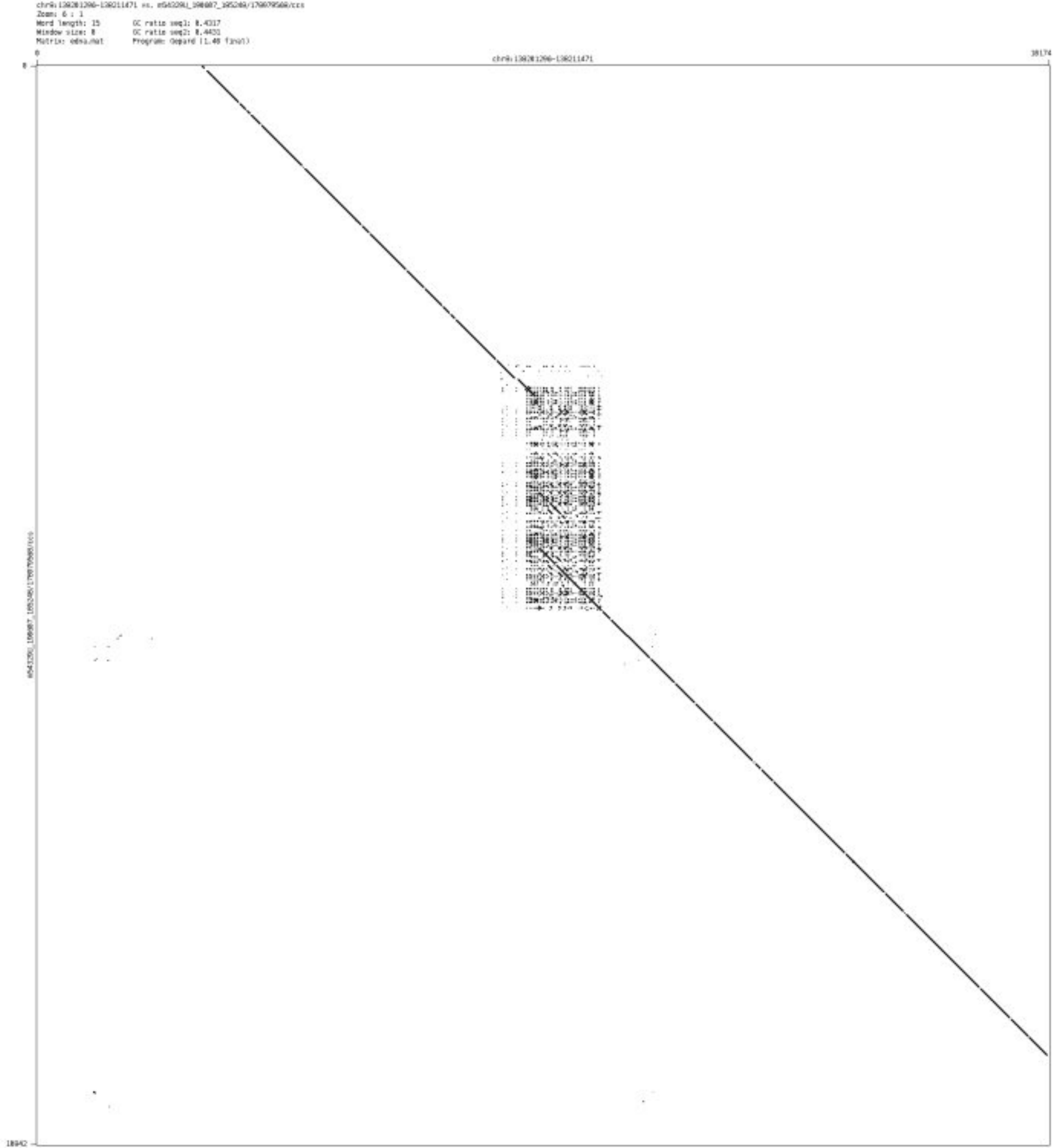

- chr9_4713910_4714160

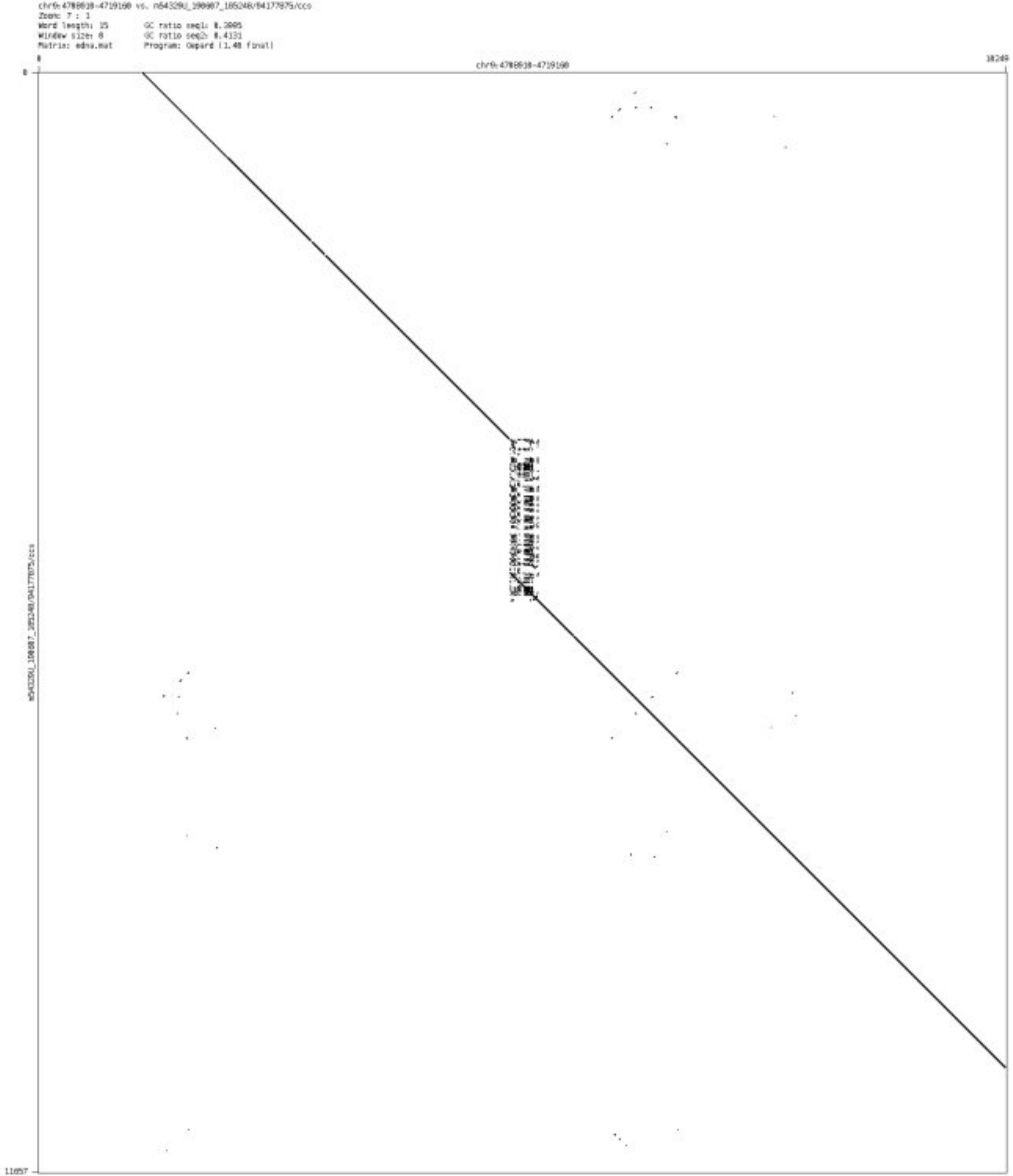

- chr9_137669378_137669456

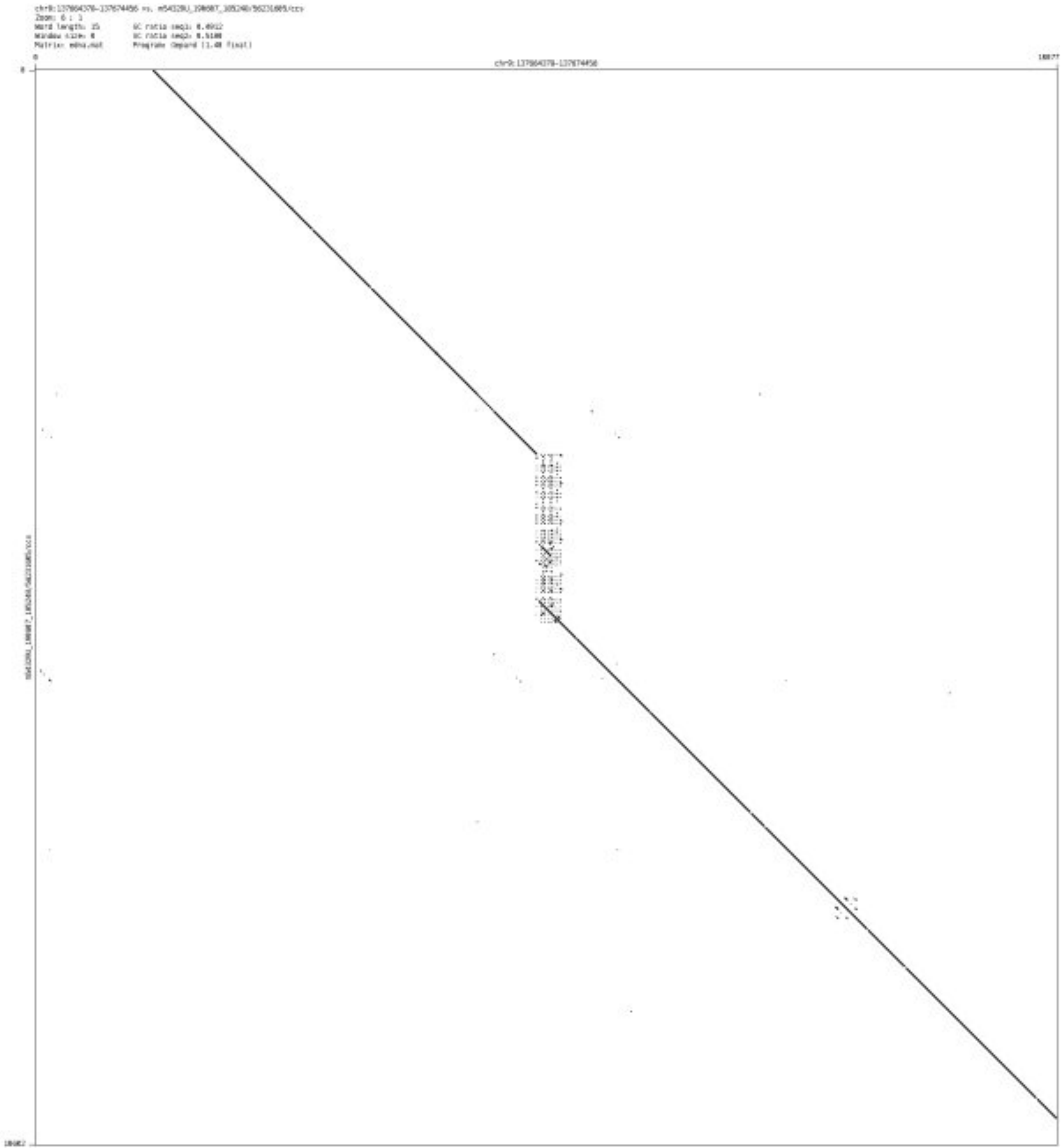

- chr10_124227125_124227273

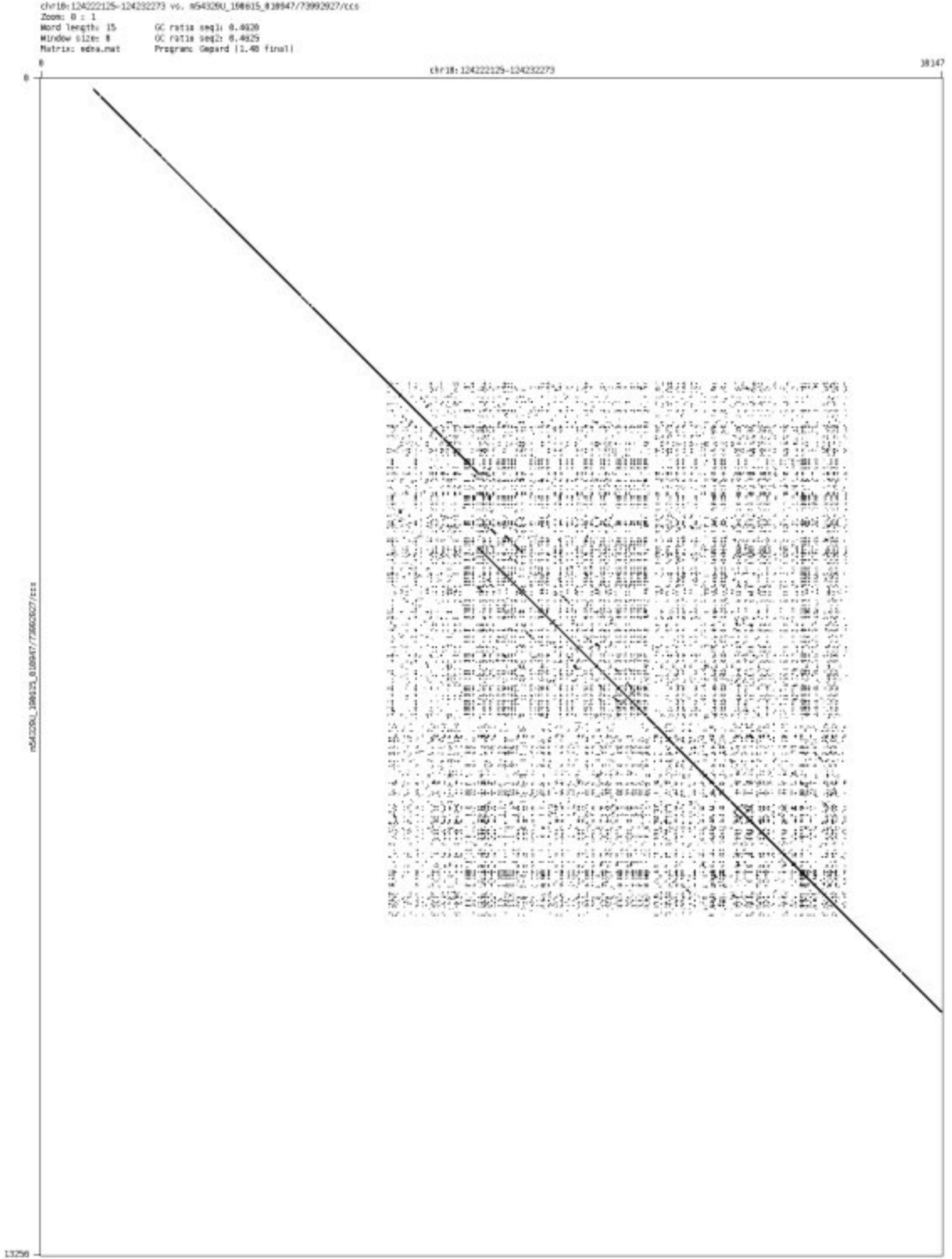

- chr10_130790080_130790697

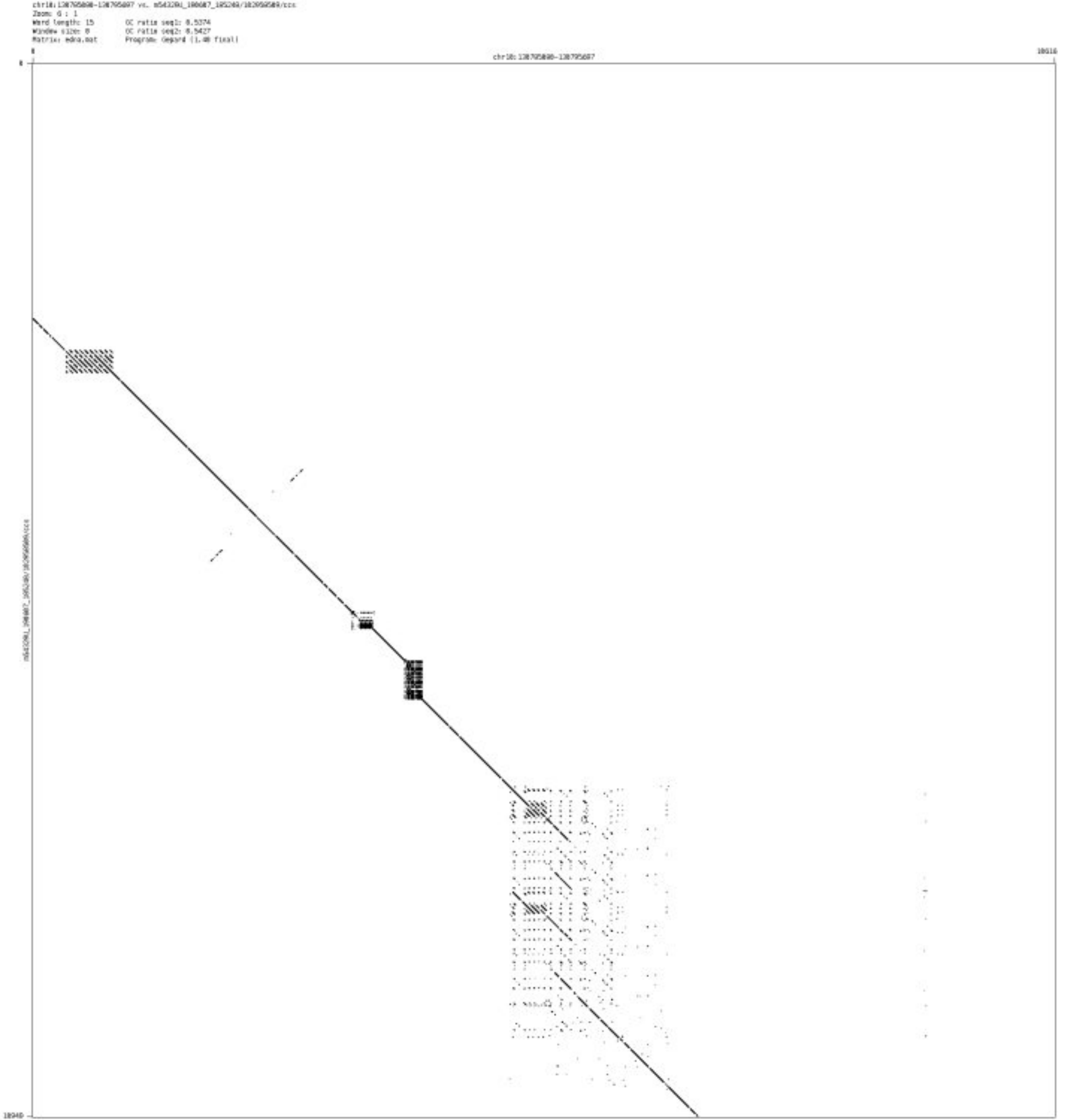

➢ chr10_131176490_131176898

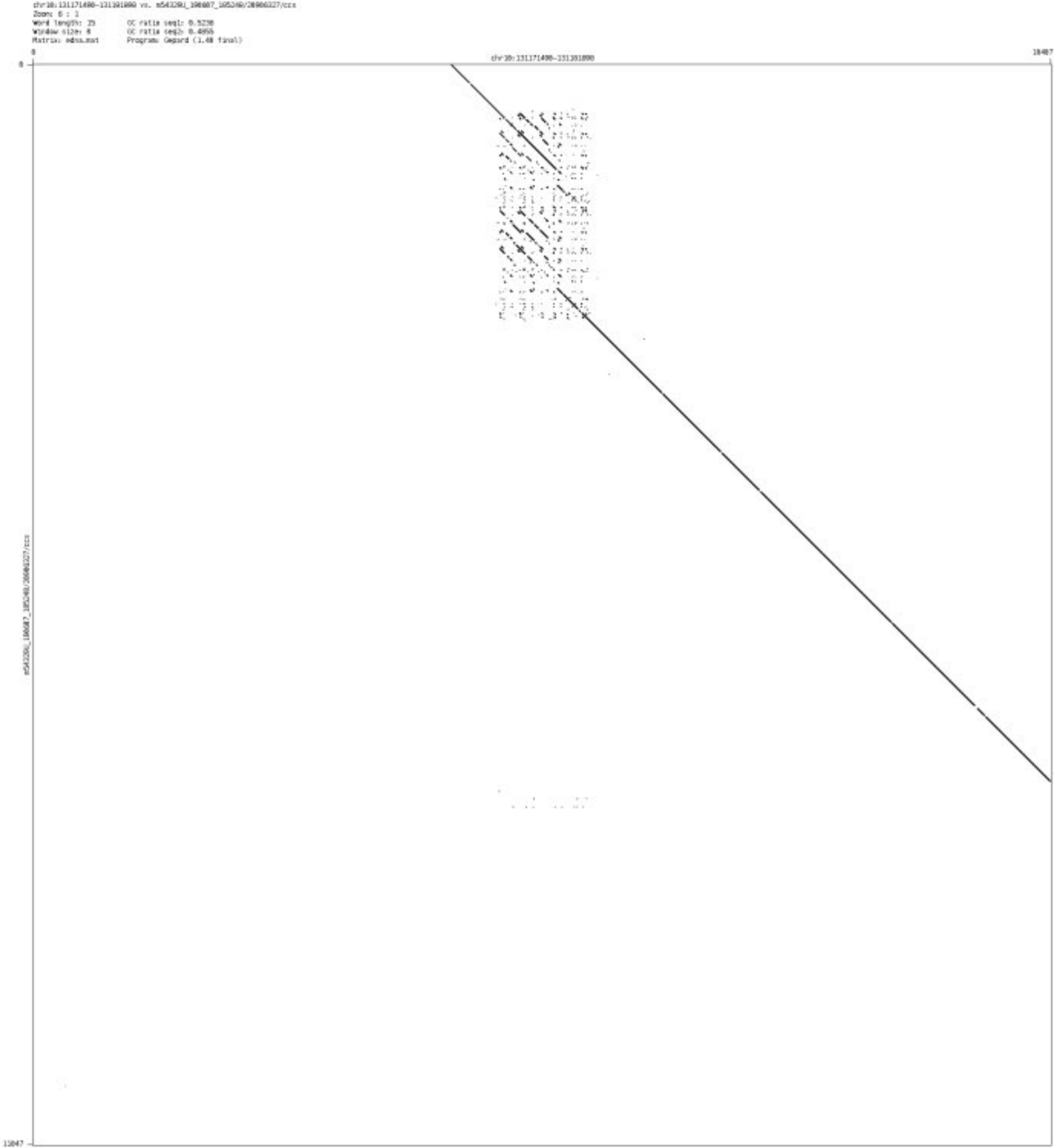

- chr10_131486500_131486696

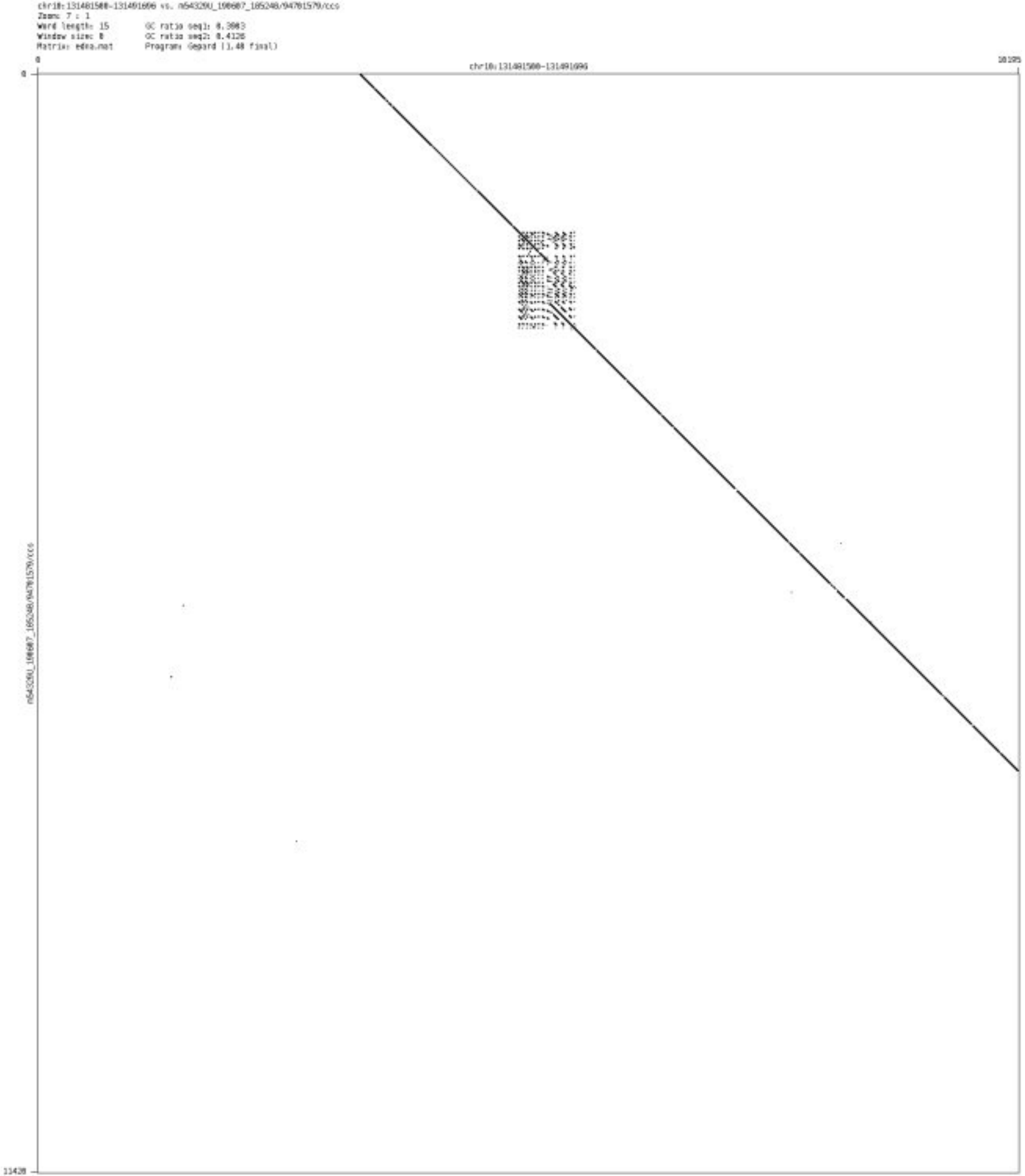

➢ chr11_68963986_68964114

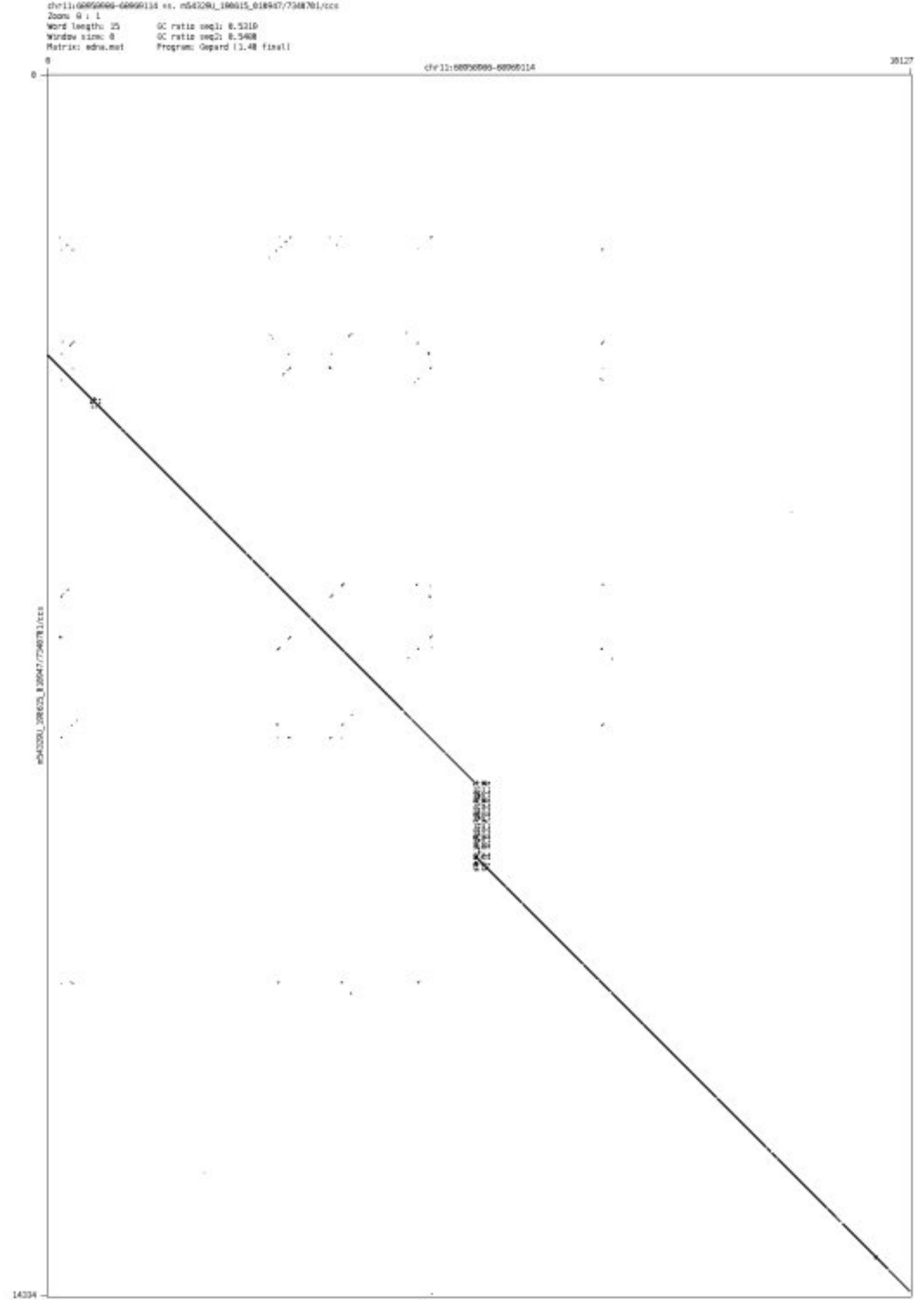

- chr11_95633307_95634471

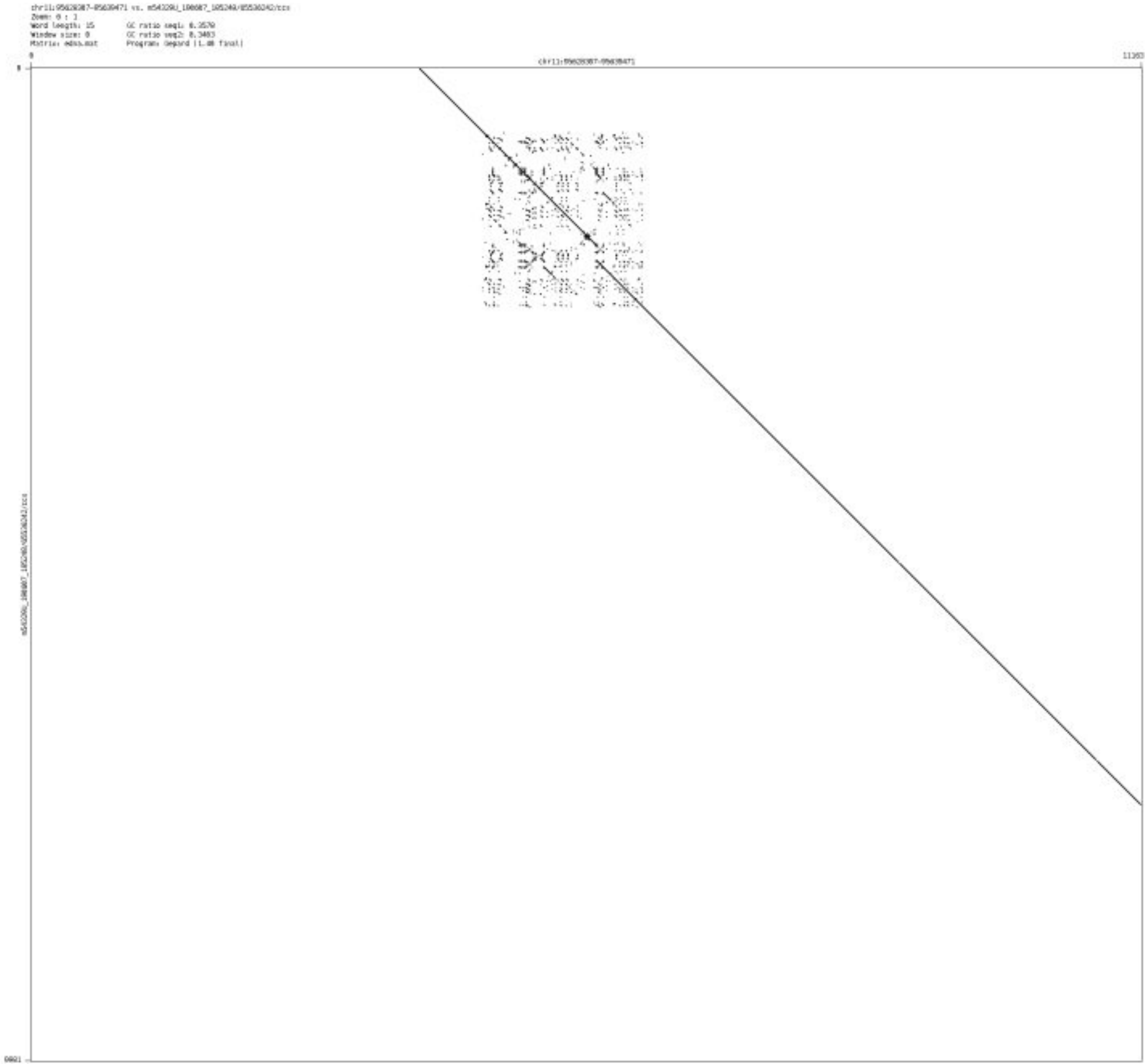

➢ chr12_120927507_120928364

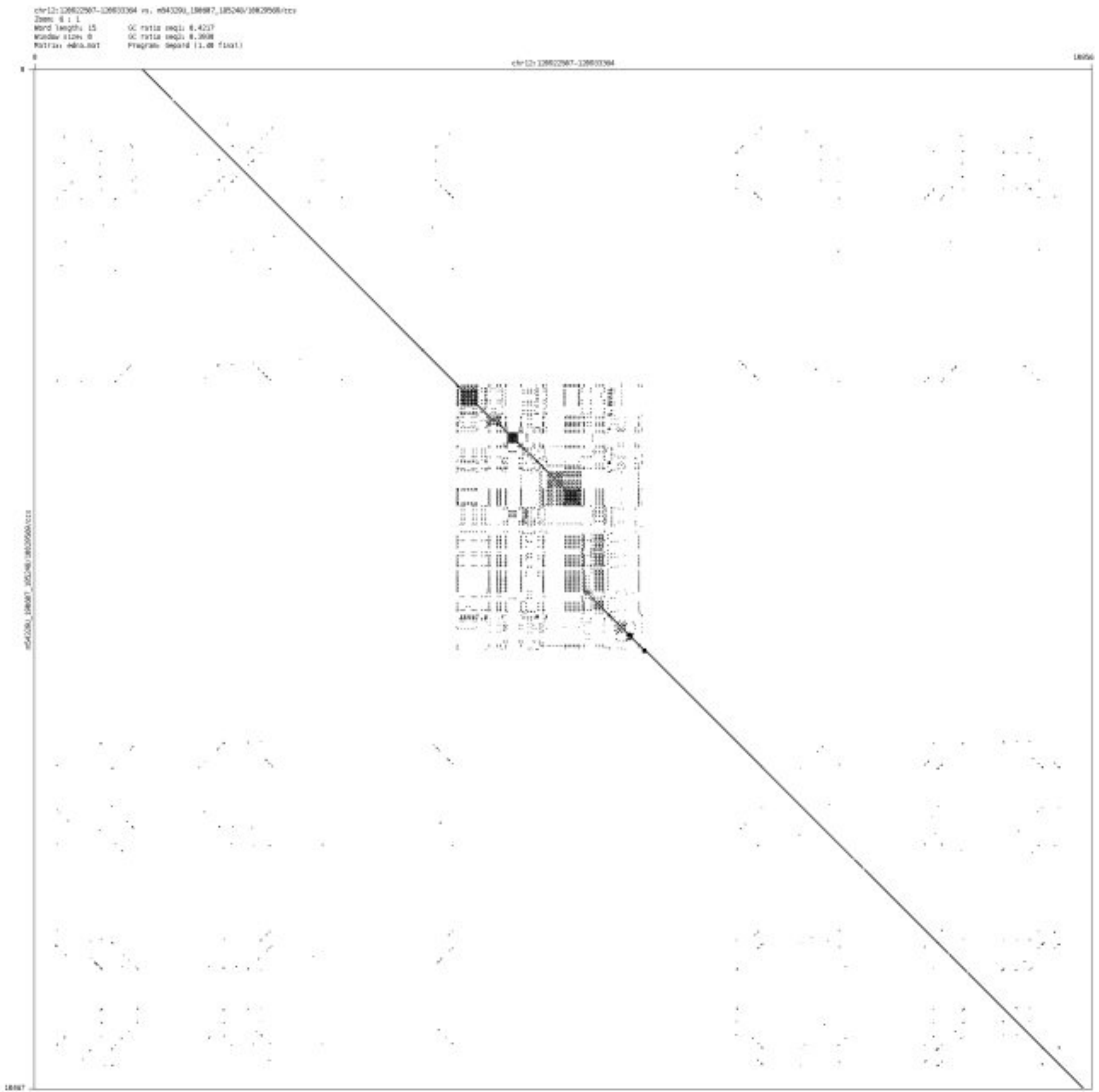

➢ chr13_35956622_35958426

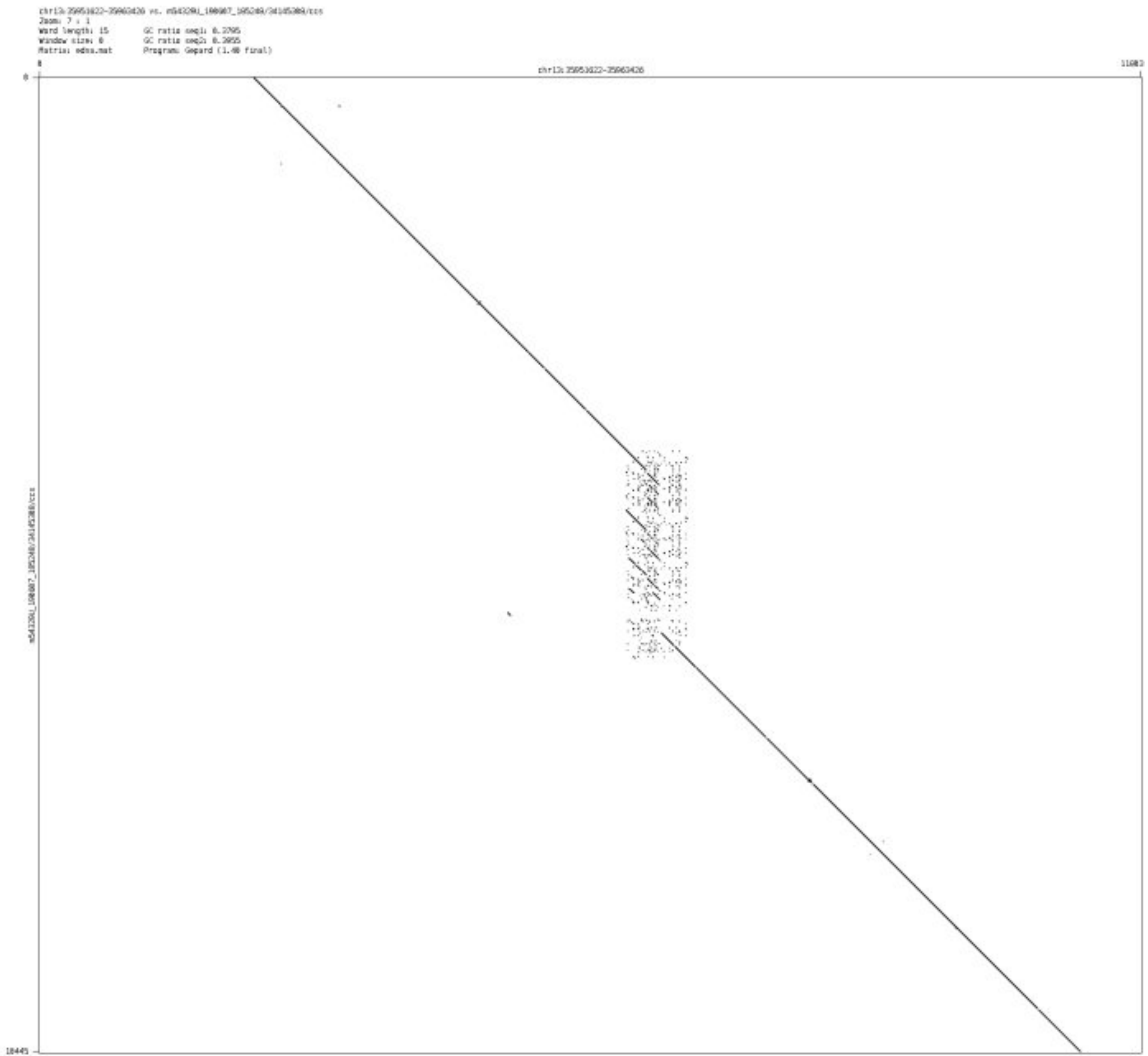

- chr13_111568050_111568256

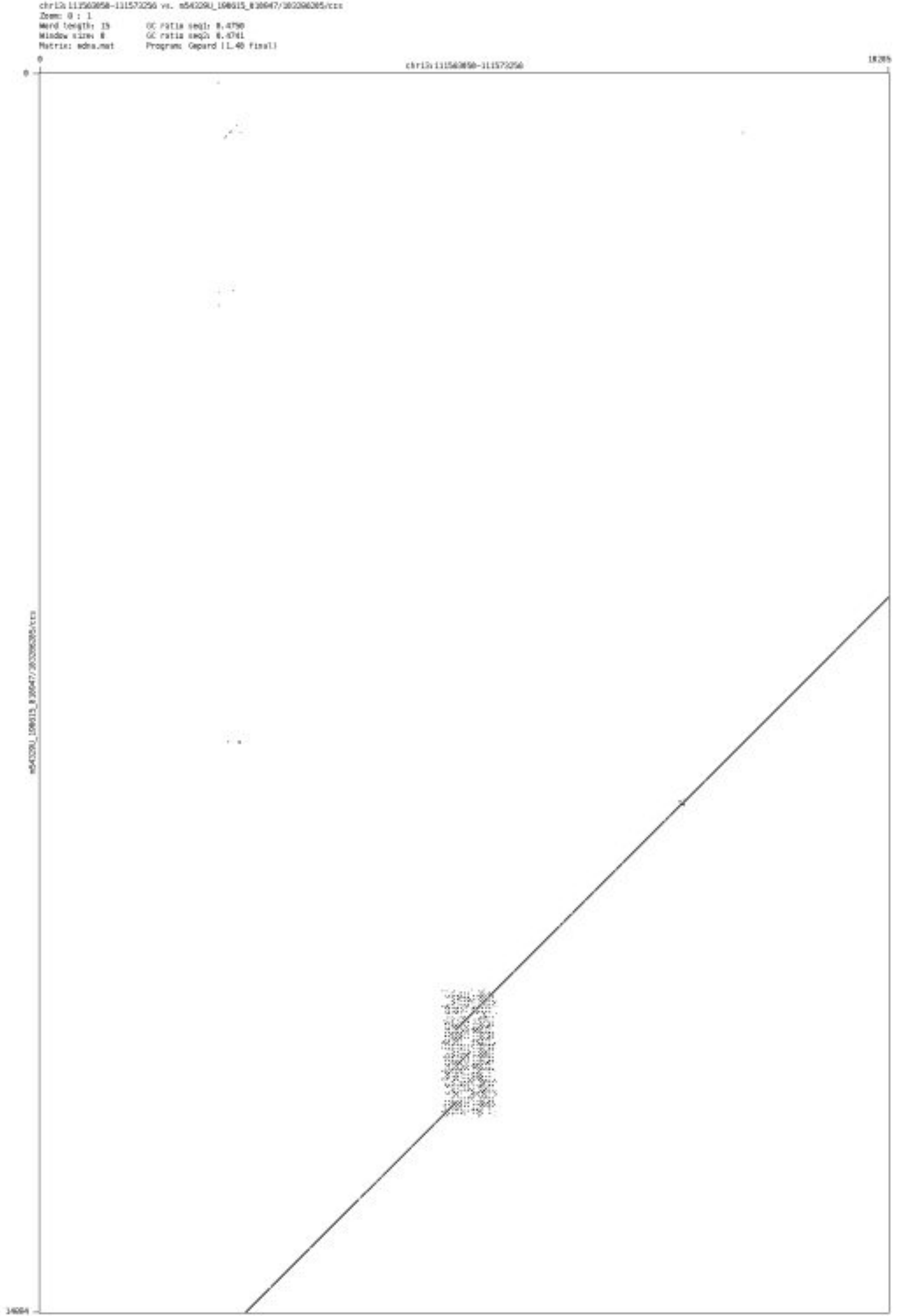

- chr15_100710334_100710411

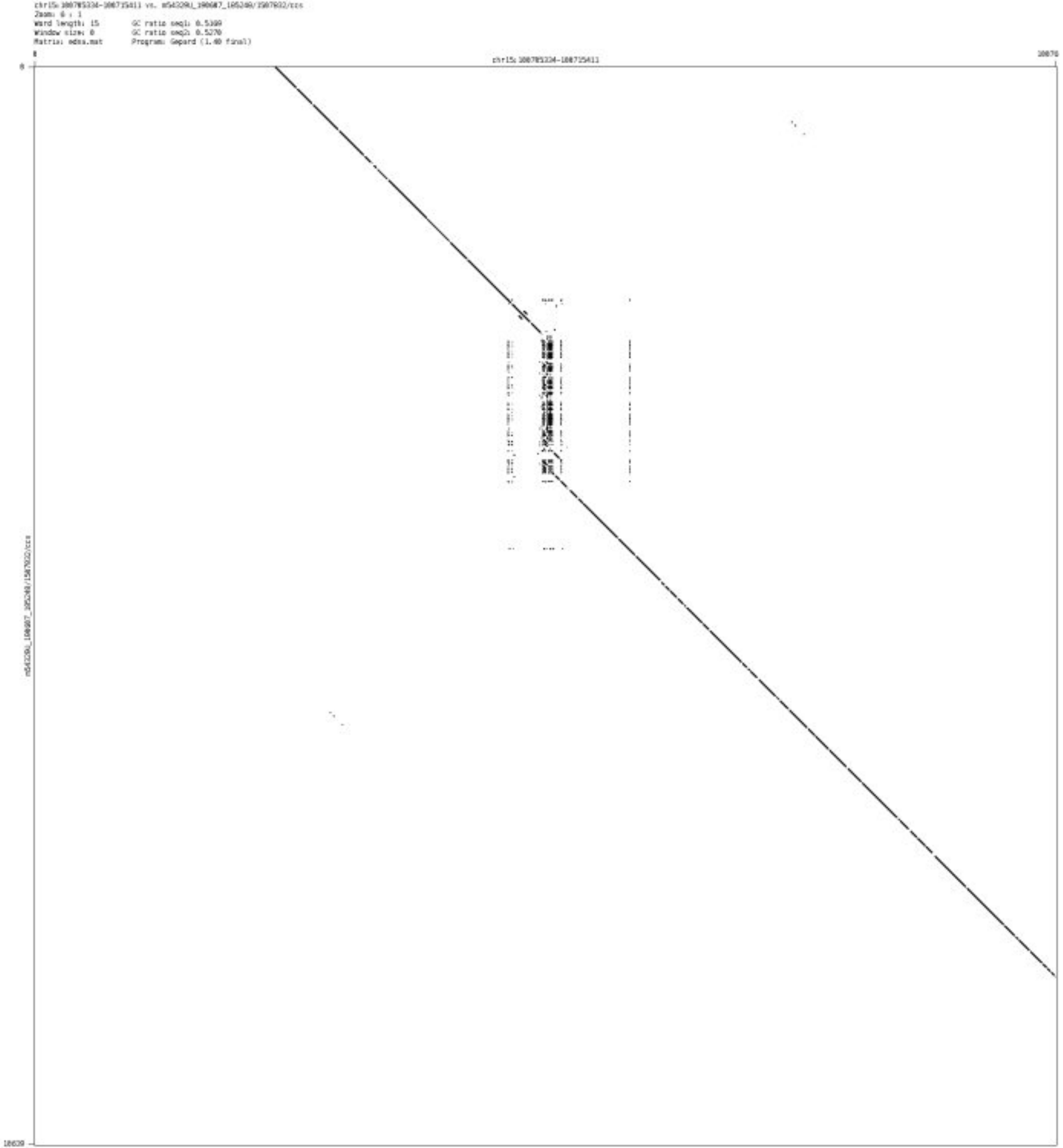

➢ chr16_894258_894382

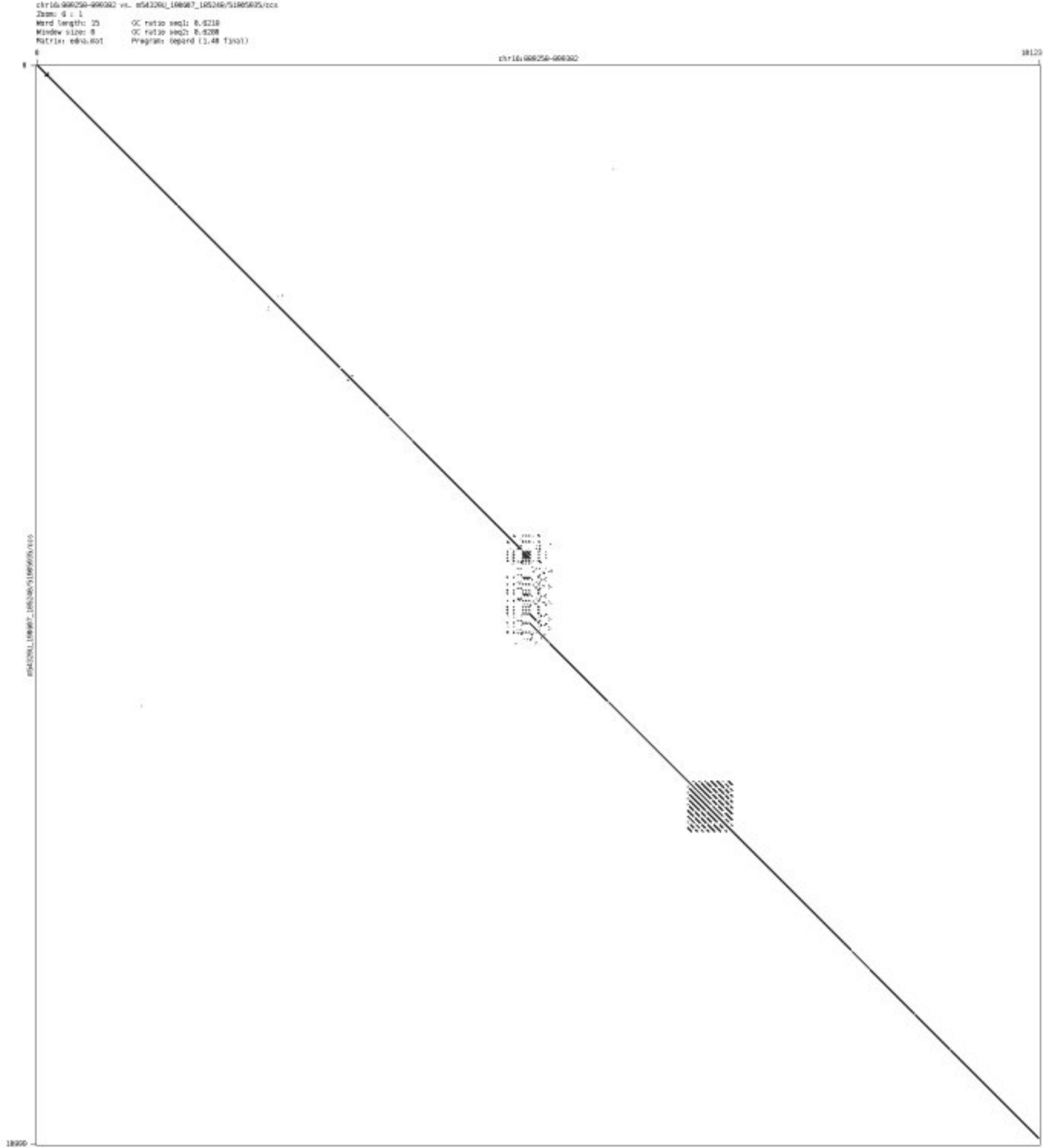

➢ chr16_86464399_86464597

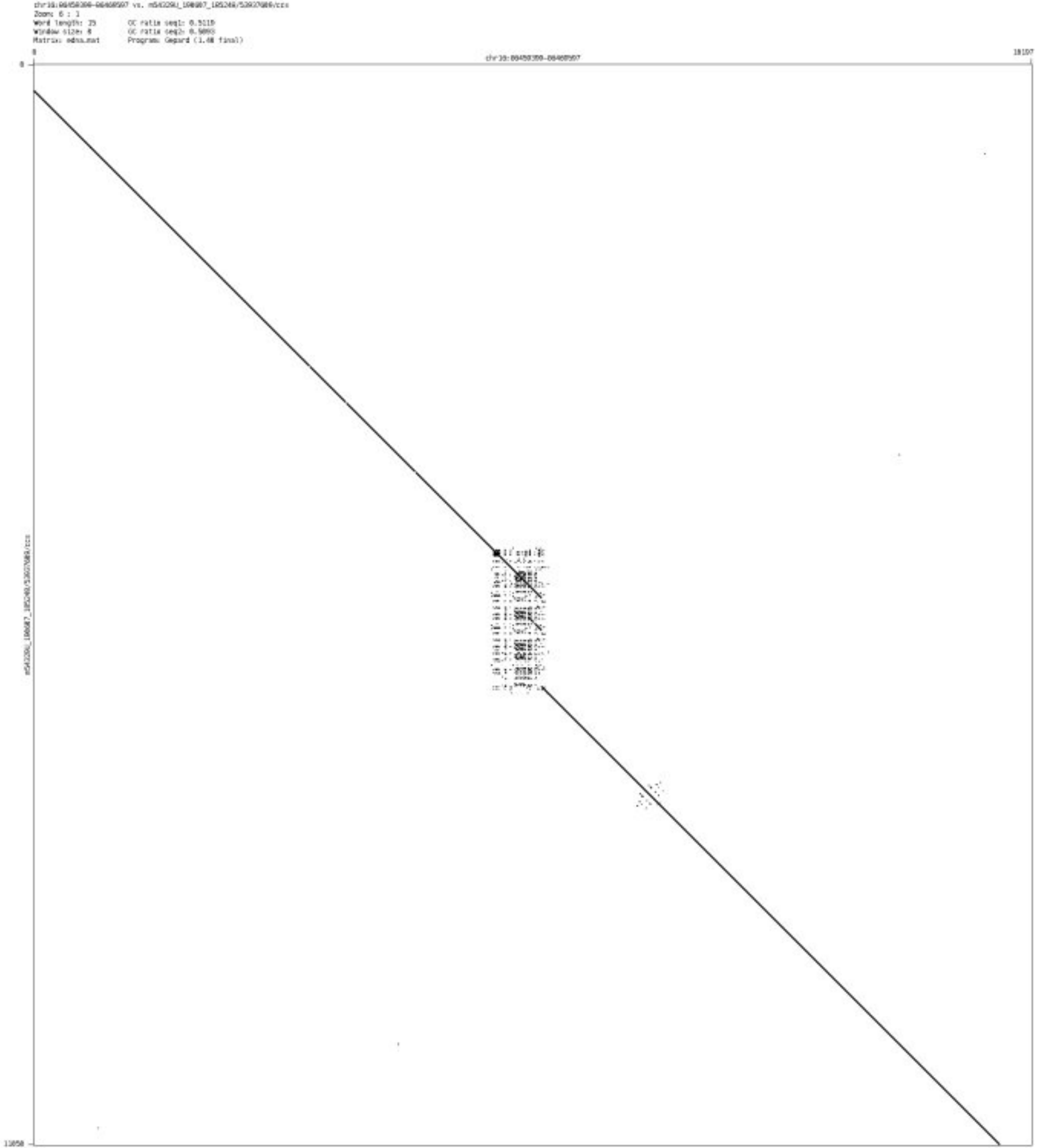

➢ chr16_87145221_87145335

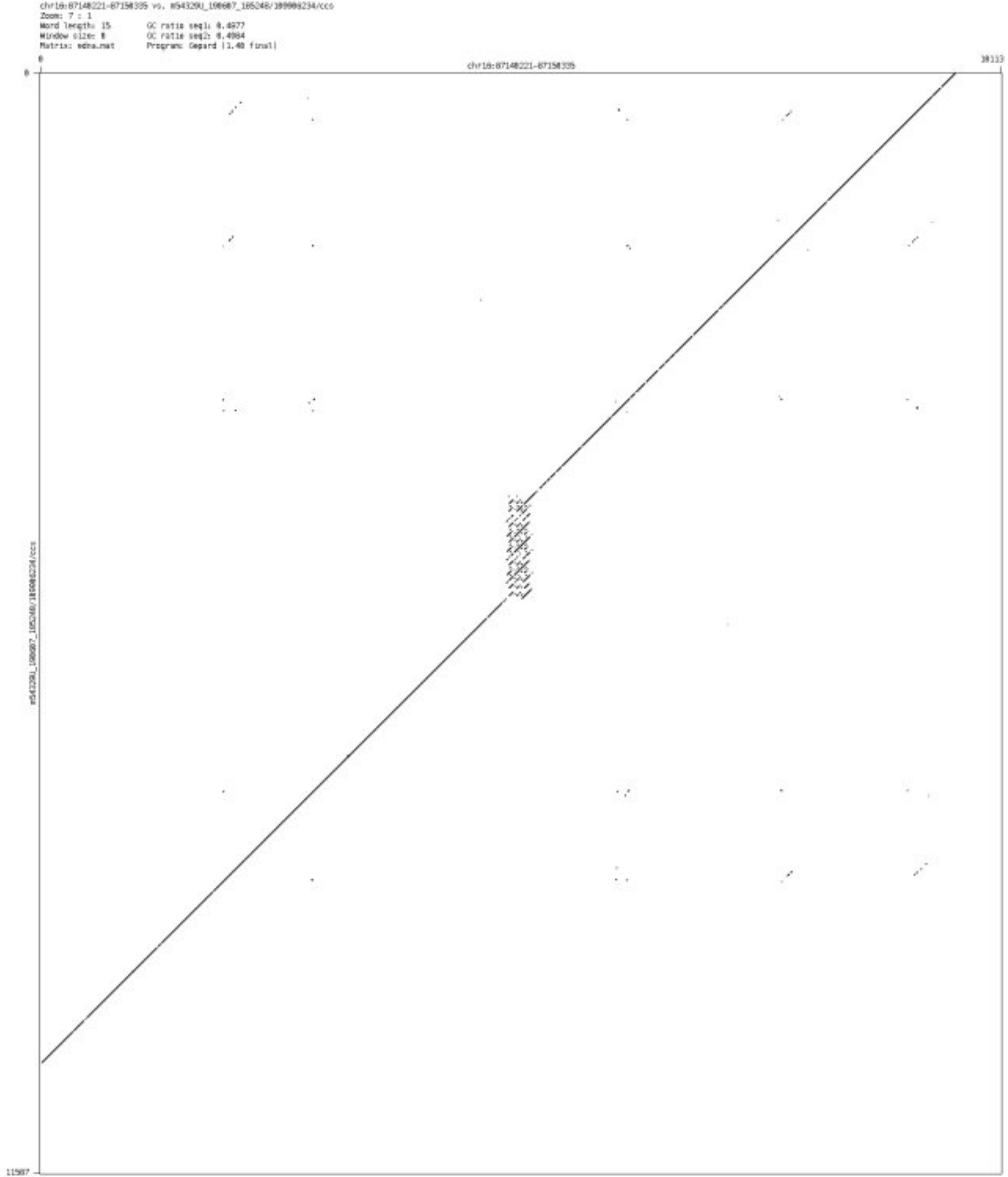

- chr17_65499952_65500292

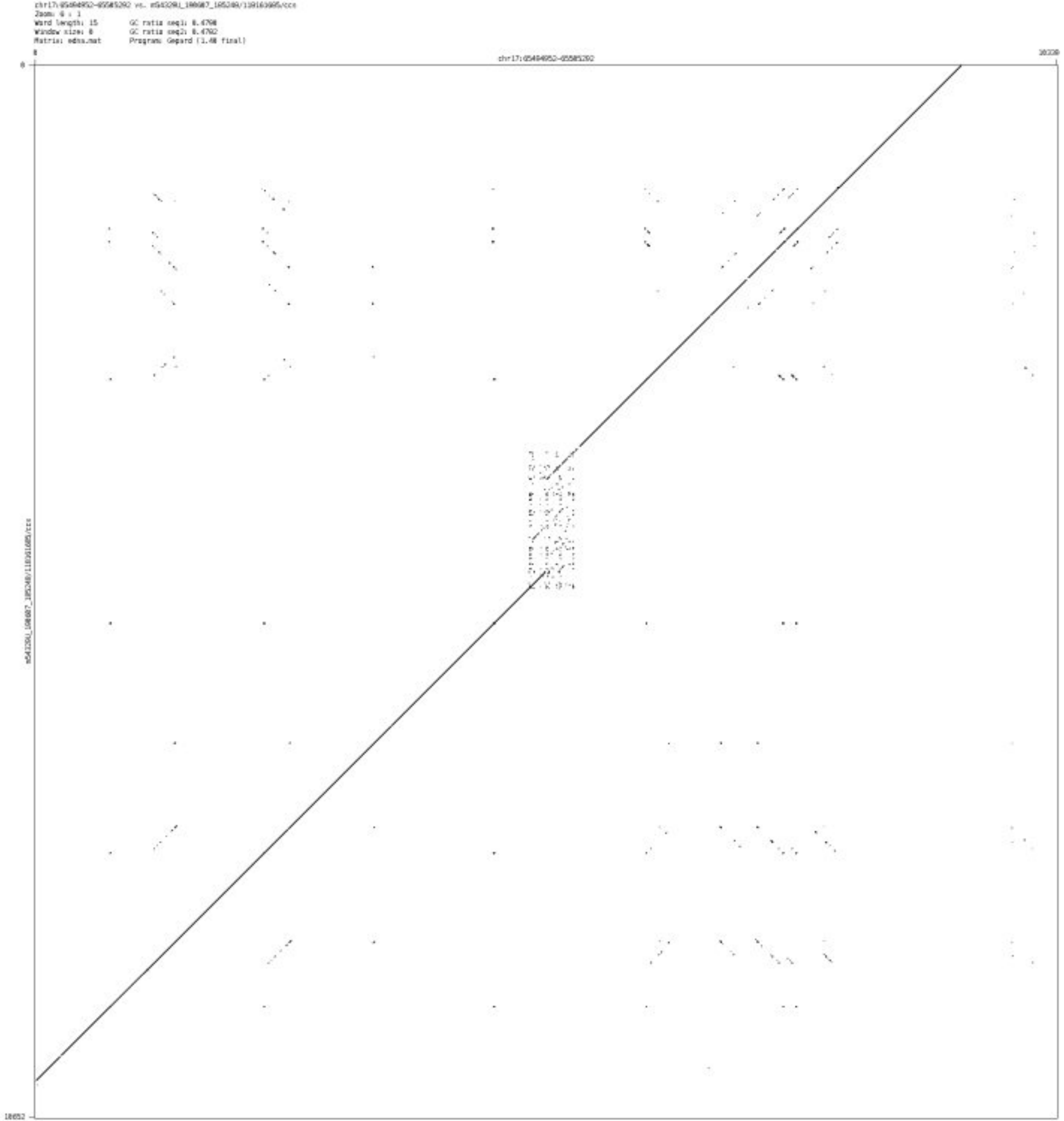

➢ chr17_79555509_79555782

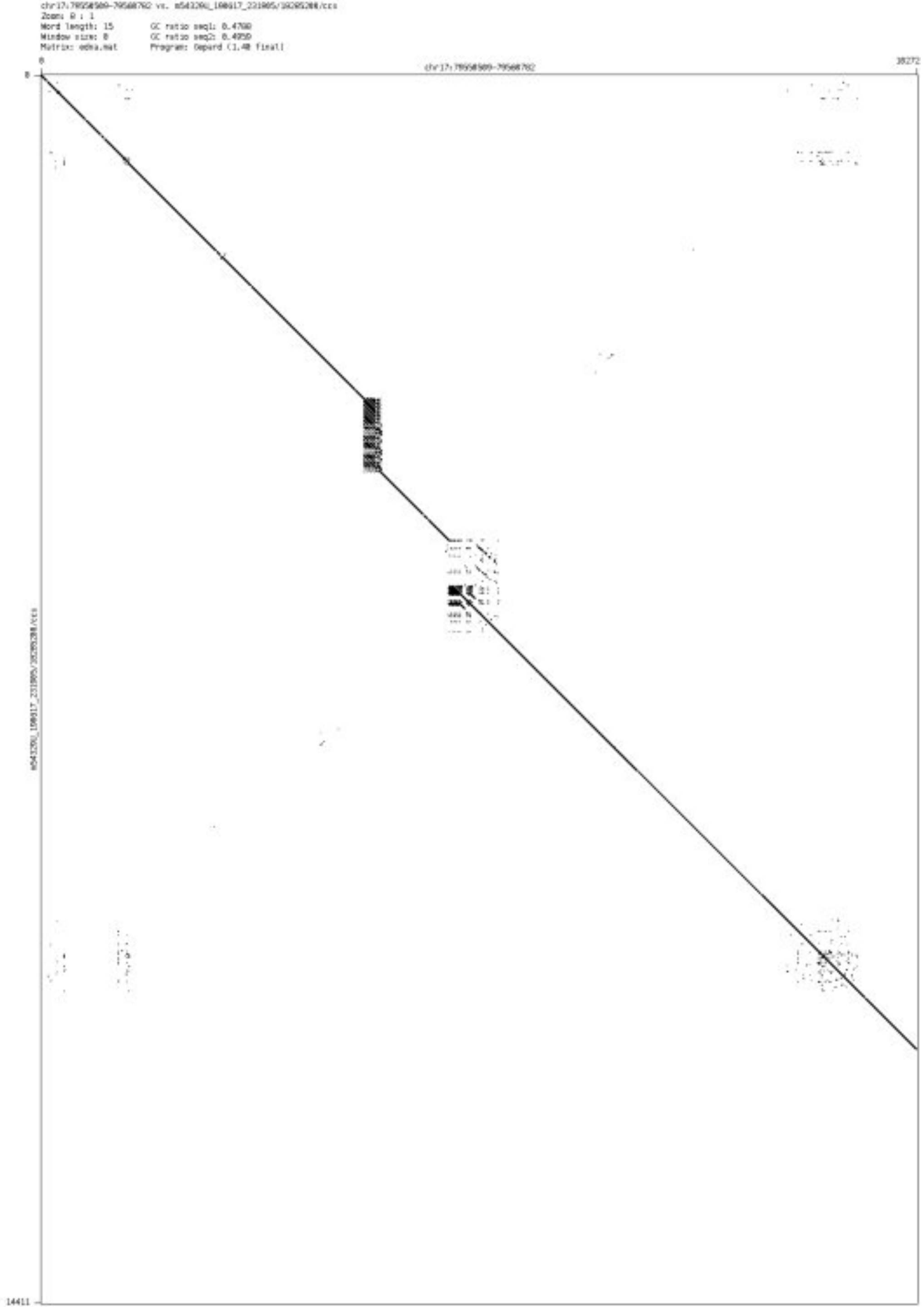

- chr17_80024327_80024606

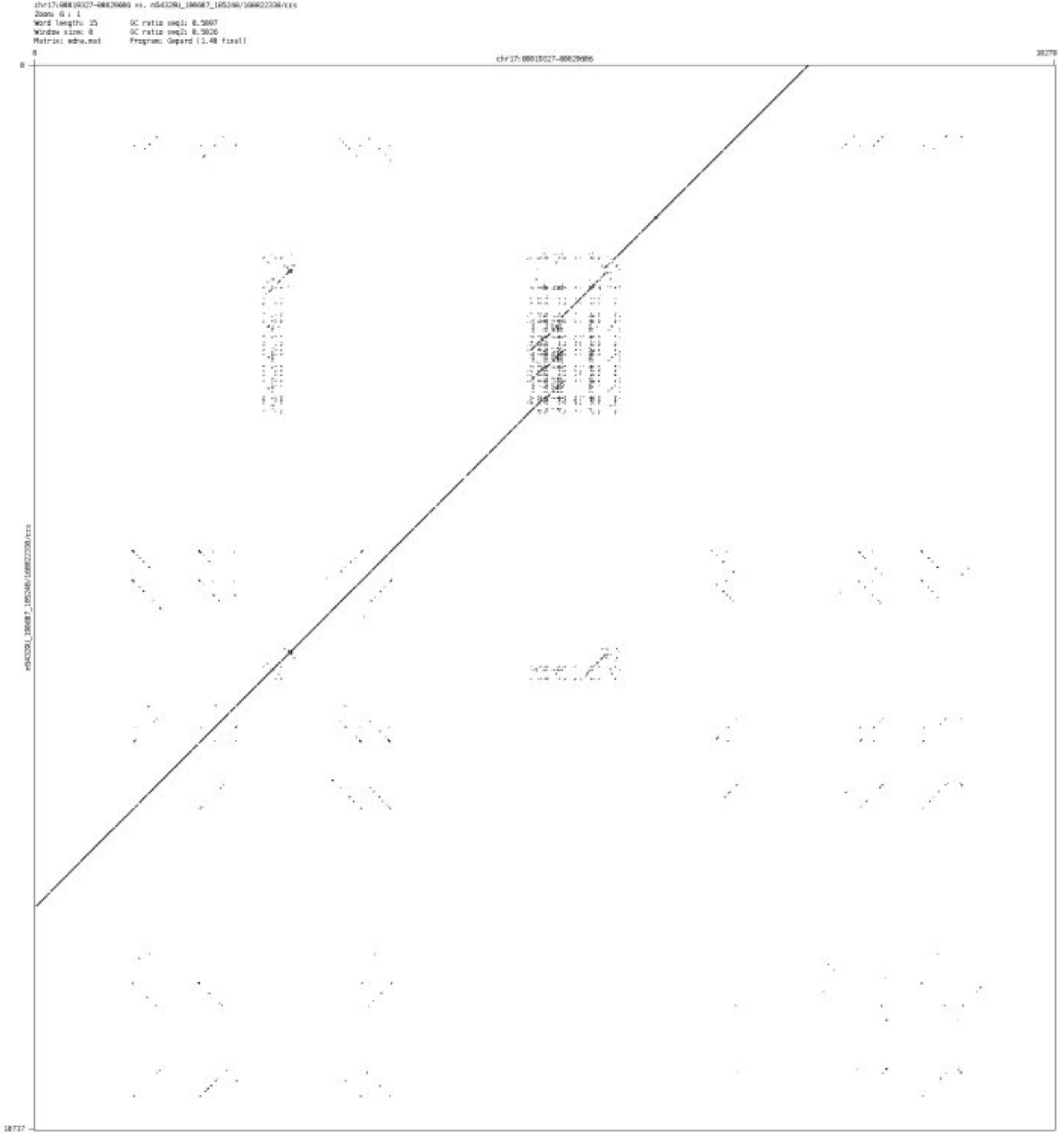

➢ chr18_78010192_78010318

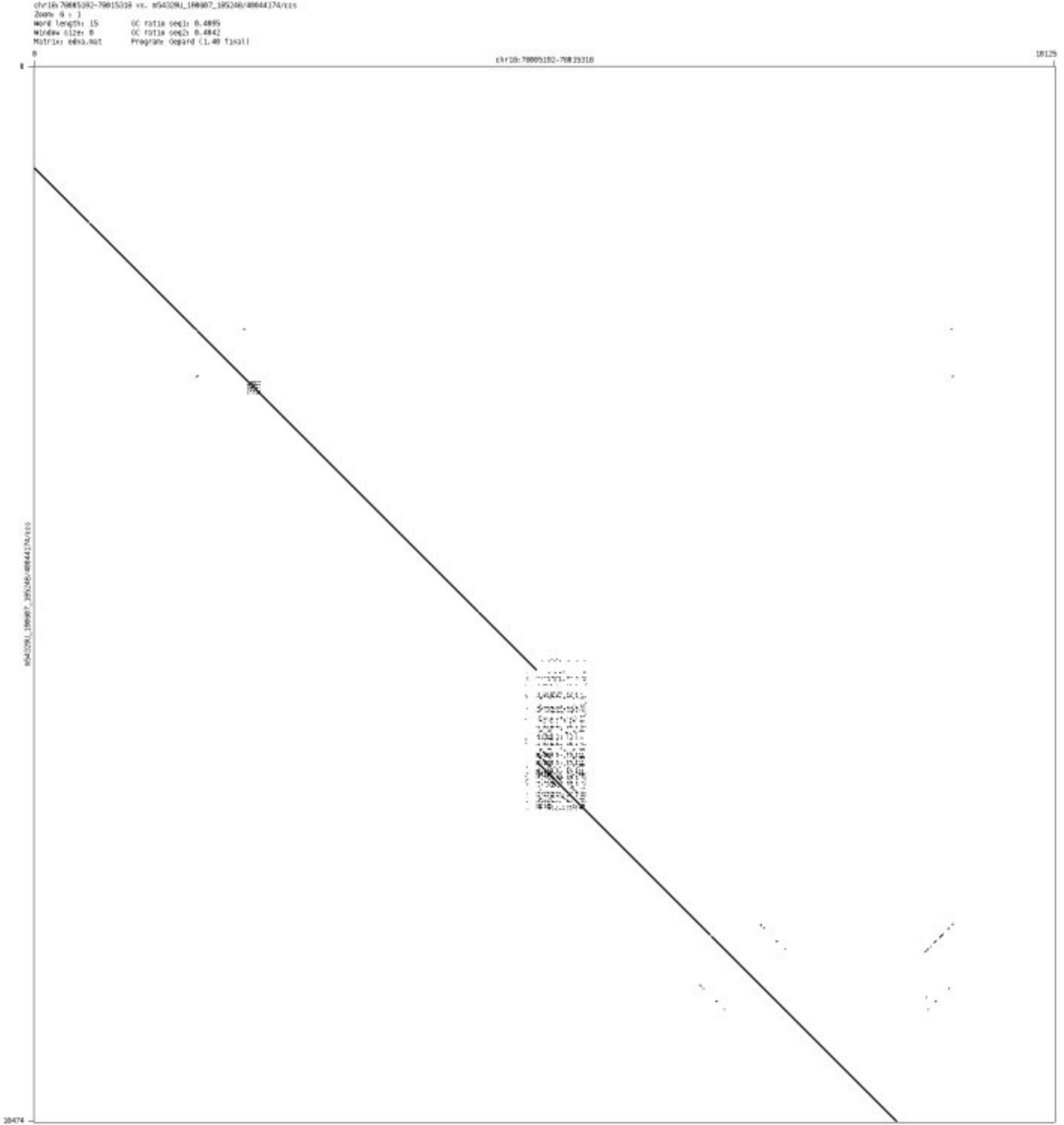

- chr19_1142124_1142241

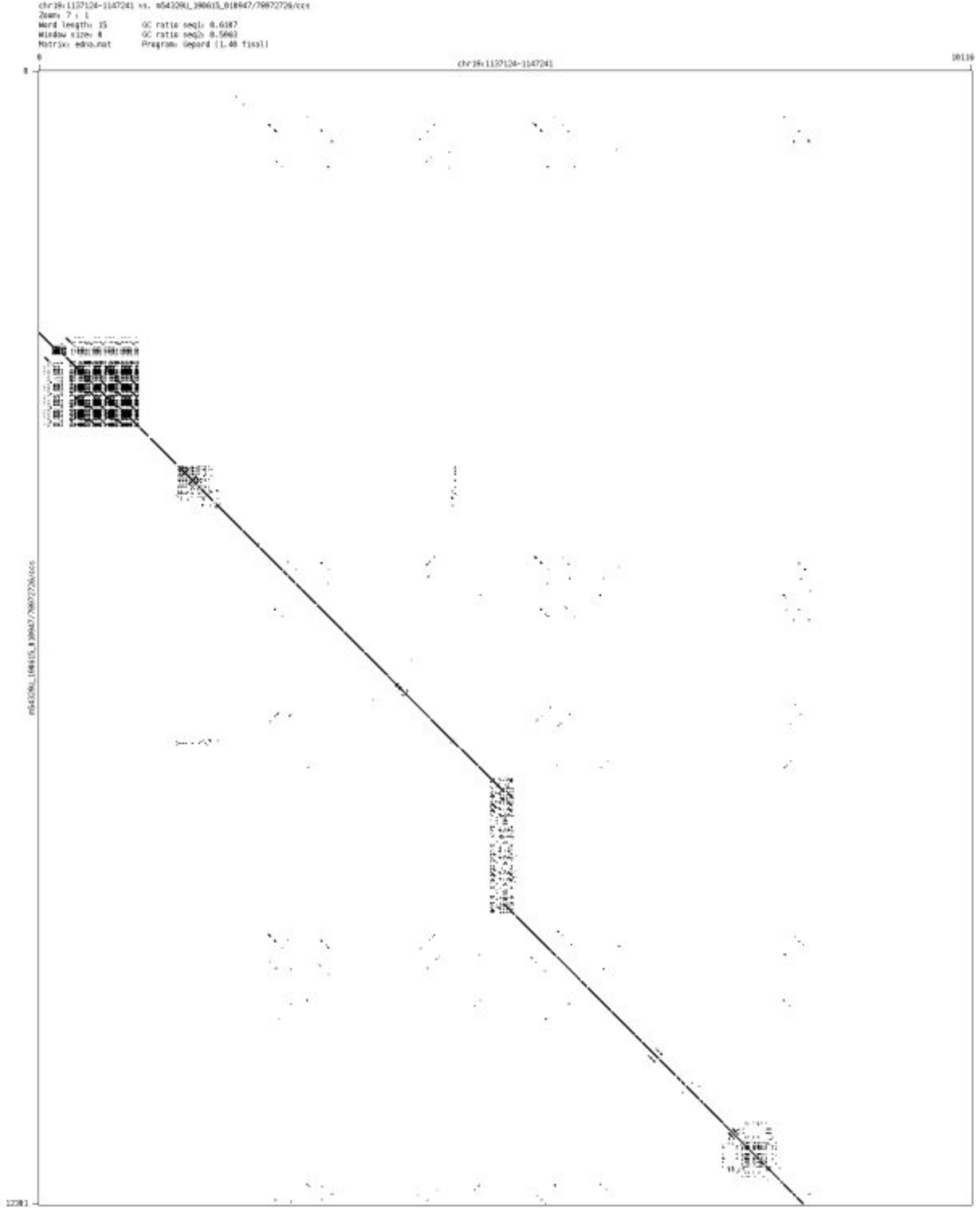

- chr19_33249731_33250285

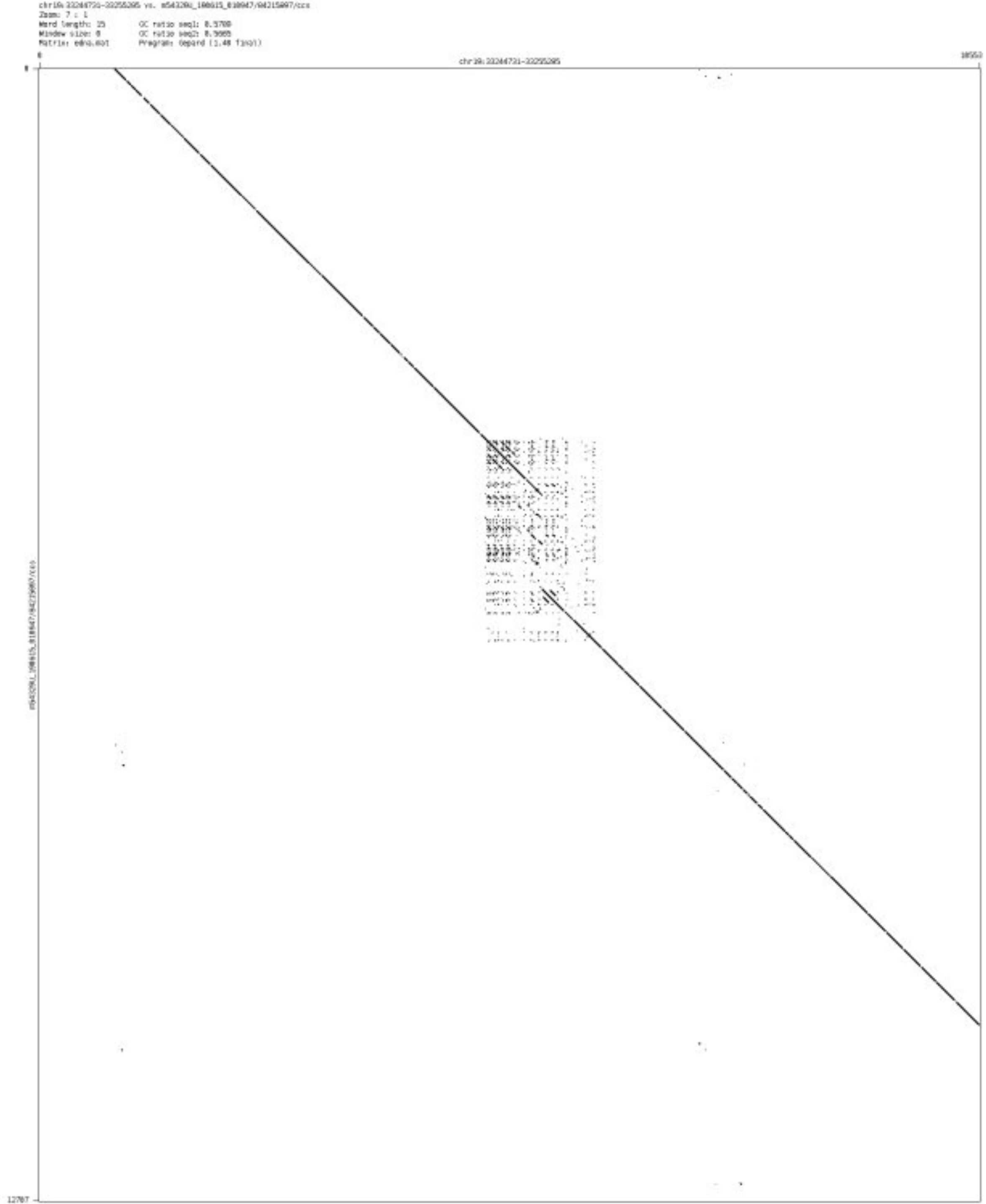

- ➢ chr20_44404913_44405031

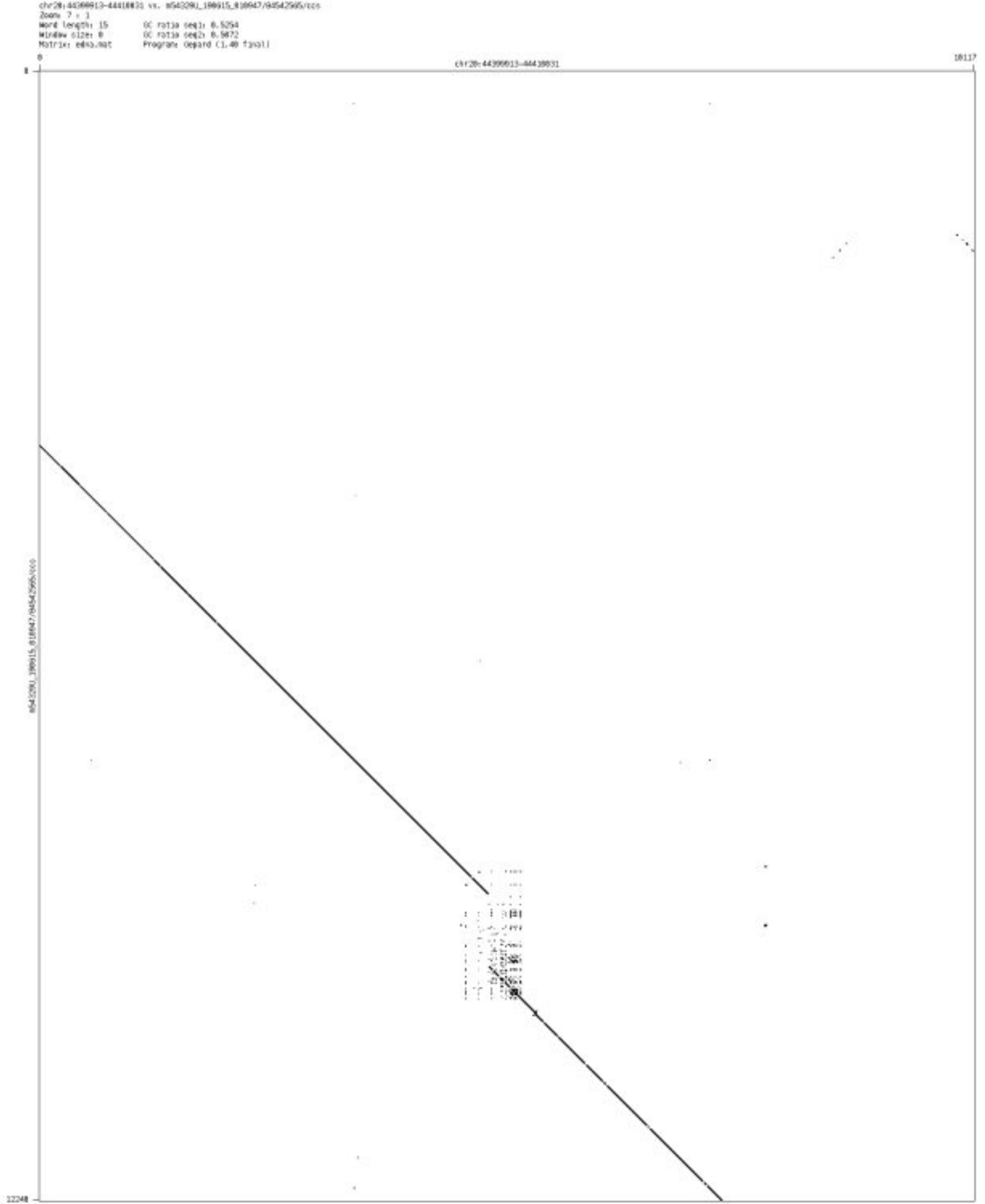

- chr20_58695019_58695331

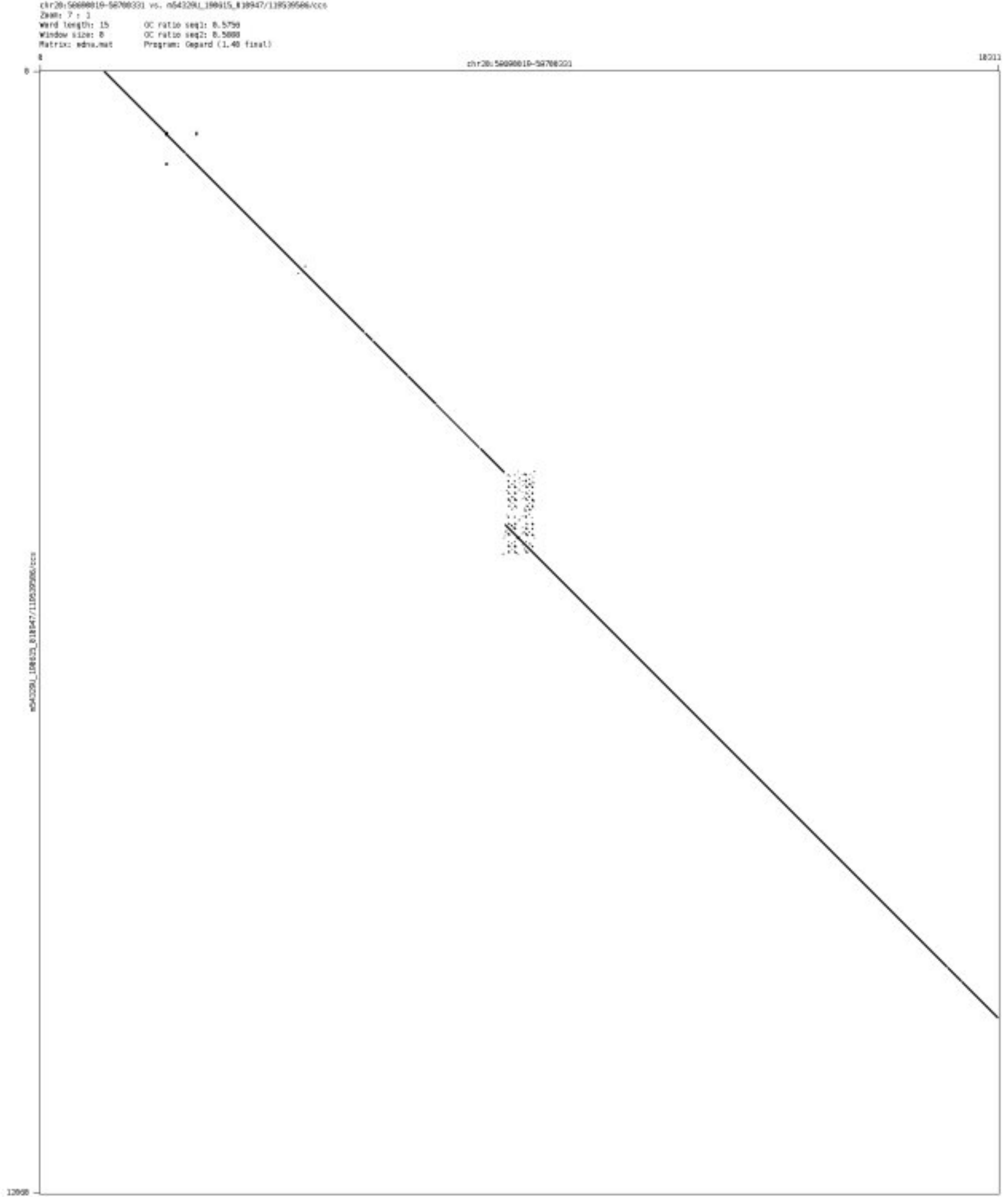

- chr20_61331578_61331692

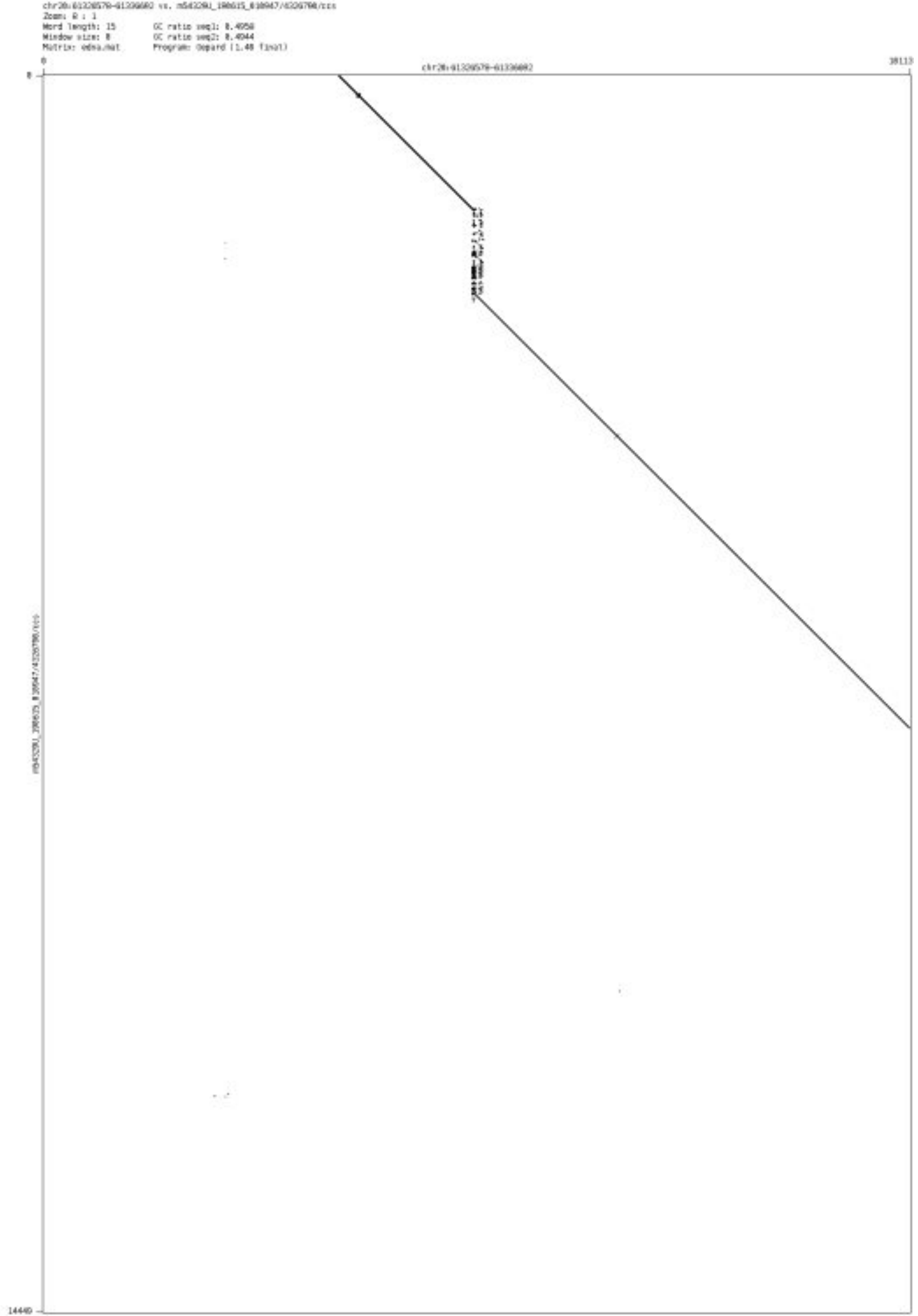

- ➢ chr20_63492299_63492537

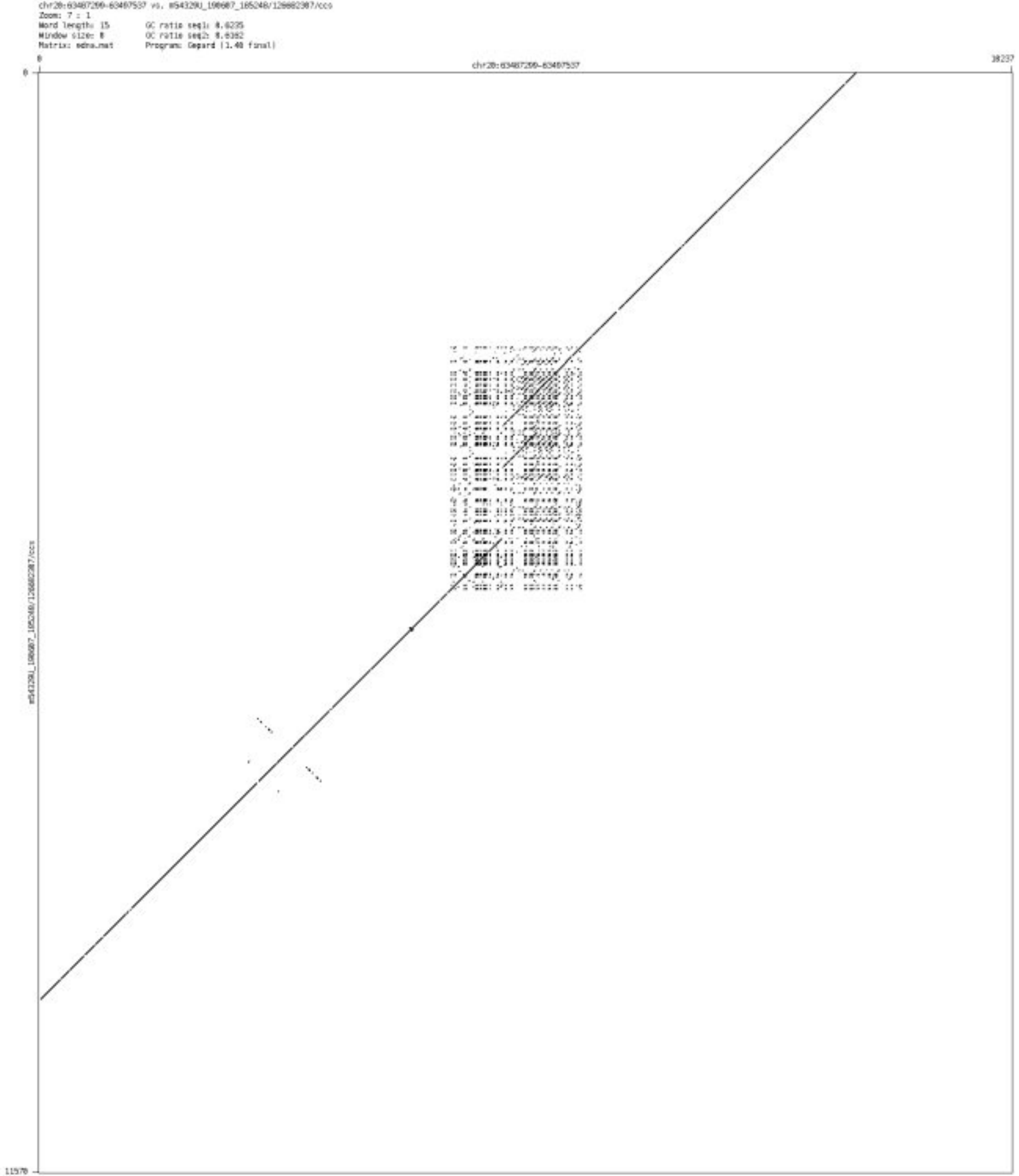

- ➢ chr21_45156103_45156277

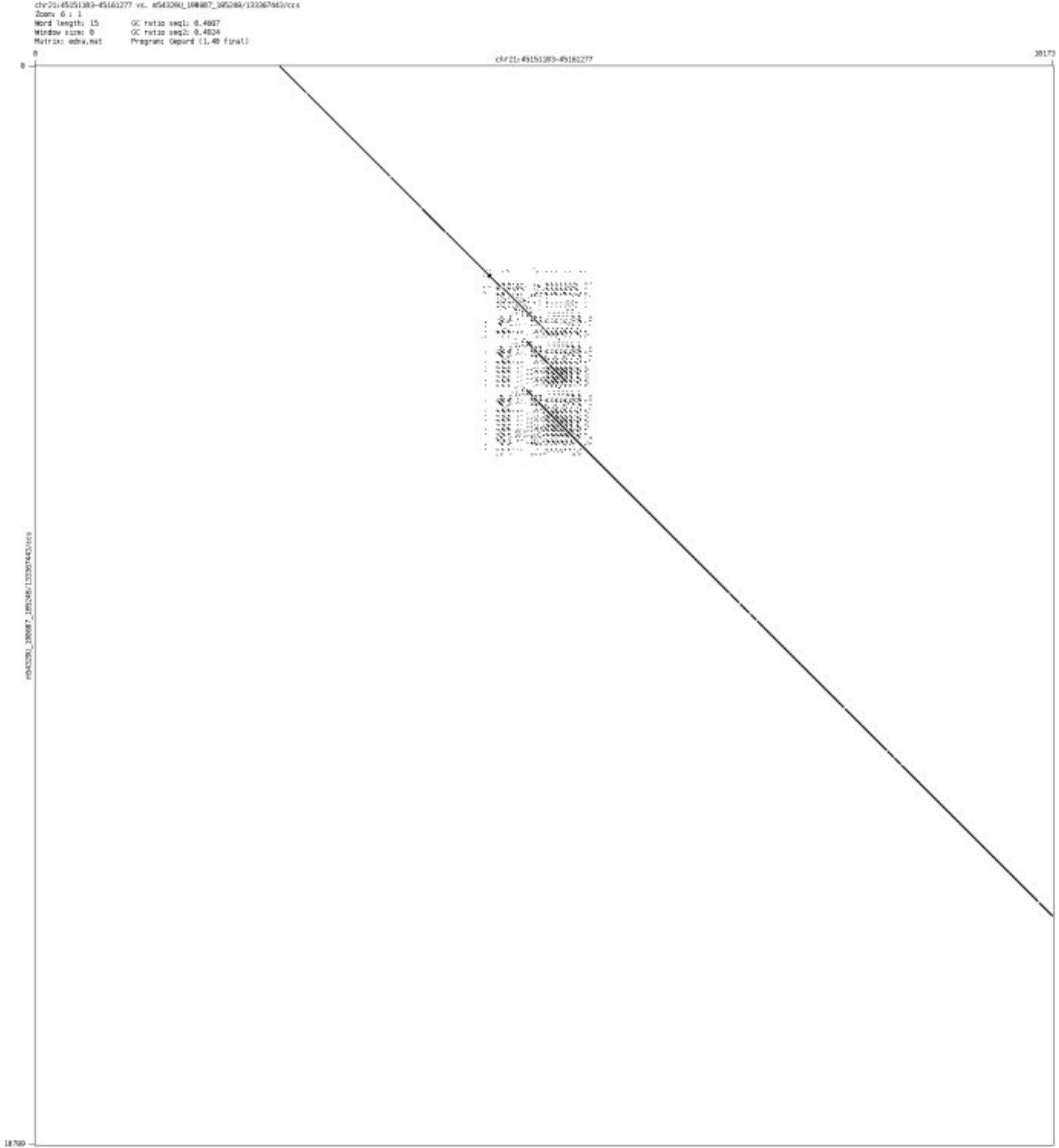

- chr21_45797476_45798540

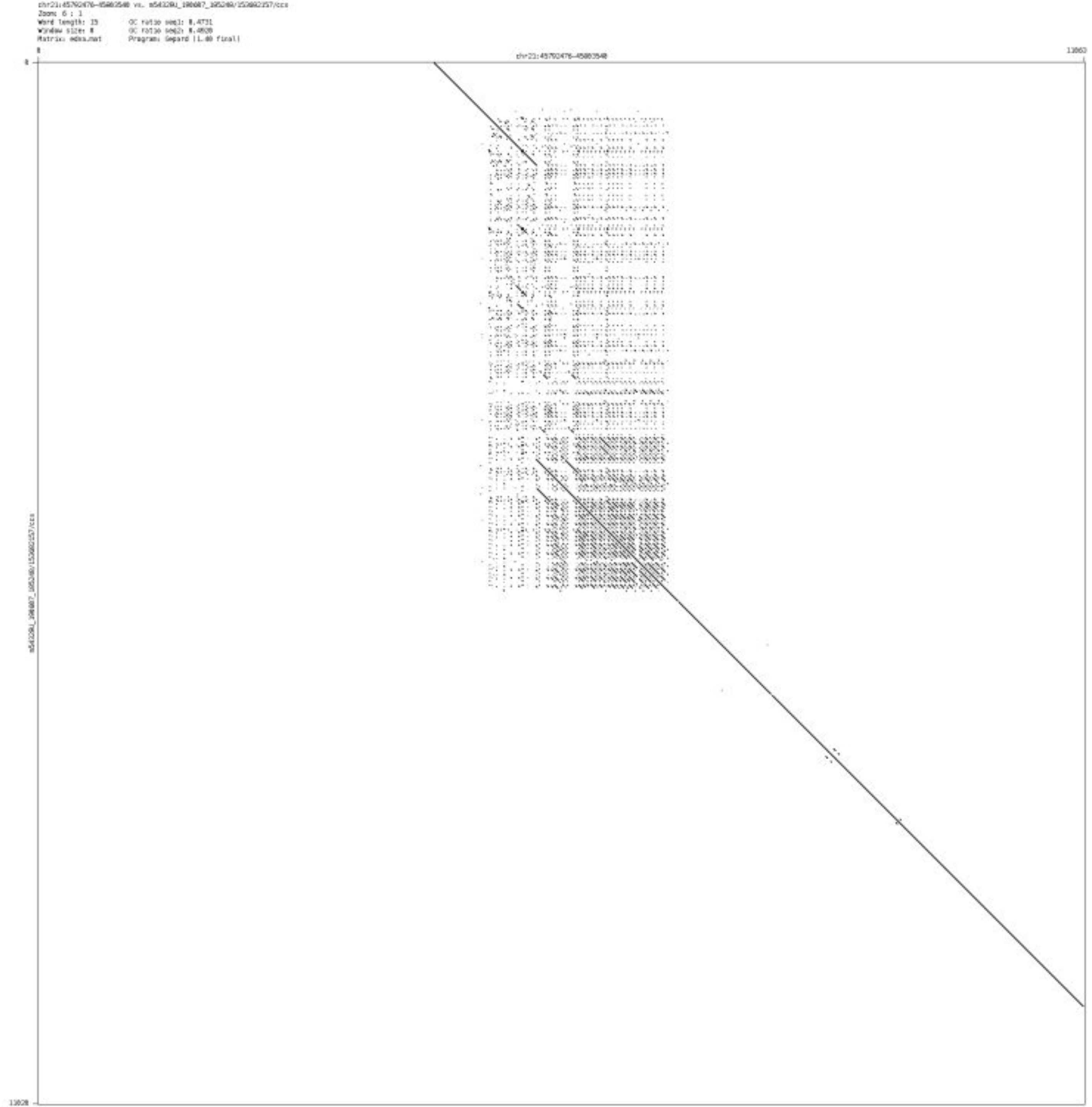

- chr22_48757225_48757402

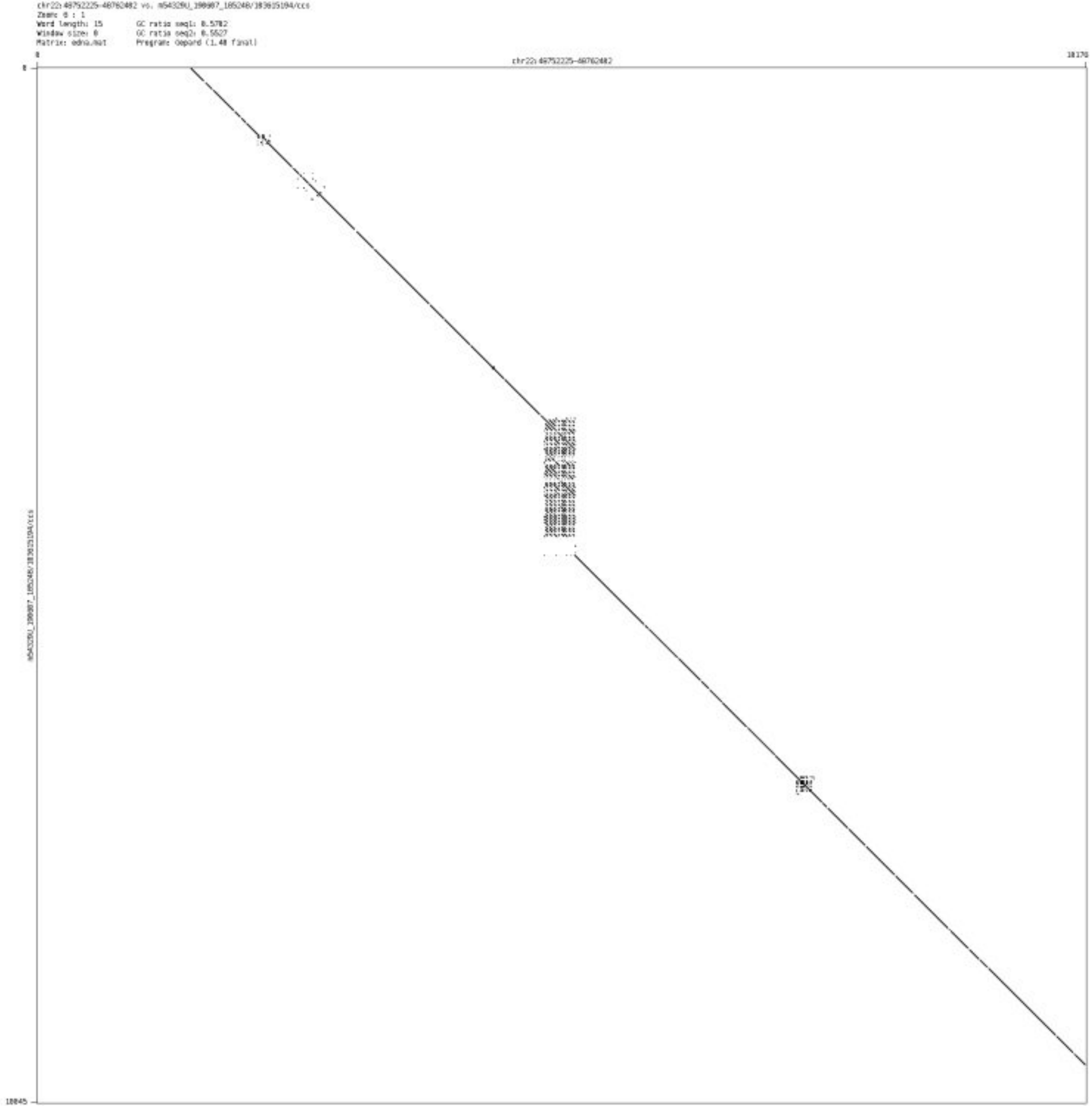

### ’Messy’ CSV-Case B

➢ chr1_1382295_1382470

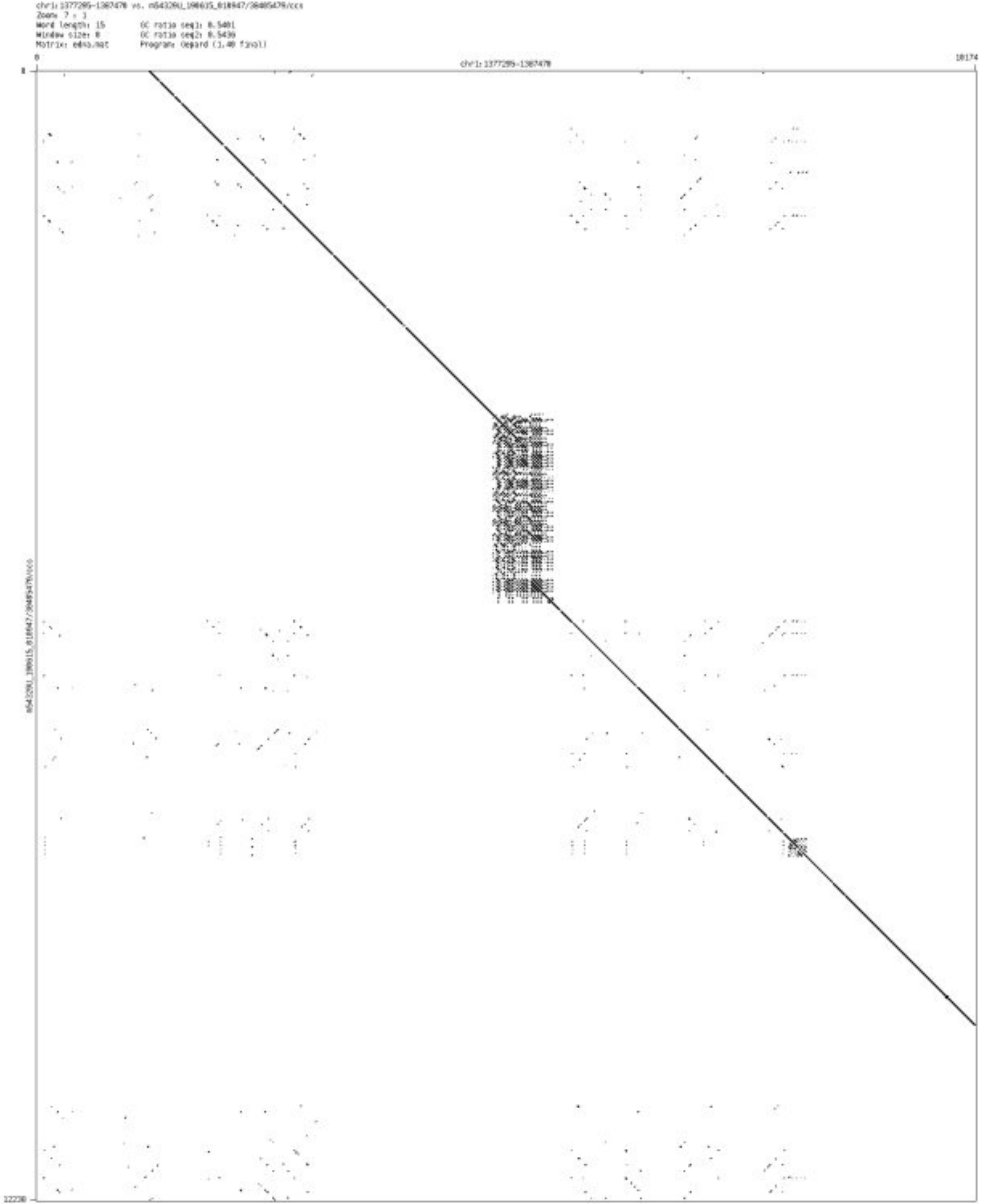

➢ chr1_2849326_2849465

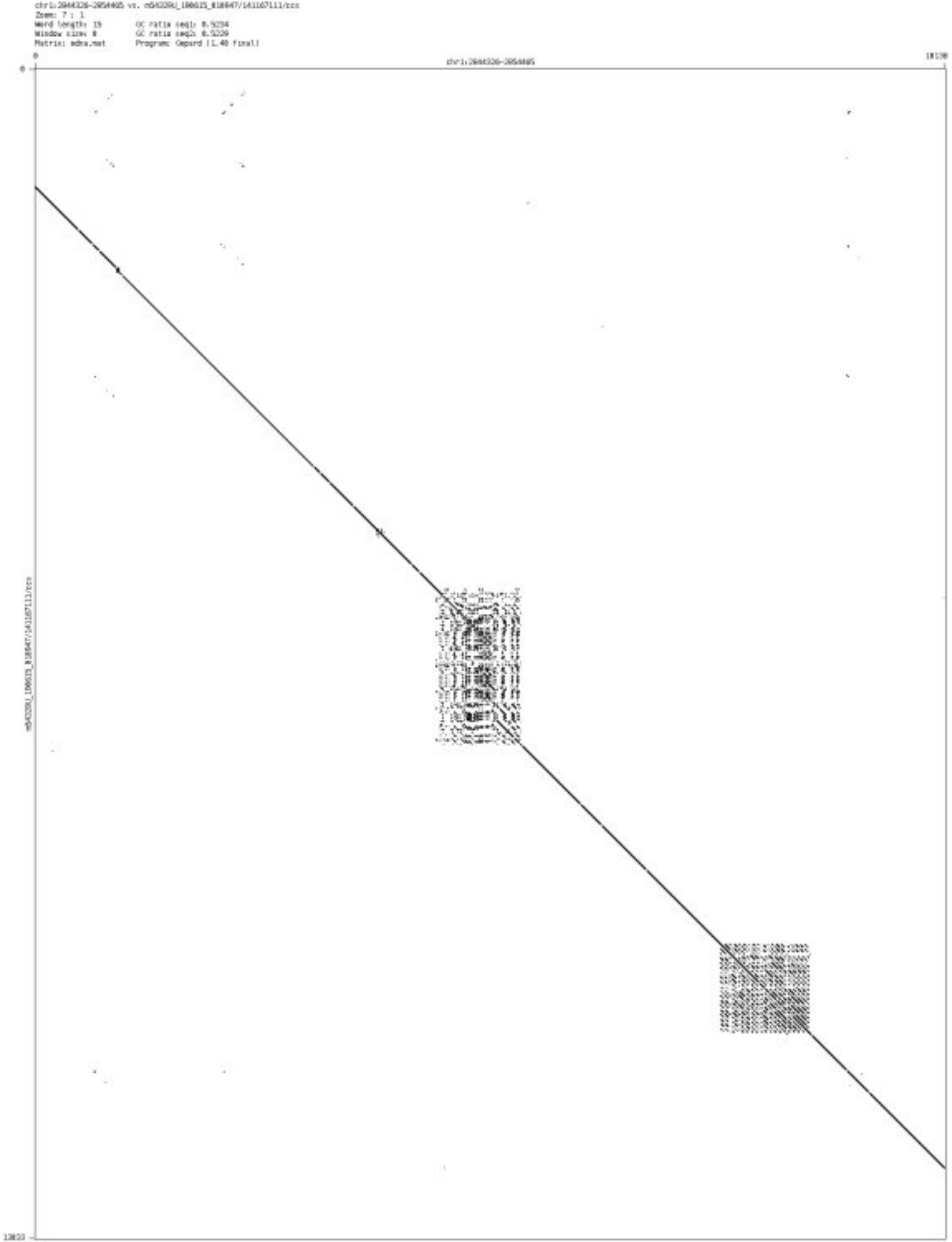

- chr1_4144504_4144634

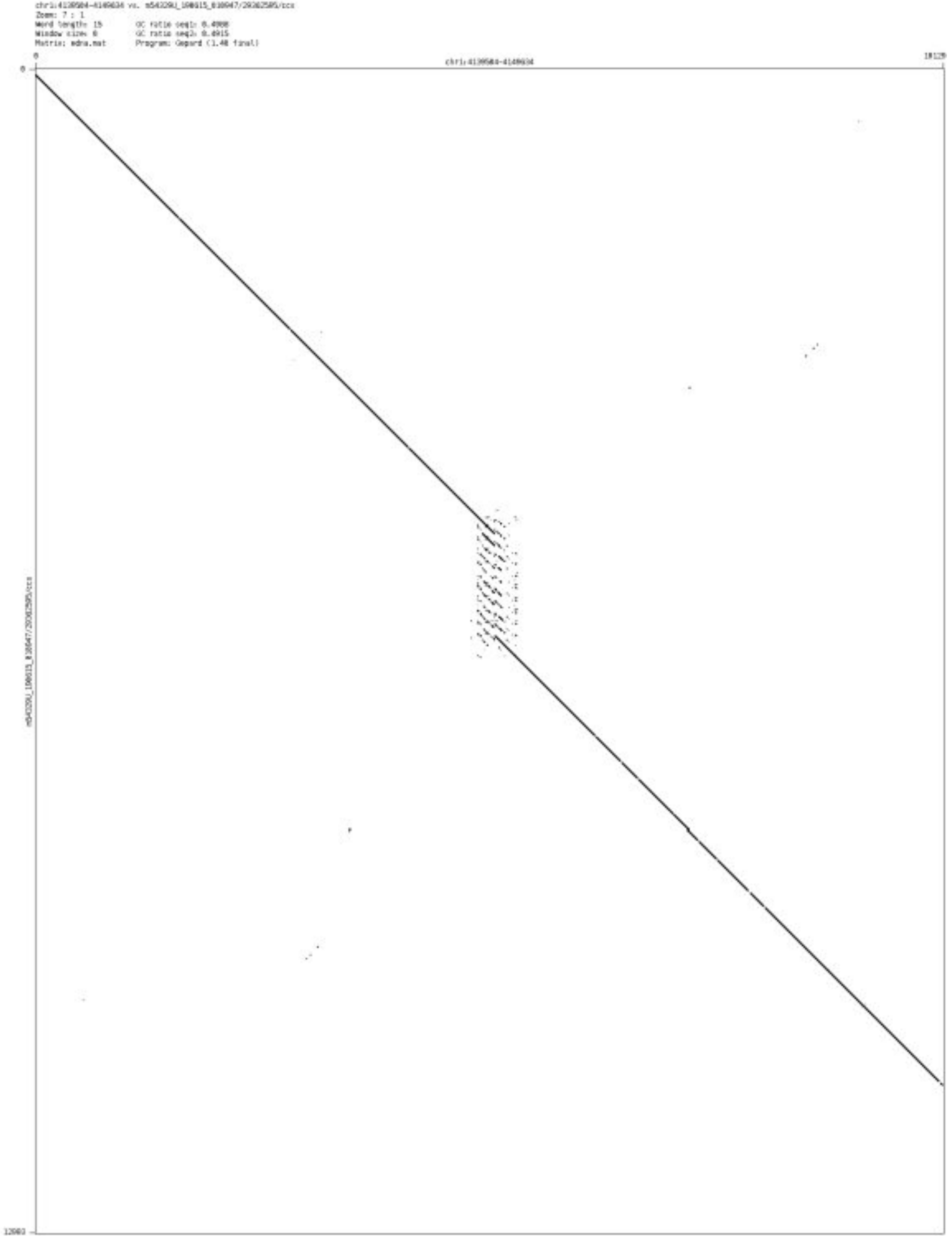

➢ chr1_167056632_167056747

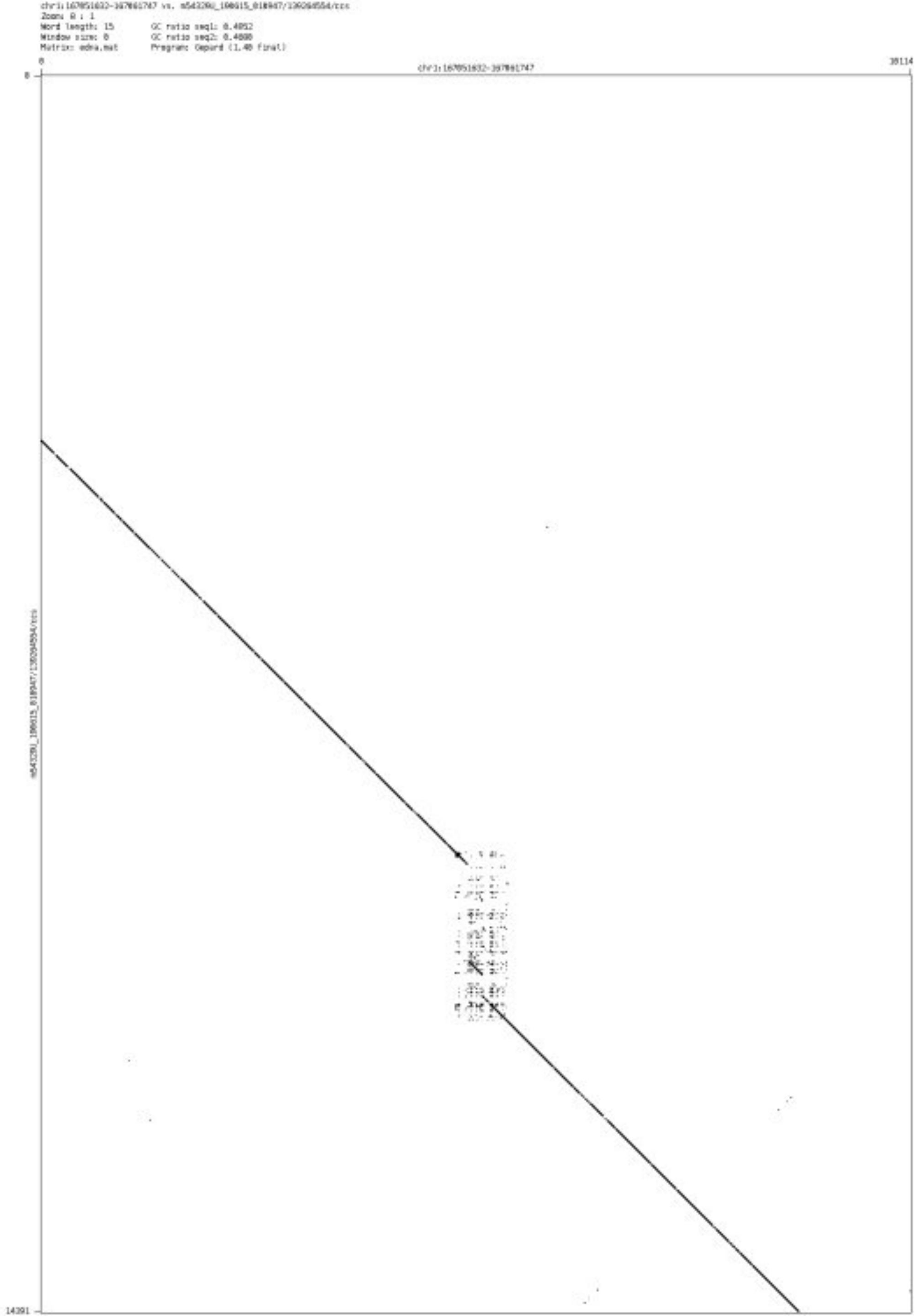

- chr1_175650824_175650946

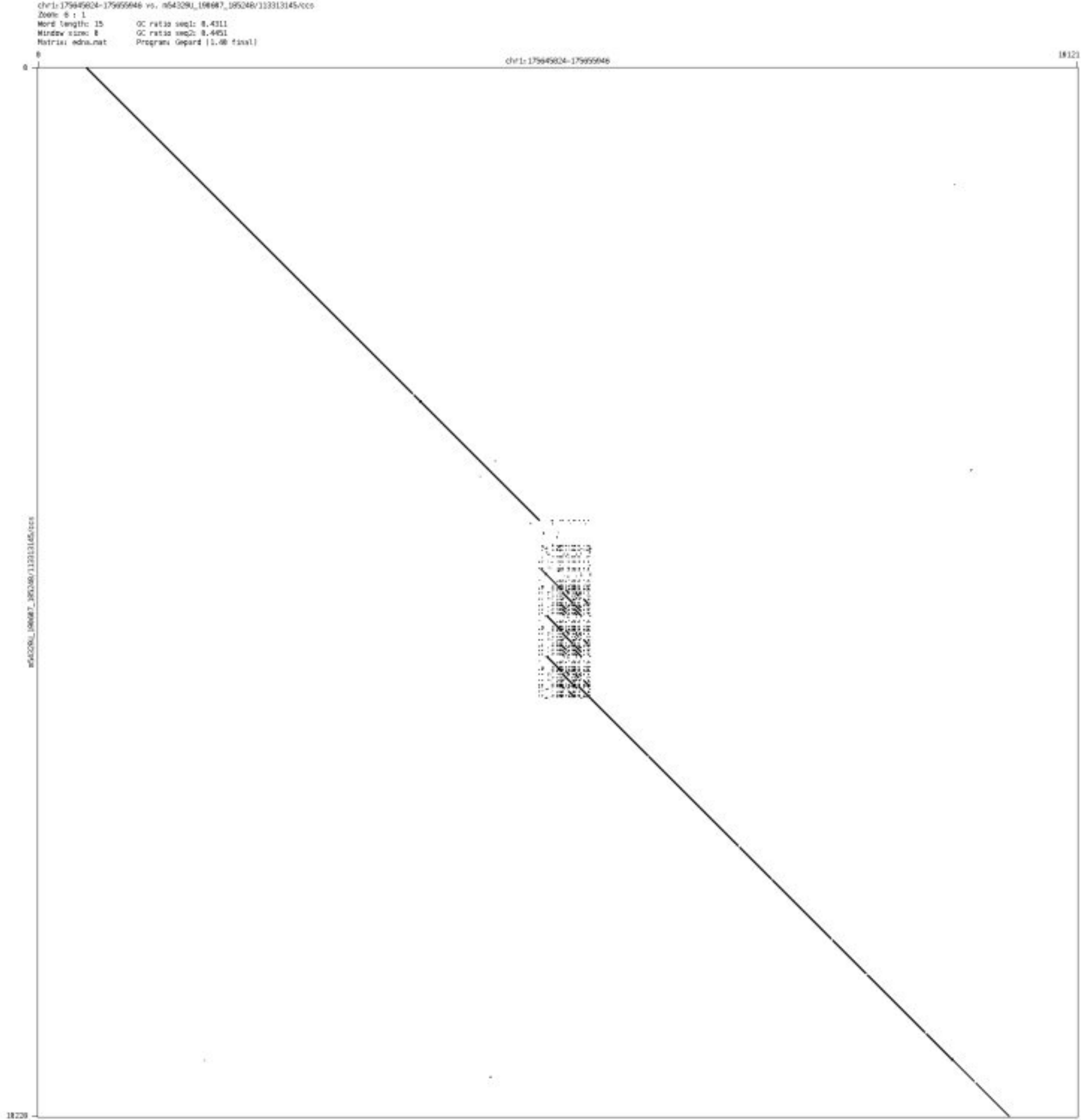

➢ chr1_202204526_202205476

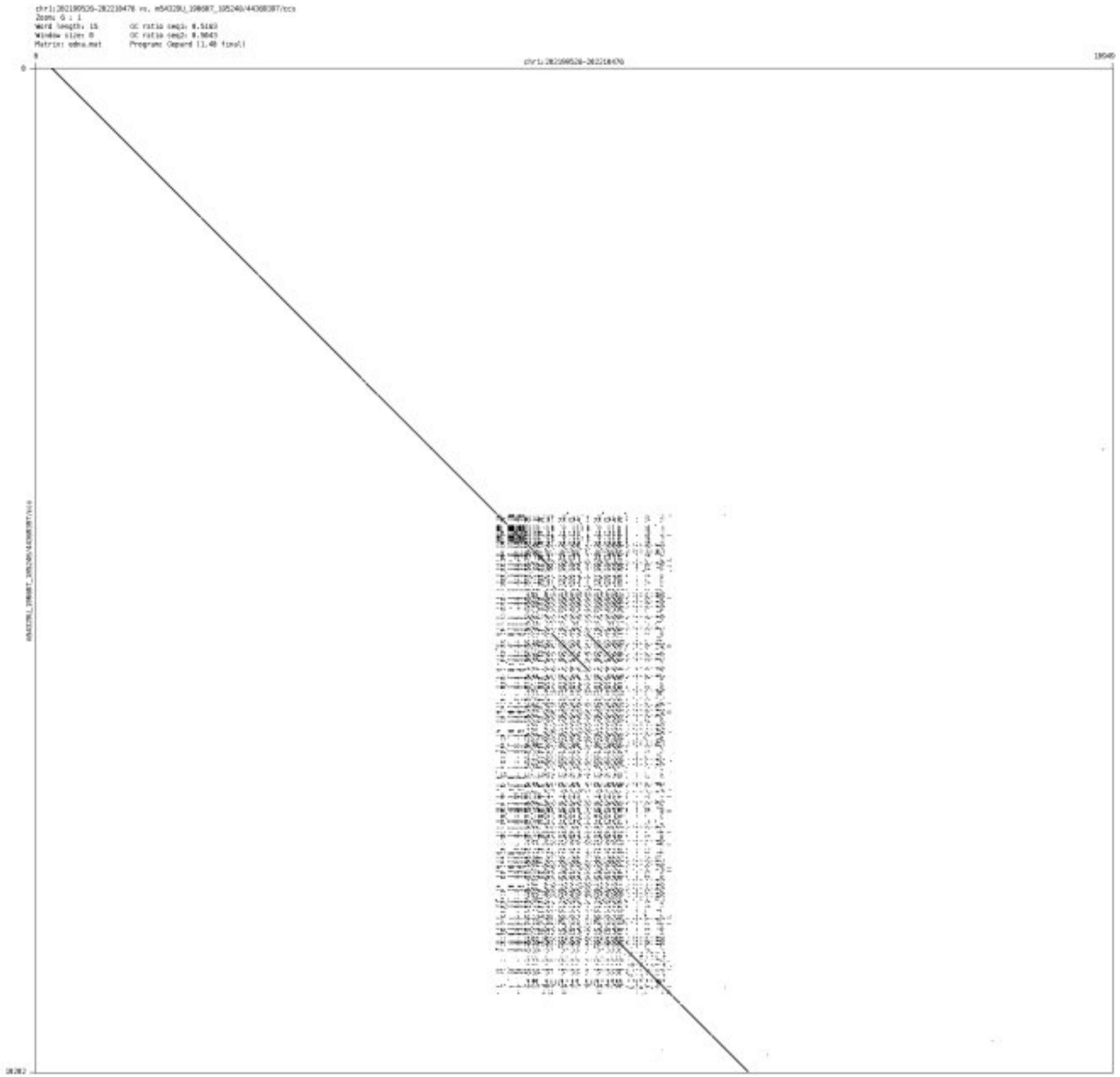

➢ chr1_243480631_243480726

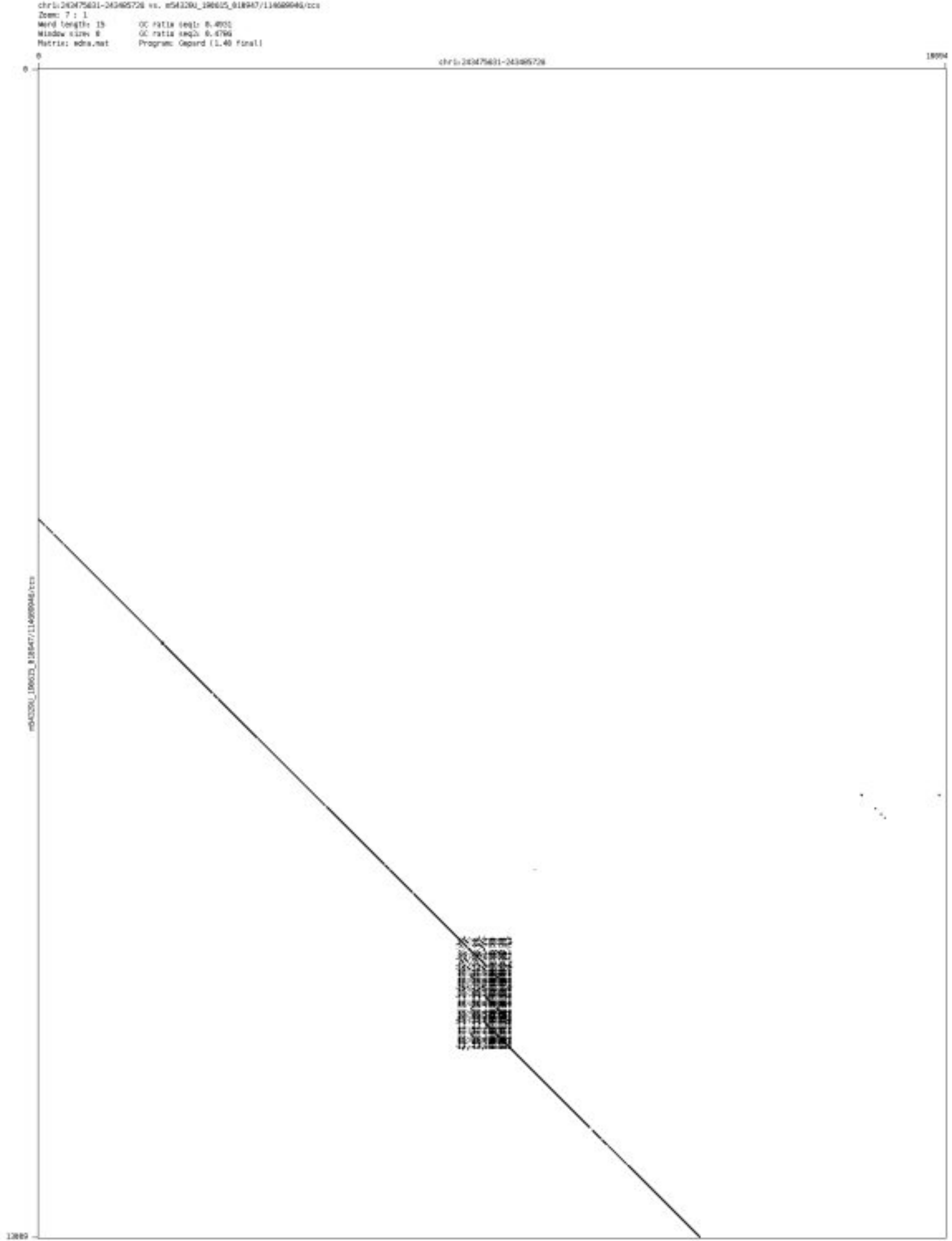

➢ chr2_3057865_3057980

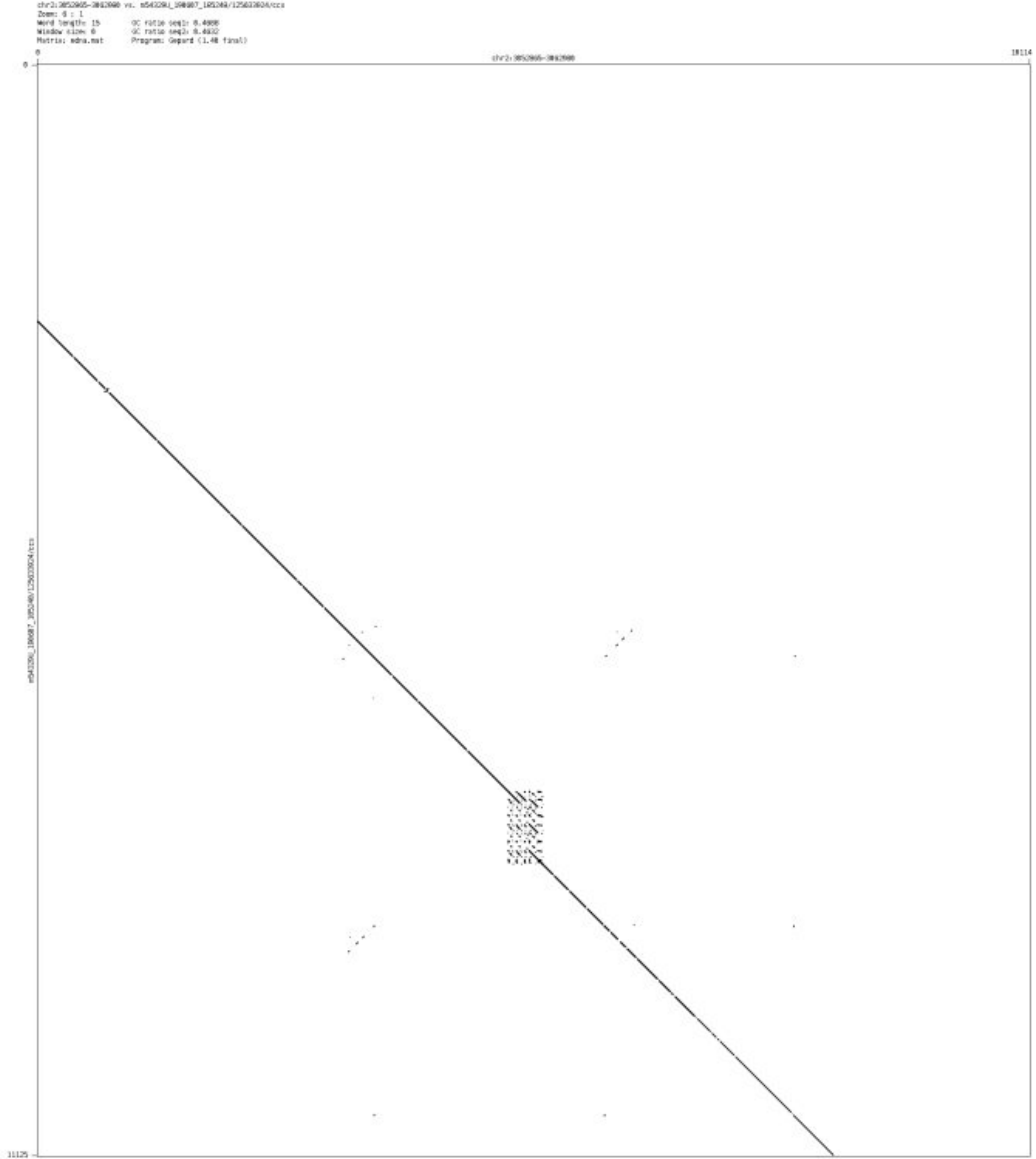

- chr2_65600866_65601053

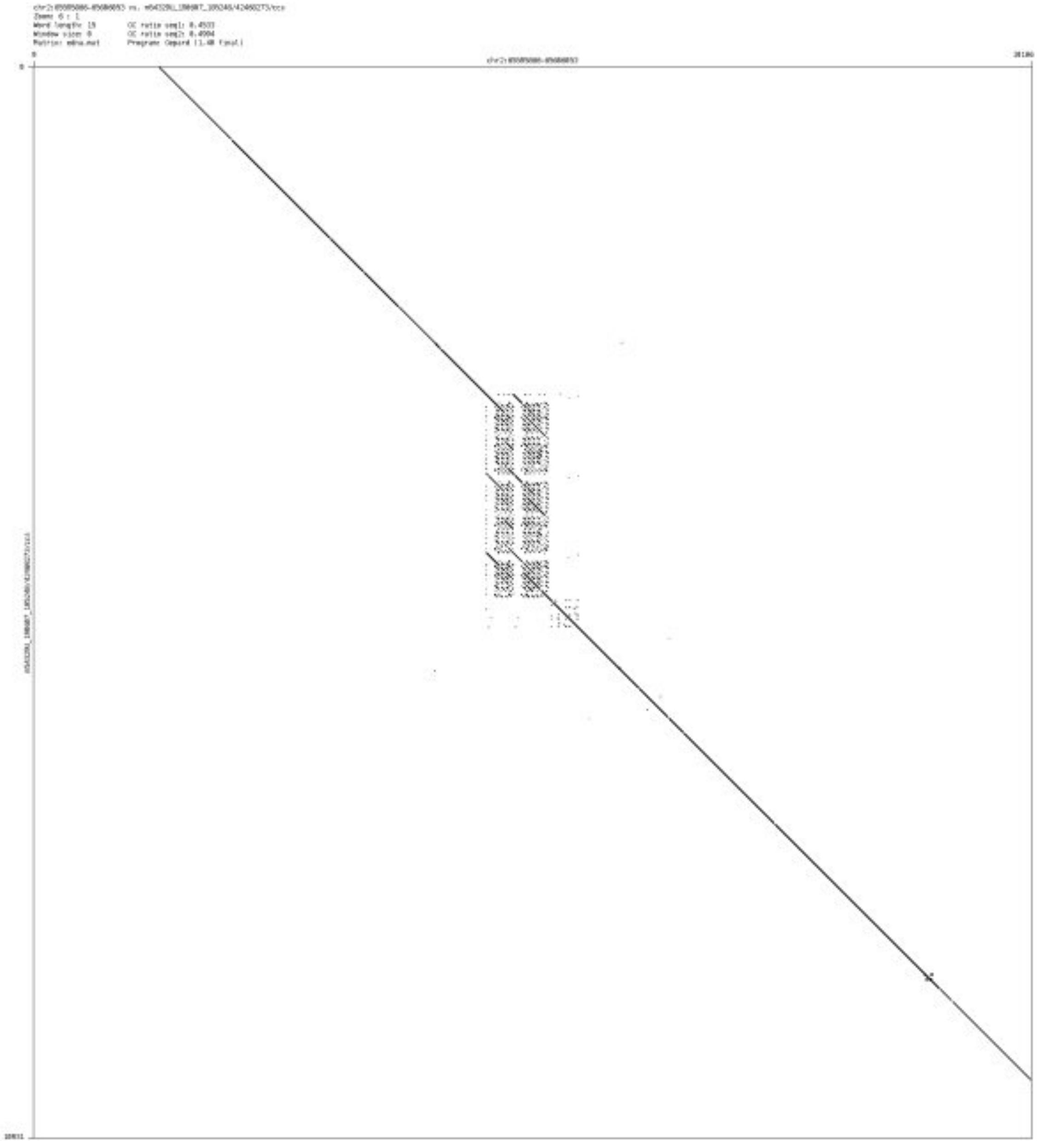

➢ chr2_71618111_71618577

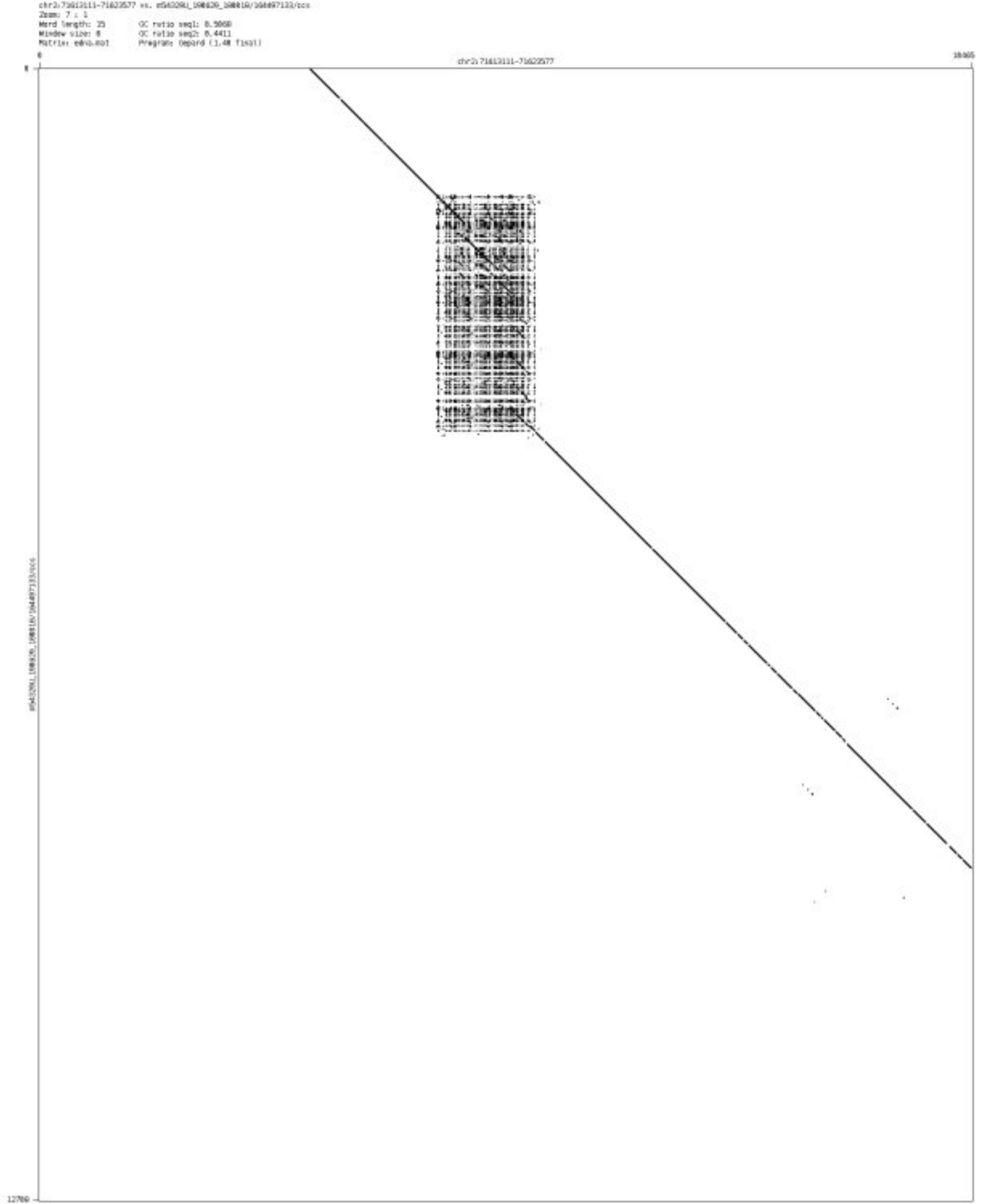

- chr2_238870308_238870448

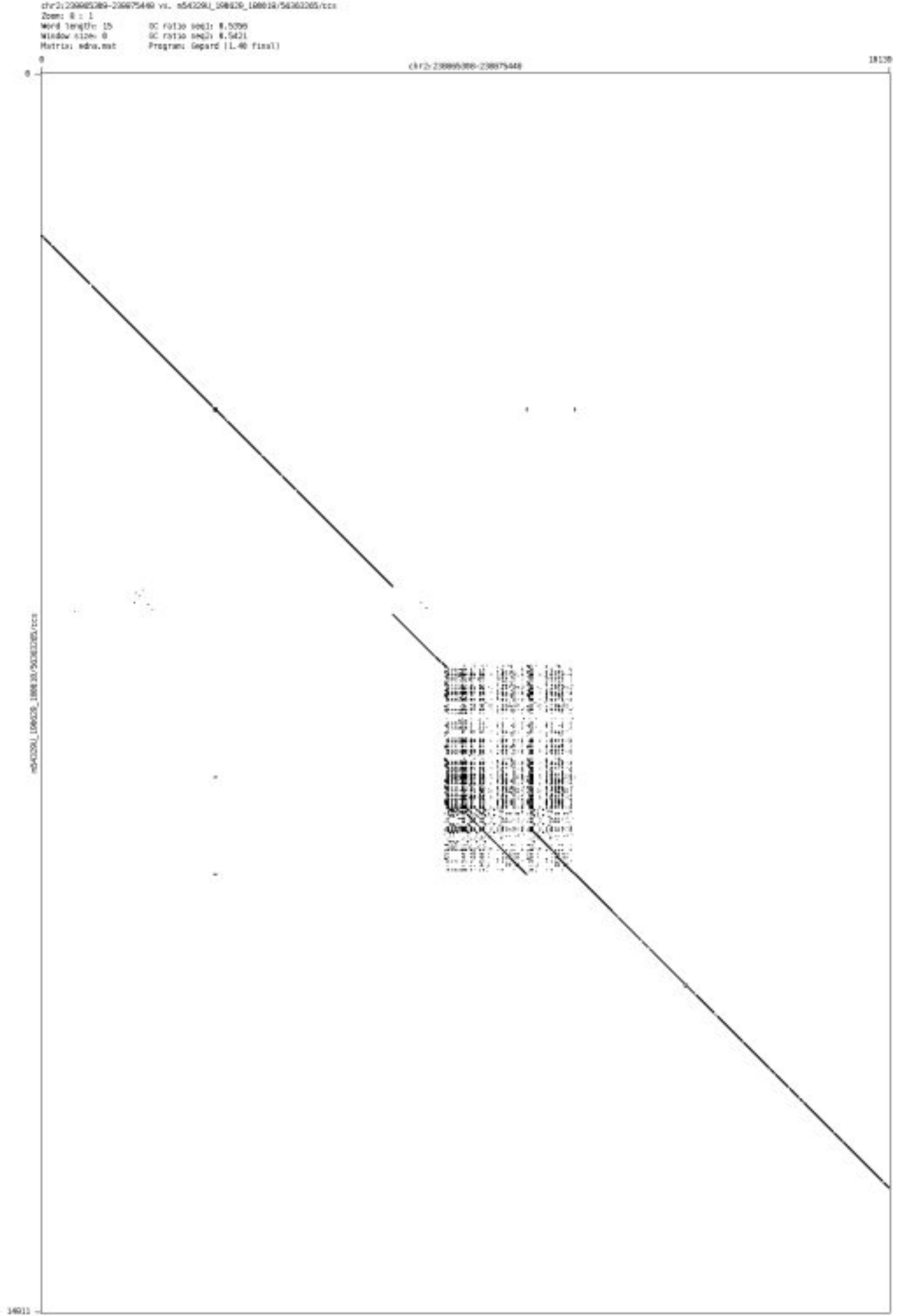

- chr2_240786636_240786828

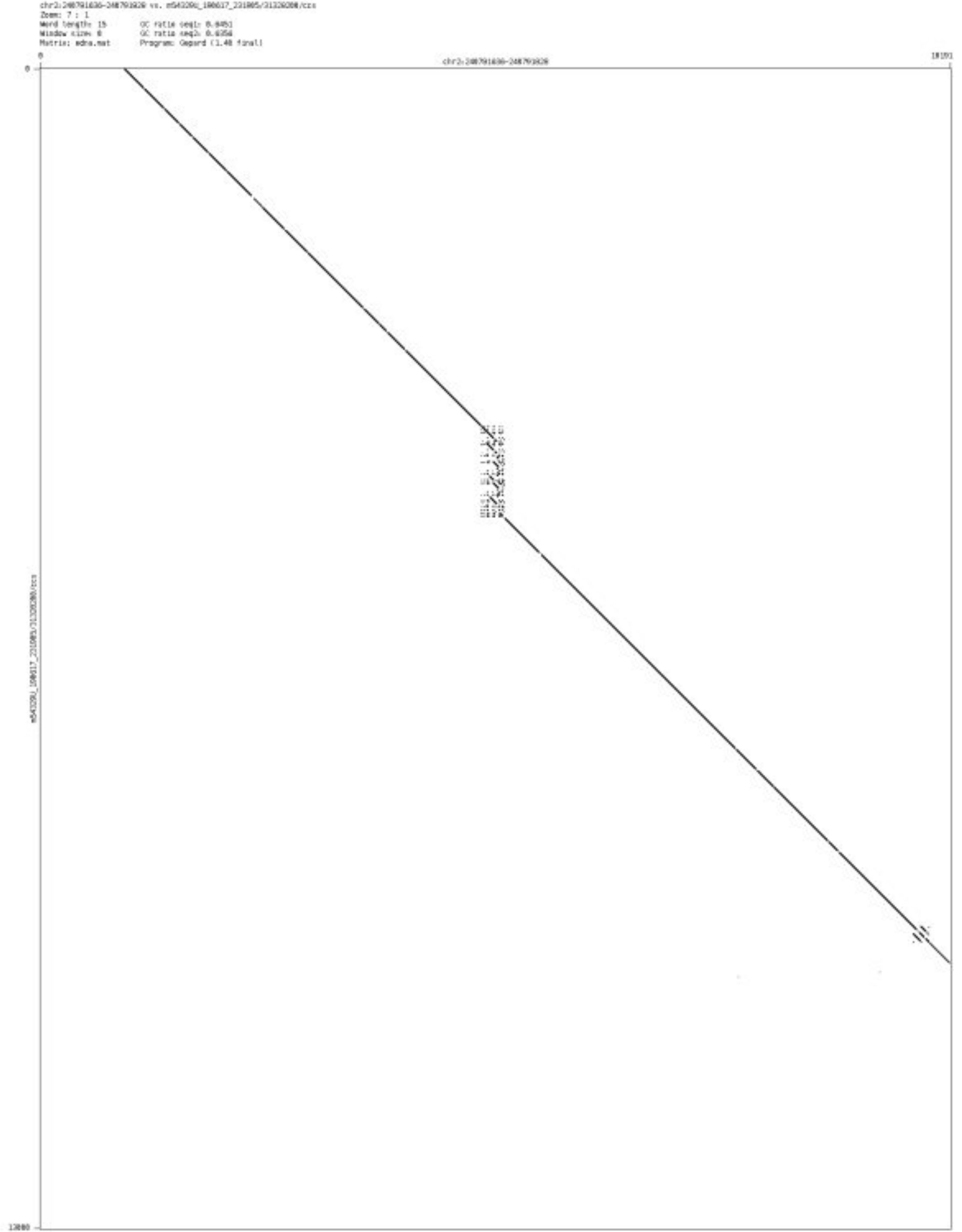

- ➢ chr3_194324598_194324699

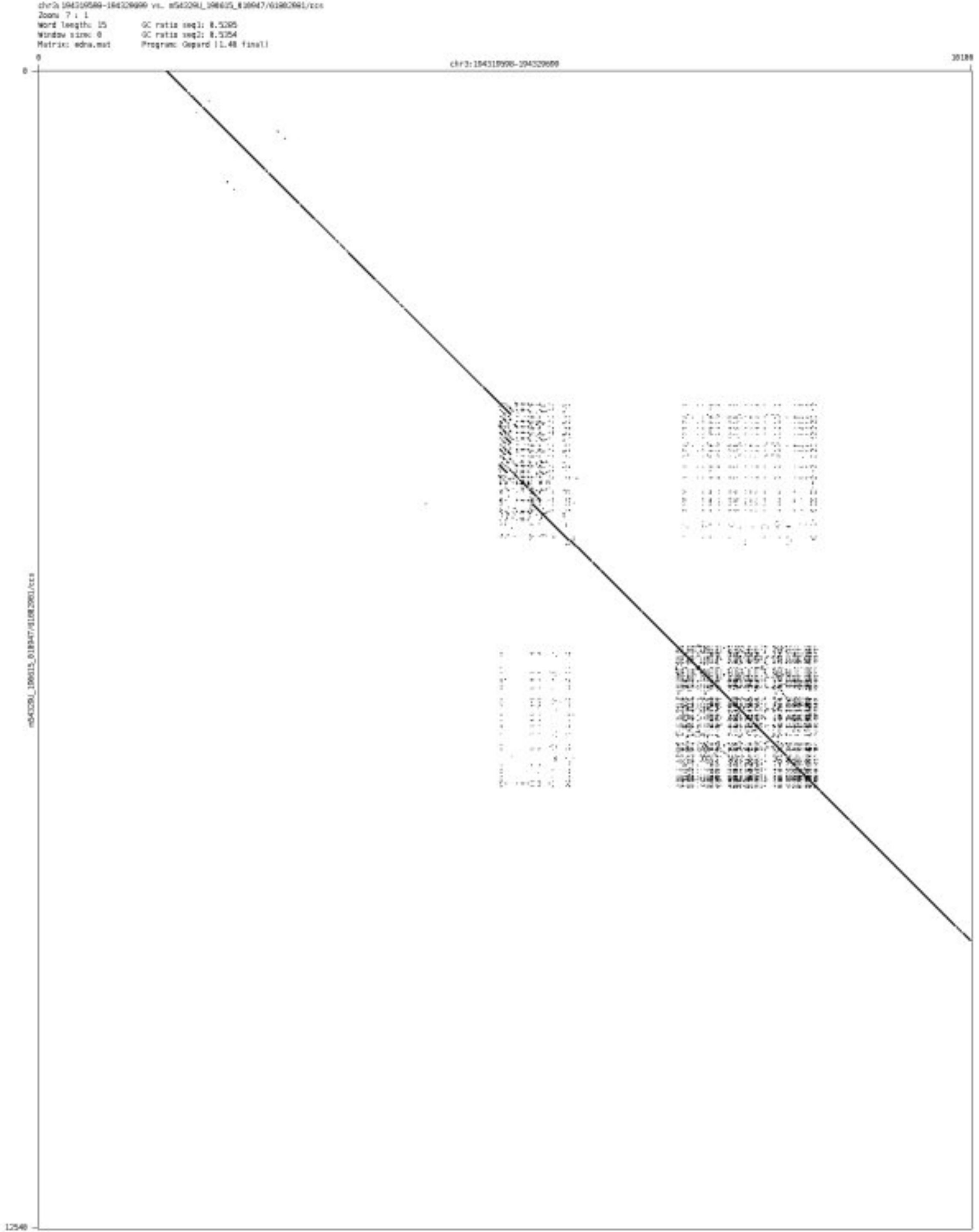

- chr3_197390213_197391660

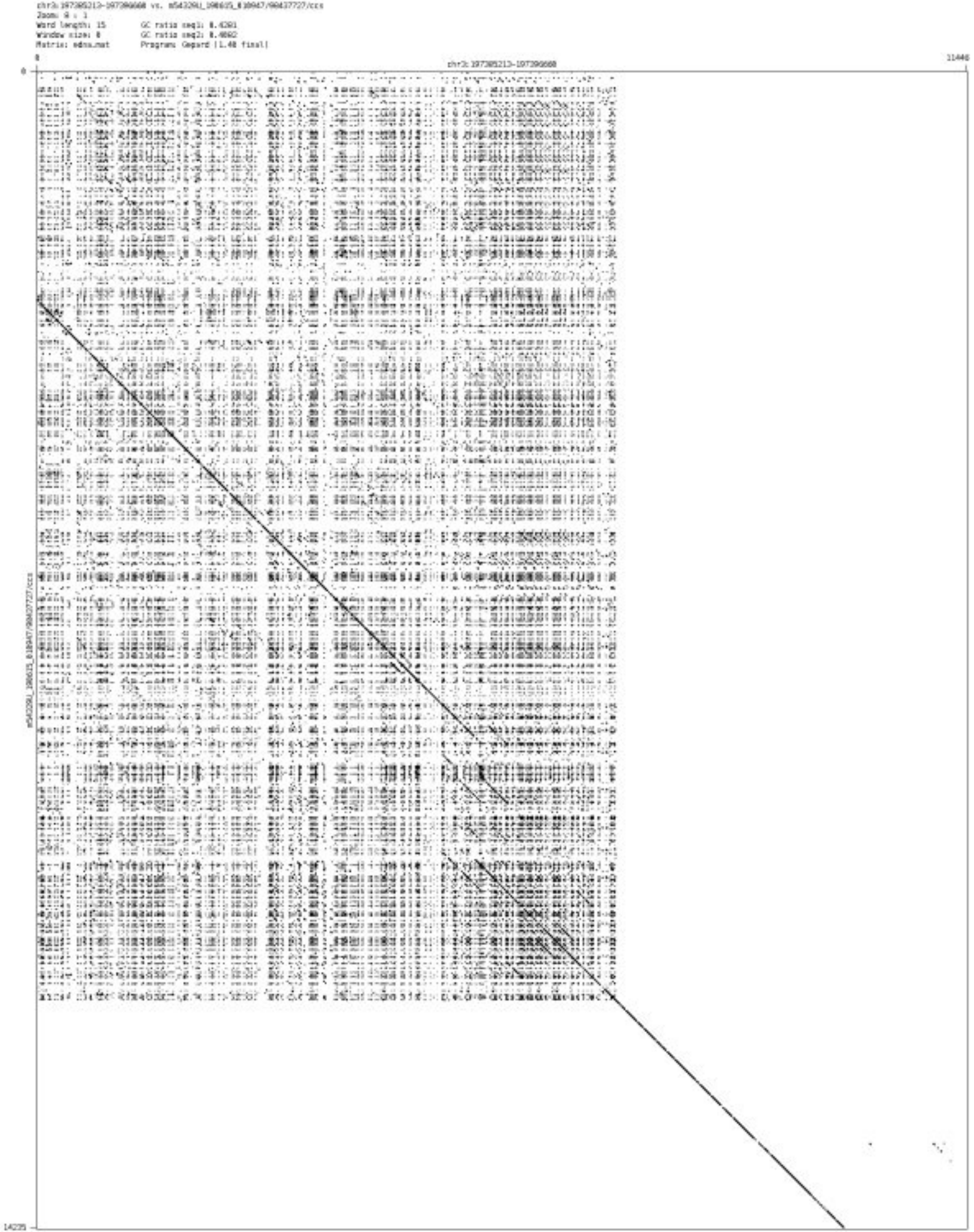

- chr3_197459802_197459965

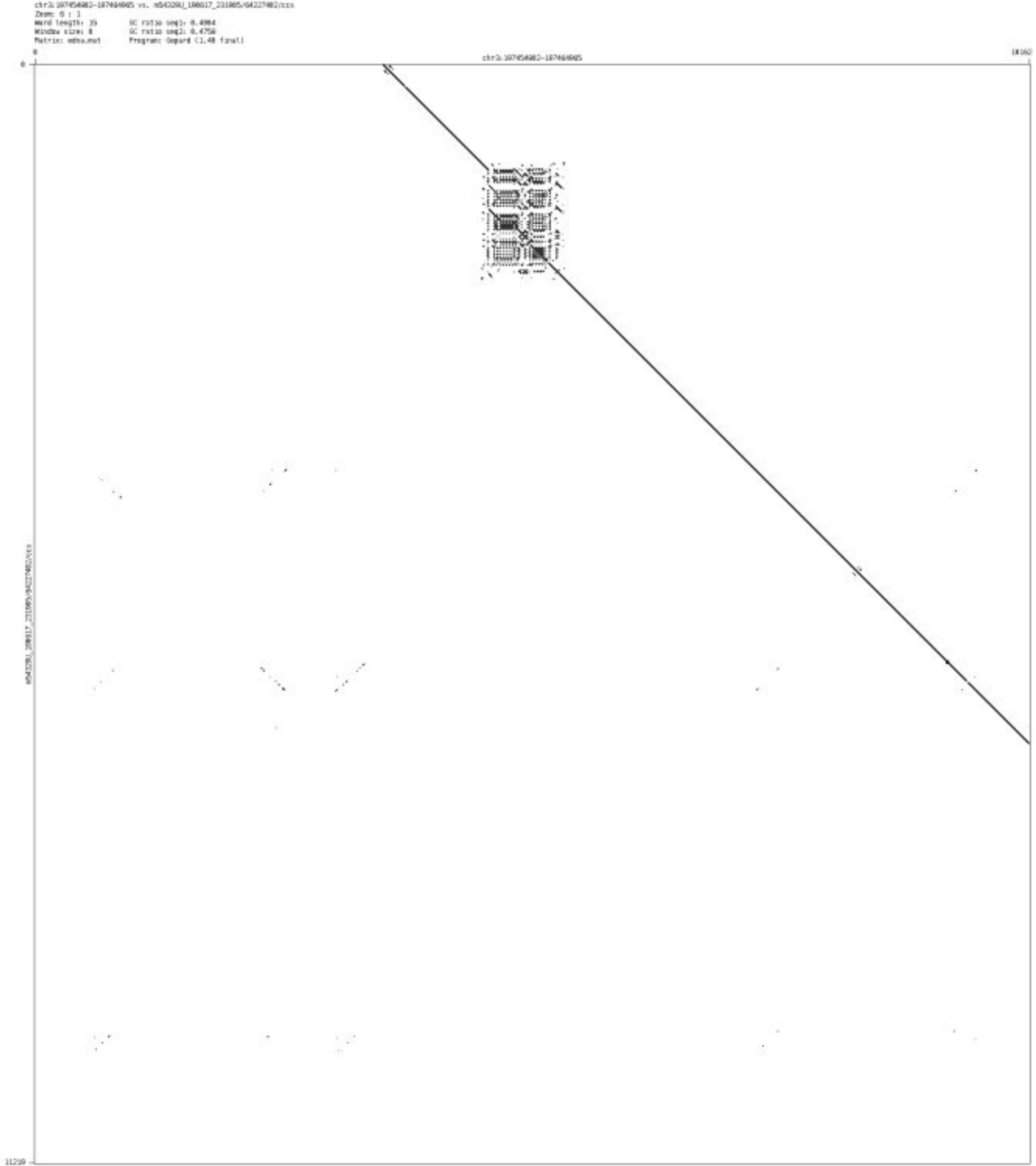

- chr4_1046301_1046664

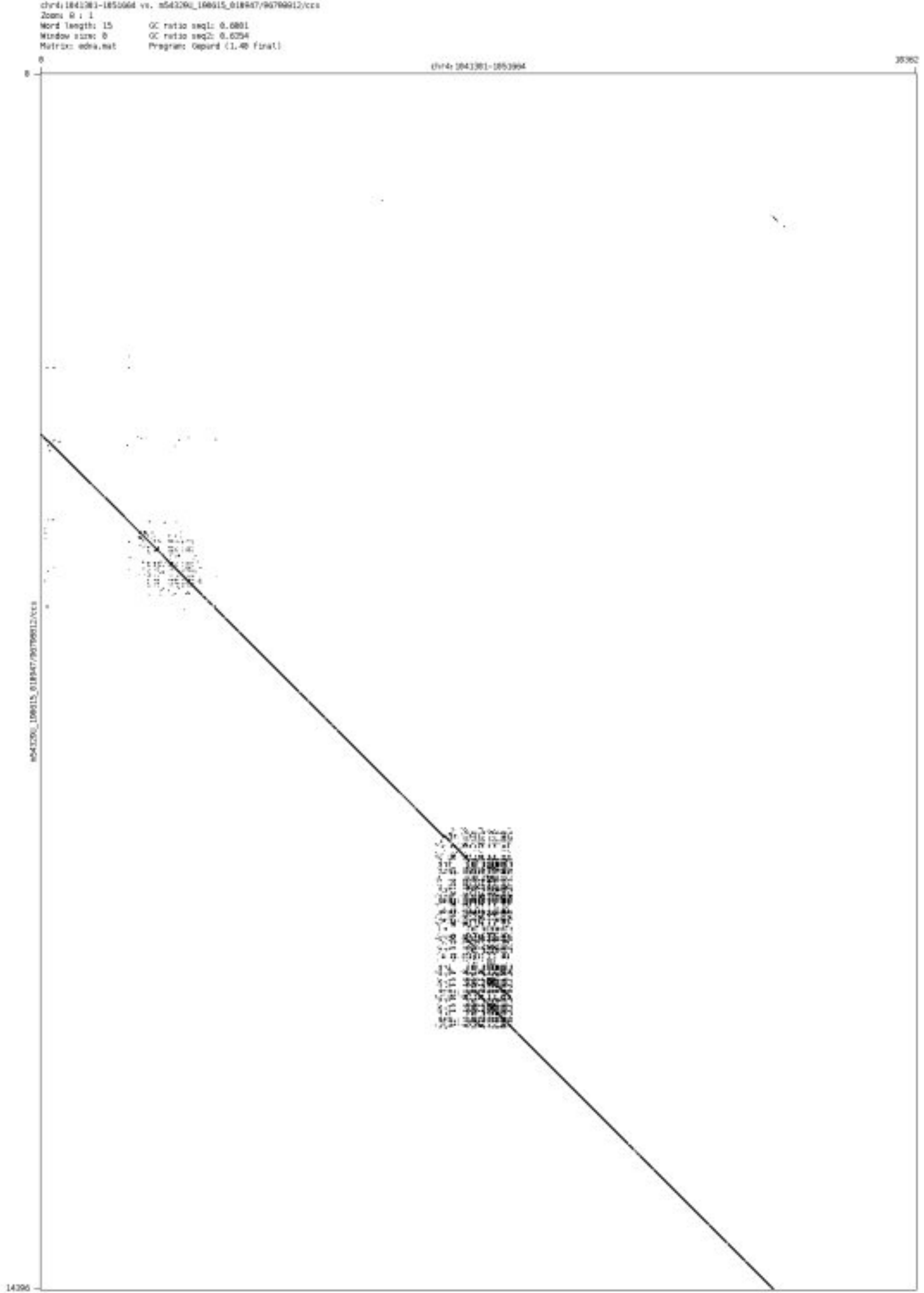

- chr4_3814944_3815102

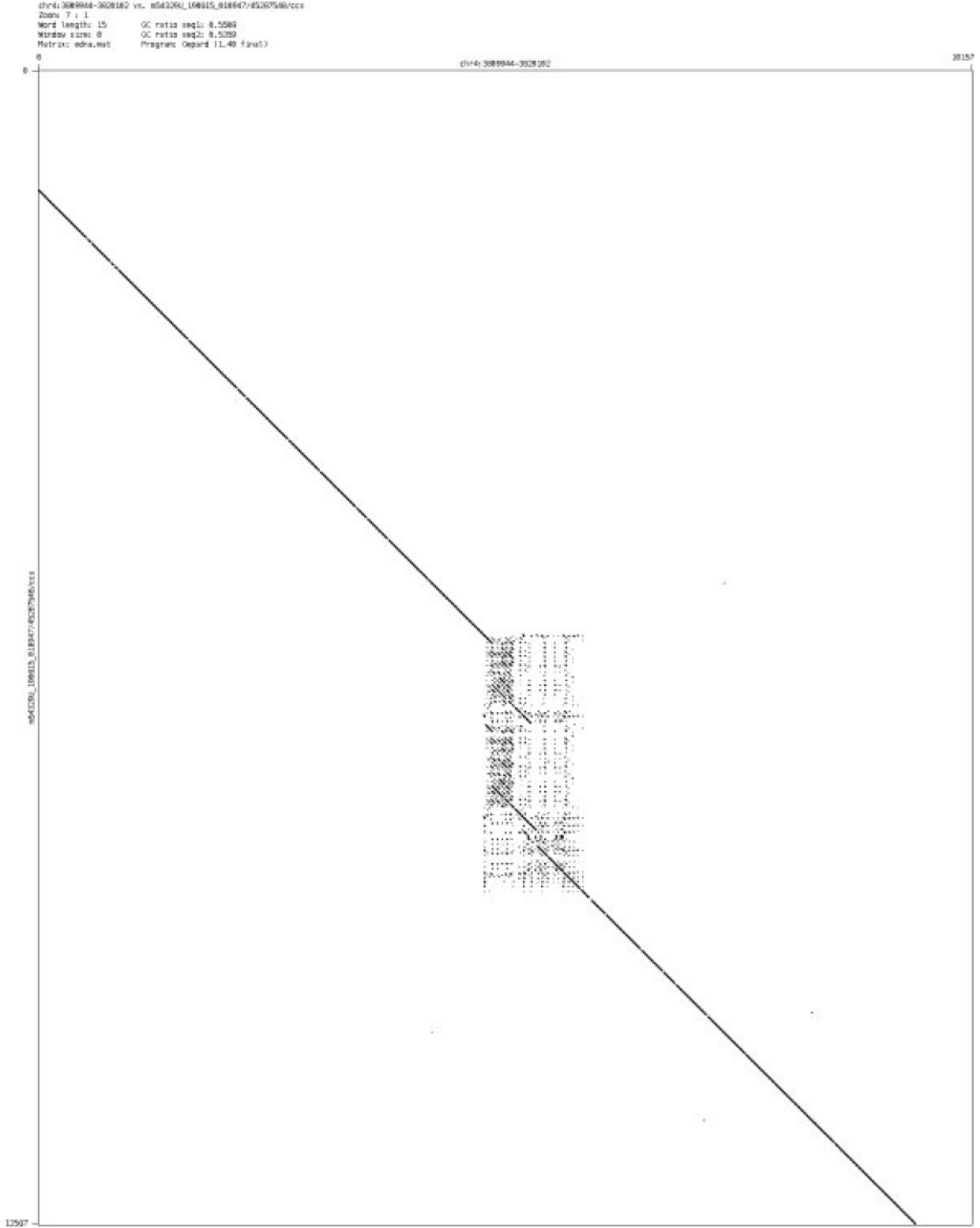

➢ chr4_40295148_40295512

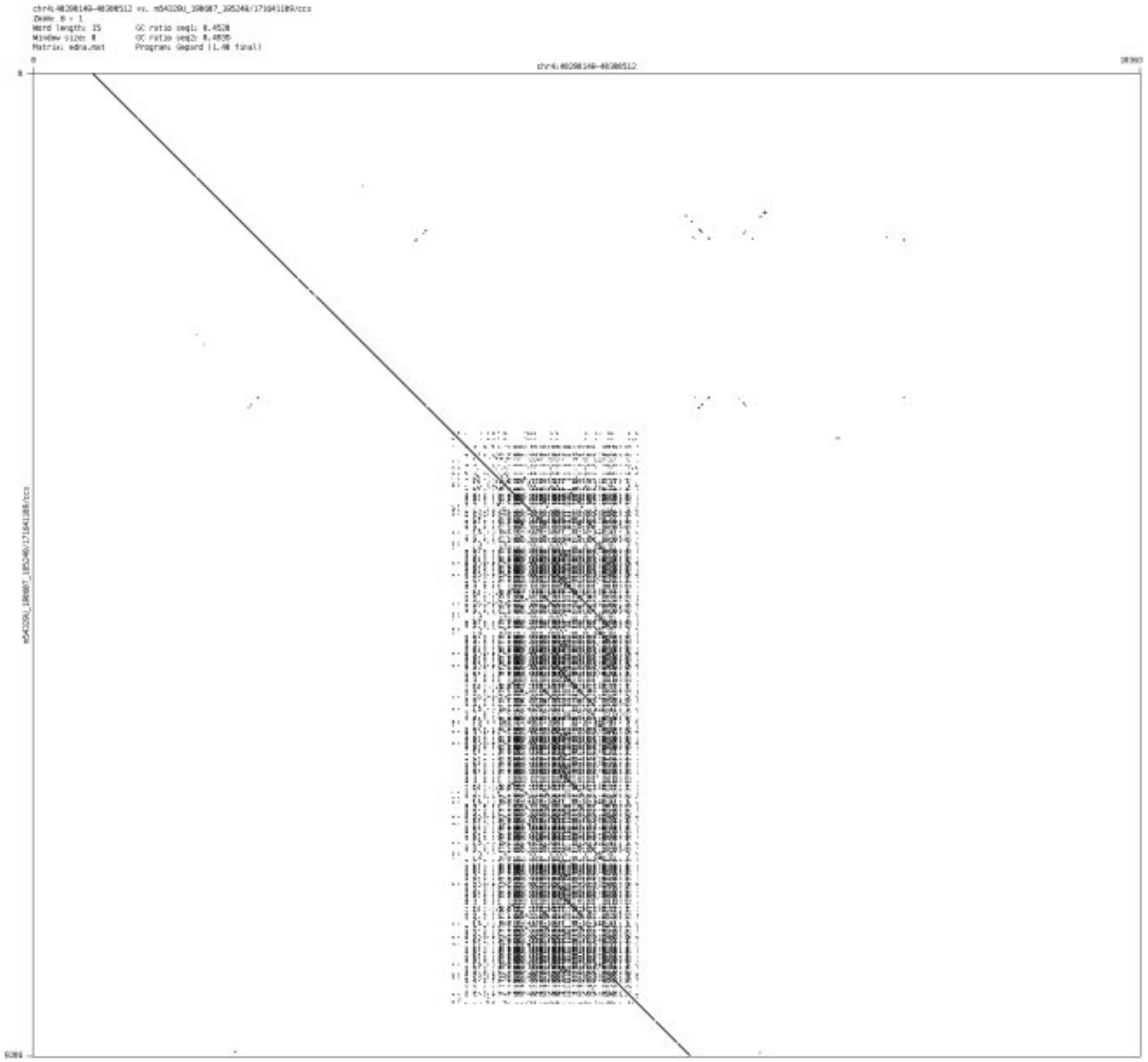

- chr4_146286299_146286463

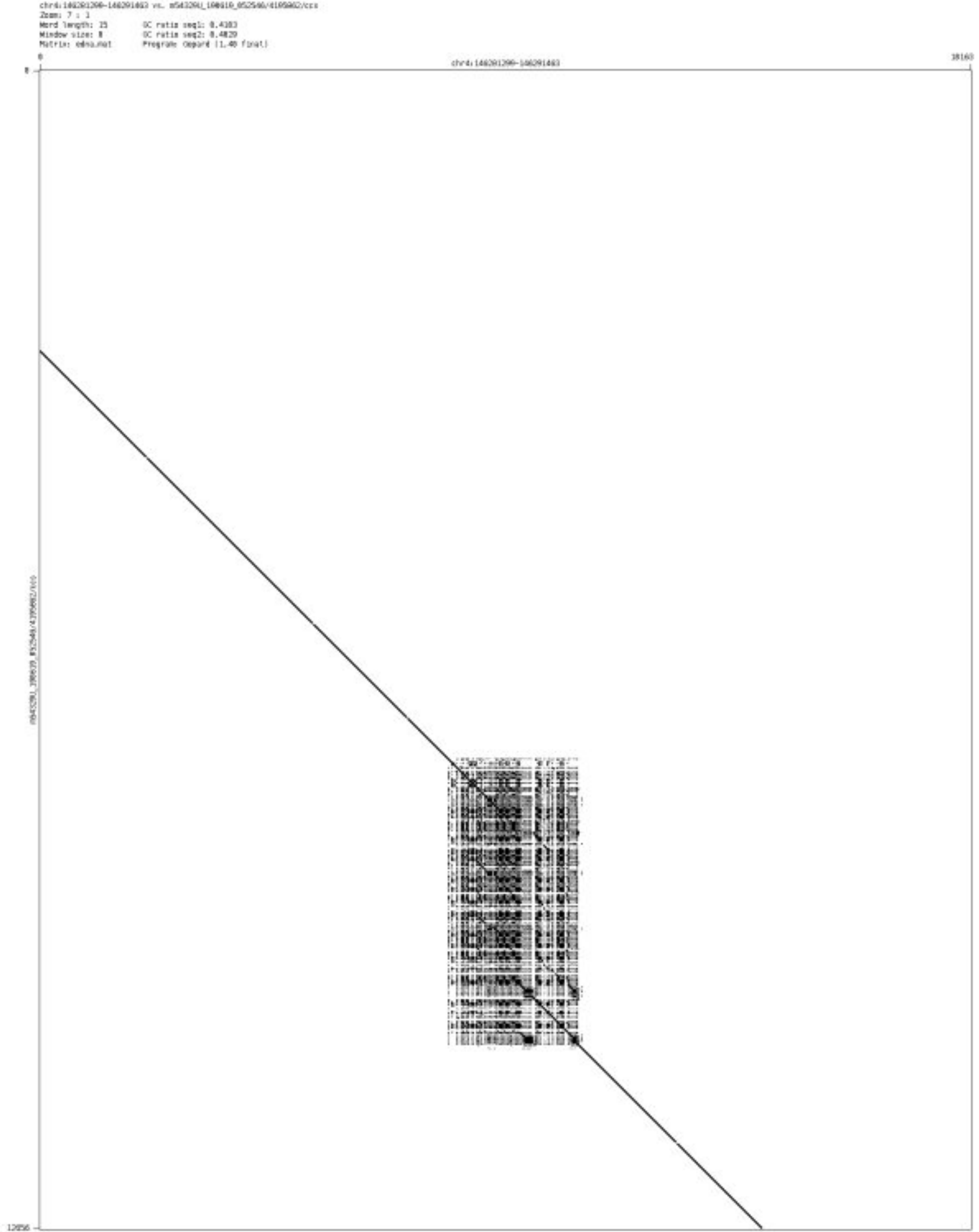

- chr5_3323810_3324066

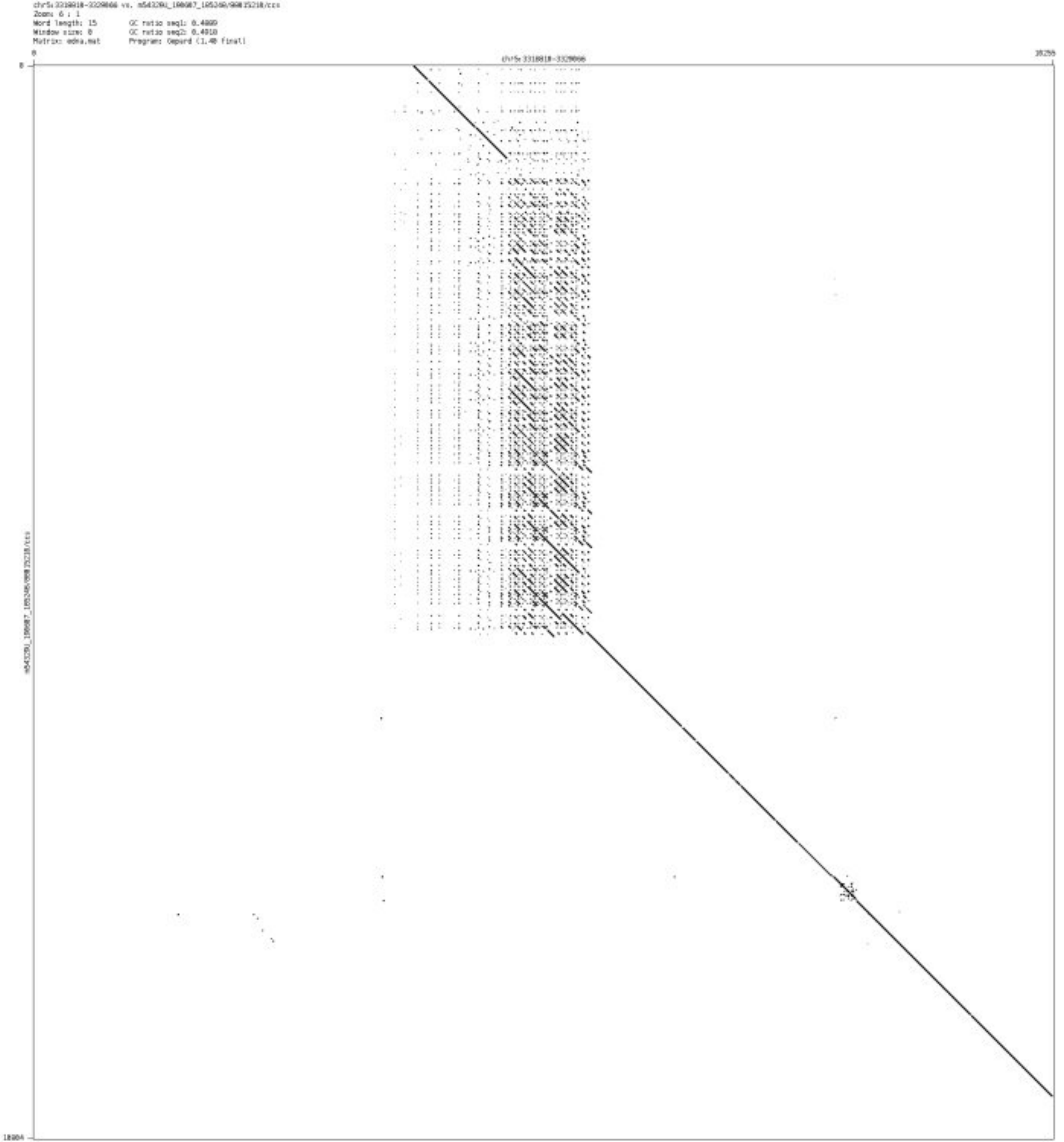

- chr5_35708616_35708706

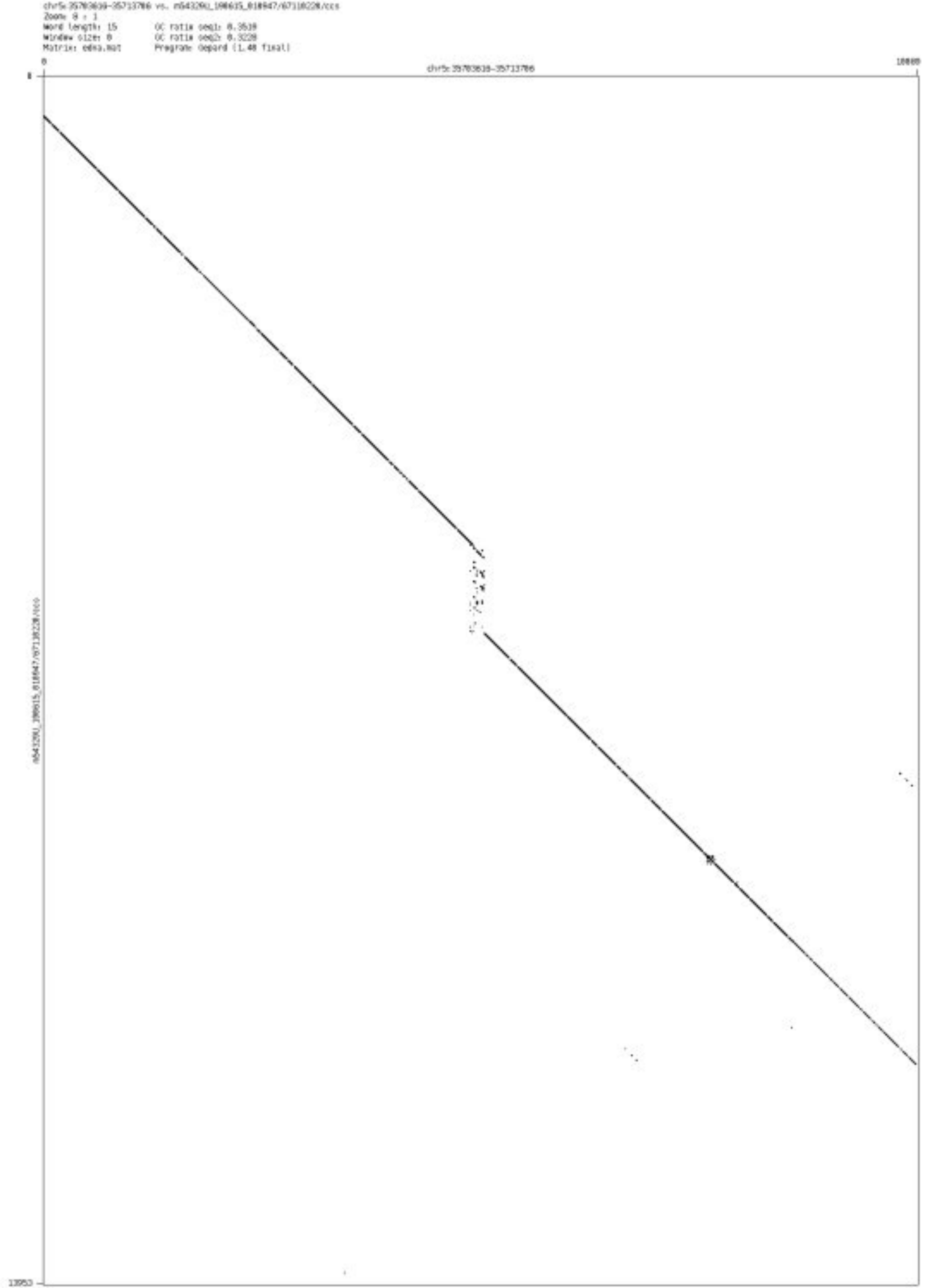

➢ chr6_3222124_3222284

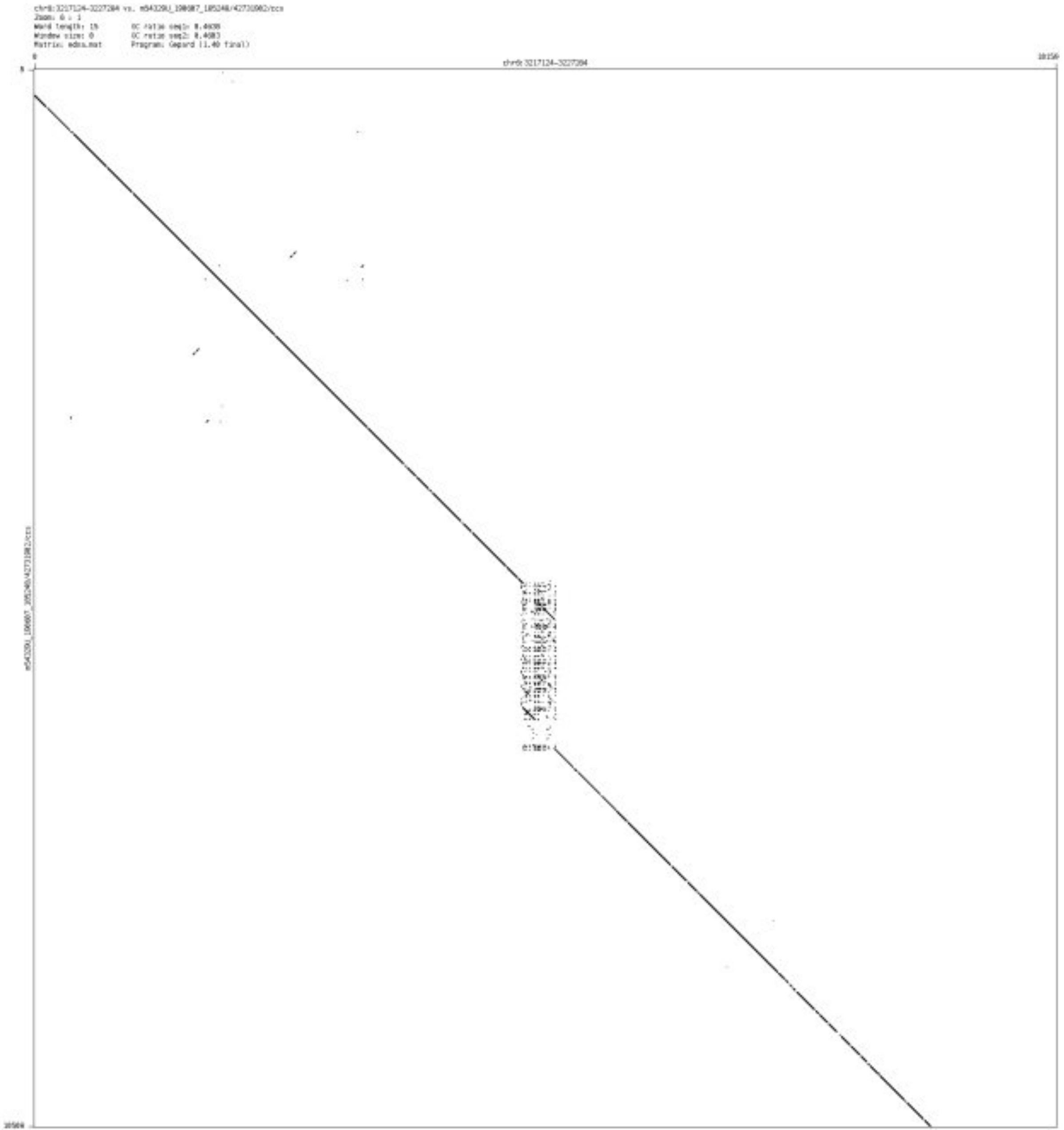

### ➢ chr6_34071455_34072974

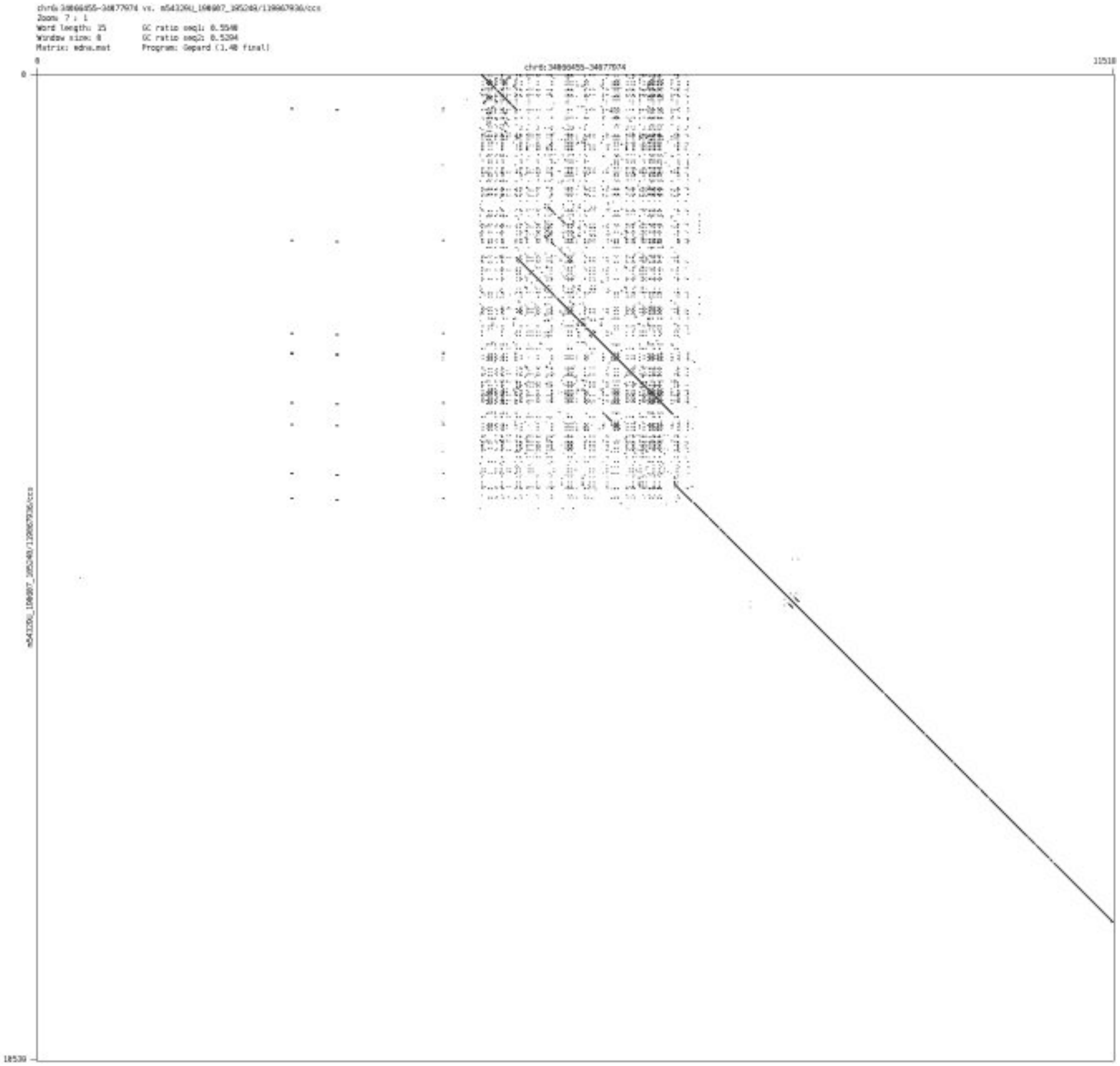

- chr6_160218975_160219097

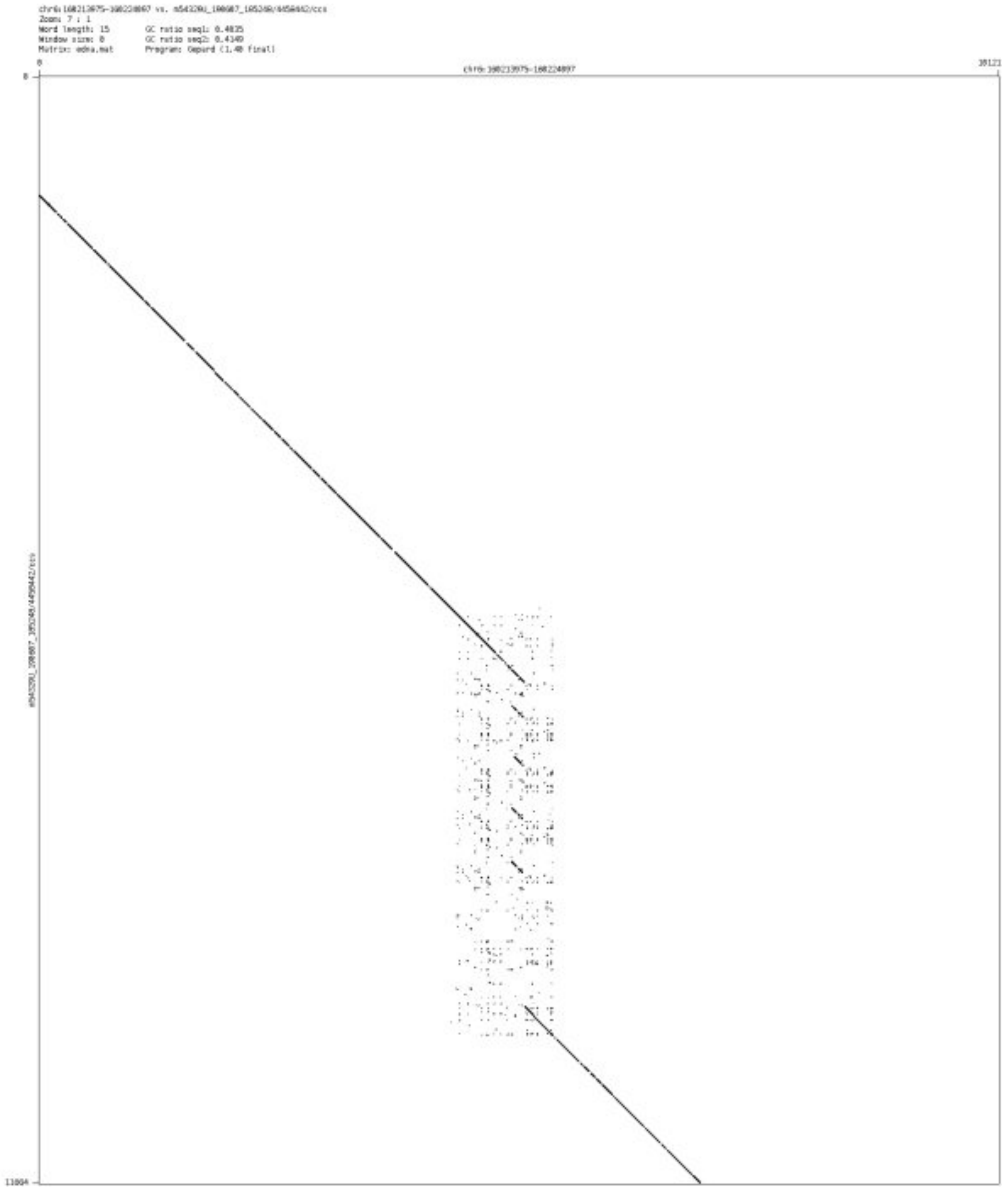

- chr6_167433114_167433256

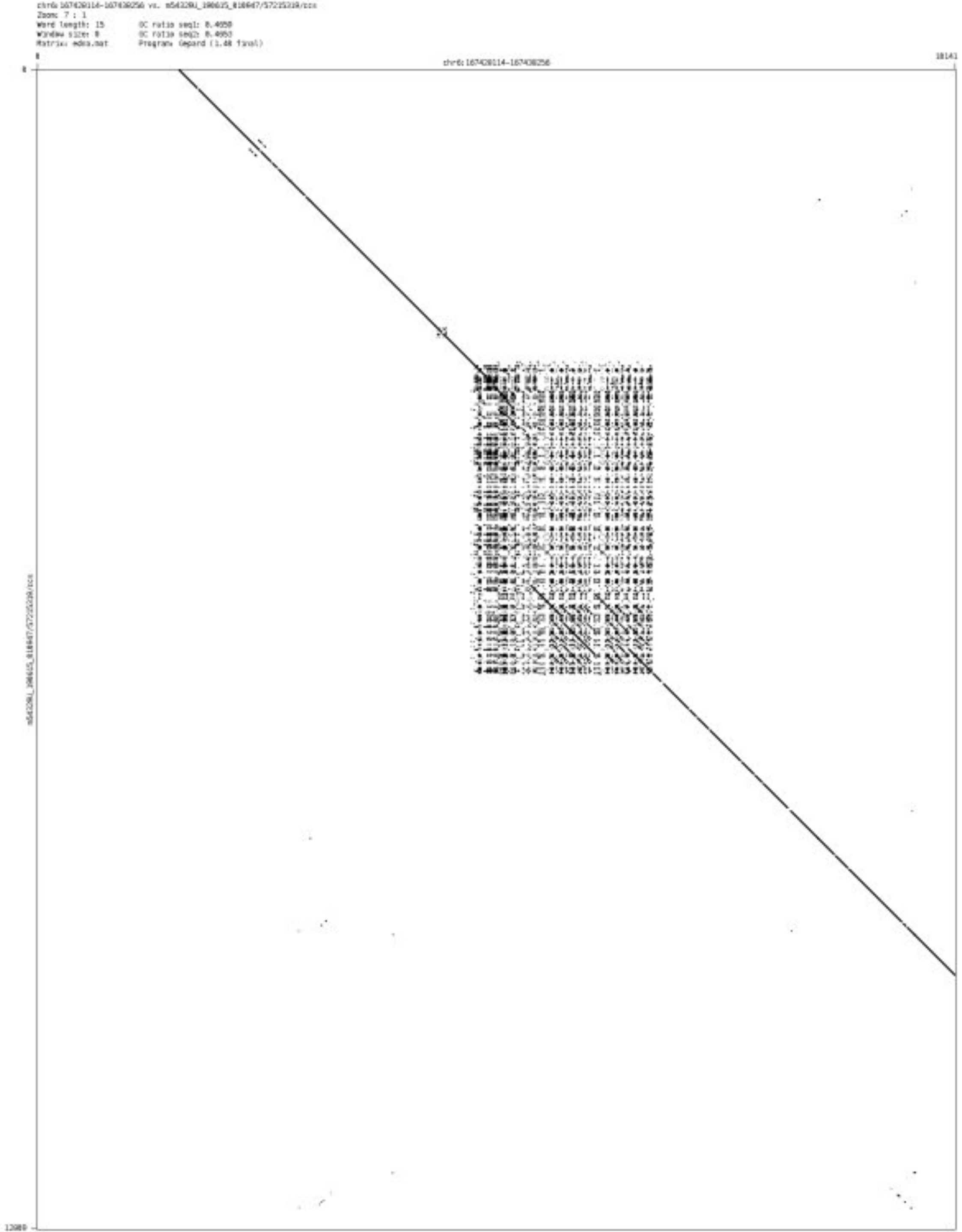

- chr6_167990058_167990217

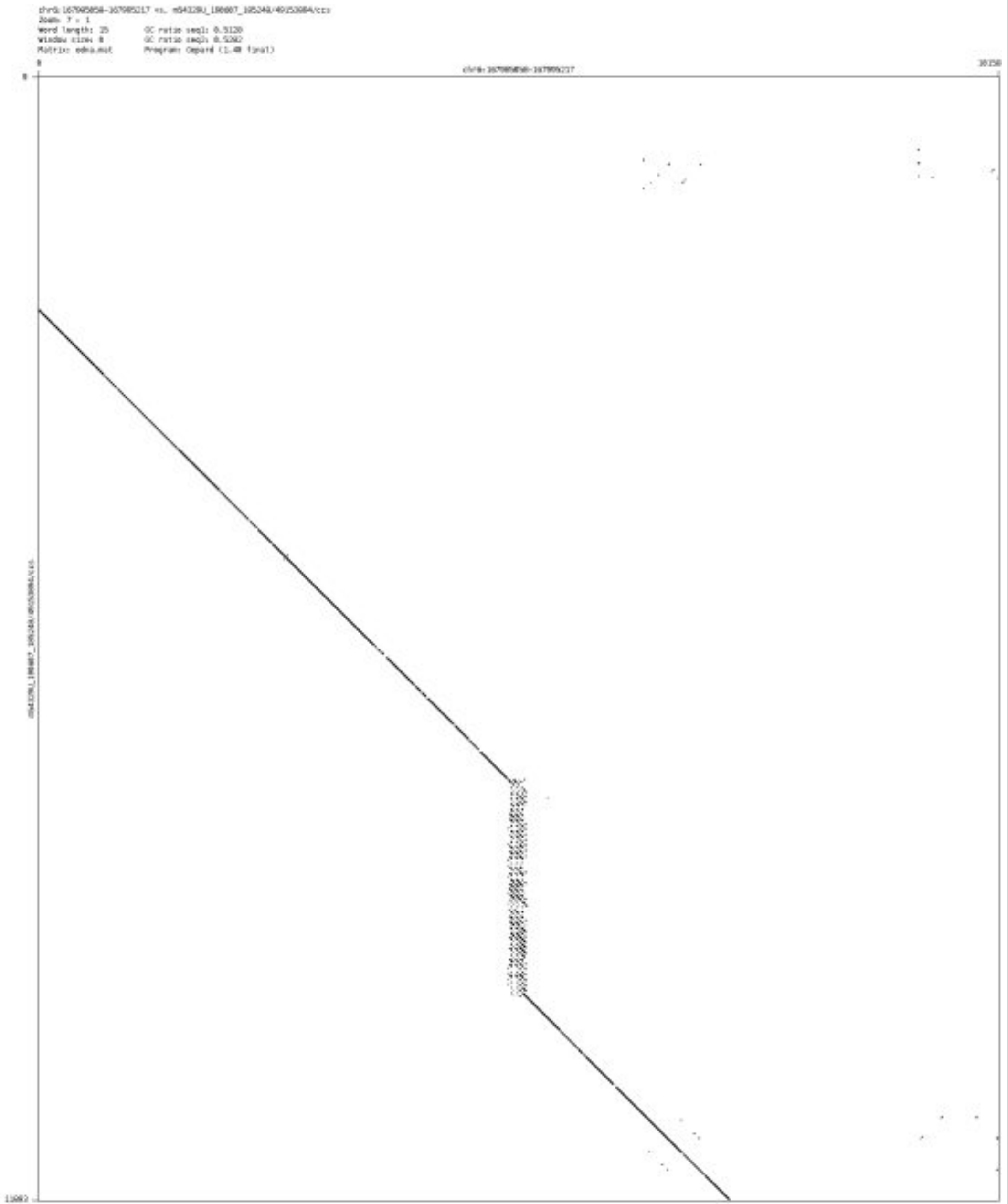

- chr6_168659501_168660054

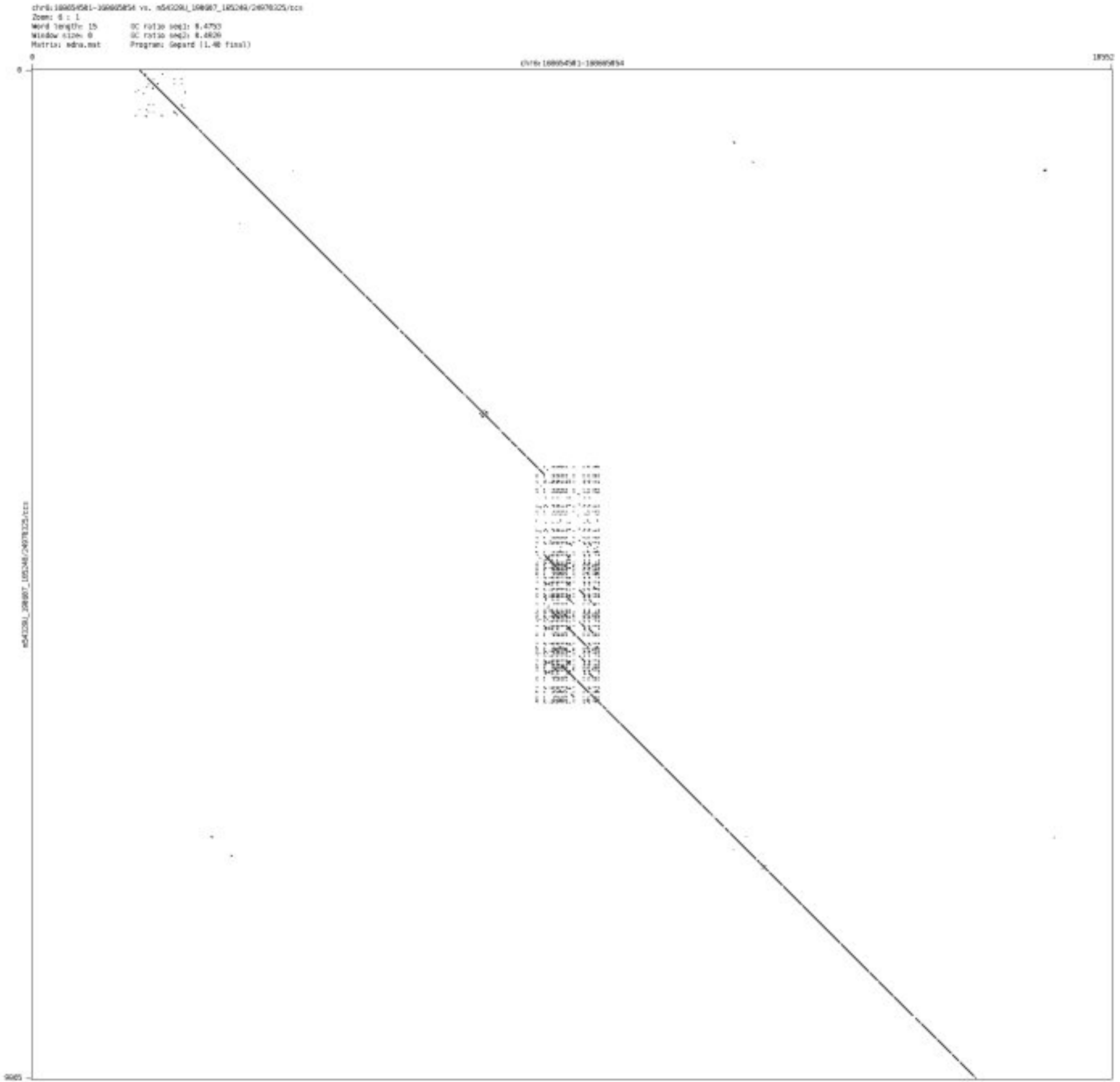

- chr7_155615228_155615438

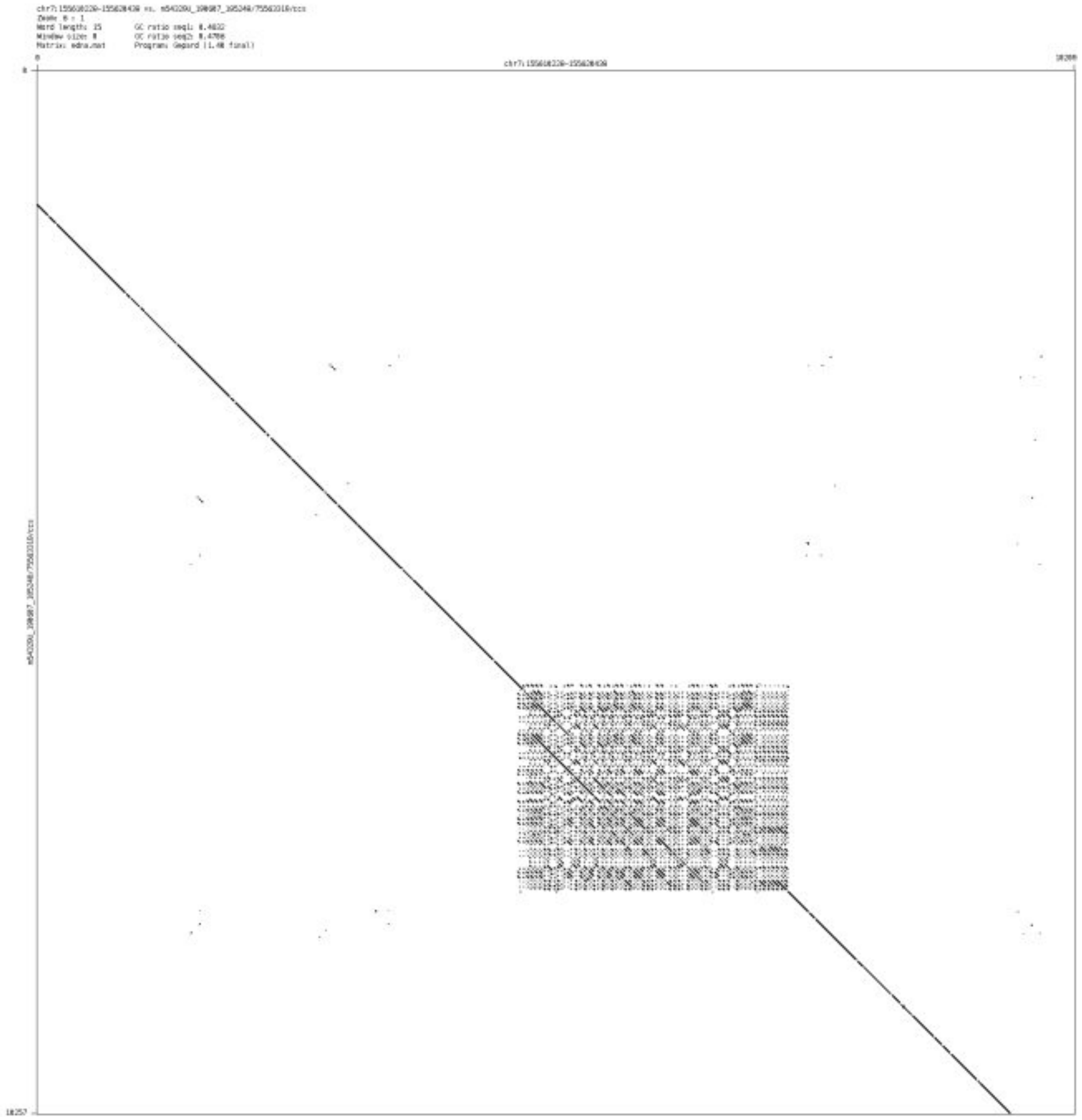

### ➢ chr7_158070331_158071546

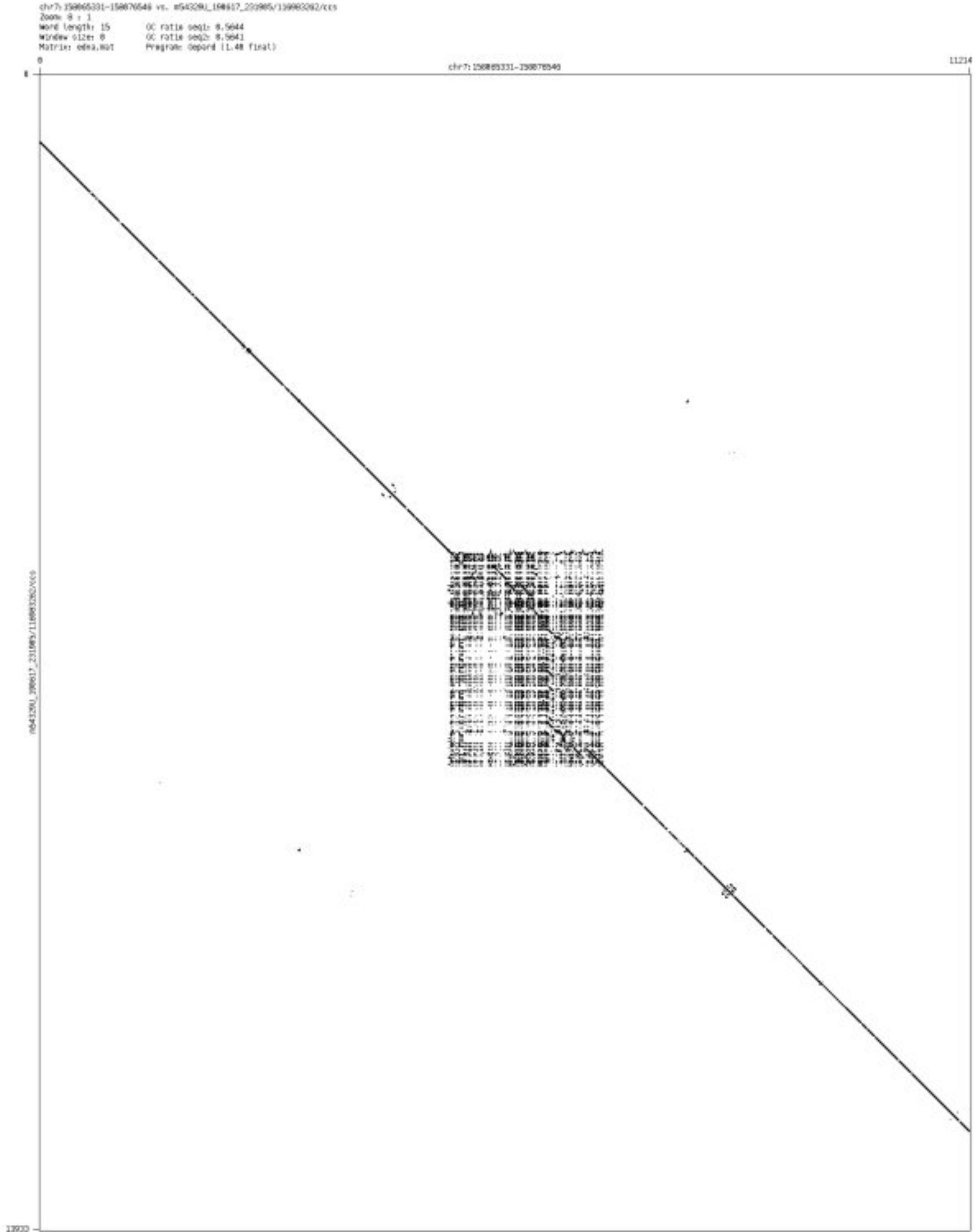

- chr8_11456240_11456367

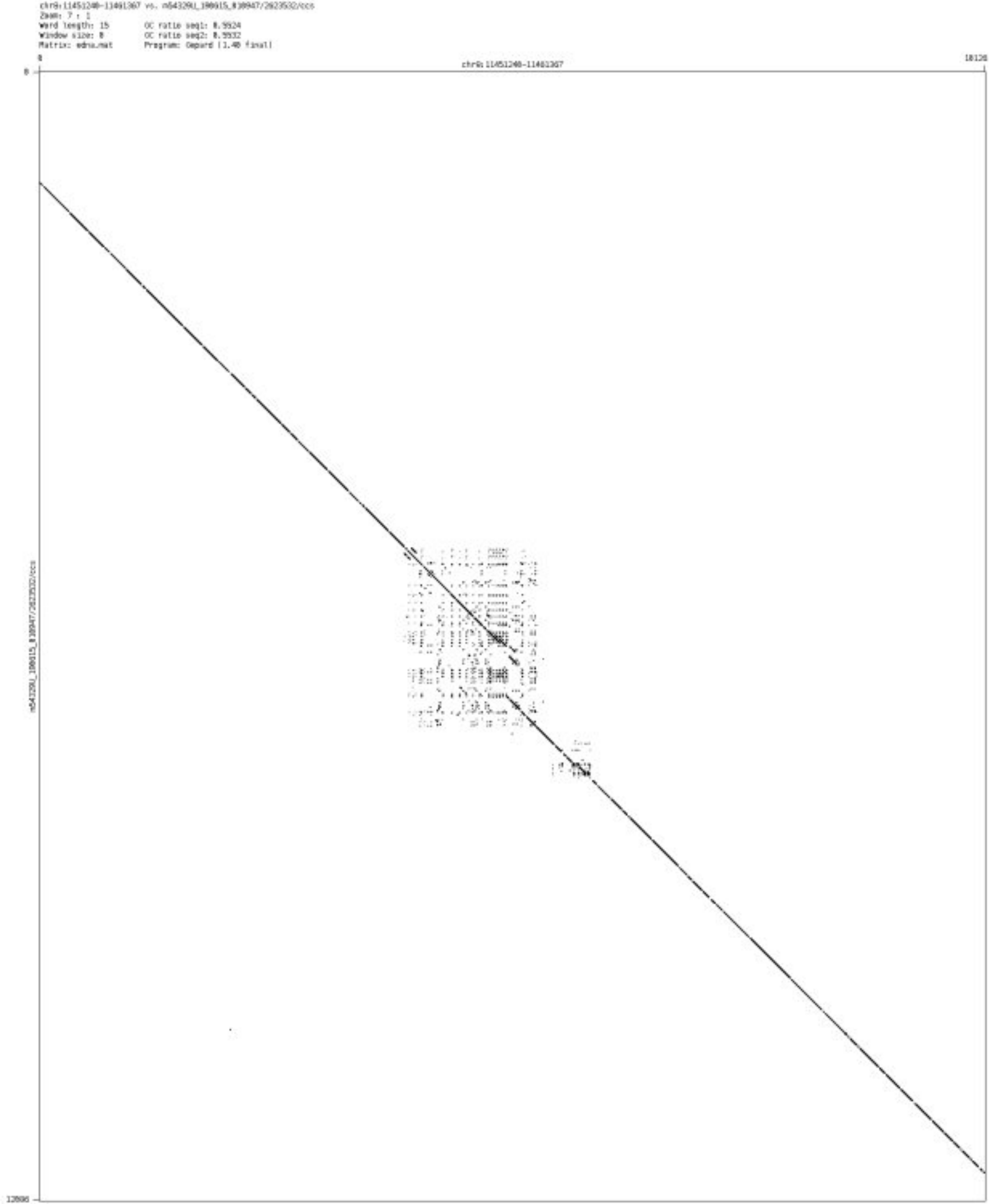

- chr8_138691548_138692228

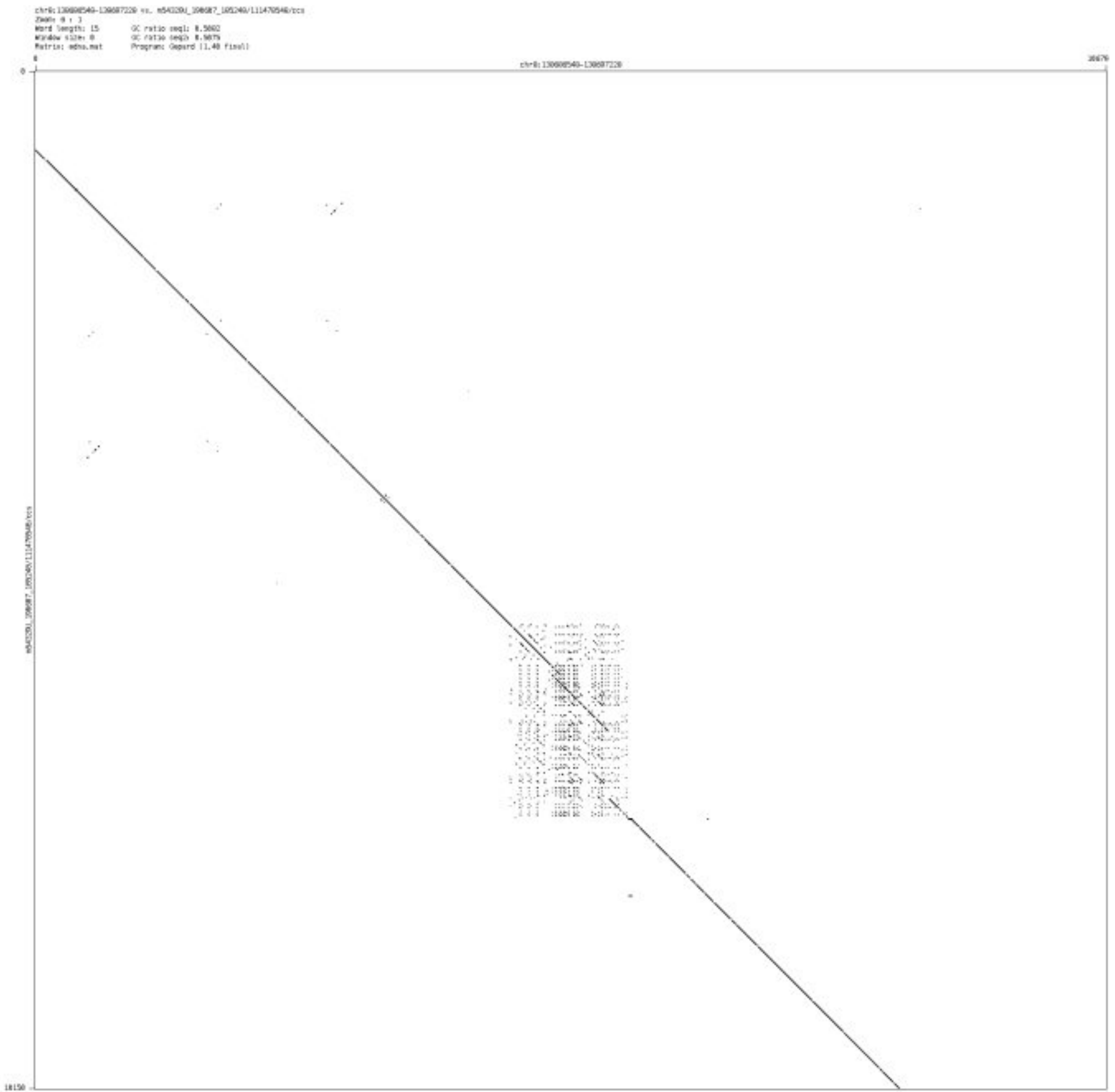

➢ chr8_141871771_141874394

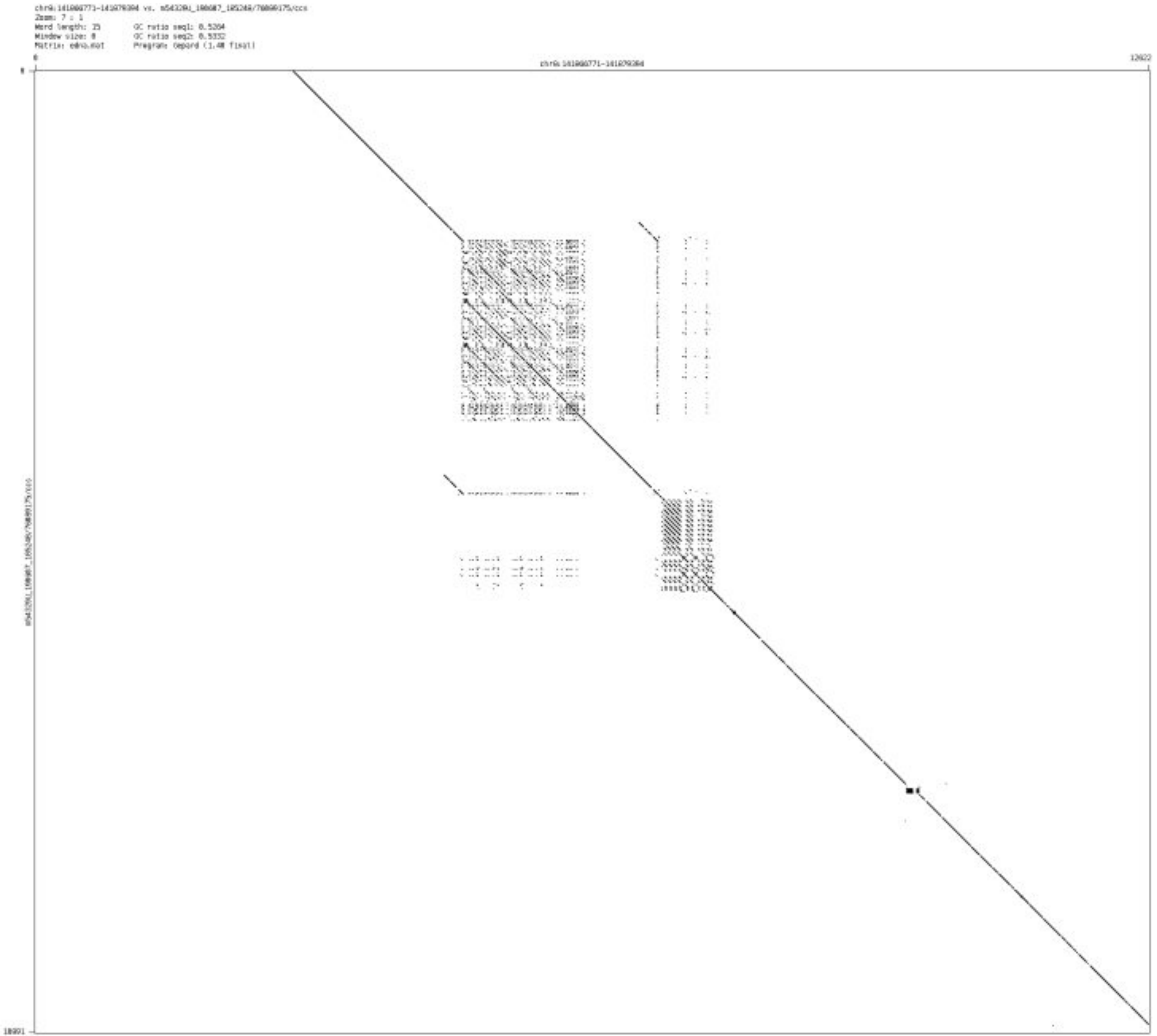

- ➢ chr8_142012915_142013093

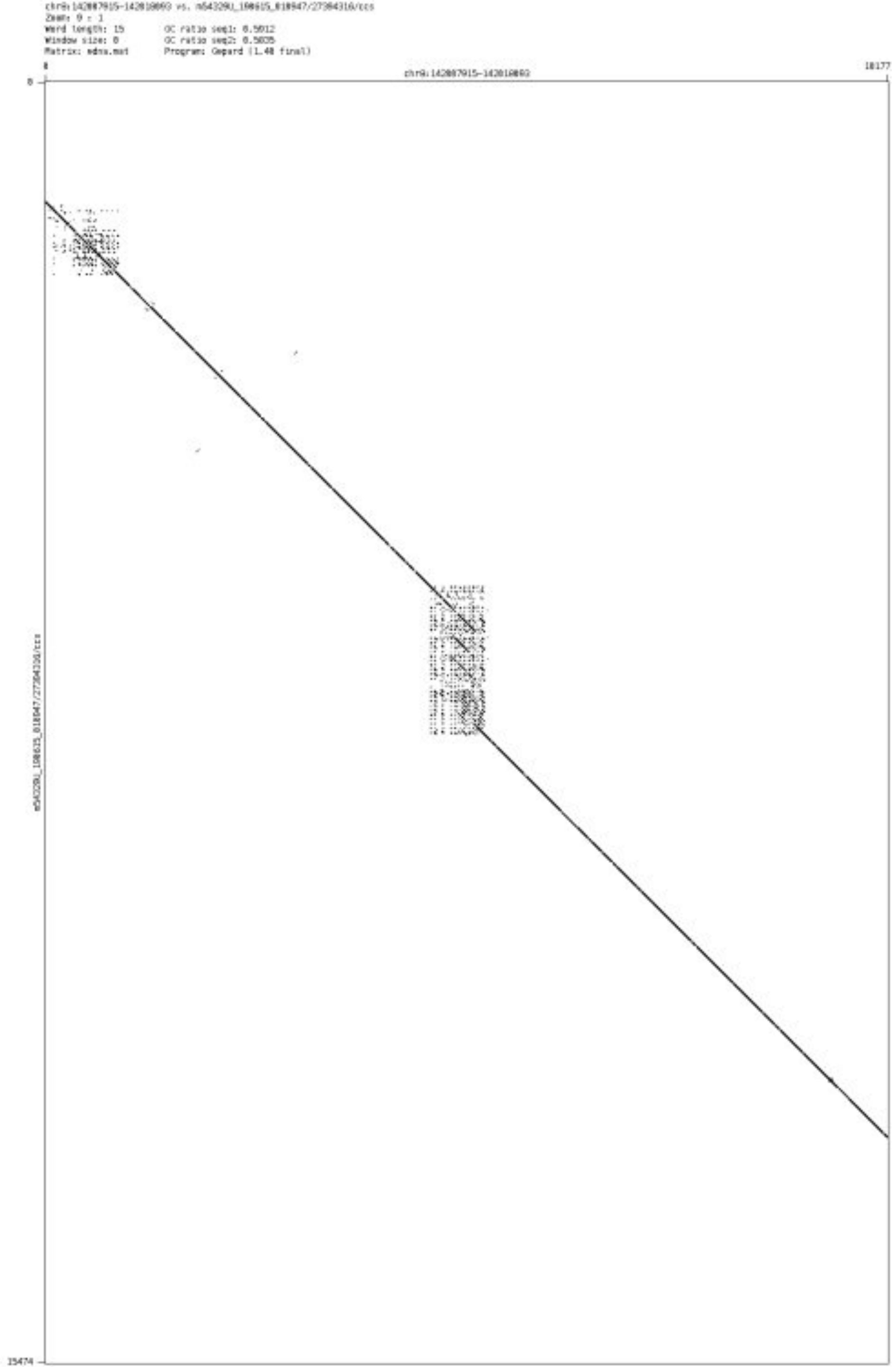

➢ chr8_144020570_144024873

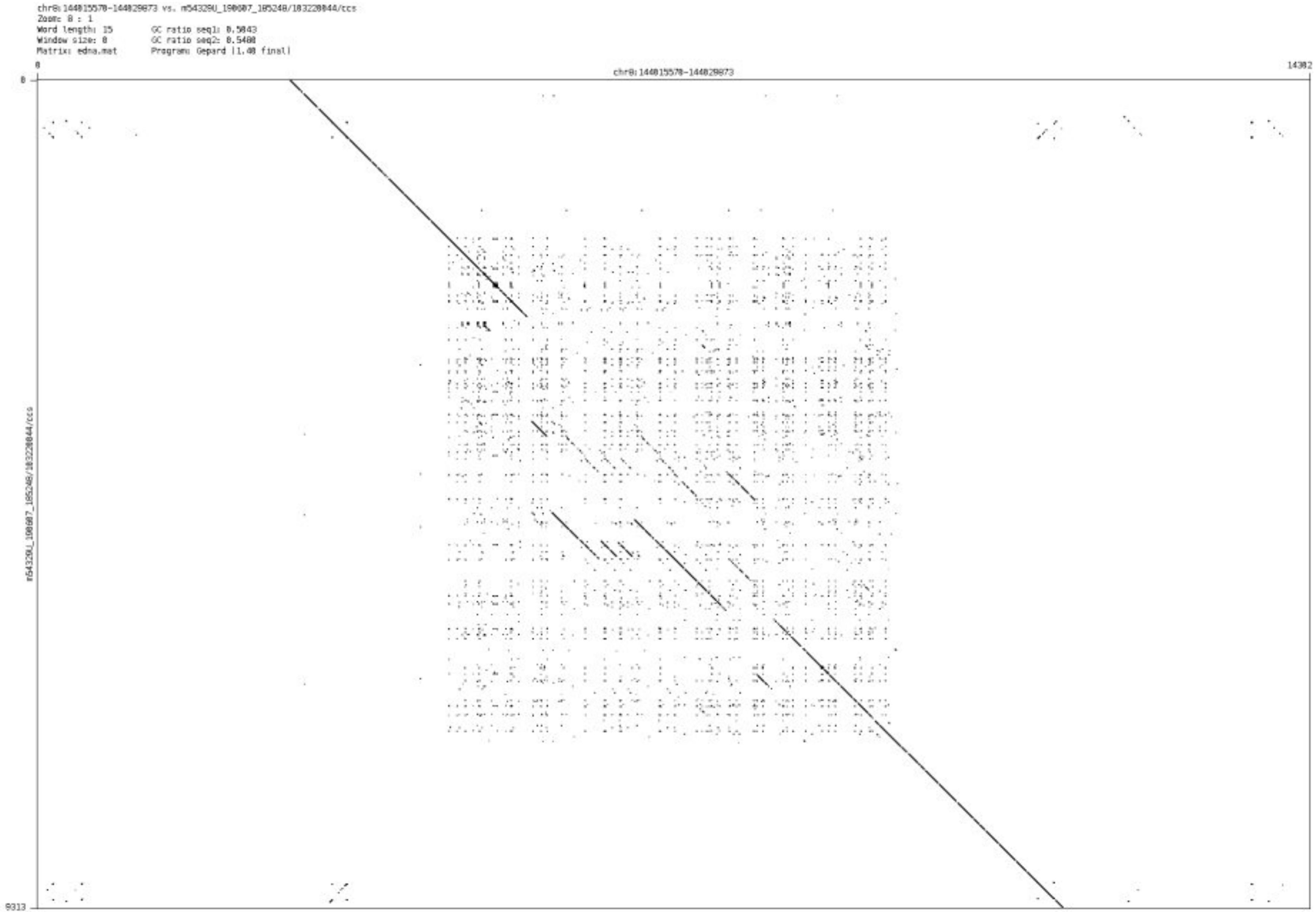

- ➢ chr8_144946952_144947229

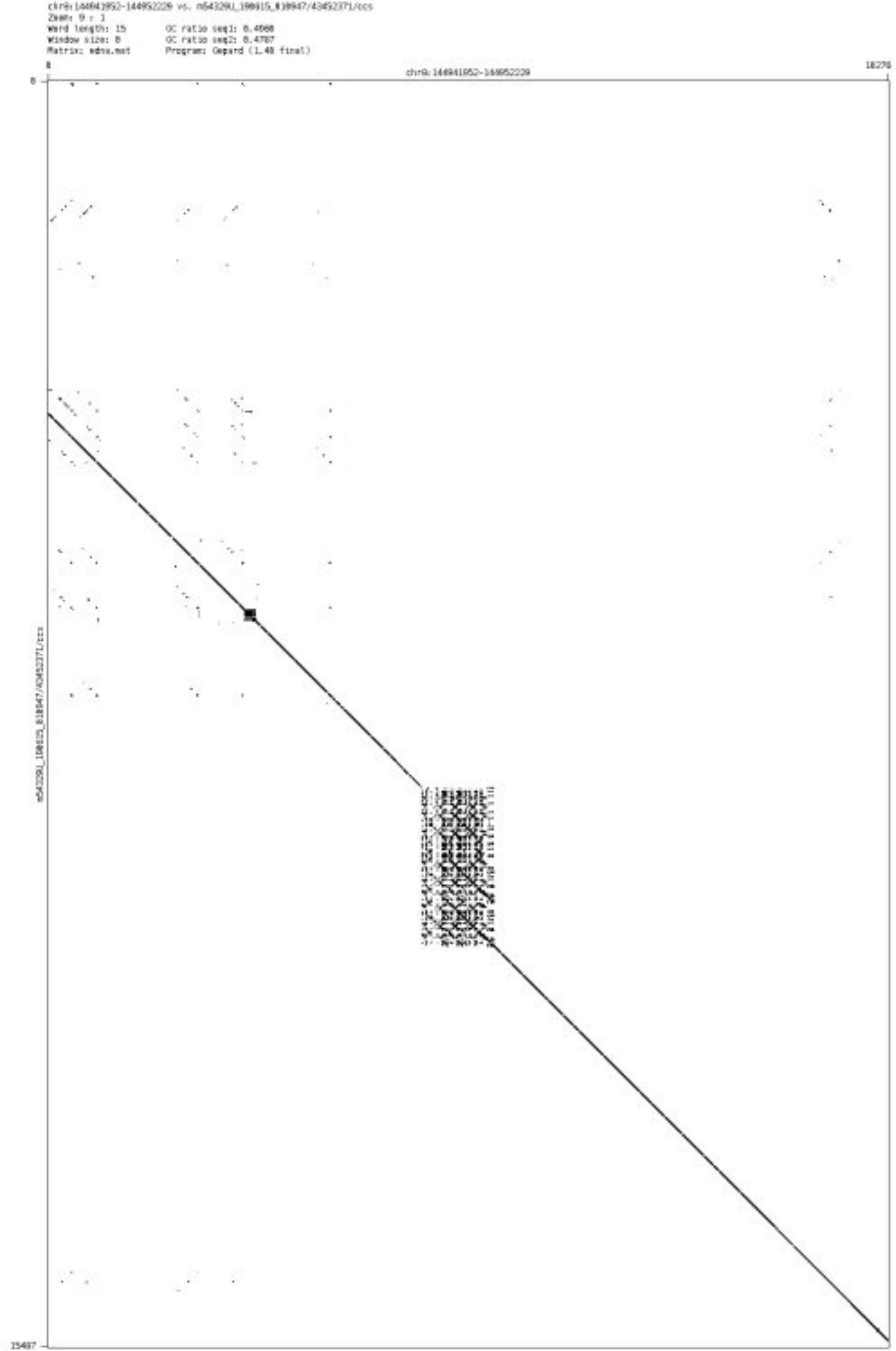

- chr9_86052967_86053698

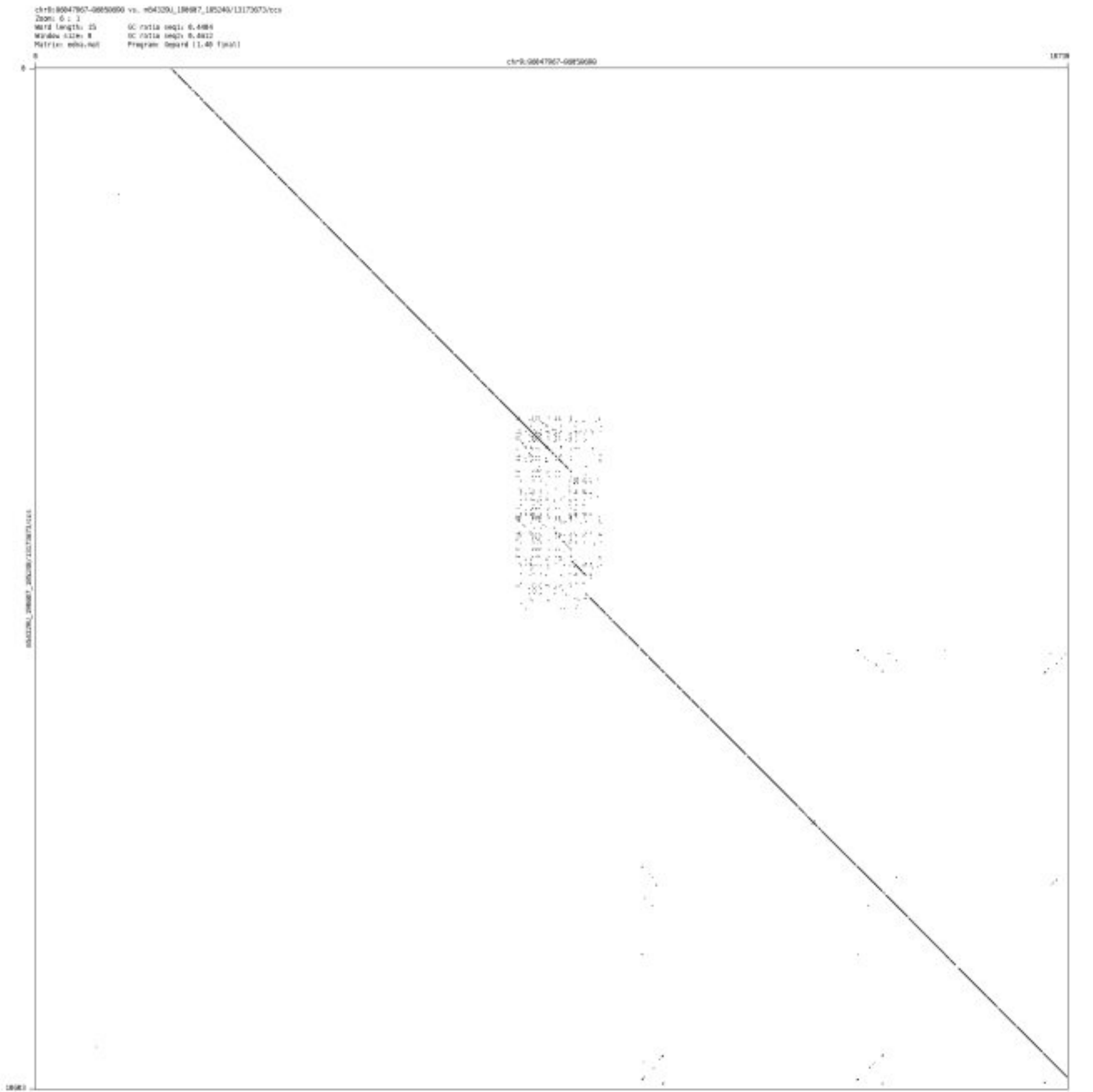

➢ chr9_89254239_89254375

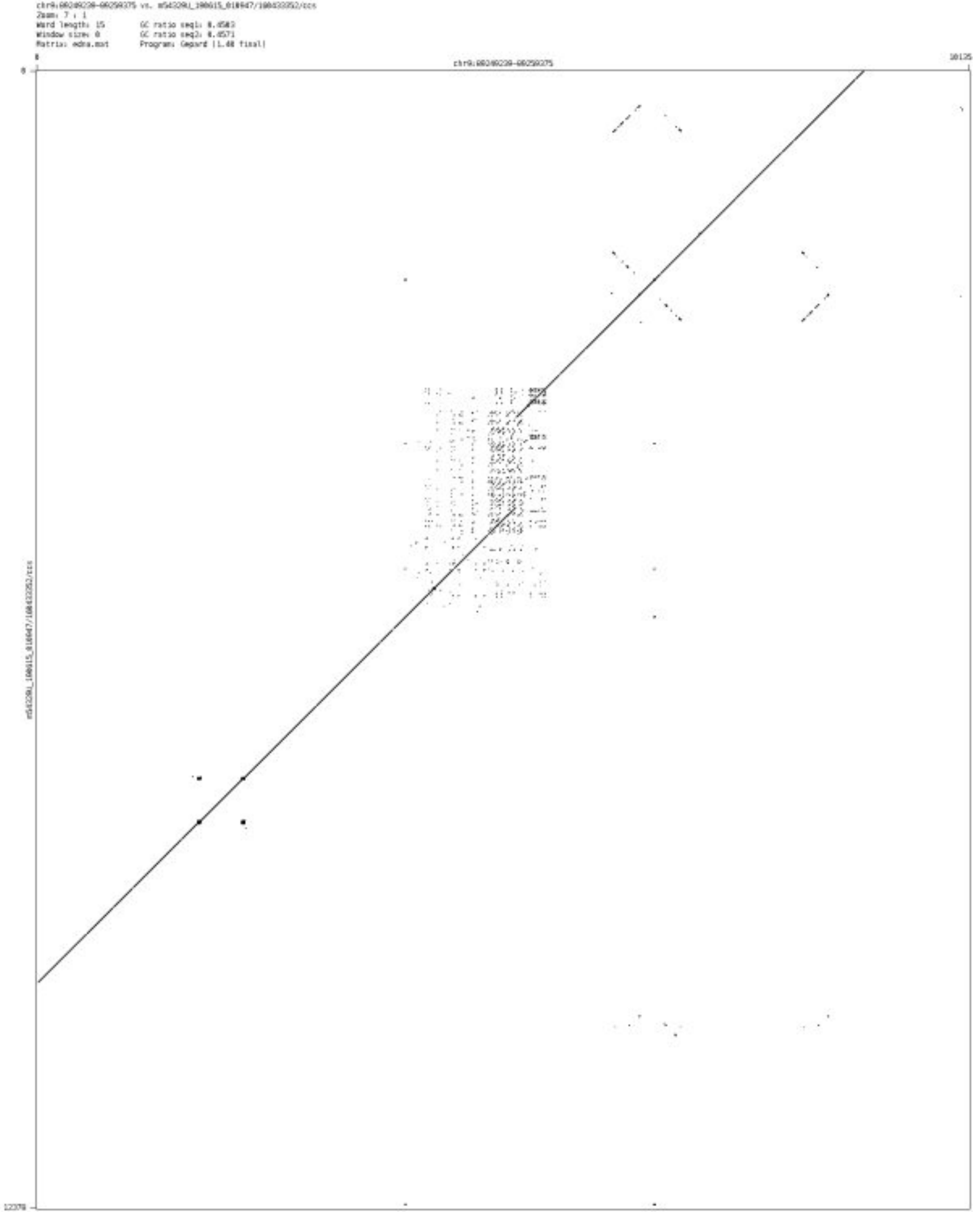

- chr9_91695537_91695651

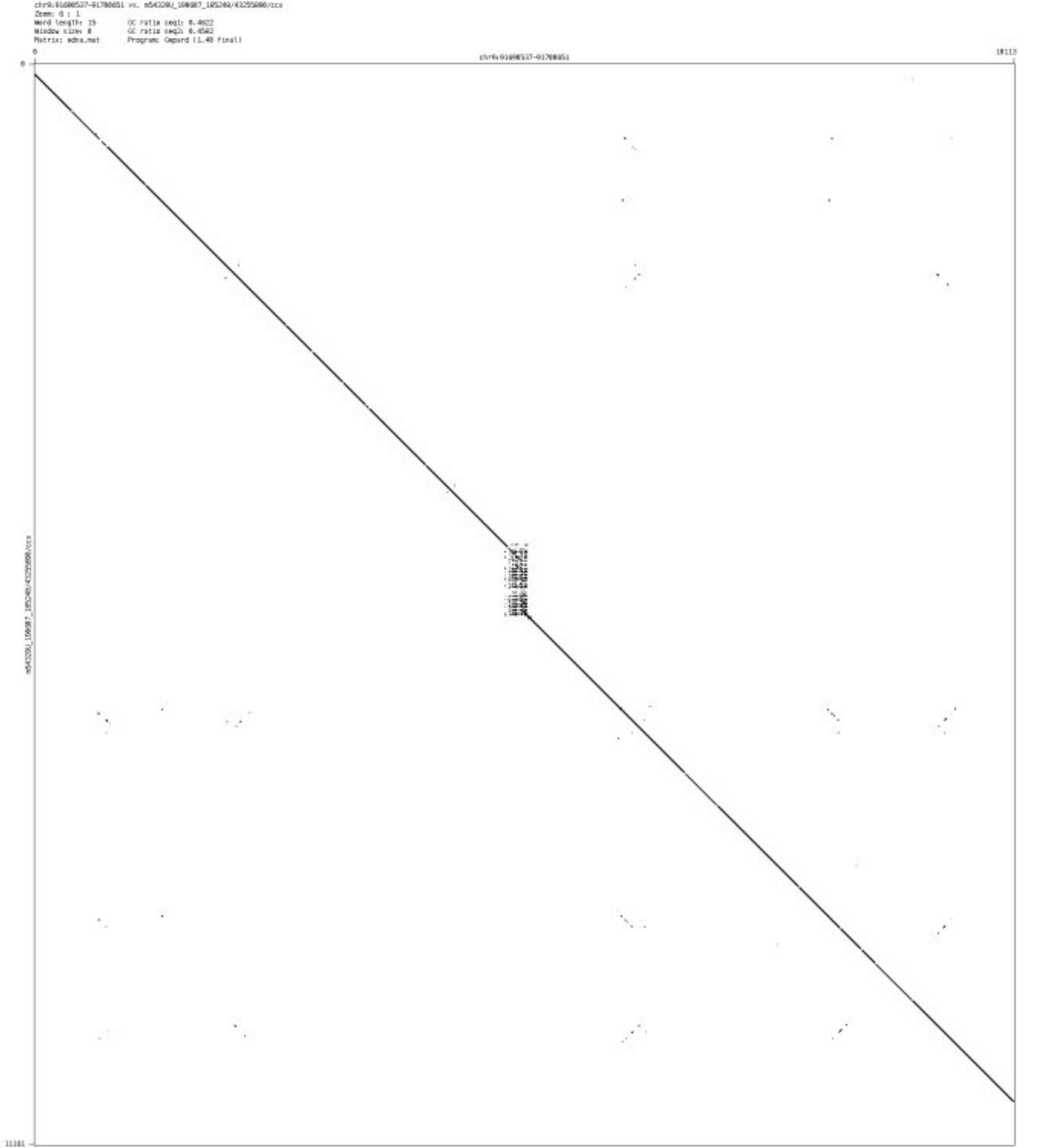

- chr9_130955139_130955274

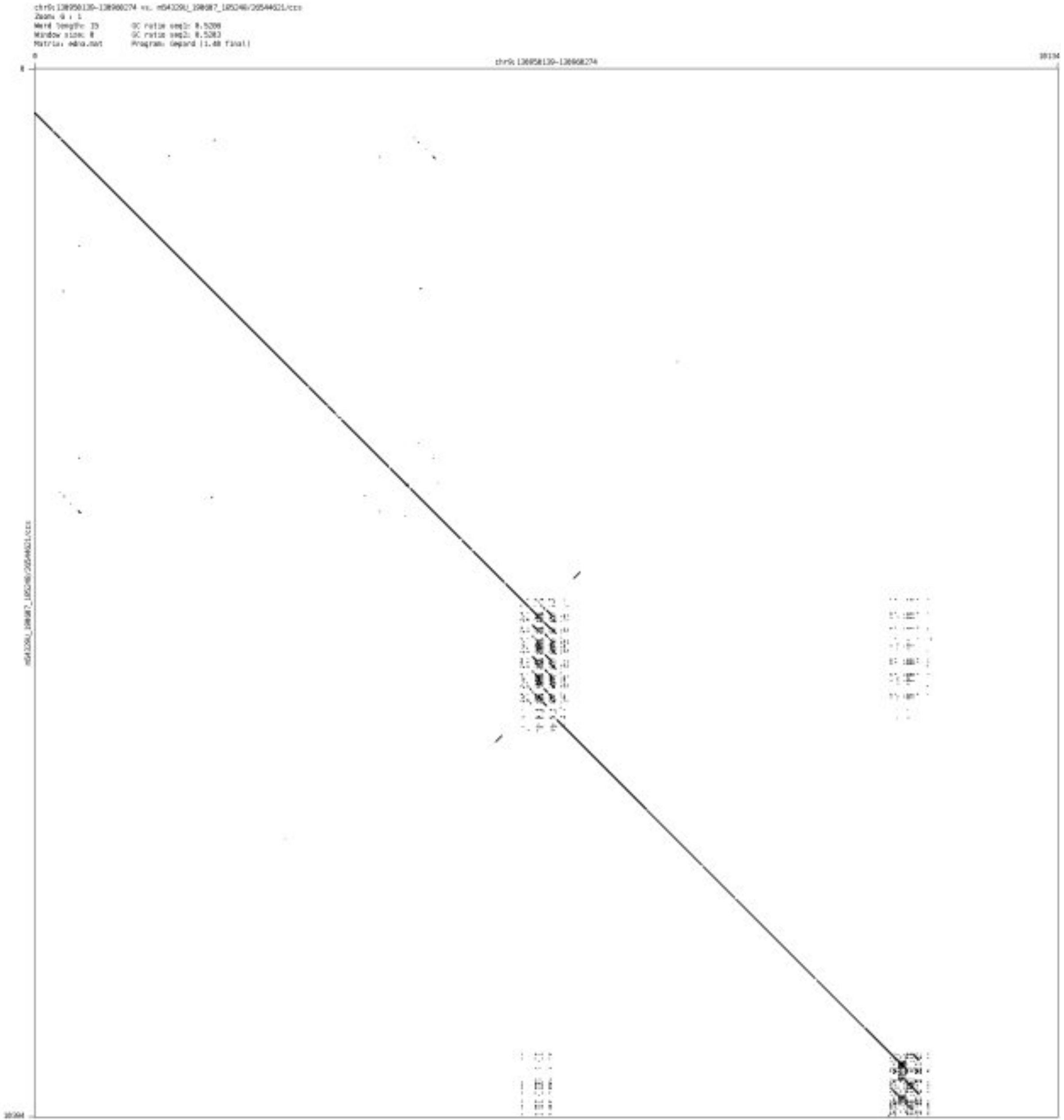

➢ chr9_134685195_134685386

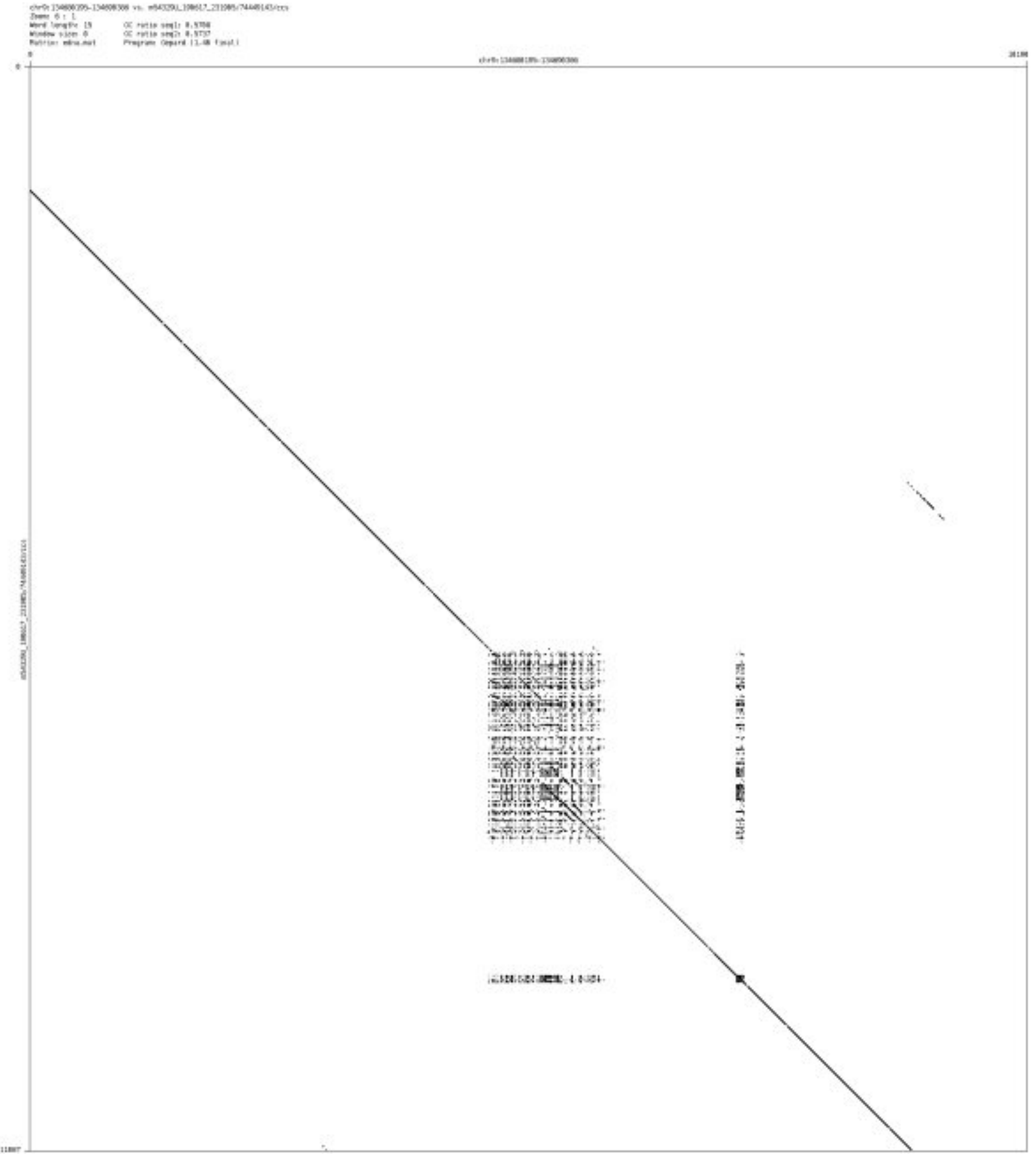

➢ chr9_136632692_136632989

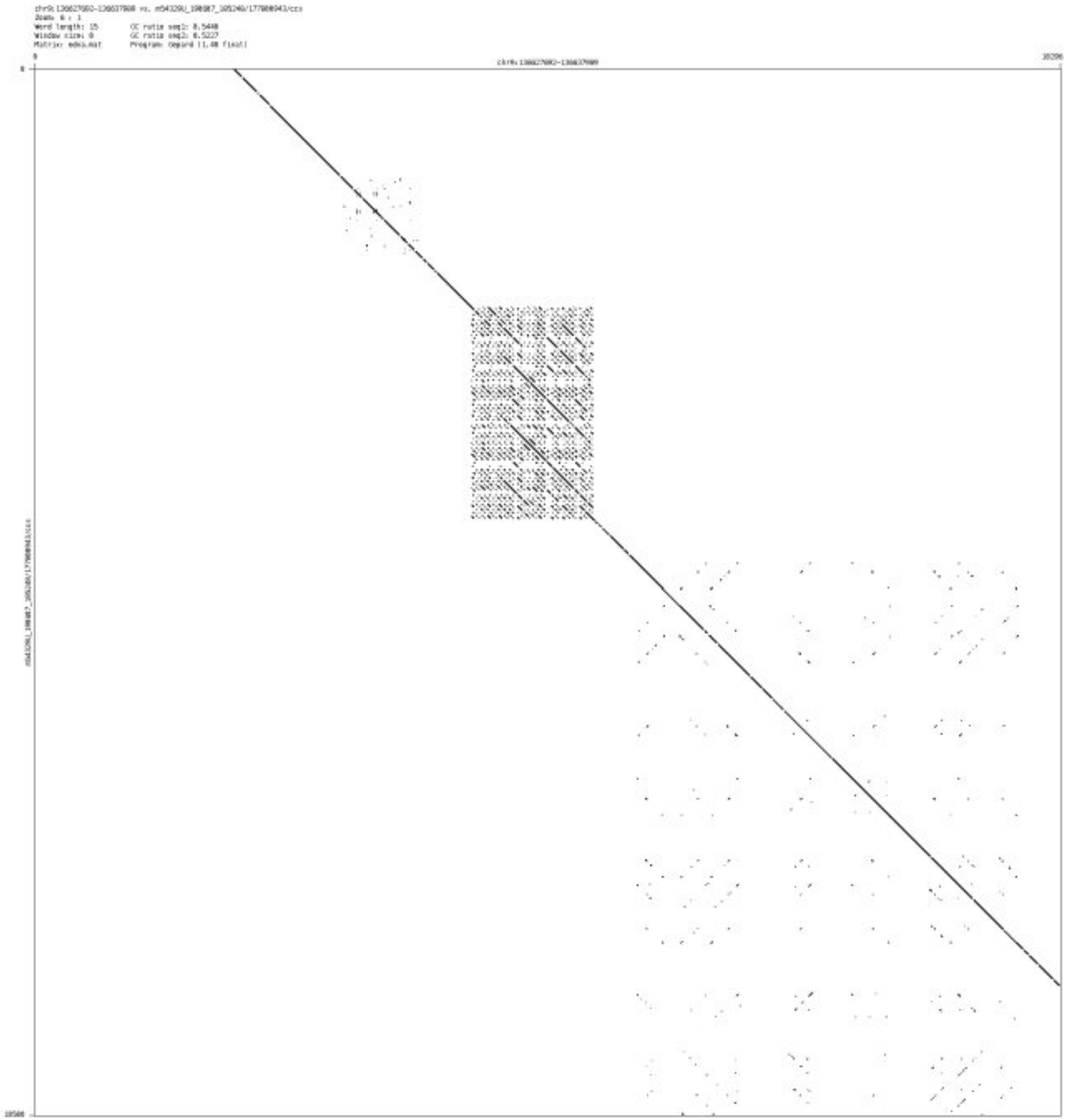

- chr9_137518162_137518324

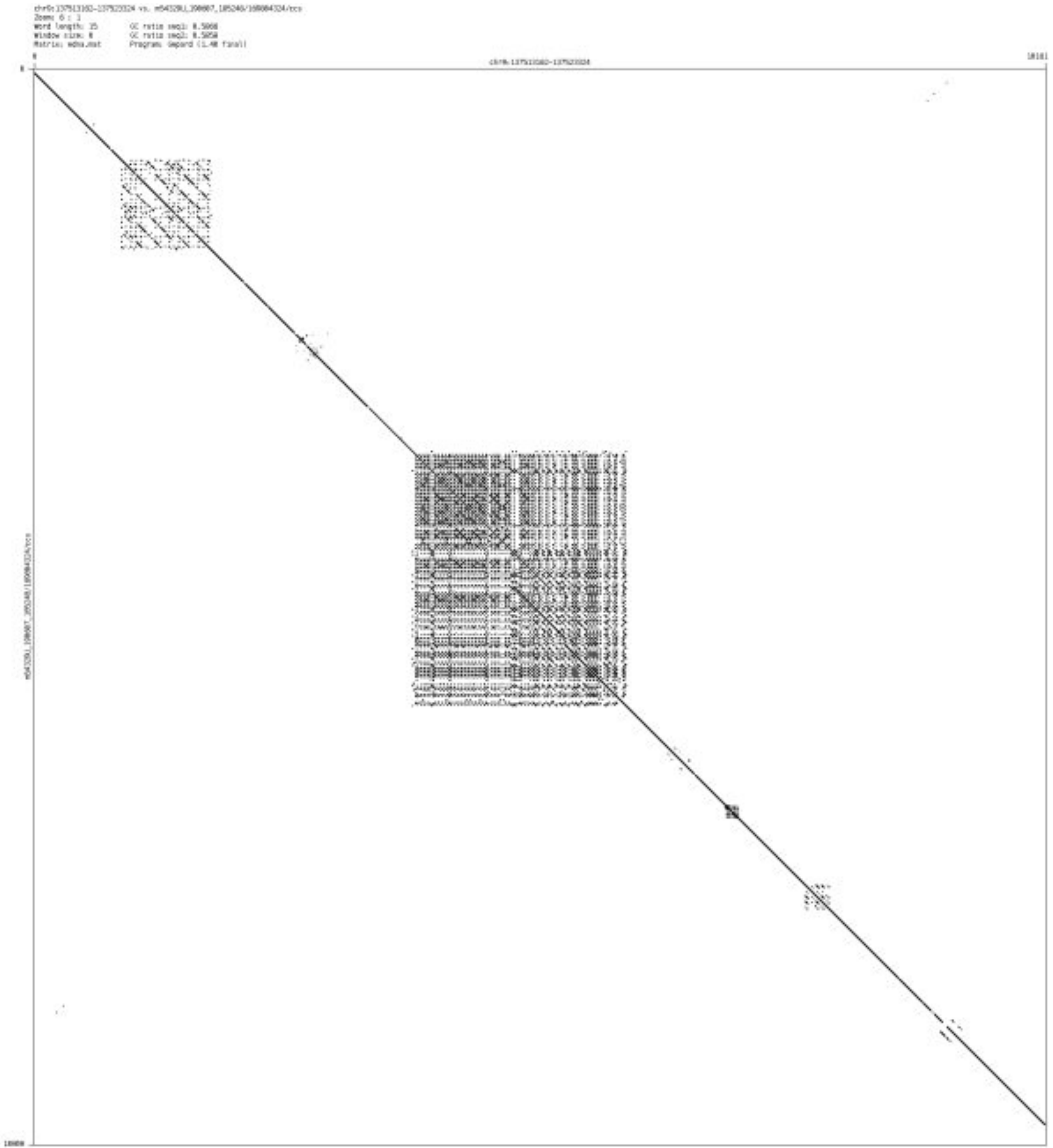

➢ chr10_735885_737771

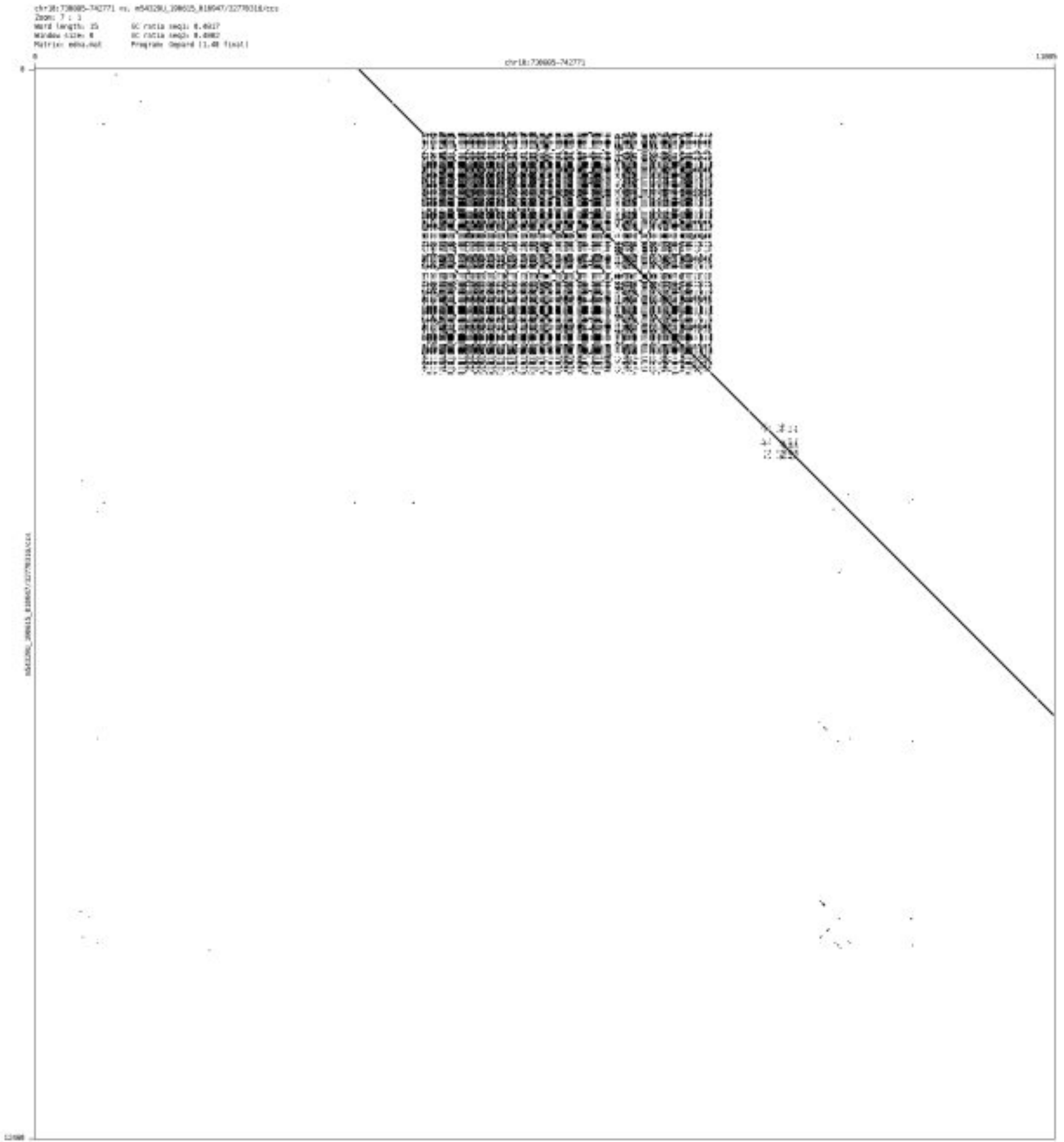

➢ chr10_5577237_5577333

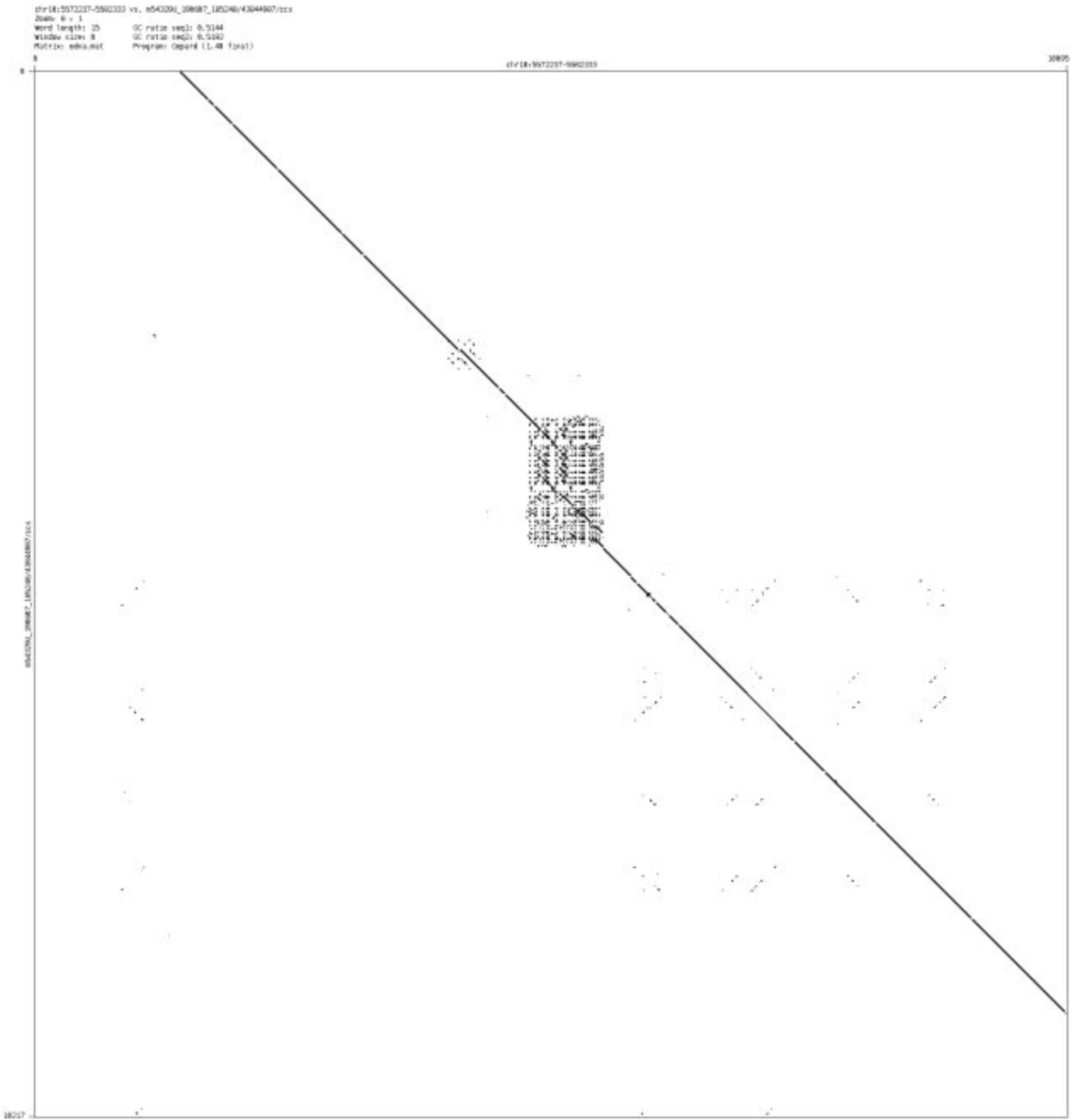

- chr10_11283968_11284497

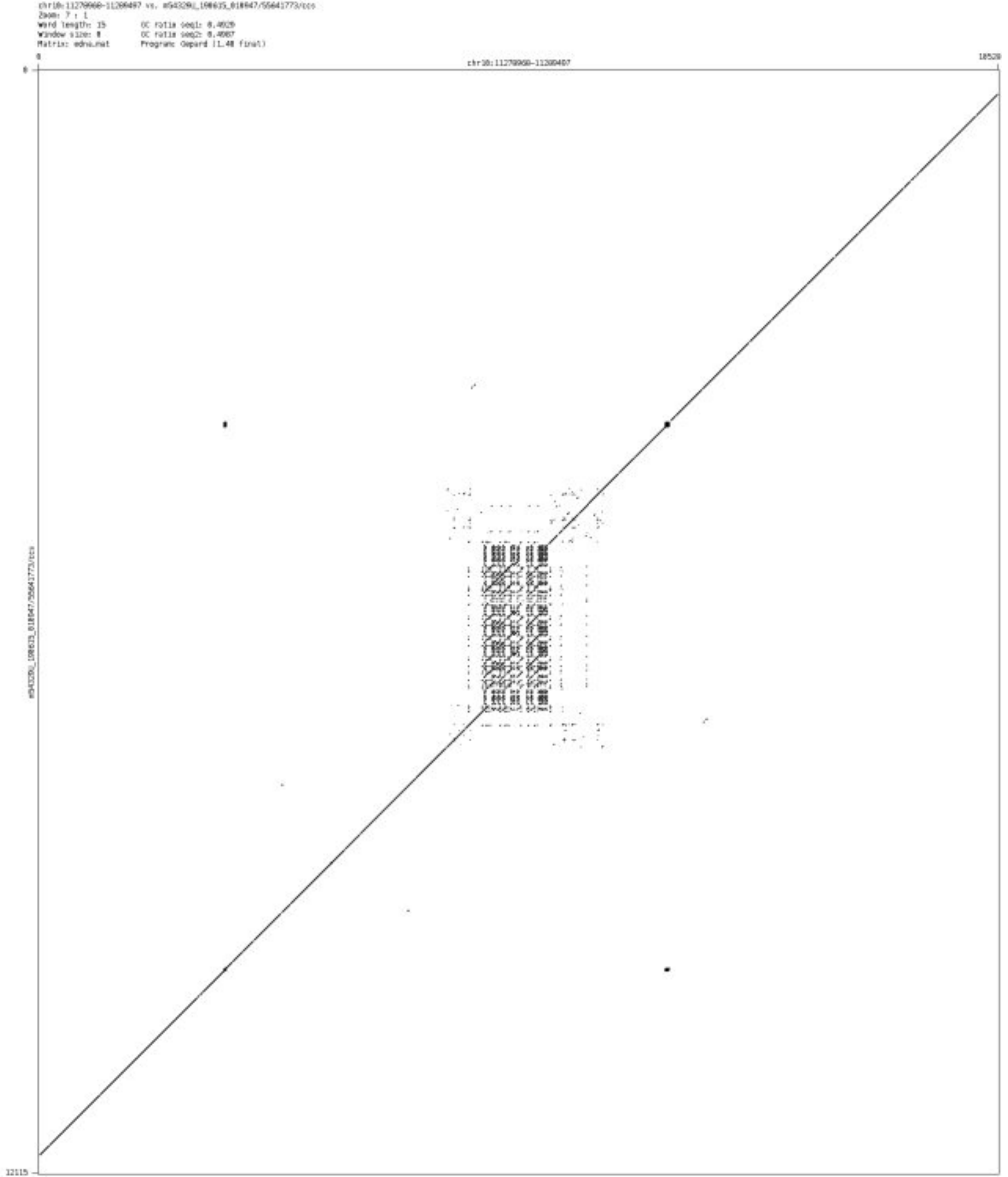

➢ chr10_12413511_12413590

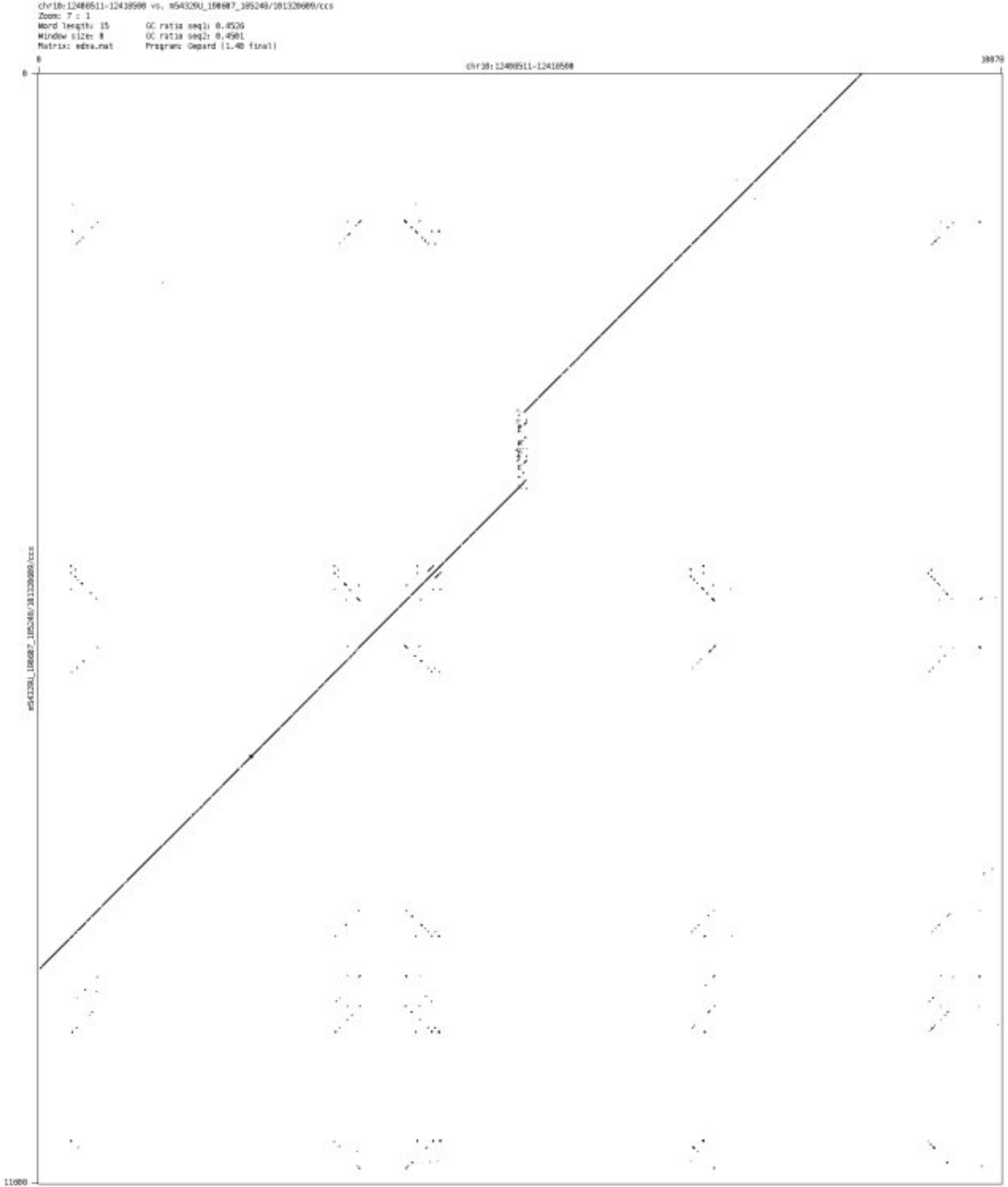

- ➢ chr10_14568488_14568677

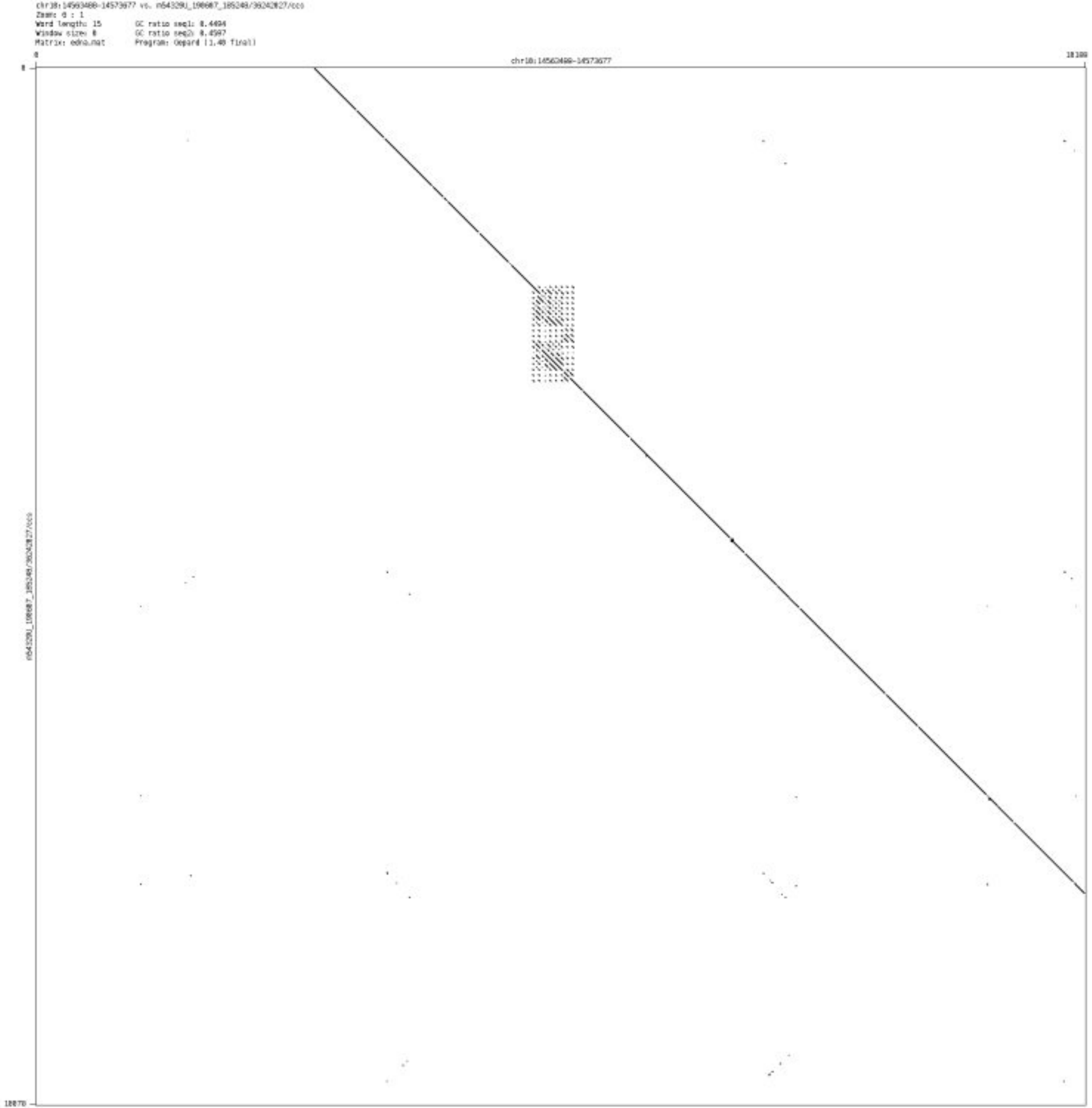

### ➢ chr10_123254394_123255170

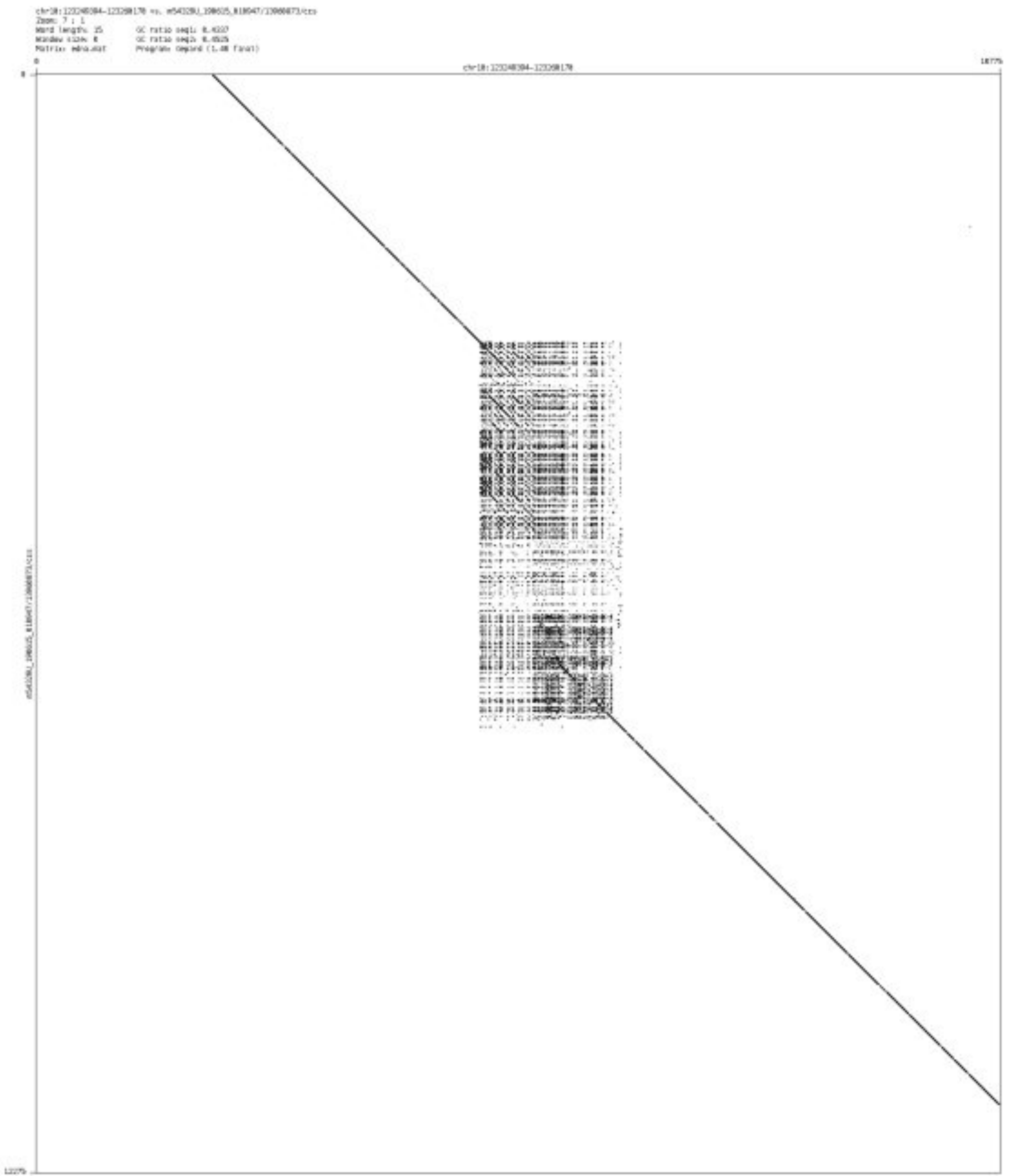

➢ chr10_132364374_132364814

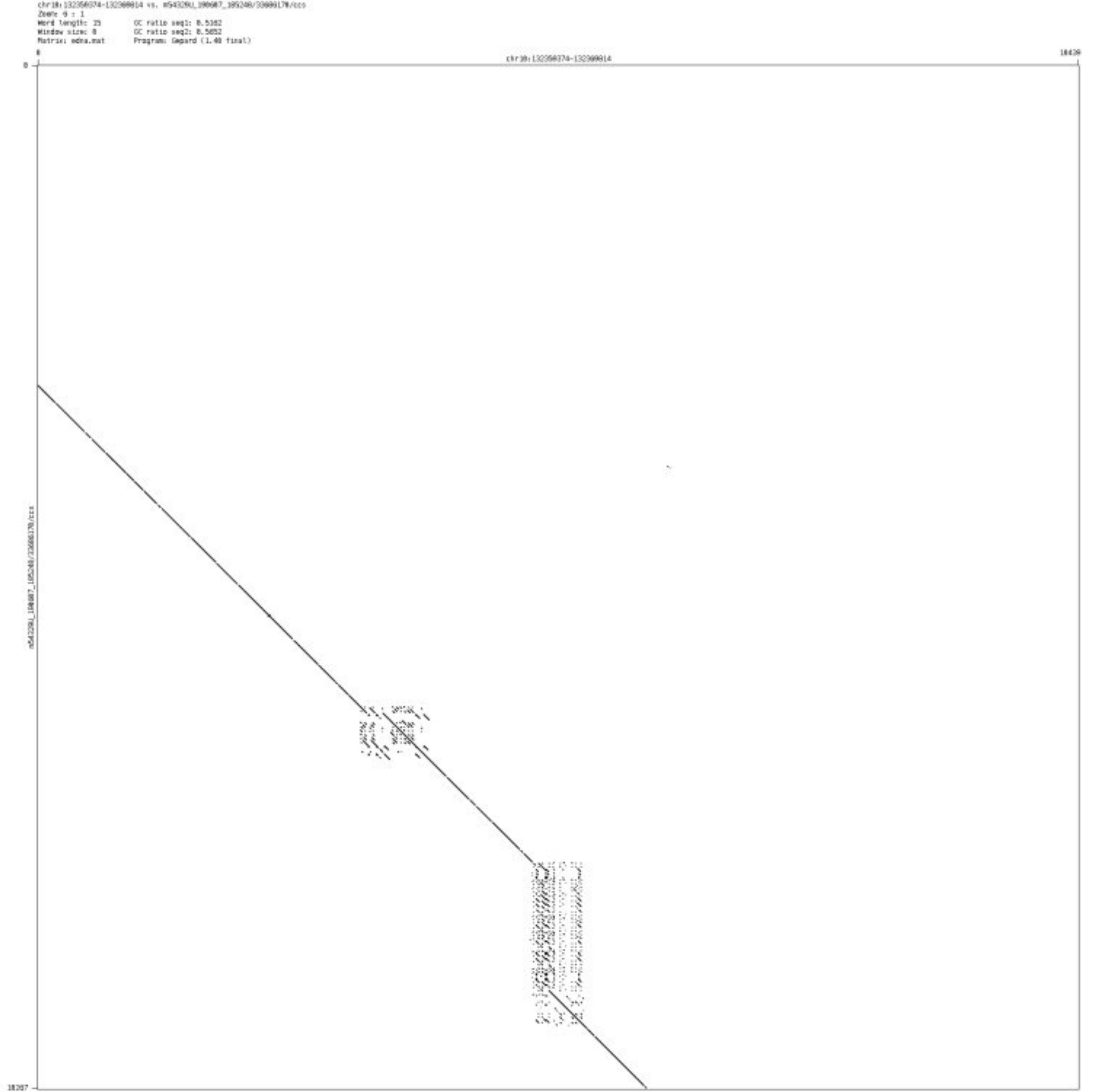

➢ chr10_133010210_133011617

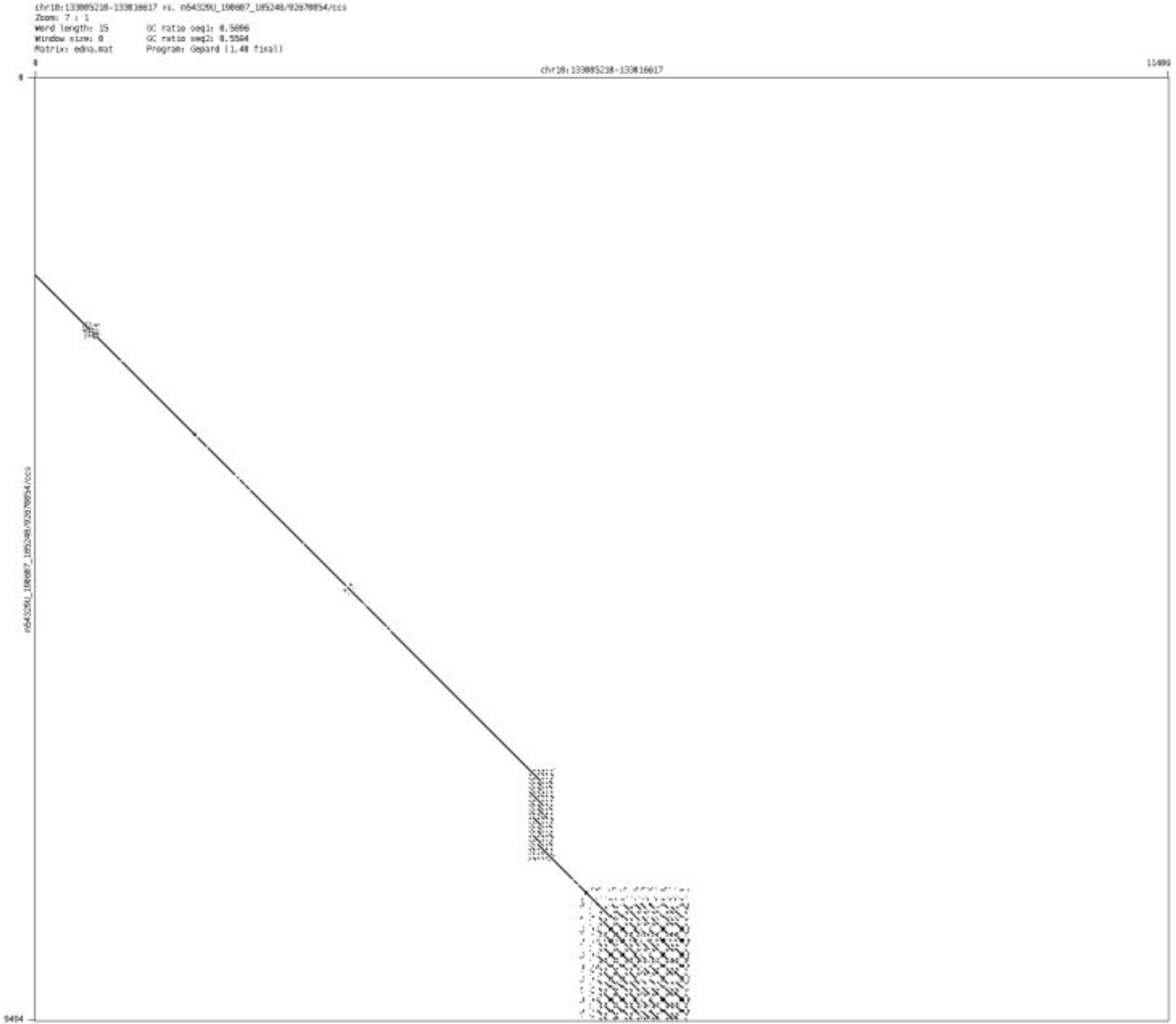

➢ chr10_133066182_133066276

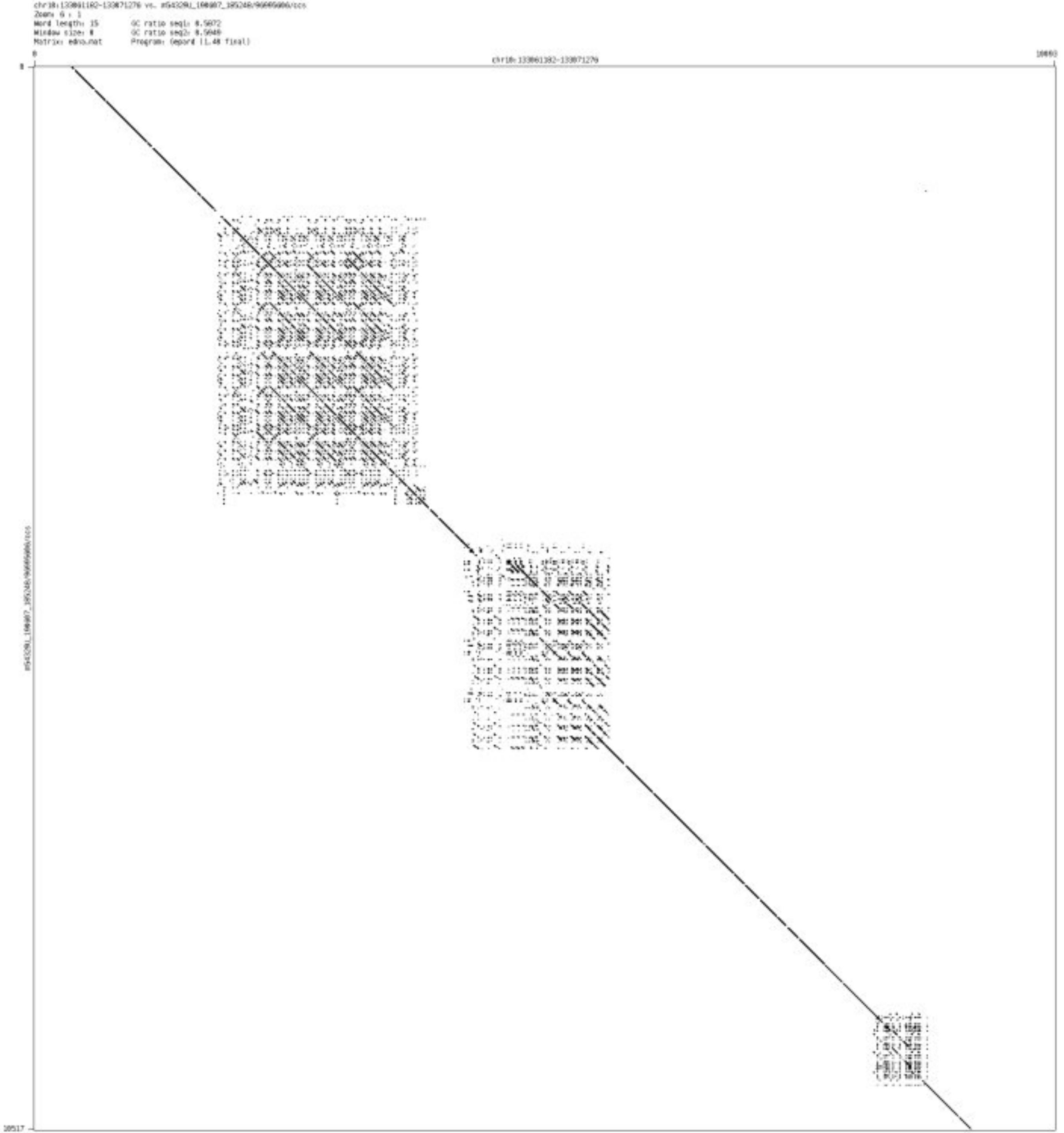

➢ chr11_71066026_71066104

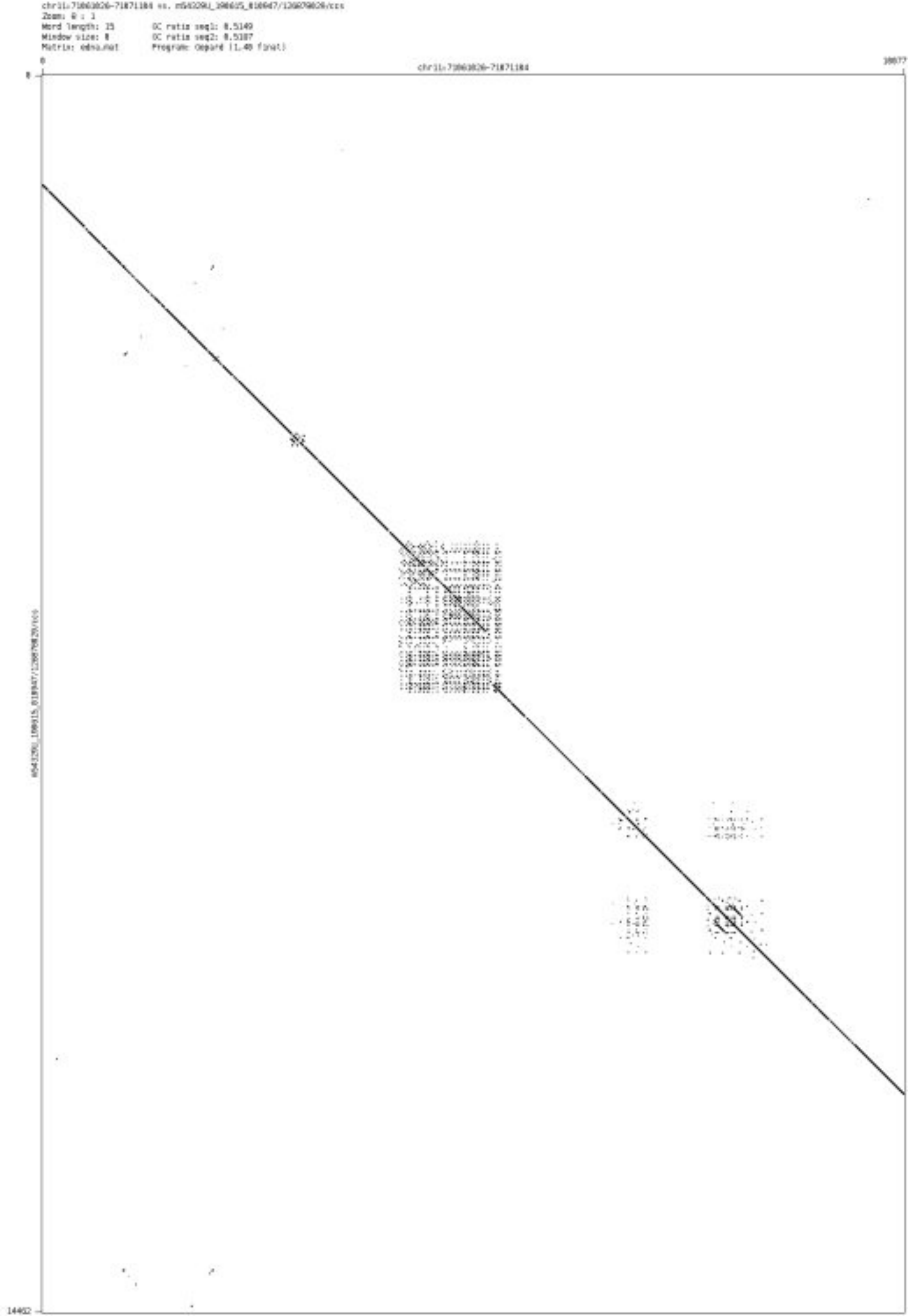

- chr11_102731115_102731273

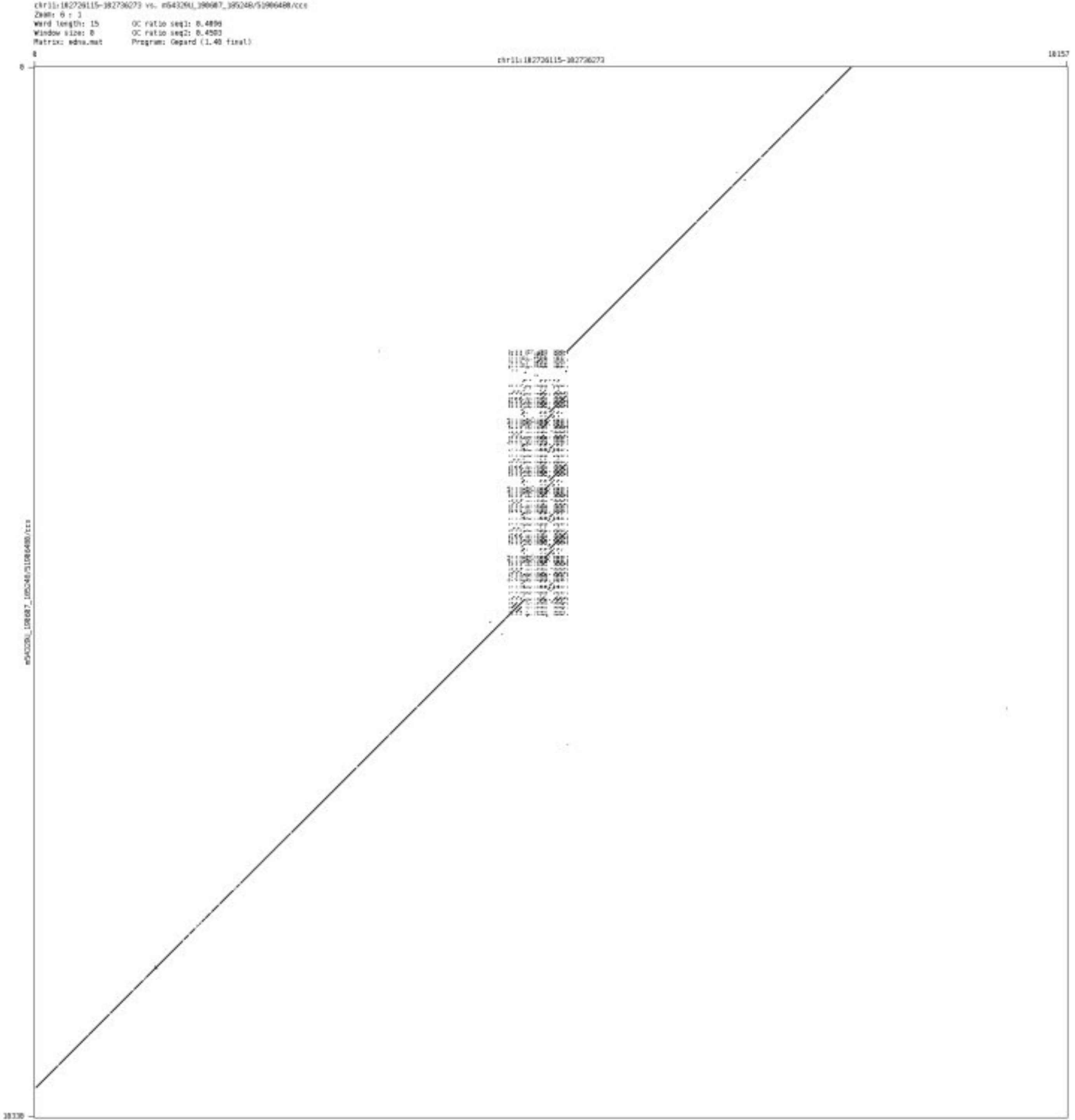

- chr11_131680439_131680801

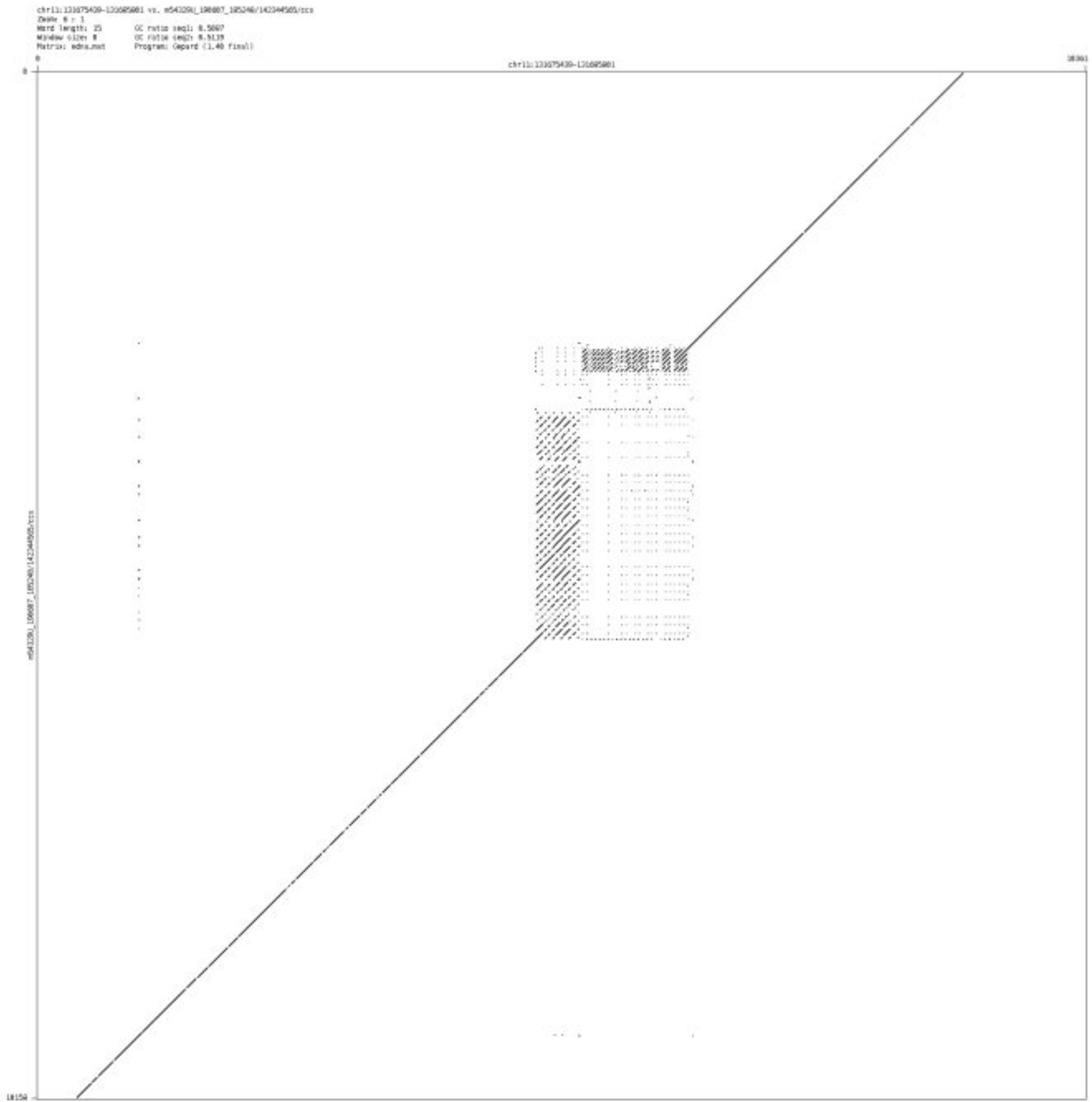

➢ chr11_133354203_133354263

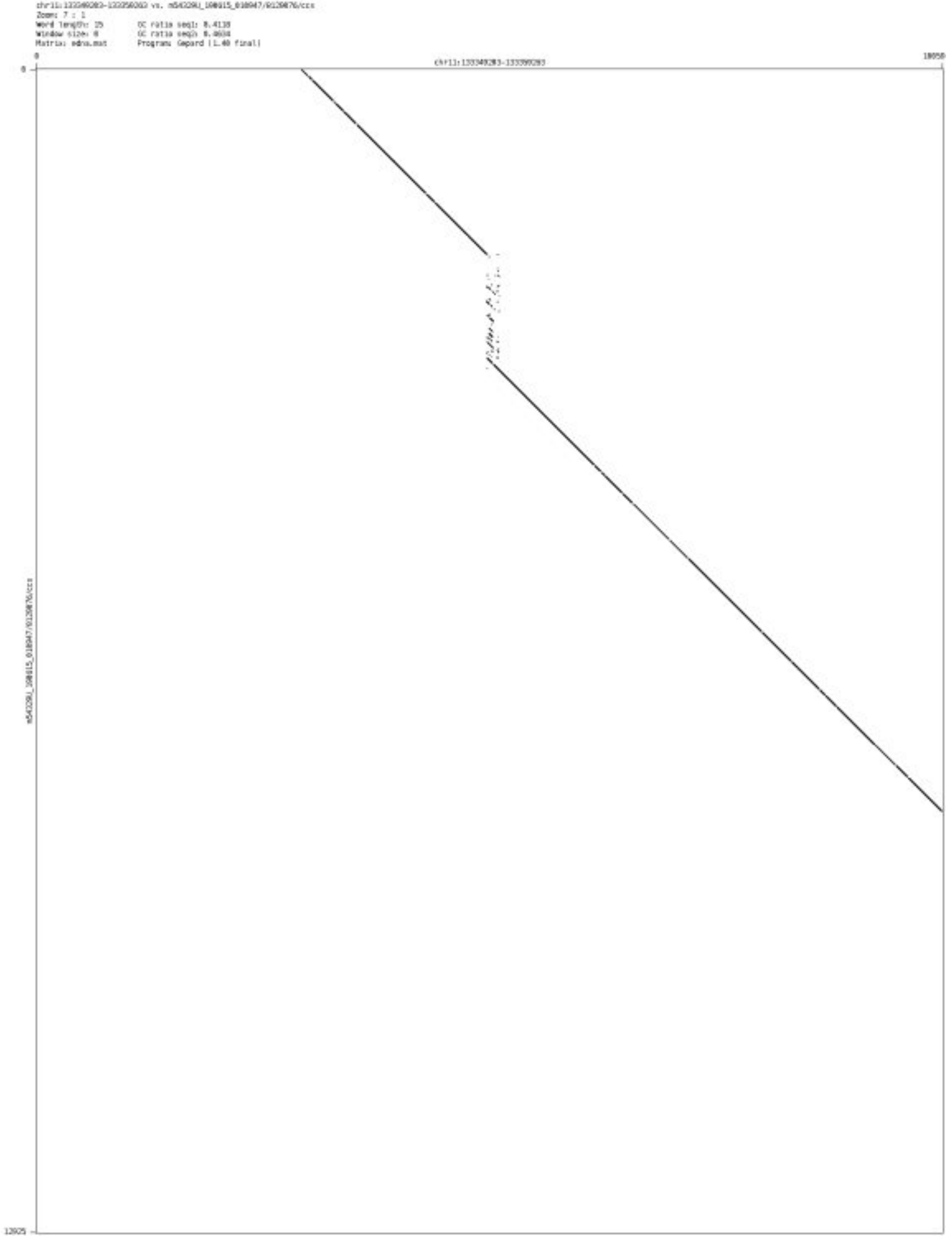

- ➢ chr12_127697075_127697231

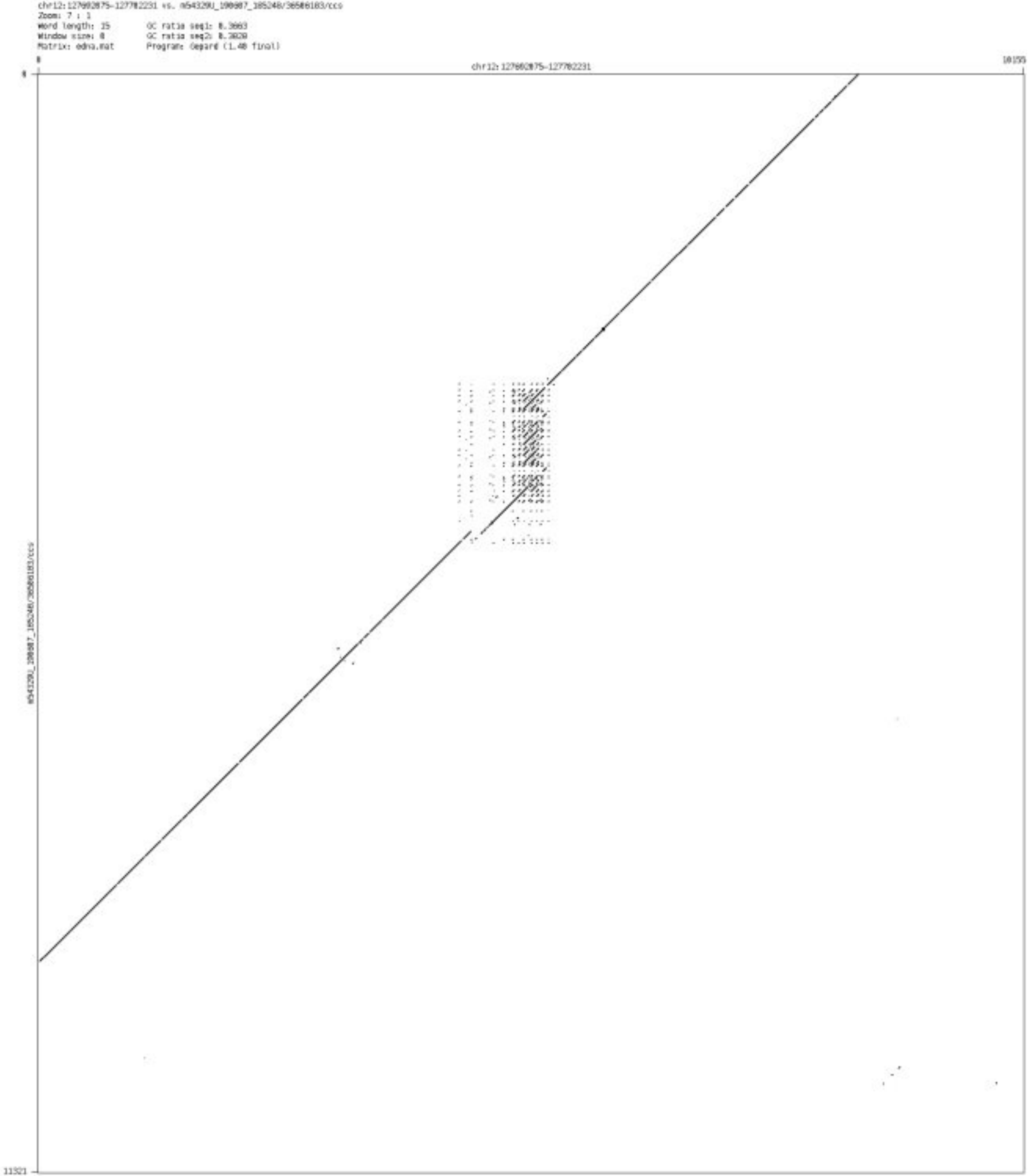

- ➢ chr12_132463913_132464505

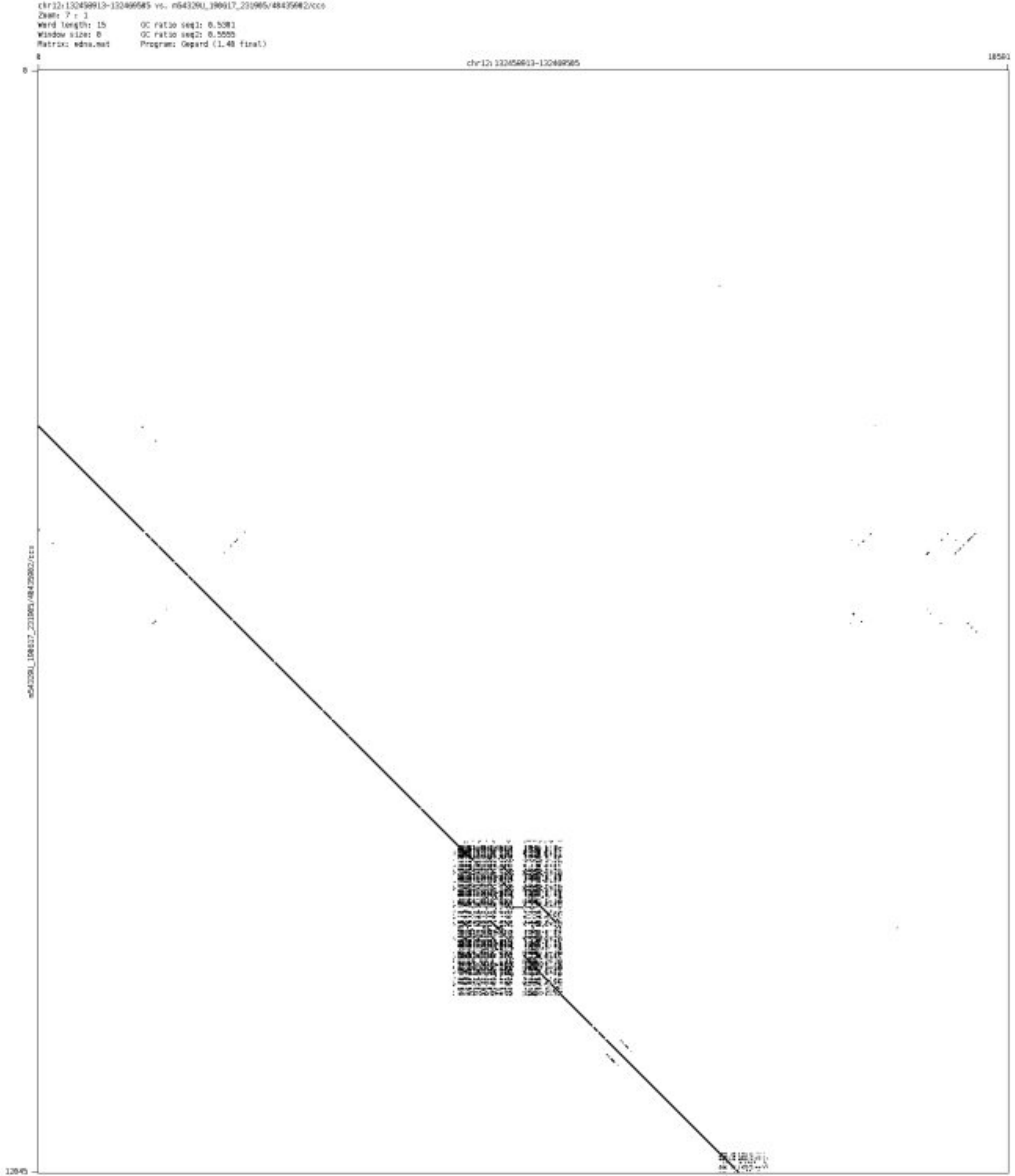

#### ➢ chr12_128708600_128708803

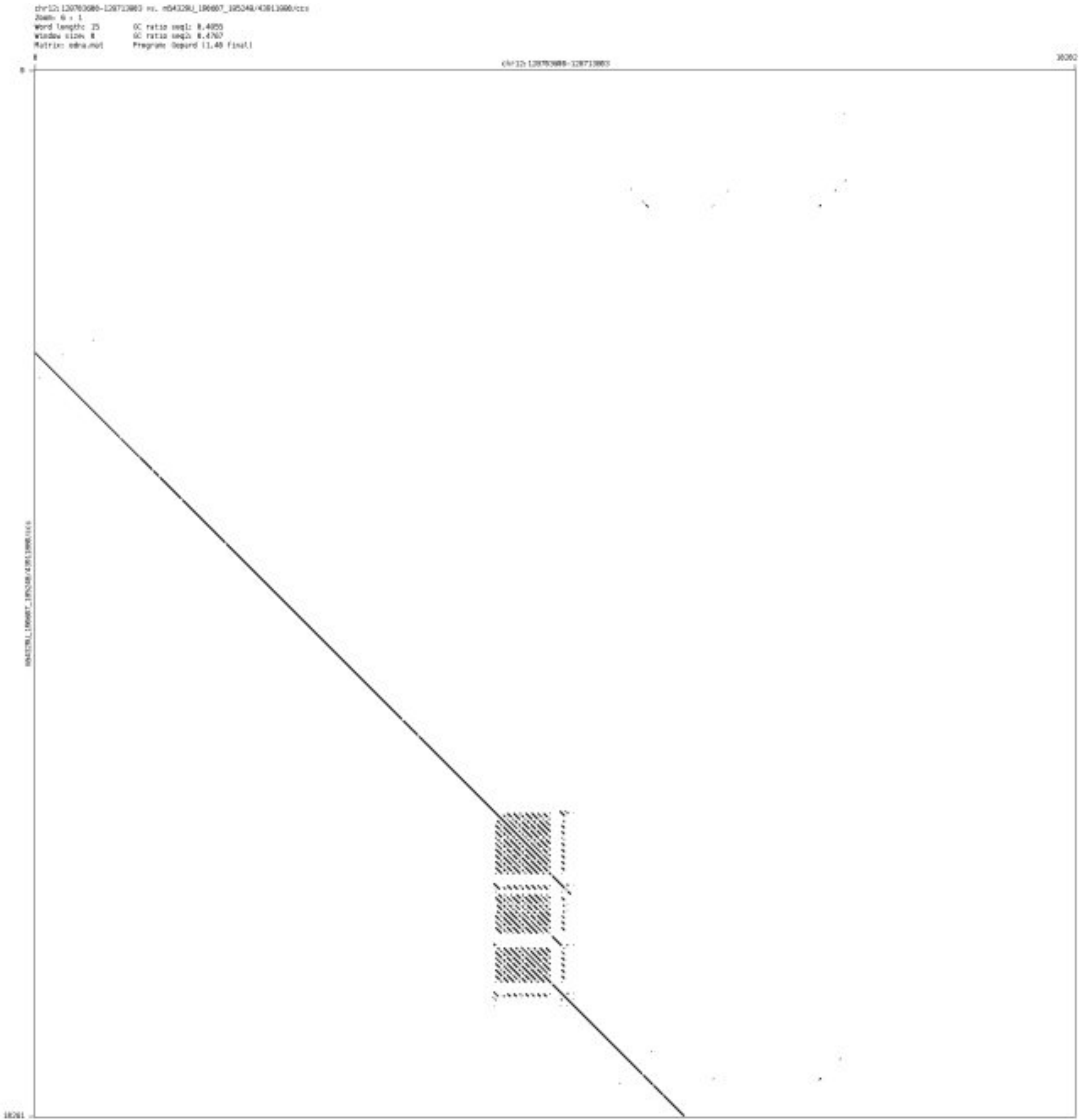

- chr13_113345937_113347871

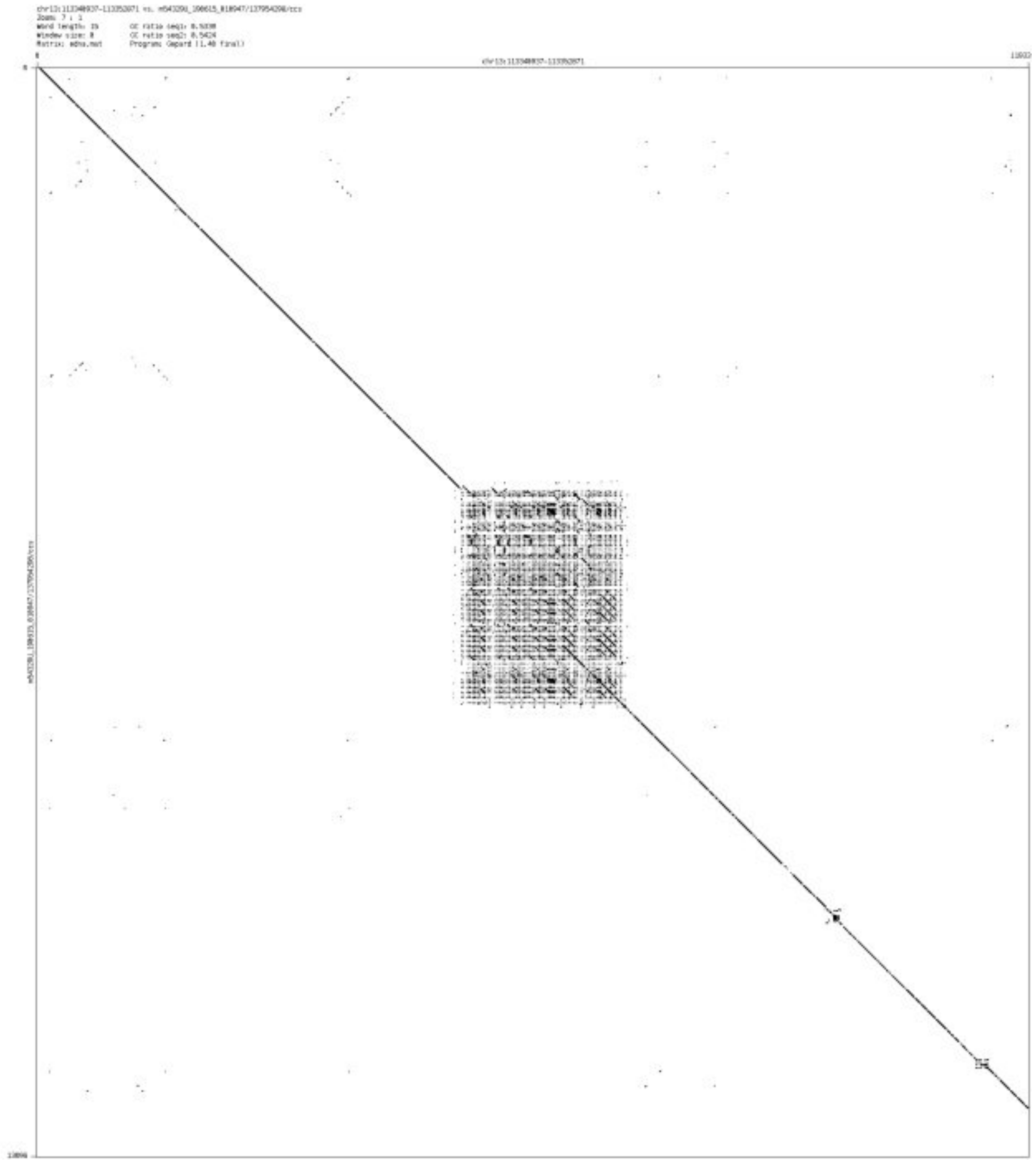

- chr13_113435956_113437202

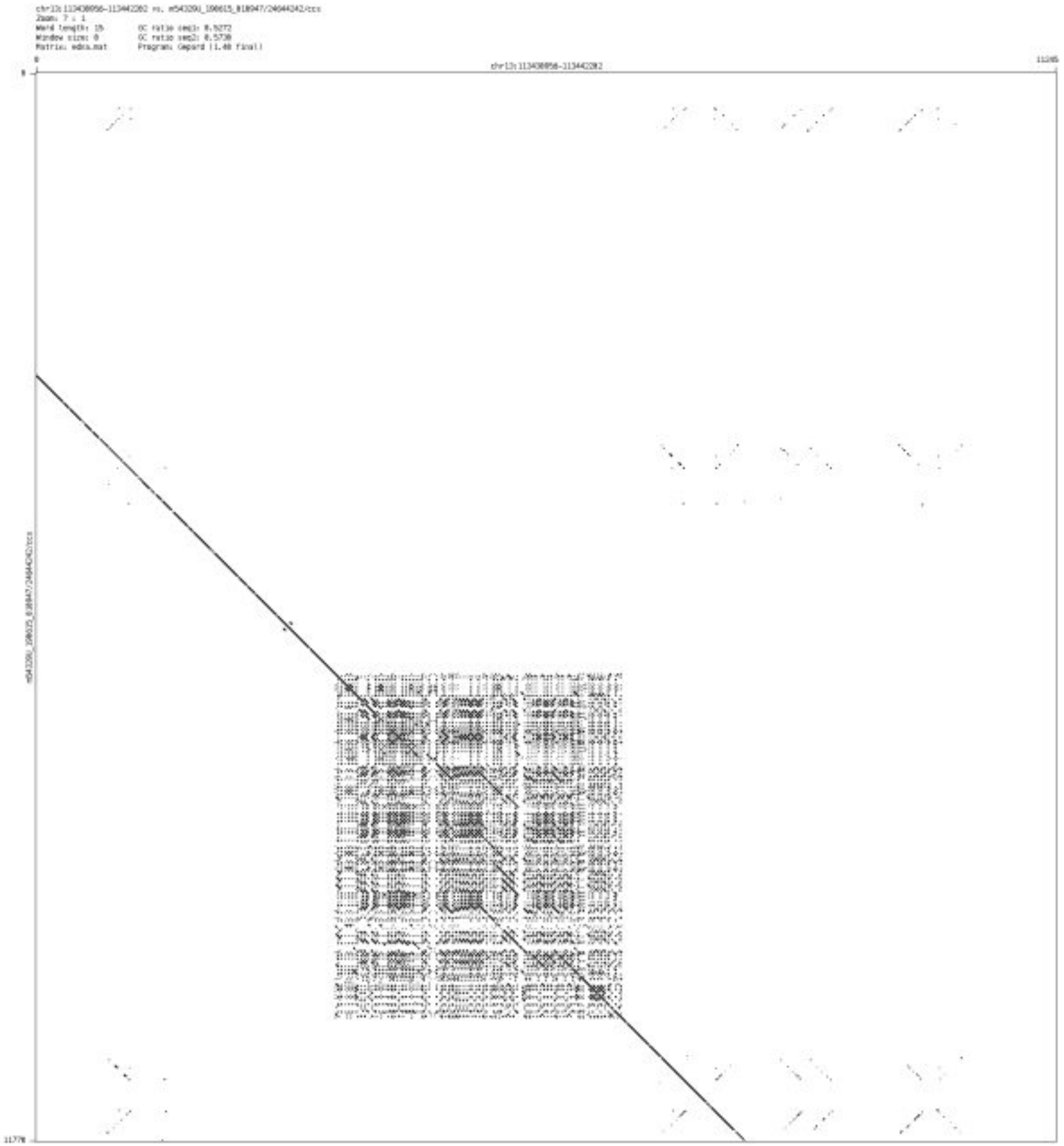

- chr14_100526791_100526935

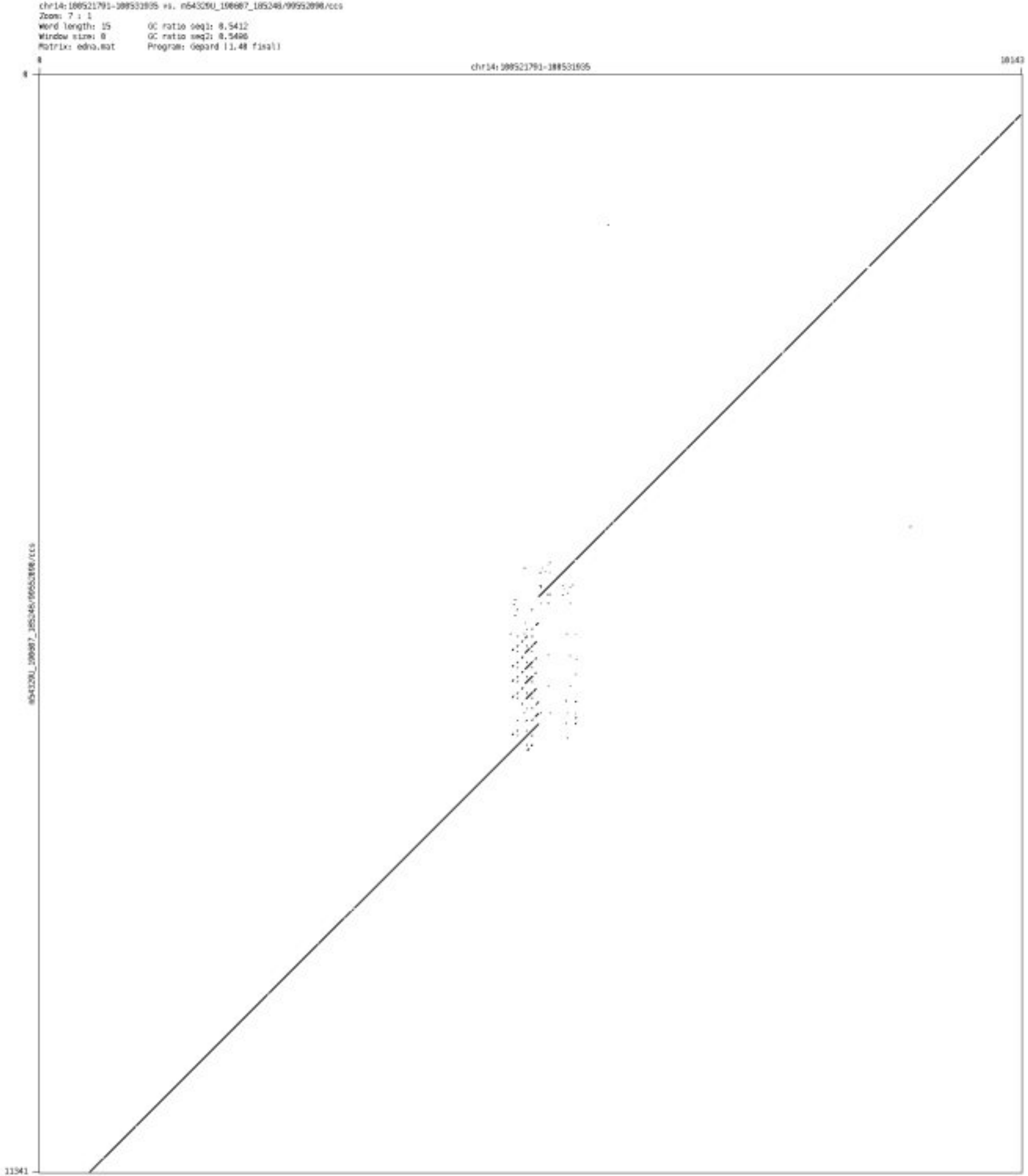

➢ chr14_104215330_104215464

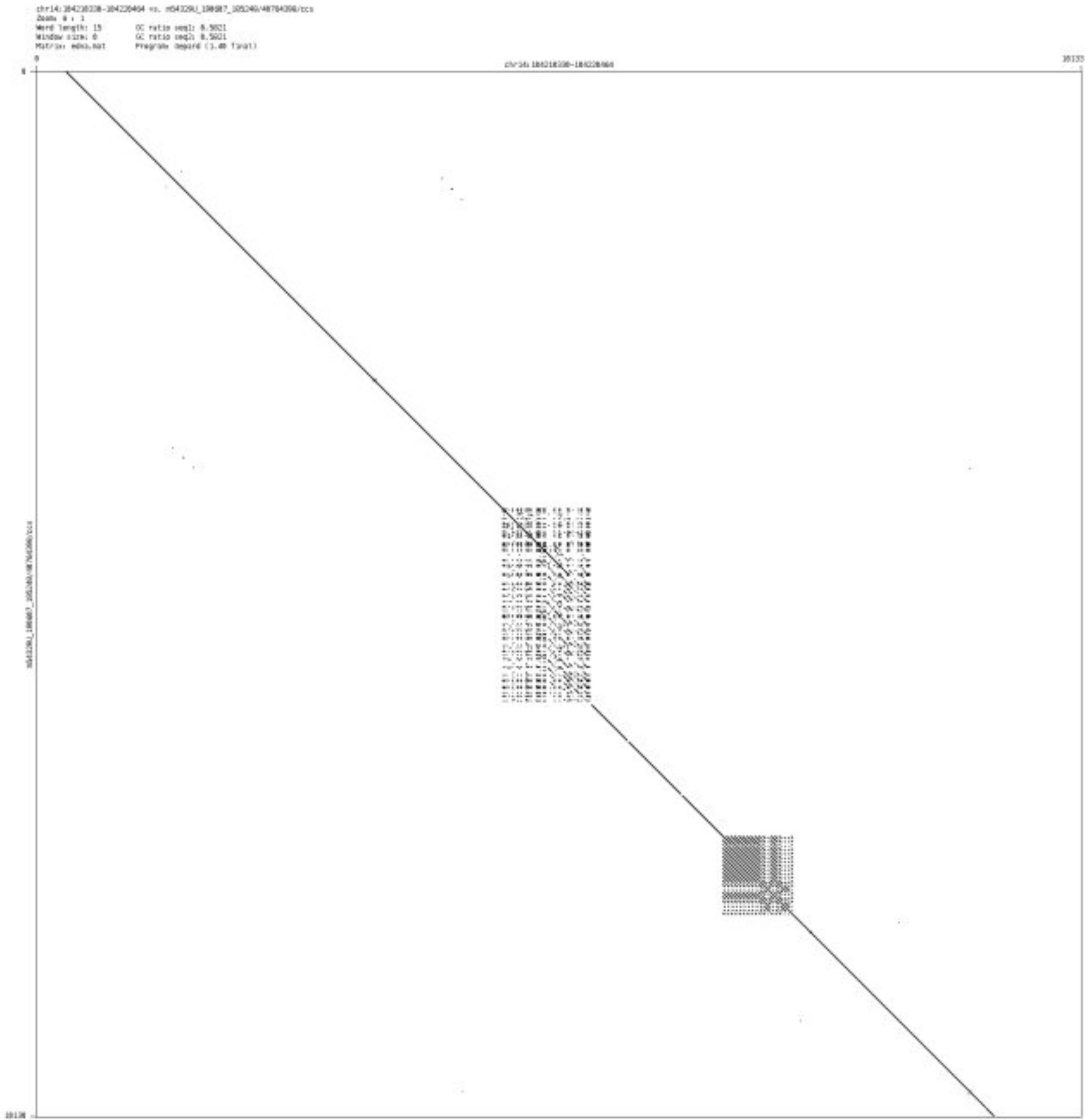

- chr15_71199689_71199869

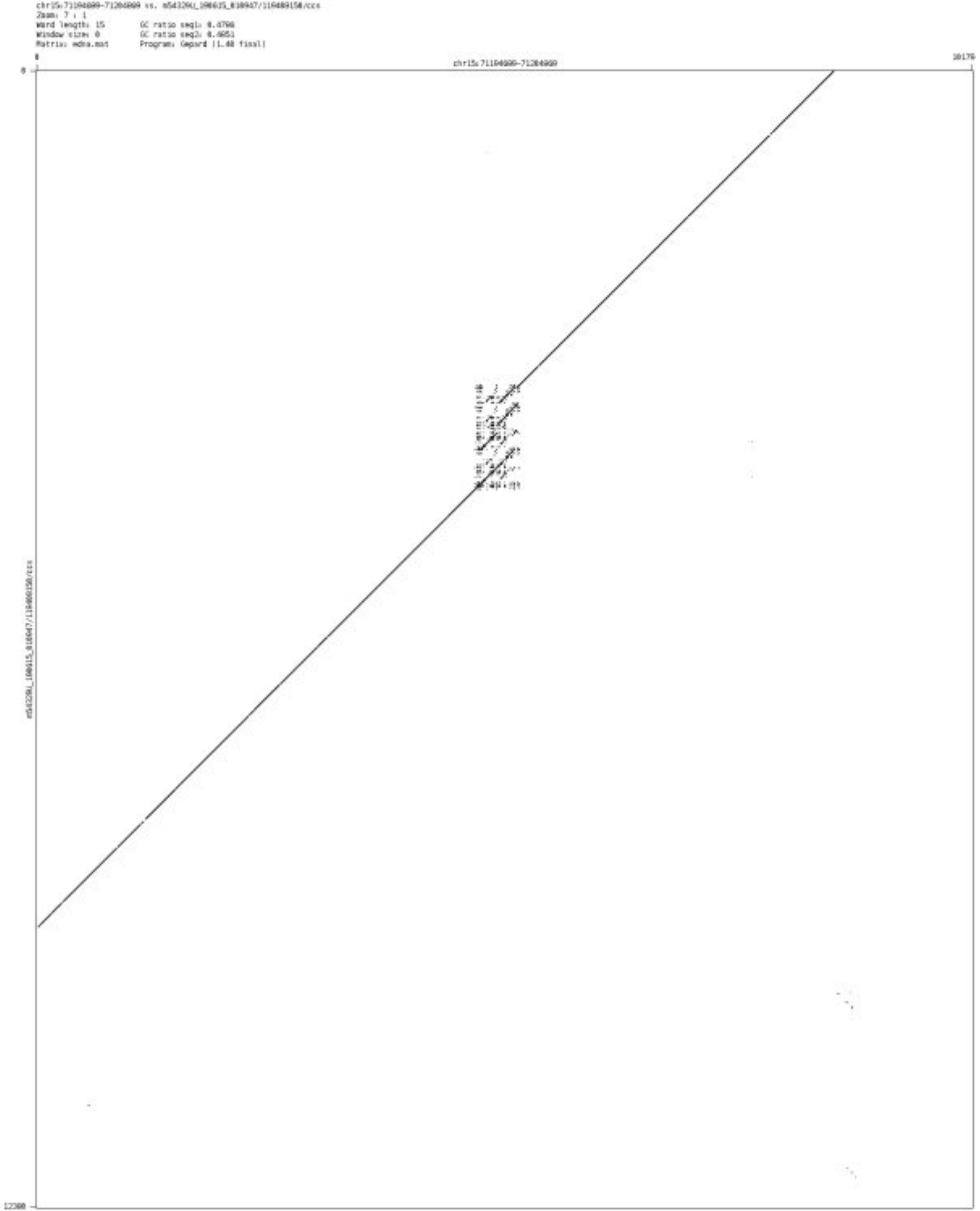

- chr16_23981983_23982098

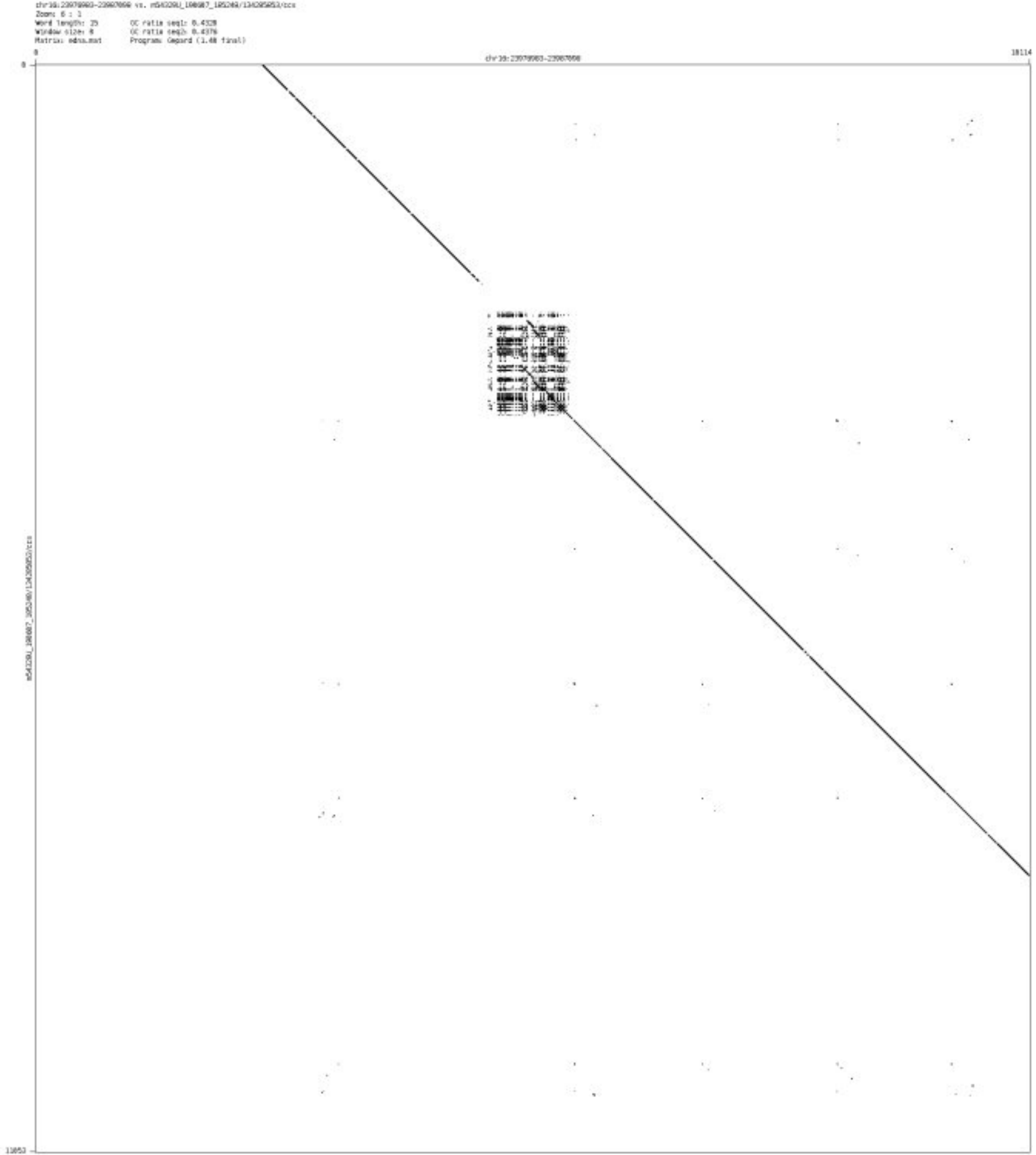

- chr16_27139102_27139213

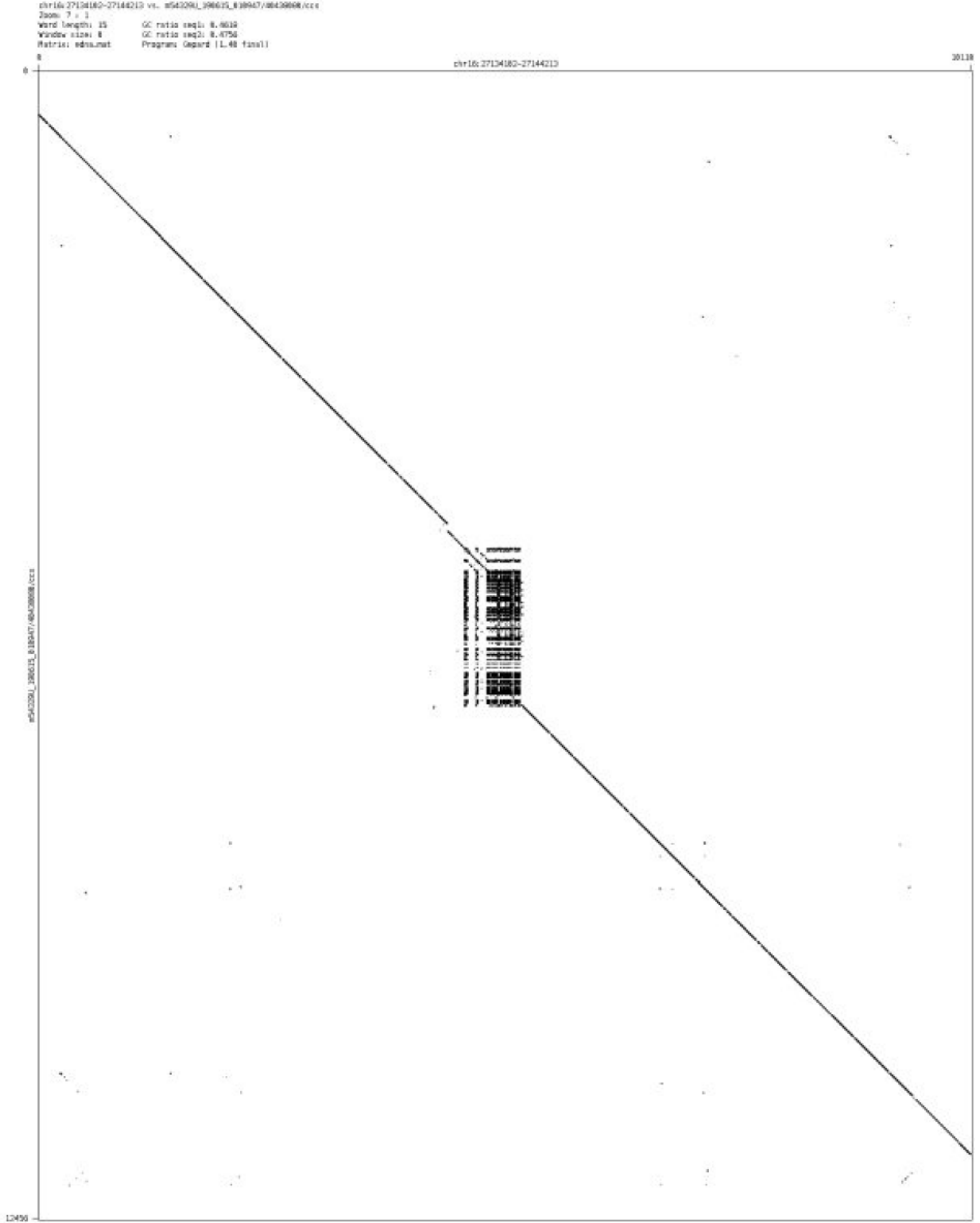

➢ chr17_314701_316802

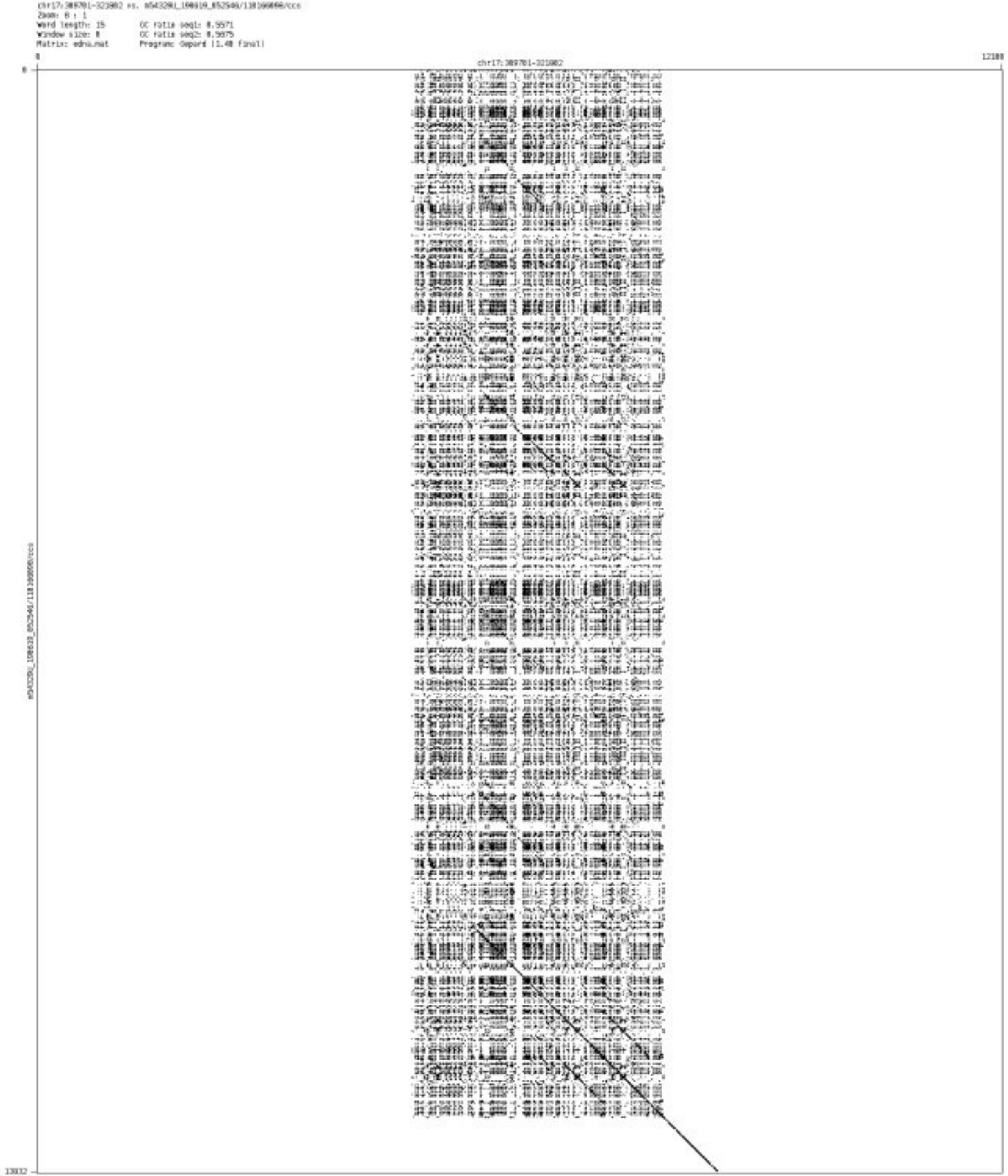

- chr17_41632760_41633721

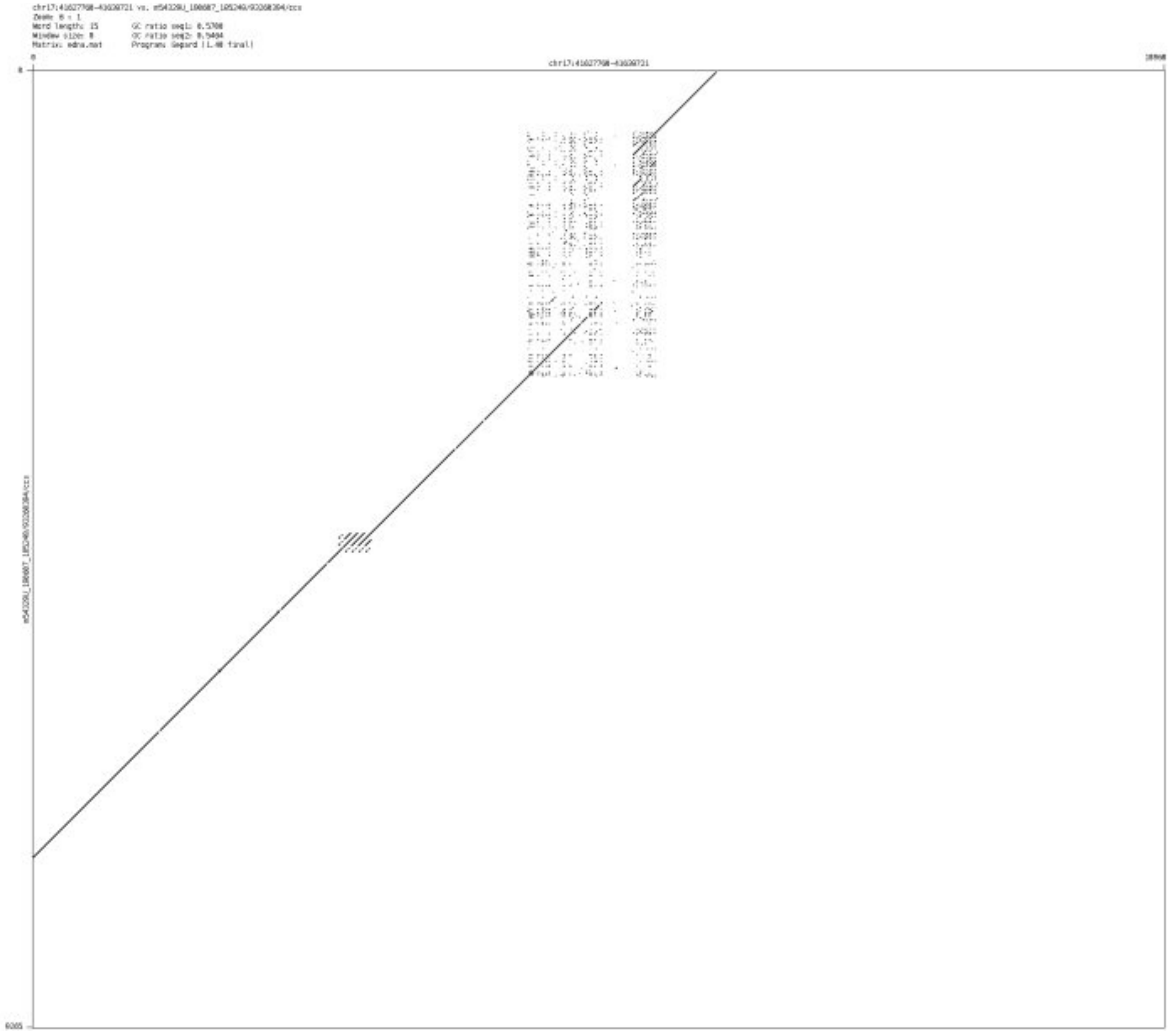

➢ chr18_13262154_13262356

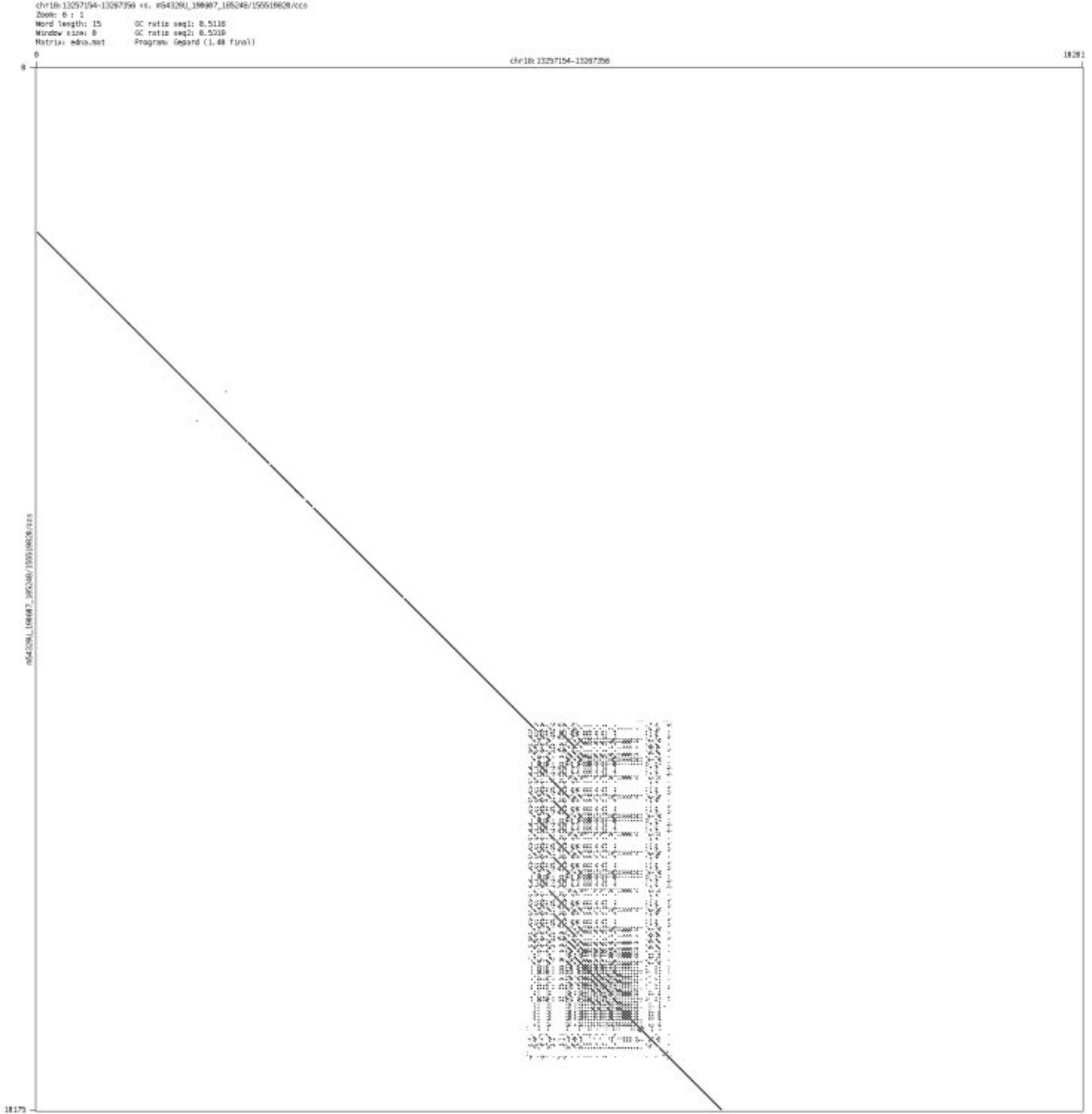

➢ chr18_76967411_76967555

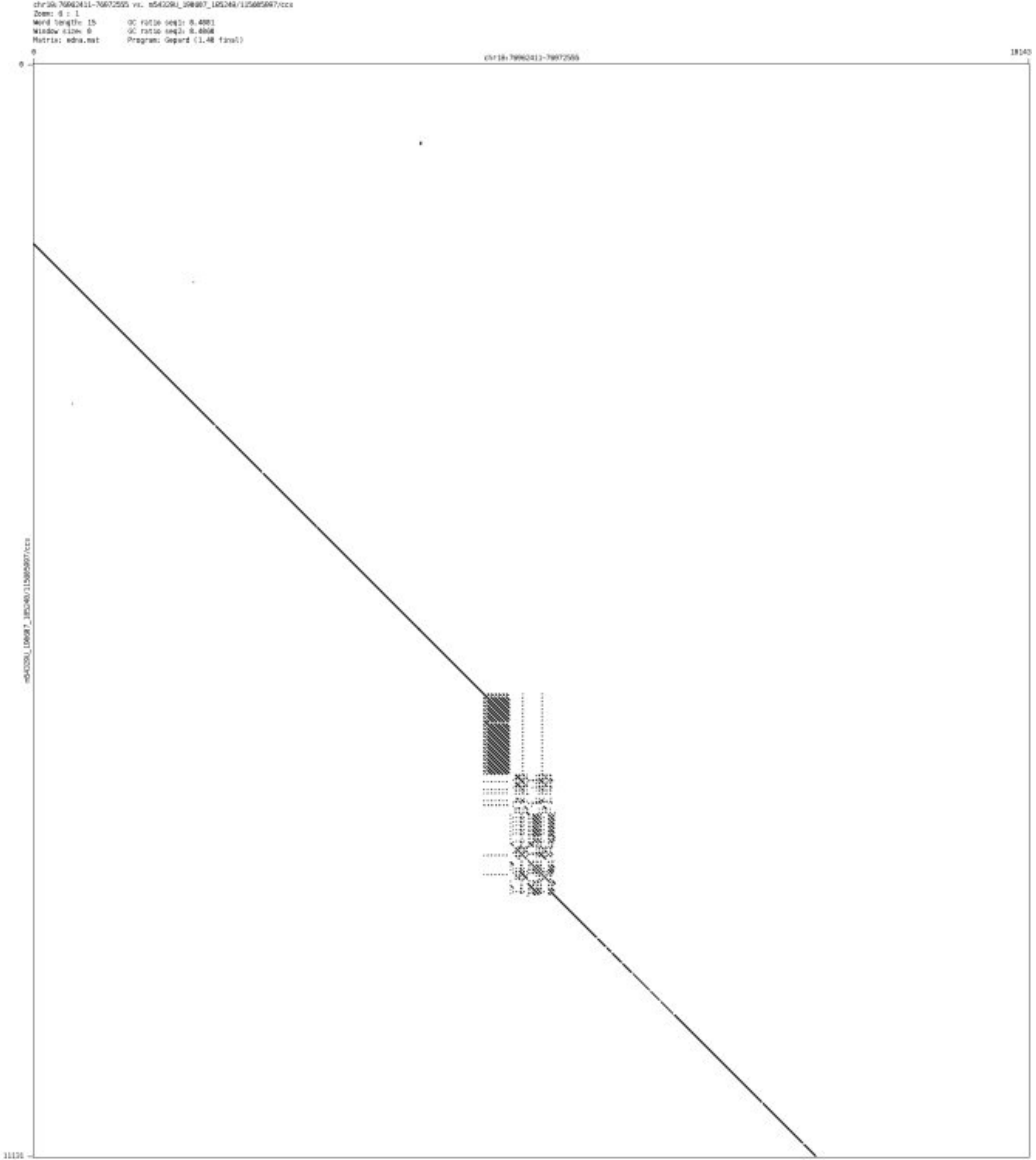

➢ chr19_879672_879791

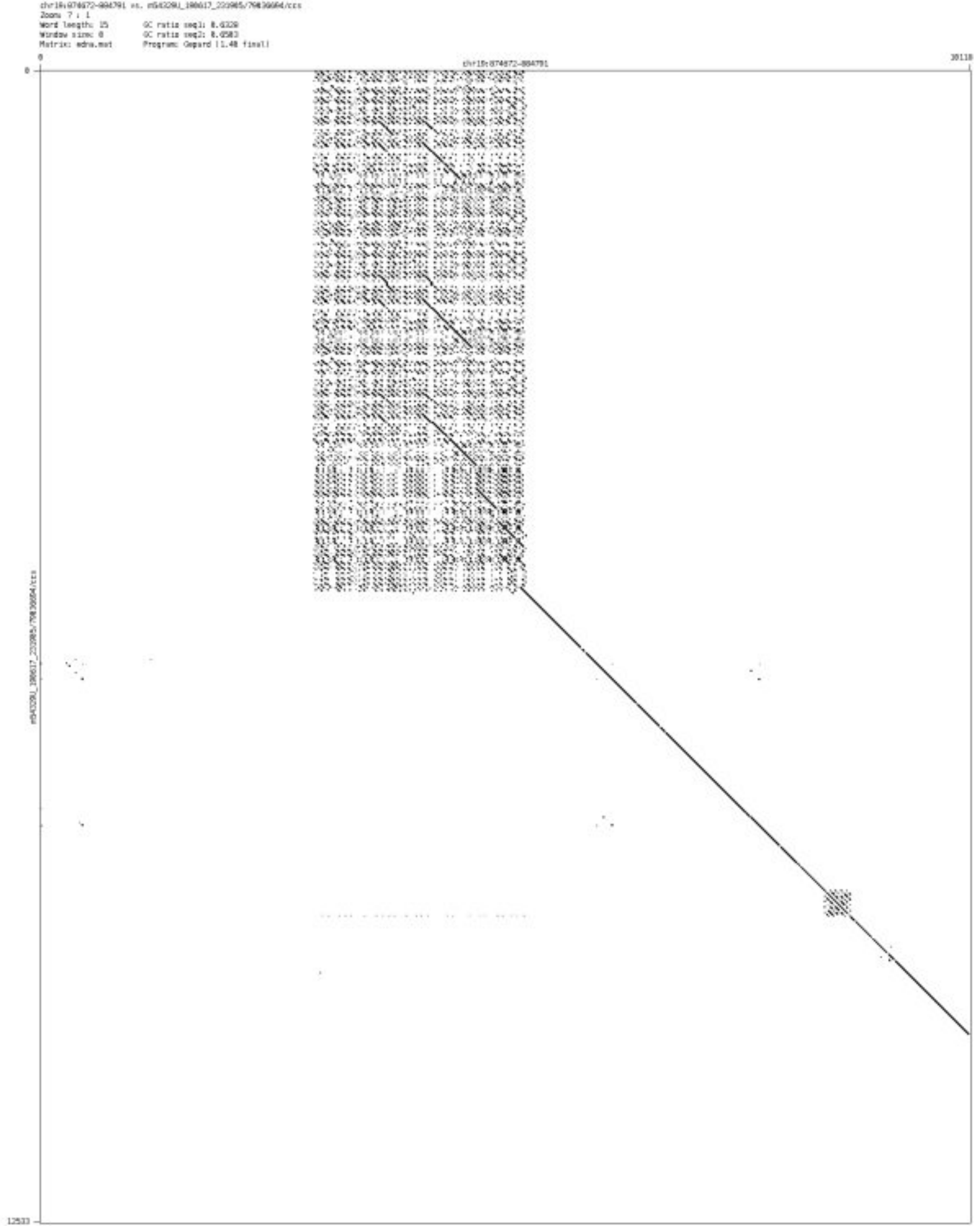

➢ chr19_4035778_4035935

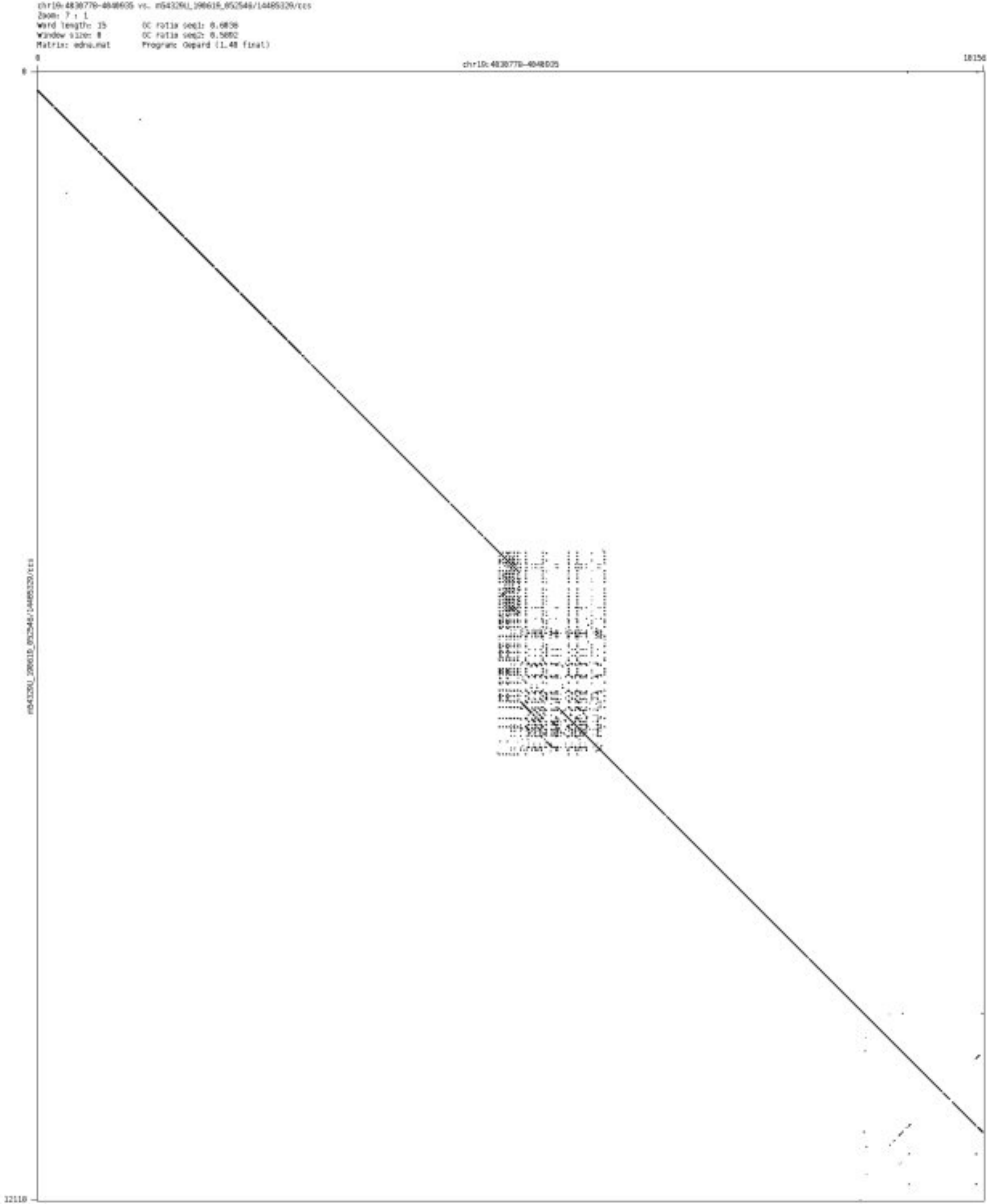

- ➢ chr19_7153130_7153347

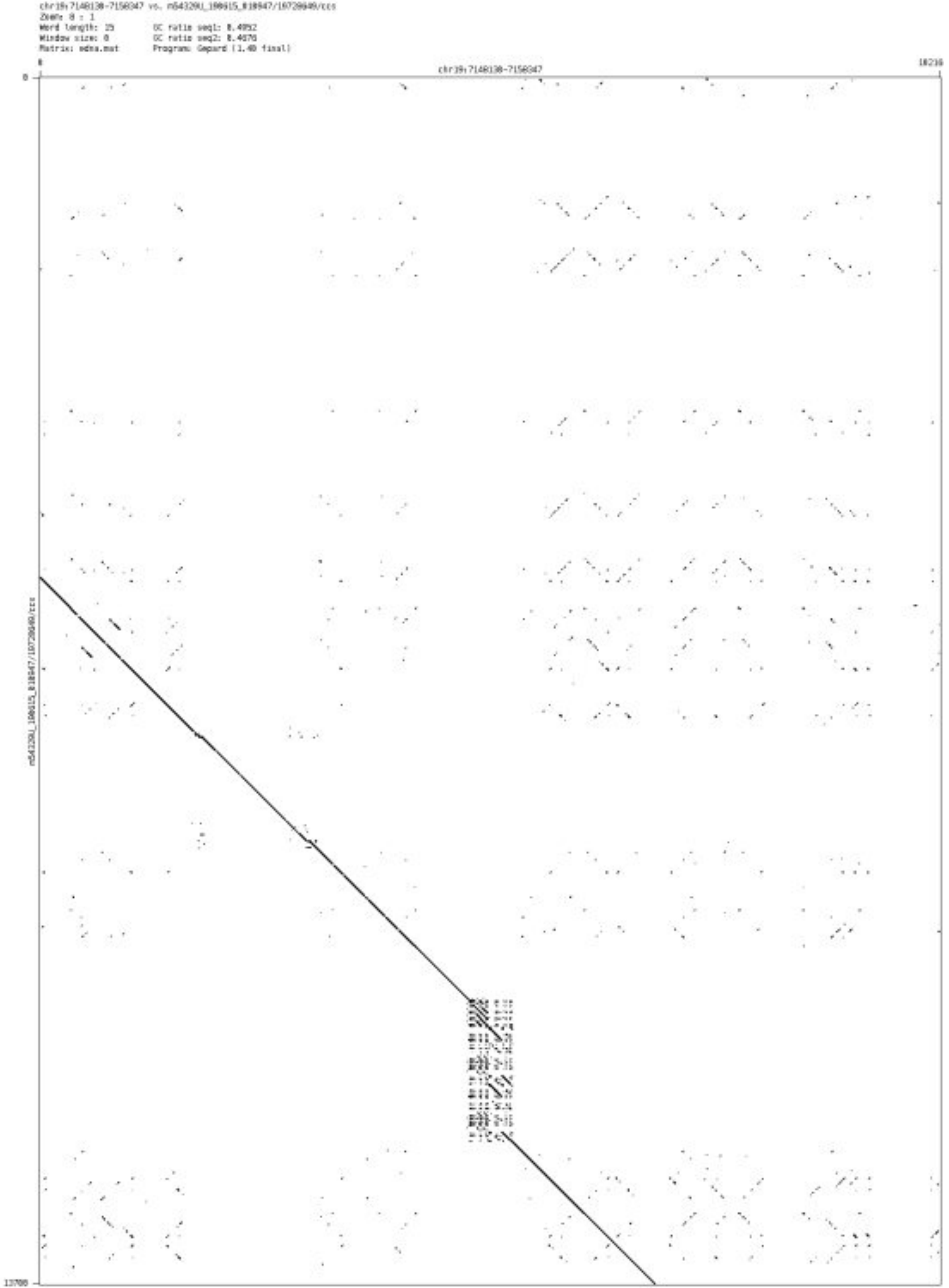

- chr19_7331323_7331681

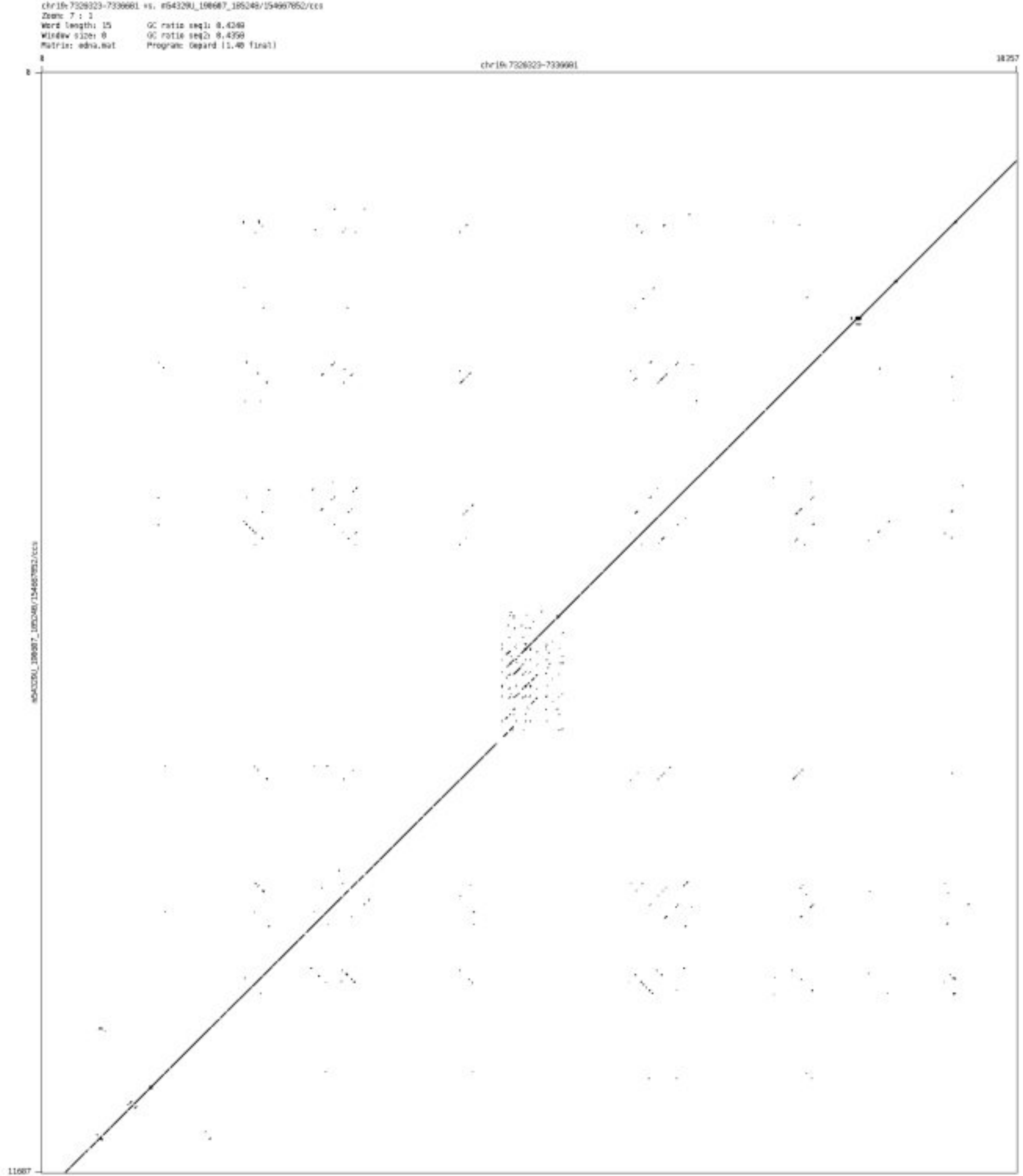

➢ chr19_12989496_12989651

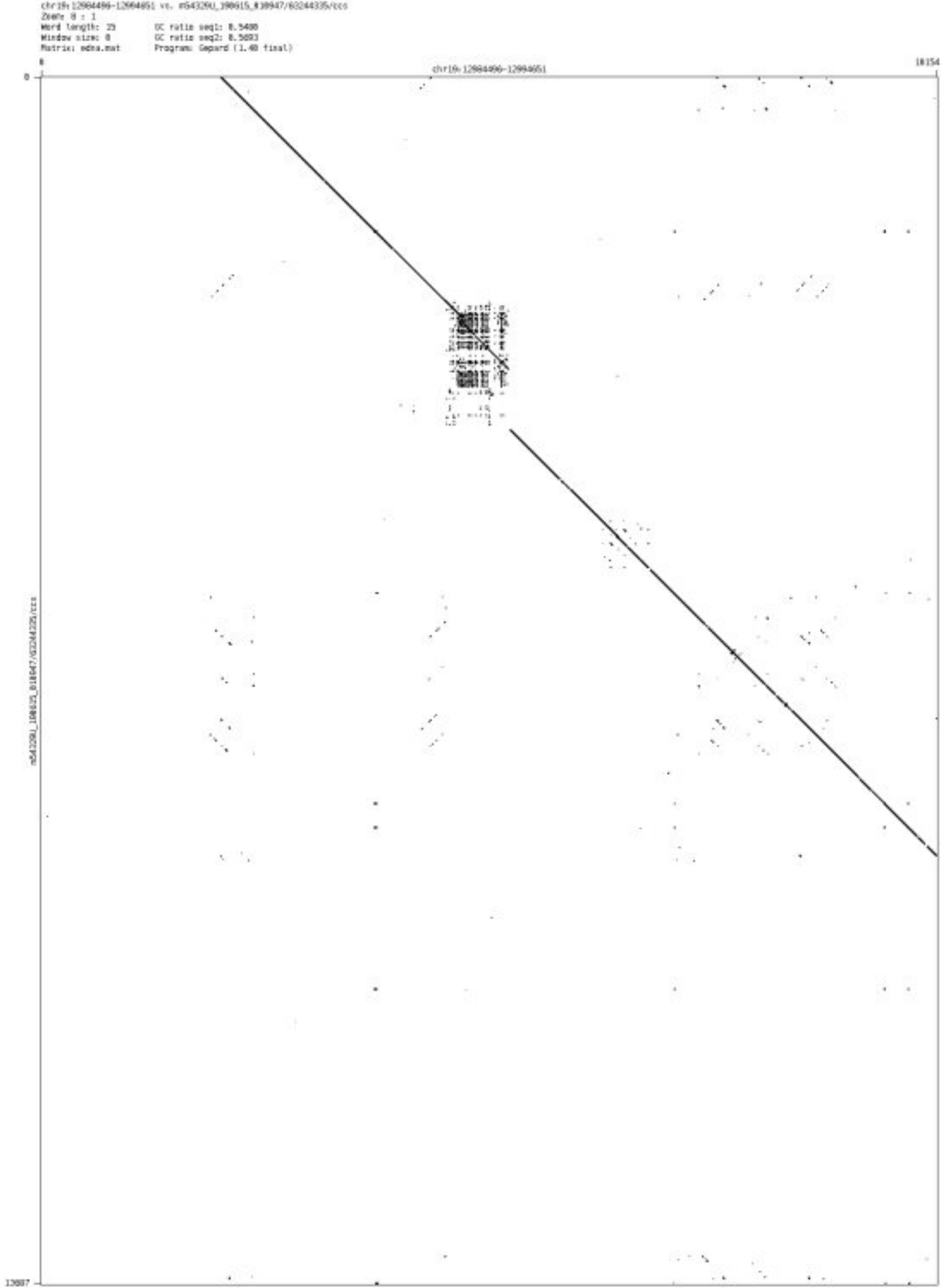

- chr20_60314368_60314789

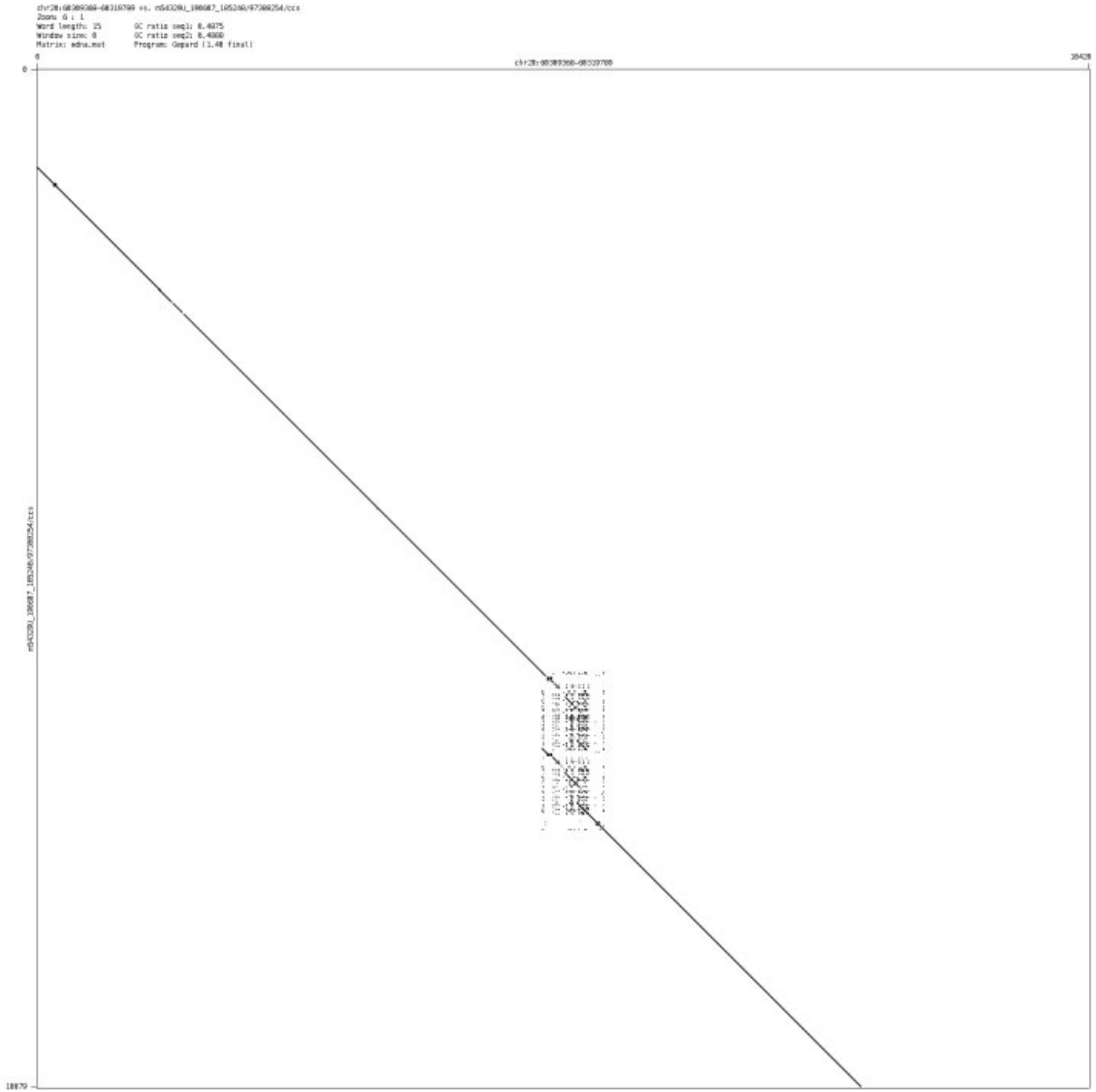

➢ chr21_33322245_33322788

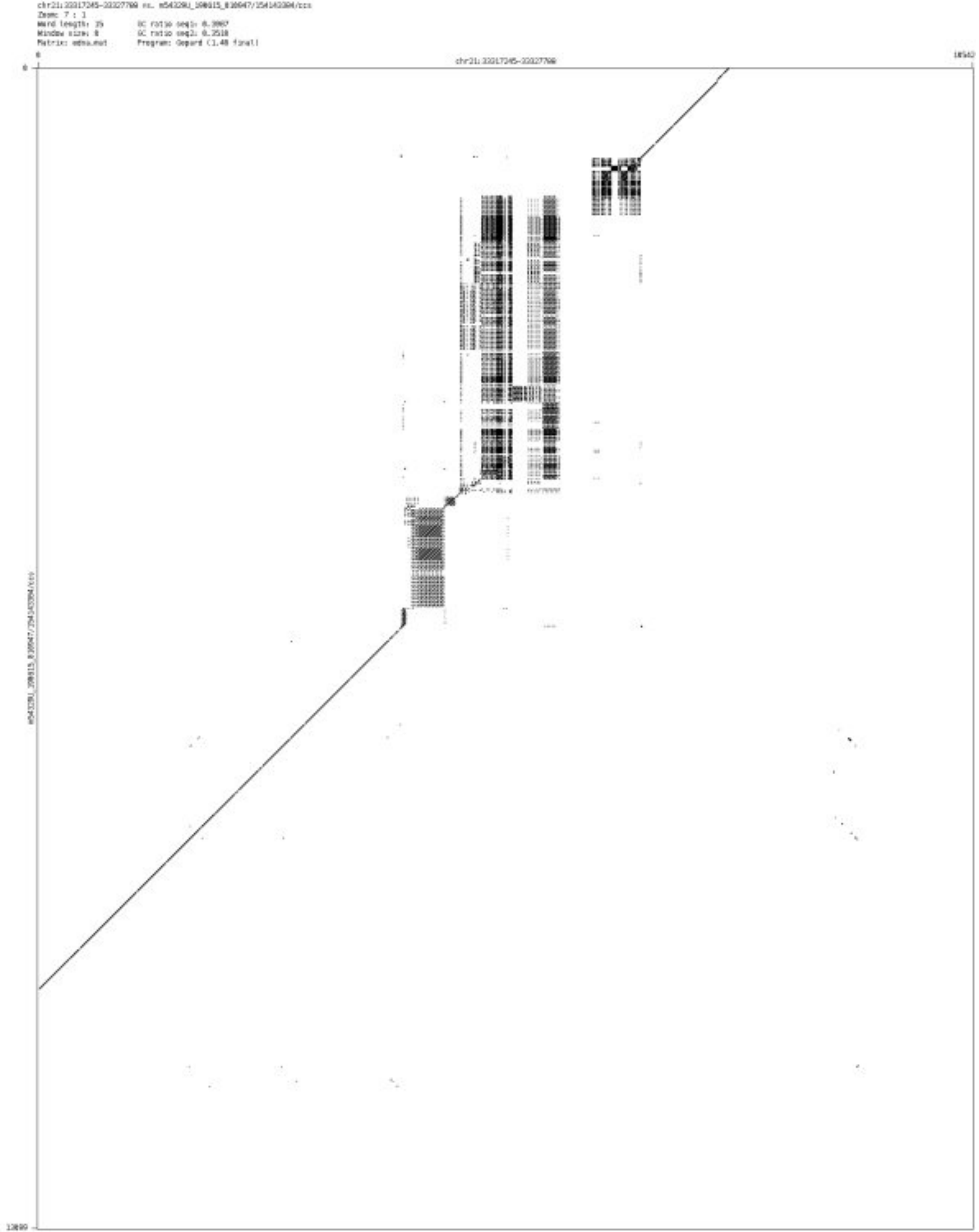

- chr21_46090975_46091314

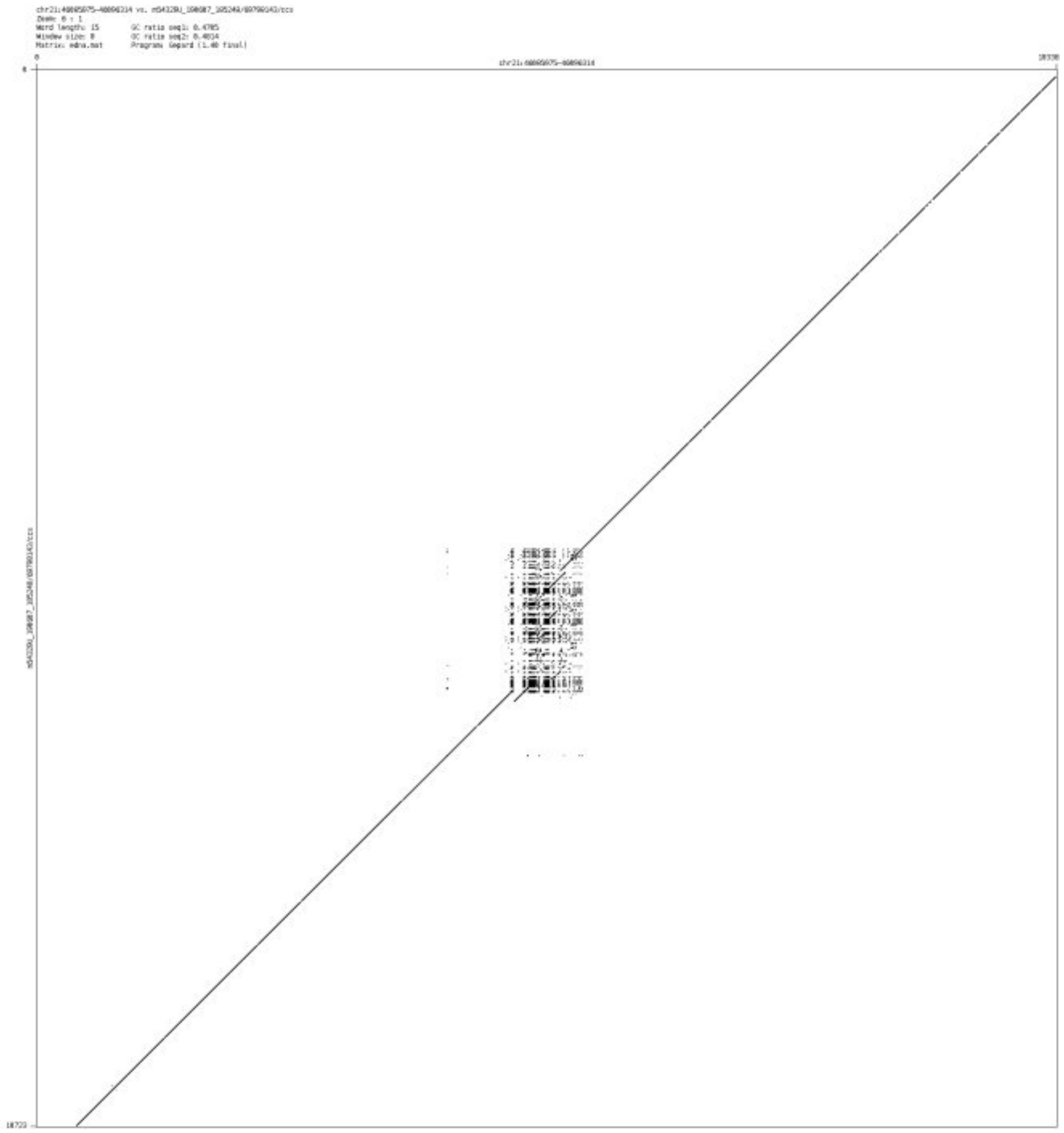

➢ chr22_18106125_18106340

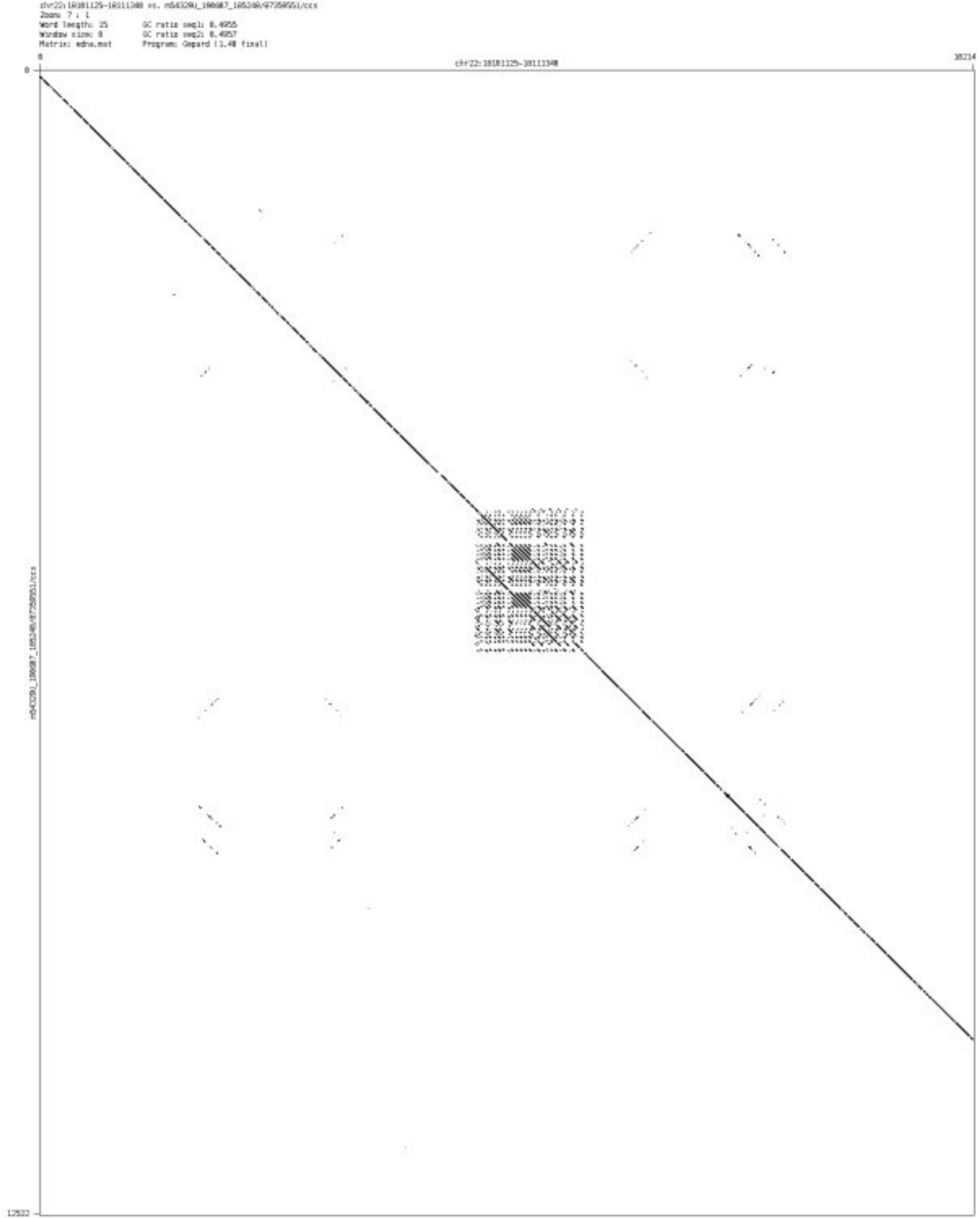

#### ➢ chr22_19202401_19202471

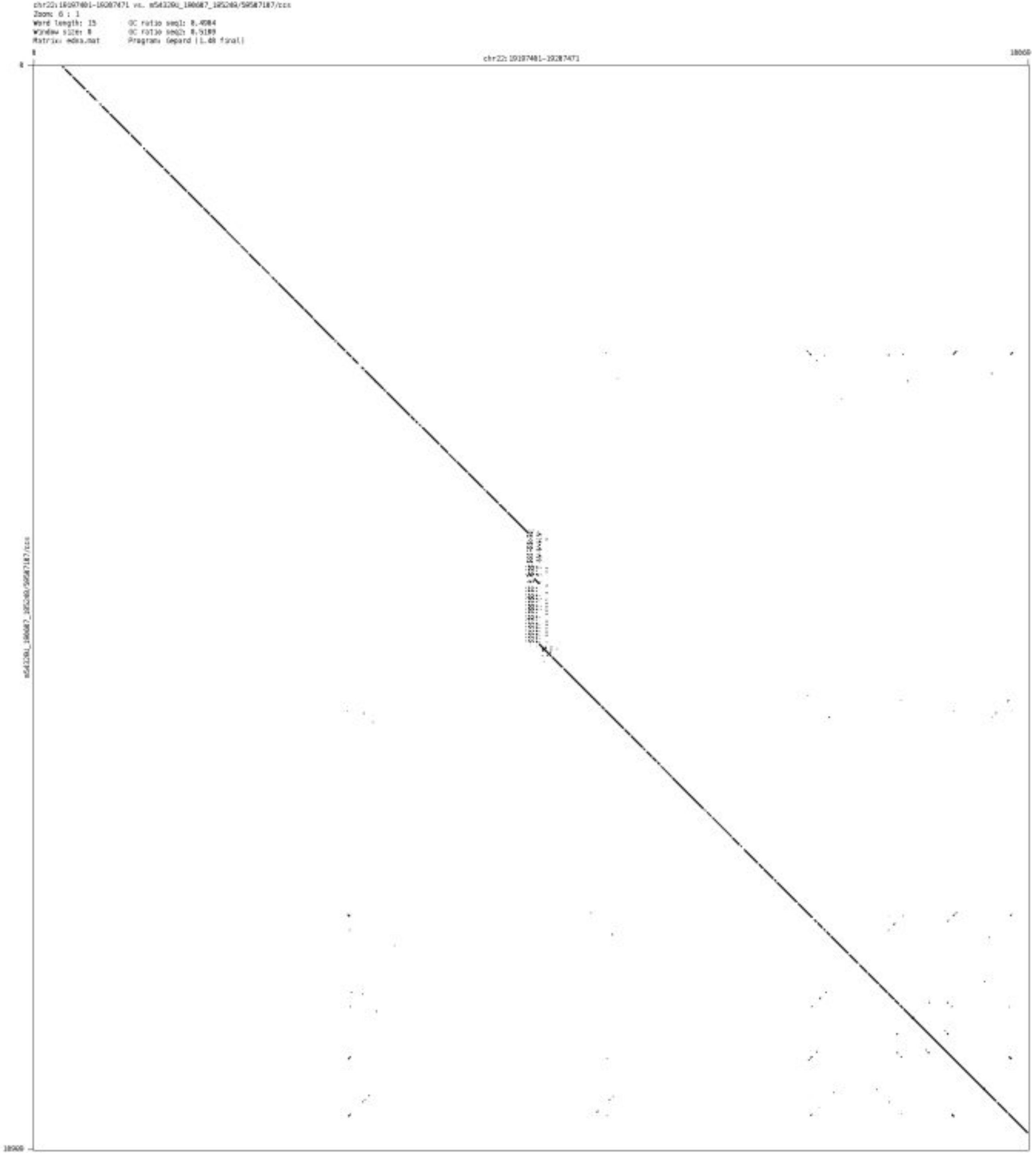

- chr22_48620822_48620943

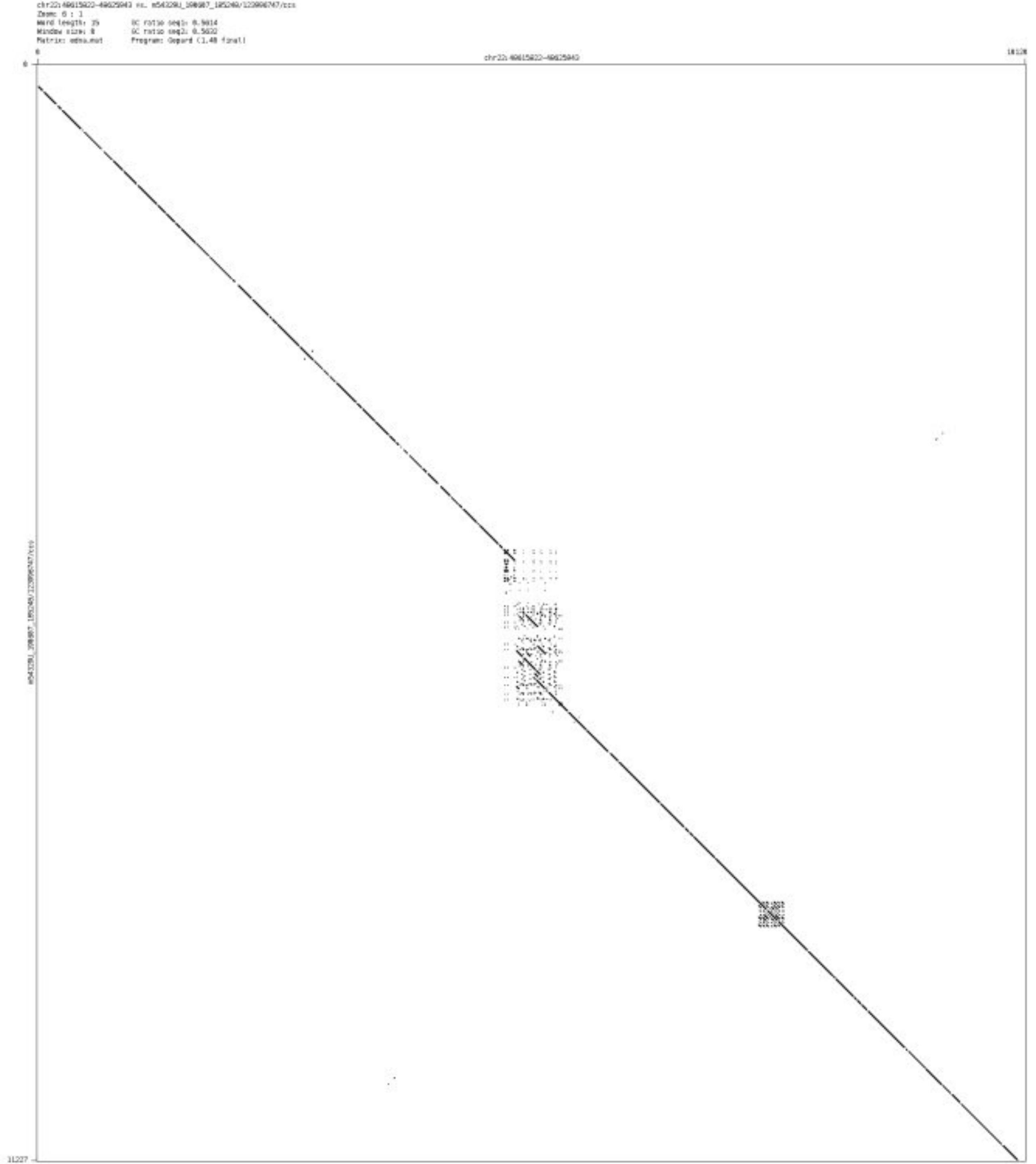

### ’Messy’ CSV-Case C

- chr3_50311835_50312092

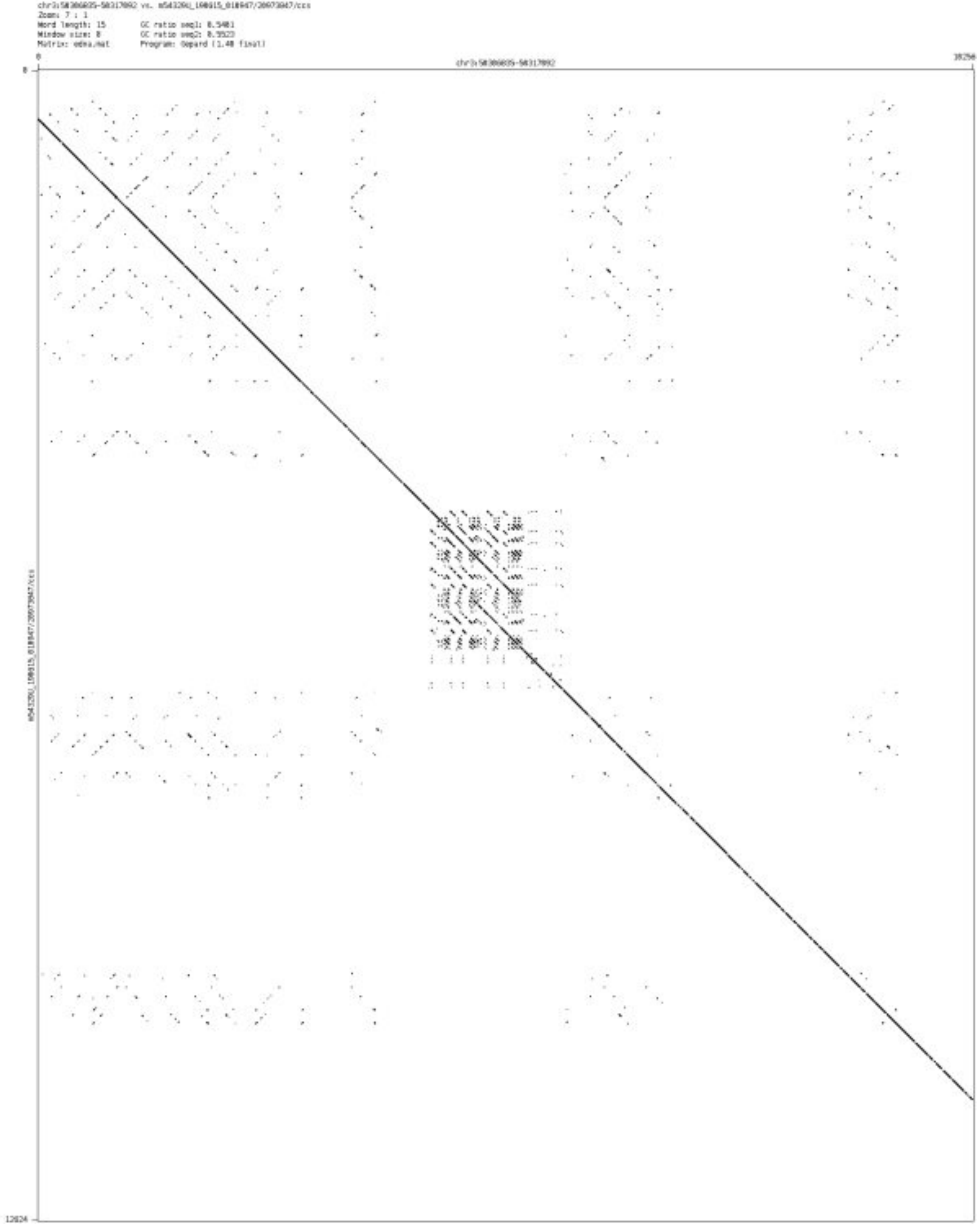

- chr5_7372546_7372667

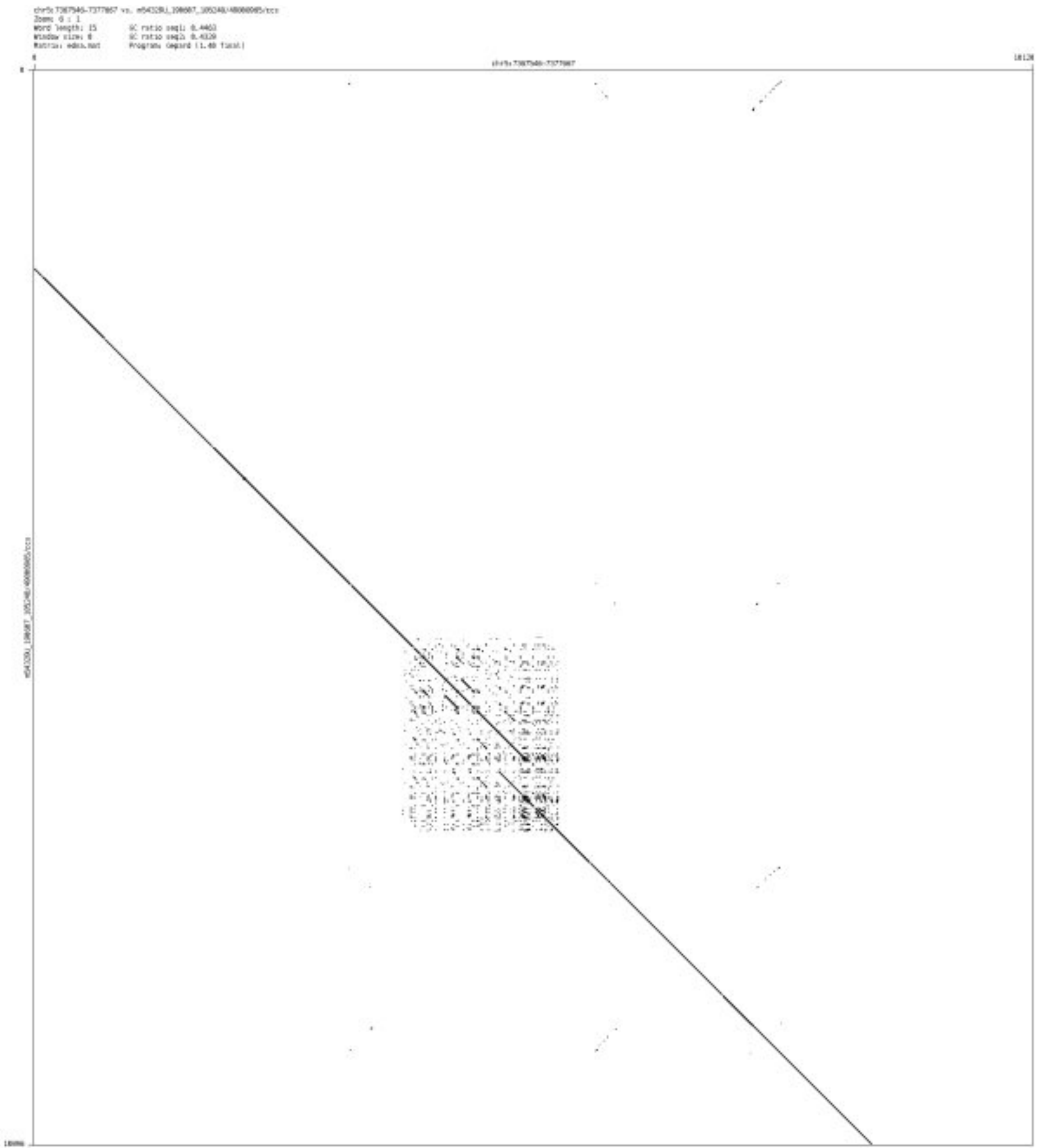

- chr7_55167302_55167528

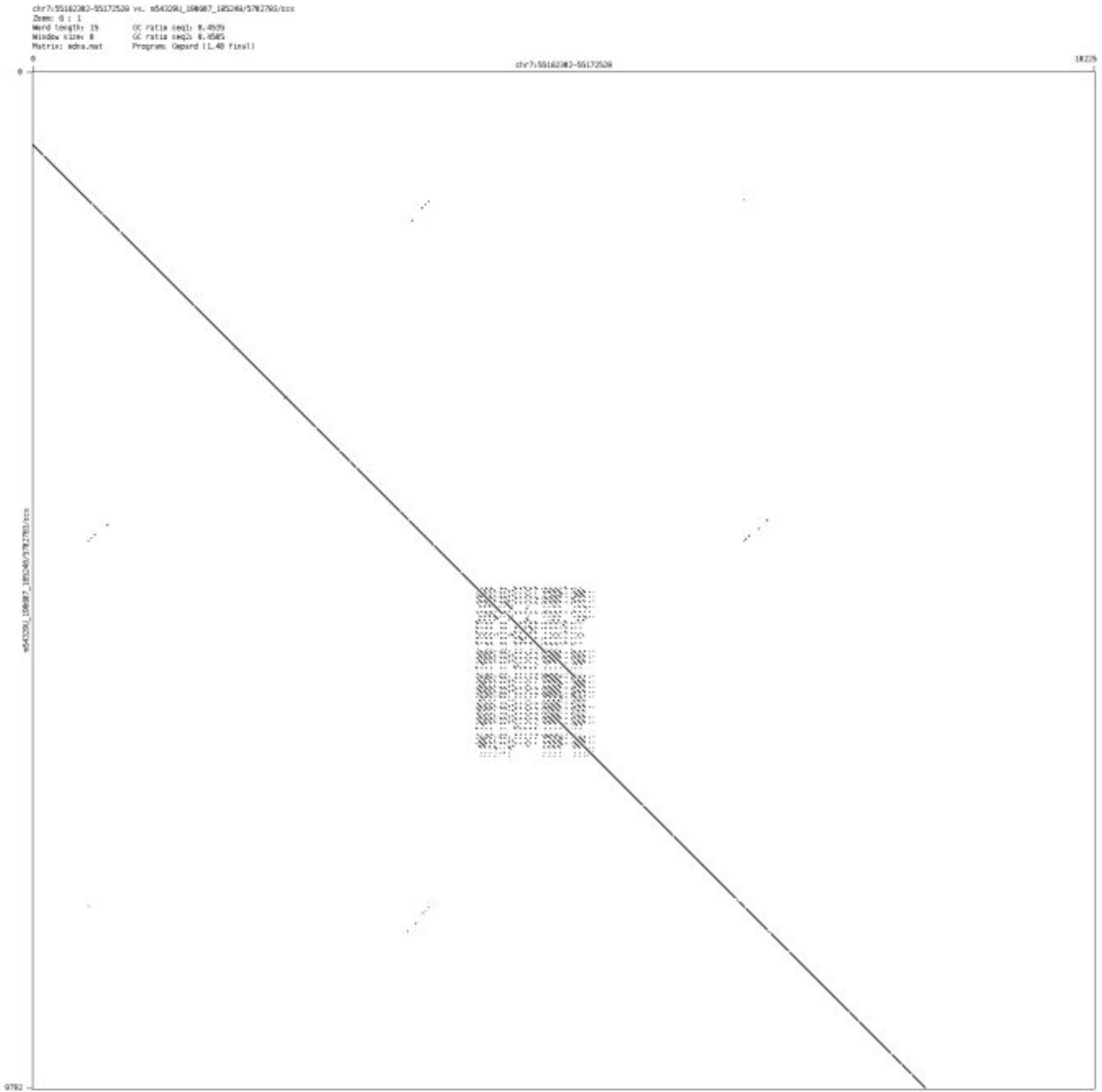

- chr7_158273882_158274358

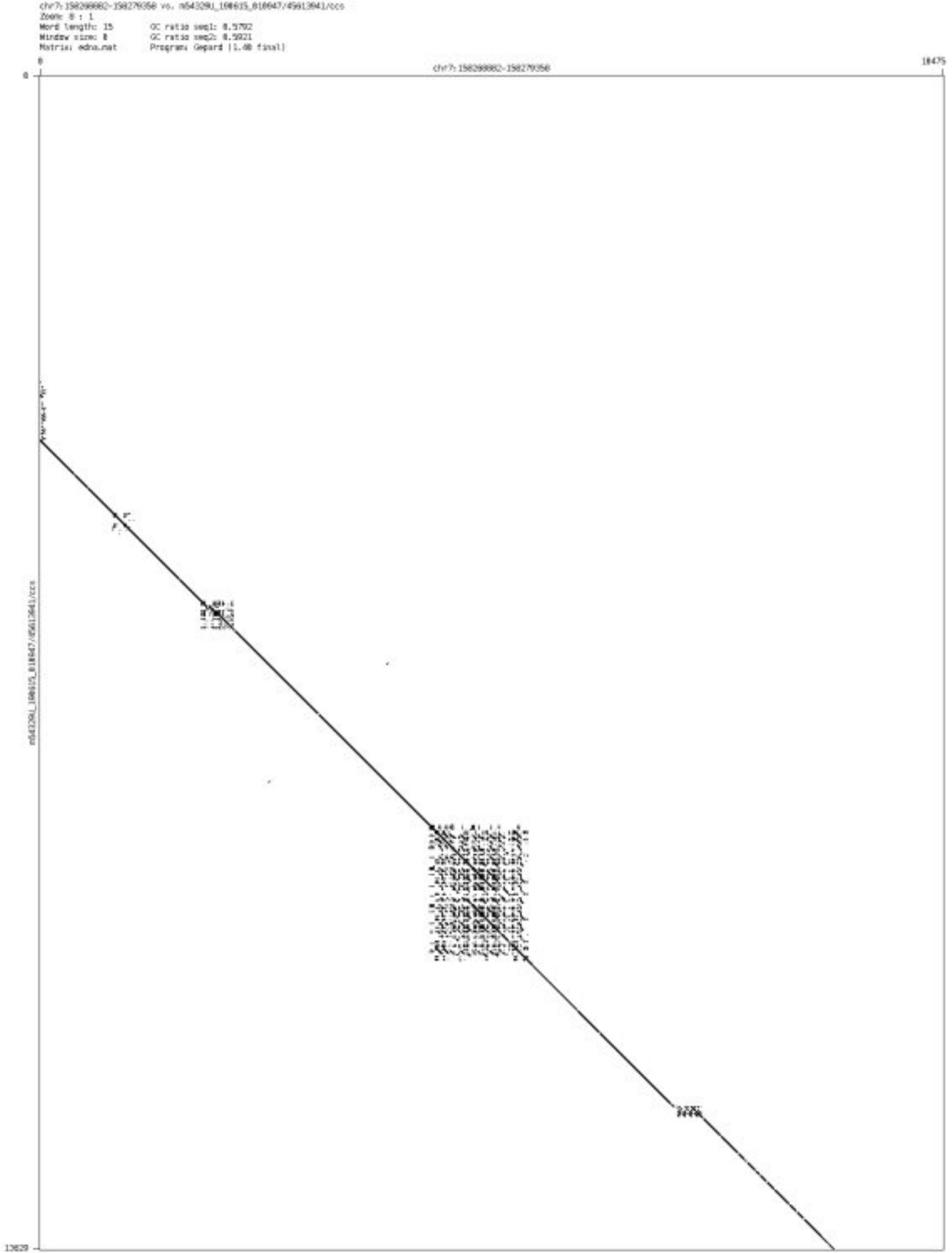

➢ chr8_1164182_1164270

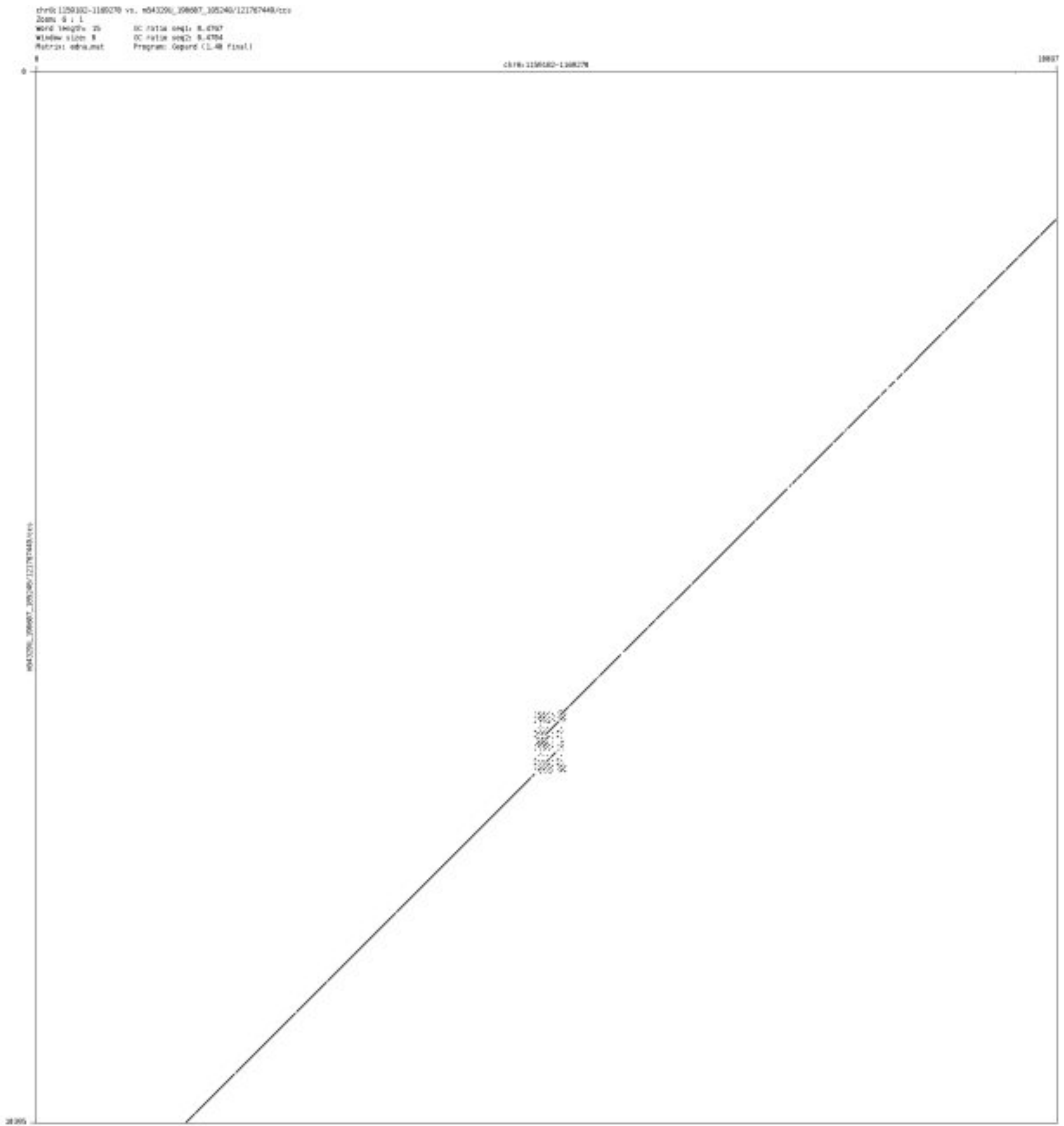

➢ chr10_132820153_132824176

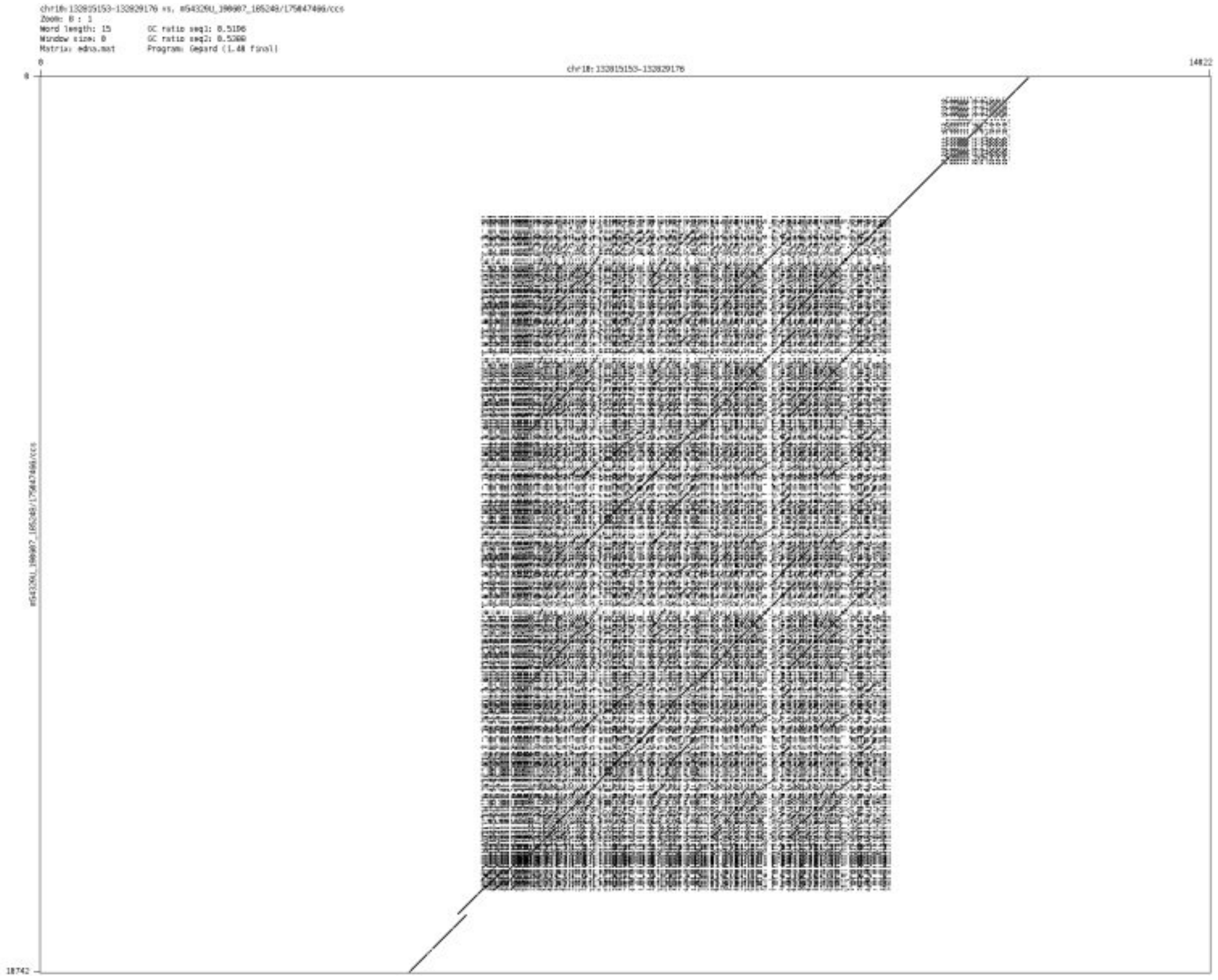

- ➢ chr11_61099653_61099916

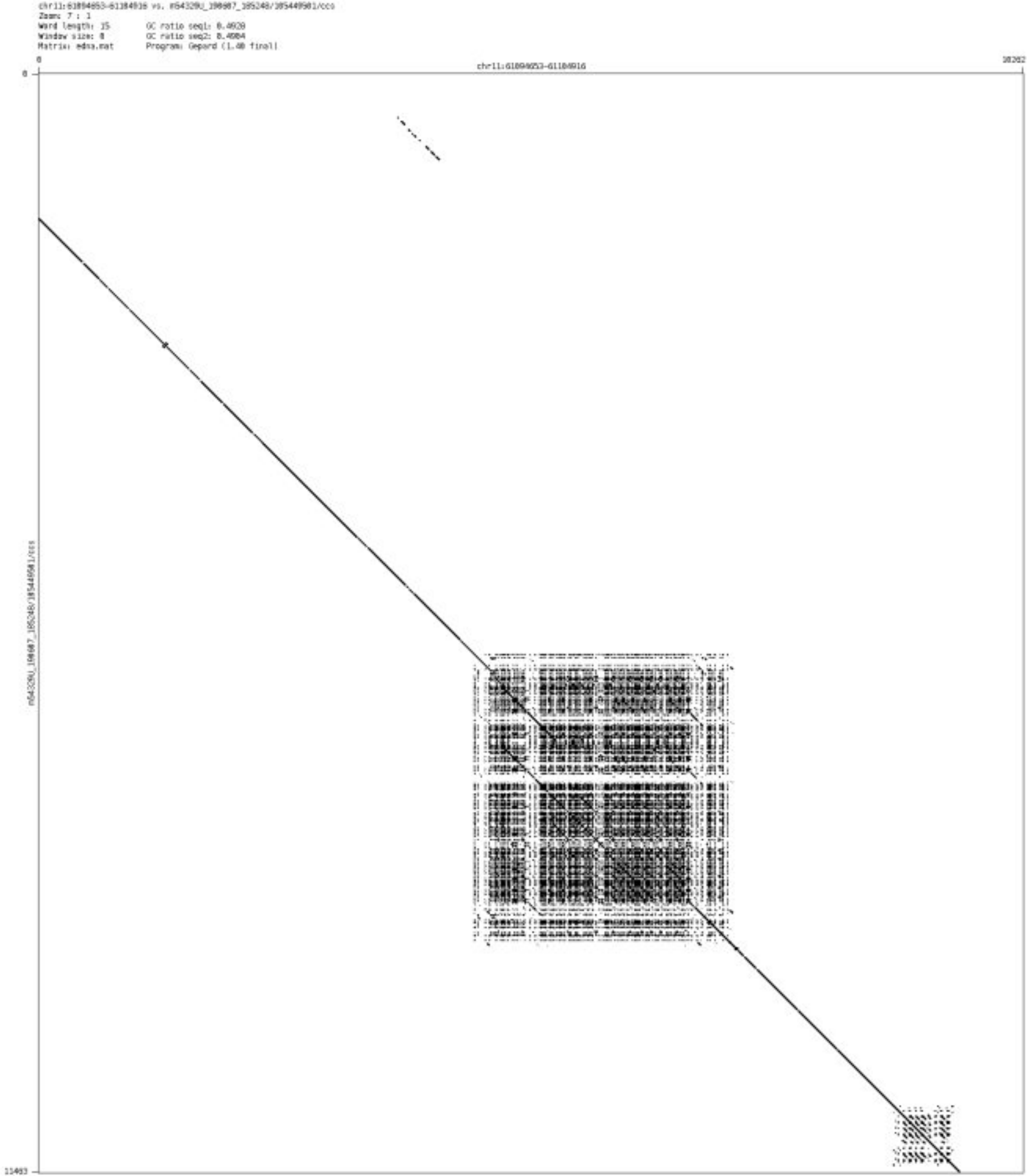

- ➢ chr11_134308286_134308429

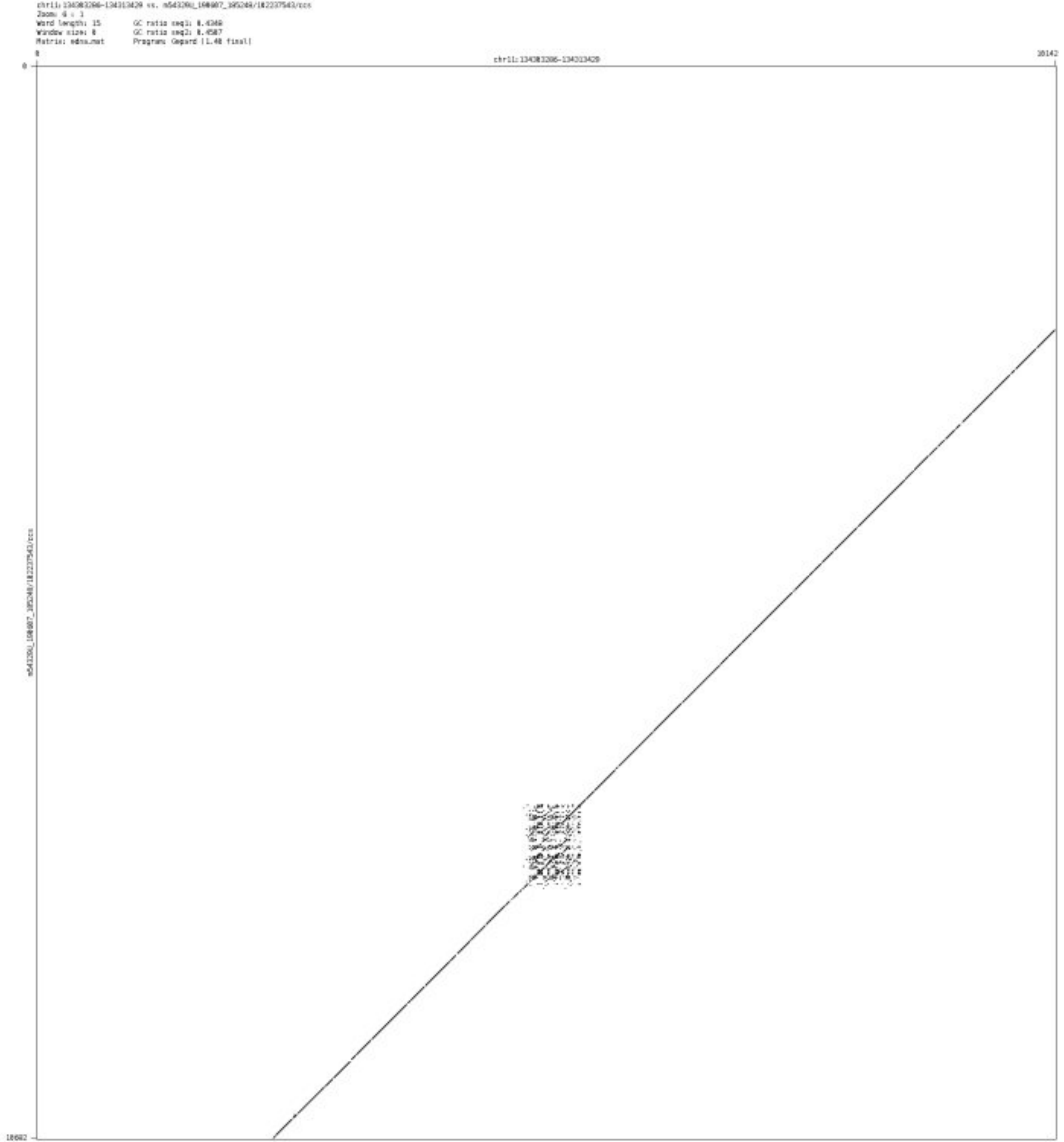

- chr12_109661571_109661755

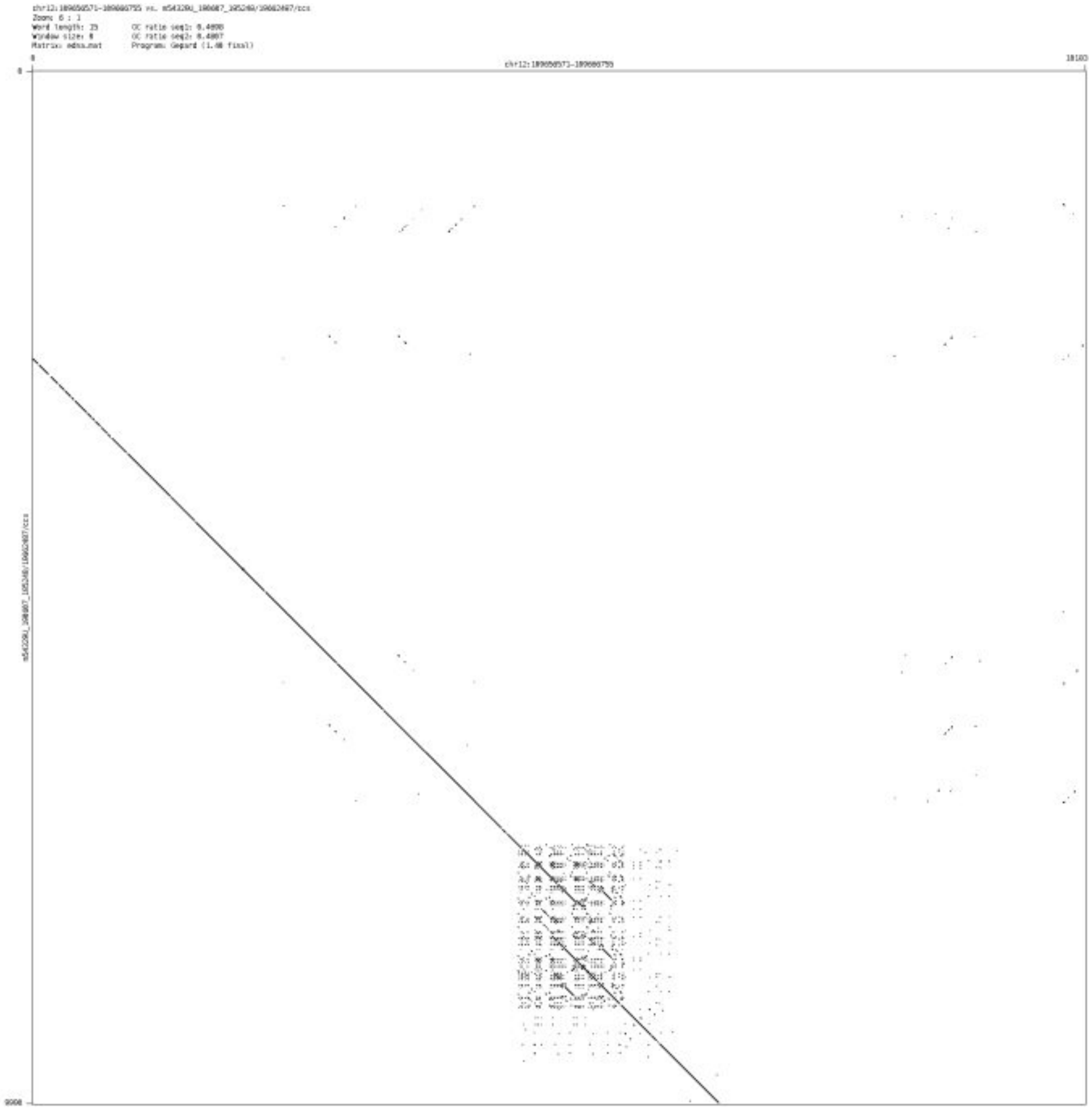

- chr12_131046207_131046337

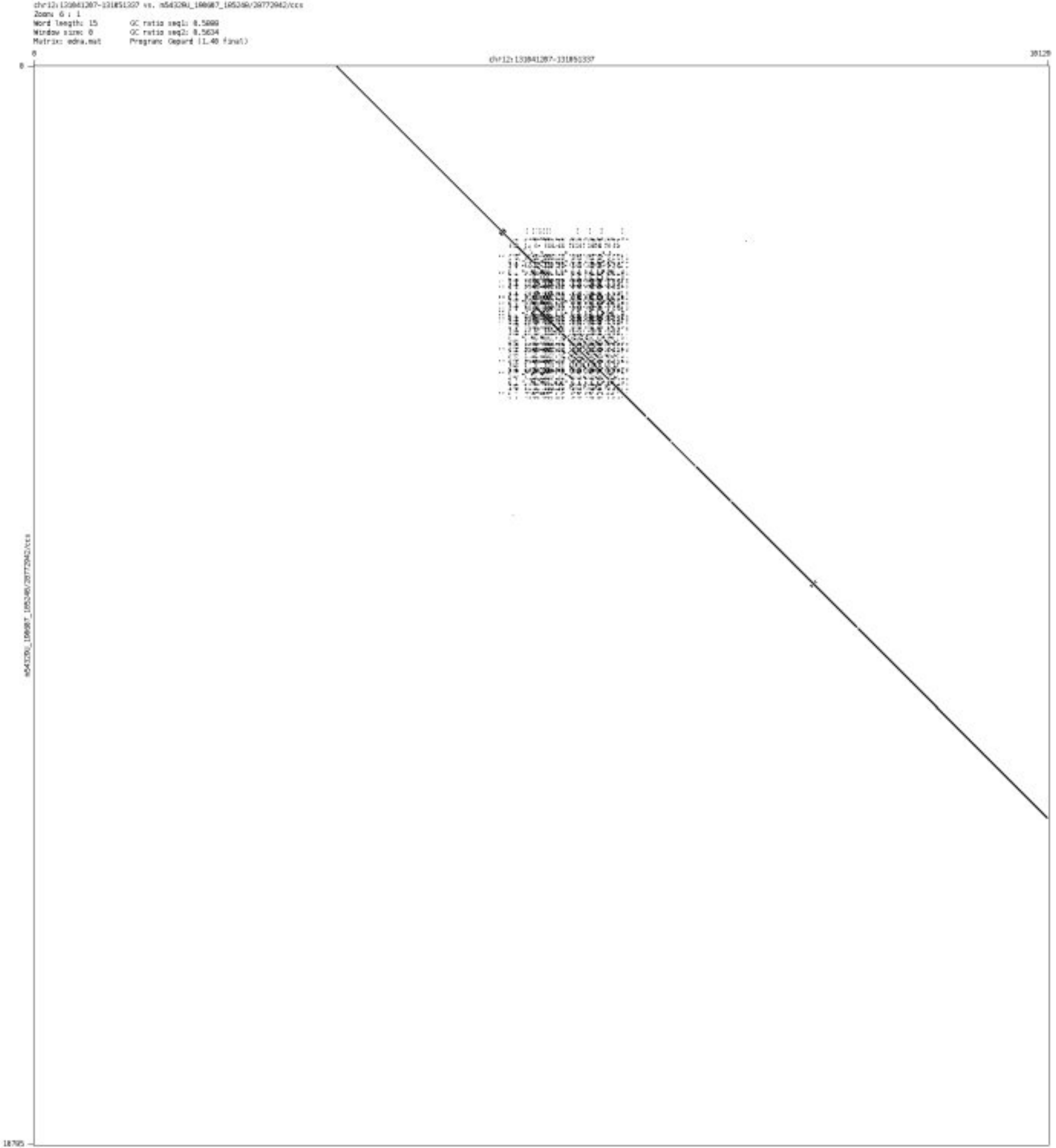

➢ chr12_132387869_132387963

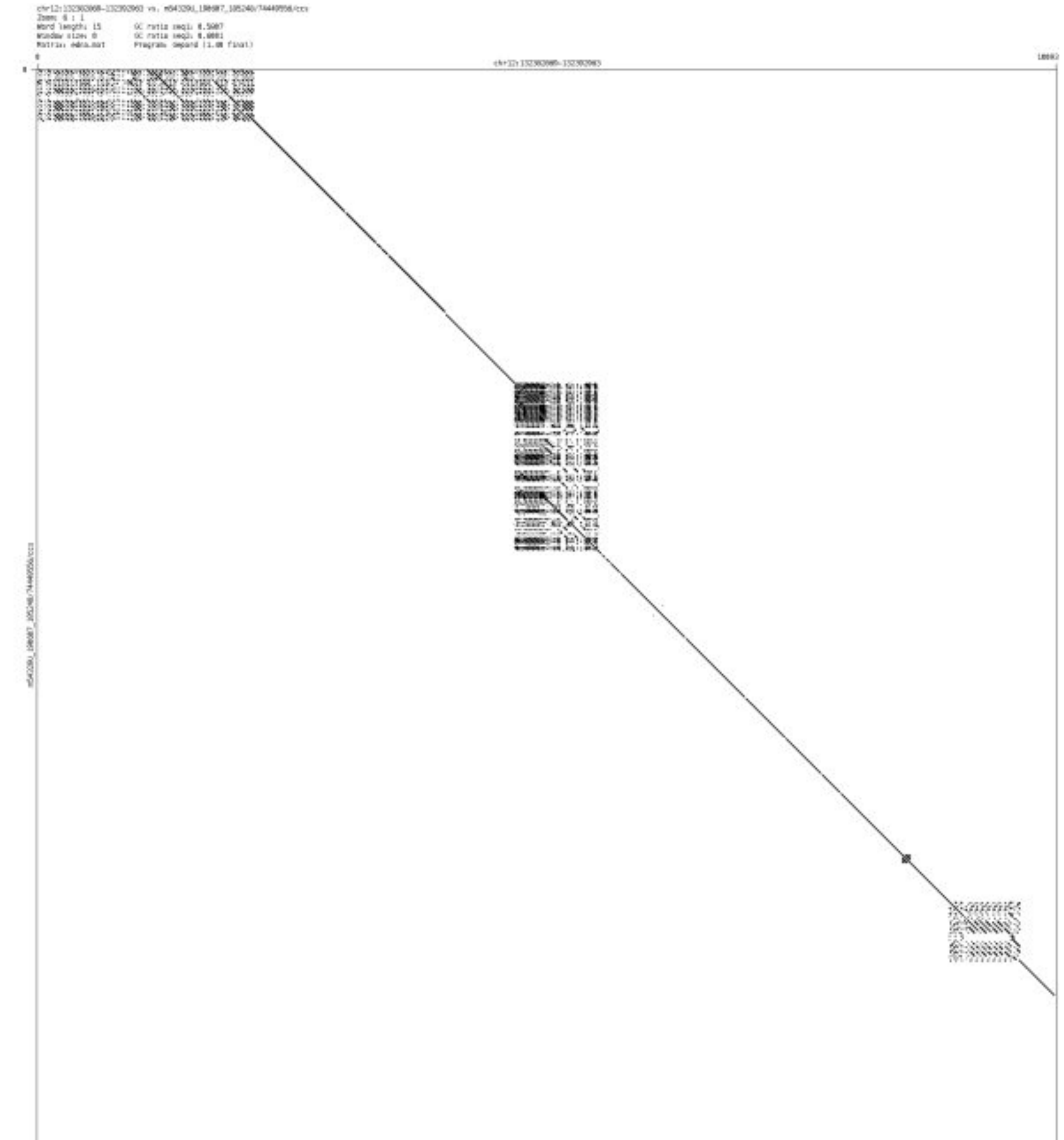

➢ chr15_91749460_91749774

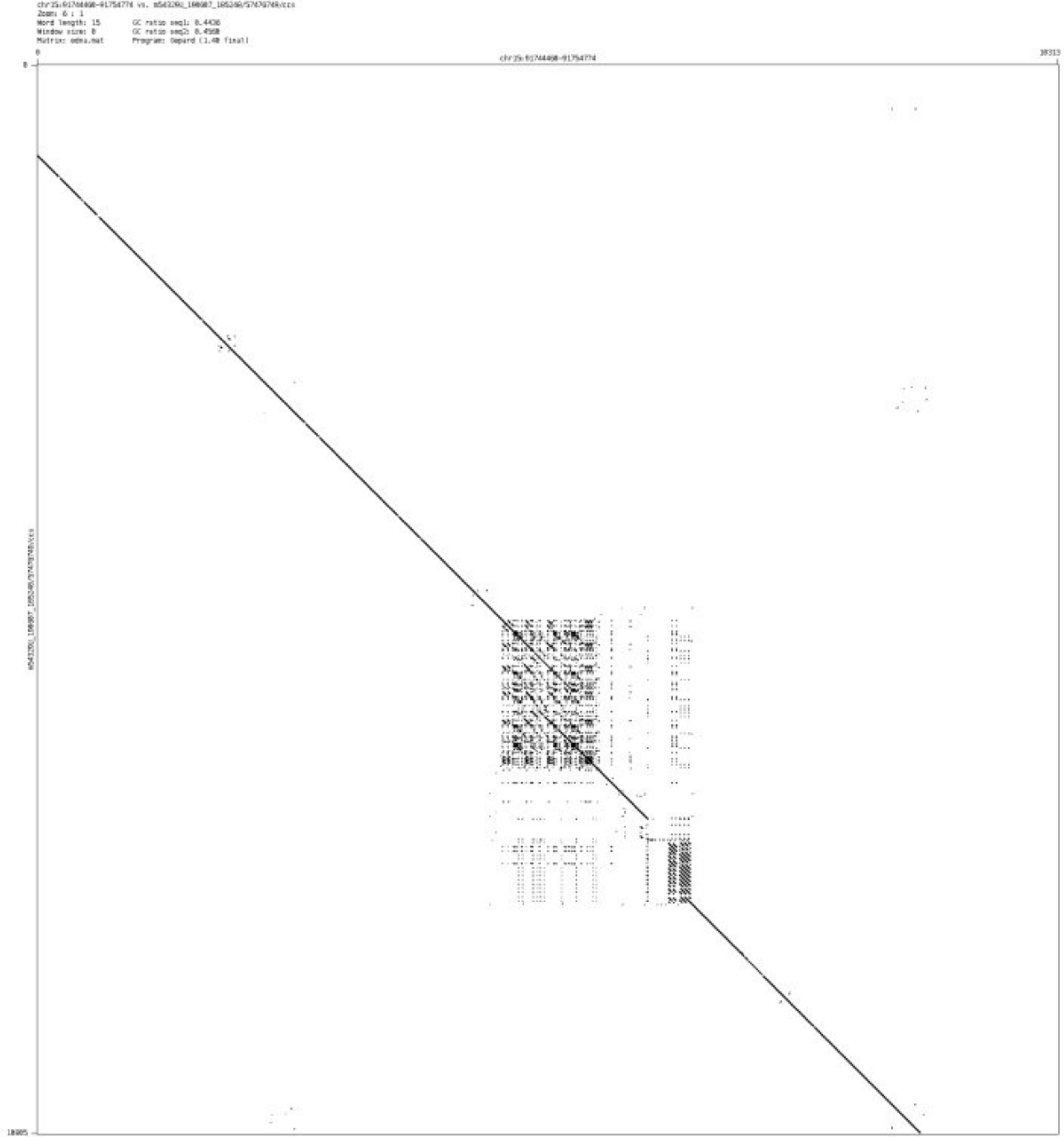

- chr16_8562755_8565066

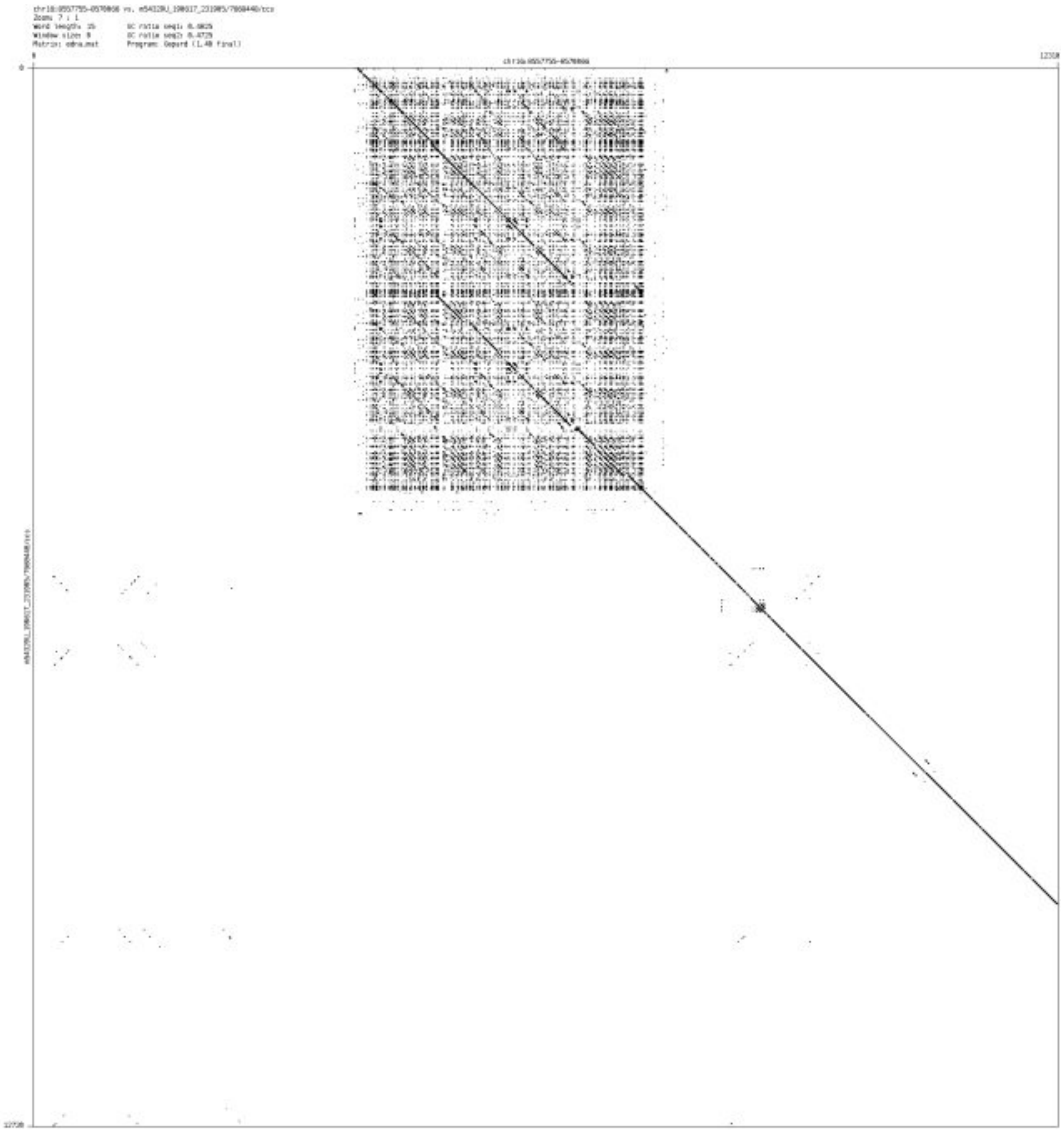

### Simple SVs-Case A

➢ chr2_240527386_240527609

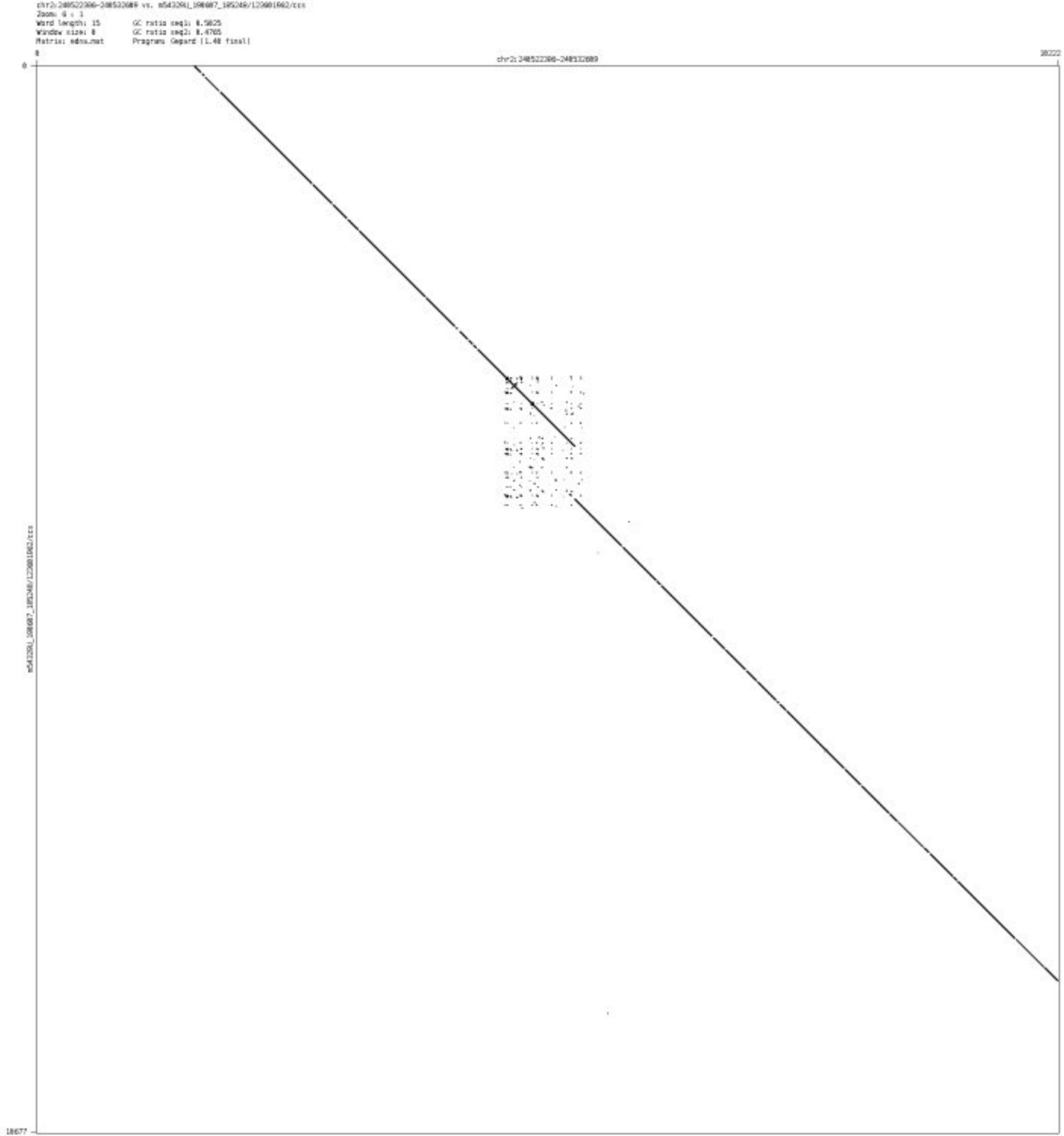

- chr6_160535459_160535555

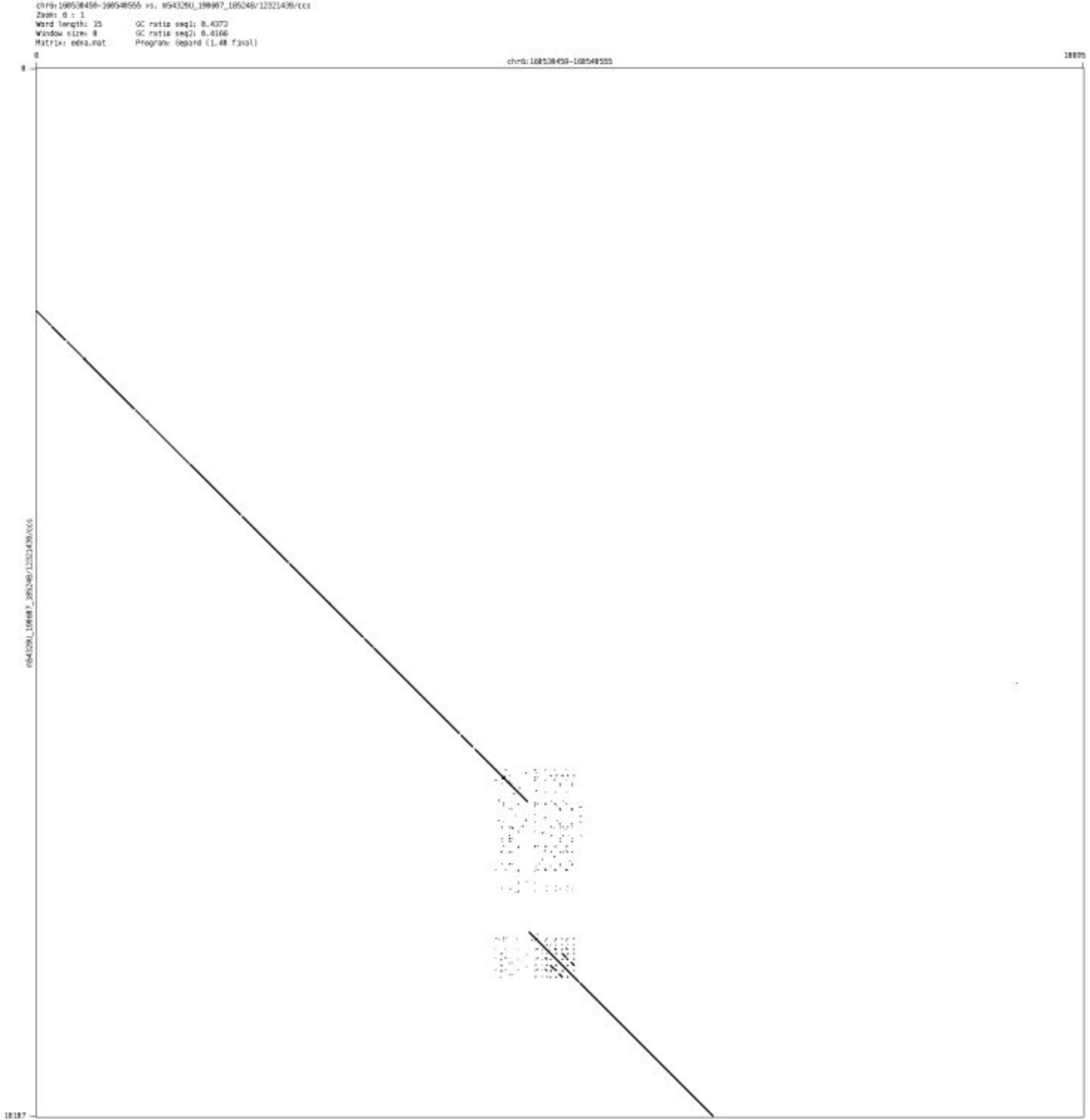

- ➢ chr9_123347689_123347773

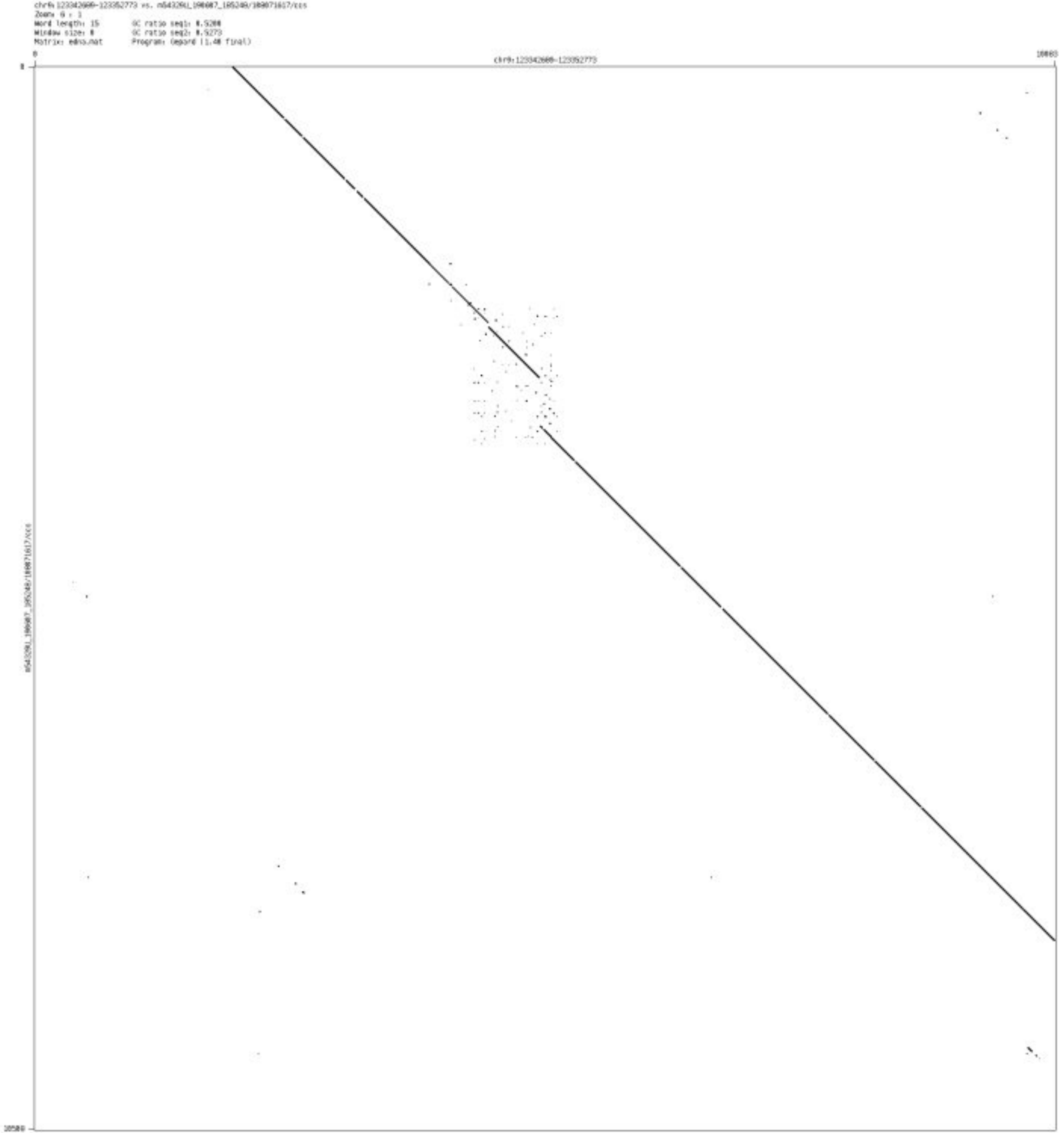

➢ chr10_7526763_7526835

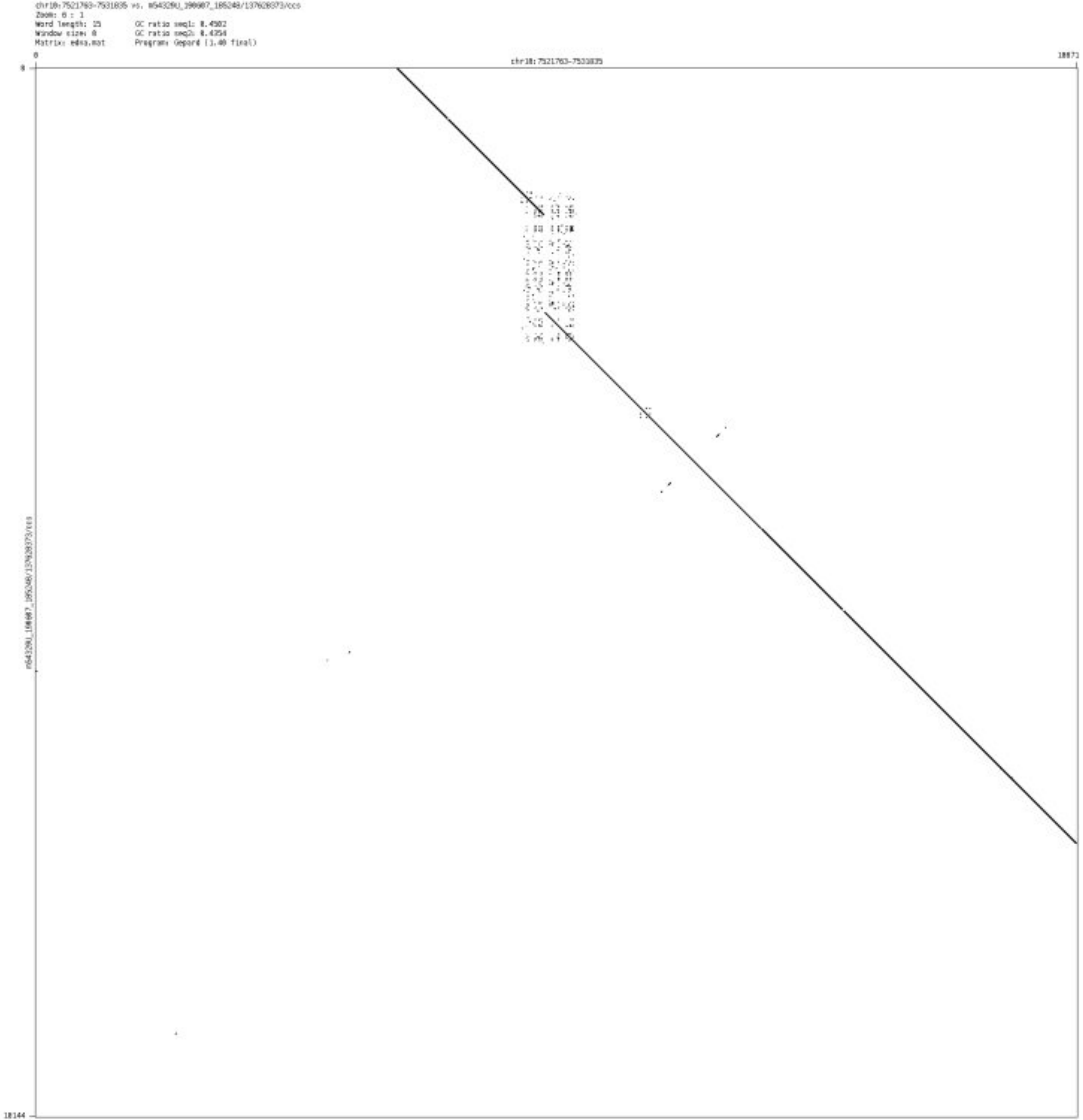

### Simple SVs-Case B

- chr1_2440414_2440528

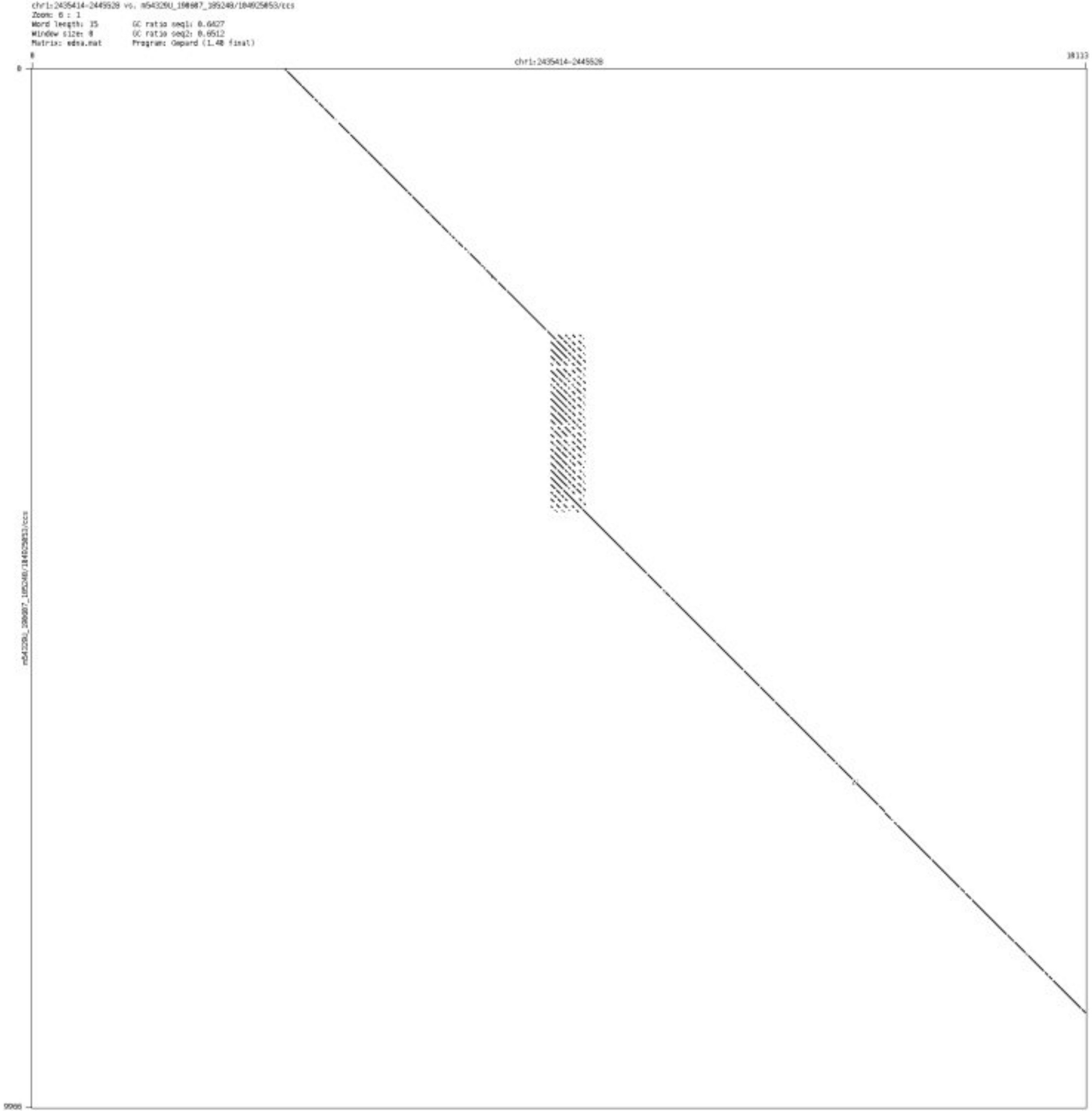

➢ chr1_3212601_3212686

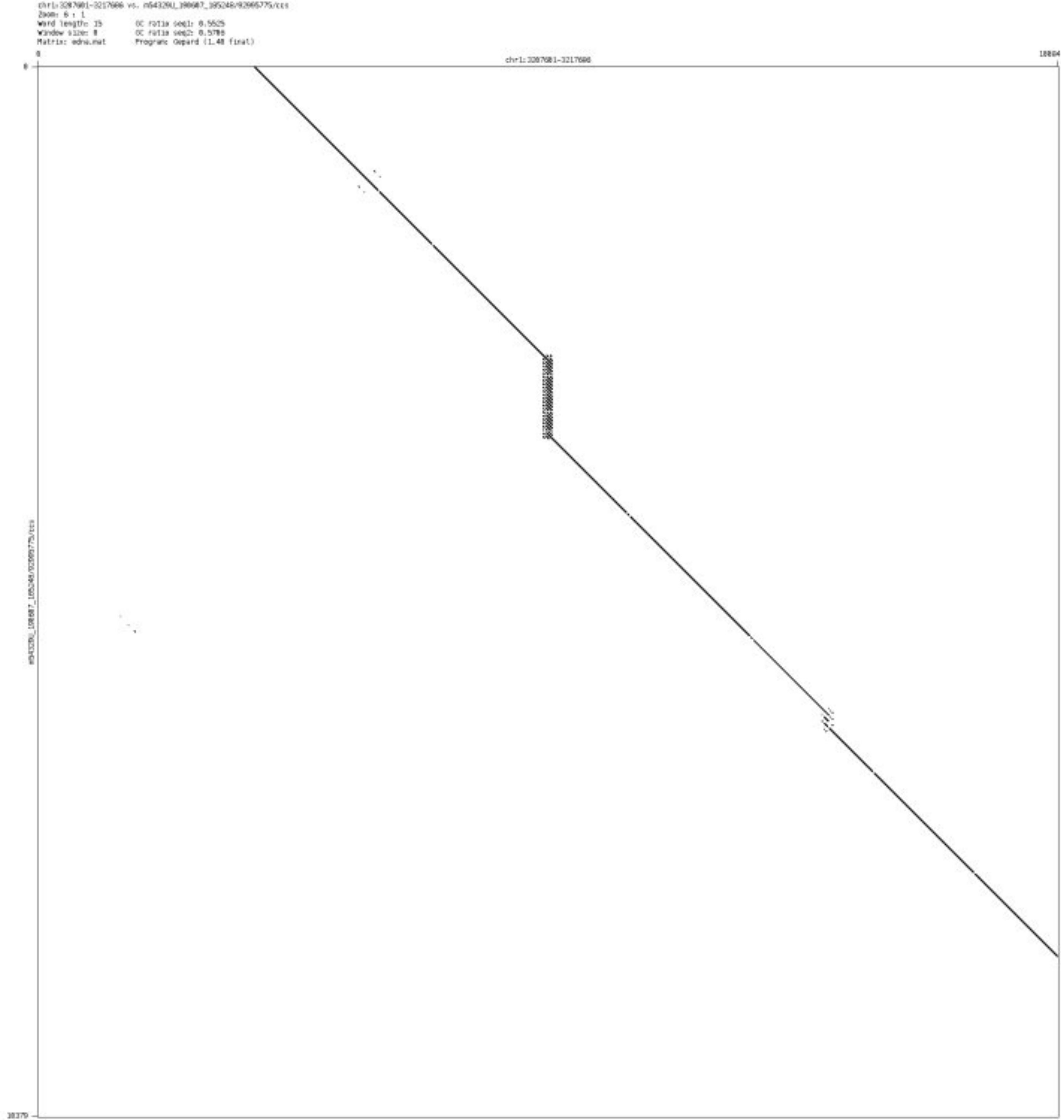

➢ chr2_238909005_238909139

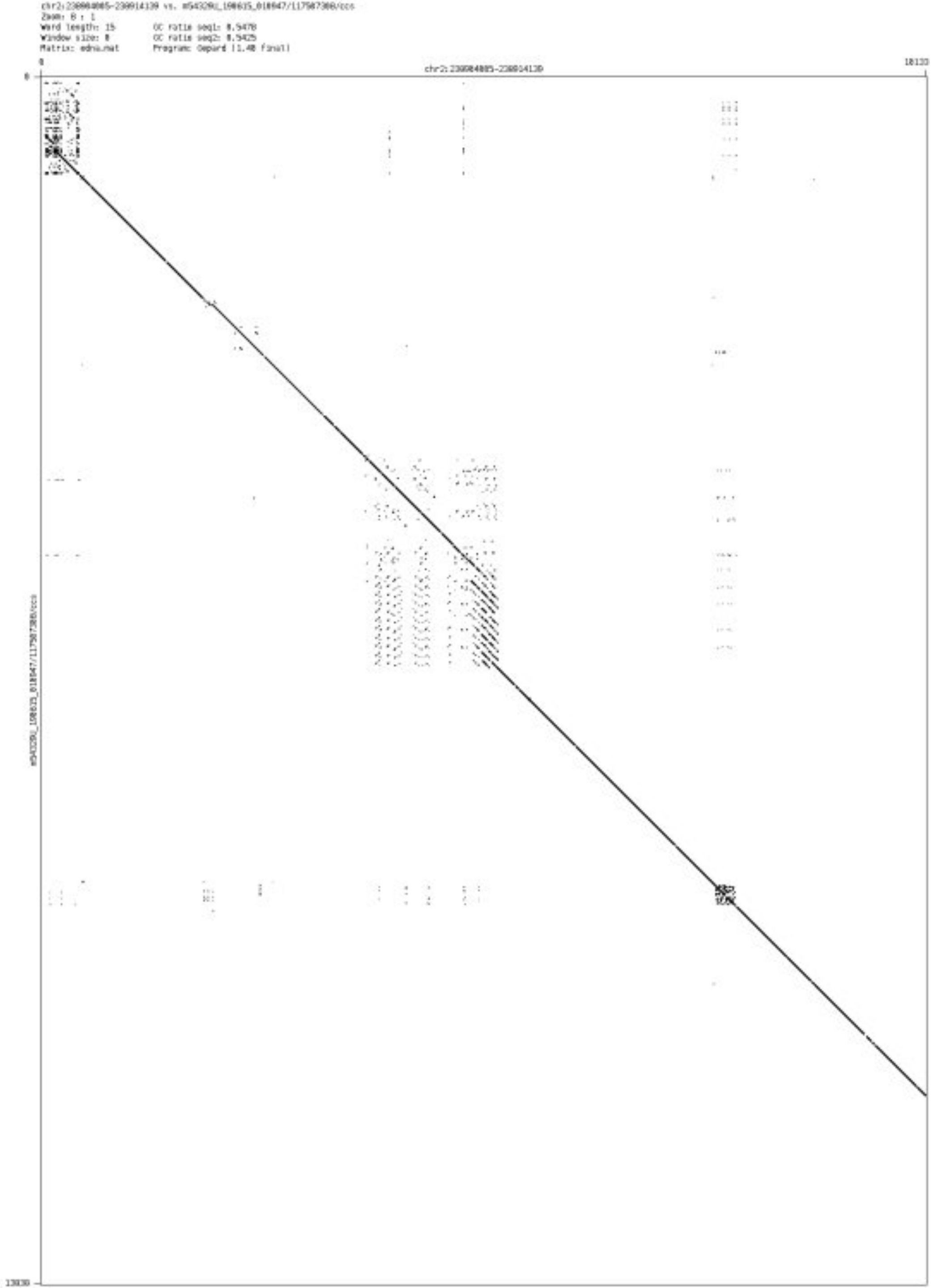

➢ chr3_77355086_77355236

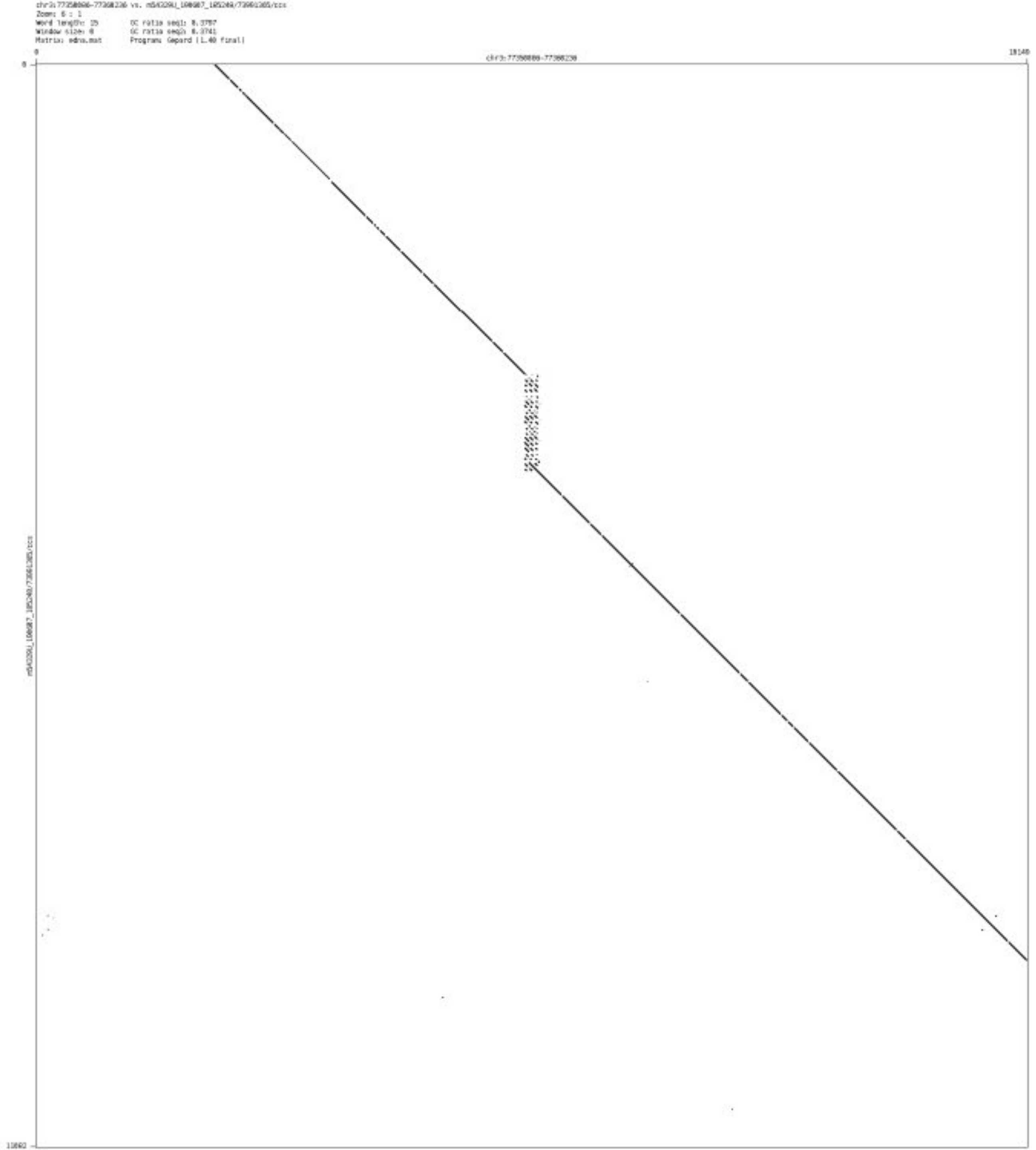

#### ➢ chr7_205261_206826

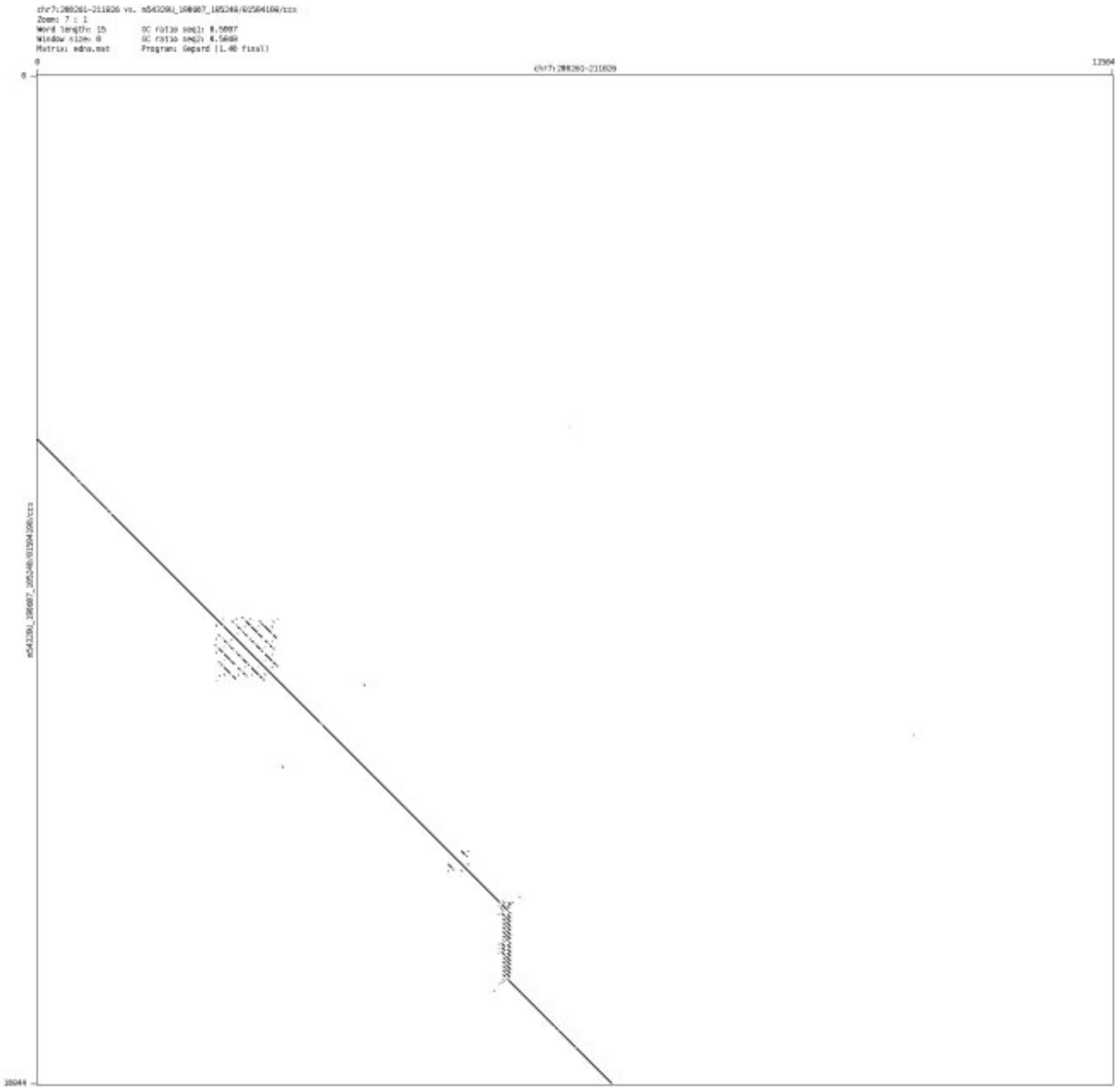

- chr7_158166484_158166596

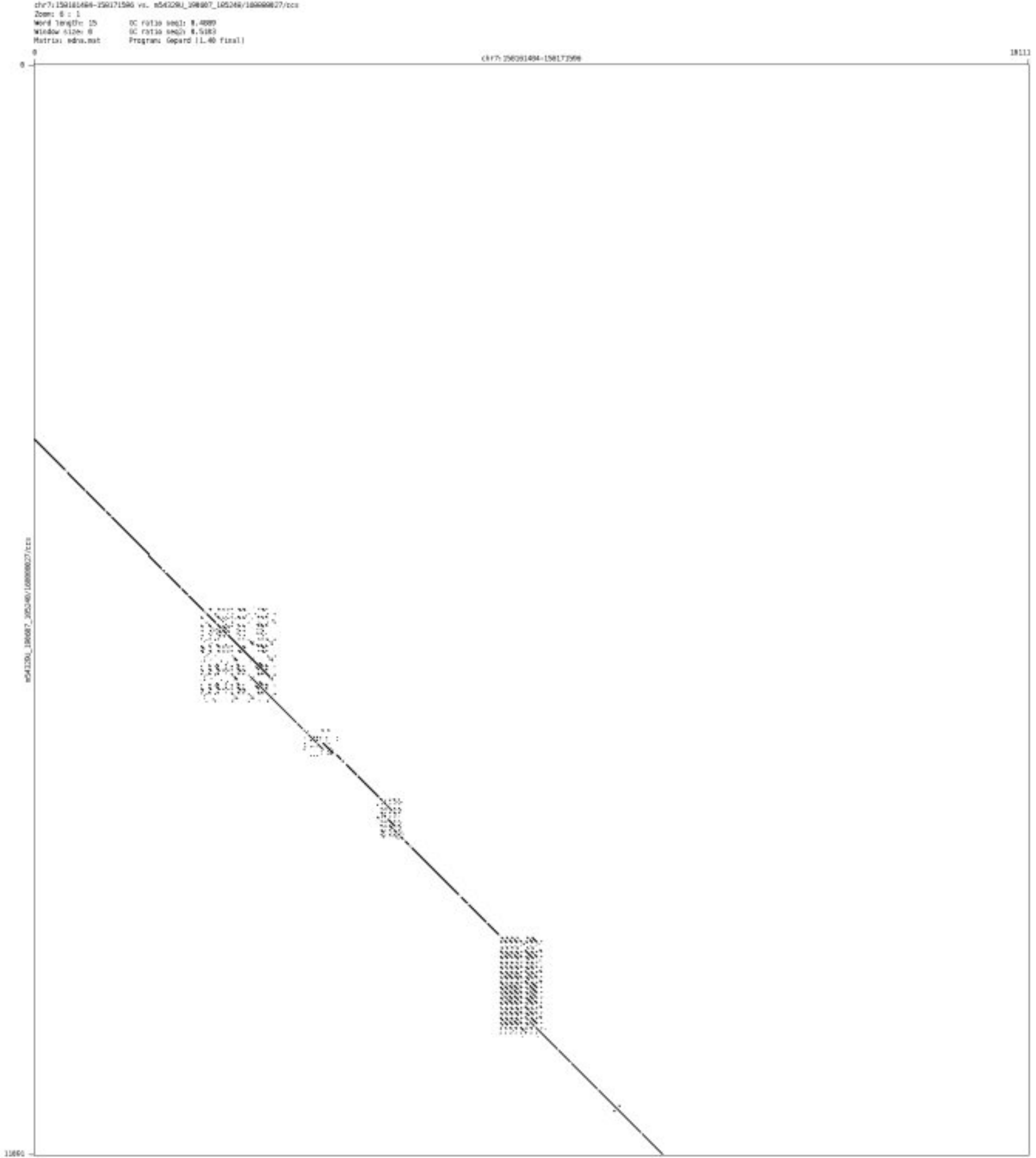

- chr10_718255_718429

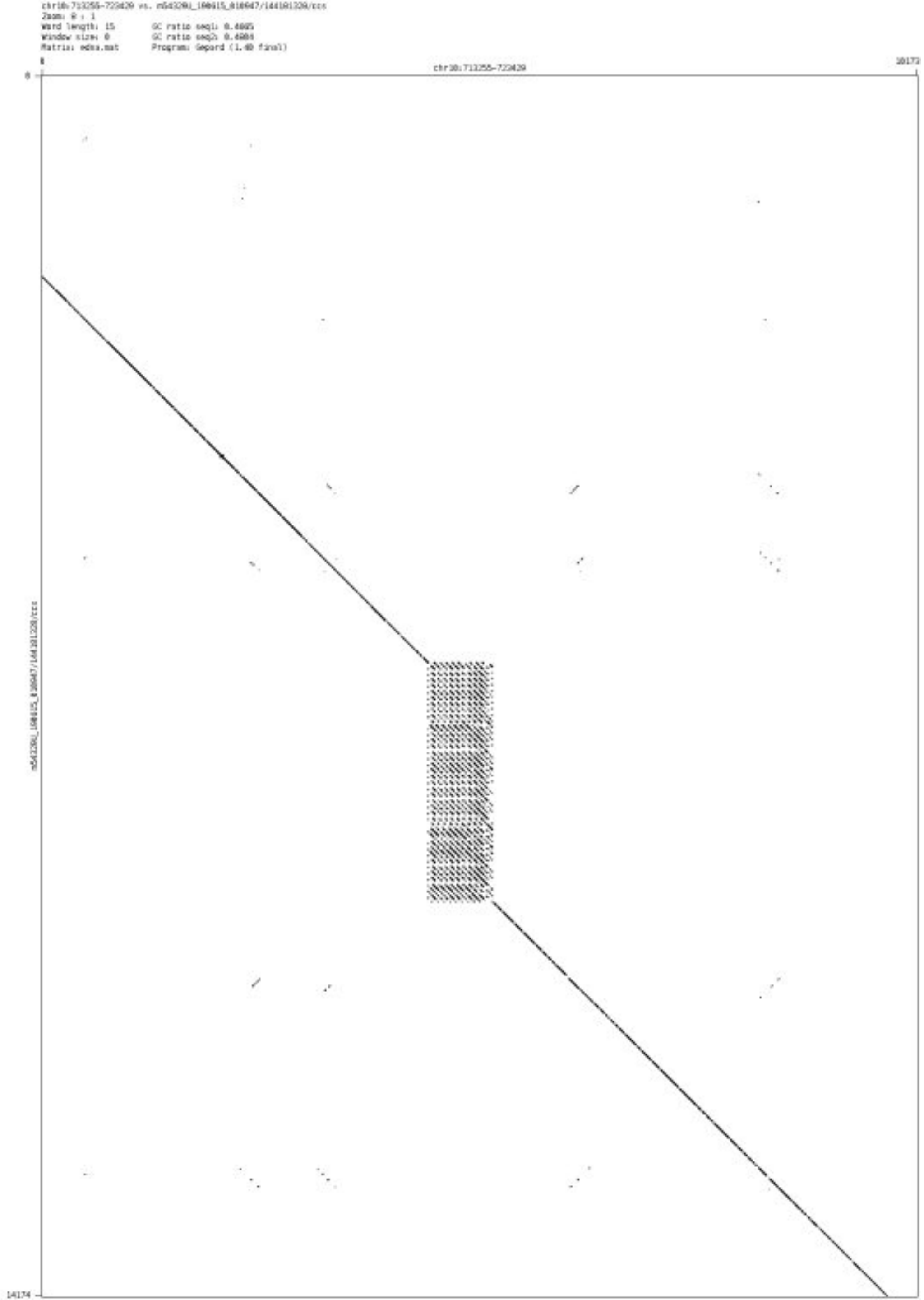

- chr11_120328683_120328976

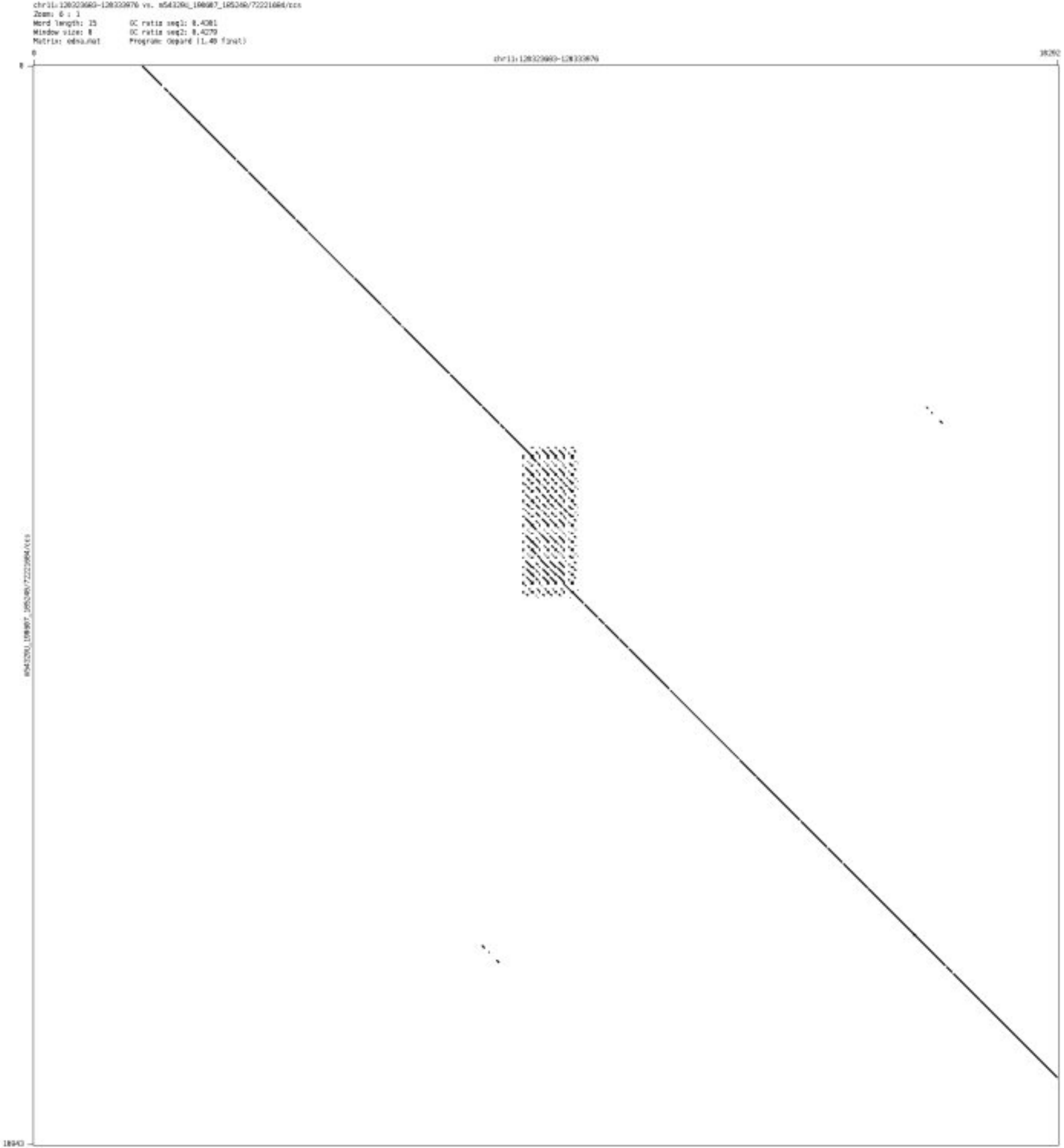

- chr13_113419003_113419055

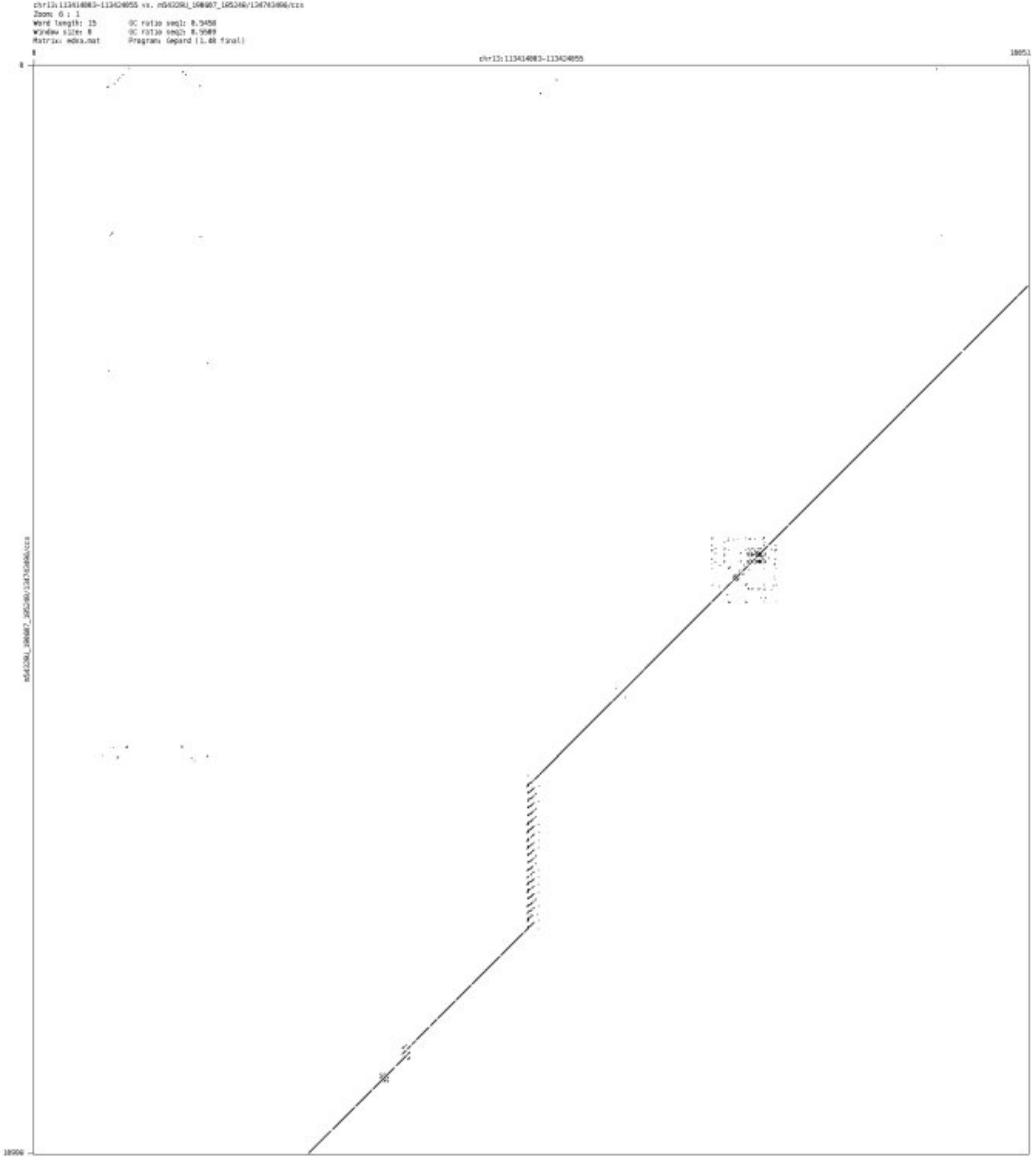

- ➢ chr16_954959_955110

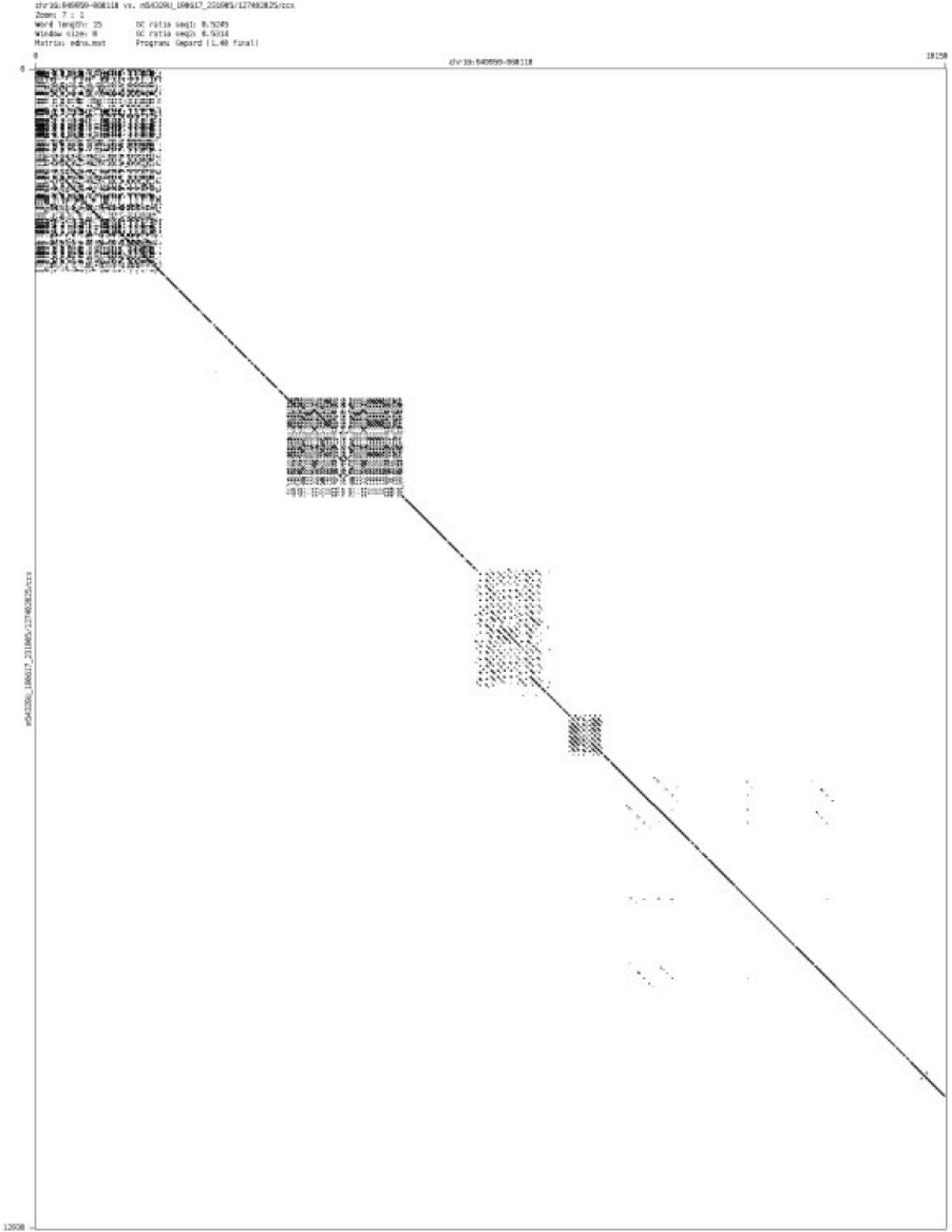

➢ chr17_952712_952801

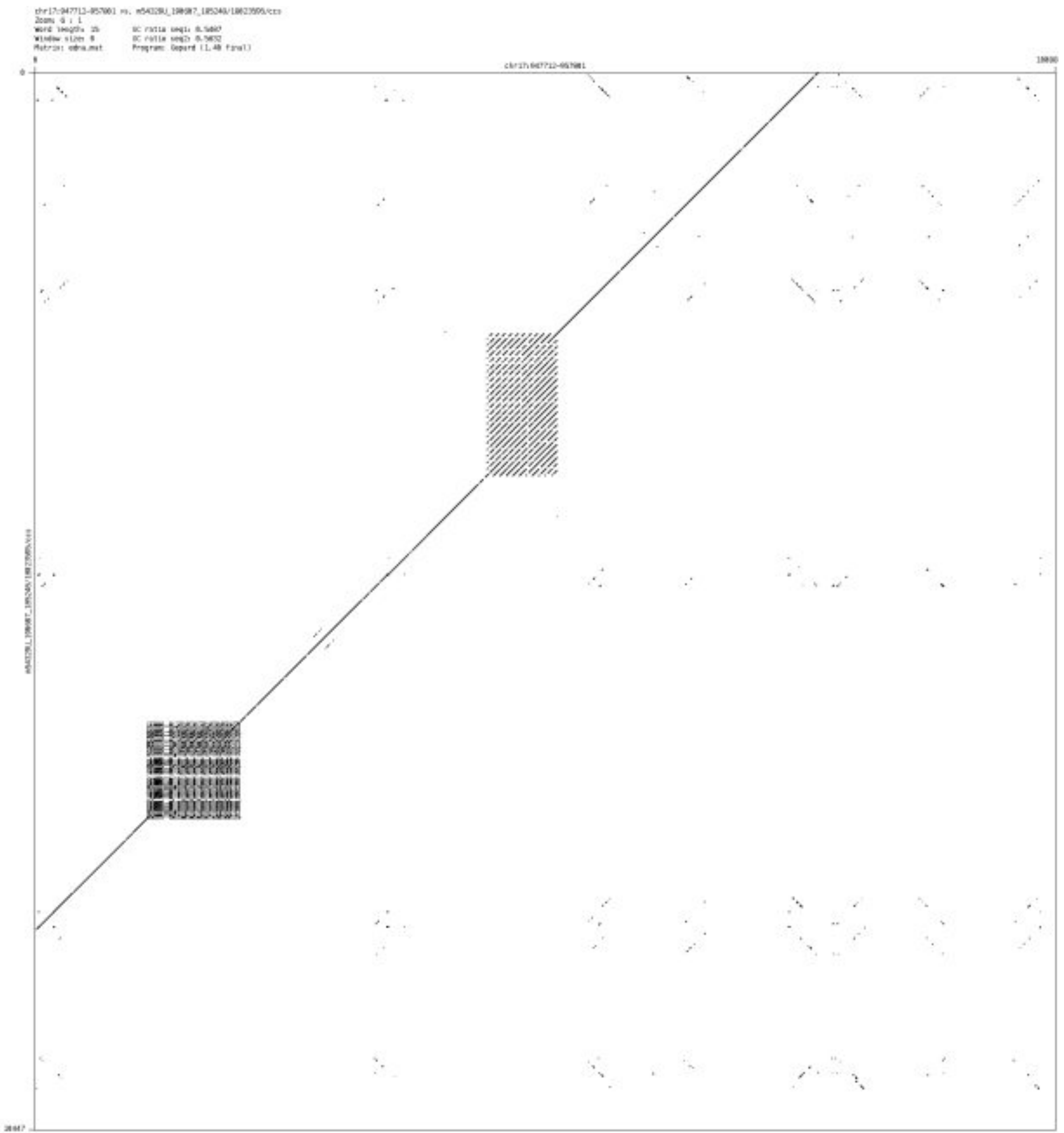

- chr20_2236338_2236455

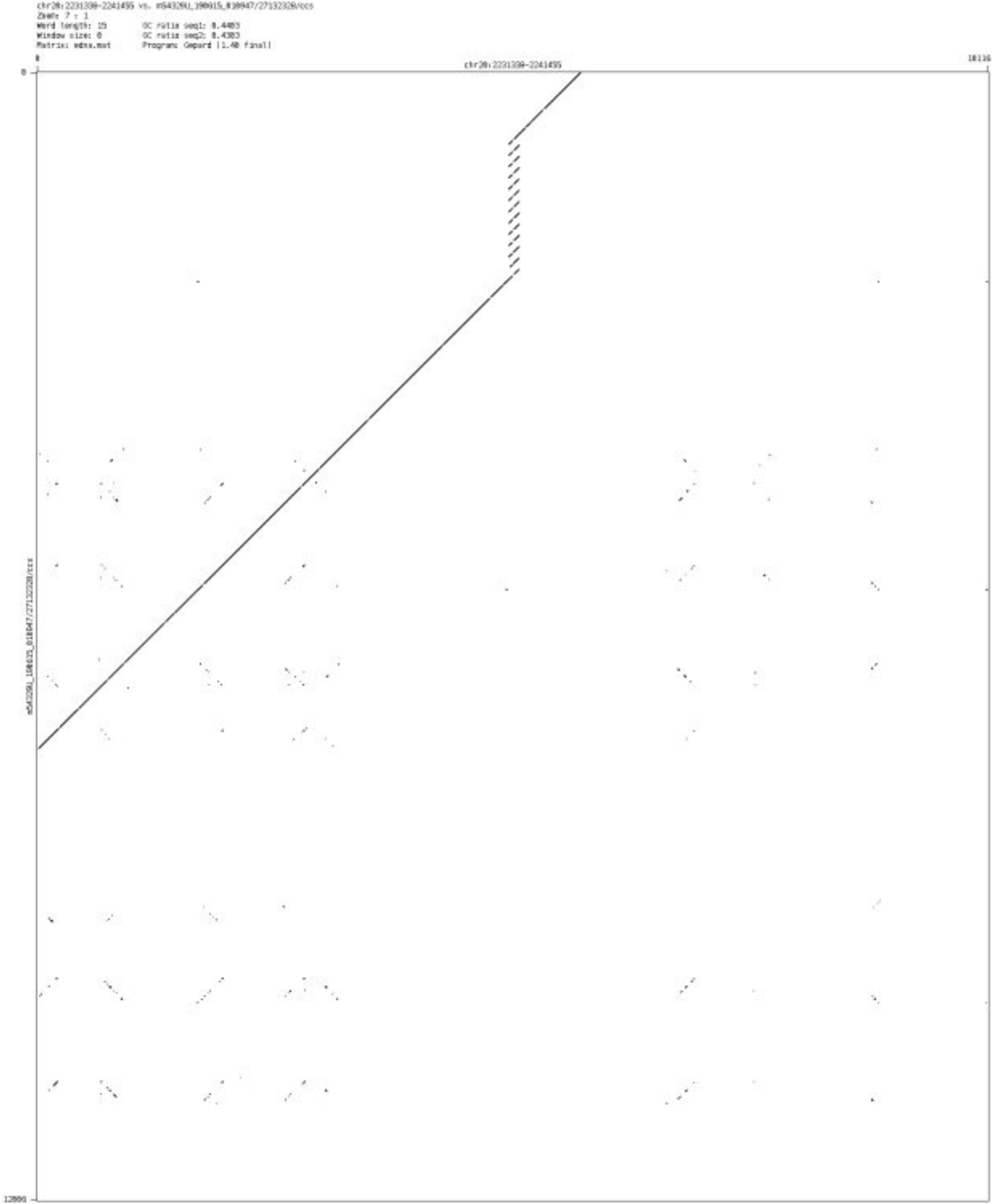

➢ chr20_61329291_61329439

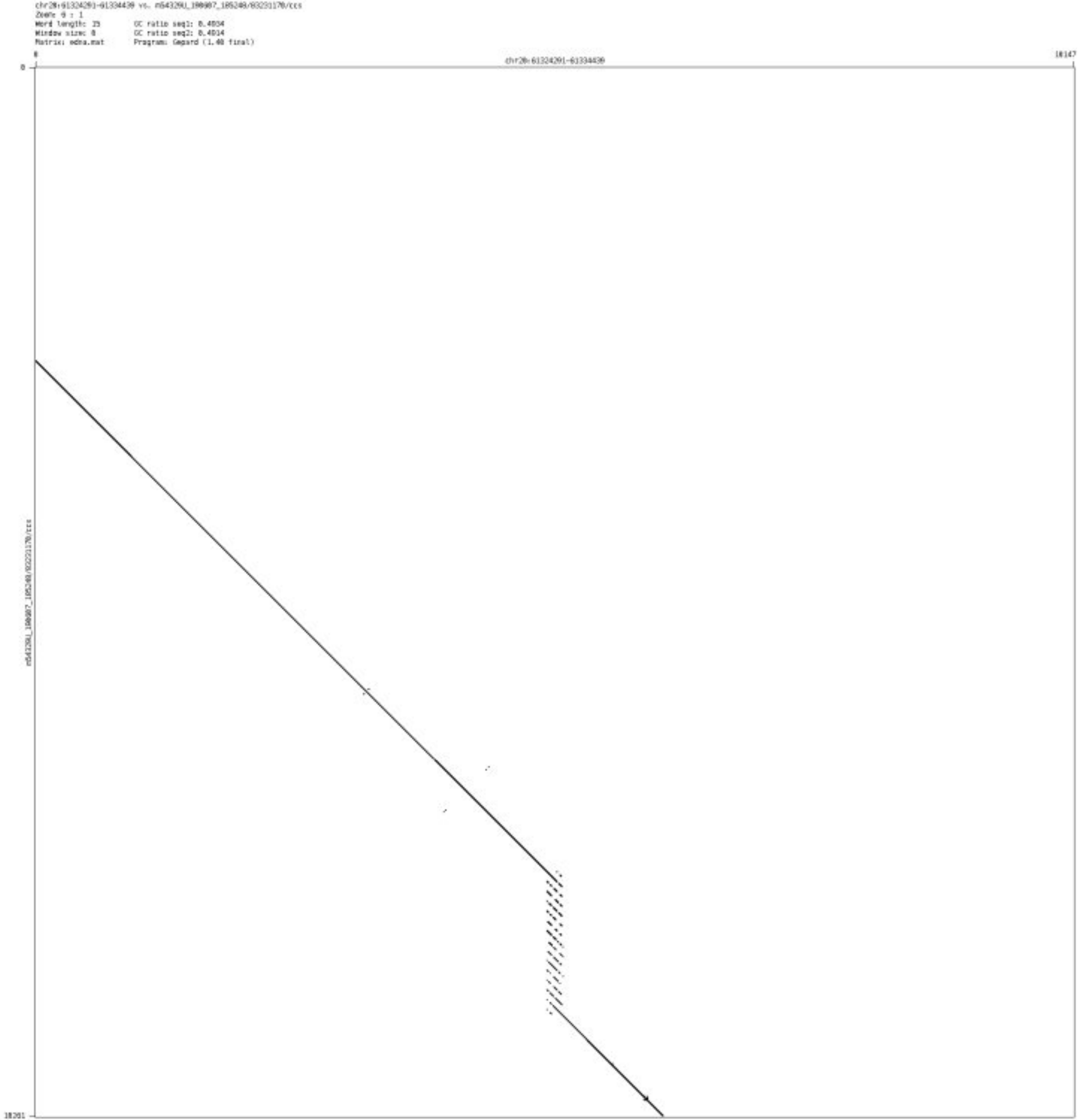

- chr21_45055043_45055102

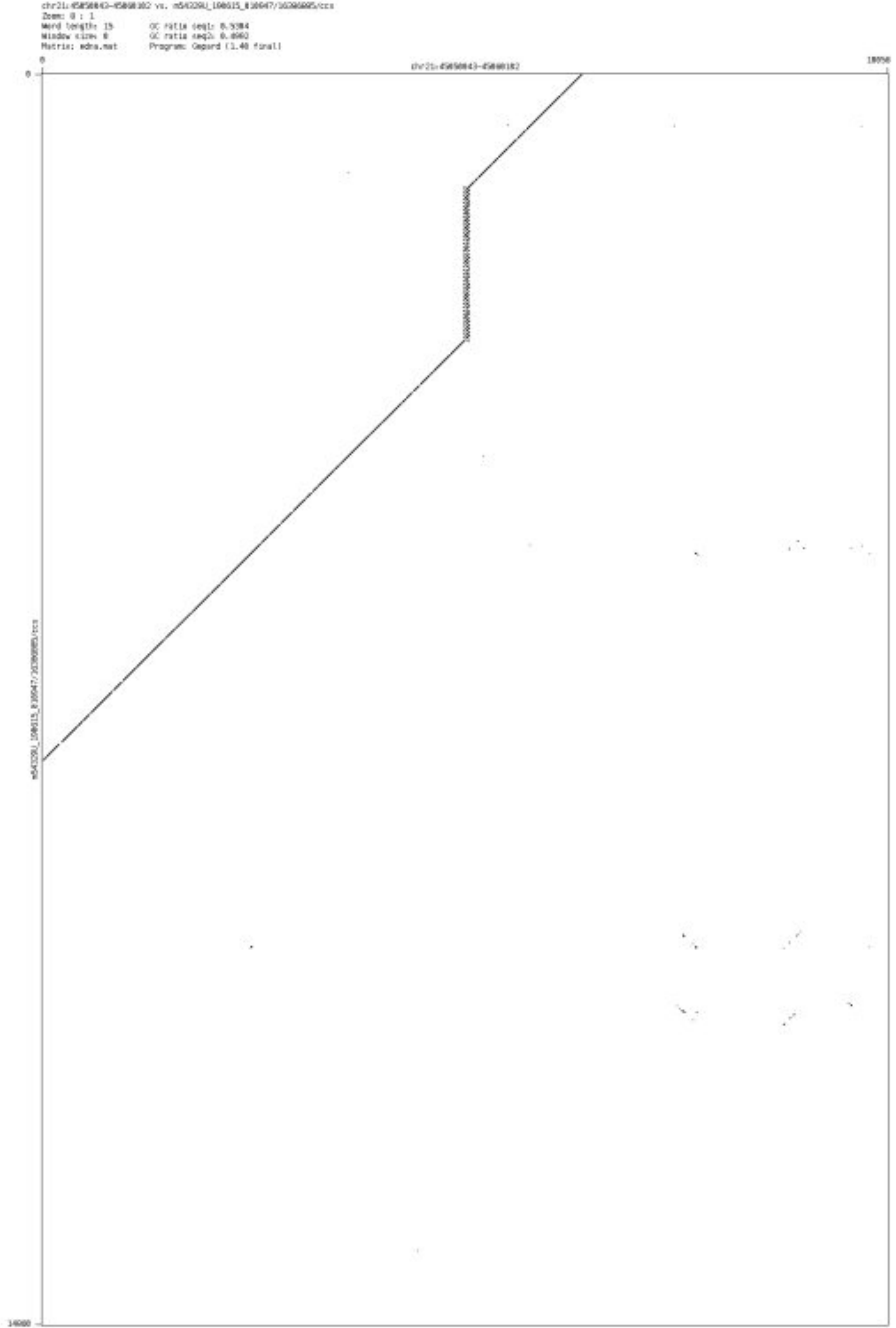

➢ chr13_113176451_113176522

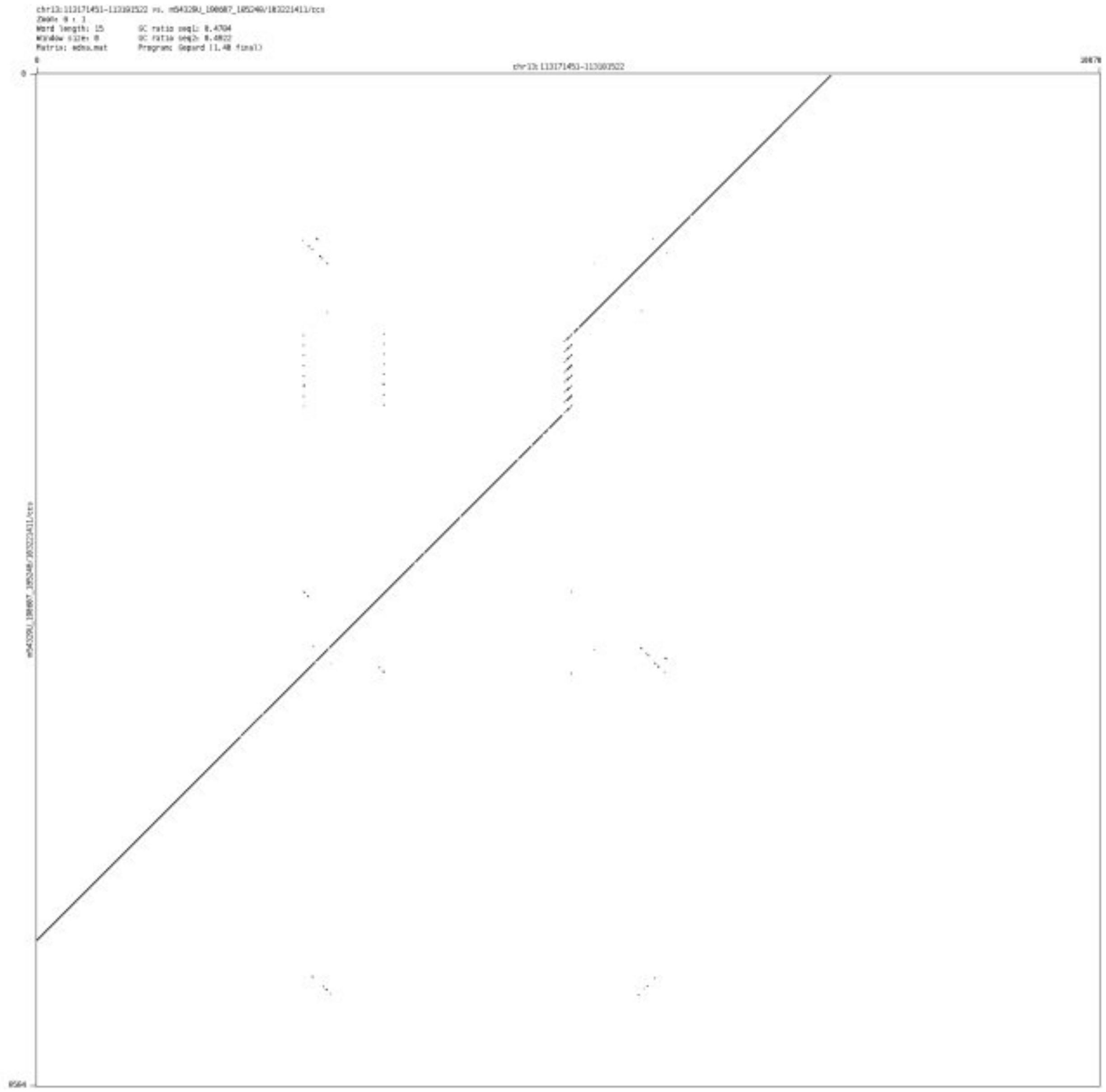

### Inconclusive events at STR/VNTR regions

- chr1_1030460_1030901

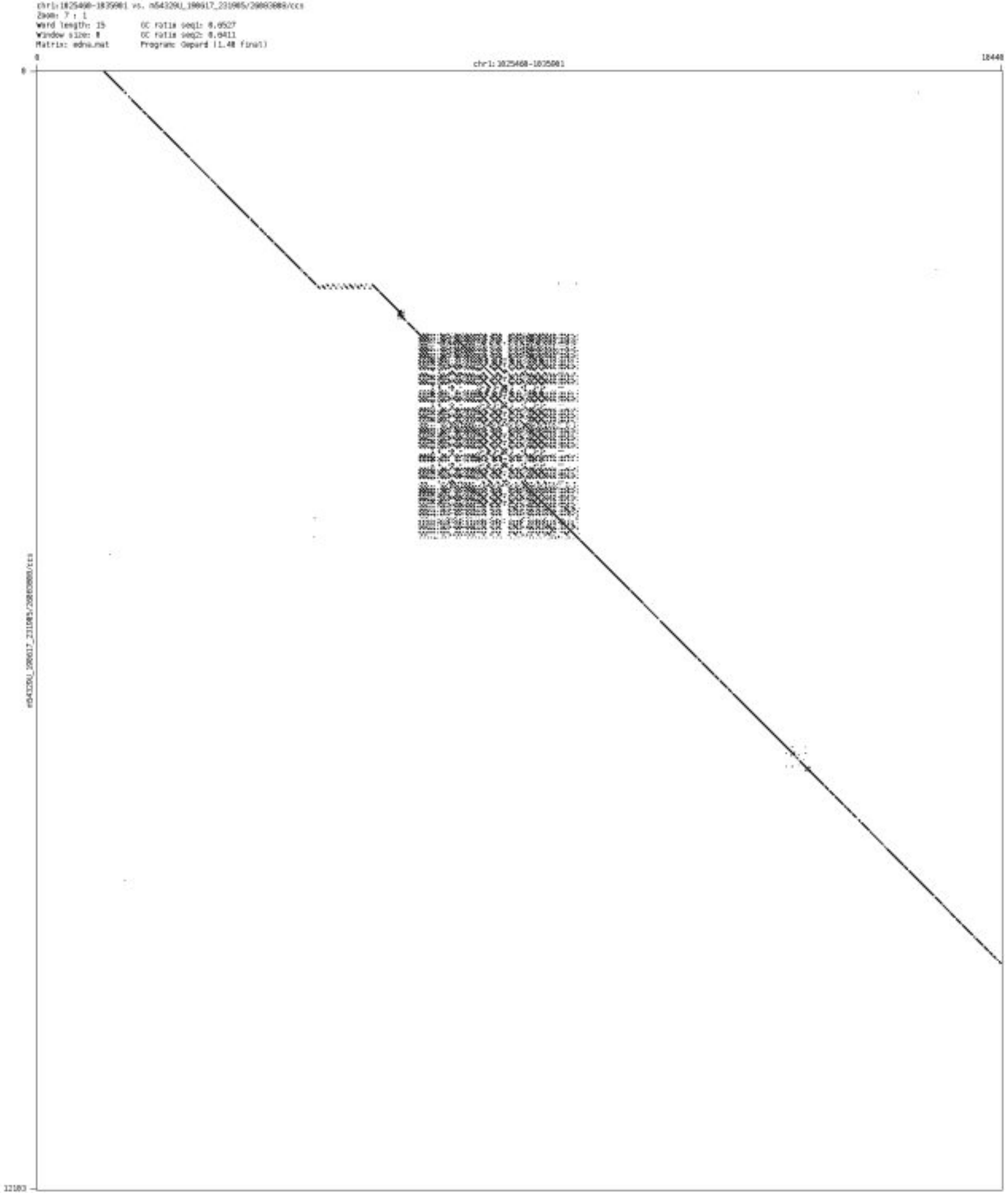

➢ chr1_5160213_5160312

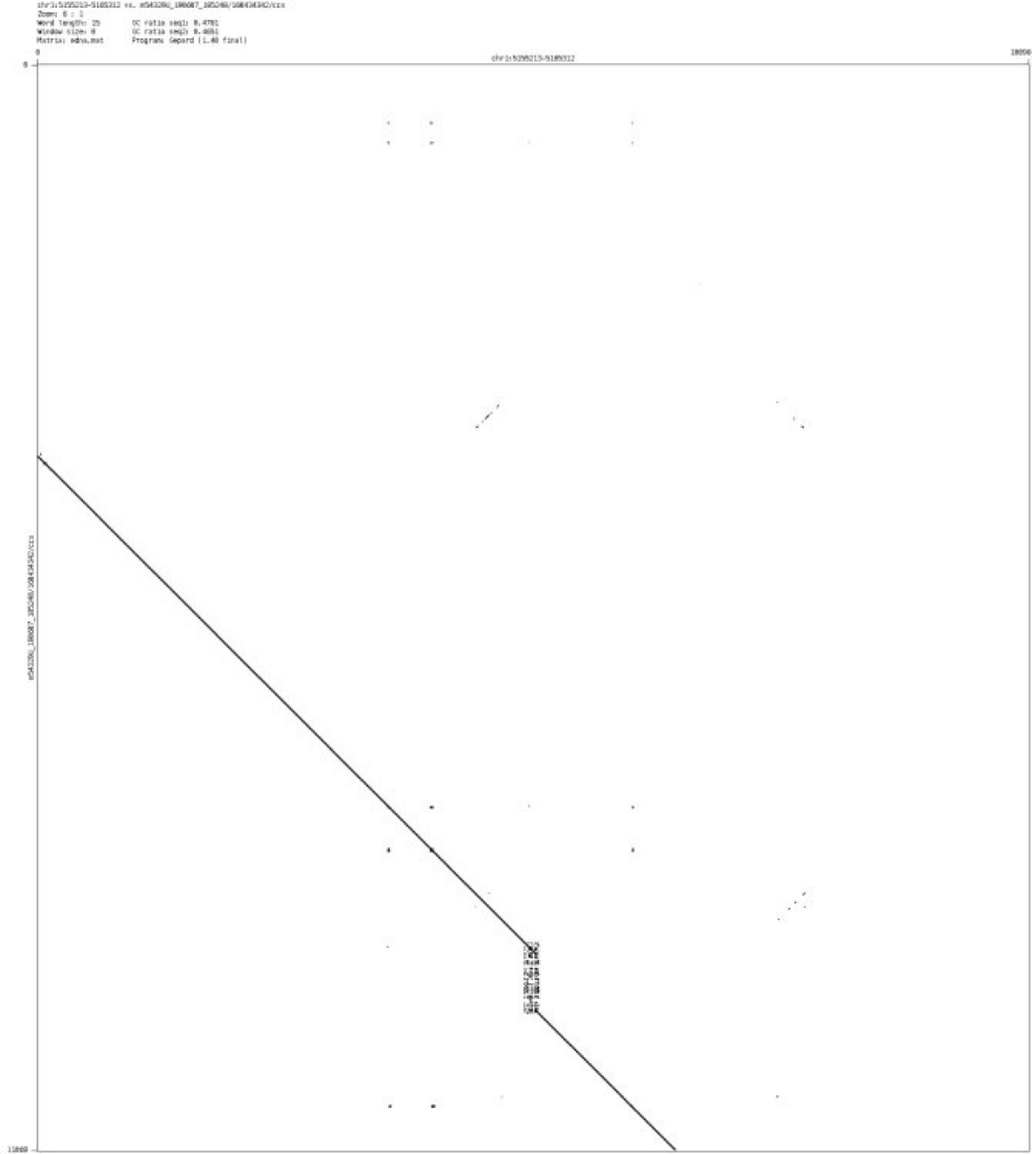

- chr1_9411149_9411257

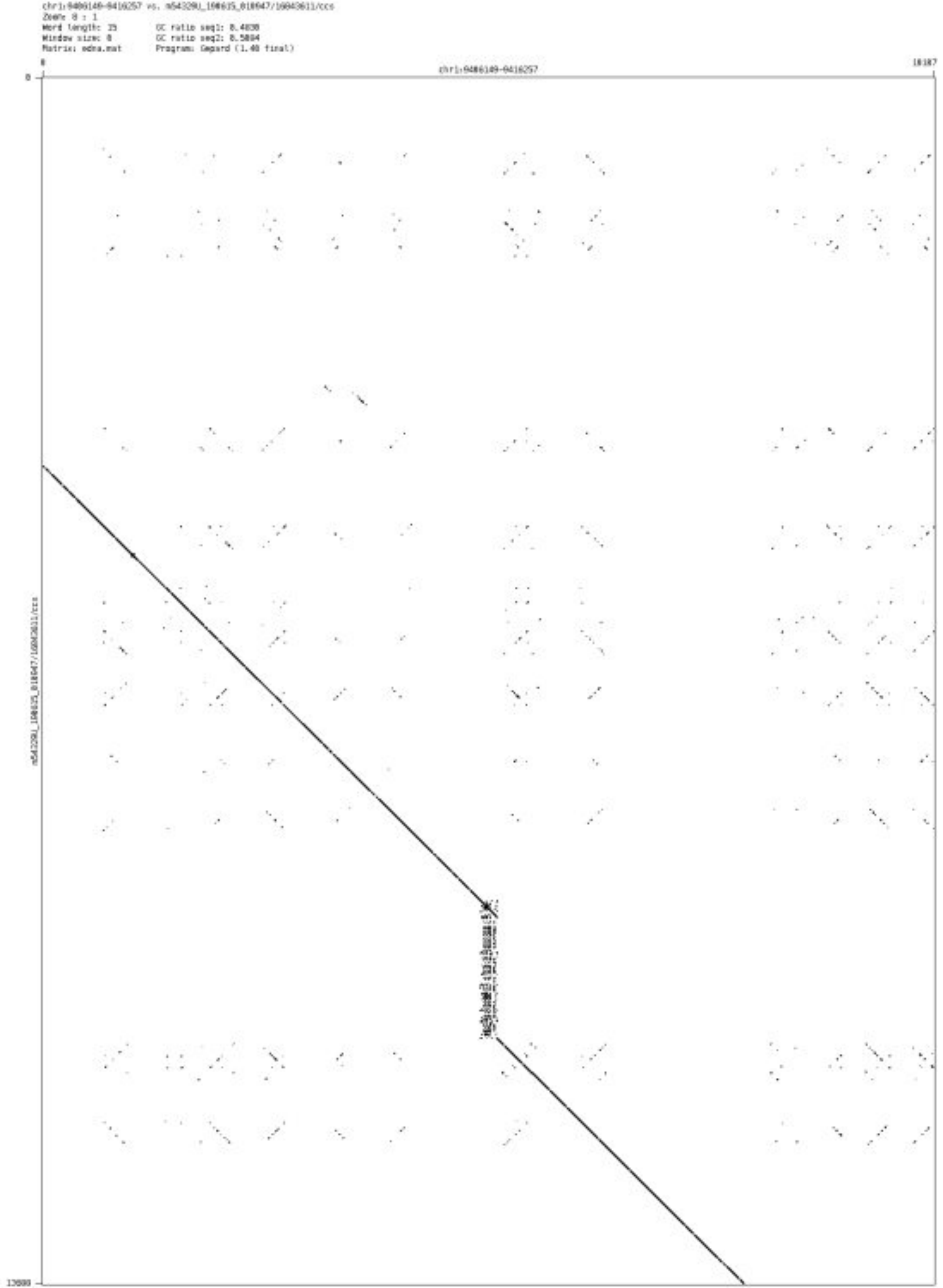

- chr1_18049823_18049961

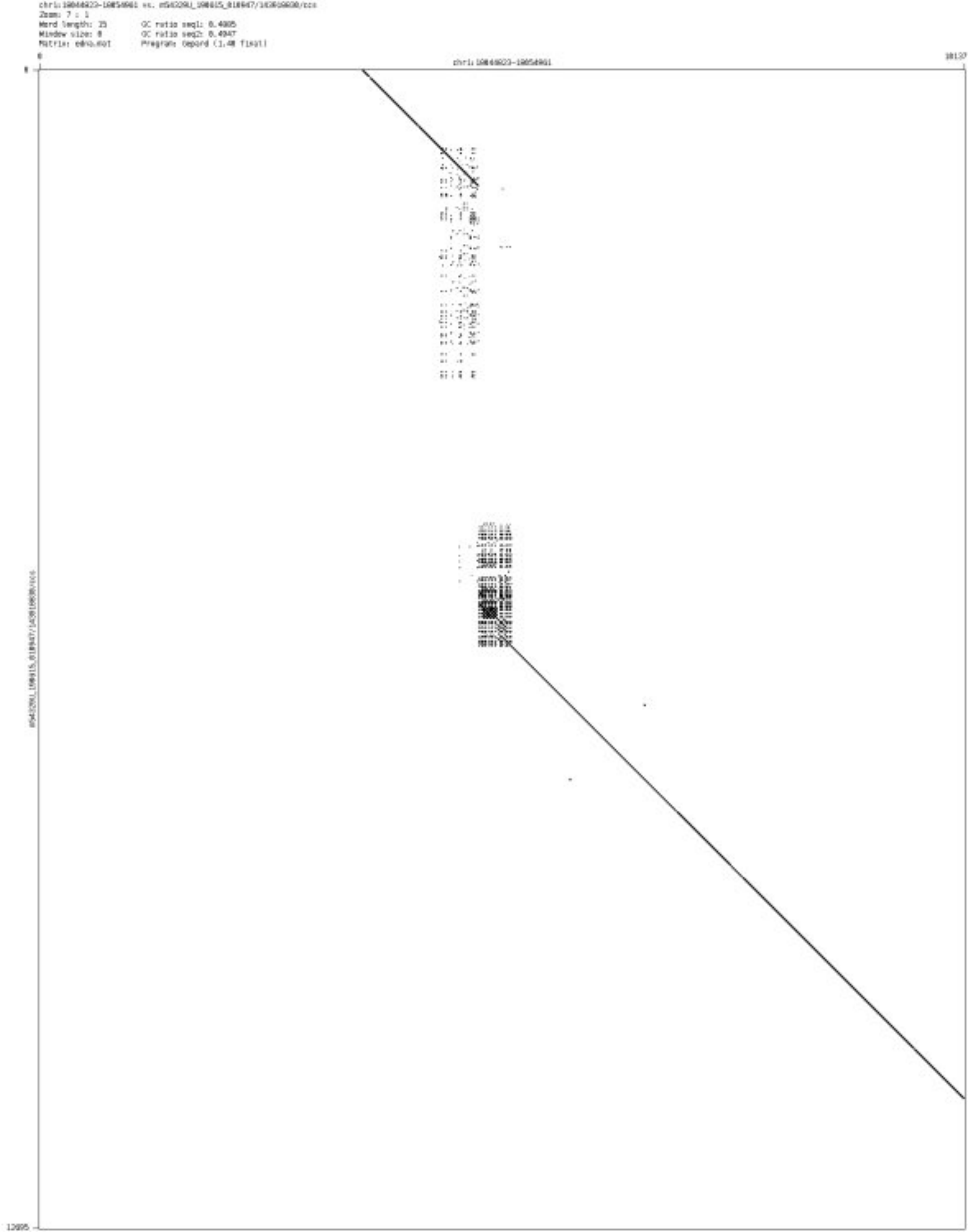

➢ chr1_18925882_18926000

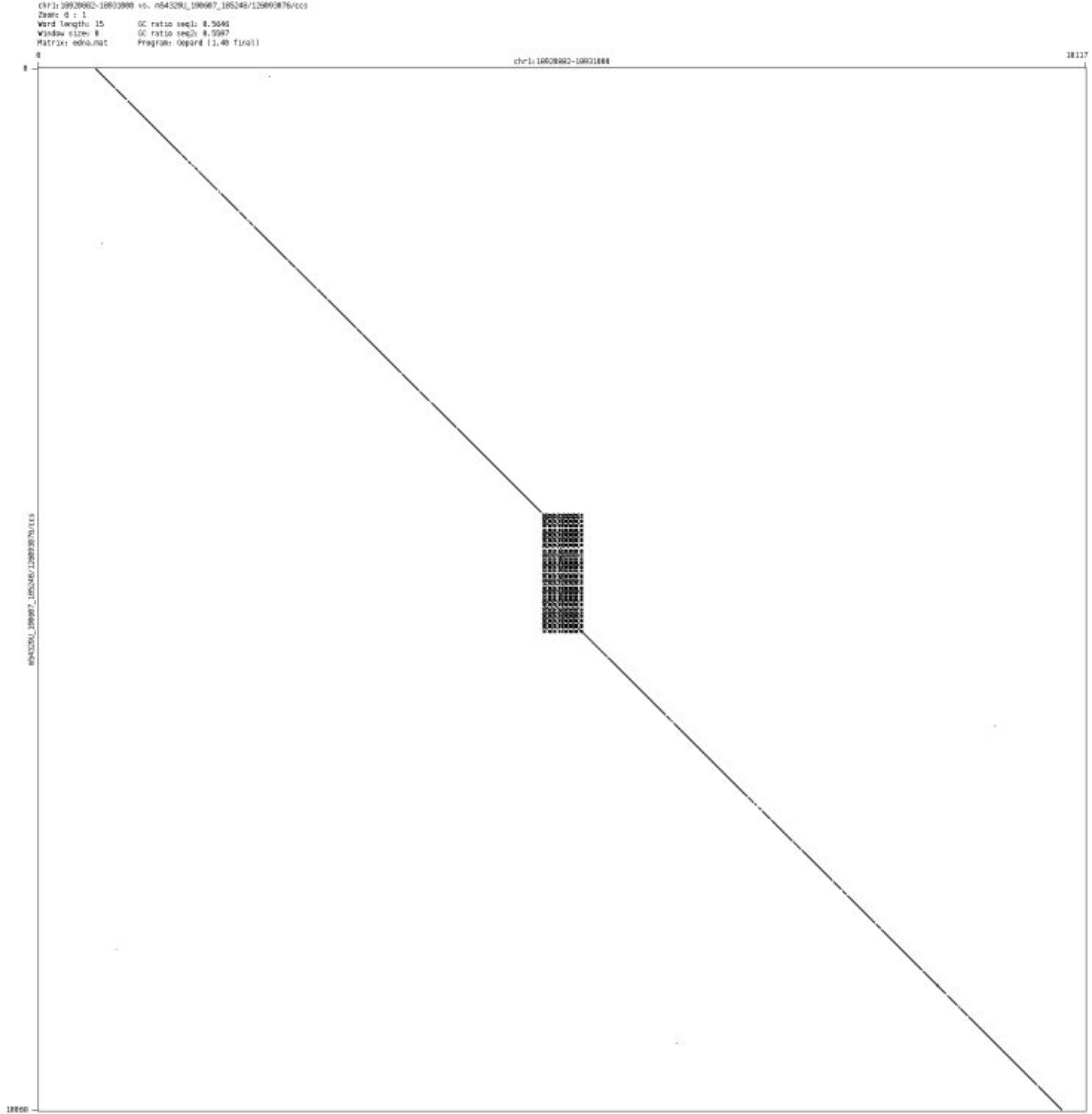

➢ chr1_23392547_23392627

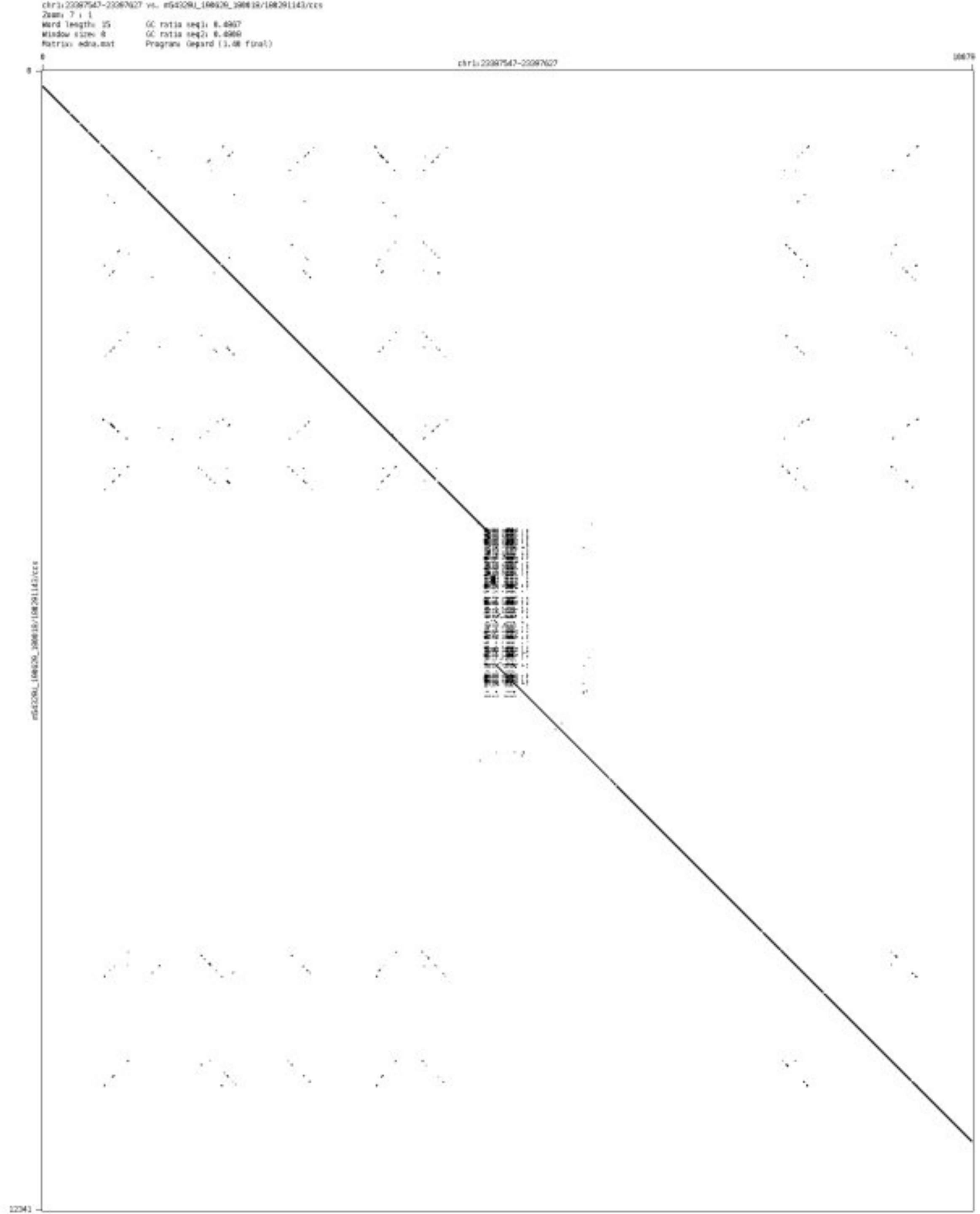

➢ chr1_29771334_29771549

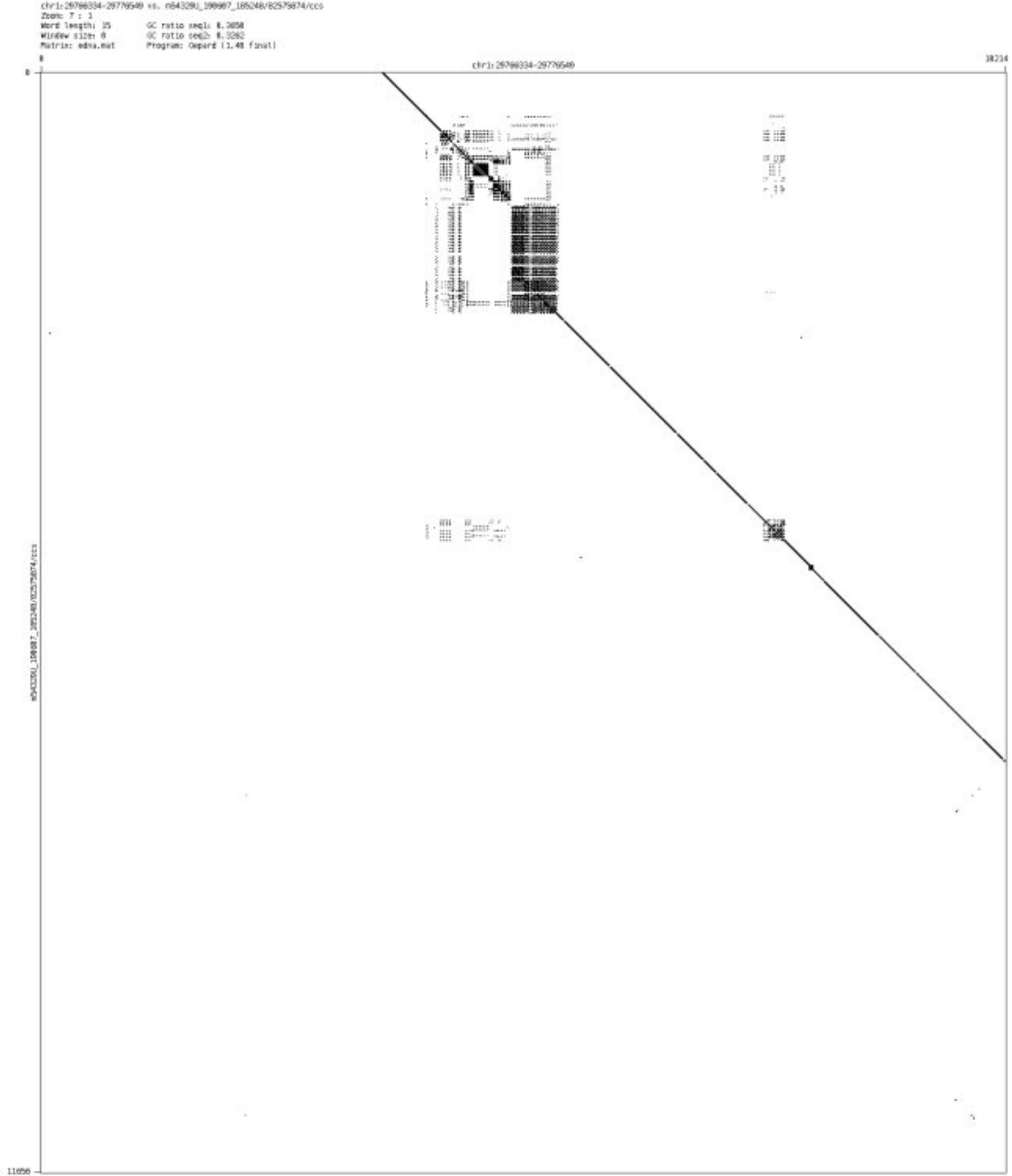

- chr1_31431806_31431981

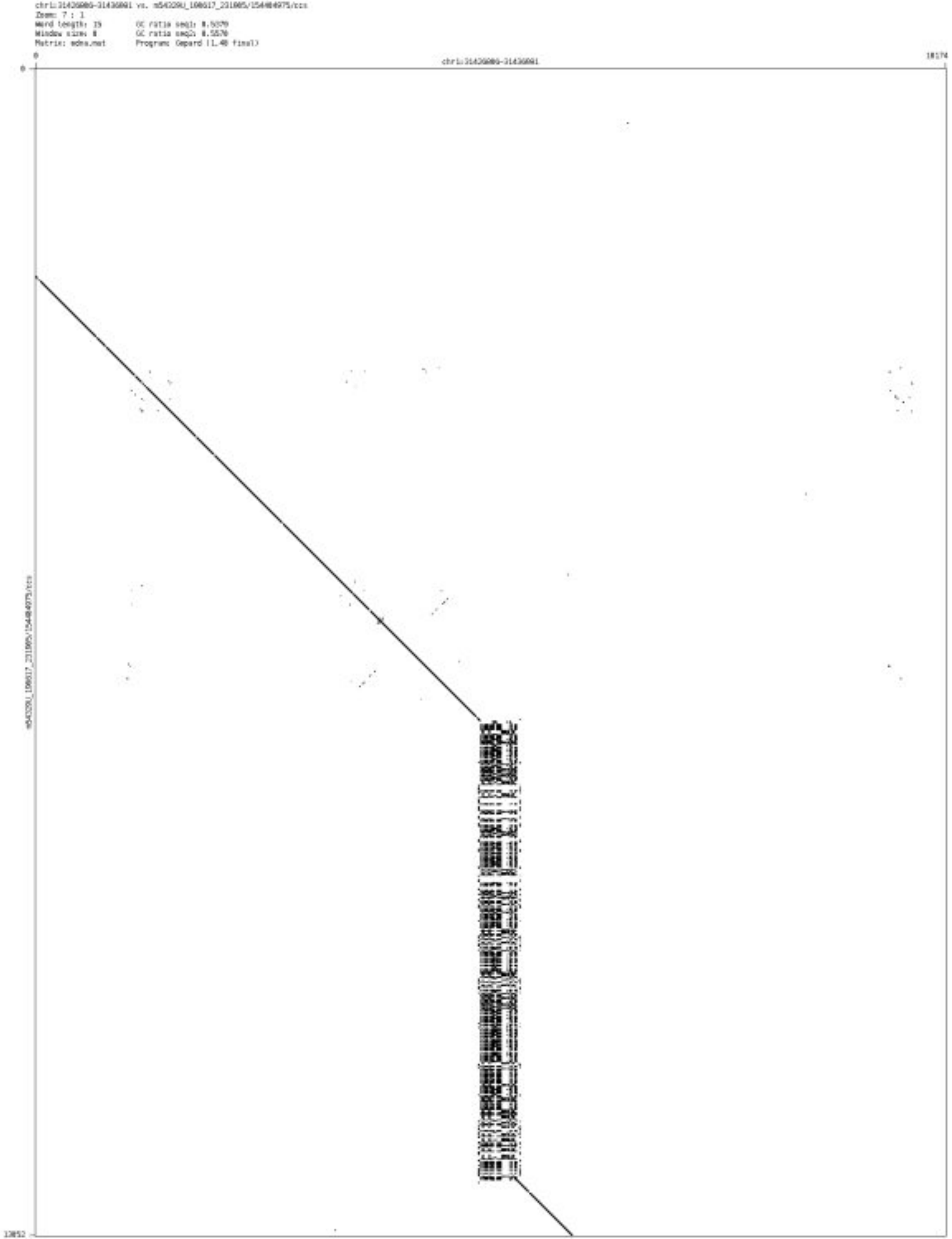

- chr1_118318808_118318888

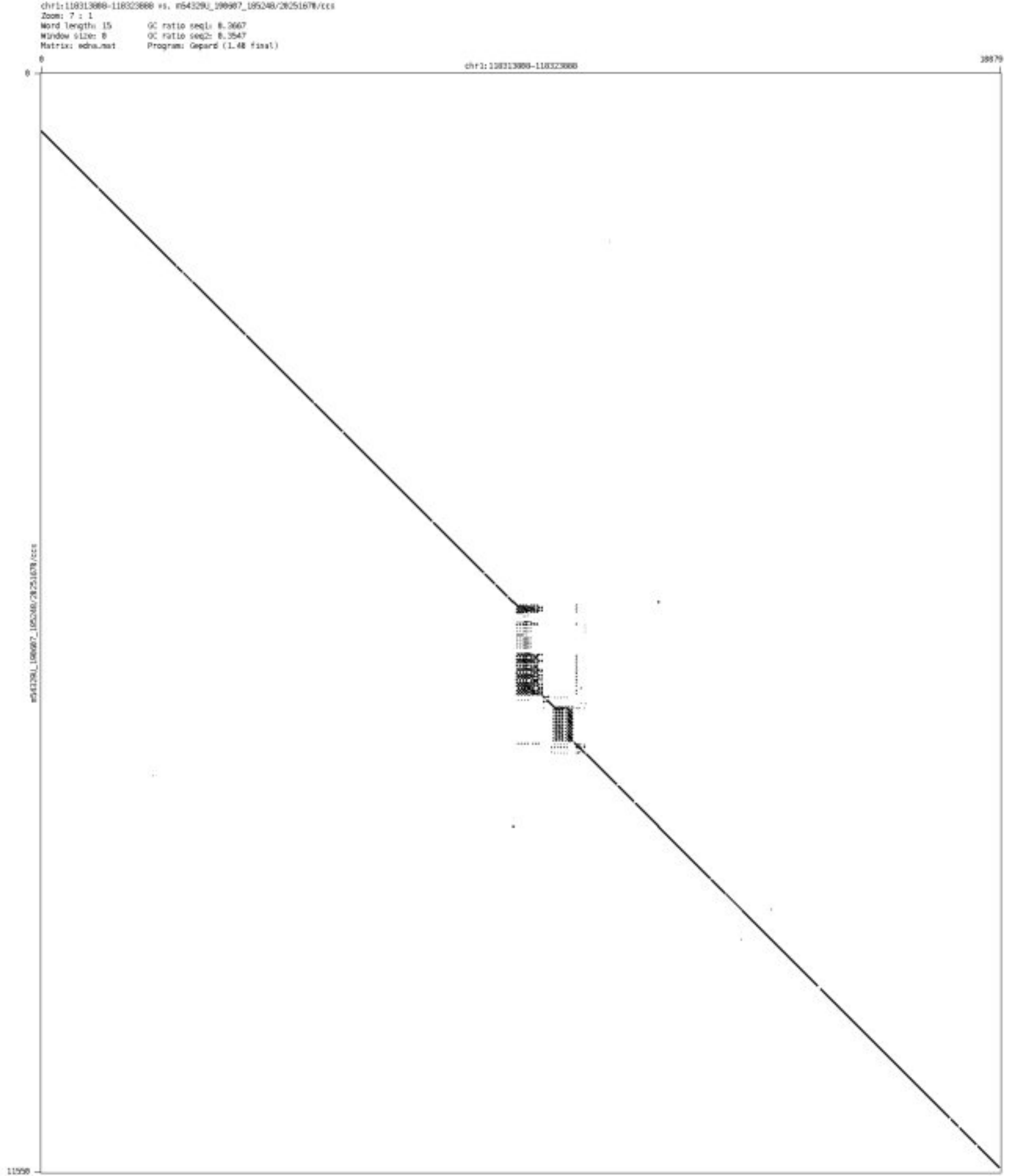

➢ chr1_144102505_144106880

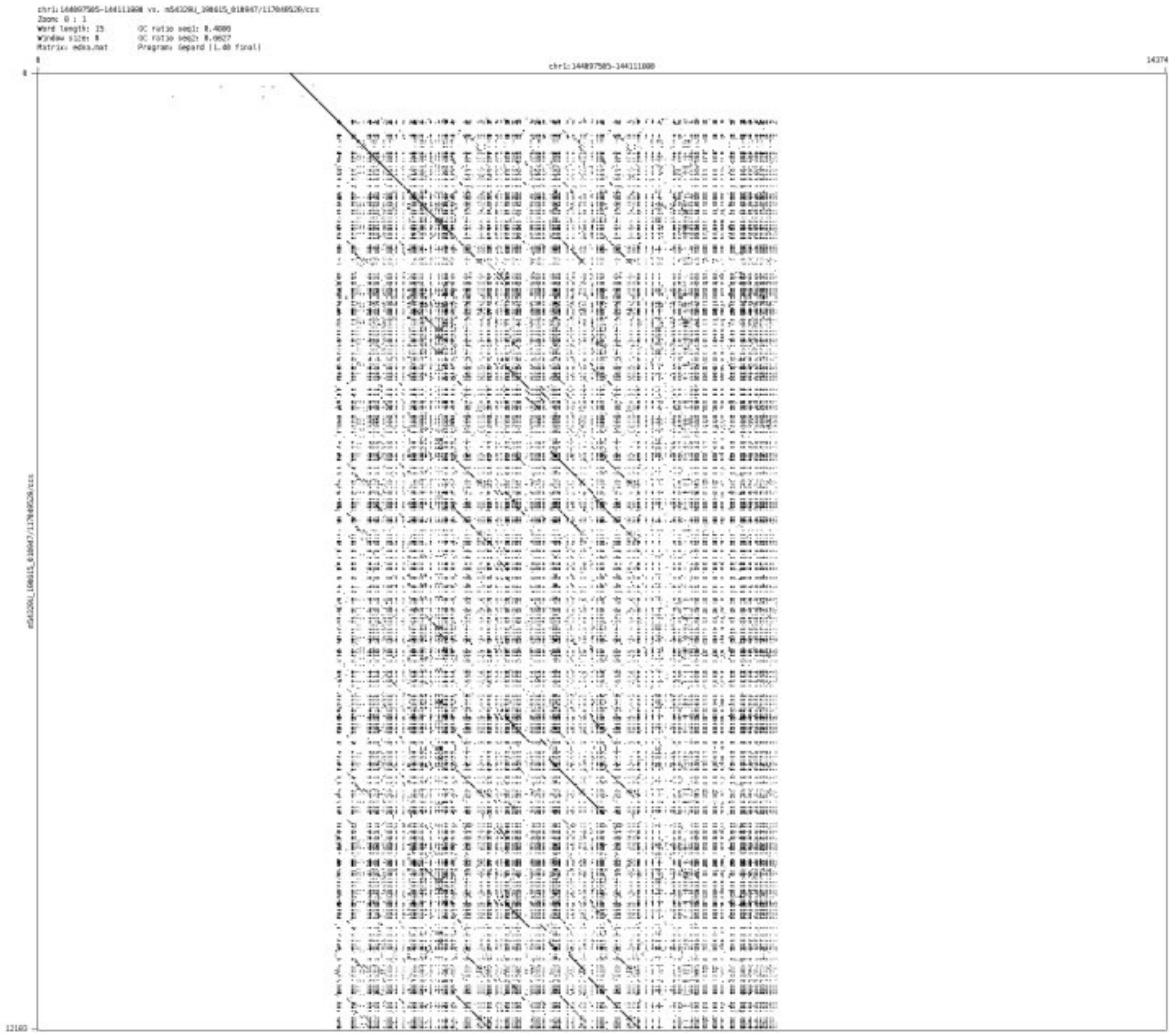

➢ chr1_172694311_172694449

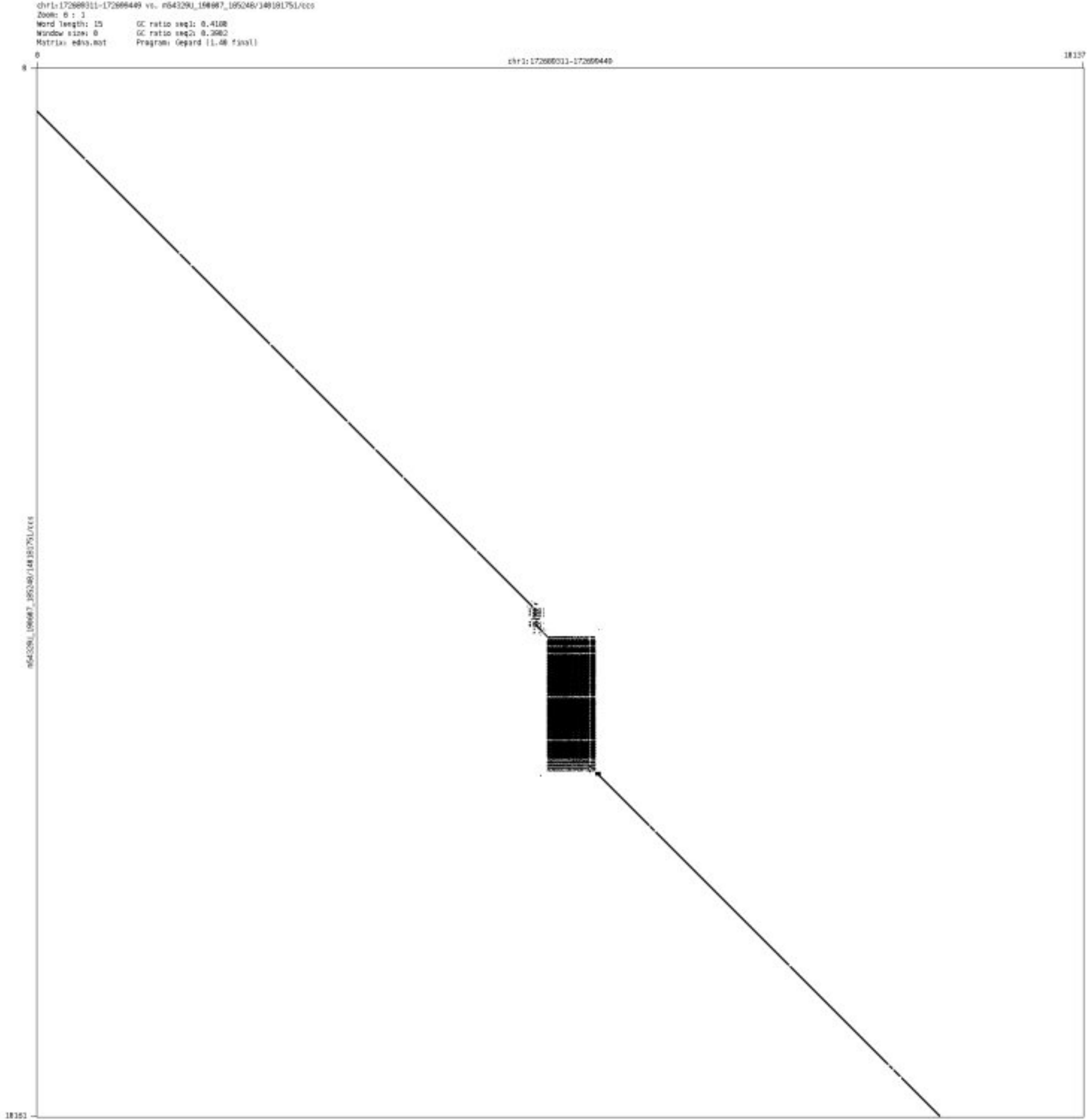

- chr1_202625025_202625085

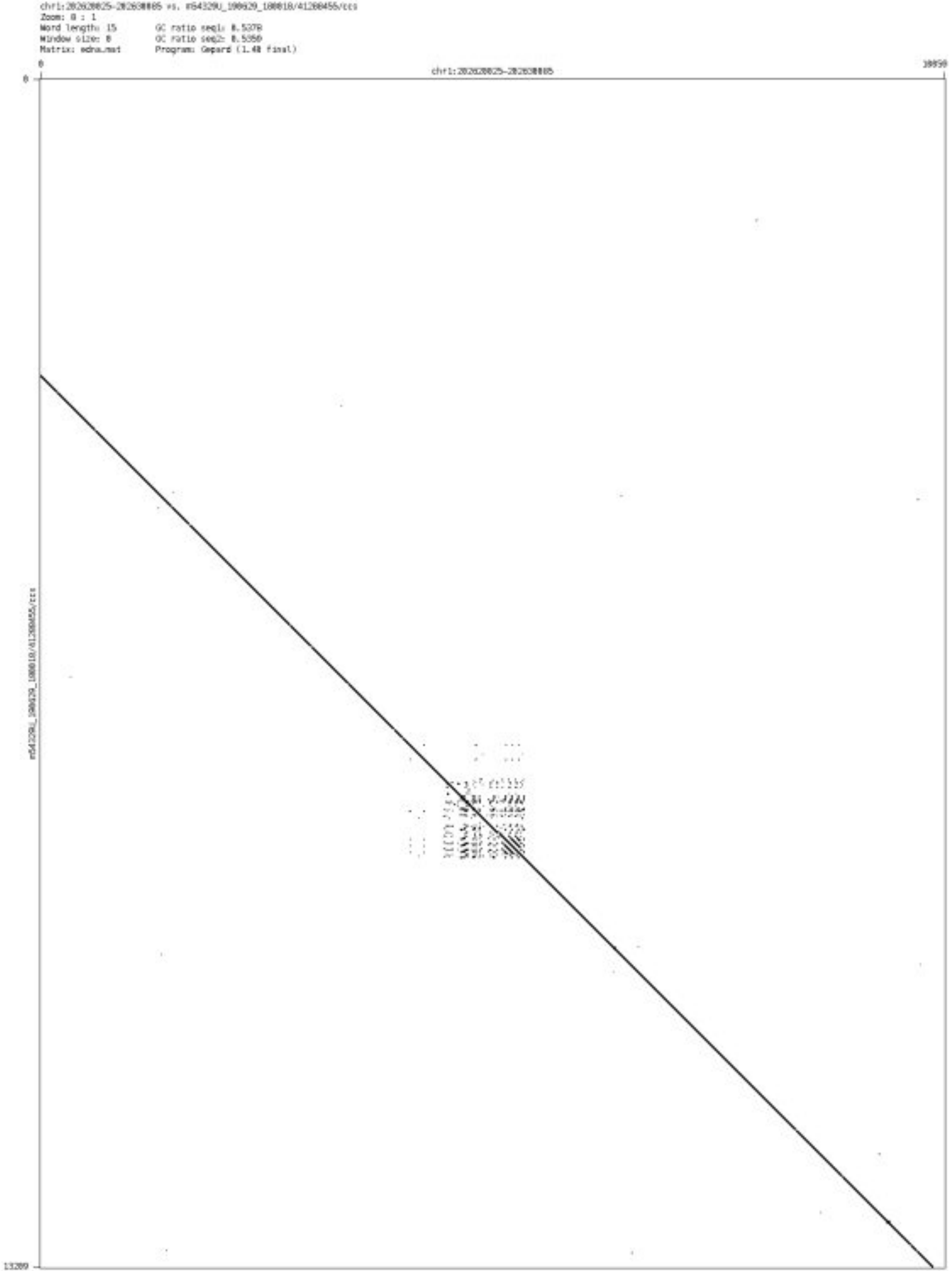

- ➢ chr2_1106532_1106732

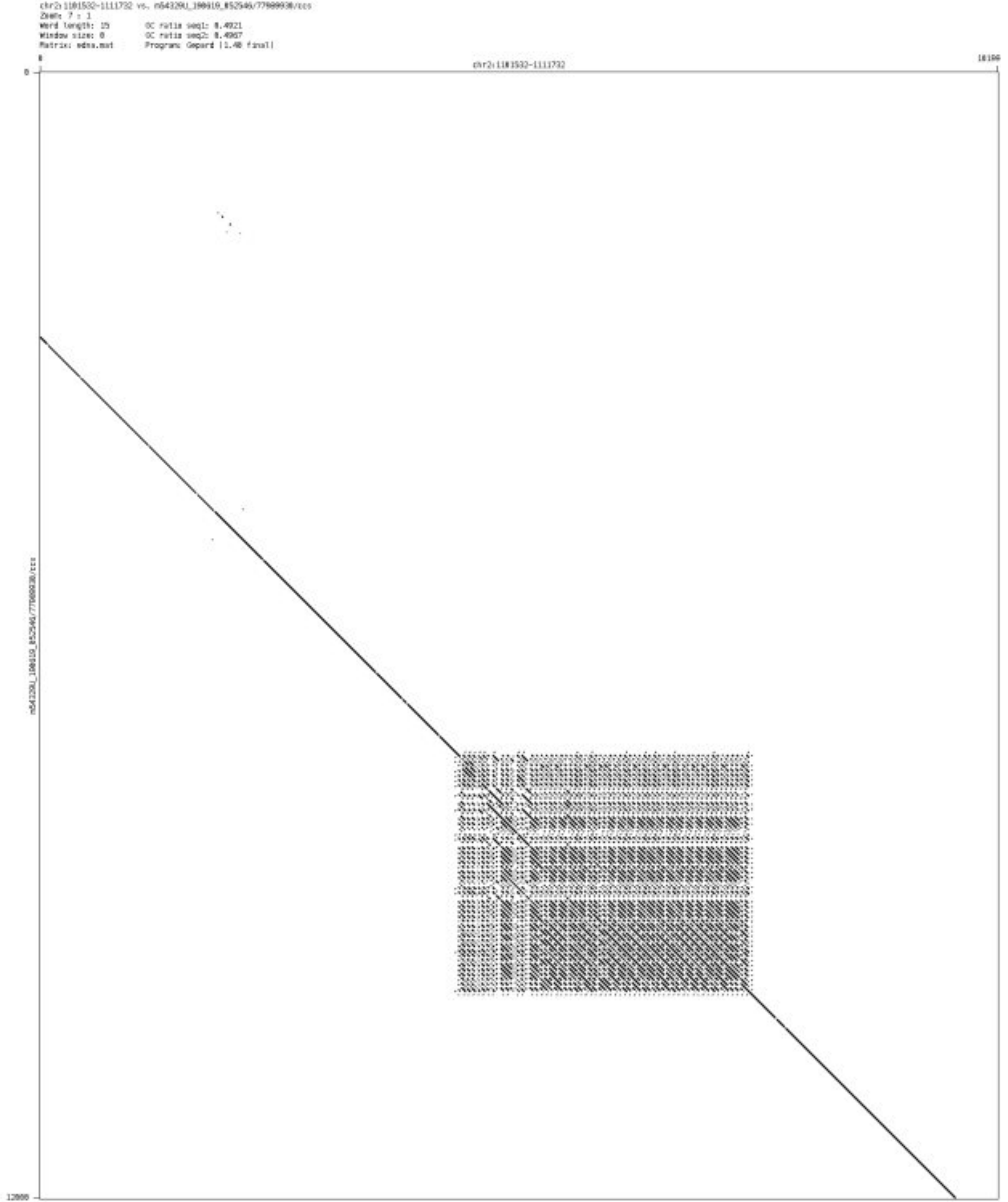

➢ chr2_1750529_1750651

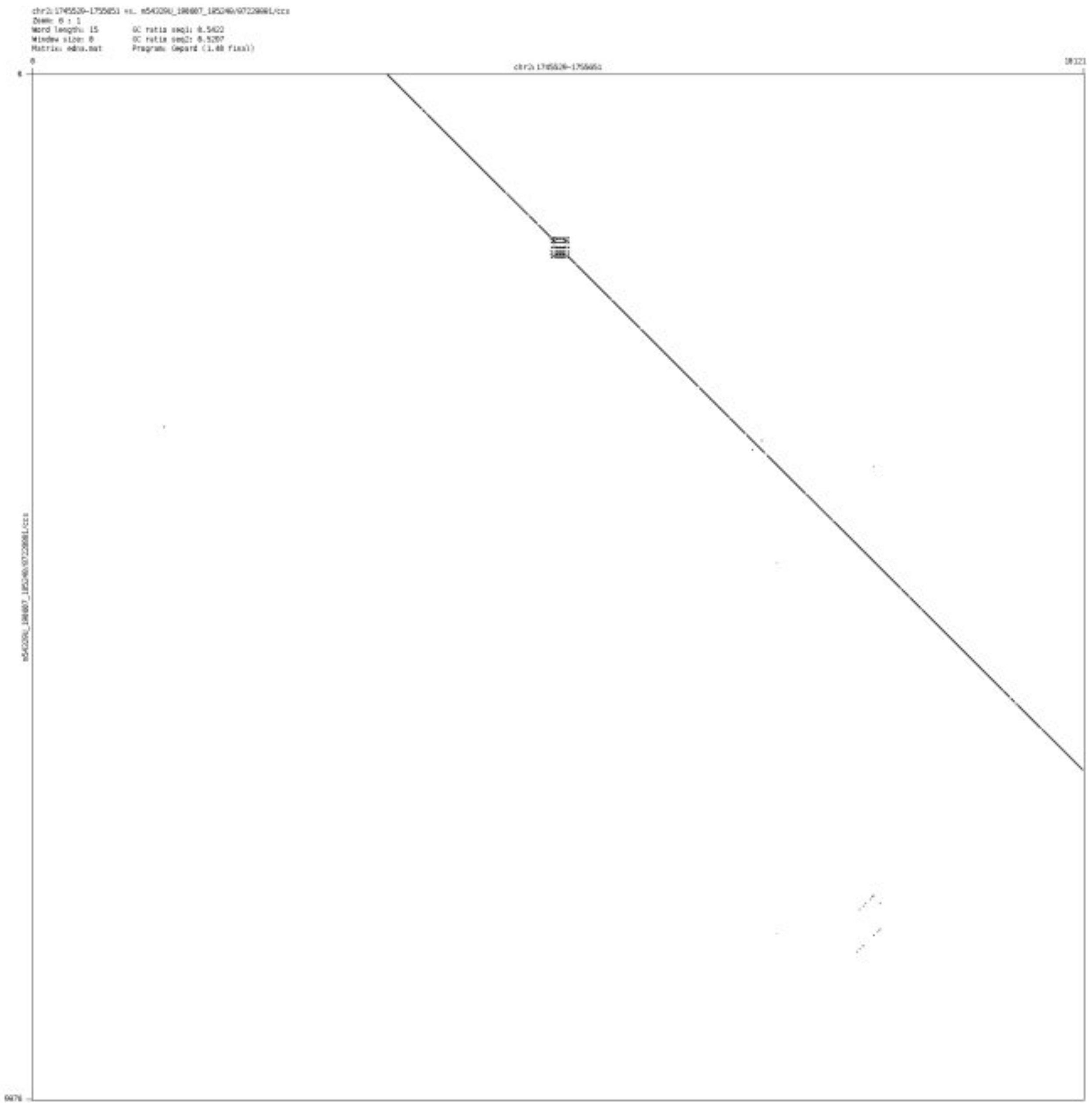

- chr2_10399253_10399369

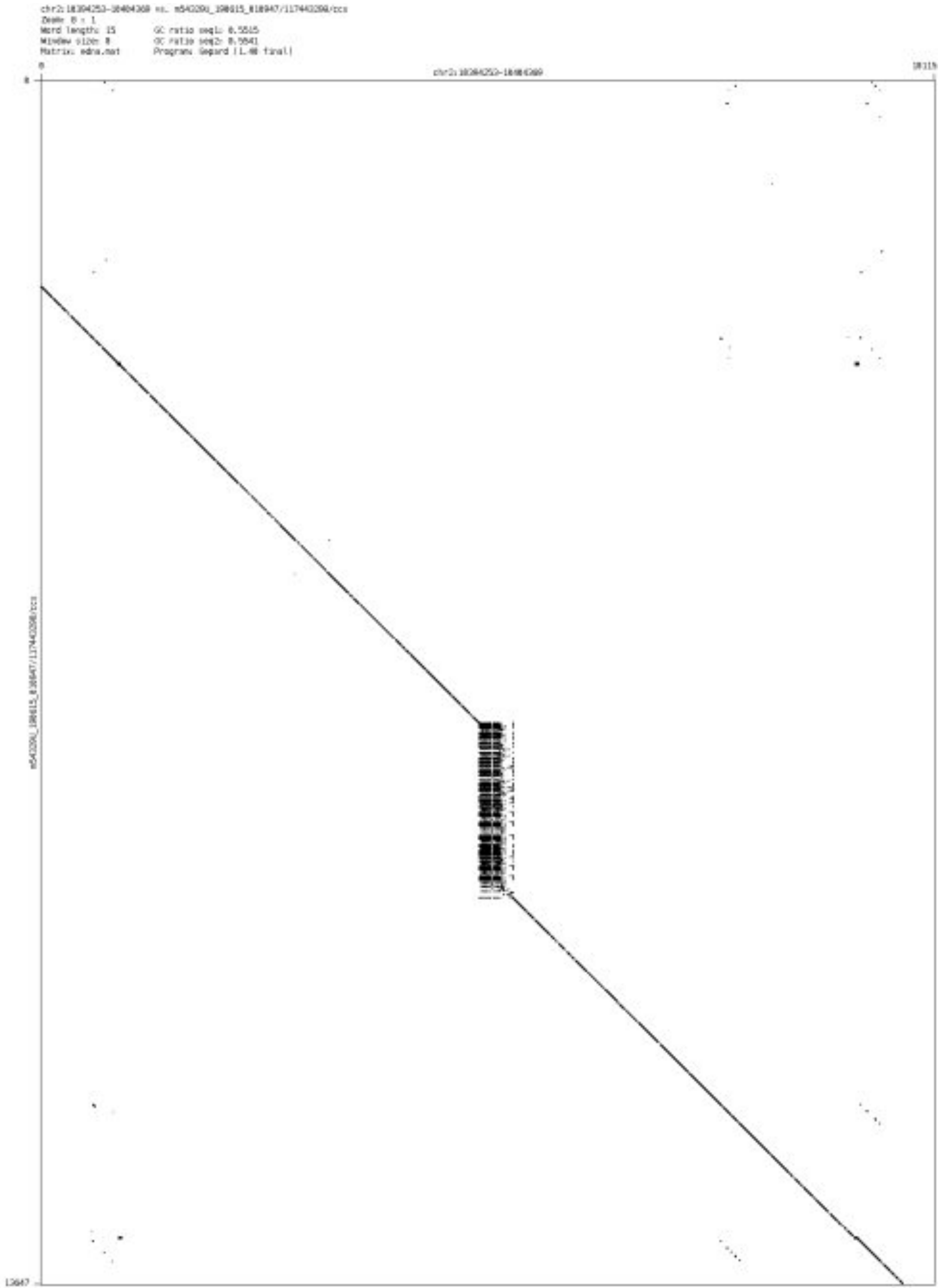

- chr2_11113534_11113628

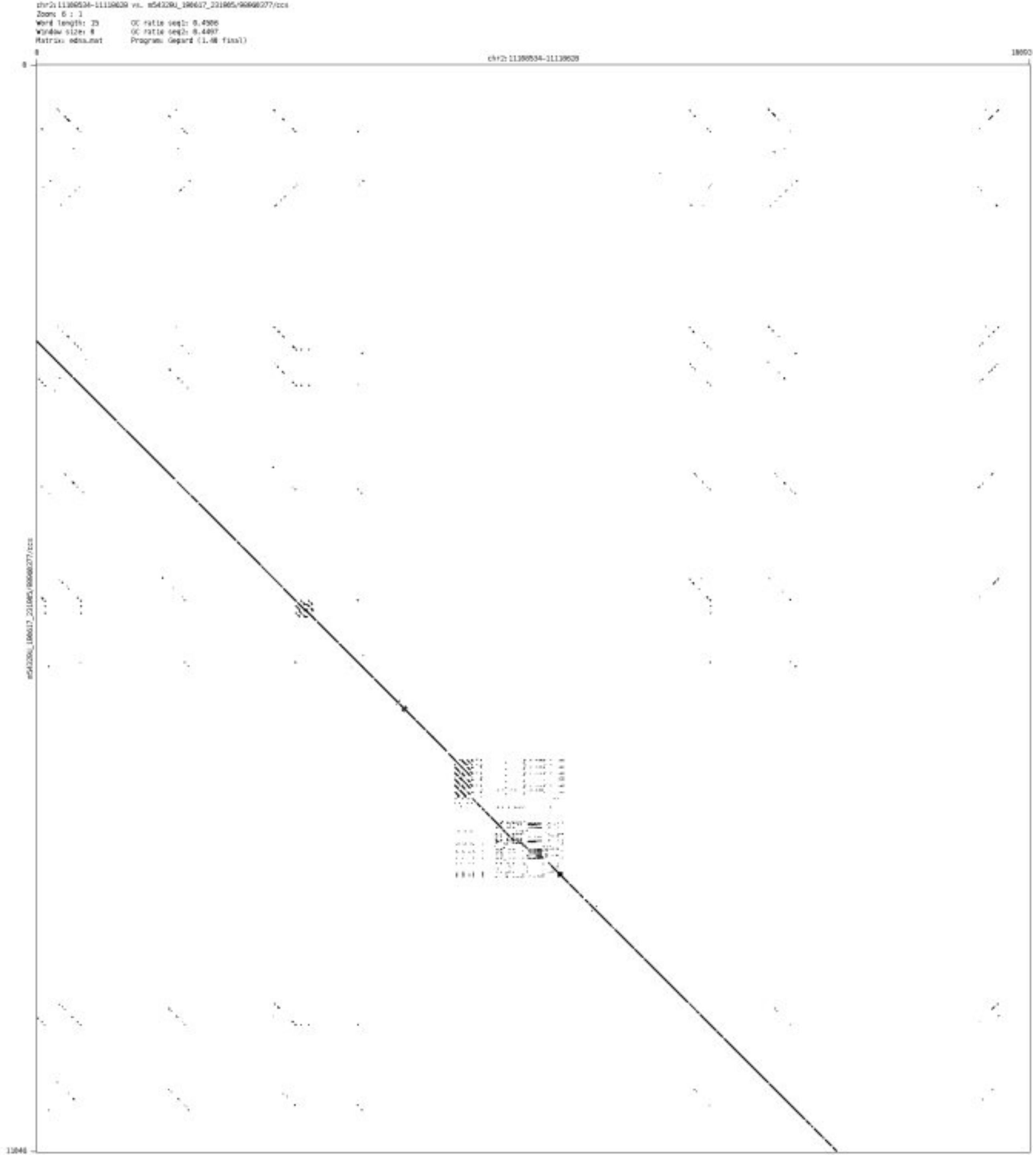

- ➢ chr2_13288455_13288641

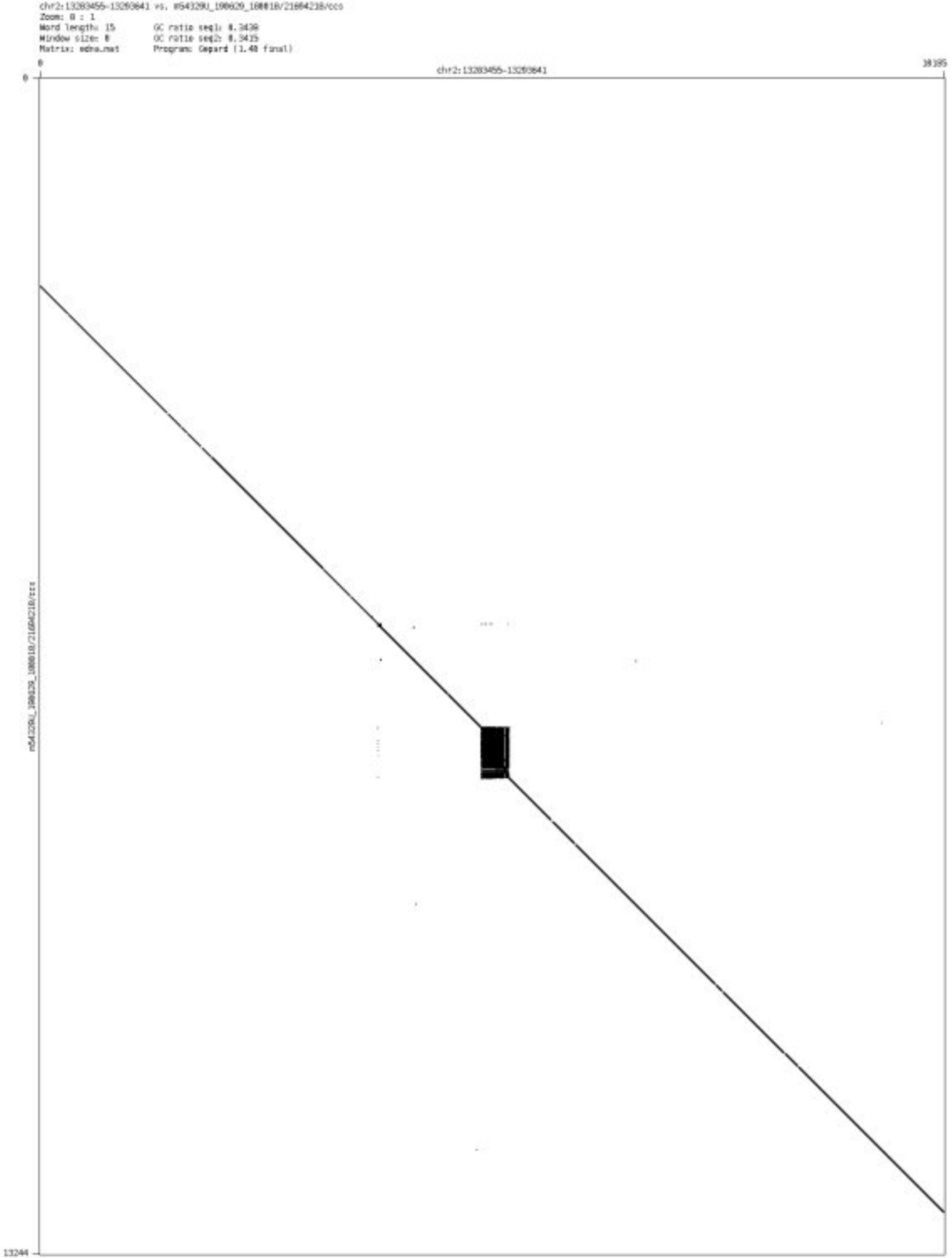

- chr2_36184130_36184311

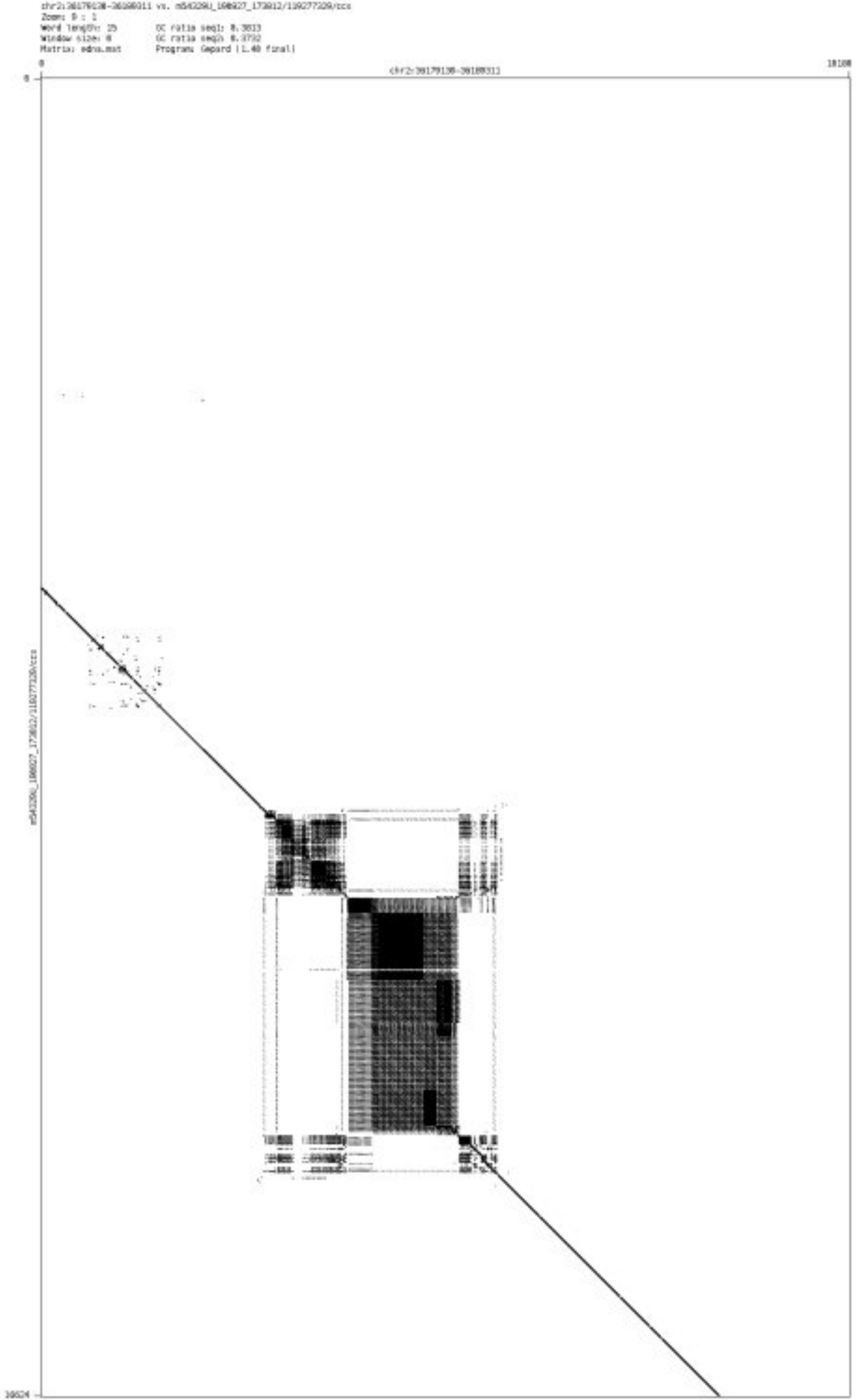

➢ chr2_60467724_60468110

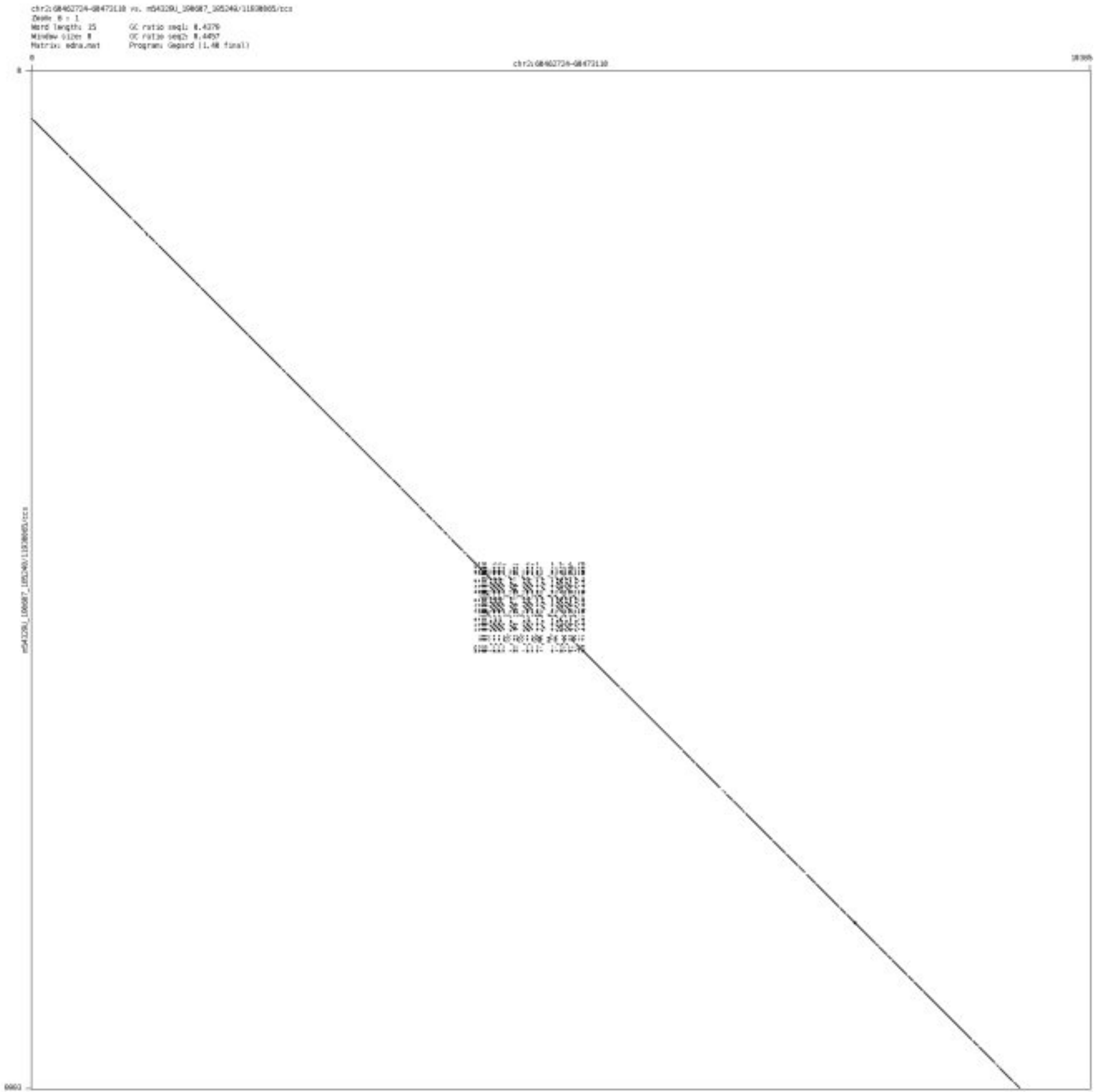

- chr2_64376016_64376165

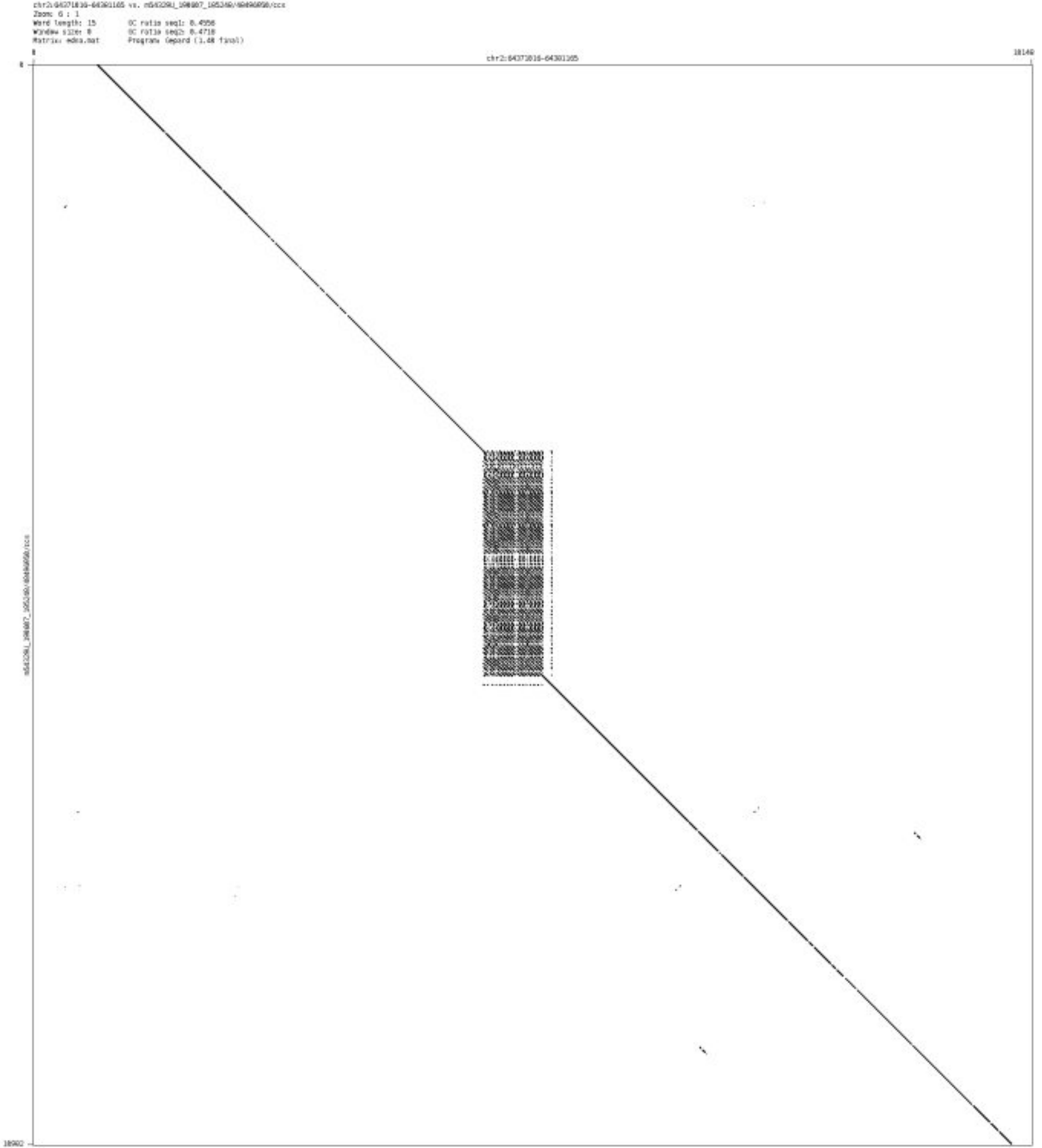

➢ chr2_83342525_83342578

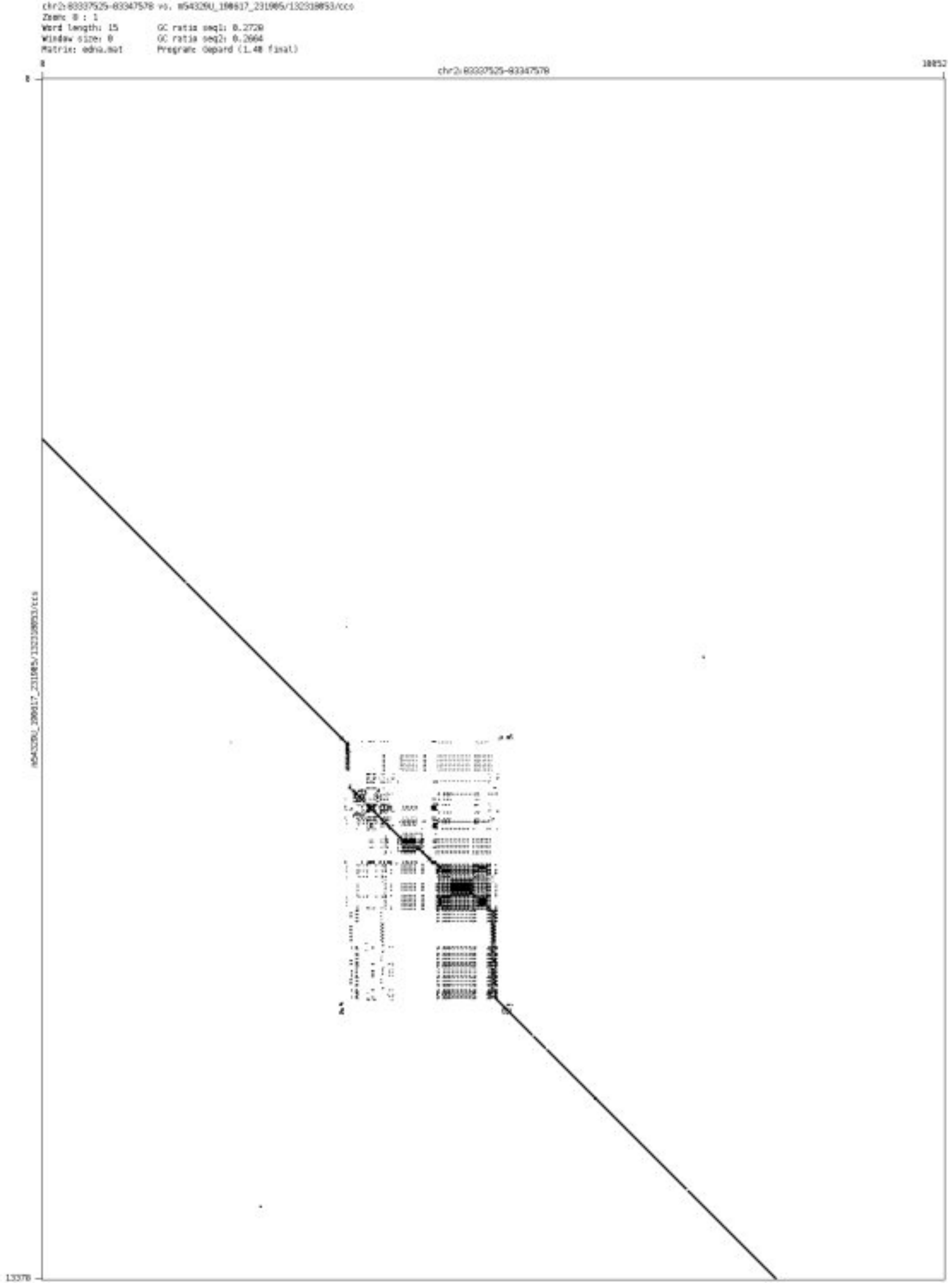

➢ chr2_119444283_119445298

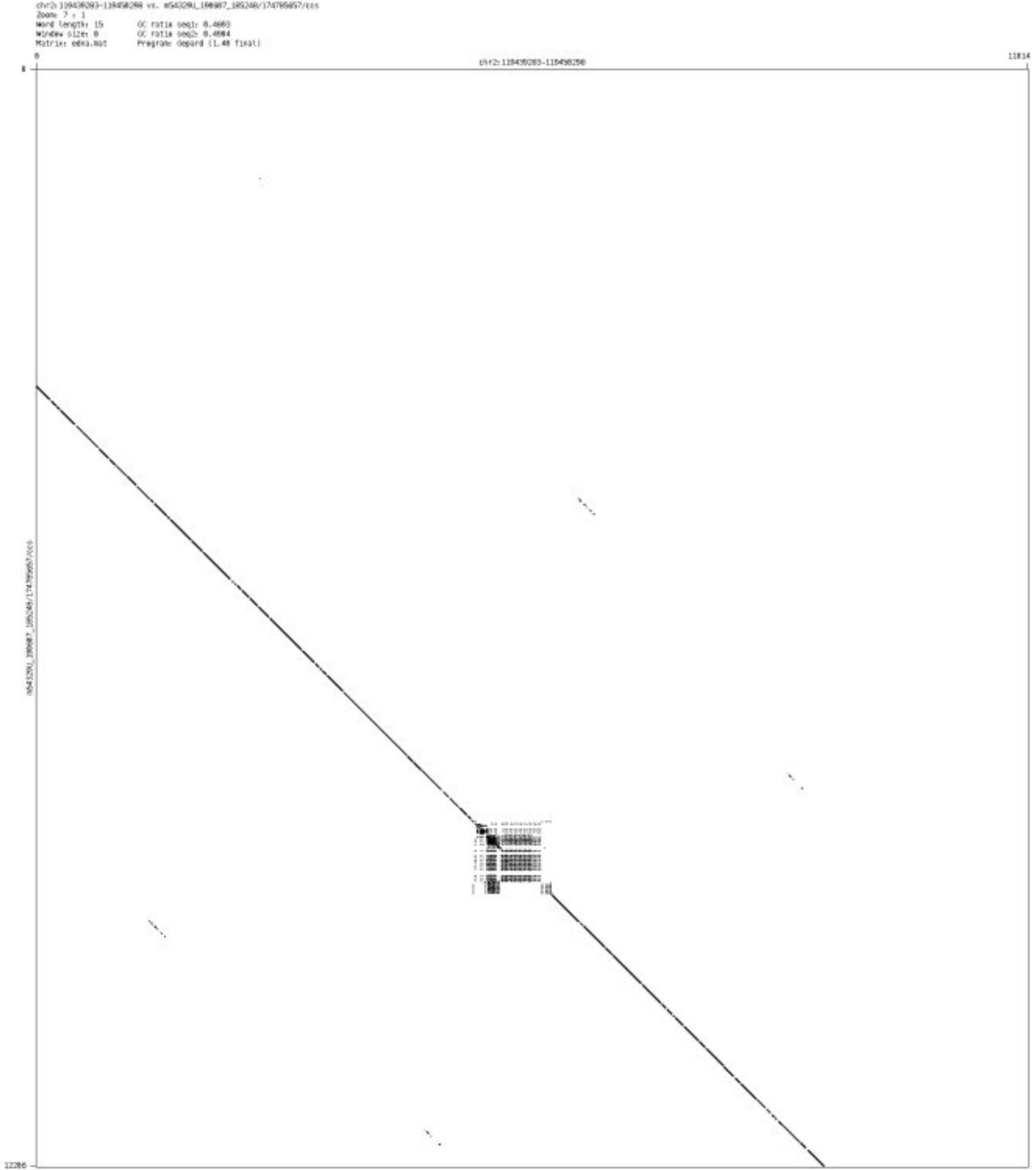

- chr2_217805487_217805794

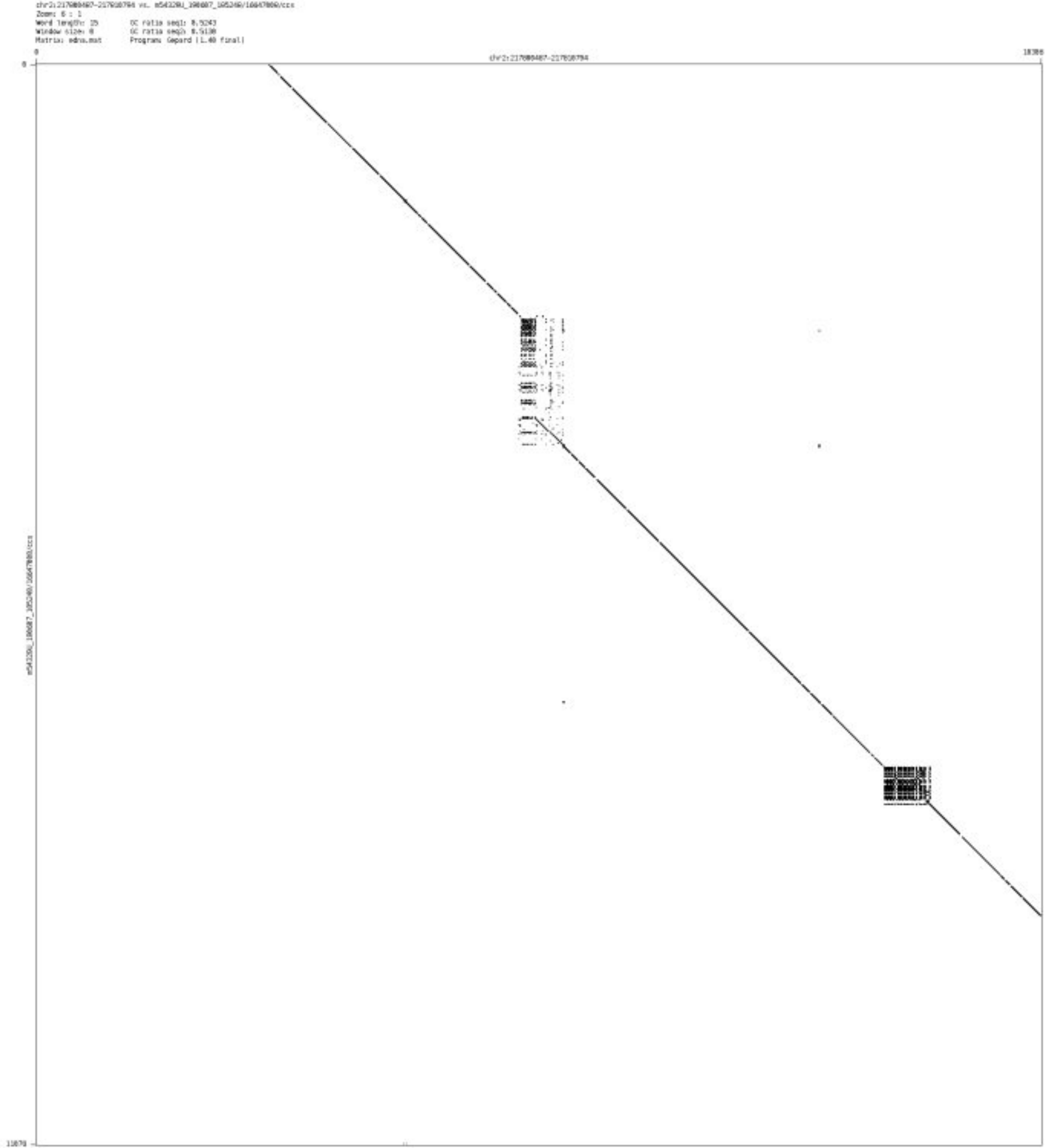

- chr2_232995035_232995577

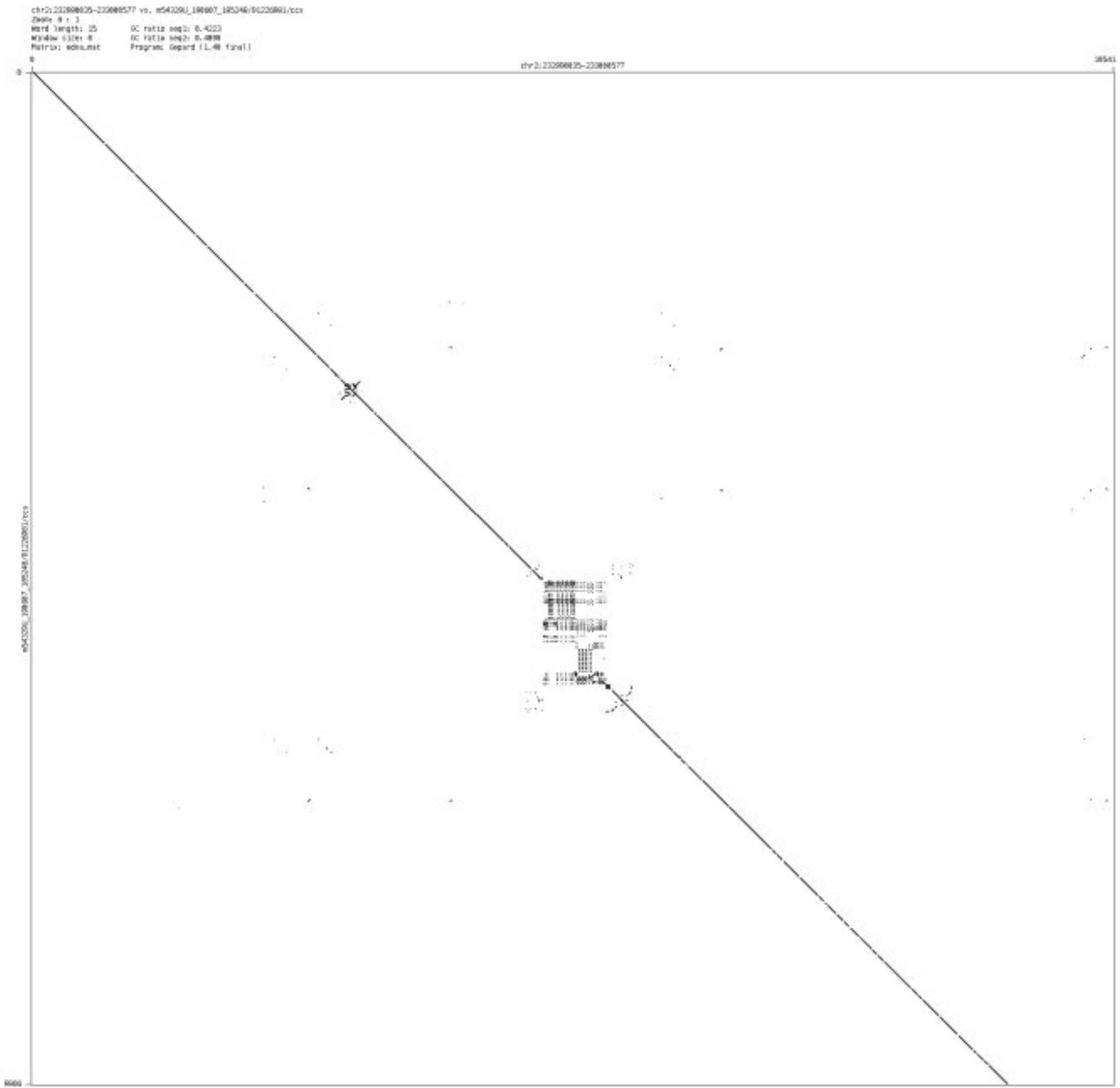

➢ chr2_240158693_240158783

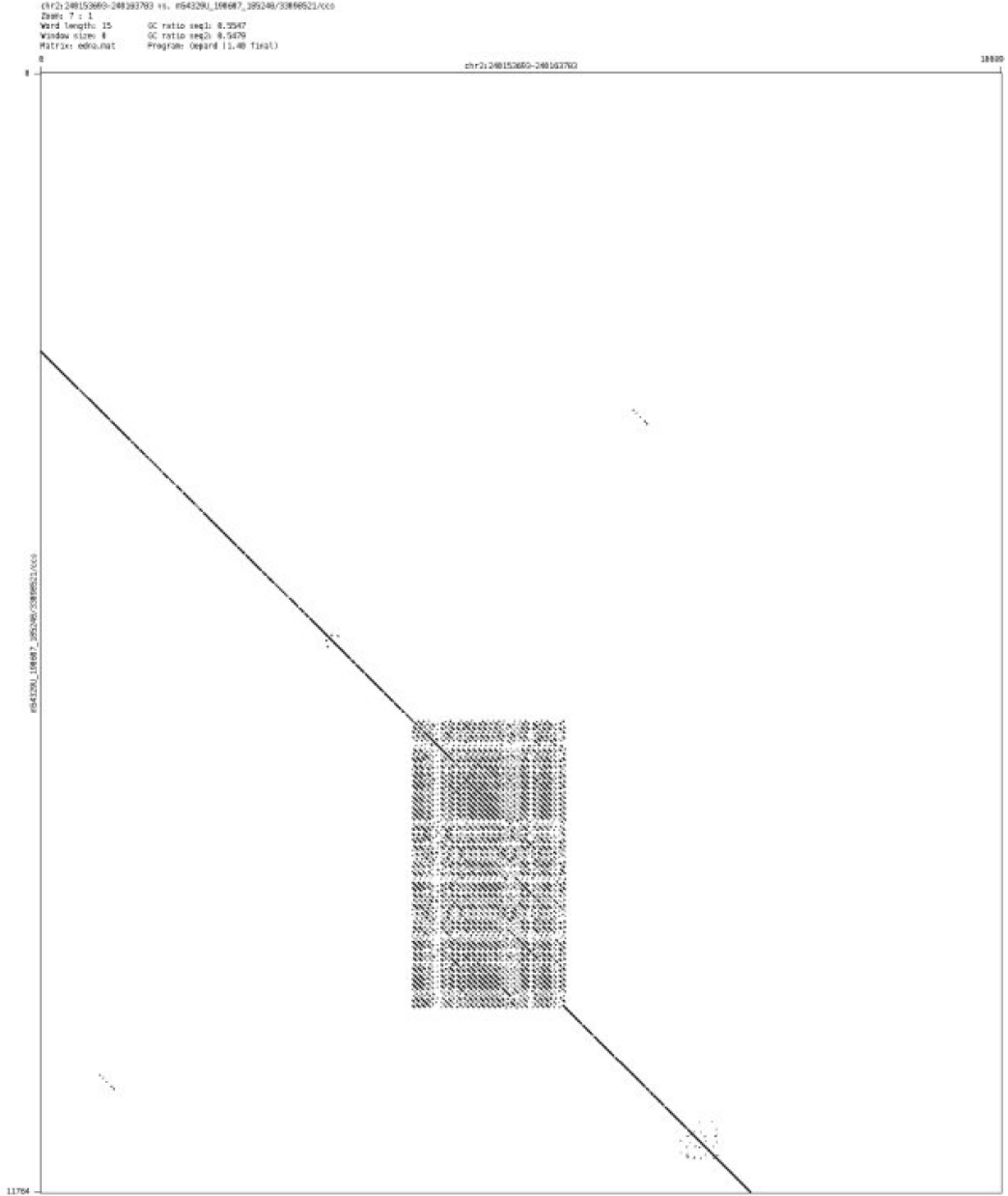

- chr2_240872289_240872460

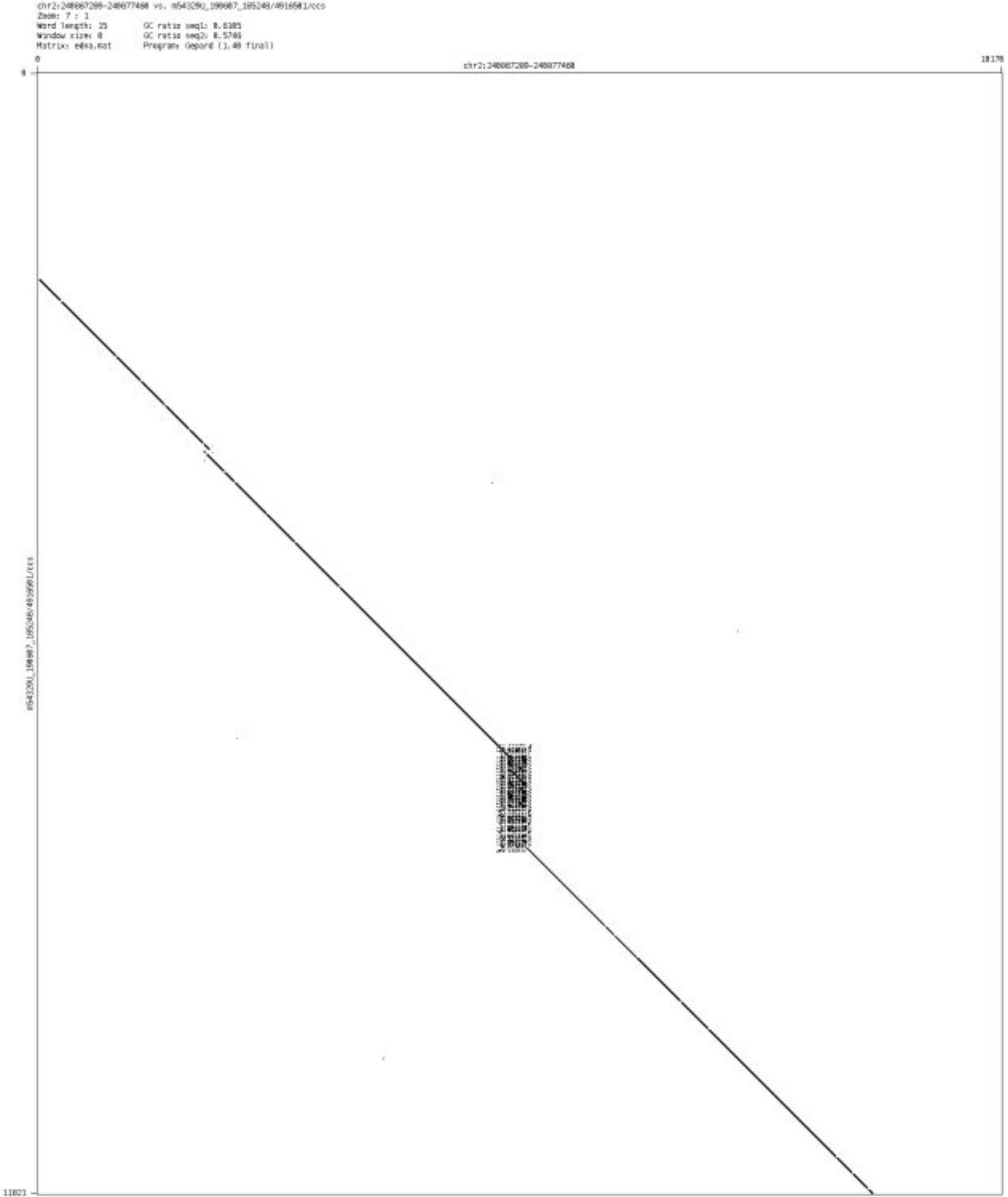

➢ chr3_1707476_1708557

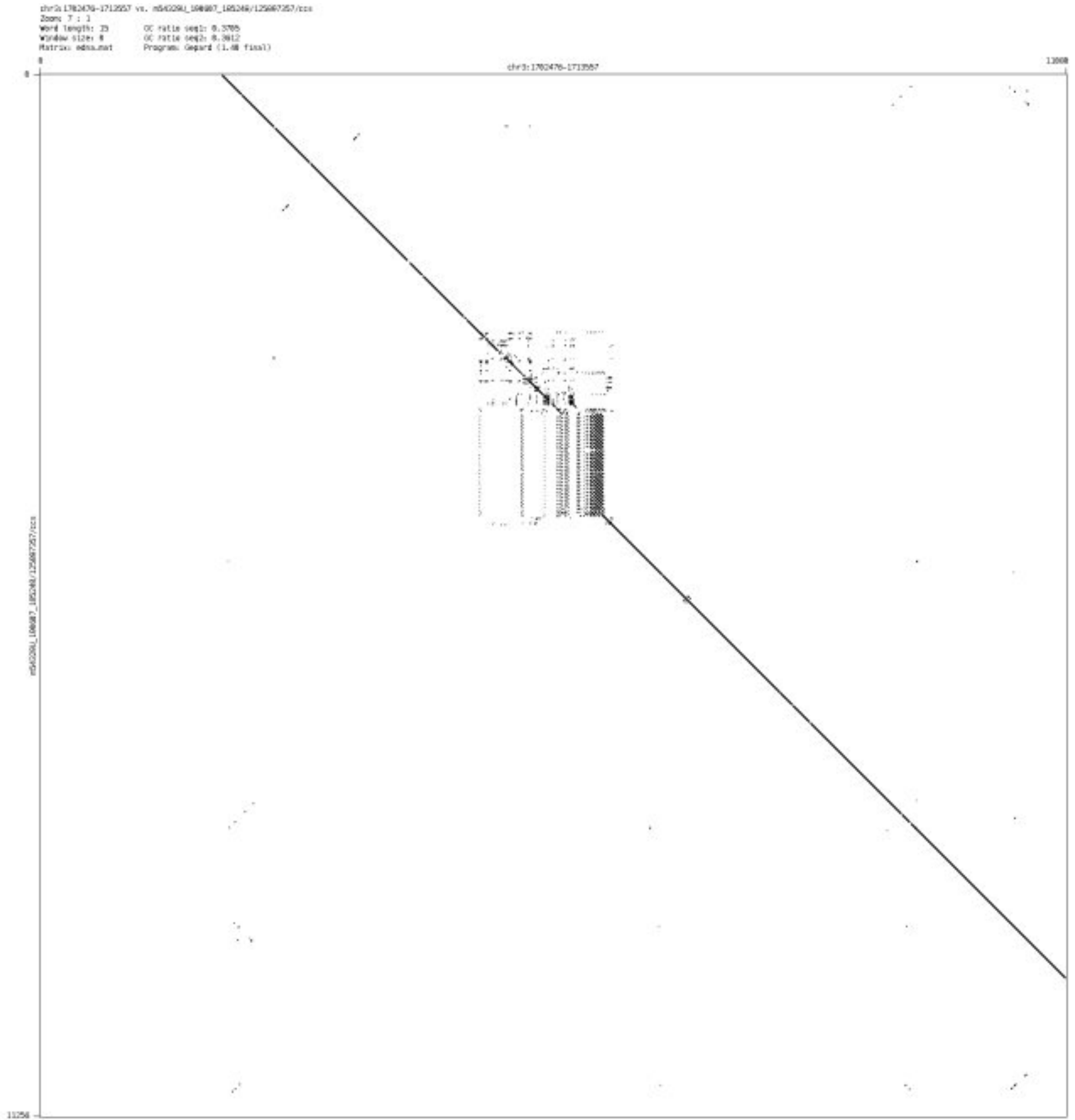

- chr3_38084451_38084565

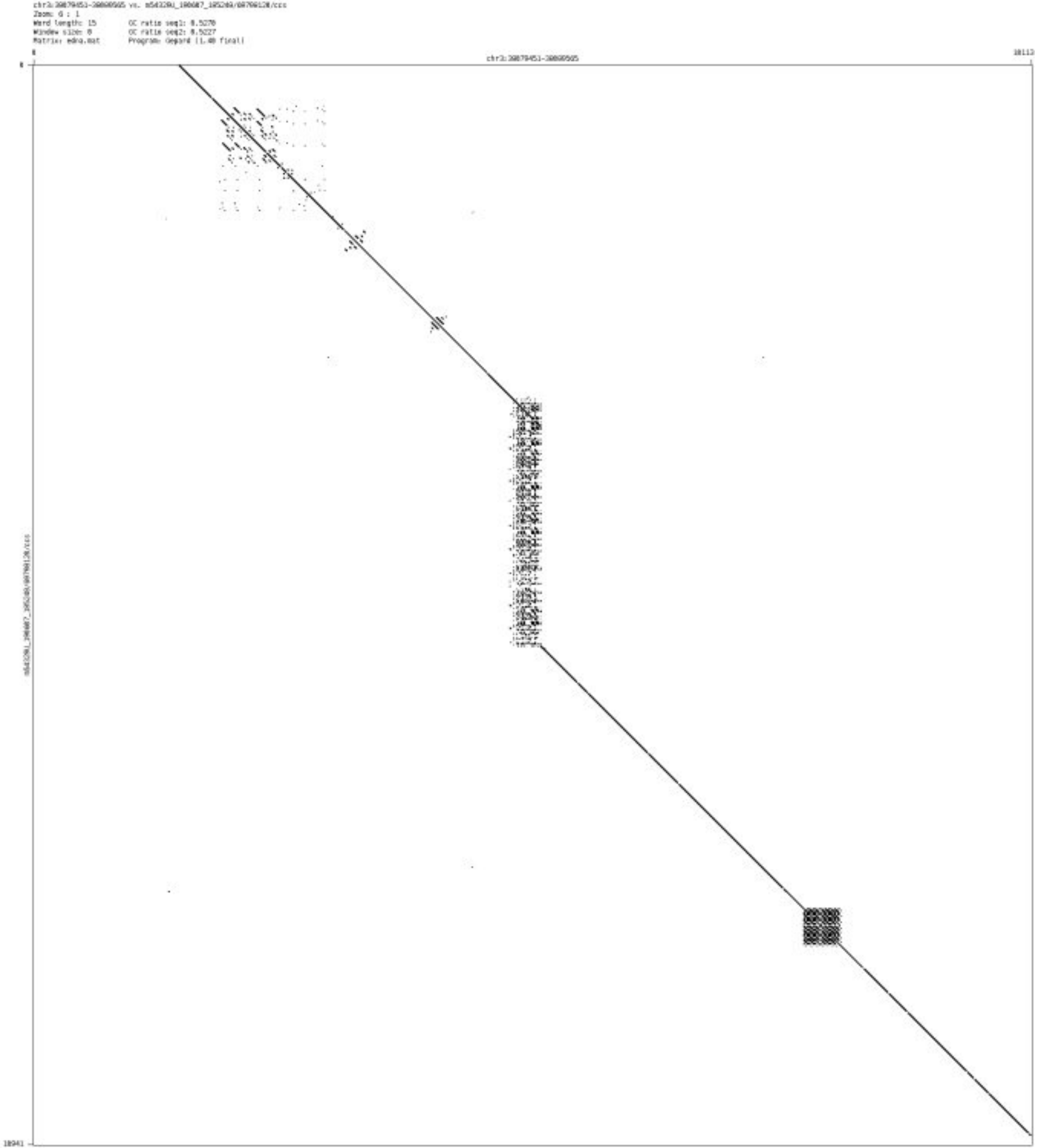

- chr3_178757959_178758511

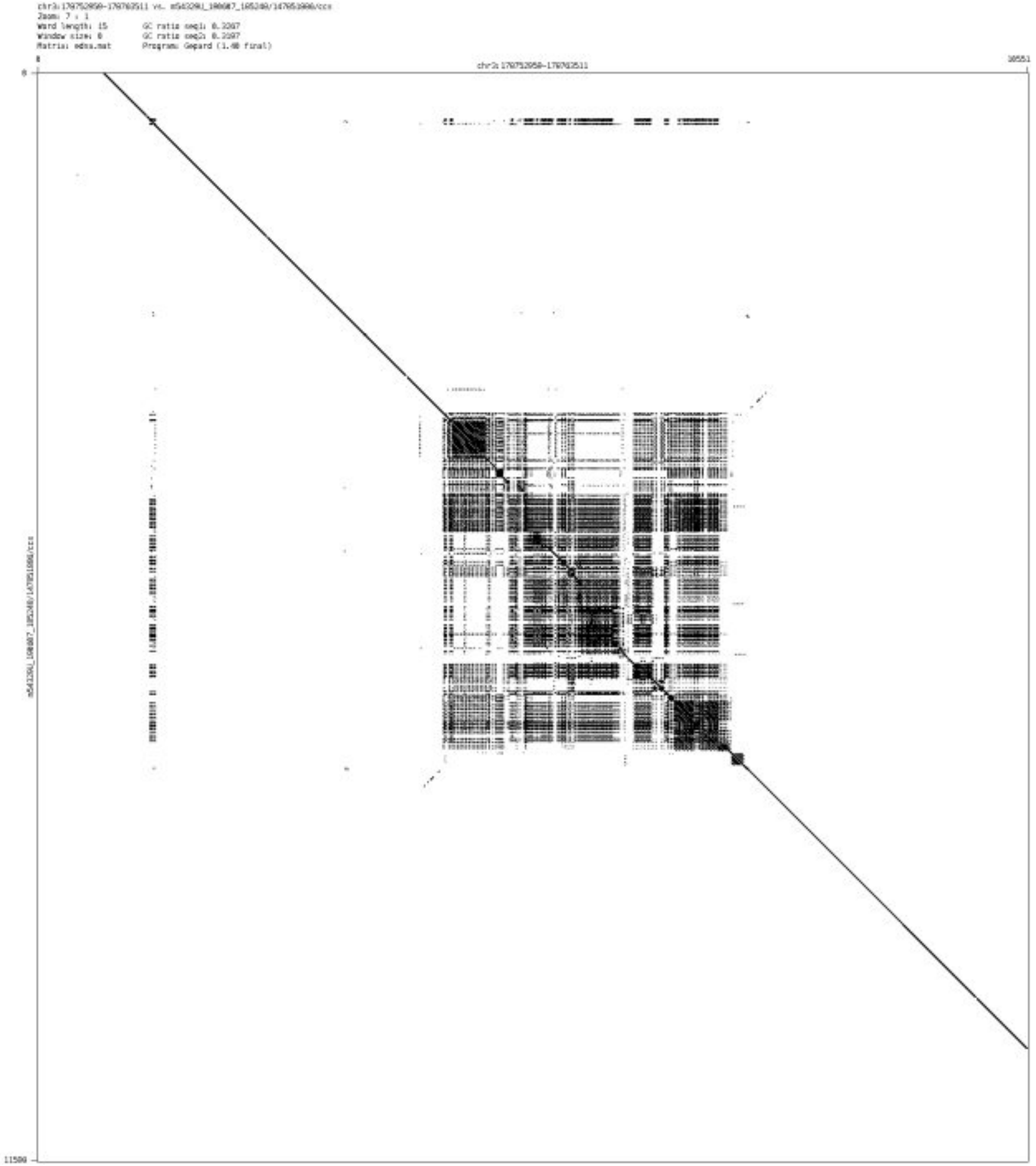

- chr3_184754687_184754818

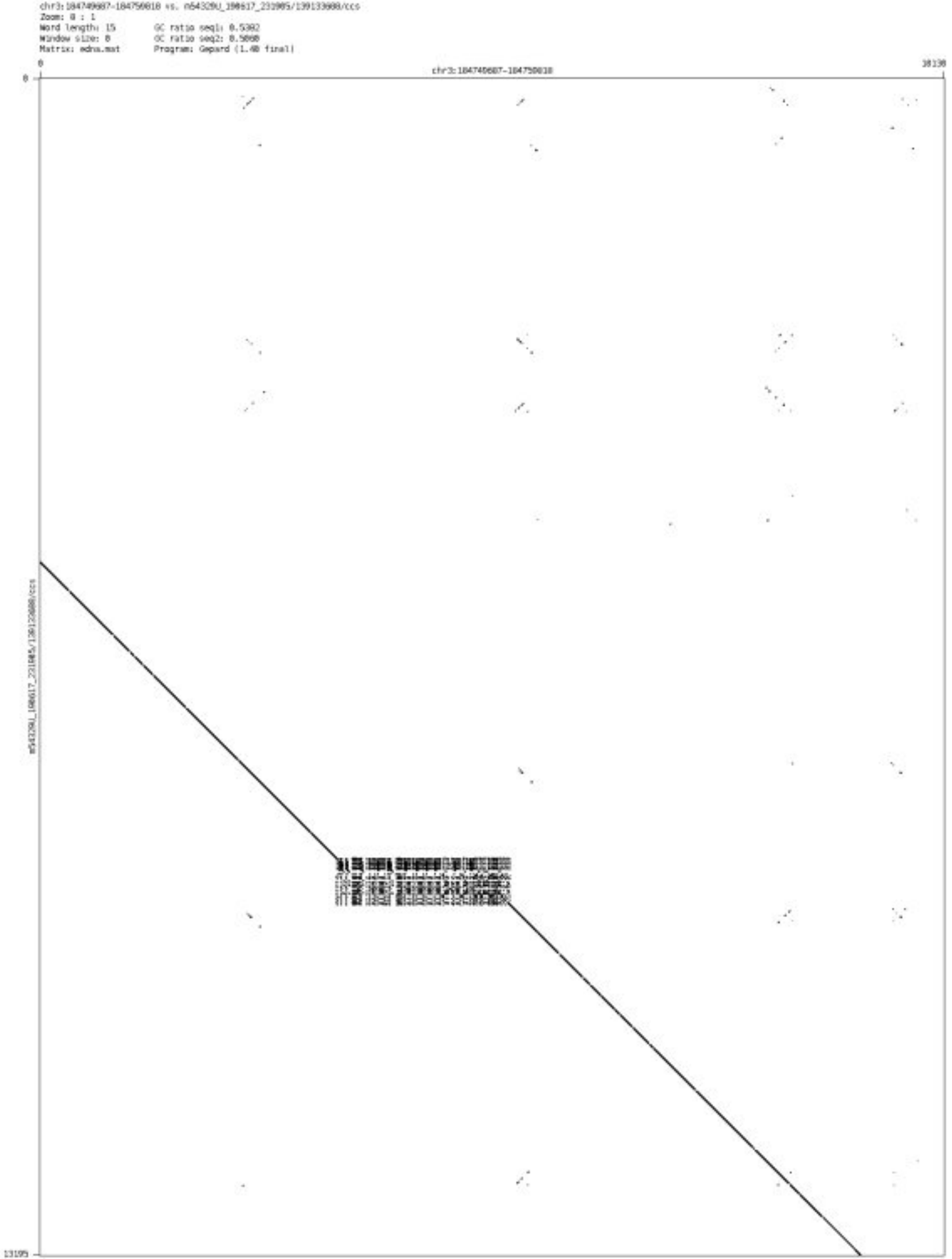

➢ chr4_555182_555337

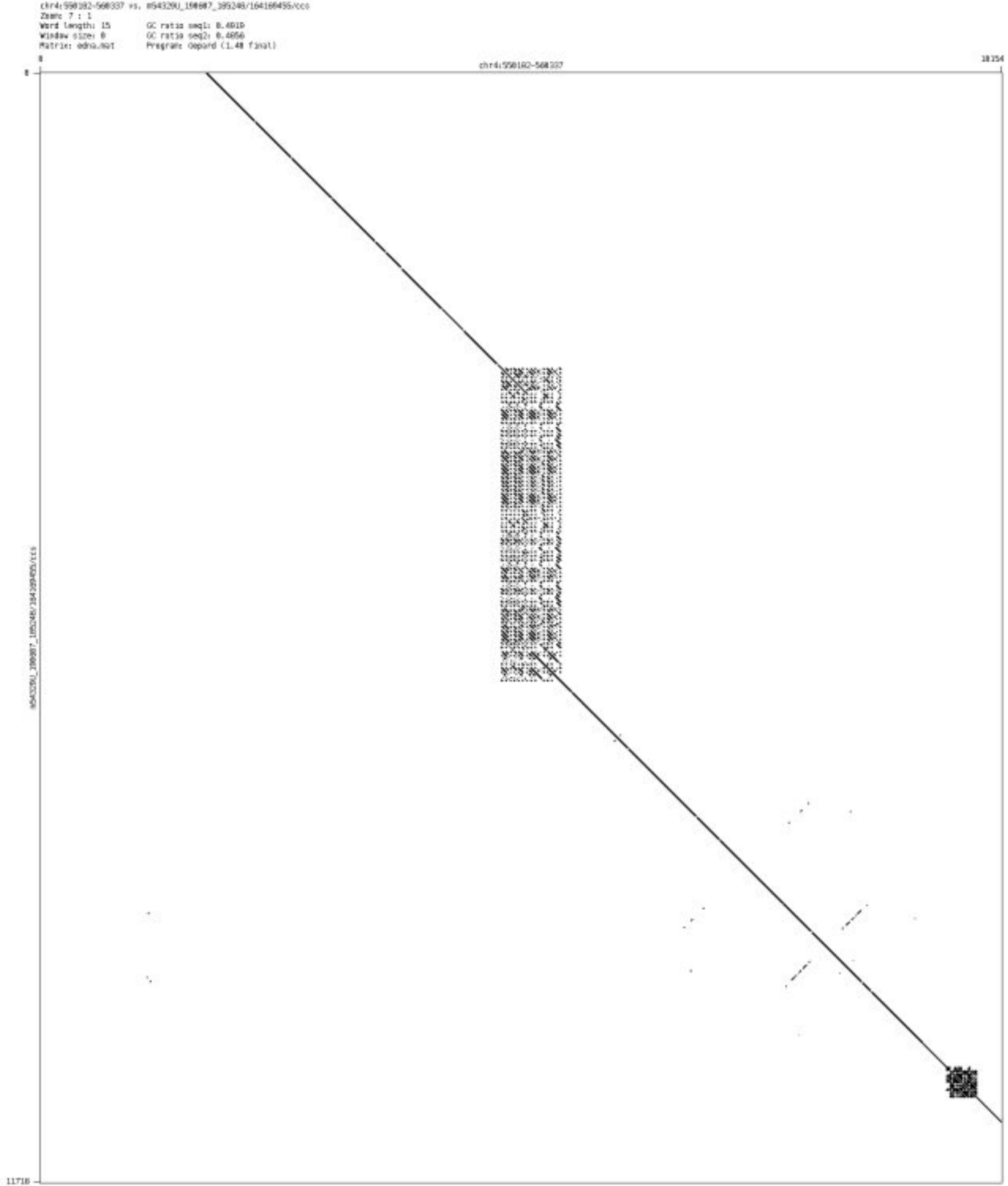

- chr4_1372721_1372812

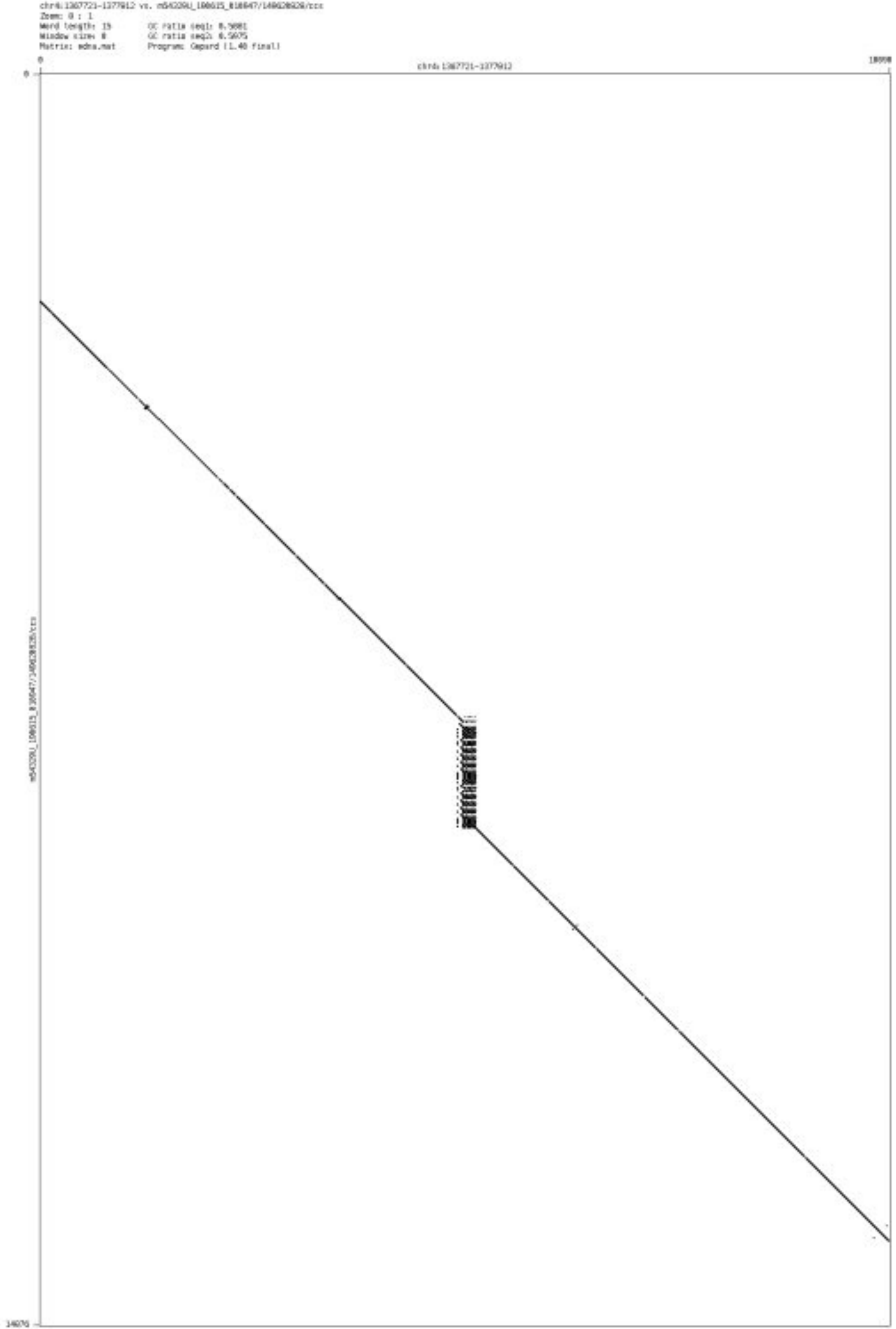

- chr4_6003932_6004226

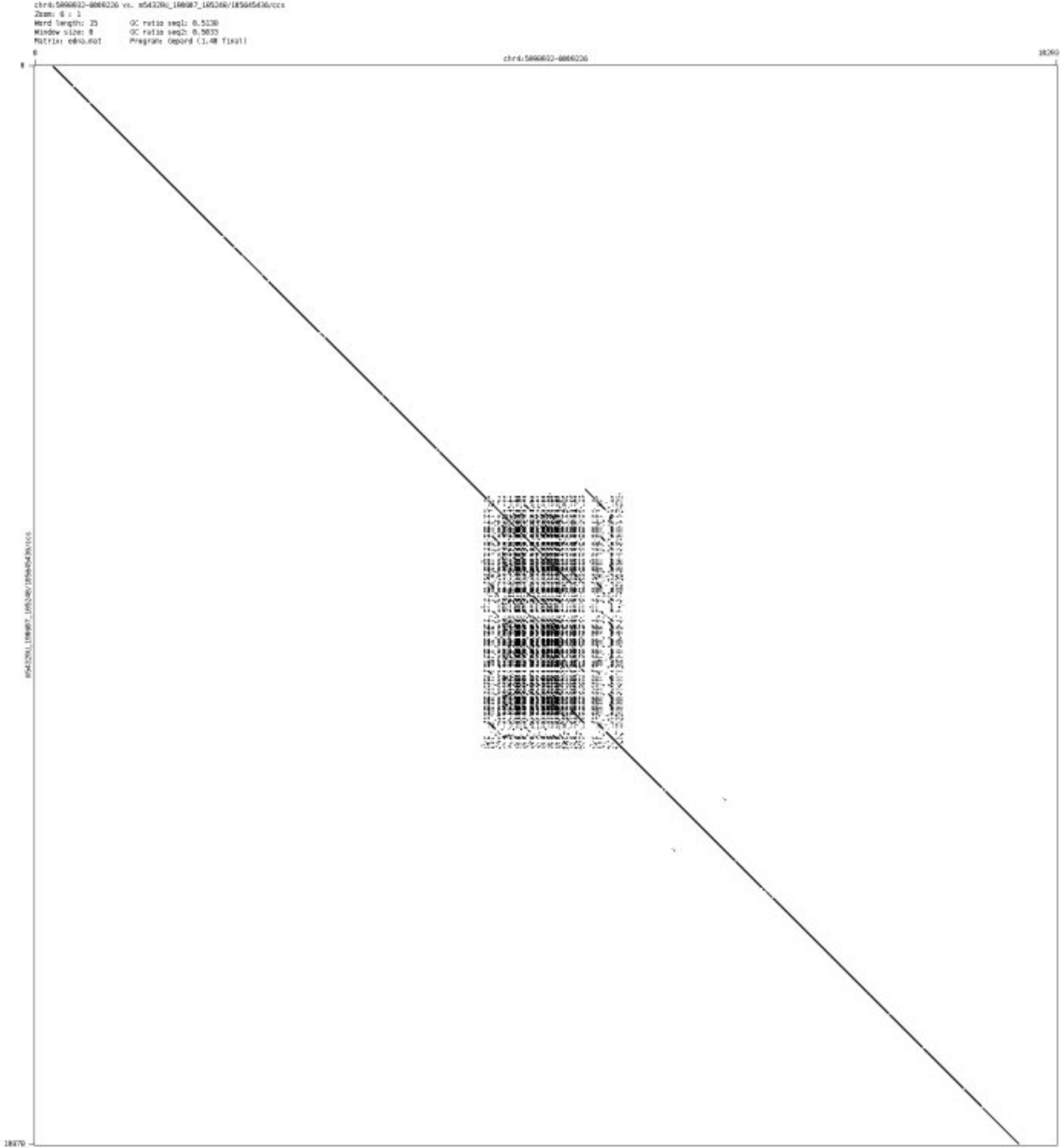

➢ chr4_7695588_7695671

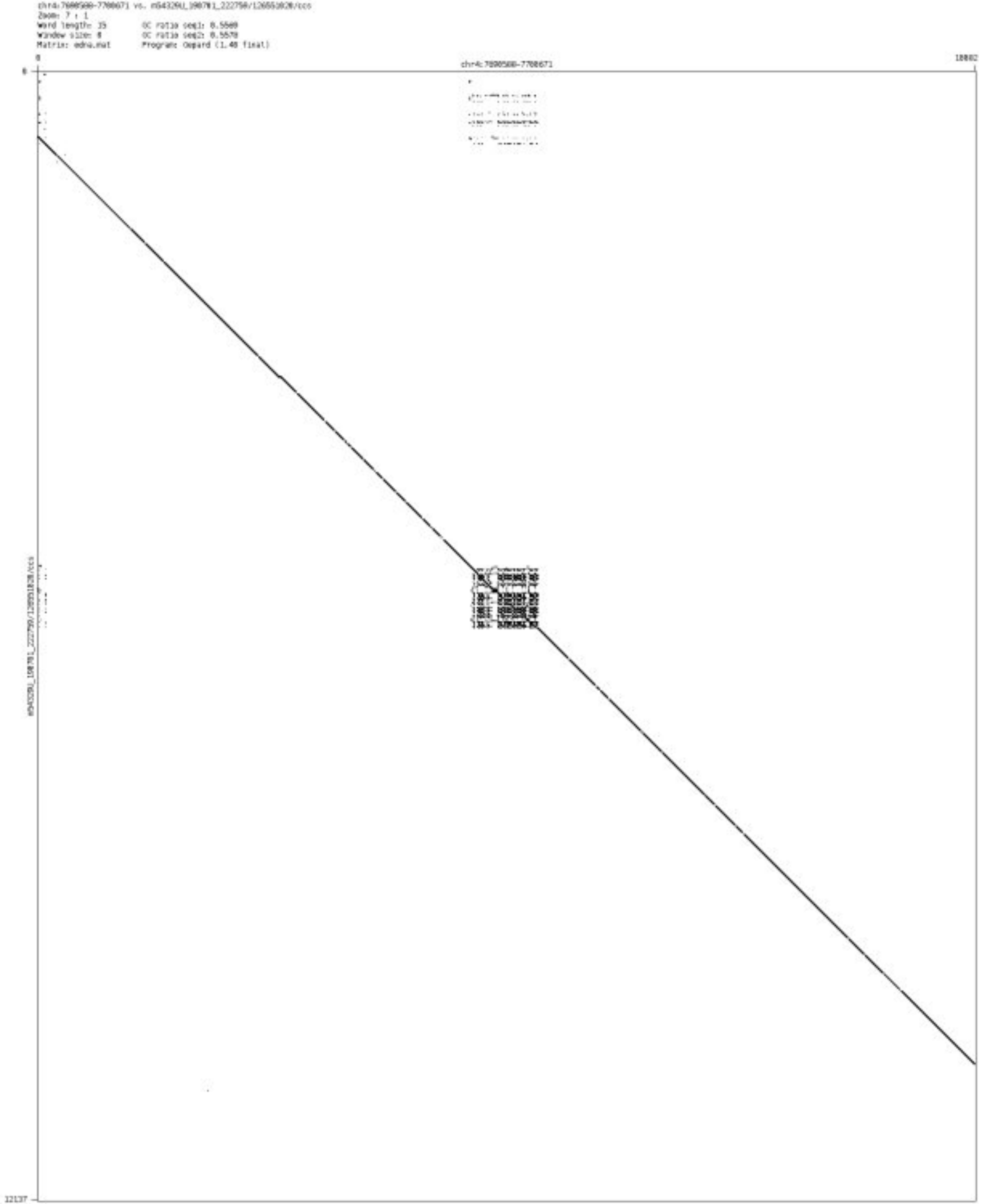

- chr4_12648458_12648926

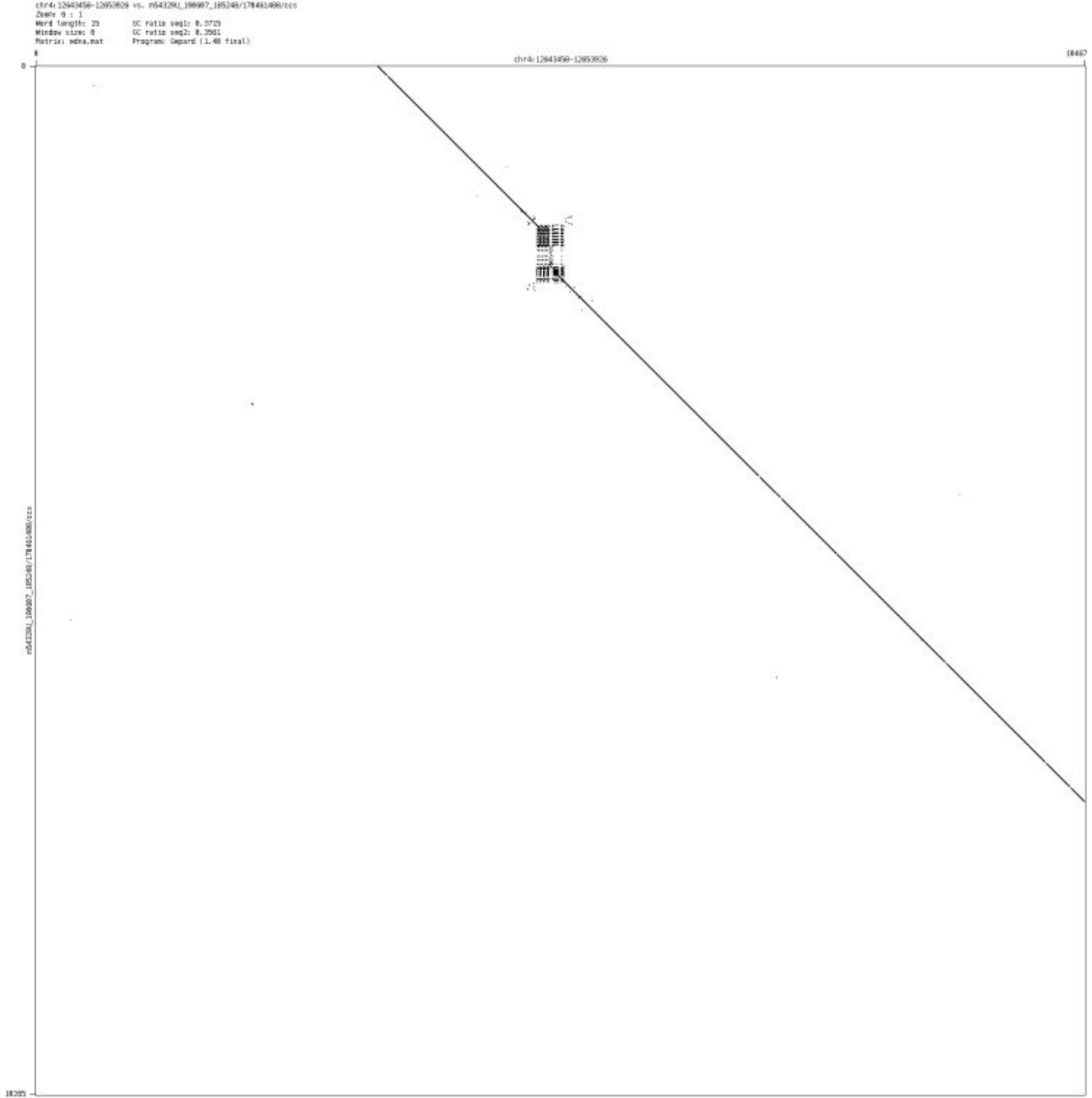

➢ chr4_37585251_37585395

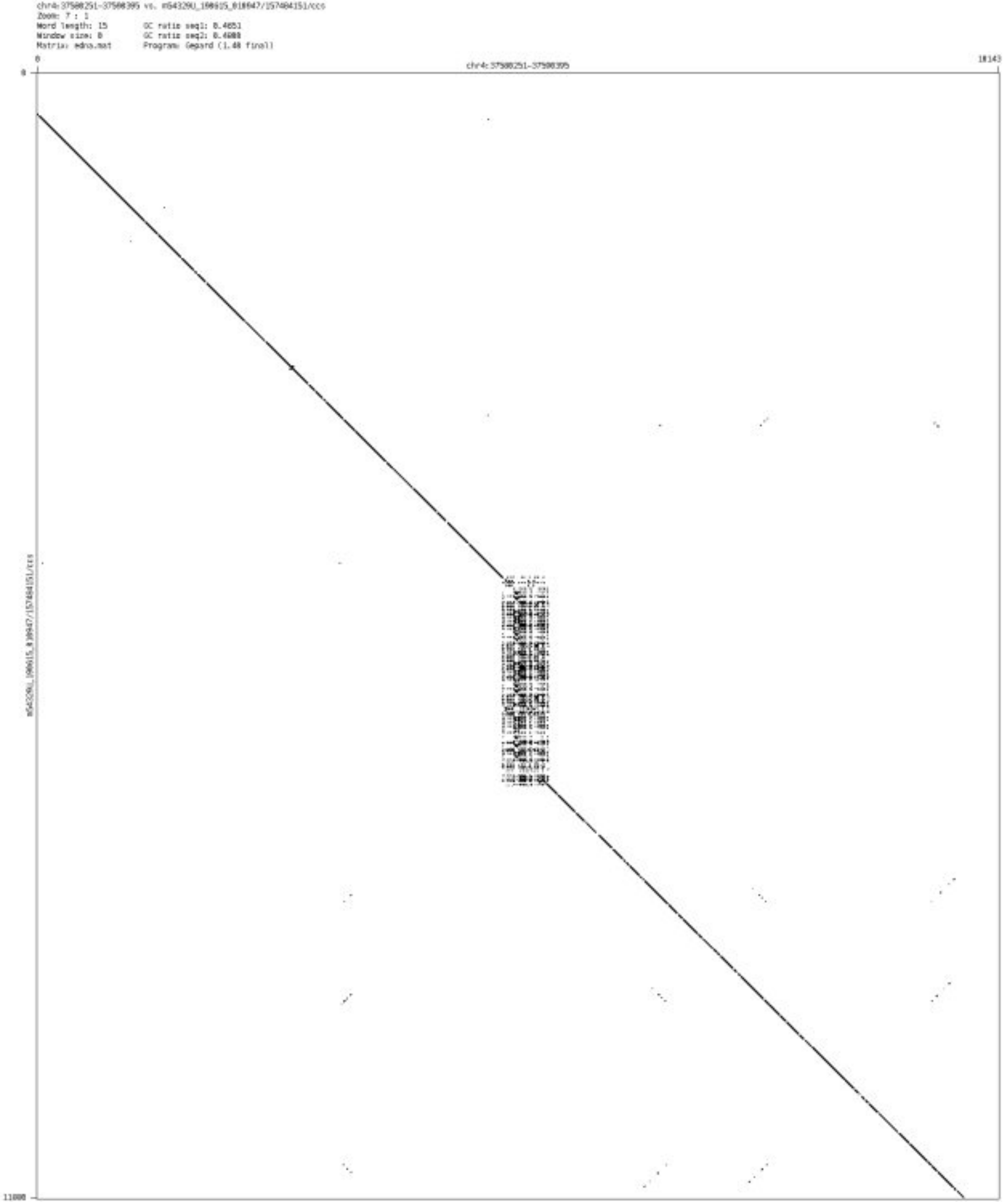

### ➢ chr4_45024776_45026675

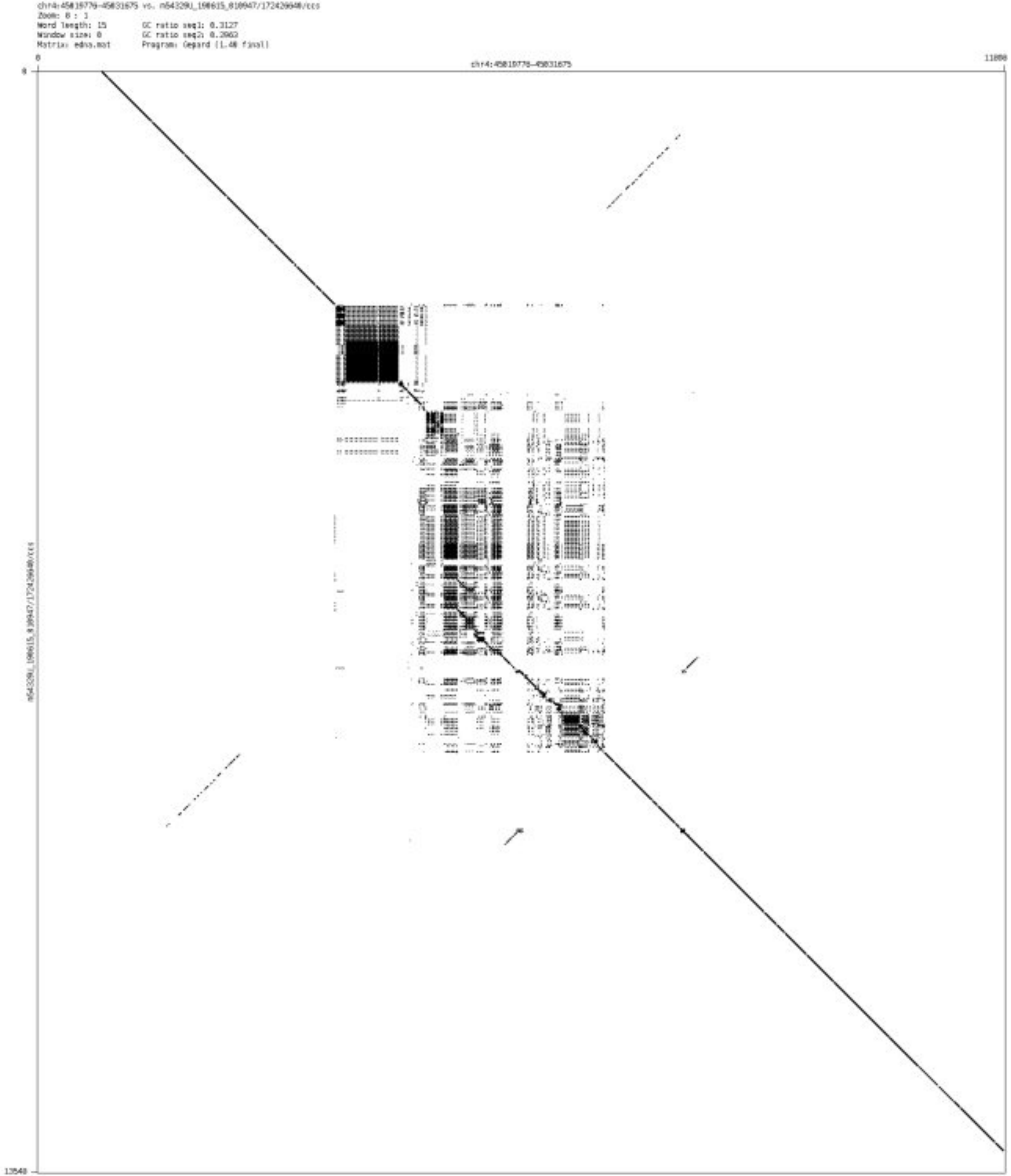

➢ chr4_72456201_72456342

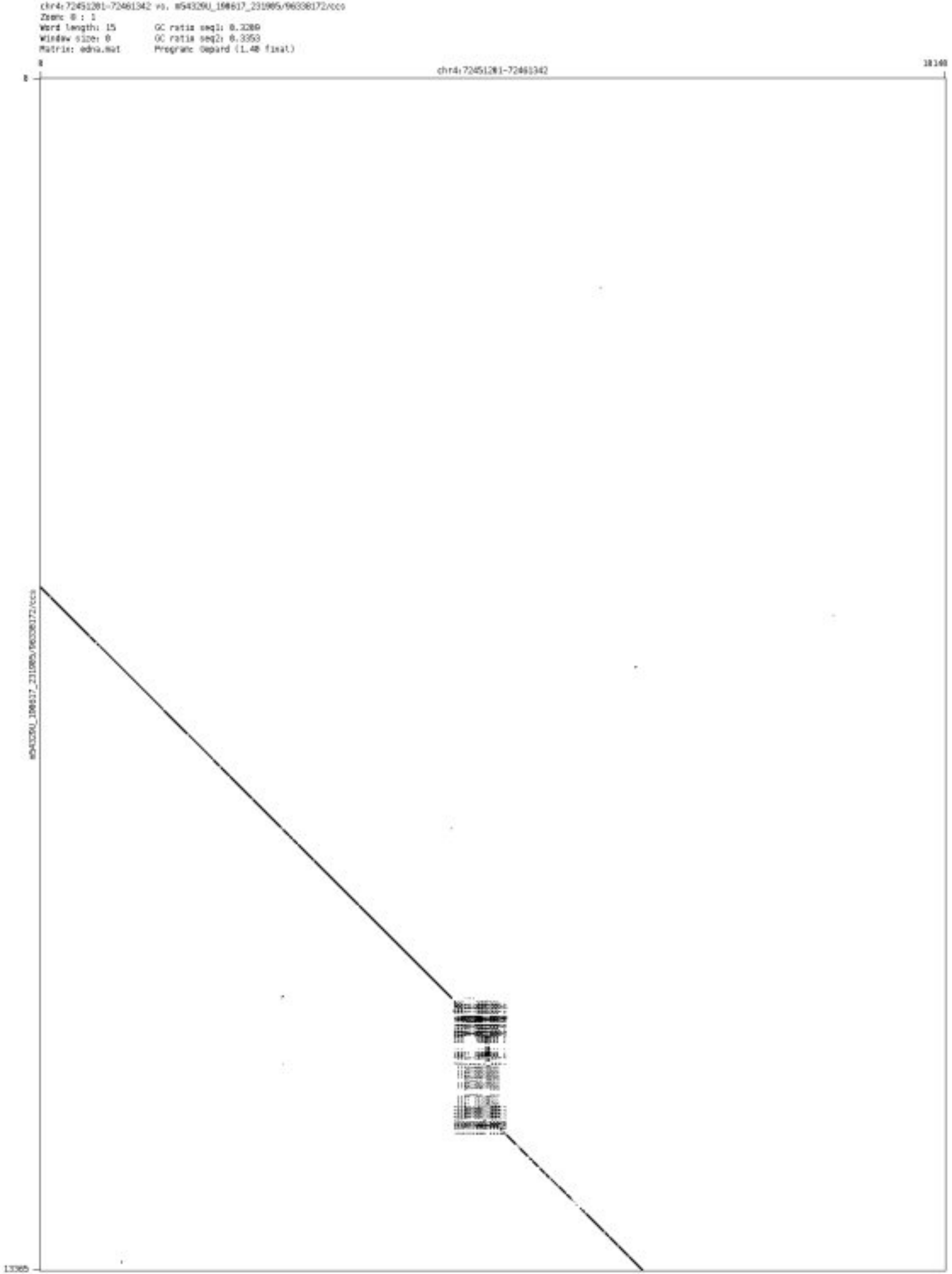

#### ➢ chr5_266746_266883

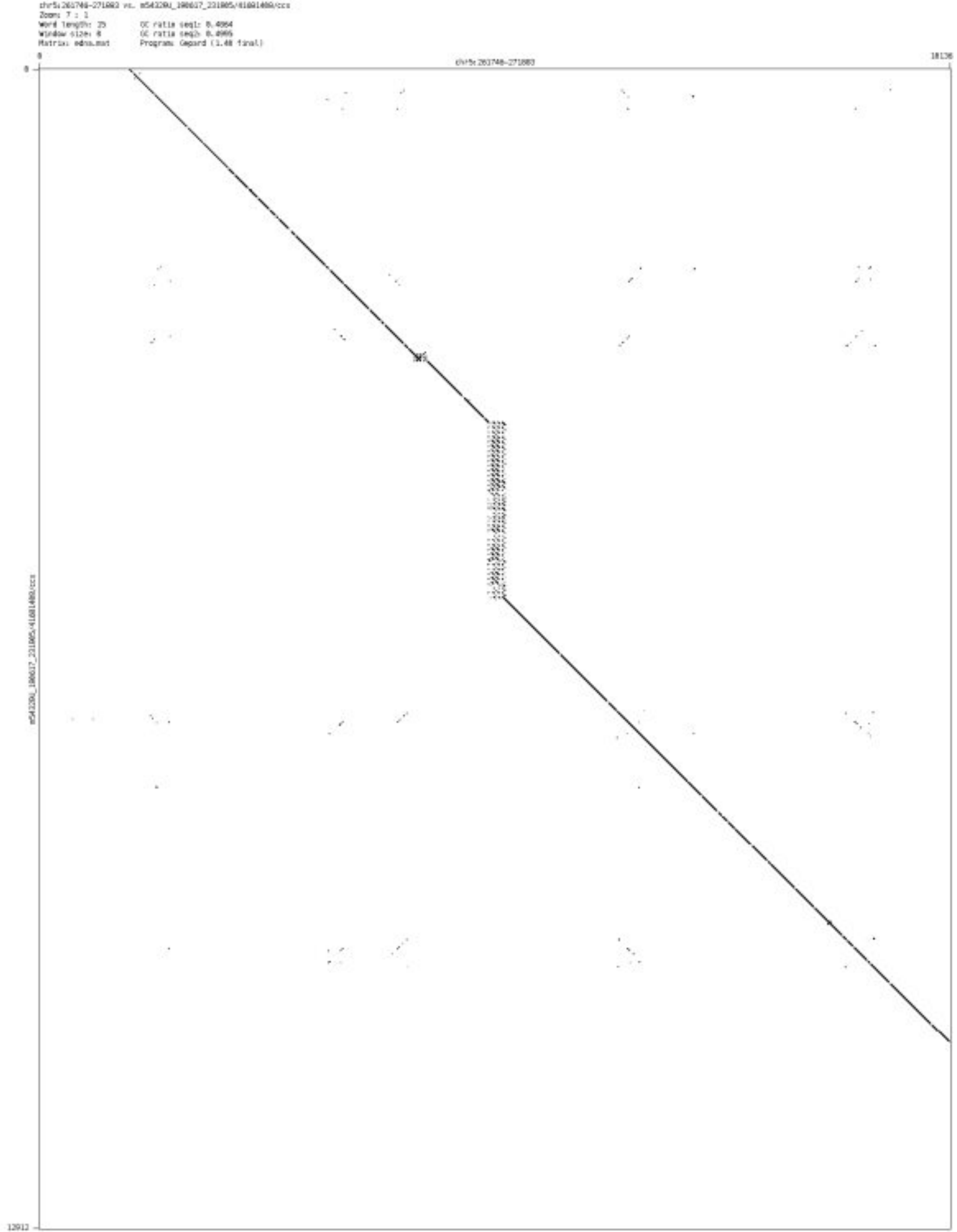

➢ chr5_415491_415686

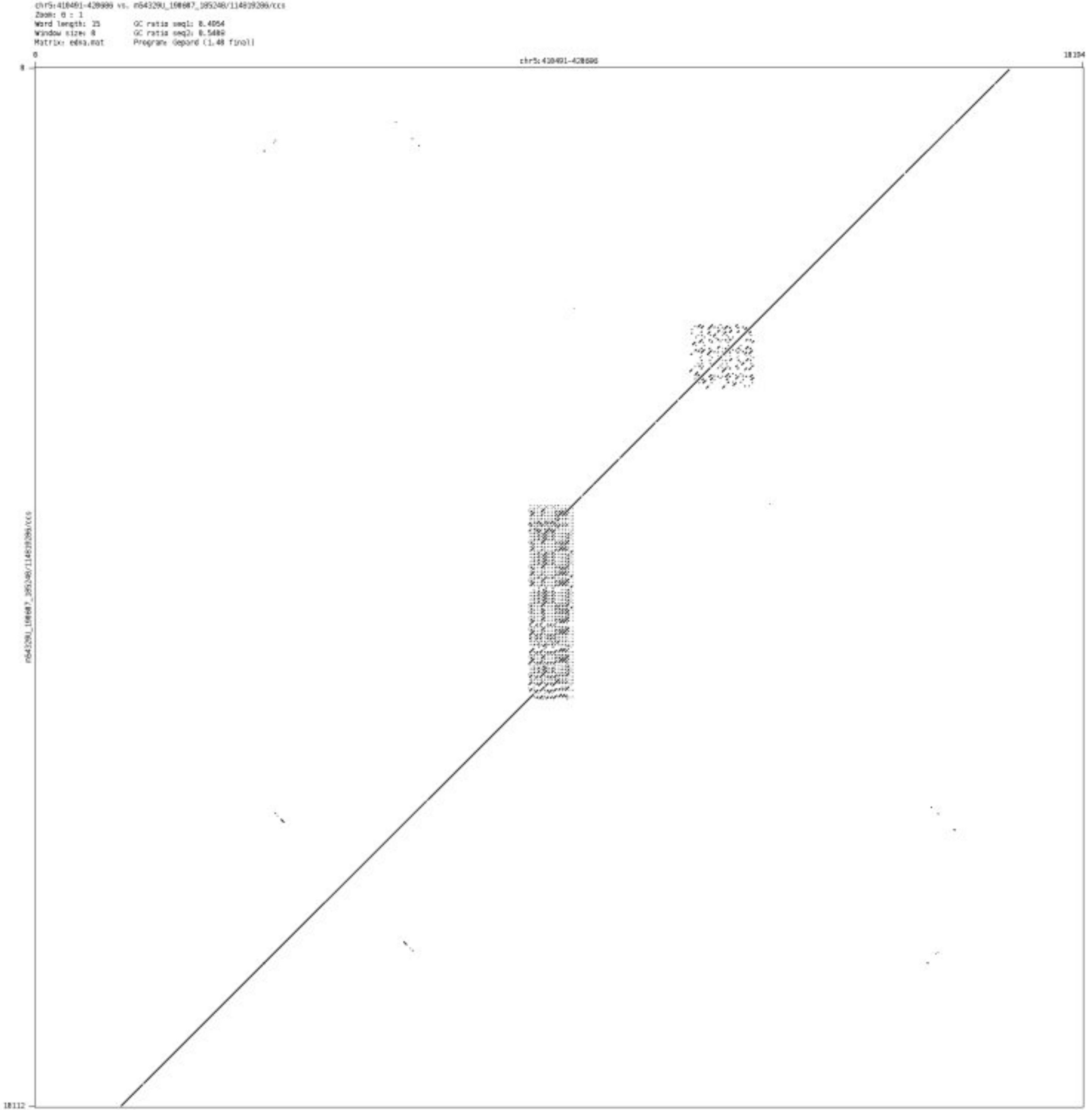

➢ chr5_628465_628658

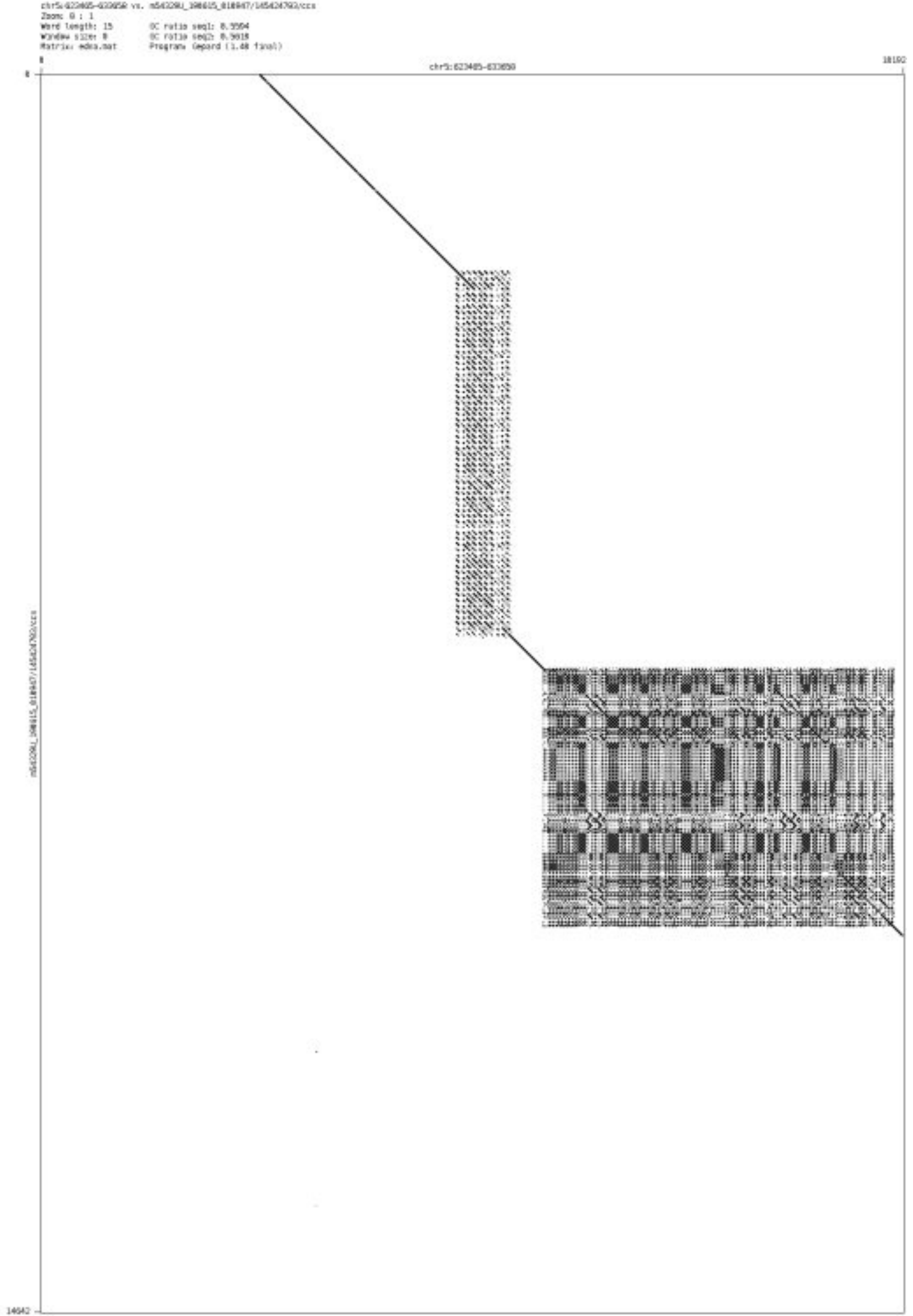

➢ chr5_1192054_1192167

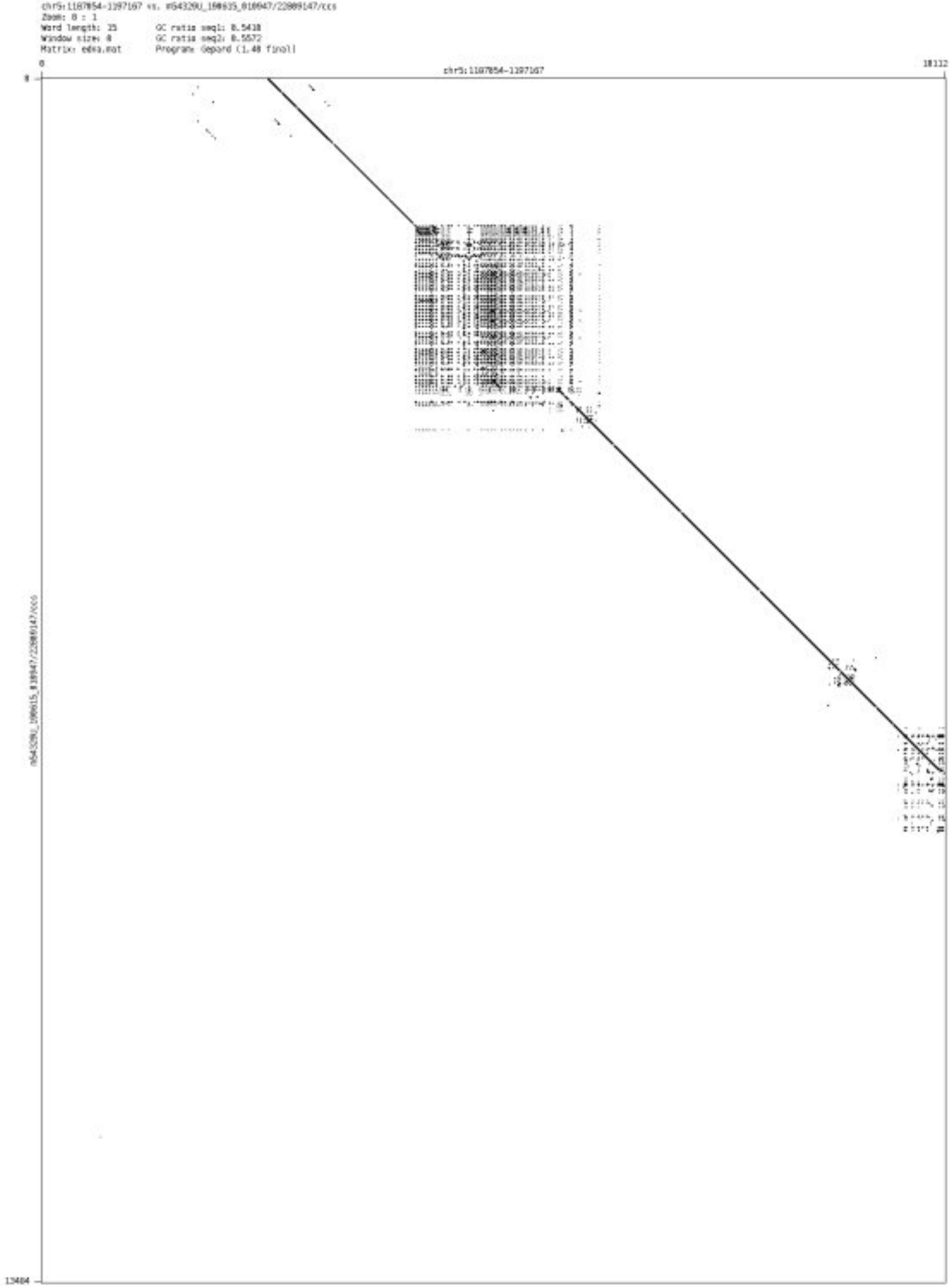

- chr5_1273419_1273547

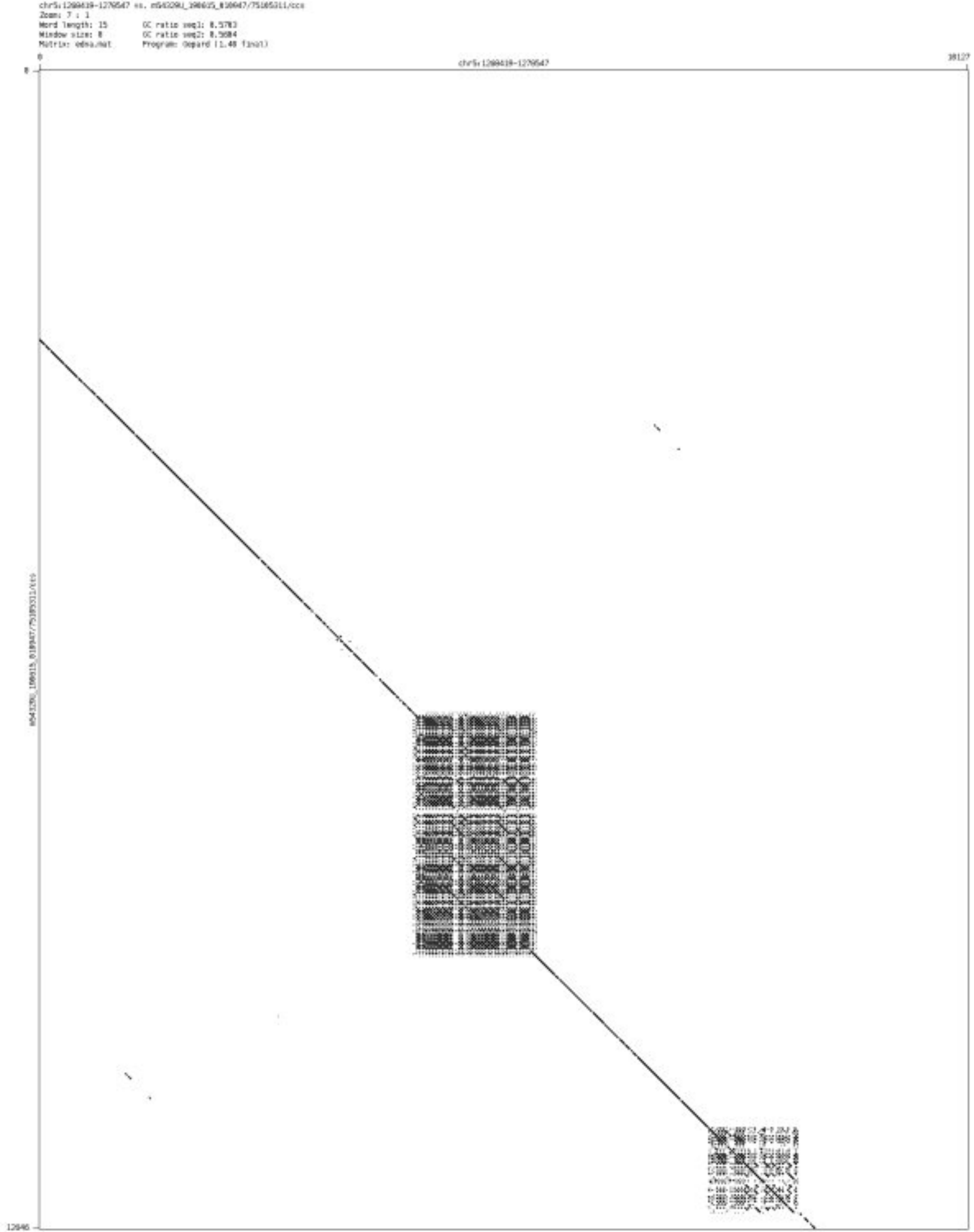

➢ chr5_1539973_1540158

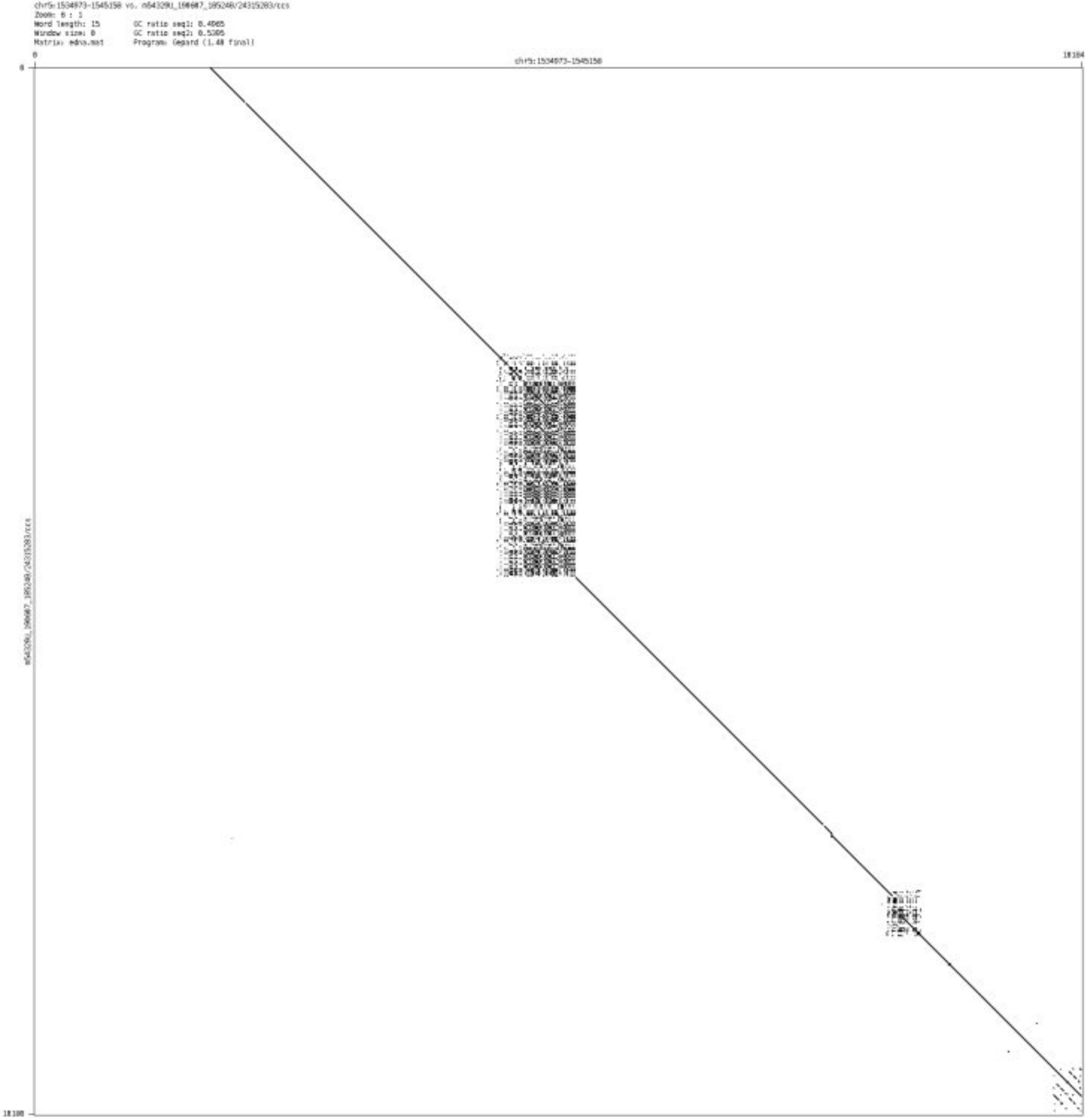

➢ chr5_5305060_5305279

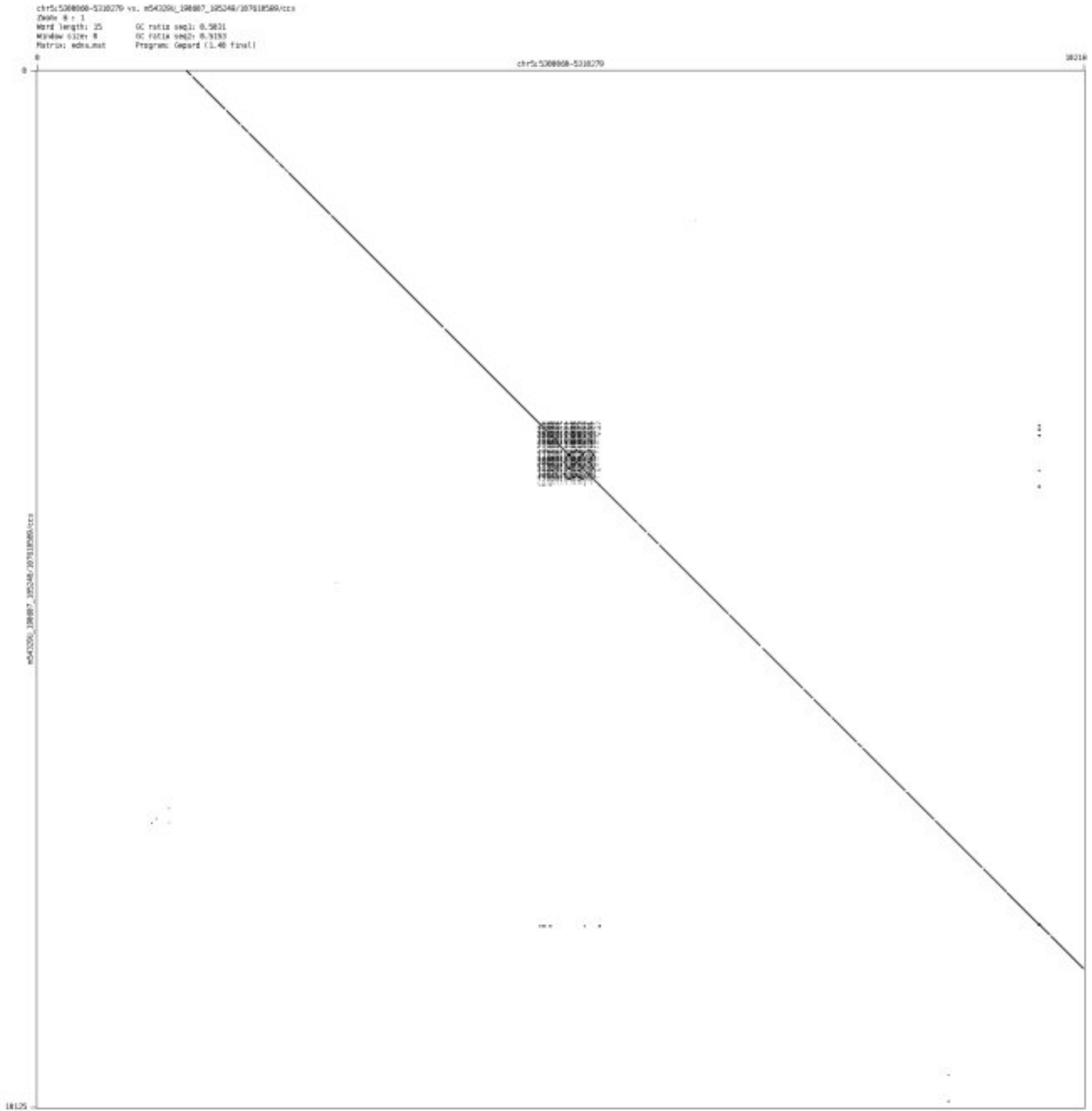

#### ➢ chr5_12425708_12426759

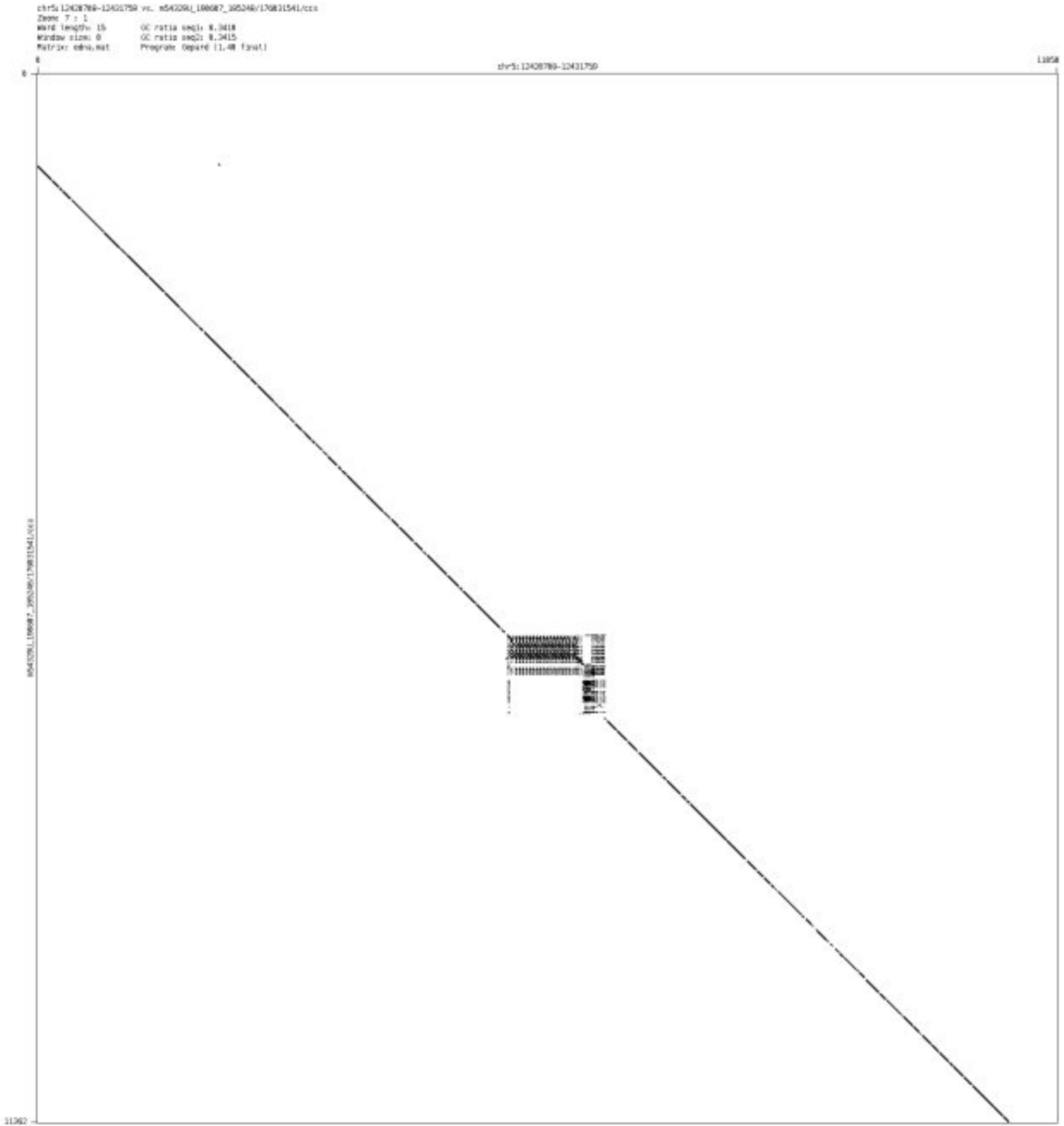

- chr5_67326226_67328703

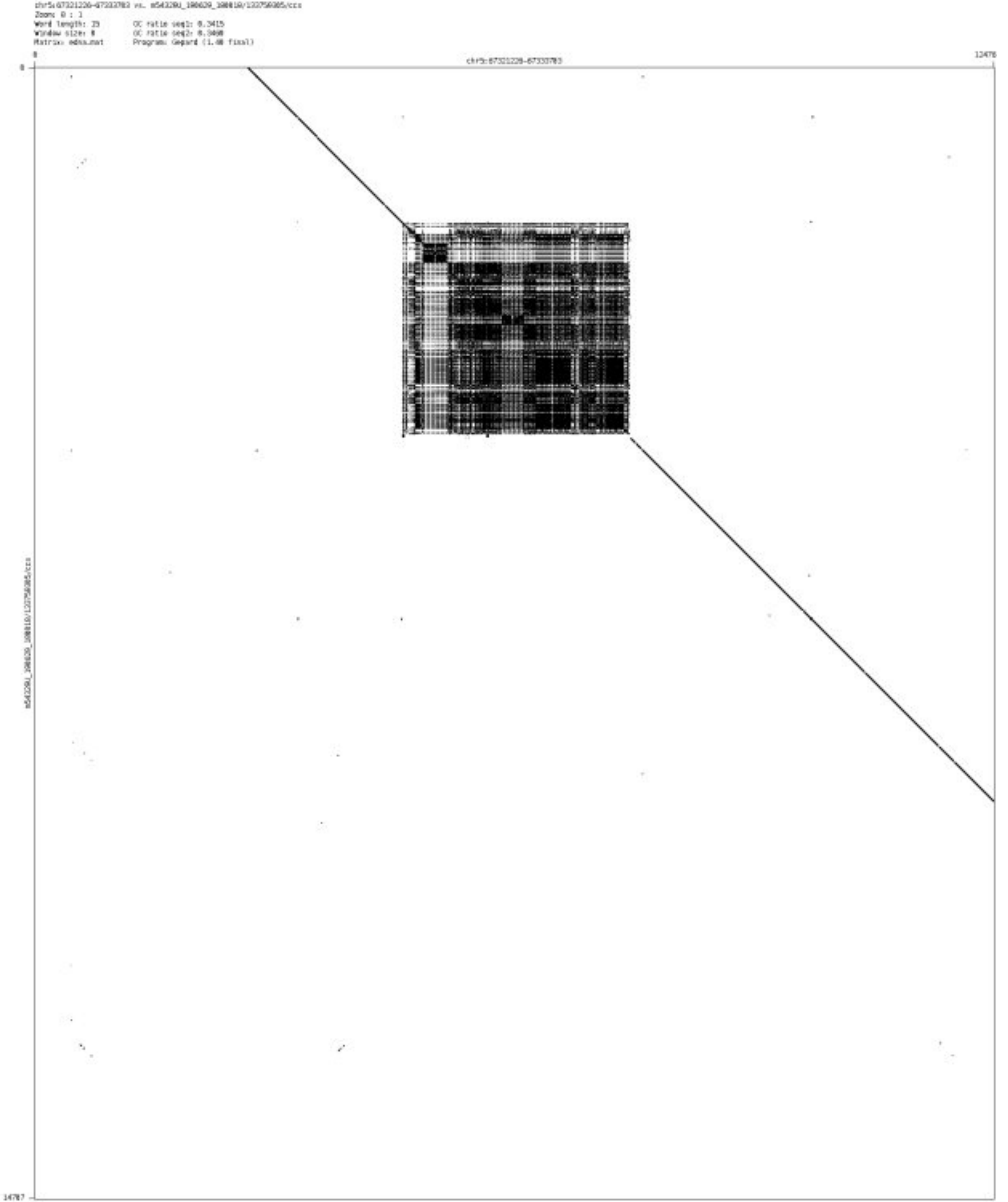

➢ chr5_80692983_80693579

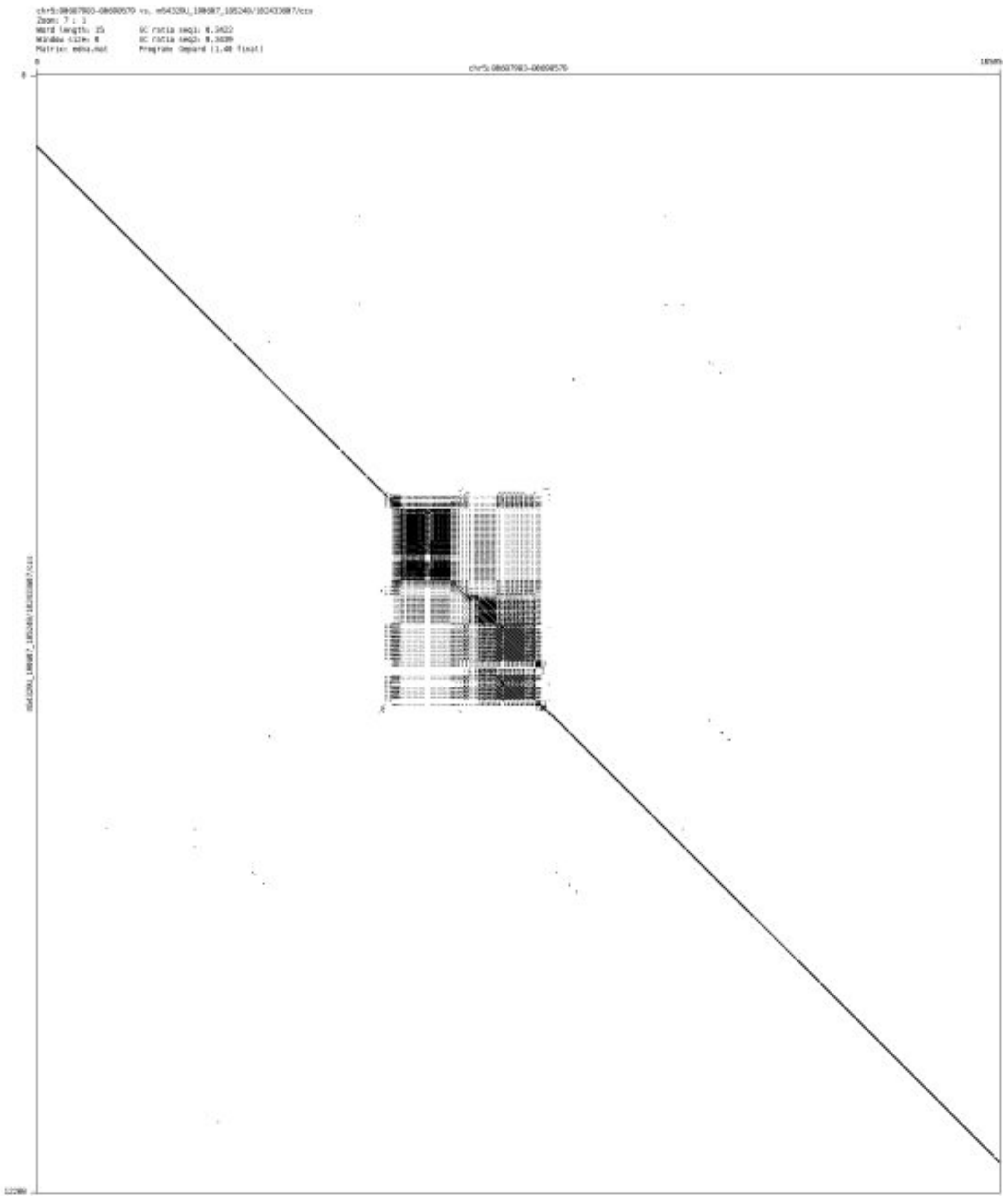

- chr5_101087938_101088040

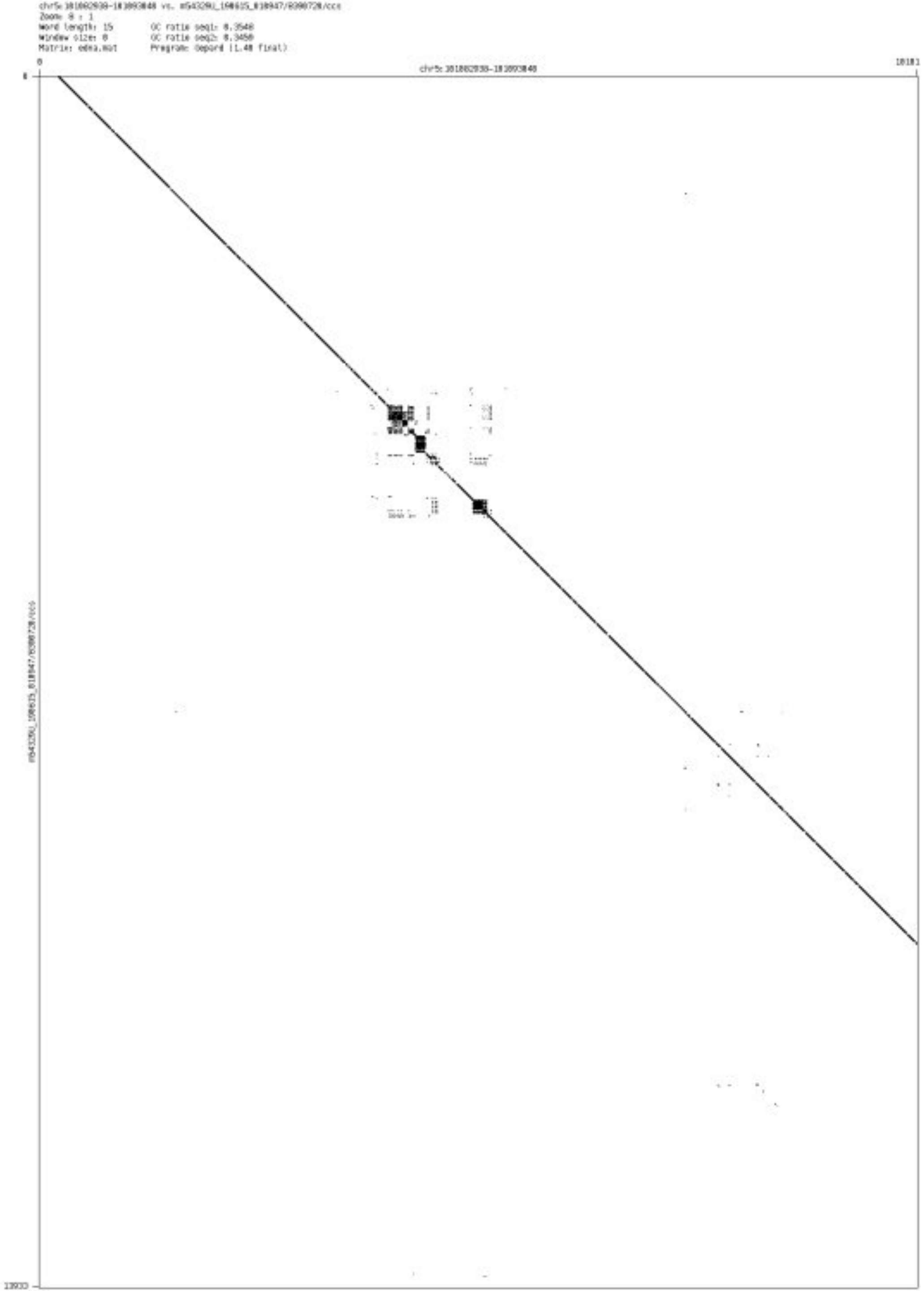

- chr5_178585514_178585658

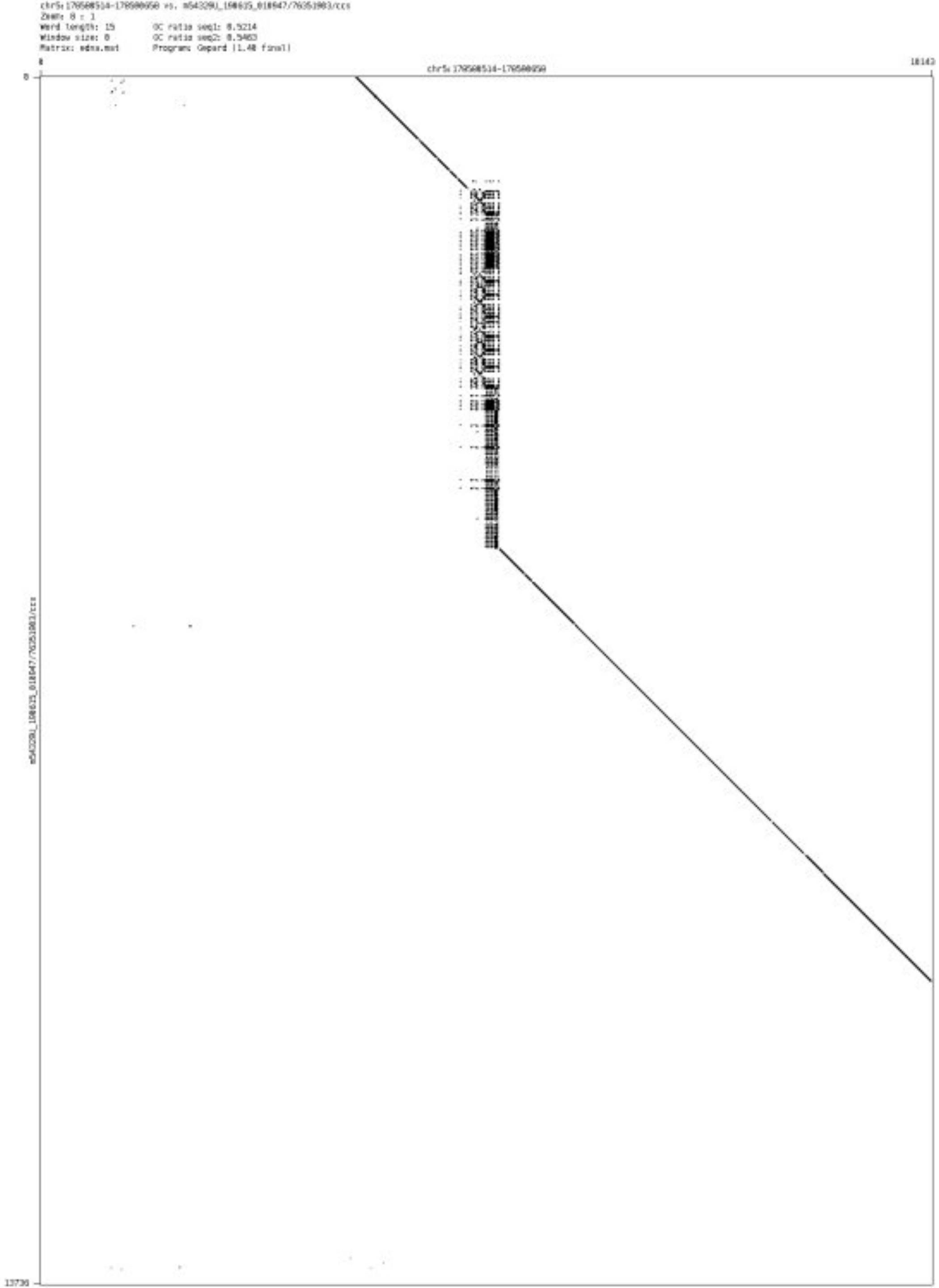

- chr5_180634930_180635915

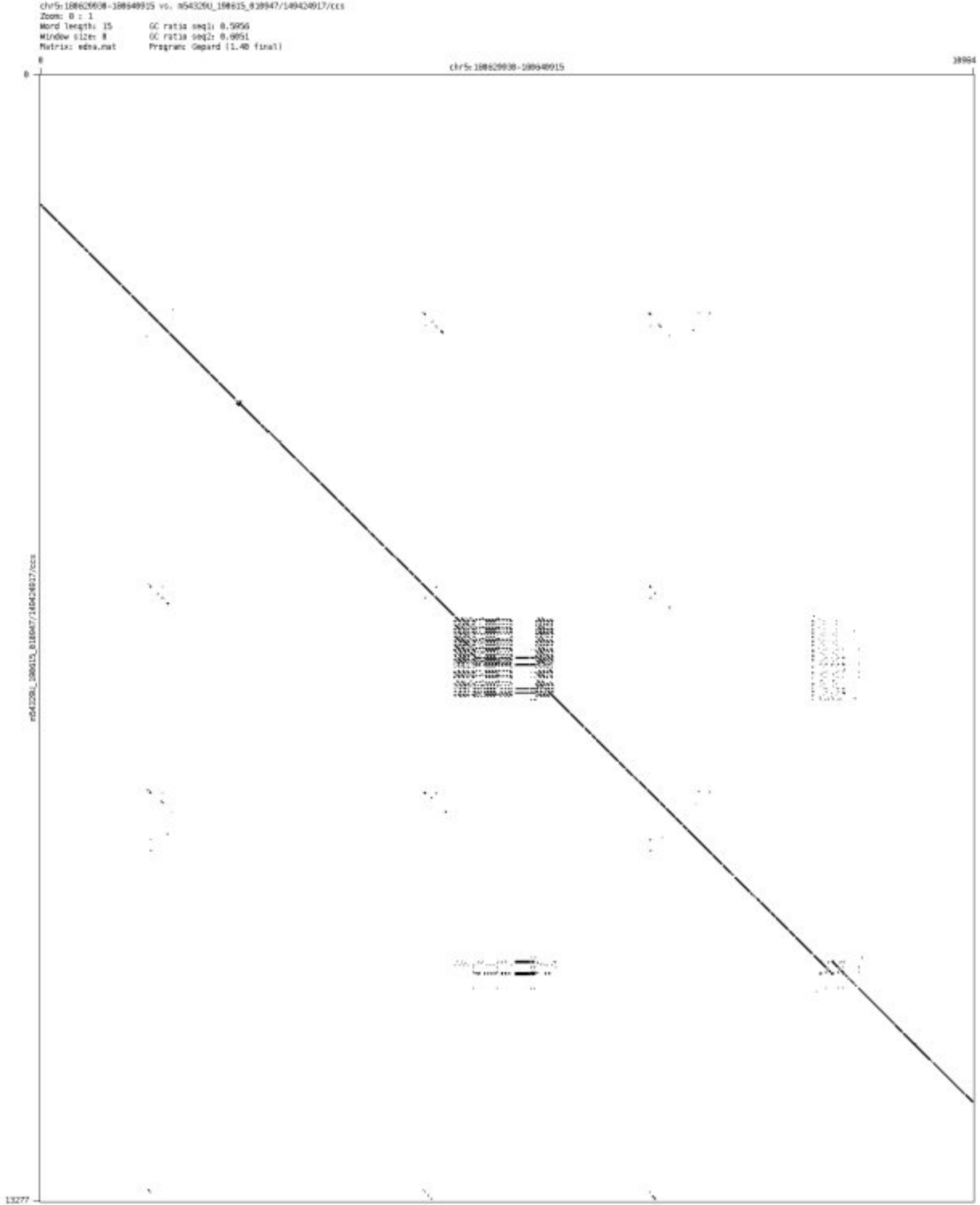

➢ chr6_372042_372248

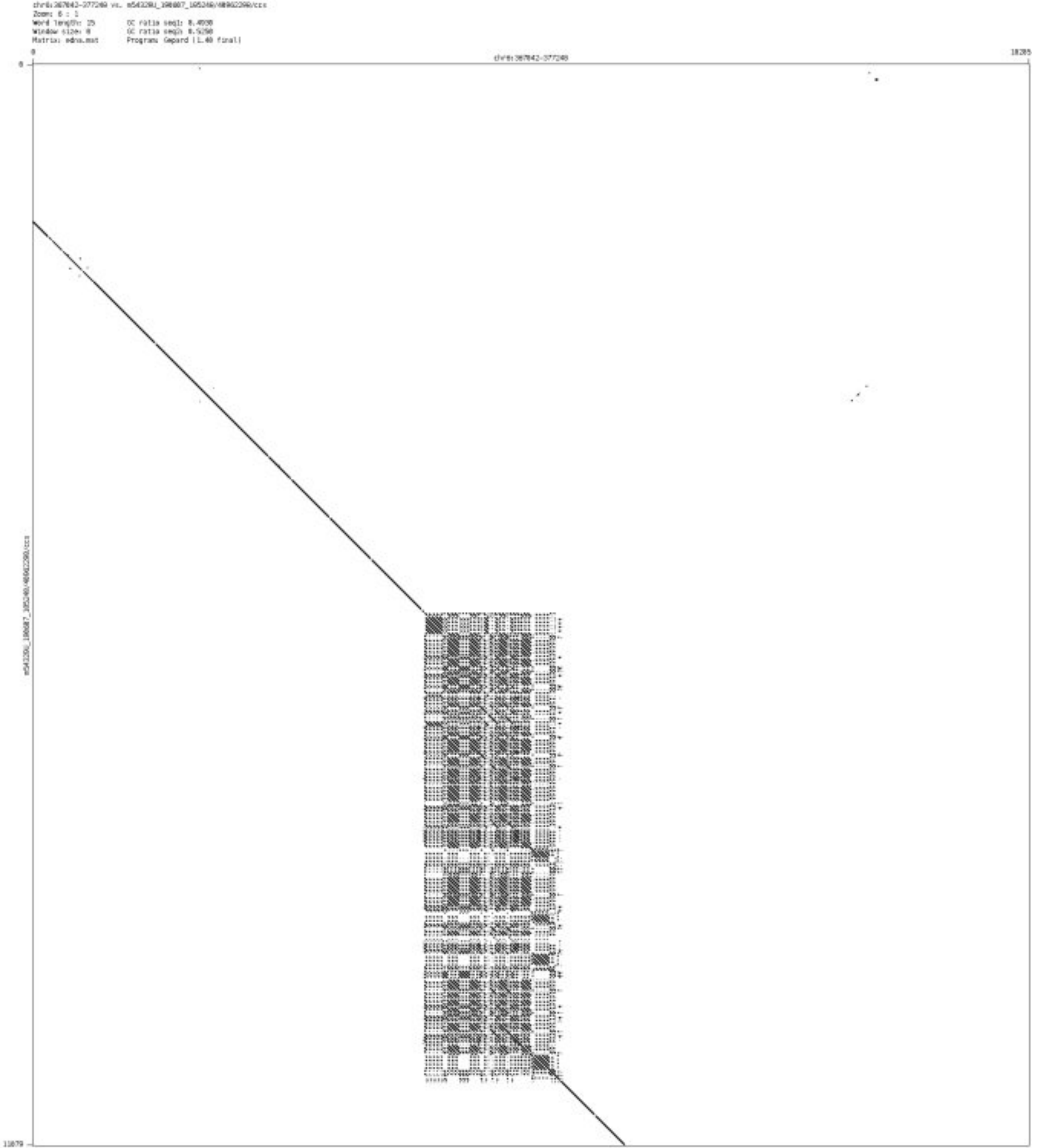

- chr6_28054362_28055446

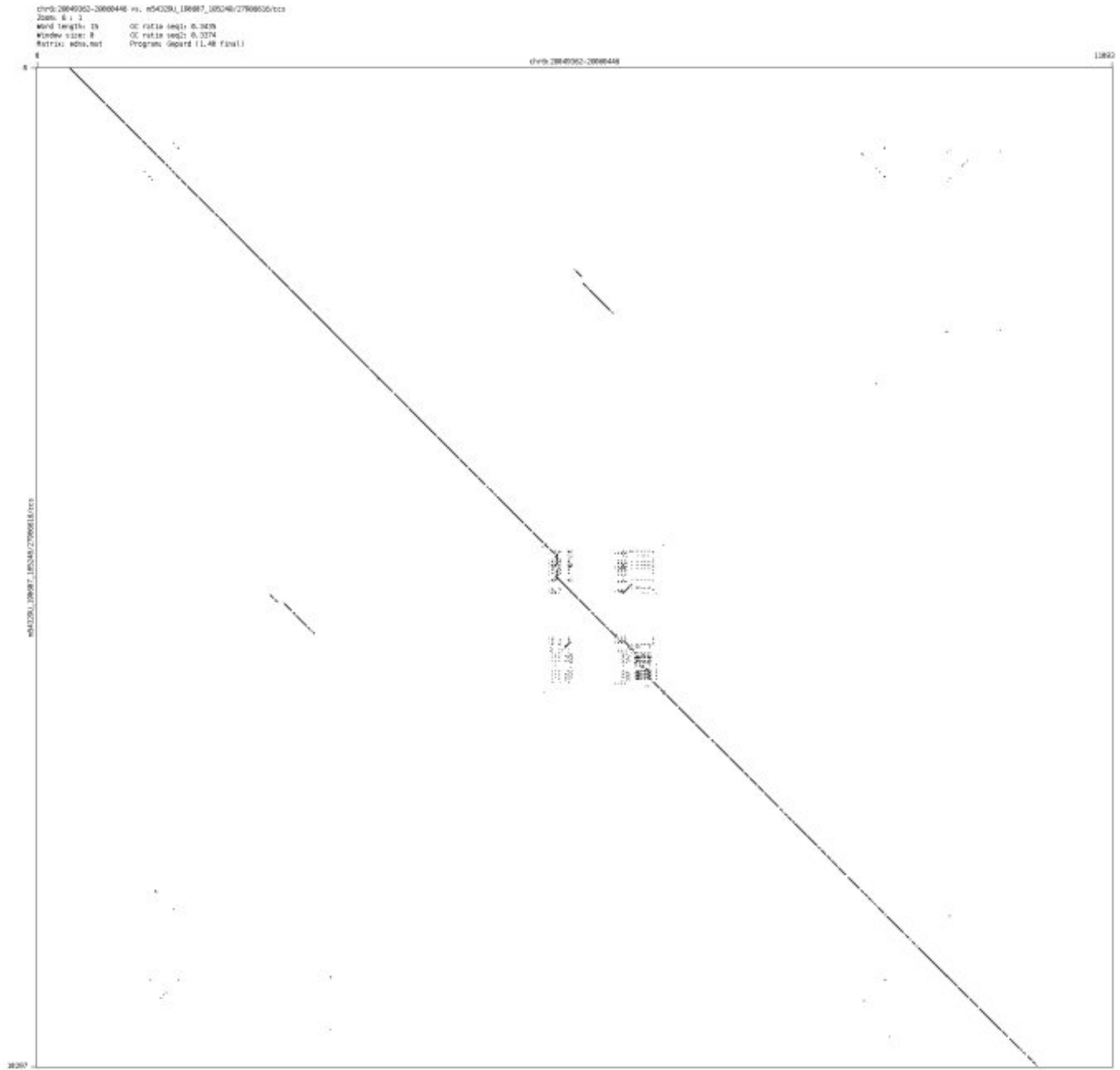

➢ chr6_31428235_31429956

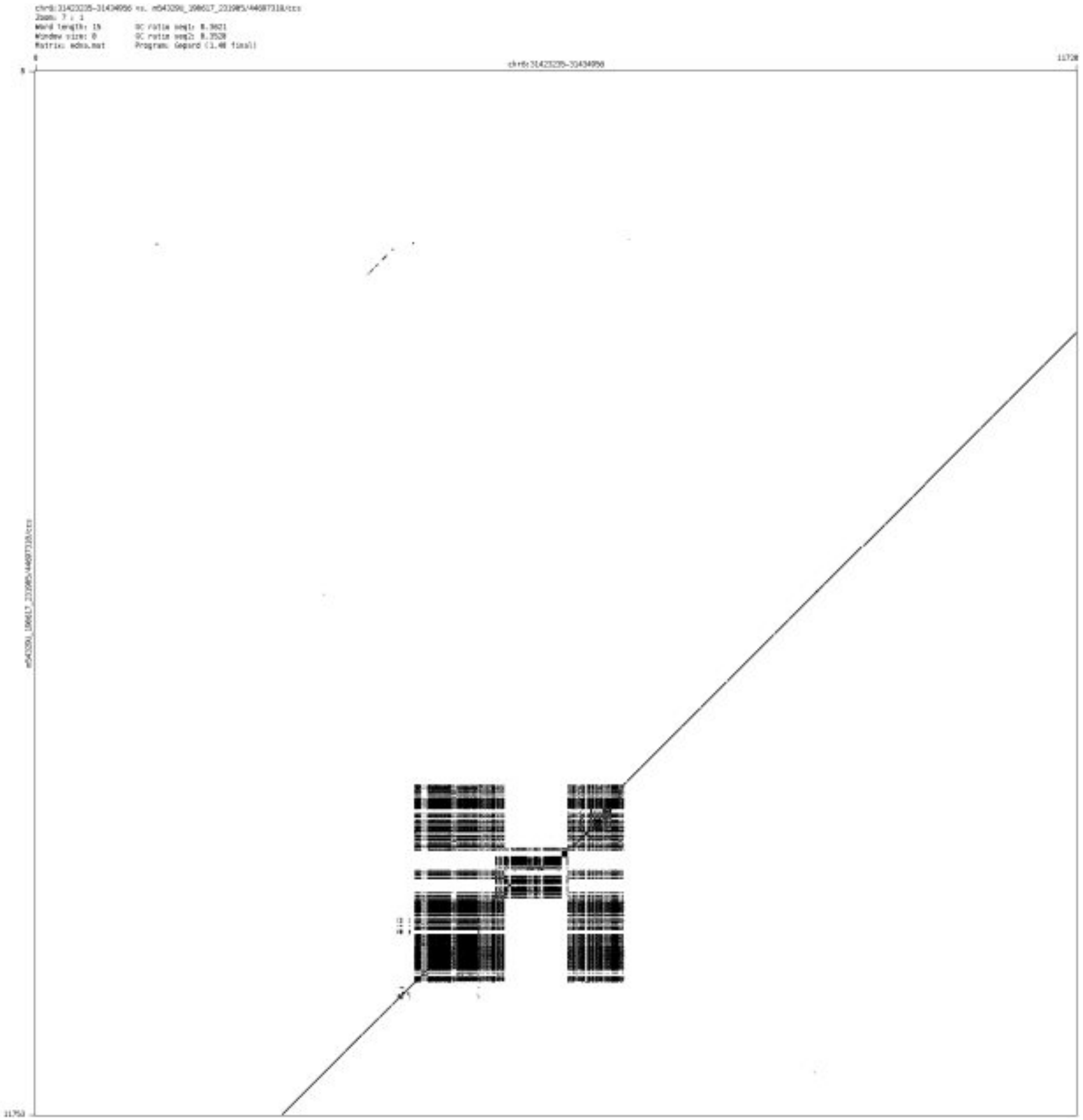

- chr6_35169659_35169798

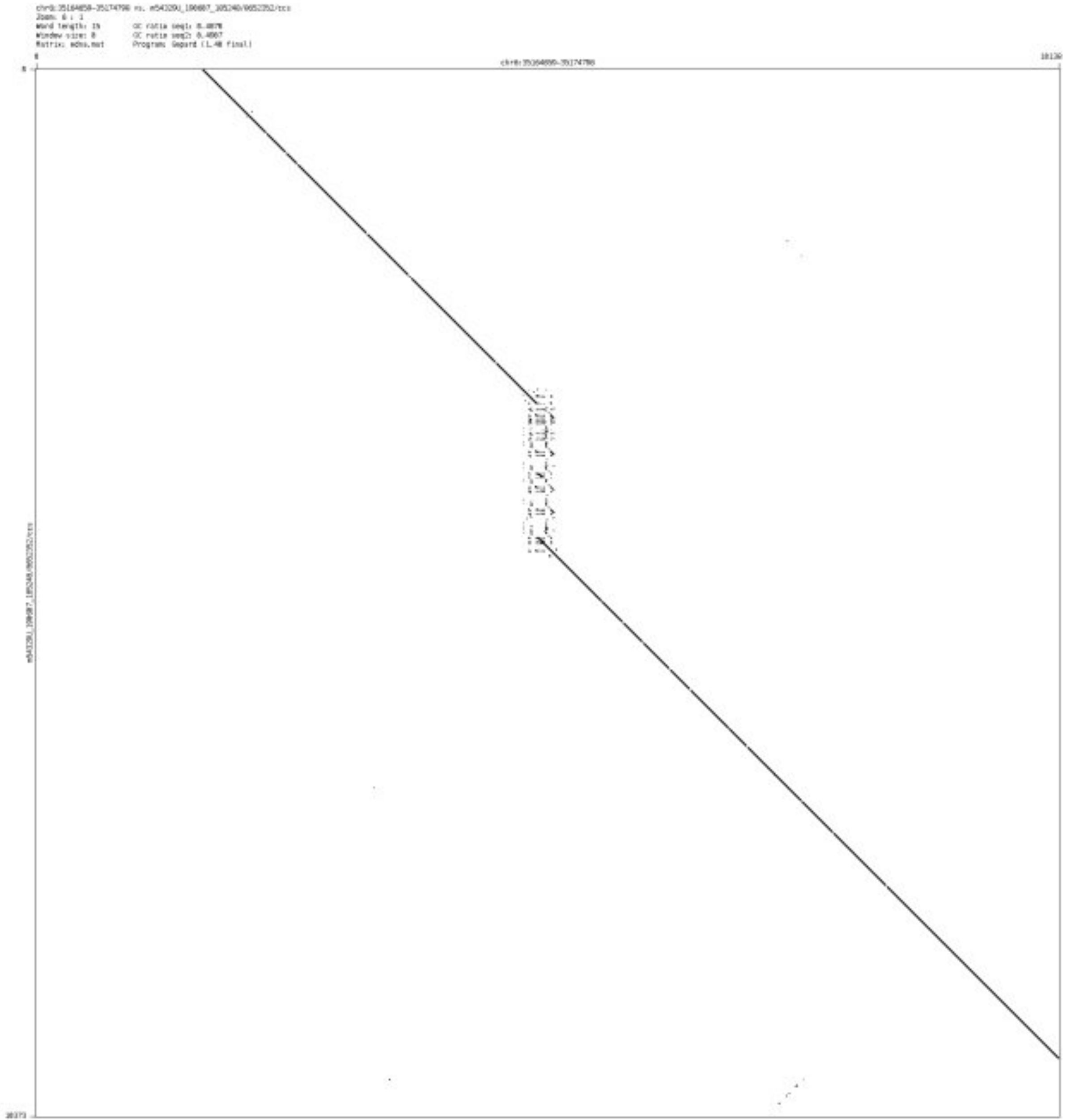

- chr6_44044655_44045175

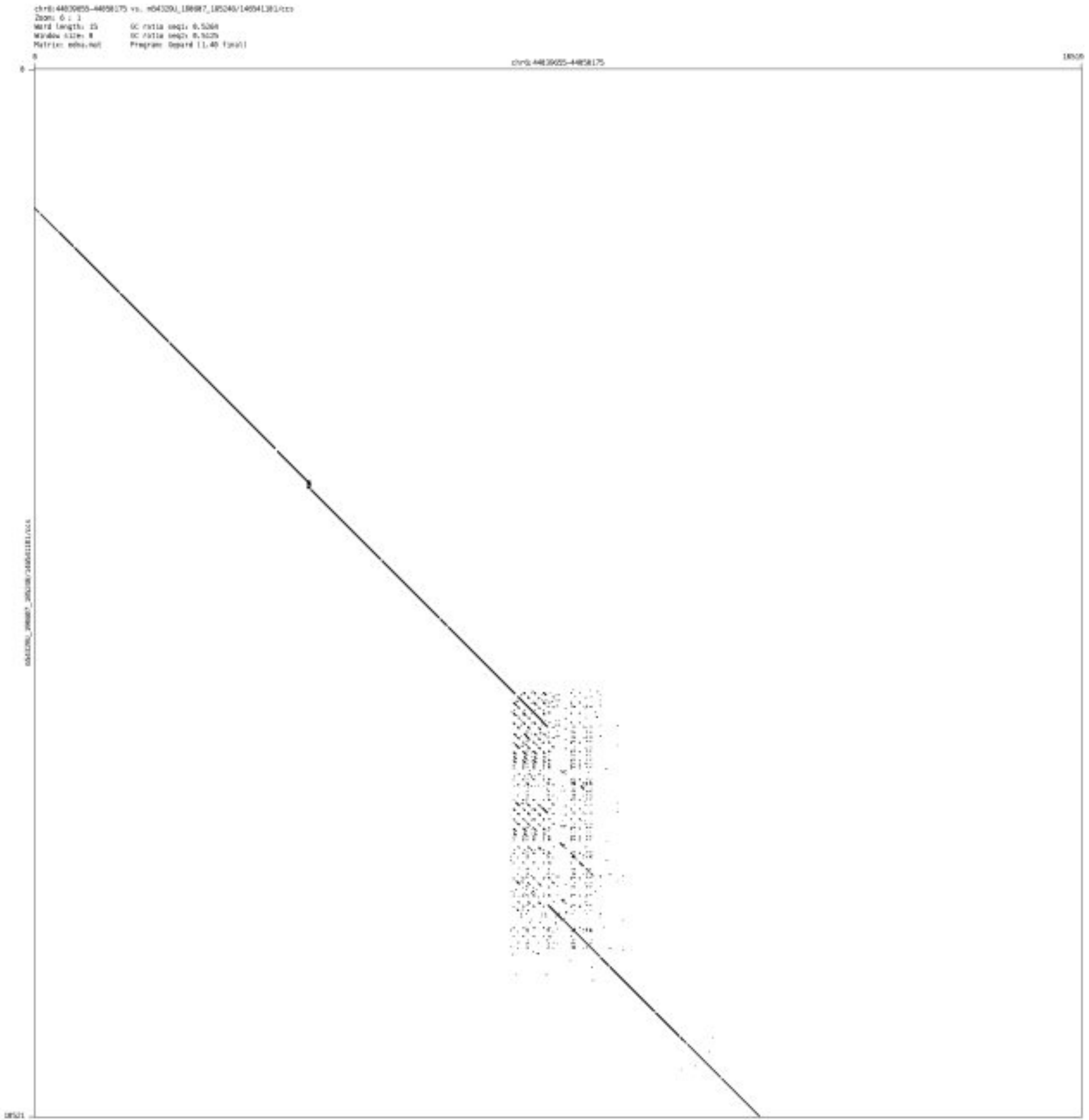

- chr6_117923609_117924524

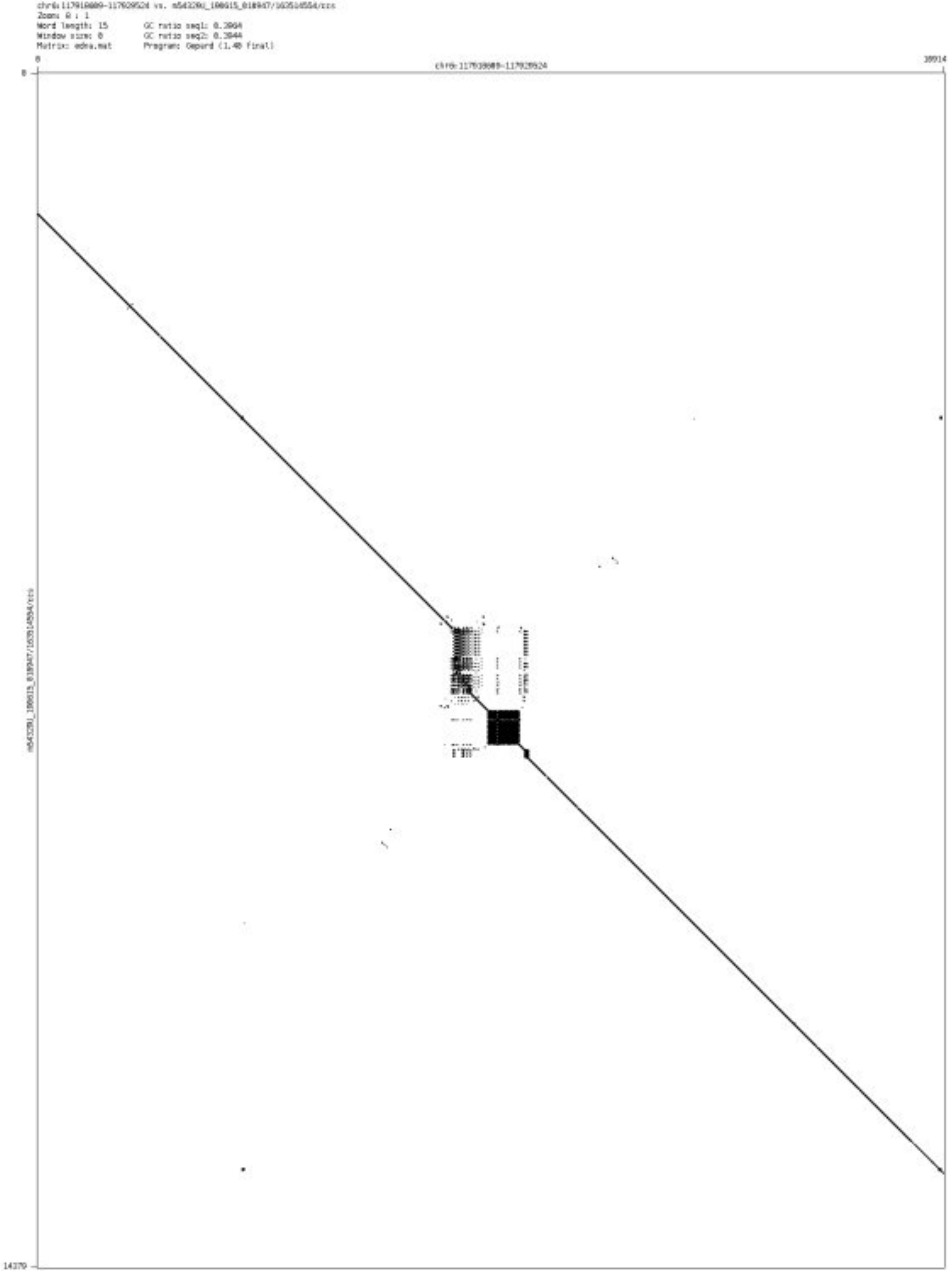

➢ chr6_118070944_118071373

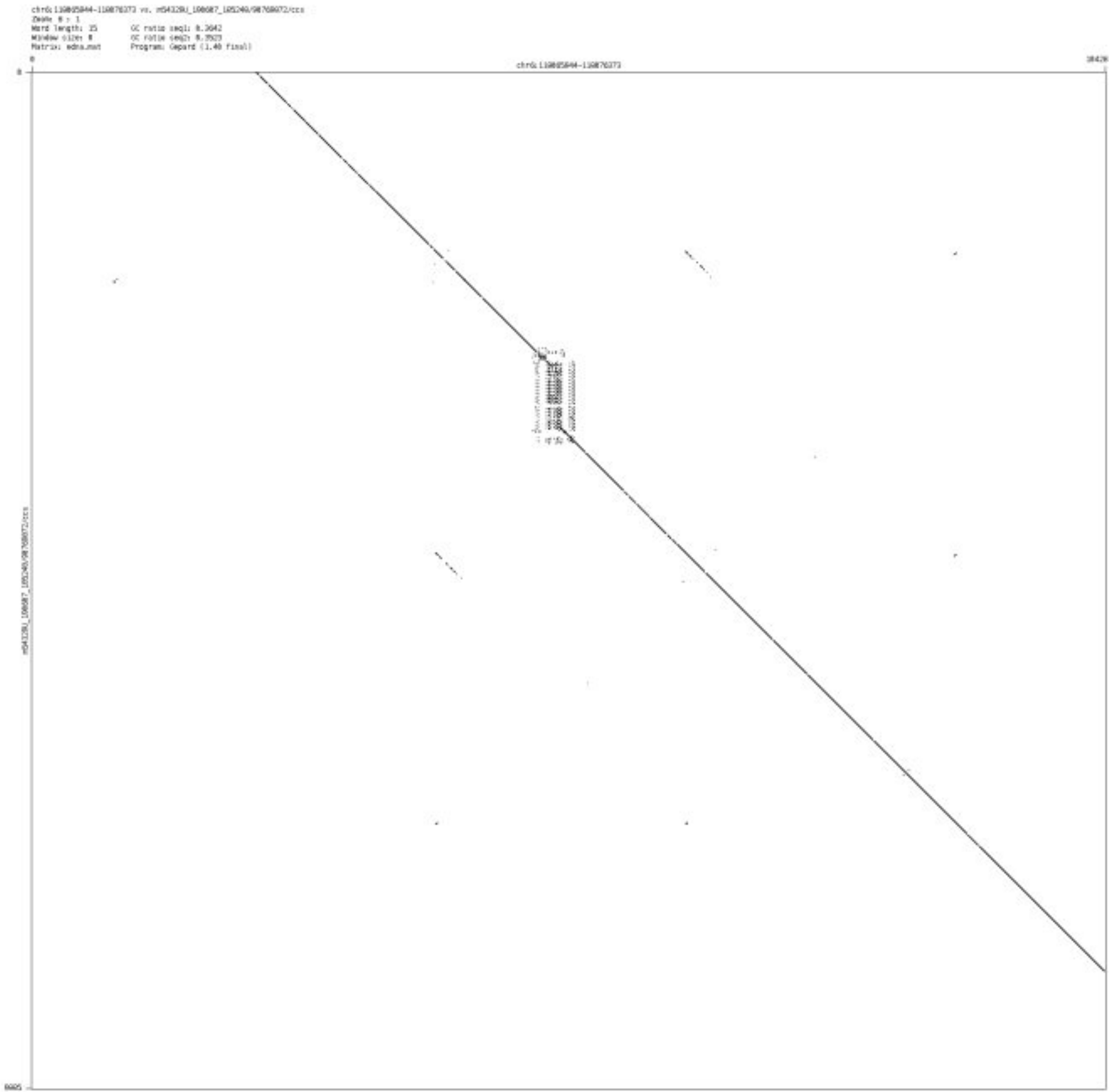

➢ chr6_132600960_132601126

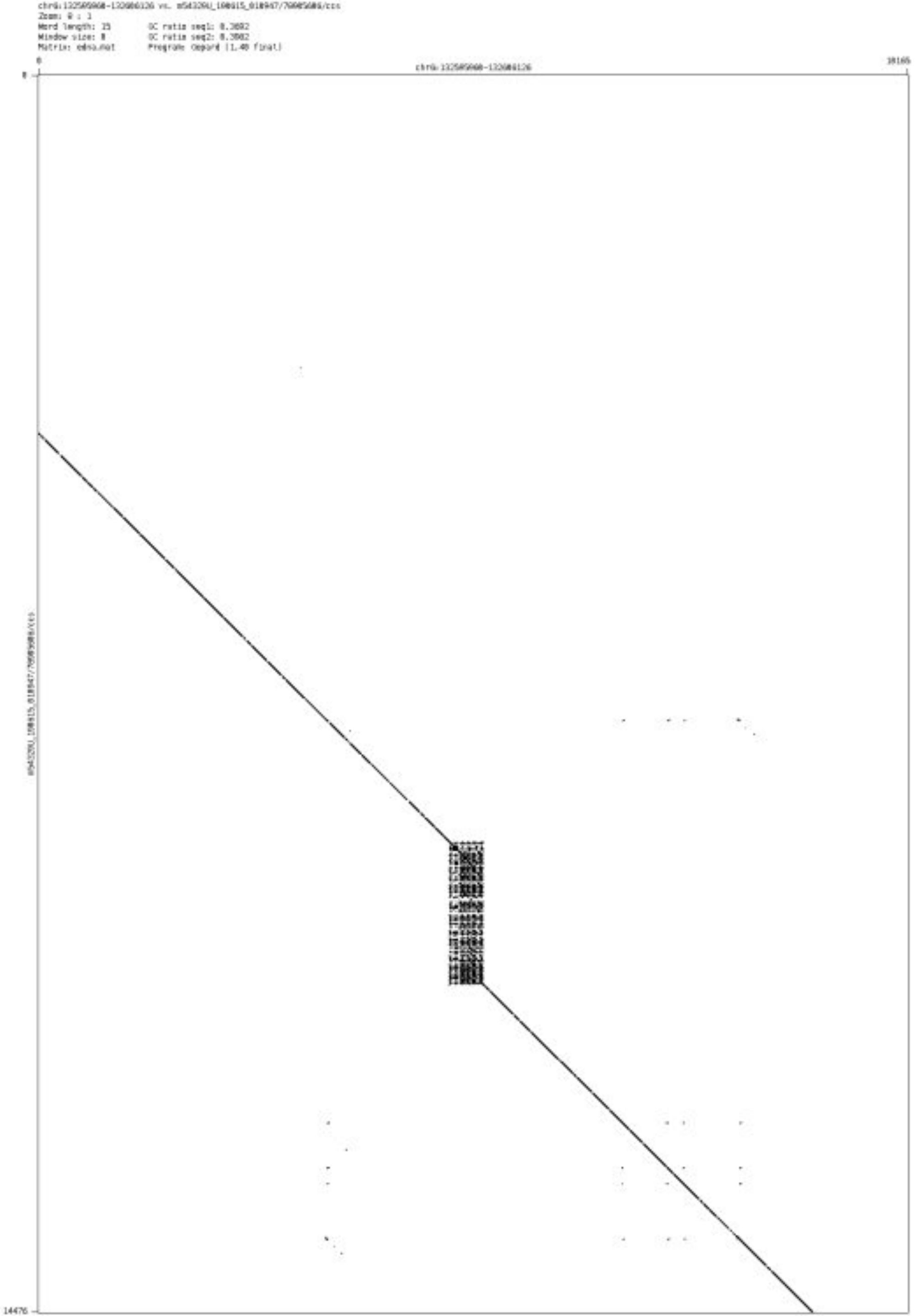

➢ chr6_154525160_154525403

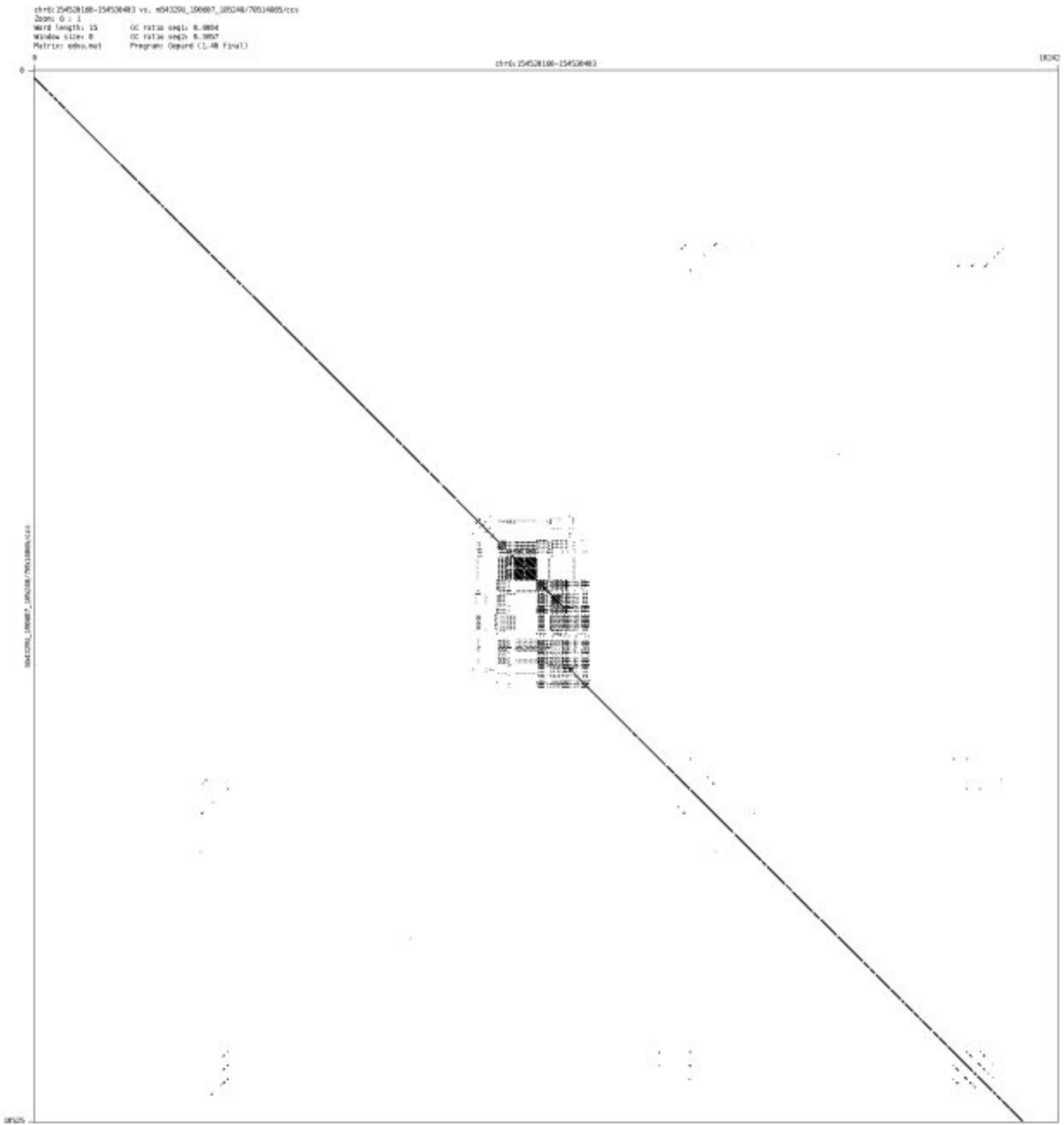

- chr6_168839698_168841851

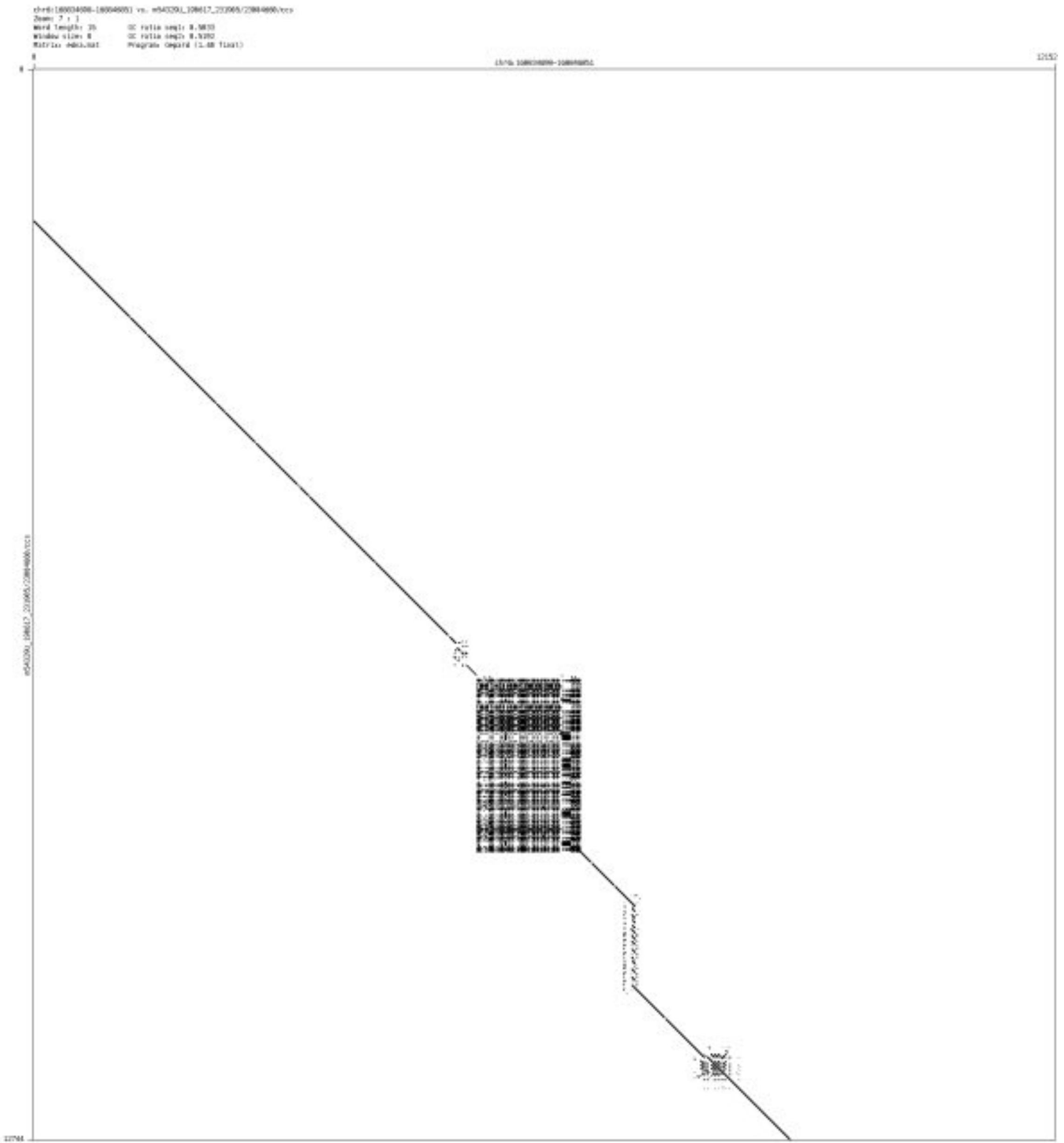

- chr6_168931371_168932485

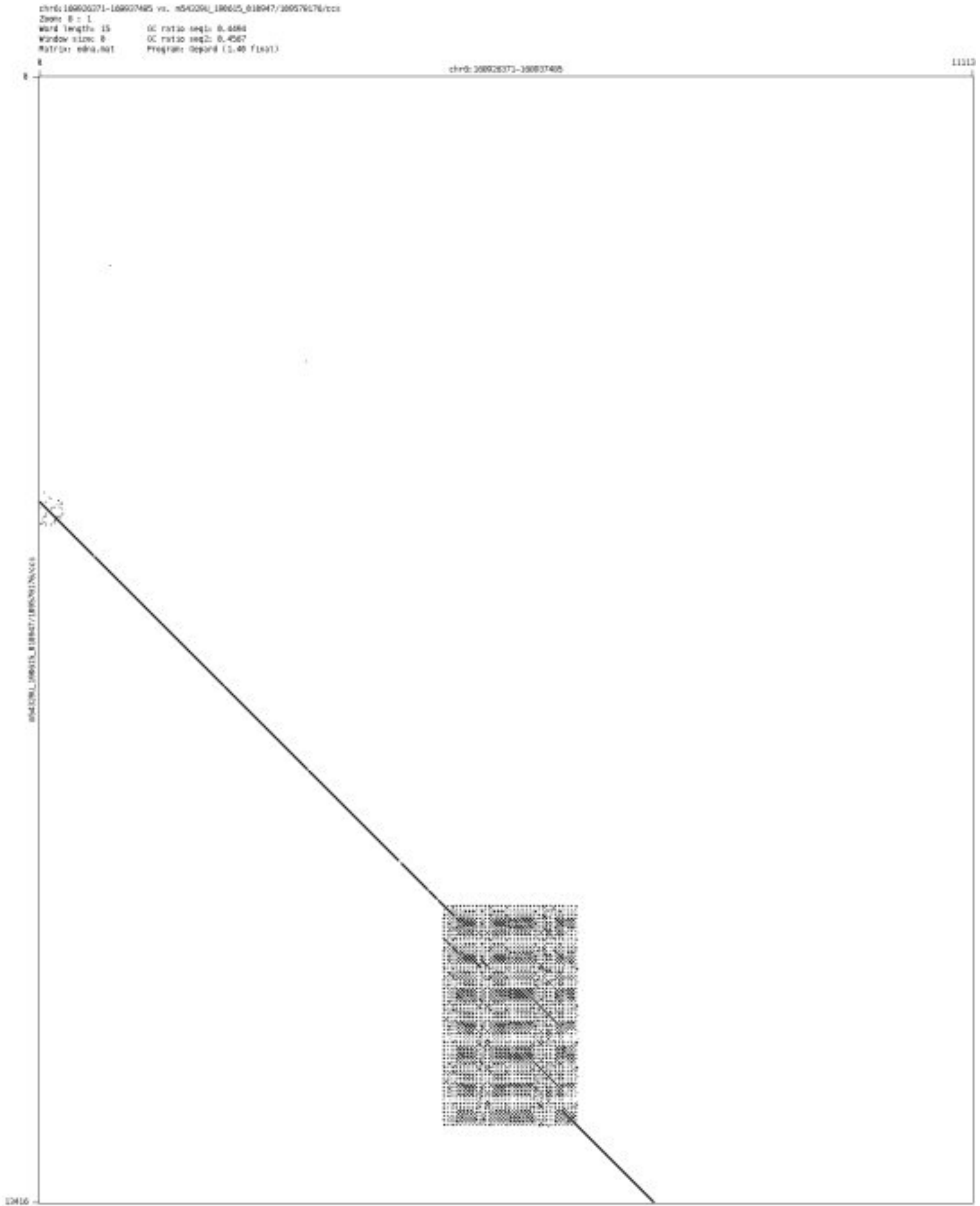

- ➢ chr6_169242827_169243076

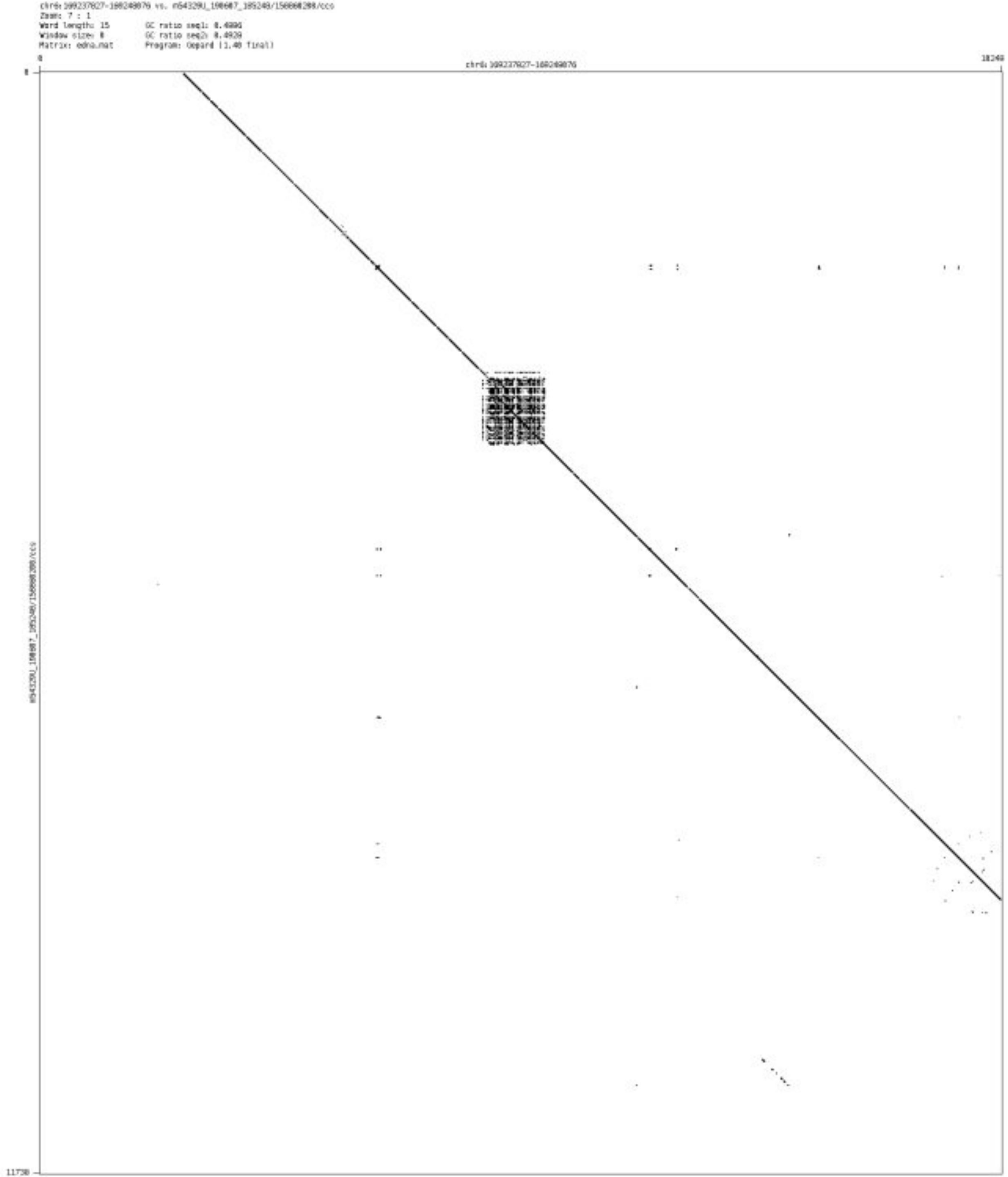

- chr6_170398827_170400228

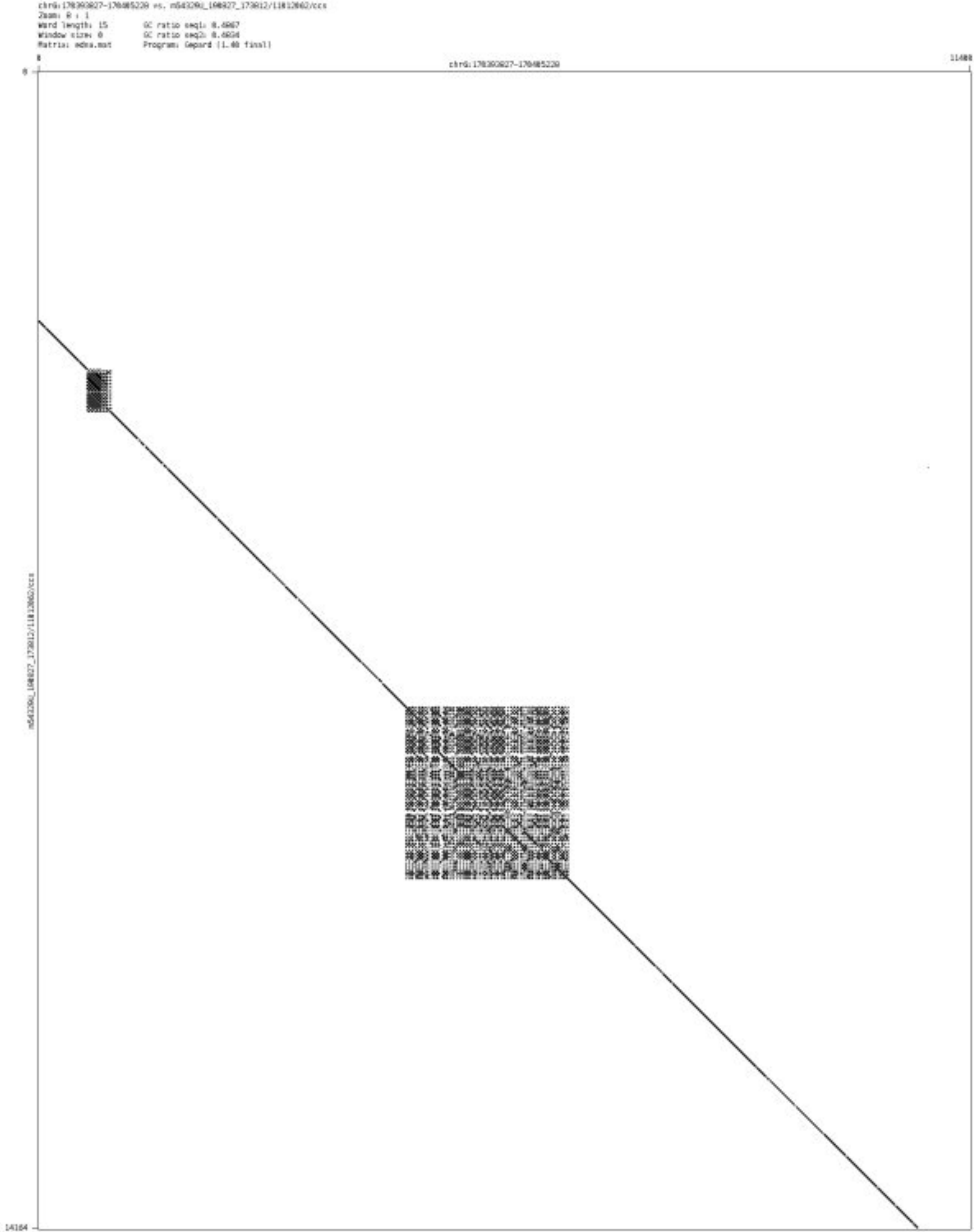

- chr7_355960_356050

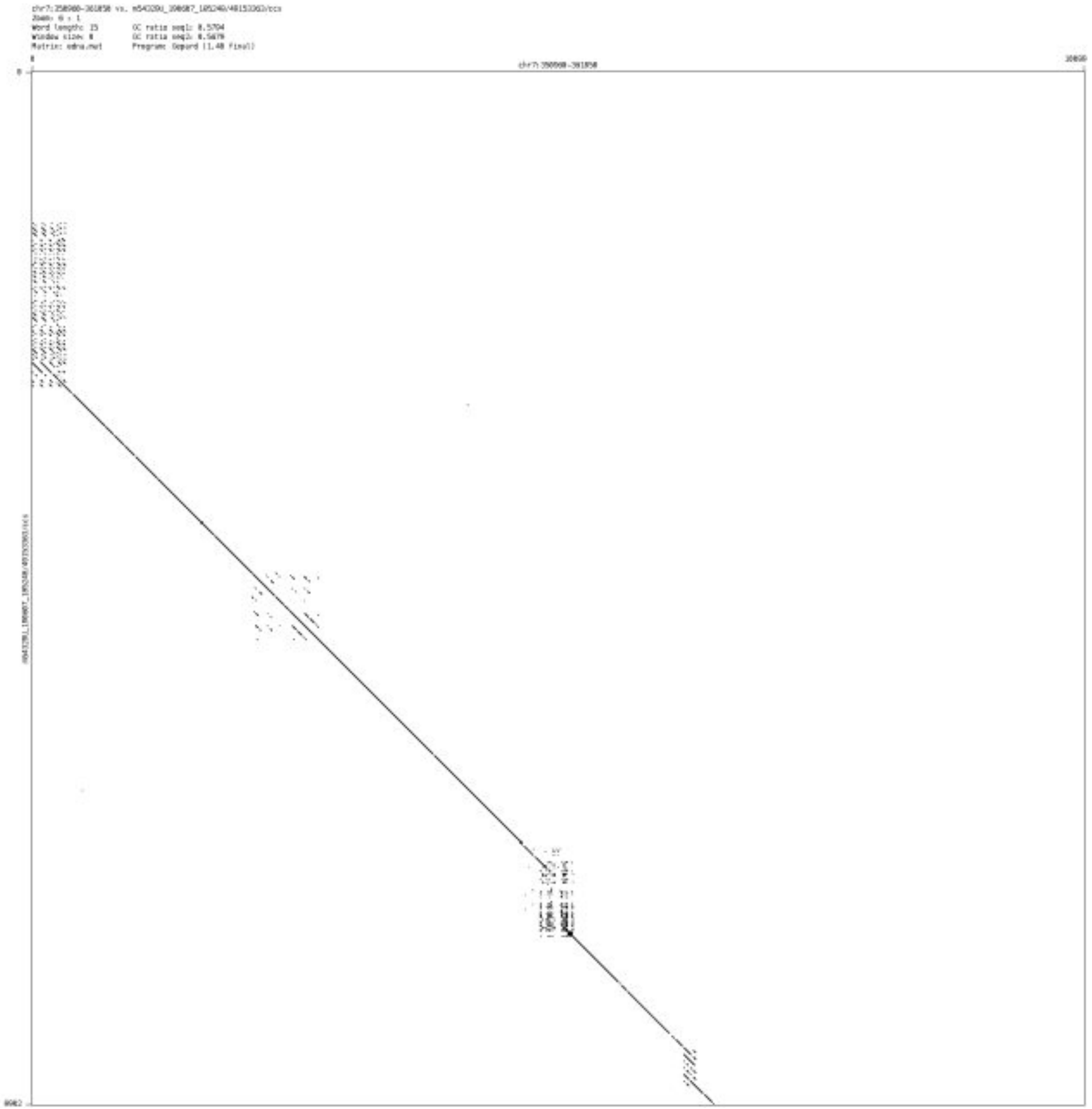

➢ chr7_588557_588704

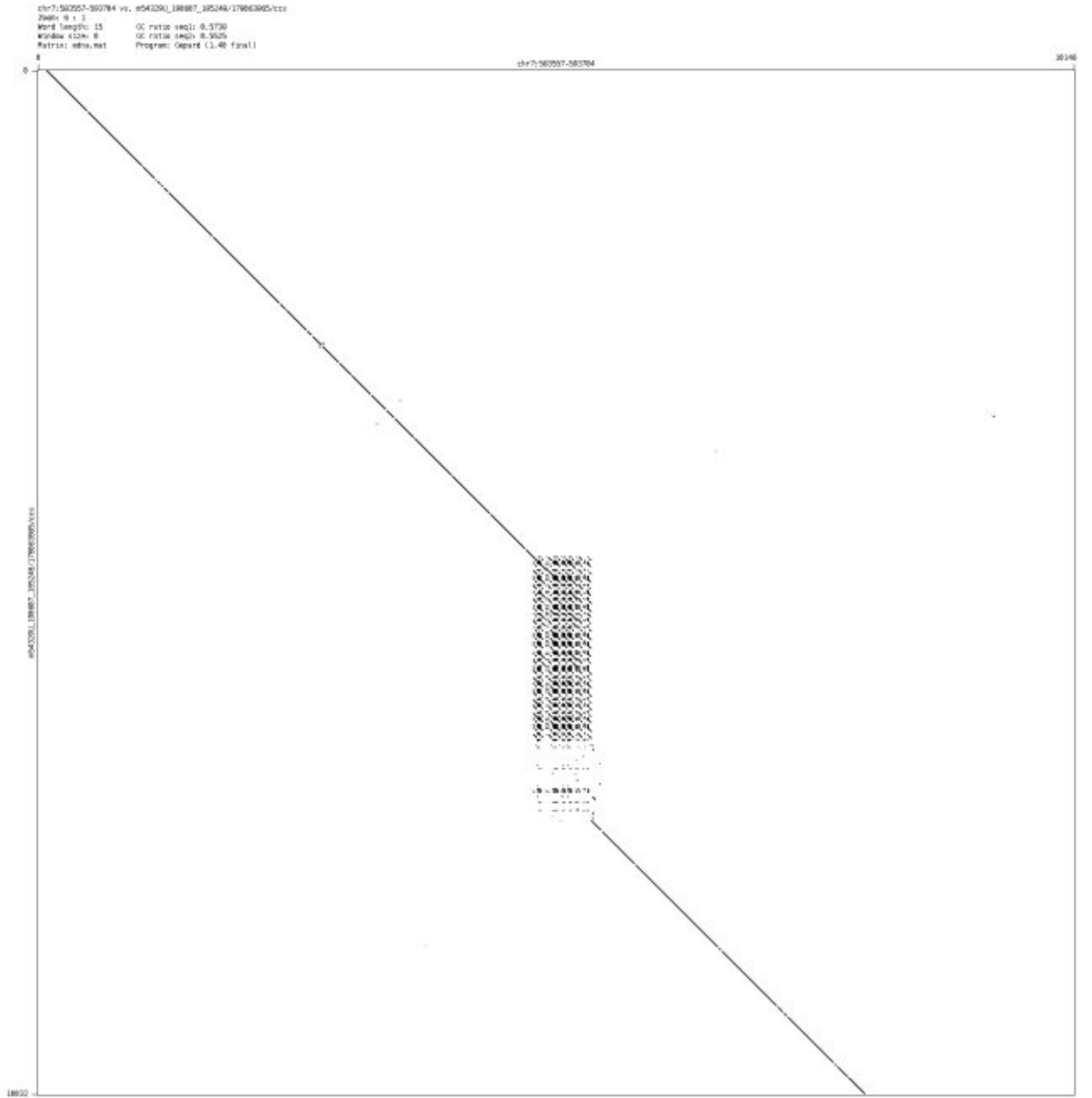

➢ chr7_905588_906538

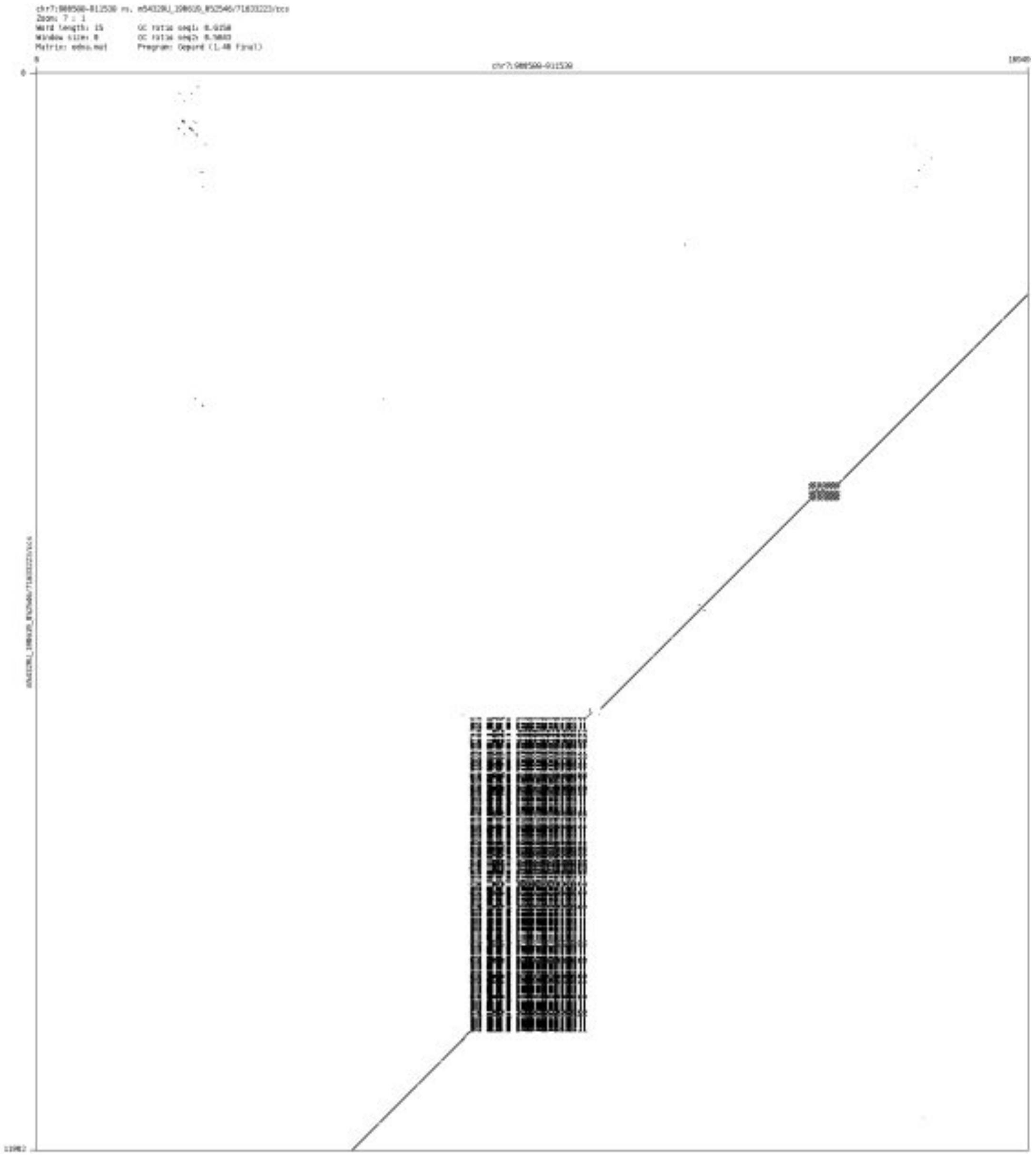

➢ chr7_1758548_1758613

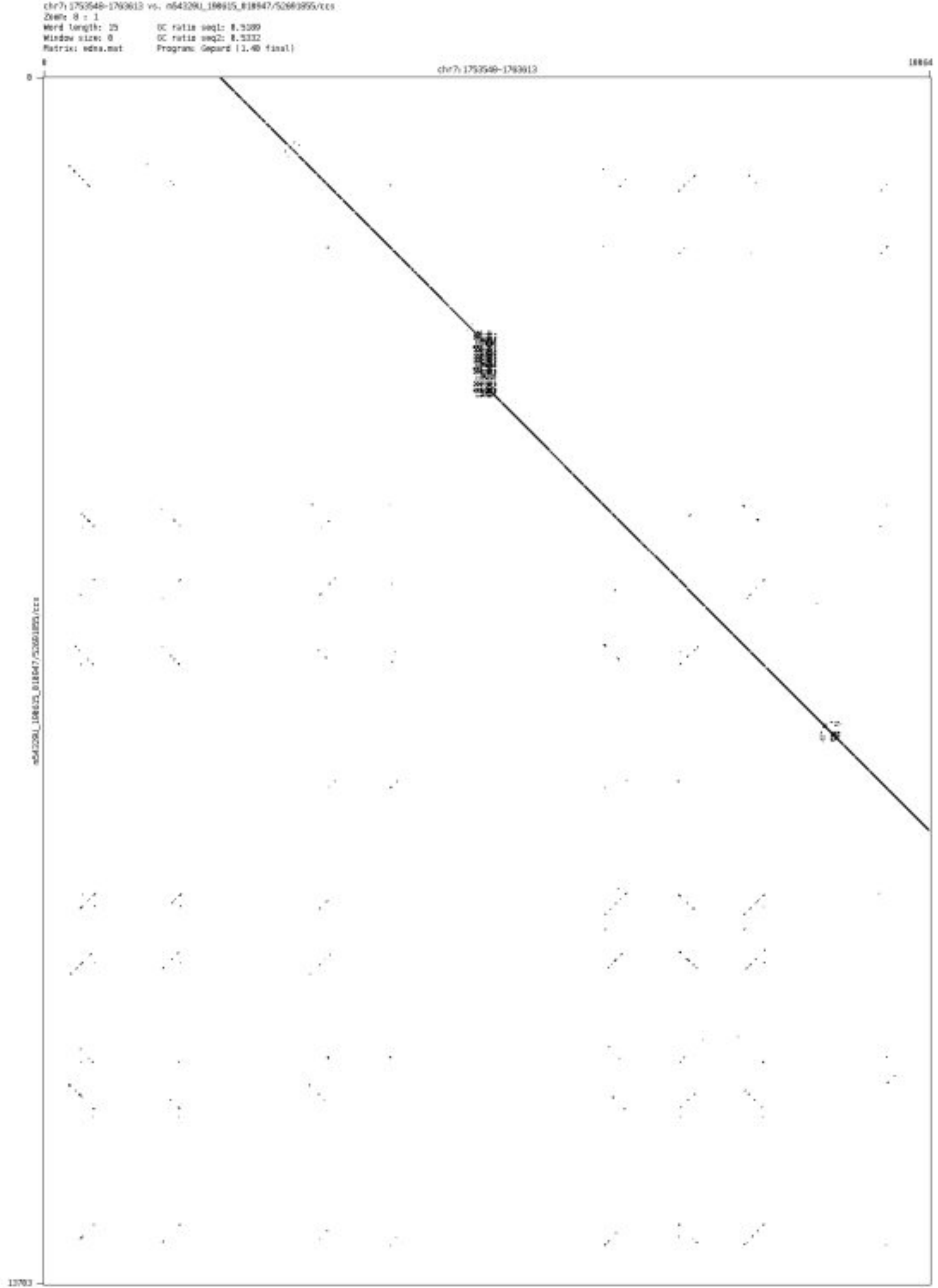

- chr7_1940931_1941009

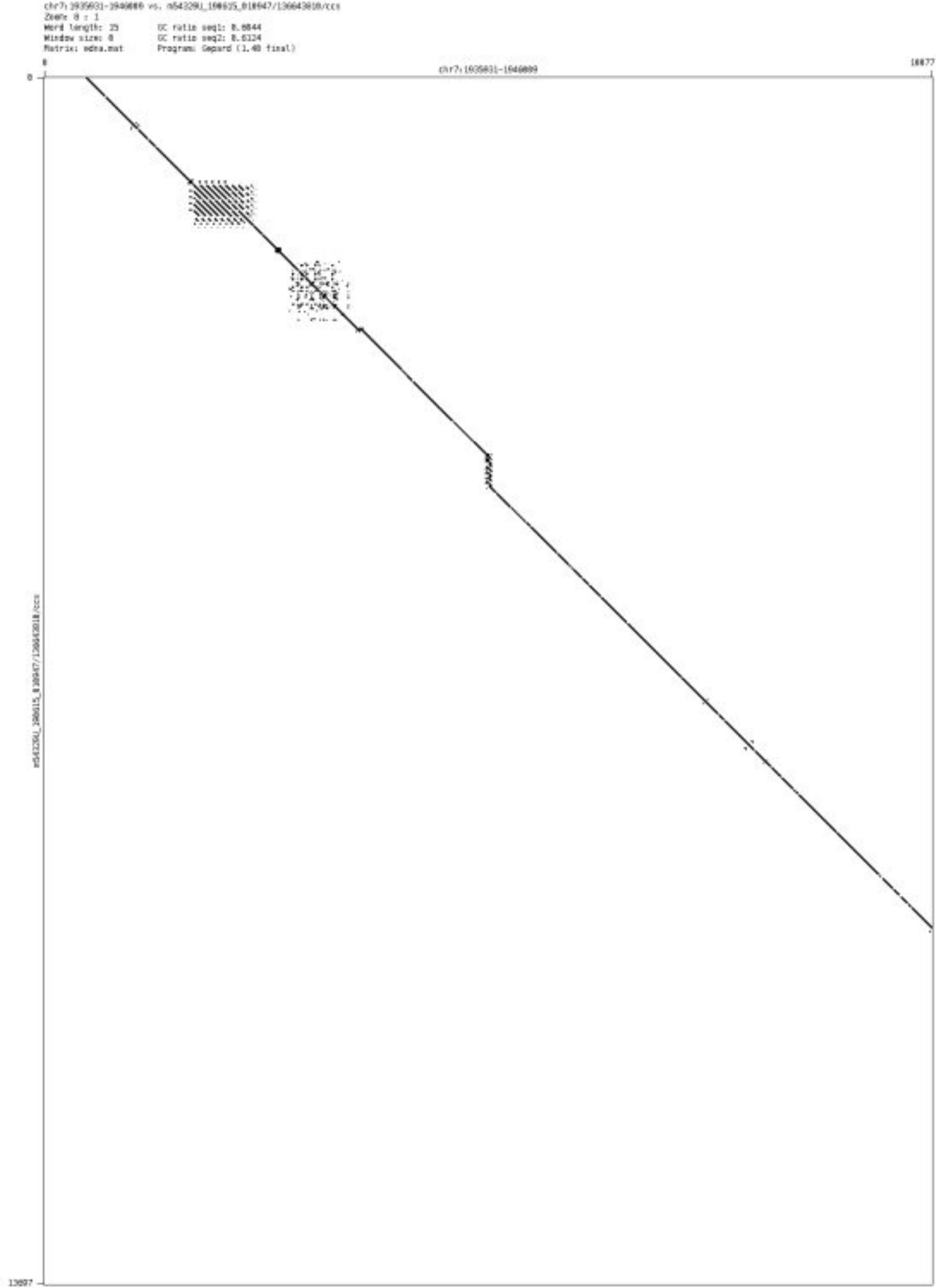

- chr7_138621939_138622106

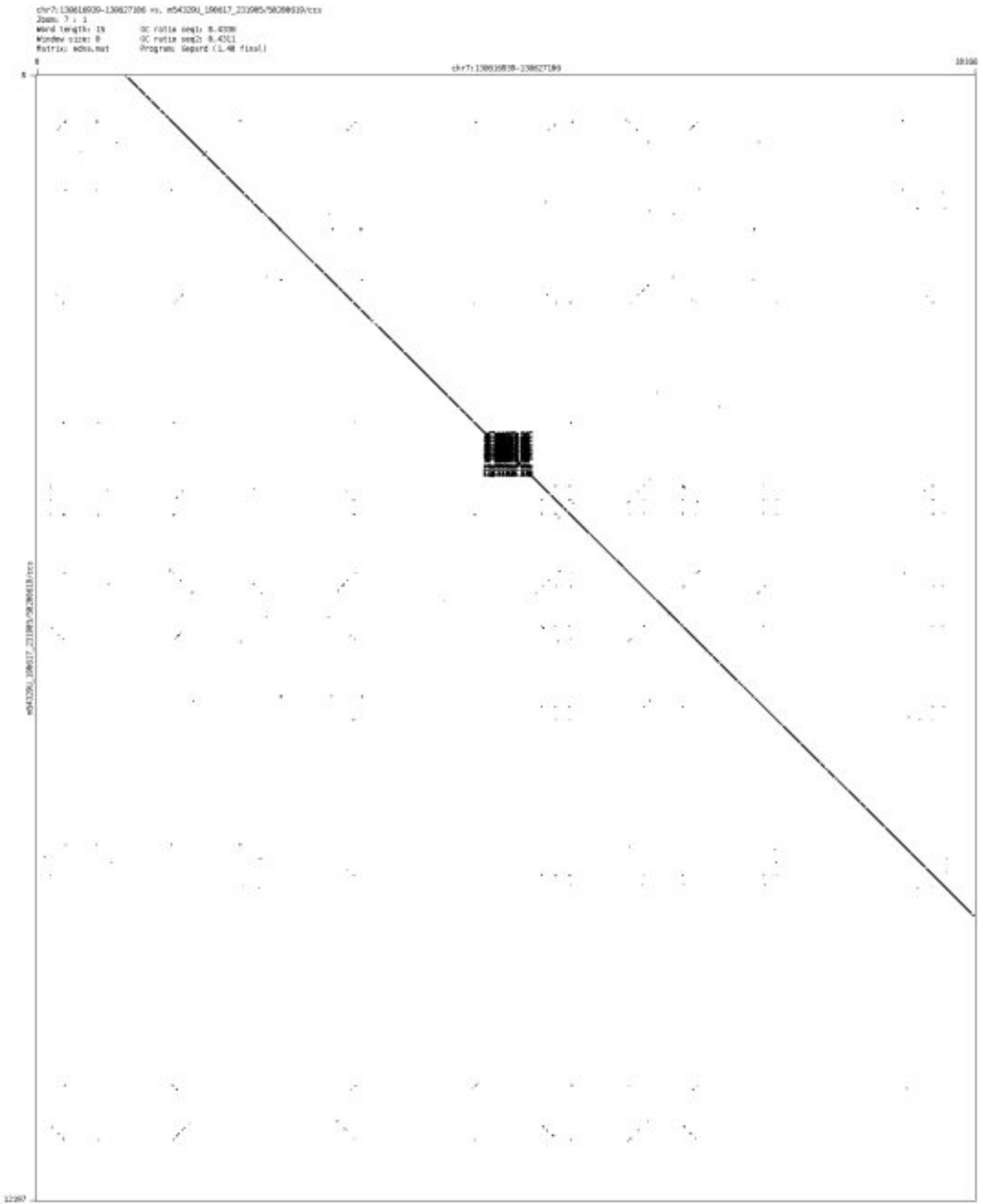

- chr7_150909790_150910469

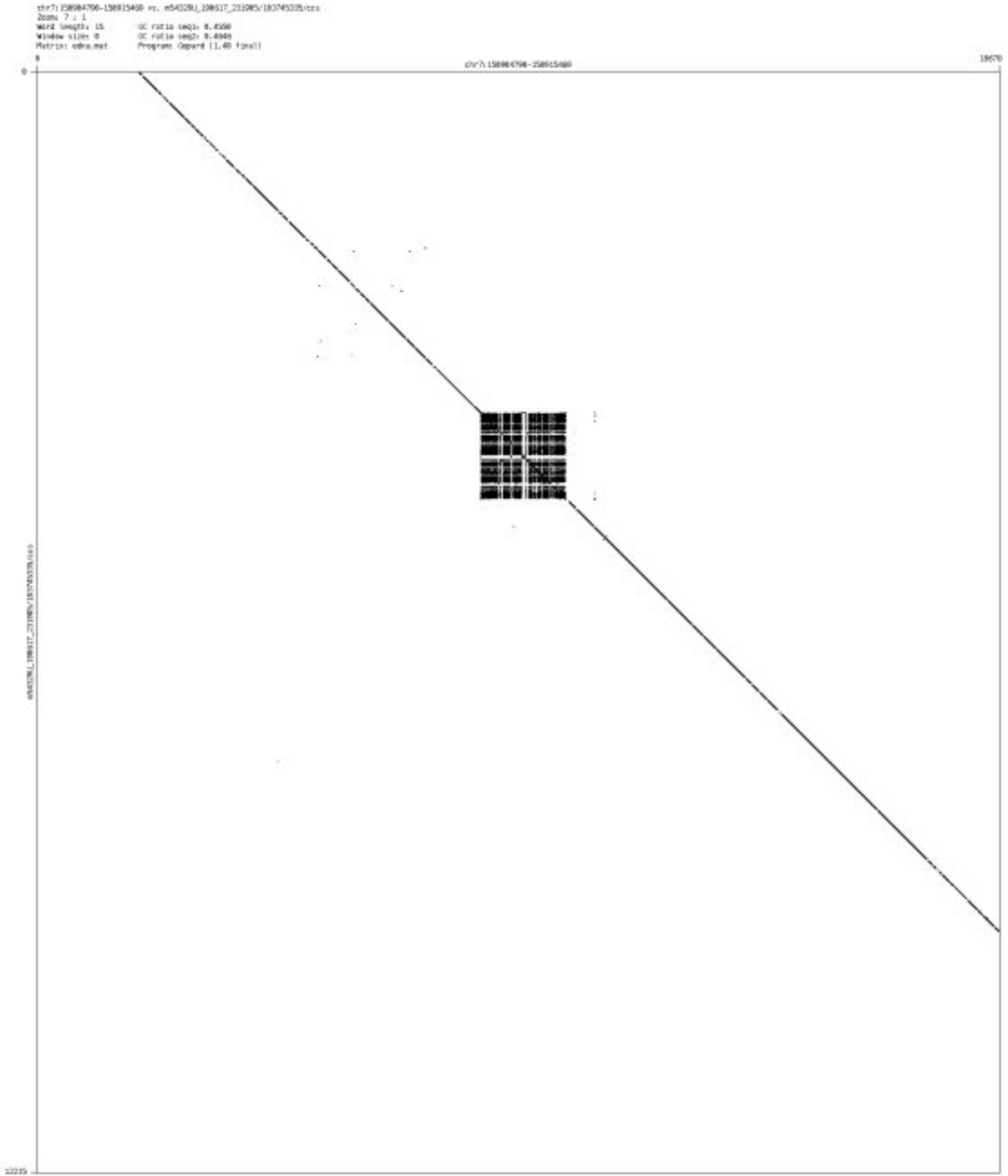

➢ chr7_151855231_151855587

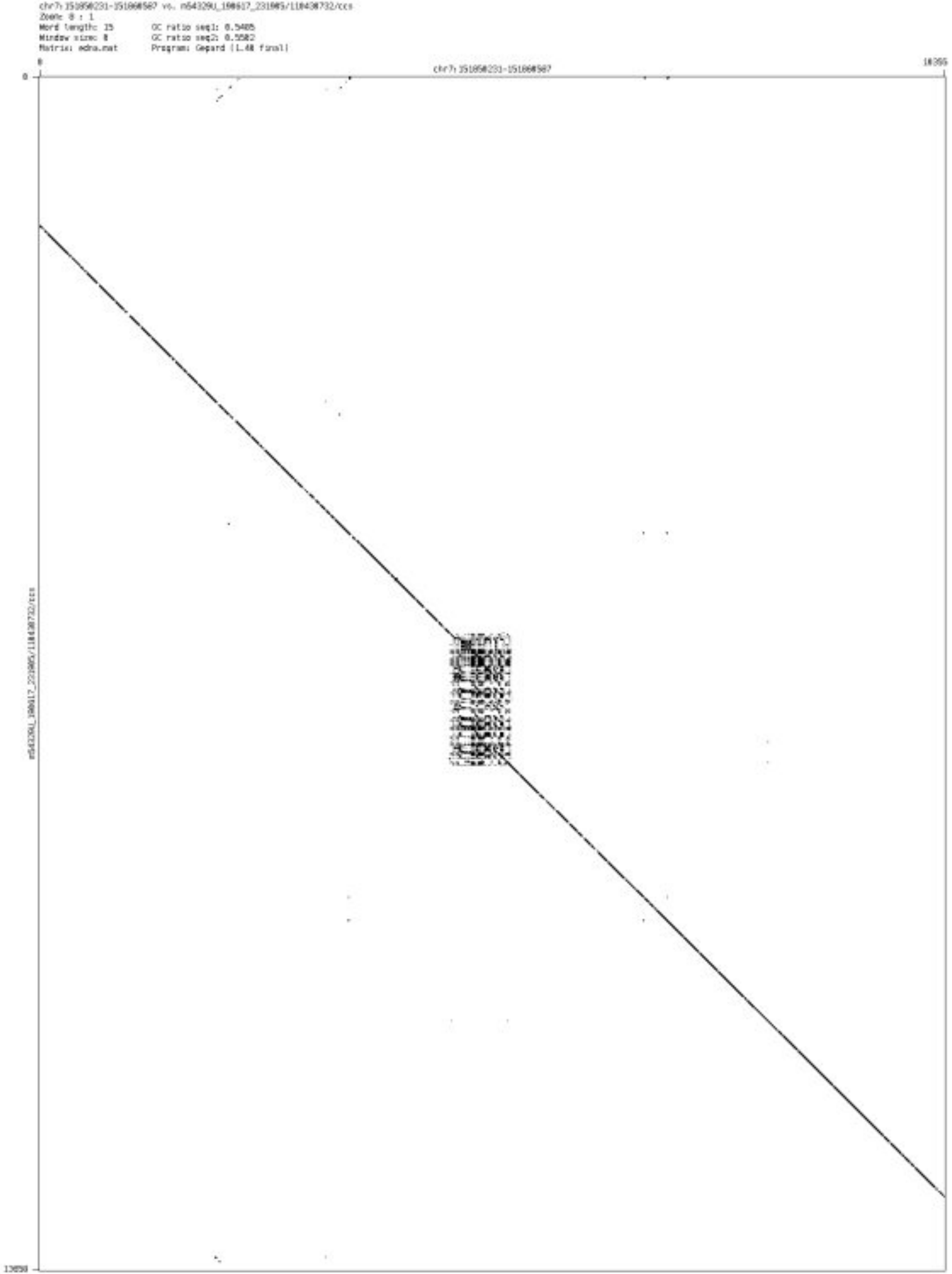

- chr7_155328712_155328857

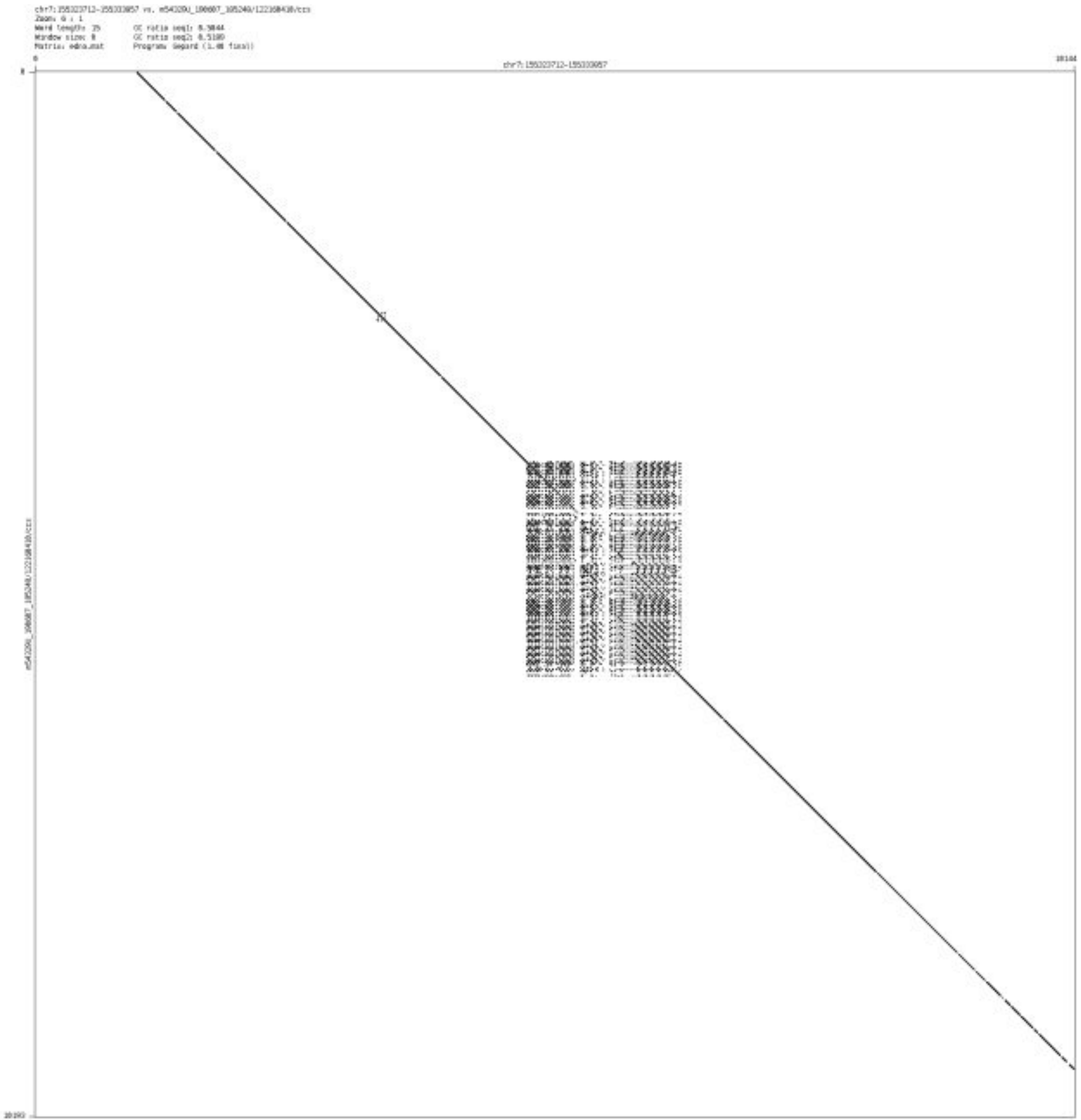

➢ chr7_155406971_155407121

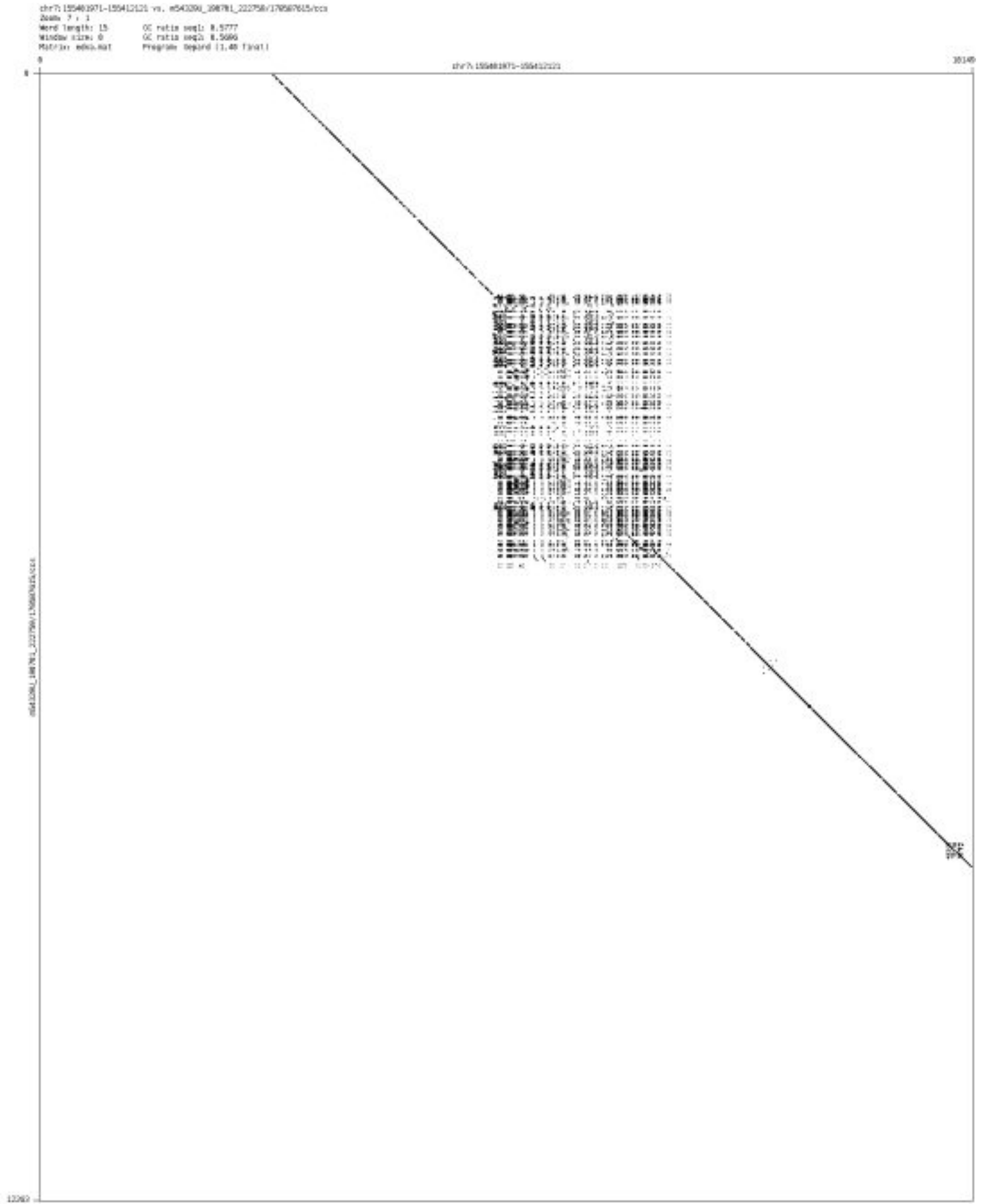

- ➢ chr7_158151379_158151489

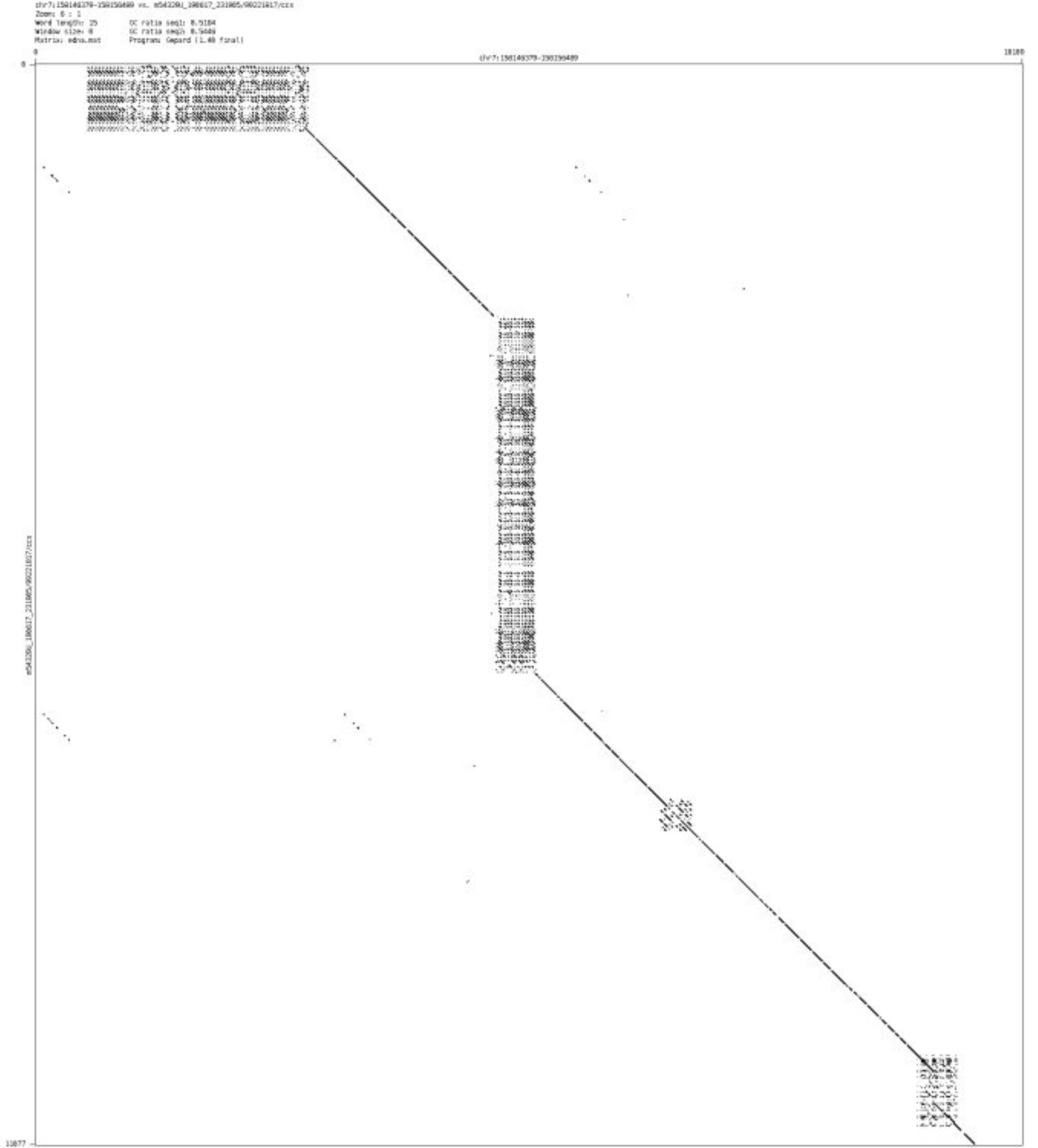

- chr7_158668126_158668212

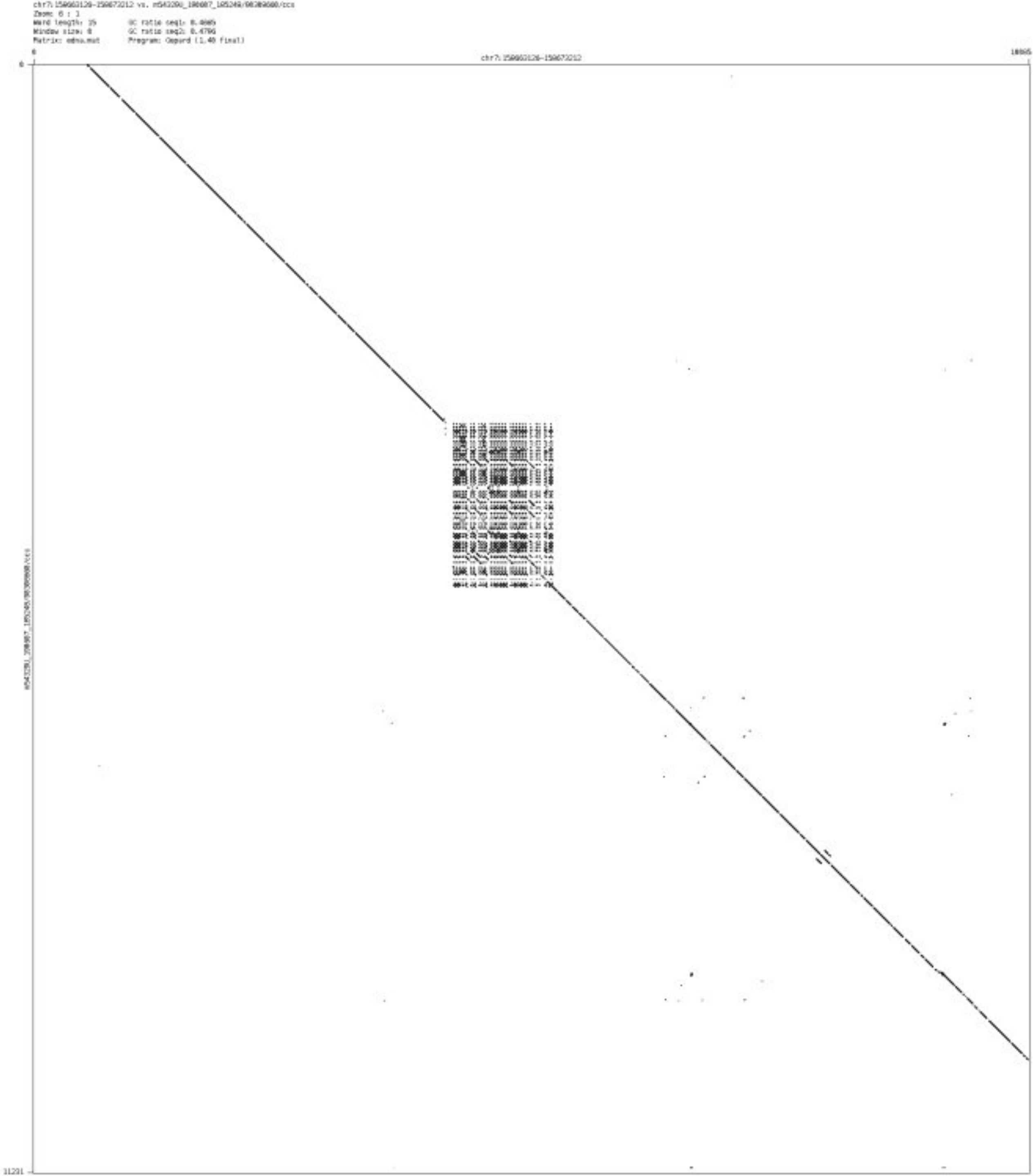

- chr7_158998436_158998524

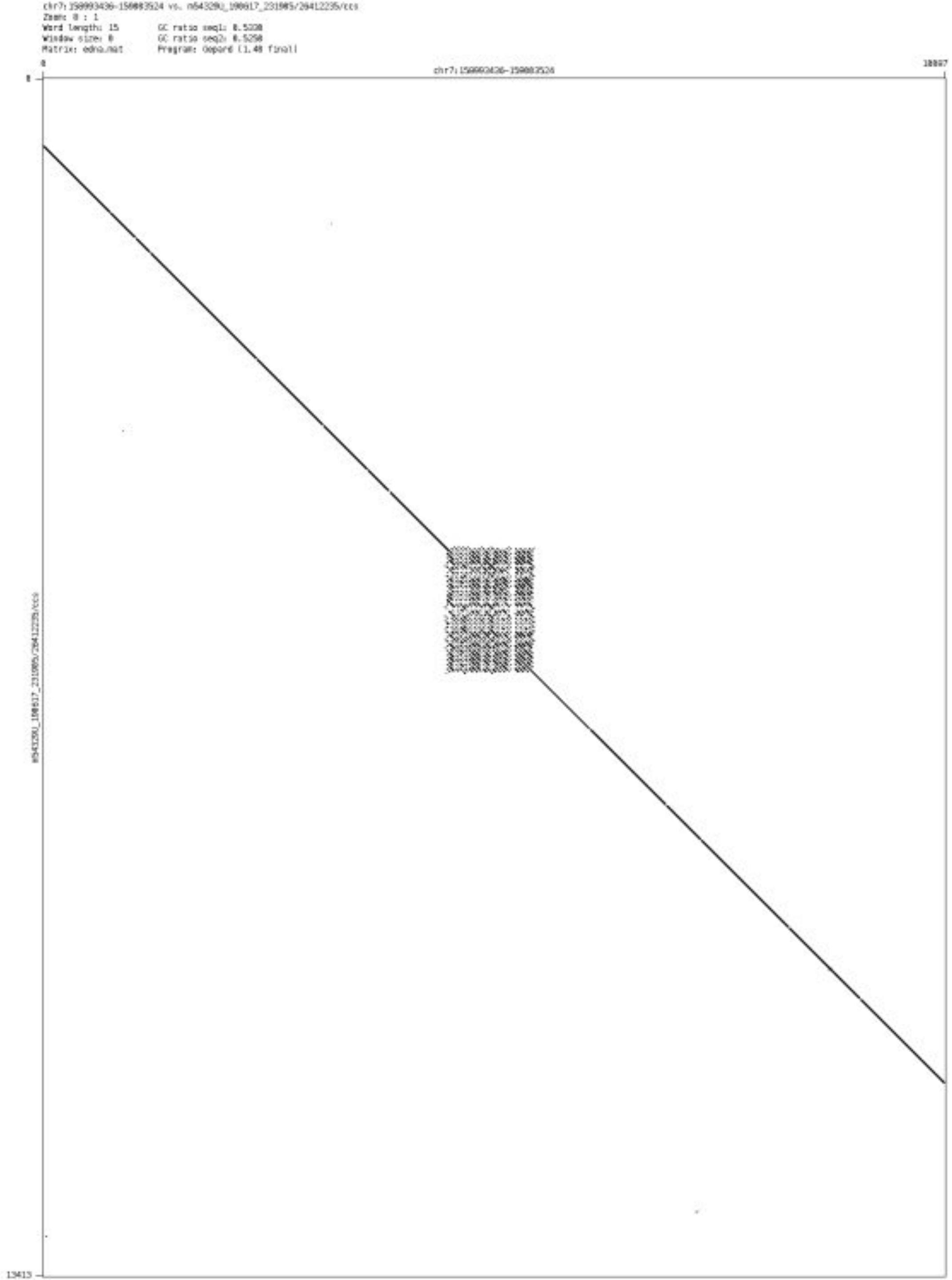

- chr8_37506281_37506358

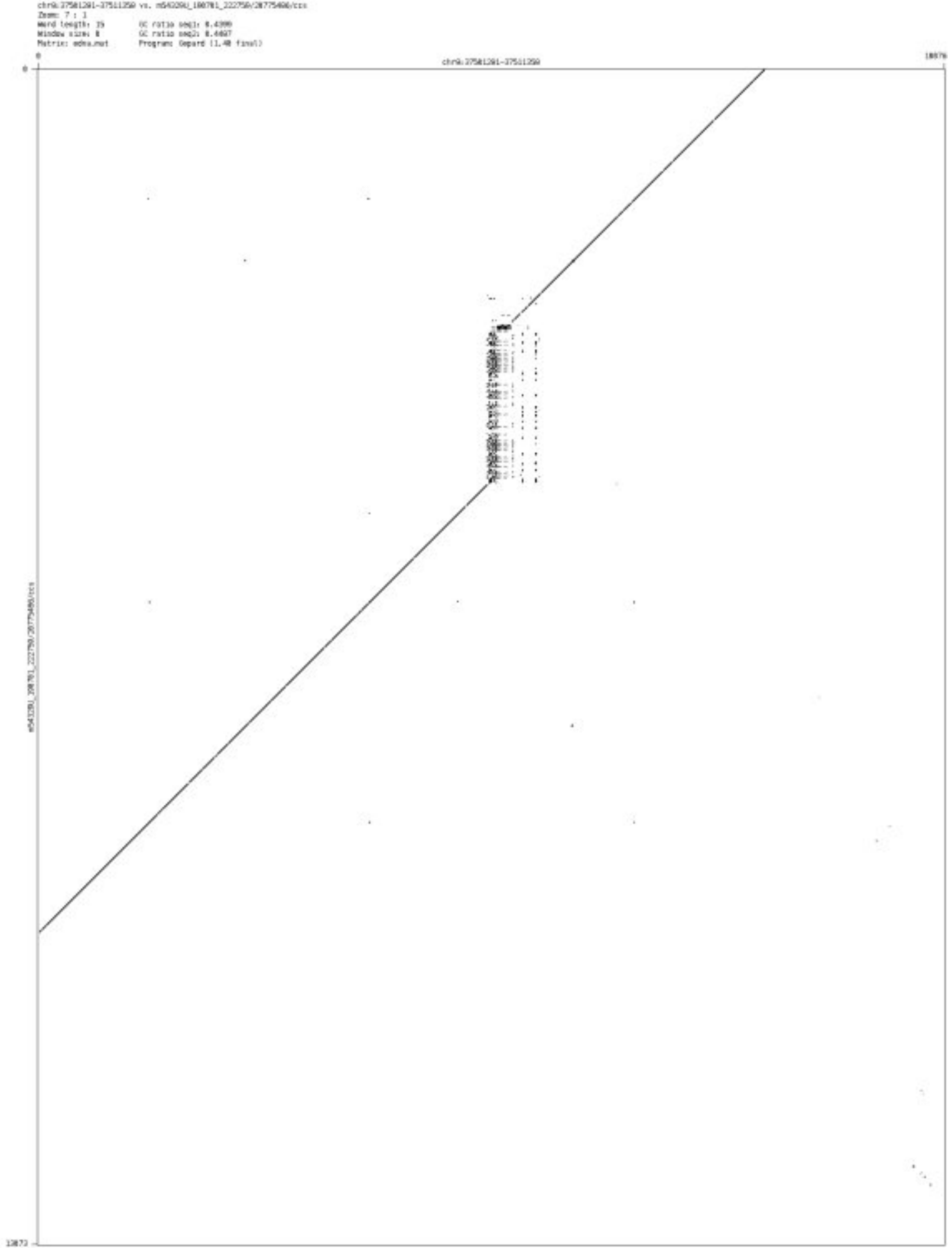

#### ➢ chr8_68906235_68906443

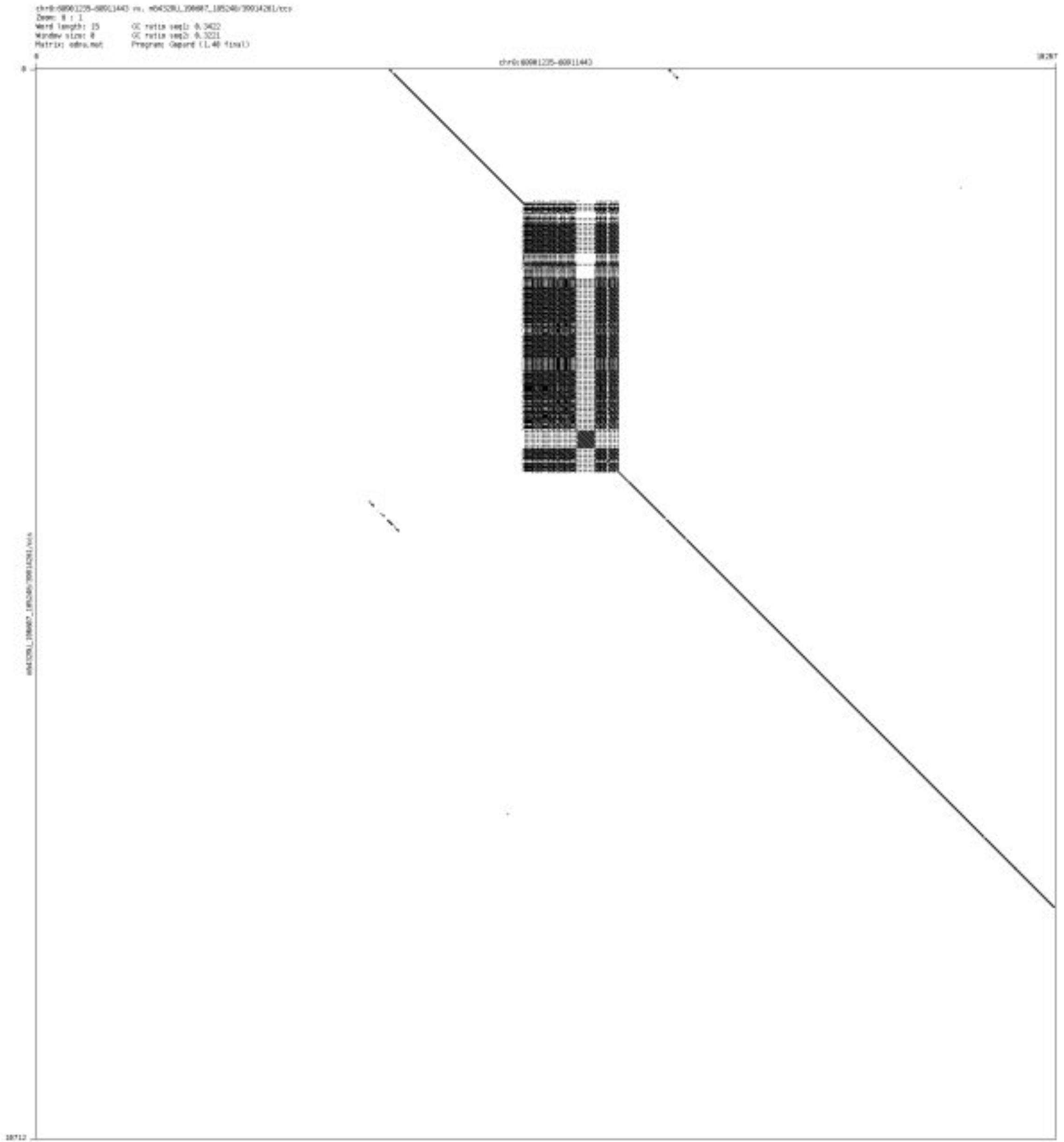

➢ chr8_76444618_76444671

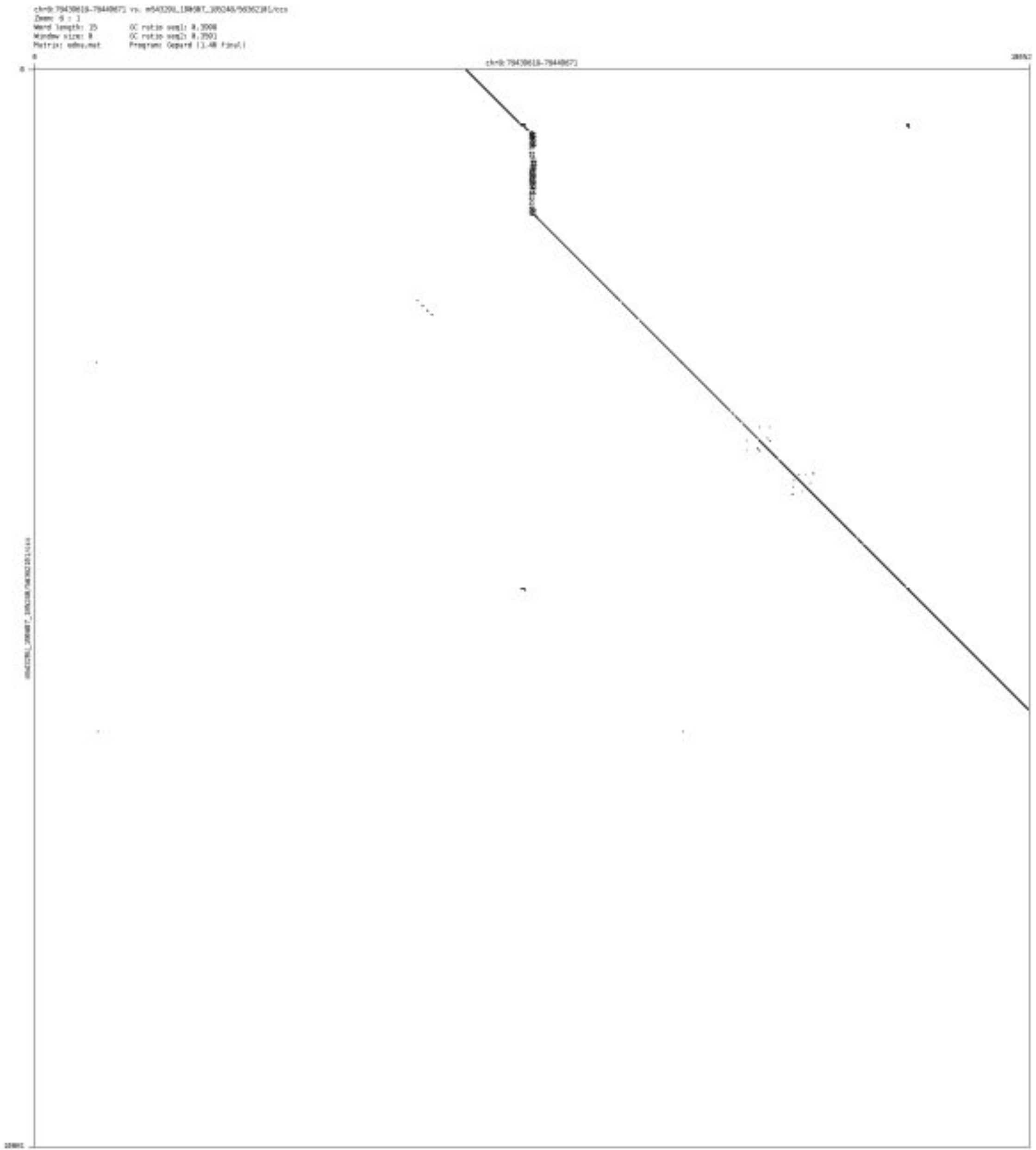

- chr8_100426632_100426724

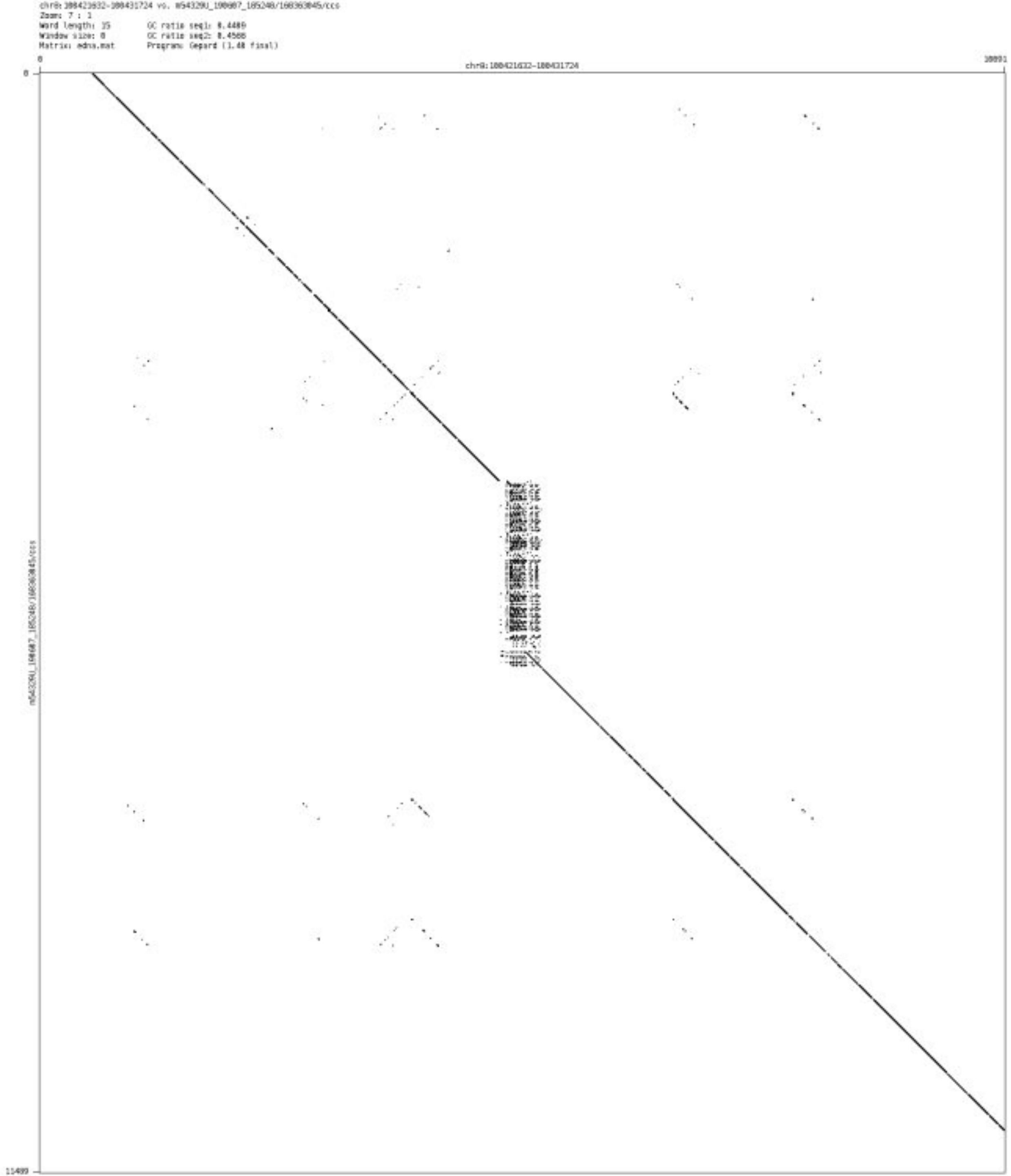

- chr8_141492292_141492474

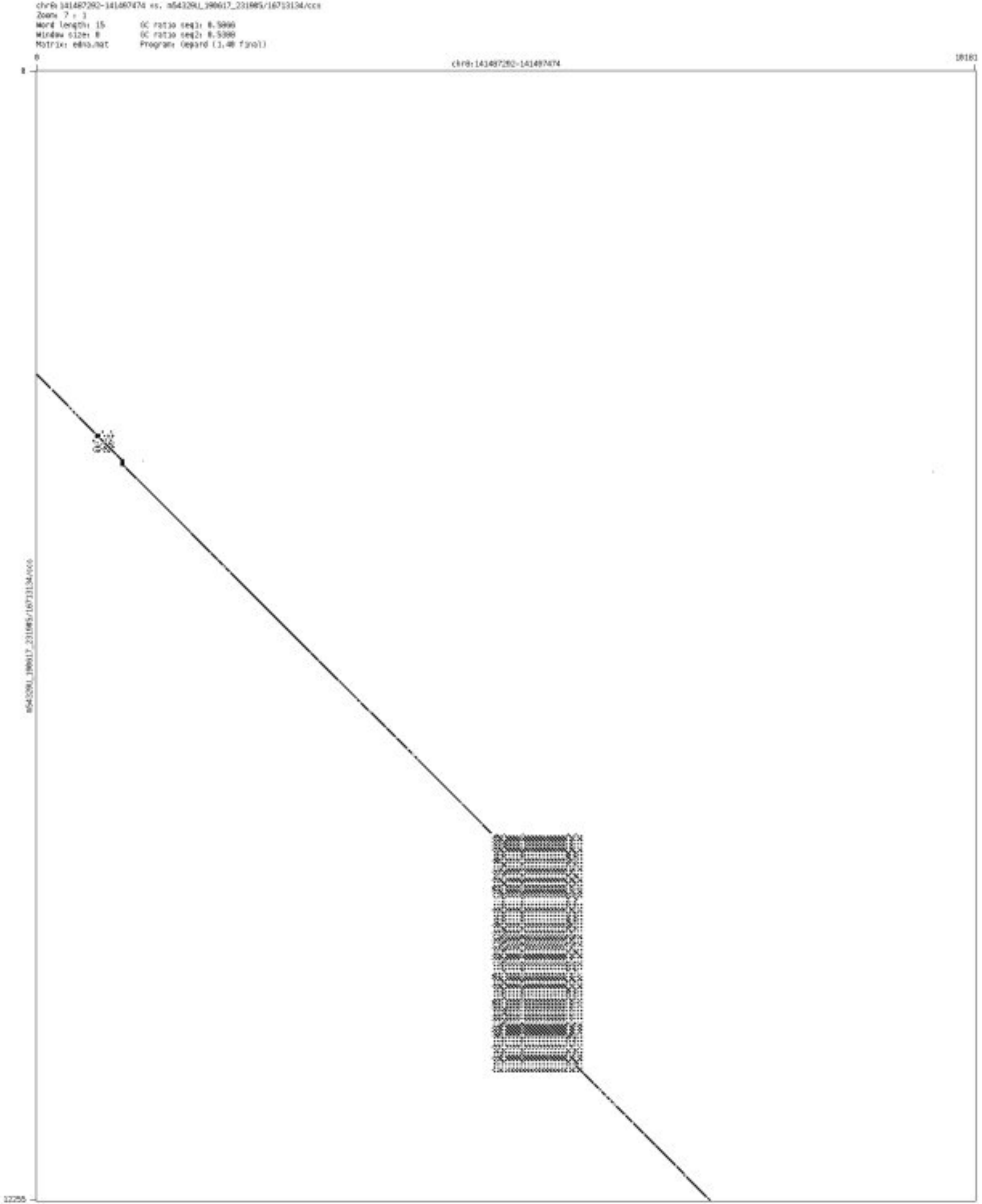

➢ chr8_143218262_143218337

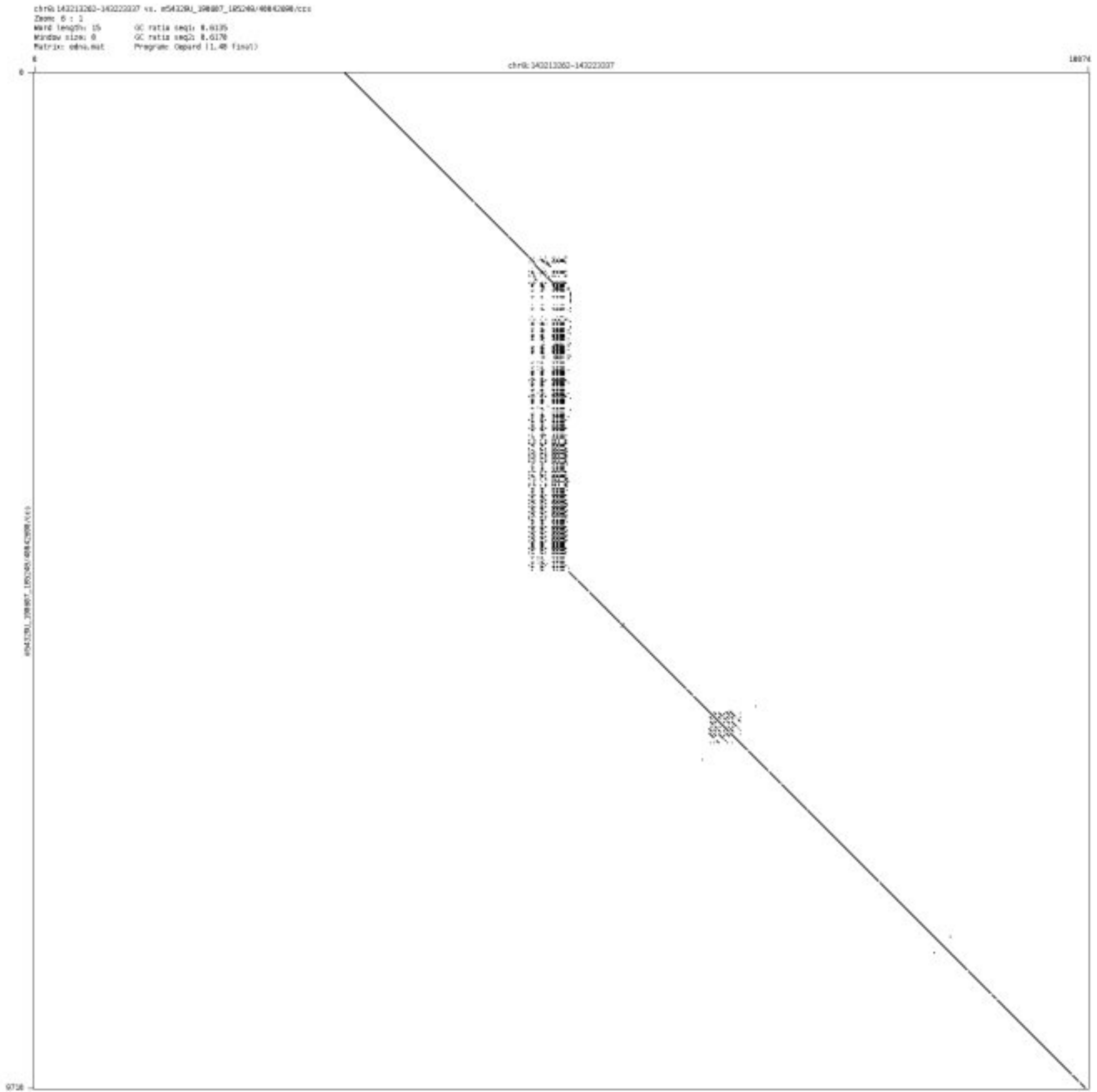

- chr8_143669463_143669602

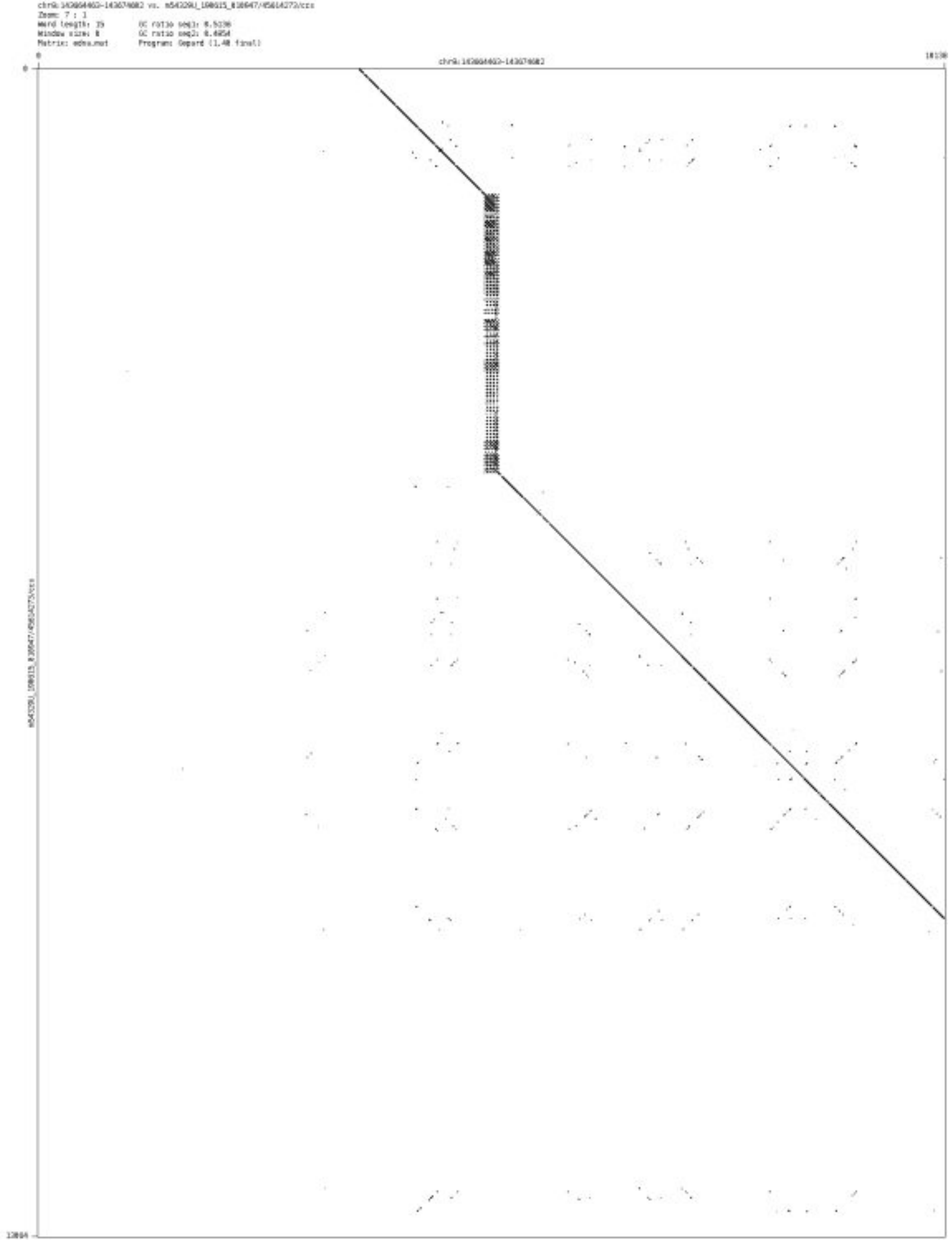

- chr8_144125777_144125935

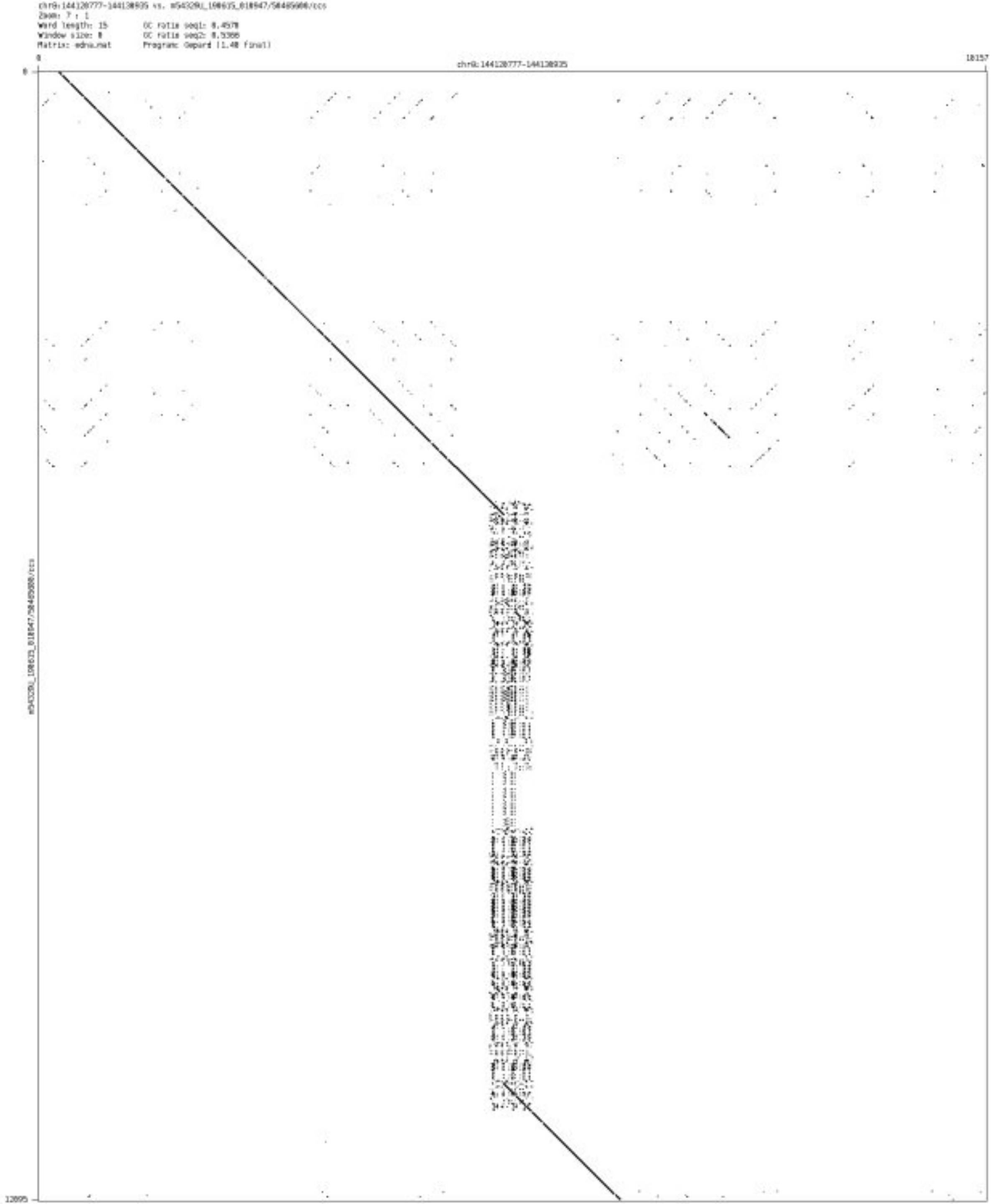

- chr8_144233387_144233472

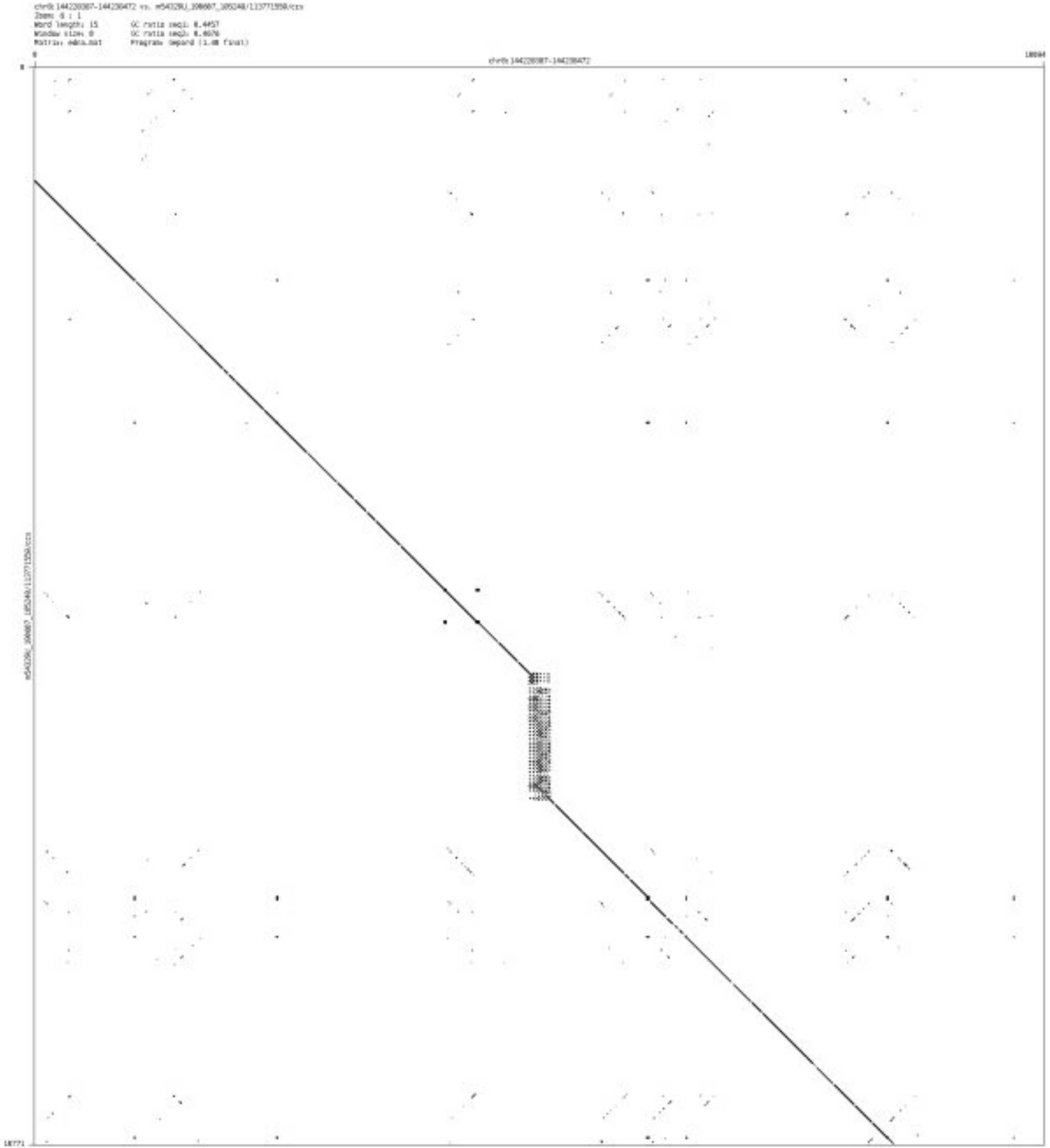

- chr8_144687804_144687929

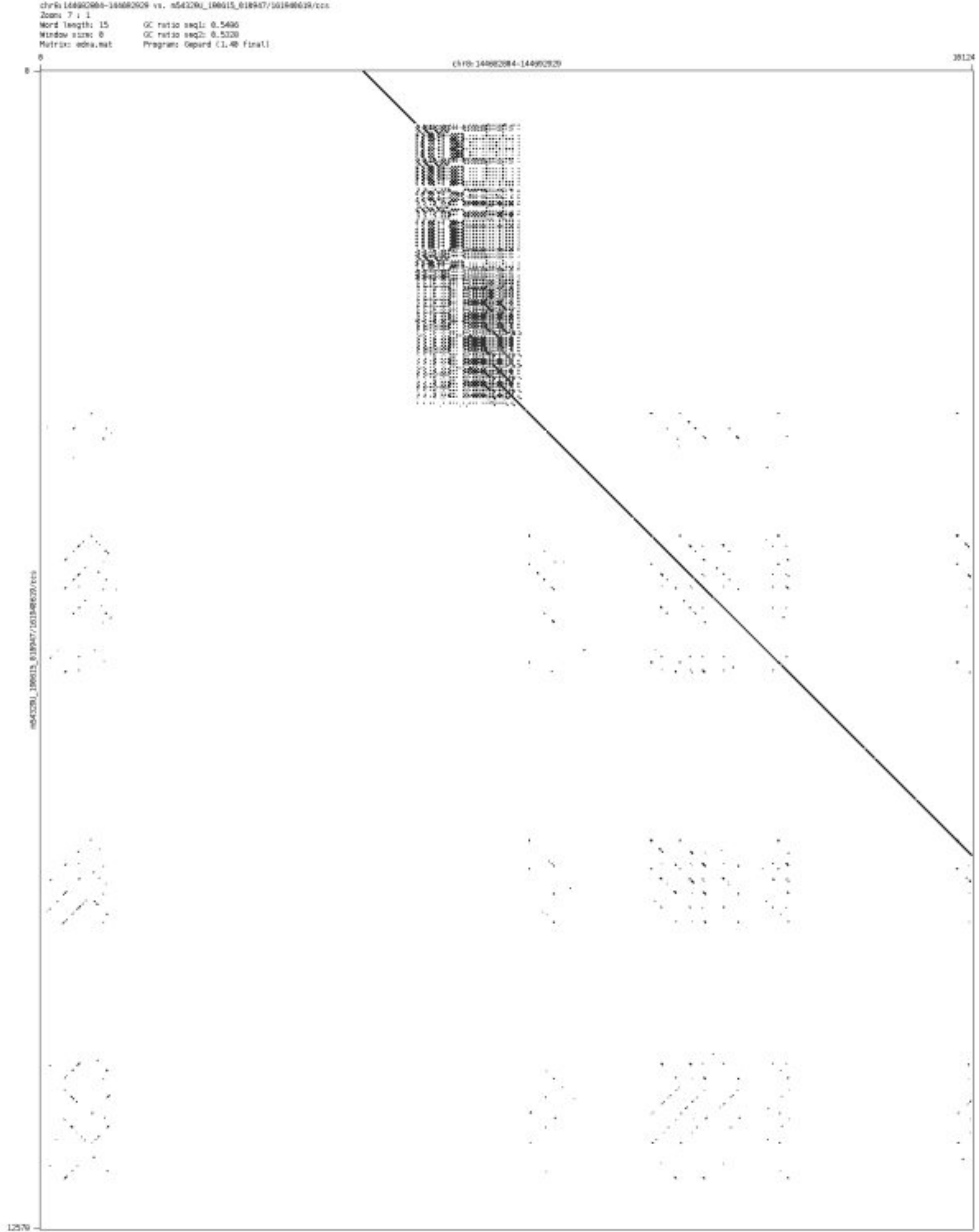

- chr9_808747_808859

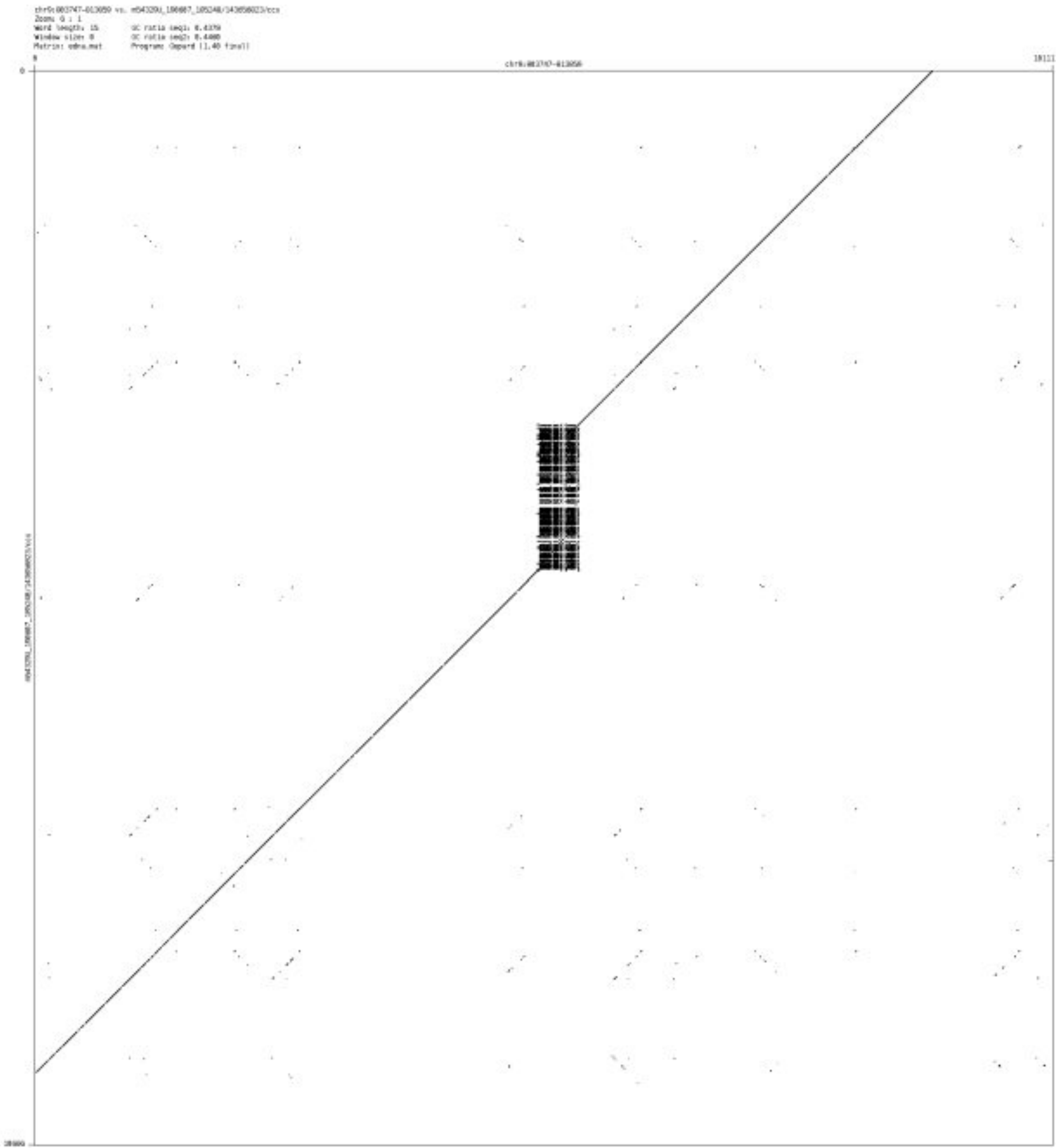

➢ chr9_15126725_15126845

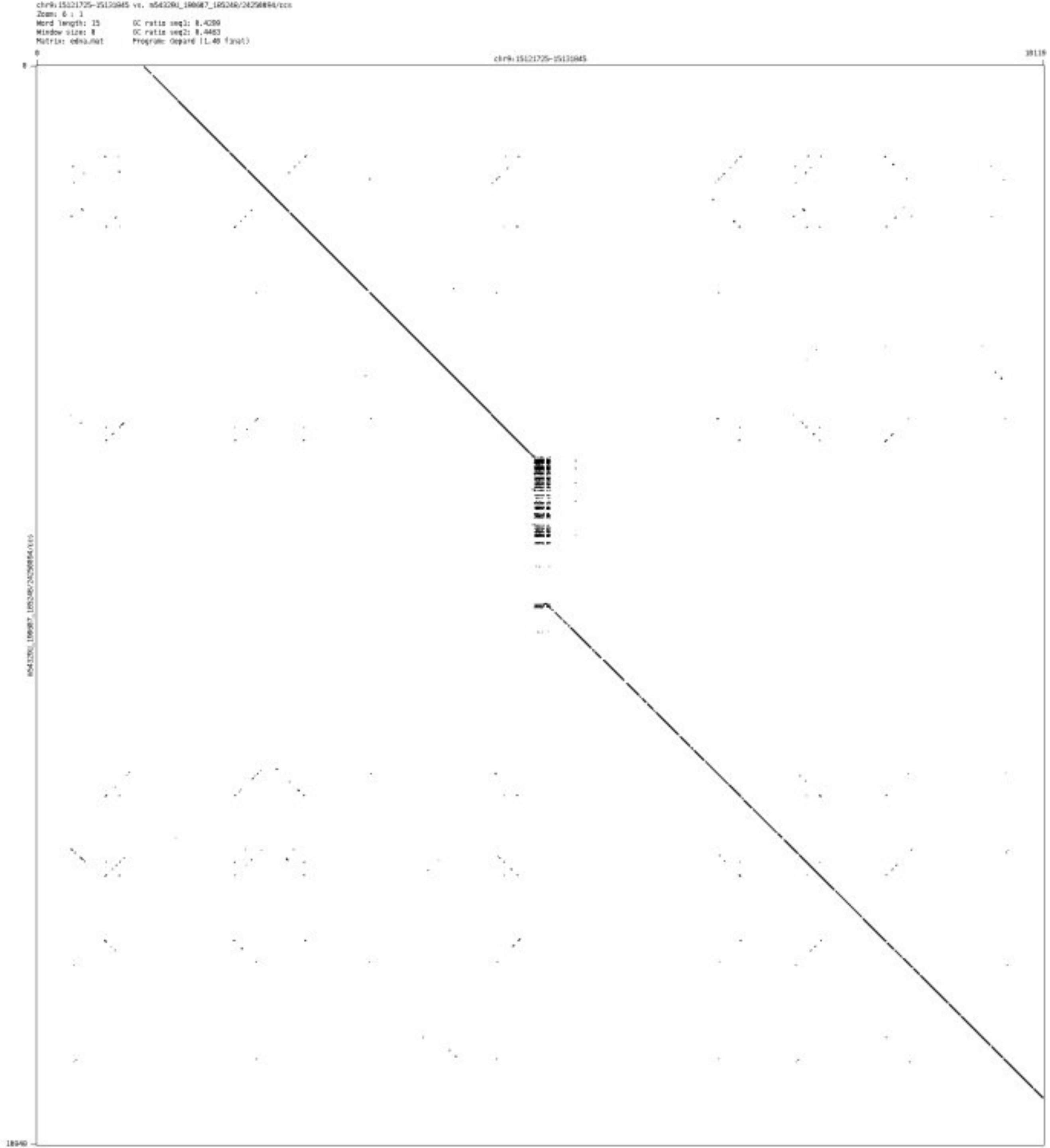

- chr9_35913547_35914174

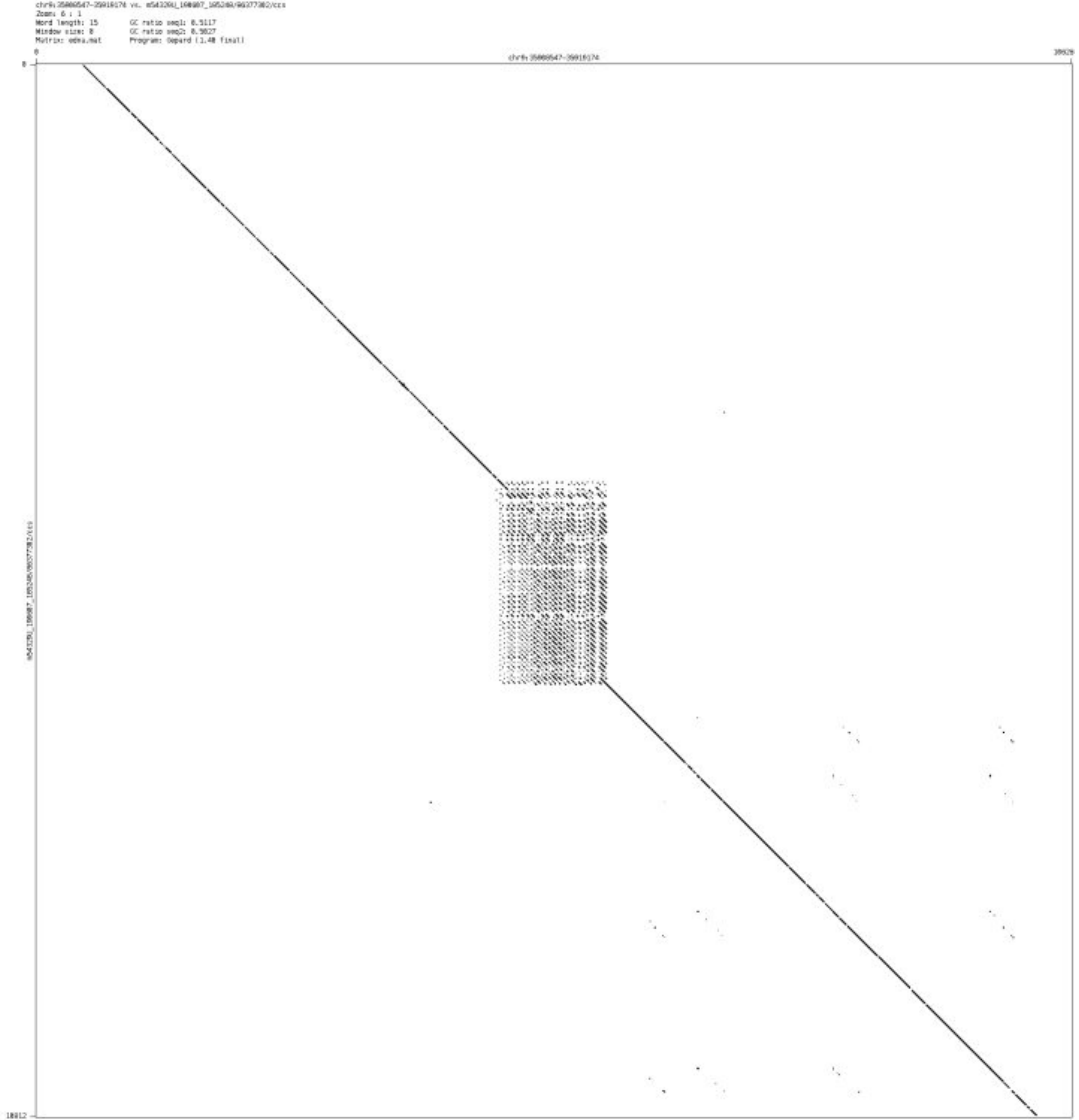

- chr9_89671543_89671681

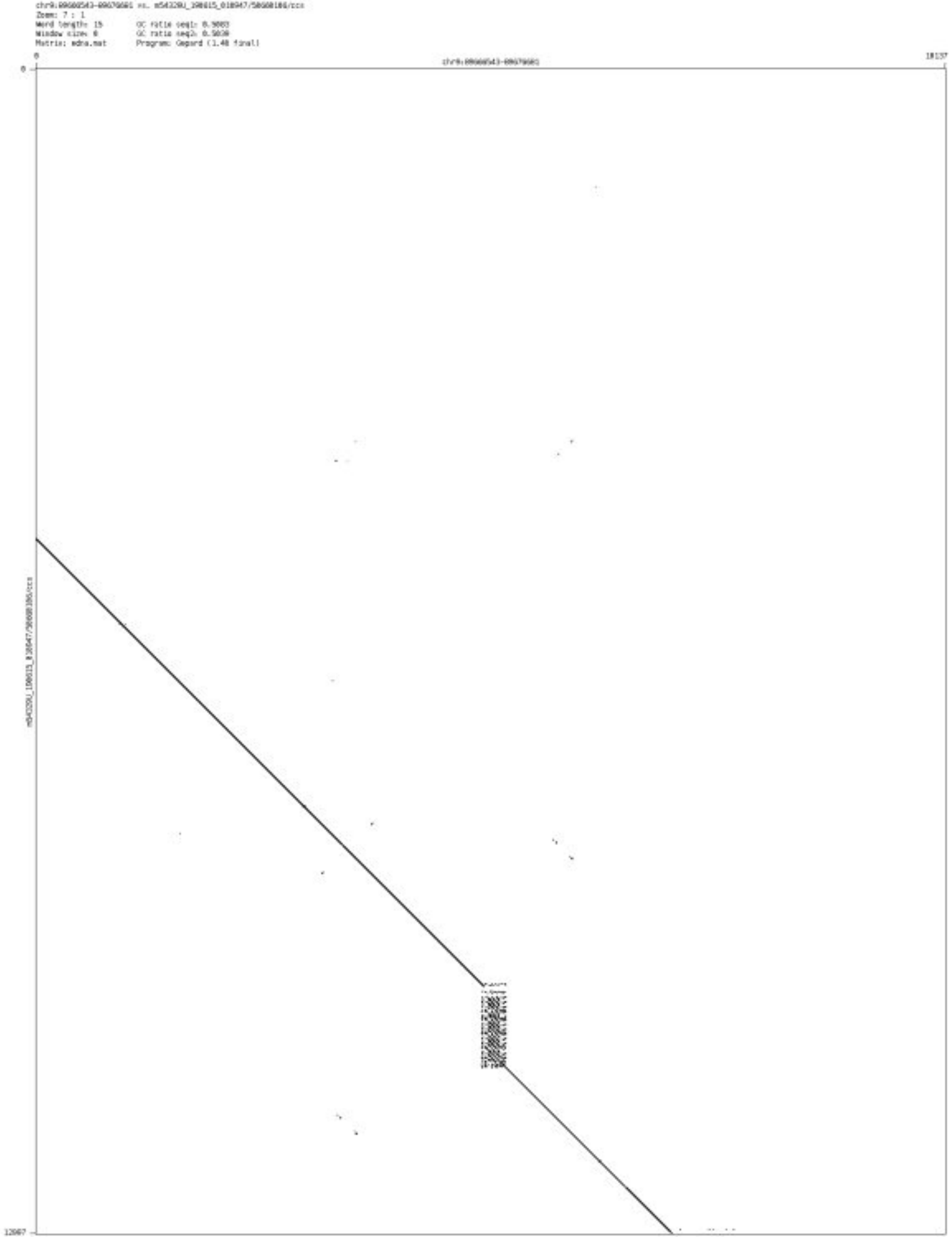

➢ chr9_137328196_137328432

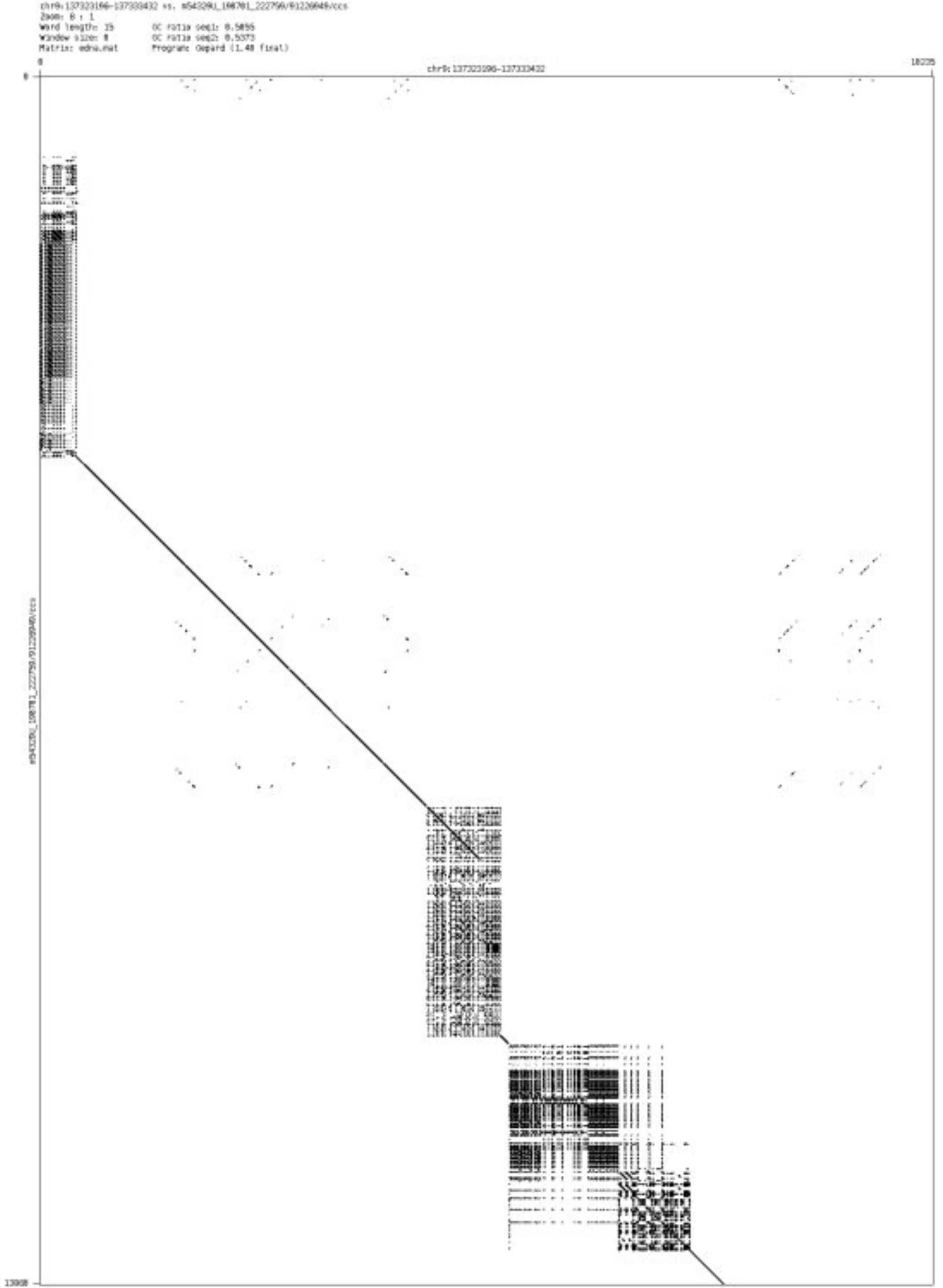

➢ chr9_137355493_137355619

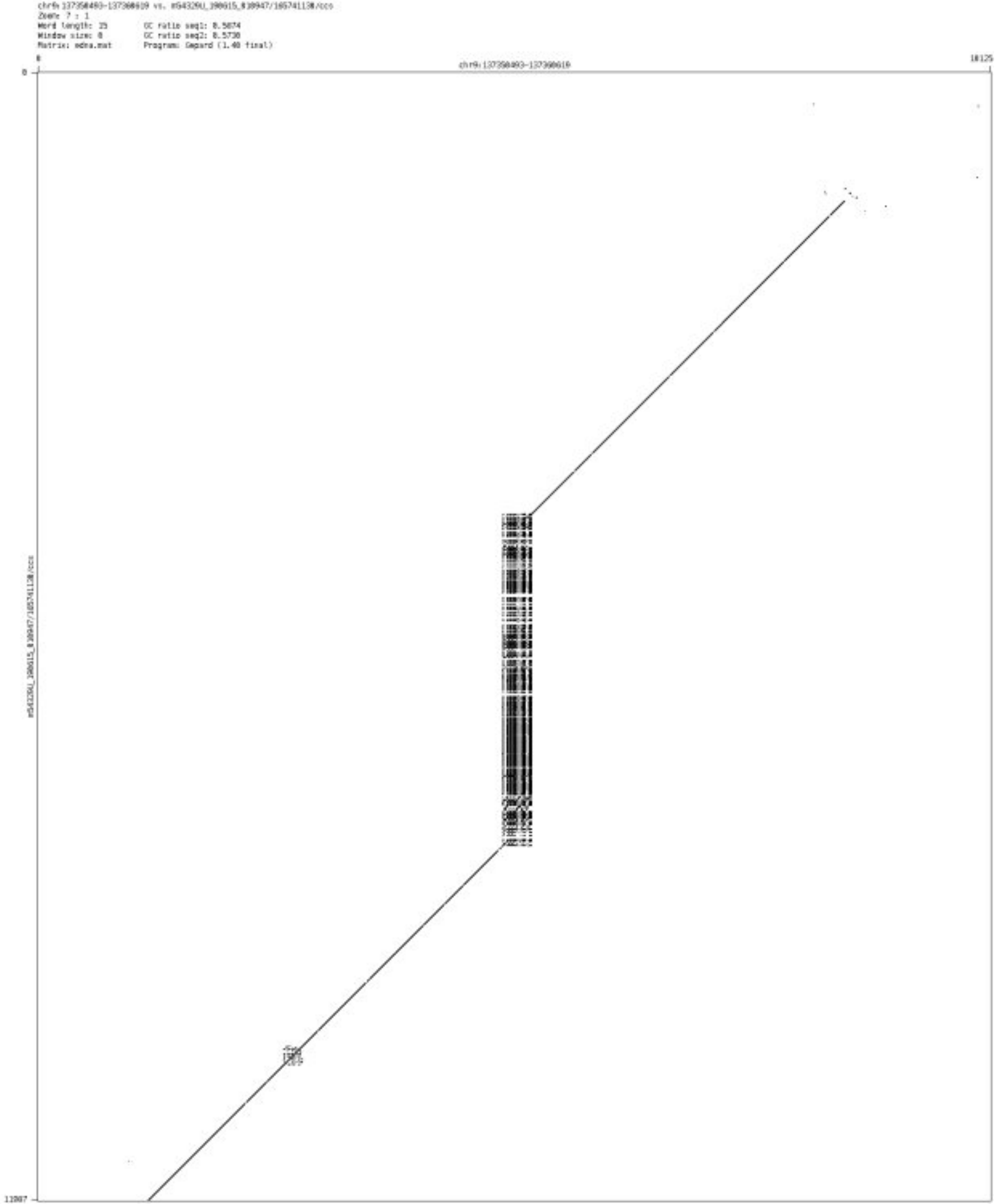

- chr9_137503837_137503947

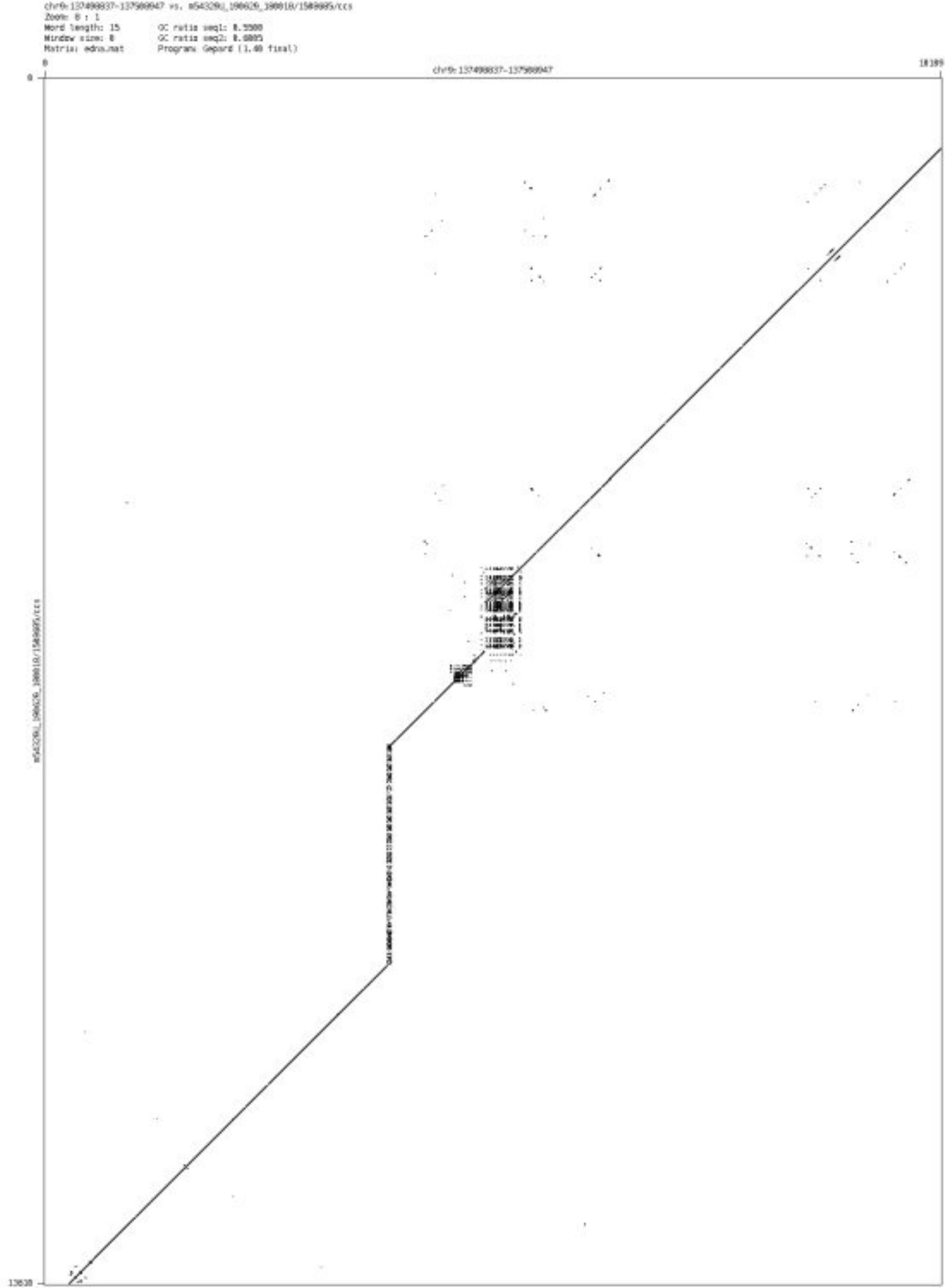

➢ chr10_5381284_5381363

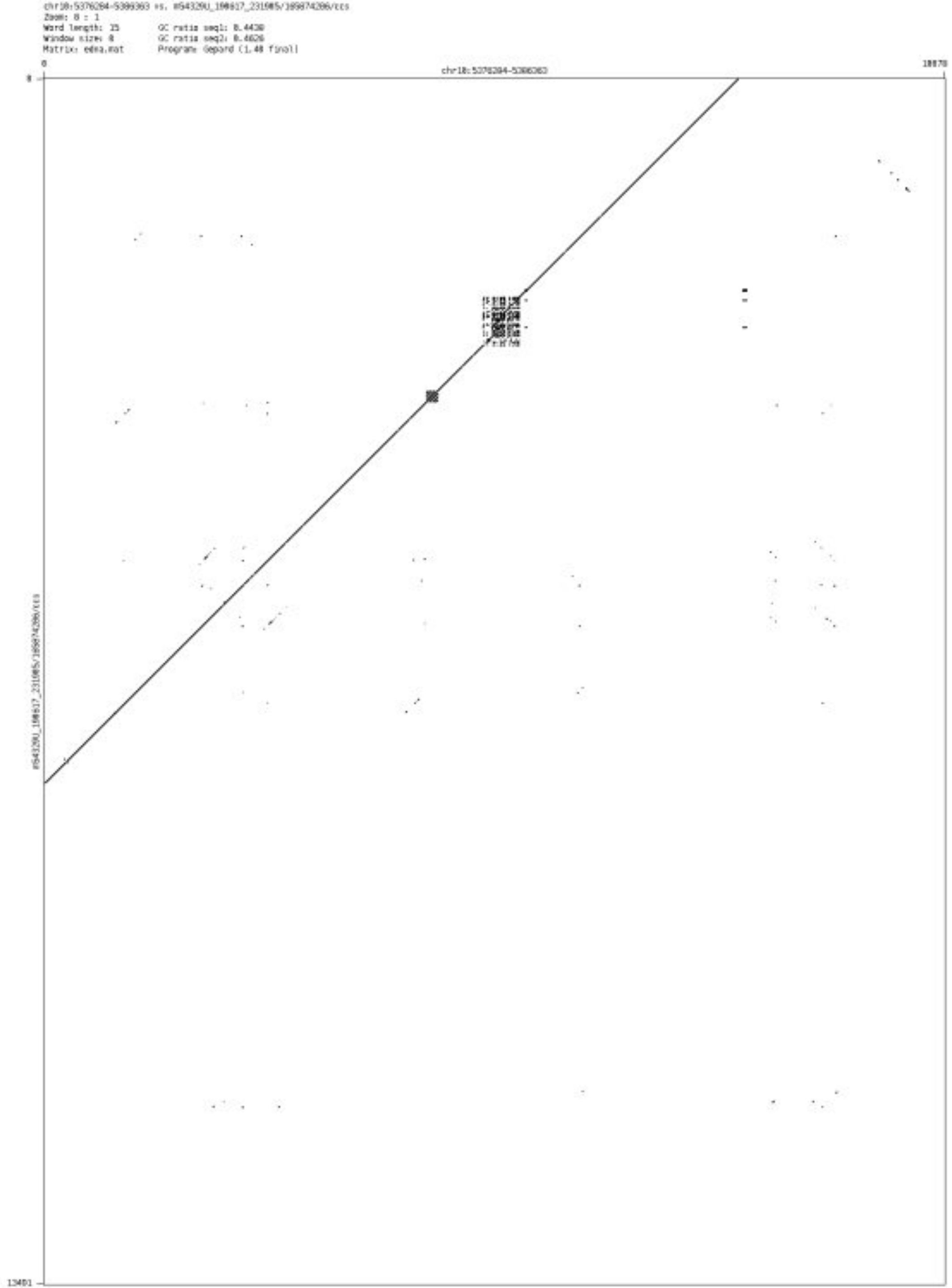

➢ chr10_60469258_60469364

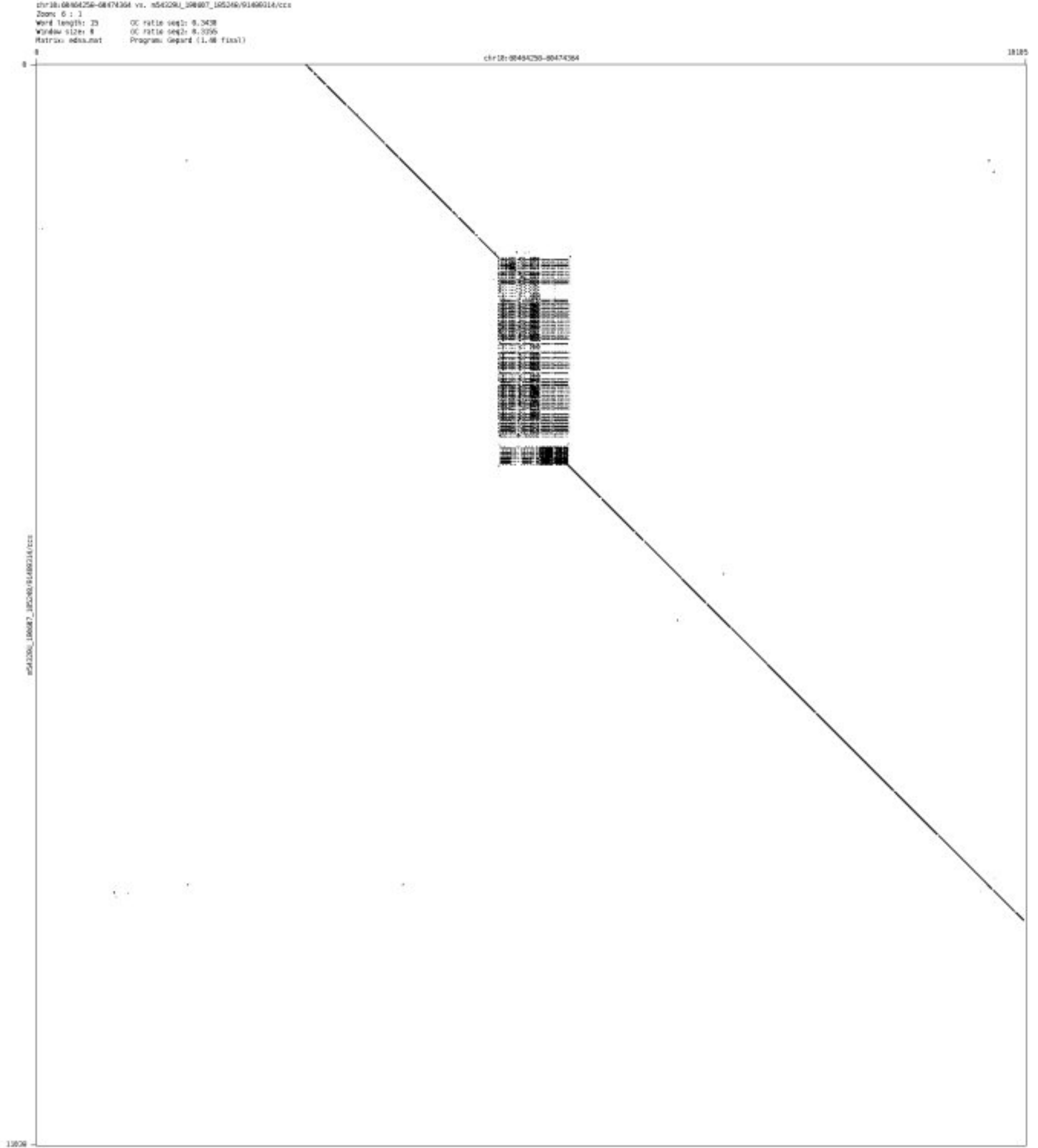

- chr10_127797498_127797602

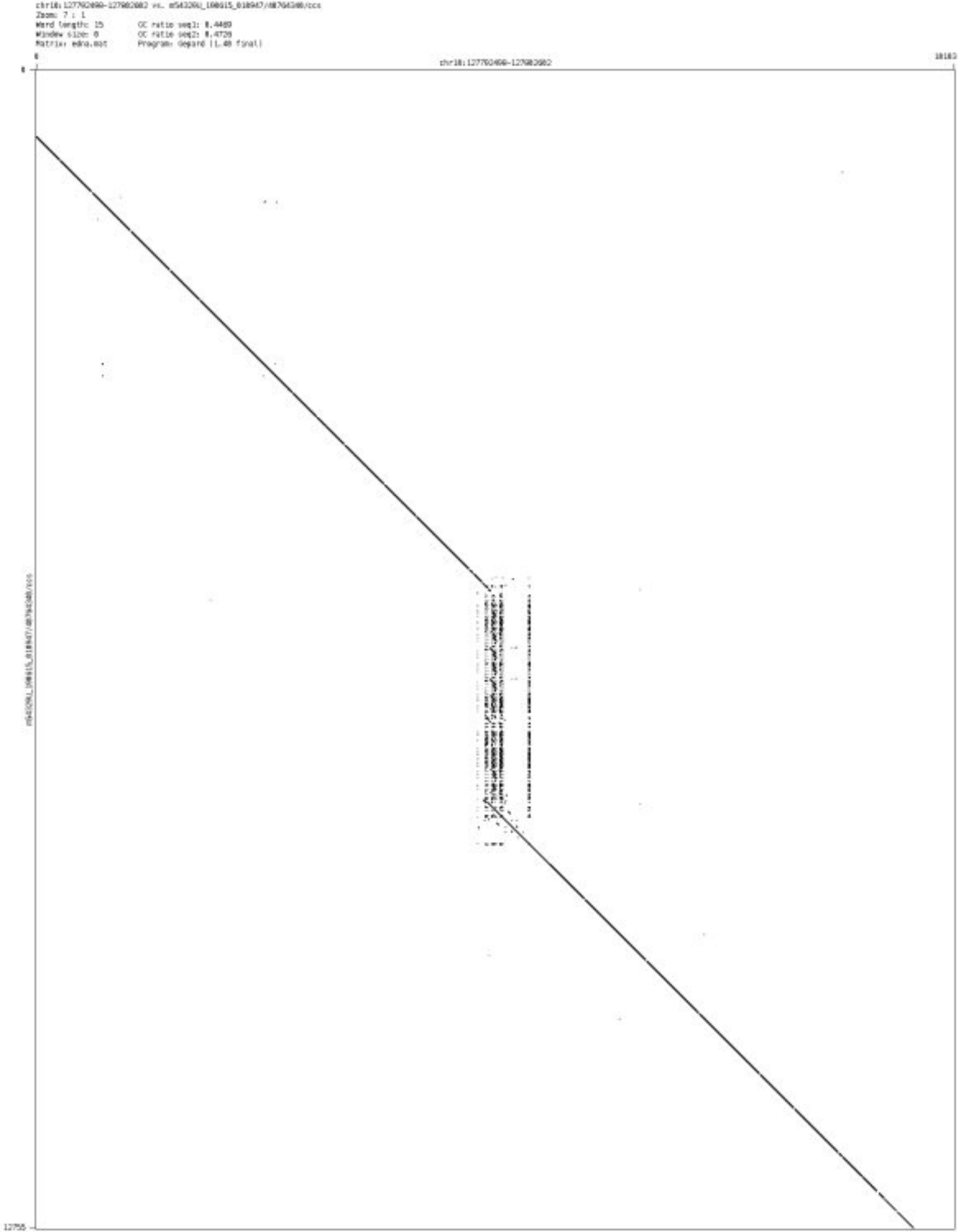

- ➢ chr10_128464484_128464594

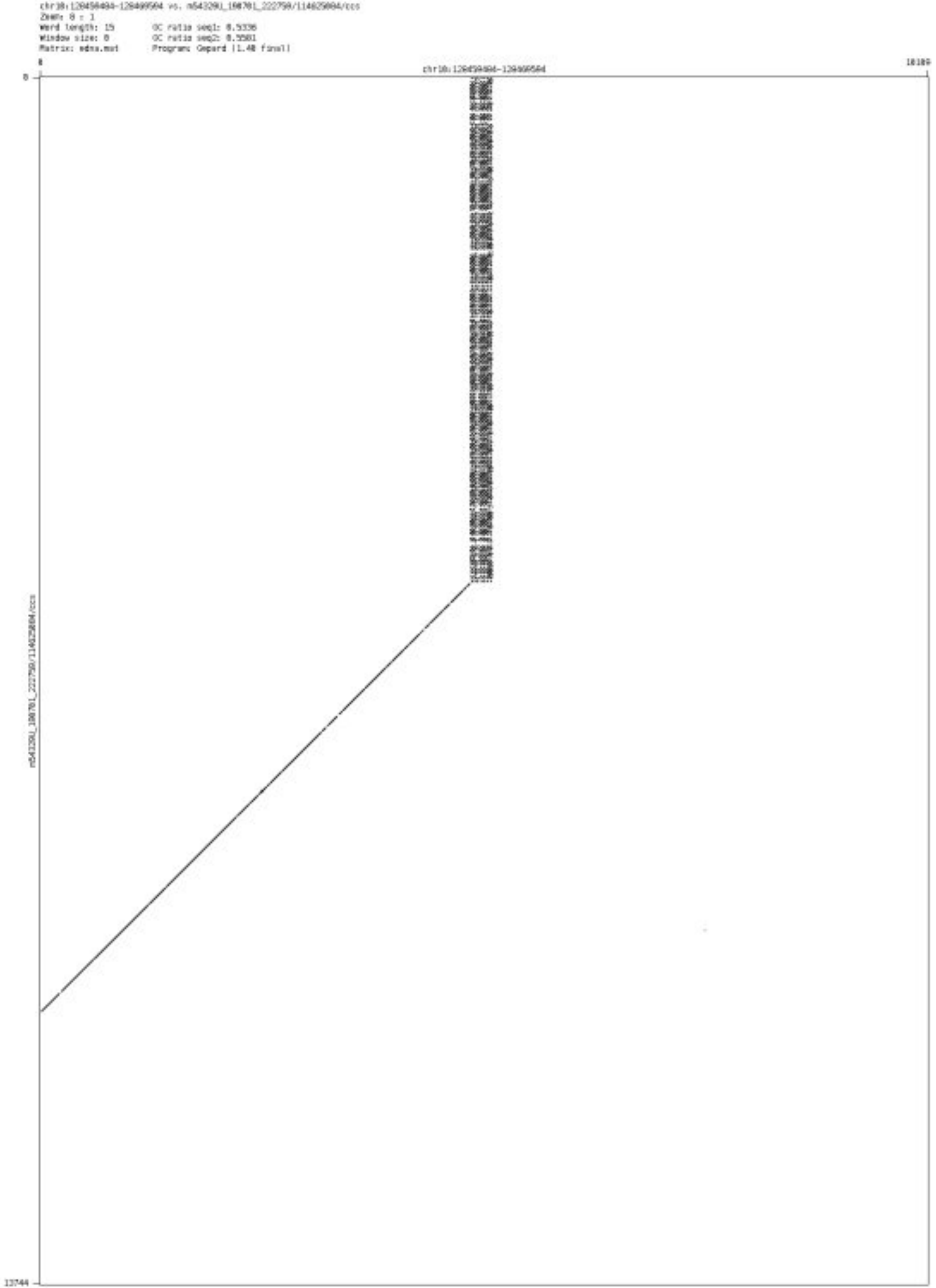

- chr10_133290814_133292455

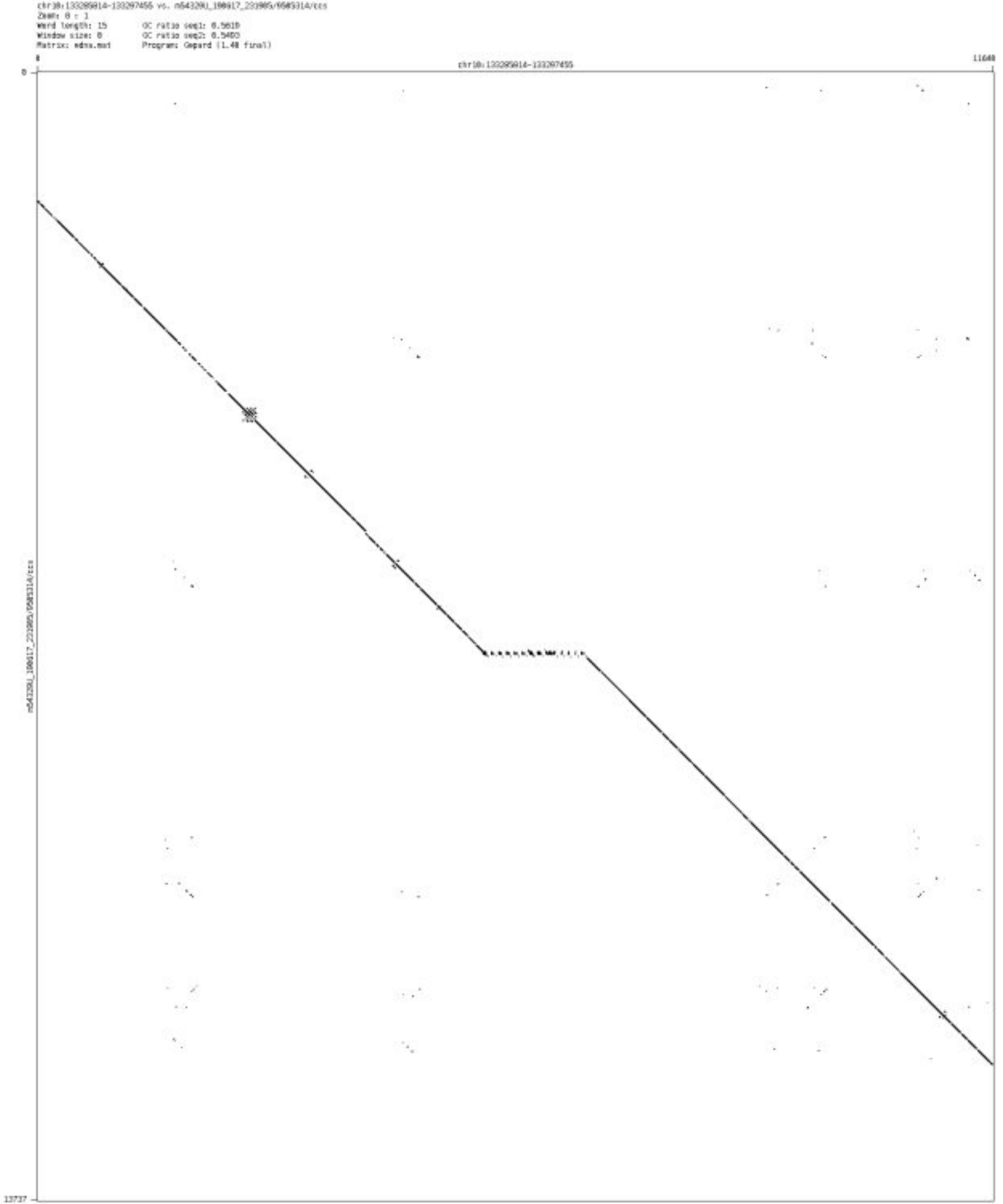

- chr11_3096979_3097064

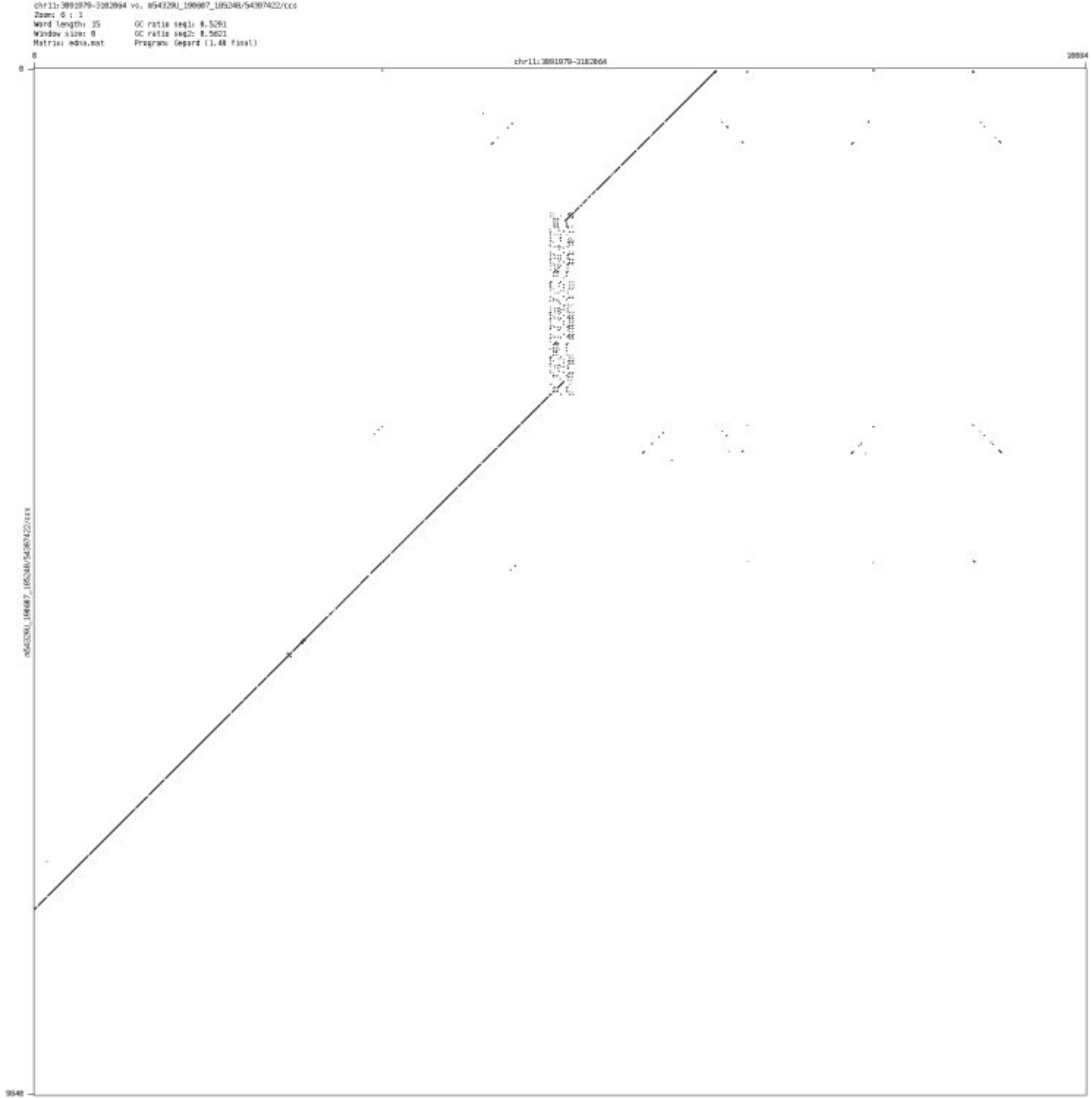

- chr11_24918873_24919036

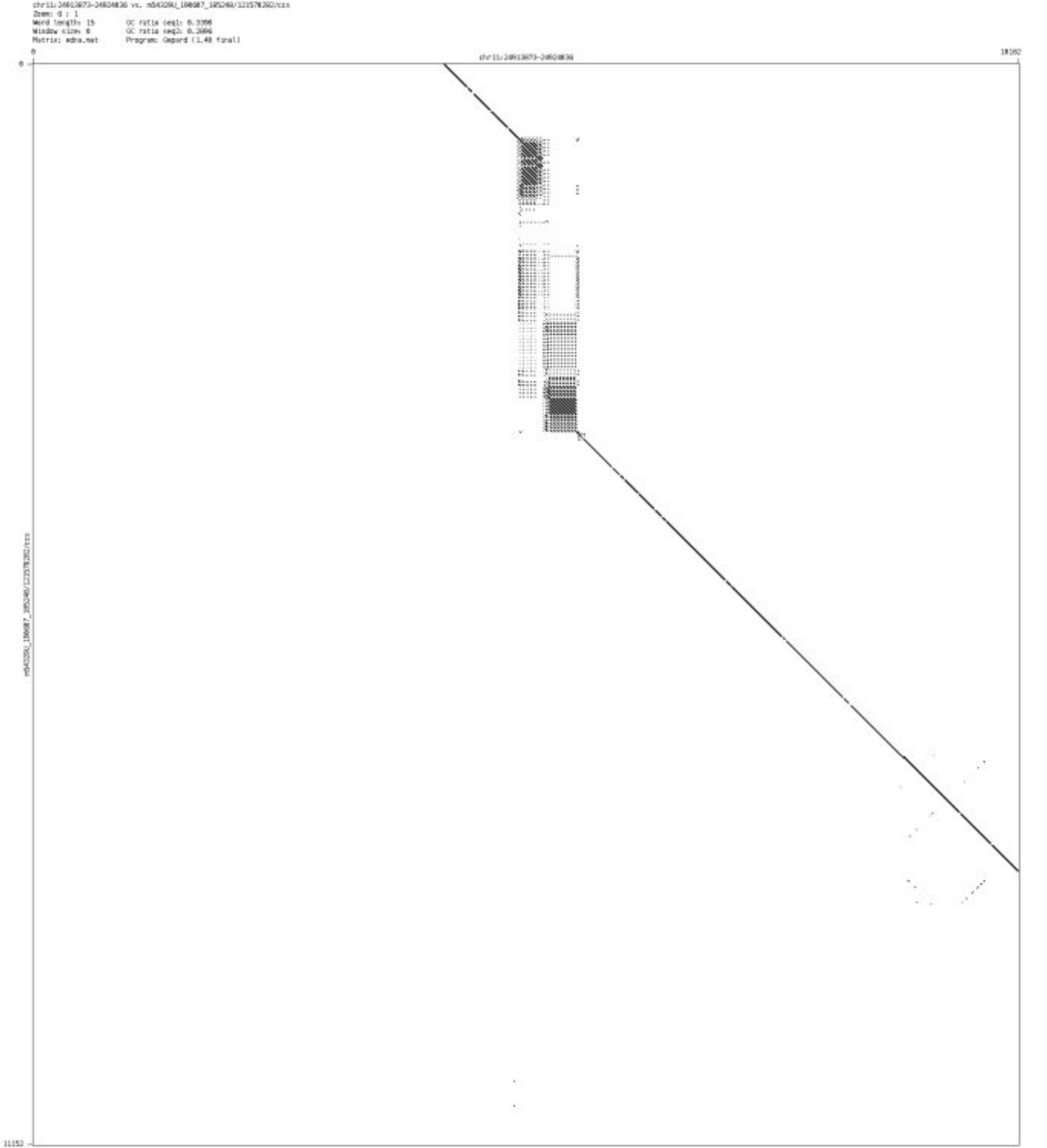

➢ chr11_36309611_36309970

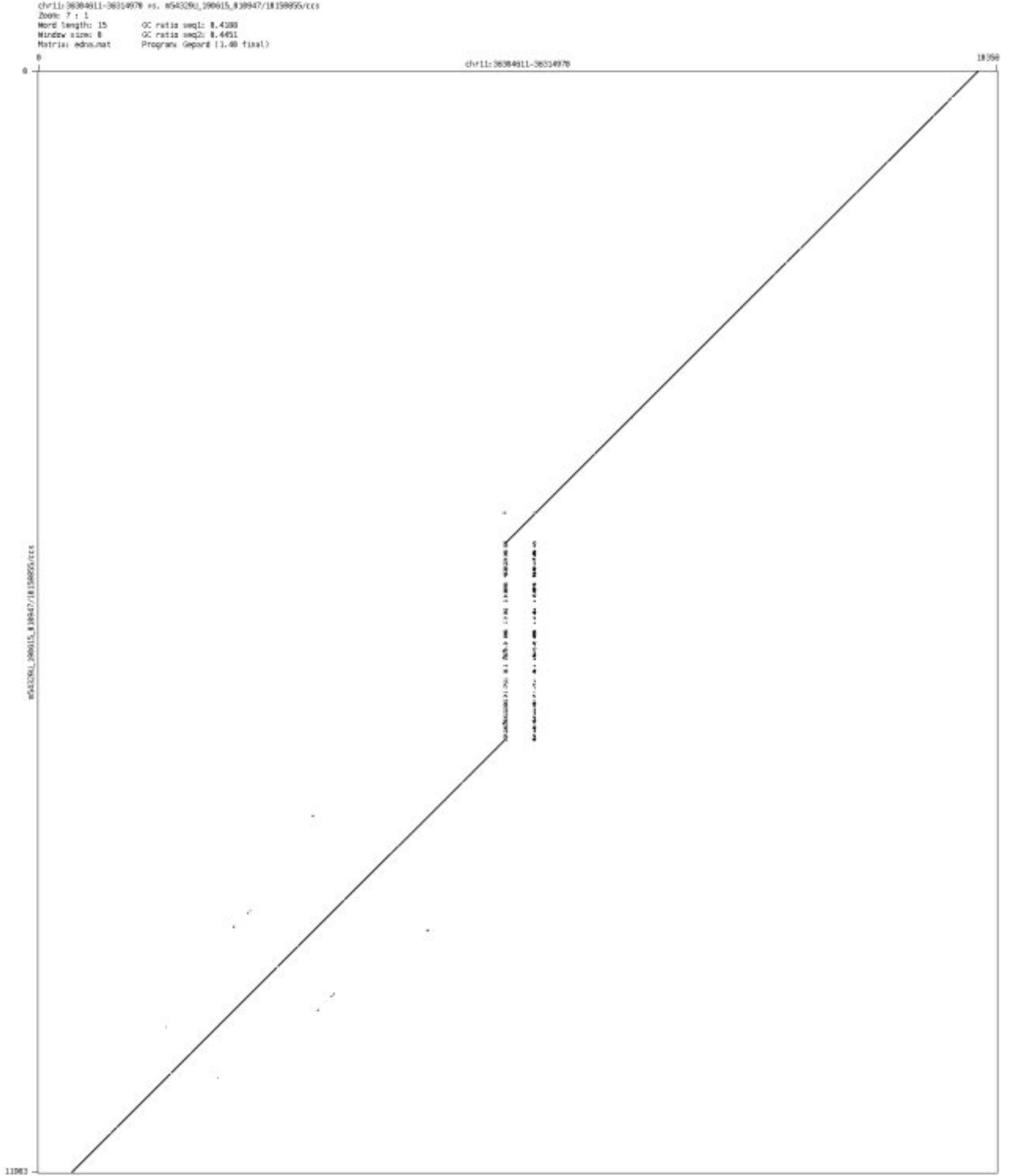

- chr11_92705384_92705891

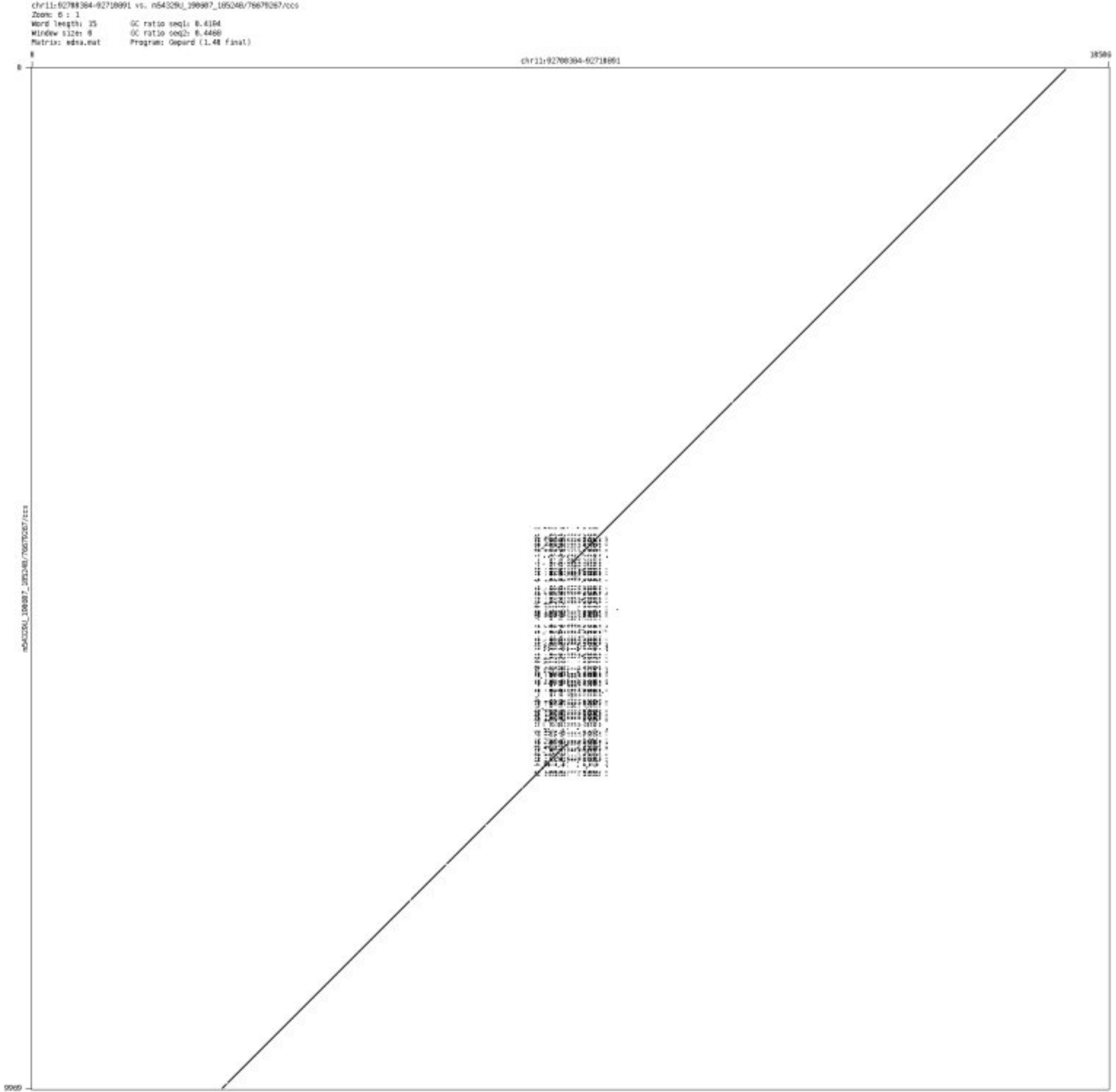

➢ chr11_122016499_122017716

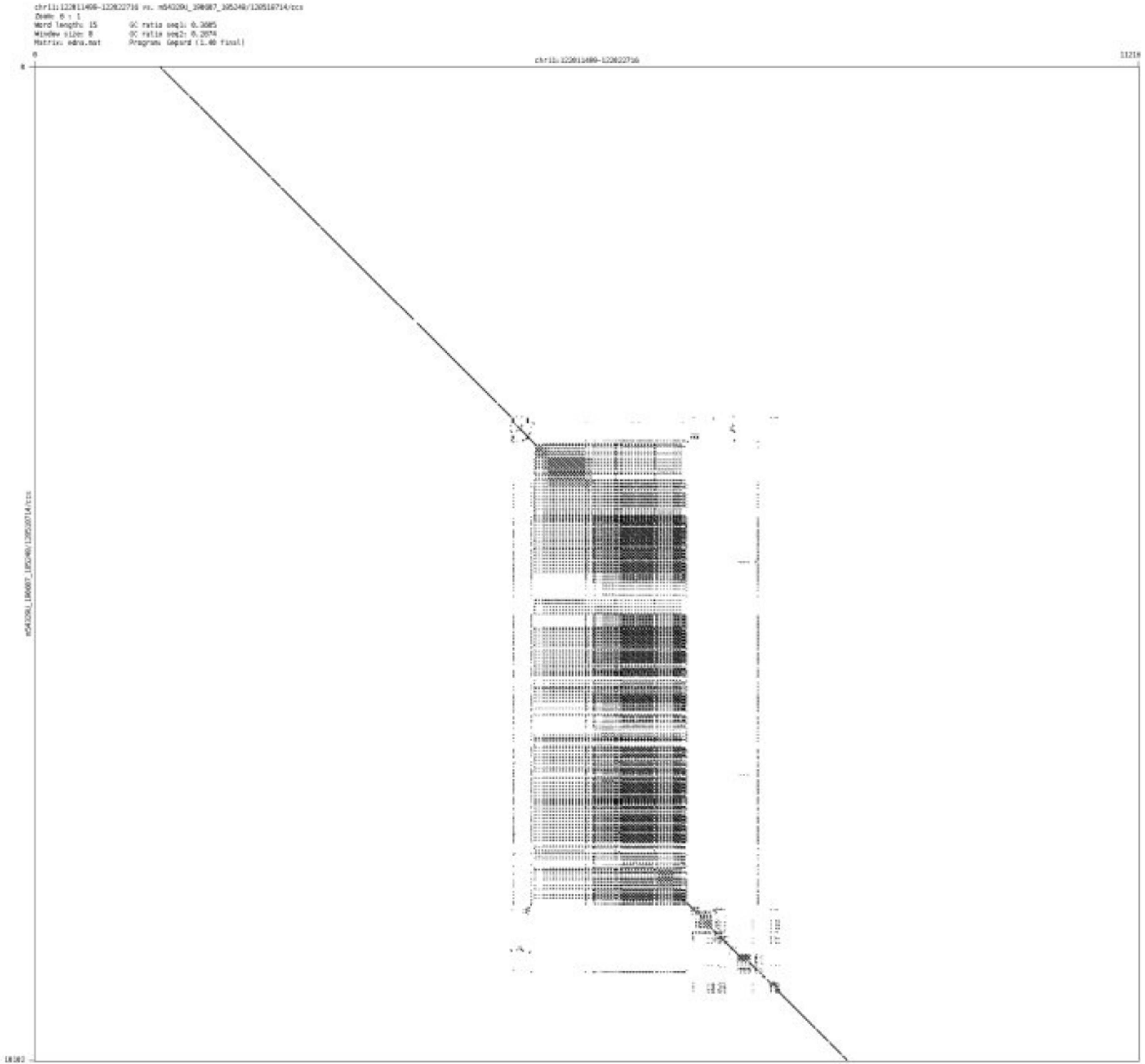

- chr11_128930543_128930892

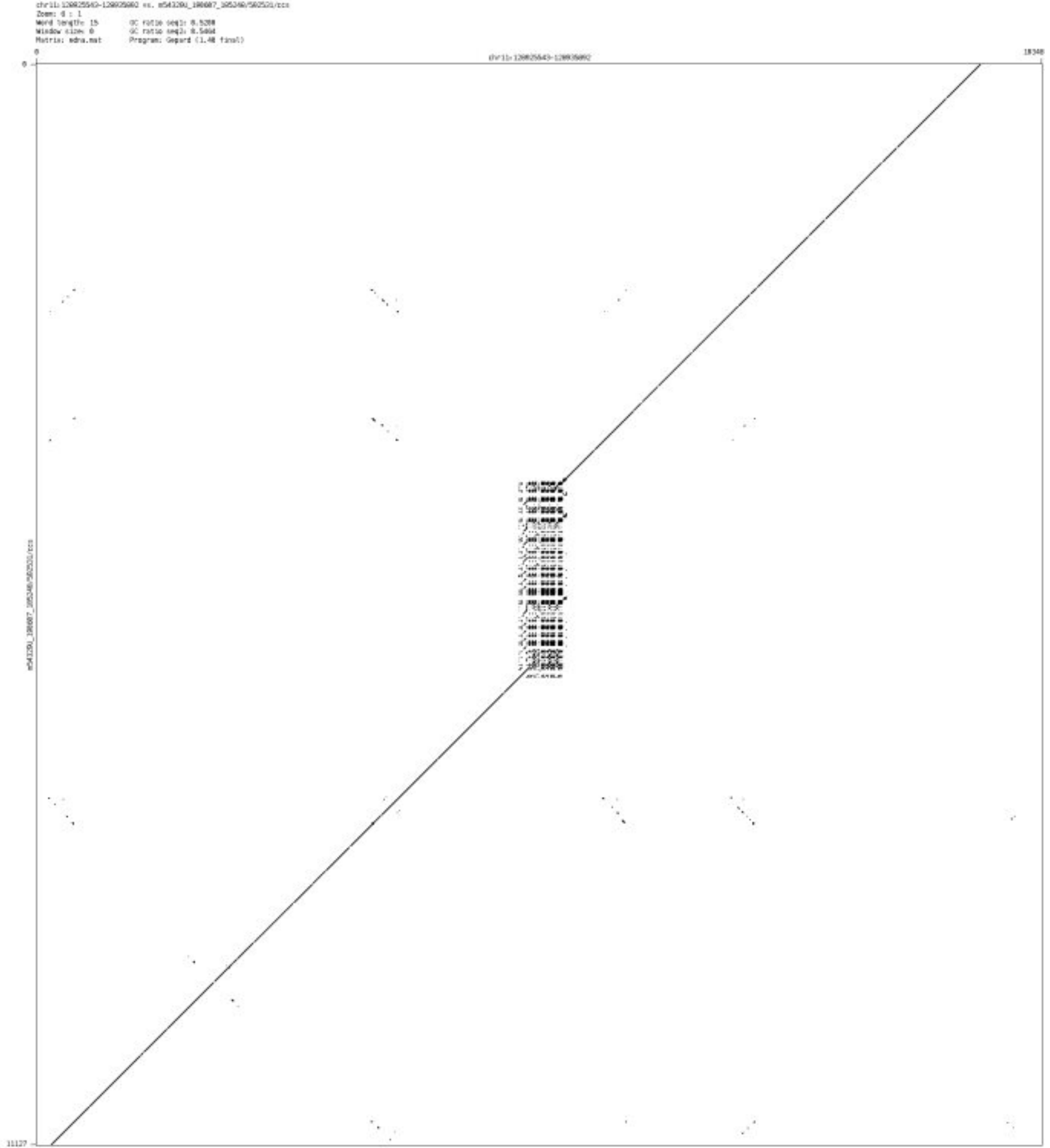

- chr12_1574361_1574528

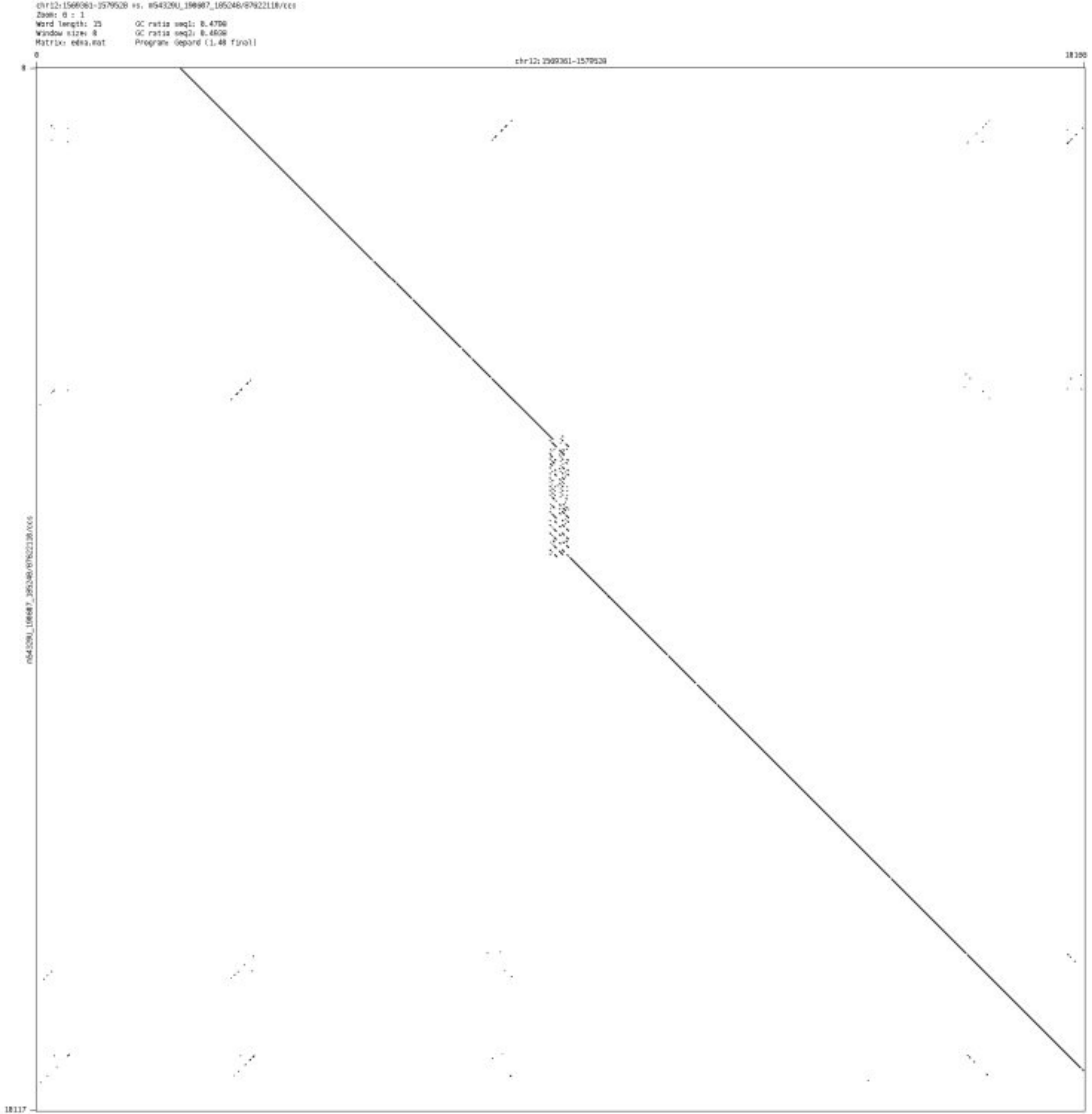

- ➢ chr12_124438819_124439422

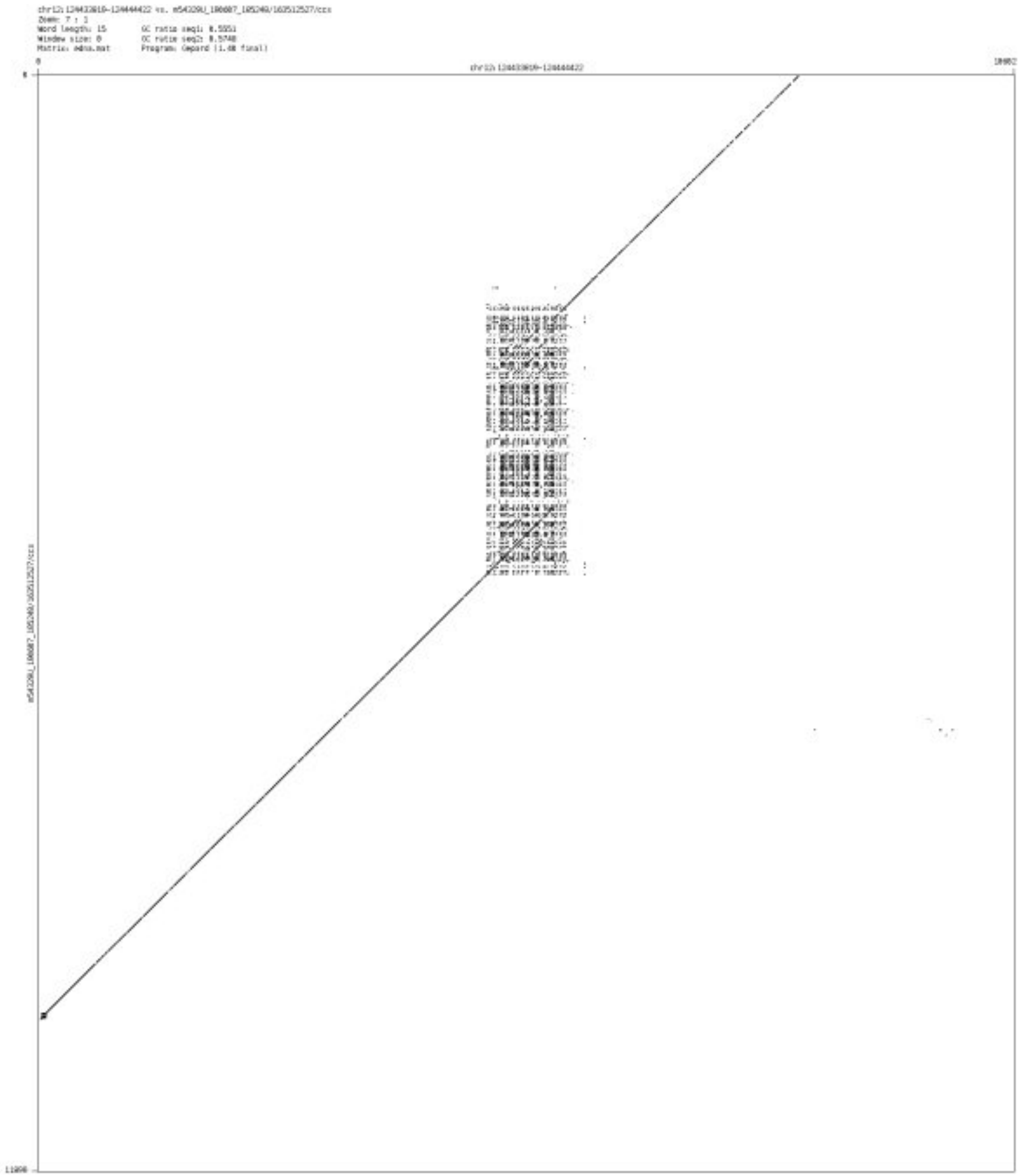

- chr12_131918765_131918869

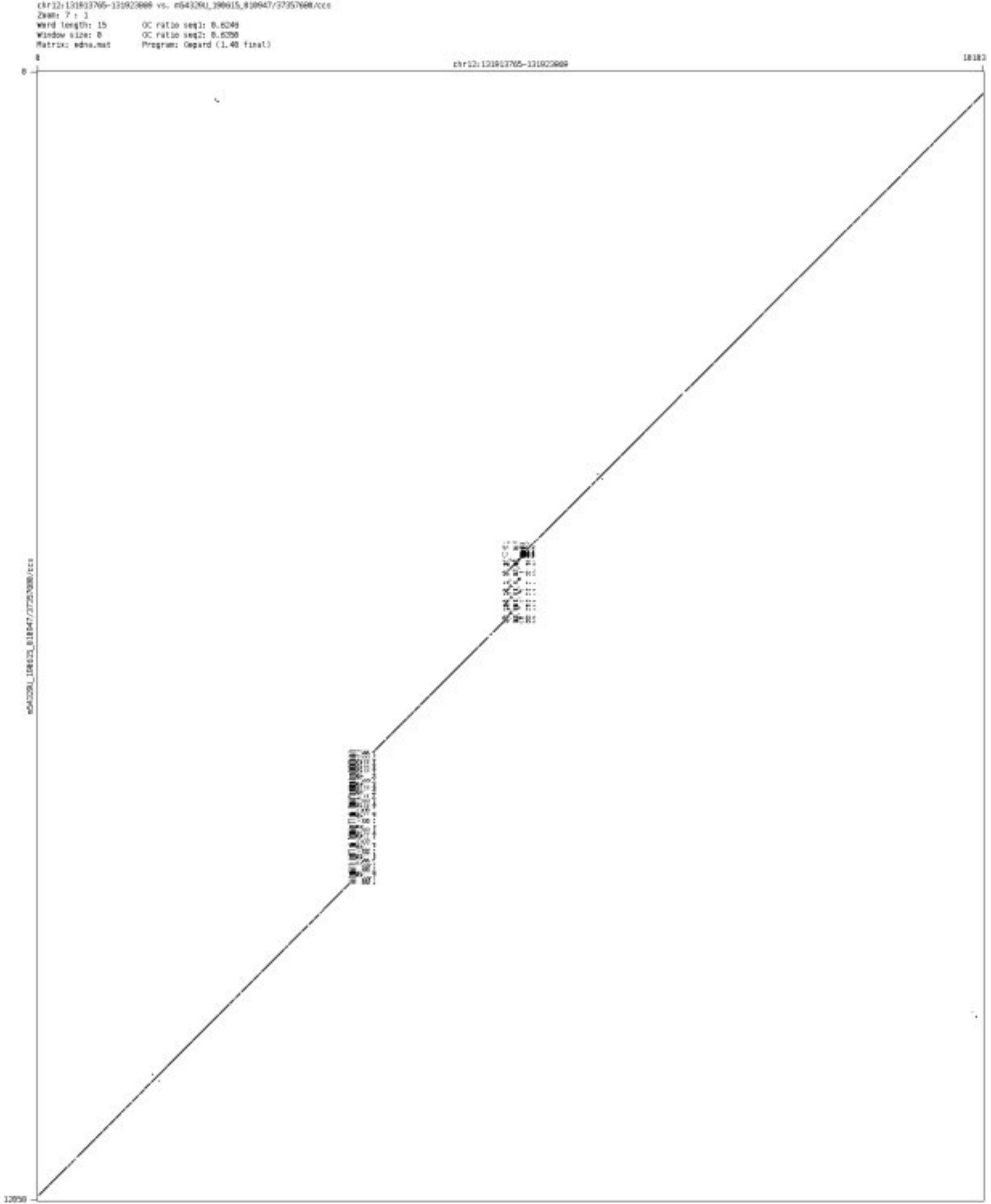

- chr13_38956678_38956810

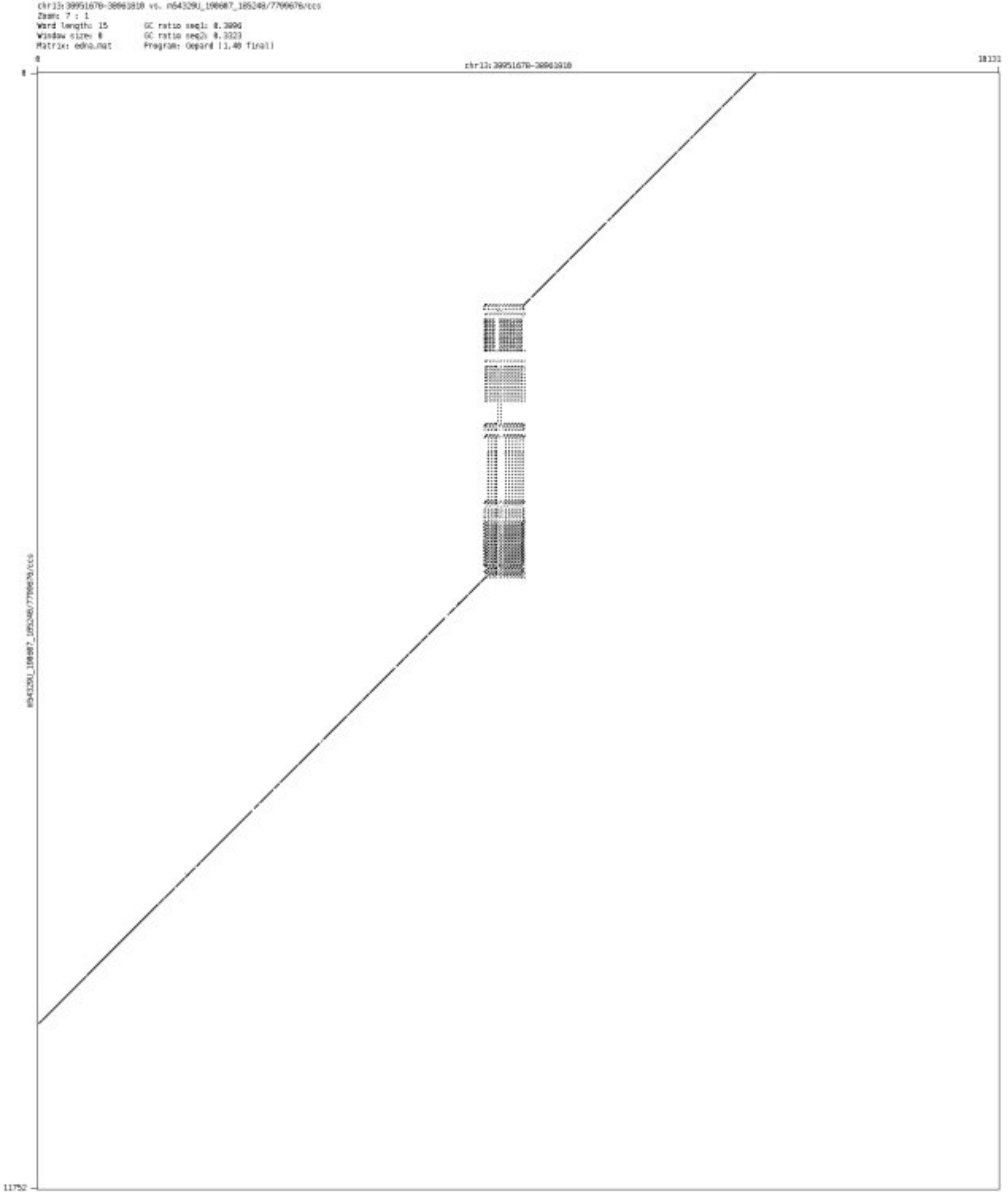

- ➢ chr13_60977009_60978048

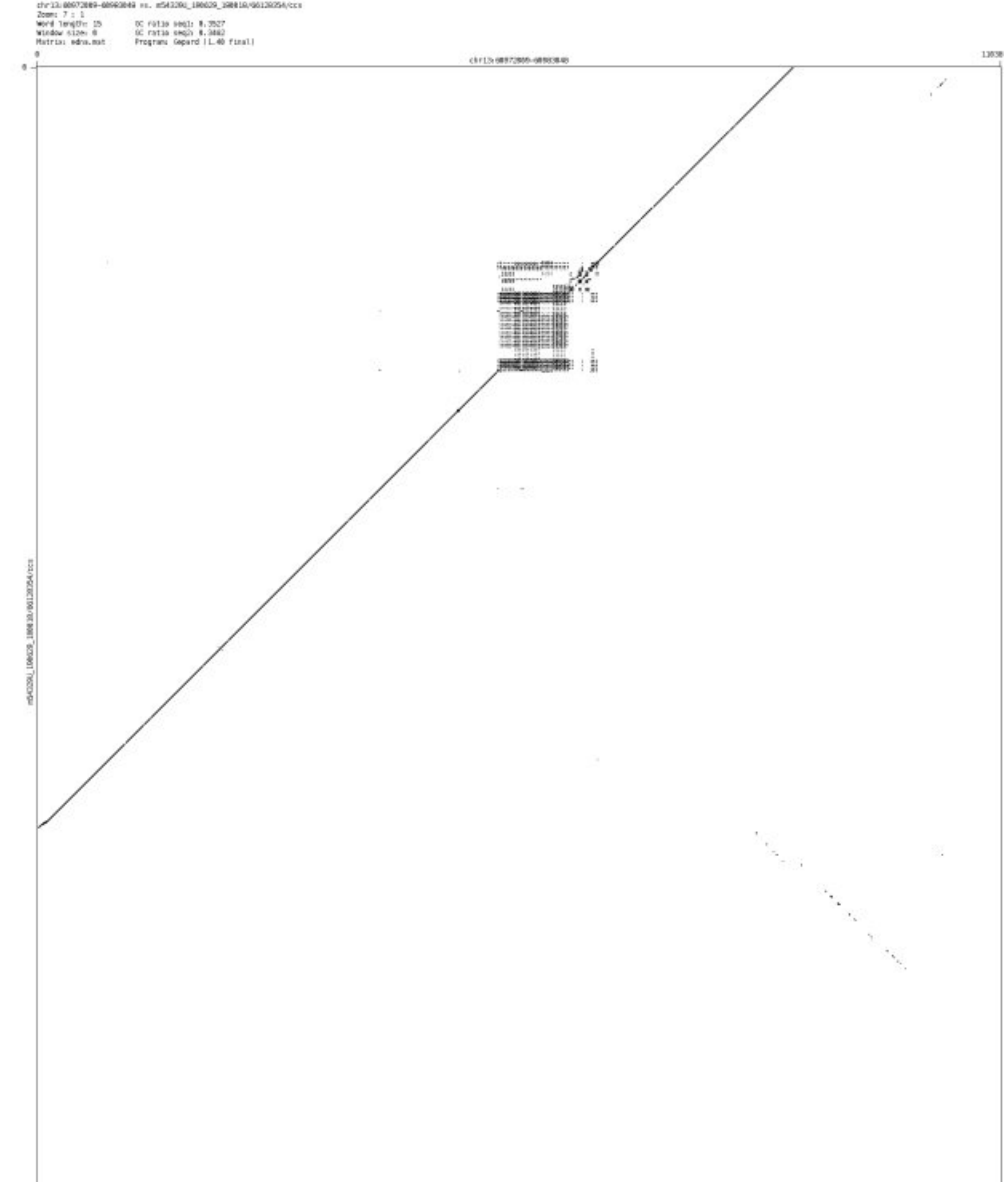

➢ chr13_71118063_71118136

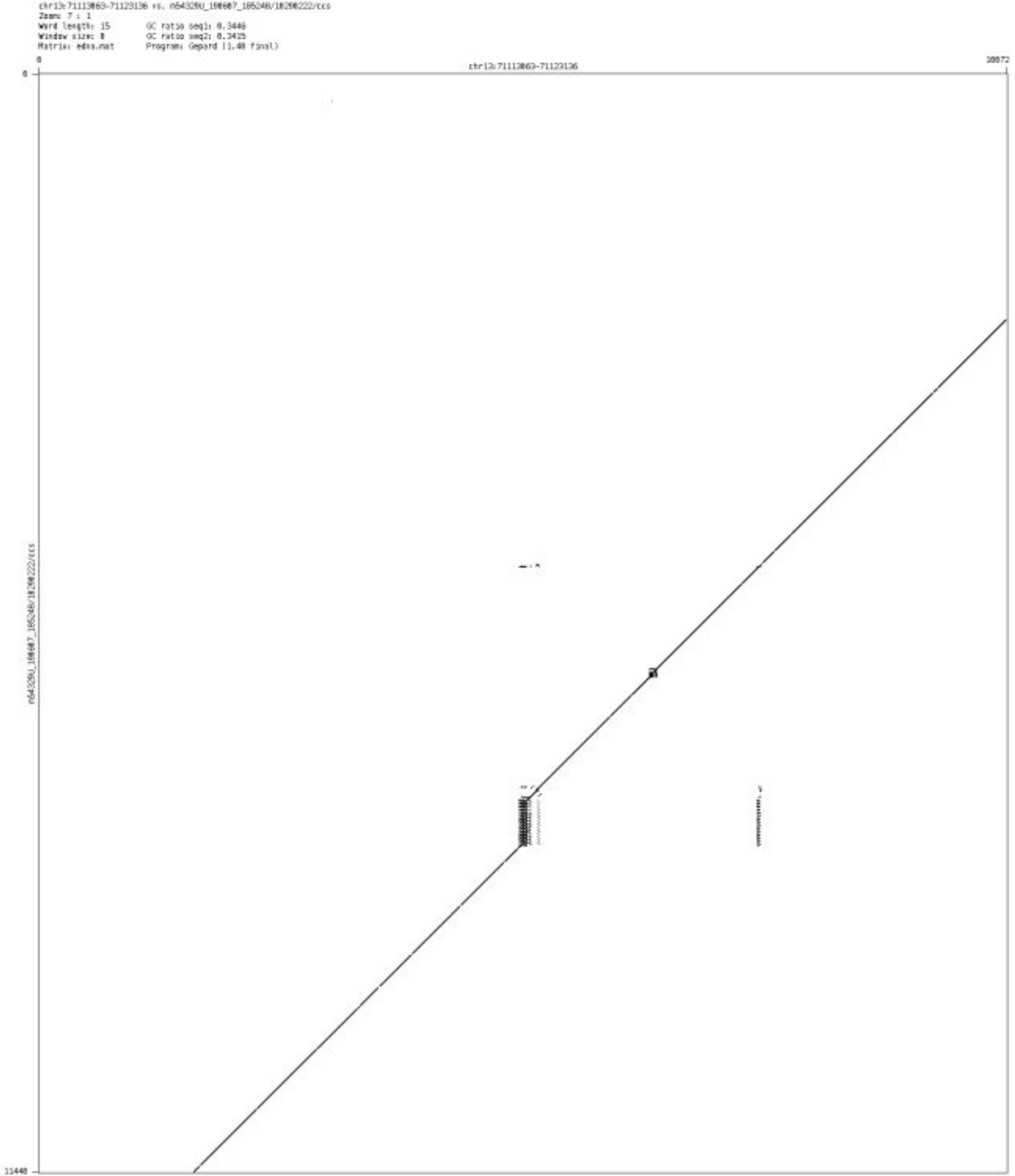

- ➢ chr13_114198570_114199832

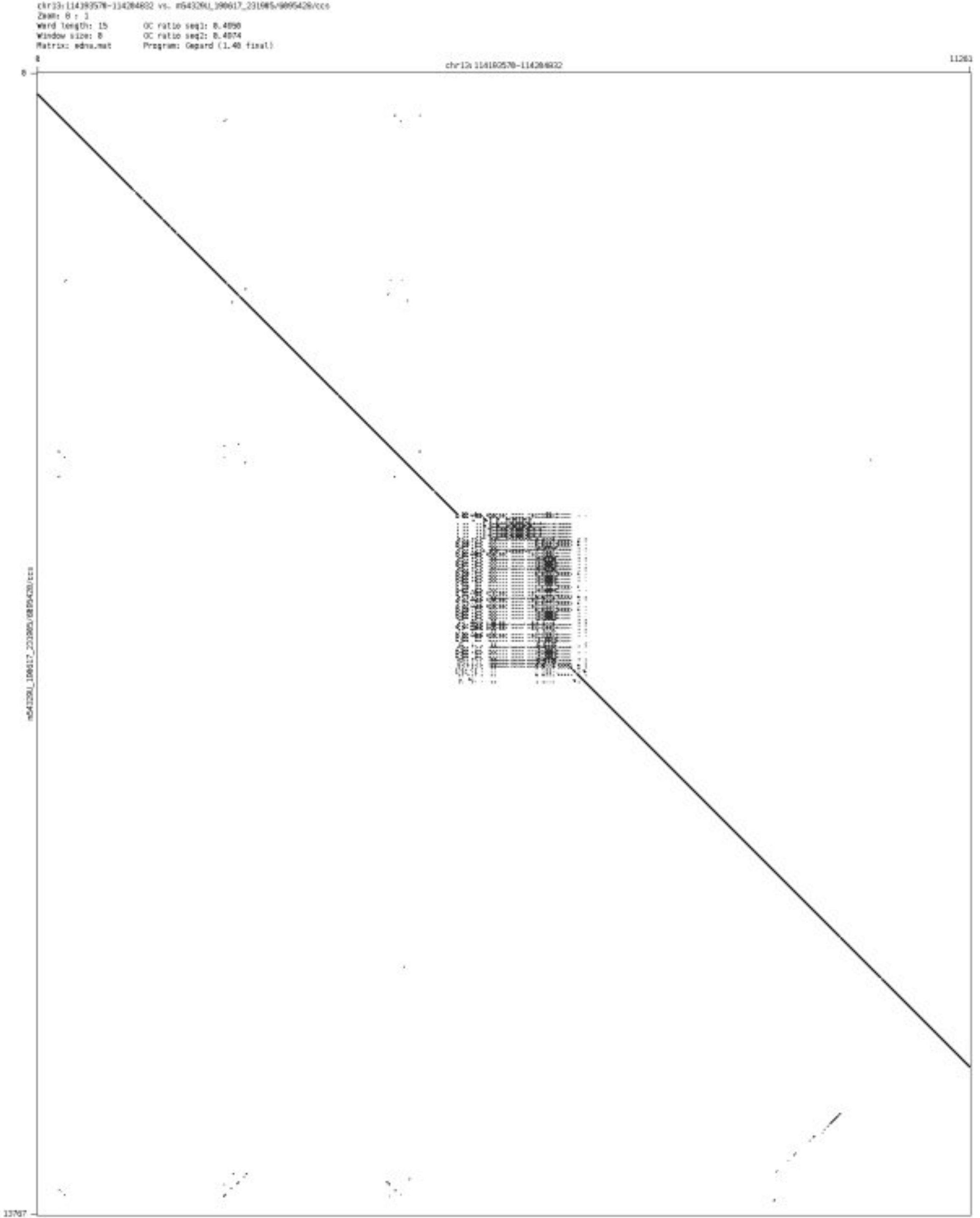

➢ chr14_35948113_35949836

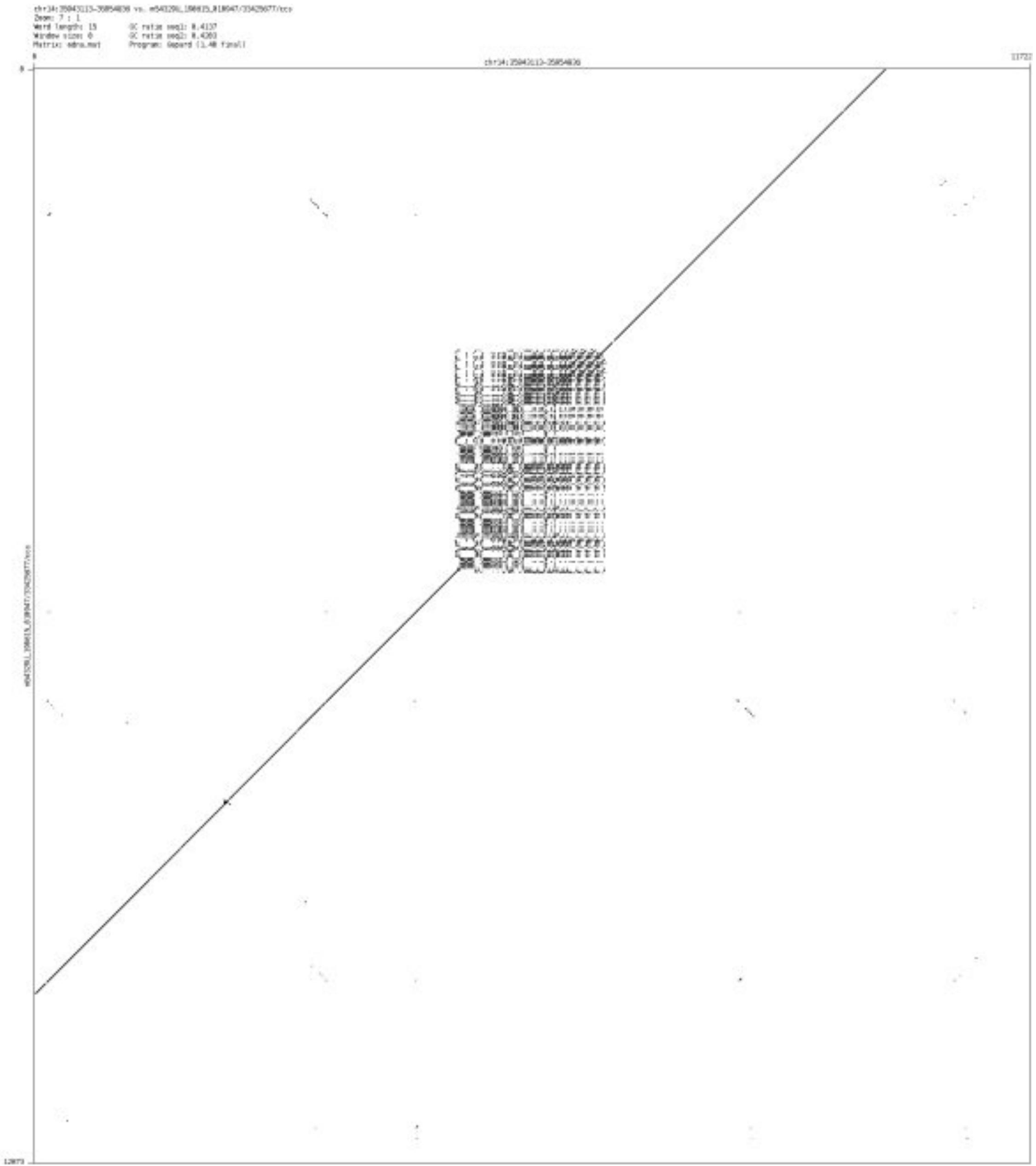

- ➢ chr14_65791578_65791692

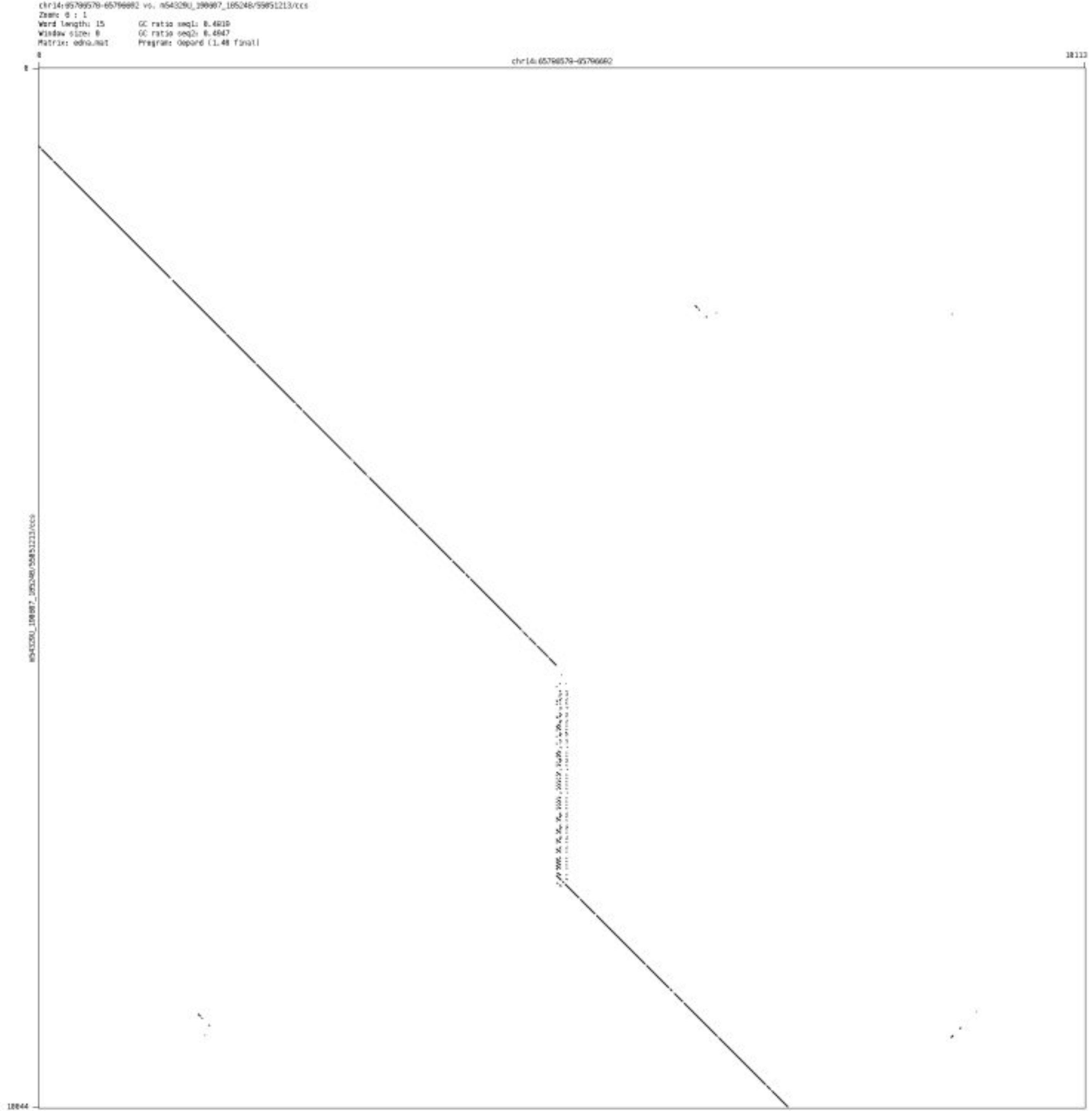

➢ chr14_91293420_91293512

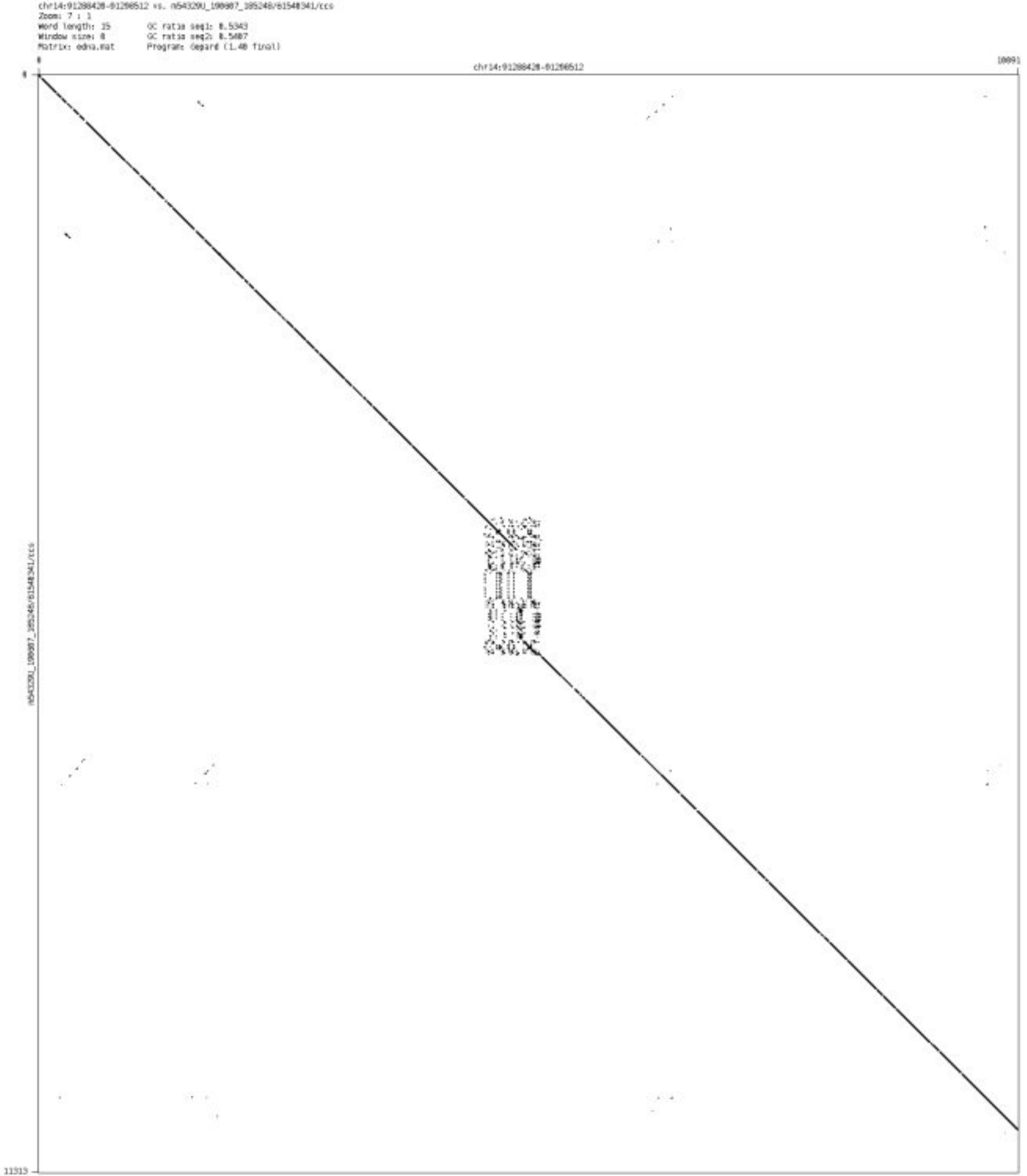

- chr14_99927890_99928038

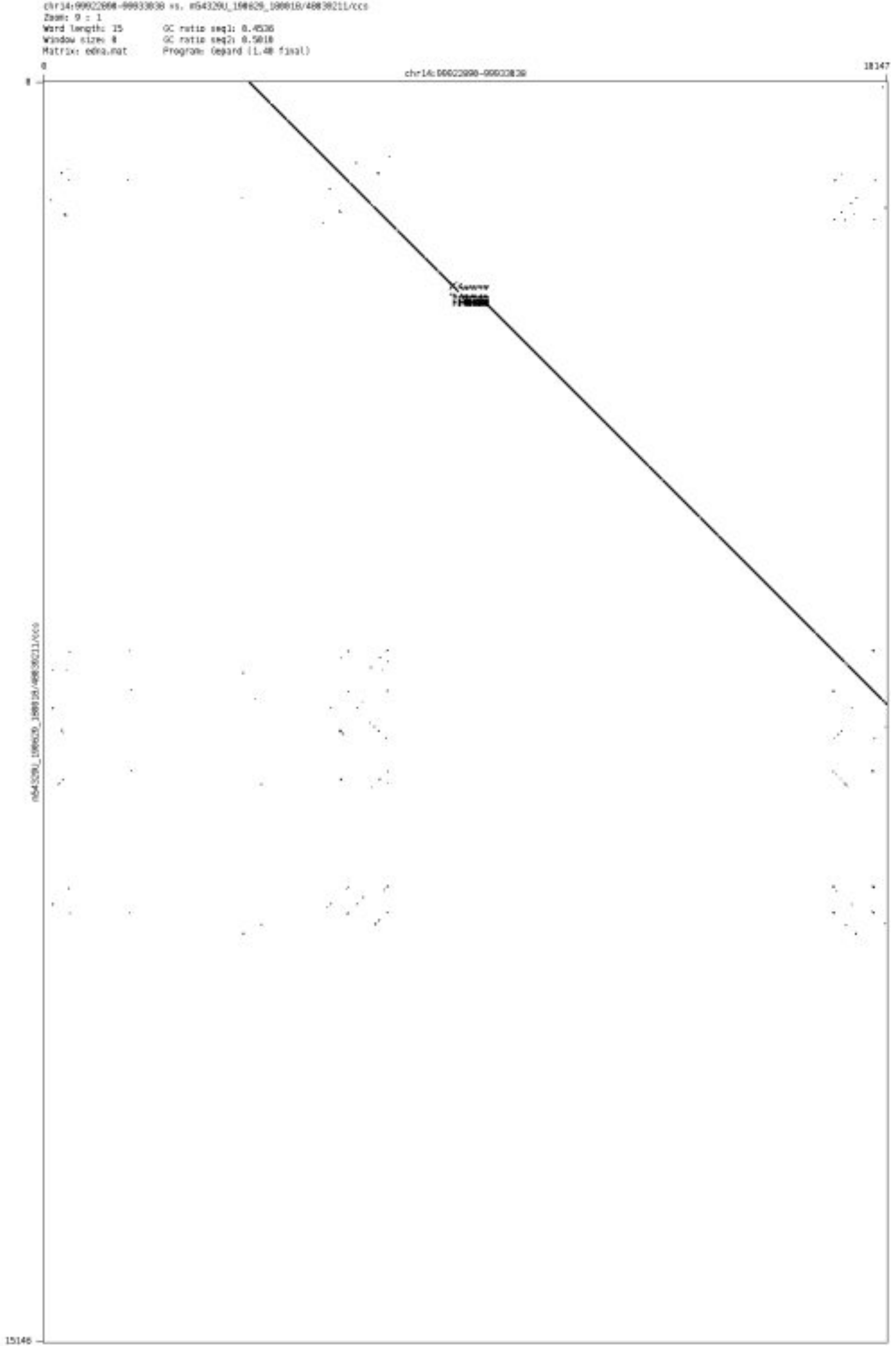

➢ chr14_100873355_100873423

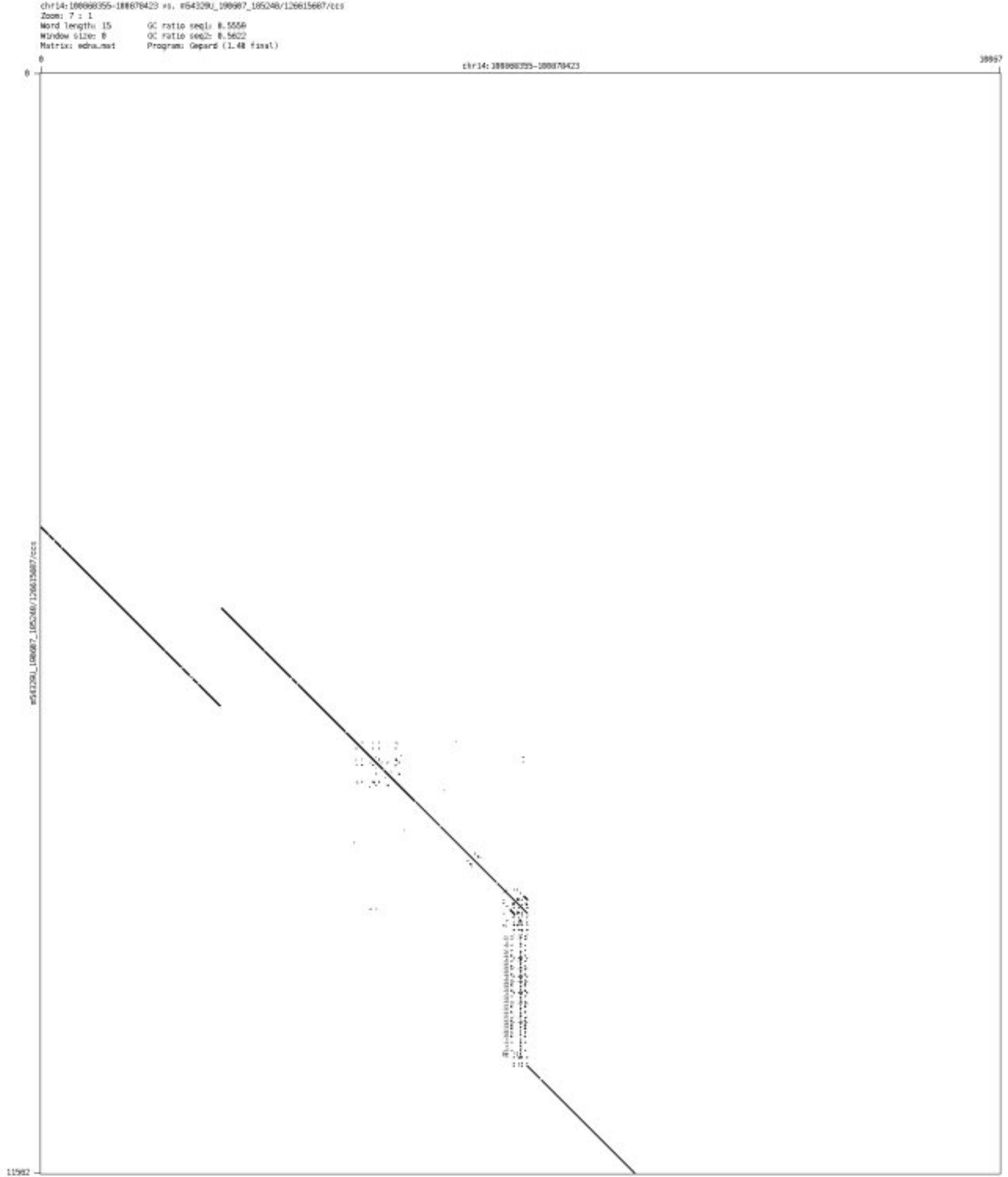

➢ chr14_103896624_103897749

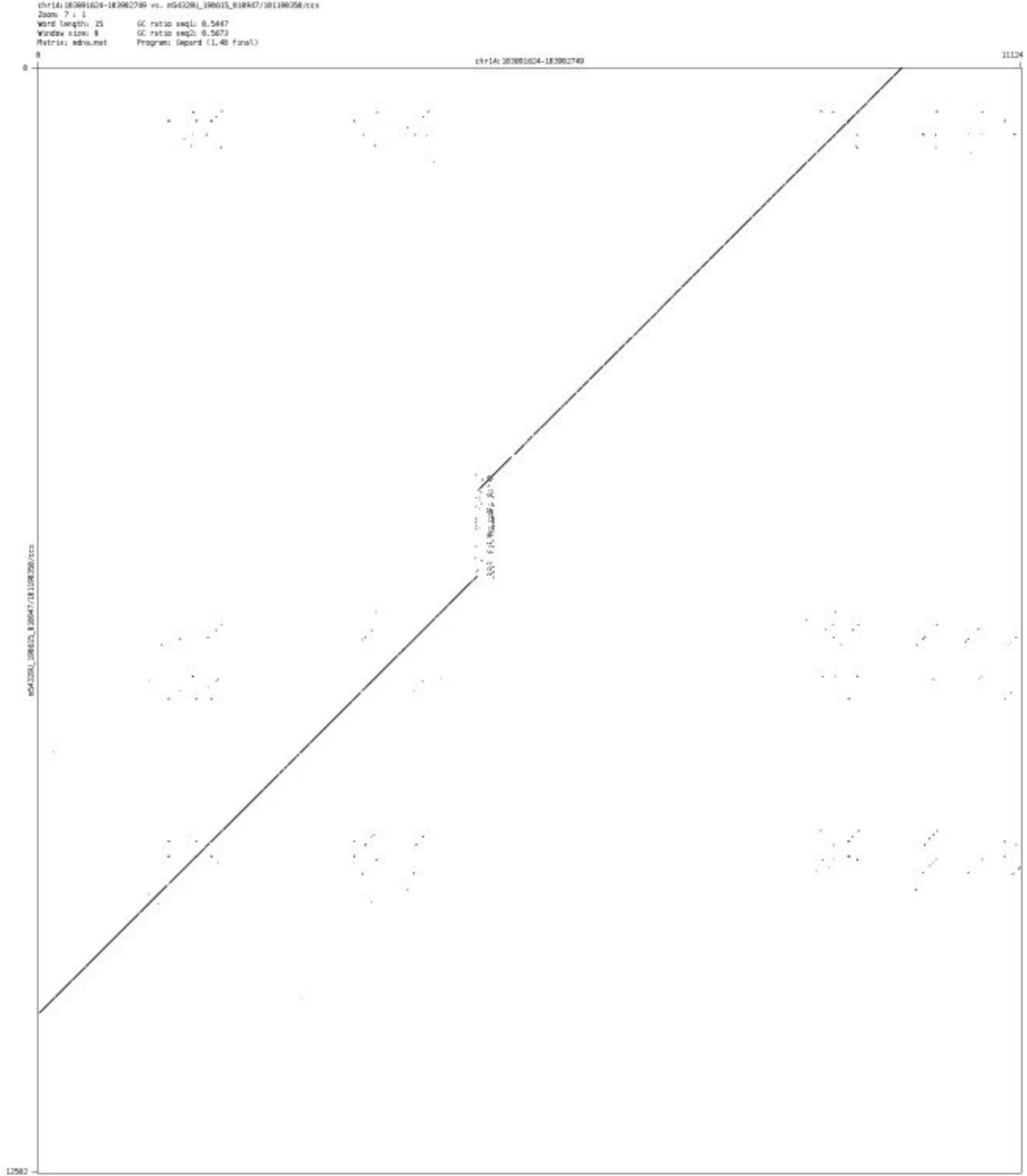

- ➢ chr14_105349868_105350042

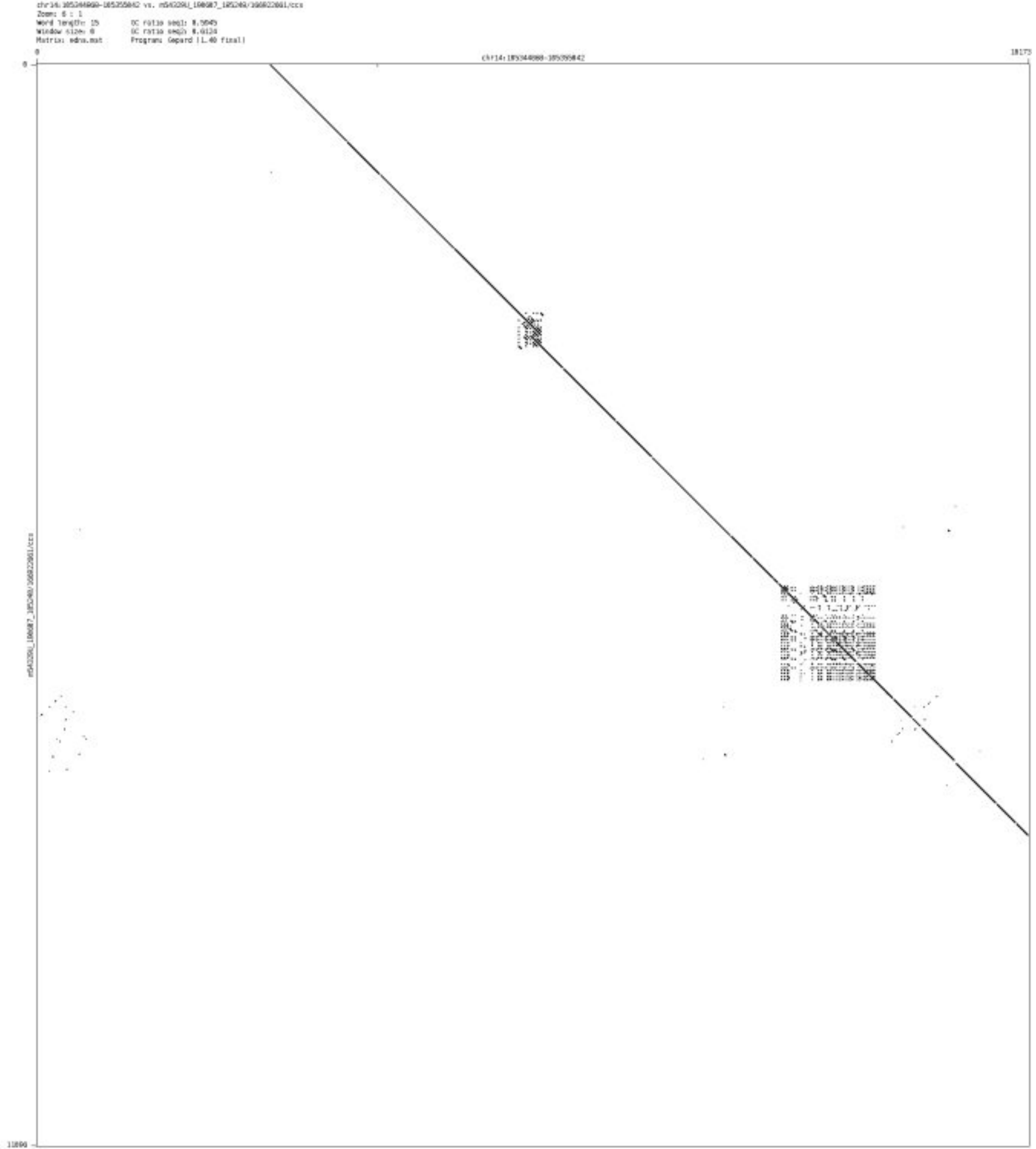

➢ chr15_30173122_30173230

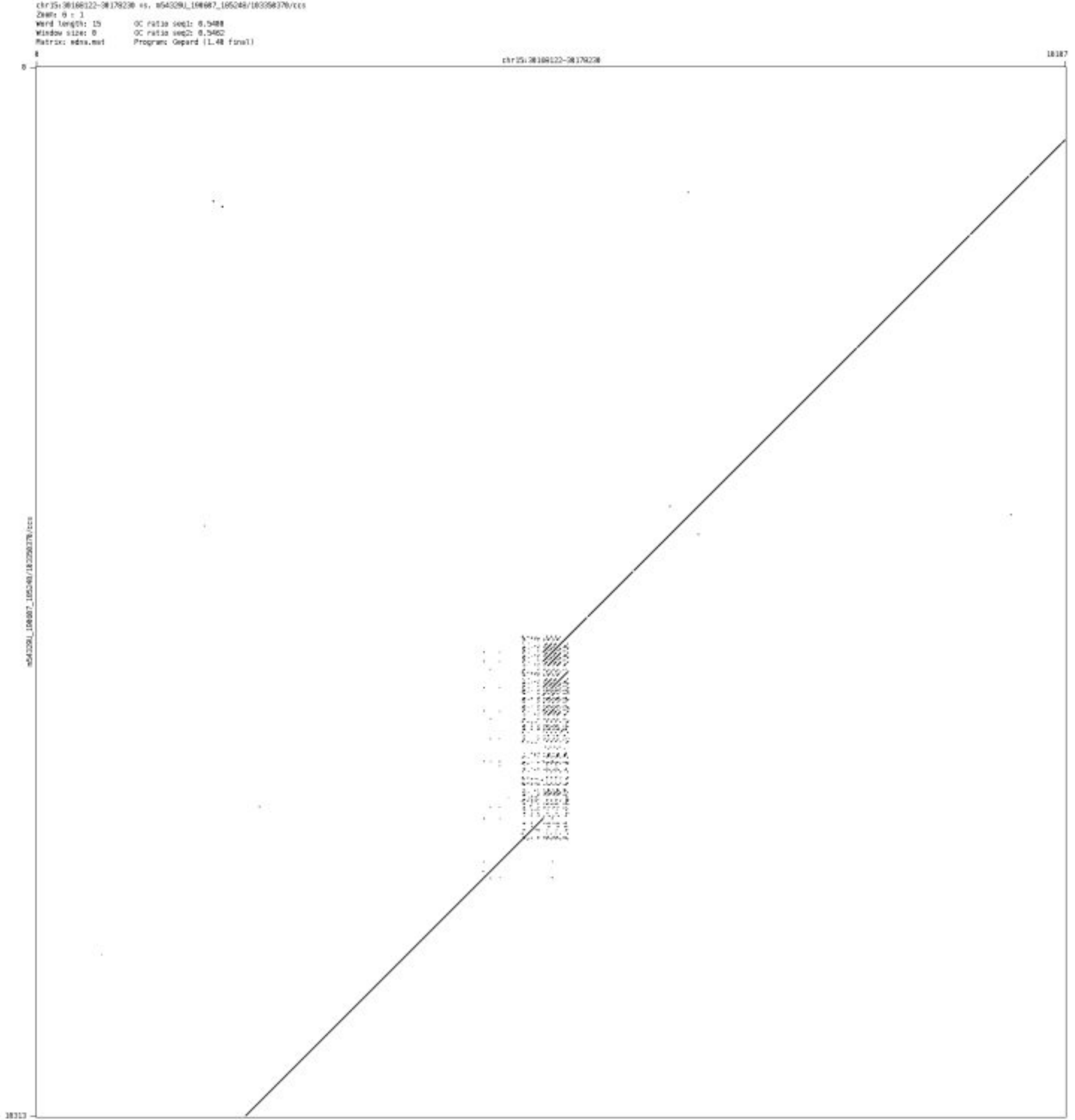

- ➢ chr15_100058267_100058336

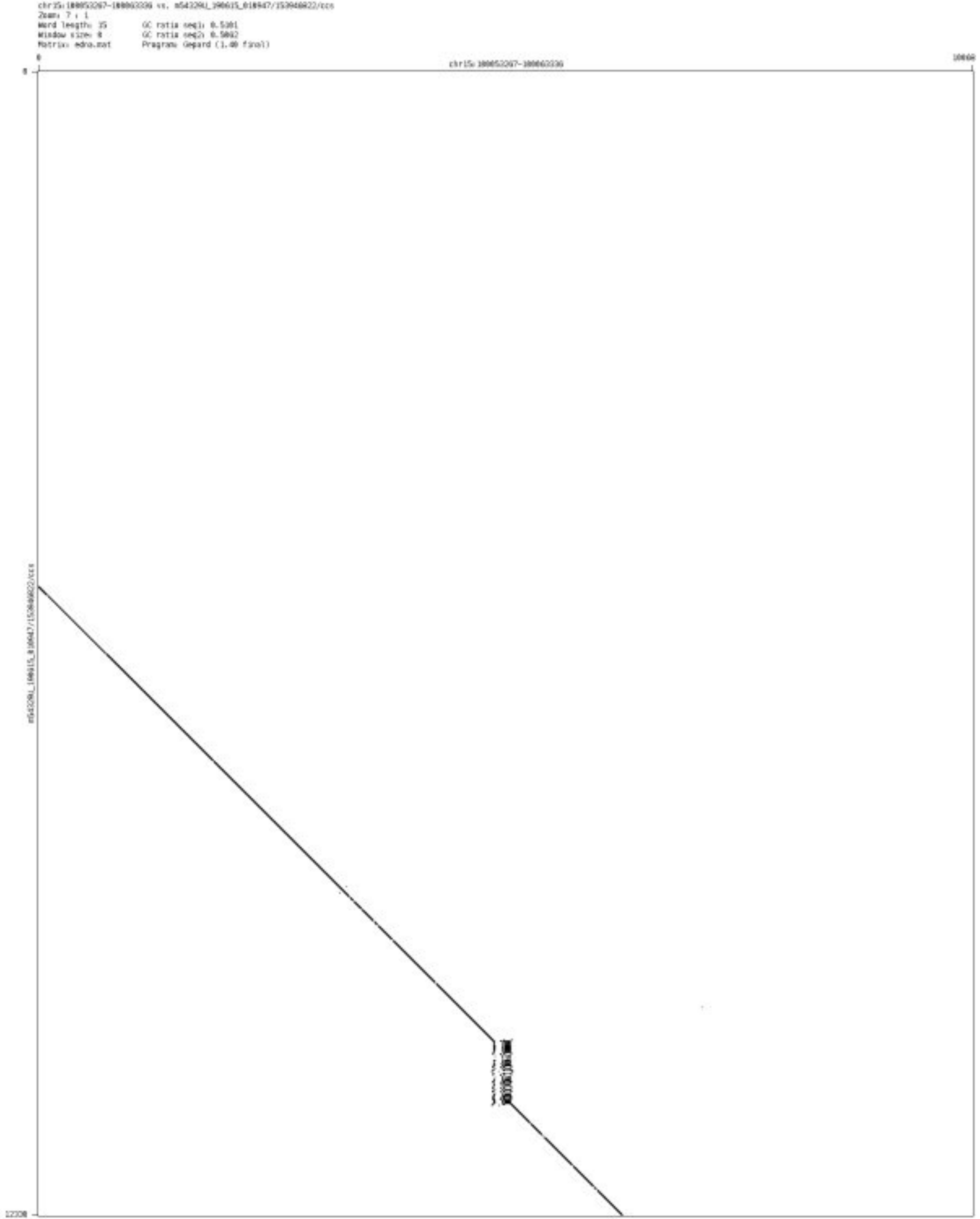

- chr16_818762_818876

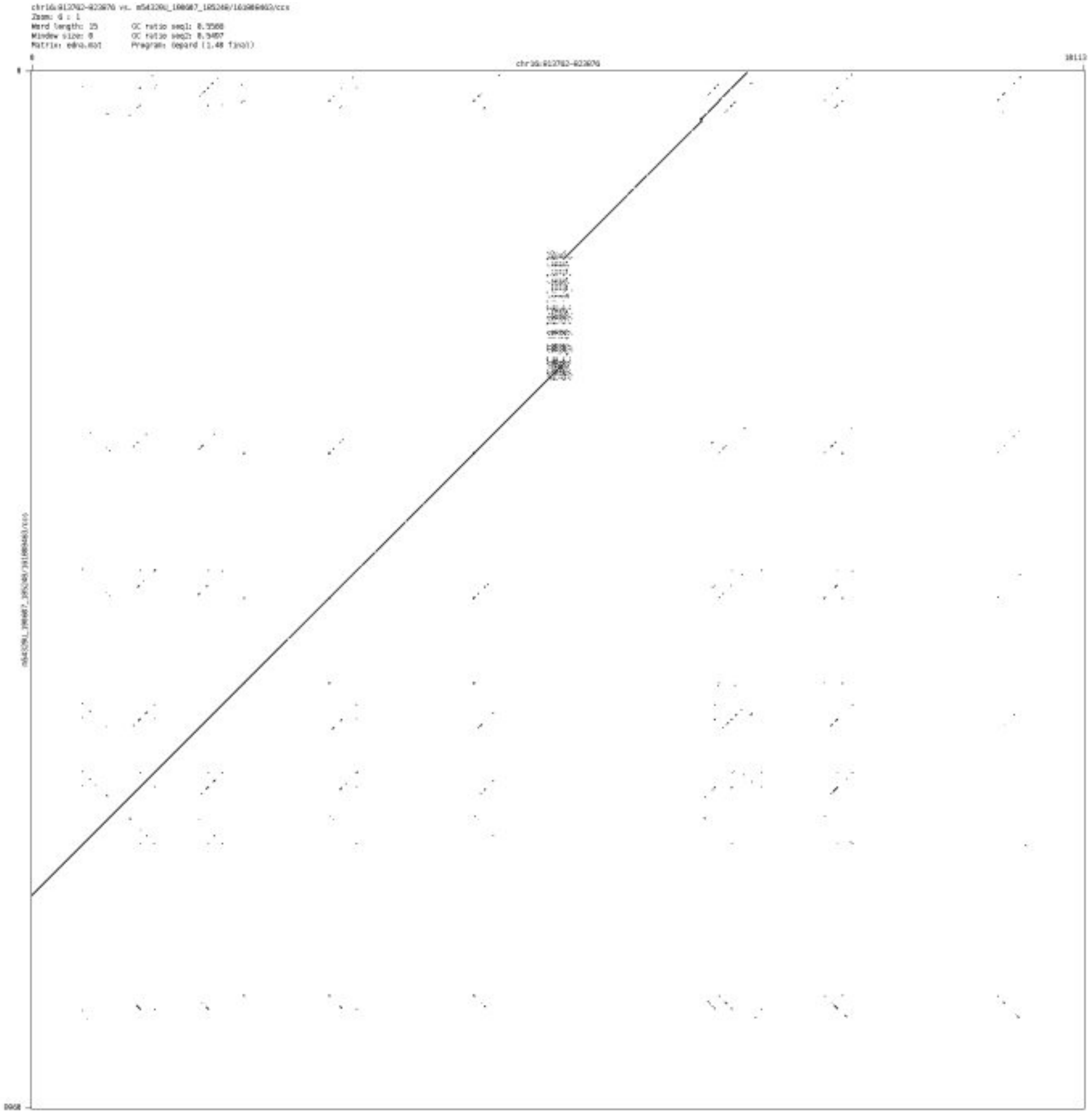

### ➢ chr16_1025314_1025497

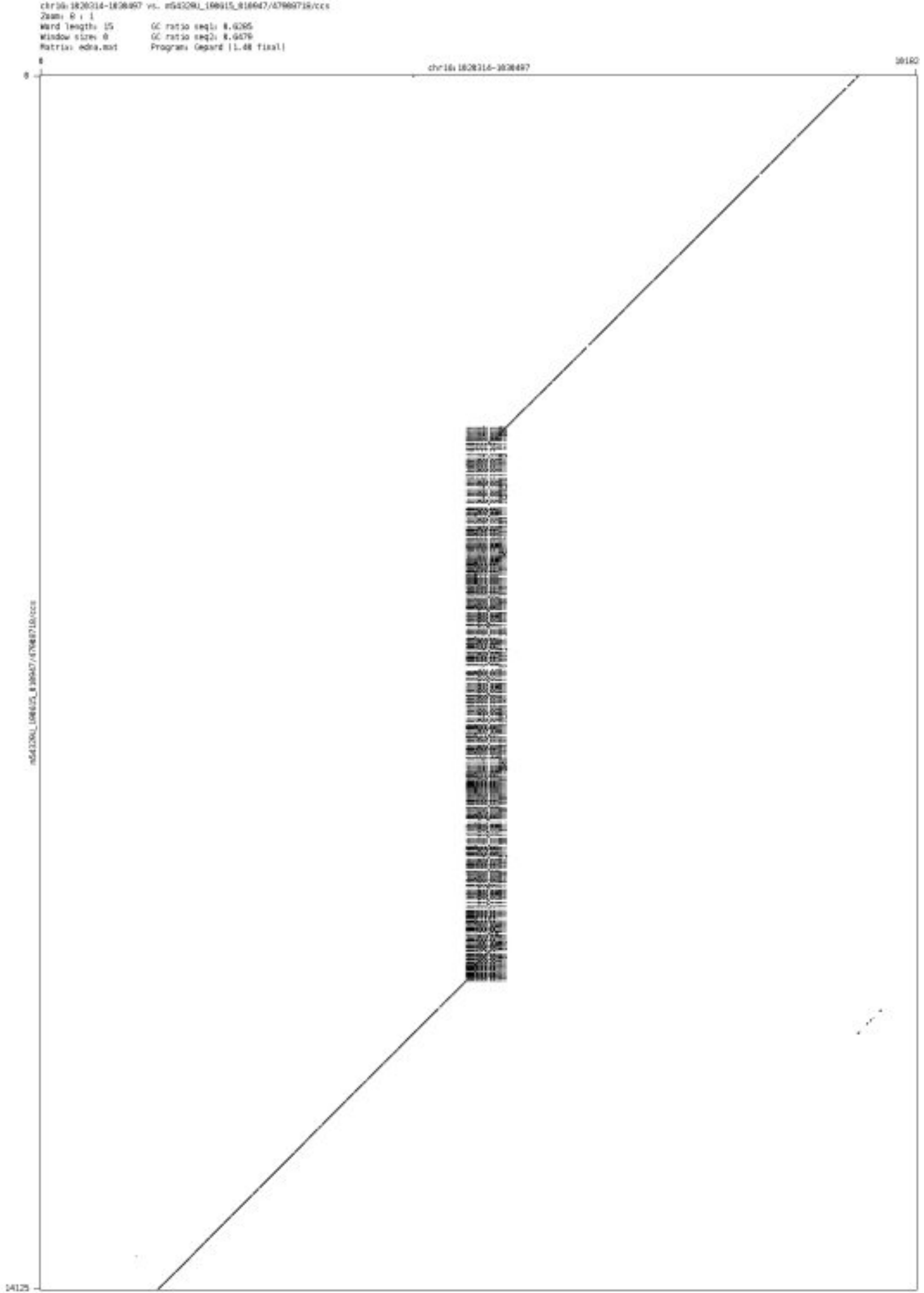

➢ chr16_1185592_1186068

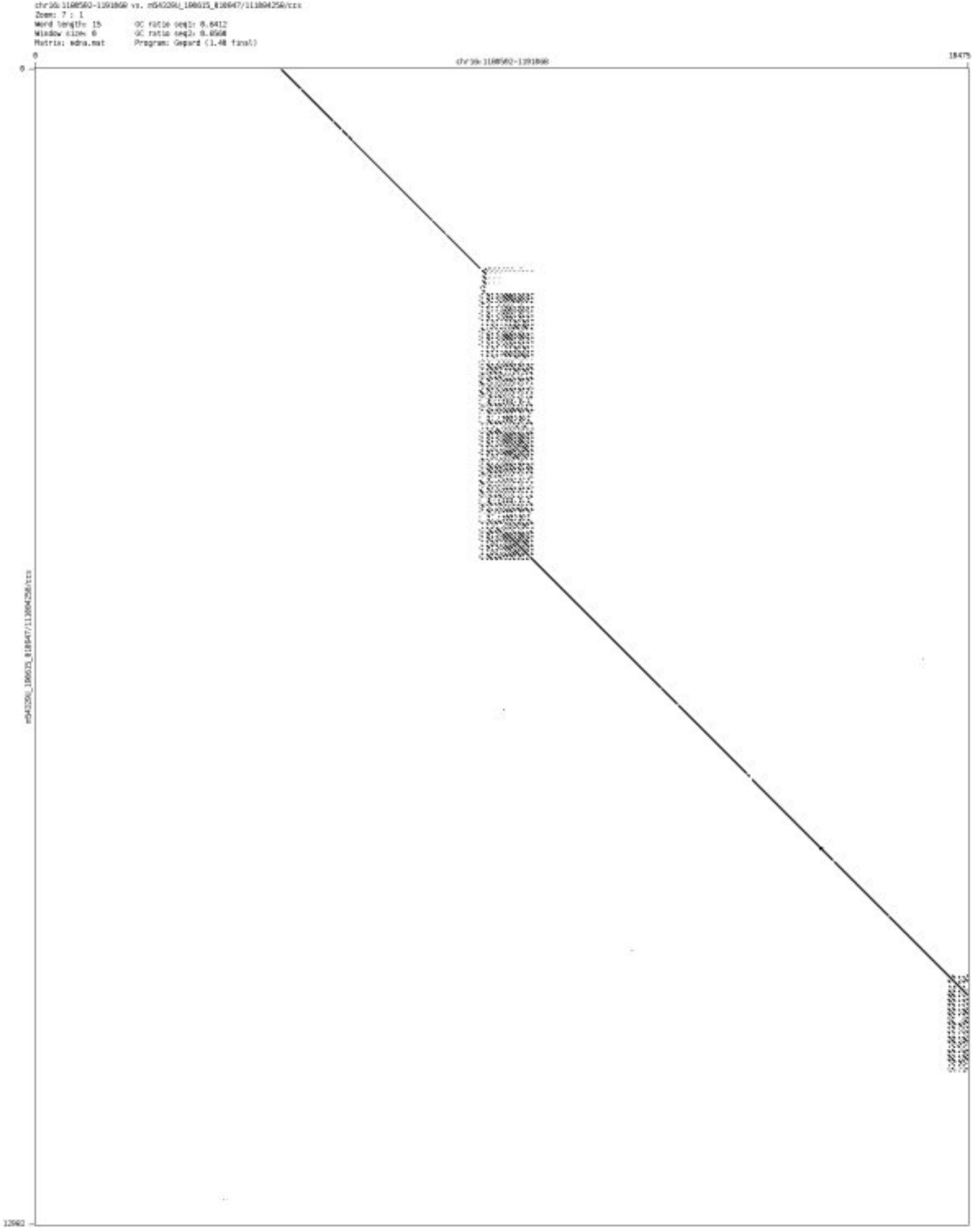

- chr16_10040023_10040092

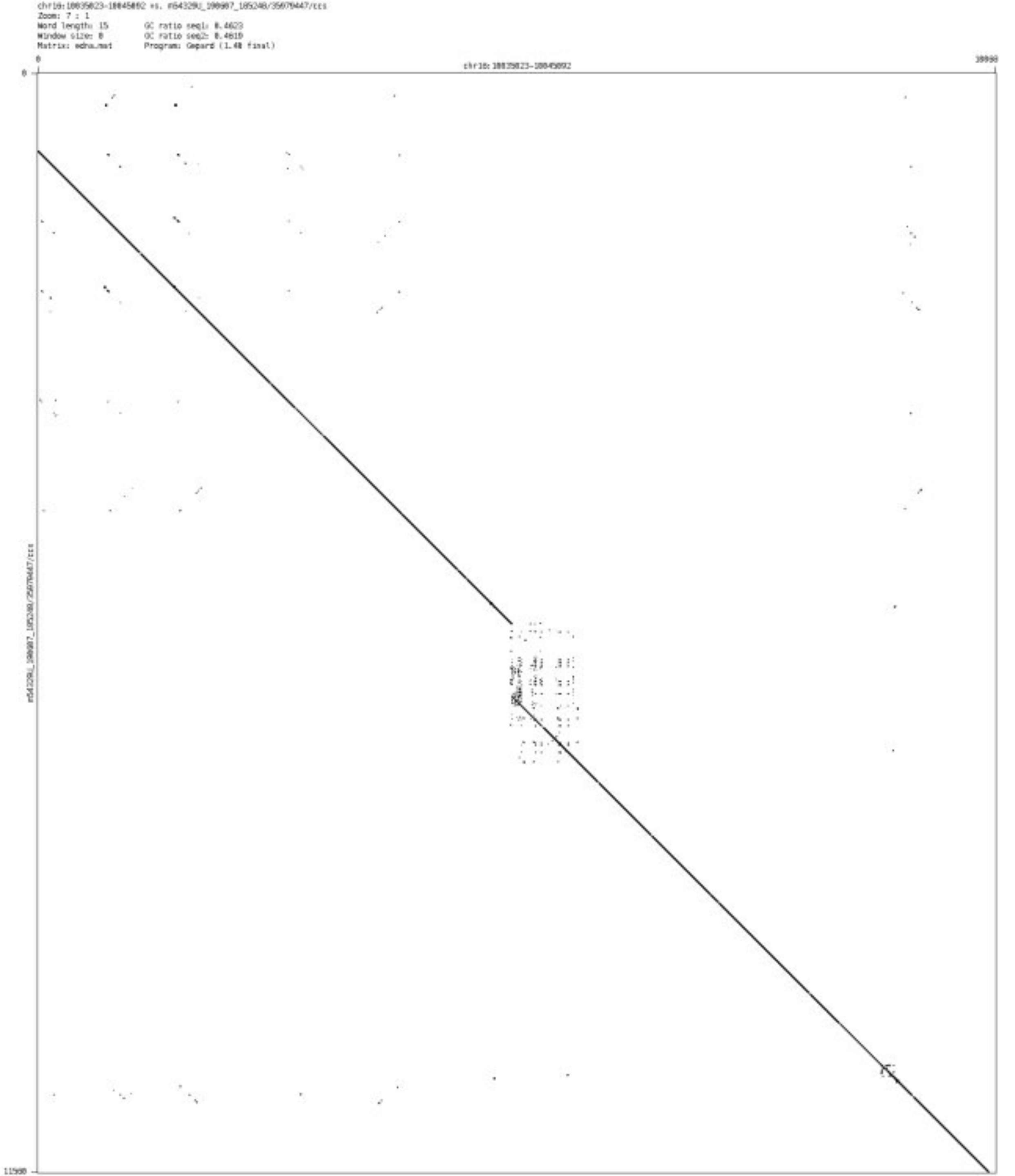

- chr16_83950511_83950871

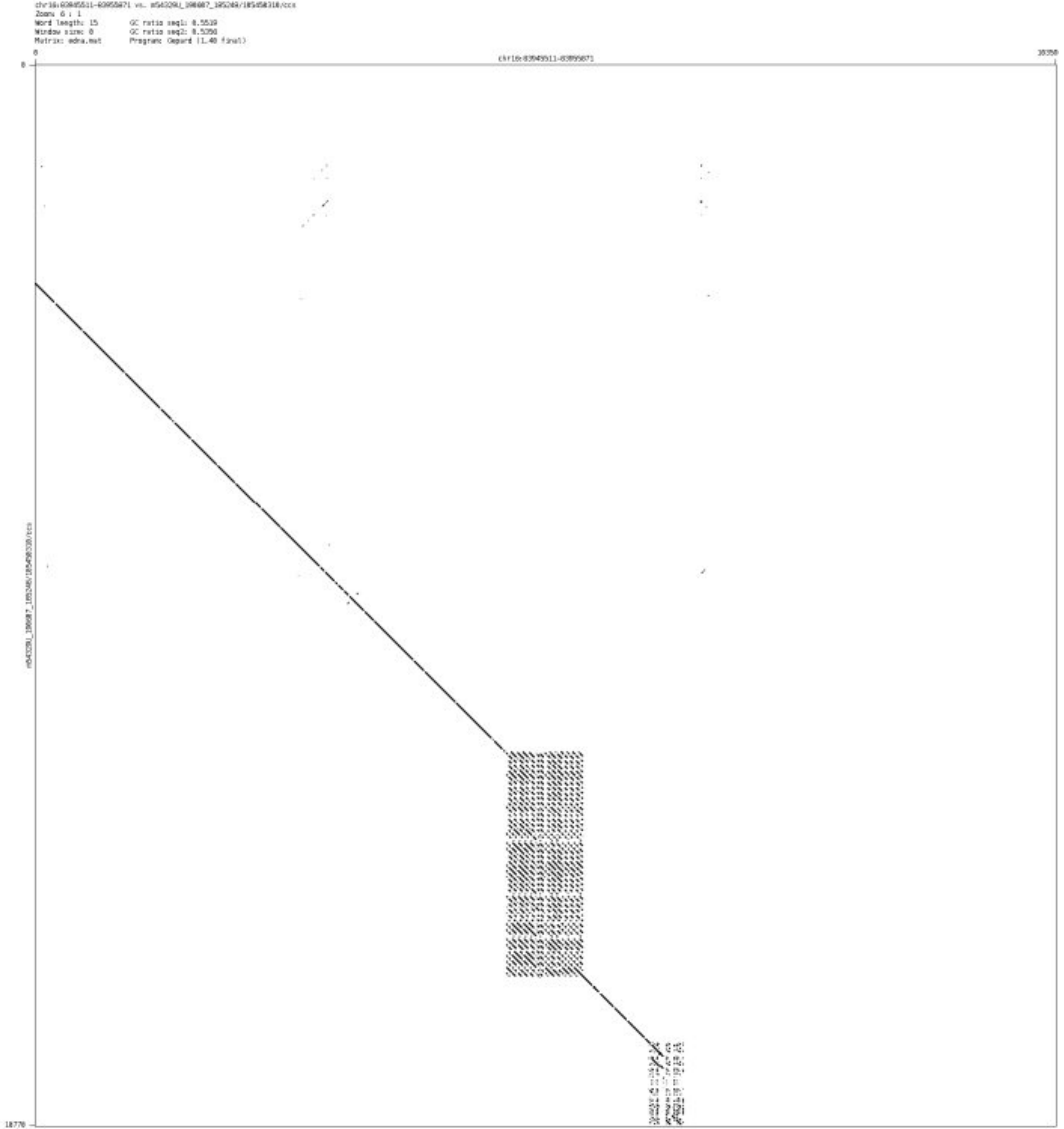

- chr16_84050534_84051508

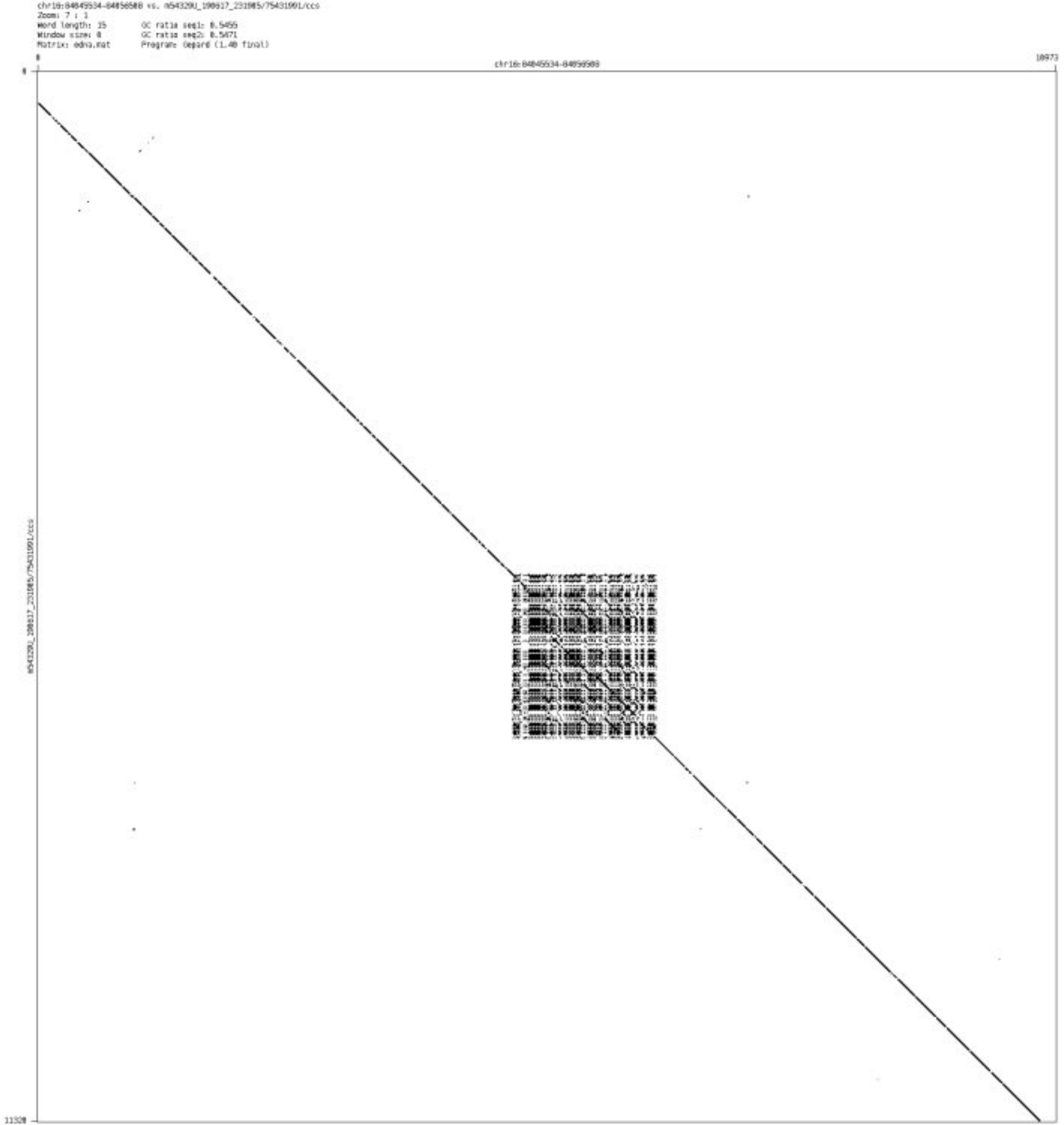

➤ chr17_809815_810302

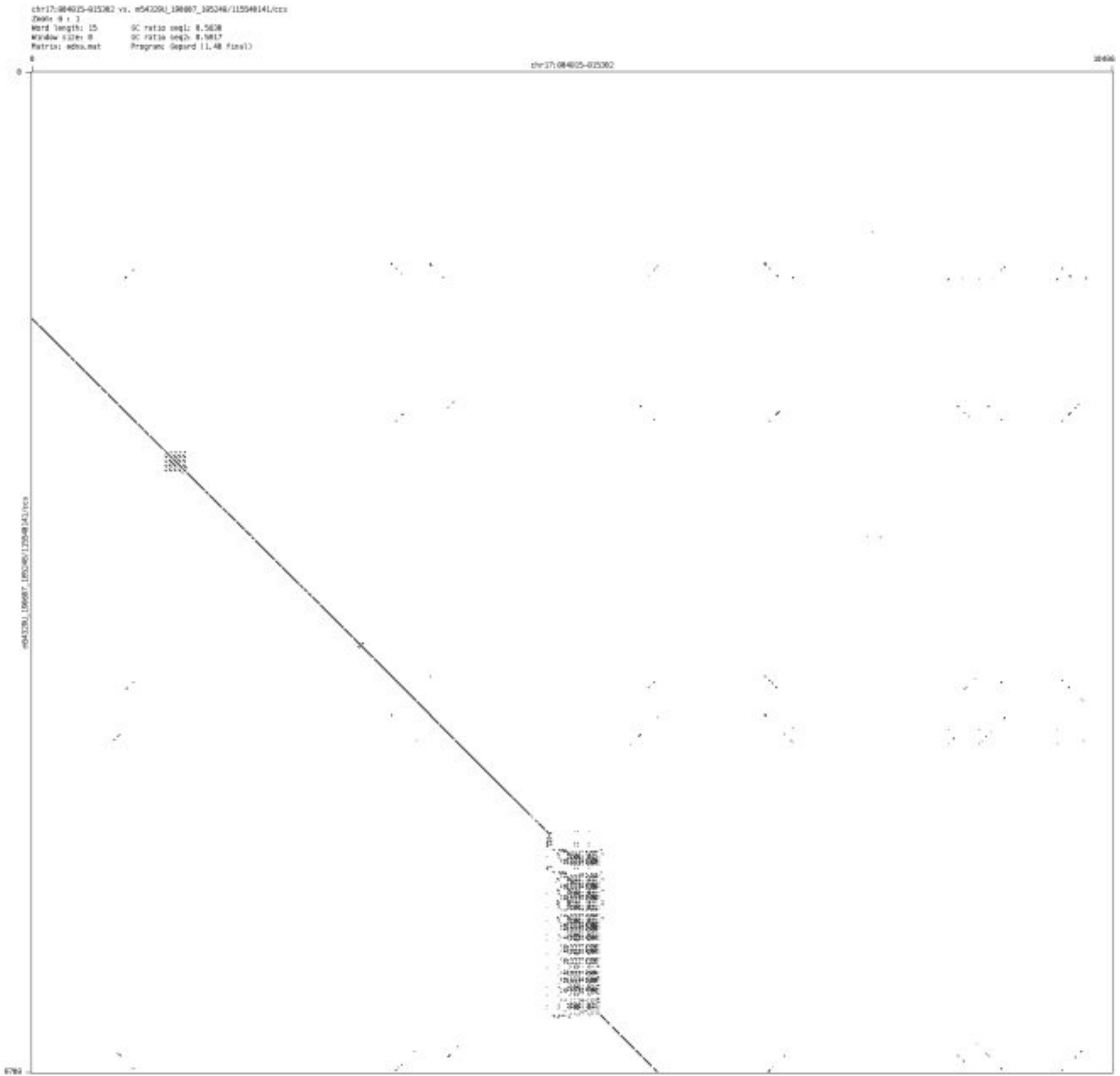

➢ chr17_841437_841560

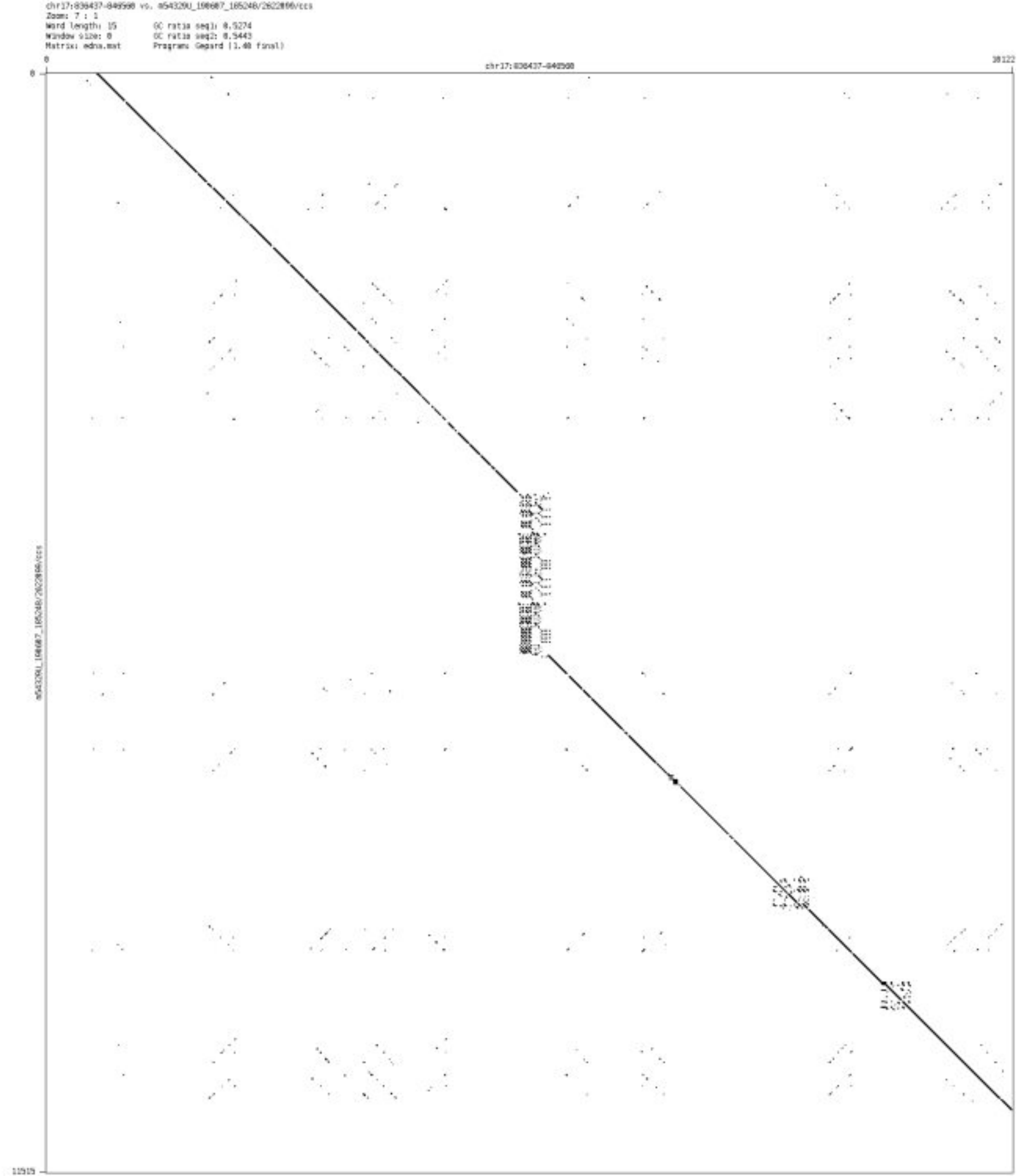

➢ chr17_1159495_1159678

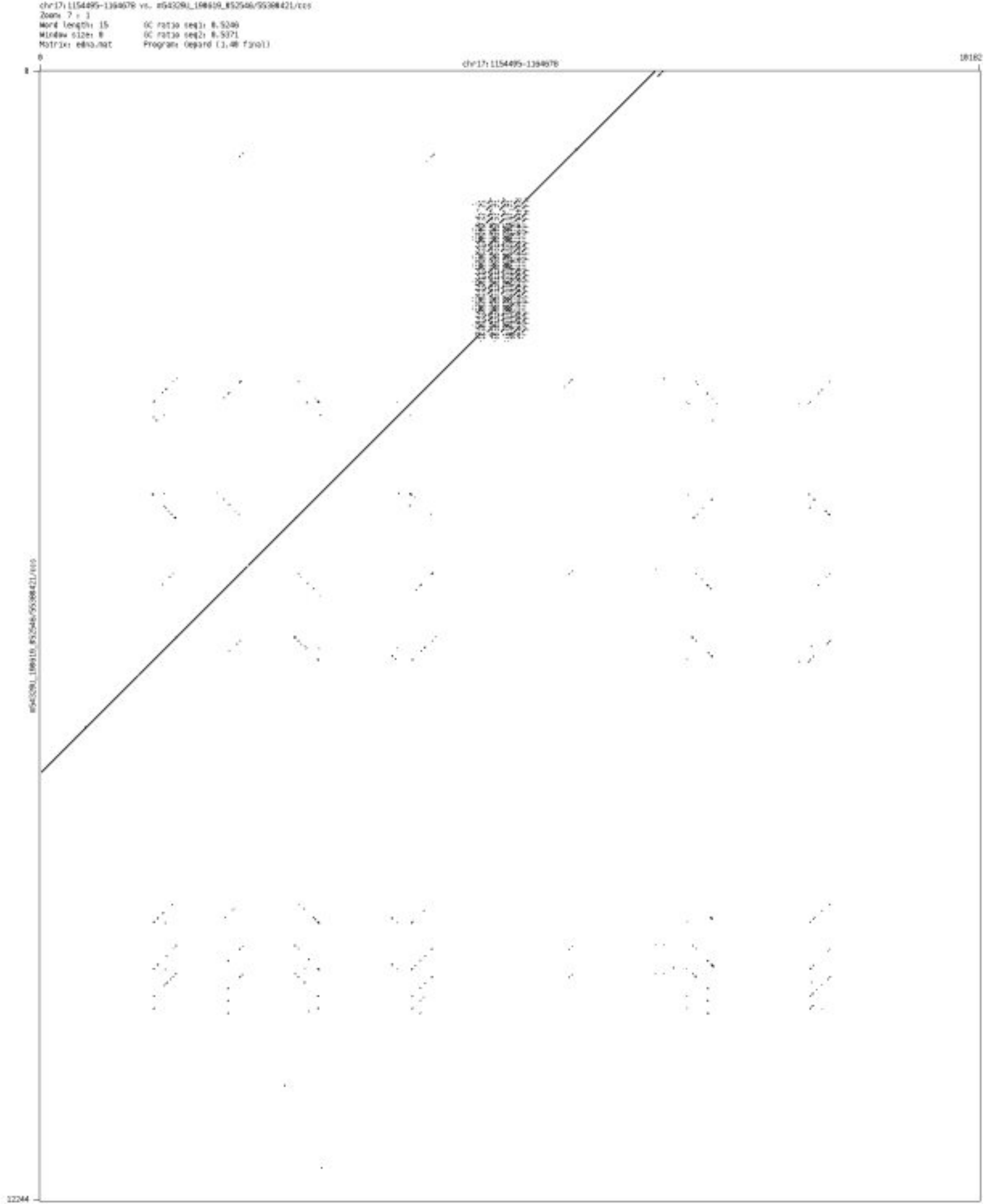

- chr17_6194135_6194242

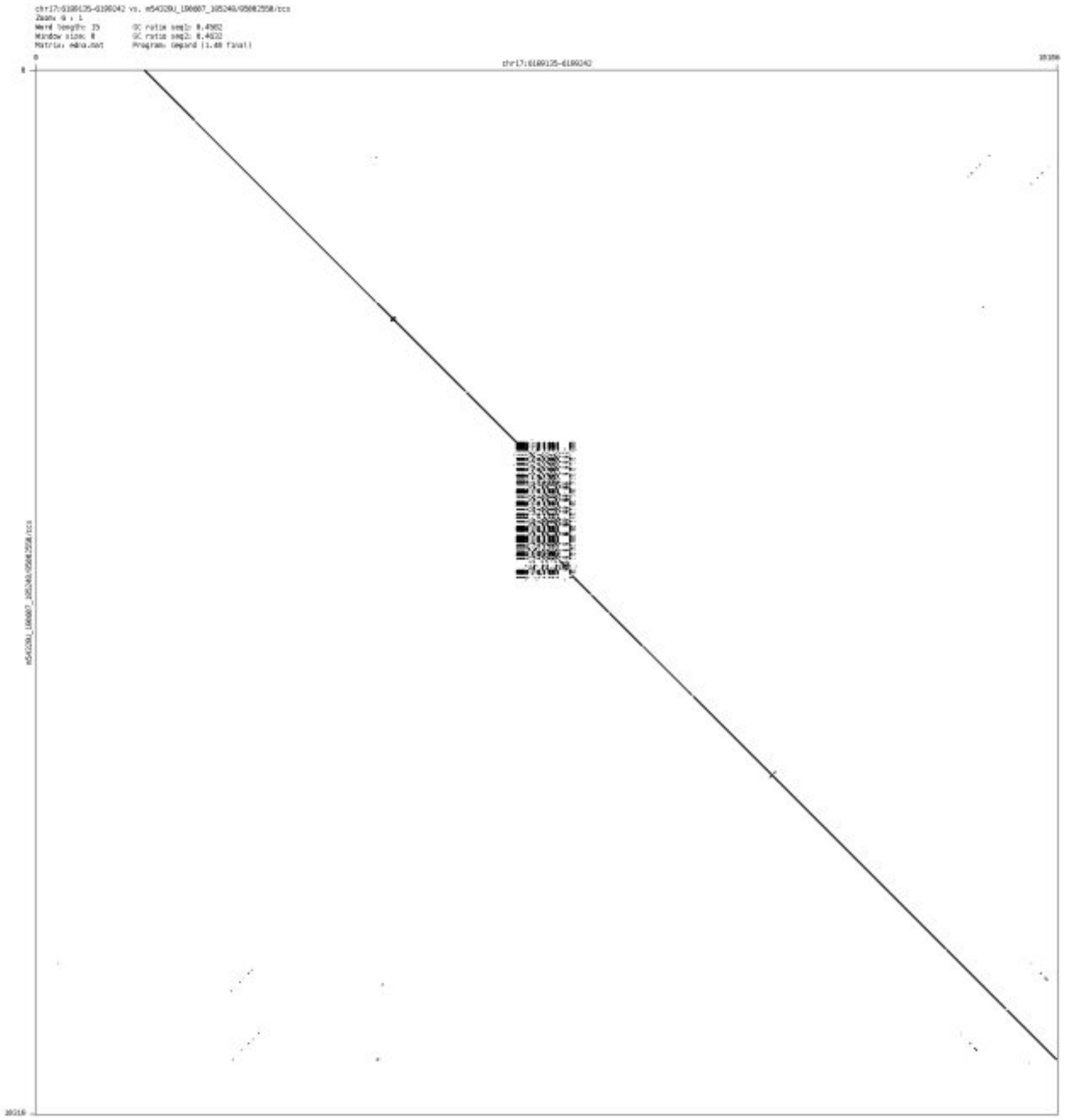

- chr17_17460134_17460311

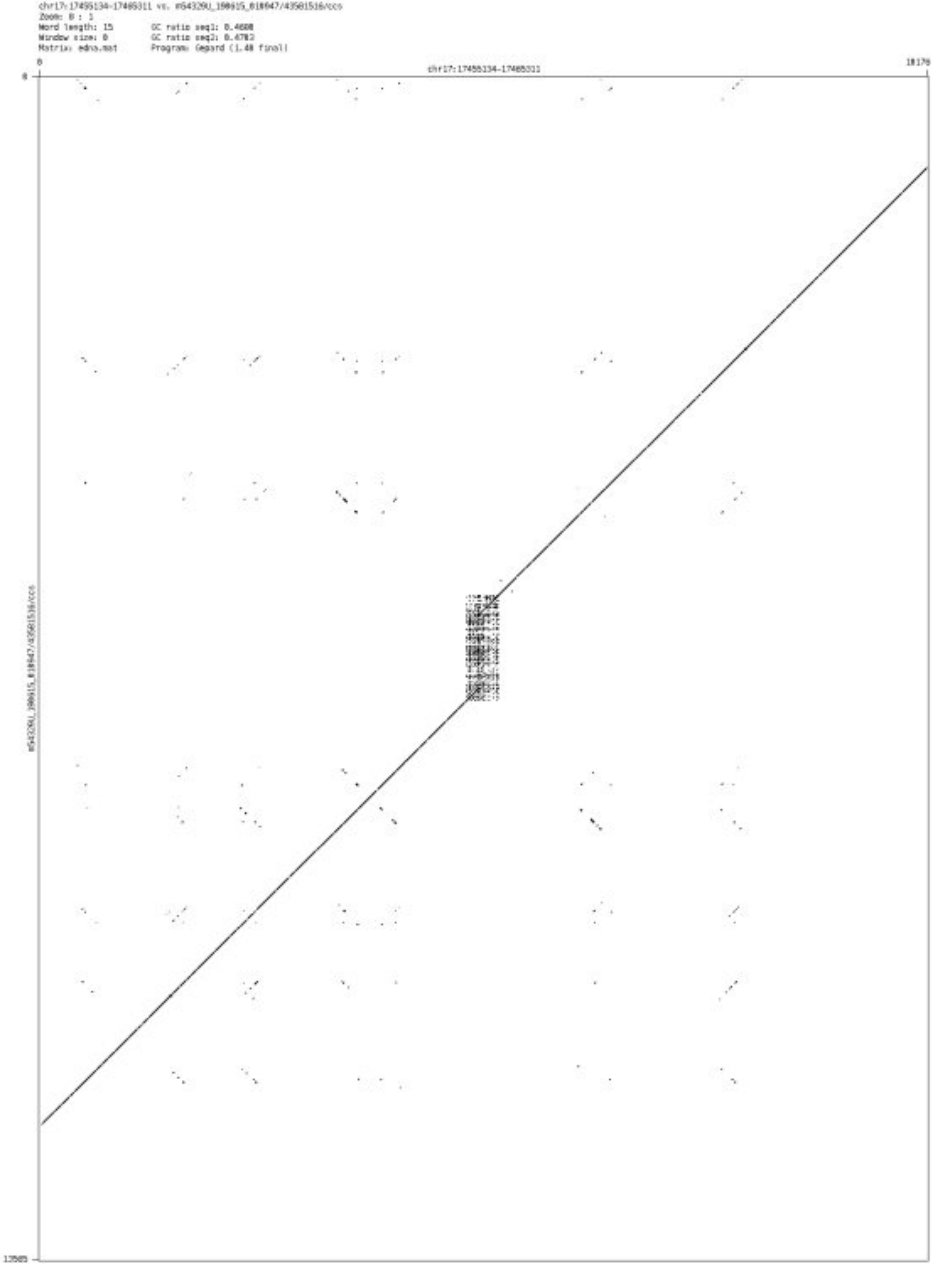

➢ chr17_56792755_56792881

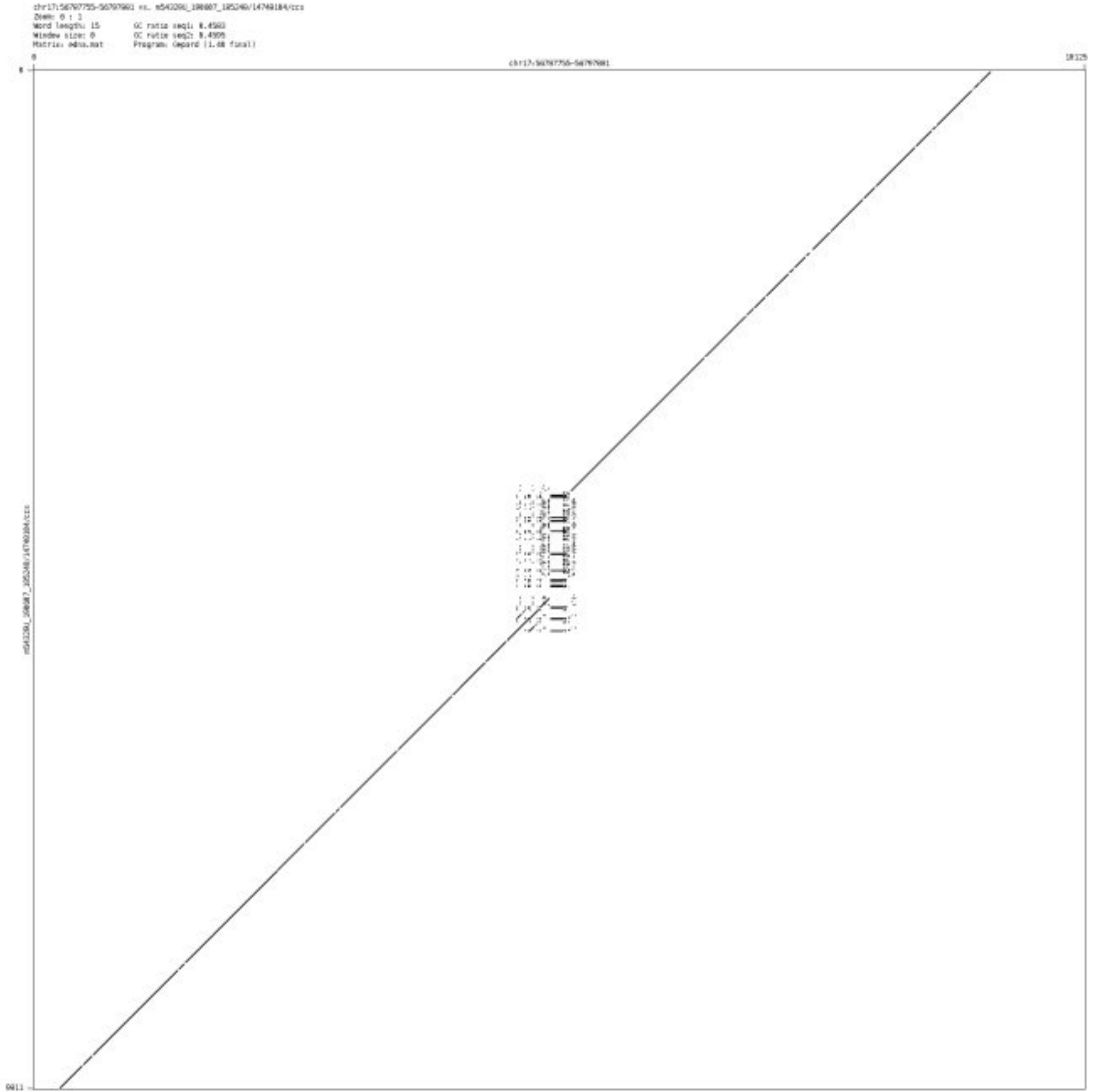

- chr17_66798674_66798865

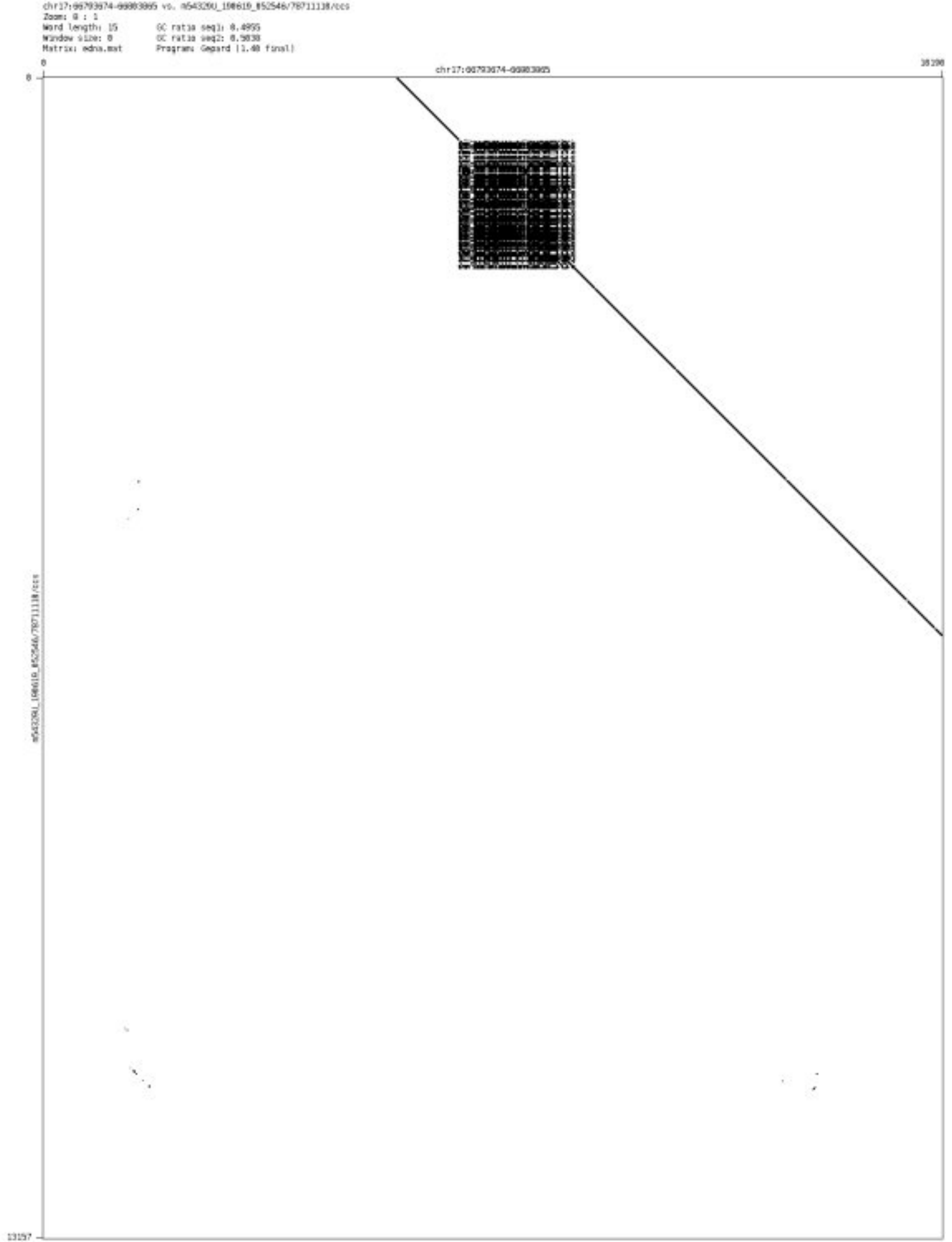

- chr17_74732756_74733512

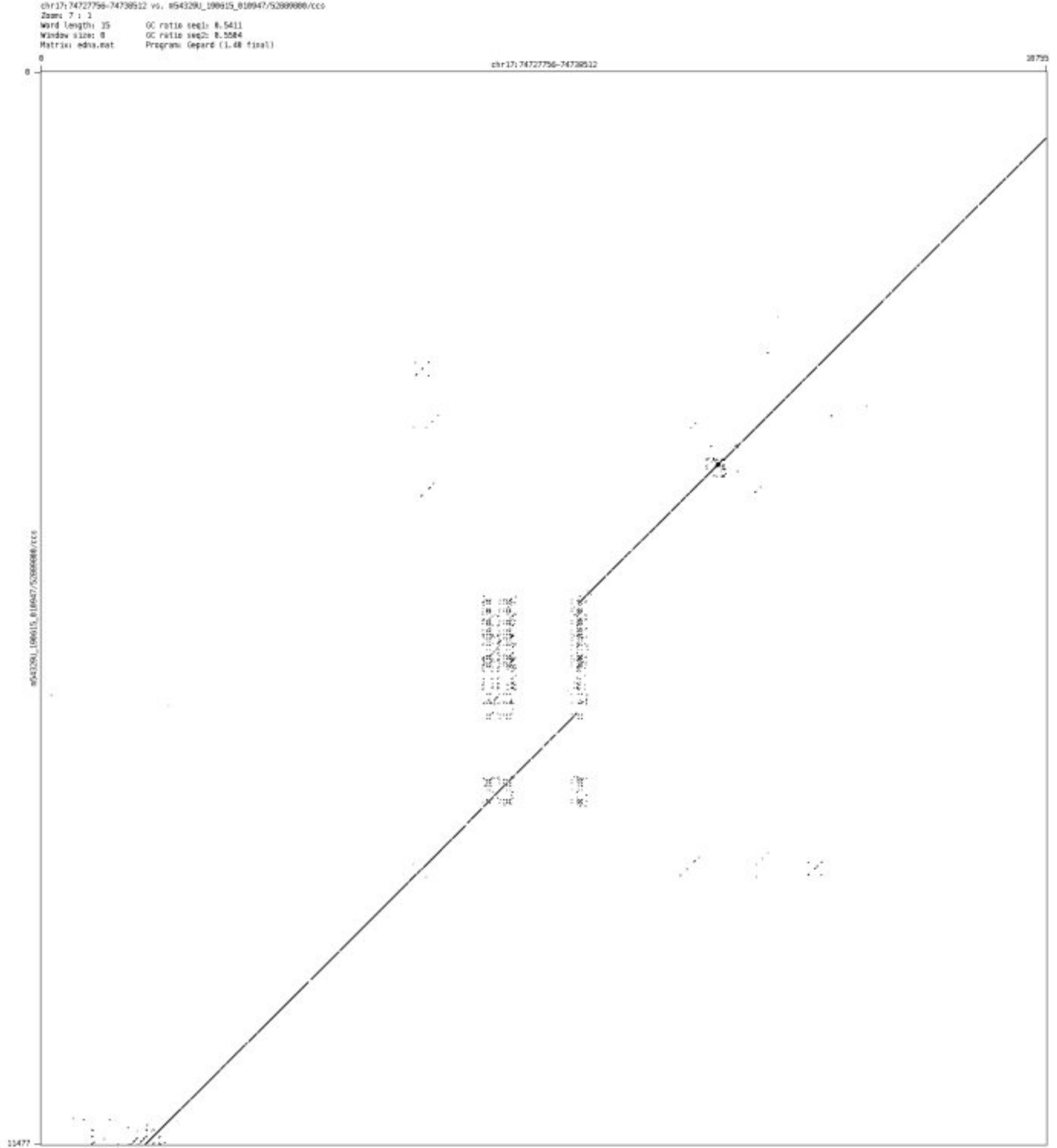

- ➢ chr17_80497958_80498077

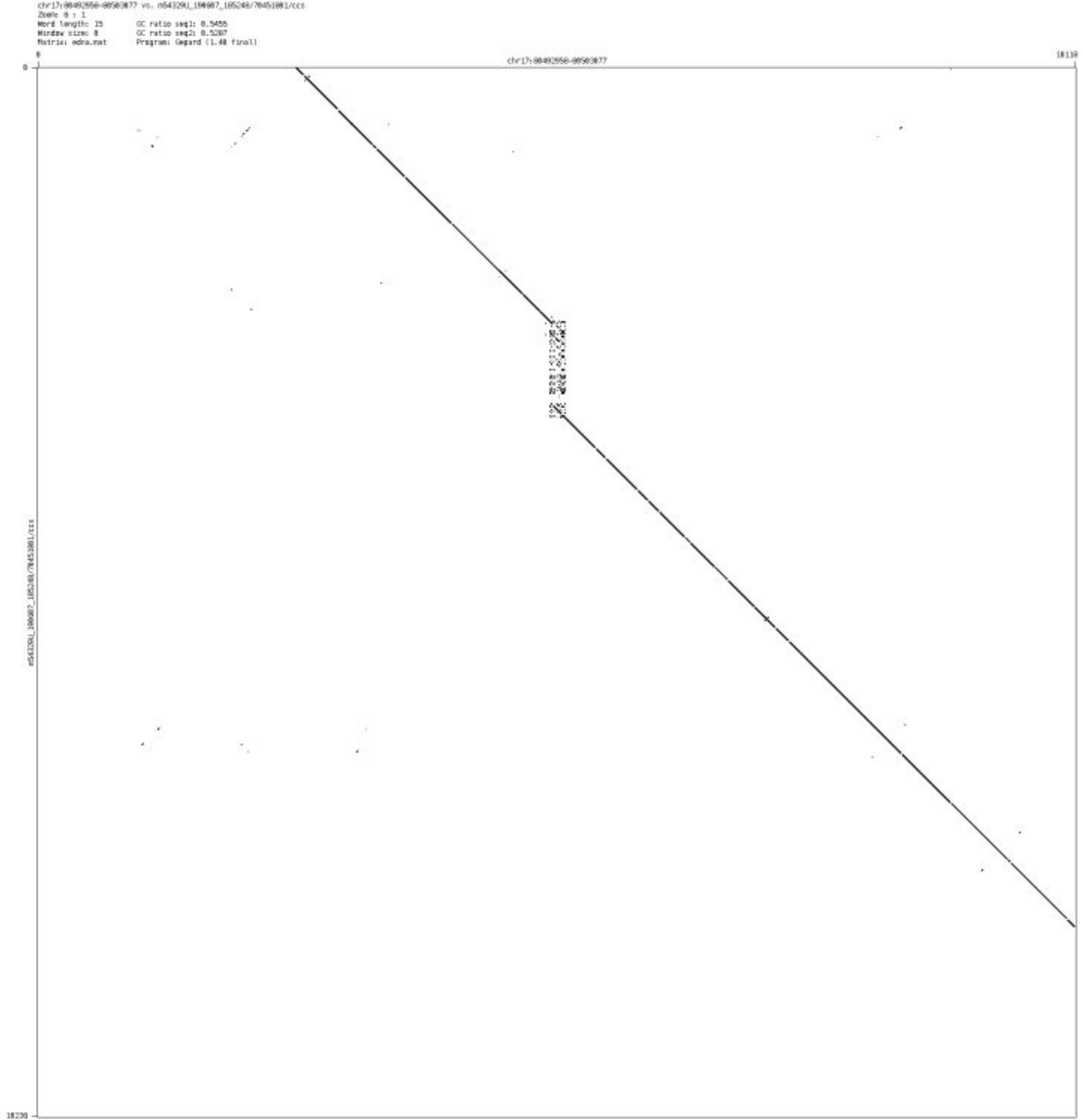

- chr17_80743782_80744966

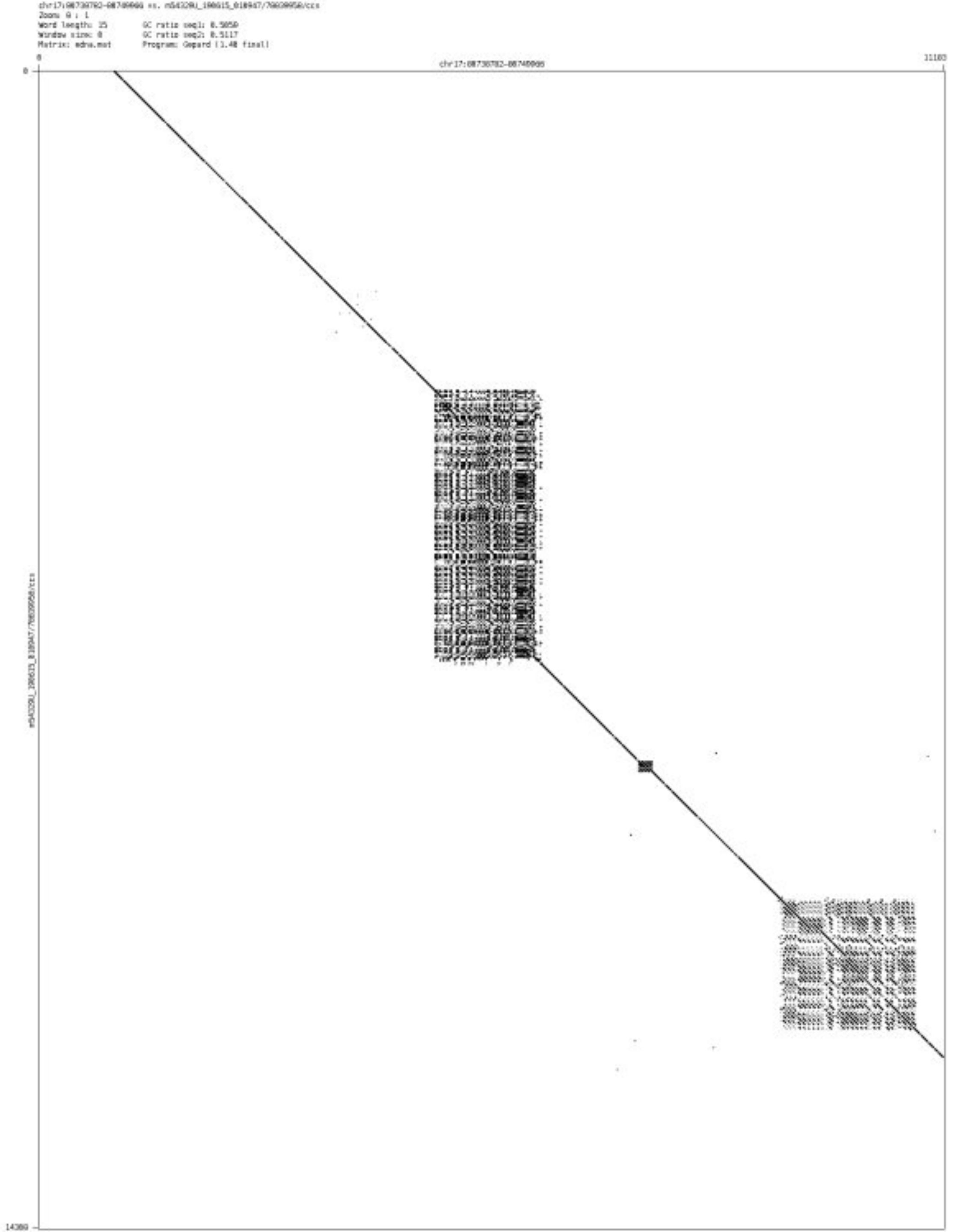

- chr17_81249381_81249456

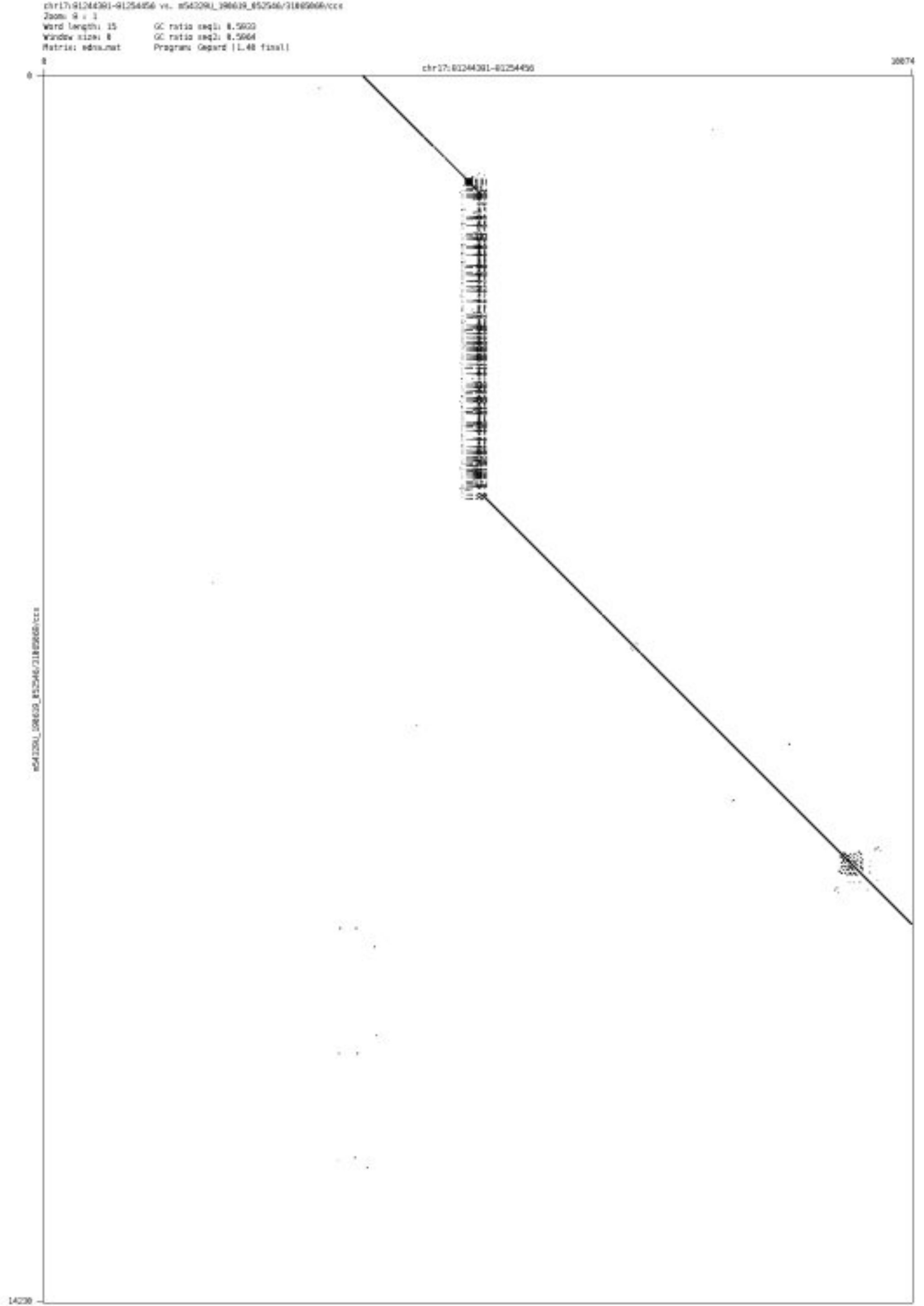

➢ chr17_81425588_81426317

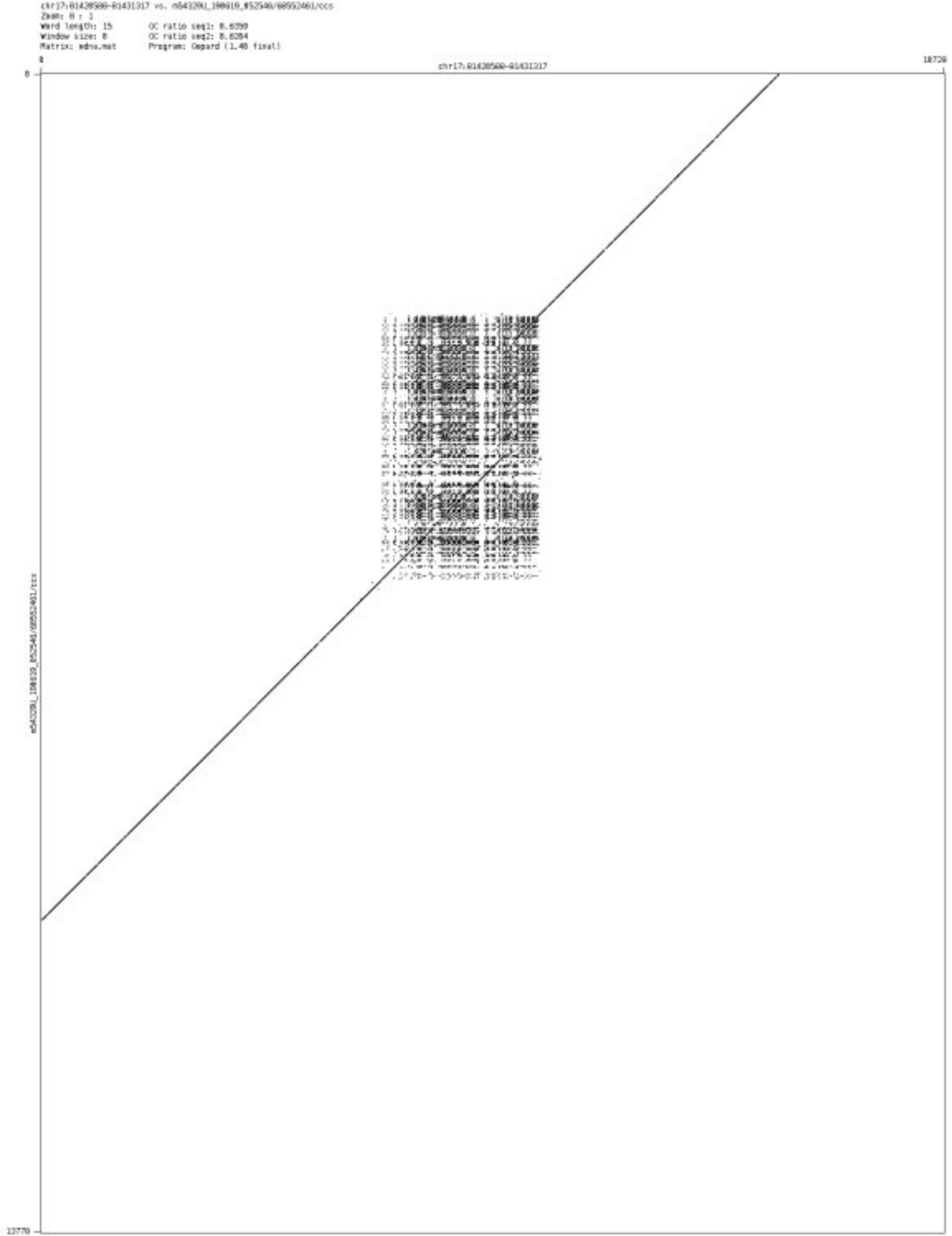

- chr18_4510874_4510952

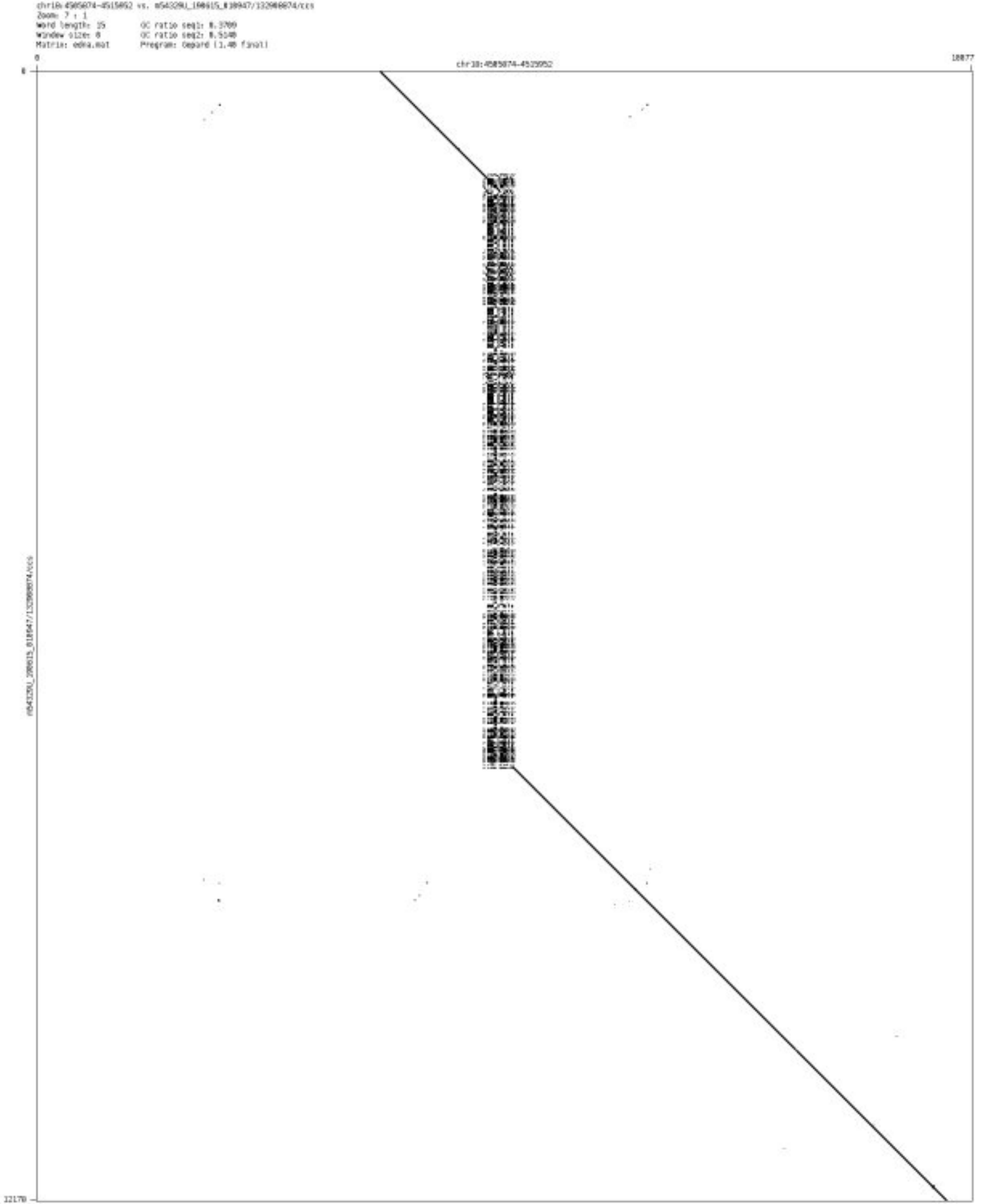

➢ chr18_22232227_22232573

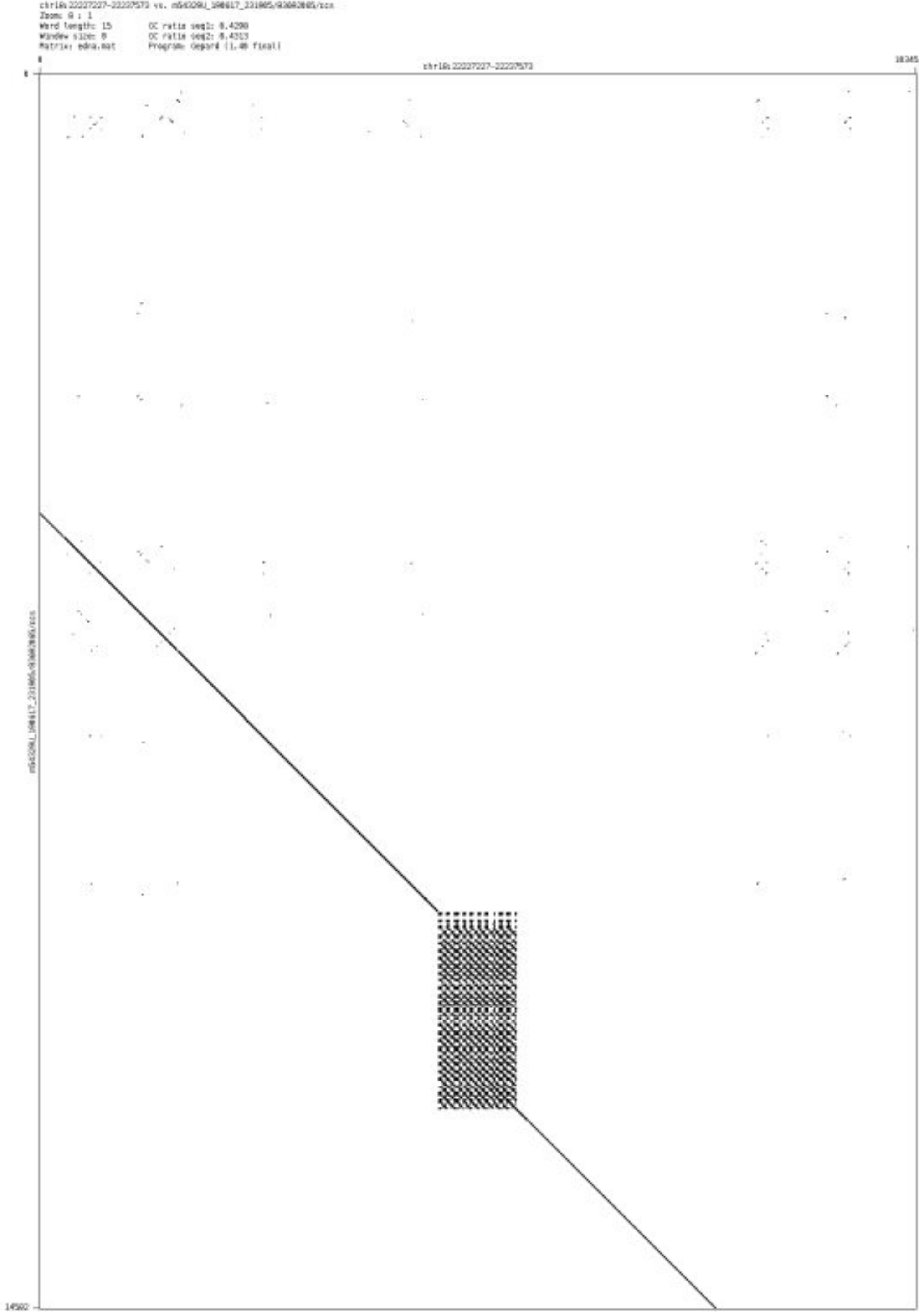

➢ chr18_36463341_36463429

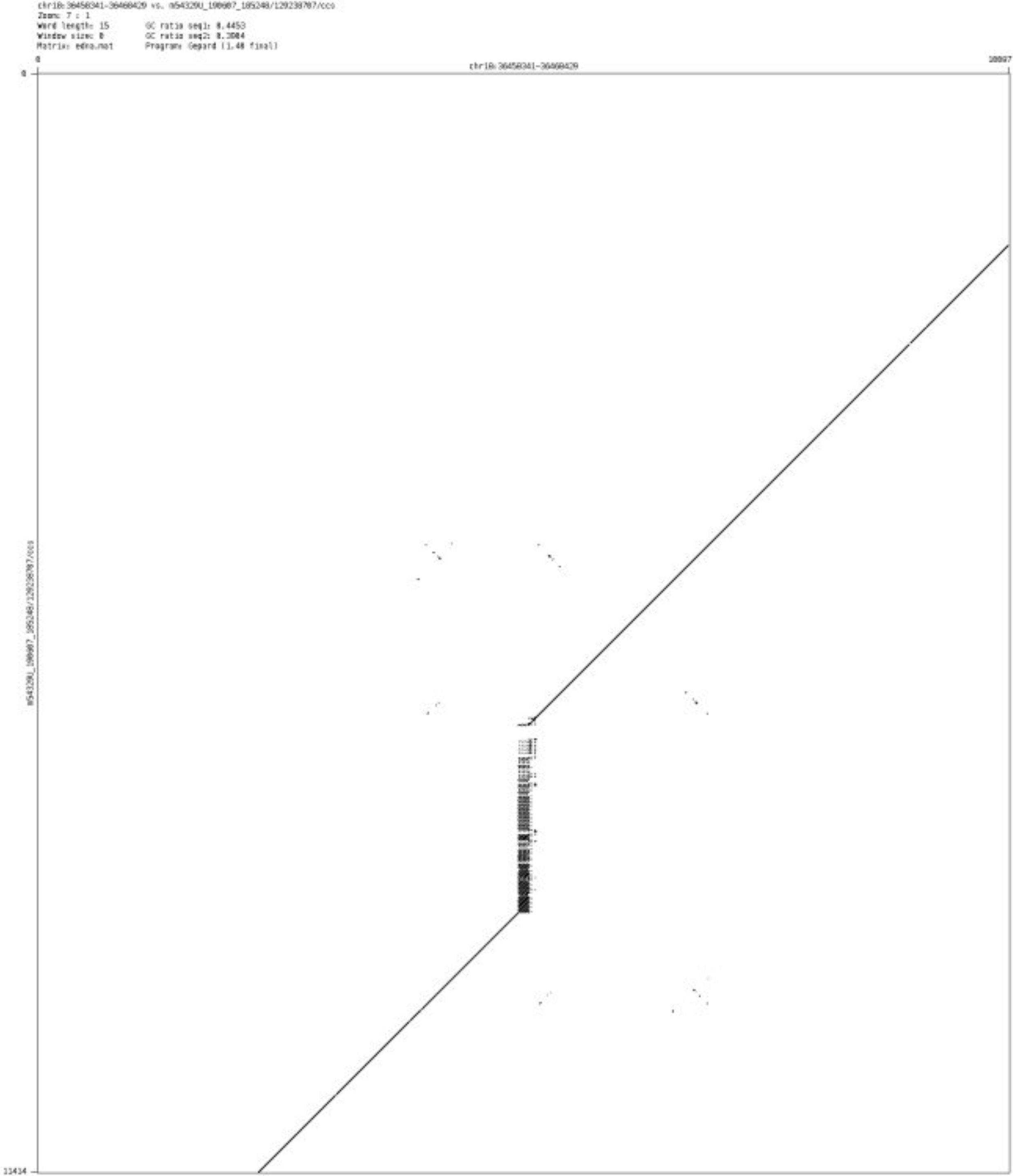

- chr18_49067249_49067378

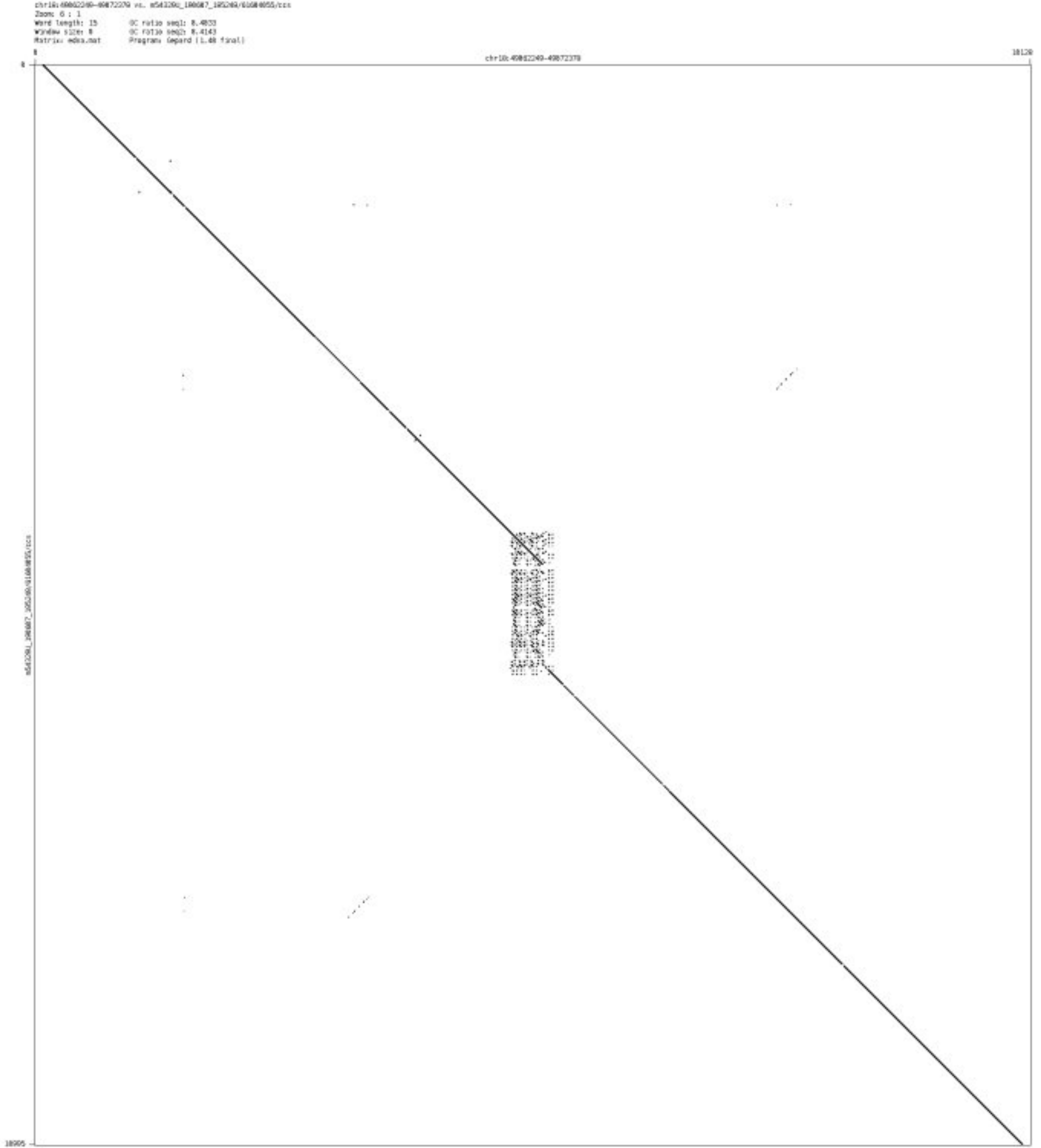

- chr18_74616915_74617116

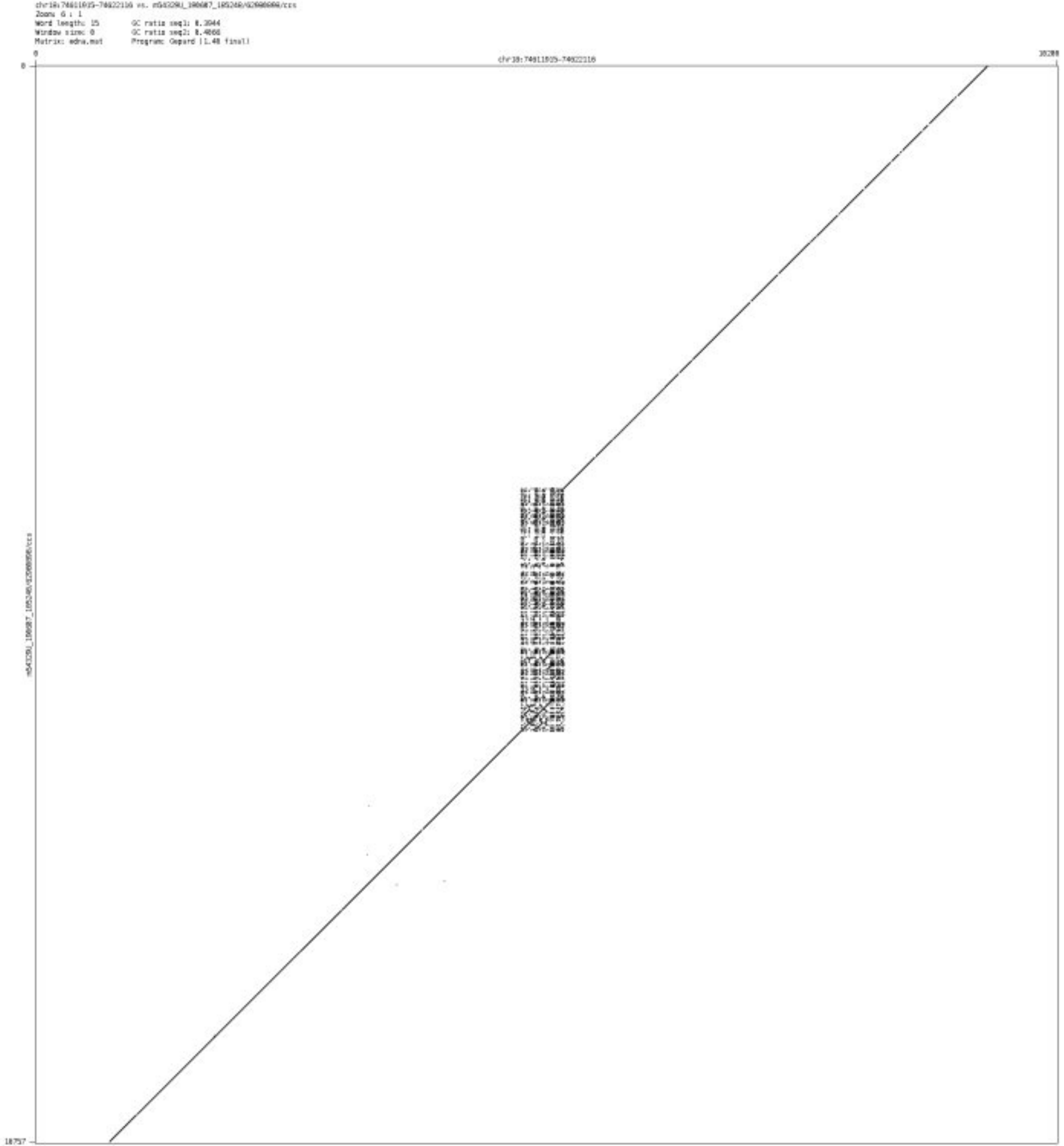

➢ chr18_76585300_76586173

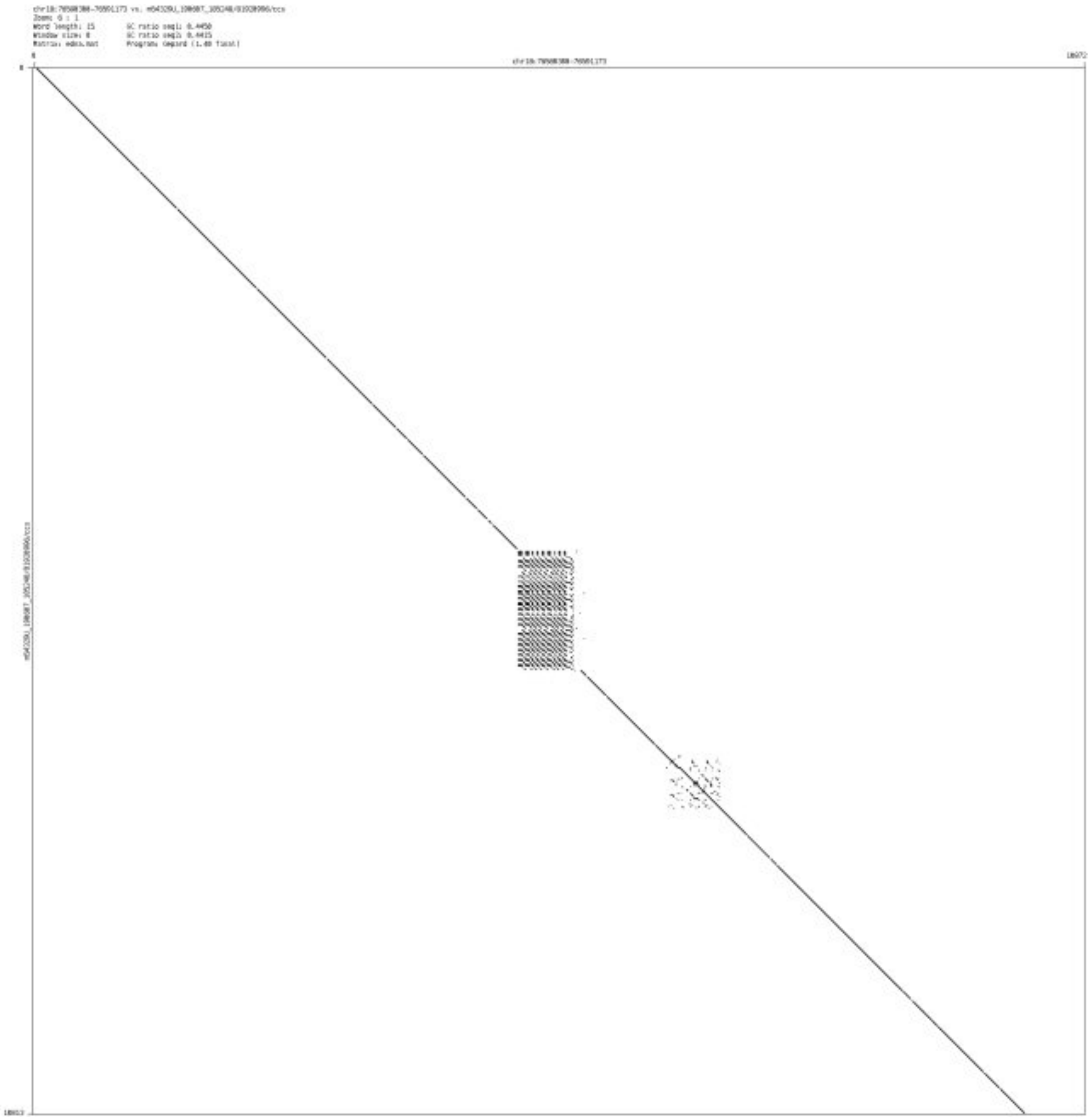

➢ chr18_78238512_78238581

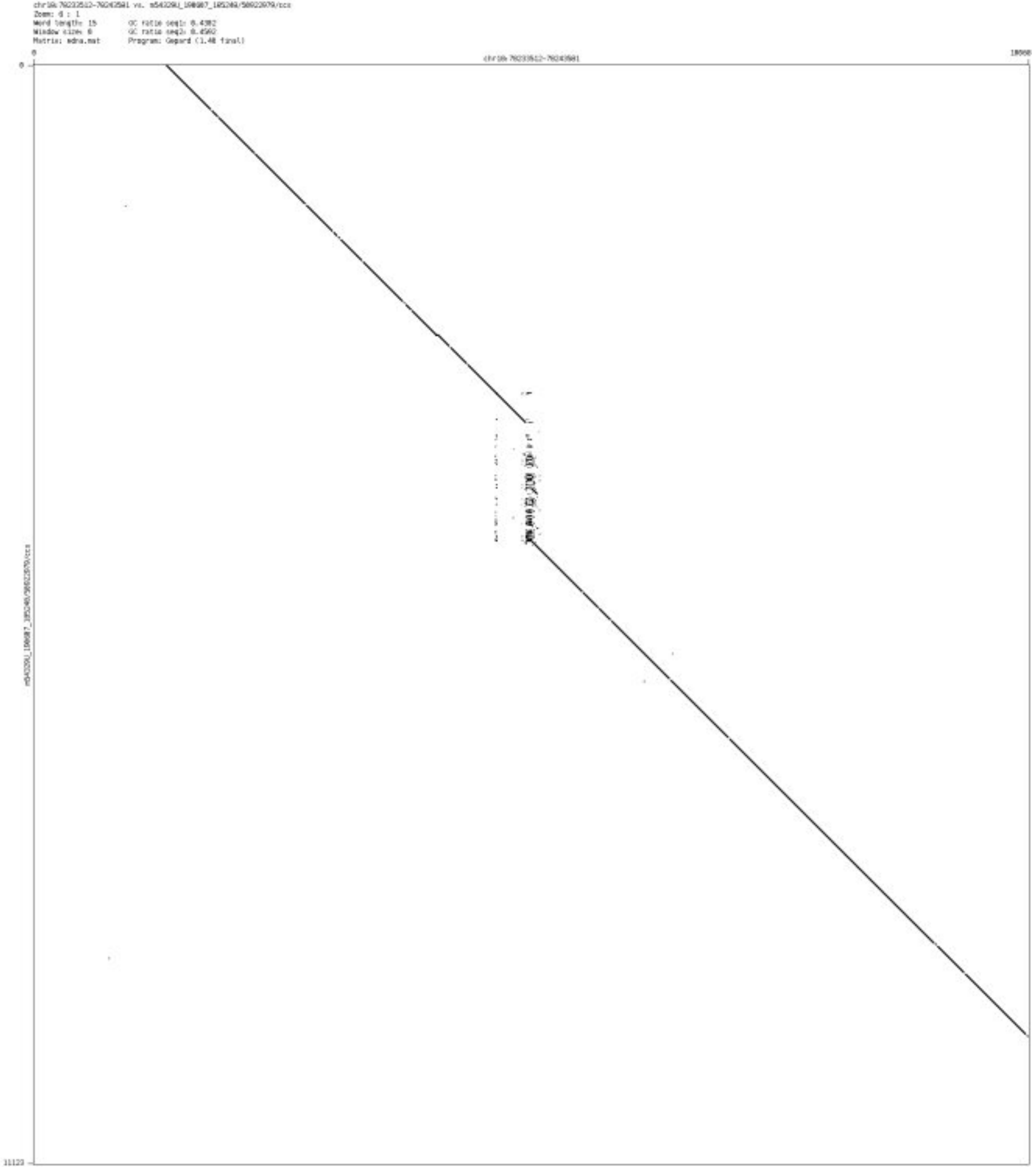

- chr18_79200704_79200757

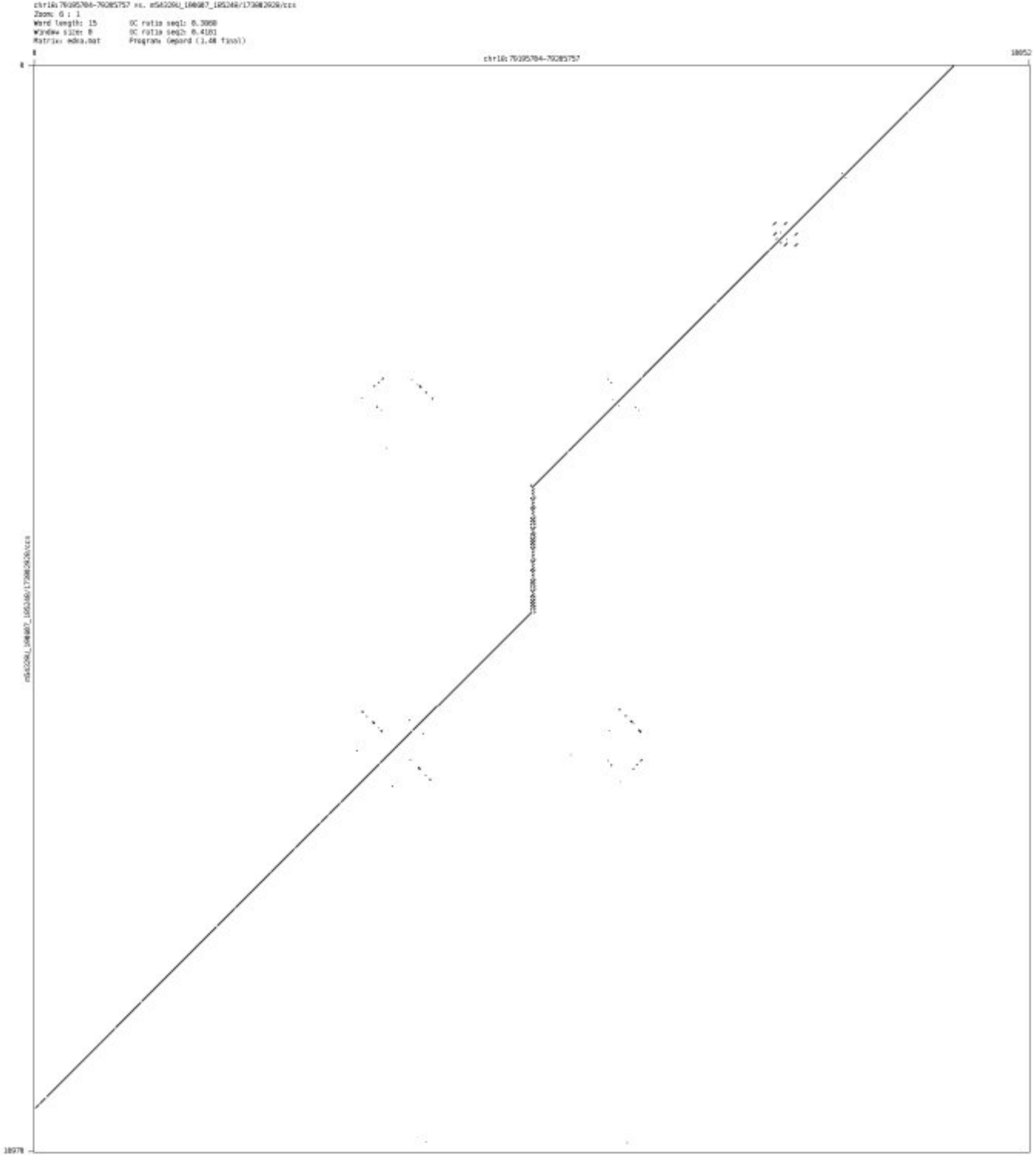

➢ chr19_350906_351902

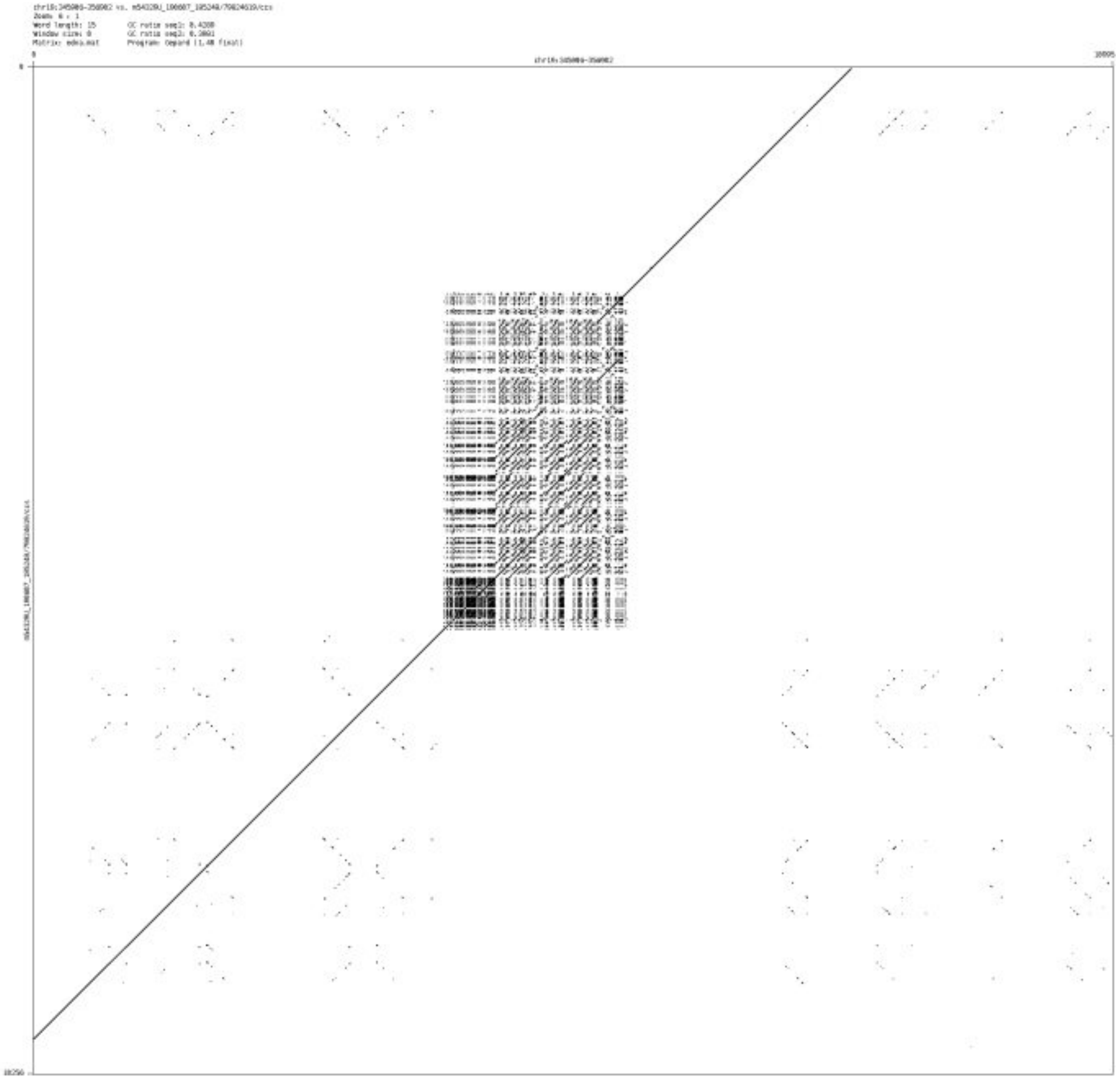

➢ chr19_835416_835497

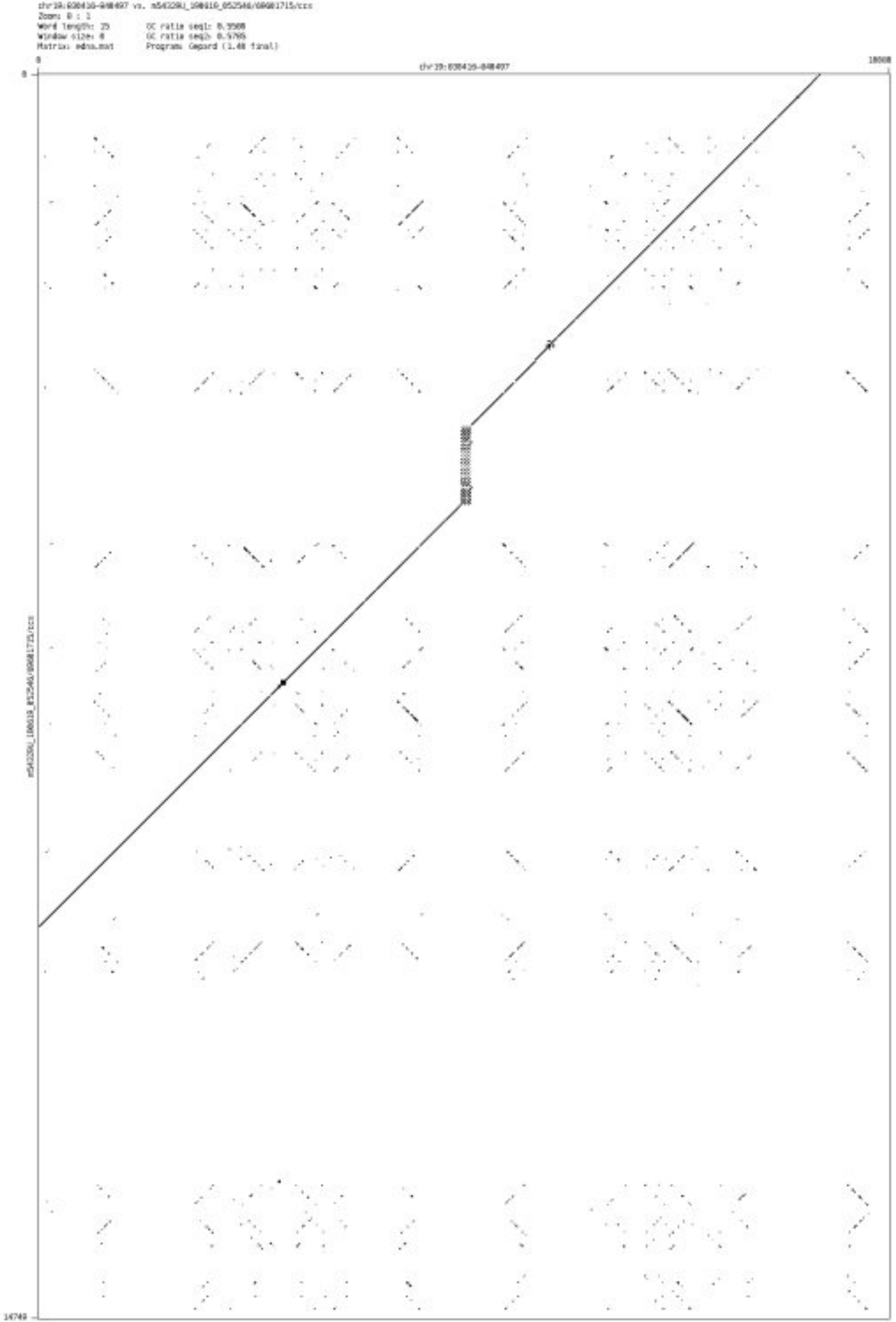

➢ chr19_1164394_1166232

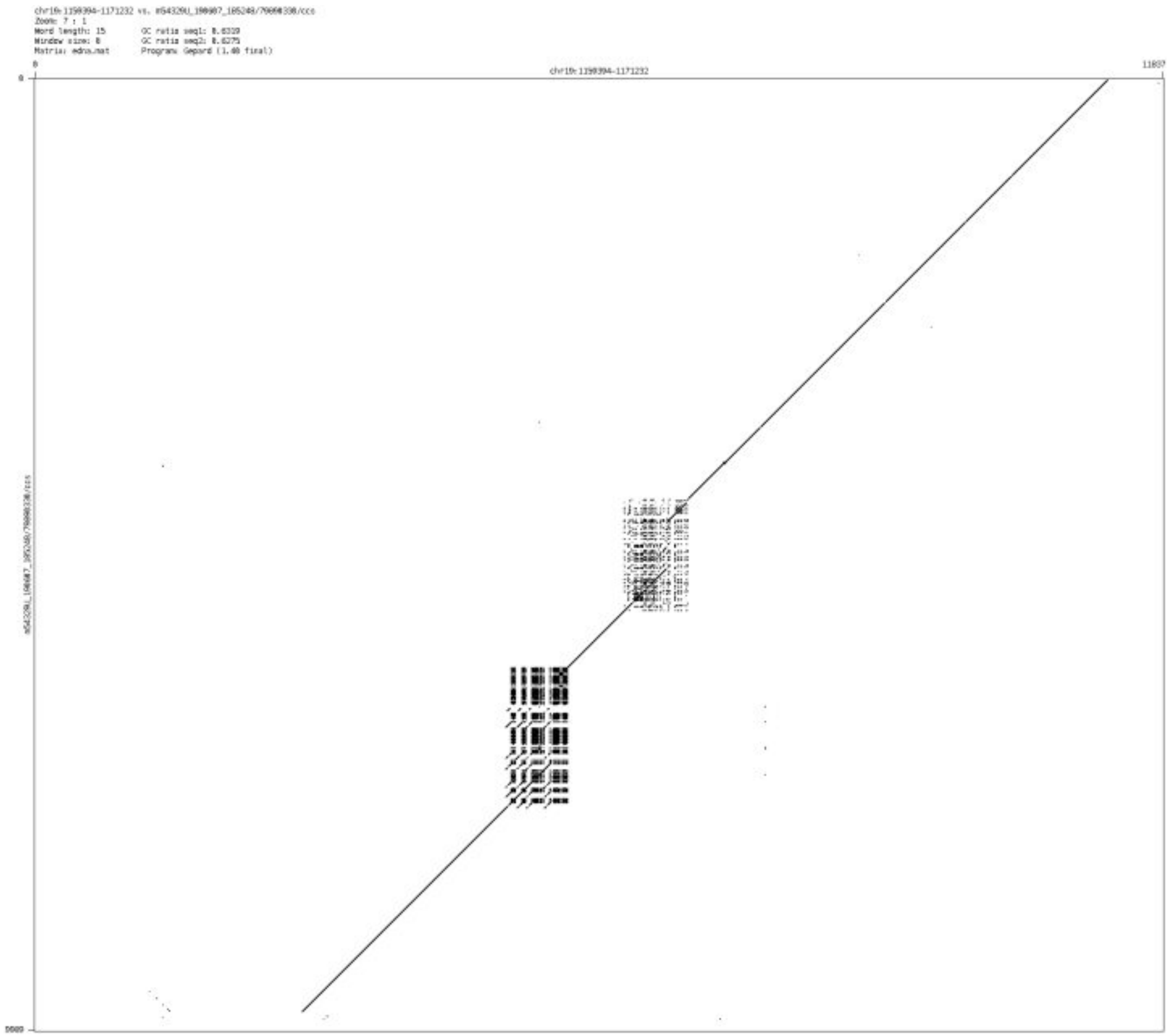

- chr19_3332439_3332577

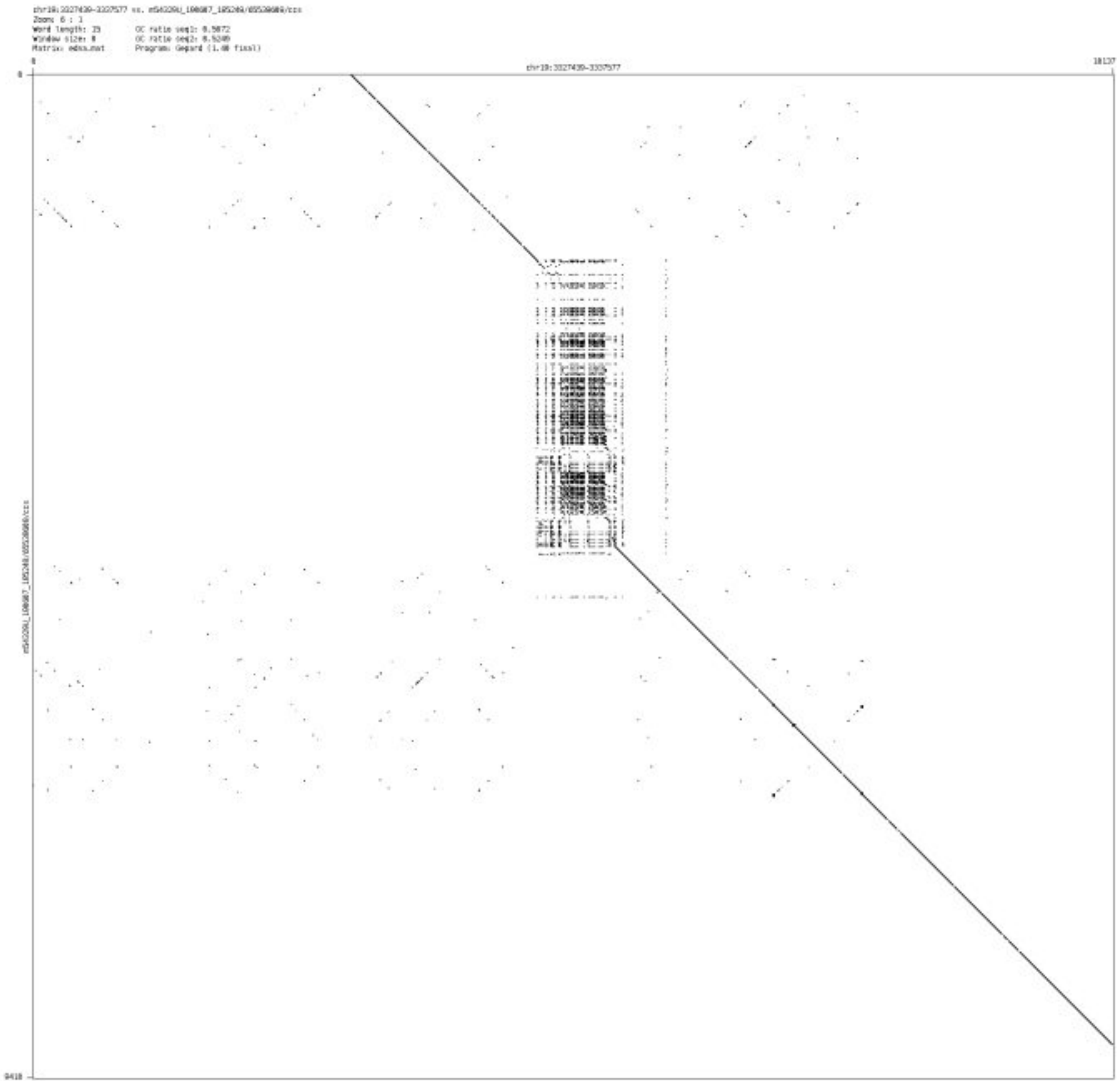

➢ chr19_6070065_6070188

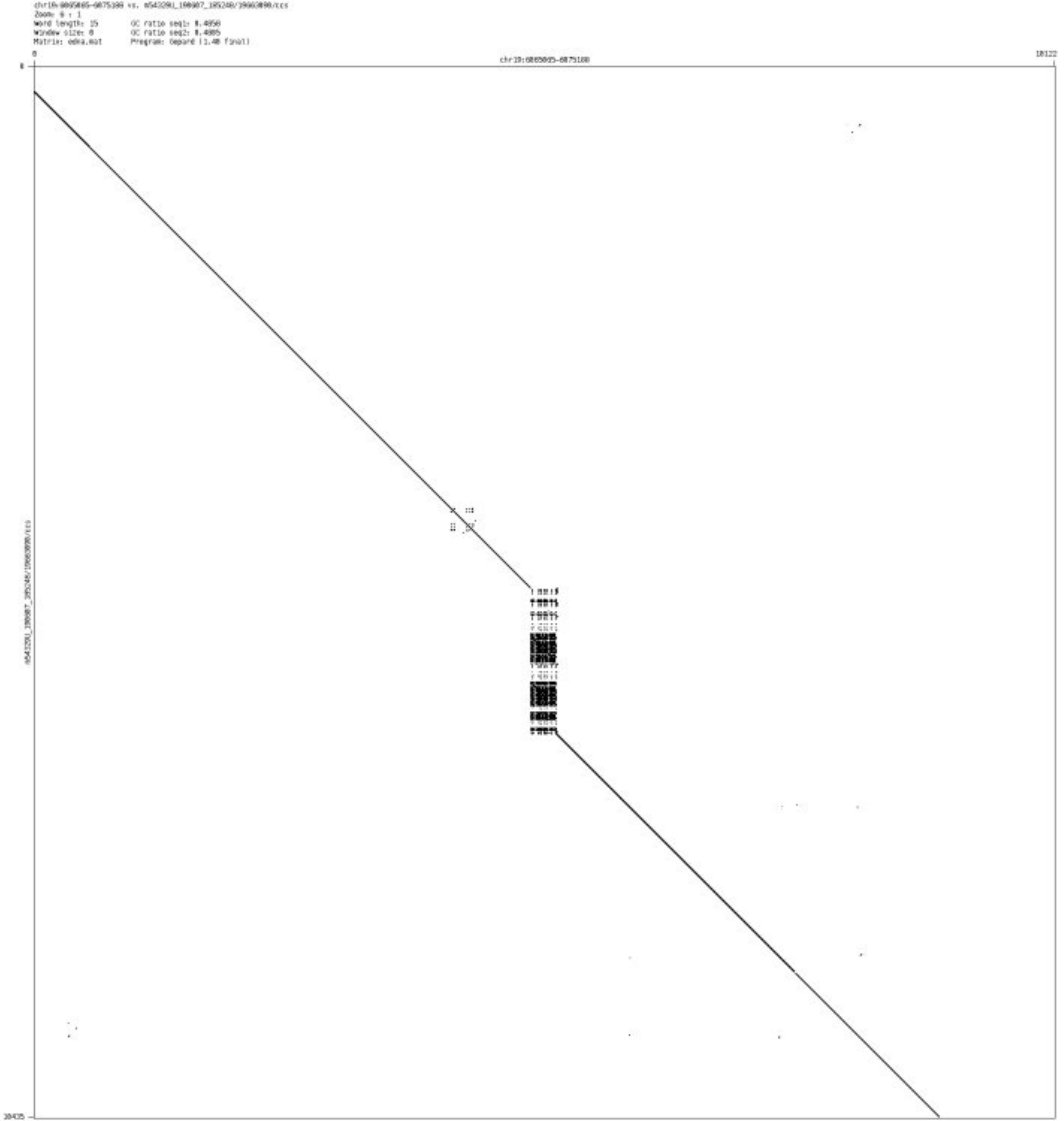

- chr19_7450388_7450641

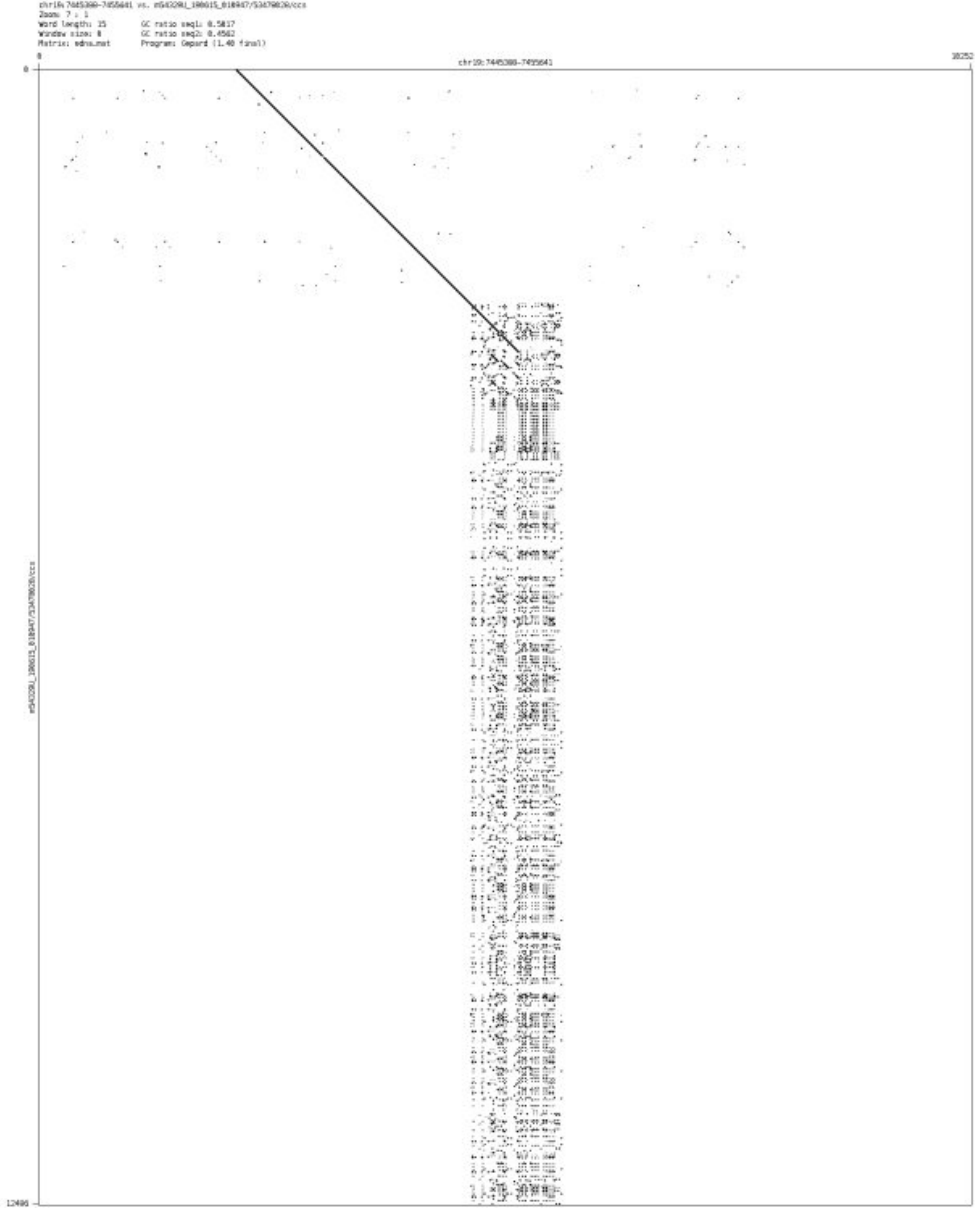

➢ chr19_28888385_28888515

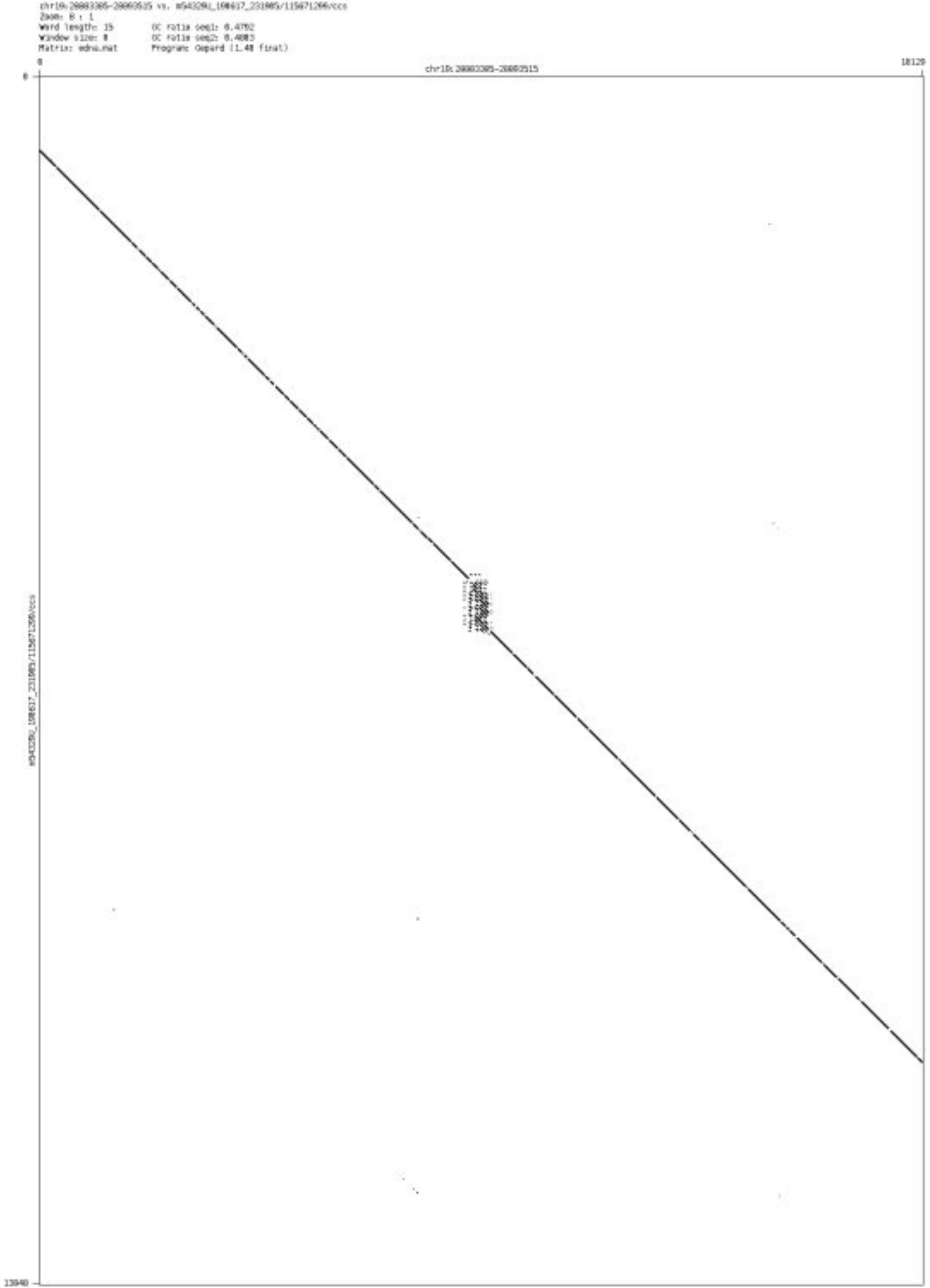

- chr19_35105945_35106010

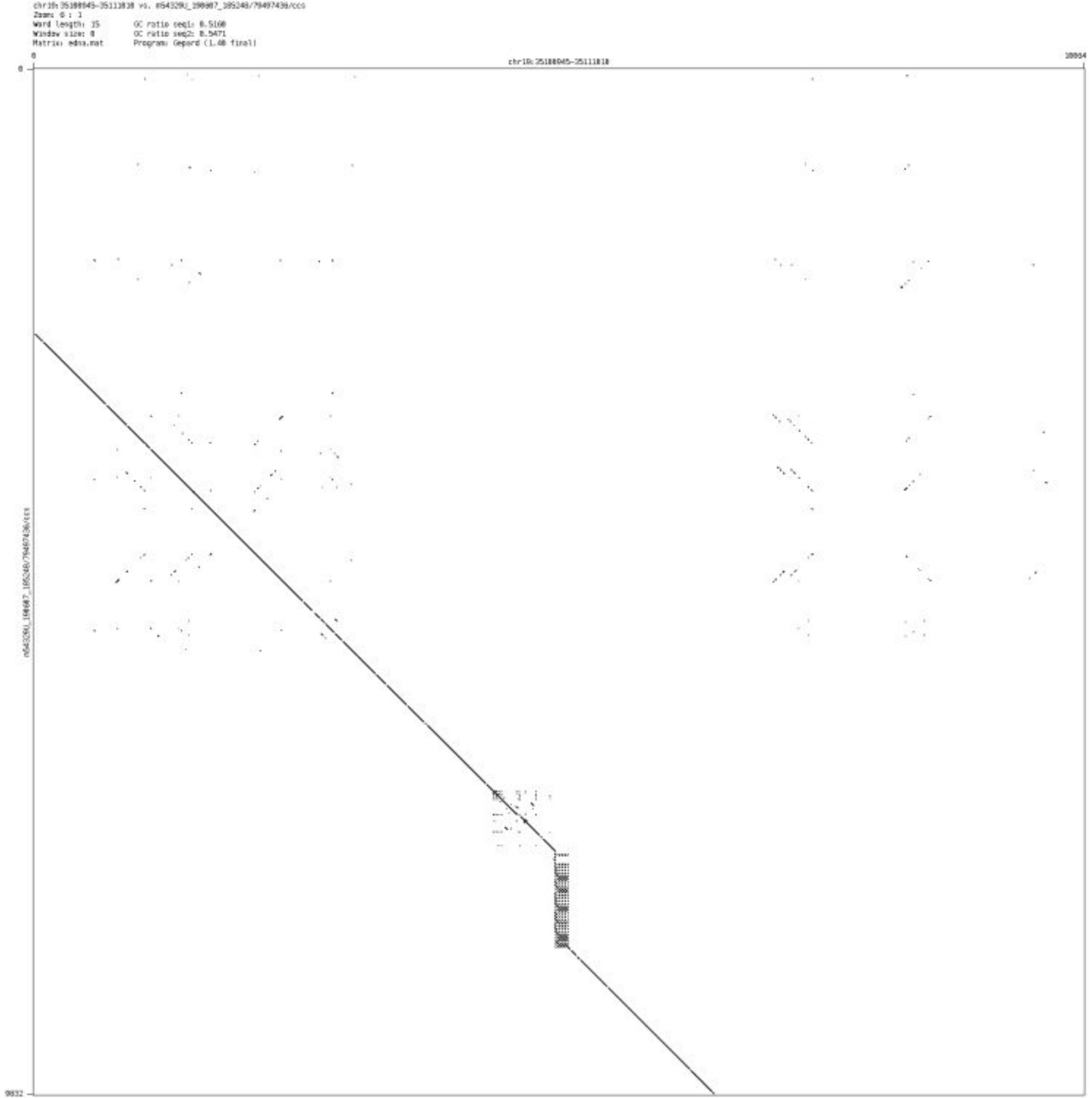

- chr19_54109797_54109852

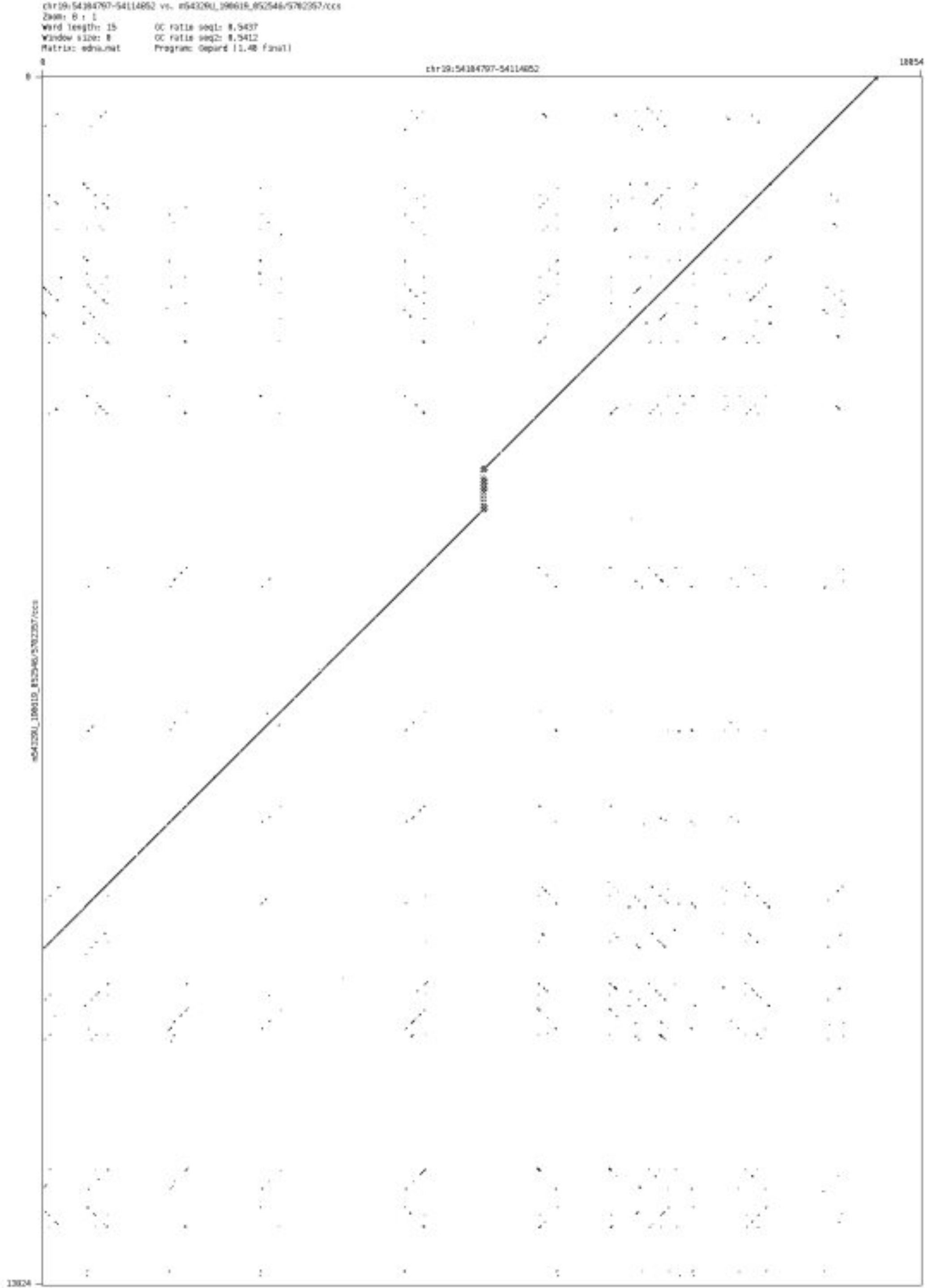

- chr19_54133743_54133869

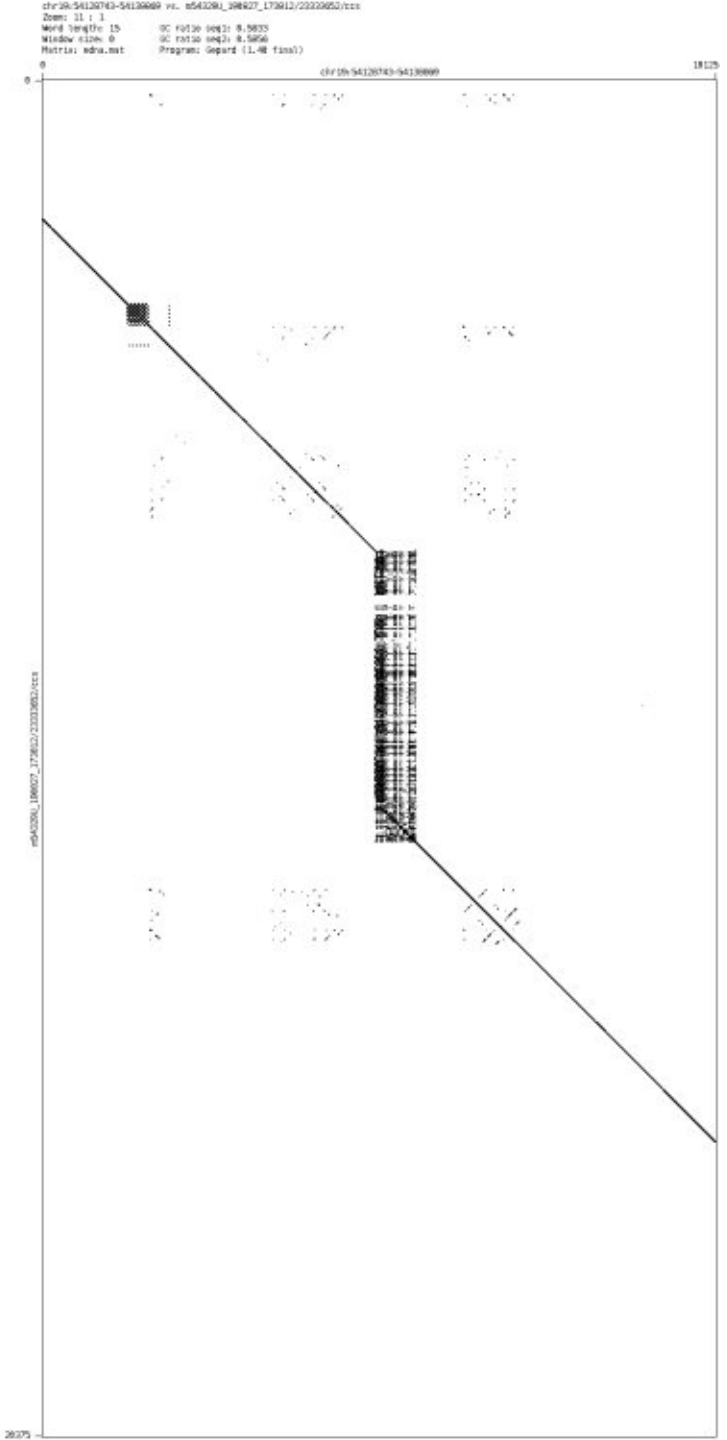

- ➢ chr19_55670439_55670523

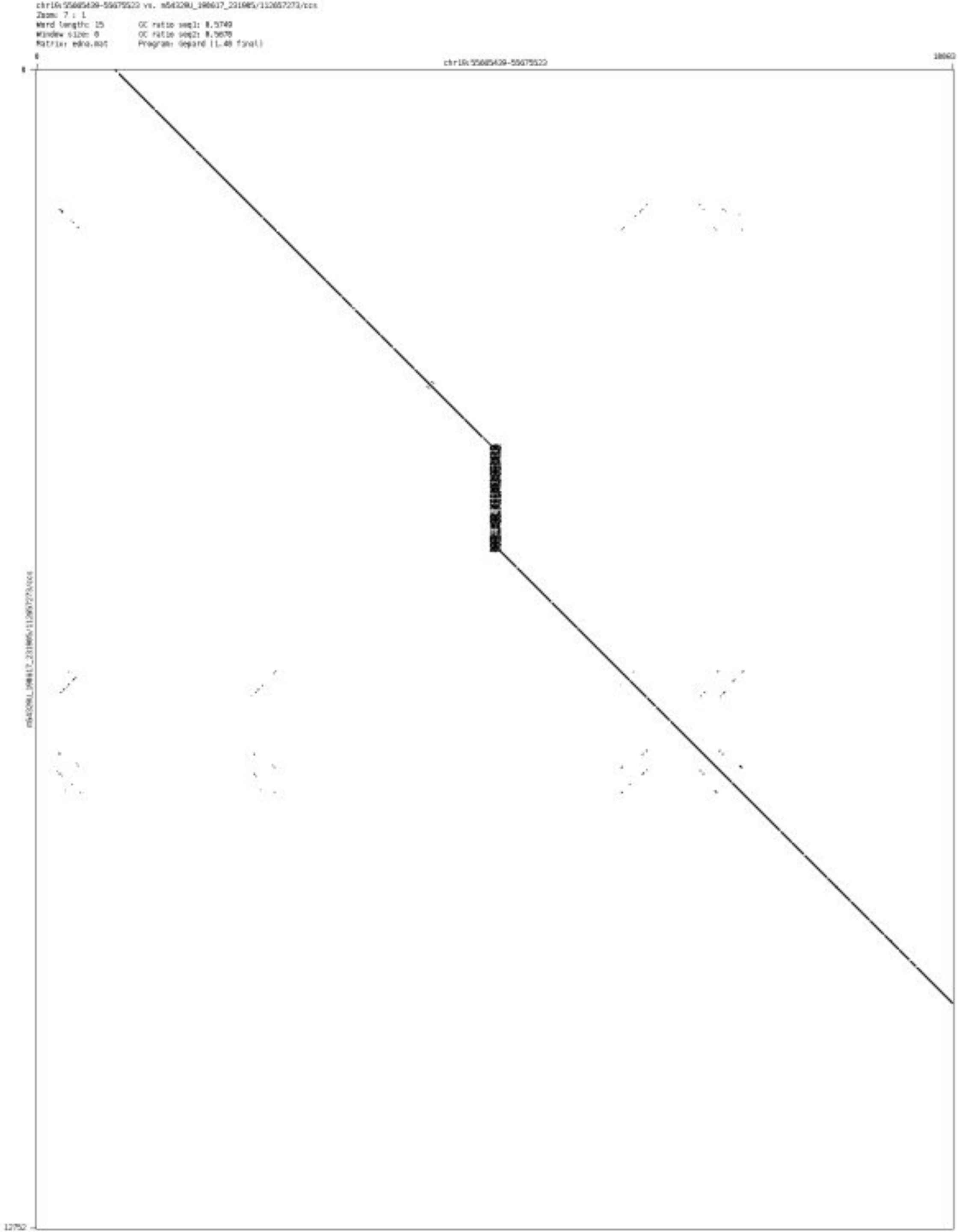

➢ chr19_56942922_56942975

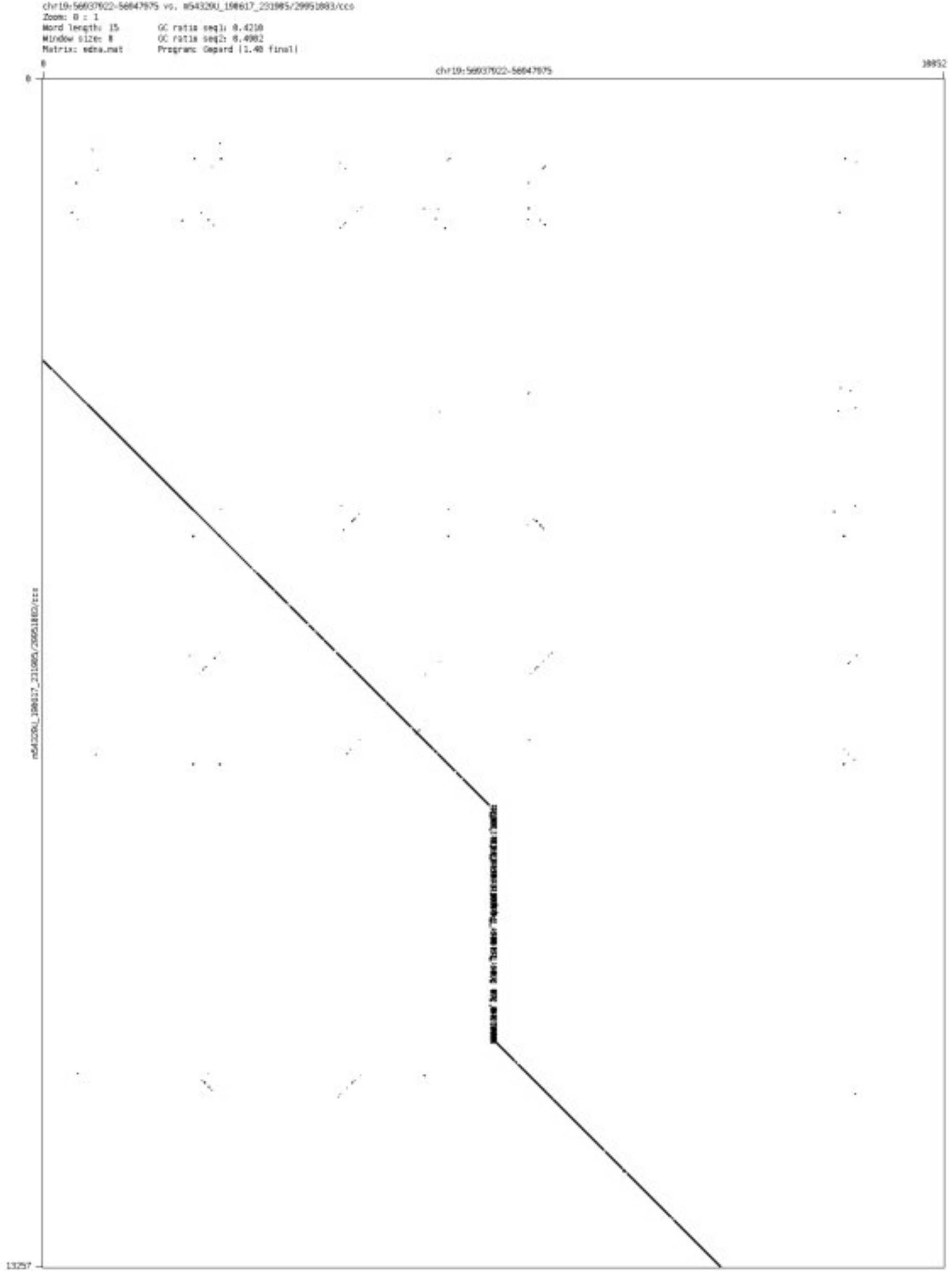

➢ chr20_13934330_13934452

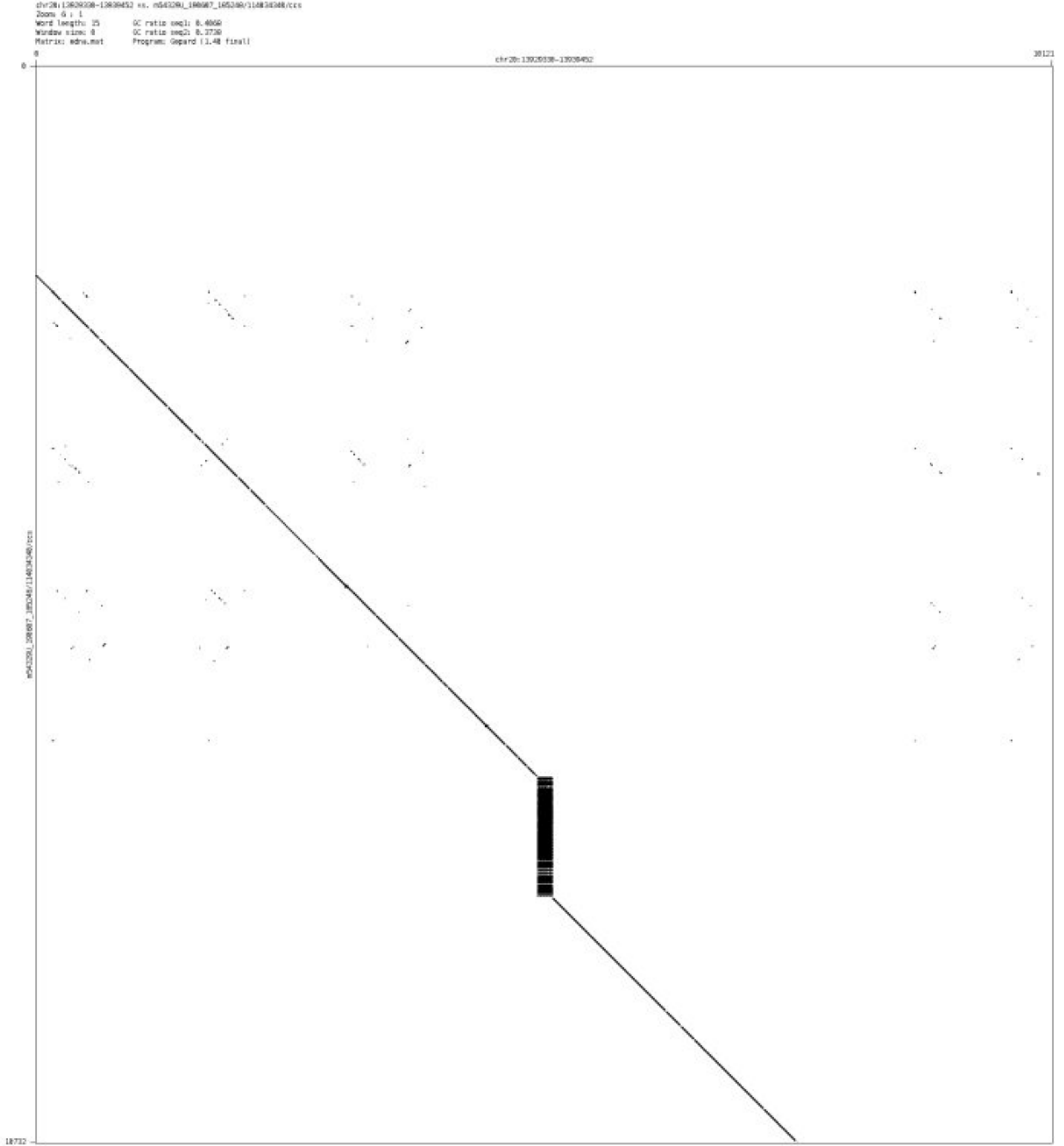

➢ chr20_19924430_19924495

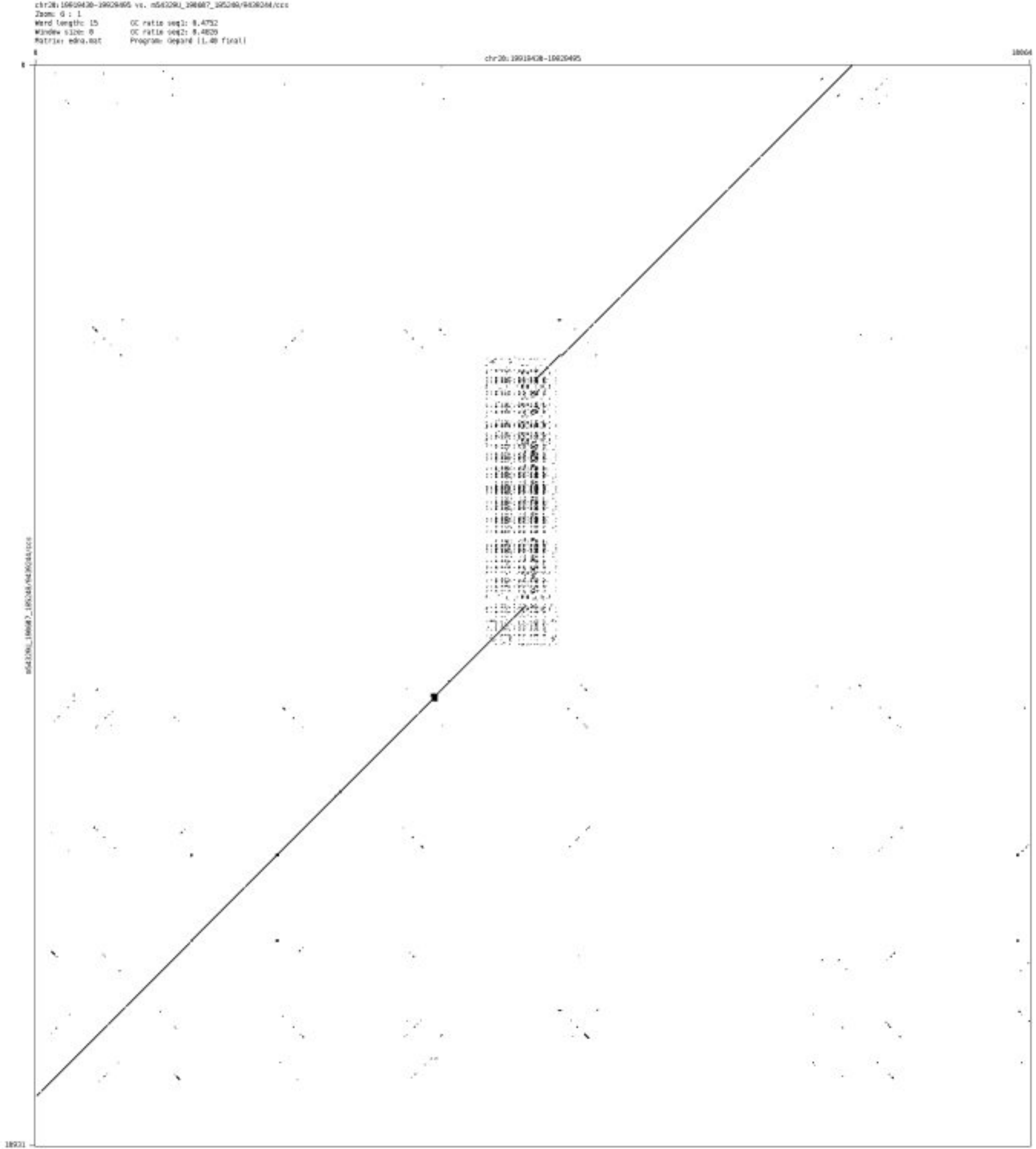

- ➢ chr20_20337356_20337505

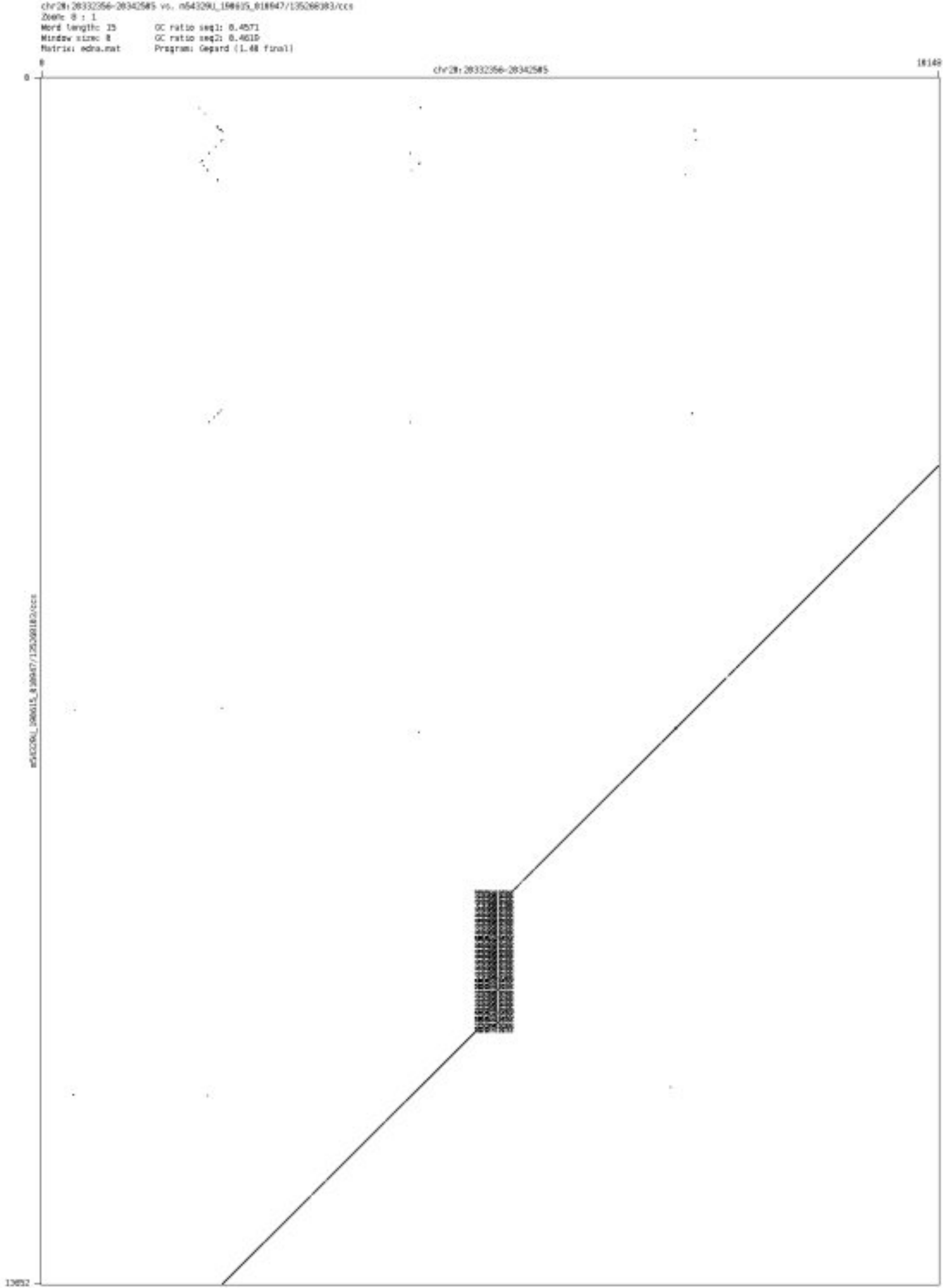

➢ chr20_20356363_20357230

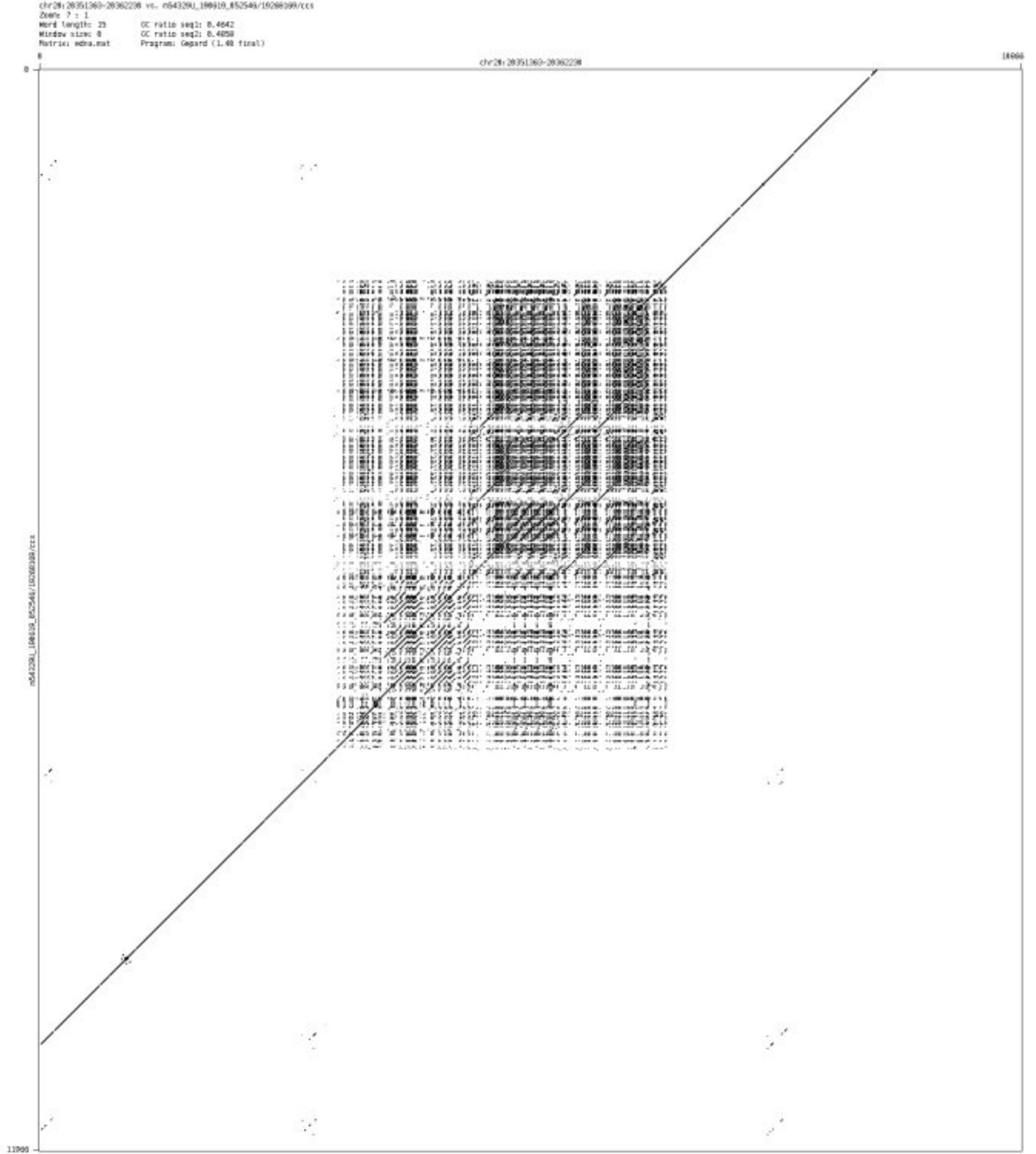

- chr20_38463938_38464116

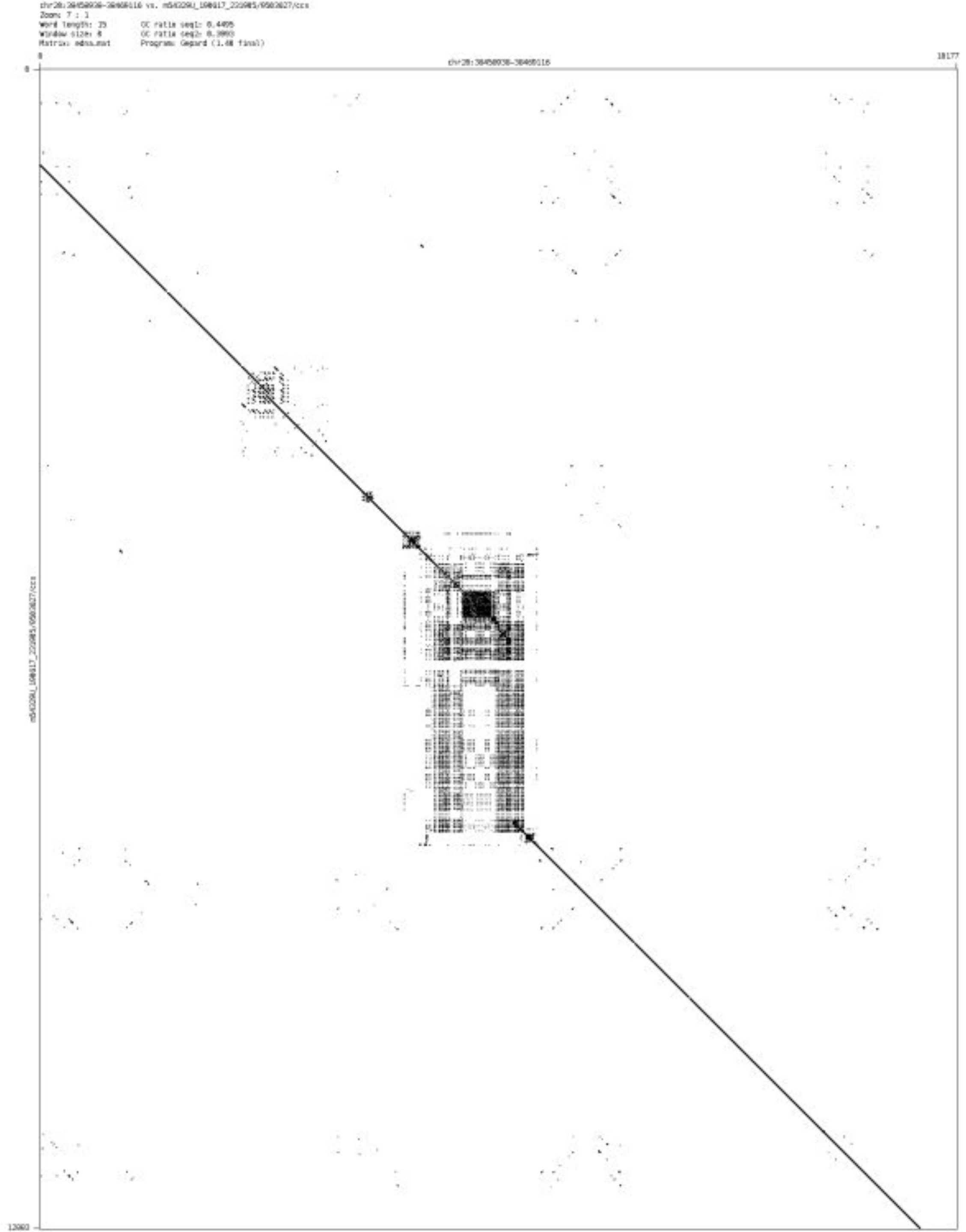

- chr20_53204104_53204269

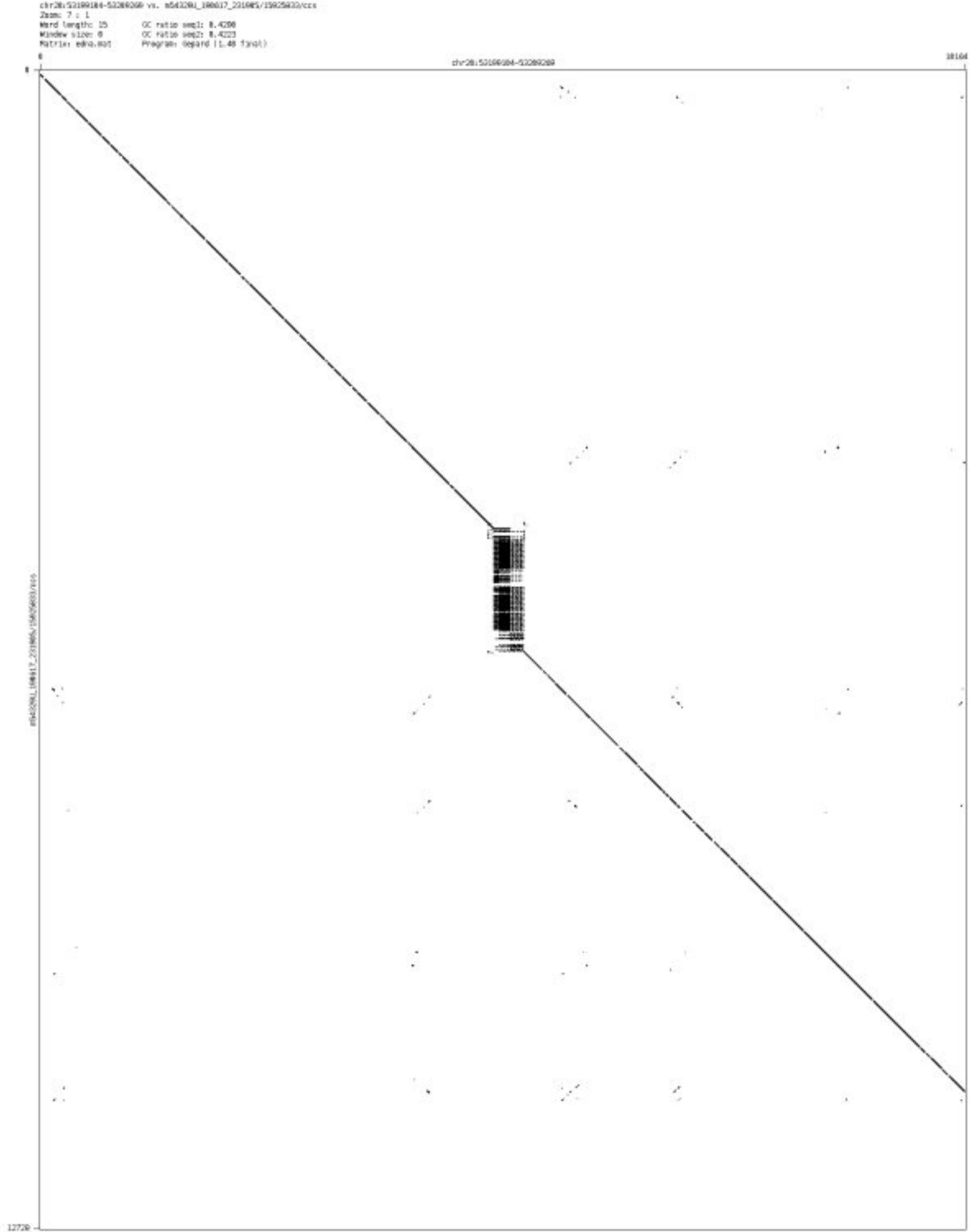

- chr20_53837740_53837850

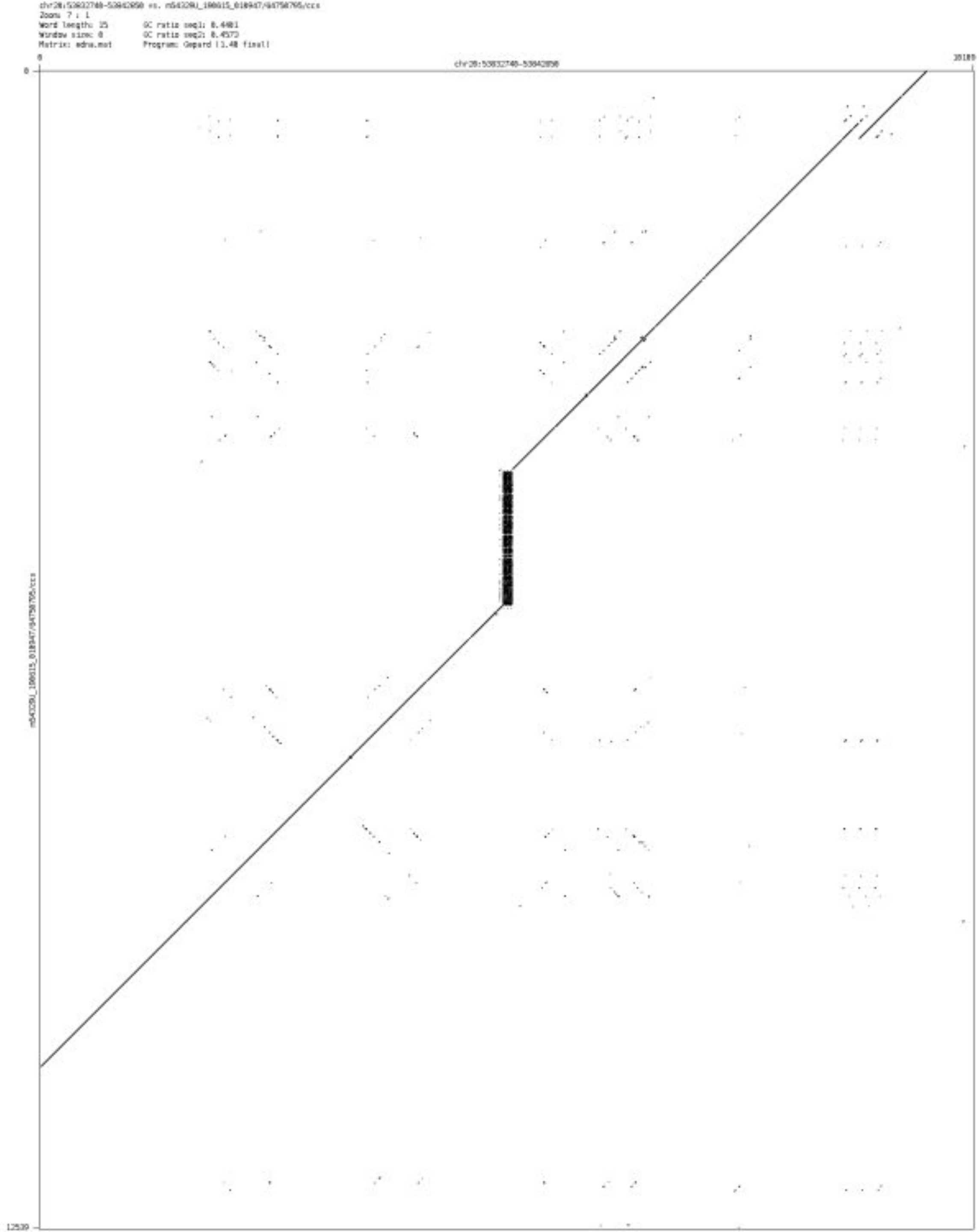

- ➢ chr20_61202088_61202234

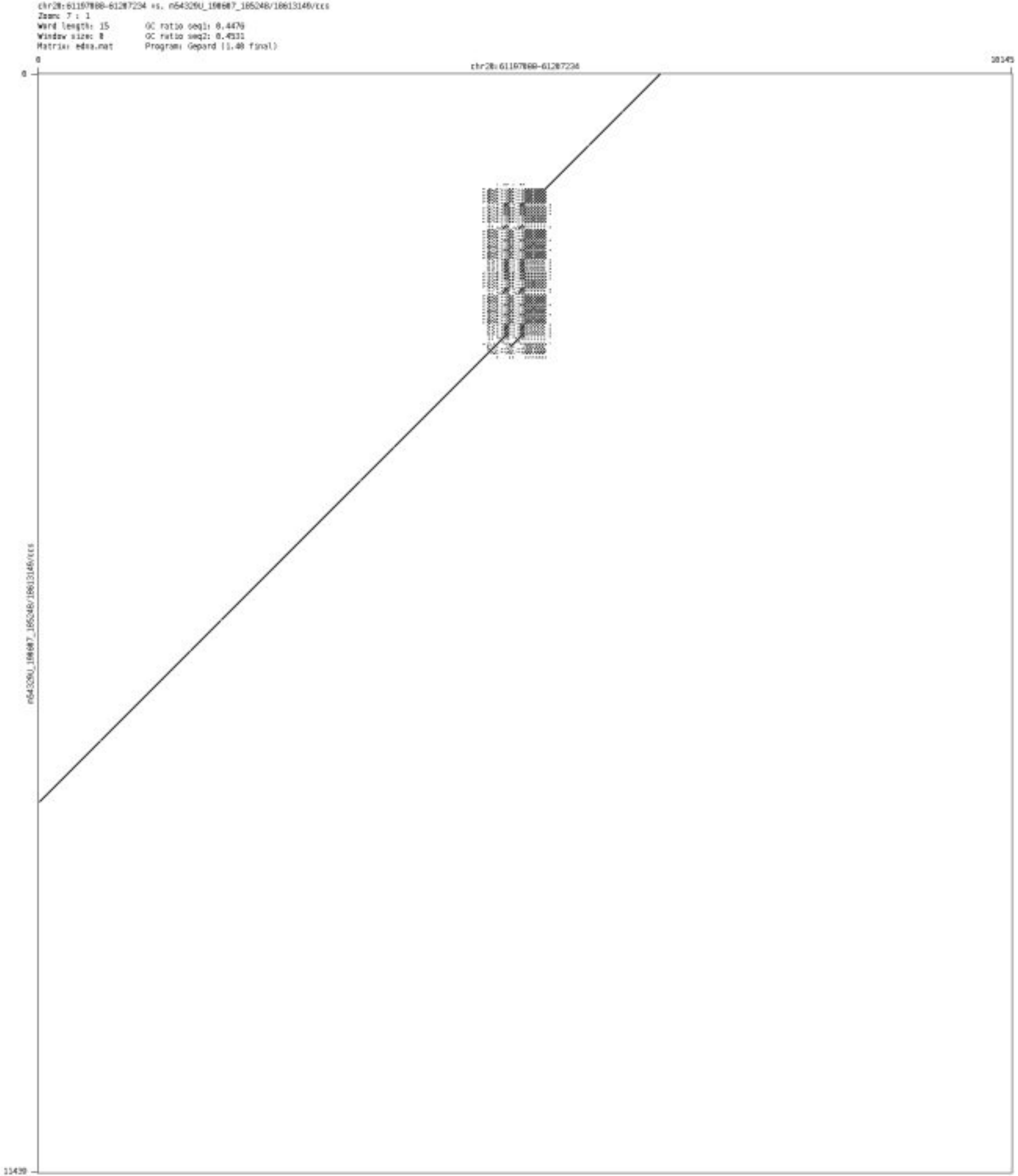

➢ chr20_61375824_61375997

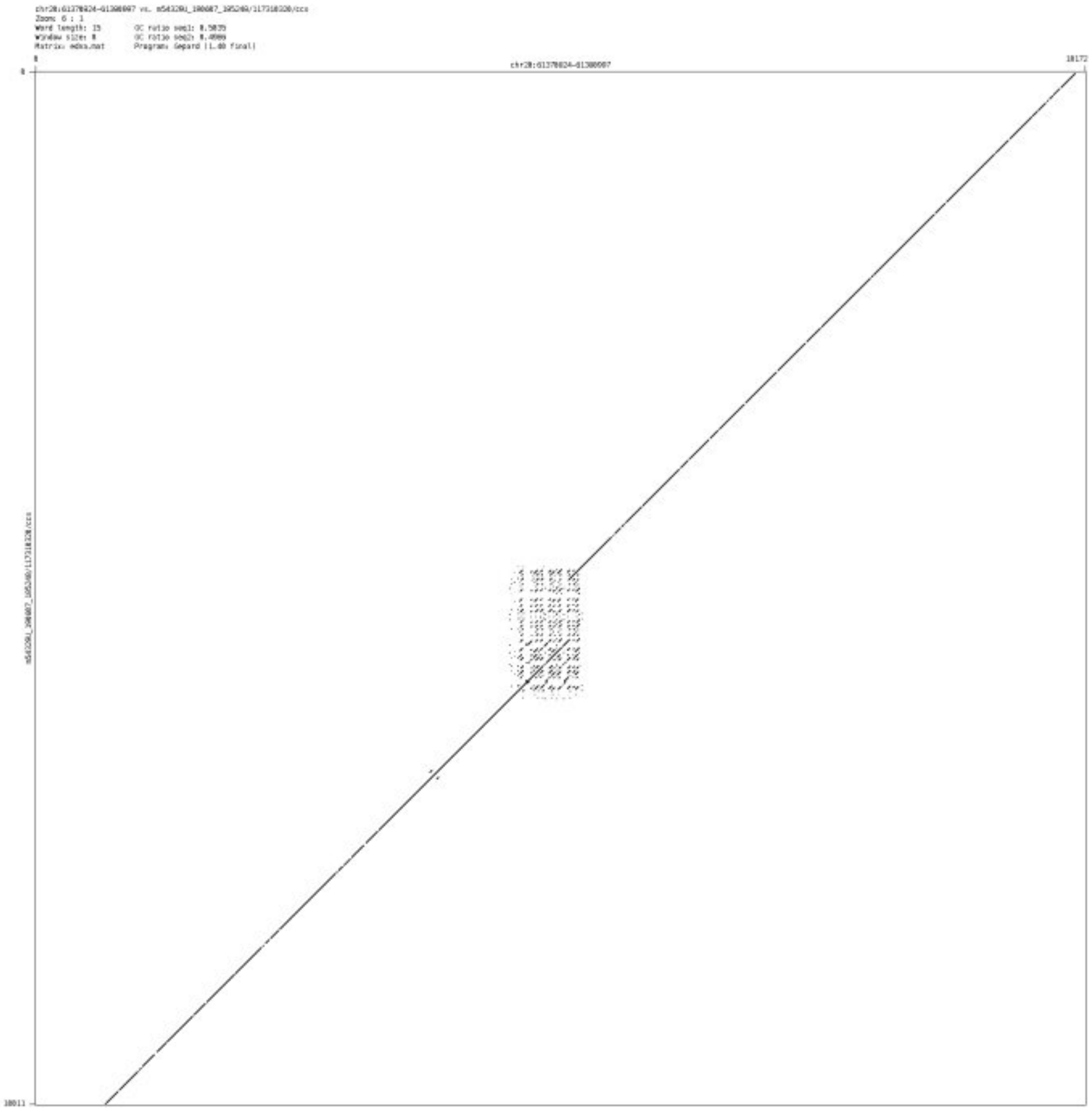

#### ➢ chr20_63693538_63693676

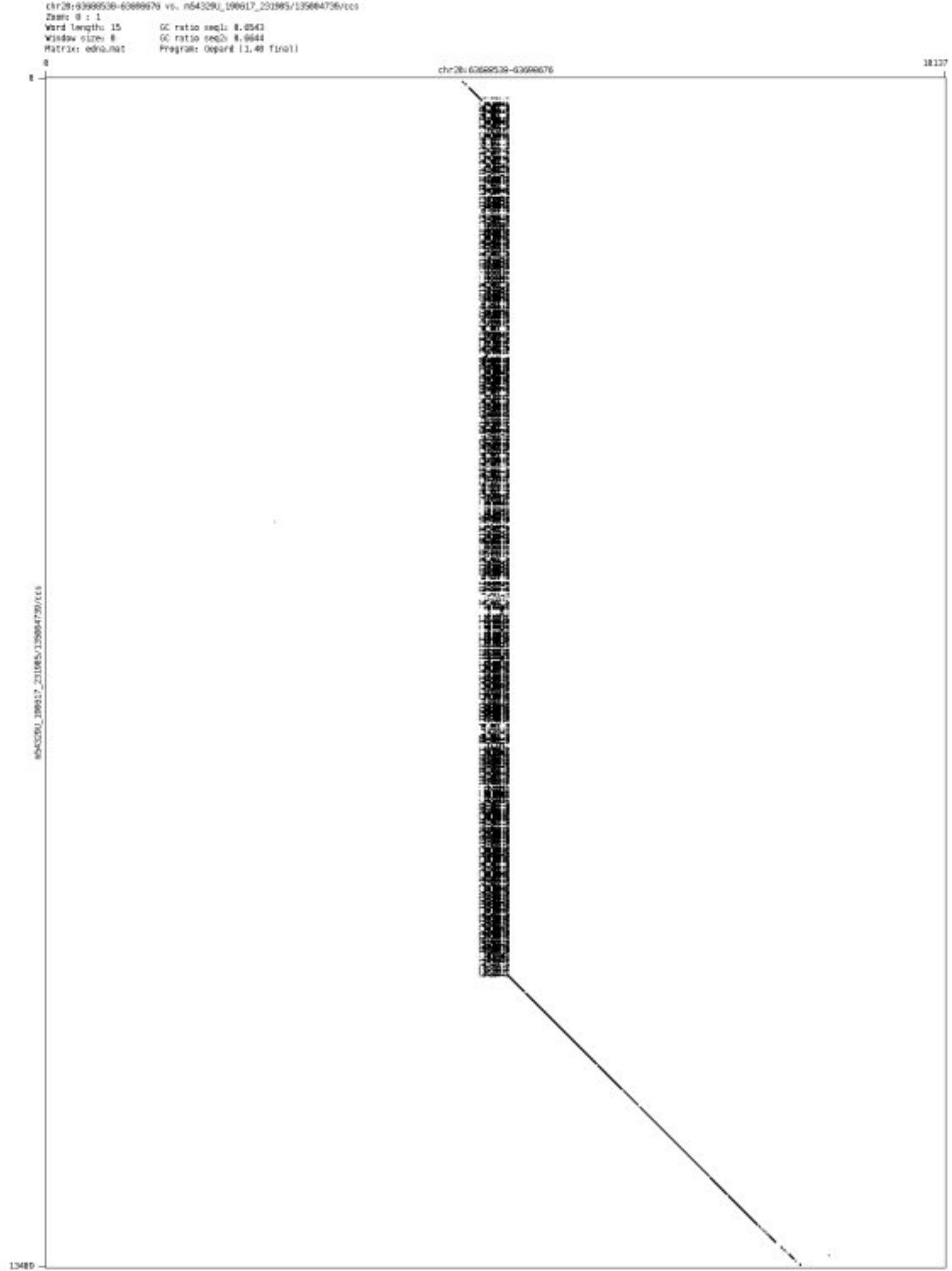

- chr20_64088663_64088895

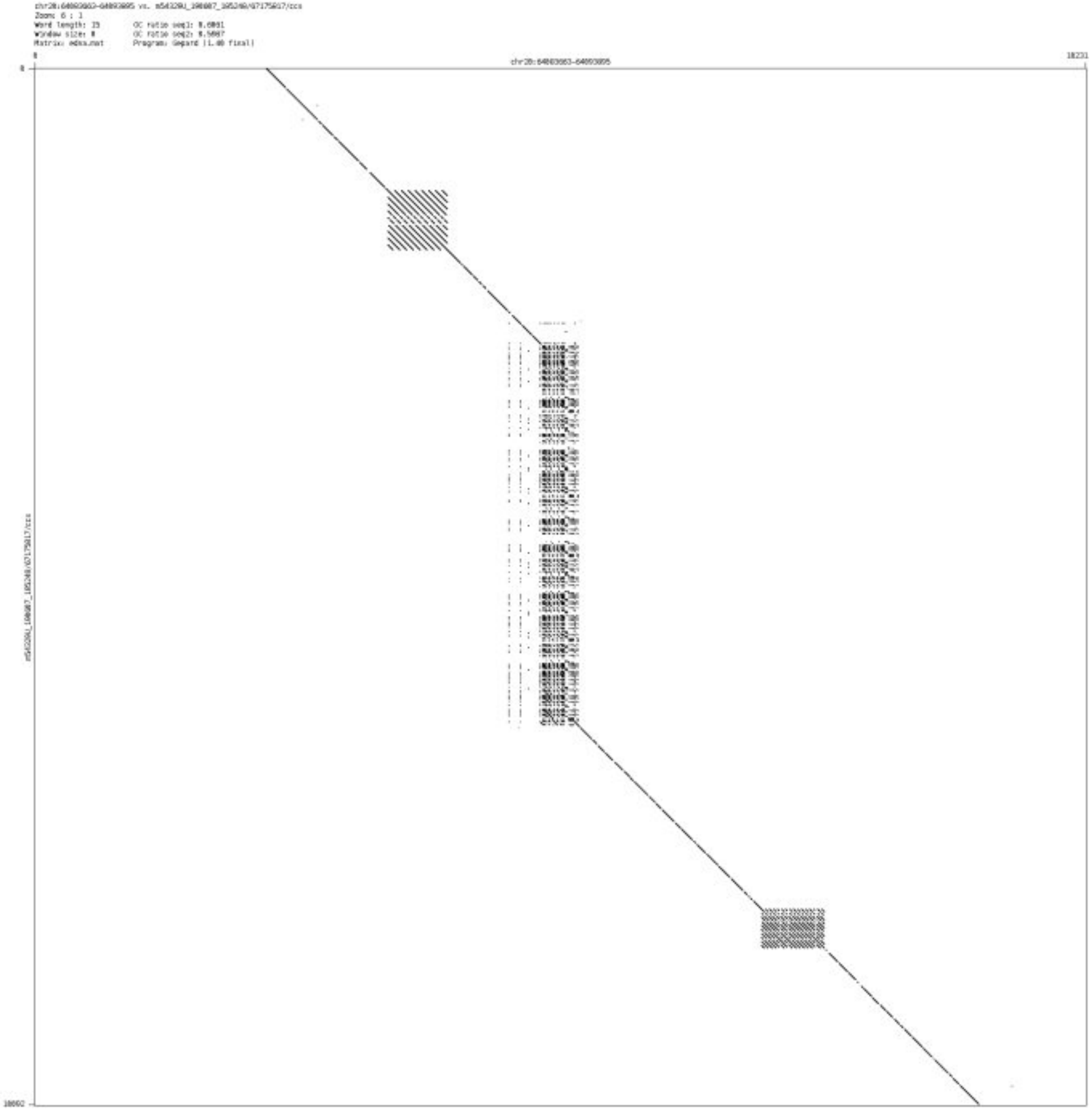

➢ chr20_64096926_64097053

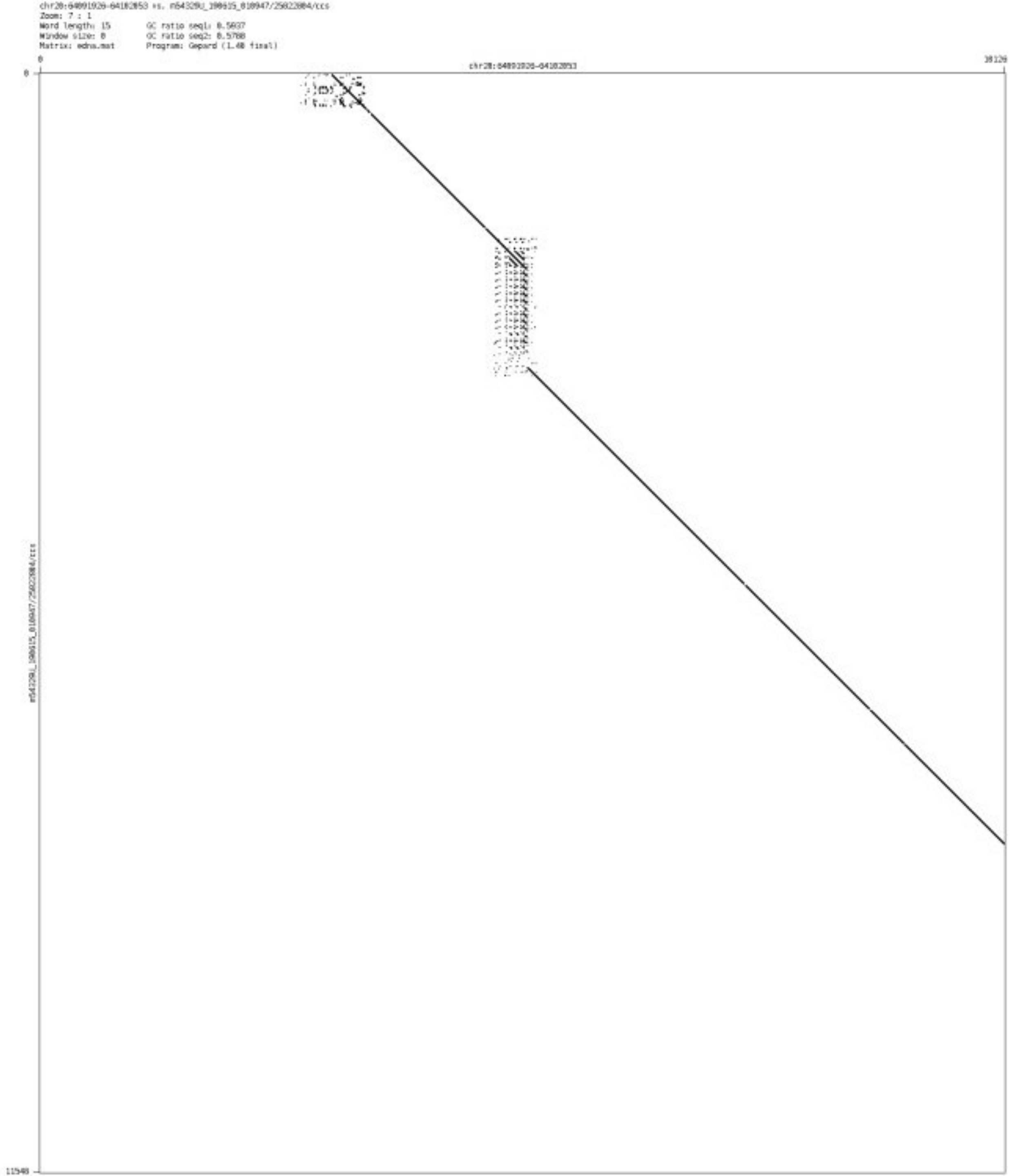

➢ chr21_27627113_27627278

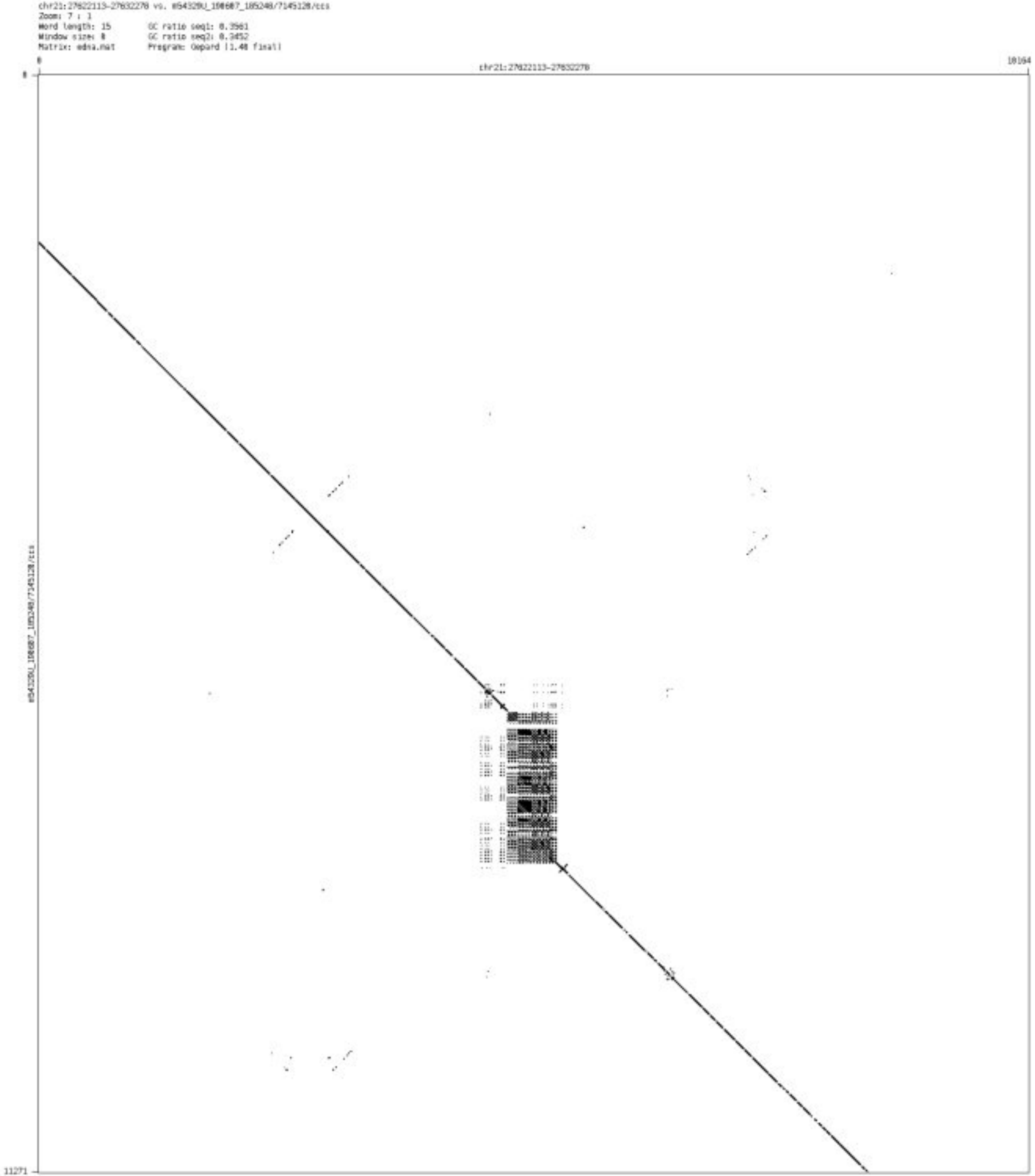

➢ chr21_31784083_31785382

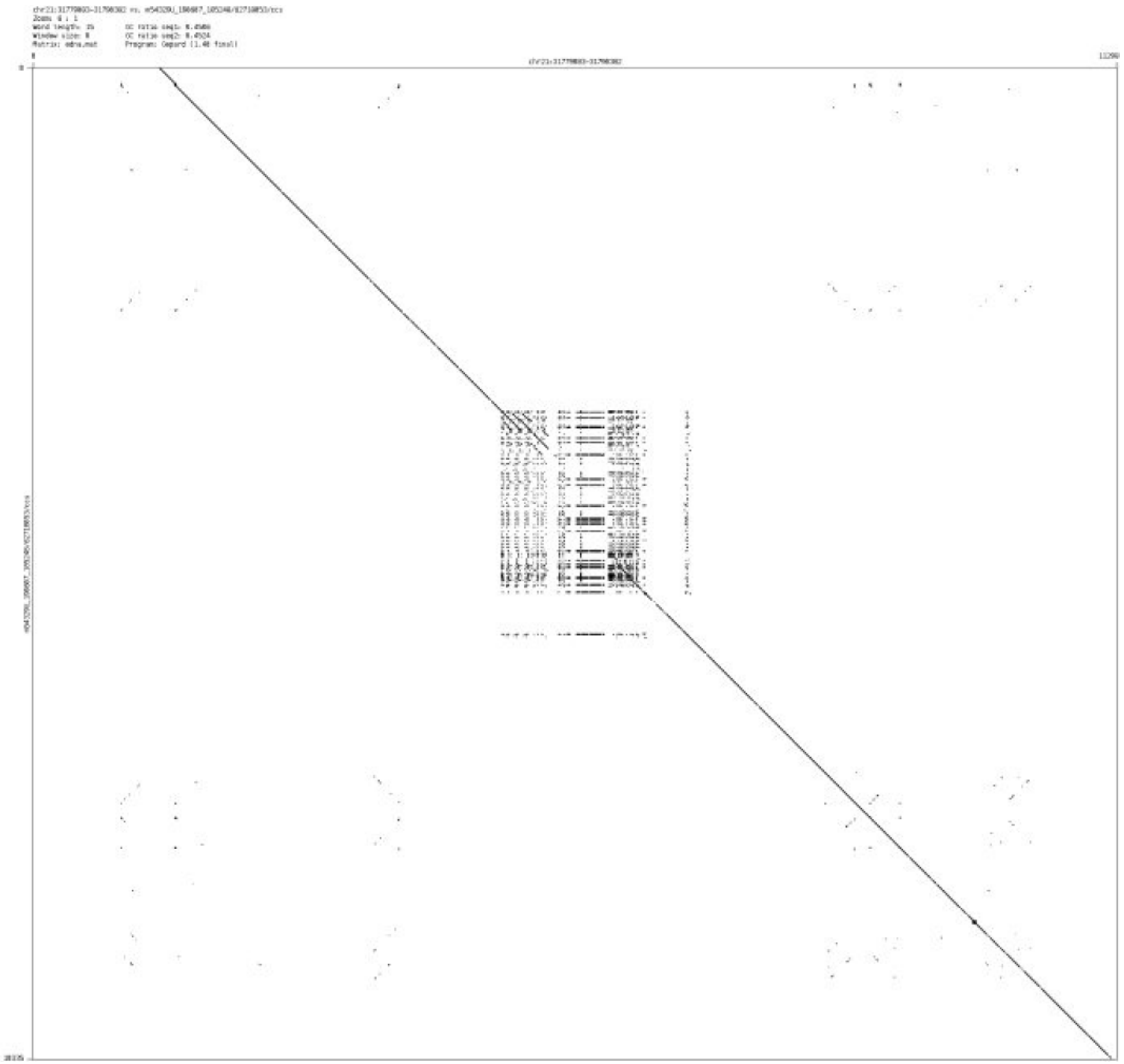

- chr21_36497141_36497270

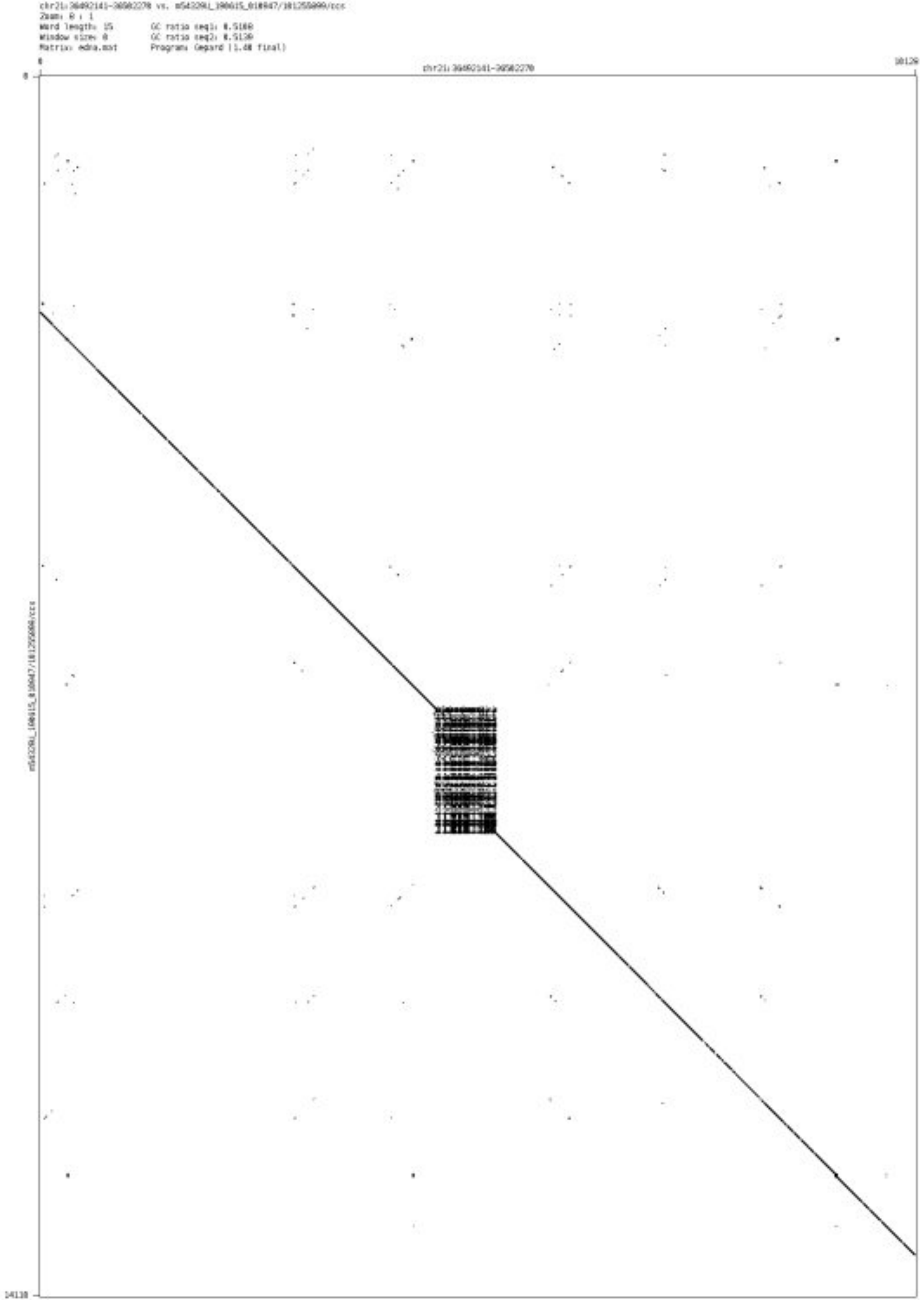

- chr21_40348162_40348388

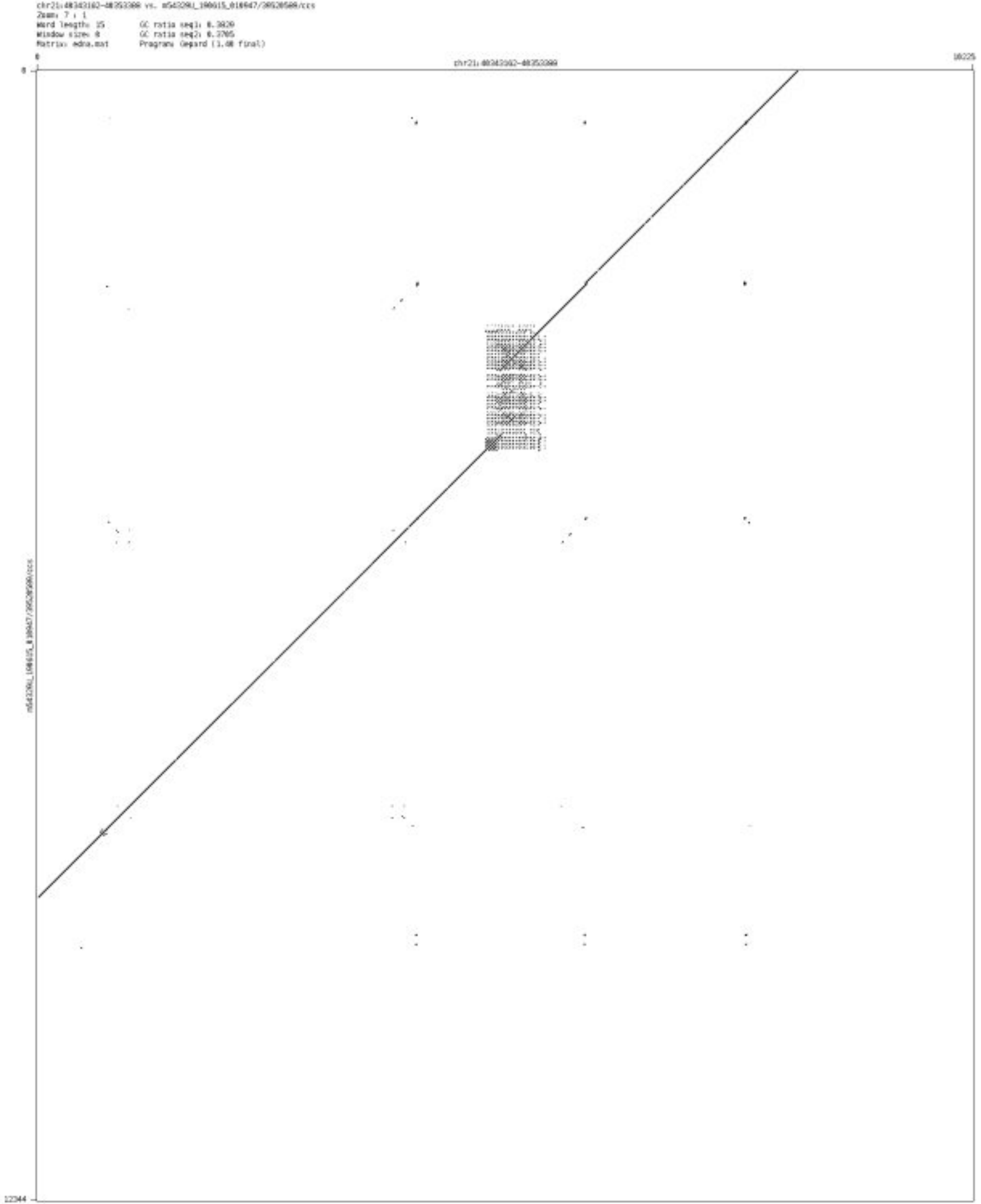

- chr21_43262539_43262718

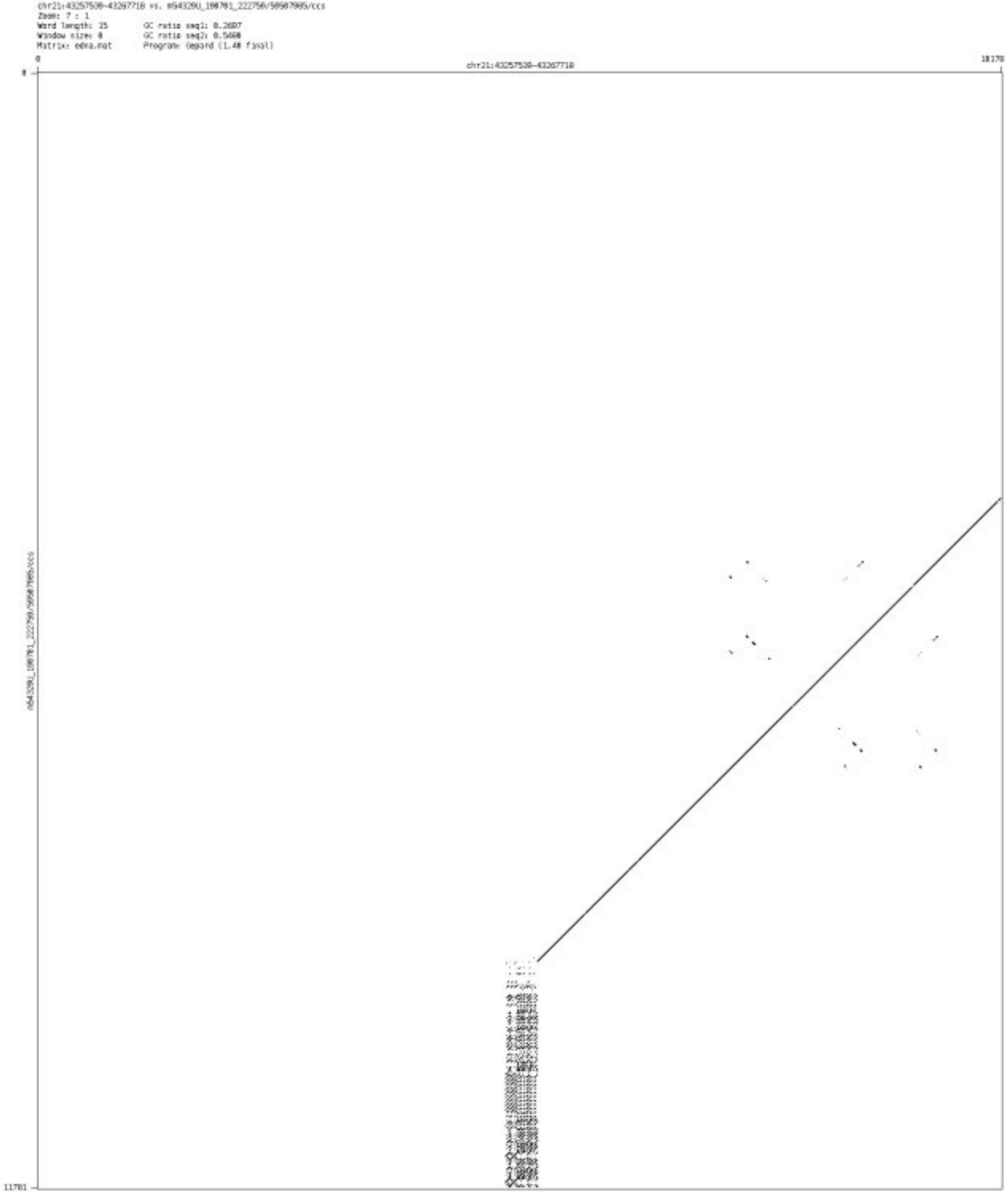

- chr21_46399240_46399382

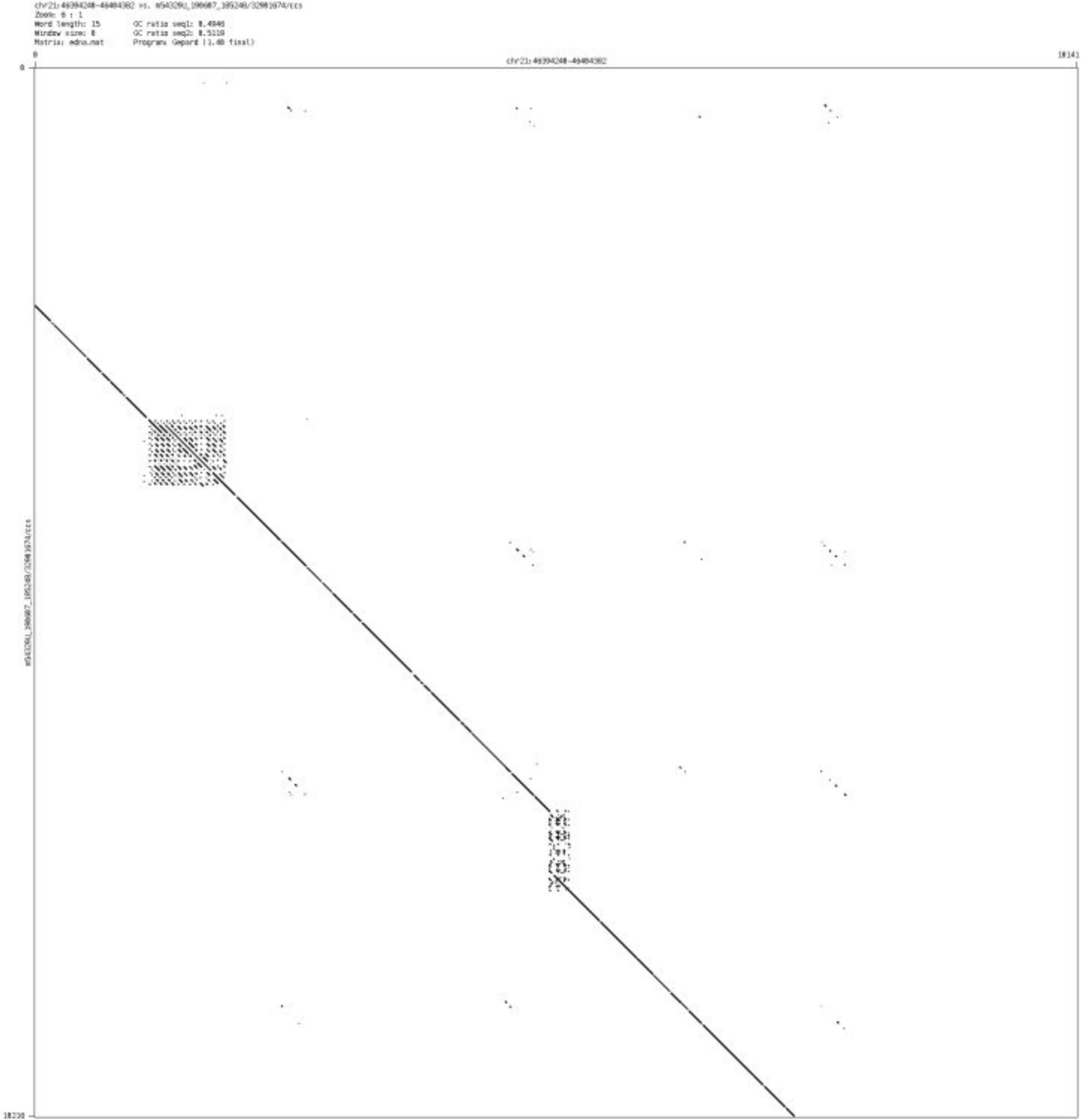

- chr21_46699871_46699959

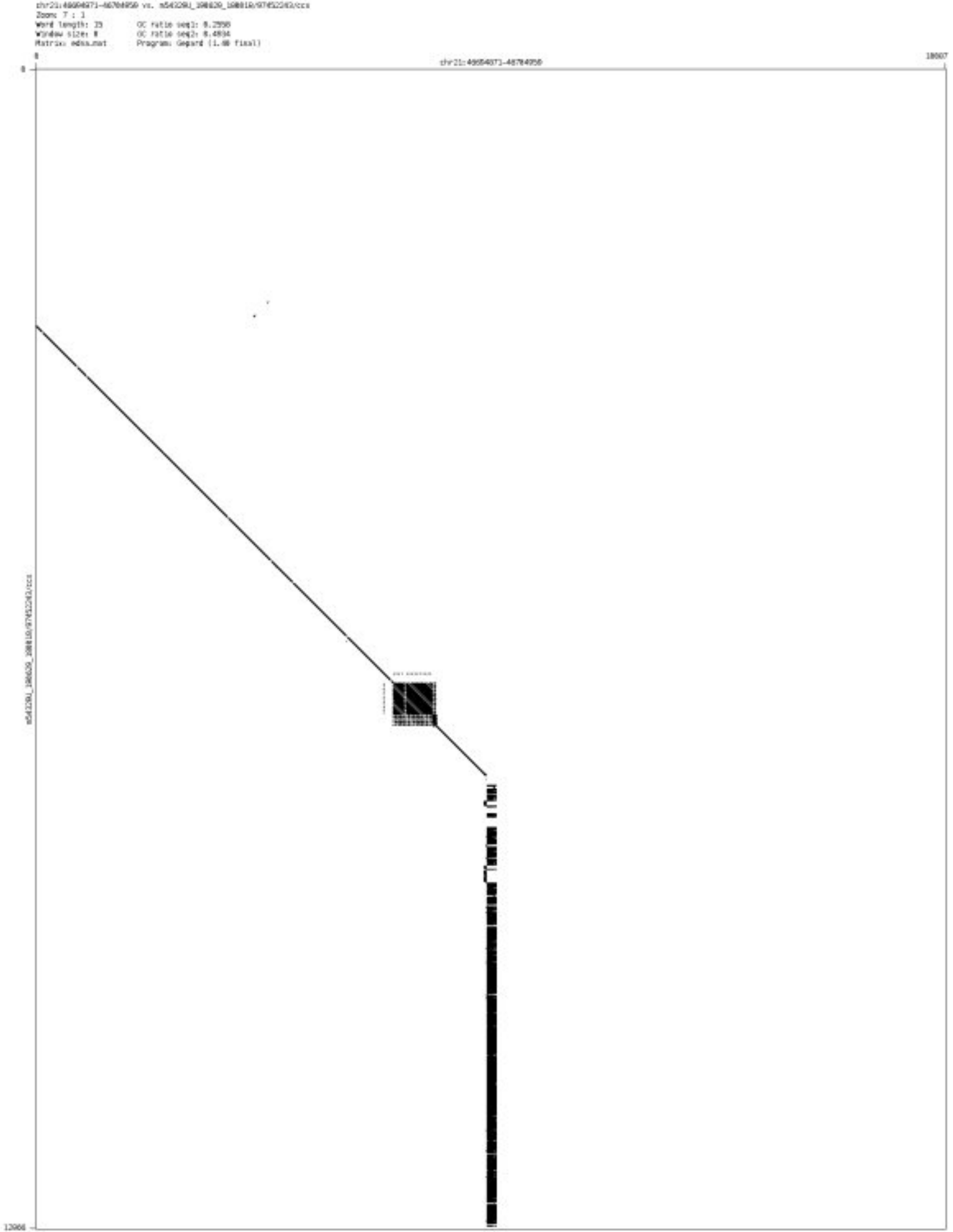

➢ chr22_16558546_16558645

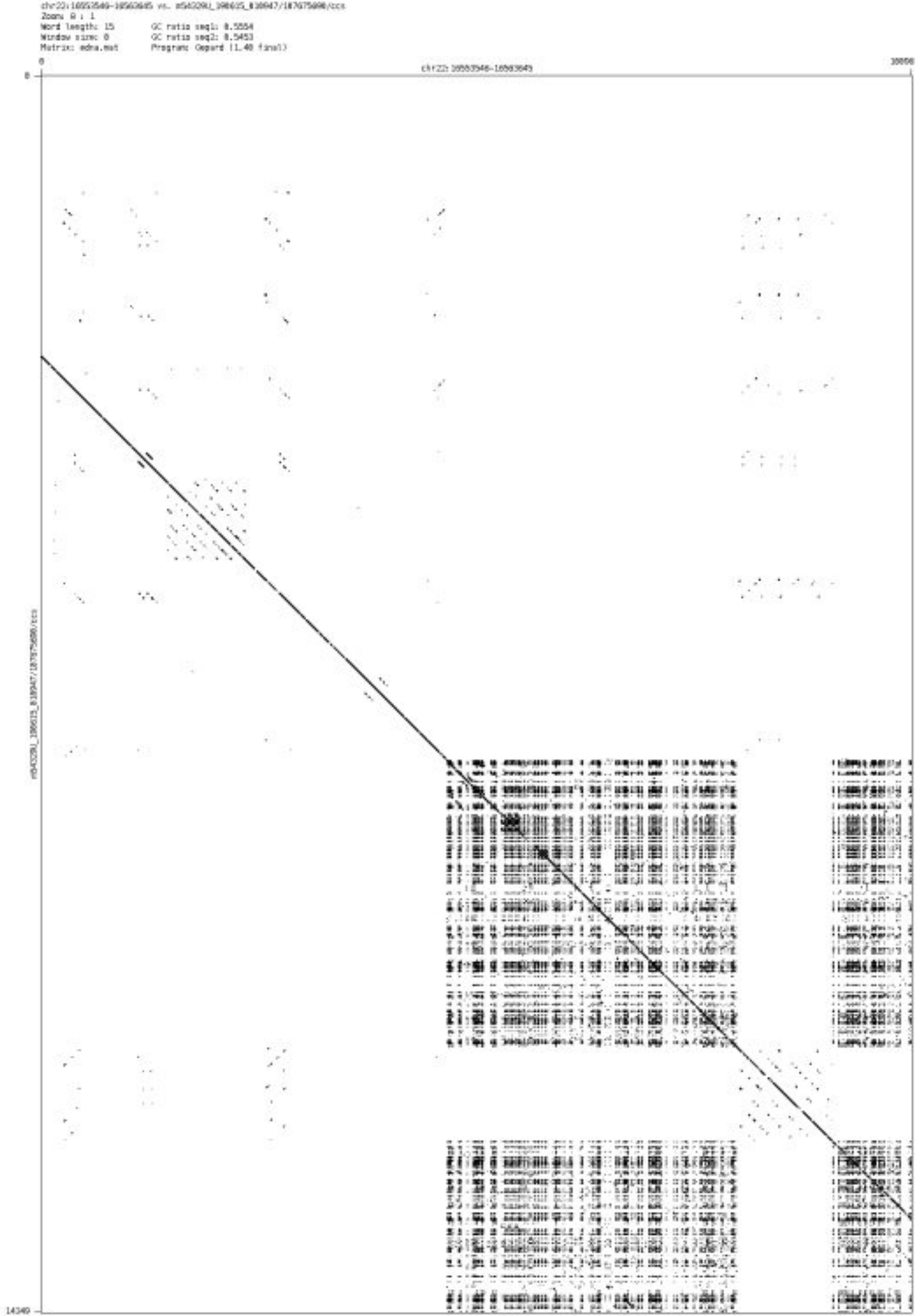

- chr22_17877099_17877318

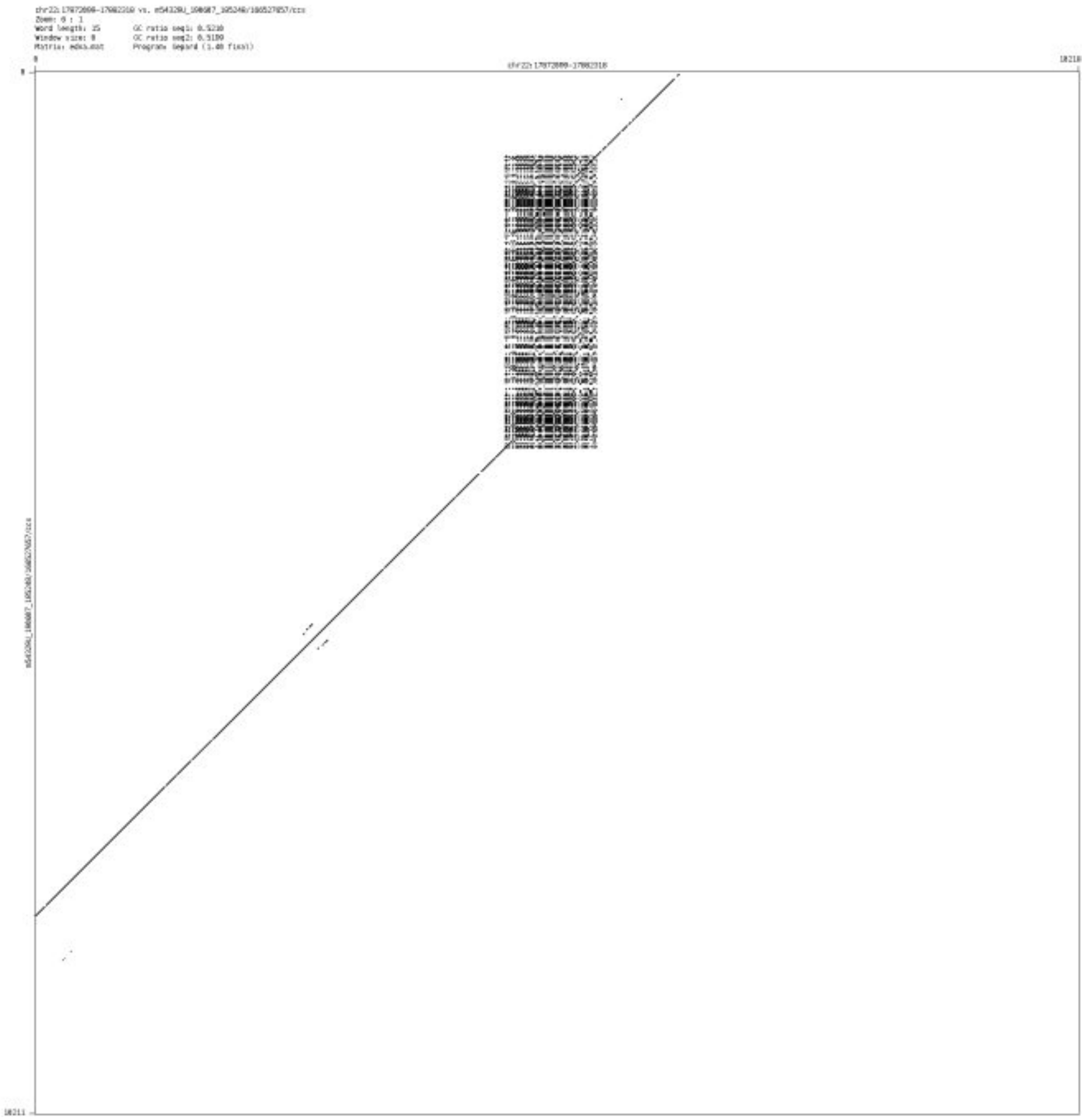

- chr22_29120197_29120382

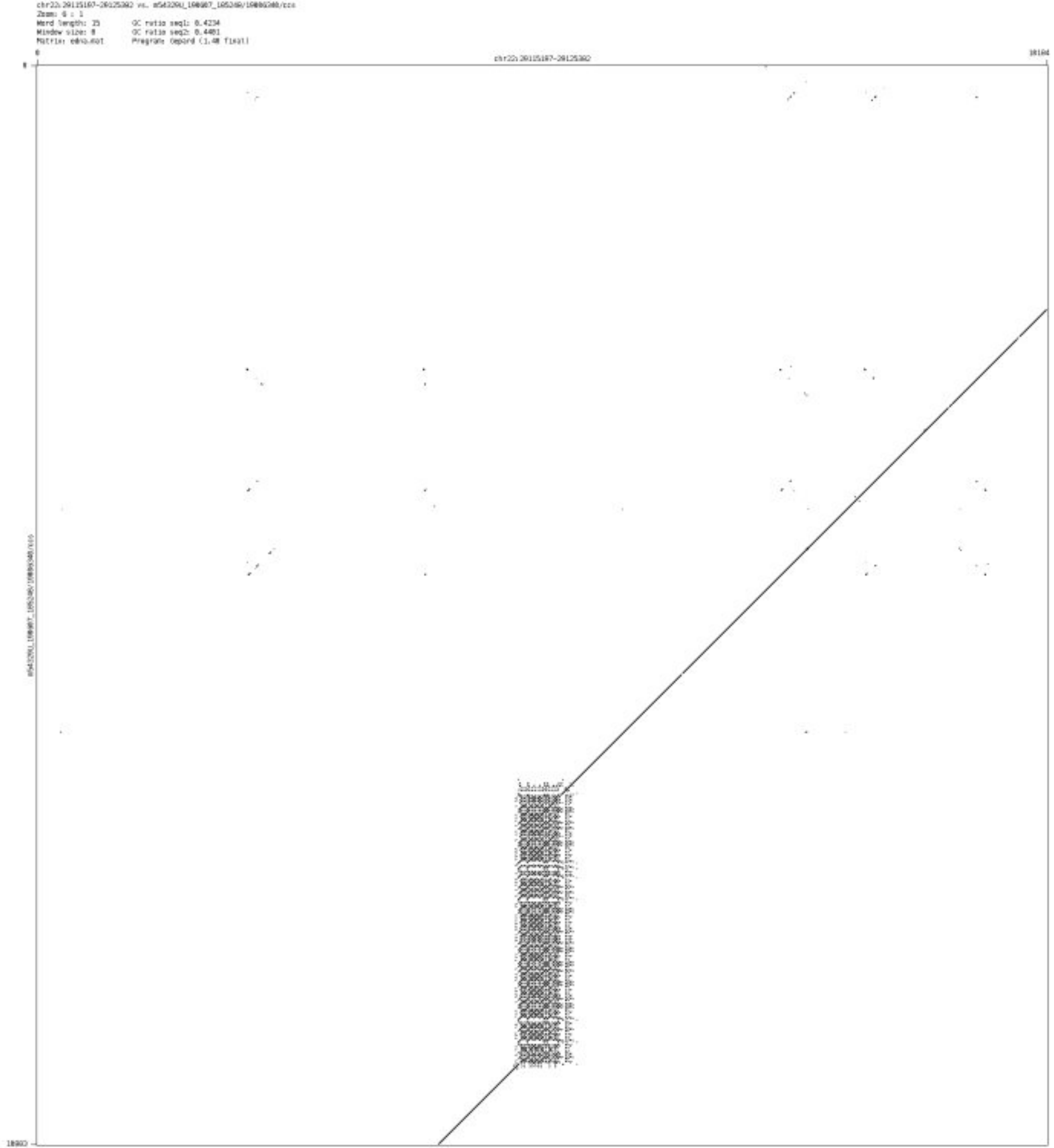

➢ chr22_42324053_42324236

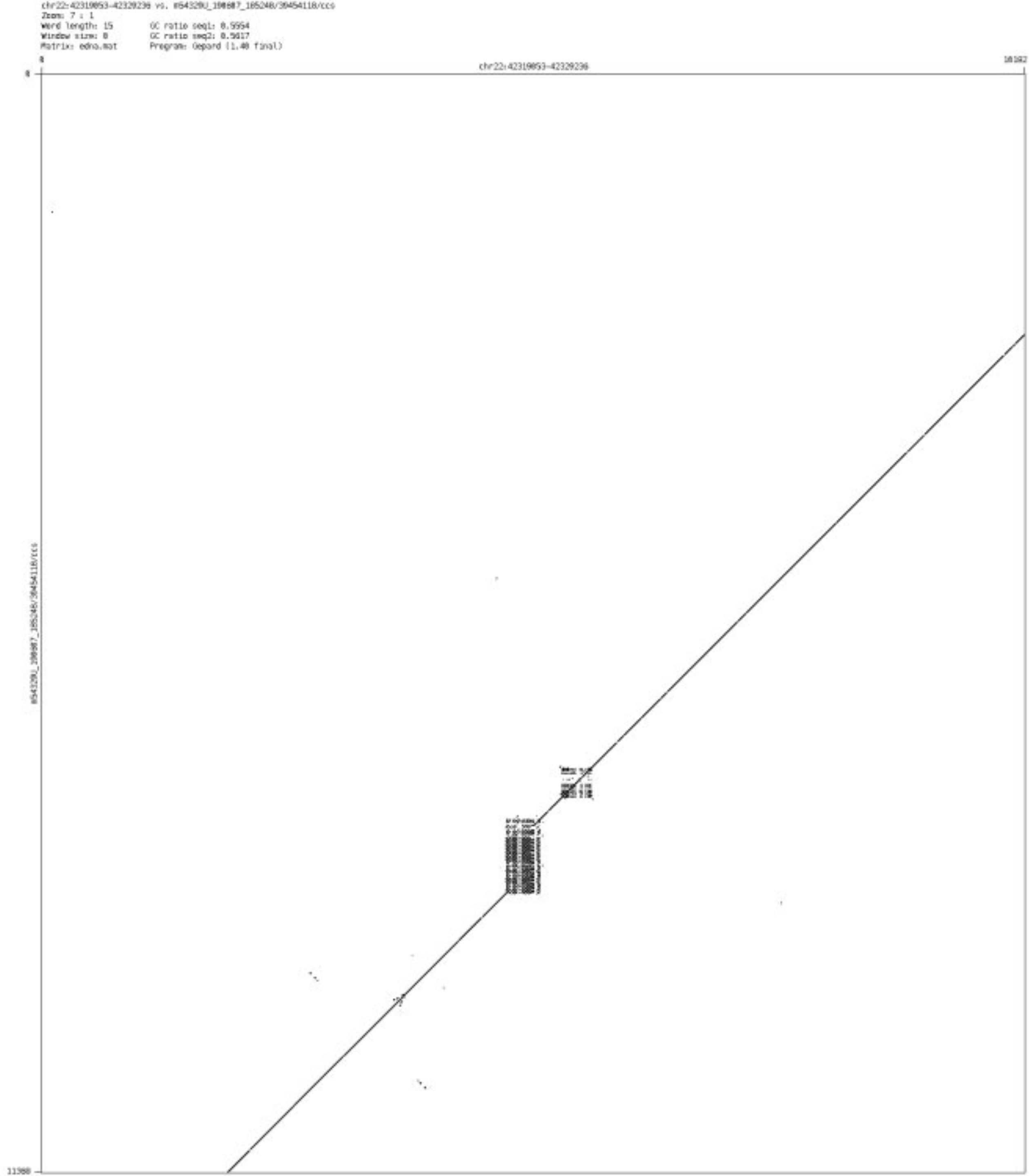

➢ chr22_43432701_43432877

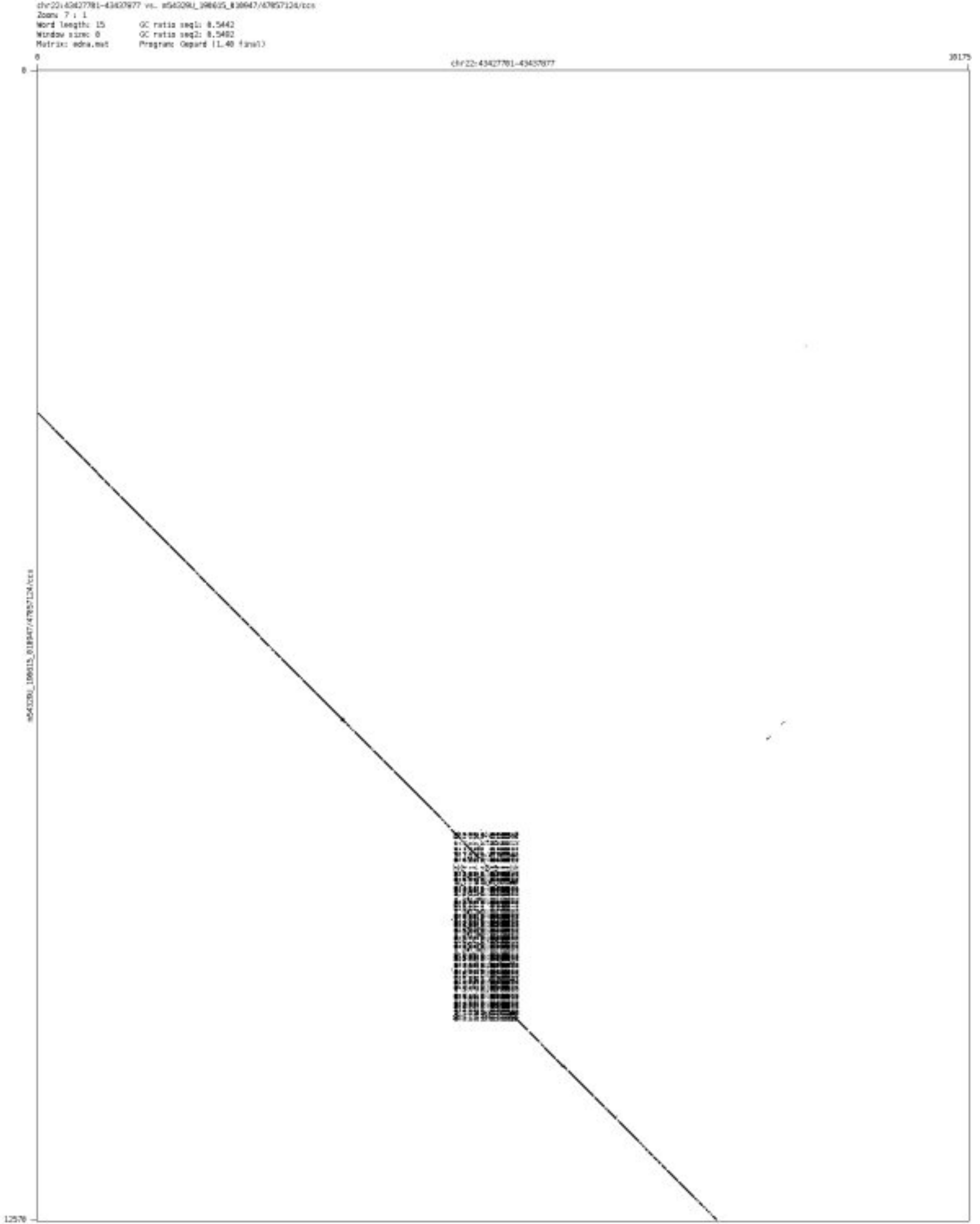

➢ chr22_47379190_47379385

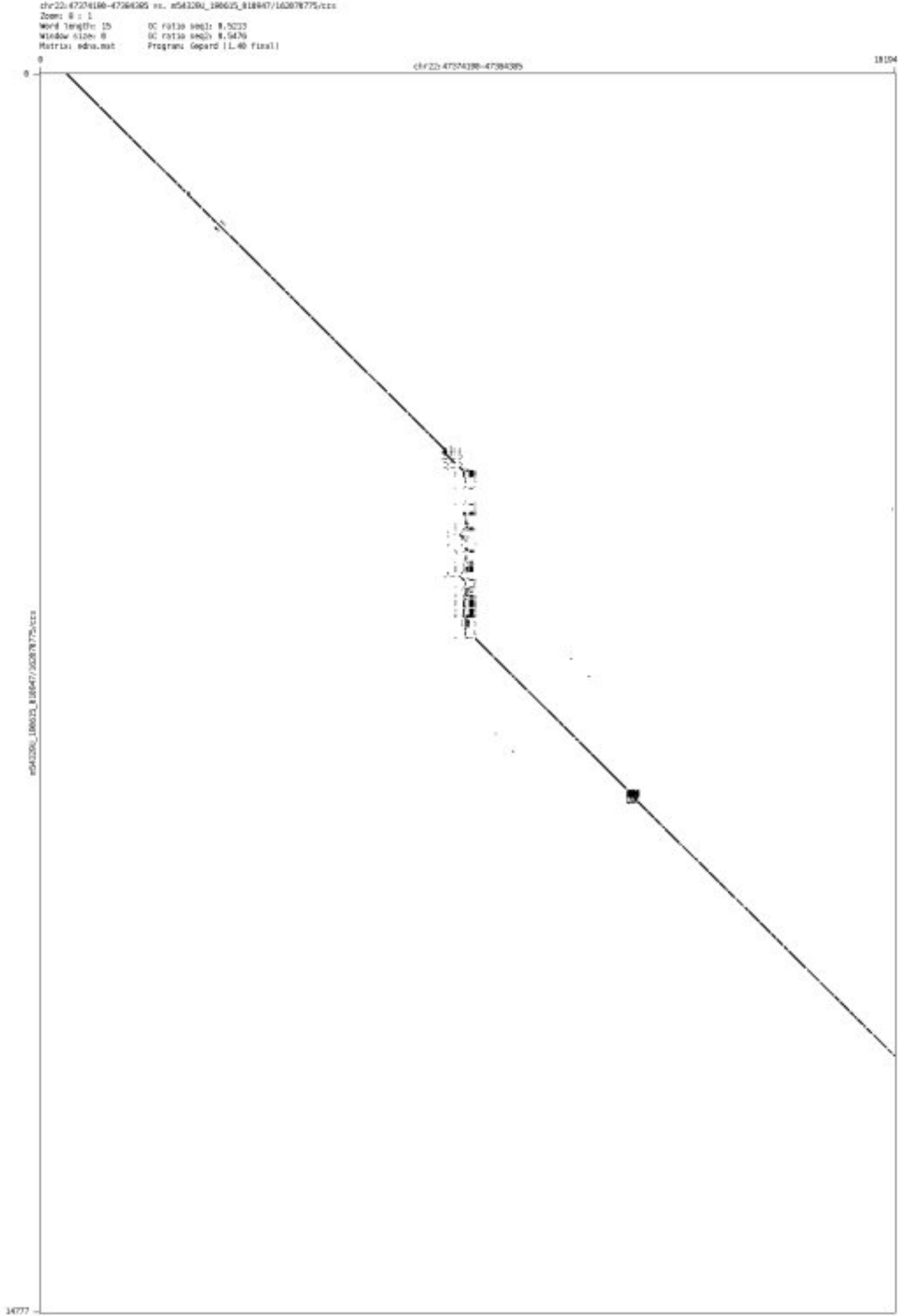

- chr22_50276306_50276417

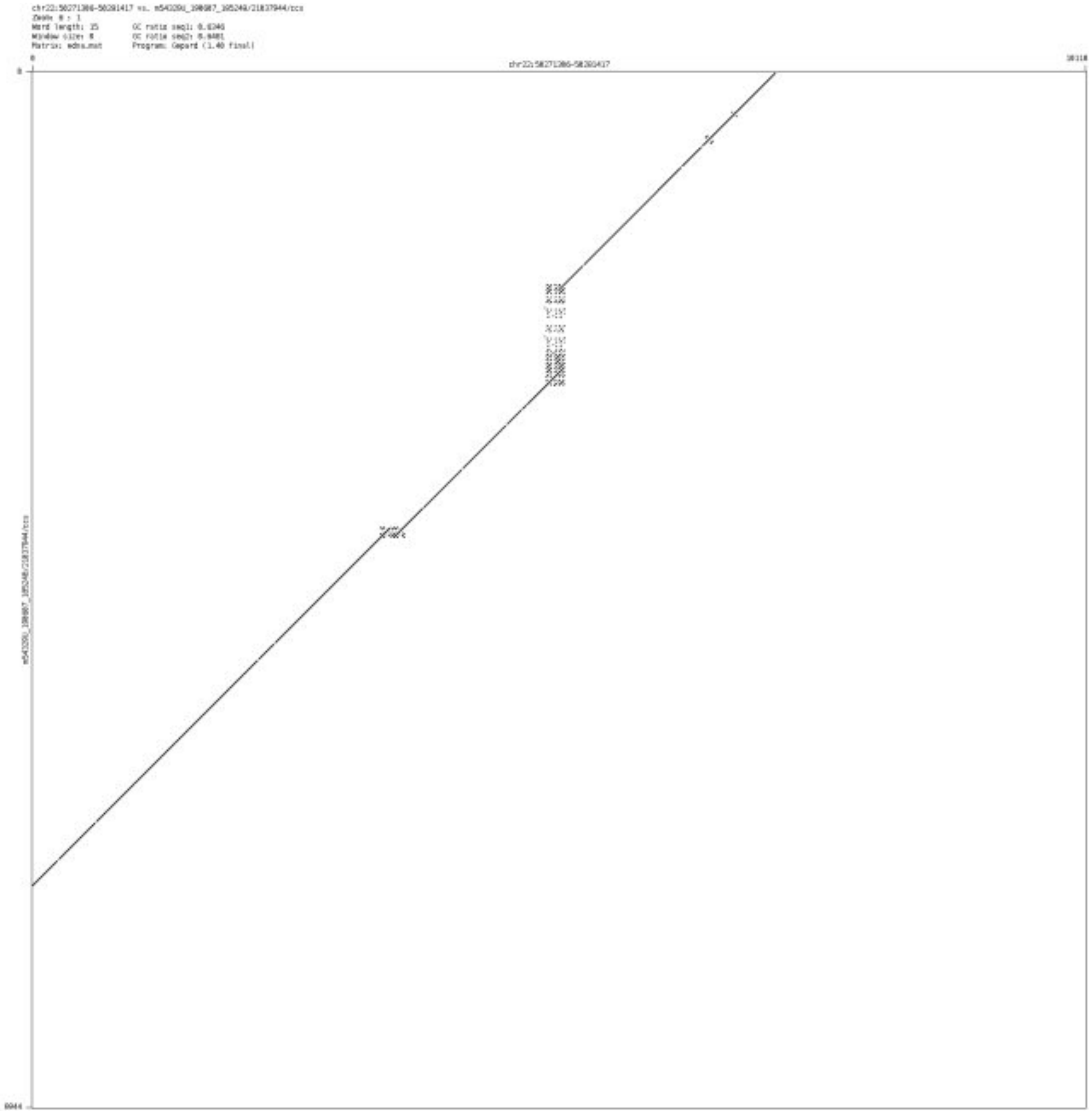

- chr22_50643752_50643941

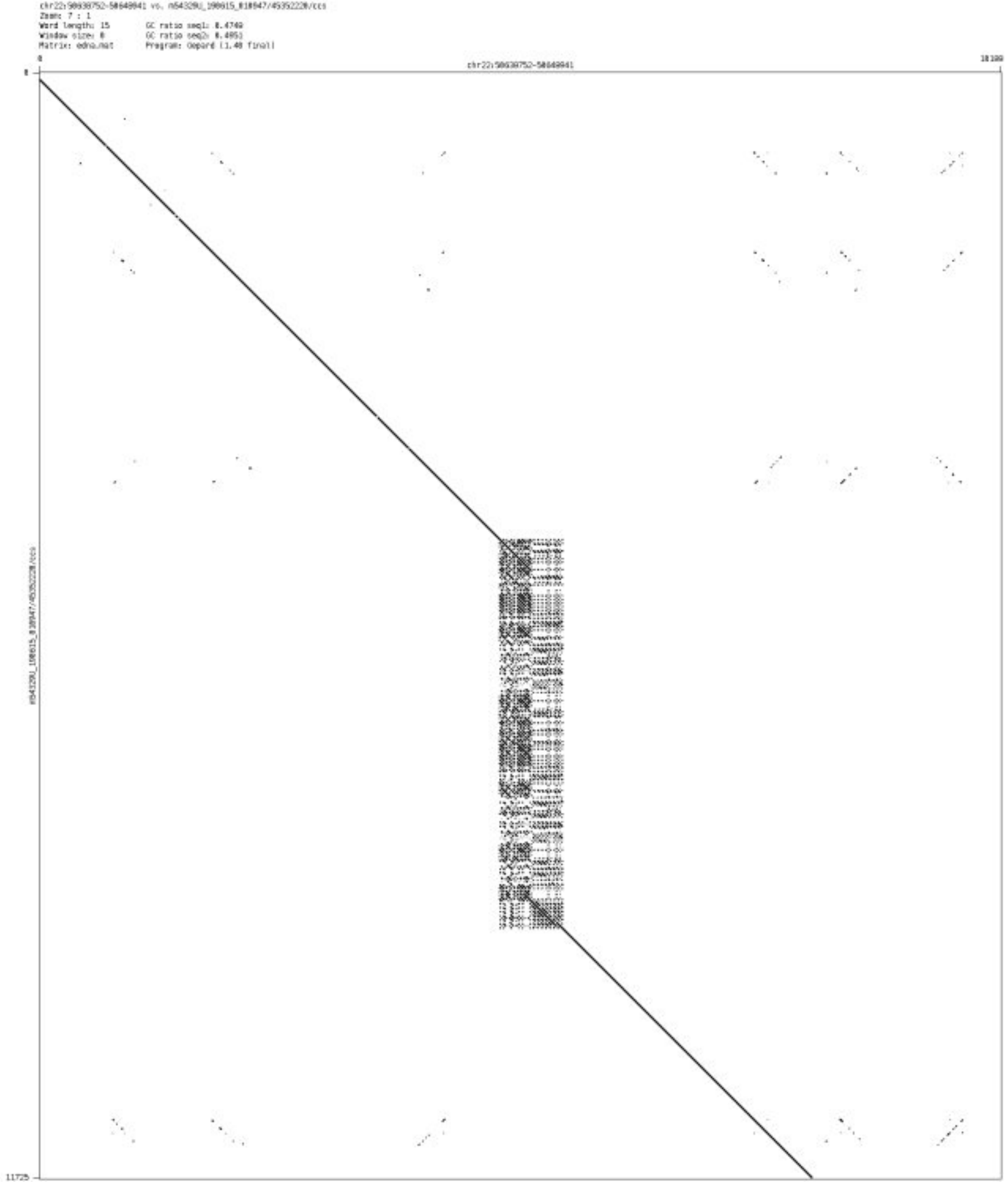

➢ chr22_50807923_50808081

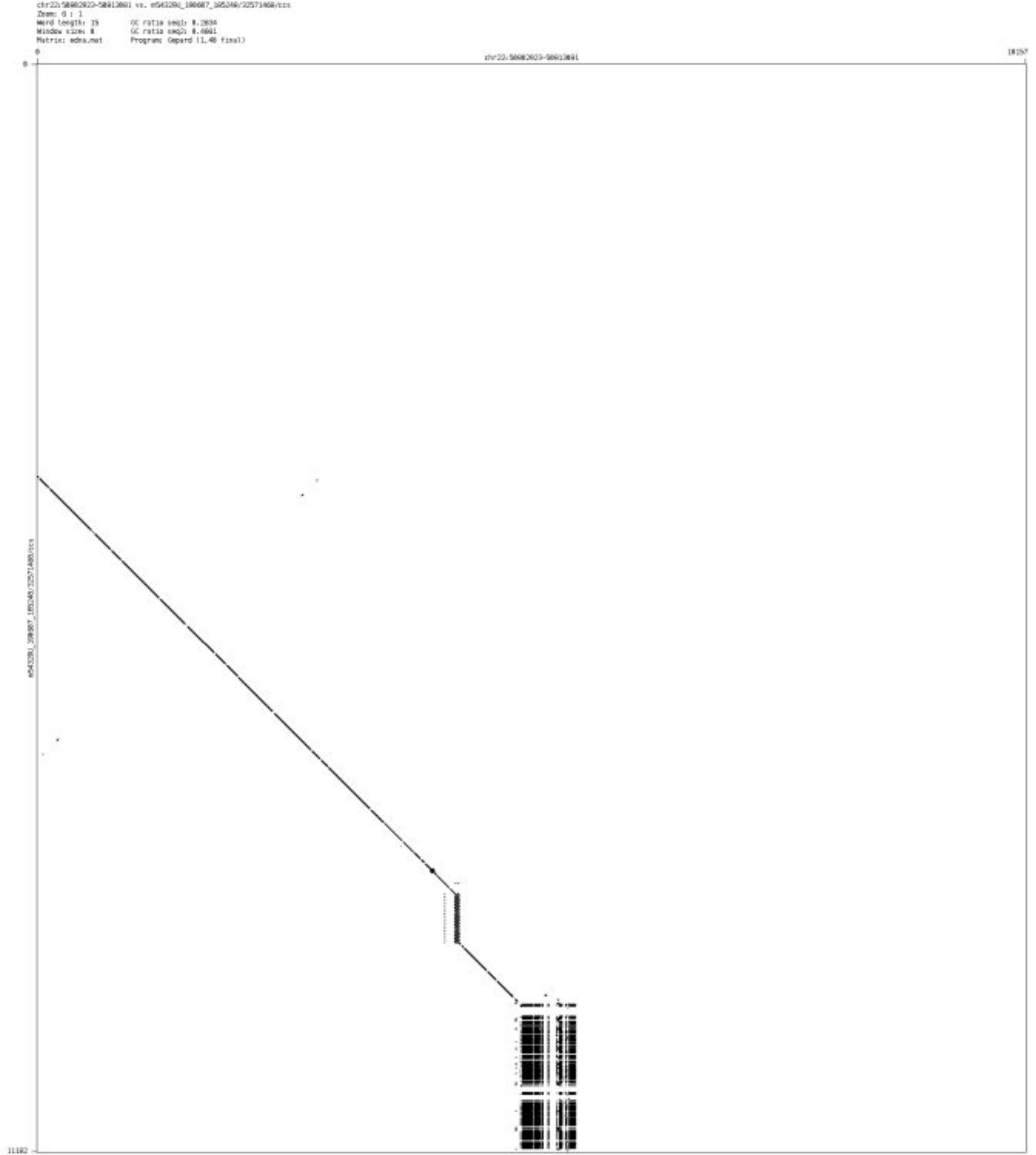
