## Supplementary File 3 for "Mako: a graph-based pattern growth approach to detect complex structural variants"

### PCR electrophoretic bands

**M:** DL5000 DNA marker. From bottom to top is: 100bp, 250bp, 500bp, 750bp, 1,000bp, 1,500bp, 2,000bp, 3,000bp and 5,000bp.
**Red arrow:** PCR product selected for sequencing with bright electrophoretic bands of targeted sizes

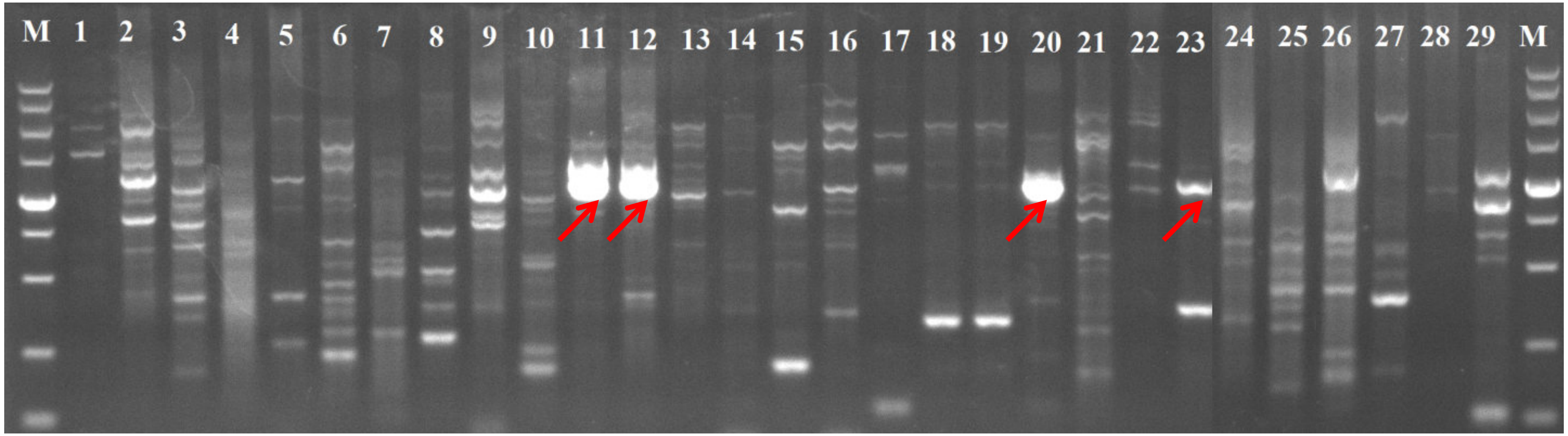

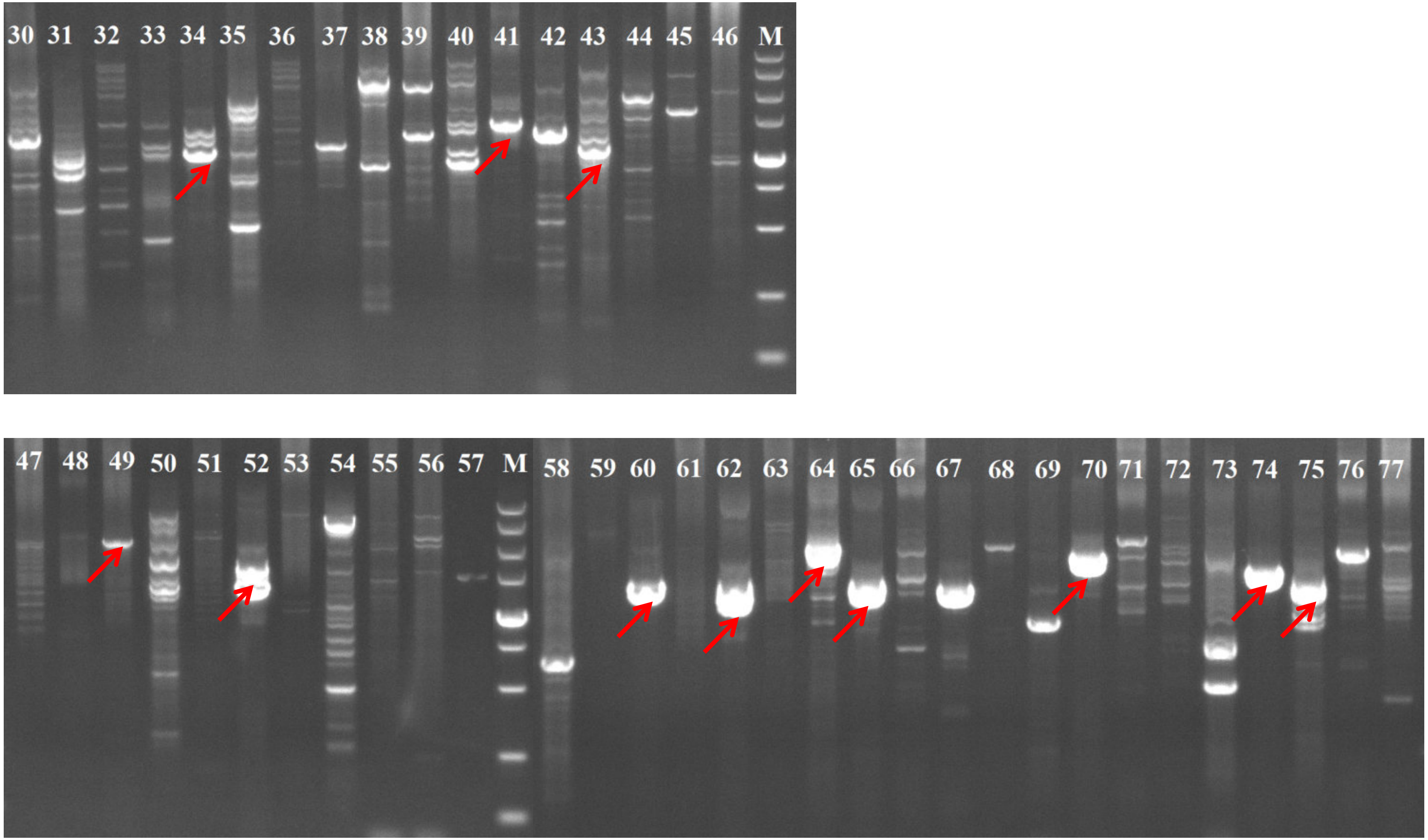
30 31 32 33 34 35 36 37 38 39 40 41 42 43 44 45 46 M
47 48 49 50 51 52 53 54 55 56 57 M 58 59 60 61 62 63 64 65 66 67 68 69 70 71 72 73 74 75 76 77

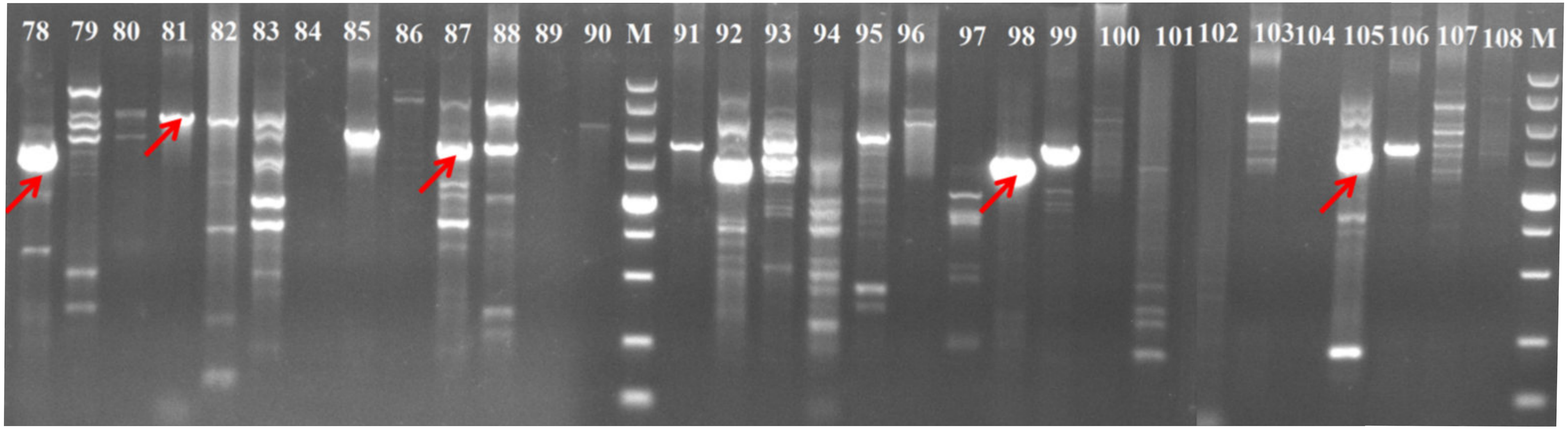


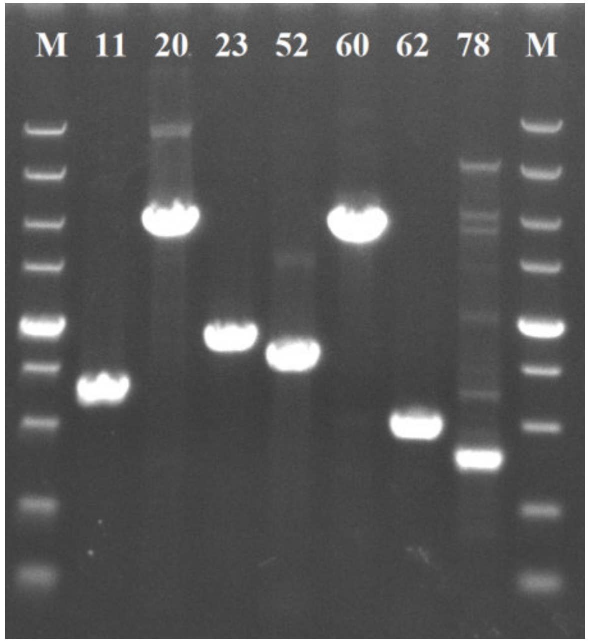


**We have designed extra primers for events 11, 20, 23, 52, 60, 62 and 78.**

### Gepard visualization of validated CSVs

### Mako chr1:81,194,398-81,195,874

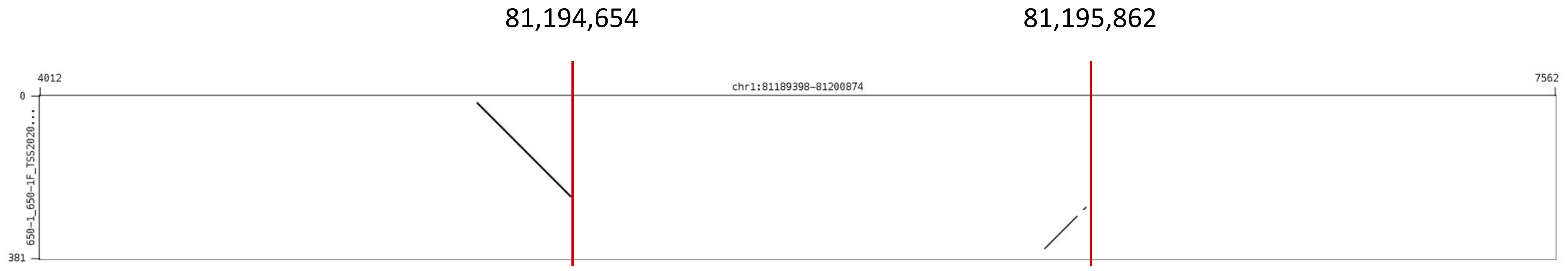

### **Mako** chr2:119,659,504-119,661,322

119,659,495

119,660,923

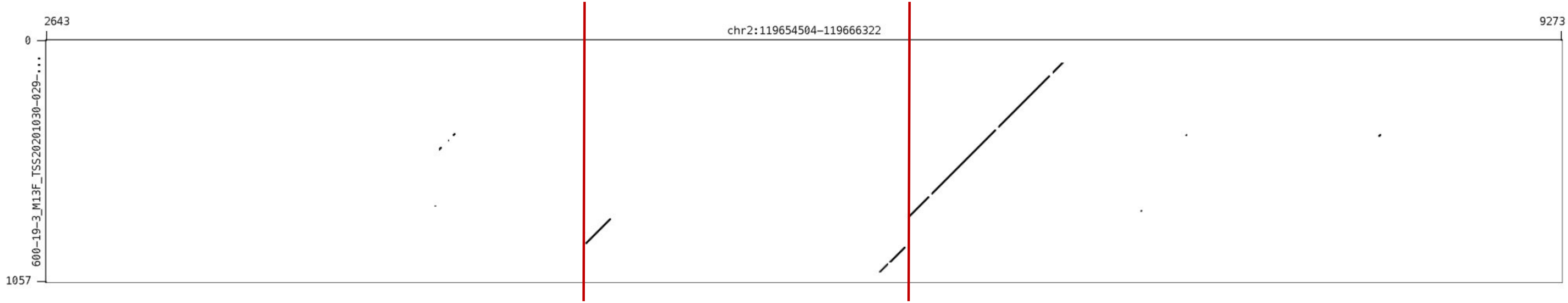

### **Mako** chr3:146,667,093-146,677,284

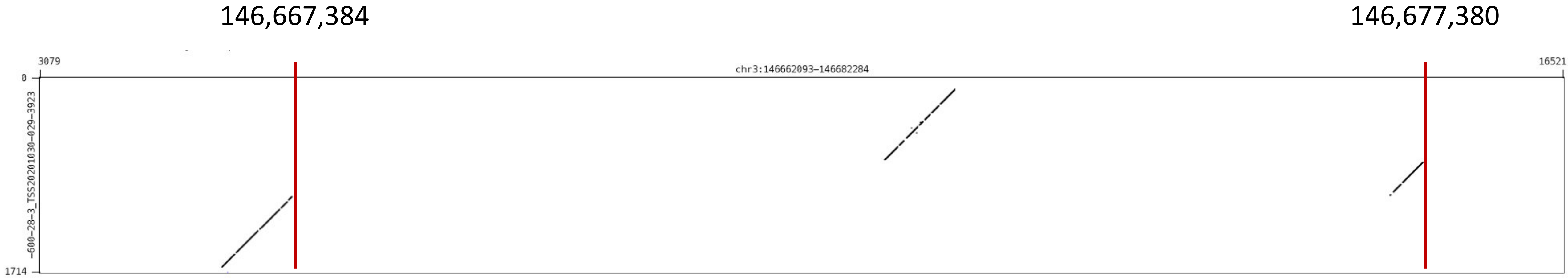

### Mako chr5:141,480,327-141,483,116

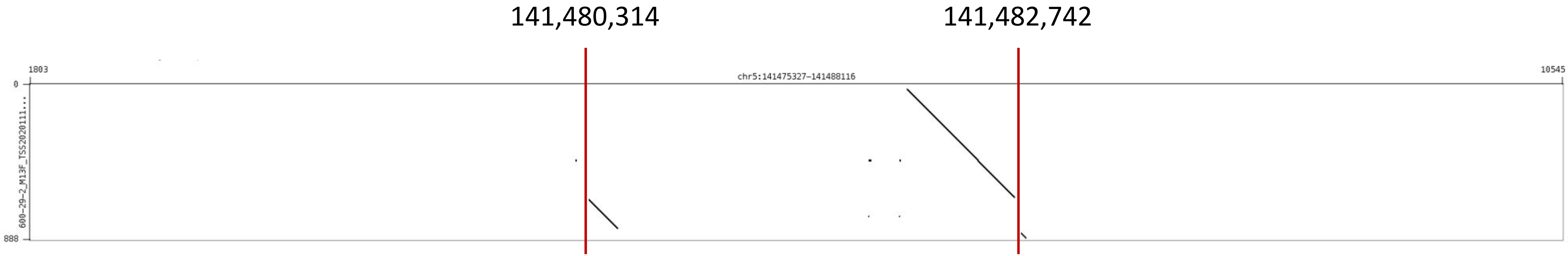

### **Mako** chr7:1,940,931-1,941,009

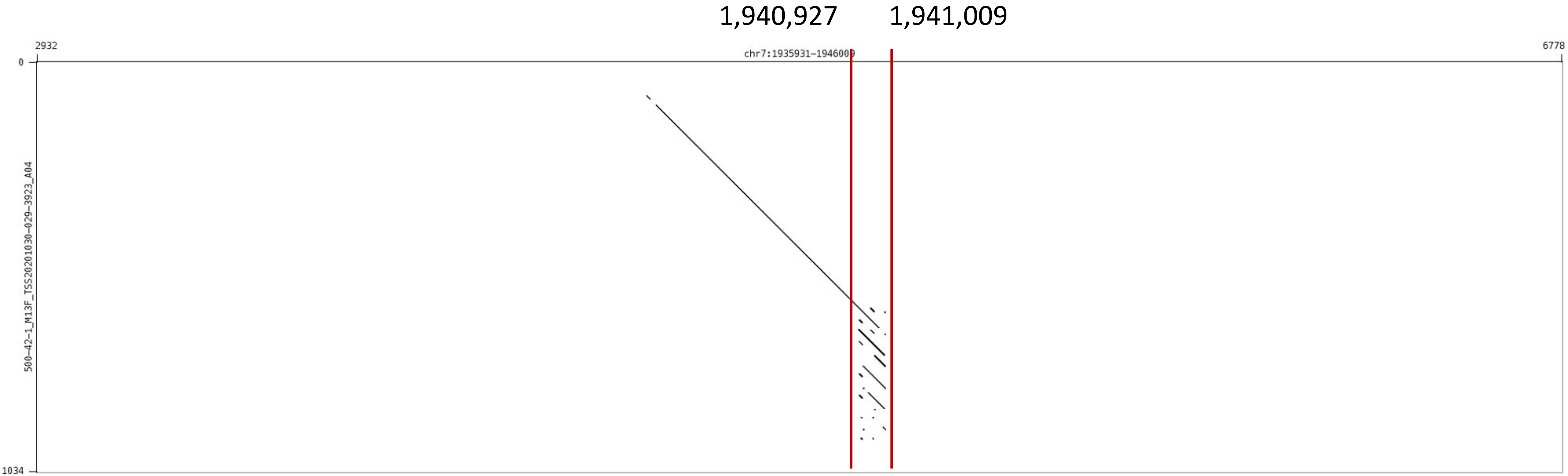

### **Mako** chr9:29,591,409-29,593,057

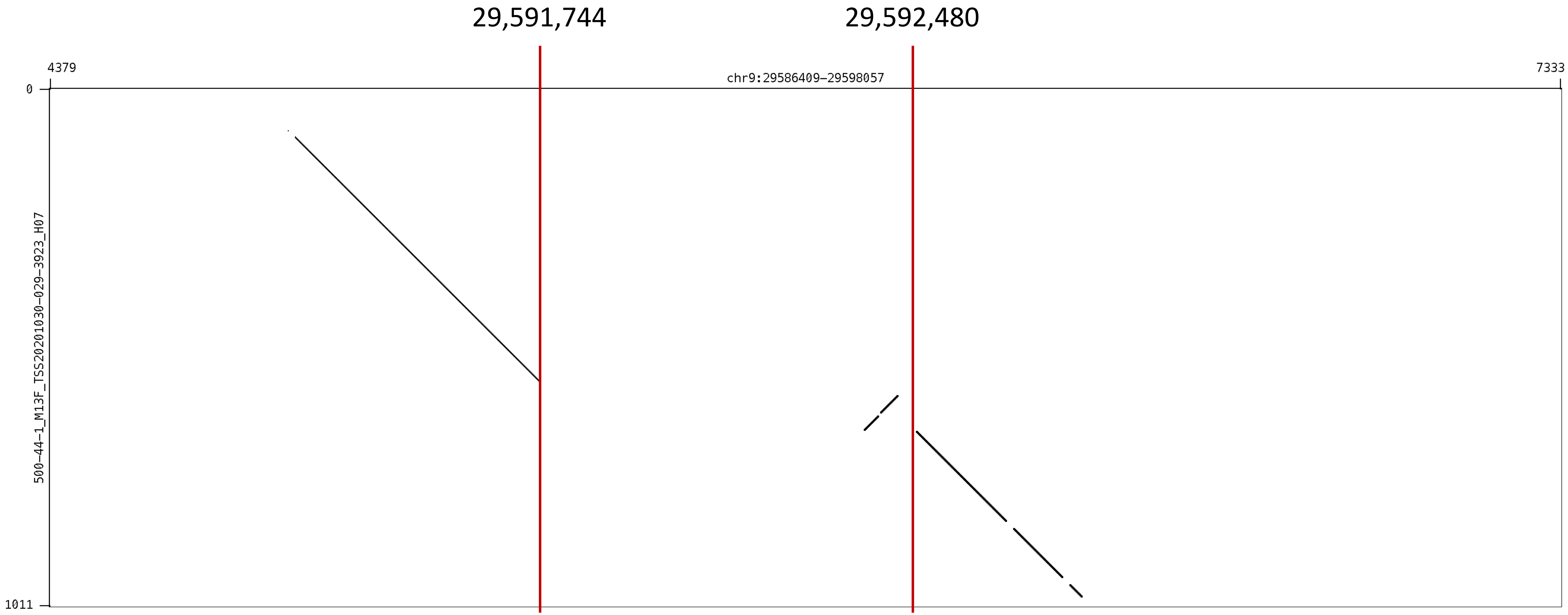

### Mako chr10:14,568,488-14,568,677

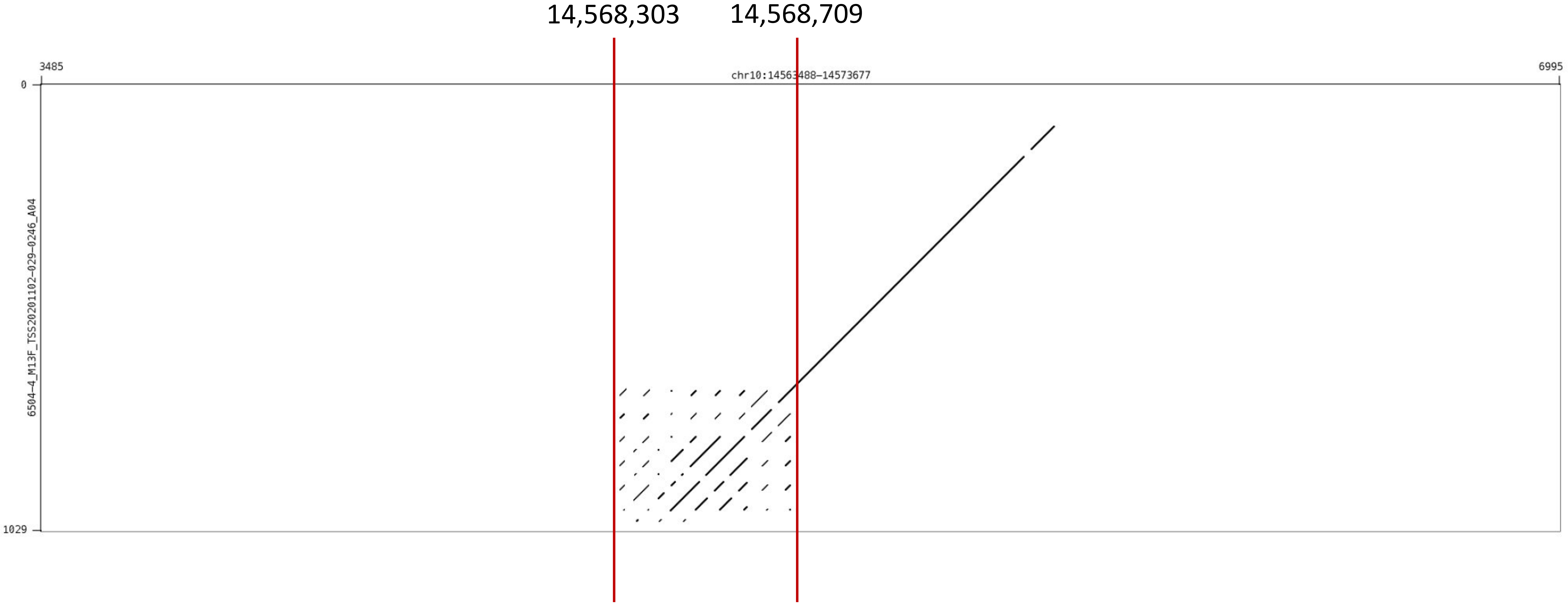

### Mako chr12:71,315,482-71,316,928

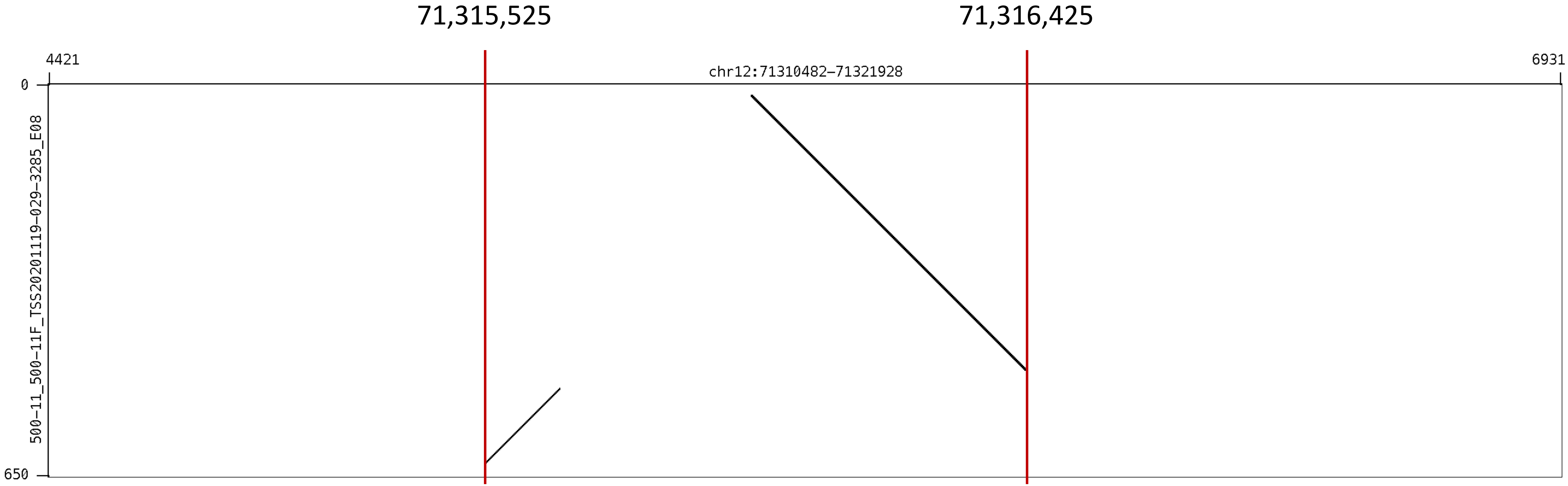

### **Mako** chr12:77,989,900-77,994,324

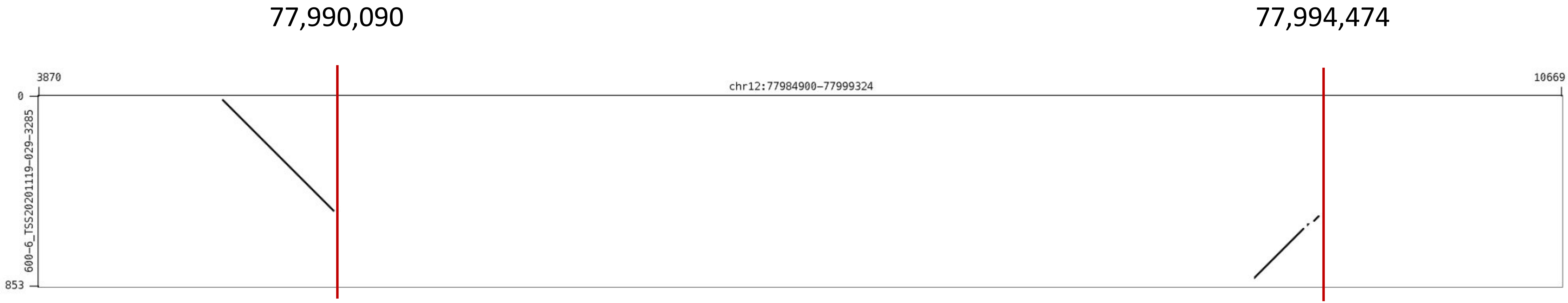

### **Mako** chr13:74,340,759-74,342,810

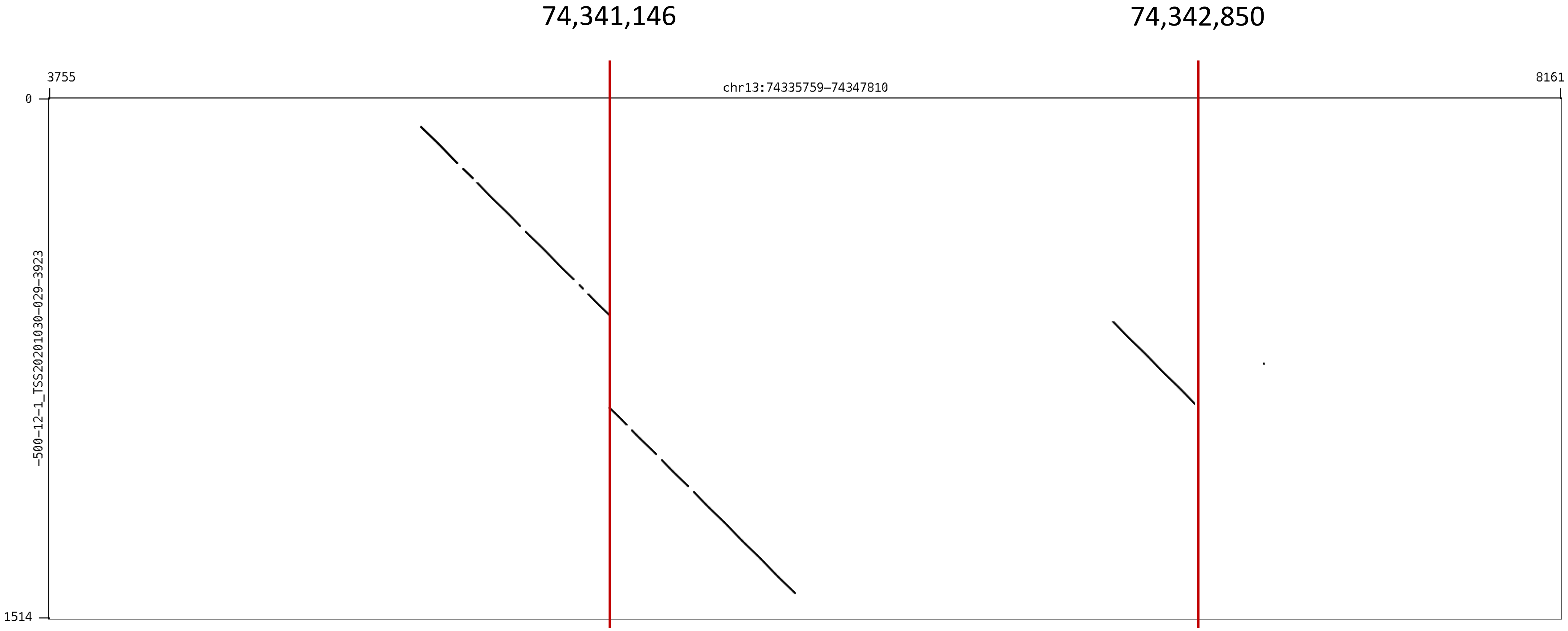

### **Mako** chr16:78,004,459-78,007,456

78,004,627

78,007,361

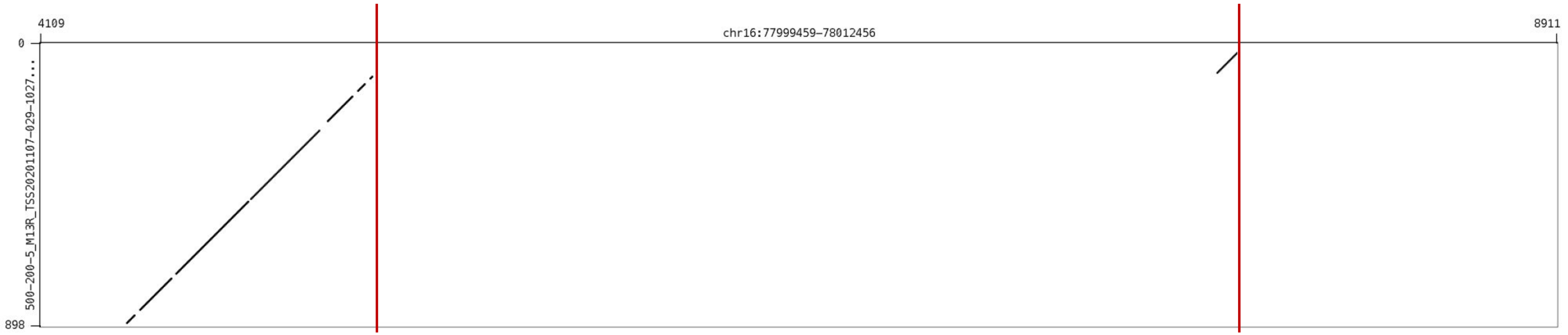

### **Mako** chr17:34,854,438-34,855,851

34,854,671 34,855,859

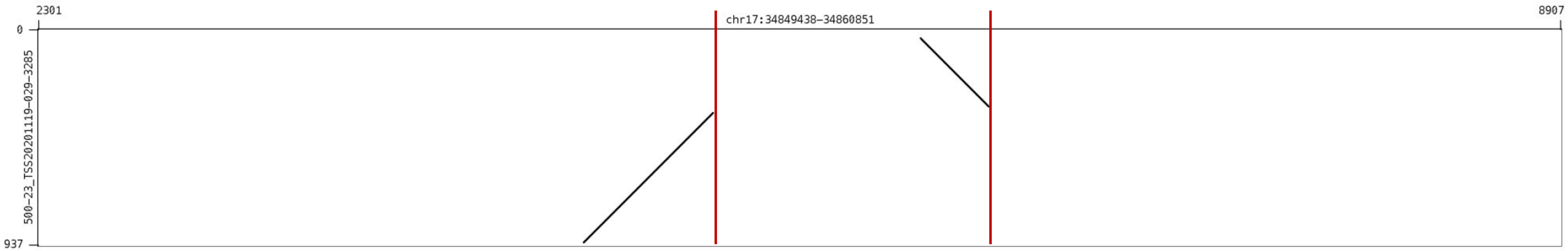

### Mako chr17:48,538,270-48,540,171

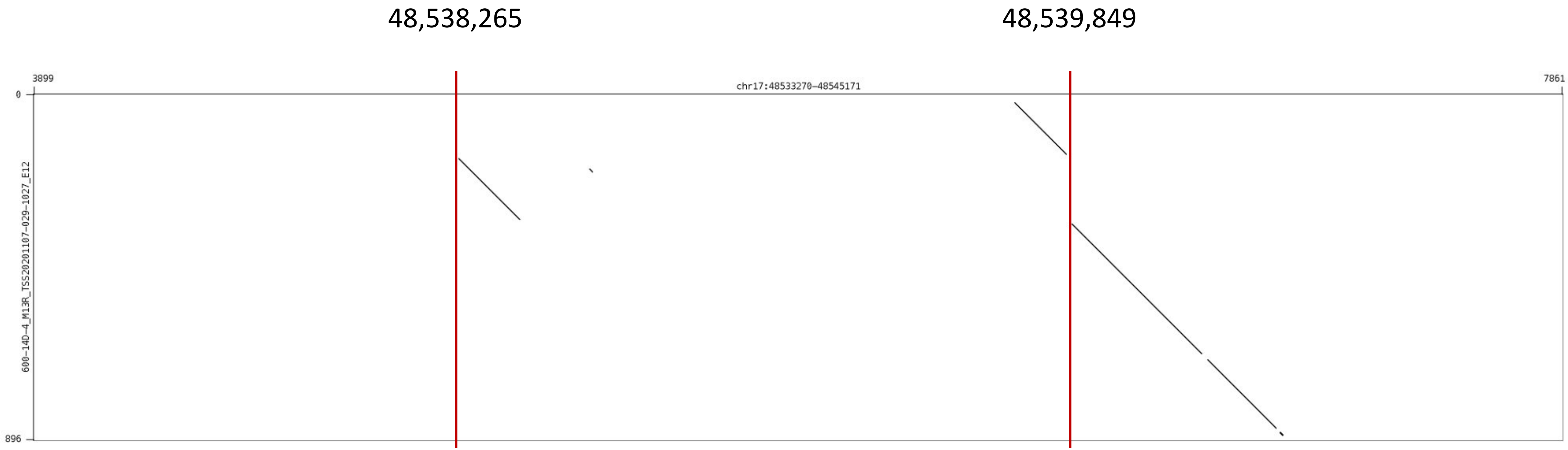

### **Mako** chr18:72,044,575-72,045,937

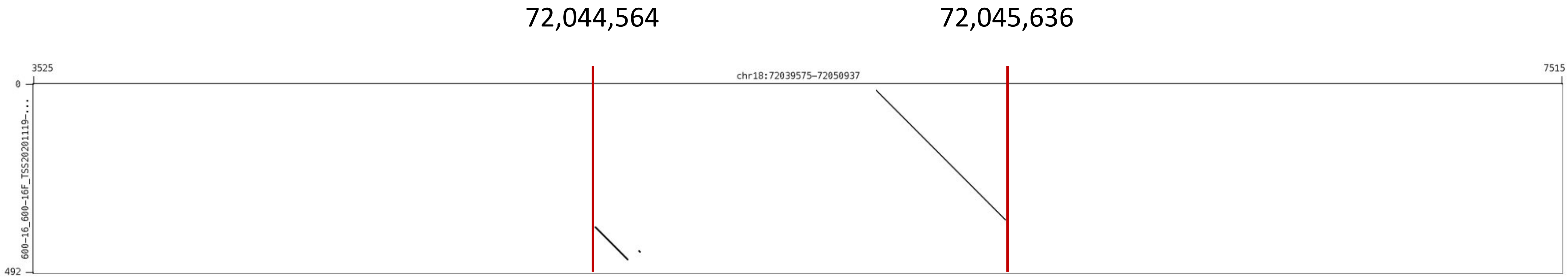

### **Mako** chr21:26,001,844-26,002,990

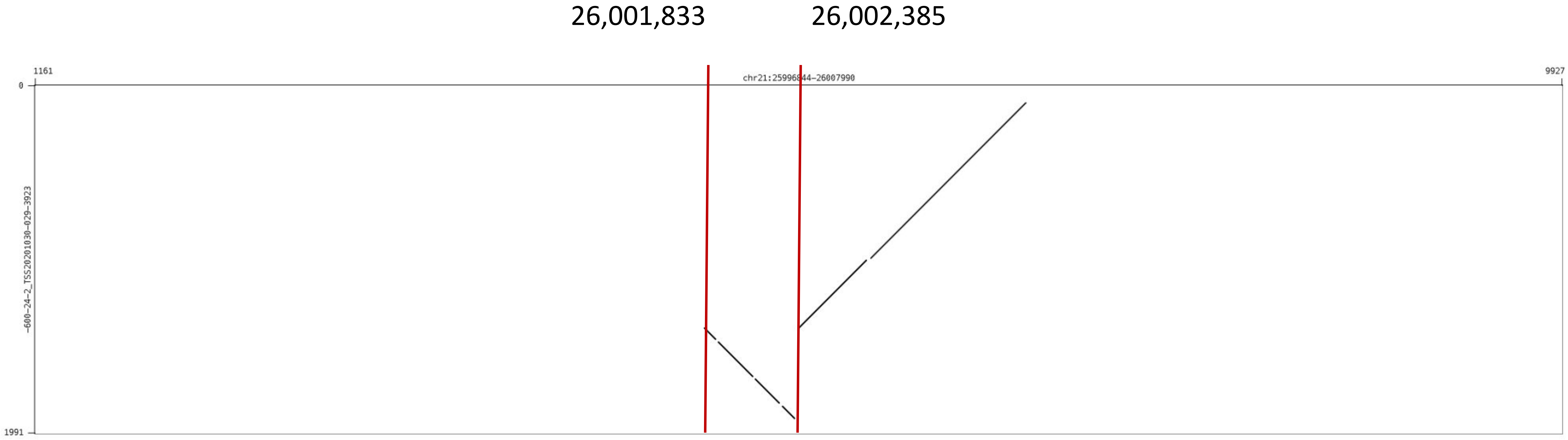
