## Supplementary Note for "Mako: a graph-based pattern growth approach to detect complex structural variants"

###### Table of contents

**I. Datasets resource**

**II. Create benchmark CSVs for real data**

**III. Performance evaluation**

**IV. Simulate complex structural variants**

**V. Experimental validation**

**VI. CSV manual inspection**

**VII. Computational validation**

**VIII. Running configurations**

#### Dataset resource

**High coverage Illumina BAM files for NA19240, HG00733 and HG00514** http://ftp.1000genomes.ebi.ac.uk/vol1/ftp/data*collections/hgsv*sv_discovery/data/

**NA19240 SVelter callset** http://ftp.1000genomes.ebi.ac.uk/vol1/ftp/data*collections/hgsv*sv*discovery/working/20160728*SVelter_UMich/

**PacBio HiFi reads for NA19240, HG00733 and HG00514** http://ftp.1000genomes.ebi.ac.uk/vol1/ftp/data_collections/HGSVC2/working/

**SK-BR-3 Illumina BAM file** http://labshare.cshl.edu/shares/schatzlab/www-data/skbr3/SKBR3*550bp*pcrFREE*S1*L001*AND*L002*R1*001.101bp.bwamem.ill.mapped.sort.bam

**SK-BR-3 SURVIRVOR merged callset** http://labshare.cshl.edu/shares/schatzlab/www-data/skbr3/SURVIVOR*1k*min2_min50bp.vcf.gz

**SK-BR-3 PacBio aligned BAM file** http://labshare.cshl.edu/shares/schatzlab/www-data/skbr3/reads*lr*skbr3.fa*ngmlr-0.2.3*mapped.bam

**HG00733 haploid HiFi assembly** http://ftp.1000genomes.ebi.ac.uk/vol1/ftp/data*collections/HGSVC2/working/20200628*HHU*assembly-results*CCS_v12/assemblies/phased/

**HG00733 Oxford Nanopore sequencing** http://ftp.1000genomes.ebi.ac.uk/vol1/ftp/data*collections/hgsv*sv*discovery/working/20181210*ONT_rebasecalled/

#### Create benchmark CSVs for real data

##### Prerequisites

Two scripts clusterSV.py and splitFasta.py, this can be found in our github repository. To run this script, the following dependencies have to be satisfied: * Python3 (version >= 3.6) * Packages: scipy, statistics, numpy, matplotlib, biopython

Gepard dotplot tool (http://cube.univie.ac.at/gepard). Java v1.8 is required to run the program.

###### Step1: clustering discovered SVs

This function is implemented as *clusterSV.py*, which takes illumina callset as input and output SV clusters. We use the median of events in the cluster as start and end for cluster coordinates.

```
python clusterSV.py -i input.bed -c chrom -w work_dir/
```

The output files are named as ***chrom.svclustersctxxx.bed*** under your working directory. ct is the cluster merge cutoff.

###### Step2: align CCS reads

This step is only for NA19240, HG00514 and HG00733. We use public available alignment file of PacBio reads for SK-BR-3.

```
# align each subread with pbmm2
pbmm2 align GRCh38.reference.fa prefix.ccs.fastq.gz prefix.pbmm2.srt.bam --preset CCS --sort -j 8 -J 8 --sample sample_name

# align each subread with NGMLR
ngmlr -t 8 GRCh38.reference.fa -q prefix.ccs.fastq.gz -o prefex.nglmr.sam
samtools sort -o prefix.nglmr.srt.bam

# Merge pbmm2 aligned BAMs
samtools merge merged.ccs.pbmm2.srt.bam *.pbmm2.srt.bam
samtools index merged.ccs.pbmm2.srt.bam

# Merge NGMLR aligned BAMs
samtools merge merged.ccs.ngmlr.srt.bam *.nglmr.srt.bam
samtools index merged.ccs.ngmlr.srt.bam
```

**Step3: prepare data for sequence dotplot**

*splitFasta.py* is self-developed script to split the multi-fastq file.

```
# Get reference sequence corresponding to each SV cluster
bedtools getfasta -fi referece.fa -bed sv_cluster.bed -fo cluster_ref.fa
# Get long reads that span each SV cluster
samtools view -b pacbio_aligned.bam sv_cluster.bed | samtools fast > cluster_reads.fa
# Split the multi-fastq file
python splitFasta.py --input cluster_reads.fa --output work_dir/
```

Finally, we obtain the reference sequence and long read sequences separated in each files.

**Step4: create sequence dotplot and inspection**

We use Gepard to visualize and create reference to reads dotplots.
`java -cp /path/to/Gepard-1.40.jar org.gepard.client.cmdline.CommandLine -seq1 cluster_ref.fa -seq2 read.fa -matrix /path/to/gepard/resou`

### Performance evaluation

The comparison is based on the idea from Truvari and Peter A.Audano's 2019 Cell paper. In general, it searches for closest event of similar size. If such predictions can be found from the compare set, we will calculate the reciprocal overlap. The comparison parameters are listed below. For more information about evaluation schema, please refer Truvari (https://github.com/spiralgenetics/truvari).

**Matching parameters**

| Parameter | Default | Definition |
|---|---|---|
| bpDist | 500 | Max distance between breakpoints of two events |
| size | 0.7 | Minimum size similarity of two events |

**Evaluation measurements**

| Metric | Definition |
|---|---|
| TP-base | Number of matching calls from the truthset |
| TP-call | Number of matching calls from the prediction set |
| FP | Number of non-matching calls from the prediction set |
| FN | Number of non-matching calls from the truthset |
| precision | TP-call / (TP-call + FP) |
| recall | TP-base / (TP-base + FN) |
| f1 | (recall * precision) / (recall + precision) |
| base cnt | Number of calls in the truthset |
| call cnt | Number of calls in the prediction set |

The script *compare.py* can be found at Mako's Github repository, which evaluates the overlaps between truthset and predictions.

```
Usage:
compare.py [commands] <parameters>

Commands:
bed: evaluate overlaps between two BED files
both: evaluate complete and unique match, all breakpoints match
base: evaluate complete and unique match
```

Please check the required inputs for different commands by running: `compare.py base -h compare.py both -h compare.py bed -h`

The evaluation function create two files.

- ***prefix.stats.txt***: contains evaluation measurements listed in the previous table.
- ***prefix.bed***: contains overlapped calls. The 1st column is the ID of observations. The 2nd column is the ID for benchmark variants. The 3rd column contains the number of observations that matches a benchmark. The 4th and 5th column are similarity info of the best match among all matches.

**For simulated data evaluation**, we use the **both** command. This will generate two statistic summary files *prefix_base.stats.txt* and *prefix_sub.stats.txt*.

- Unique-interval match results are from *prefix_base.stats.txt*.
- All-breakpoint match results are from *prefix_sub.stats.txt*.

**For real data evaluation**, we use the **bed** command. The VCF file is firstly converted to BED file. This command will only produce the *prefix_base.stats.txt*.

The VCF file conversion of SVelter, GRIDSS and TARDIS follows the filtering process, and is obtained through *convert.py*. The first four columns of output BED file **must** be chrom, start, end and length. ``` Usage: convert.py [options]

Options: -h --help show this help message and exit -s SAMPLE Sample name (yri, skbr3) -t TOOL Detection method (gridss, svelter, tardis) -v VCF Path to VCF file -b BED Path to output BED file ```

### Simulate complex structural variants

We use the idea of VISOR to simulate complex rearrangement and create a modified script simulate.py to create complex events. VISOR is a tool that can simulate simple SVs at different haplotypes which includes deletion, insertion, tandem duplication, translocation copy-paste and more information can be found in the Github (https://github.com/davidebolo1993/VISOR). Dependencies are listed below to run simulations * Python (>=3.6) * Python package for simulation data: pysam, pybedtools, pyfaidx, wgsim, bwa.

We recommend to use the python3.6 virtual environment created by Conda. All of these packages can be installed through Conda. Please refer VISOR for details.

#### Simulate reported complex events

We first simulate some reported complex events by other studies, including deletion flanked by inversion, inverted duplication, dispersed duplication and etc. Please see figure below for details. This is done on chr1 as described in steps.

***Note:*** We only randomly select and combine from deletion, inversion, inverted tandem duplication, tandem duplication. Translocation copy-paste are not modified. Meanwhile, transolcation copy-paste are treated inverted and dispersed duplication. The BED file of this simulation can be found in our Github.

###### Step1: Random generate SV regions

```
# Create random regions
Rscript randomregion.r -d chrom.dim.tsv -n 300 -l 5000 -s 500 -x chr1.exclude.bed -v 'deletion,inverion,tandem duplication,inverted tandem duplication,translocation coyp-paste' -r '35:35:5:10:15' | sortBed > chr1.random.bed
```

Under your working directory, you will find generated random regions in BED files.

###### Step2: Create nested events

We next randomly add events to those generated in step2. The added events involve deletion, inversion, inverted tandem duplication and tandem duplication. For example, deletion associated with inversions can be simulated as below, which is similar to SURVIVOR. If deletion is generate in Step1, inversions are randomly added events.

> chr1 20000000 20001000 inversion chr1 20001001 21000000 deletion chr1 21000001 210001000 inversion

The below command will generate two files: 1) added.SVs.bed, this saves all simple events; 2) added.SVs.nested.bed contains the nested events and will be used as the truth set, each nested event contains at least two simple events.

```
# Simulate reported CSVs with known command
python simluate.py known -i chr1.random.bed -w /path/to/work_dir -c chr1
```

###### Step3: Simulate the variation genome with CSVs

In this step, we will add created events to the genome with SNPs created at step1. Only one haplotype for nested events, the other one only has SNPs.

```
Python simulate.py sim —g templatewithsnp/chr1.fa —bed chr1.added.SVs.bed  —o variation_genome/ -c chr1
```

###### Step4: Start simulate reads

Once the variation genome has been created for each chromosome, we can start to simulate and align reads with VISOR.

These steps are required for creating input for reads simulation.

```
# Get chromosome size information of variation genome and get the maximum dimension for further simulation
cd variation_genome/ && find . -name "*.fa" -exec samtools faidx {} \; && cut -f1,2 *.fai ../reference.fa.fai > haplochroms.dim.tsv
cat haplochroms.dim.tsv | sort  | awk '$2 > maxvals[$1] {lines[$1]=$0; maxvals[$1]=$2} END { for (tag in lines) print lines[tag] }' > maxdi
ms.tsv
awk 'OFS=FS="\t"''{print $1, "0", $2, "100.0", "100.0"}' maxdims.tsv > ../shorts.laser.simple.bed
```

All required files are prepared, we can start reads simulation and alignment.

```
VISOR SHORtS –g chr1.fa –s variation_genome/ -bed shorts.laser.simple.bed –o bam_out/ -threads 7 –c 10
```

##### Randomized CSVs simulation

We only simulate from chr1 to chr22. For each chromosome, we follow the above steps. And in this part, we randomly made combination of different simple event to create the complex events. Before simulation, we need to create a configuration file for each chromosome as listed below. The configure file and CSV file can be found at our Github.

###### Create a configure file

This file contains the initial settings for each chromosome, but can be modified.

```
# Create configure file for simulation
python simulate.py config -f reference.fa.fai -n 4500 -w working_dir/ -l 5000 -s 500
```

The file ***sim_chrom.config.txt*** will be used later and has to be under working_dir.

###### Simulation

Create basic operations for each chromosome independently from chr1 to chr22

```
# Create basic operations
simulate random -w ./working_dir/
# Random select and combine basic operations
simulate add -i ./working_dir/chr1.bed -w /working_dir/ -c chr1
# Add create nested events to the genome
simulate sim -g /path/to/reference.fa -c chr1 -bed chr1.added_SVs.bed -o /path/to/variation_genome/
```

Once the variation genome is created, we follow the step mentioned above to simulate short reads.

### Experimental validation

#### Methods

##### PCR development

The following pipeline has been applied to the PCR assay development. Firstly, the genomic sequence of a 500 bp region next to each SV breakpoint was extracted from UCSC Genome Browser on GRCh38/hg38 Assembly. Secondly, Primer3 Plus (Untergasser et al 2007) is used to compute a set of primer pairs flanking the breakpoint for these regions. 200bp from each side of breakpoint will be excluded from the primer design to avoid the potential uncertainty of breakpoints. Thirdly, the quality score of the primers were checked using Netprimer (PREMIER Biosoft International, Palo Alto, CA) software. The primer would not be used if the quality score was less than 80%. Fourthly, all primer pairs were tested for their uniqueness across the human genome using In Silico PCR from UCSC Genome Browser. BLAT search were also performed at the same time to make sure all primer candidates have only one hit in the human genome. Lastly, NCBI 1000 Genome Browser was used to check if there were any SNPs in the primer or probe binding region. If this does not result in a valid primer pair, the size of the regions for which primers are designed was increased from 500 bp to 750 bp and all process were repeated to search for primers.

##### PCR and Sanger Sequencing

PCR amplifications were performed in 15 $\mu$l reactions using DNA Engine Peltier thermal cycler and C1000 Touch™ thermal cycler (BioRad). Each PCR reaction contained 50 ng of template DNA; 1X PrimeSTAR GXL PCR buffer (1 mM MgCl2) 0.2 mM dNTPs, 100 nM of each primer, and 0.375 U PrimeSTAR® GXL Taq DNA polymerase (Takara). PCR reactions were performed under the following conditions: 32 cycles of denaturation at 98°C for 10 seconds, annealing at 55°C for 15 seconds, and extension at 68°C for 45 seconds to 3 min 45 seconds depending on the predicted PCR amplicon size, followed by a final extension at 68°C for 5 minutes.

PCR products were electrophoresed on a pre-cast gel or a submerged agarose gel. For pre-cast gels, aliquots of 2 $\mu$l of PCR product were electrophoresed in 1.2% or 2% E-gels® containing SYBR (Invitrogen) and visualized with an E-Gel Imager® (Invitrogen). To prepare for Sanger sequencing, 5 ul PCR product was mixed with 2 ul ExoSAP-IT (AppliedBiosystems). Samples were incubated at 37°C for 15 minutes then 80°C for 15 minutes using a C1000 Touch thermal cycler.

For submerged agarose gels, PCR reactions were run in duplicate and the entire PCR product was electrophoresed in 1% agarose gels (1× TAE) containing 0.1 $\mu$g/ml SYBR Gold

(Molecular Probes Inc.) for 45-120 minutes at 100-120V. DNA fragments were visualized on a ChemiDoc™ MP (BioRad) using a UV light and filter. The target band was cut from the gel and purified using the NucleoSpin® Gel and PCR Clean-Up kit (Macherey-Nagel) following the manufacturer's instructions. The DNA concentrations were measured using the Qubit® 2.0 (ThermoFisher). The optimal DNA concentration for Sanger sequencing would be between 10-20 ng/µl. Samples were then sent to Eton Bioscience to be sequenced.

#### Results

##### Validation of all CSVs detected by Mako

We evaluated the accuracy of CSV detection given by Mako of HG00733, which contains 609 autosomes CSVs. Due to the large number of primers to desgin, we used the Primer3 command line options with defualt parameters except the parameter ***SEQUENCE PRIMER PAIR OK REGION LIST***. This parameter is set based on the region 200bp outside the breakpoint but within the extended flanking region. Table 2 shows the validated CSVs and PacBio refined types.

| Chrom | Start | End | PacBio Type |
|---|---|---|---|
| chr1 | 81,194,398 | 81,195,874 | invDup |
| chr2 | 119,659,504 | 119,661,322 | insDup |
| chr3 | 146,667,093 | 146,677,284 | delDisDup |
| chr5 | 141,480,327 | 141,483,116 | disDup |
| chr7 | 1,940,931 | 1,941,009 | insDup |
| chr9 | 29,591,409 | 29,593,057 | delINV |
| chr10 | 14,568,488 | 14,568,677 | insDup |
| chr12 | 71,315,482 | 71,316,928 | invDup |
| chr12 | 77,989,900 | 77,994,324 | invDup |
| chr13 | 74,340,759 | 74,342,810 | disDup |
| chr16 | 78,004,459 | 78,007,456 | disDup |
| chr17 | 34,854,438 | 34,855,851 | invDup |
| chr17 | 48,538,270 | 48,540,171 | disDup |
| chr18 | 72,044,575 | 72,045,937 | disDup |
| chr21 | 26,001,844 | 26,002,990 | delINV |

**Table 1.** Summary of experimental validation of PCR succeed CSVs.

##### Validation of CSVs at coding regions

After evaluating all CSVs given by Mako, we further validated some gene overlapping CSVs as follows. A detailed analysis of Sanger results of these events are shown in **Figure 1-4**.

| Sample | Gene | SV Type | Validation Strategy | Status |
|---|---|---|---|---|
| HG00514 | CARD6 | DEL+INV | Sanger sequencing | validated |
| HG00514 | SORBS1 (CAP) | INS+DUP | Sanger sequencing | validated |
| HG00713 | SORBS1 (CAP) | INS+DUP | Sanger sequencing | validated |
| NA19240 | C4BPA | DEL+INV | Sanger sequencing | validated |
| NA19240 | TSPAN8 | INS+INV | Sanger sequencing | not validated |
| HG00514 | SPOCK3 | DEL+INV | Sanger sequencing | validated |
| NA19240 | MMP8 | INS+INV | Sanger sequencing | not validated |

**Table 2.** Summary of experimental validation of novel complex SVs

The Sanger sequencing not only validated the SV events predicted by the Mako algorithm, but not deciphered the exact breakpoints. For example, Mako algorithm predicted a 751bp deletion from the C4BPA gene, and meanwhile a 153bp fragment upstream of this deletion was inverted (**Figure 1**). Our Sanger was able to identify the exact breakpoints for both predicted deletion and inversion events, with a few basepairs away from the predicted breakpoints (**Figure 2**). Interestingly, around 90% of CEU population has deletion (44.4% heterozygous and 45.4% homozygous deletions) at this locus based on the 1000 Genomes dataset (Sudmant et al 2015).

In some other cases such as the CARD6 gene in the sample HG00514, Mako algorithm was able to identify the exact breakpoints. In this gene Mako algorithm predicted a 2941bp deletion together with a 440bp fragment inversely inserted upstream region of the deletion (**Figure 3**). The Sanger identified the deletion breakpoints and the left inversion breakpoints a few basepairs away from the predicted ones. However, the validated right breakpoint from the inversion exactly matches the prediction, indicating the high sensitivity of the Mako algorithm (**Figure 4**).

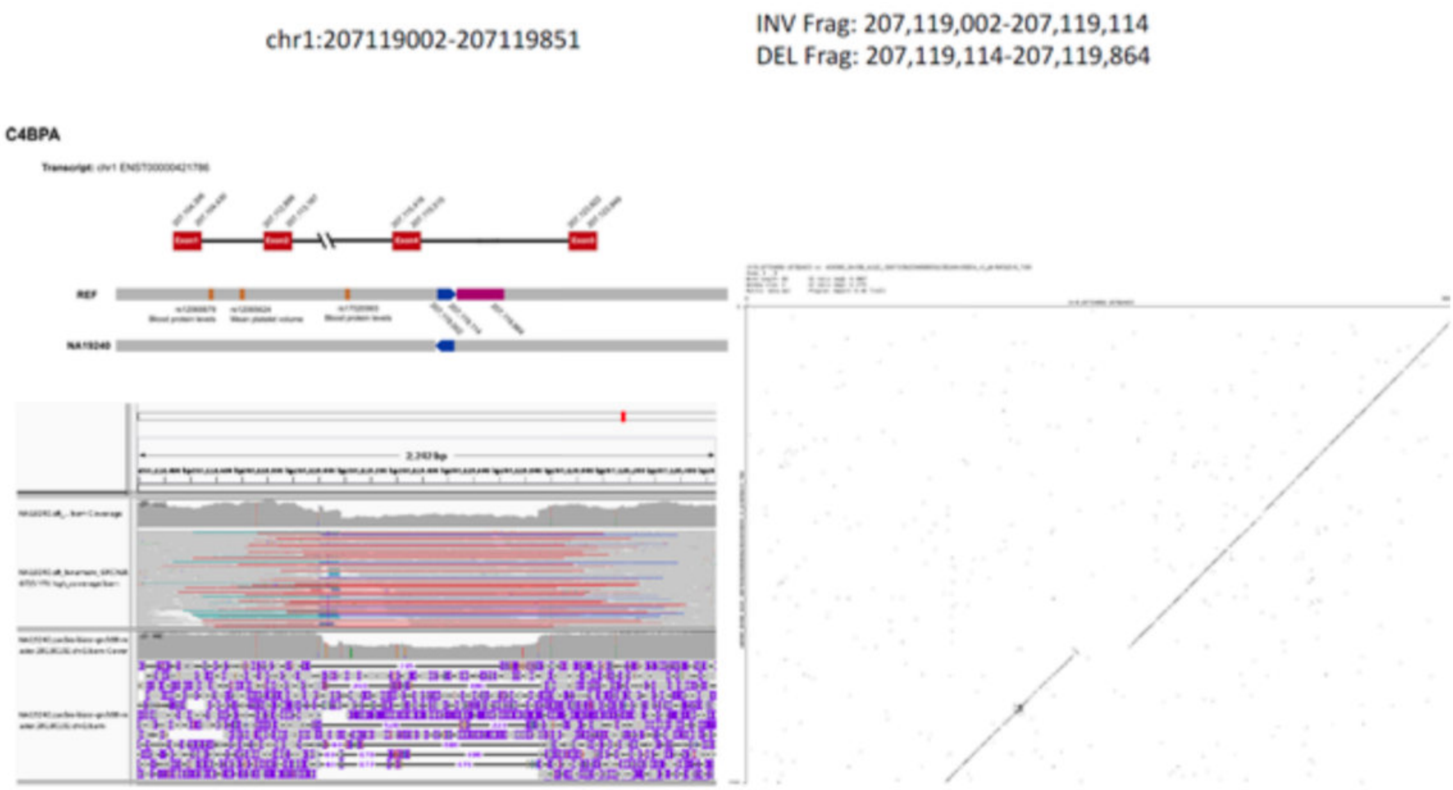


**Figure N1.** Sanger sequencing validation of a deletion and inversion event.

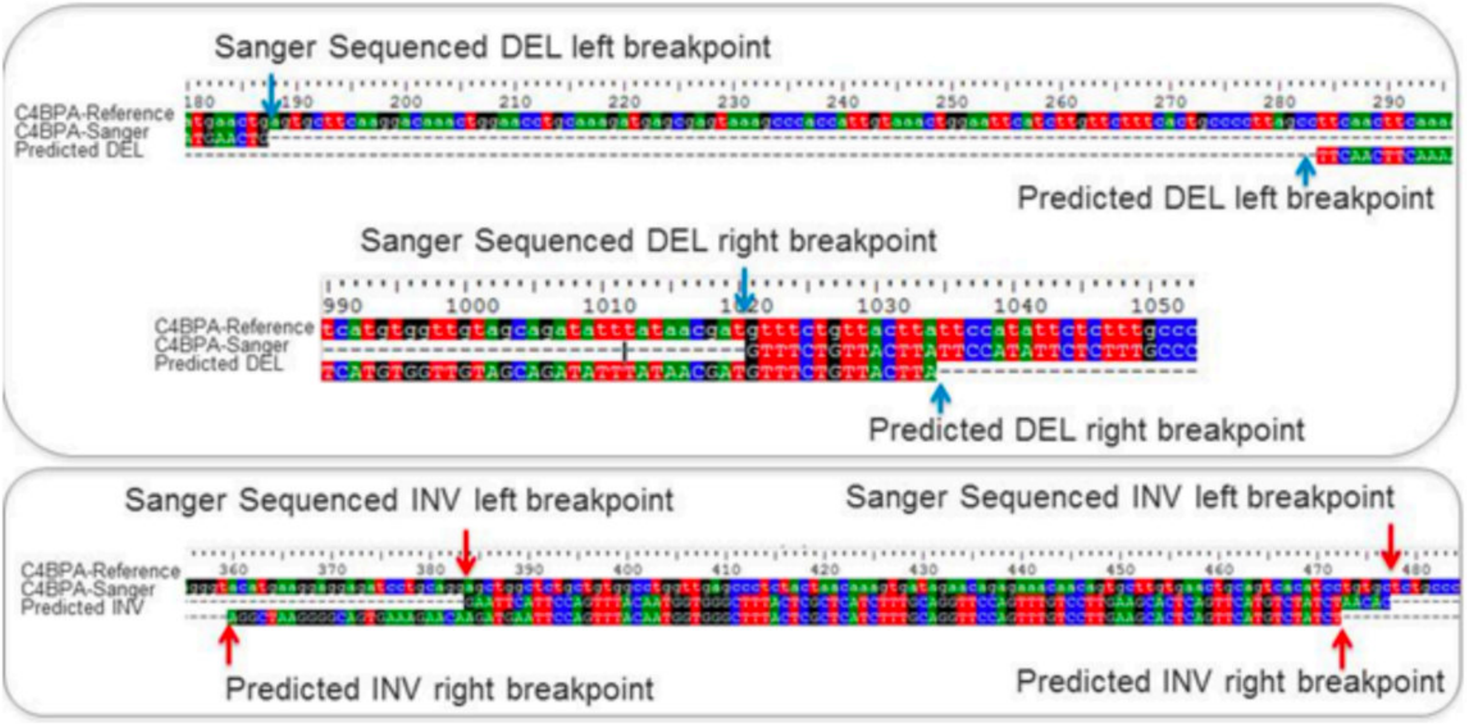


**Figure N2.** Sanger sequencing validation on deletion and inversion at C4BPA gene from NA19240.

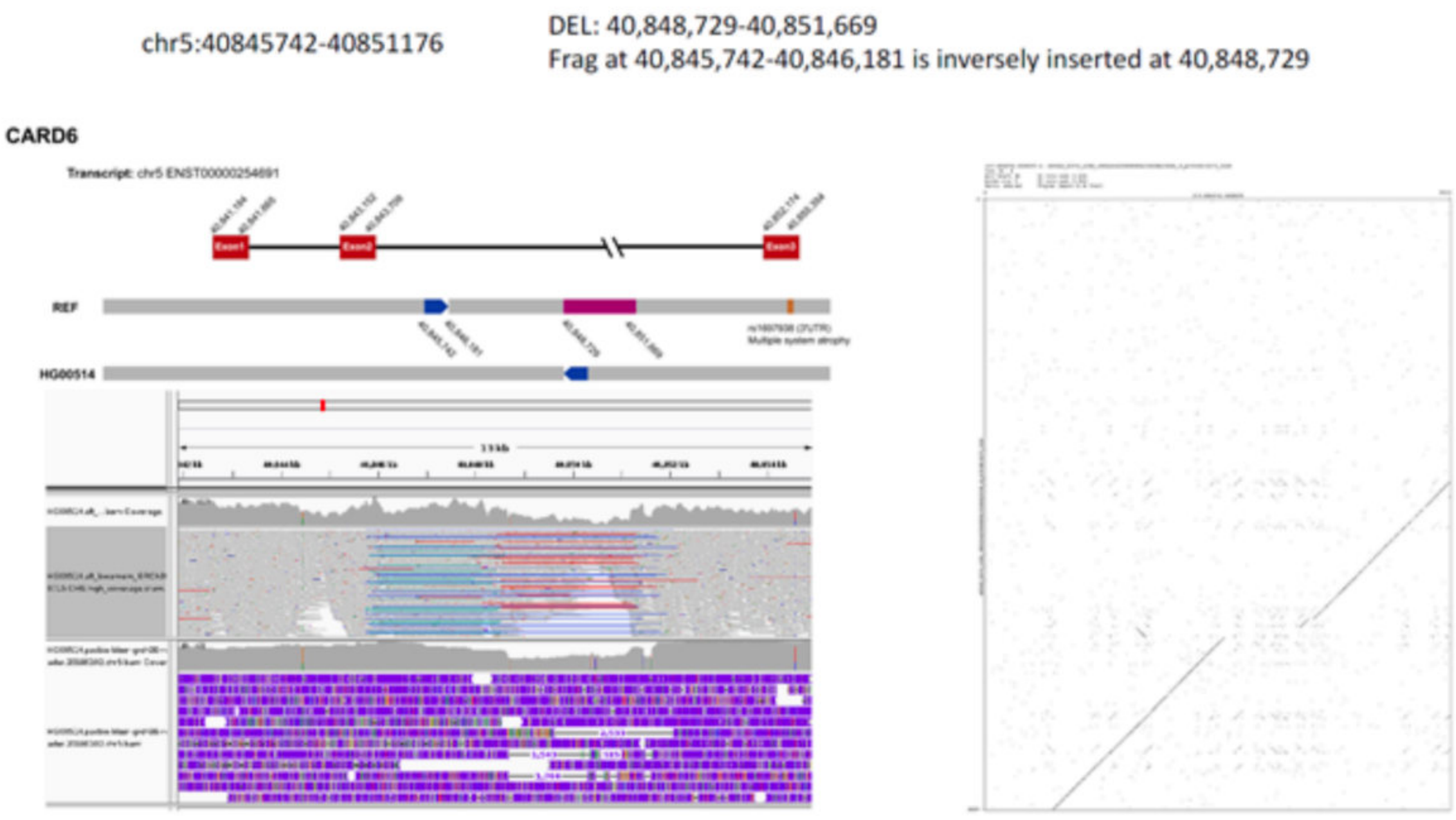


**Figure N3.** Sanger sequencing validation on deletion and inverted duplication event.

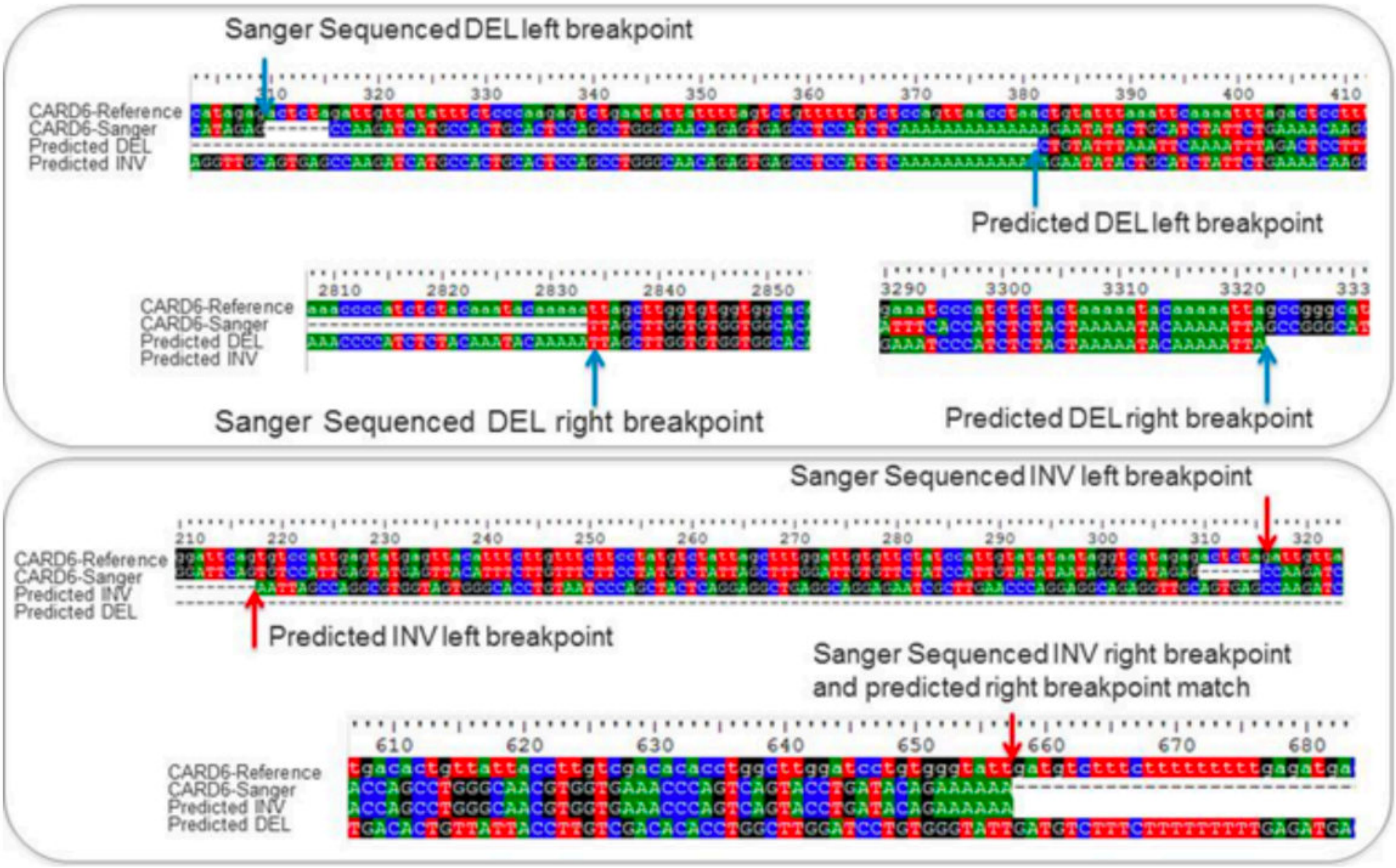


**Figure N4.** Sanger sequencing validation on deletion and inversion at CARD6 gene from HG00514.

#### CSV manual inspection

###### External sources

1. PacBio long-read sequencing (HGSVC)

2. Gepard (https://github.com/univieCUBE/gepard).

###### Major steps

1. Filtering Mako's CSVs in excludable regions for each sample. The GRCh38 exclude regions are provided by SVelter (https://github.com/mills-lab/svelter/tree/master/Support/GRCh38).
2. We extract each PacBio reads with SAMtools and create the sequence Dotplot with Gepard. We use k=15 to build create the sequence Dotplot. In total, we created in average 30 Dotplots for each event.

###### Dotplot inspection

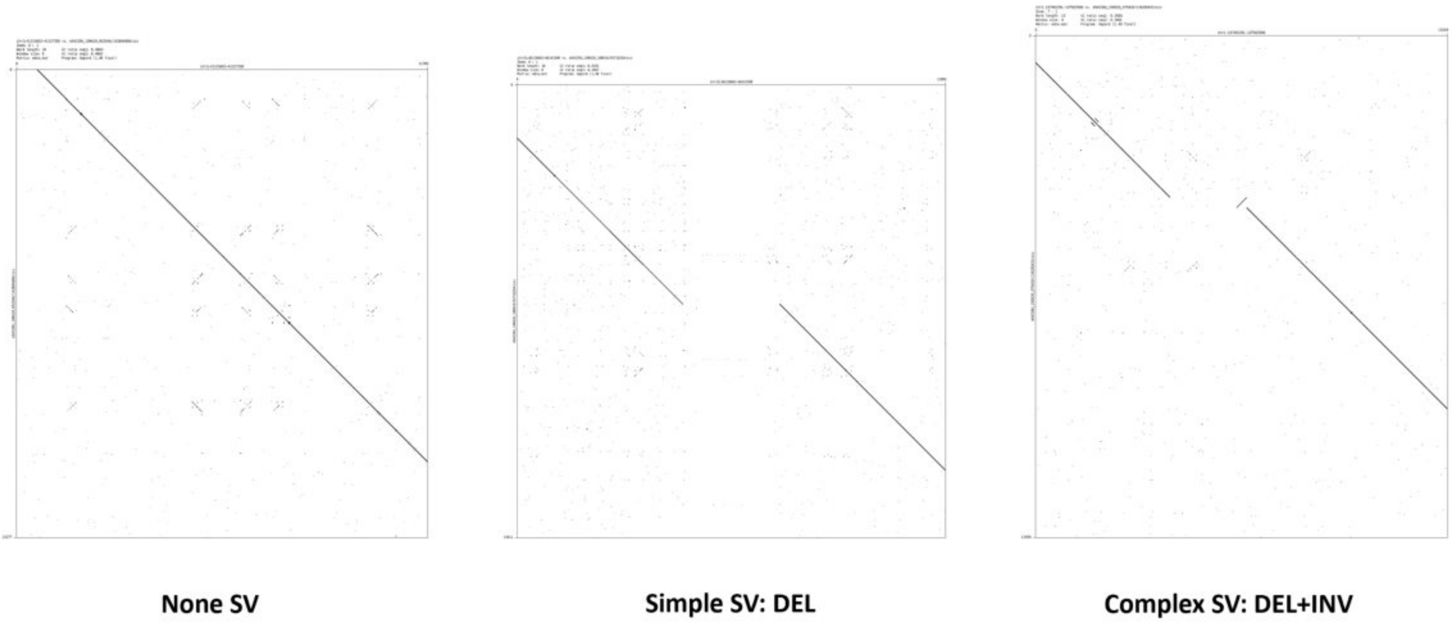


#### Computational validation

##### Methods

###### External sources

1. VaPoR (https://github.com/mills-lab/vapor).
2. Assemblies alignment of HiFi haploid reads (http://ftp.1000genomes.ebi.ac.uk/vol1/ftp/data*collections/HGSVC2/working/20200628*HHU*assembly-results*CCS*v12/haploid*reads/).
3. K-Mer match Python script.

###### Vapor ONT validation

```
VaPoR bed --sv-input csvs.bed --output-path /path/to/vapor_out/ --reference /path/to/reference.fa --pacbio-input /path/to/contig_align.bam
```

###### Kmer HiFi contig validation

The source code can be found in our GitHub repo.

##### Results

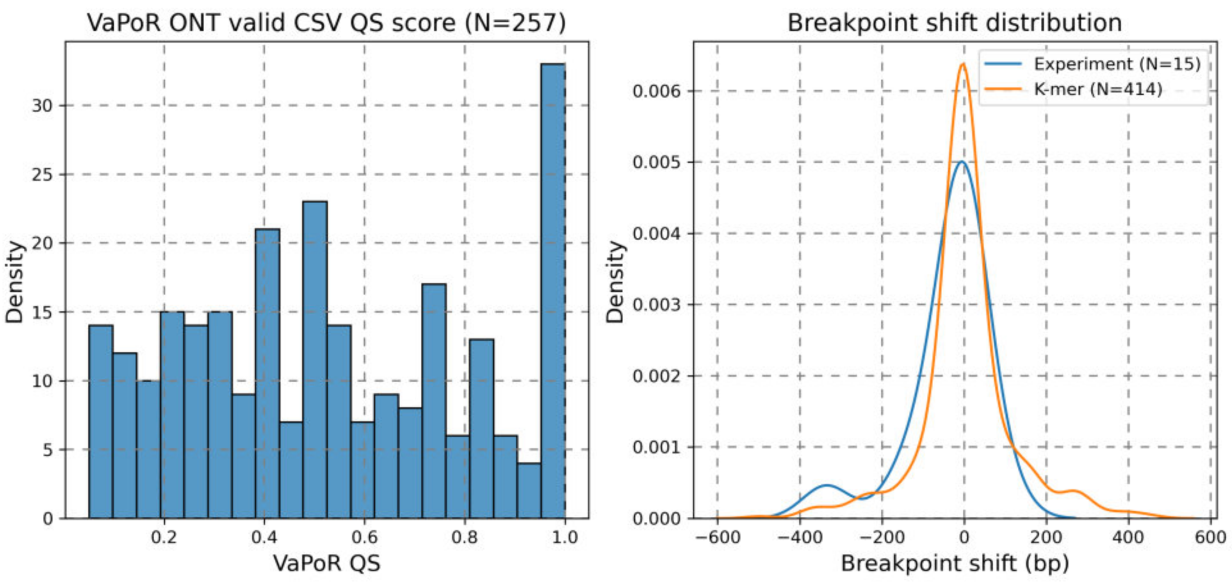


### Running configurations

#### Mako

##### Running dependencies

Mako is a standalone java program. Python is only used to create CSV benchmarks and simulate CSVs. * Java (jdk >= 1.8) * htsjdk: https://github.com/samtools/htsjdk

###### Step1: Create configuration file

**NOTE:** BAM file **NEED** to be under work*dir, and please use absolute path for work*dir

```
# Get Mako configuration file
python process.py config -b /path/to/sample.bam -w /path/to/work_dir/ -n 30000 -s sampleName -f /path/to/ref.fa.fai

# Example of a config file for simulation data
mean:499
stdev:50
readlen:150
workDir:/path/to/work_dir/
bam:/path/to/sample.bam
name:sim
```

###### Step2: SV detection

Using config file create in previous step. The config file usually names as sampleName.mako.cfg.

```
java -jar Mako.jar -R /path/to/your/reference.fa -F /path/to/your/sampleName.mako.cfg
```

###### Step3: convert Mako output to VCF format

```
python ParseMako.py tovcf -m sampleName_mako_calls.txt -o sampleName_mako.vcf -r /path/to/ref.fa -s sampleName
```

The default output is ***sampleName.mako.sites.txt***.

#### Manta

Version=1.6.0

```
# Create Manta workflow
${MANTA_INSTALL_PATH}/bin/configManta.py \
--bam sample.bam \
--referenceFasta GRCh38.fa \
--runDir ${MANTA_ANALYSIS_PATH}--bam

# Run workflow
python ${MANTA_ANALYSIS_PATH}/runWorkflow.py -j 8
```

#### SVelter

Version=0.1

```
svelter.py Setup --reference GRCh38.fa --workdir /working/directory/ --support ../Support/GRCh38/

svelter.py NullModel --sample /absolute/path/of/sample.bam --workdir /working/directory

svelter.py BPSearch --sample /absolute/path/of/sample.bam --workdir /working/directory

svelter.py BPIntegrate --sample /absolute/path/of/sample.bam --workdir /working/directory

svelter.py SVPredict --sample sample.bam --workdir /working/directory --bp-file sample.bam/sample.bam.txt

svelter.py SVIntegrate --workdir /working/directory --prefix output  --input-path path/of/output/from/Step4
```

#### Lumpy

Version=0.2.13

```
# Add read group tag to simulated BAM file
samtootls addreplacerg -r ID:wgs_sim -r LB:visor_sim -r SM:wgs_sim -o simrg.srt.bam sim.srt.bam
samtools index simrg.srt.bam

# Extract the discordant paired-end alignments.
samtools view -b -F 1294 sample.bam > sample.discordants.unsorted.bam
# Extract the split-read alignments

samtools view -h sample.bam \
    | scripts/extractSplitReads_BwaMem -i stdin \
    | samtools view -Sb - \
    > sample.splitters.unsorted.bam

# Sort both alignments
samtools sort sample.discordants.unsorted.bam sample.discordants
samtools sort sample.splitters.unsorted.bam sample.splitters

# Call SVs
lumpyexpress \
    -B sample.bam \
    -S sample.splitters.bam \
    -D sample.discordants.bam \
    -o sample.vcf
```

#### GRIDSS

Version=2.6.2

```
bash /path/to/gridss.sh --reference GRCh38.fa --output sample.vcf.gz --assembly sample.assembly.bam --threads 1 --jar /path/to/gridss-2.6.2
-gridss-jar-with-dependencies.jar --workingdir /path/to/workdir sample.bam
```

#### TARDIS

Version=1.0.6

First download the sonic file of GRCh38 (GRCh38_1kg.sonic)

```
tardis -i /path/to/sample.bam --ref /path/to/reference.fa --sonic /path/to/GRCh38_1kg.sonic --out /path/to/tardis.vcf --threads 6
```
